## Supplementary material for "Comparing rates of introgression in parasitic feather lice with differing dispersal capabilities": OUT_sppIDerDepthPlot-covBins_Cosp_Cobuc_11_2_2016_2.pdf (from another version of this preprint: version 7, 2020-03-31)

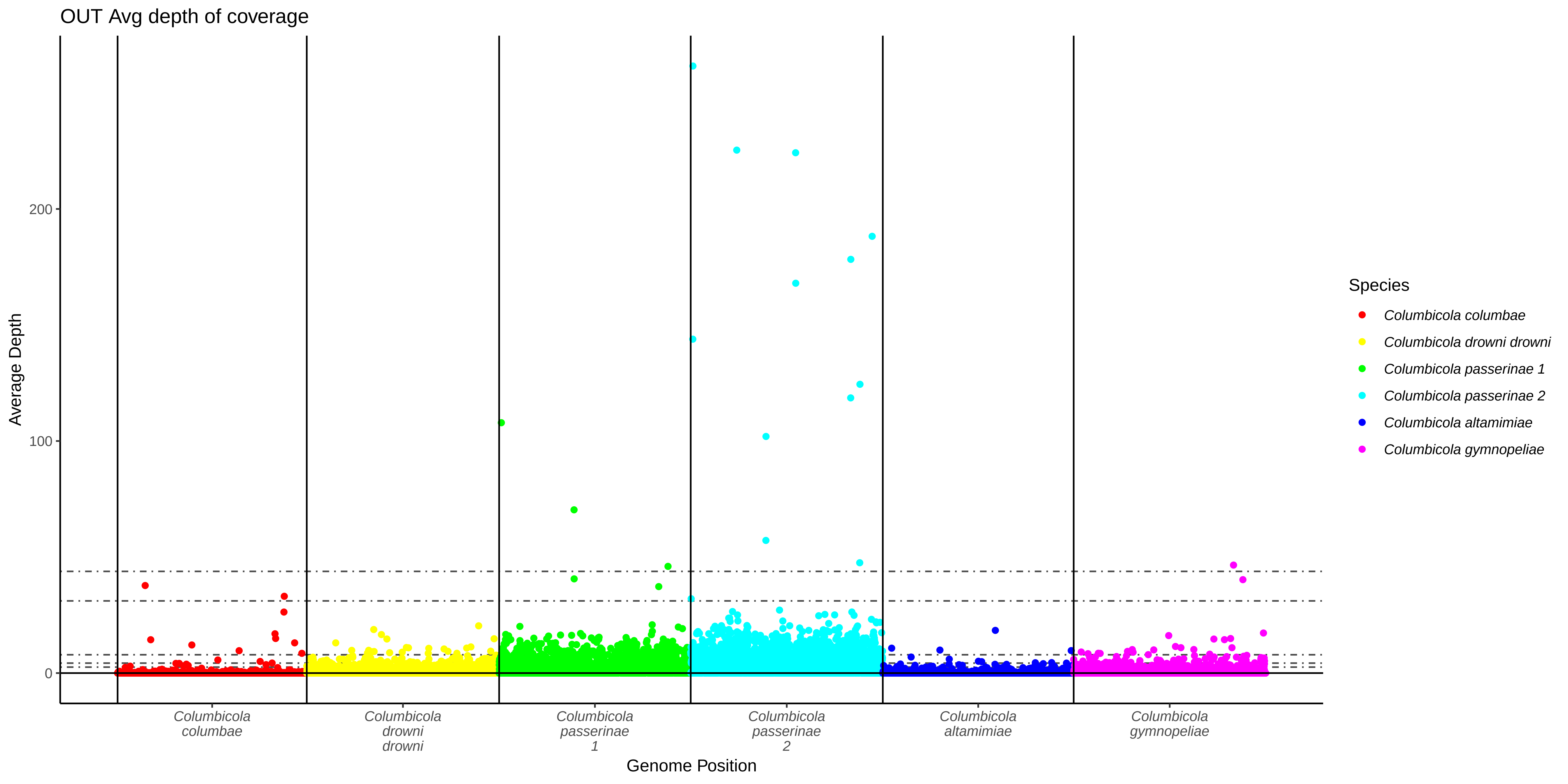
OUT Avg depth of coverage
Average Depth
200
100
0
Columbicola columbae
Columbicola drowni drowni
Columbicola passerinae 1
Columbicola passerinae 2
Columbicola altamimiae
Columbicola gymnopeliae
Genome Position
Species
Columbicola columbae
Columbicola drowni drowni
Columbicola passerinae 1
Columbicola passerinae 2
Columbicola altamimiae
Columbicola gymnopeliae

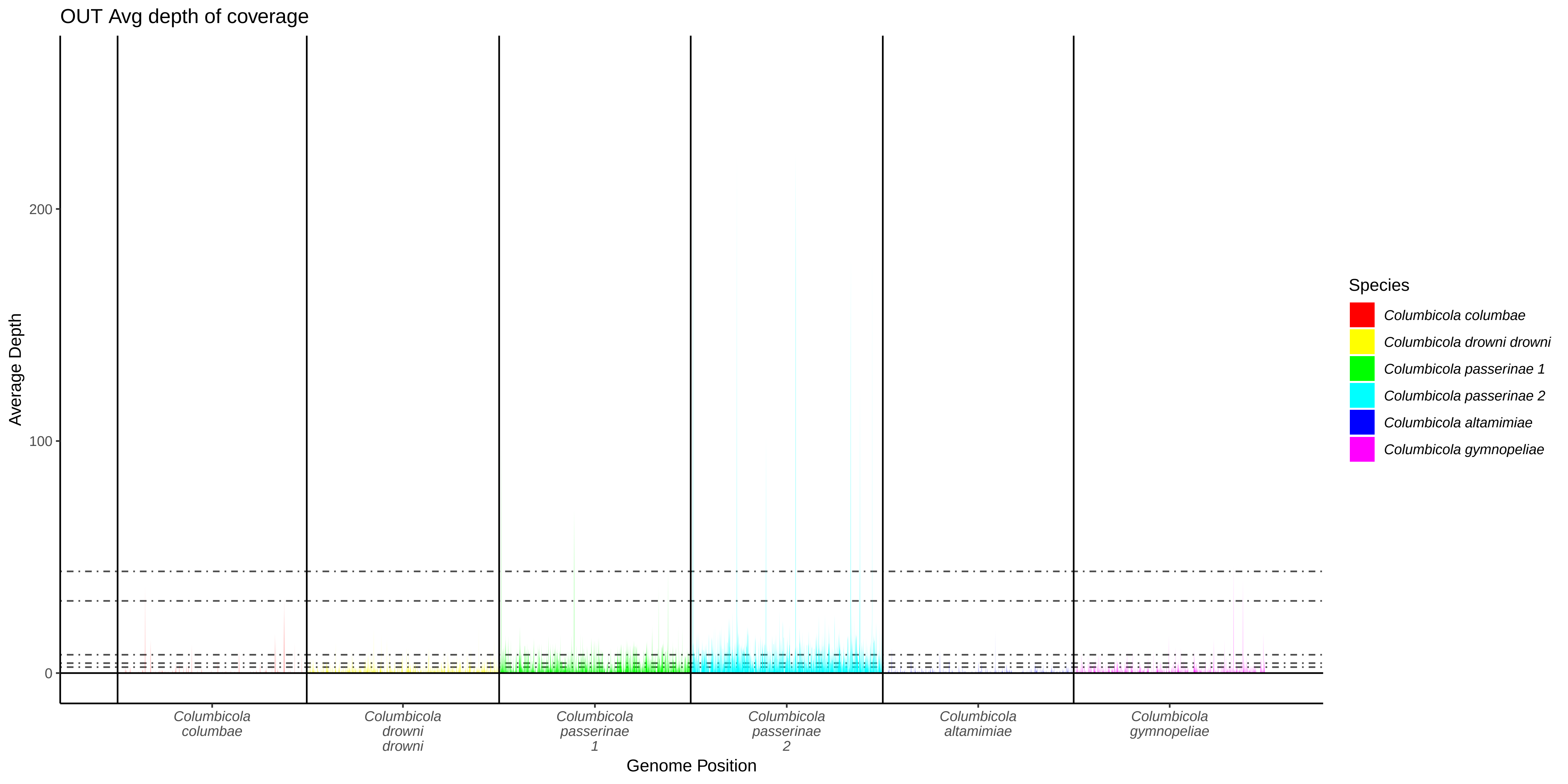

OUT Avg depth of coverage
Average Depth
200
100
0
Columbicola columbae
Columbicola drowni drowni
Columbicola passerinae 1
Columbicola passerinae 2
Columbicola altamimiae
Columbicola gymnopeliae
Genome Position
Species
Columbicola columbae
Columbicola drowni drowni
Columbicola passerinae 1
Columbicola passerinae 2
Columbicola altamimiae
Columbicola gymnopeliae

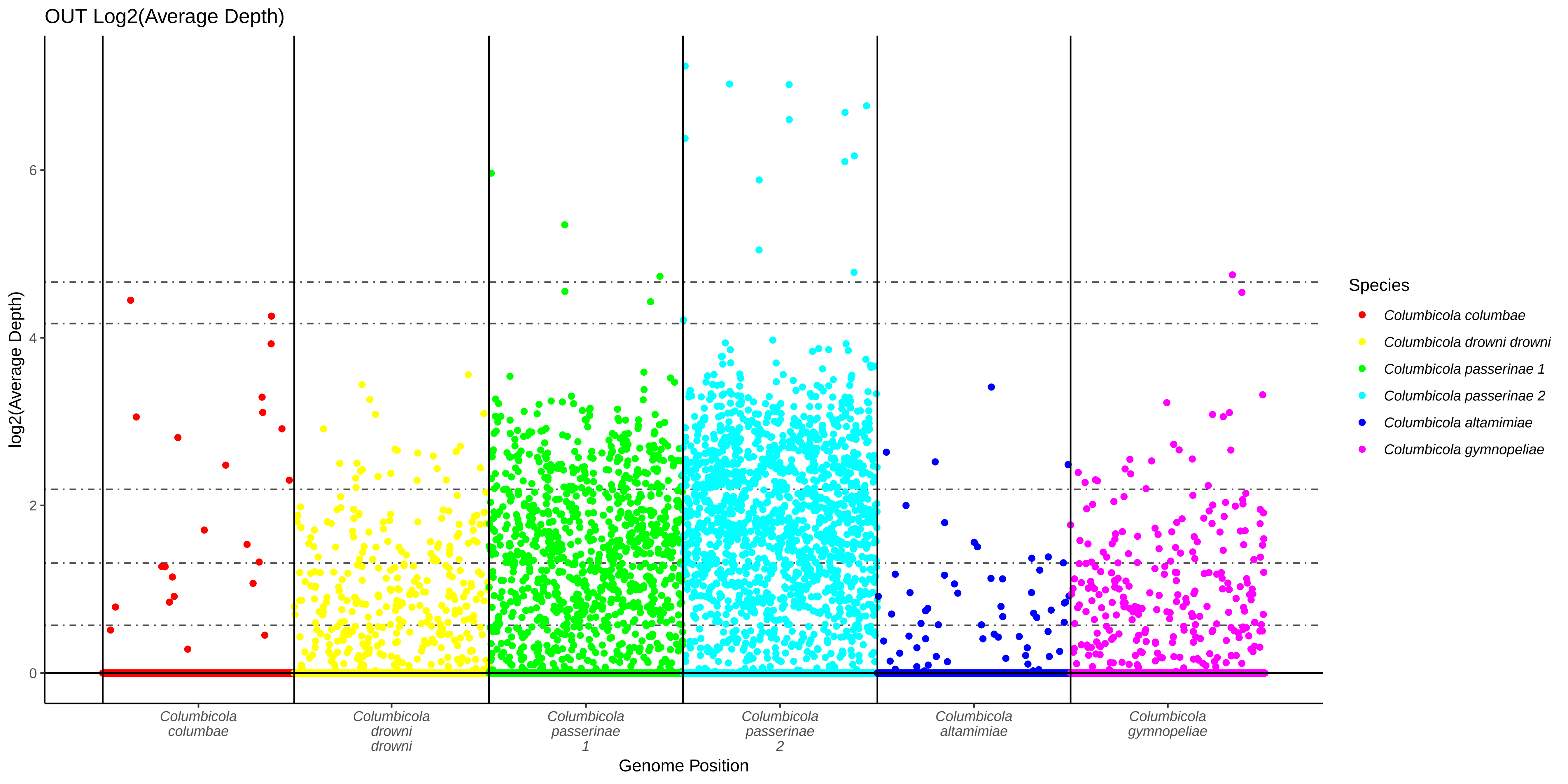

OUT Log2(Average Depth)
log2(Average Depth)
0
2
4
6
Genome Position
Columbicola columbae
Columbicola drowni drowni
Columbicola passerinae 1
Columbicola passerinae 2
Columbicola altamimiae
Columbicola gymnopeliae
Species
Columbicola columbae
Columbicola drowni drowni
Columbicola passerinae 1
Columbicola passerinae 2
Columbicola altamimiae
Columbicola gymnopeliae

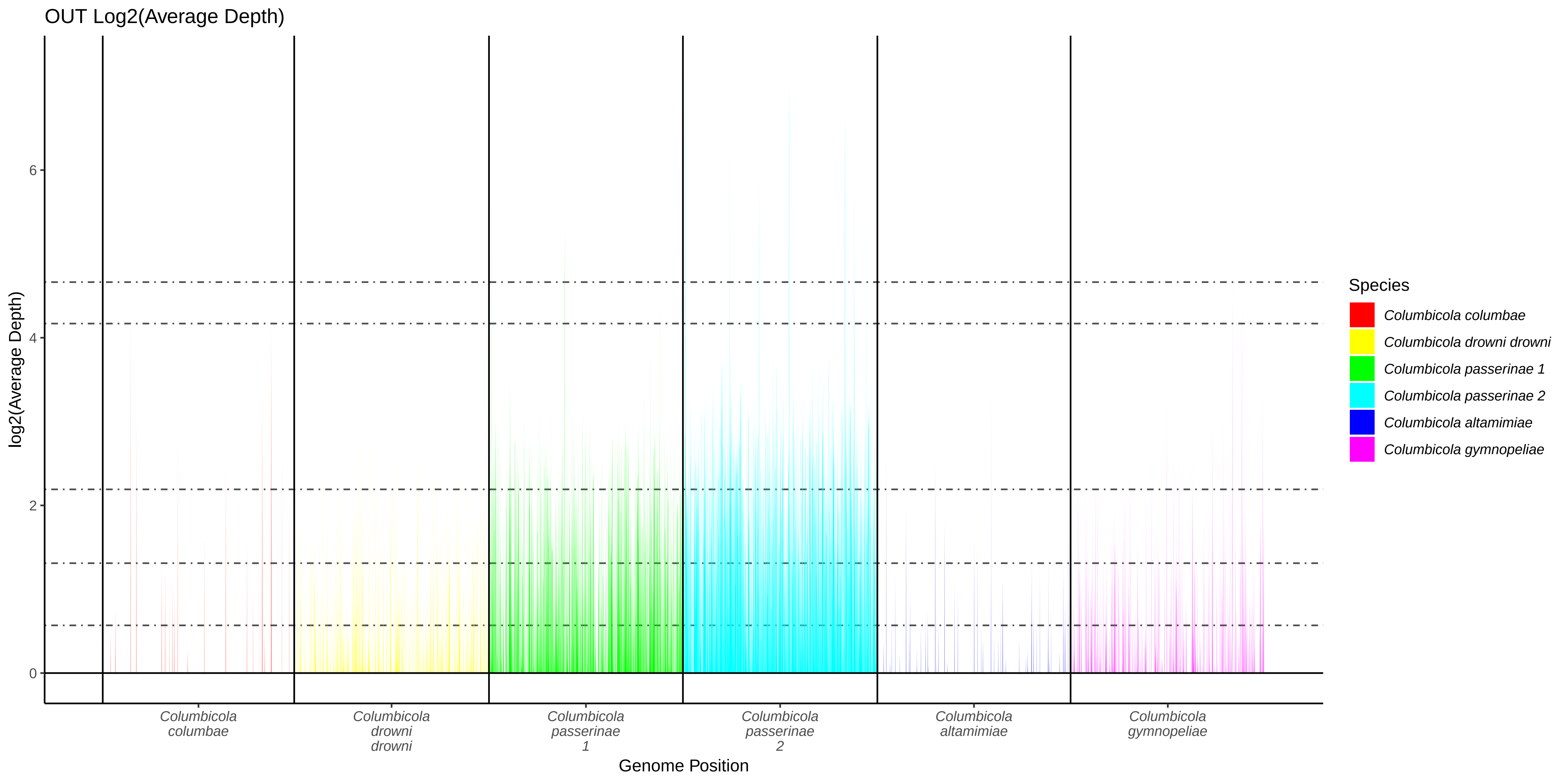
OUT Log2(Average Depth)
log2(Average Depth)
6
4
2
0
Columbicola columbae
Columbicola drowni drowni
Columbicola passerinae 1
Columbicola passerinae 2
Columbicola altamimiae
Columbicola gymnopeliae
Genome Position
Species
Columbicola columbae
Columbicola drowni drowni
Columbicola passerinae 1
Columbicola passerinae 2
Columbicola altamimiae
Columbicola gymnopeliae

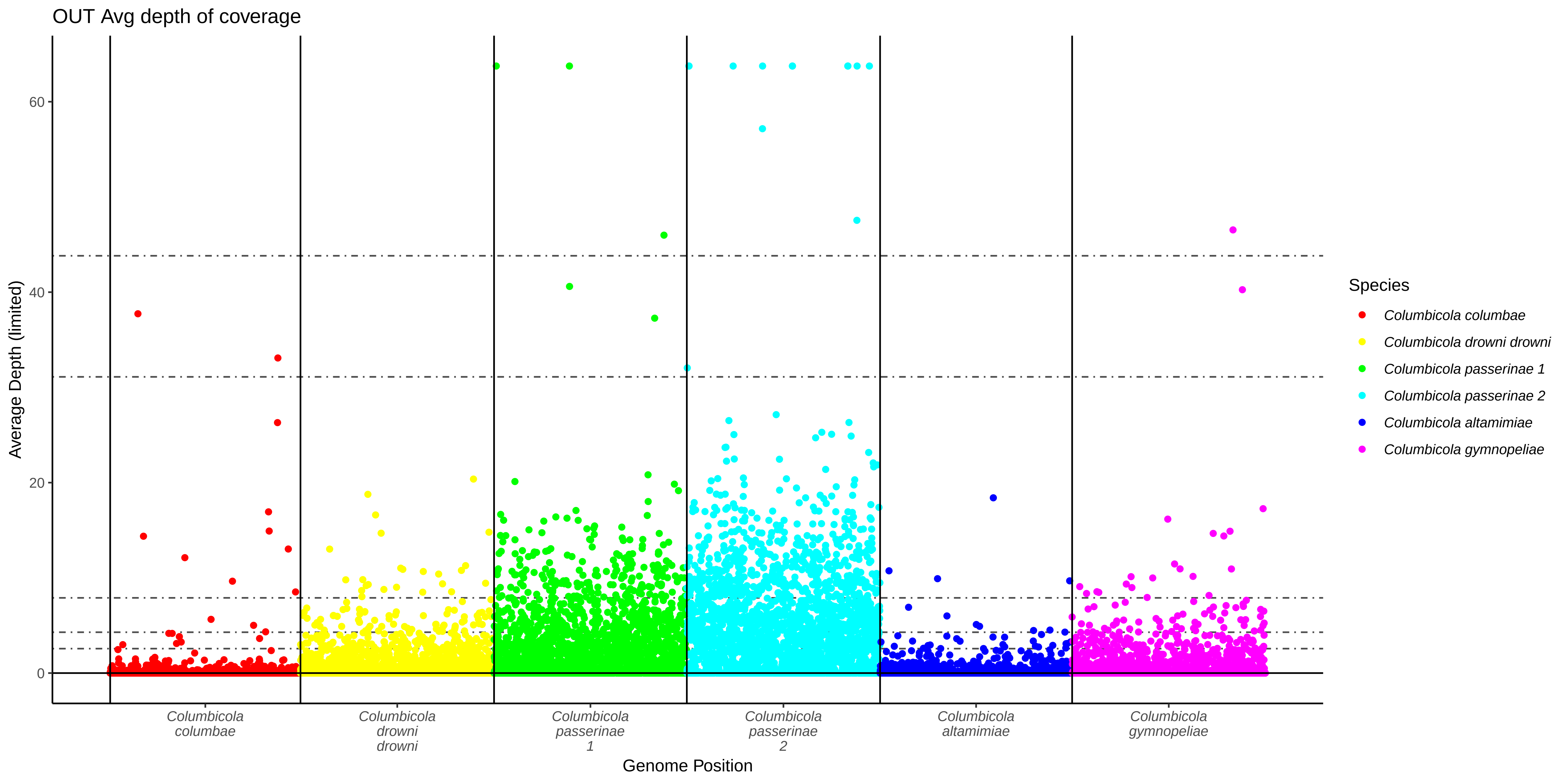
OUT Avg depth of coverage
Average Depth (limited)
0
20
40
60
Genome Position
Columbicola columbae
Columbicola drowni drowni
Columbicola passerinae 1
Columbicola passerinae 2
Columbicola altamimiae
Columbicola gymnopeliae
Species
Columbicola columbae
Columbicola drowni drowni
Columbicola passerinae 1
Columbicola passerinae 2
Columbicola altamimiae
Columbicola gymnopeliae

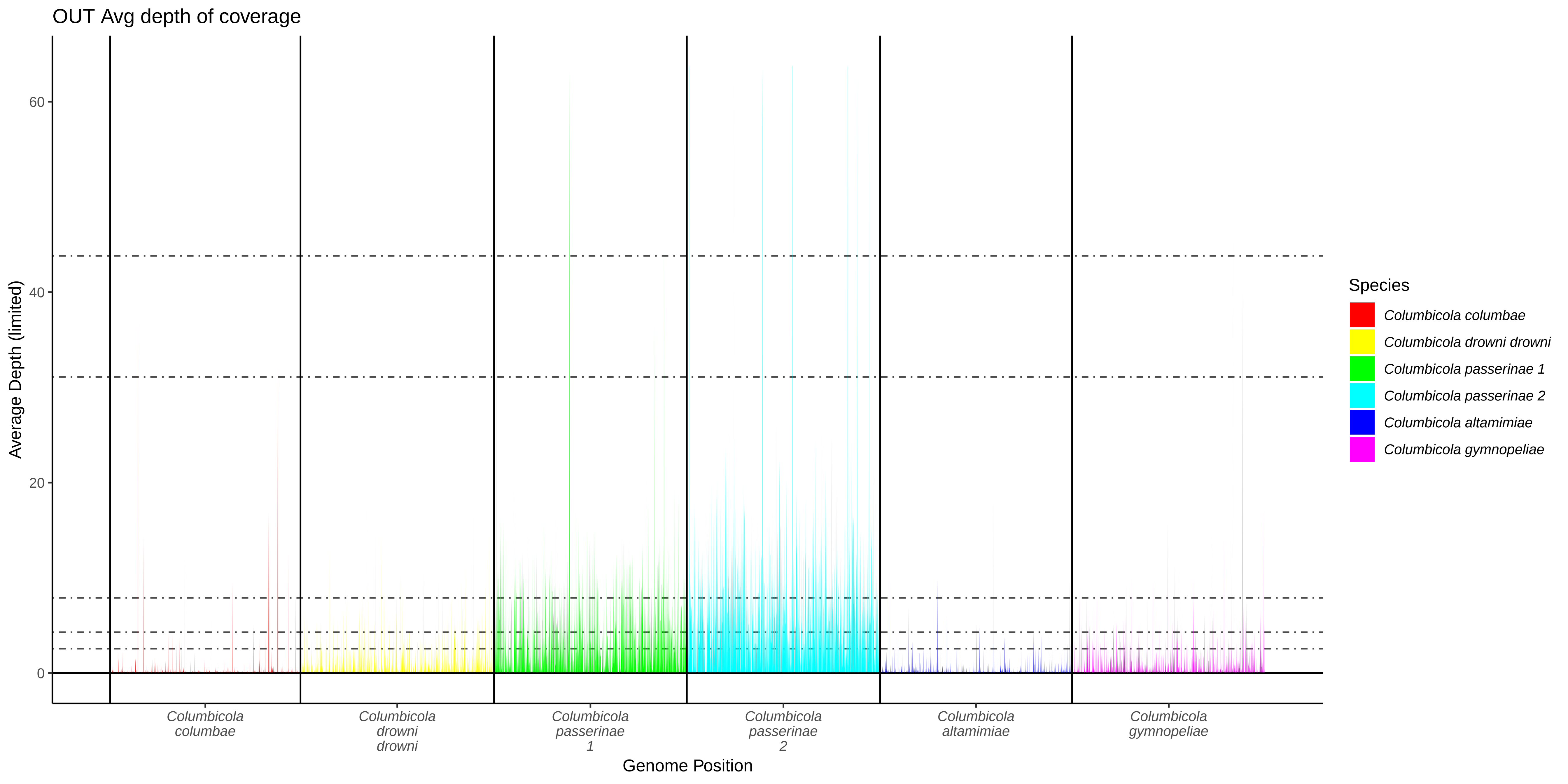

OUT Avg depth of coverage
Average Depth (limited)
0
20
40
60
Columbicola columbae
Columbicola drowni drowni
Columbicola passerinae 1
Columbicola passerinae 2
Columbicola altamimiae
Columbicola gymnopeliae
Genome Position
Species
Columbicola columbae
Columbicola drowni drowni
Columbicola passerinae 1
Columbicola passerinae 2
Columbicola altamimiae
Columbicola gymnopeliae

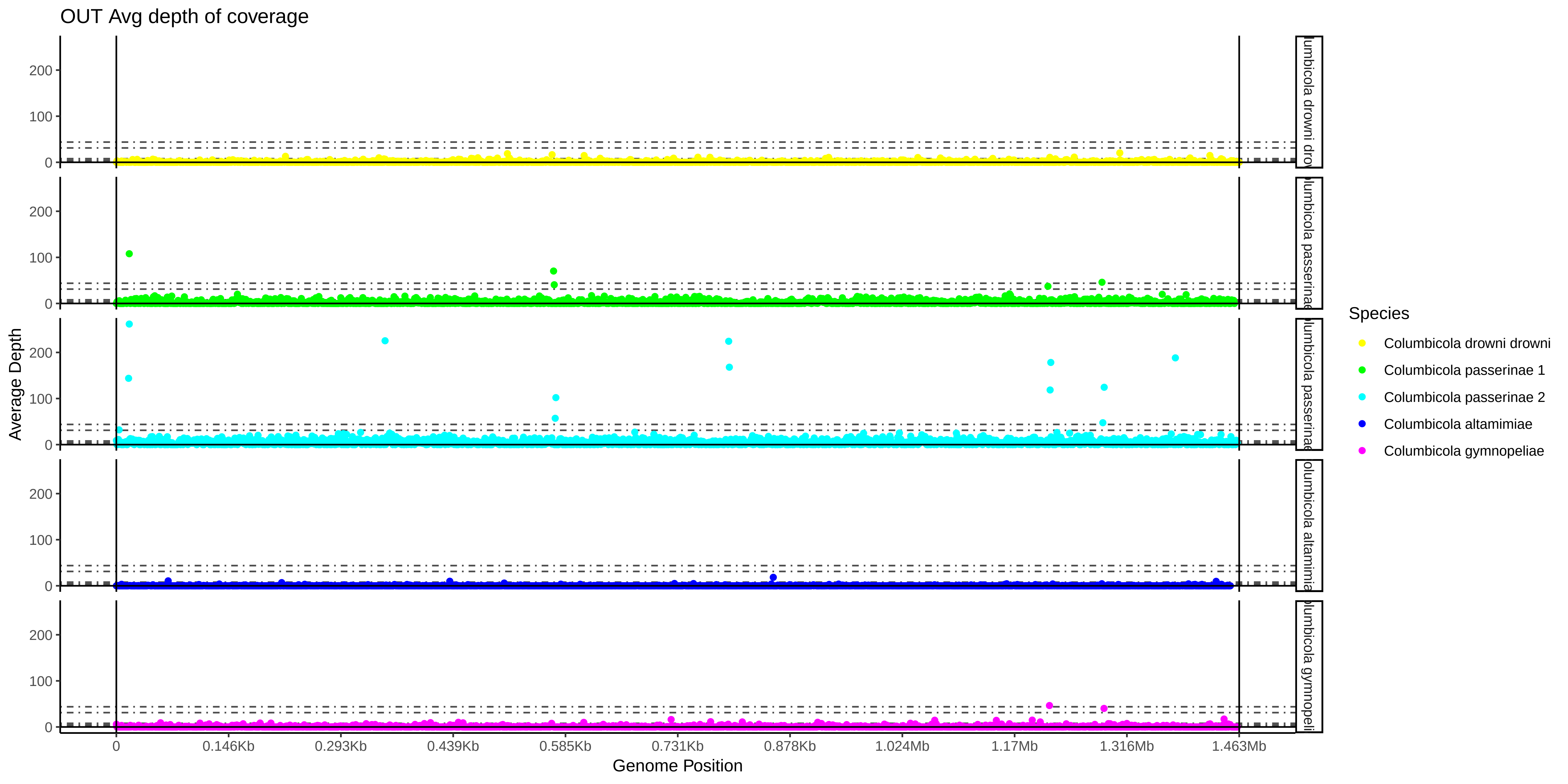
OUT Avg depth of coverage
Average Depth
Genome Position
0
100
200
0
0.146Kb
0.293Kb
0.439Kb
0.585Kb
0.731Kb
0.878Kb
1.024Mb
1.17Mb
1.316Mb
1.463Mb
umbicola drowni dro
umbicola passerinae
umbicola passerinae
olumbicola altamimia
umbicola gymnopeli
Species
Columbicola drowni drowni
Columbicola passerinae 1
Columbicola passerinae 2
Columbicola altamimiae
Columbicola gymnopeliae

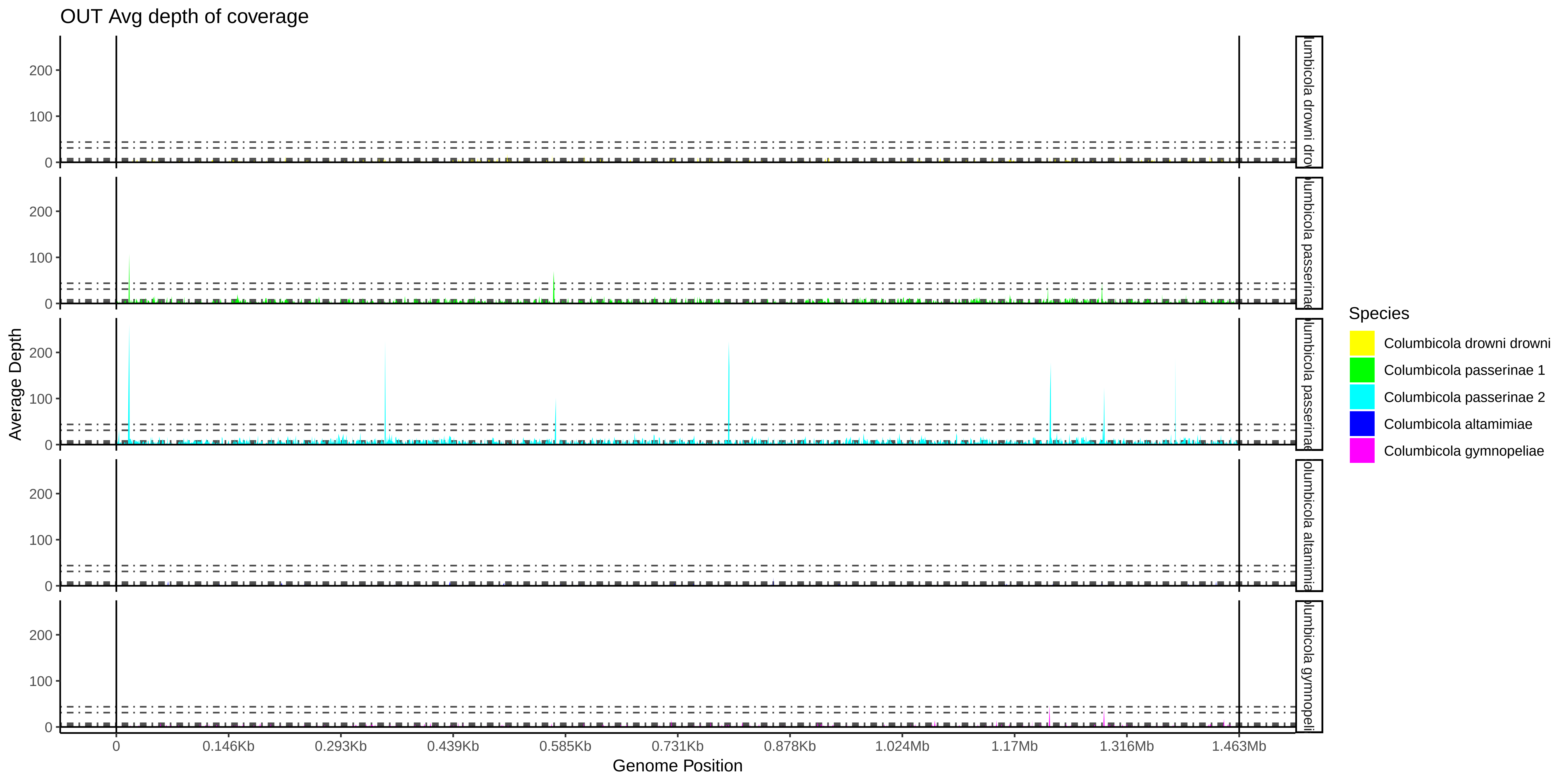
OUT Avg depth of coverage
Average Depth
Genome Position
0
100
200
0
0.146Kb
0.293Kb
0.439Kb
0.585Kb
0.731Kb
0.878Kb
1.024Mb
1.17Mb
1.316Mb
1.463Mb
Species
Columbicola drowni drowni
Columbicola passerinae 1
Columbicola passerinae 2
Columbicola altamimiae
Columbicola gymnopeliae

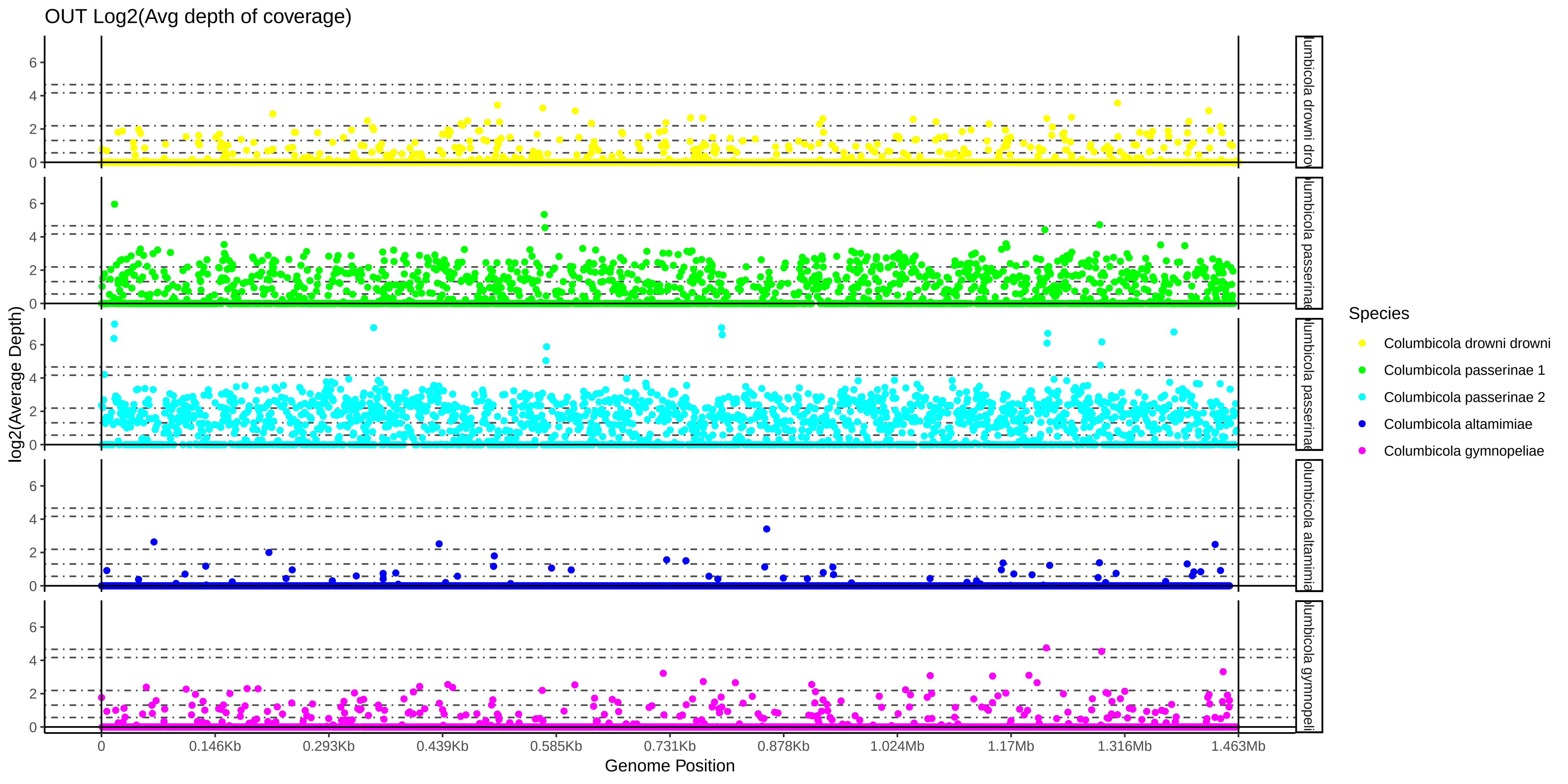

OUT Log2(Avg depth of coverage)
log2(Average Depth)
Genome Position
0
0.146Kb
0.293Kb
0.439Kb
0.585Kb
0.731Kb
0.878Kb
1.024Mb
1.17Mb
1.316Mb
1.463Mb
0
2
4
6
umbicola drowni dro
umbicola passerinae
umbicola passerinae
olumbicola altamimia
umbicola gymnopeli
Species
Columbicola drowni drowni
Columbicola passerinae 1
Columbicola passerinae 2
Columbicola altamimiae
Columbicola gymnopeliae

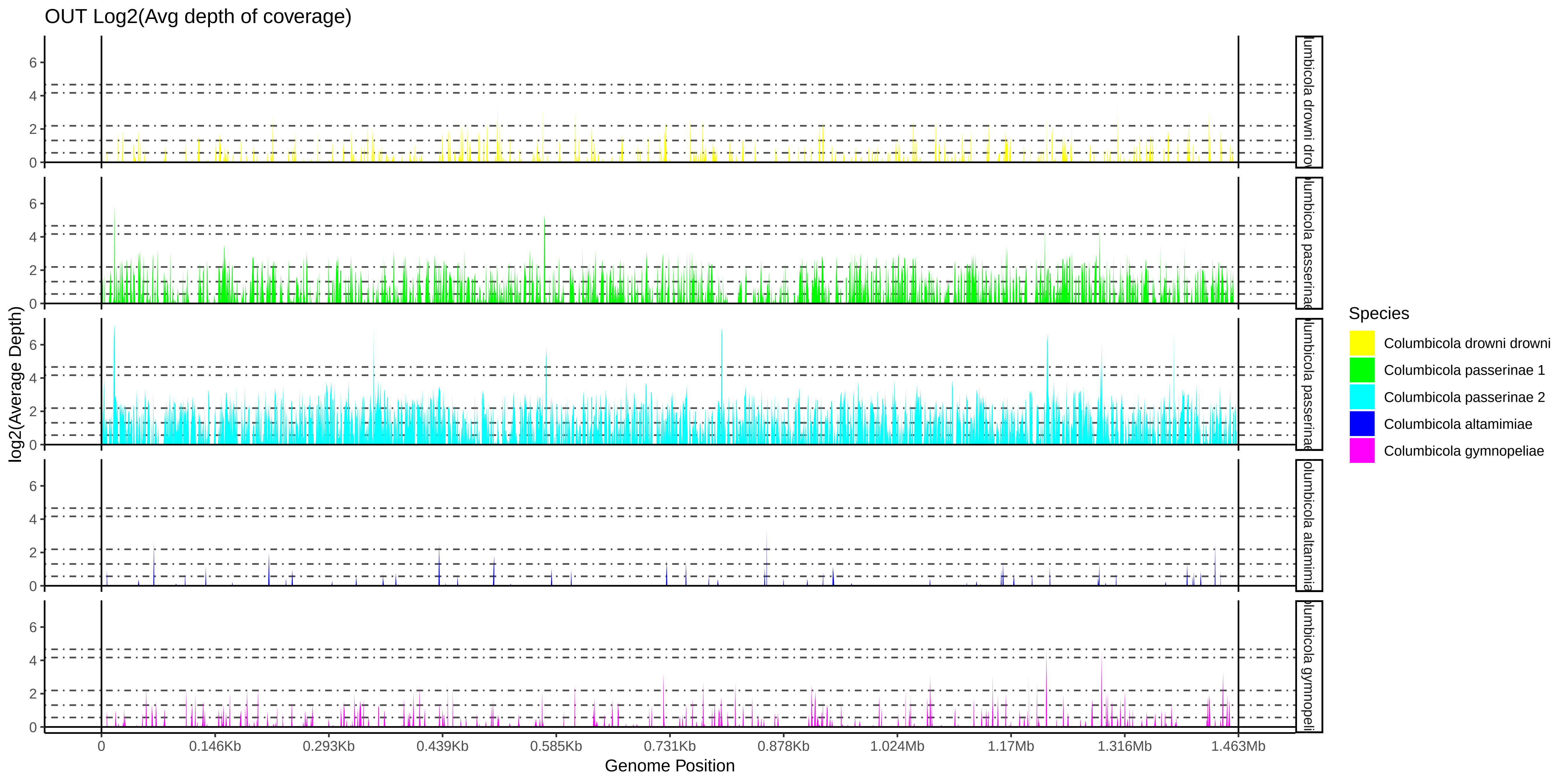

OUT Log2(Avg depth of coverage)
log2(Average Depth)
Genome Position
0
0.146Kb
0.293Kb
0.439Kb
0.585Kb
0.731Kb
0.878Kb
1.024Mb
1.17Mb
1.316Mb
1.463Mb
umbicola drowni dro
umbicola passerinae
umbicola passerinae
olumbicola altamimia
umbicola gymnopeli
Species
Columbicola drowni drowni
Columbicola passerinae 1
Columbicola passerinae 2
Columbicola altamimiae
Columbicola gymnopeliae

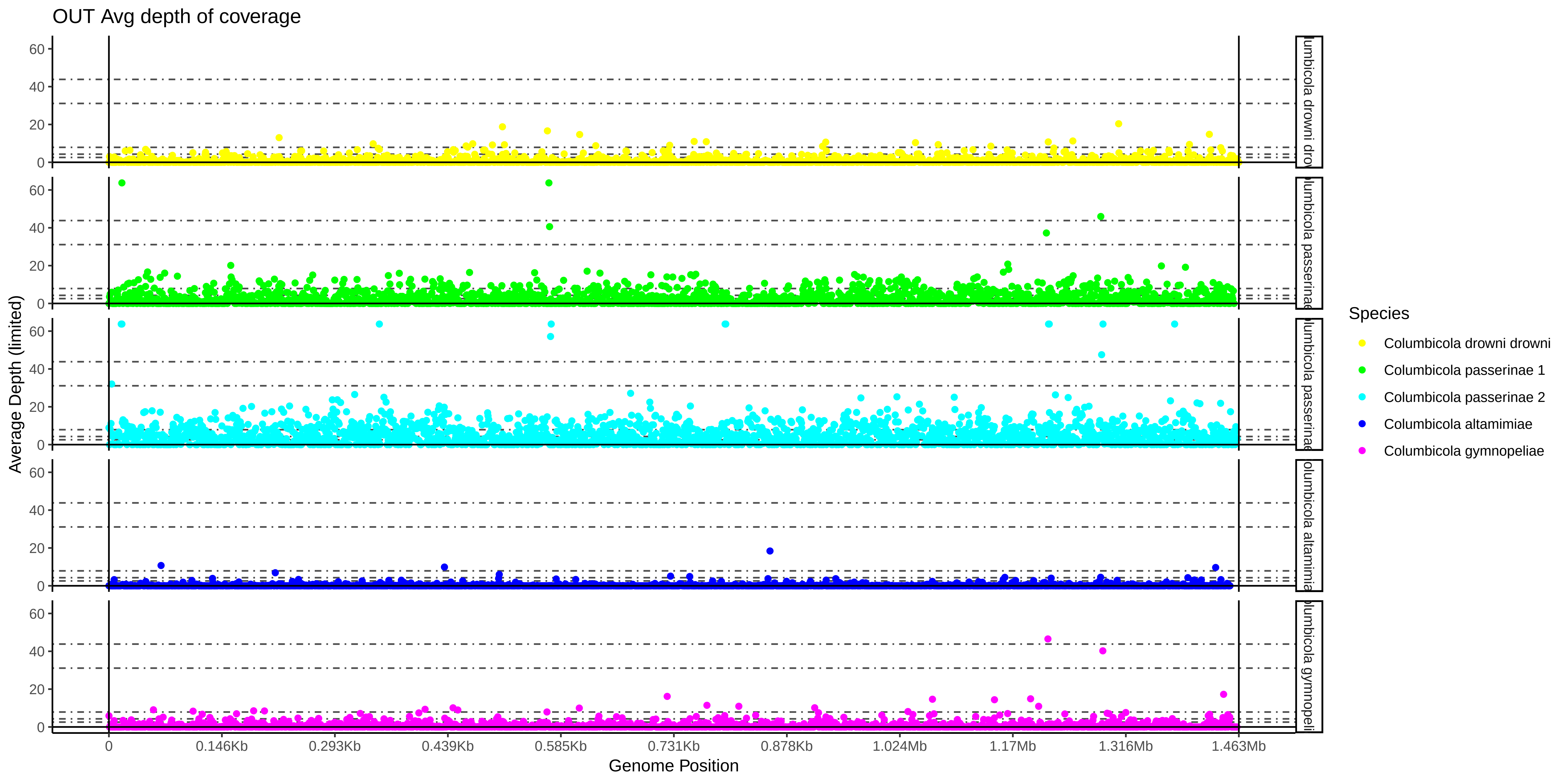

OUT Avg depth of coverage
Average Depth (limited)
Genome Position
0
20
40
60
0.146Kb
0.293Kb
0.439Kb
0.585Kb
0.731Kb
0.878Kb
1.024Mb
1.17Mb
1.316Mb
1.463Mb
umbicola drowni dro
umbicola passerinae
umbicola passerinae
olumbicola altamimia
umbicola gymnopeli
Species
Columbicola drowni drowni
Columbicola passerinae 1
Columbicola passerinae 2
Columbicola altamimiae
Columbicola gymnopeliae

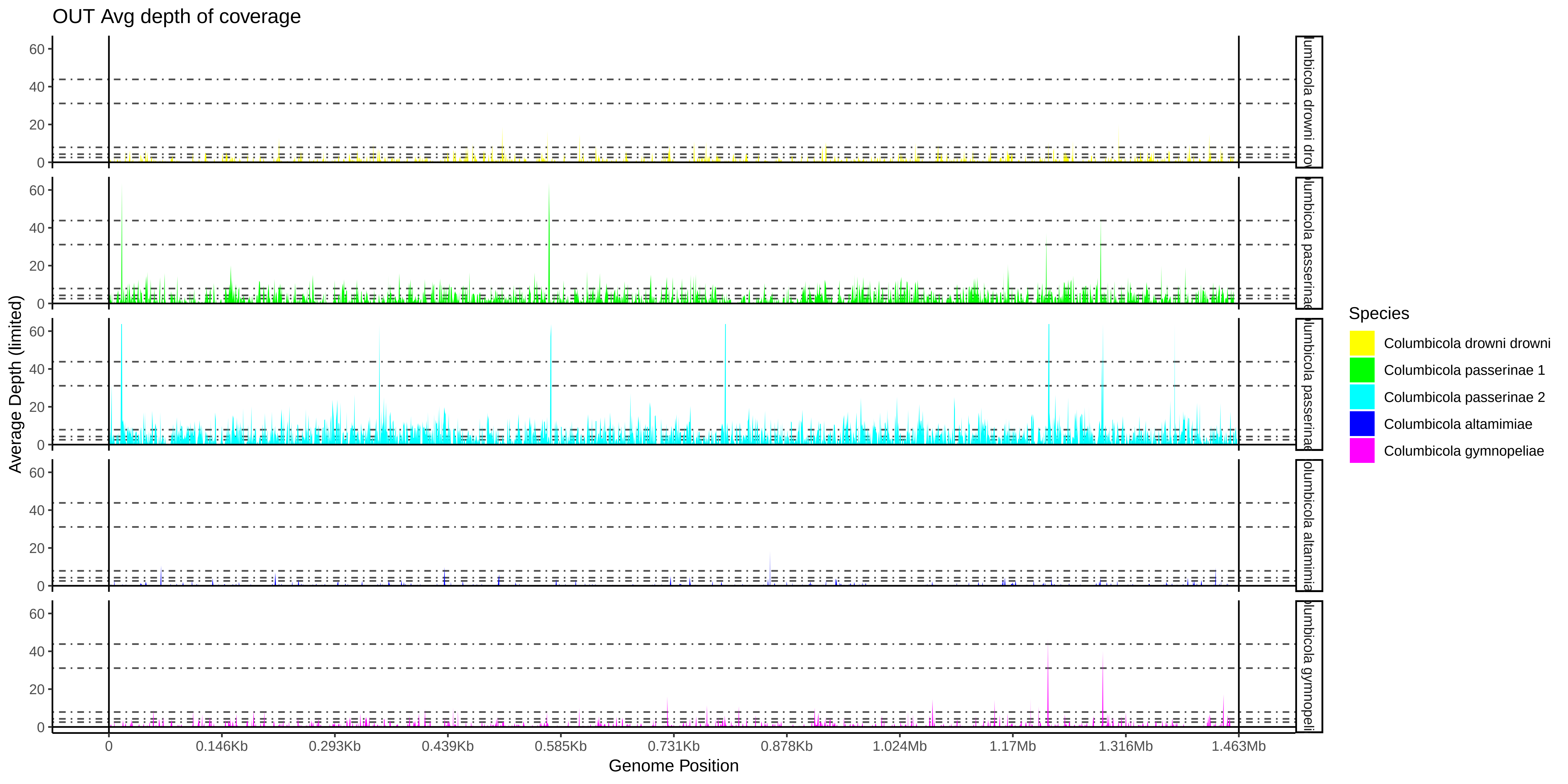

OUT Avg depth of coverage
Average Depth (limited)
Genome Position
0
0.146Kb
0.293Kb
0.439Kb
0.585Kb
0.731Kb
0.878Kb
1.024Mb
1.17Mb
1.316Mb
1.463Mb
0
20
40
60
umbicola drowni dro
lumbicola passerinae
lumbicola passerinae
olumbicola altamimia
umbicola gymnopeli
Species
Columbicola drowni drowni
Columbicola passerinae 1
Columbicola passerinae 2
Columbicola altamimiae
Columbicola gymnopeliae
