## Supplementary material for "Comparing rates of introgression in parasitic feather lice with differing dispersal capabilities": OUT_sppIDerDepthPlot-covBins_Cosp_Comin_11_2_2016_4.pdf (from another version of this preprint: version 7, 2020-03-31)

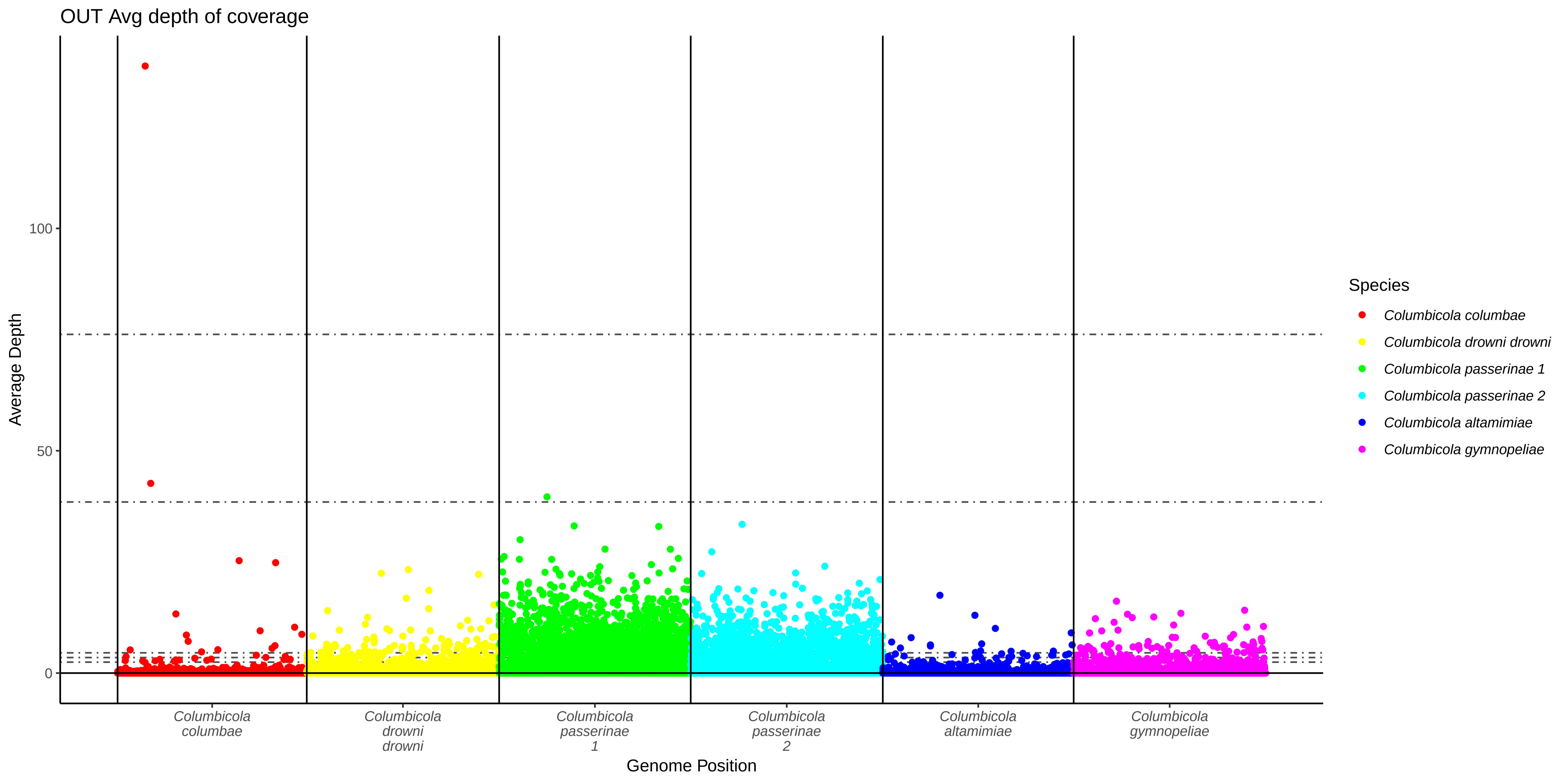
OUT Avg depth of coverage
Average Depth
100
50
0
Columbicola columbae
Columbicola drowni drowni
Columbicola passerinae 1
Columbicola passerinae 2
Columbicola altamimiae
Columbicola gymnopeliae
Genome Position
Species
Columbicola columbae
Columbicola drowni drowni
Columbicola passerinae 1
Columbicola passerinae 2
Columbicola altamimiae
Columbicola gymnopeliae

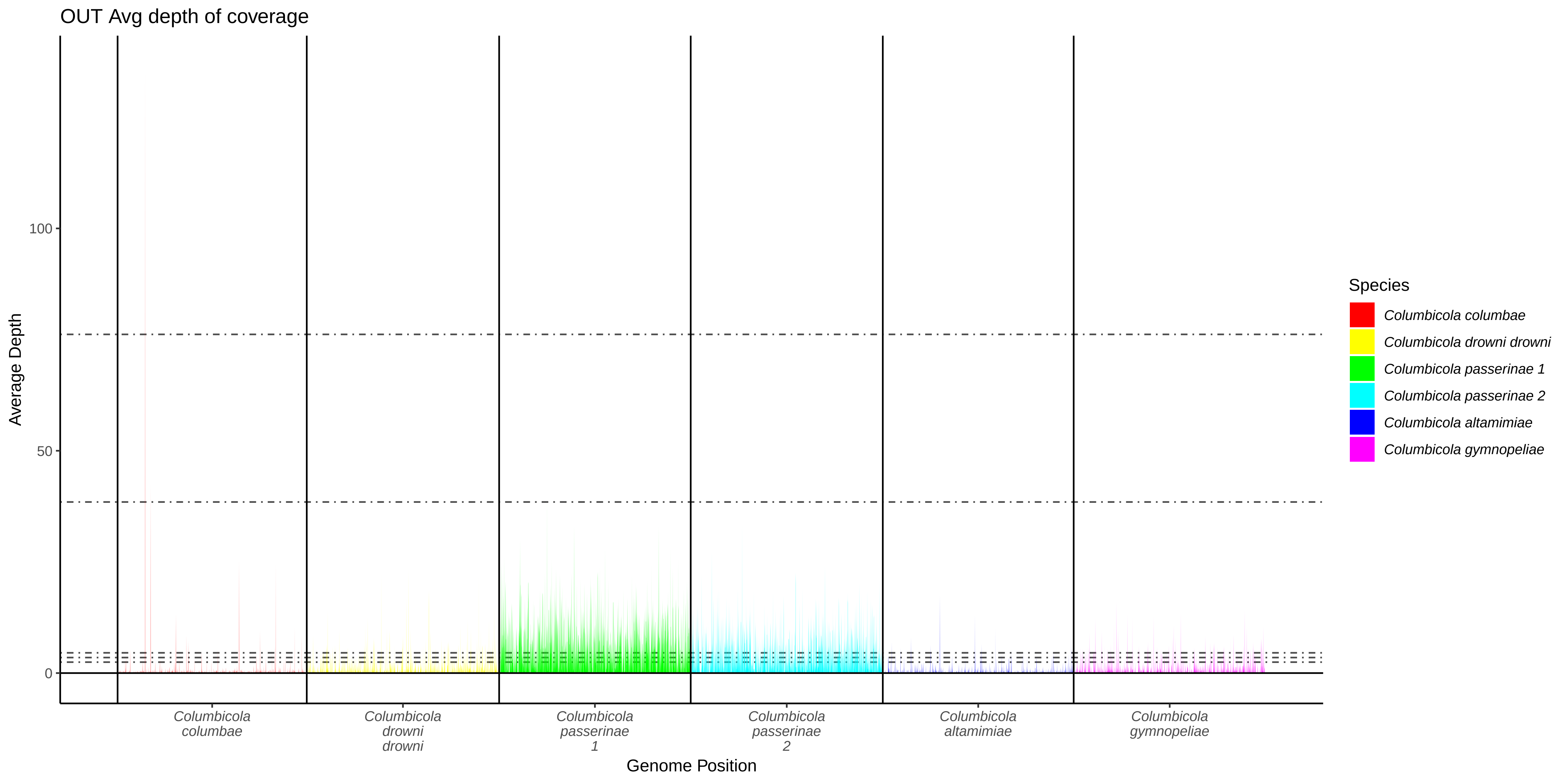
OUT Avg depth of coverage
Average Depth
100
50
0
Columbicola columbae
Columbicola drowni drowni
Columbicola passerinae 1
Columbicola passerinae 2
Columbicola altamimiae
Columbicola gymnopeliae
Genome Position
Species
Columbicola columbae
Columbicola drowni drowni
Columbicola passerinae 1
Columbicola passerinae 2
Columbicola altamimiae
Columbicola gymnopeliae

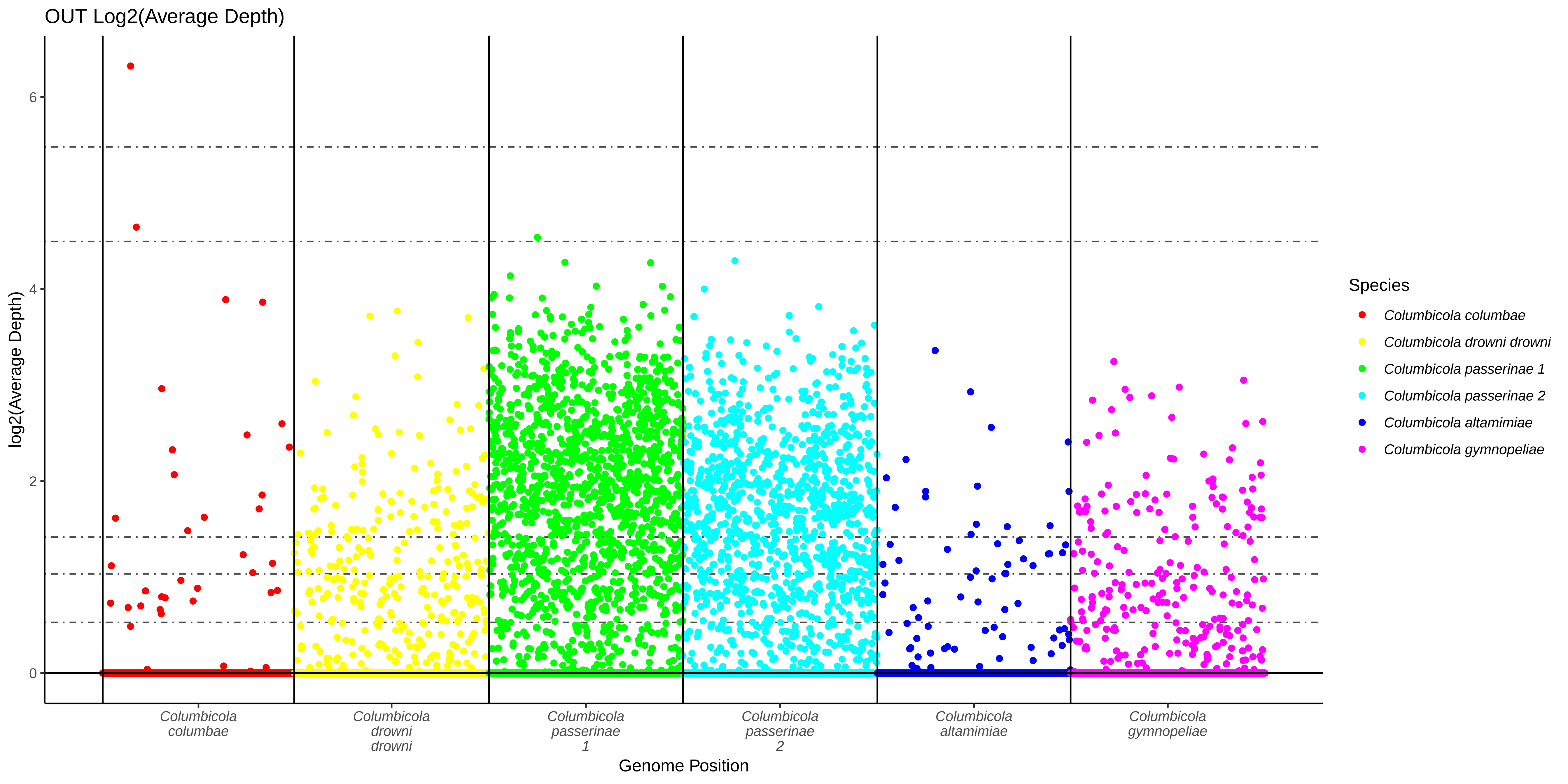

OUT Log2(Average Depth)
log2(Average Depth)
6
4
2
0
Columbicola columbae
Columbicola drowni drowni
Columbicola passerinae 1
Columbicola passerinae 2
Columbicola altamimiae
Columbicola gymnopeliae
Genome Position
Species
Columbicola columbae
Columbicola drowni drowni
Columbicola passerinae 1
Columbicola passerinae 2
Columbicola altamimiae
Columbicola gymnopeliae

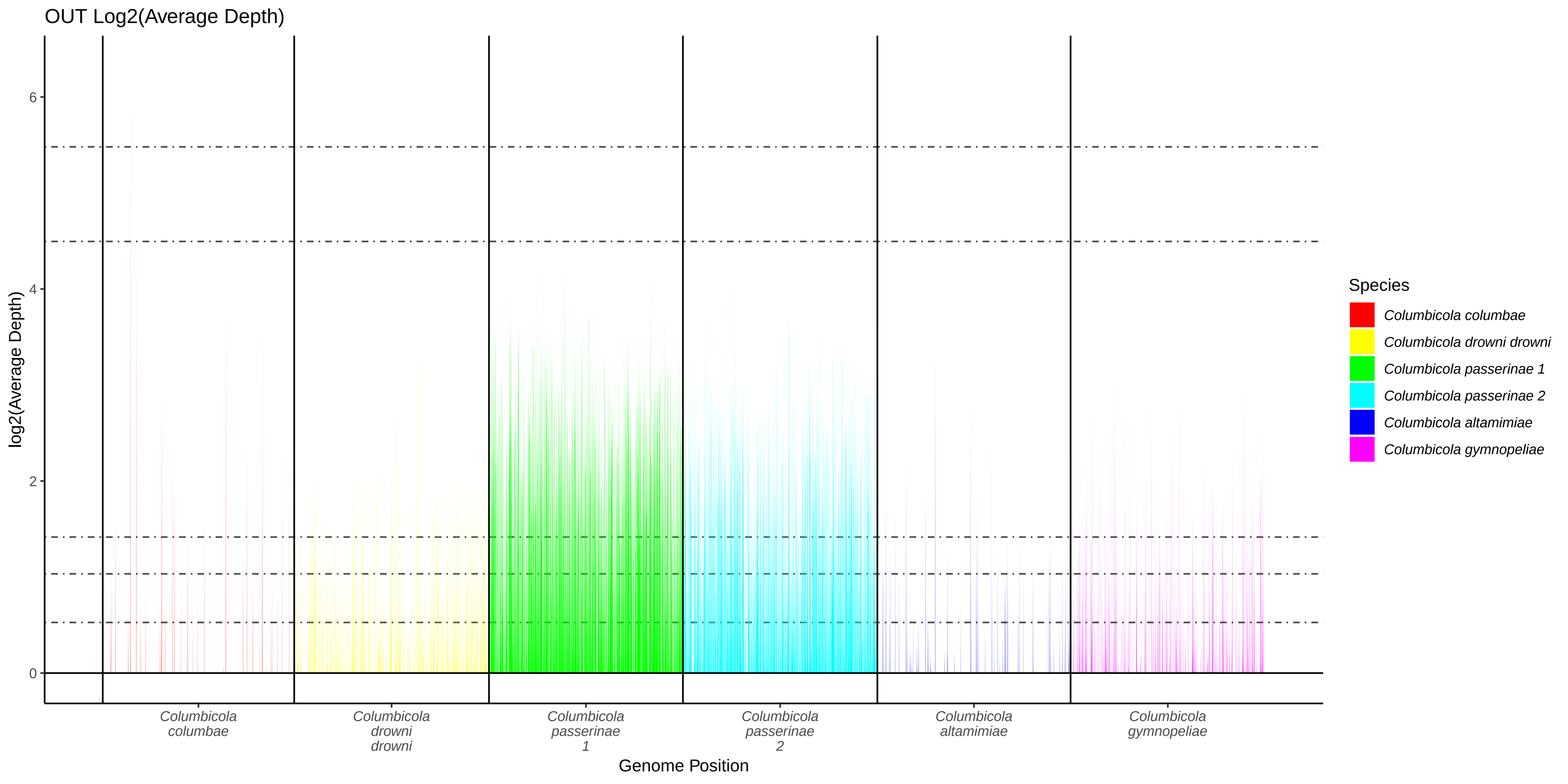

OUT Log2(Average Depth)
log2(Average Depth)
0
2
4
6
Columbicola columbae
Columbicola drowni drowni
Columbicola passerinae 1
Columbicola passerinae 2
Columbicola altamimiae
Columbicola gymnopeliae
Genome Position
Species
Columbicola columbae
Columbicola drowni drowni
Columbicola passerinae 1
Columbicola passerinae 2
Columbicola altamimiae
Columbicola gymnopeliae

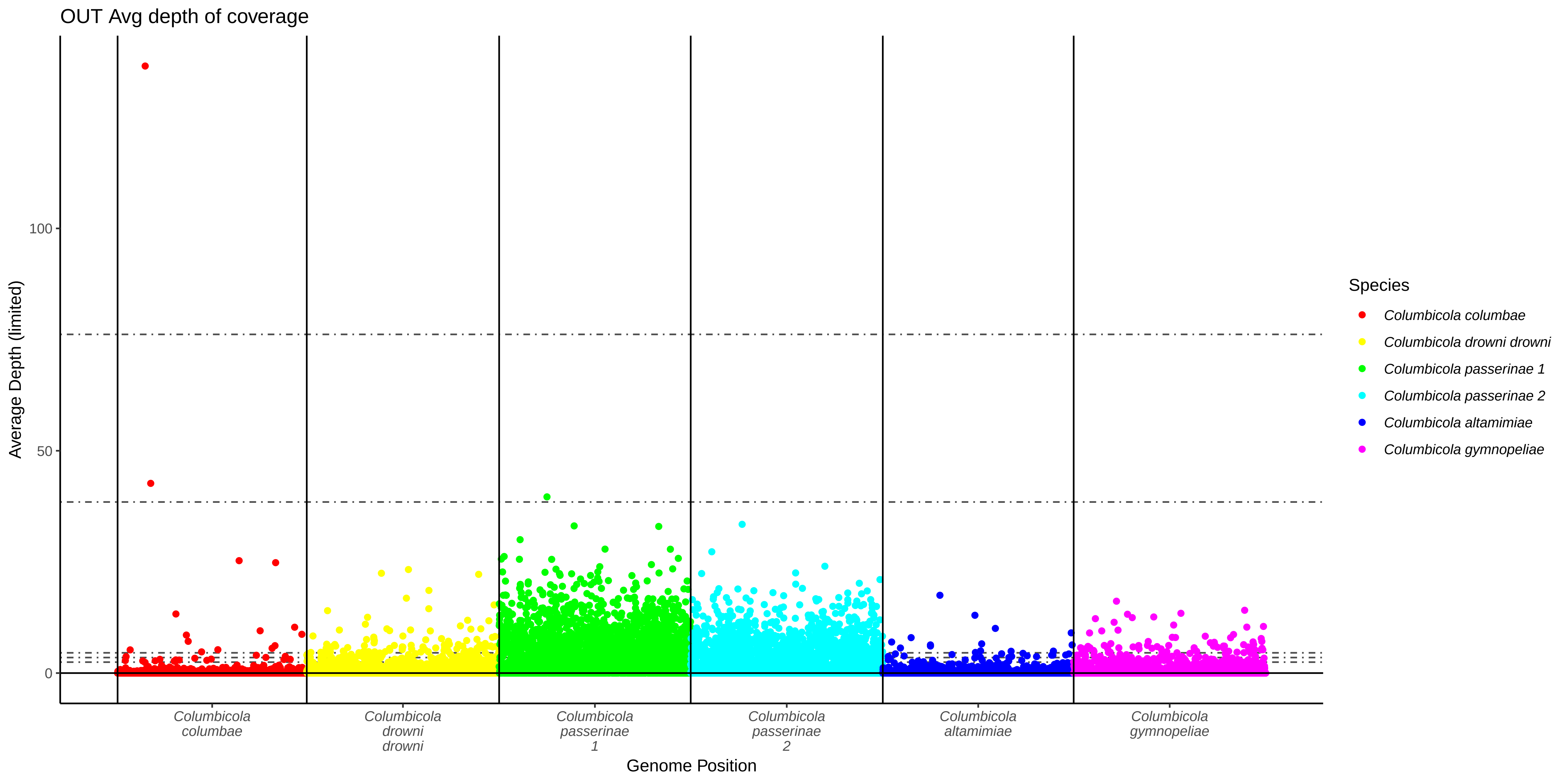
OUT Avg depth of coverage
Average Depth (limited)
100
50
0
Columbicola columbae
Columbicola drowni drowni
Columbicola passerinae 1
Columbicola passerinae 2
Columbicola altamimiae
Columbicola gymnopeliae
Genome Position
Species
Columbicola columbae
Columbicola drowni drowni
Columbicola passerinae 1
Columbicola passerinae 2
Columbicola altamimiae
Columbicola gymnopeliae

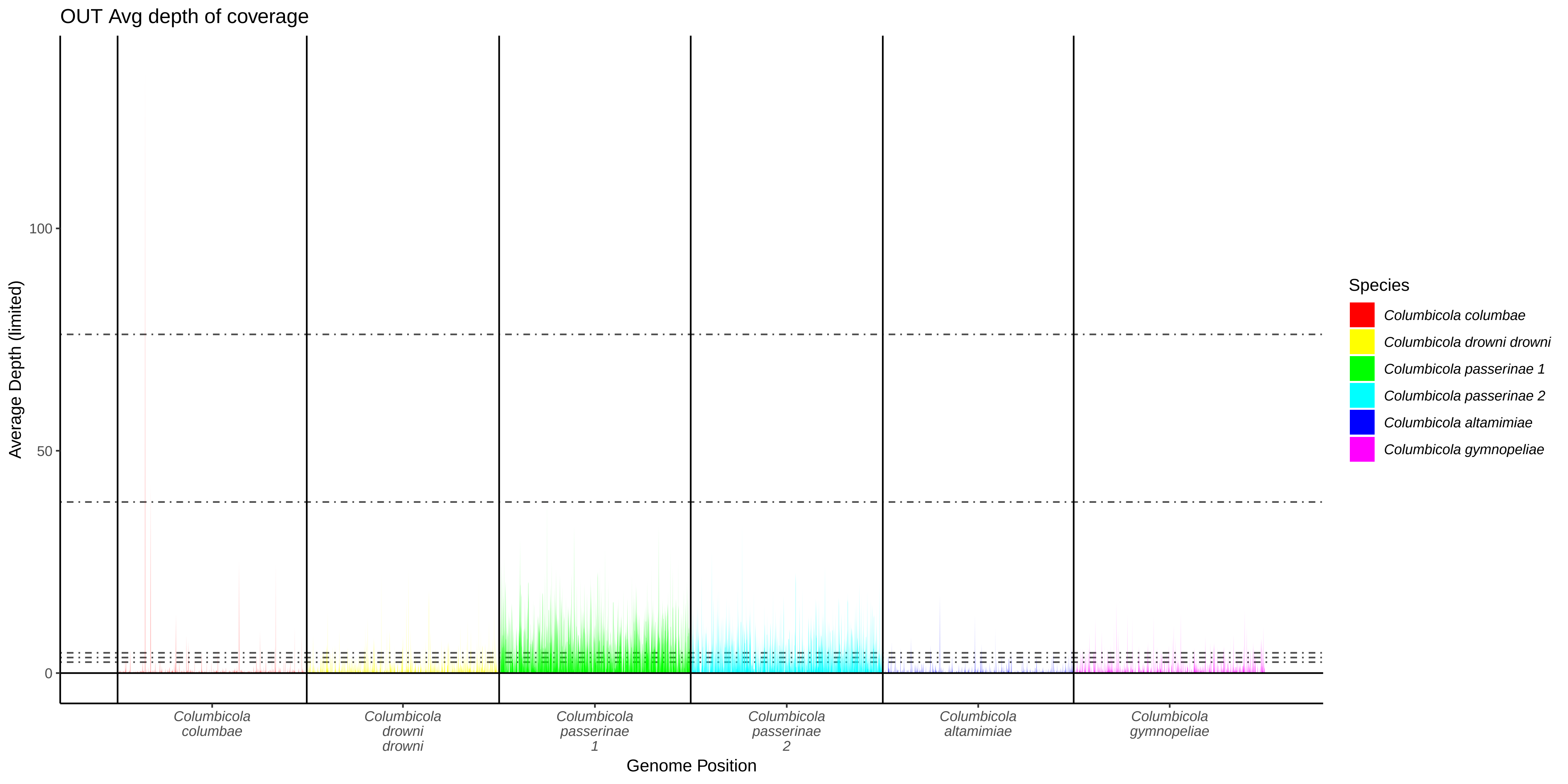
OUT Avg depth of coverage
Average Depth (limited)
100
50
0
Columbicola columbae
Columbicola drowni drowni
Columbicola passerinae 1
Columbicola passerinae 2
Columbicola altamimiae
Columbicola gymnopeliae
Genome Position
Species
Columbicola columbae
Columbicola drowni drowni
Columbicola passerinae 1
Columbicola passerinae 2
Columbicola altamimiae
Columbicola gymnopeliae

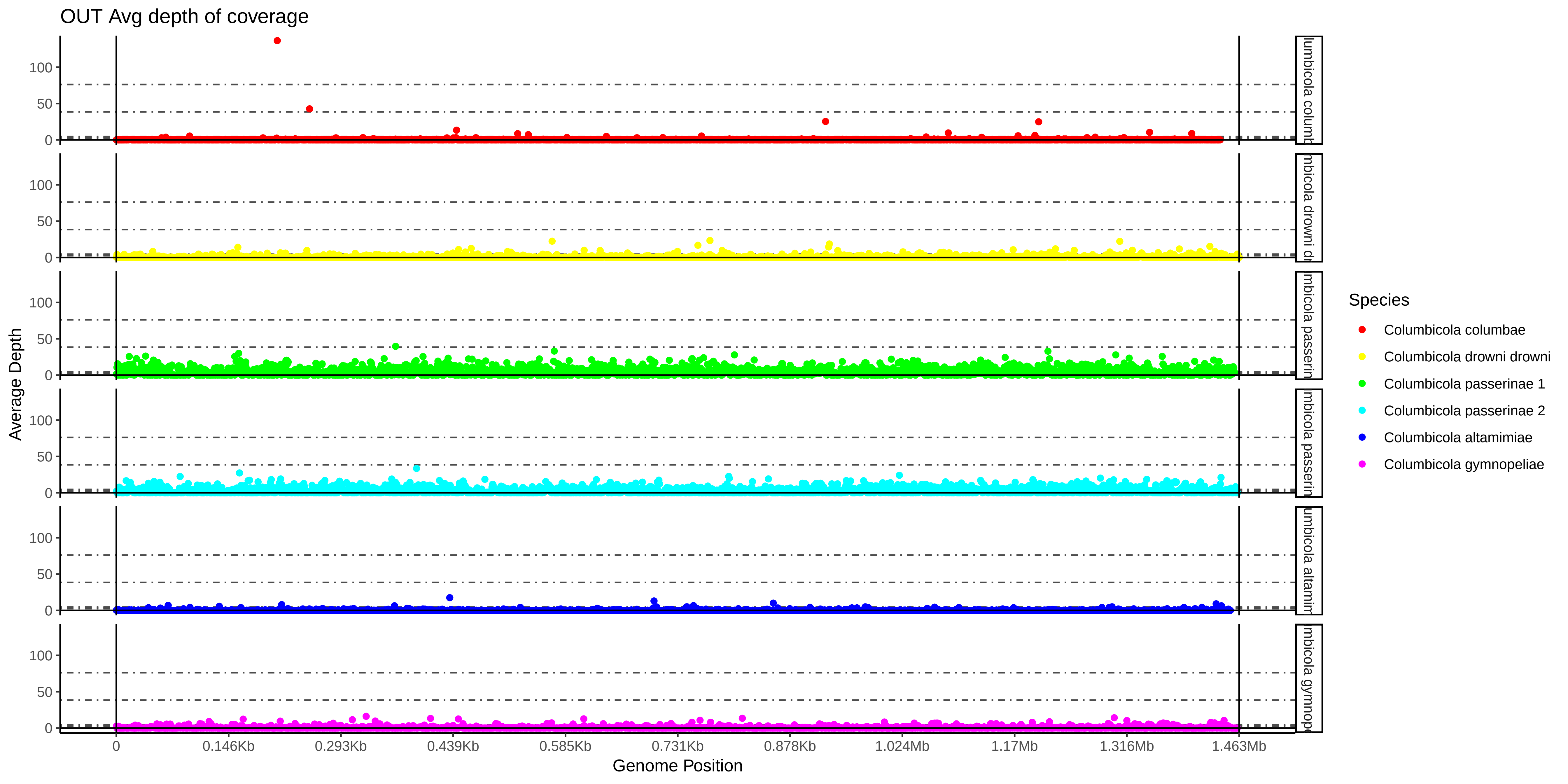

OUT Avg depth of coverage
Average Depth
Genome Position
0
50
100
0
0.146Kb
0.293Kb
0.439Kb
0.585Kb
0.731Kb
0.878Kb
1.024Mb
1.17Mb
1.316Mb
1.463Mb
lumbicola columb
mbicola drowni dr
mbicola passerin
mbicola passerin
umbicola altamim
mbicola gymnope
Species
Columbicola columbae
Columbicola drowni drowni
Columbicola passerinae 1
Columbicola passerinae 2
Columbicola altamimiae
Columbicola gymnopeliae

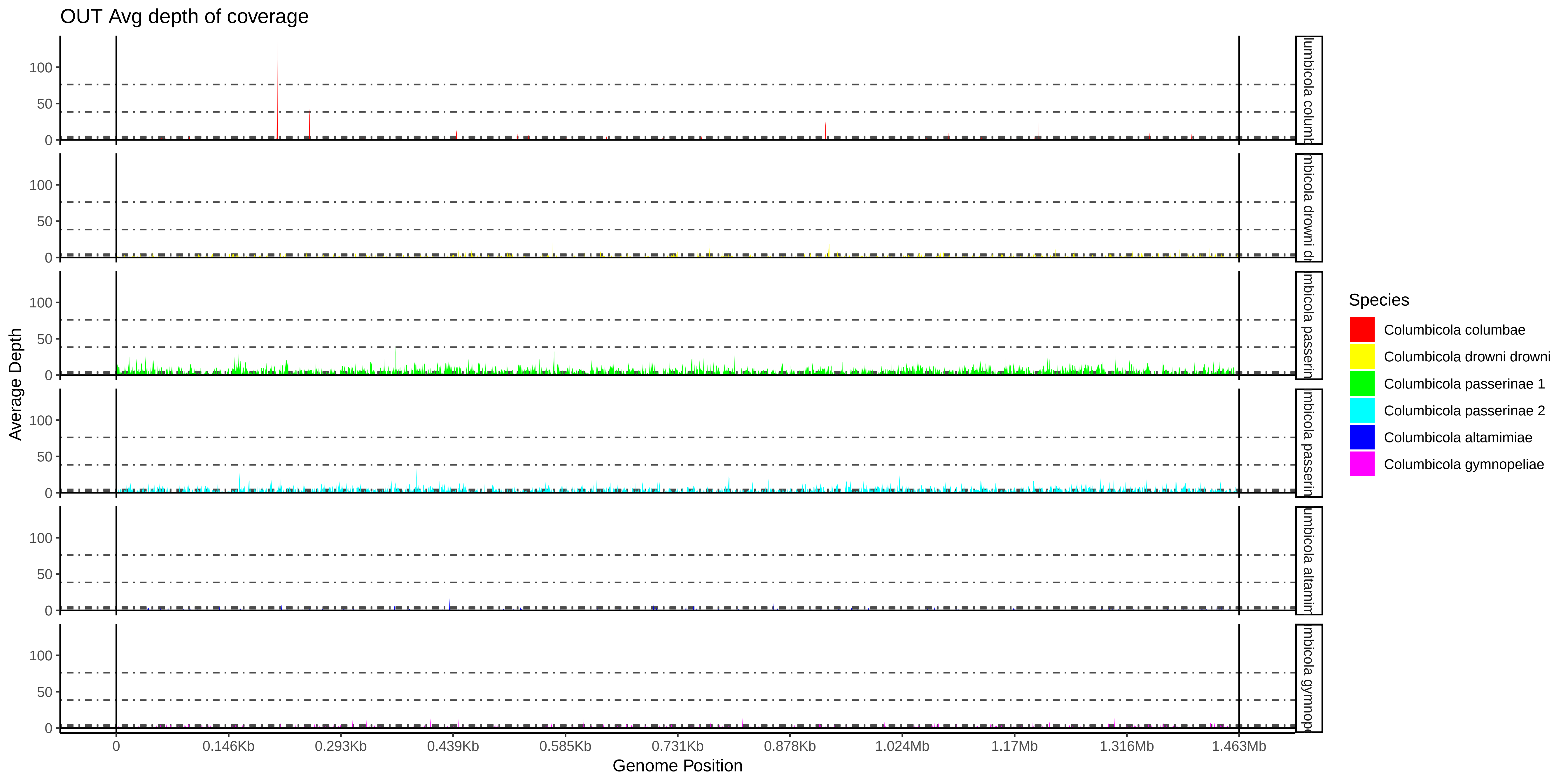

OUT Avg depth of coverage
Average Depth
Genome Position
0
50
100
0
0.146Kb
0.293Kb
0.439Kb
0.585Kb
0.731Kb
0.878Kb
1.024Mb
1.17Mb
1.316Mb
1.463Mb
umbicola columb
mbicola drowni dr
mbicola passerin
mbicola passerin
umbicola altamim
mbicola gymnop
Species
Columbicola columbae
Columbicola drowni drowni
Columbicola passerinae 1
Columbicola passerinae 2
Columbicola altamimiae
Columbicola gymnopeliae

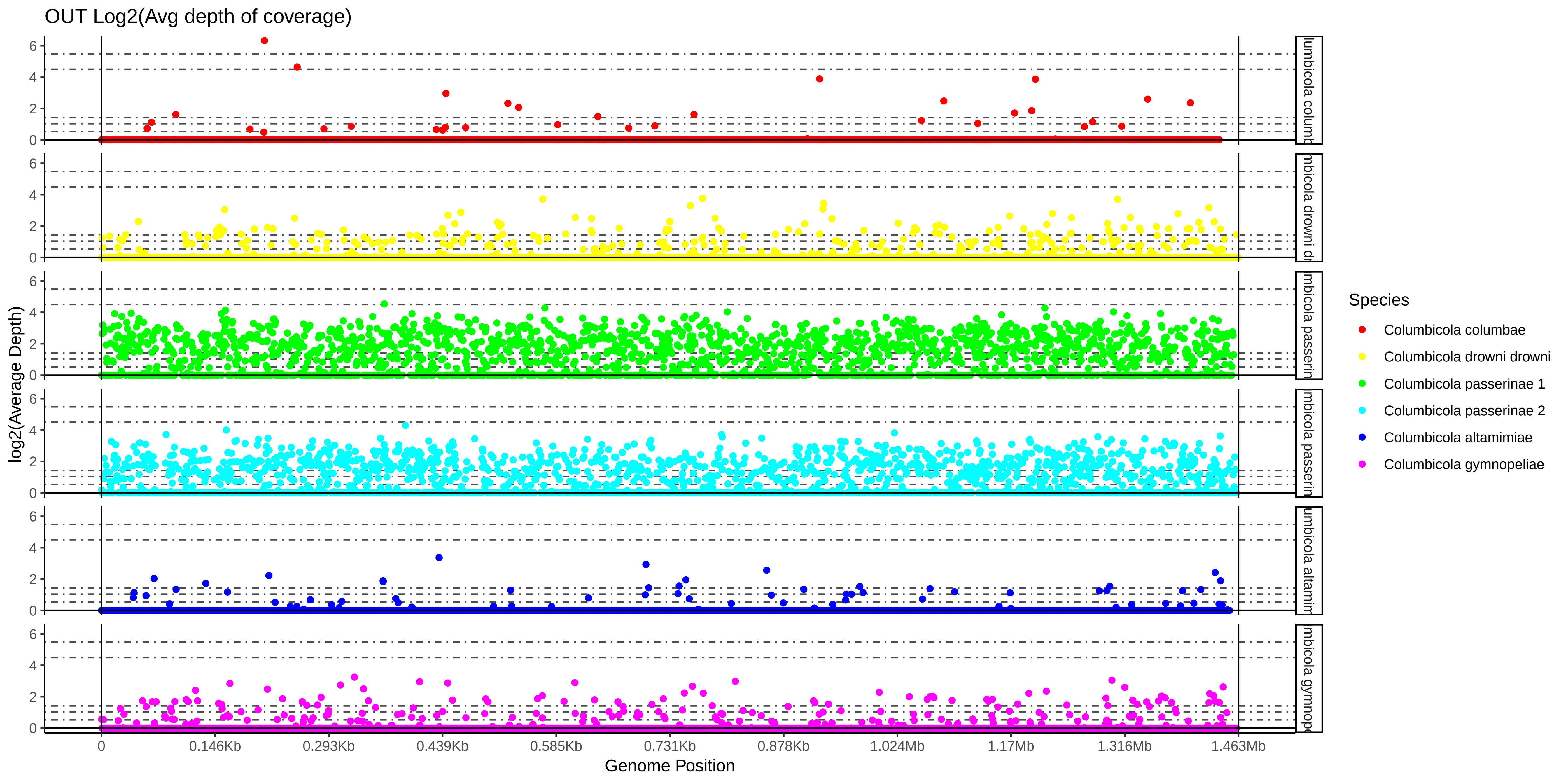

OUT Log2(Avg depth of coverage)
log2(Average Depth)
Genome Position
0
0.146Kb
0.293Kb
0.439Kb
0.585Kb
0.731Kb
0.878Kb
1.024Mb
1.17Mb
1.316Mb
1.463Mb
Species
Columbicola columbae
Columbicola drowni drowni
Columbicola passerinae 1
Columbicola passerinae 2
Columbicola altamimiae
Columbicola gymnopeliae

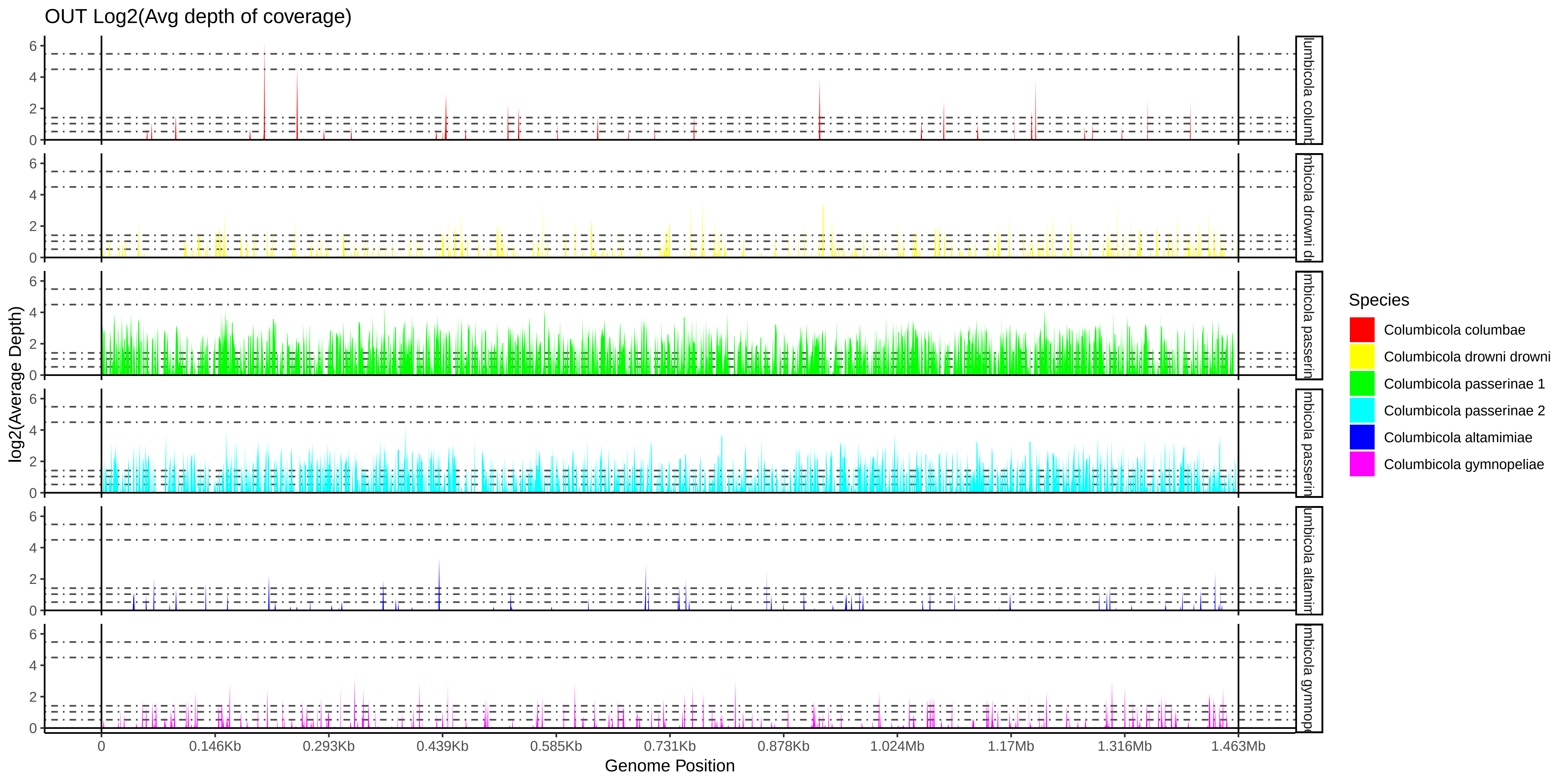
OUT Log2(Avg depth of coverage)
log2(Average Depth)
Genome Position
0
0.146Kb
0.293Kb
0.439Kb
0.585Kb
0.731Kb
0.878Kb
1.024Mb
1.17Mb
1.316Mb
1.463Mb
Species
Columbicola columbae
Columbicola drowni drowni
Columbicola passerinae 1
Columbicola passerinae 2
Columbicola altamimiae
Columbicola gymnopeliae

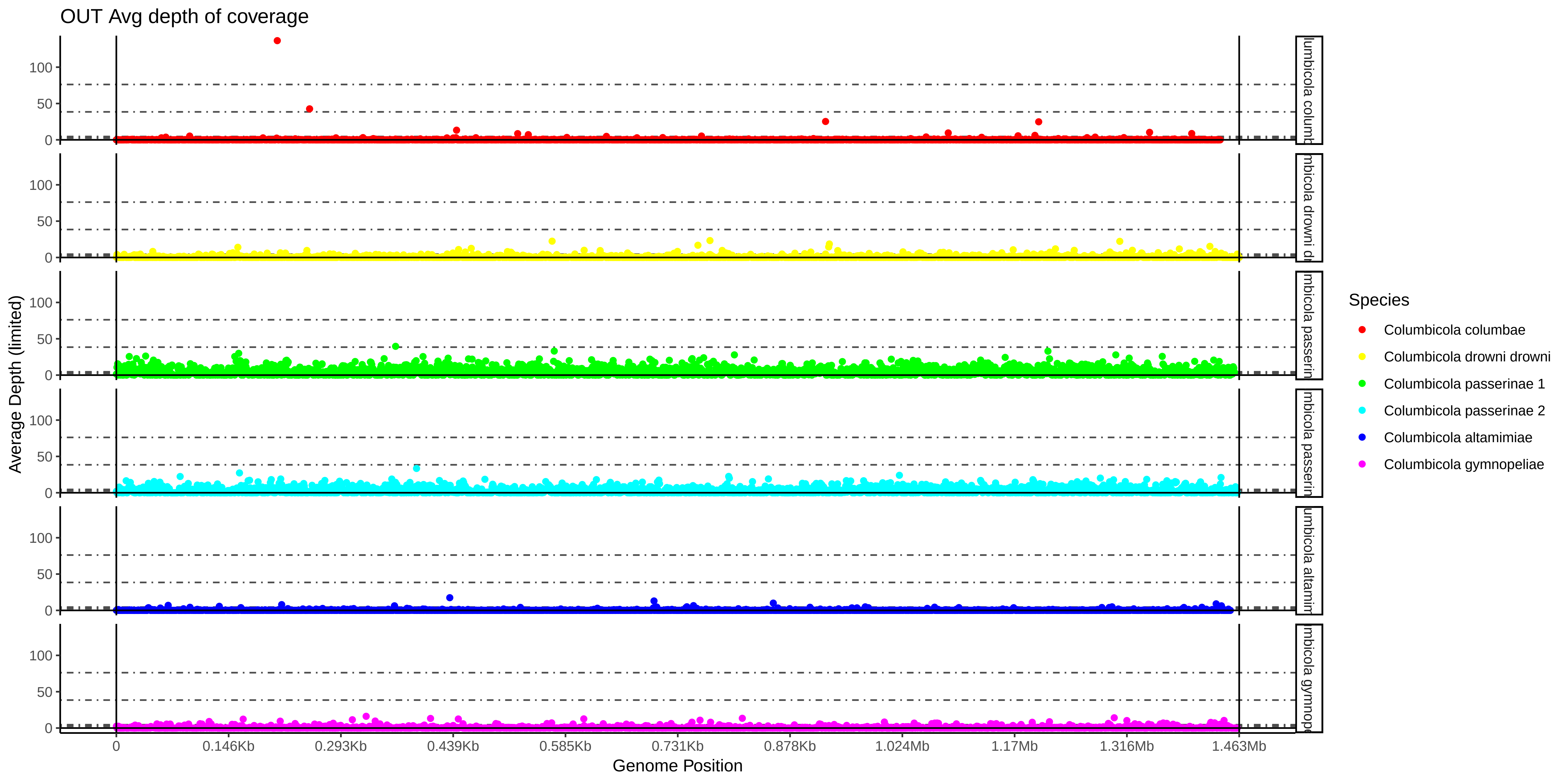

OUT Avg depth of coverage
Average Depth (limited)
Genome Position
0
50
100
0
0.146Kb
0.293Kb
0.439Kb
0.585Kb
0.731Kb
0.878Kb
1.024Mb
1.17Mb
1.316Mb
1.463Mb
Species
Columbicola columbae
Columbicola drowni drowni
Columbicola passerinae 1
Columbicola passerinae 2
Columbicola altamimiae
Columbicola gymnopeliae

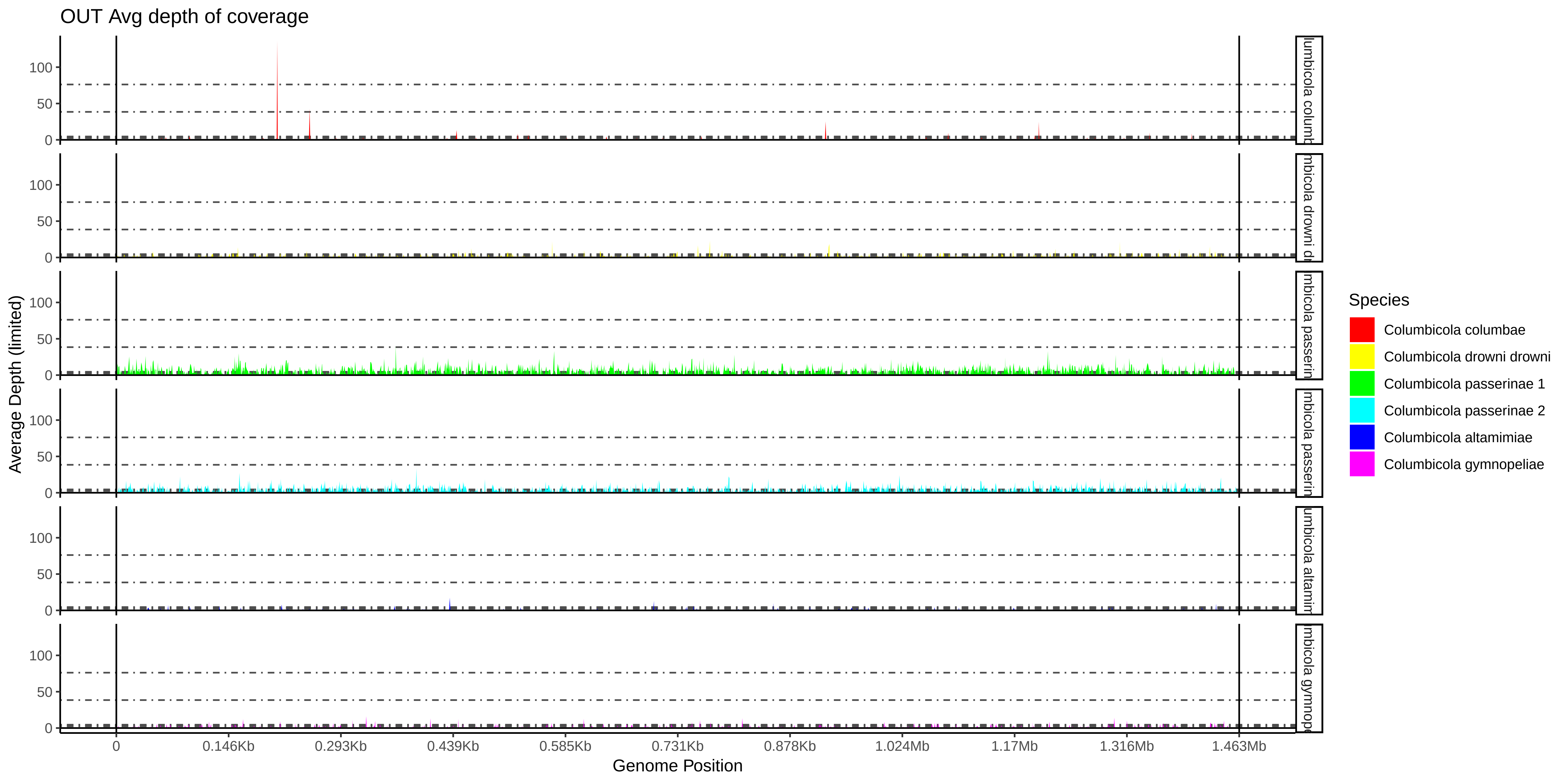

OUT Avg depth of coverage
Average Depth (limited)
Genome Position
0
0.146Kb
0.293Kb
0.439Kb
0.585Kb
0.731Kb
0.878Kb
1.024Mb
1.17Mb
1.316Mb
1.463Mb
Species
Columbicola columbae
Columbicola drowni drowni
Columbicola passerinae 1
Columbicola passerinae 2
Columbicola altamimiae
Columbicola gymnopeliae
