## Supplementary material for "Comparing rates of introgression in parasitic feather lice with differing dispersal capabilities": OUT_sppIDerDepthPlot-covBins_Cosp_Copic_11_2_2016_1.pdf (from another version of this preprint: version 7, 2020-03-31)

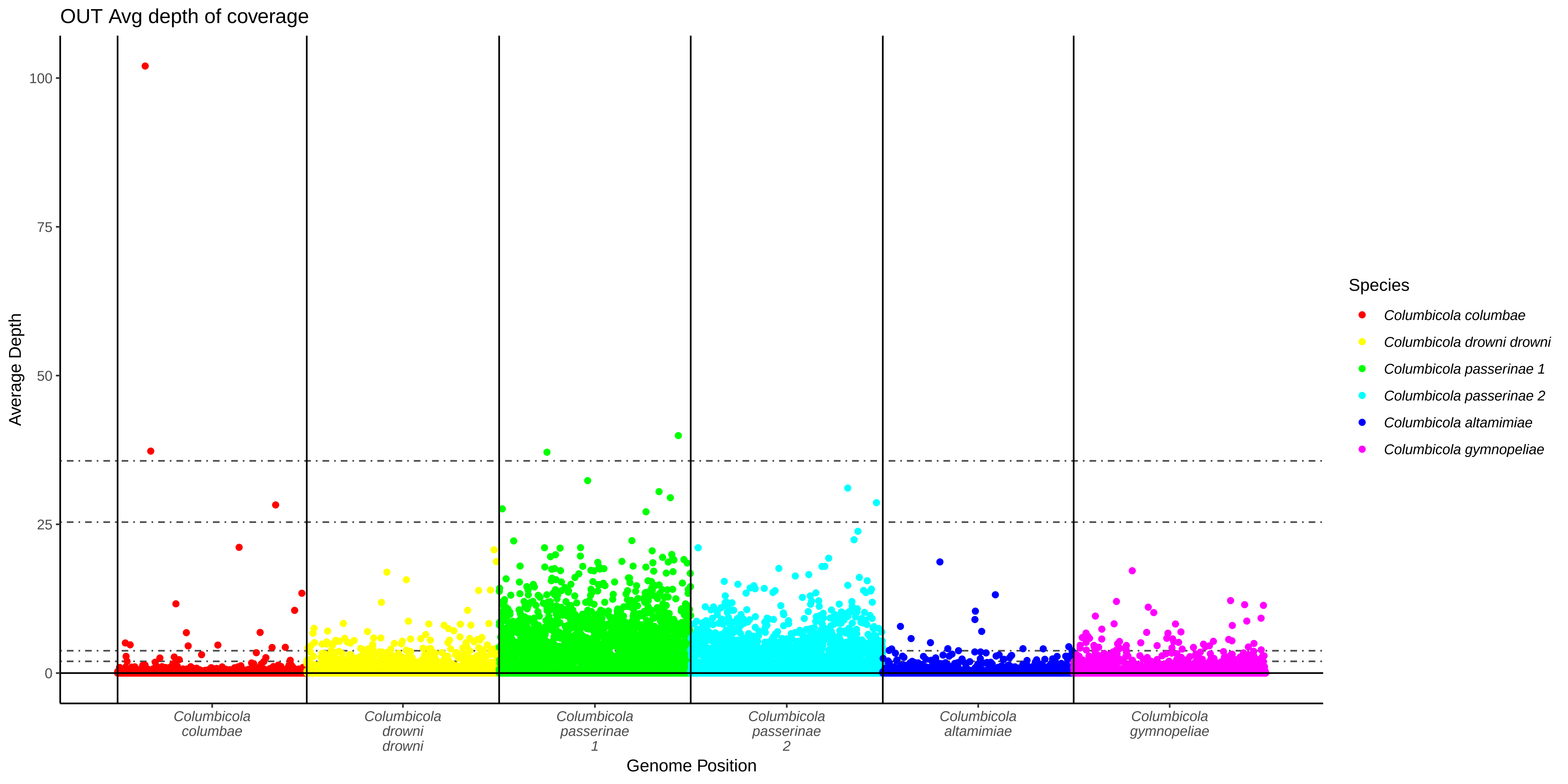

OUT Avg depth of coverage
Average Depth
100
75
50
25
0
Columbicola columbae
Columbicola drowni drowni
Columbicola passerinae 1
Columbicola passerinae 2
Columbicola altamimiae
Columbicola gymnopeliae
Genome Position
Species
Columbicola columbae
Columbicola drowni drowni
Columbicola passerinae 1
Columbicola passerinae 2
Columbicola altamimiae
Columbicola gymnopeliae

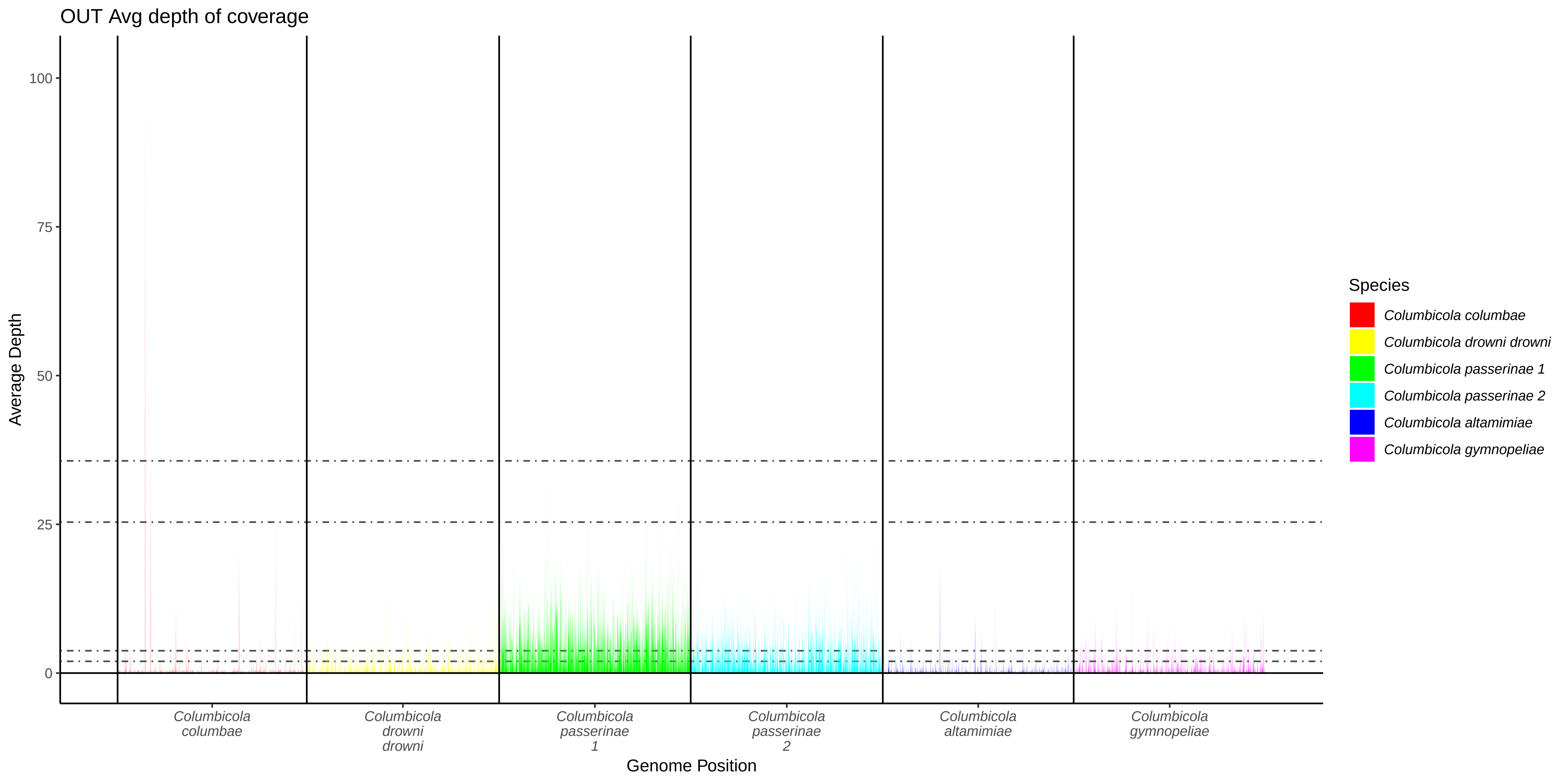

OUT Avg depth of coverage
Average Depth
100
75
50
25
0
Genome Position
Columbicola columbae
Columbicola drowni drowni
Columbicola passerinae 1
Columbicola passerinae 2
Columbicola altamimiae
Columbicola gymnopeliae
Species
Columbicola columbae
Columbicola drowni drowni
Columbicola passerinae 1
Columbicola passerinae 2
Columbicola altamimiae
Columbicola gymnopeliae

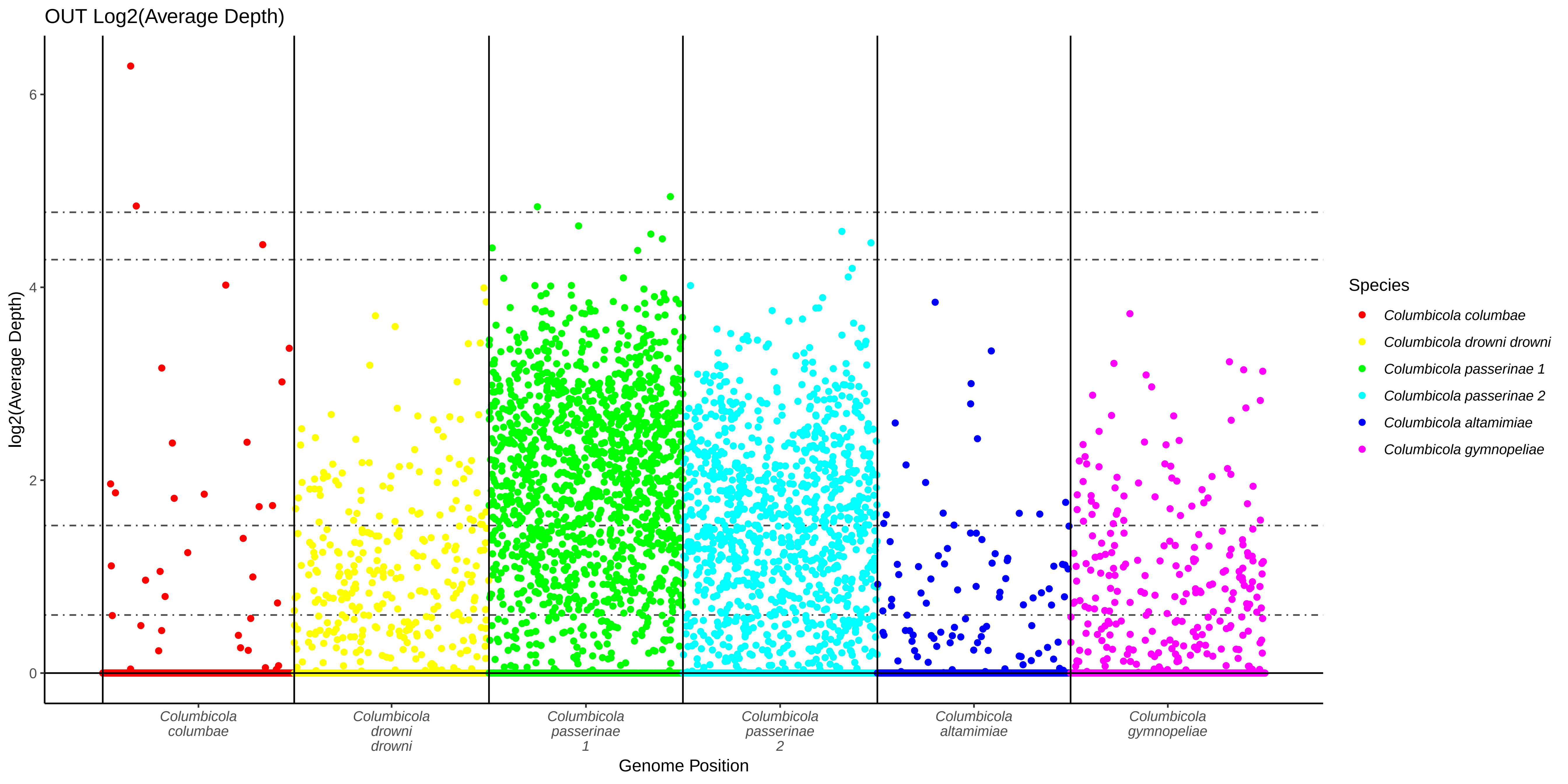

OUT Log2(Average Depth)
log2(Average Depth)
6
4
2
0
Columbicola columbae
Columbicola drowni drowni
Columbicola passerinae 1
Columbicola passerinae 2
Columbicola altamimiae
Columbicola gymnopeliae
Genome Position
Species
Columbicola columbae
Columbicola drowni drowni
Columbicola passerinae 1
Columbicola passerinae 2
Columbicola altamimiae
Columbicola gymnopeliae

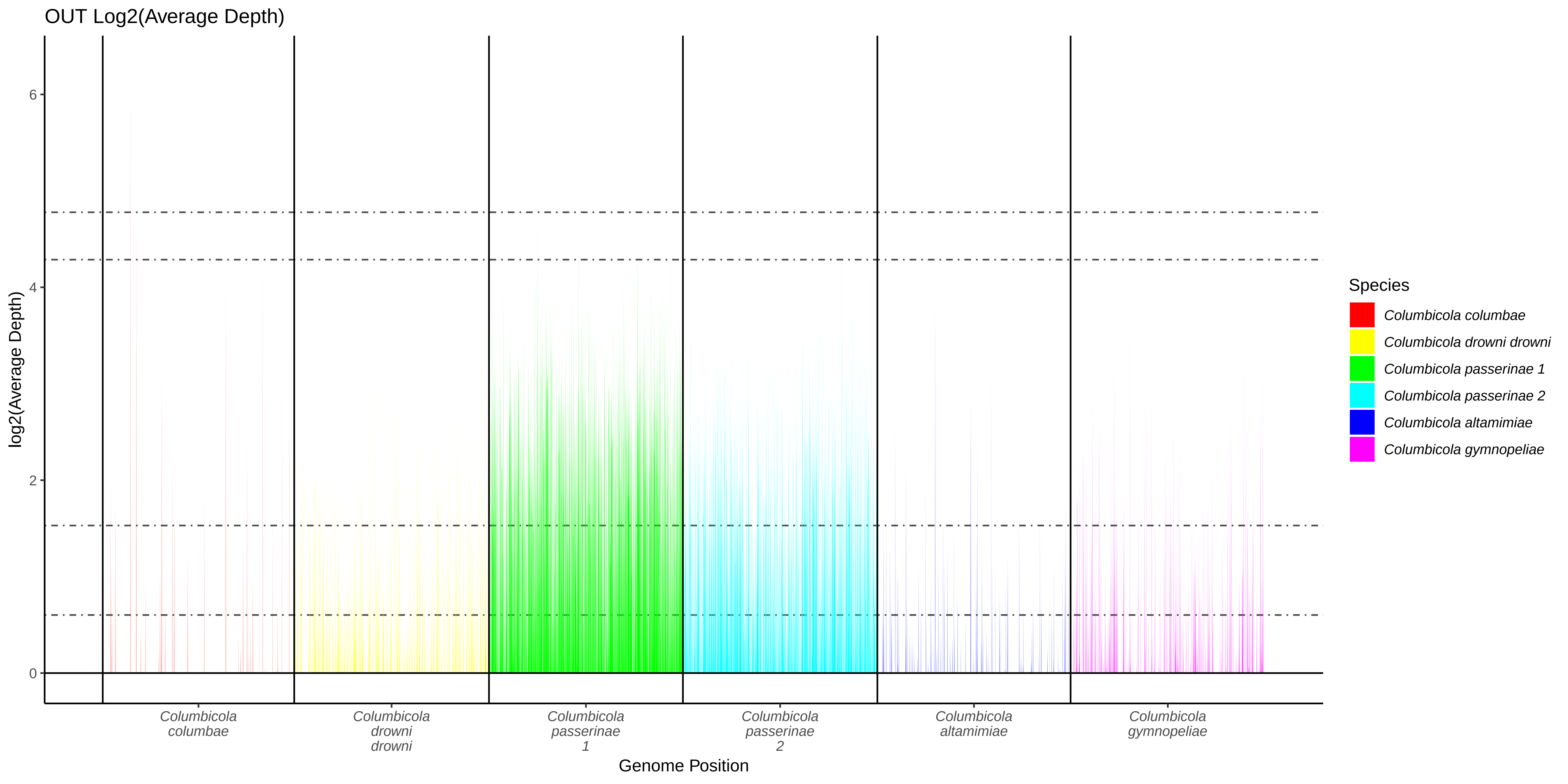
OUT Log2(Average Depth)
log2(Average Depth)
0
2
4
6
Species
Columbicola columbae
Columbicola drowni drowni
Columbicola passerinae 1
Columbicola passerinae 2
Columbicola altamimiae
Columbicola gymnopeliae
Columbicola columbae
Columbicola drowni drowni
Columbicola passerinae 1
Columbicola passerinae 2
Columbicola altamimiae
Columbicola gymnopeliae
Genome Position

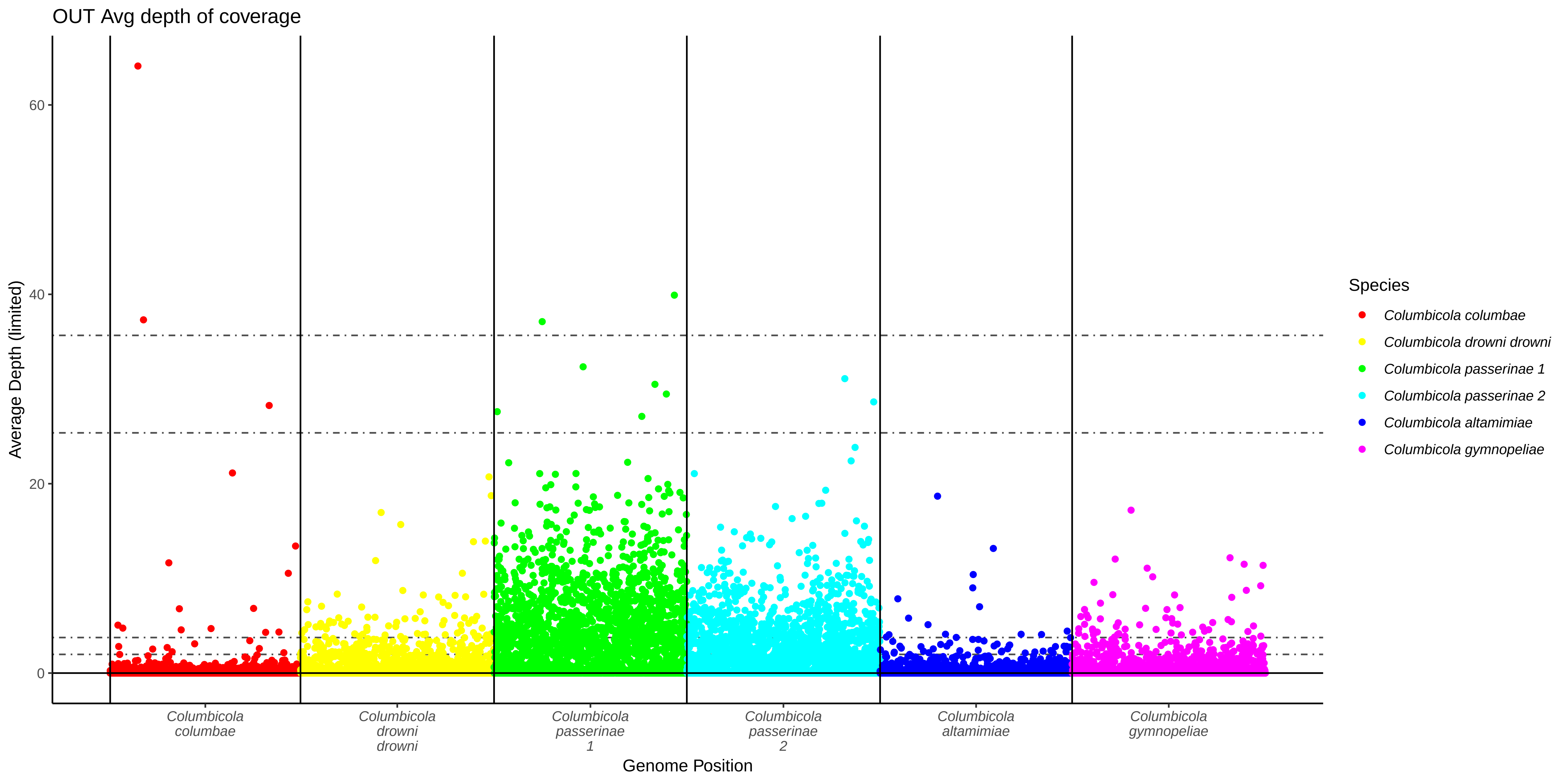
OUT Avg depth of coverage
Average Depth (limited)
60
40
20
0
Columbicola columbae
Columbicola drowni drowni
Columbicola passerinae 1
Columbicola passerinae 2
Columbicola altamimiae
Columbicola gymnopeliae
Genome Position
Species
Columbicola columbae
Columbicola drowni drowni
Columbicola passerinae 1
Columbicola passerinae 2
Columbicola altamimiae
Columbicola gymnopeliae

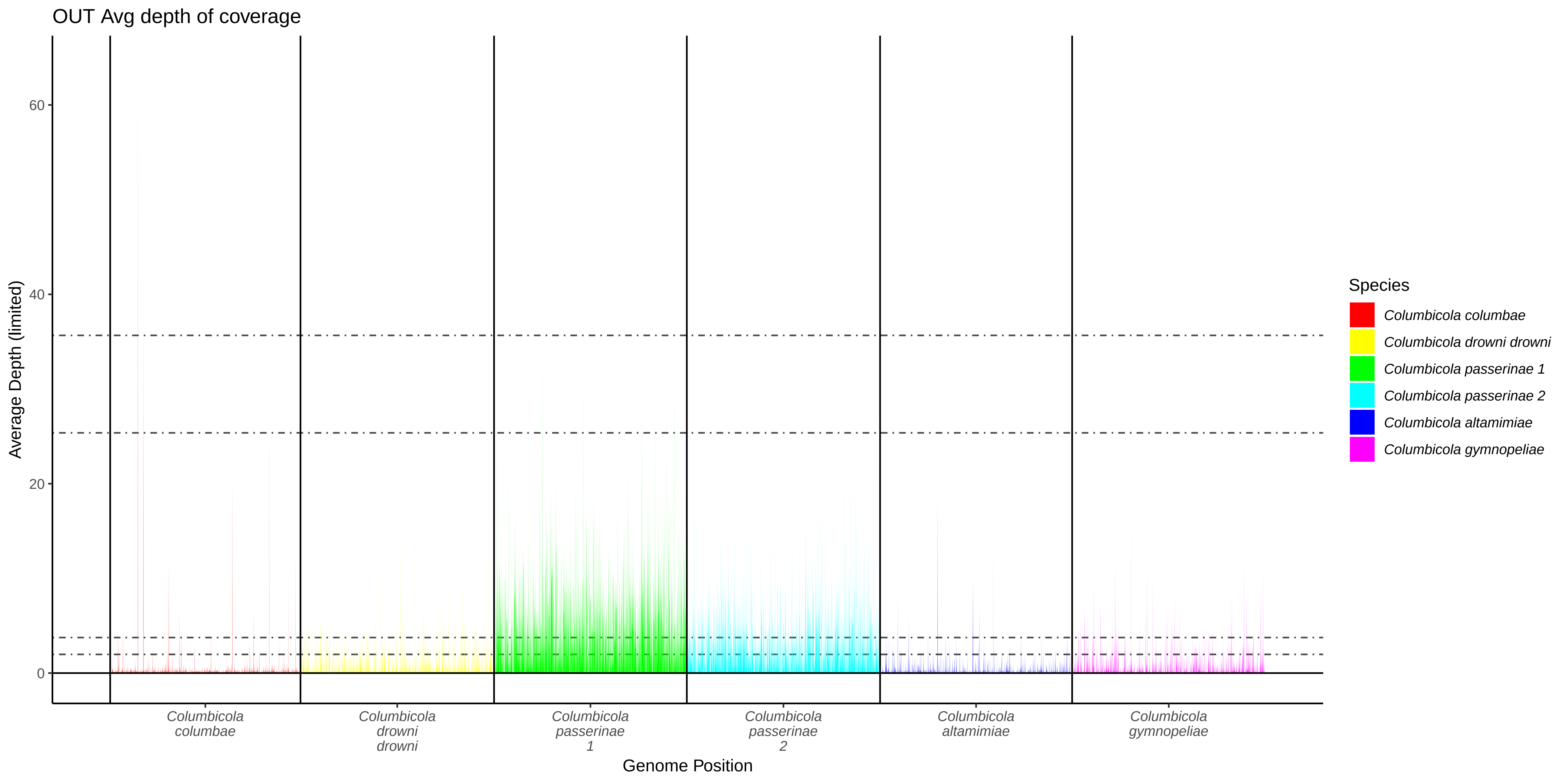

OUT Avg depth of coverage
Average Depth (limited)
0
20
40
60
Columbicola columbae
Columbicola drowni drowni
Columbicola passerinae 1
Columbicola passerinae 2
Columbicola altamimiae
Columbicola gymnopeliae
Genome Position
Species
Columbicola columbae
Columbicola drowni drowni
Columbicola passerinae 1
Columbicola passerinae 2
Columbicola altamimiae
Columbicola gymnopeliae

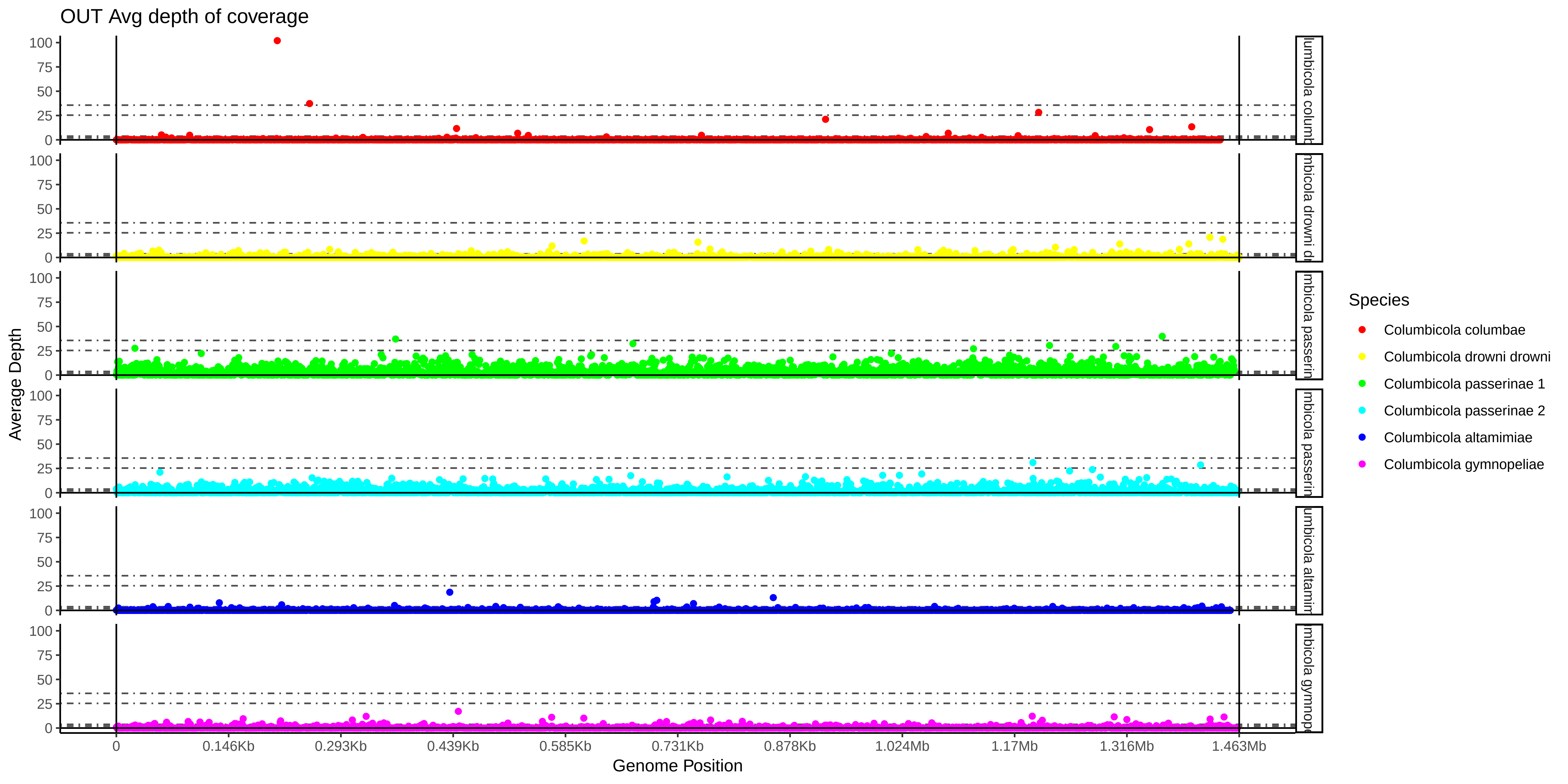

OUT Avg depth of coverage
Average Depth
Genome Position
0
0.146Kb
0.293Kb
0.439Kb
0.585Kb
0.731Kb
0.878Kb
1.024Mb
1.17Mb
1.316Mb
1.463Mb
100
75
50
25
0
umbicola columb
mbicola drowni dr
mbicola passerin
mbicola passerin
umbicola altamim
mbicola gymnope
Species
Columbicola columbae
Columbicola drowni drowni
Columbicola passerinae 1
Columbicola passerinae 2
Columbicola altamimiae
Columbicola gymnopeliae

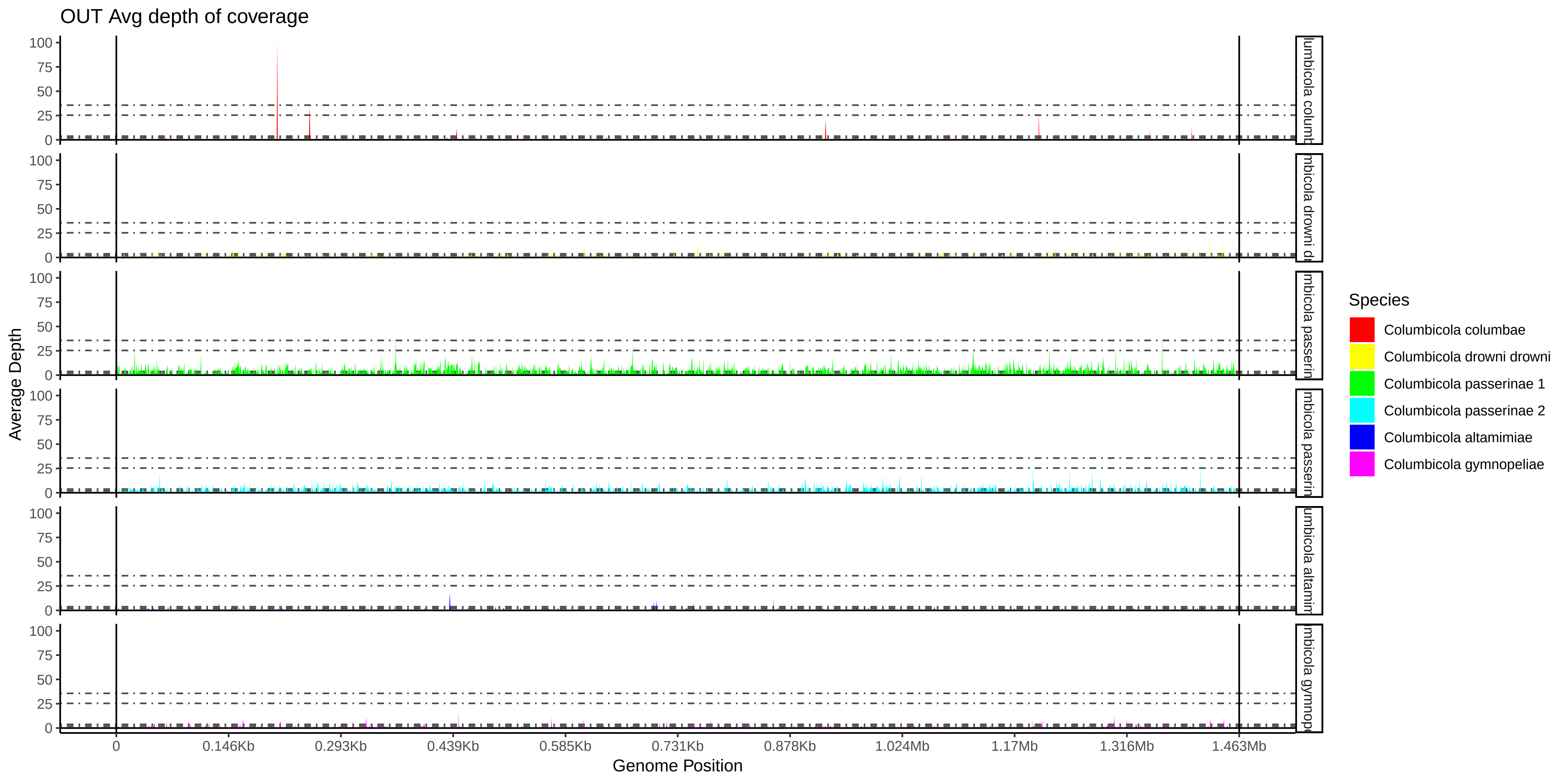

OUT Avg depth of coverage
Average Depth
0
25
50
75
100
umbicola columb
mbicola drowni dr
mbicola passerin
mbicola passerin
umbicola altamim
mbicola gymnope
0
0.146Kb
0.293Kb
0.439Kb
0.585Kb
0.731Kb
0.878Kb
1.024Mb
1.17Mb
1.316Mb
1.463Mb
Genome Position
Species
Columbicola columbae
Columbicola drowni drowni
Columbicola passerinae 1
Columbicola passerinae 2
Columbicola altamimiae
Columbicola gymnopeliae

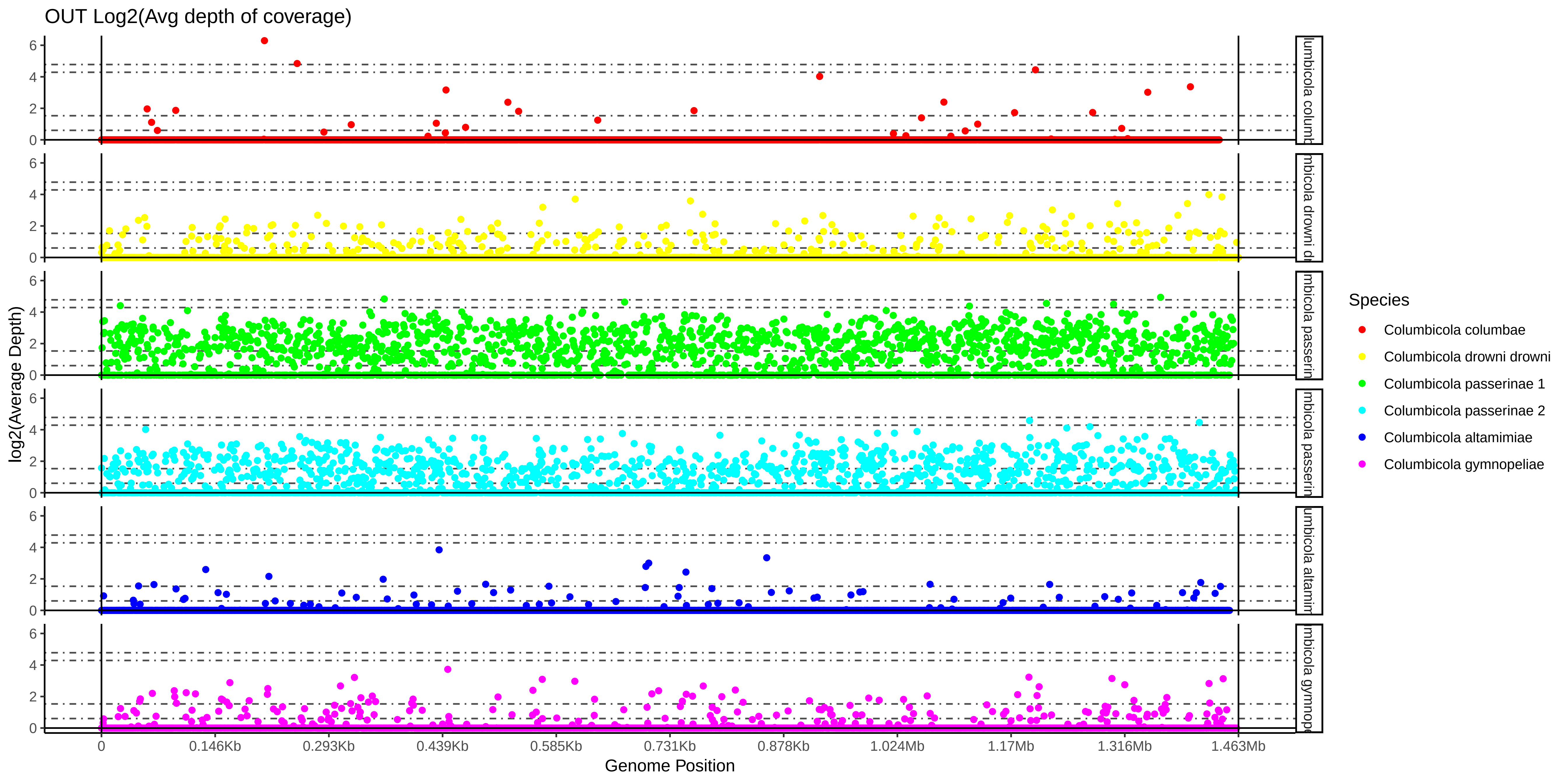
OUT Log2(Avg depth of coverage)
log2(Average Depth)
Genome Position
0
0.146Kb
0.293Kb
0.439Kb
0.585Kb
0.731Kb
0.878Kb
1.024Mb
1.17Mb
1.316Mb
1.463Mb
umbicola columb
mbicola drowni d
mbicola passerin
mbicola passerin
umbicola altamim
mbicola gymnop
Species
Columbicola columbae
Columbicola drowni drowni
Columbicola passerinae 1
Columbicola passerinae 2
Columbicola altamimiae
Columbicola gymnopeliae

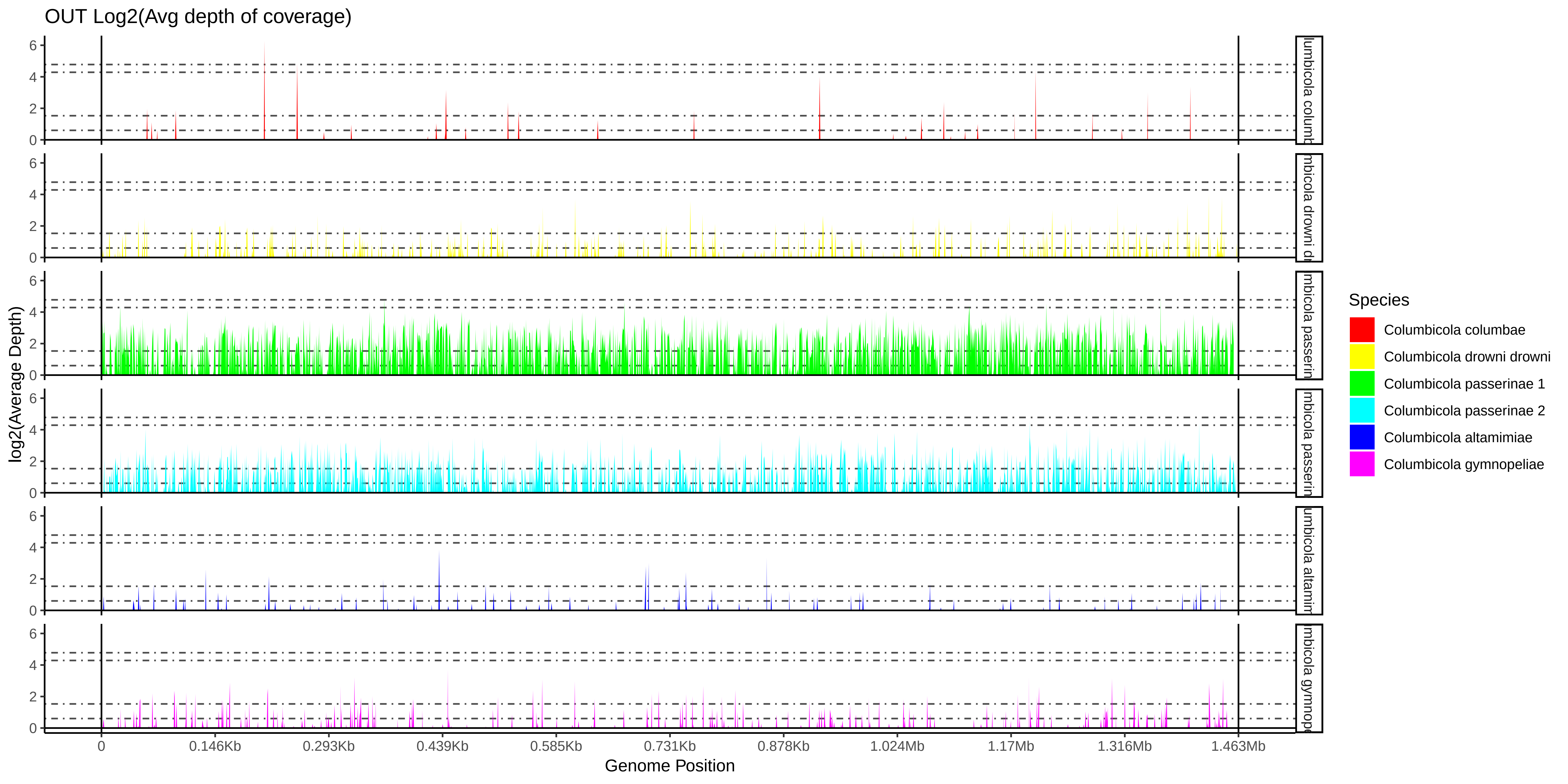

OUT Log2(Avg depth of coverage)
log2(Average Depth)
0
2
4
6
0
0.146Kb
0.293Kb
0.439Kb
0.585Kb
0.731Kb
0.878Kb
1.024Mb
1.17Mb
1.316Mb
1.463Mb
Genome Position
Species
Columbicola columbae
Columbicola drowni drowni
Columbicola passerinae 1
Columbicola passerinae 2
Columbicola altamimiae
Columbicola gymnopeliae

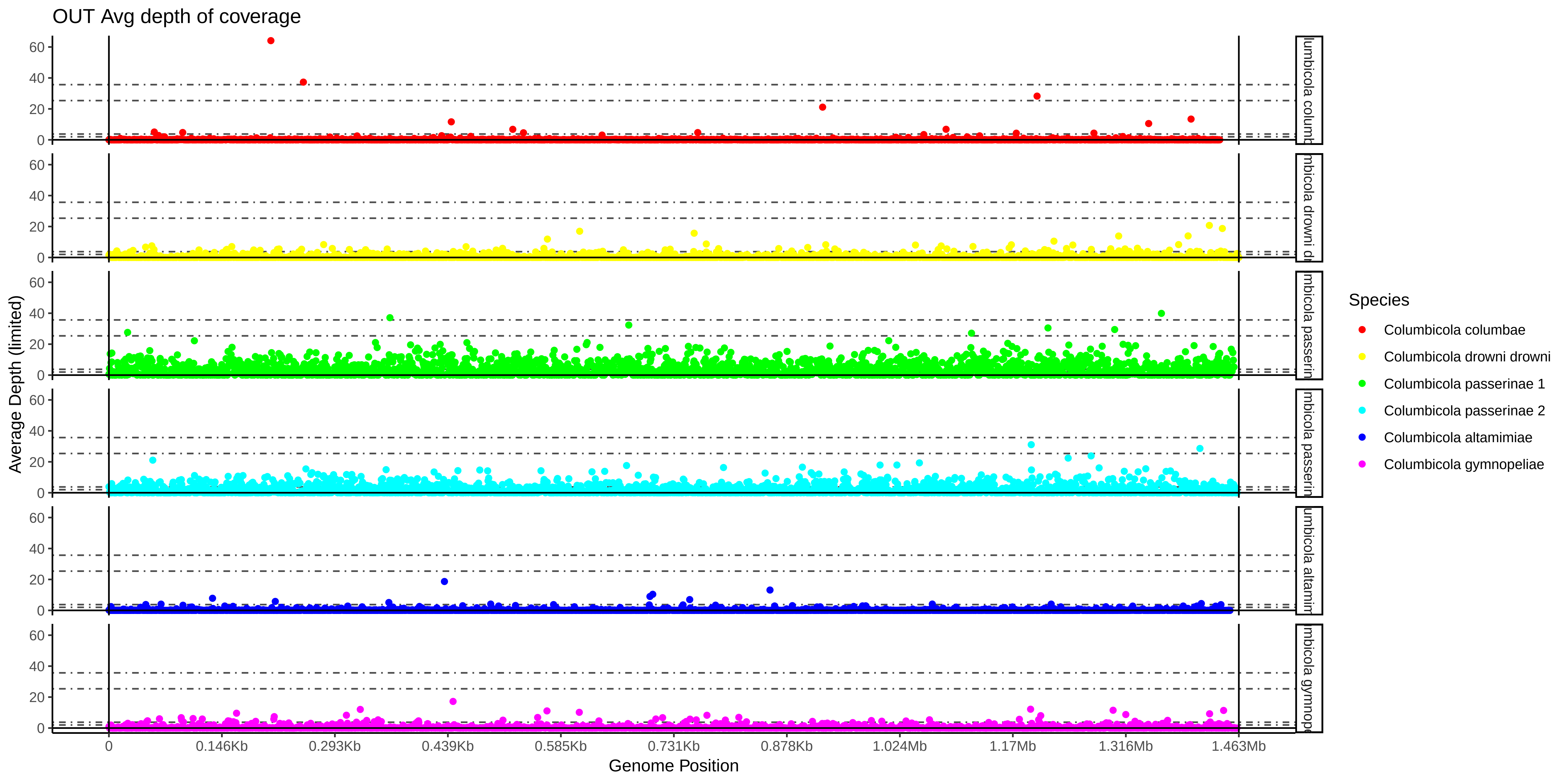

OUT Avg depth of coverage
Average Depth (limited)
Genome Position
0
0.146Kb
0.293Kb
0.439Kb
0.585Kb
0.731Kb
0.878Kb
1.024Mb
1.17Mb
1.316Mb
1.463Mb
0
20
40
60
Species
Columbicola columbae
Columbicola drowni drowni
Columbicola passerinae 1
Columbicola passerinae 2
Columbicola altamimiae
Columbicola gymnopeliae

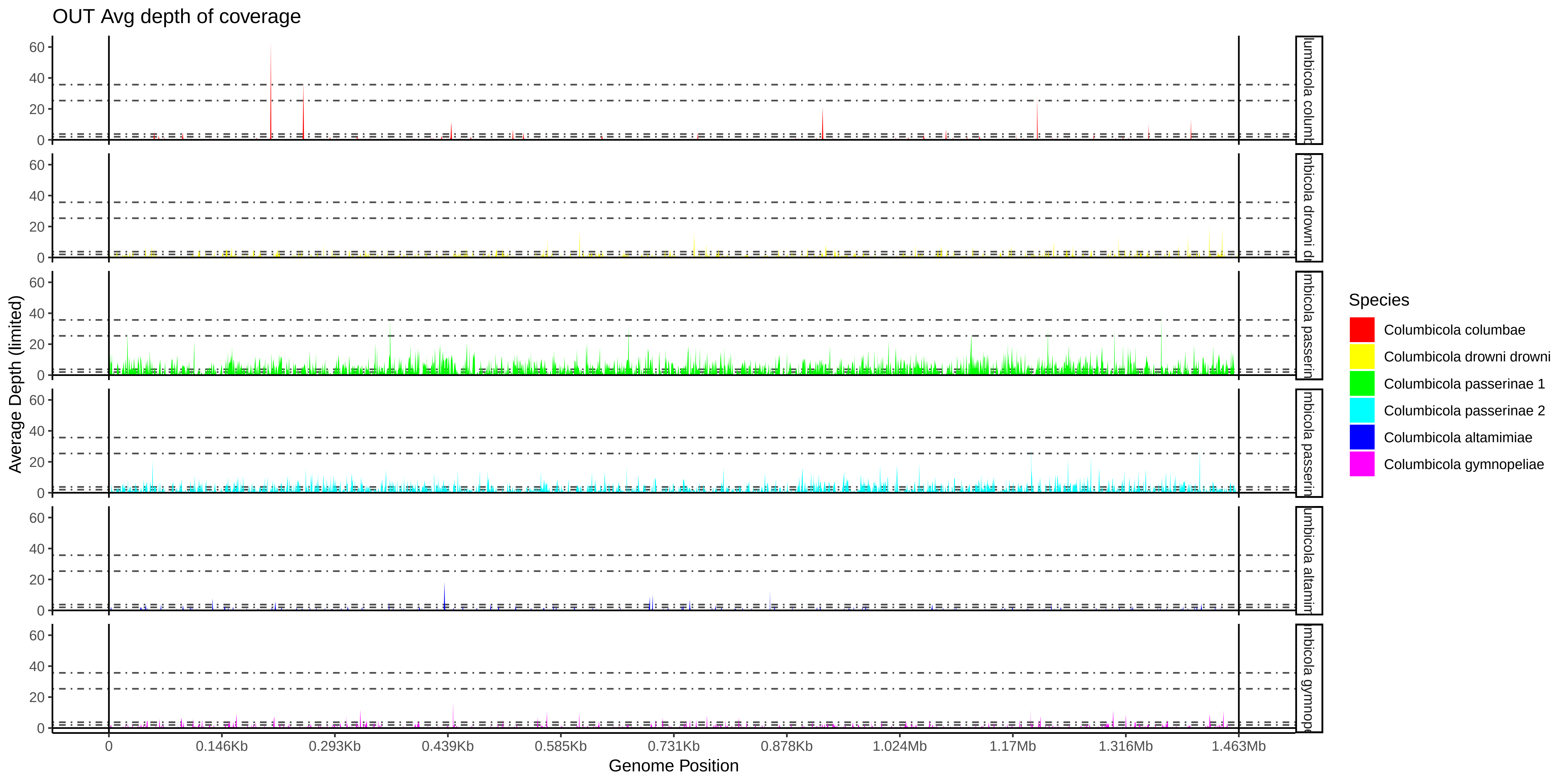

OUT Avg depth of coverage
Average Depth (limited)
Genome Position
0
0.146Kb
0.293Kb
0.439Kb
0.585Kb
0.731Kb
0.878Kb
1.024Mb
1.17Mb
1.316Mb
1.463Mb
0
20
40
60
umbicola columb
mbicola drowni d
mbicola passerin
mbicola passerin
umbicola altamim
mbicola gymnope
Species
Columbicola columbae
Columbicola drowni drowni
Columbicola passerinae 1
Columbicola passerinae 2
Columbicola altamimiae
Columbicola gymnopeliae
