## Supplementary material for "Comparing rates of introgression in parasitic feather lice with differing dispersal capabilities": OUT_sppIDerDepthPlot-covBins_Cosp_Cotal_11_7_2016_2.pdf (from another version of this preprint: version 7, 2020-03-31)

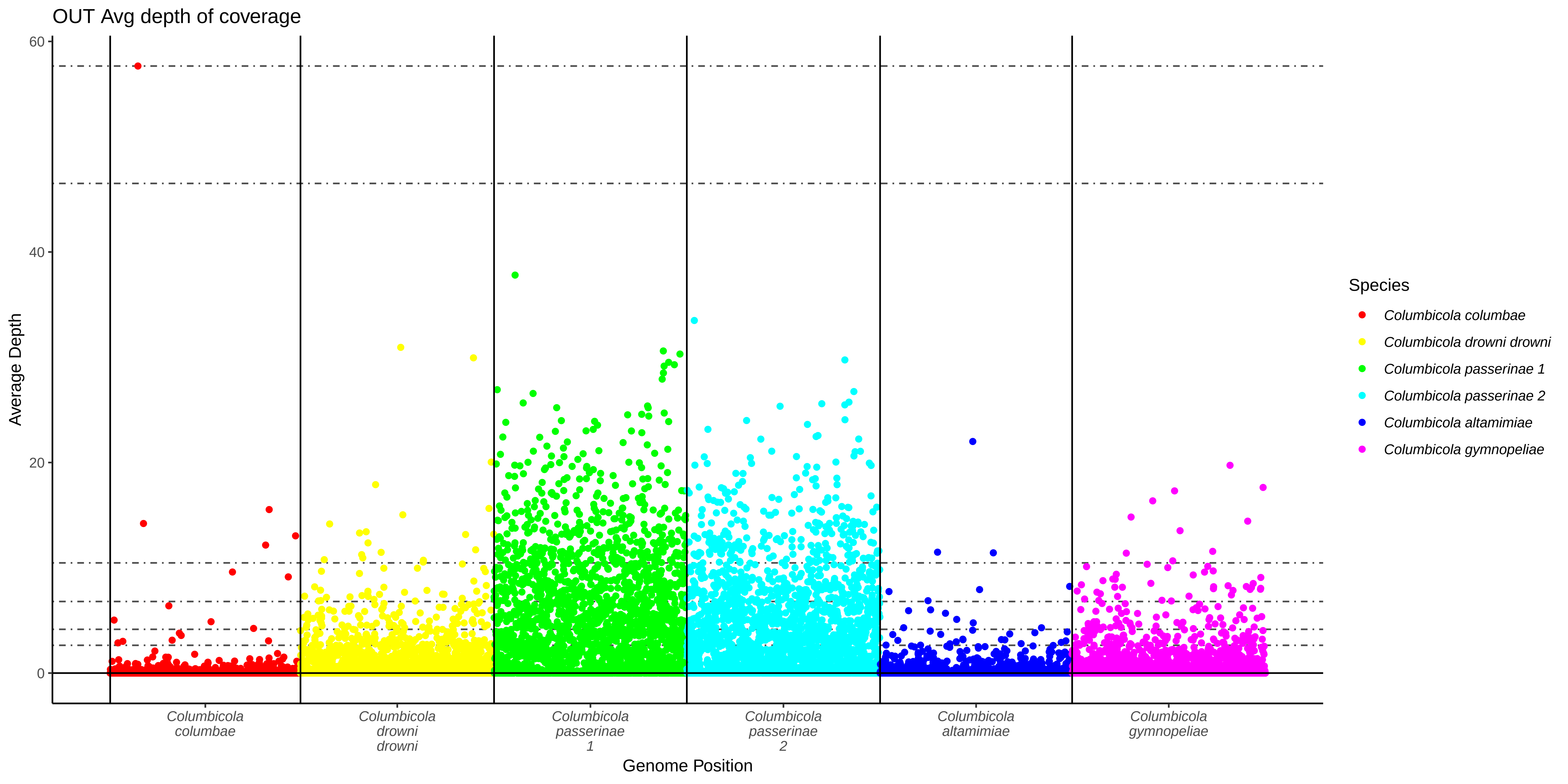

OUT Avg depth of coverage
Average Depth
60
40
20
0
Columbicola columbae
Columbicola drowni drowni
Columbicola passerinae 1
Columbicola passerinae 2
Columbicola altamimiae
Columbicola gymnopeliae
Genome Position
Species
Columbicola columbae
Columbicola drowni drowni
Columbicola passerinae 1
Columbicola passerinae 2
Columbicola altamimiae
Columbicola gymnopeliae

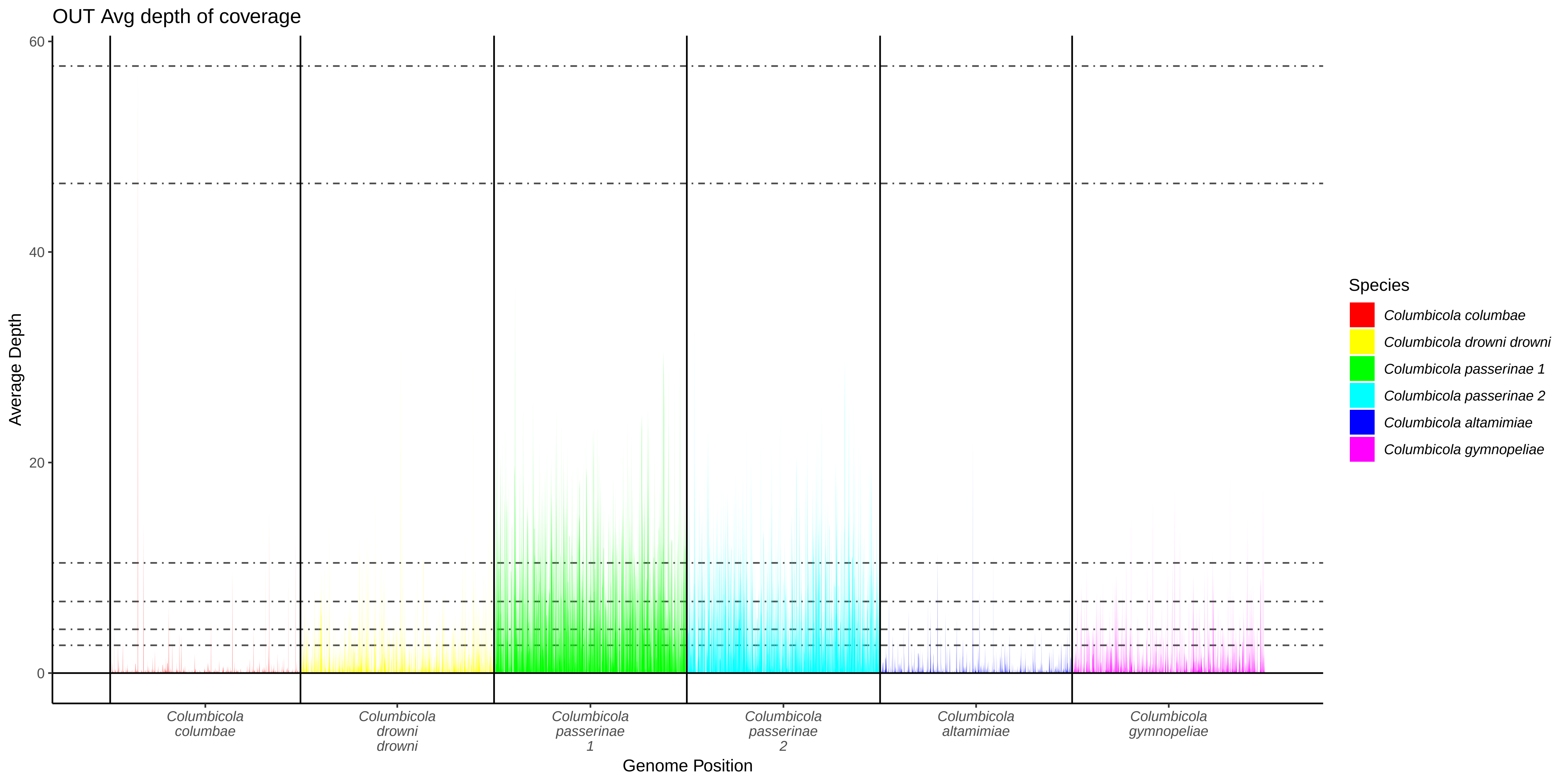

OUT Avg depth of coverage
Average Depth
0
20
40
60
Columbicola columbae
Columbicola drowni drowni
Columbicola passerinae 1
Columbicola passerinae 2
Columbicola altamimiae
Columbicola gymnopeliae
Genome Position
Species
Columbicola columbae
Columbicola drowni drowni
Columbicola passerinae 1
Columbicola passerinae 2
Columbicola altamimiae
Columbicola gymnopeliae

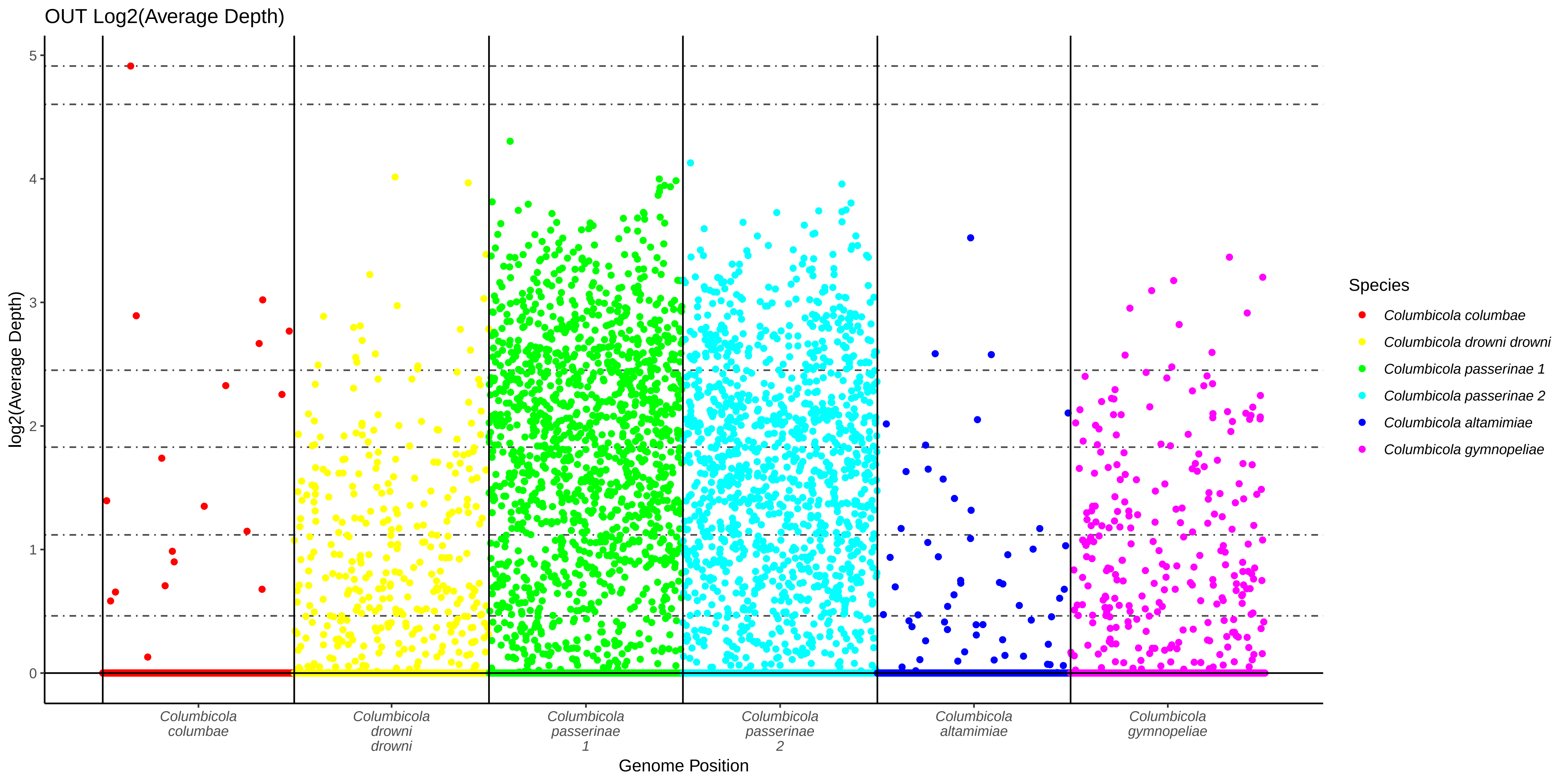

OUT Log2(Average Depth)
log2(Average Depth)
0
1
2
3
4
5
Columbicola columbae
Columbicola drowni drowni
Columbicola passerinae 1
Columbicola passerinae 2
Columbicola altamimiae
Columbicola gymnopeliae
Genome Position
Species
Columbicola columbae
Columbicola drowni drowni
Columbicola passerinae 1
Columbicola passerinae 2
Columbicola altamimiae
Columbicola gymnopeliae

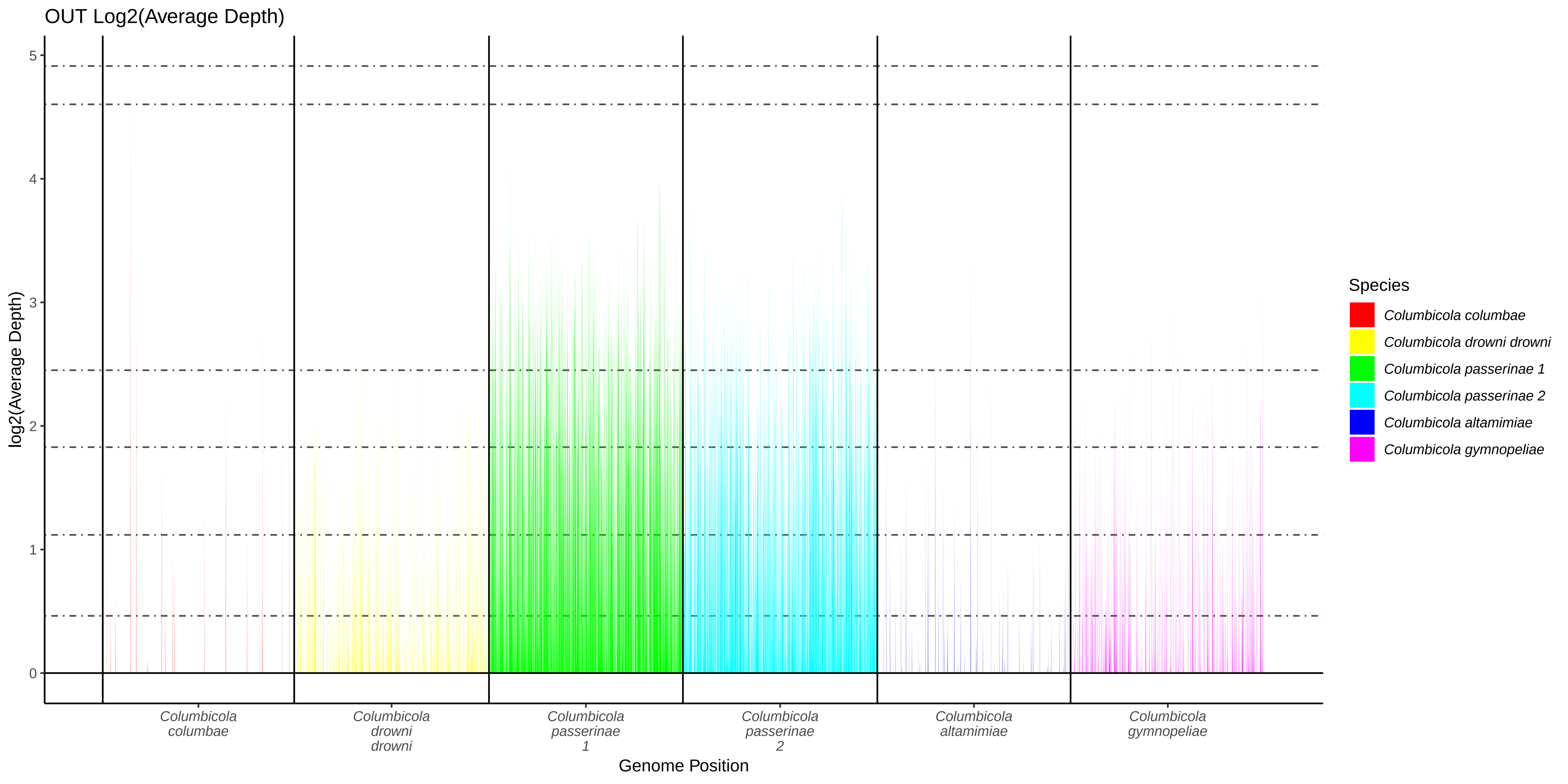
OUT Log2(Average Depth)
log2(Average Depth)
0
1
2
3
4
5
Columbicola columbae
Columbicola drowni drowni
Columbicola passerinae 1
Columbicola passerinae 2
Columbicola altamimiae
Columbicola gymnopeliae
Genome Position
Species
Columbicola columbae
Columbicola drowni drowni
Columbicola passerinae 1
Columbicola passerinae 2
Columbicola altamimiae
Columbicola gymnopeliae

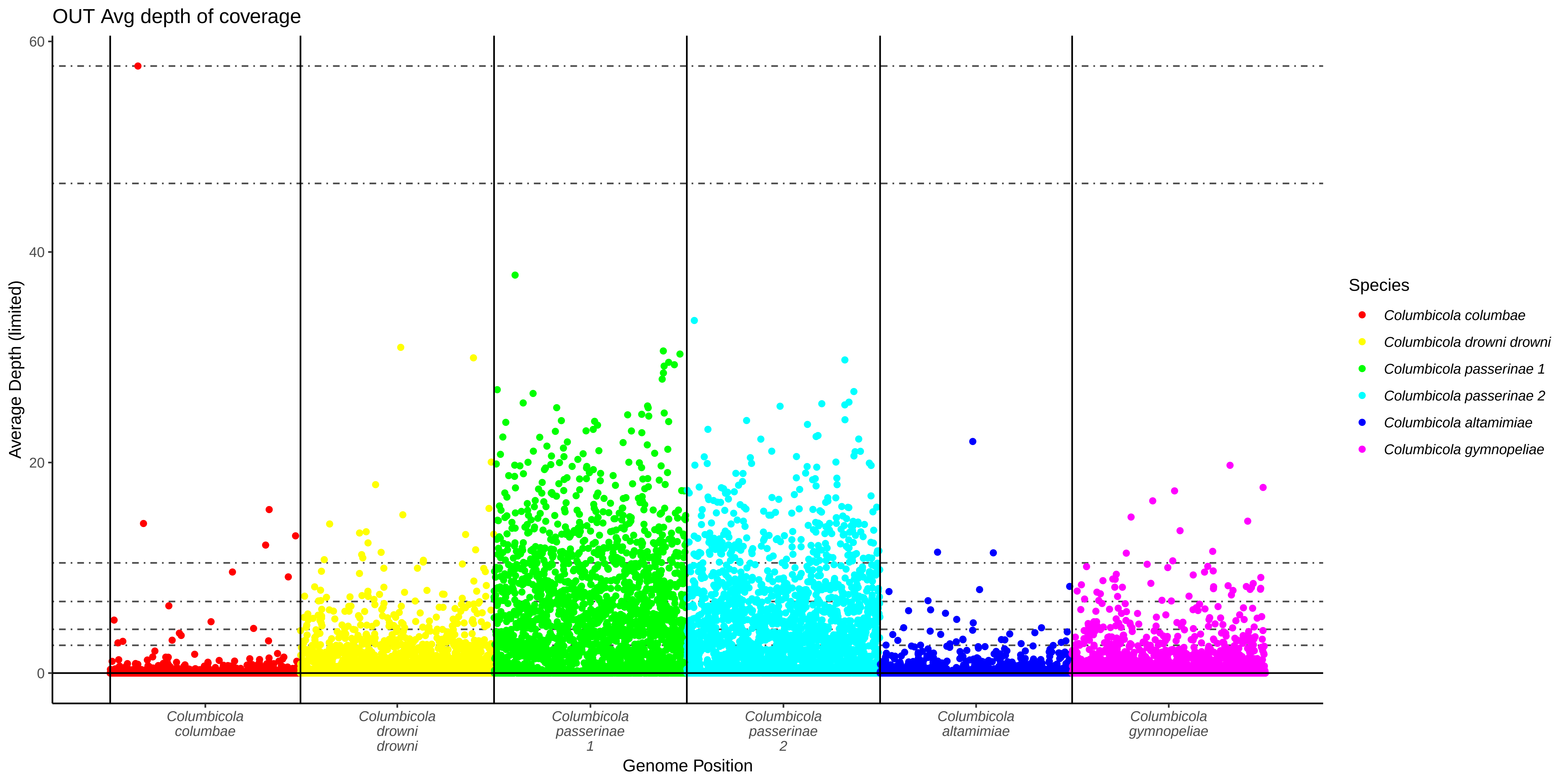

OUT Avg depth of coverage
60
40
20
0
Average Depth (limited)
Columbicola columbae
Columbicola drowni drowni
Columbicola passerinae 1
Columbicola passerinae 2
Columbicola altamimiae
Columbicola gymnopeliae
Genome Position
Species
Columbicola columbae
Columbicola drowni drowni
Columbicola passerinae 1
Columbicola passerinae 2
Columbicola altamimiae
Columbicola gymnopeliae

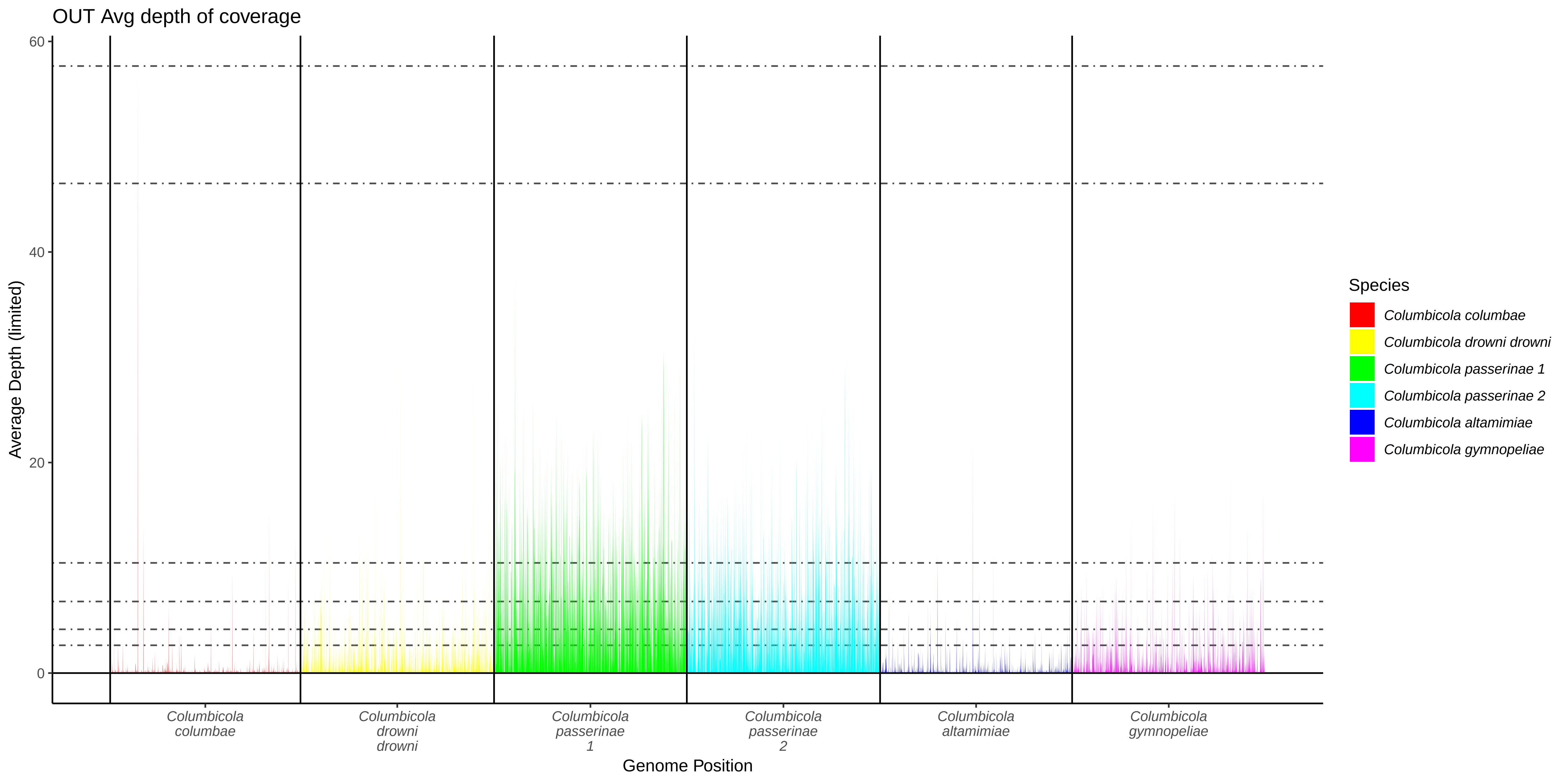
OUT Avg depth of coverage
Average Depth (limited)
60
40
20
0
Columbicola columbae
Columbicola drowni drowni
Columbicola passerinae 1
Columbicola passerinae 2
Columbicola altamimiae
Columbicola gymnopeliae
Genome Position
Species
Columbicola columbae
Columbicola drowni drowni
Columbicola passerinae 1
Columbicola passerinae 2
Columbicola altamimiae
Columbicola gymnopeliae

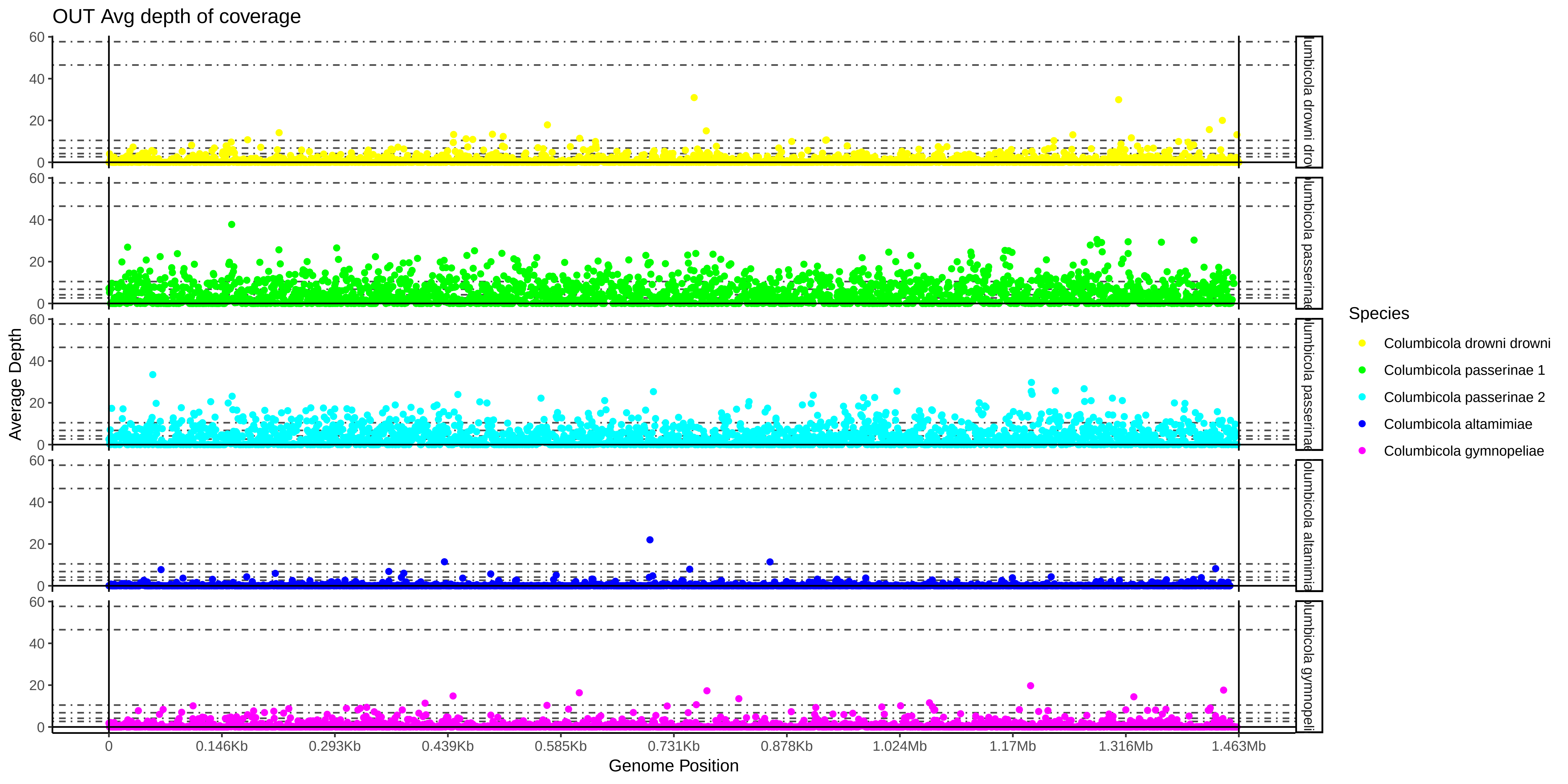

OUT Avg depth of coverage
Average Depth
Genome Position
0
0.146Kb
0.293Kb
0.439Kb
0.585Kb
0.731Kb
0.878Kb
1.024Mb
1.17Mb
1.316Mb
1.463Mb
Species
Columbicola drowni drowni
Columbicola passerinae 1
Columbicola passerinae 2
Columbicola altamimiae
Columbicola gymnopeliae

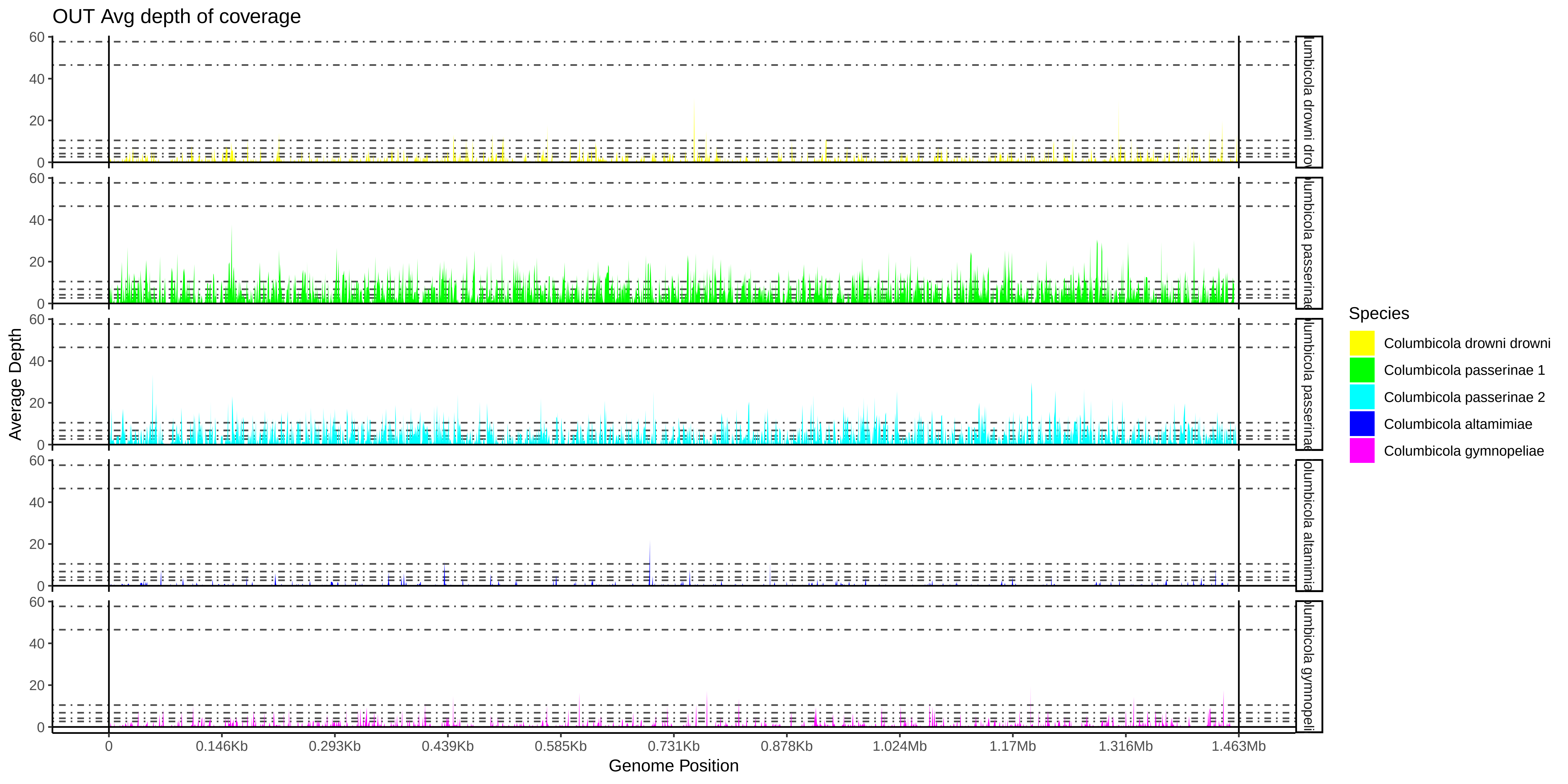
OUT Avg depth of coverage
Average Depth
Genome Position
0
0.146Kb
0.293Kb
0.439Kb
0.585Kb
0.731Kb
0.878Kb
1.024Mb
1.17Mb
1.316Mb
1.463Mb
60
40
20
0
Species
Columbicola drowni drowni
Columbicola passerinae 1
Columbicola passerinae 2
Columbicola altamimiae
Columbicola gymnopeliae

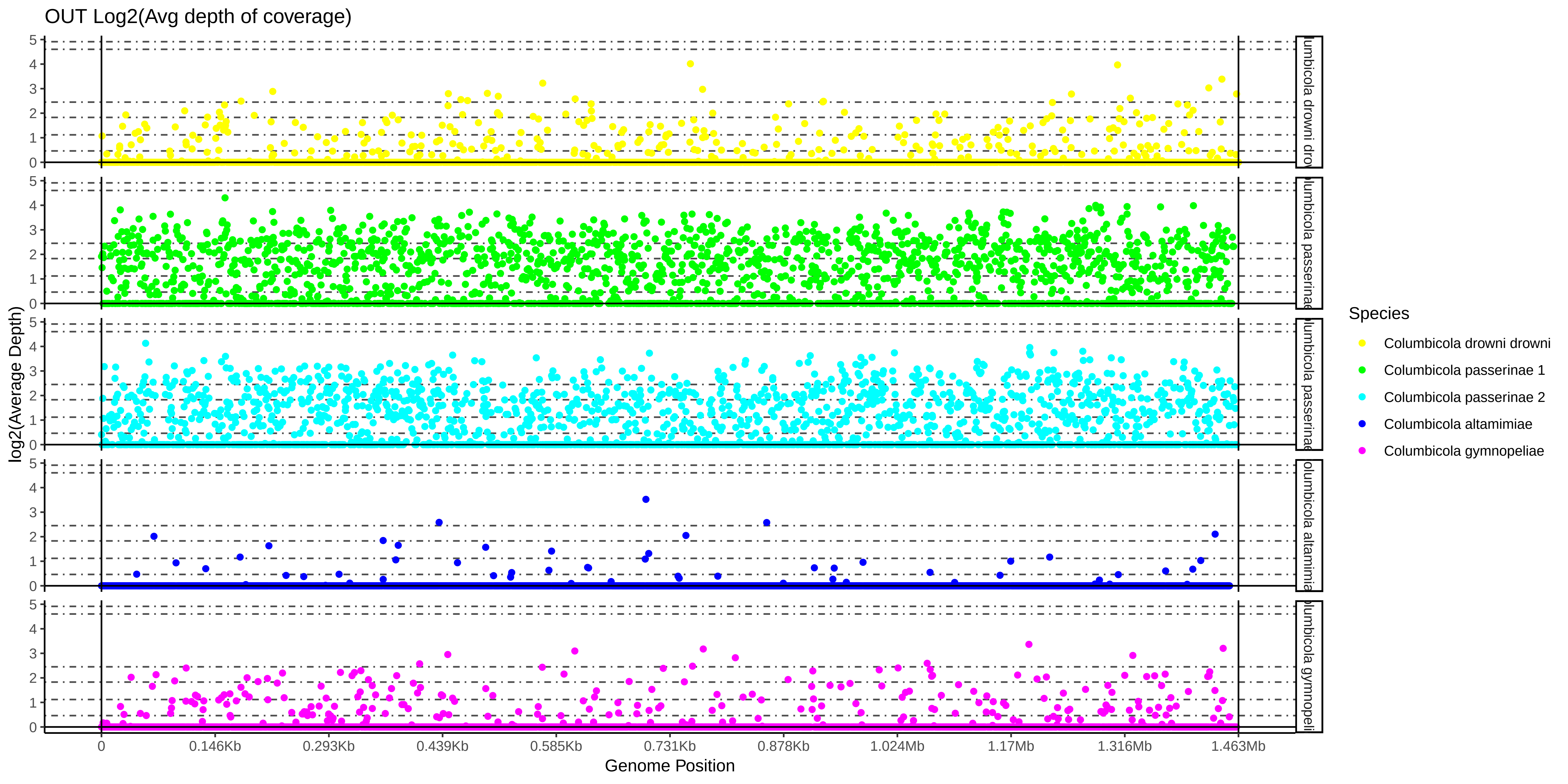

OUT Log2(Avg depth of coverage)
log2(Average Depth)
Genome Position
0
0.146Kb
0.293Kb
0.439Kb
0.585Kb
0.731Kb
0.878Kb
1.024Mb
1.17Mb
1.316Mb
1.463Mb
Species
Columbicola drowni drowni
Columbicola passerinae 1
Columbicola passerinae 2
Columbicola altamimiae
Columbicola gymnopeliae

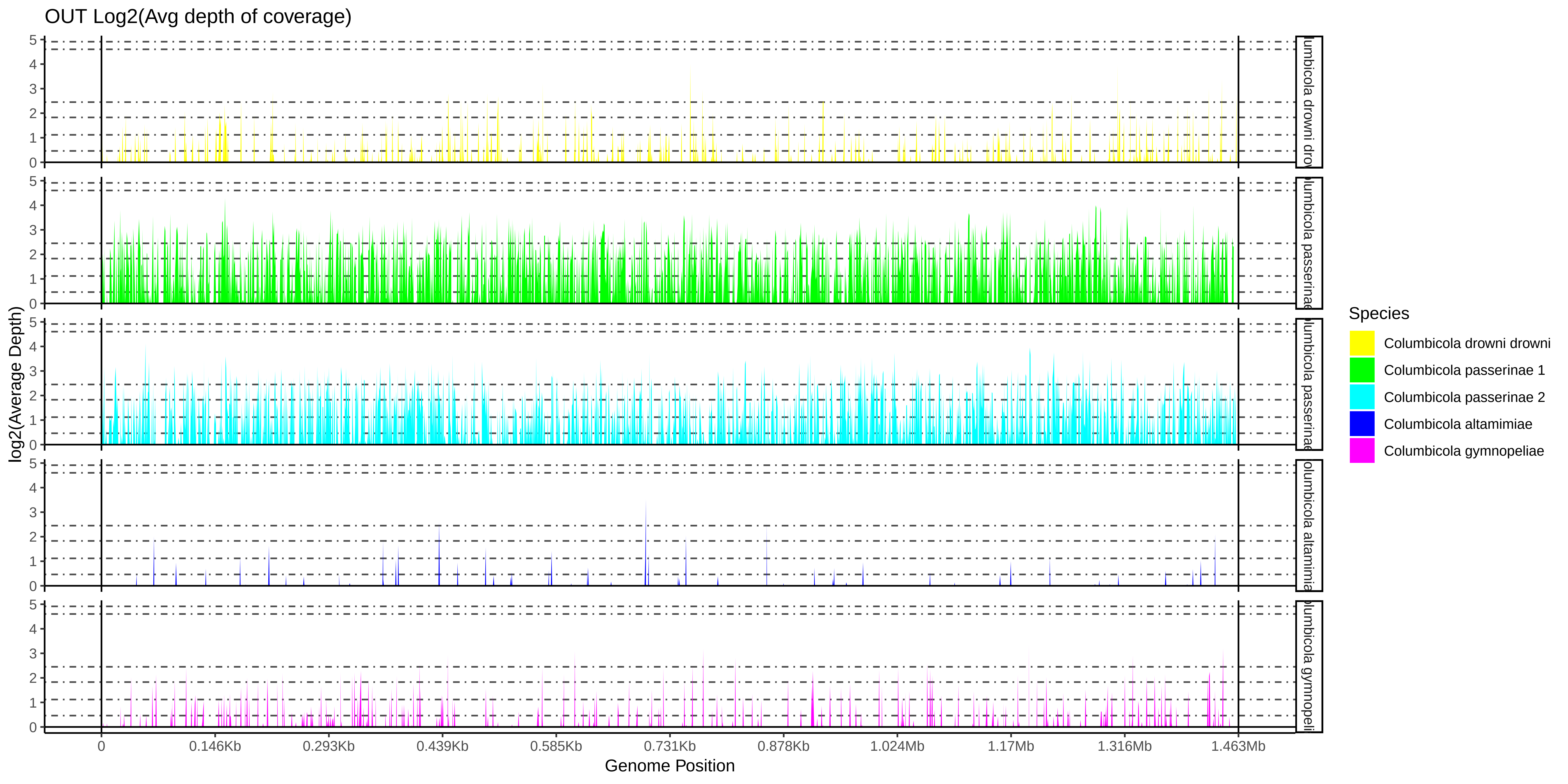
OUT Log2(Avg depth of coverage)
log2(Average Depth)
Genome Position
0
0.146Kb
0.293Kb
0.439Kb
0.585Kb
0.731Kb
0.878Kb
1.024Mb
1.17Mb
1.316Mb
1.463Mb
Species
Columbicola drowni drowni
Columbicola passerinae 1
Columbicola passerinae 2
Columbicola altamimiae
Columbicola gymnopeliae

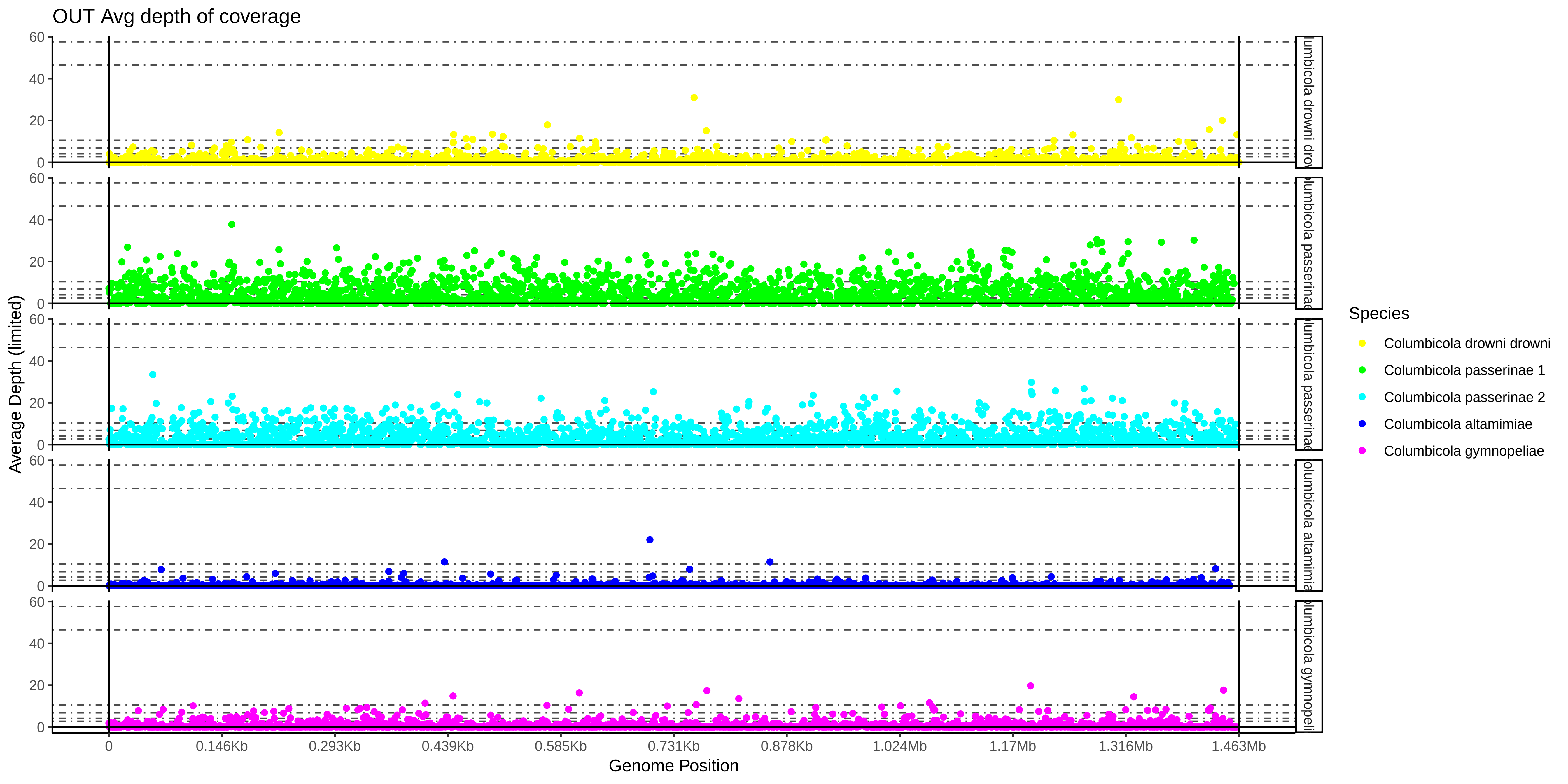

OUT Avg depth of coverage
Average Depth (limited)
Genome Position
0
0.146Kb
0.293Kb
0.439Kb
0.585Kb
0.731Kb
0.878Kb
1.024Mb
1.17Mb
1.316Mb
1.463Mb
Species
Columbicola drowni drowni
Columbicola passerinae 1
Columbicola passerinae 2
Columbicola altamimiae
Columbicola gymnopeliae

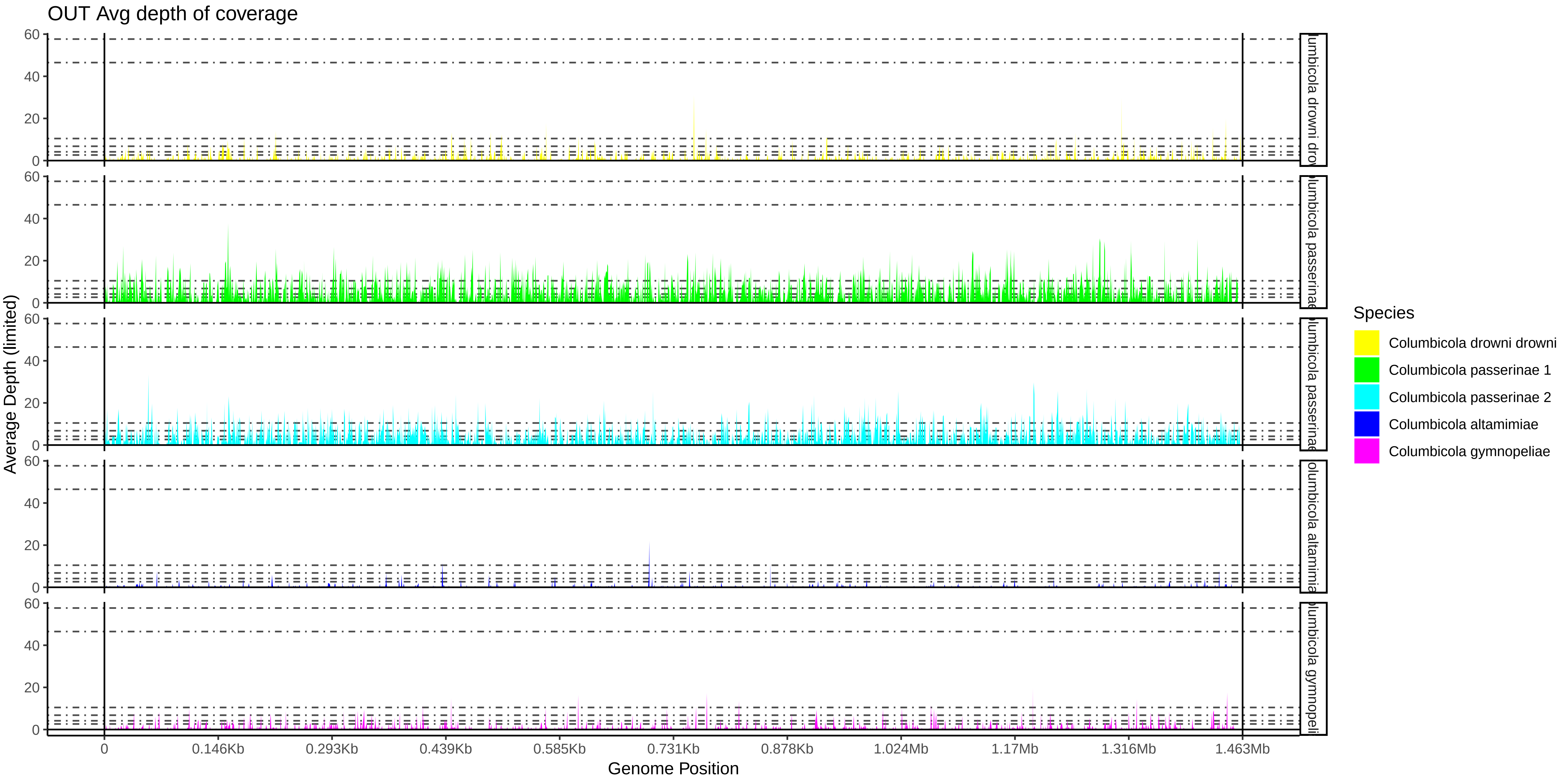

OUT Avg depth of coverage
Average Depth (limited)
Genome Position
0
0.146Kb
0.293Kb
0.439Kb
0.585Kb
0.731Kb
0.878Kb
1.024Mb
1.17Mb
1.316Mb
1.463Mb
Species
Columbicola drowni drowni
Columbicola passerinae 1
Columbicola passerinae 2
Columbicola altamimiae
Columbicola gymnopeliae
