## Supplementary material for "Comparing rates of introgression in parasitic feather lice with differing dispersal capabilities": OUT_sppIDerDepthPlot-covBins_Pheur1_Comin_3_14_2016_9.pdf (from another version of this preprint: version 7, 2020-03-31)

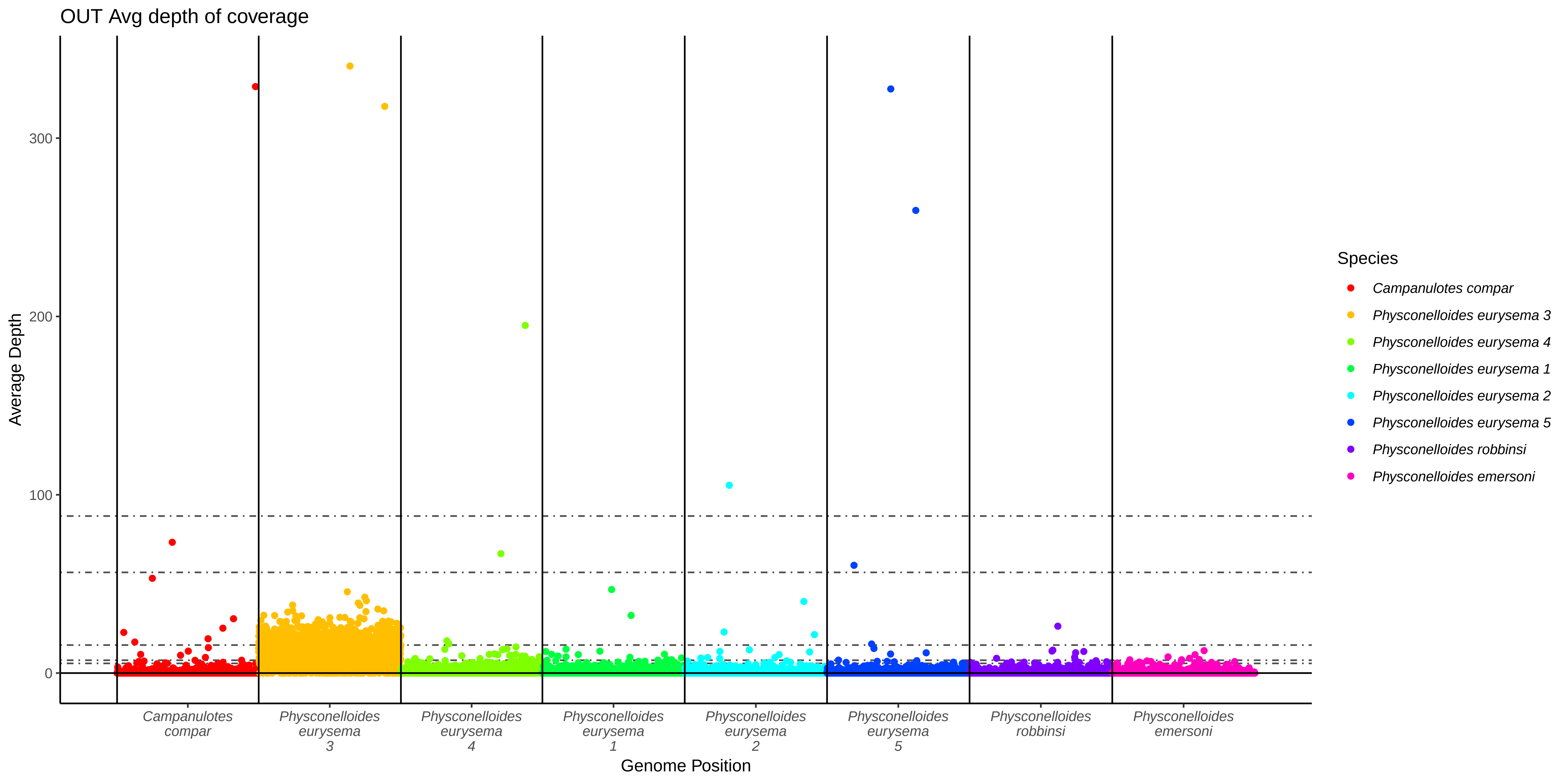

OUT Avg depth of coverage
Average Depth
300
200
100
0
Campanulotes compar
Physconelloides eurysema 3
Physconelloides eurysema 4
Physconelloides eurysema 1
Physconelloides eurysema 2
Physconelloides eurysema 5
Physconelloides robbinsi
Physconelloides emersoni
Genome Position
Species
Campanulotes compar
Physconelloides eurysema 3
Physconelloides eurysema 4
Physconelloides eurysema 1
Physconelloides eurysema 2
Physconelloides eurysema 5
Physconelloides robbinsi
Physconelloides emersoni

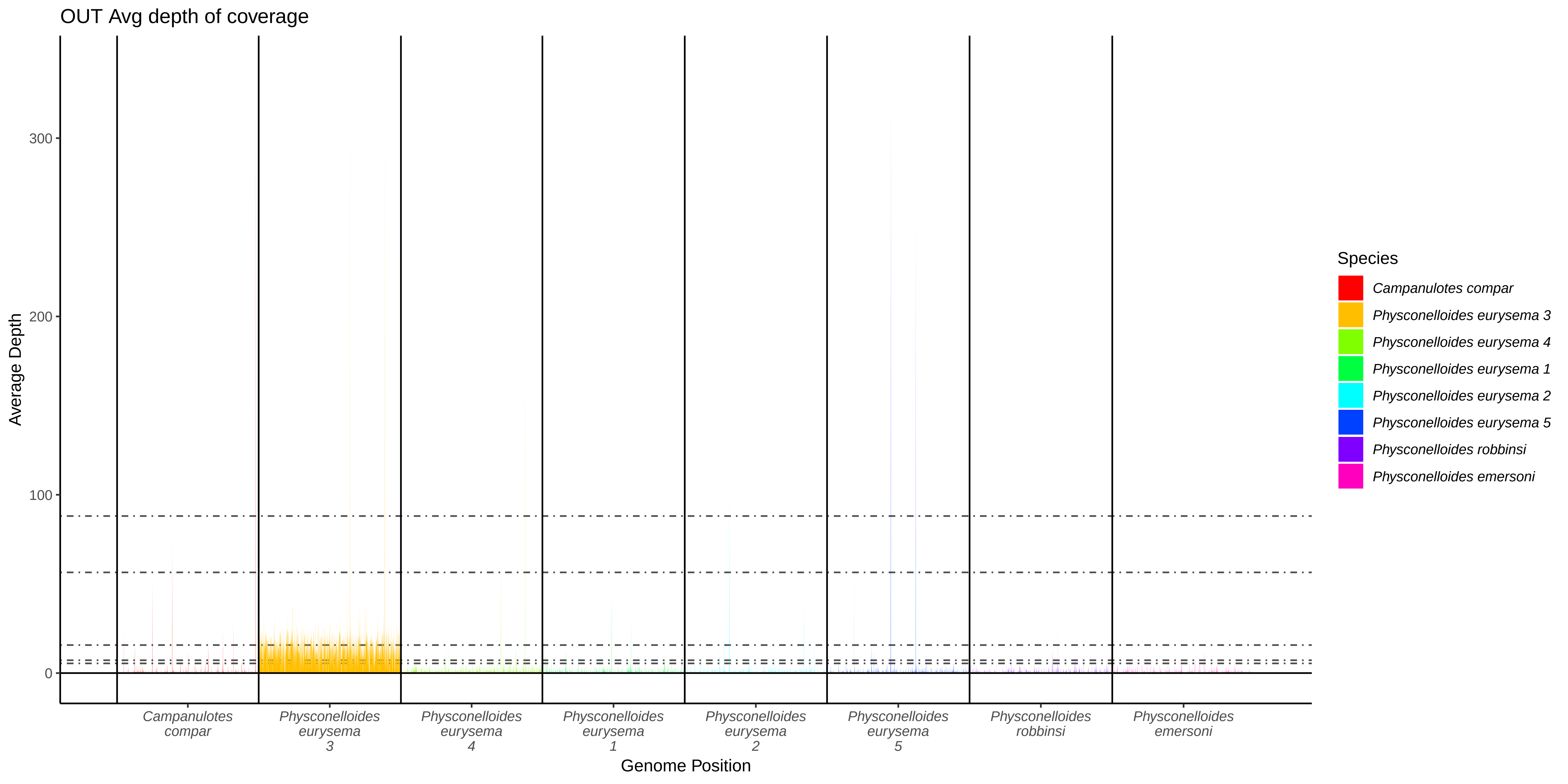
OUT Avg depth of coverage
Average Depth
300
200
100
0
Campanulotes compar
Physconelloides eurysema 3
Physconelloides eurysema 4
Physconelloides eurysema 1
Physconelloides eurysema 2
Physconelloides eurysema 5
Physconelloides robbinsi
Physconelloides emersoni
Genome Position
Species
Campanulotes compar
Physconelloides eurysema 3
Physconelloides eurysema 4
Physconelloides eurysema 1
Physconelloides eurysema 2
Physconelloides eurysema 5
Physconelloides robbinsi
Physconelloides emersoni

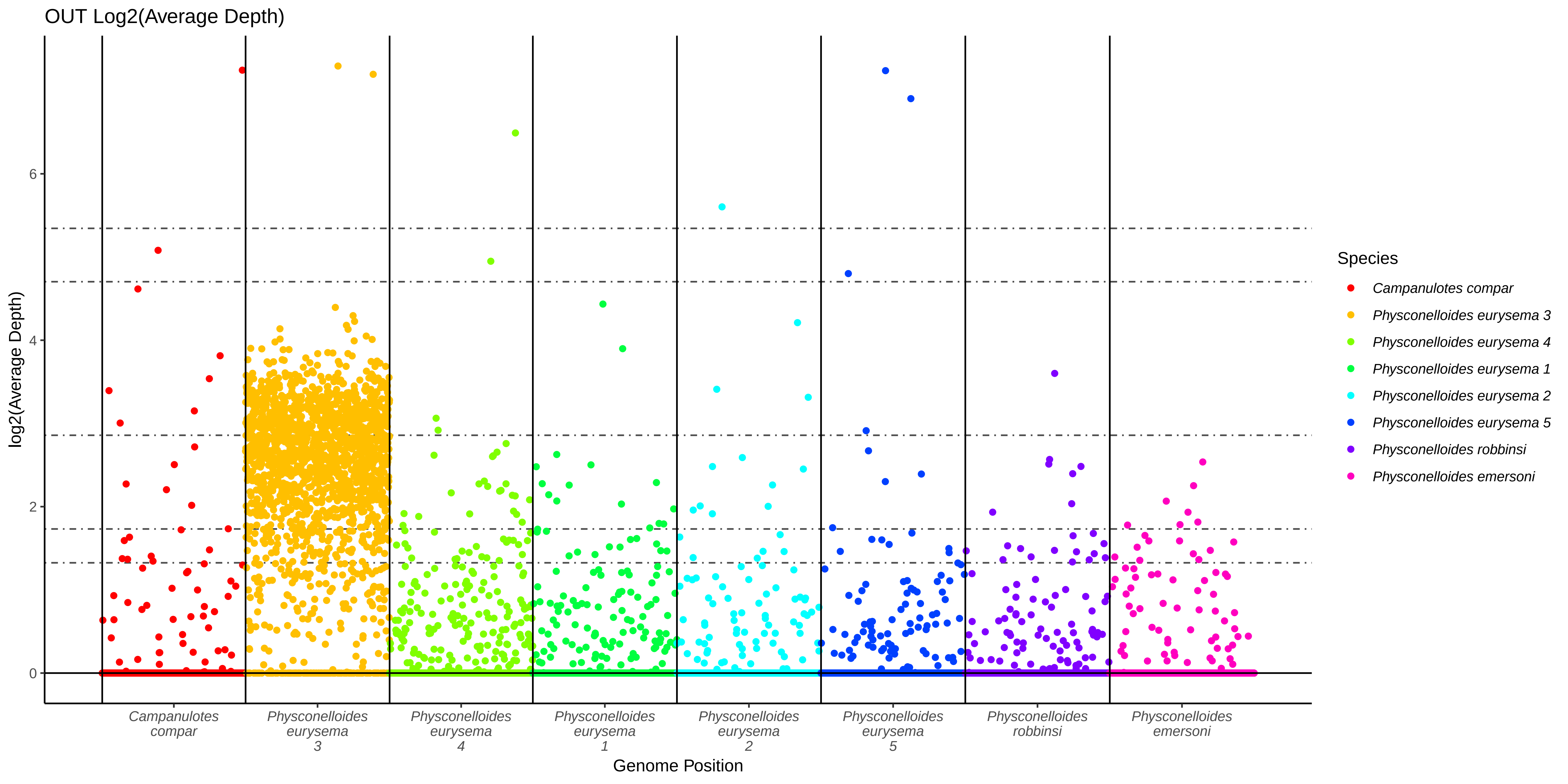

OUT Log2(Average Depth)
log2(Average Depth)
0
2
4
6
Genome Position
Campanulotes compar
Physconelloides eurysema 3
Physconelloides eurysema 4
Physconelloides eurysema 1
Physconelloides eurysema 2
Physconelloides eurysema 5
Physconelloides robbinsi
Physconelloides emersoni
Species
Campanulotes compar
Physconelloides eurysema 3
Physconelloides eurysema 4
Physconelloides eurysema 1
Physconelloides eurysema 2
Physconelloides eurysema 5
Physconelloides robbinsi
Physconelloides emersoni

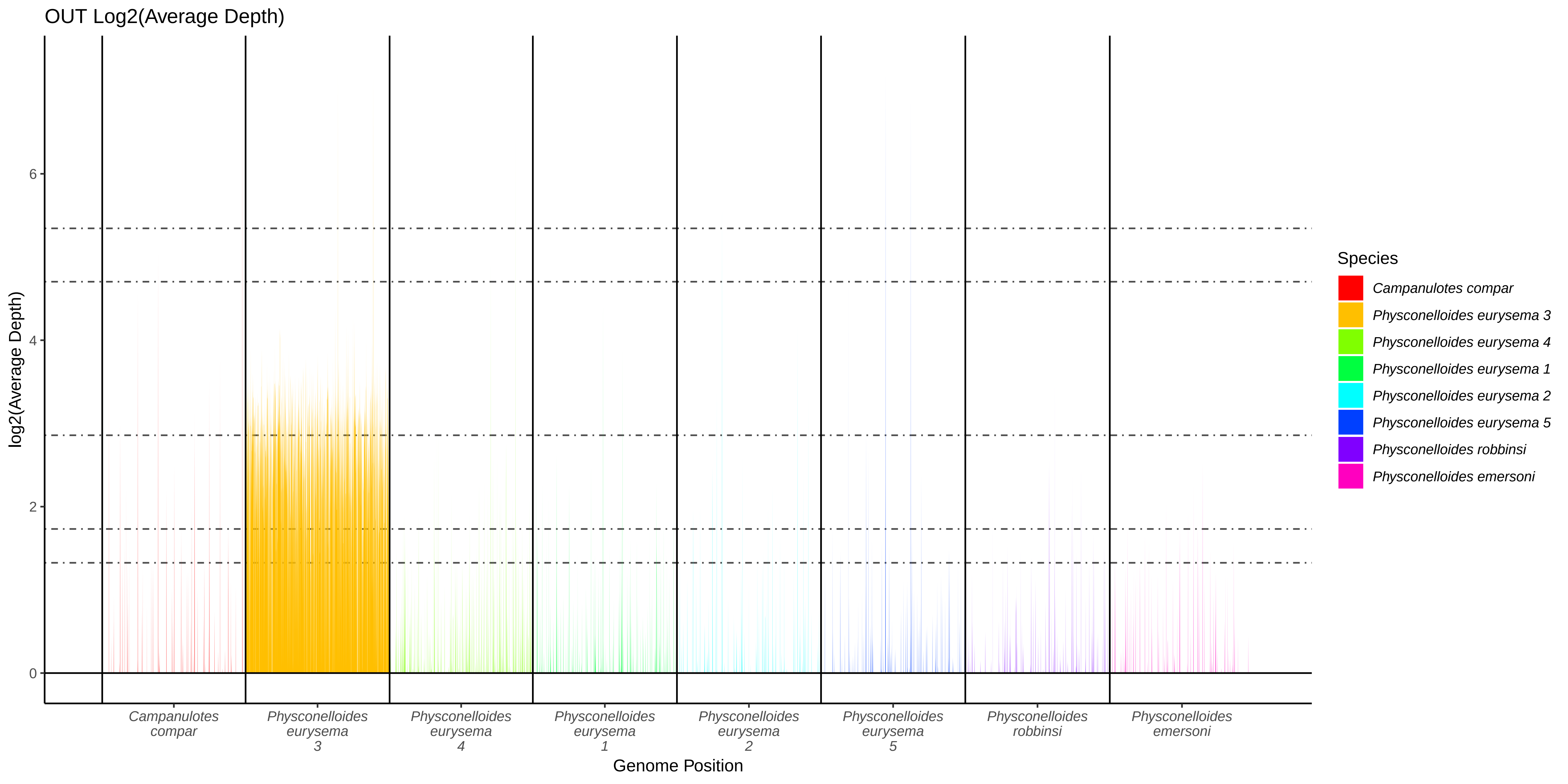
OUT Log2(Average Depth)
log2(Average Depth)
6
4
2
0
Campanulotes compar
Physconelloides eurysema 3
Physconelloides eurysema 4
Physconelloides eurysema 1
Physconelloides eurysema 2
Physconelloides eurysema 5
Physconelloides robbinsi
Physconelloides emersoni
Genome Position
Species
Campanulotes compar
Physconelloides eurysema 3
Physconelloides eurysema 4
Physconelloides eurysema 1
Physconelloides eurysema 2
Physconelloides eurysema 5
Physconelloides robbinsi
Physconelloides emersoni

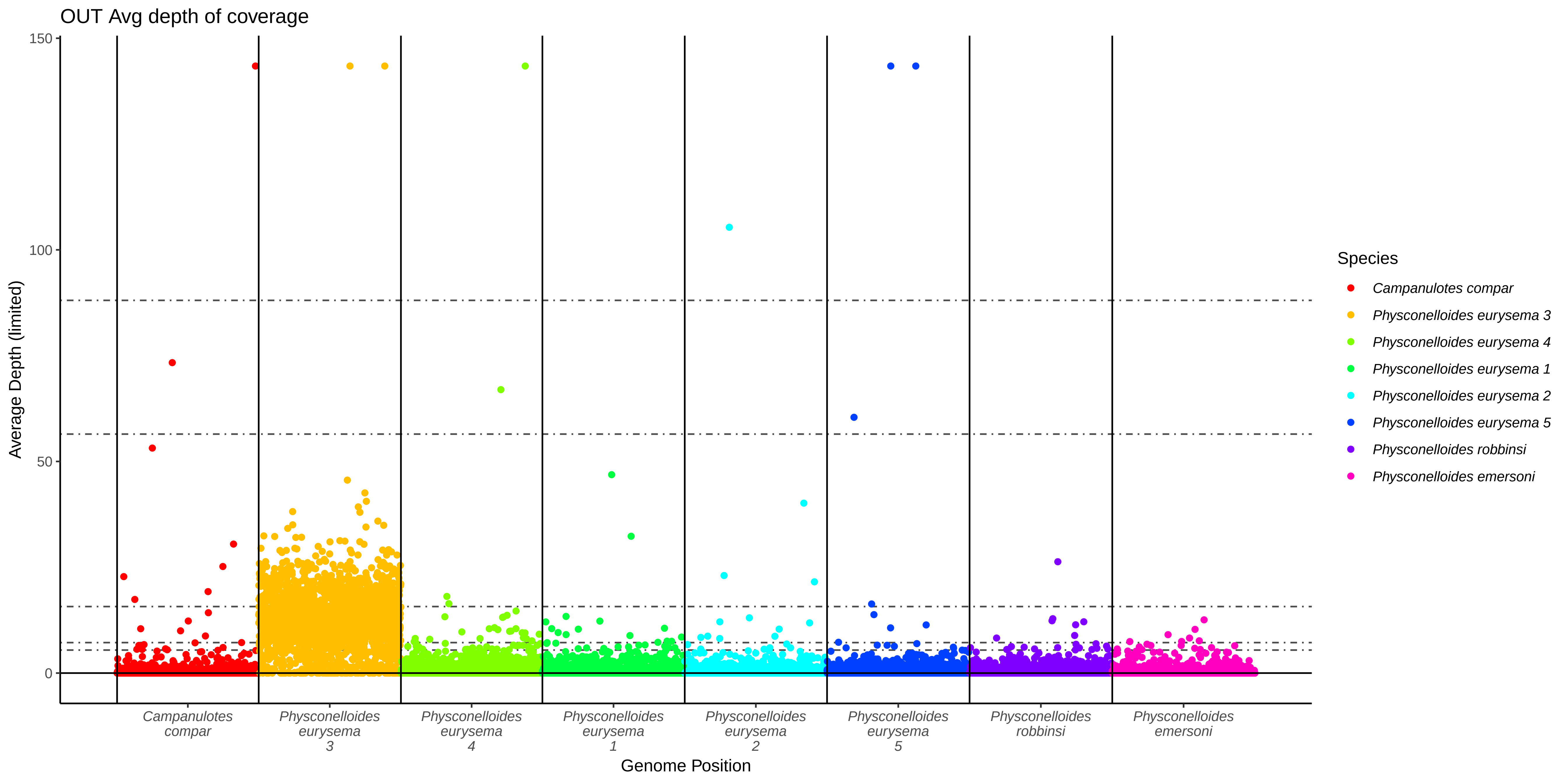
OUT Avg depth of coverage
Average Depth (limited)
150
100
50
0
Campanulotes compar
Physconelloides eurysema 3
Physconelloides eurysema 4
Physconelloides eurysema 1
Physconelloides eurysema 2
Physconelloides eurysema 5
Physconelloides robbinsi
Physconelloides emersoni
Genome Position
Species
Campanulotes compar
Physconelloides eurysema 3
Physconelloides eurysema 4
Physconelloides eurysema 1
Physconelloides eurysema 2
Physconelloides eurysema 5
Physconelloides robbinsi
Physconelloides emersoni

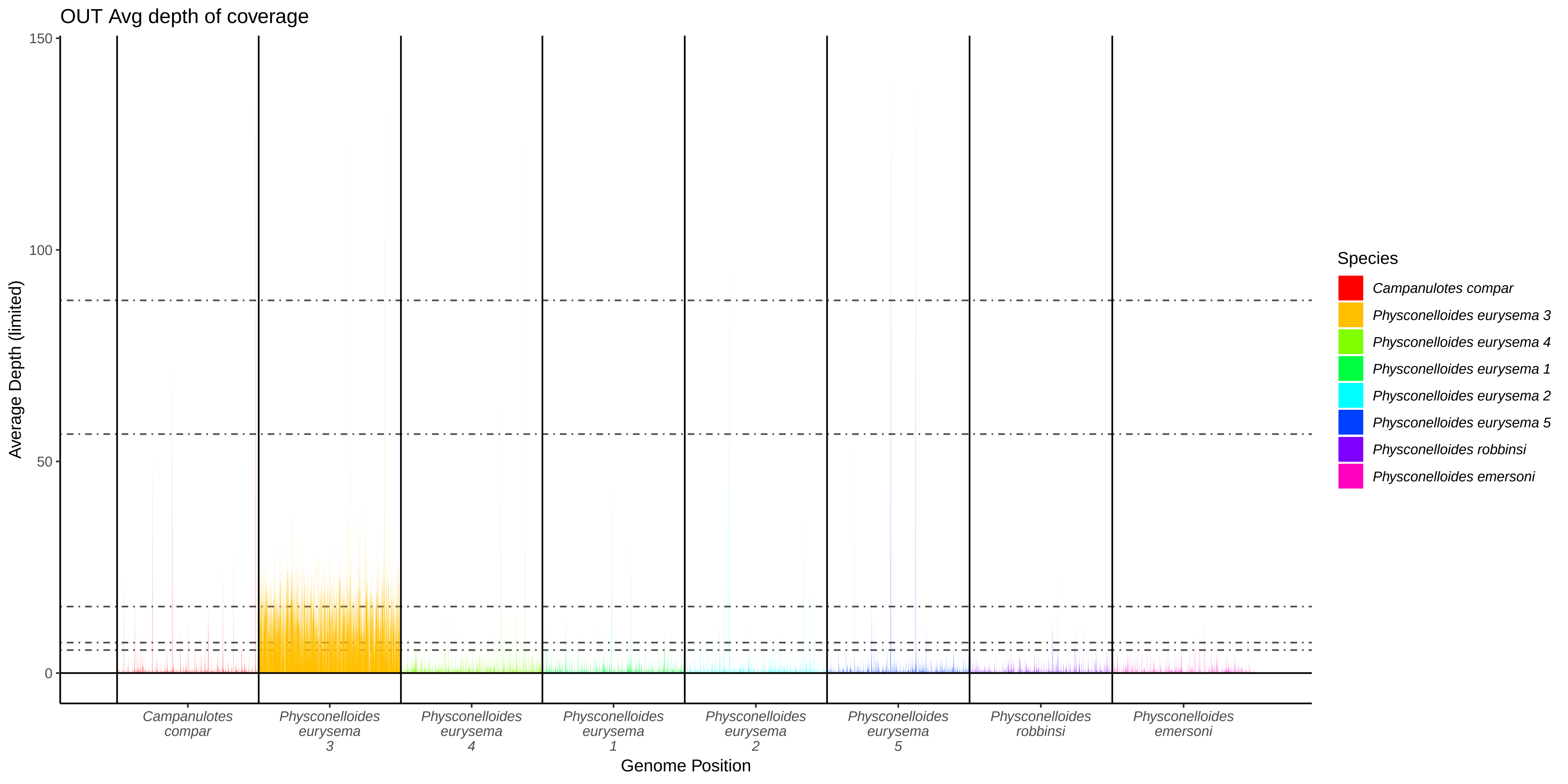

OUT Avg depth of coverage
150
100
50
0
Average Depth (limited)
Genome Position
Campanulotes compar
Physconelloides eurysema 3
Physconelloides eurysema 4
Physconelloides eurysema 1
Physconelloides eurysema 2
Physconelloides eurysema 5
Physconelloides robbinsi
Physconelloides emersoni
Species
Campanulotes compar
Physconelloides eurysema 3
Physconelloides eurysema 4
Physconelloides eurysema 1
Physconelloides eurysema 2
Physconelloides eurysema 5
Physconelloides robbinsi
Physconelloides emersoni

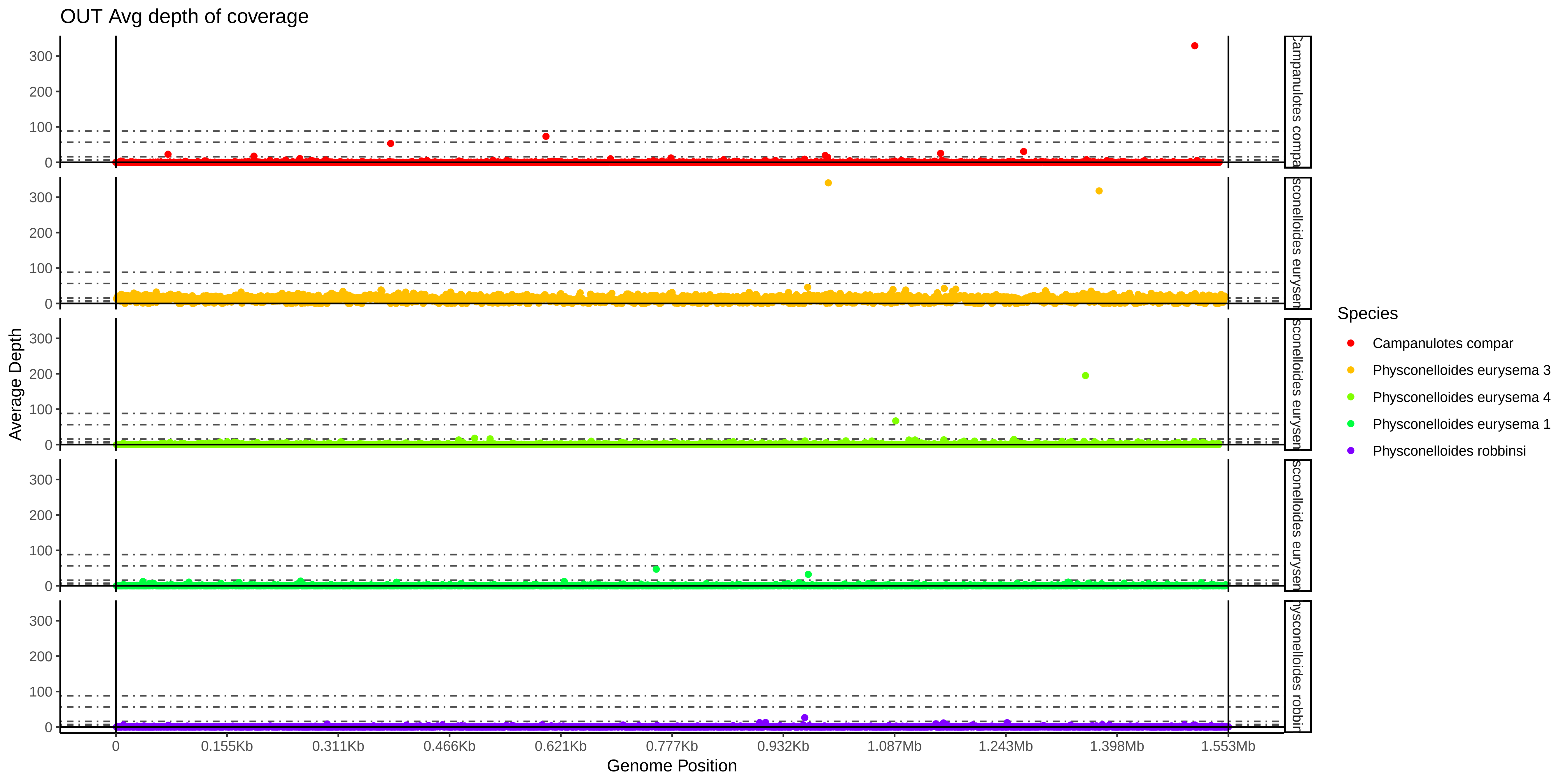

OUT Avg depth of coverage
Average Depth
Genome Position
0
100
200
300
0
0.155Kb
0.311Kb
0.466Kb
0.621Kb
0.777Kb
0.932Kb
1.087Mb
1.243Mb
1.398Mb
1.553Mb
Campanulotes compa
sconelloides eurysen
sconelloides eurysen
sconelloides eurysen
ysconelloides robbin
Species
Campanulotes compar
Physconelloides eurysema 3
Physconelloides eurysema 4
Physconelloides eurysema 1
Physconelloides robbinsi

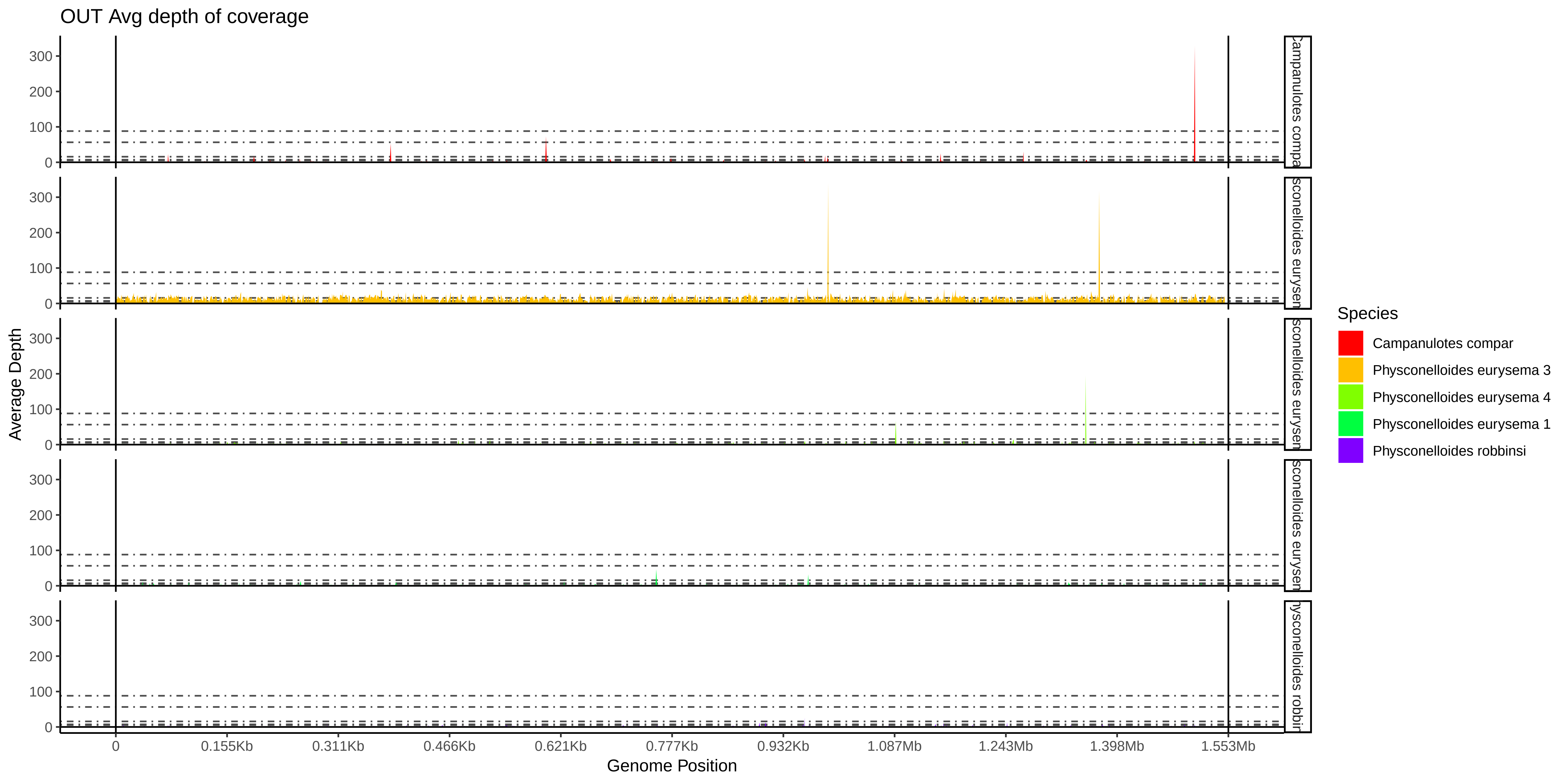
OUT Avg depth of coverage
Average Depth
Genome Position
0
100
200
300
0.155Kb
0.311Kb
0.466Kb
0.621Kb
0.777Kb
0.932Kb
1.087Mb
1.243Mb
1.398Mb
1.553Mb
Species
Campanulotes compar
Physconelloides eurysema 3
Physconelloides eurysema 4
Physconelloides eurysema 1
Physconelloides robbinsi

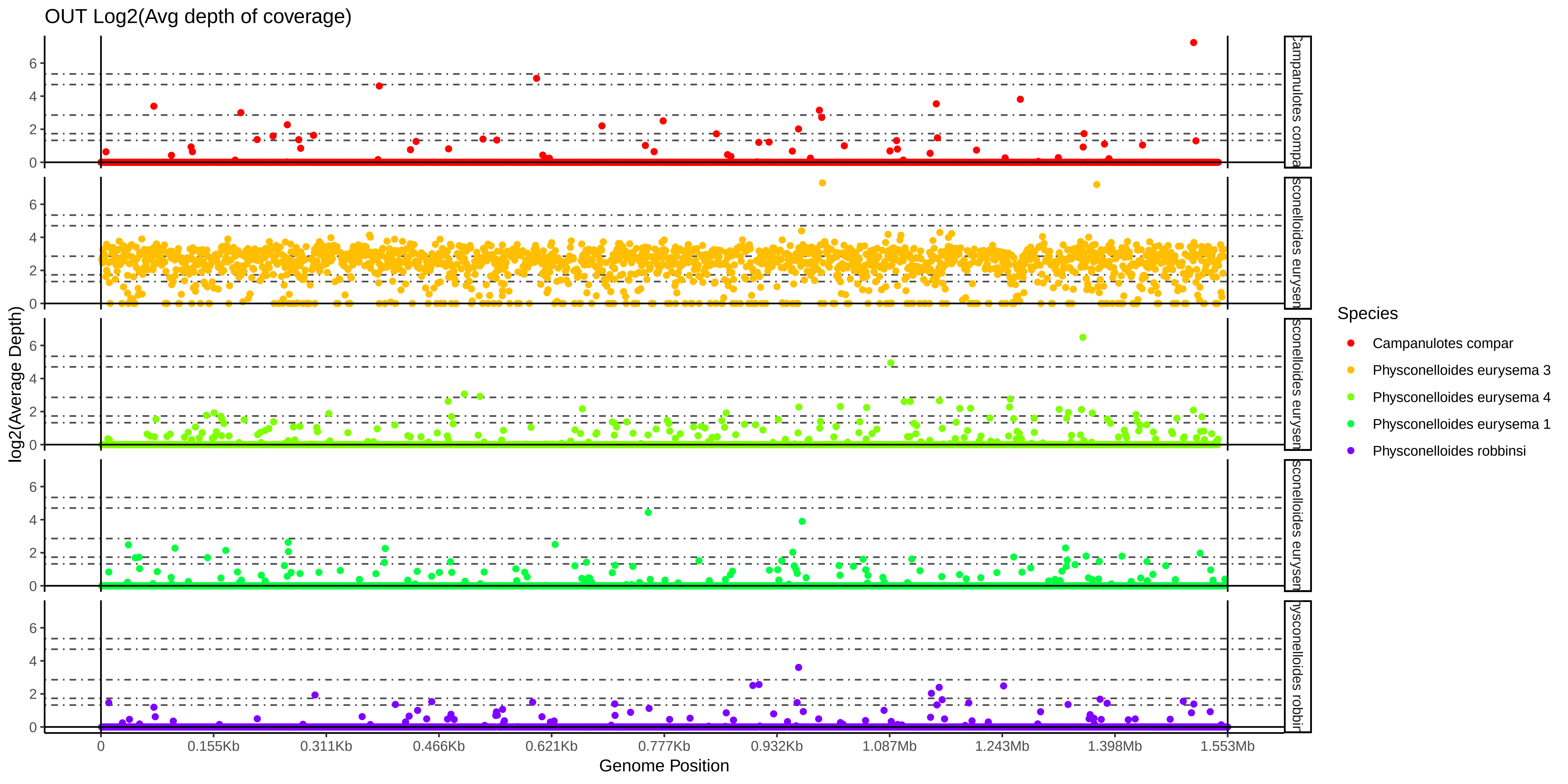
OUT Log2(Avg depth of coverage)
log2(Average Depth)
Genome Position
0
0.155Kb
0.311Kb
0.466Kb
0.621Kb
0.777Kb
0.932Kb
1.087Mb
1.243Mb
1.398Mb
1.553Mb
Species
Campanulotes compar
Physconelloides eurysema 3
Physconelloides eurysema 4
Physconelloides eurysema 1
Physconelloides robbinsi

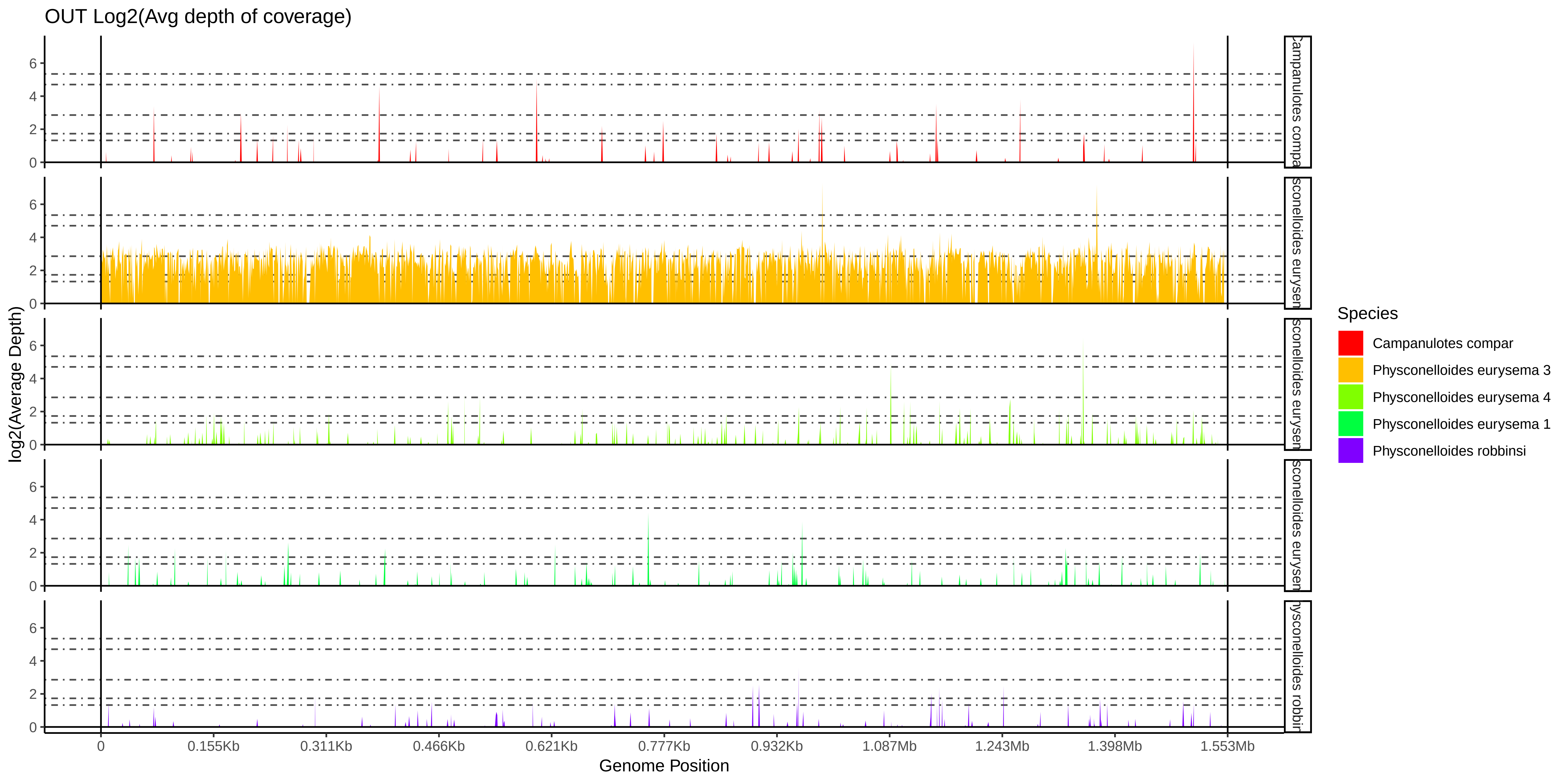

OUT Log2(Avg depth of coverage)
log2(Average Depth)
Genome Position
0
0.155Kb
0.311Kb
0.466Kb
0.621Kb
0.777Kb
0.932Kb
1.087Mb
1.243Mb
1.398Mb
1.553Mb
Campanulotes compa
sconelloides eurysen
sconelloides eurysen
sconelloides eurysen
ysconelloides robbin
Species
Campanulotes compar
Physconelloides eurysema 3
Physconelloides eurysema 4
Physconelloides eurysema 1
Physconelloides robbinsi

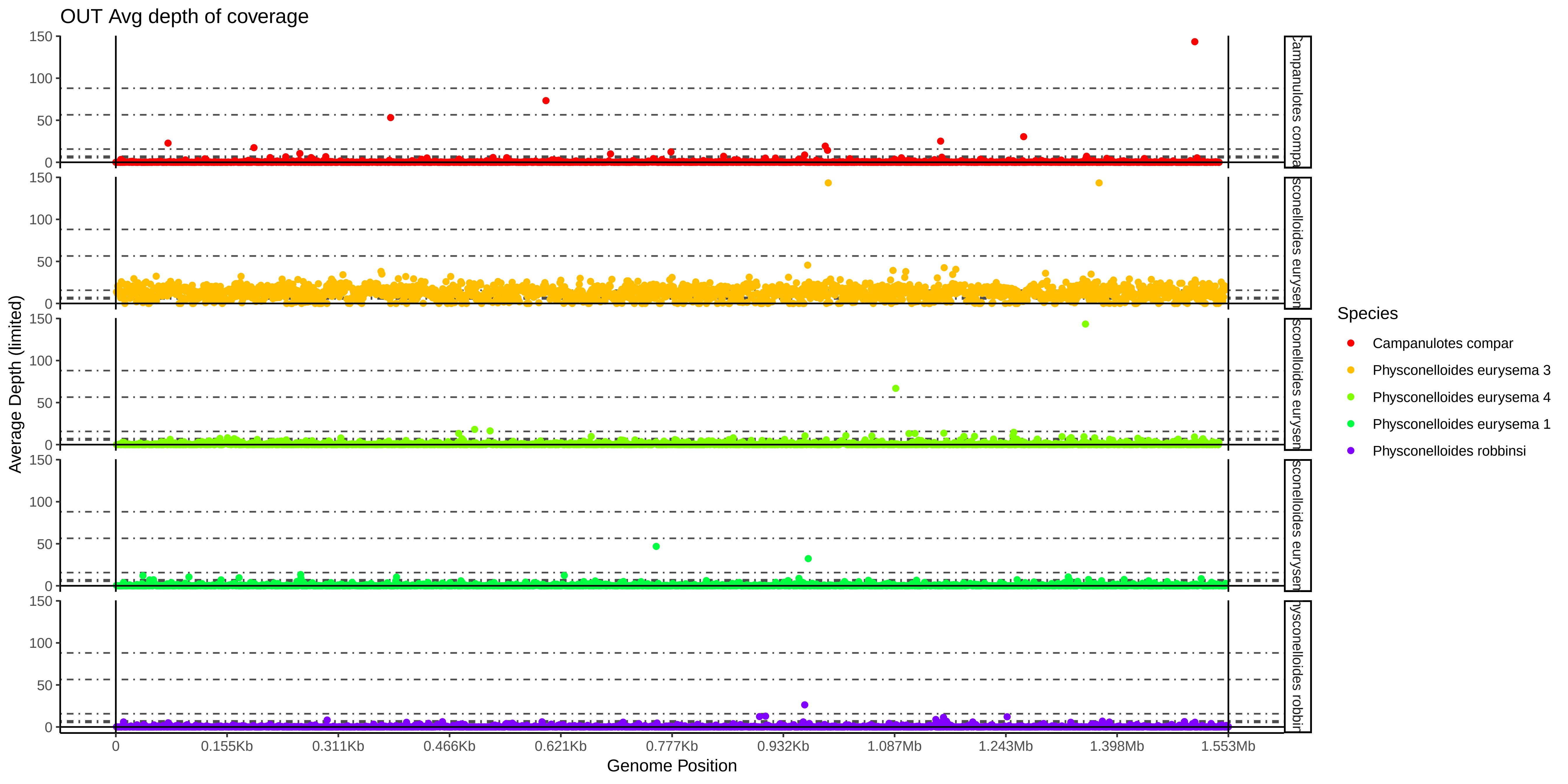
OUT Avg depth of coverage
Average Depth (limited)
150
100
50
0
0
0.155Kb
0.311Kb
0.466Kb
0.621Kb
0.777Kb
0.932Kb
1.087Mb
1.243Mb
1.398Mb
1.553Mb
Genome Position
Campanulotes compa
sconelloides eurysen
sconelloides eurysen
sconelloides eurysen
ysconelloides robbin
Species
Campanulotes compar
Physconelloides eurysema 3
Physconelloides eurysema 4
Physconelloides eurysema 1
Physconelloides robbinsi

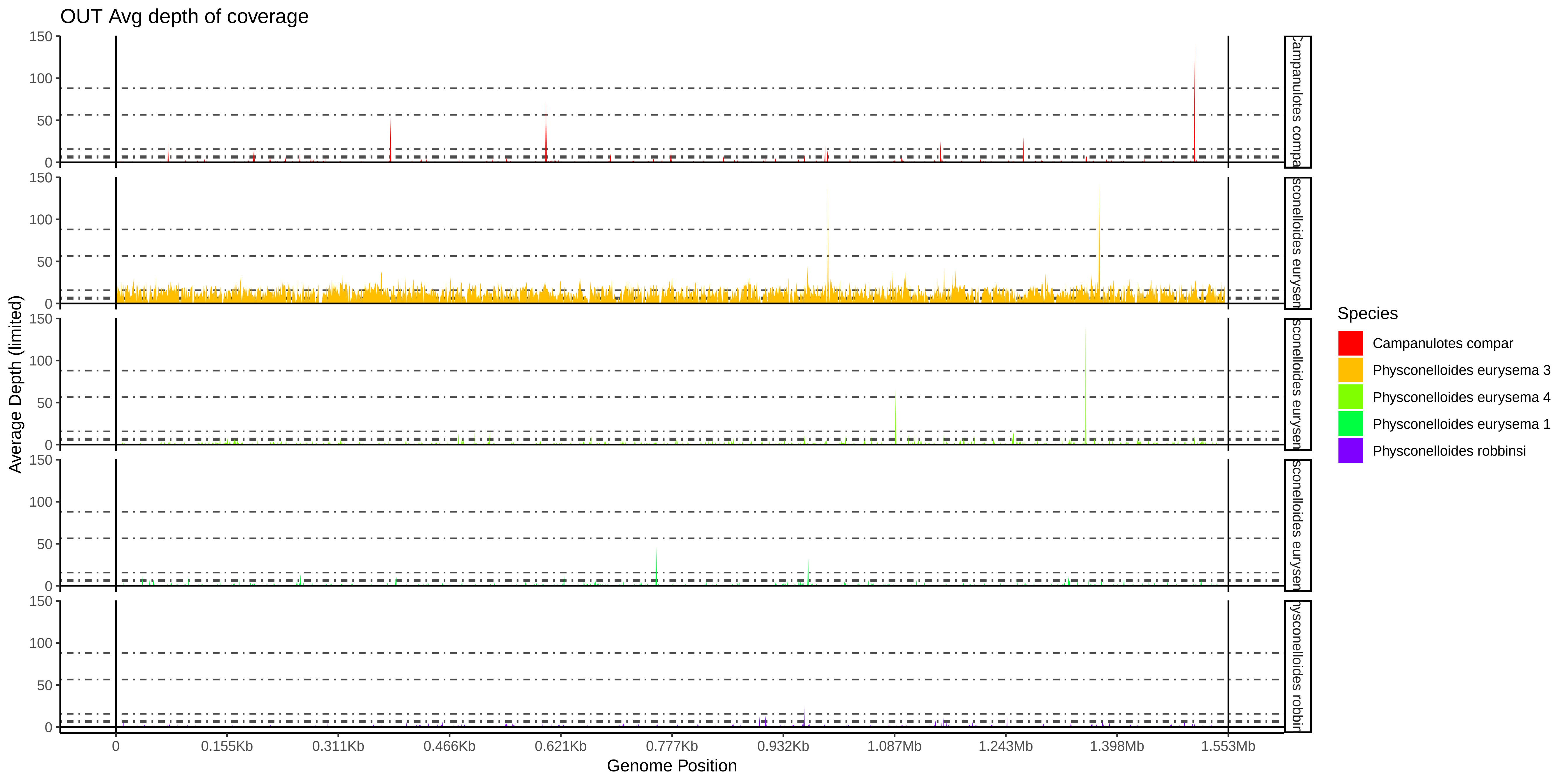

OUT Avg depth of coverage
Average Depth (limited)
Genome Position
0
0.155Kb
0.311Kb
0.466Kb
0.621Kb
0.777Kb
0.932Kb
1.087Mb
1.243Mb
1.398Mb
1.553Mb
0
50
100
150
Campanulotes compa
sconelloides eurysen
sconelloides eurysen
sconelloides eurysen
ysconelloides robbin
Species
Campanulotes compar
Physconelloides eurysema 3
Physconelloides eurysema 4
Physconelloides eurysema 1
Physconelloides robbinsi
