## Supplementary material for "Comparing rates of introgression in parasitic feather lice with differing dispersal capabilities": OUT_sppIDerDepthPlot-covBins_Pheur2_Cocru_3_16_2016_9.pdf (from another version of this preprint: version 7, 2020-03-31)

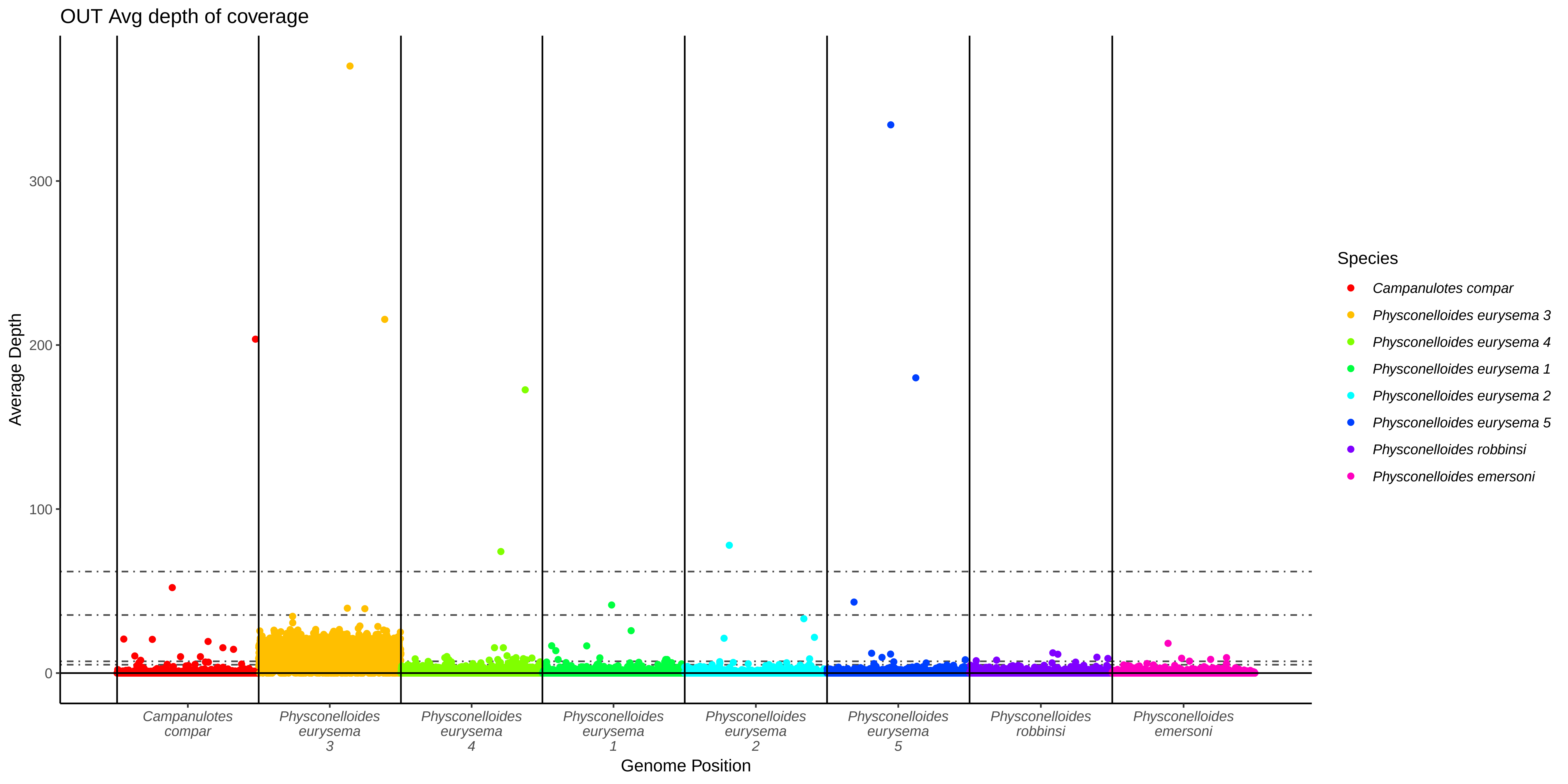

OUT Avg depth of coverage
Average Depth
300
200
100
0
Campanulotes compar
Physconelloides eurysema 3
Physconelloides eurysema 4
Physconelloides eurysema 1
Physconelloides eurysema 2
Physconelloides eurysema 5
Physconelloides robbinsi
Physconelloides emersoni
Genome Position
Species
Campanulotes compar
Physconelloides eurysema 3
Physconelloides eurysema 4
Physconelloides eurysema 1
Physconelloides eurysema 2
Physconelloides eurysema 5
Physconelloides robbinsi
Physconelloides emersoni

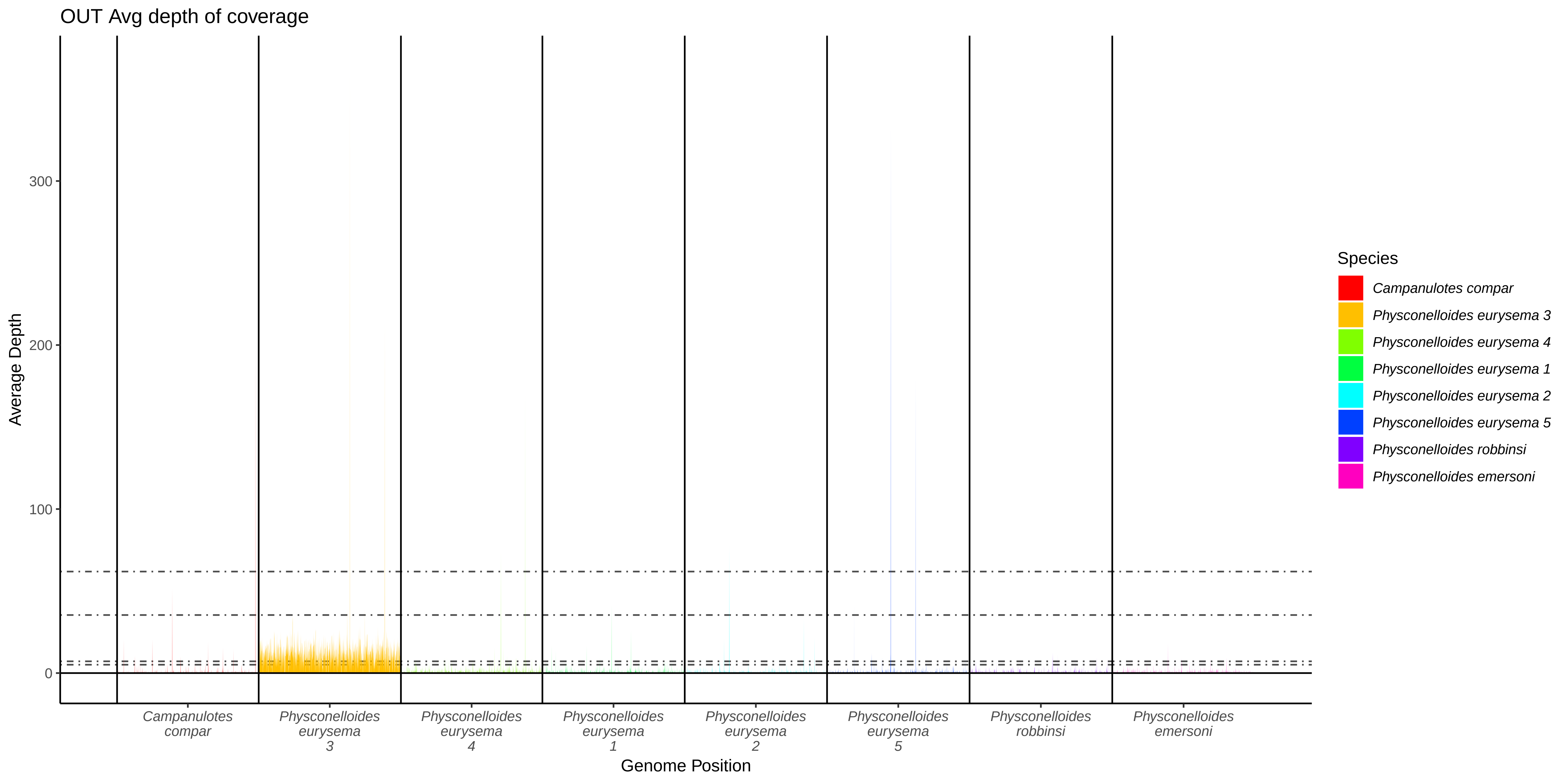
OUT Avg depth of coverage
Average Depth
300
200
100
0
Campanulotes compar
Physconelloides eurysema 3
Physconelloides eurysema 4
Physconelloides eurysema 1
Physconelloides eurysema 2
Physconelloides eurysema 5
Physconelloides robbinsi
Physconelloides emersoni
Genome Position
Species
Campanulotes compar
Physconelloides eurysema 3
Physconelloides eurysema 4
Physconelloides eurysema 1
Physconelloides eurysema 2
Physconelloides eurysema 5
Physconelloides robbinsi
Physconelloides emersoni

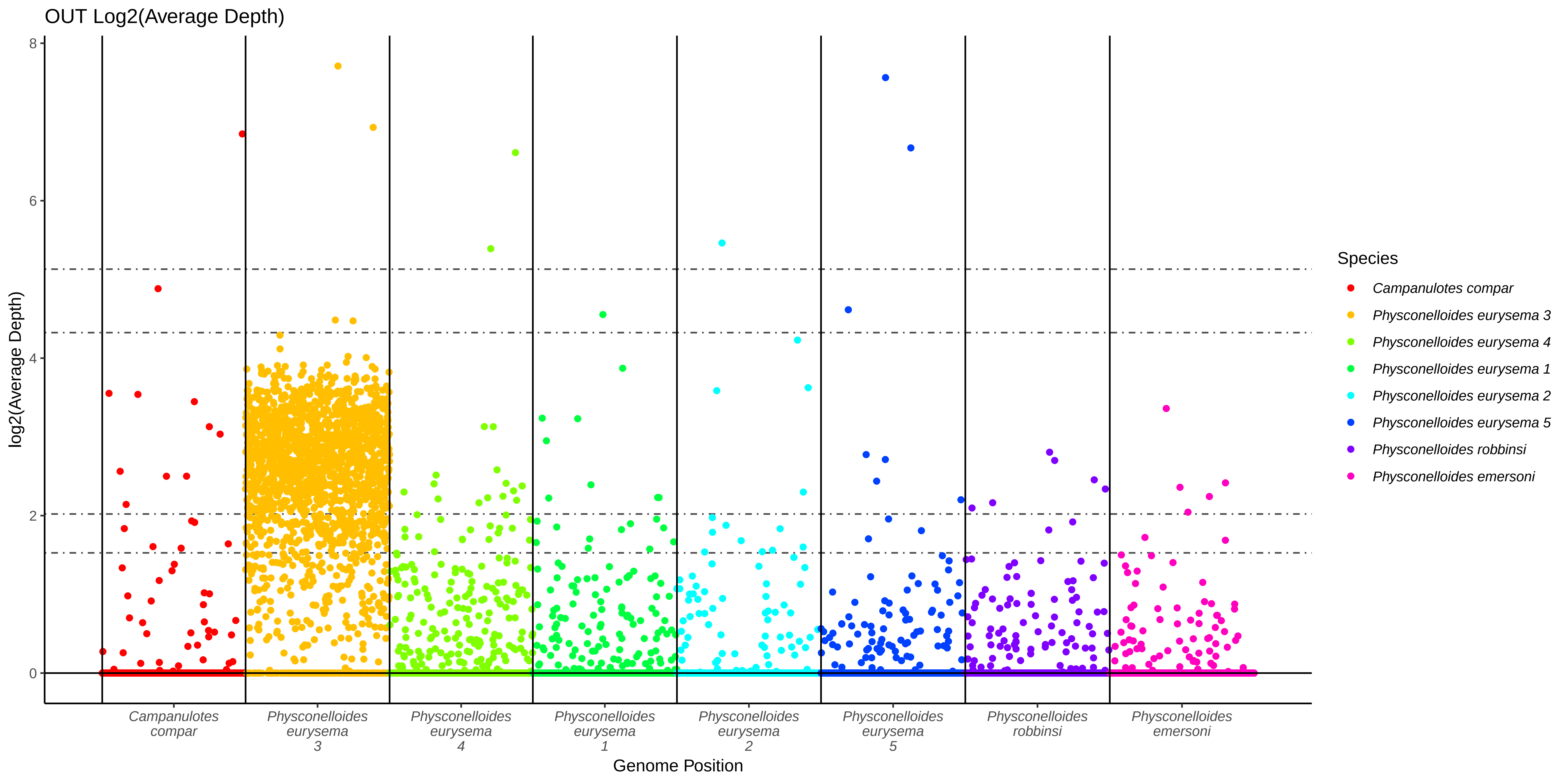

OUT Log2(Average Depth)
log2(Average Depth)
0
2
4
6
8
Campanulotes compar
Physconelloides eurysema 3
Physconelloides eurysema 4
Physconelloides eurysema 1
Physconelloides eurysema 2
Physconelloides eurysema 5
Physconelloides robbinsi
Physconelloides emersoni
Genome Position
Species
Campanulotes compar
Physconelloides eurysema 3
Physconelloides eurysema 4
Physconelloides eurysema 1
Physconelloides eurysema 2
Physconelloides eurysema 5
Physconelloides robbinsi
Physconelloides emersoni

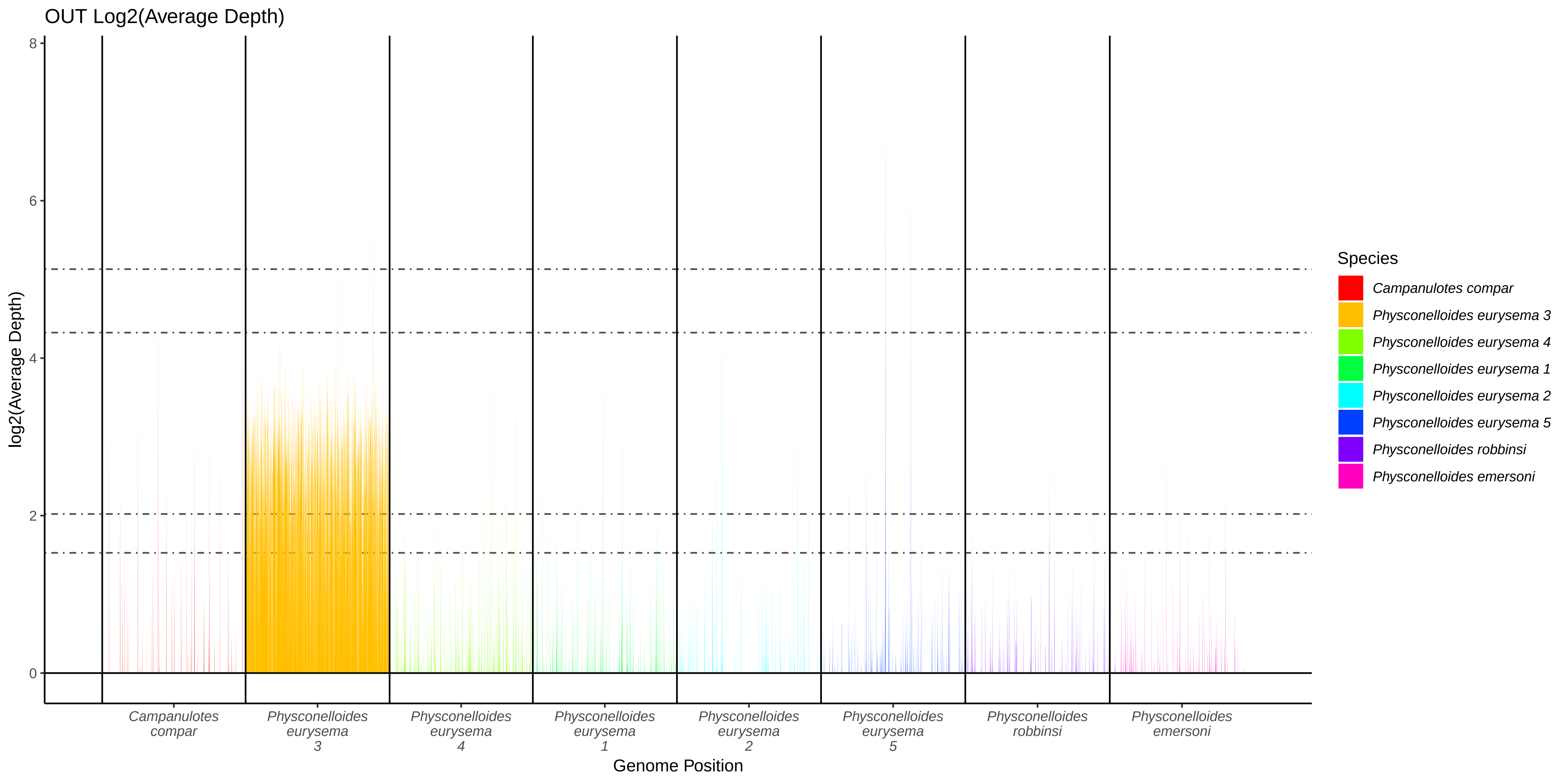
OUT Log2(Average Depth)
log2(Average Depth)
8
6
4
2
0
Campanulotes compar
Physconelloides eurysema 3
Physconelloides eurysema 4
Physconelloides eurysema 1
Physconelloides eurysema 2
Physconelloides eurysema 5
Physconelloides robbinsi
Physconelloides emersoni
Genome Position
Species
Campanulotes compar
Physconelloides eurysema 3
Physconelloides eurysema 4
Physconelloides eurysema 1
Physconelloides eurysema 2
Physconelloides eurysema 5
Physconelloides robbinsi
Physconelloides emersoni

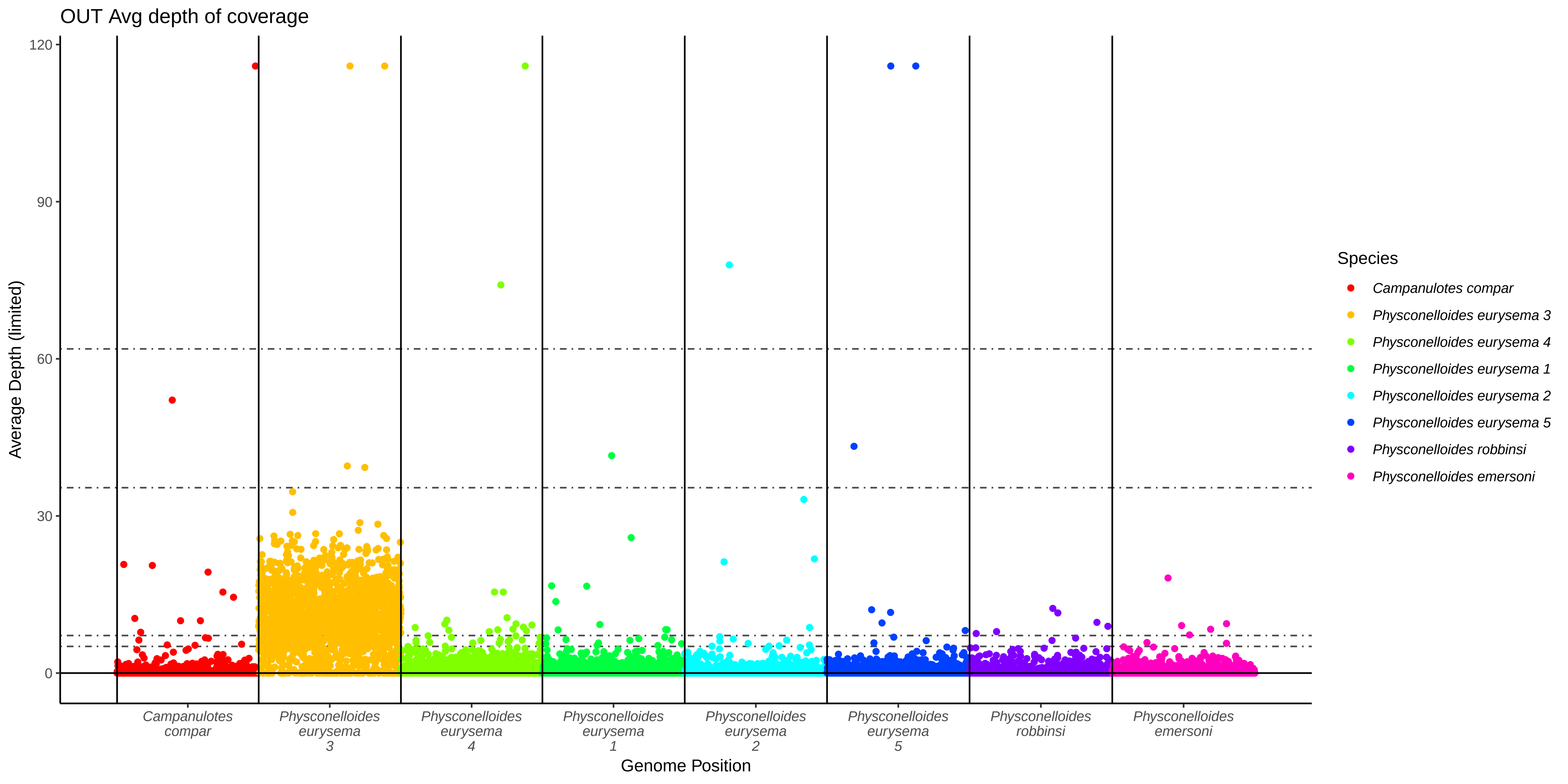

OUT Avg depth of coverage
Average Depth (limited)
120
90
60
30
0
Campanulotes compar
Physconelloides eurysema 3
Physconelloides eurysema 4
Physconelloides eurysema 1
Physconelloides eurysema 2
Physconelloides eurysema 5
Physconelloides robbinsi
Physconelloides emersoni
Genome Position
Species
Campanulotes compar
Physconelloides eurysema 3
Physconelloides eurysema 4
Physconelloides eurysema 1
Physconelloides eurysema 2
Physconelloides eurysema 5
Physconelloides robbinsi
Physconelloides emersoni

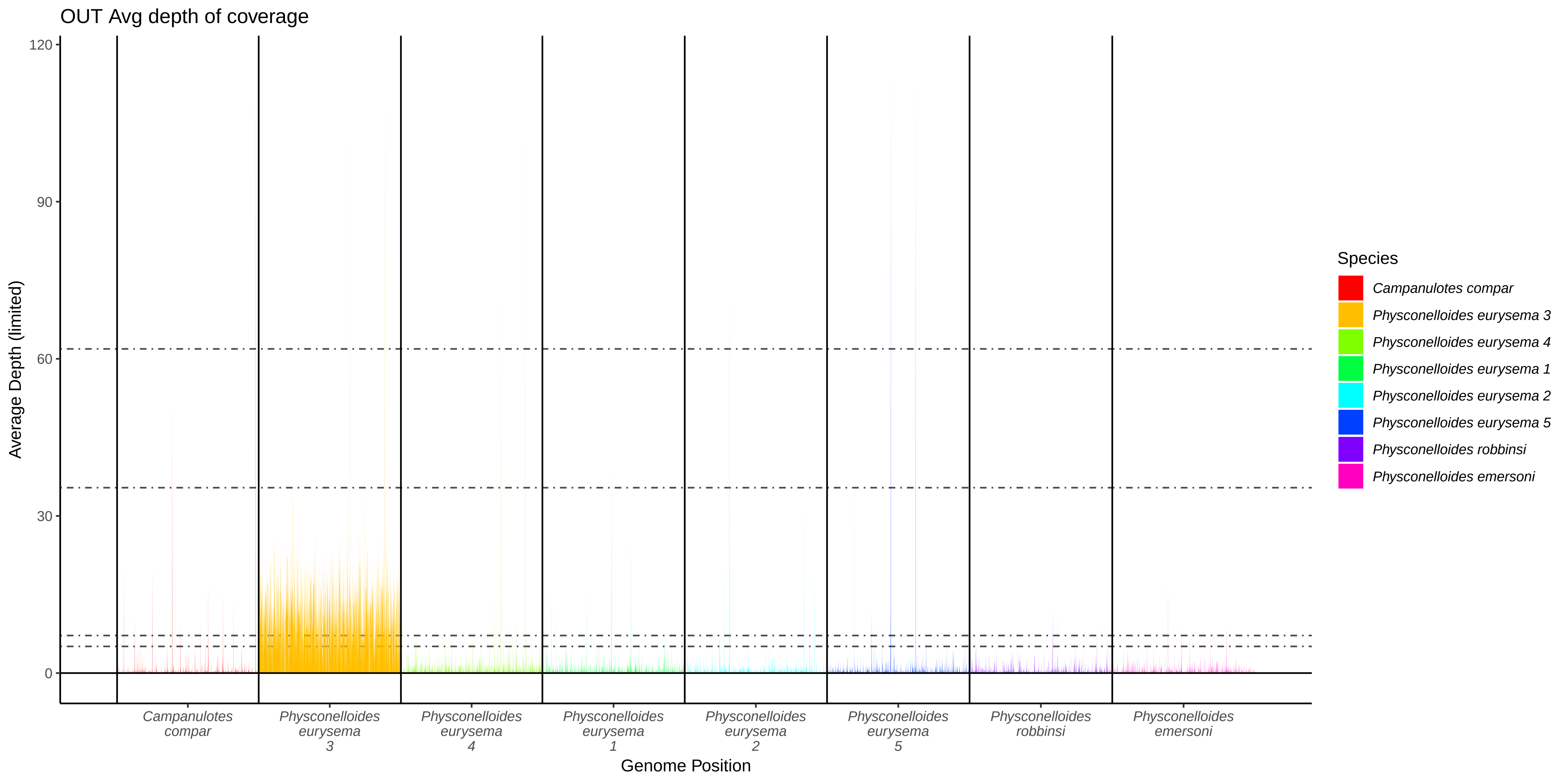

OUT Avg depth of coverage
Average Depth (limited)
120
90
60
30
0
Campanulotes compar
Physconelloides eurysema 3
Physconelloides eurysema 4
Physconelloides eurysema 1
Physconelloides eurysema 2
Physconelloides eurysema 5
Physconelloides robbinsi
Physconelloides emersoni
Genome Position
Species
Campanulotes compar
Physconelloides eurysema 3
Physconelloides eurysema 4
Physconelloides eurysema 1
Physconelloides eurysema 2
Physconelloides eurysema 5
Physconelloides robbinsi
Physconelloides emersoni

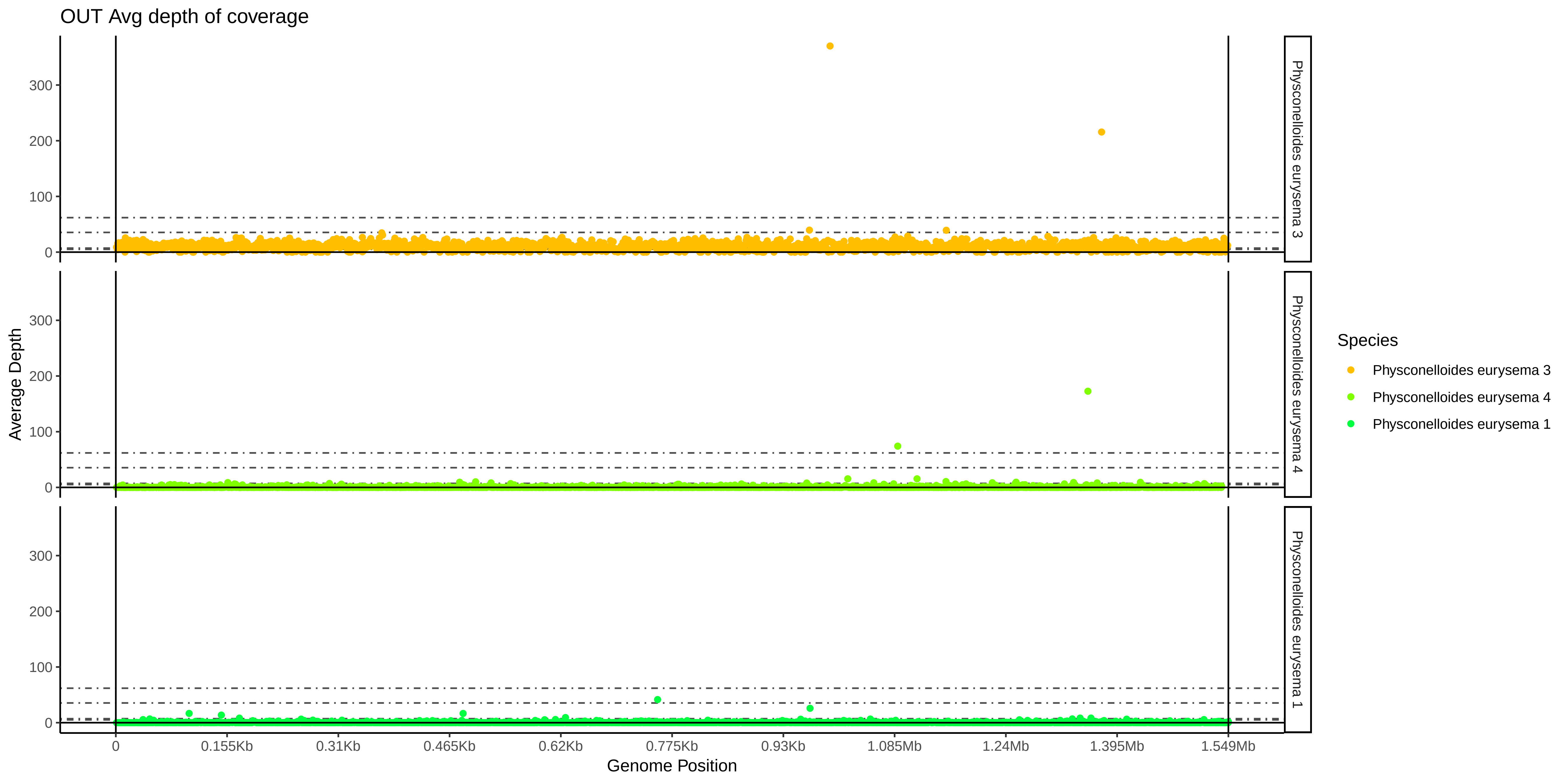

OUT Avg depth of coverage
Average Depth
300
200
100
0
Physconelloides eurysema 3
Physconelloides eurysema 4
Physconelloides eurysema 1
0
0.155Kb
0.31Kb
0.465Kb
0.62Kb
0.775Kb
0.93Kb
1.085Mb
1.24Mb
1.395Mb
1.549Mb
Genome Position
Species
Physconelloides eurysema 3
Physconelloides eurysema 4
Physconelloides eurysema 1

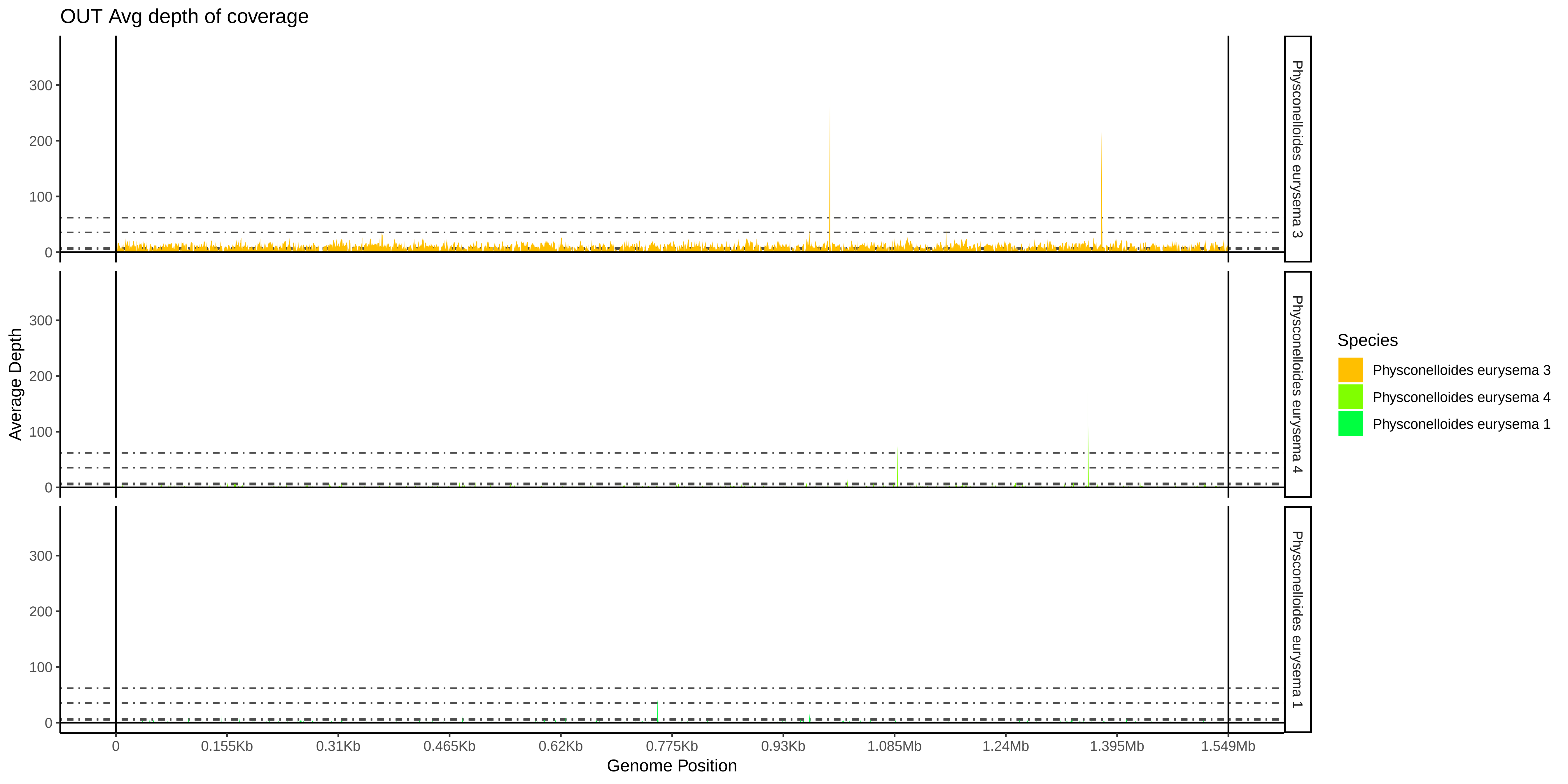

OUT Avg depth of coverage
Average Depth
0
100
200
300
Physconelloides eurysema 3
Physconelloides eurysema 4
Physconelloides eurysema 1
0
0.155Kb
0.31Kb
0.465Kb
0.62Kb
0.775Kb
0.93Kb
1.085Mb
1.24Mb
1.395Mb
1.549Mb
Genome Position
Species
Physconelloides eurysema 3
Physconelloides eurysema 4
Physconelloides eurysema 1

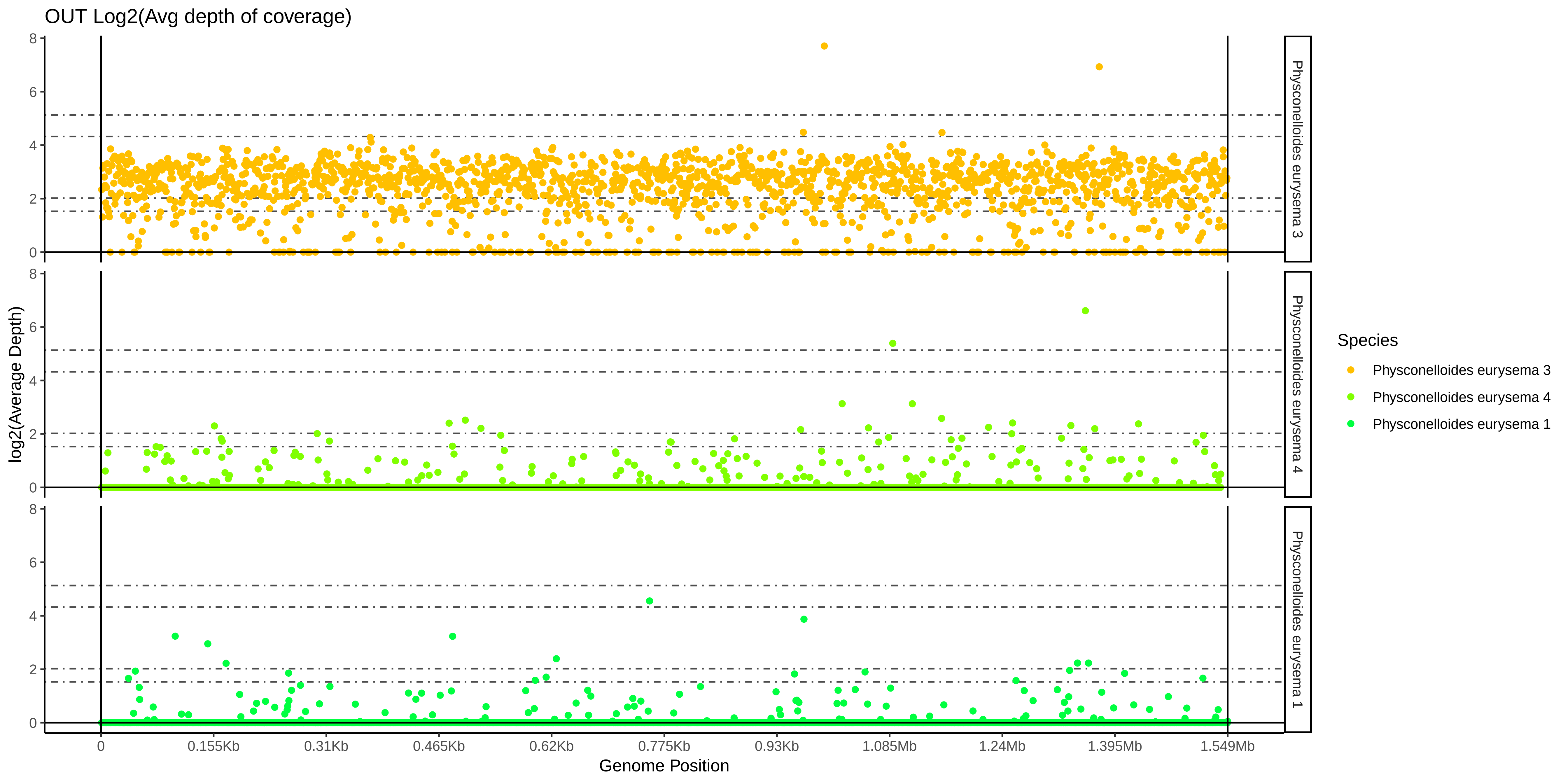

OUT Log2(Avg depth of coverage)
log2(Average Depth)
Genome Position
0
0.155Kb
0.31Kb
0.465Kb
0.62Kb
0.775Kb
0.93Kb
1.085Mb
1.24Mb
1.395Mb
1.549Mb
Physconelloides eurysema 3
Physconelloides eurysema 4
Physconelloides eurysema 1
Species
Physconelloides eurysema 3
Physconelloides eurysema 4
Physconelloides eurysema 1

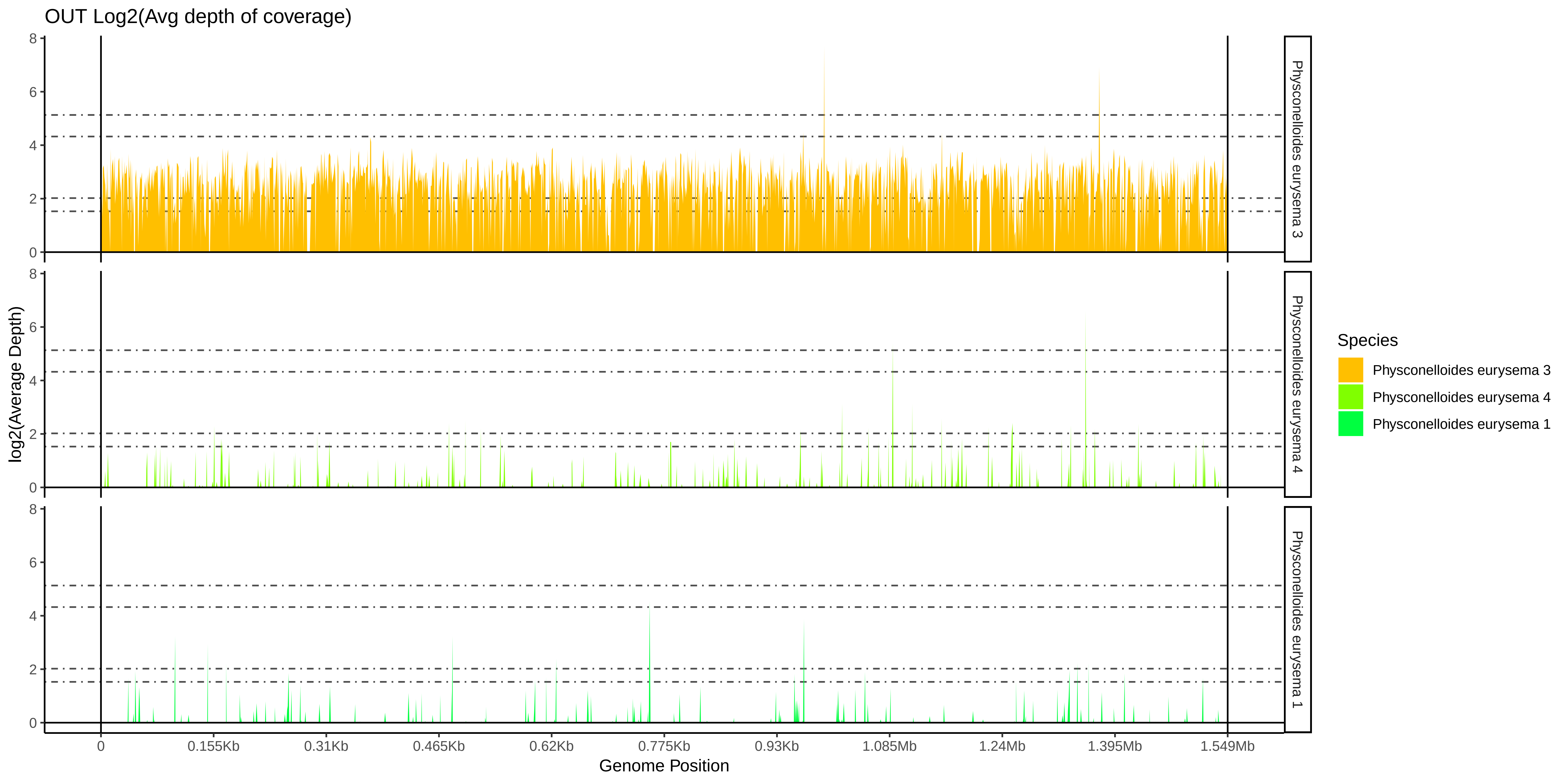

OUT Log2(Avg depth of coverage)
log2(Average Depth)
0
2
4
6
8
Physconelloides eurysema 3
Physconelloides eurysema 4
Physconelloides eurysema 1
0
0.155Kb
0.31Kb
0.465Kb
0.62Kb
0.775Kb
0.93Kb
1.085Mb
1.24Mb
1.395Mb
1.549Mb
Genome Position
Species
Physconelloides eurysema 3
Physconelloides eurysema 4
Physconelloides eurysema 1

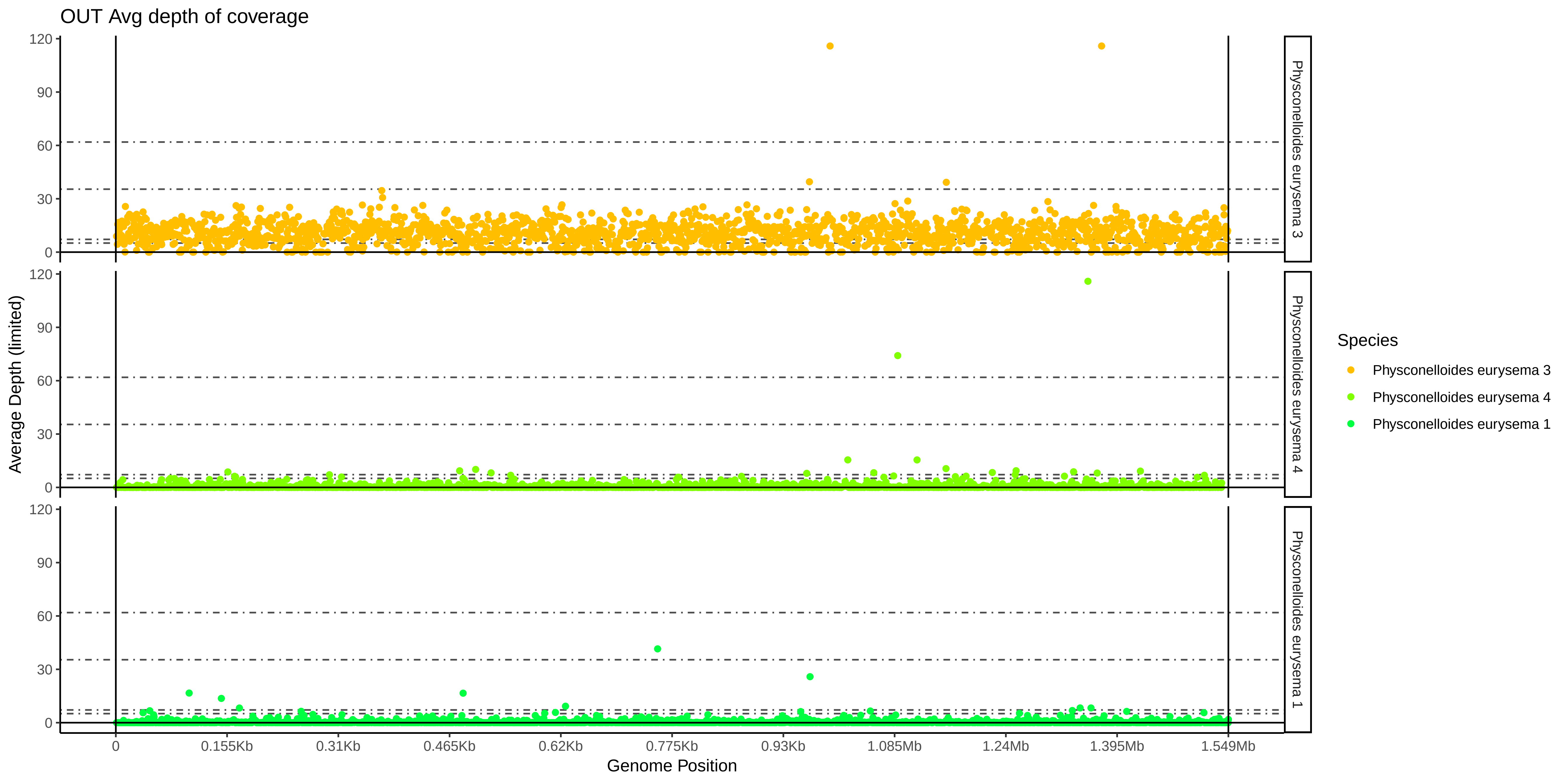
OUT Avg depth of coverage
Average Depth (limited)
Genome Position
0
0.155Kb
0.31Kb
0.465Kb
0.62Kb
0.775Kb
0.93Kb
1.085Mb
1.24Mb
1.395Mb
1.549Mb
0
30
60
90
120
Physconelloides eurysema 3
Physconelloides eurysema 4
Physconelloides eurysema 1
Species
Physconelloides eurysema 3
Physconelloides eurysema 4
Physconelloides eurysema 1

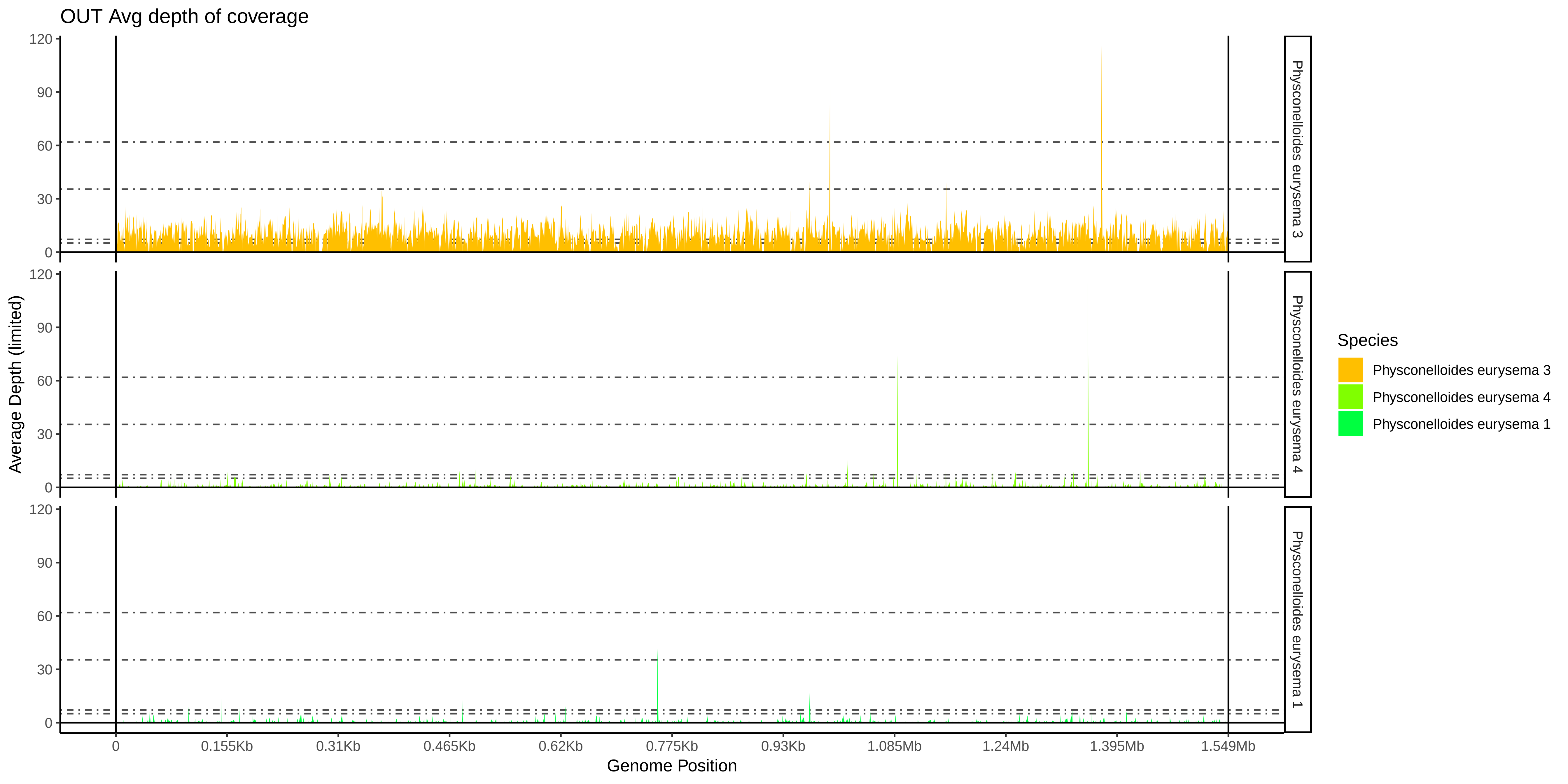

OUT Avg depth of coverage
Average Depth (limited)
120
90
60
30
0
Physconelloides eurysema 3
Physconelloides eurysema 4
Physconelloides eurysema 1
0
0.155Kb
0.31Kb
0.465Kb
0.62Kb
0.775Kb
0.93Kb
1.085Mb
1.24Mb
1.395Mb
1.549Mb
Genome Position
Species
Physconelloides eurysema 3
Physconelloides eurysema 4
Physconelloides eurysema 1
