## Supplementary material for "Comparing rates of introgression in parasitic feather lice with differing dispersal capabilities": OUT_sppIDerDepthPlot-covBins_Pheur_Copas_3_16_2016_7.pdf (from another version of this preprint: version 7, 2020-03-31)

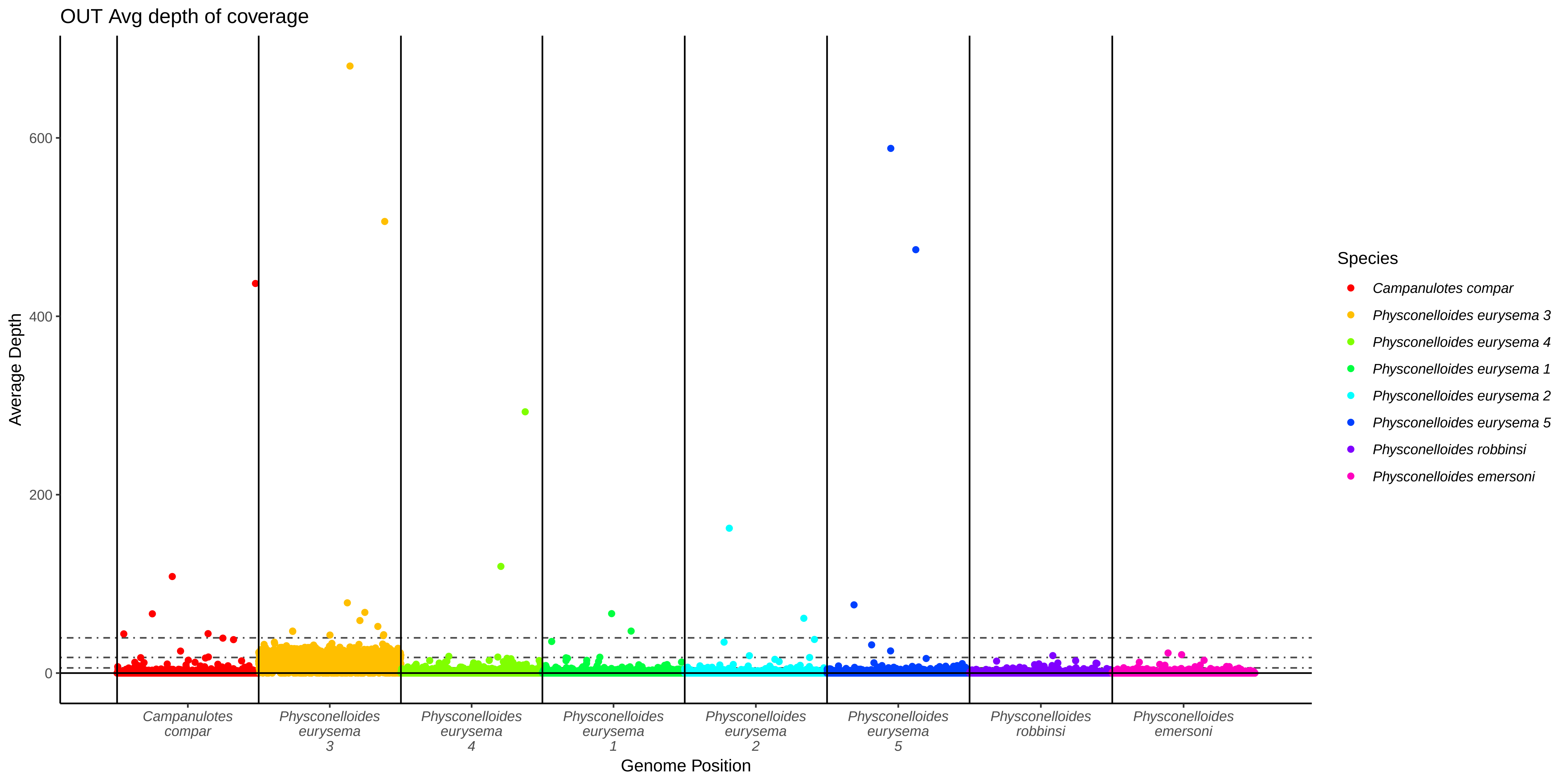
OUT Avg depth of coverage
Average Depth
600
400
200
0
Campanulotes compar
Physconelloides eurysema 3
Physconelloides eurysema 4
Physconelloides eurysema 1
Physconelloides eurysema 2
Physconelloides eurysema 5
Physconelloides robbinsi
Physconelloides emersoni
Genome Position
Species
Campanulotes compar
Physconelloides eurysema 3
Physconelloides eurysema 4
Physconelloides eurysema 1
Physconelloides eurysema 2
Physconelloides eurysema 5
Physconelloides robbinsi
Physconelloides emersoni

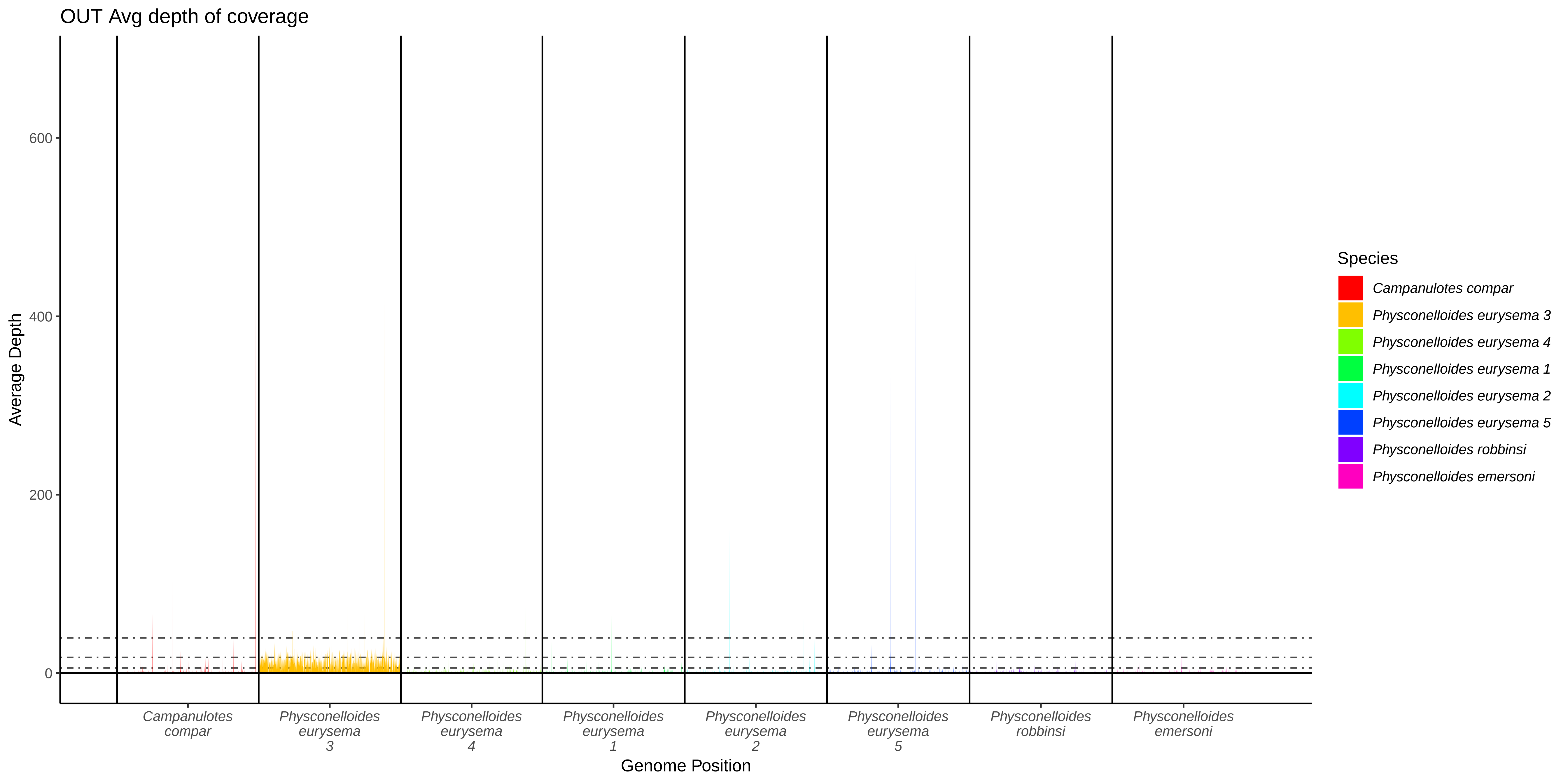

OUT Avg depth of coverage
Average Depth
600
400
200
0
Campanulotes compar
Physconelloides eurysema 3
Physconelloides eurysema 4
Physconelloides eurysema 1
Physconelloides eurysema 2
Physconelloides eurysema 5
Physconelloides robbinsi
Physconelloides emersoni
Genome Position
Species
Campanulotes compar
Physconelloides eurysema 3
Physconelloides eurysema 4
Physconelloides eurysema 1
Physconelloides eurysema 2
Physconelloides eurysema 5
Physconelloides robbinsi
Physconelloides emersoni

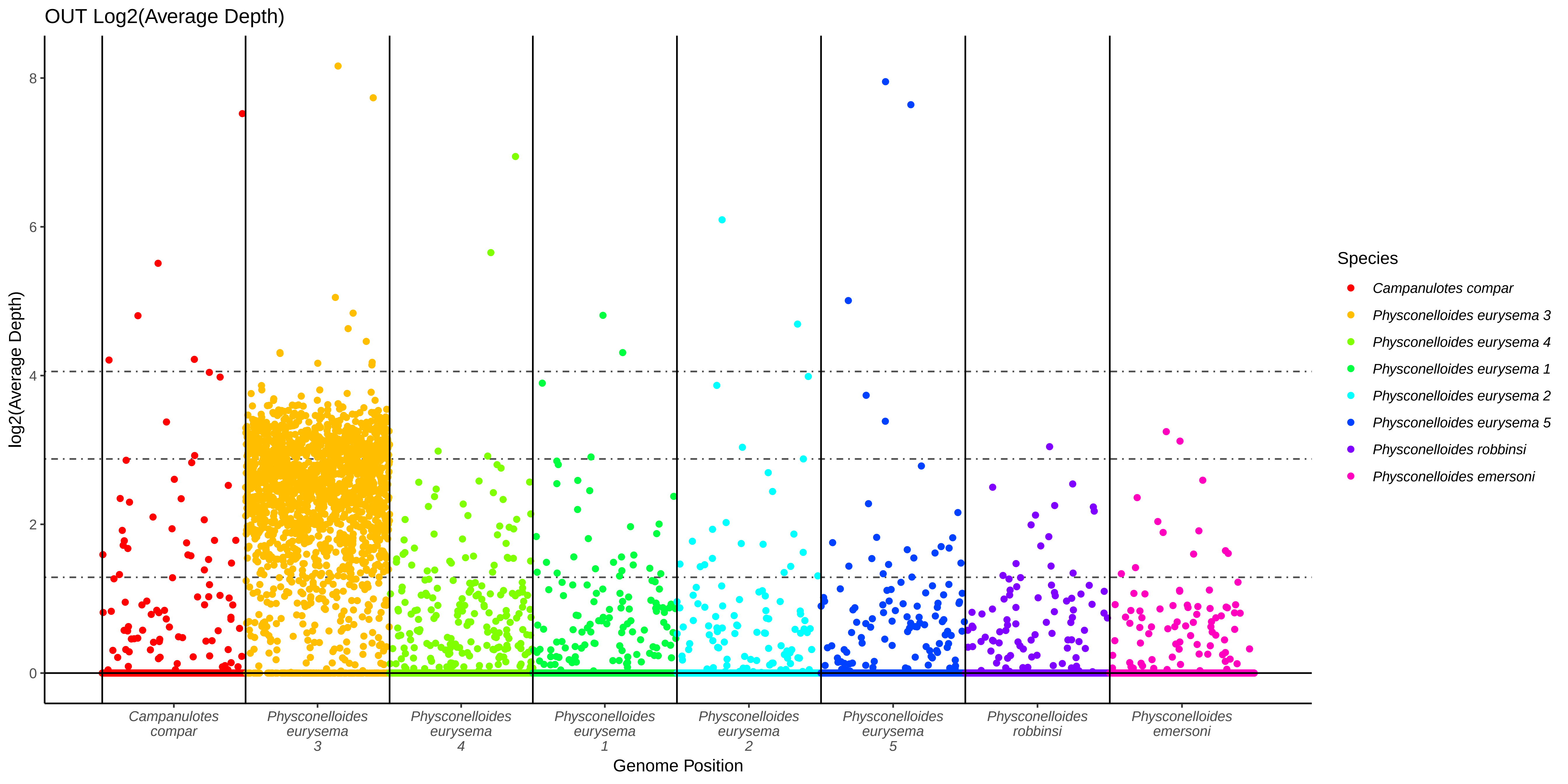
OUT Log2(Average Depth)
log2(Average Depth)
0
2
4
6
8
Genome Position
Campanulotes compar
Physconelloides eurysema 3
Physconelloides eurysema 4
Physconelloides eurysema 1
Physconelloides eurysema 2
Physconelloides eurysema 5
Physconelloides robbinsi
Physconelloides emersoni
Species
Campanulotes compar
Physconelloides eurysema 3
Physconelloides eurysema 4
Physconelloides eurysema 1
Physconelloides eurysema 2
Physconelloides eurysema 5
Physconelloides robbinsi
Physconelloides emersoni

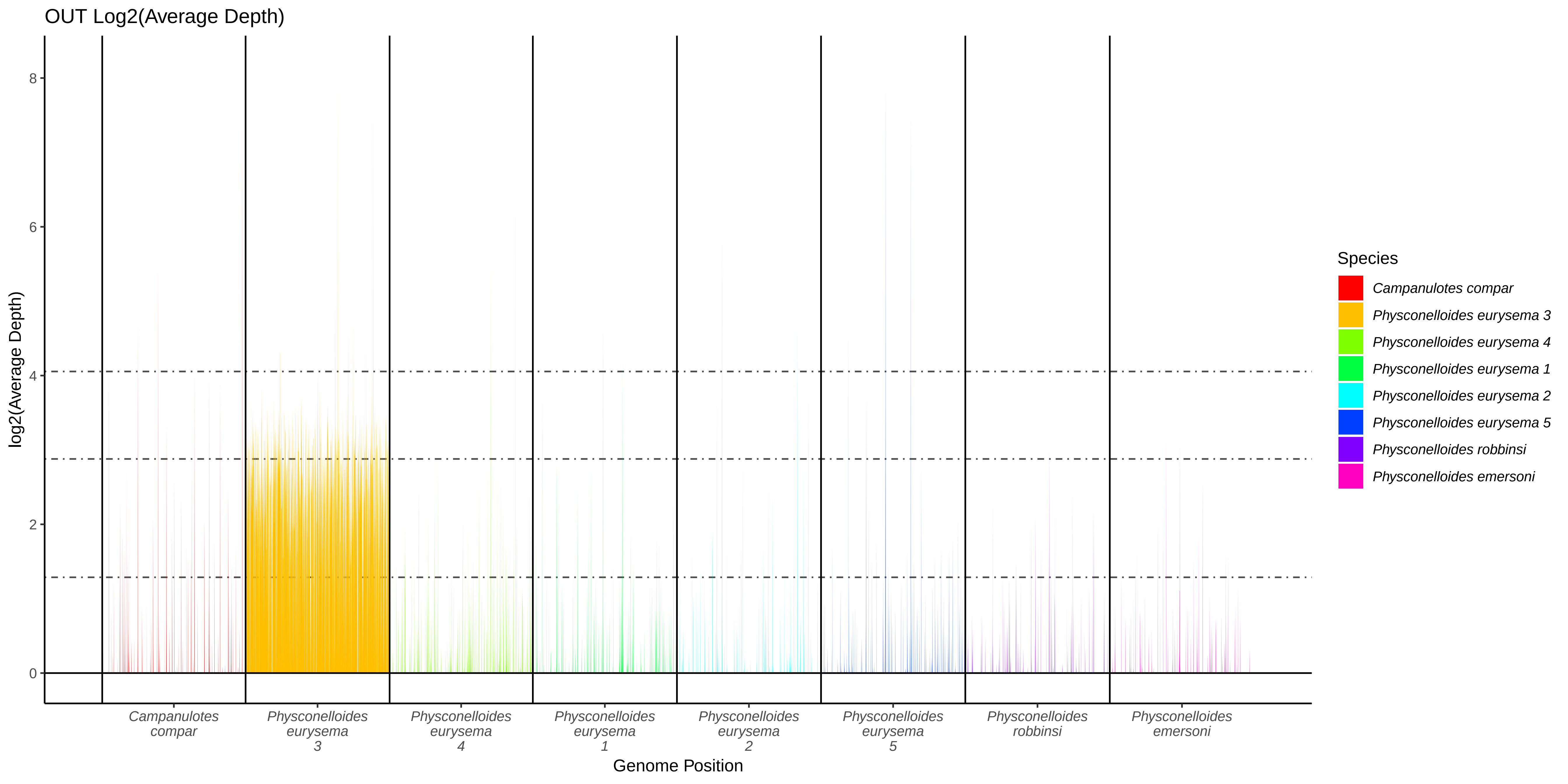

OUT Log2(Average Depth)
log2(Average Depth)
0
2
4
6
8
Genome Position
Campanulotes compar
Physconelloides eurysema 3
Physconelloides eurysema 4
Physconelloides eurysema 1
Physconelloides eurysema 2
Physconelloides eurysema 5
Physconelloides robbinsi
Physconelloides emersoni
Species
Campanulotes compar
Physconelloides eurysema 3
Physconelloides eurysema 4
Physconelloides eurysema 1
Physconelloides eurysema 2
Physconelloides eurysema 5
Physconelloides robbinsi
Physconelloides emersoni

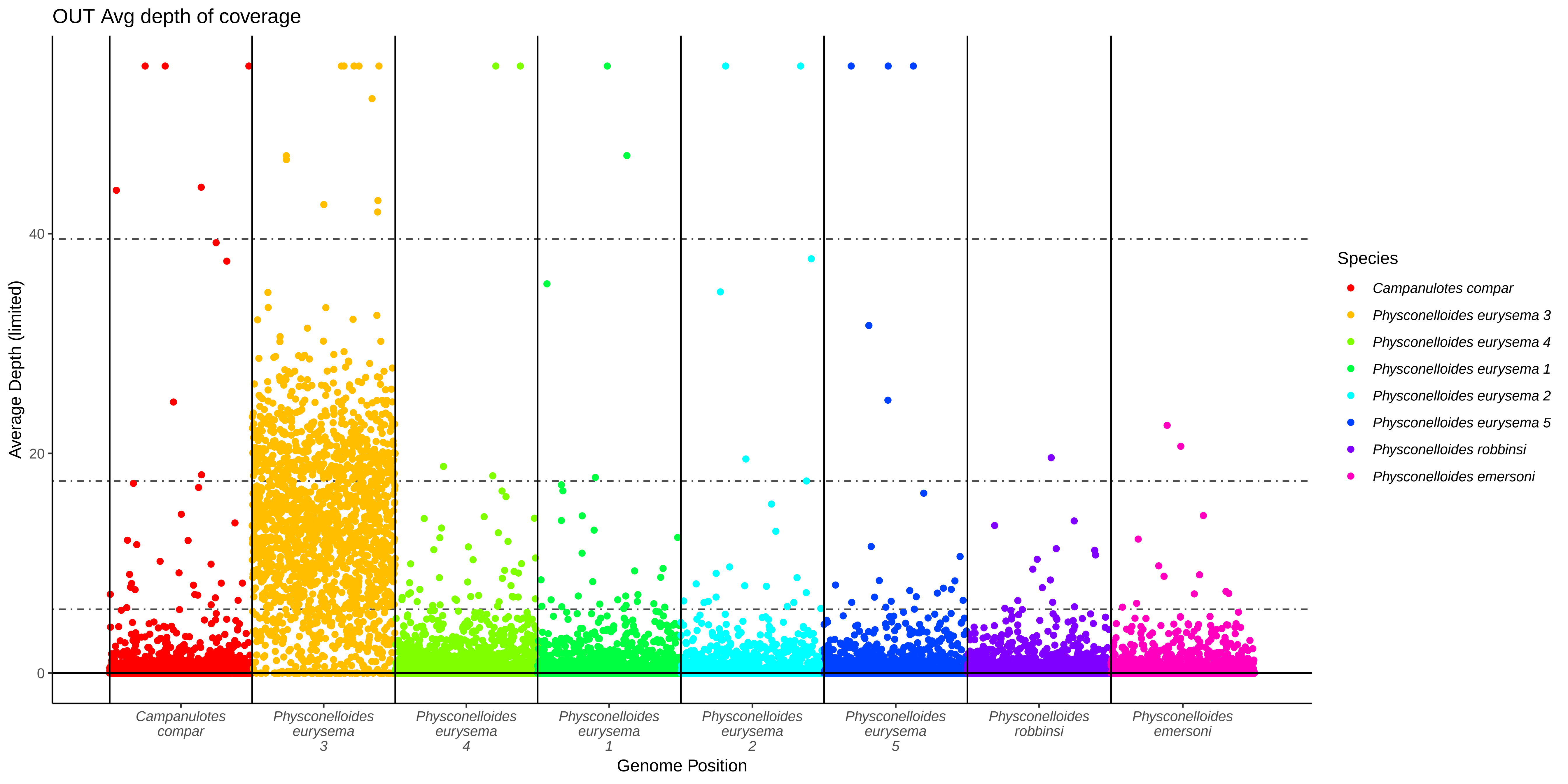
OUT Avg depth of coverage
Average Depth (limited)
0
20
40
Genome Position
Campanulotes compar
Physconelloides eurysema 3
Physconelloides eurysema 4
Physconelloides eurysema 1
Physconelloides eurysema 2
Physconelloides eurysema 5
Physconelloides robbinsi
Physconelloides emersoni
Species
Campanulotes compar
Physconelloides eurysema 3
Physconelloides eurysema 4
Physconelloides eurysema 1
Physconelloides eurysema 2
Physconelloides eurysema 5
Physconelloides robbinsi
Physconelloides emersoni

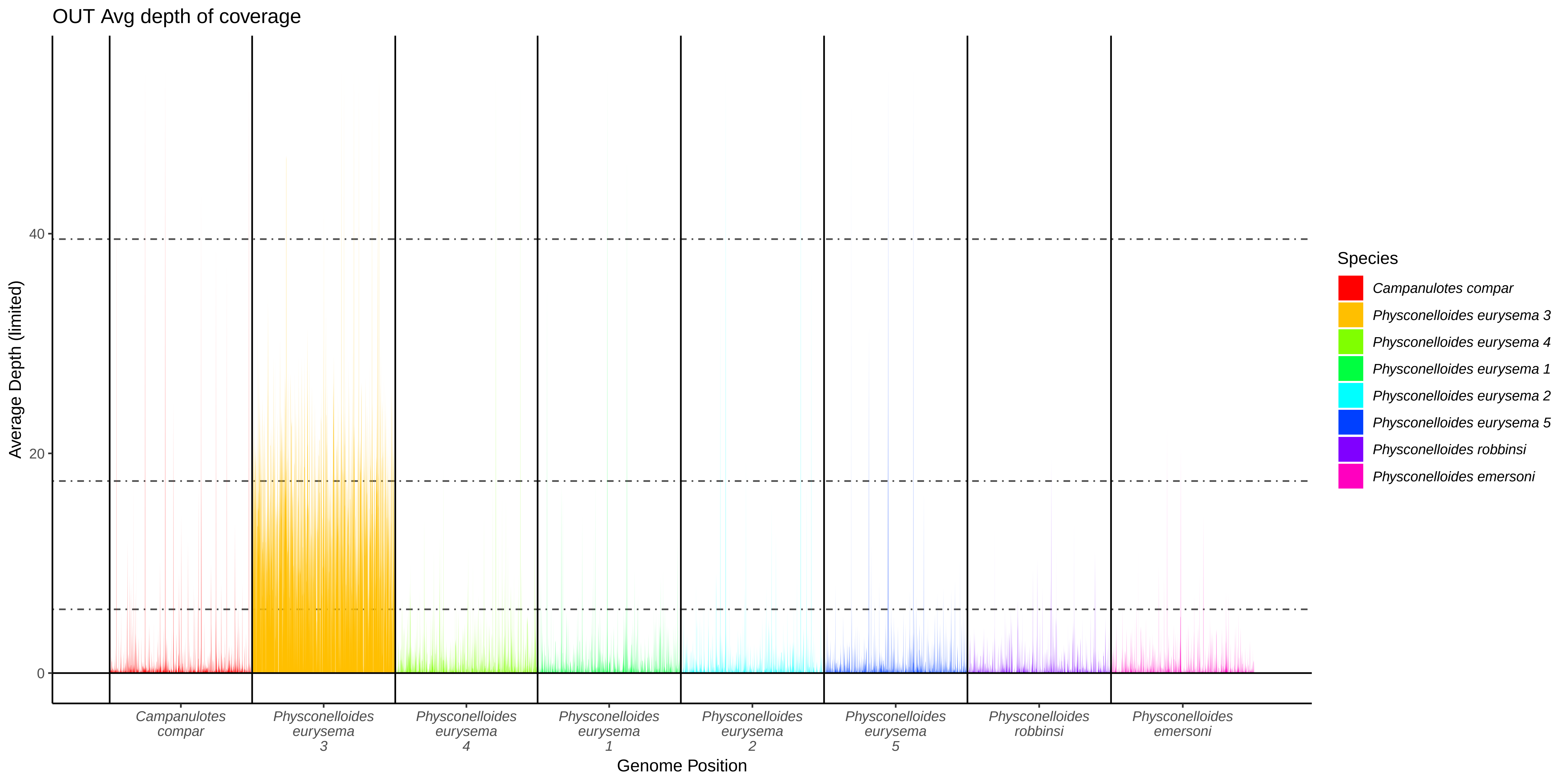
OUT Avg depth of coverage
Average Depth (limited)
40
20
0
Campanulotes compar
Physconelloides eurysema 3
Physconelloides eurysema 4
Physconelloides eurysema 1
Physconelloides eurysema 2
Physconelloides eurysema 5
Physconelloides robbinsi
Physconelloides emersoni
Genome Position
Species
Campanulotes compar
Physconelloides eurysema 3
Physconelloides eurysema 4
Physconelloides eurysema 1
Physconelloides eurysema 2
Physconelloides eurysema 5
Physconelloides robbinsi
Physconelloides emersoni

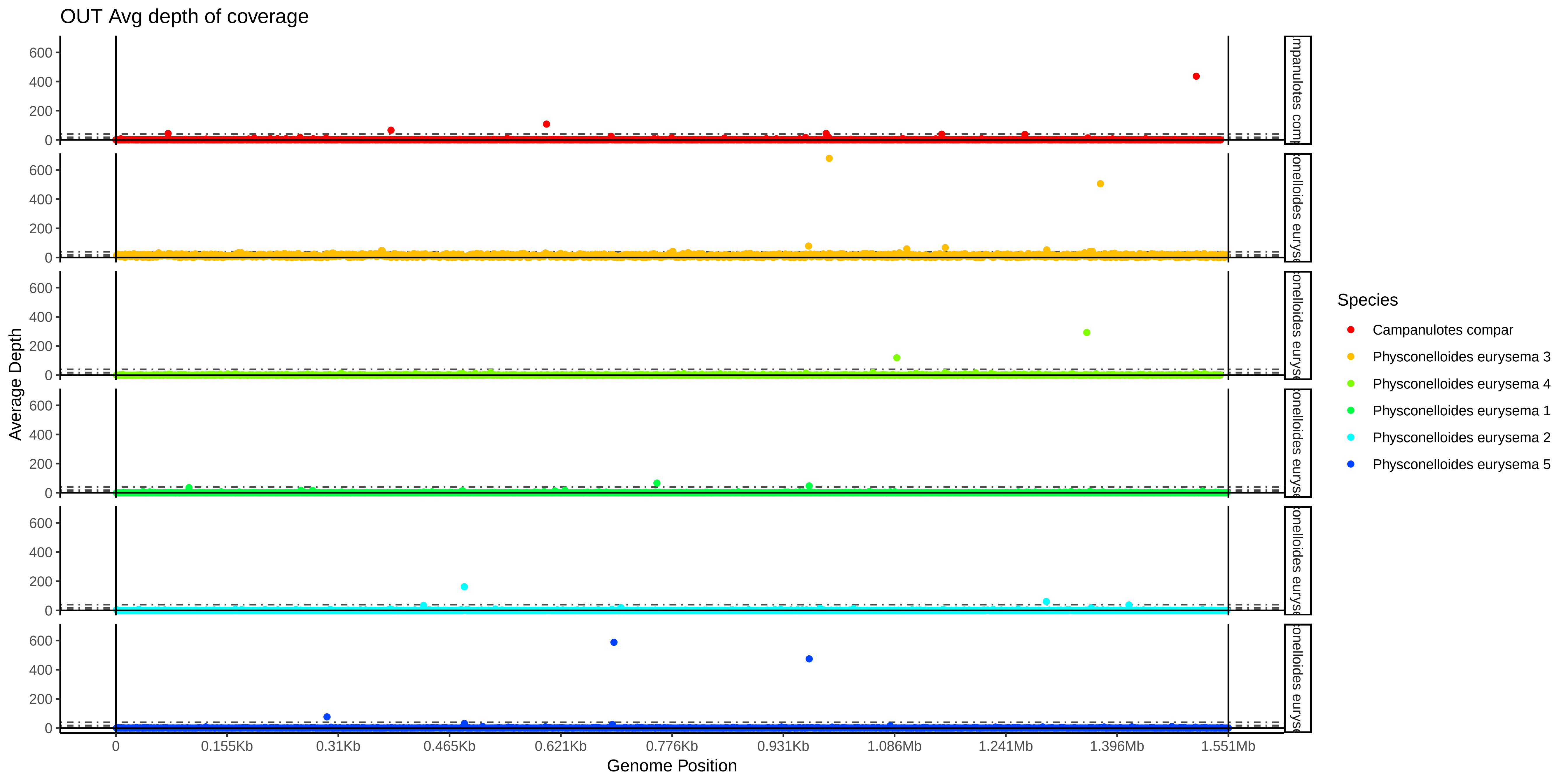
OUT Avg depth of coverage
Average Depth
Genome Position
0
200
400
600
0
0.155Kb
0.31Kb
0.465Kb
0.621Kb
0.776Kb
0.931Kb
1.086Mb
1.241Mb
1.396Mb
1.551Mb
Species
Campanulotes compar
Physconelloides eurysema 3
Physconelloides eurysema 4
Physconelloides eurysema 1
Physconelloides eurysema 2
Physconelloides eurysema 5

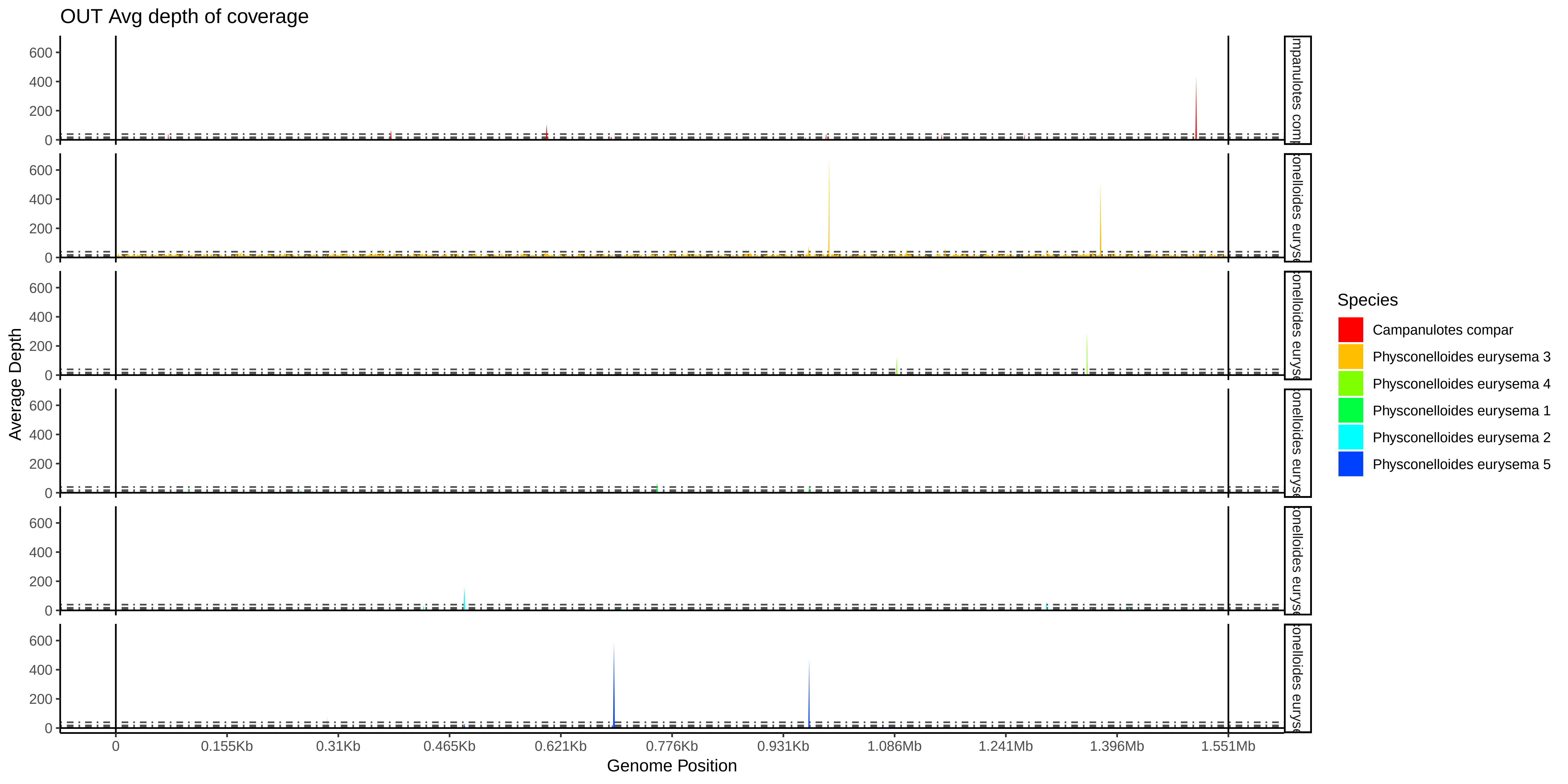
OUT Avg depth of coverage
Average Depth
Genome Position
0
0.155Kb
0.31Kb
0.465Kb
0.621Kb
0.776Kb
0.931Kb
1.086Mb
1.241Mb
1.396Mb
1.551Mb
0
200
400
600
Species
Campanulotes compar
Physconelloides eurysema 3
Physconelloides eurysema 4
Physconelloides eurysema 1
Physconelloides eurysema 2
Physconelloides eurysema 5

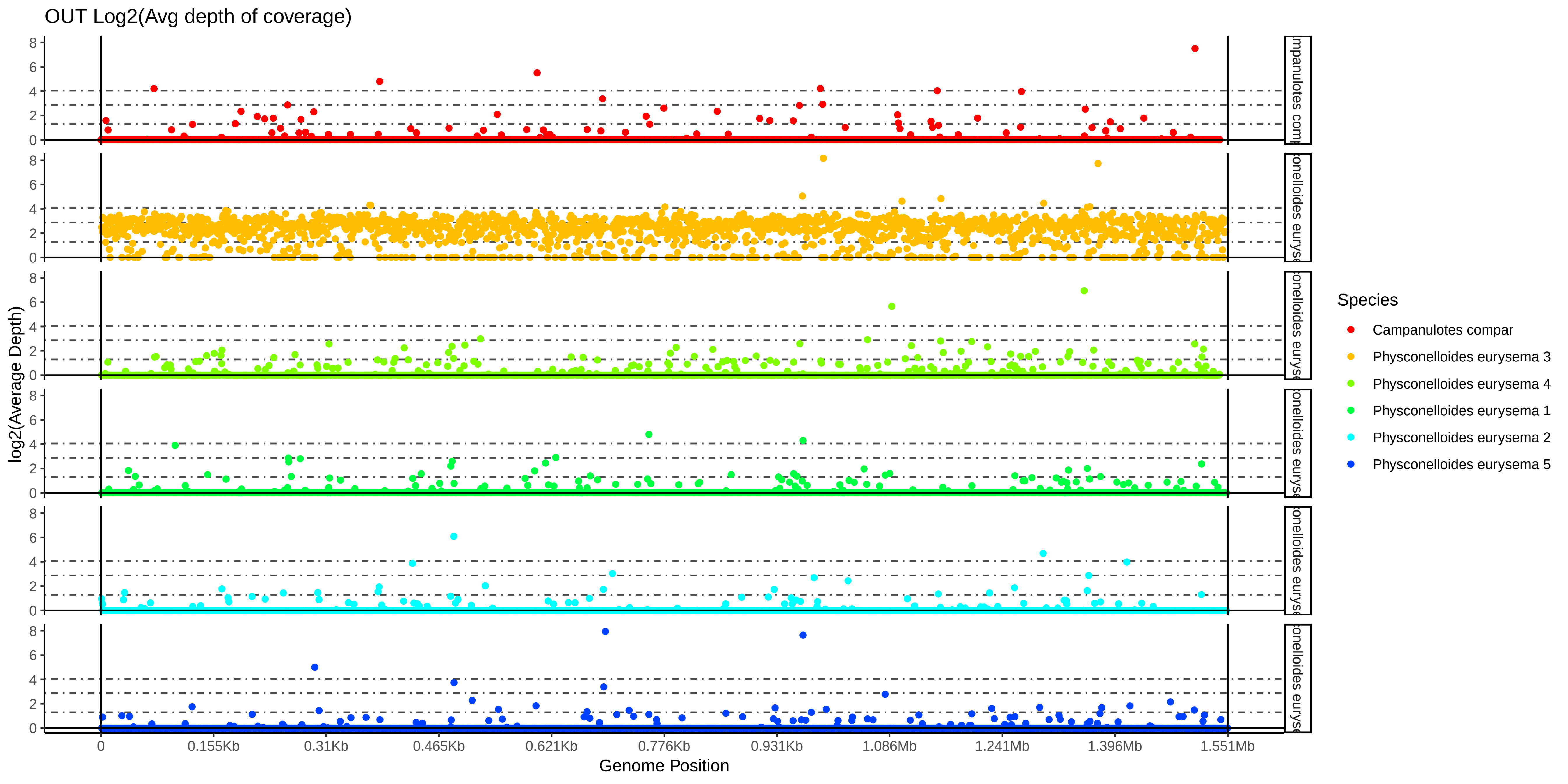

OUT Log2(Avg depth of coverage)
log2(Average Depth)
Genome Position
0
0.155Kb
0.31Kb
0.465Kb
0.621Kb
0.776Kb
0.931Kb
1.086Mb
1.241Mb
1.396Mb
1.551Mb
Species
Campanulotes compar
Physconelloides eurysema 3
Physconelloides eurysema 4
Physconelloides eurysema 1
Physconelloides eurysema 2
Physconelloides eurysema 5

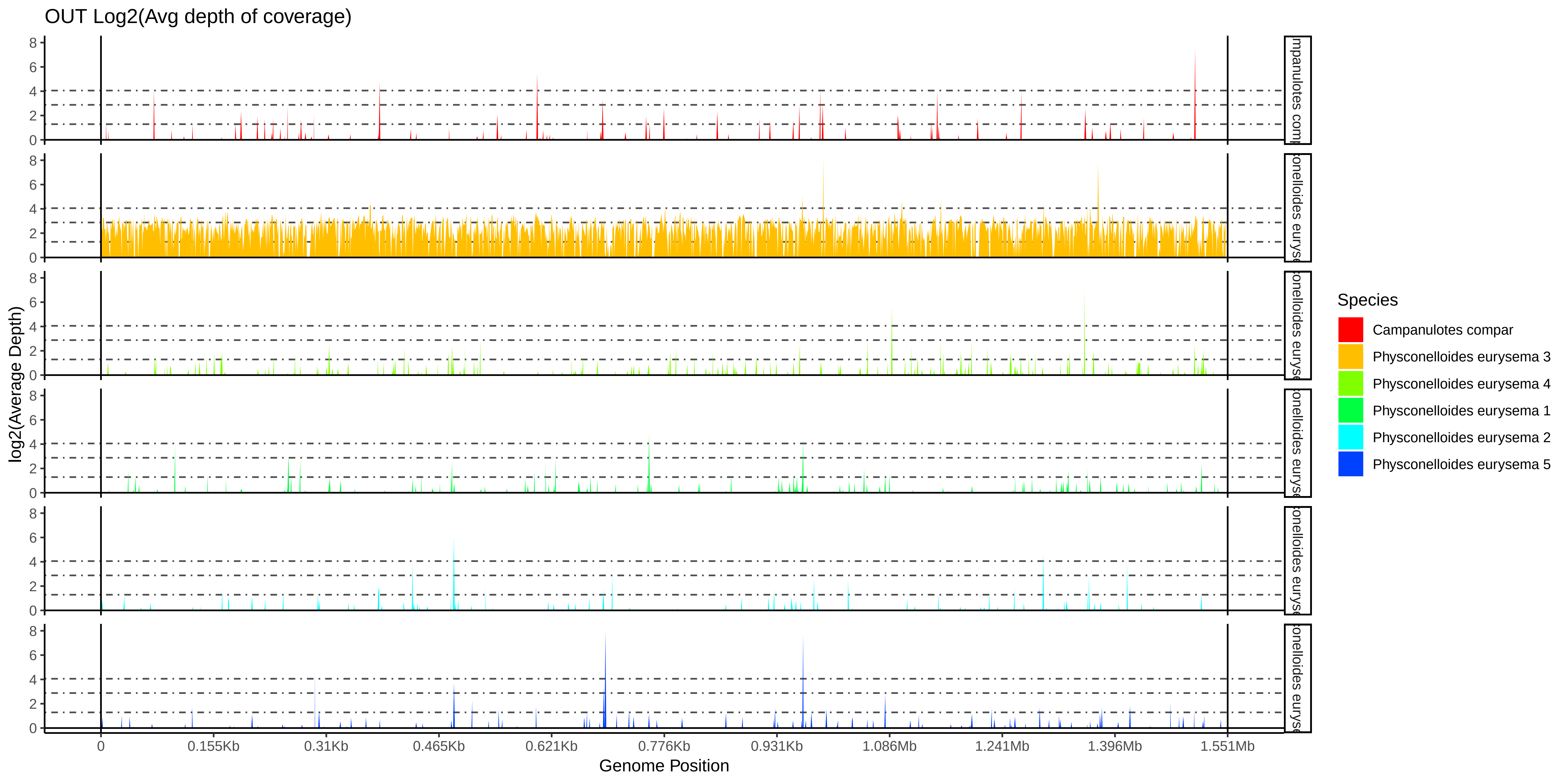

OUT Log2(Avg depth of coverage)
log2(Average Depth)
Genome Position
0
0.155Kb
0.31Kb
0.465Kb
0.621Kb
0.776Kb
0.931Kb
1.086Mb
1.241Mb
1.396Mb
1.551Mb
Species
Campanulotes compar
Physconelloides eurysema 3
Physconelloides eurysema 4
Physconelloides eurysema 1
Physconelloides eurysema 2
Physconelloides eurysema 5

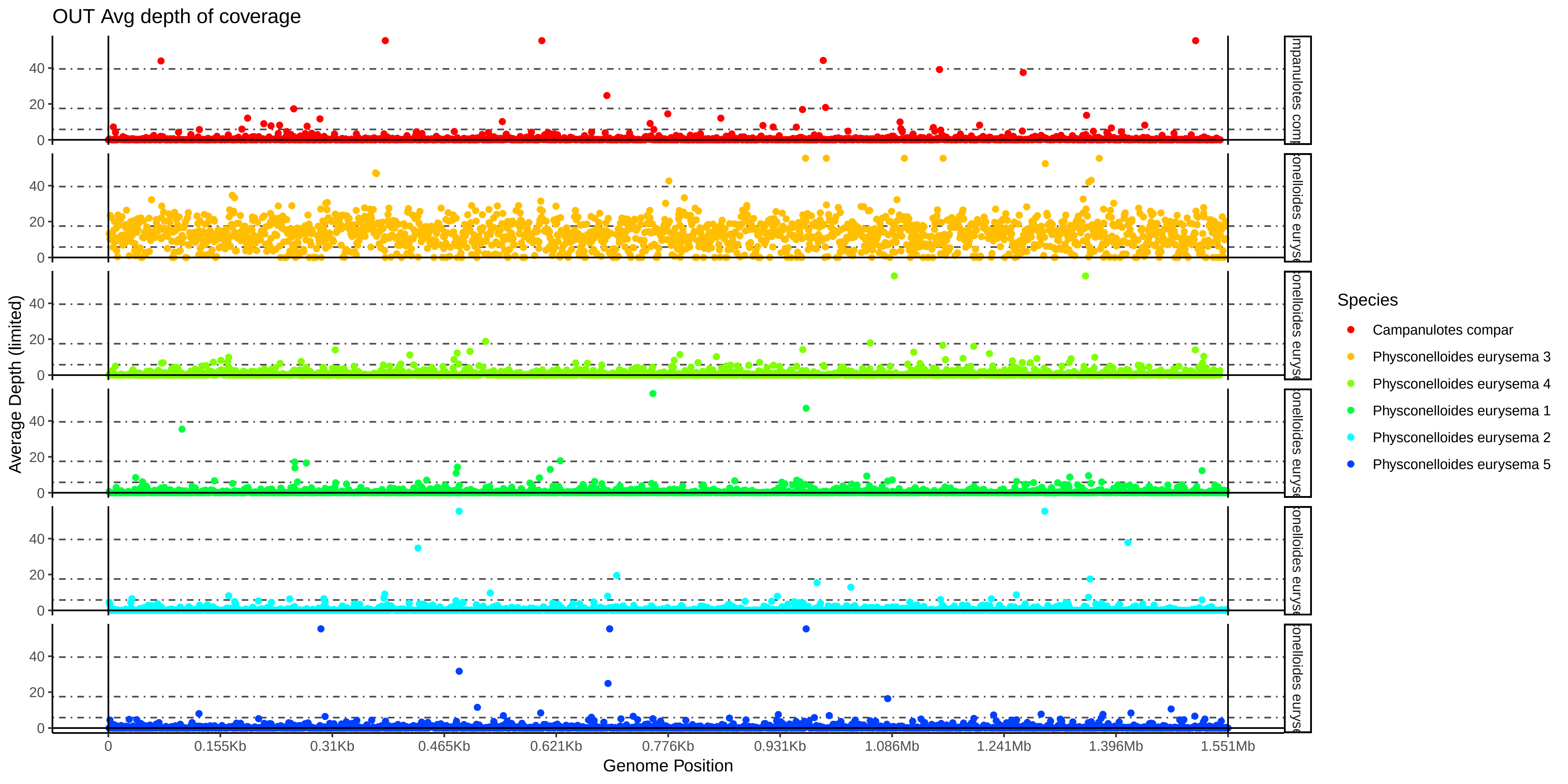
OUT Avg depth of coverage
Average Depth (limited)
Genome Position
0
20
40
0
0.155Kb
0.31Kb
0.465Kb
0.621Kb
0.776Kb
0.931Kb
1.086Mb
1.241Mb
1.396Mb
1.551Mb
Species
Campanulotes compar
Physconelloides eurysema 3
Physconelloides eurysema 4
Physconelloides eurysema 1
Physconelloides eurysema 2
Physconelloides eurysema 5

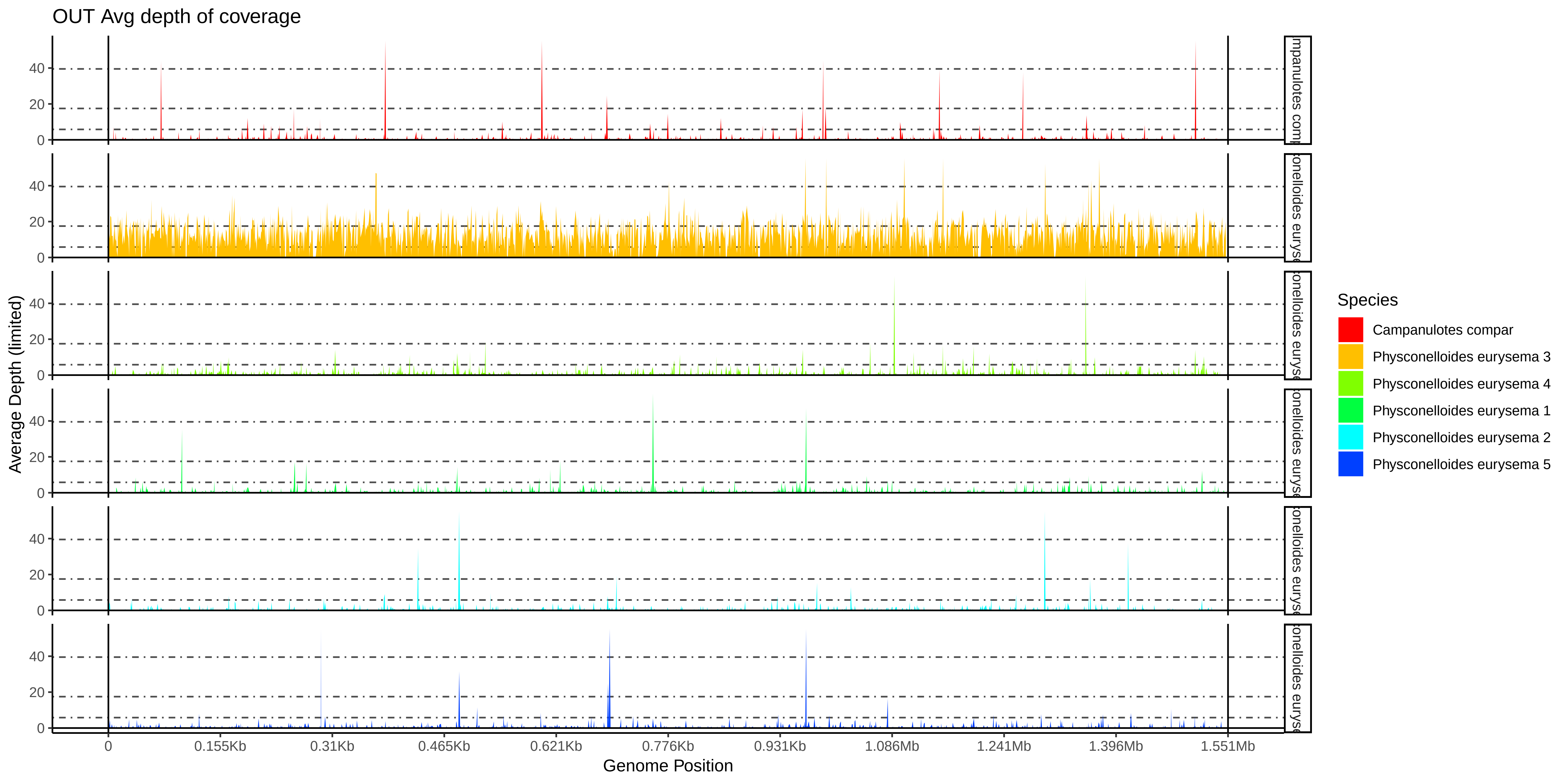

OUT Avg depth of coverage
Average Depth (limited)
Genome Position
0
20
40
0
0.155Kb
0.31Kb
0.465Kb
0.621Kb
0.776Kb
0.931Kb
1.086Mb
1.241Mb
1.396Mb
1.551Mb
Species
Campanulotes compar
Physconelloides eurysema 3
Physconelloides eurysema 4
Physconelloides eurysema 1
Physconelloides eurysema 2
Physconelloides eurysema 5
