## Supplementary material for "Comparing rates of introgression in parasitic feather lice with differing dispersal capabilities": OUT_sppIDerDepthPlot-covBins_Pheur_Cosqu_3_14_2016_1.pdf (from another version of this preprint: version 7, 2020-03-31)

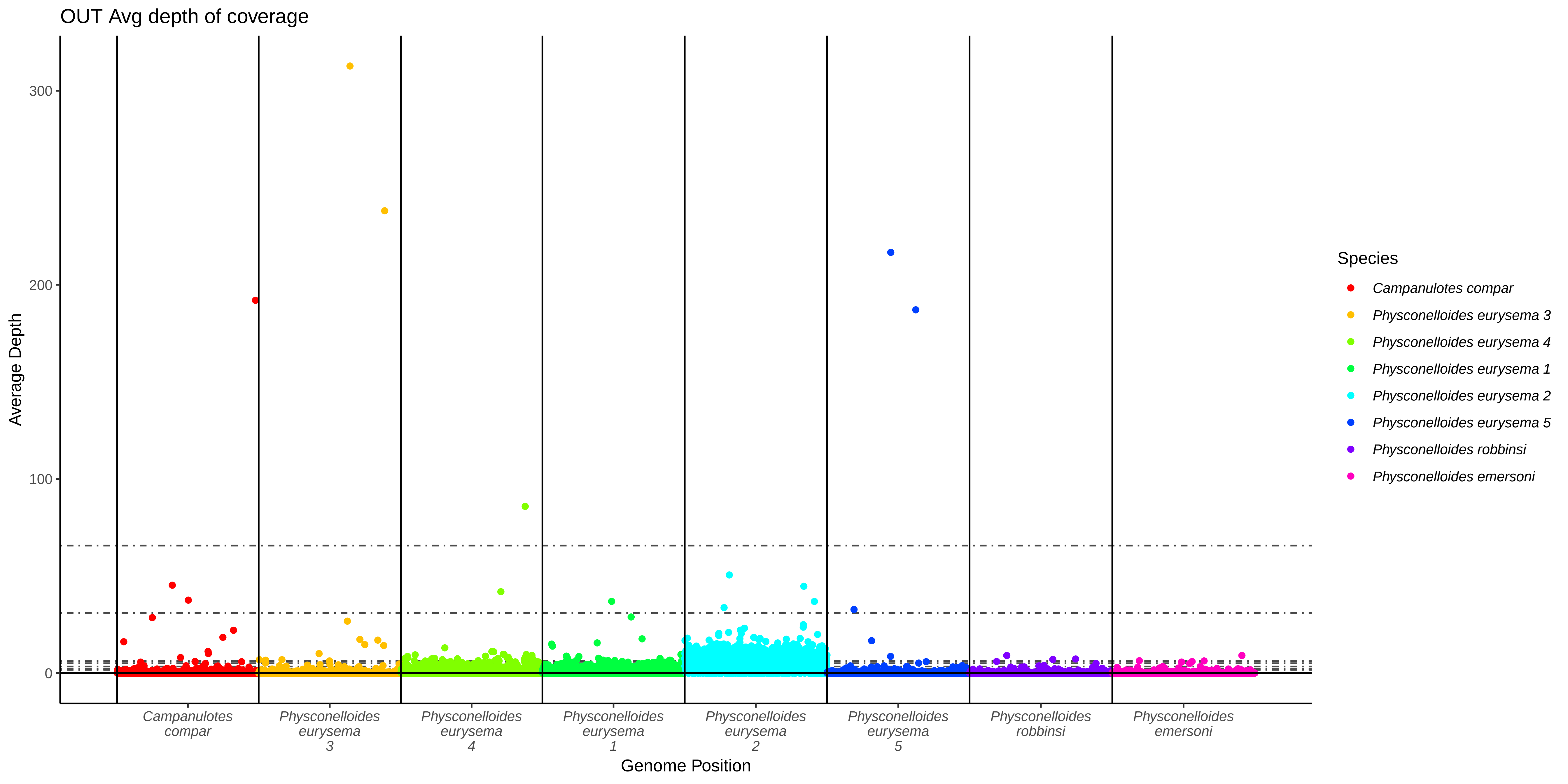

OUT Avg depth of coverage
Average Depth
300
200
100
0
Campanulotes compar
Physconelloides eurysema 3
Physconelloides eurysema 4
Physconelloides eurysema 1
Physconelloides eurysema 2
Physconelloides eurysema 5
Physconelloides robbinsi
Physconelloides emersoni
Genome Position
Species
Campanulotes compar
Physconelloides eurysema 3
Physconelloides eurysema 4
Physconelloides eurysema 1
Physconelloides eurysema 2
Physconelloides eurysema 5
Physconelloides robbinsi
Physconelloides emersoni

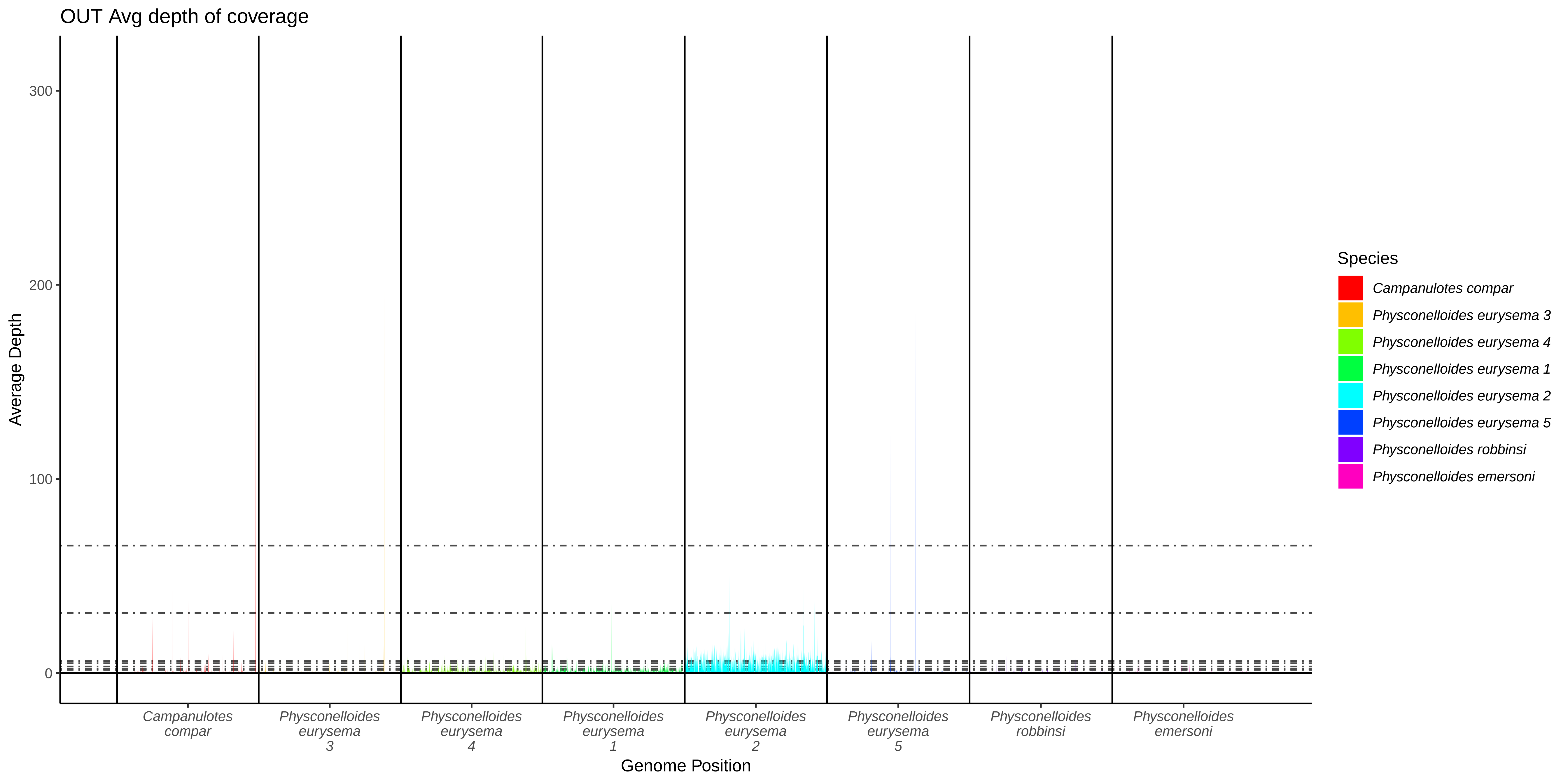
OUT Avg depth of coverage
300
200
100
0
Average Depth
Campanulotes compar
Physconelloides eurysema 3
Physconelloides eurysema 4
Physconelloides eurysema 1
Physconelloides eurysema 2
Physconelloides eurysema 5
Physconelloides robbinsi
Physconelloides emersoni
Genome Position
Species
Campanulotes compar
Physconelloides eurysema 3
Physconelloides eurysema 4
Physconelloides eurysema 1
Physconelloides eurysema 2
Physconelloides eurysema 5
Physconelloides robbinsi
Physconelloides emersoni

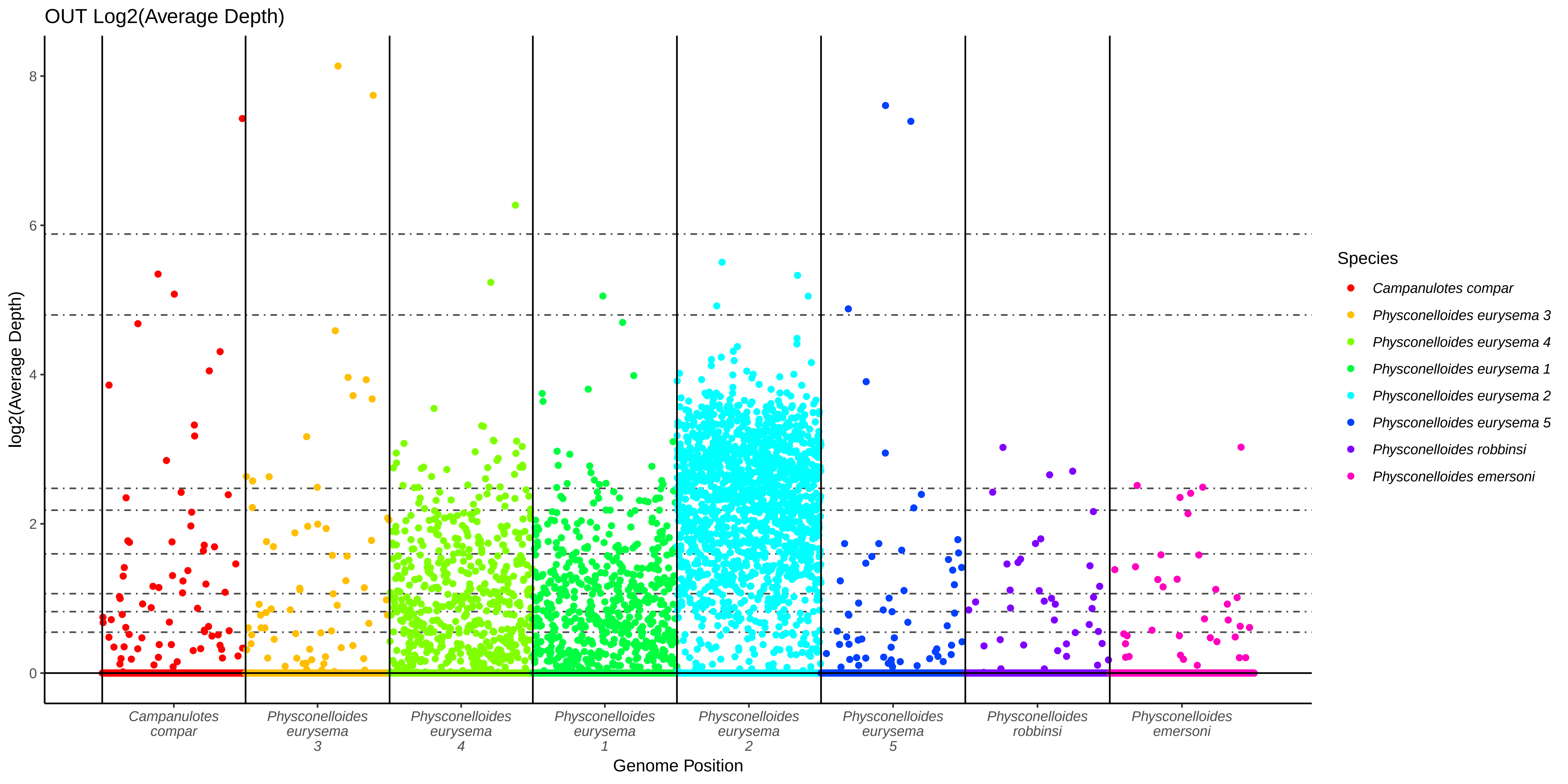
OUT Log2(Average Depth)
log2(Average Depth)
0
2
4
6
8
Campanulotes compar
Physconelloides eurysema 3
Physconelloides eurysema 4
Physconelloides eurysema 1
Physconelloides eurysema 2
Physconelloides eurysema 5
Physconelloides robbinsi
Physconelloides emersoni
Genome Position
Species
Campanulotes compar
Physconelloides eurysema 3
Physconelloides eurysema 4
Physconelloides eurysema 1
Physconelloides eurysema 2
Physconelloides eurysema 5
Physconelloides robbinsi
Physconelloides emersoni

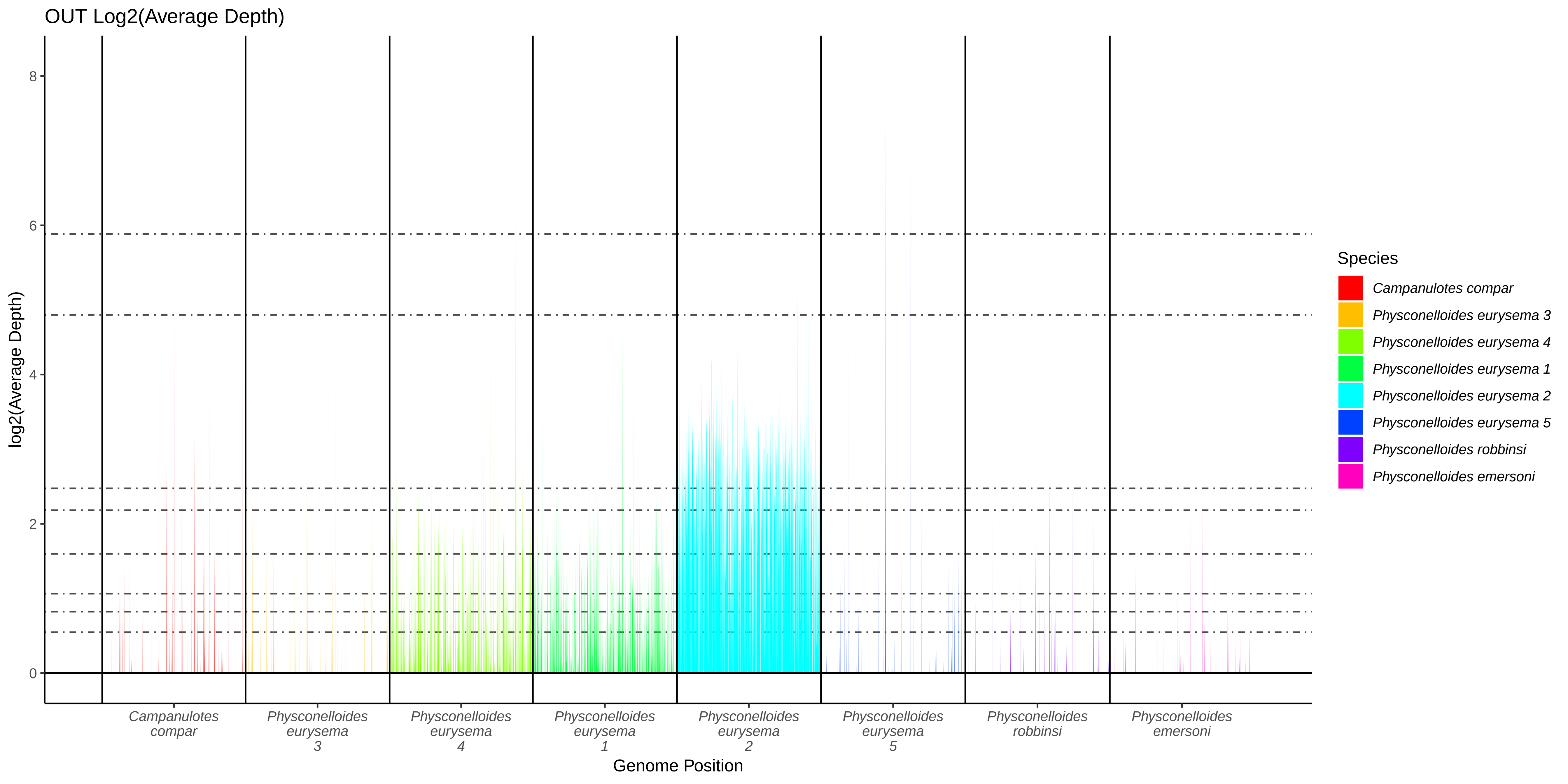

OUT Log2(Average Depth)
log2(Average Depth)
8
6
4
2
0
Campanulotes compar
Physconelloides eurysema 3
Physconelloides eurysema 4
Physconelloides eurysema 1
Physconelloides eurysema 2
Physconelloides eurysema 5
Physconelloides robbinsi
Physconelloides emersoni
Genome Position
Species
Campanulotes compar
Physconelloides eurysema 3
Physconelloides eurysema 4
Physconelloides eurysema 1
Physconelloides eurysema 2
Physconelloides eurysema 5
Physconelloides robbinsi
Physconelloides emersoni

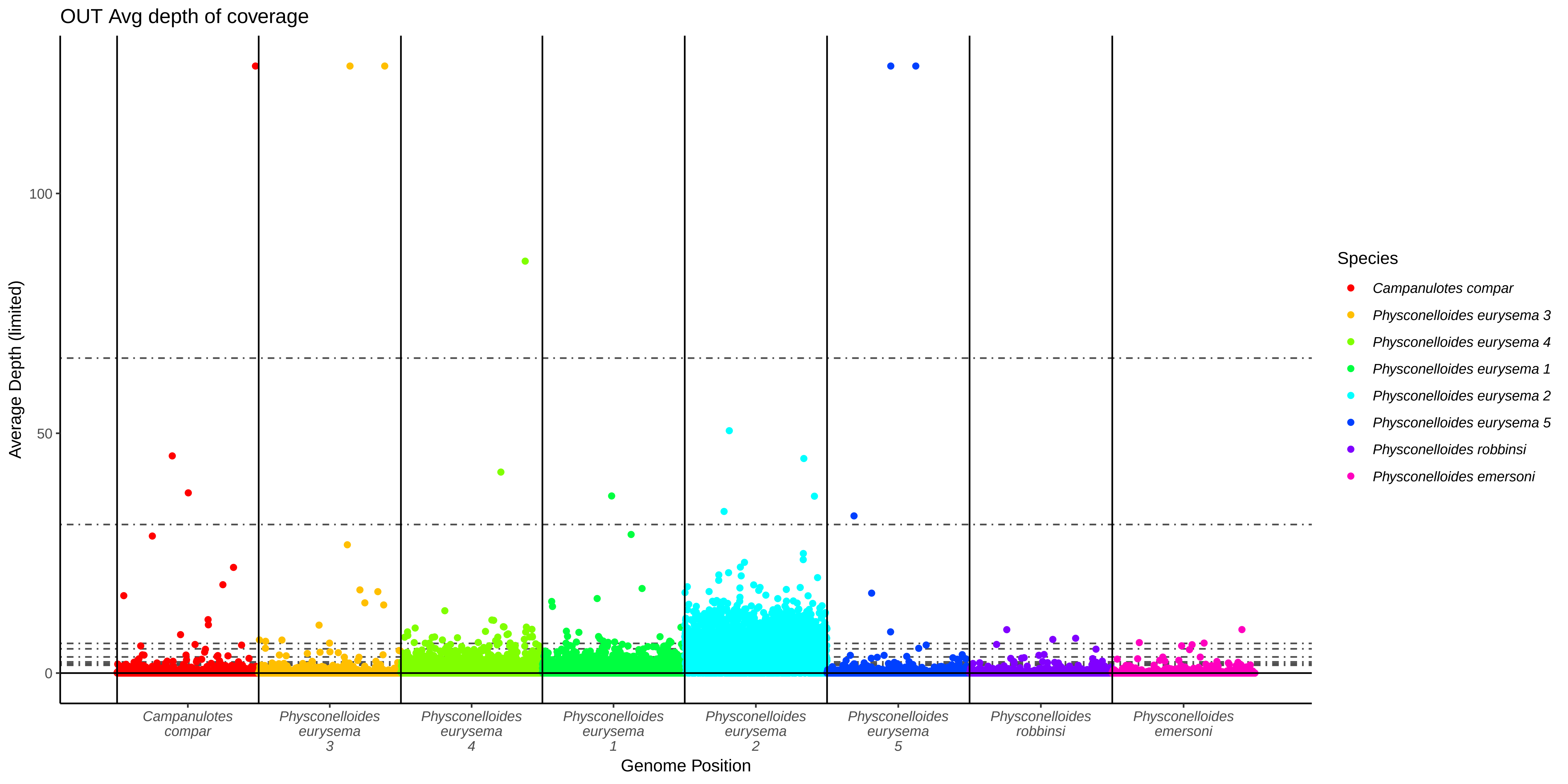

OUT Avg depth of coverage
Average Depth (limited)
100
50
0
Campanulotes compar
Physconelloides eurysema 3
Physconelloides eurysema 4
Physconelloides eurysema 1
Physconelloides eurysema 2
Physconelloides eurysema 5
Physconelloides robbinsi
Physconelloides emersoni
Genome Position
Species
Campanulotes compar
Physconelloides eurysema 3
Physconelloides eurysema 4
Physconelloides eurysema 1
Physconelloides eurysema 2
Physconelloides eurysema 5
Physconelloides robbinsi
Physconelloides emersoni

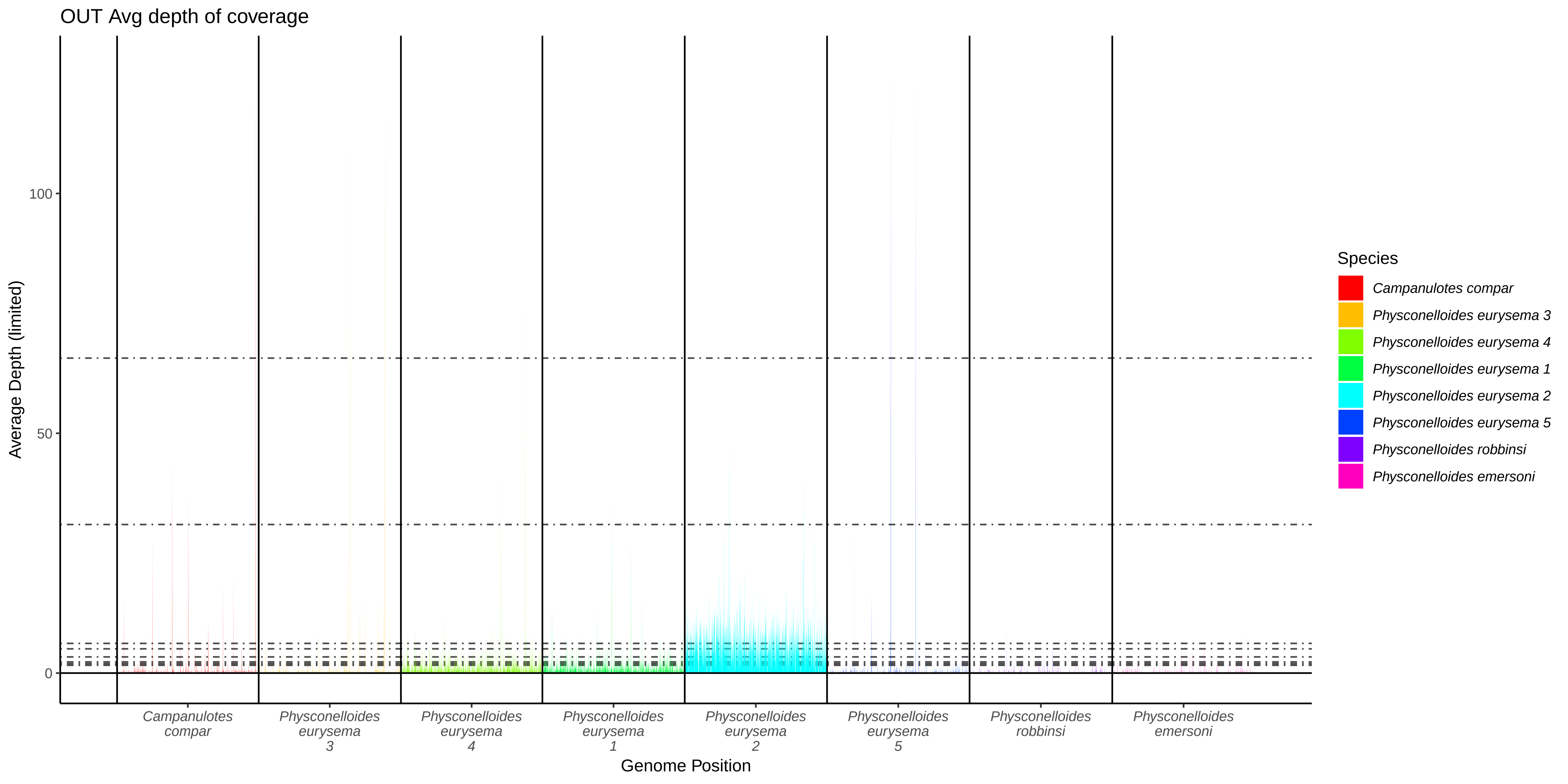
OUT Avg depth of coverage
Average Depth (limited)
100
50
0
Campanulotes compar
Physconelloides eurysema 3
Physconelloides eurysema 4
Physconelloides eurysema 1
Physconelloides eurysema 2
Physconelloides eurysema 5
Physconelloides robbinsi
Physconelloides emersoni
Genome Position
Species
Campanulotes compar
Physconelloides eurysema 3
Physconelloides eurysema 4
Physconelloides eurysema 1
Physconelloides eurysema 2
Physconelloides eurysema 5
Physconelloides robbinsi
Physconelloides emersoni

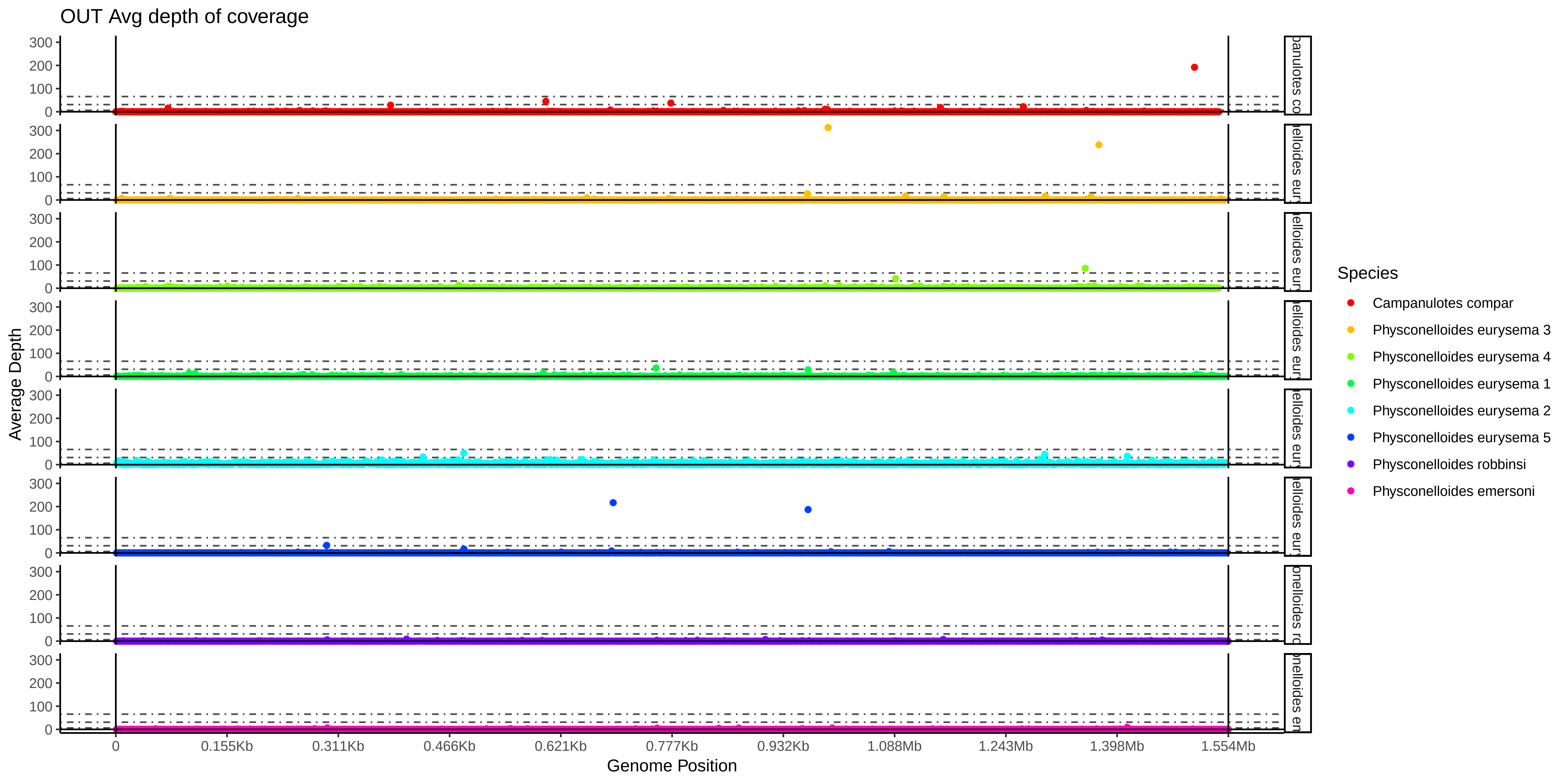

OUT Avg depth of coverage
Average Depth
Genome Position
0
0.155Kb
0.311Kb
0.466Kb
0.621Kb
0.777Kb
0.932Kb
1.088Mb
1.243Mb
1.398Mb
1.554Mb
300
200
100
0
Species
Campanulotes compar
Physconelloides eurysema 3
Physconelloides eurysema 4
Physconelloides eurysema 1
Physconelloides eurysema 2
Physconelloides eurysema 5
Physconelloides robbinsi
Physconelloides emersoni

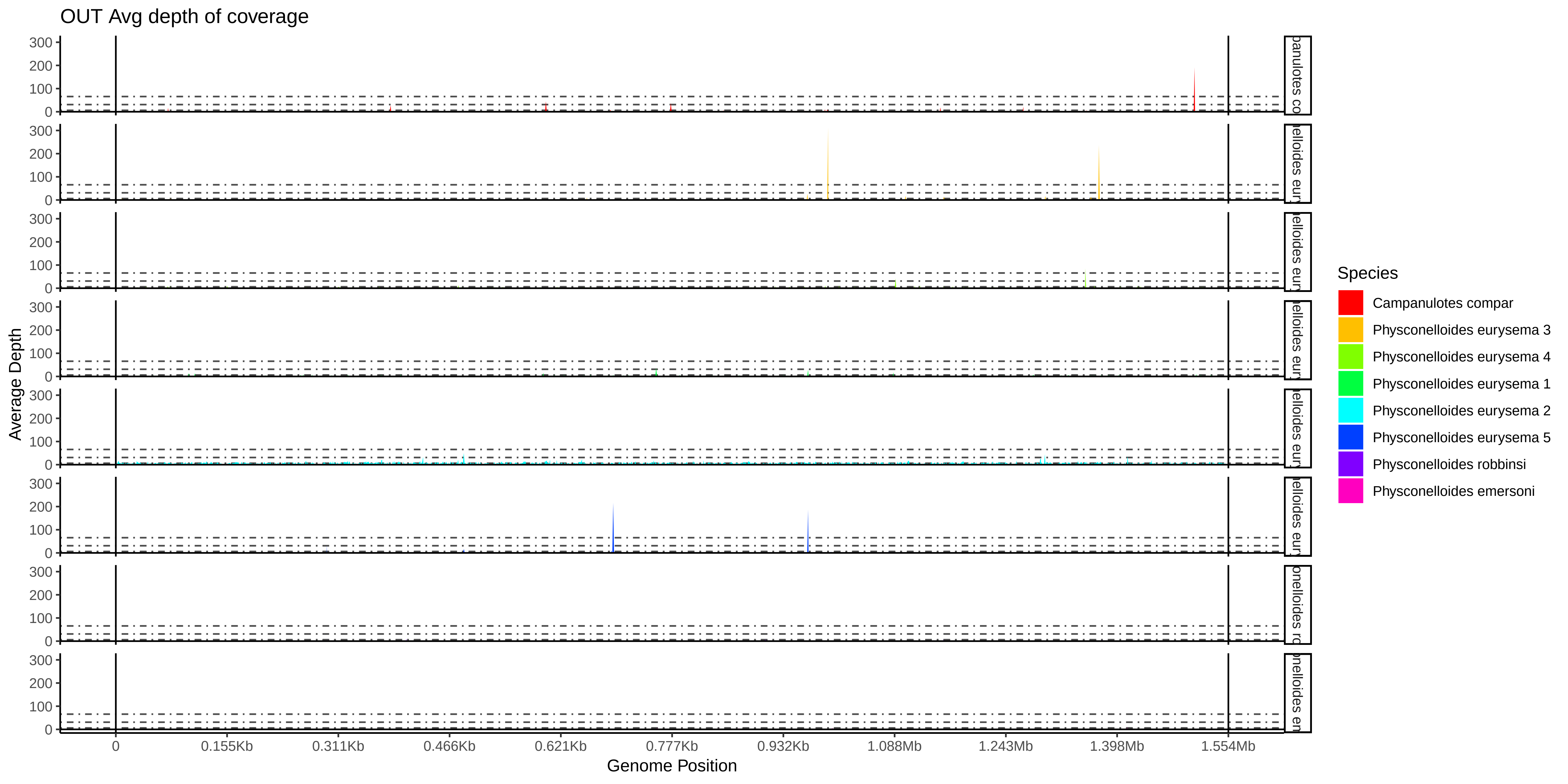

OUT Avg depth of coverage
Average Depth
300
200
100
0
Genome Position
0
0.155Kb
0.311Kb
0.466Kb
0.621Kb
0.777Kb
0.932Kb
1.088Mb
1.243Mb
1.398Mb
1.554Mb
Species
Campanulotes compar
Physconelloides eurysema 3
Physconelloides eurysema 4
Physconelloides eurysema 1
Physconelloides eurysema 2
Physconelloides eurysema 5
Physconelloides robbinsi
Physconelloides emersoni

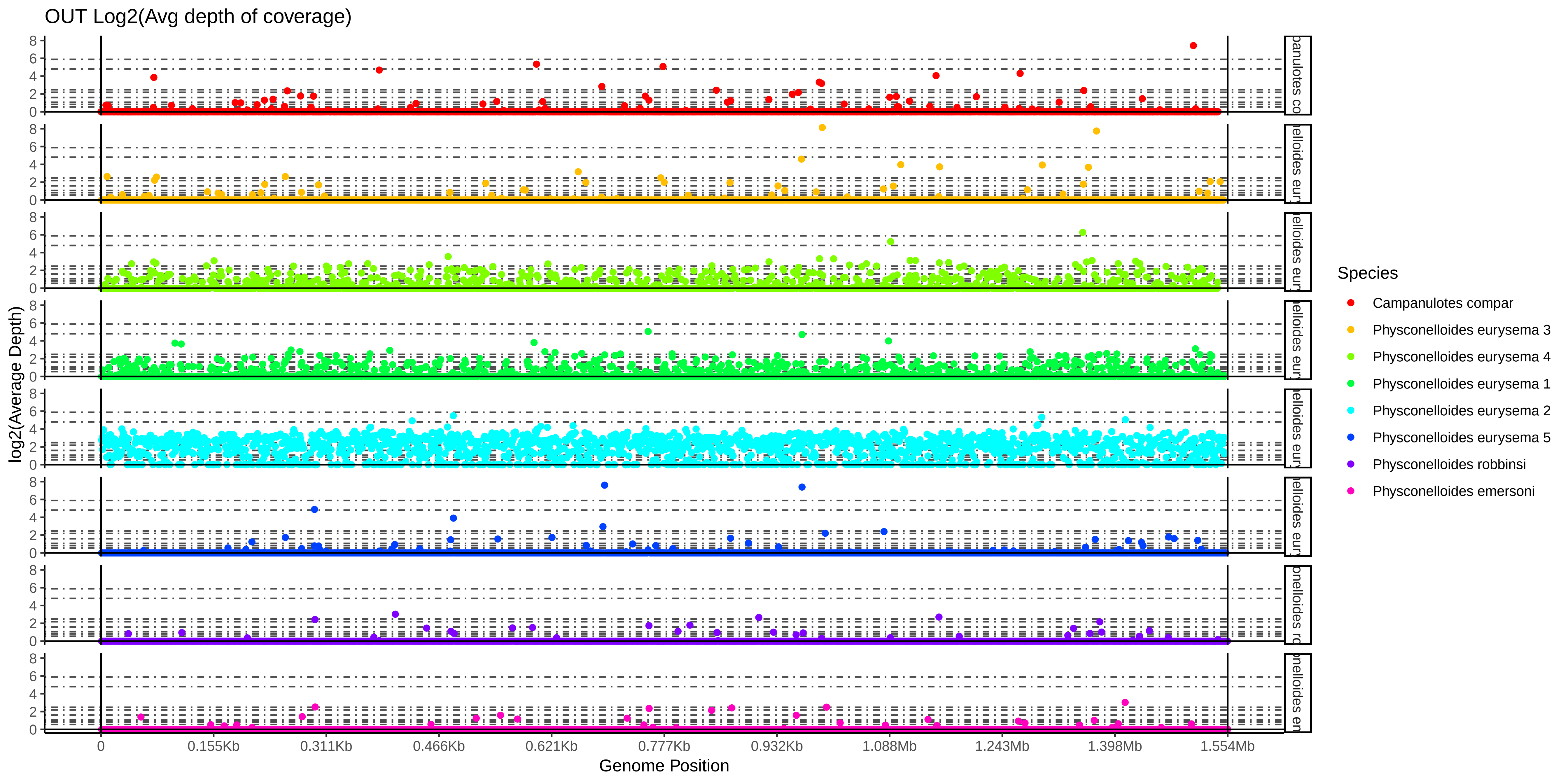

OUT Log2(Avg depth of coverage)
log2(Average Depth)
Genome Position
0
0.155Kb
0.311Kb
0.466Kb
0.621Kb
0.777Kb
0.932Kb
1.088Mb
1.243Mb
1.398Mb
1.554Mb
Species
Campanulotes compar
Physconelloides eurysema 3
Physconelloides eurysema 4
Physconelloides eurysema 1
Physconelloides eurysema 2
Physconelloides eurysema 5
Physconelloides robbinsi
Physconelloides emersoni

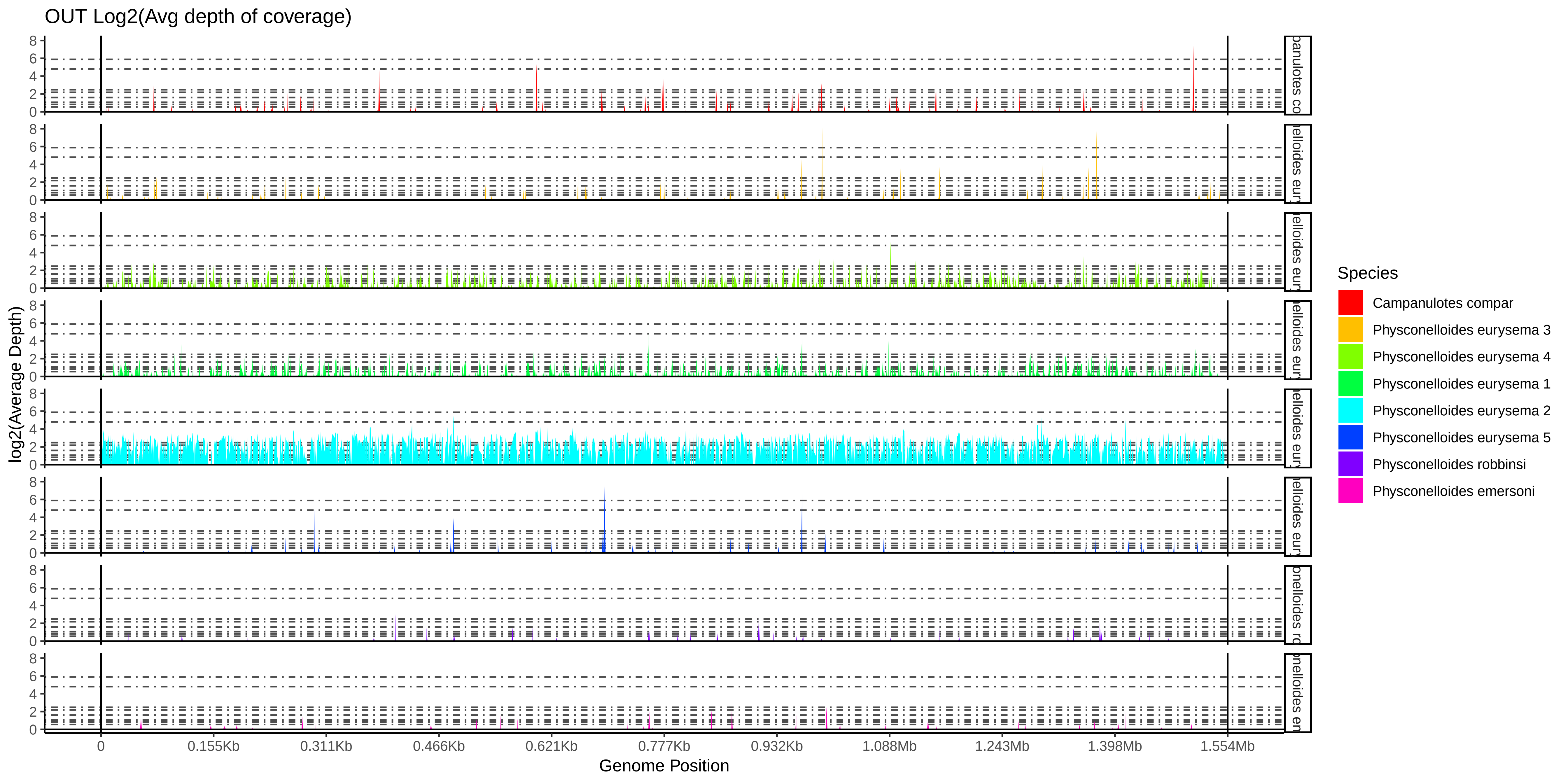

OUT Log2(Avg depth of coverage)
log2(Average Depth)
Genome Position
0
0.155Kb
0.311Kb
0.466Kb
0.621Kb
0.777Kb
0.932Kb
1.088Mb
1.243Mb
1.398Mb
1.554Mb
Species
Campanulotes compar
Physconelloides eurysema 3
Physconelloides eurysema 4
Physconelloides eurysema 1
Physconelloides eurysema 2
Physconelloides eurysema 5
Physconelloides robbinsi
Physconelloides emersoni

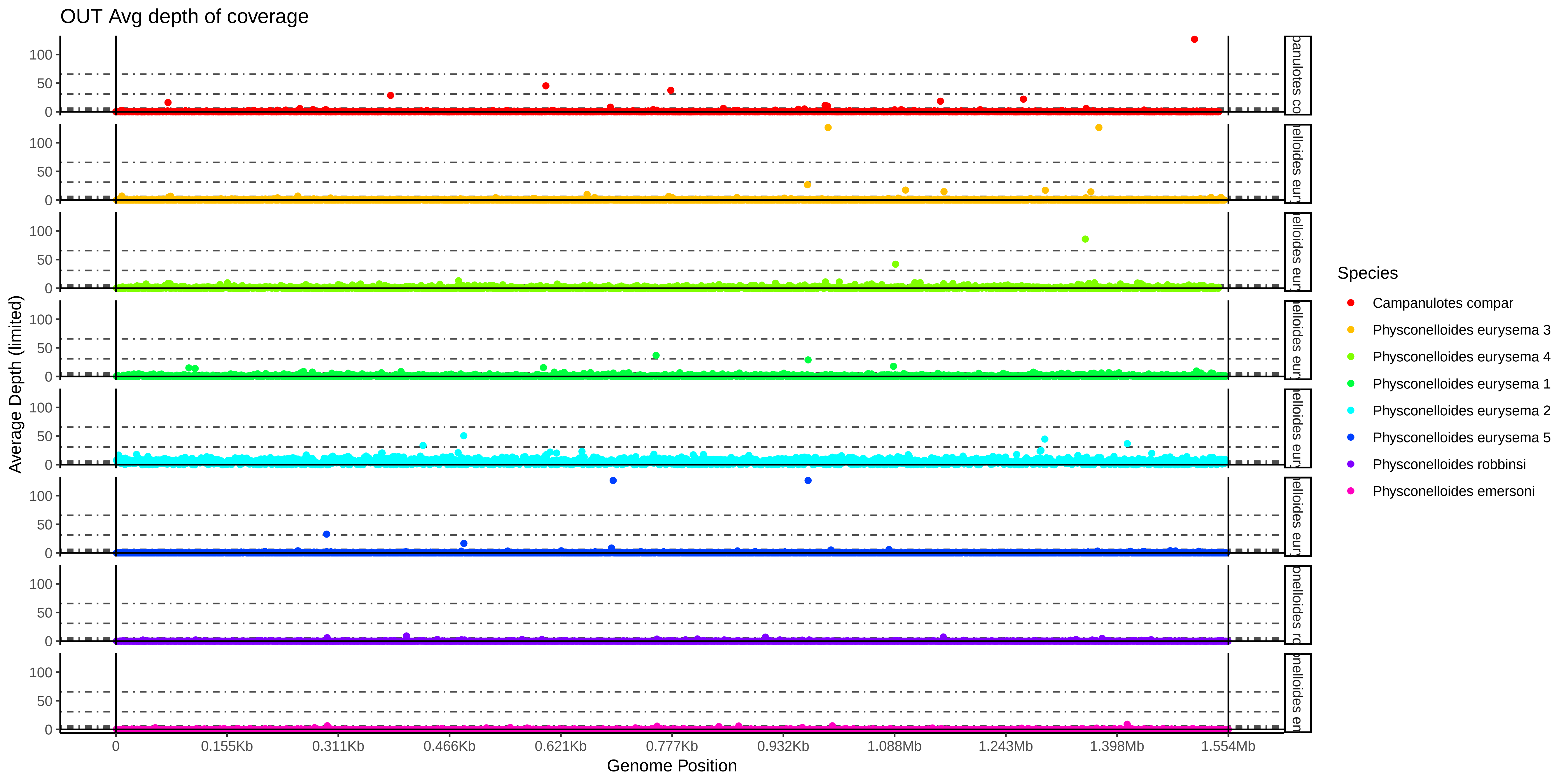

OUT Avg depth of coverage
Average Depth (limited)
Genome Position
0
0.155Kb
0.311Kb
0.466Kb
0.621Kb
0.777Kb
0.932Kb
1.088Mb
1.243Mb
1.398Mb
1.554Mb
100
50
0
Species
Campanulotes compar
Physconelloides eurysema 3
Physconelloides eurysema 4
Physconelloides eurysema 1
Physconelloides eurysema 2
Physconelloides eurysema 5
Physconelloides robbinsi
Physconelloides emersoni

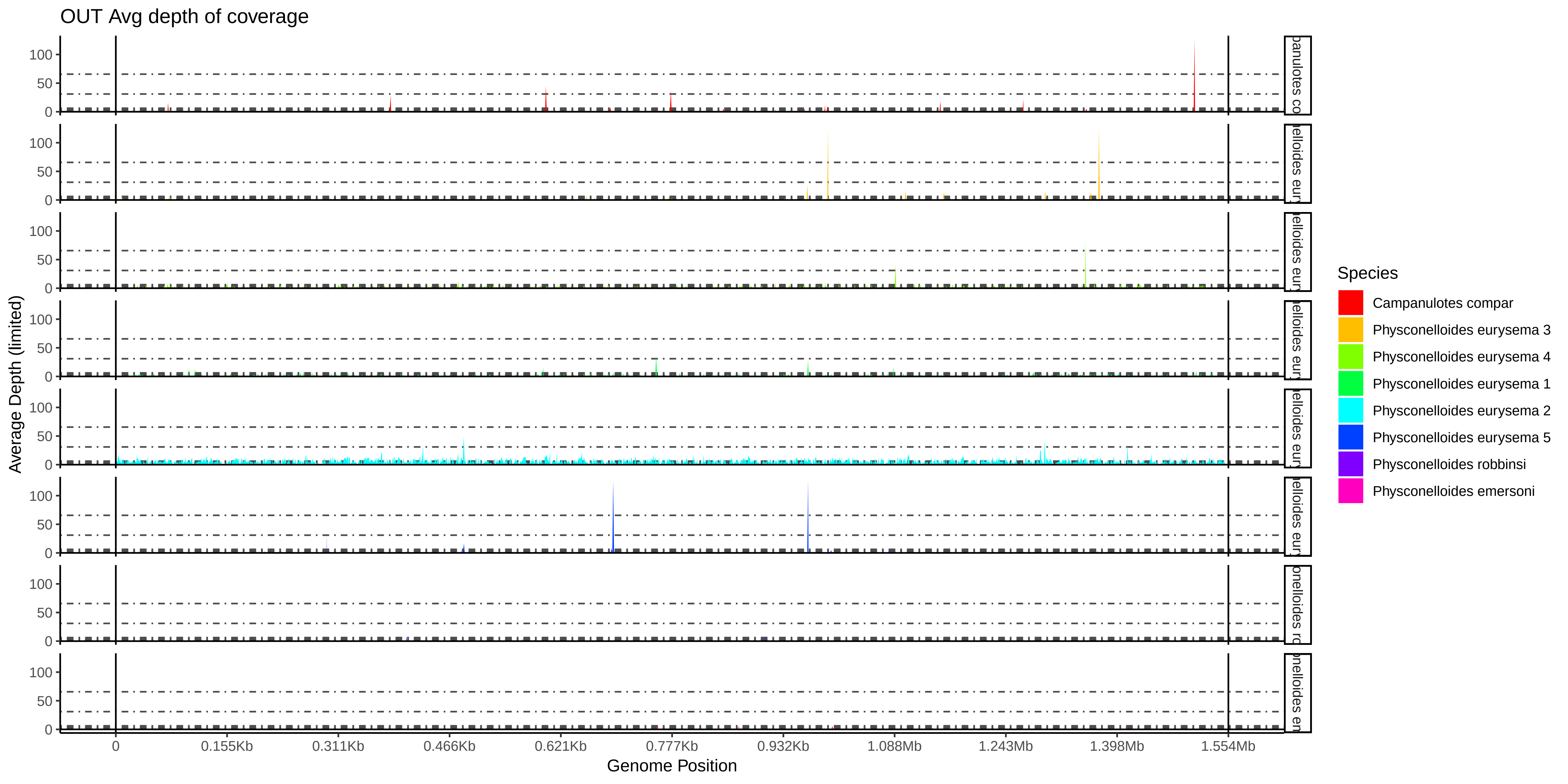
OUT Avg depth of coverage
Average Depth (limited)
Genome Position
0
50
100
0.155Kb
0.311Kb
0.466Kb
0.621Kb
0.777Kb
0.932Kb
1.088Mb
1.243Mb
1.398Mb
1.554Mb
Species
Campanulotes compar
Physconelloides eurysema 3
Physconelloides eurysema 4
Physconelloides eurysema 1
Physconelloides eurysema 2
Physconelloides eurysema 5
Physconelloides robbinsi
Physconelloides emersoni
