## Supplementary material for "Comparing rates of introgression in parasitic feather lice with differing dispersal capabilities": OUT_sppIDerDepthPlot-covBins_Phrob_Mecec_3_16_2016_10.pdf (from another version of this preprint: version 7, 2020-03-31)

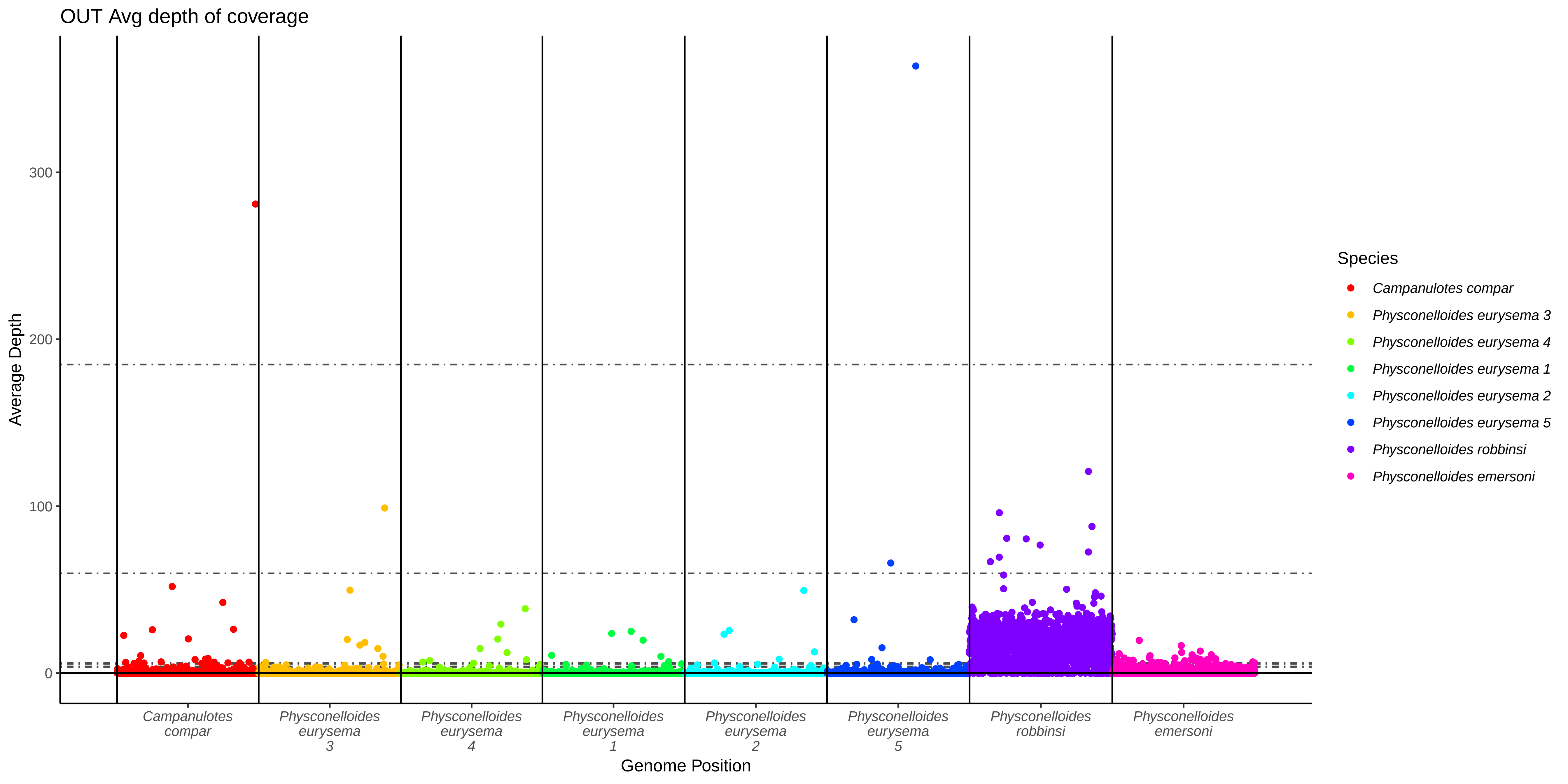
OUT Avg depth of coverage
Average Depth
0
100
200
300
Genome Position
Campanulotes compar
Physconelloides eurysema 3
Physconelloides eurysema 4
Physconelloides eurysema 1
Physconelloides eurysema 2
Physconelloides eurysema 5
Physconelloides robbinsi
Physconelloides emersoni
Species
Campanulotes compar
Physconelloides eurysema 3
Physconelloides eurysema 4
Physconelloides eurysema 1
Physconelloides eurysema 2
Physconelloides eurysema 5
Physconelloides robbinsi
Physconelloides emersoni

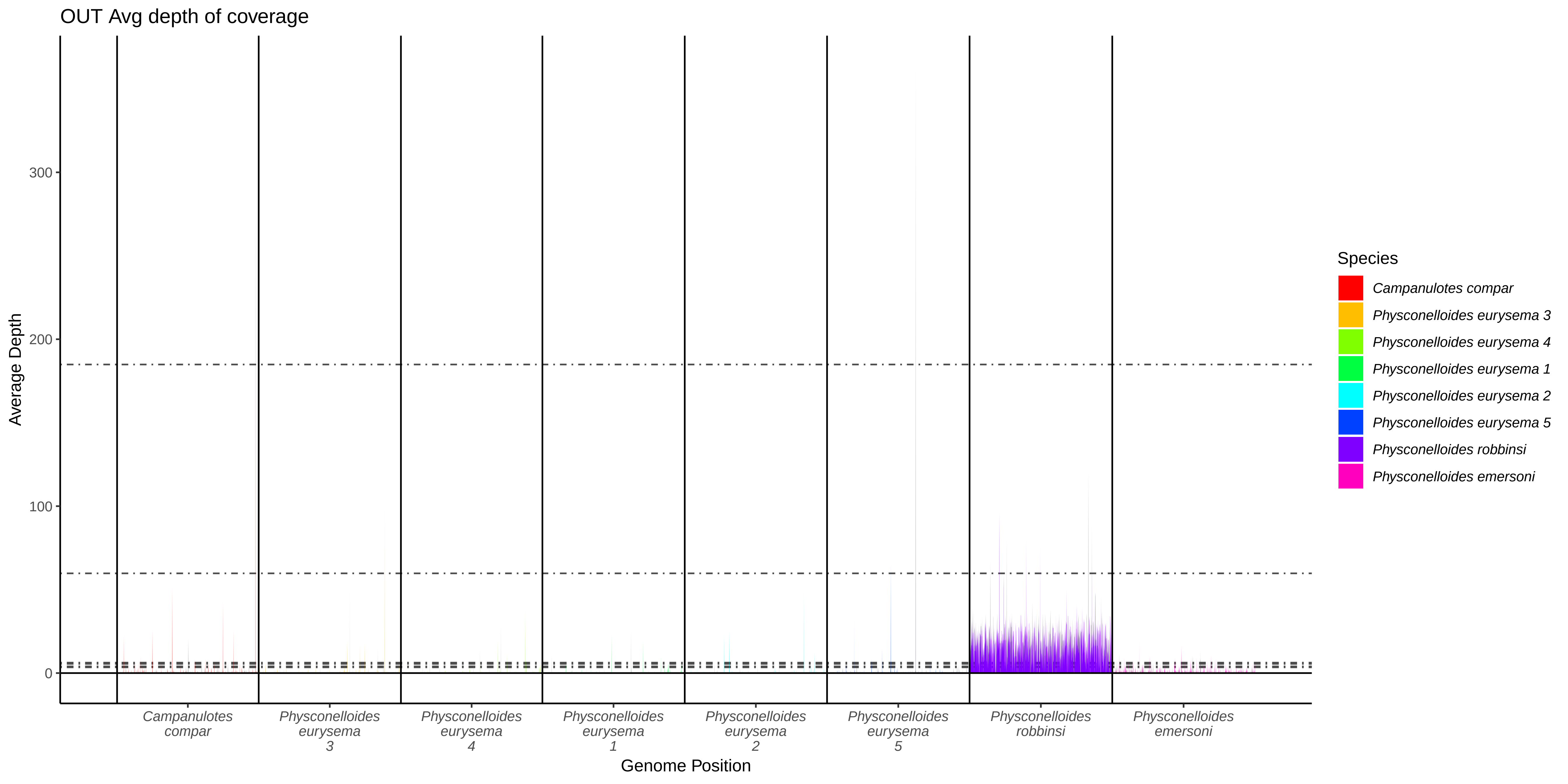

OUT Avg depth of coverage
Average Depth
300
200
100
0
Campanulotes compar
Physconelloides eurysema 3
Physconelloides eurysema 4
Physconelloides eurysema 1
Physconelloides eurysema 2
Physconelloides eurysema 5
Physconelloides robbinsi
Physconelloides emersoni
Genome Position
Species
Campanulotes compar
Physconelloides eurysema 3
Physconelloides eurysema 4
Physconelloides eurysema 1
Physconelloides eurysema 2
Physconelloides eurysema 5
Physconelloides robbinsi
Physconelloides emersoni

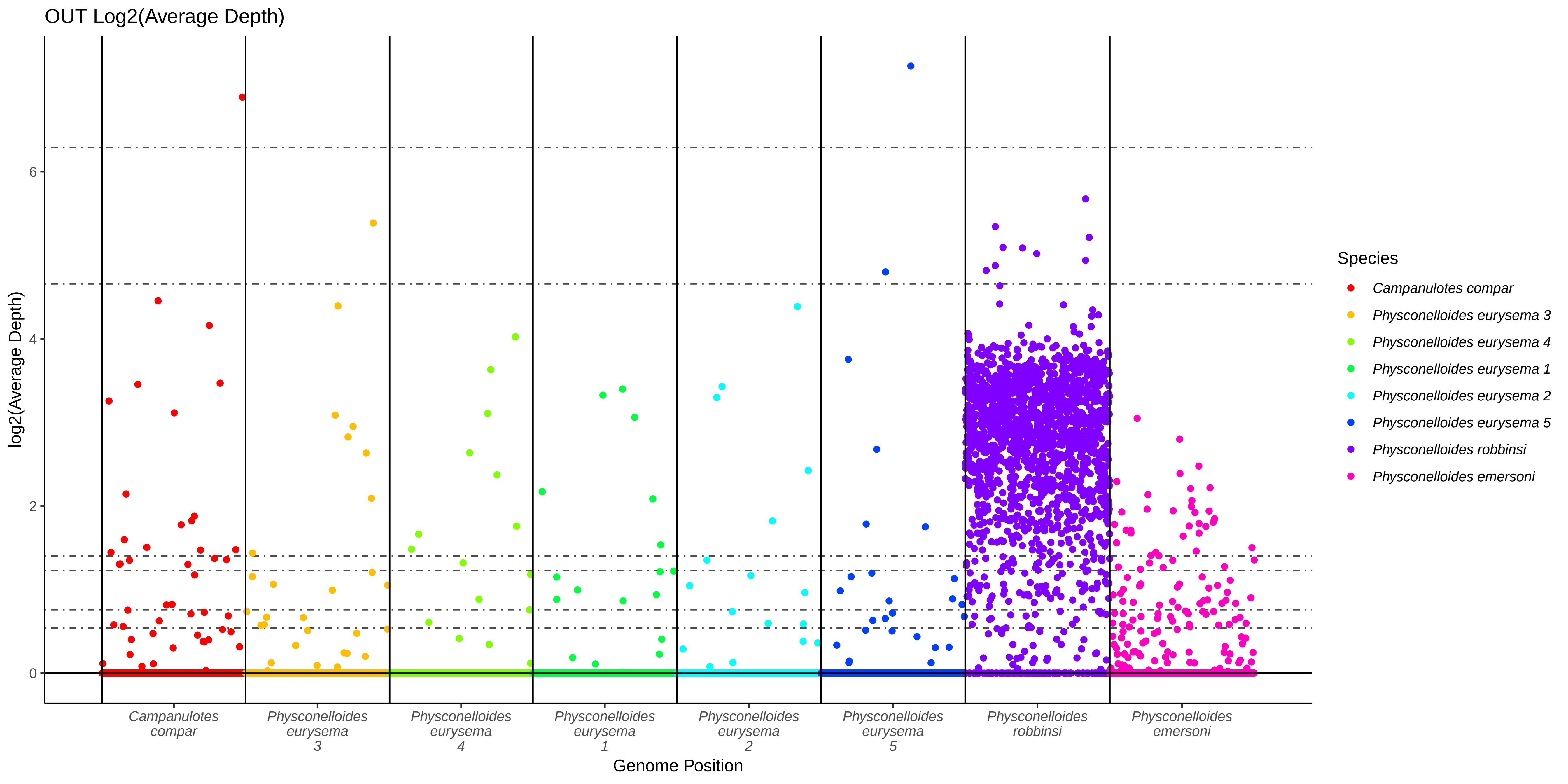
OUT Log2(Average Depth)
log2(Average Depth)
0
2
4
6
Campanulotes compar
Physconelloides eurysema 3
Physconelloides eurysema 4
Physconelloides eurysema 1
Physconelloides eurysema 2
Physconelloides eurysema 5
Physconelloides robbinsi
Physconelloides emersoni
Genome Position
Species
Campanulotes compar
Physconelloides eurysema 3
Physconelloides eurysema 4
Physconelloides eurysema 1
Physconelloides eurysema 2
Physconelloides eurysema 5
Physconelloides robbinsi
Physconelloides emersoni

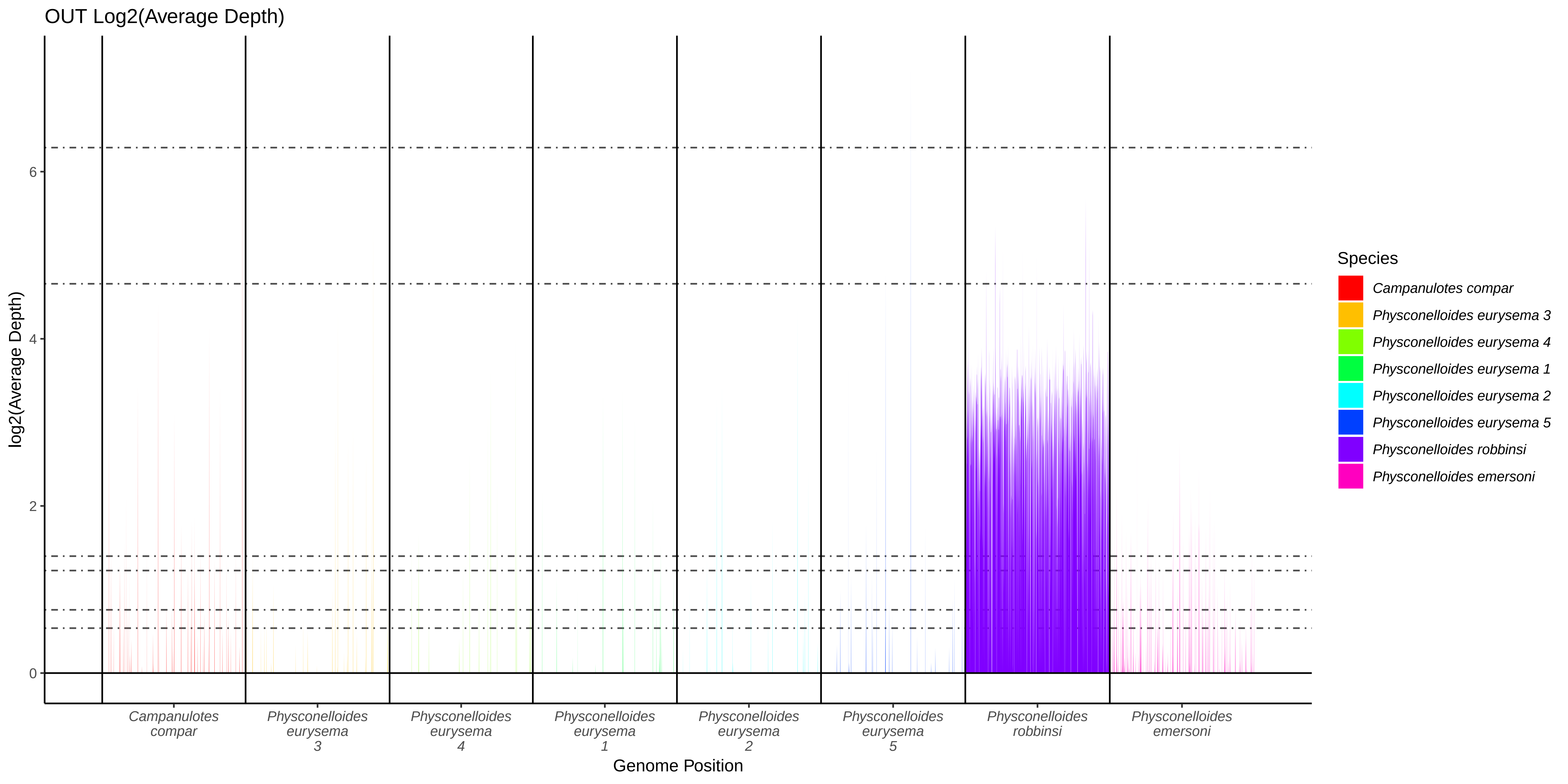

OUT Log2(Average Depth)
log2(Average Depth)
0
2
4
6
Campanulotes compar
Physconelloides eurysema 3
Physconelloides eurysema 4
Physconelloides eurysema 1
Physconelloides eurysema 2
Physconelloides eurysema 5
Physconelloides robbinsi
Physconelloides emersoni
Genome Position
Species
Campanulotes compar
Physconelloides eurysema 3
Physconelloides eurysema 4
Physconelloides eurysema 1
Physconelloides eurysema 2
Physconelloides eurysema 5
Physconelloides robbinsi
Physconelloides emersoni

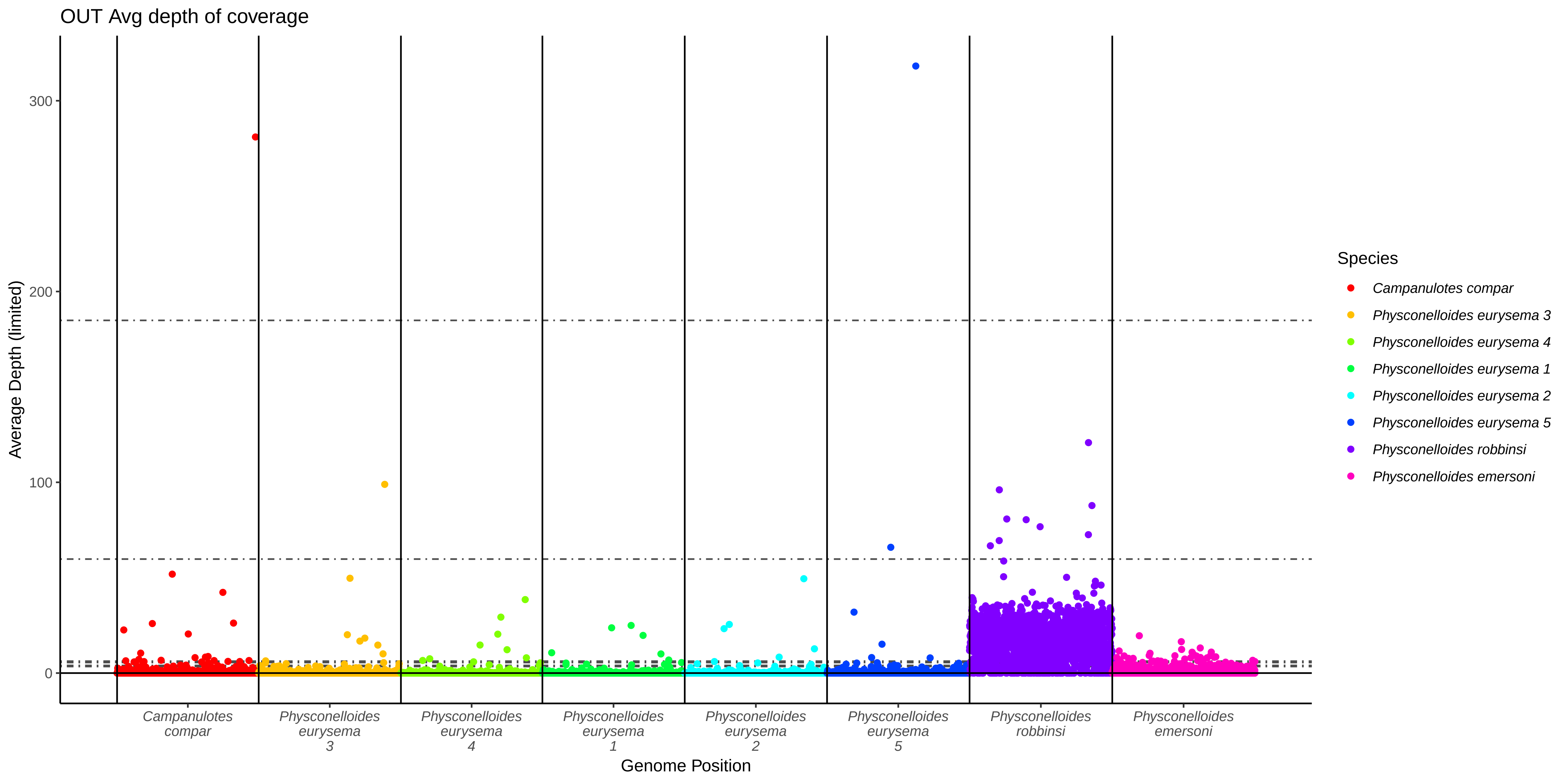
OUT Avg depth of coverage
Average Depth (limited)
300
200
100
0
Campanulotes compar
Physconelloides eurysema 3
Physconelloides eurysema 4
Physconelloides eurysema 1
Physconelloides eurysema 2
Physconelloides eurysema 5
Physconelloides robbinsi
Physconelloides emersoni
Genome Position
Species
Campanulotes compar
Physconelloides eurysema 3
Physconelloides eurysema 4
Physconelloides eurysema 1
Physconelloides eurysema 2
Physconelloides eurysema 5
Physconelloides robbinsi
Physconelloides emersoni

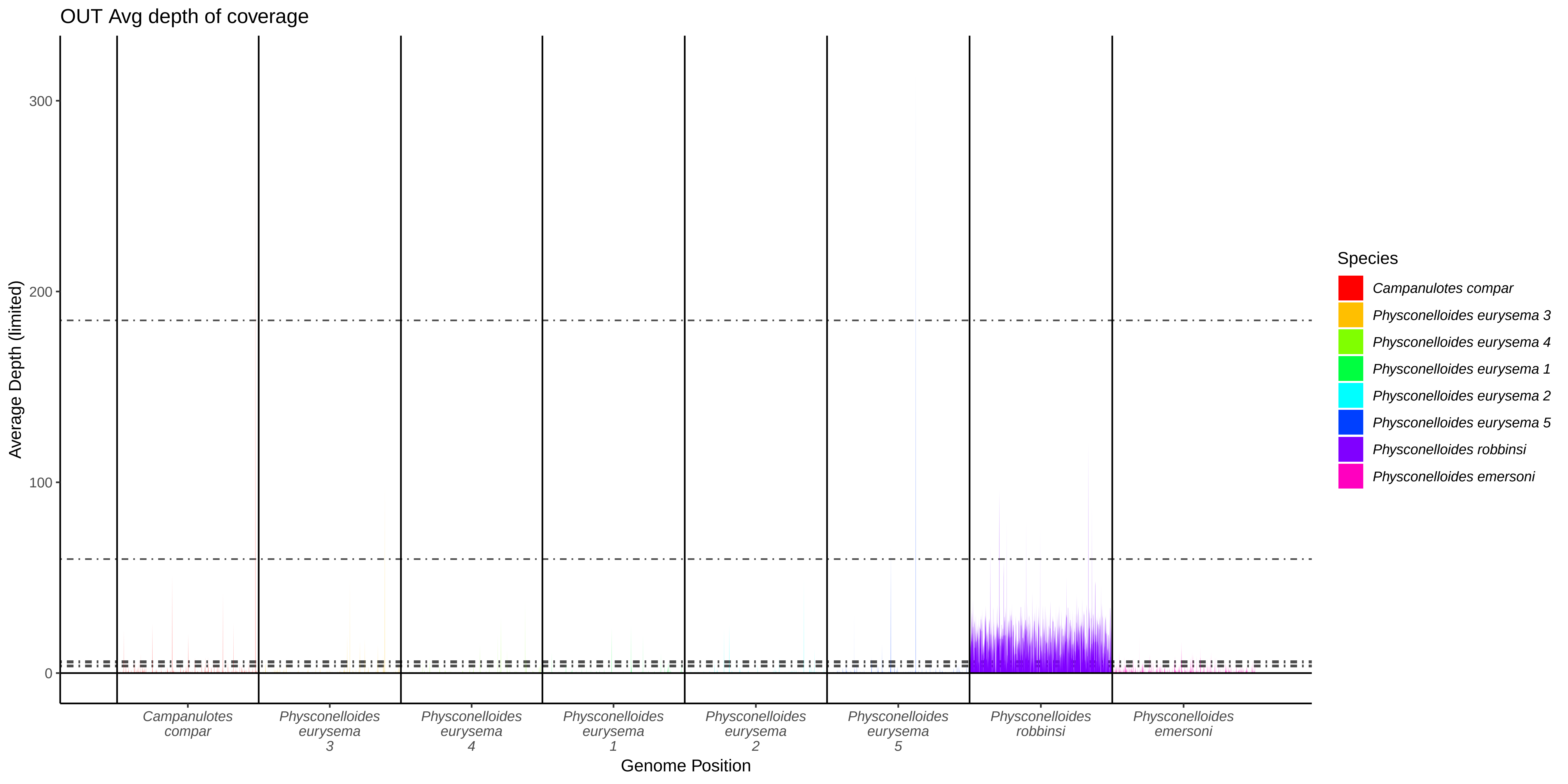

OUT Avg depth of coverage
Average Depth (limited)
300
200
100
0
Campanulotes compar
Physconelloides eurysema 3
Physconelloides eurysema 4
Physconelloides eurysema 1
Physconelloides eurysema 2
Physconelloides eurysema 5
Physconelloides robbinsi
Physconelloides emersoni
Genome Position
Species
Campanulotes compar
Physconelloides eurysema 3
Physconelloides eurysema 4
Physconelloides eurysema 1
Physconelloides eurysema 2
Physconelloides eurysema 5
Physconelloides robbinsi
Physconelloides emersoni

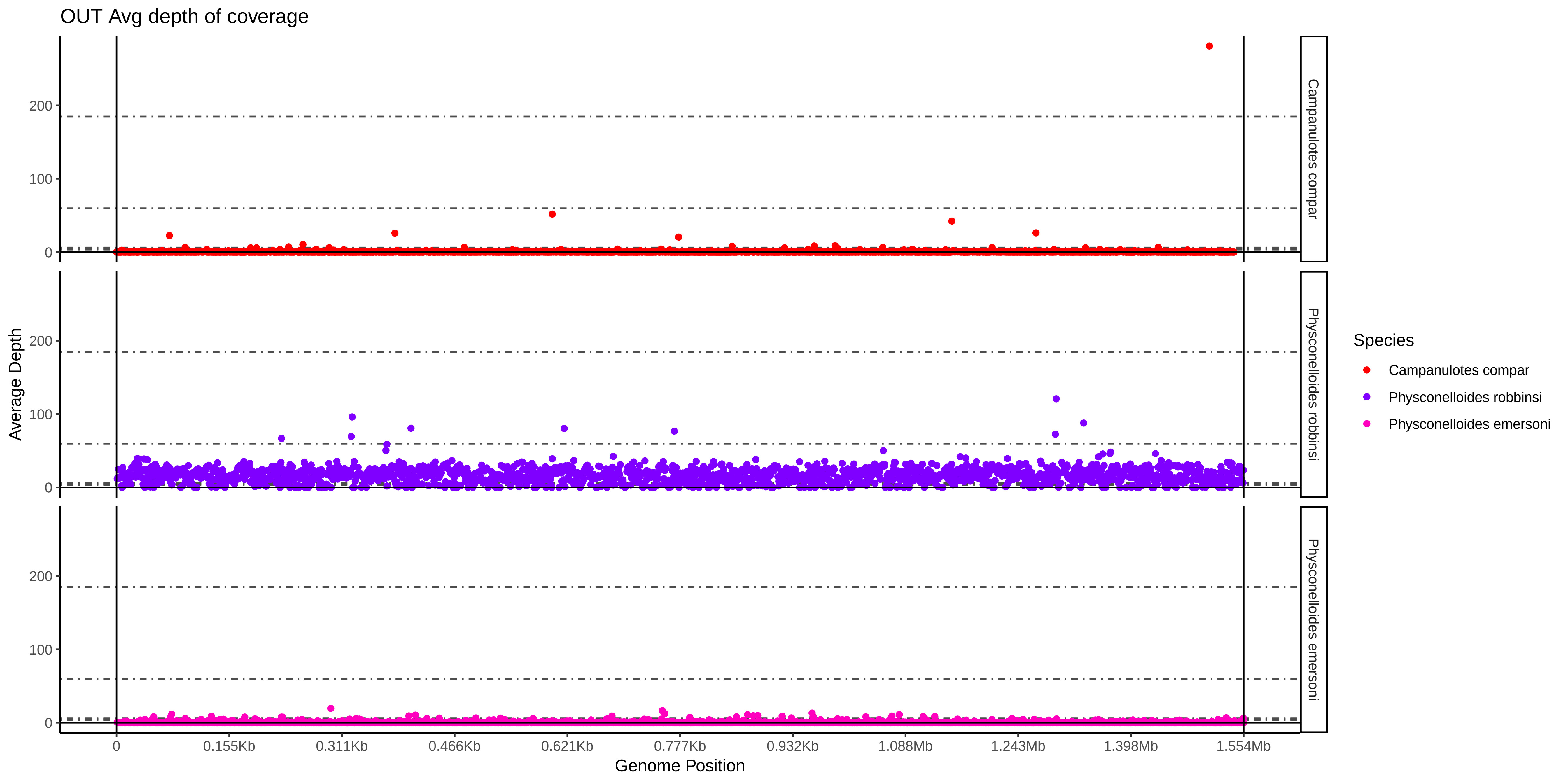

OUT Avg depth of coverage
Average Depth
Genome Position
0
100
200
0
0.155Kb
0.311Kb
0.466Kb
0.621Kb
0.777Kb
0.932Kb
1.088Mb
1.243Mb
1.398Mb
1.554Mb
Campanulotes compar
Physconelloides robbinsi
Physconelloides emersoni
Species
Campanulotes compar
Physconelloides robbinsi
Physconelloides emersoni

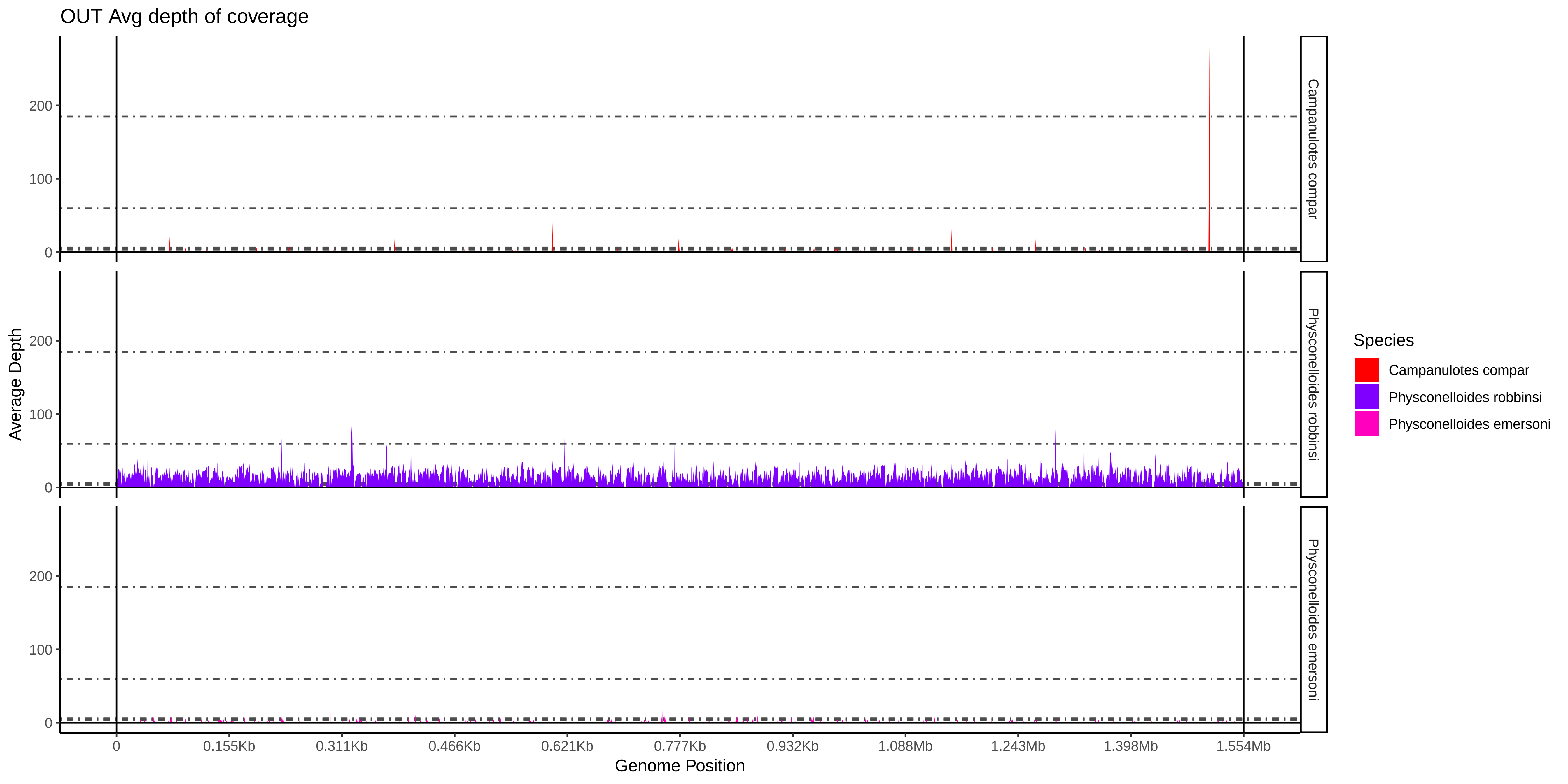

OUT Avg depth of coverage
Average Depth
Genome Position
0
100
200
0
0.155Kb
0.311Kb
0.466Kb
0.621Kb
0.777Kb
0.932Kb
1.088Mb
1.243Mb
1.398Mb
1.554Mb
Campanulotes compar
Physconelloides robbinsi
Physconelloides emersoni
Species
Campanulotes compar
Physconelloides robbinsi
Physconelloides emersoni

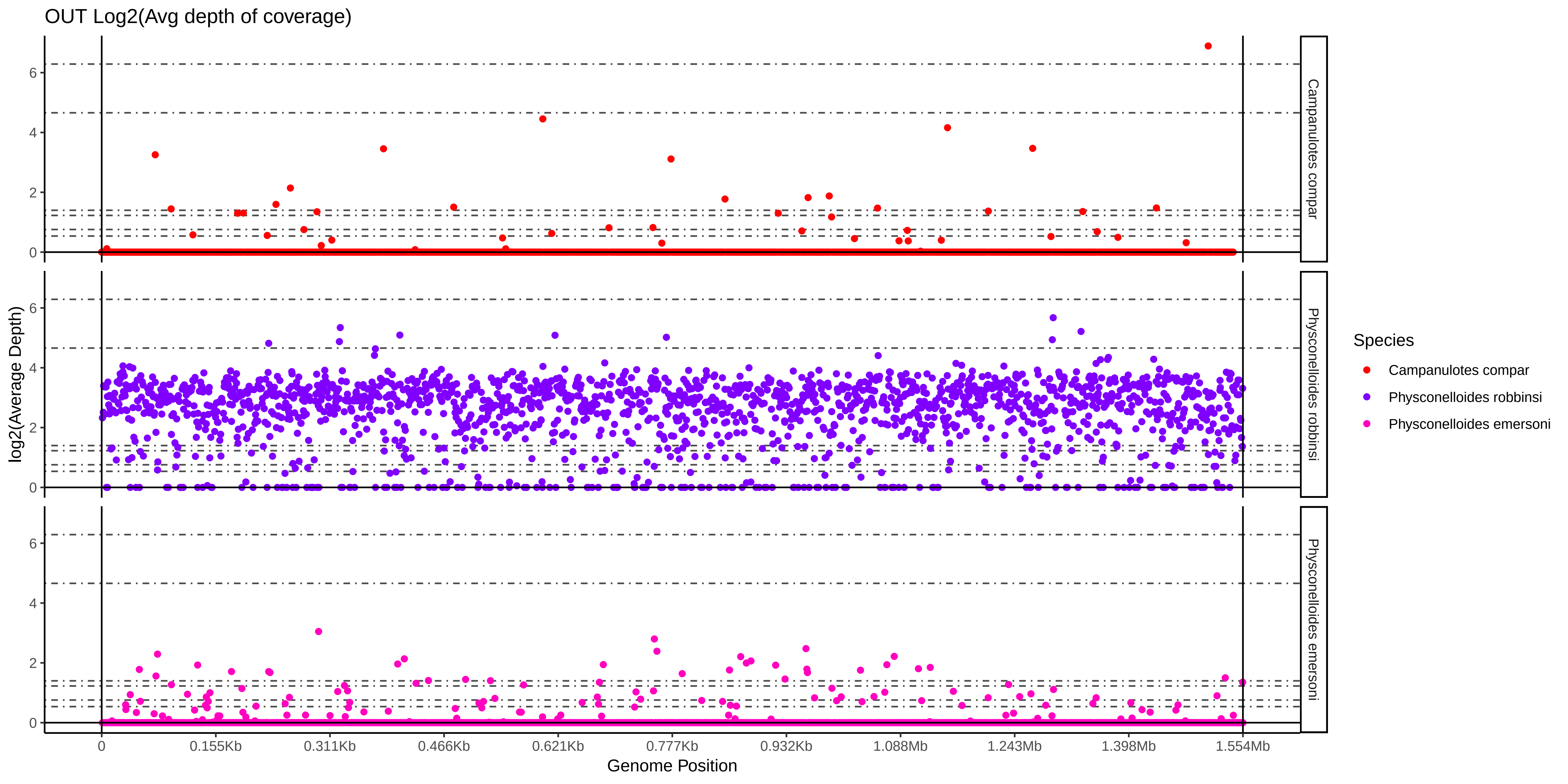

OUT Log2(Avg depth of coverage)
log2(Average Depth)
0
2
4
6
Campanulotes compar
Physconelloides robbinsi
Physconelloides emersoni
0
0.155Kb
0.311Kb
0.466Kb
0.621Kb
0.777Kb
0.932Kb
1.088Mb
1.243Mb
1.398Mb
1.554Mb
Genome Position
Species
Campanulotes compar
Physconelloides robbinsi
Physconelloides emersoni

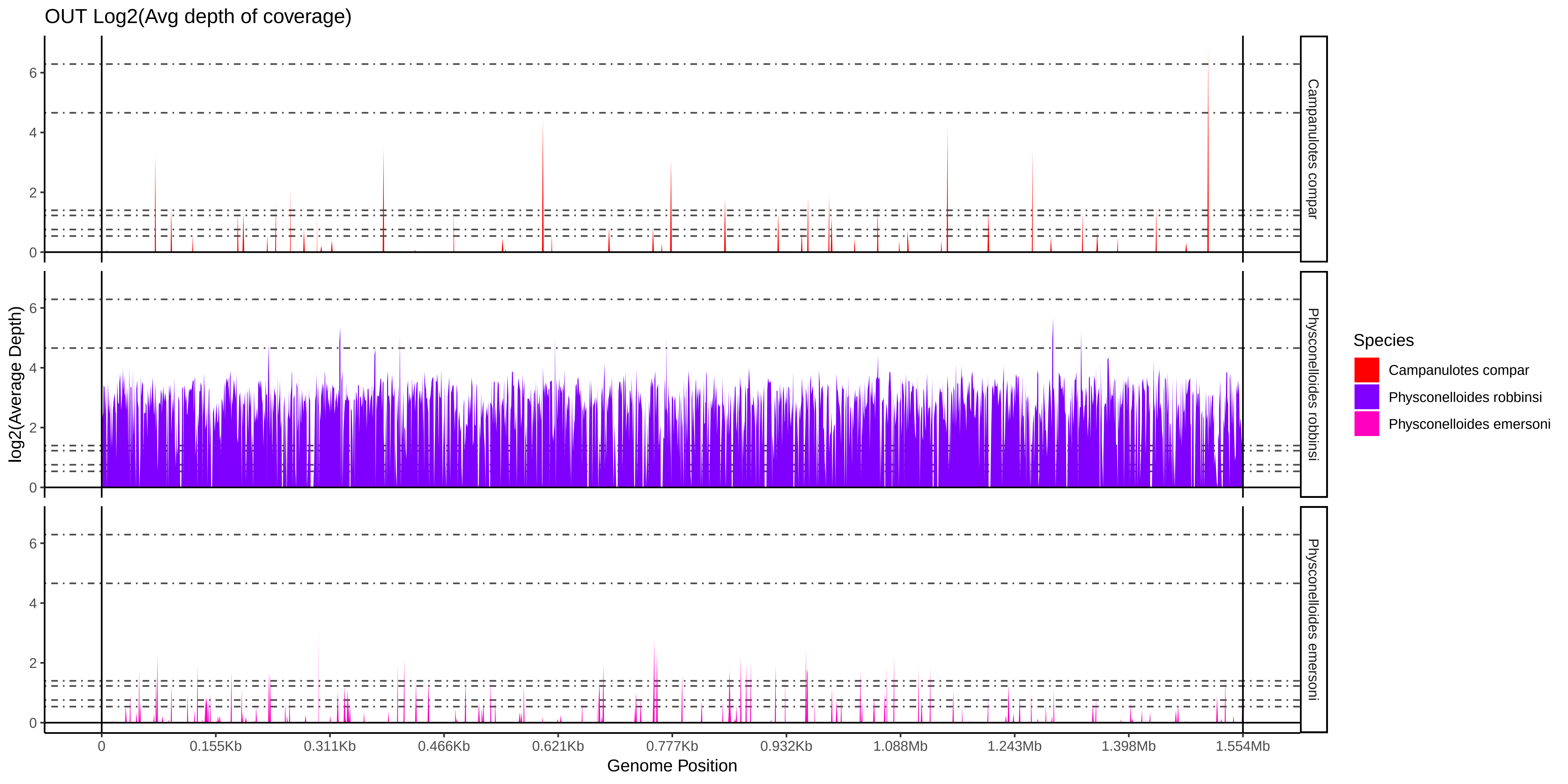

OUT Log2(Avg depth of coverage)
log2(Average Depth)
0
2
4
6
Campanulotes compar
Physconelloides robbinsi
Physconelloides emersoni
0
0.155Kb
0.311Kb
0.466Kb
0.621Kb
0.777Kb
0.932Kb
1.088Mb
1.243Mb
1.398Mb
1.554Mb
Genome Position
Species
Campanulotes compar
Physconelloides robbinsi
Physconelloides emersoni

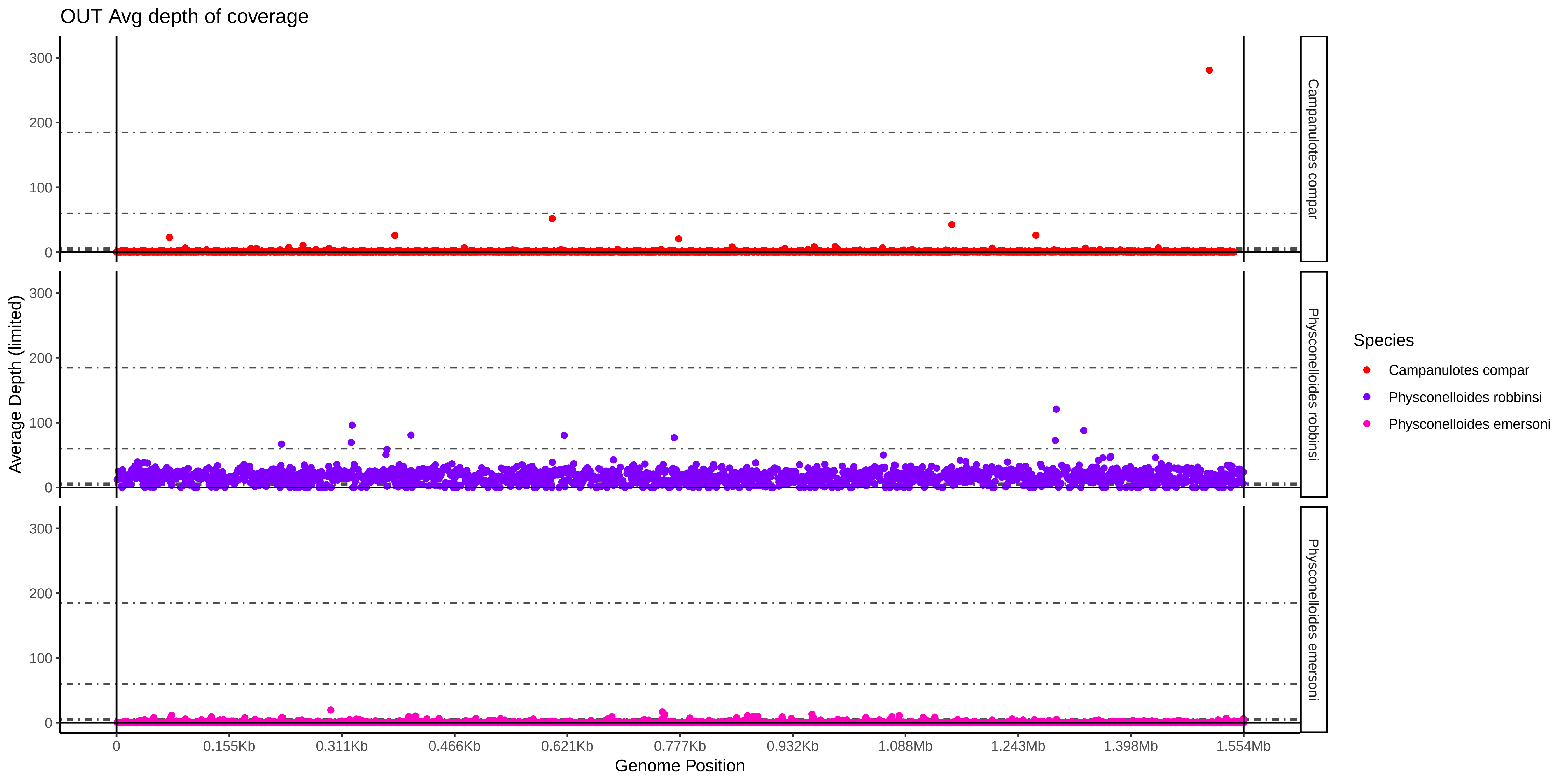
OUT Avg depth of coverage
Average Depth (limited)
300
200
100
0
Campanulotes compar
Physconelloides robbinsi
Physconelloides emersoni
0
0.155Kb
0.311Kb
0.466Kb
0.621Kb
0.777Kb
0.932Kb
1.088Mb
1.243Mb
1.398Mb
1.554Mb
Genome Position
Species
Campanulotes compar
Physconelloides robbinsi
Physconelloides emersoni

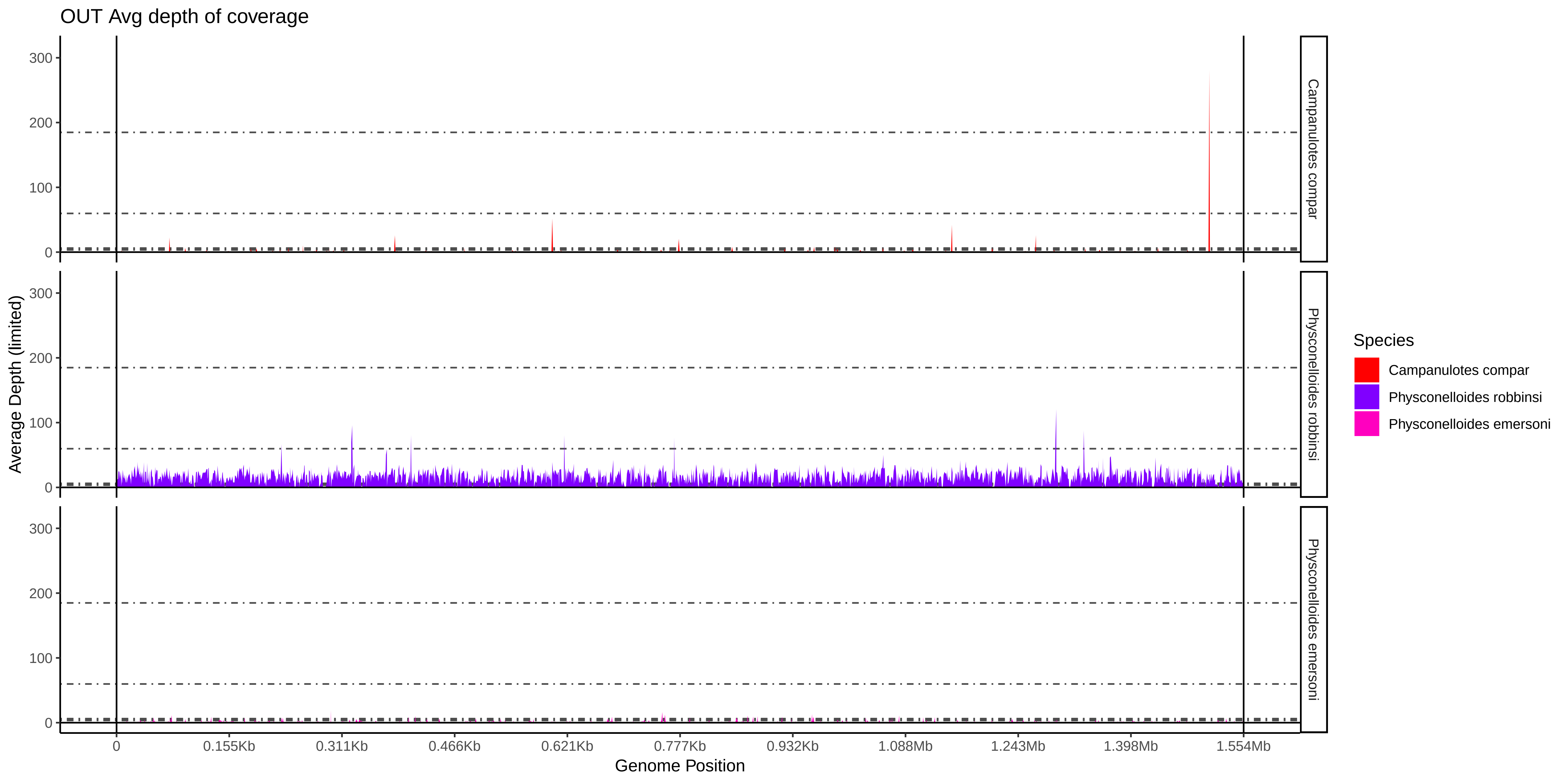

OUT Avg depth of coverage
Average Depth (limited)
300
200
100
0
Campanulotes compar
Physconelloides robbinsi
Physconelloides emersoni
0
0.155Kb
0.311Kb
0.466Kb
0.621Kb
0.777Kb
0.932Kb
1.088Mb
1.243Mb
1.398Mb
1.554Mb
Genome Position
Species
Campanulotes compar
Physconelloides robbinsi
Physconelloides emersoni
