## Supplementary material for "Comparing rates of introgression in parasitic feather lice with differing dispersal capabilities": supplementary.html (from another version of this preprint: version 7, 2020-03-31)

Table S1. Results of the GLMs of each iteration of the R simulation.

| Iteration | F | P-value |
| --- | --- | --- |
| 1 | 15.22433 | 0.0002474 |
| 2 | 13.64784 | 0.0004843 |
| 3 | 14.69258 | 0.0003097 |
| 4 | 16.47615 | 0.0001470 |
| 5 | 17.23495 | 0.0001077 |
| 6 | 16.29551 | 0.0001584 |
| 7 | 15.75052 | 0.0001985 |
| 8 | 15.26520 | 0.0002432 |
| 9 | 15.68247 | 0.0002042 |
| 10 | 14.58683 | 0.0003239 |
| 11 | 14.39204 | 0.0003519 |
| 12 | 16.78047 | 0.0001297 |
| 13 | 16.06236 | 0.0001744 |
| 14 | 16.05789 | 0.0001747 |
| 15 | 14.26203 | 0.0003720 |
| 16 | 13.41263 | 0.0005362 |
| 17 | 13.80318 | 0.0004529 |
| 18 | 16.33000 | 0.0001561 |
| 19 | 15.24110 | 0.0002457 |
| 20 | 15.03614 | 0.0002678 |
| 21 | 15.21652 | 0.0002482 |
| 22 | 14.87141 | 0.0002871 |
| 23 | 15.02394 | 0.0002692 |
| 24 | 16.83916 | 0.0001266 |
| 25 | 15.90099 | 0.0001865 |
| 26 | 14.32882 | 0.0003615 |
| 27 | 17.12290 | 0.0001128 |
| 28 | 14.97626 | 0.0002747 |
| 29 | 15.28071 | 0.0002416 |
| 30 | 15.67908 | 0.0002045 |
| 31 | 15.41777 | 0.0002281 |
| 32 | 15.31815 | 0.0002379 |
| 33 | 12.31456 | 0.0008673 |
| 34 | 15.79259 | 0.0001951 |
| 35 | 15.19614 | 0.0002504 |
| 36 | 14.85553 | 0.0002890 |
| 37 | 15.36649 | 0.0002331 |
| 38 | 15.89650 | 0.0001868 |
| 39 | 15.42534 | 0.0002274 |
| 40 | 15.10801 | 0.0002598 |
| 41 | 14.03679 | 0.0004096 |
| 42 | 16.80676 | 0.0001283 |
| 43 | 14.33955 | 0.0003599 |
| 44 | 16.14082 | 0.0001688 |
| 45 | 16.53914 | 0.0001432 |
| 46 | 17.64362 | 0.0000913 |
| 47 | 16.54421 | 0.0001429 |
| 48 | 16.77791 | 0.0001299 |
| 49 | 15.22370 | 0.0002475 |
| 50 | 14.62479 | 0.0003187 |
| 51 | 15.02217 | 0.0002694 |
| 52 | 15.99477 | 0.0001793 |
| 53 | 15.29044 | 0.0002406 |
| 54 | 16.55918 | 0.0001421 |
| 55 | 13.32214 | 0.0005576 |
| 56 | 15.82790 | 0.0001922 |
| 57 | 12.96108 | 0.0006527 |
| 58 | 16.08644 | 0.0001727 |
| 59 | 16.05871 | 0.0001747 |
| 60 | 18.04664 | 0.0000776 |
| 61 | 13.76757 | 0.0004599 |
| 62 | 15.72709 | 0.0002005 |
| 63 | 15.55908 | 0.0002150 |
| 64 | 15.08537 | 0.0002623 |
| 65 | 11.51995 | 0.0012361 |
| 66 | 14.26142 | 0.0003721 |
| 67 | 15.08217 | 0.0002627 |
| 68 | 15.79173 | 0.0001951 |
| 69 | 15.93359 | 0.0001840 |
| 70 | 14.34038 | 0.0003597 |
| 71 | 15.09003 | 0.0002618 |
| 72 | 15.70749 | 0.0002021 |
| 73 | 14.95929 | 0.0002766 |
| 74 | 15.16627 | 0.0002535 |
| 75 | 17.04410 | 0.0001165 |
| 76 | 16.69256 | 0.0001345 |
| 77 | 16.34819 | 0.0001549 |
| 78 | 11.48875 | 0.0012535 |
| 79 | 16.56616 | 0.0001416 |
| 80 | 12.98989 | 0.0006445 |
| 81 | 15.07607 | 0.0002633 |
| 82 | 15.01286 | 0.0002705 |
| 83 | 14.67114 | 0.0003125 |
| 84 | 13.92249 | 0.0004302 |
| 85 | 16.64480 | 0.0001371 |
| 86 | 16.42326 | 0.0001502 |
| 87 | 14.06429 | 0.0004048 |
| 88 | 15.80926 | 0.0001937 |
| 89 | 17.71282 | 0.0000888 |
| 90 | 15.62020 | 0.0002096 |
| 91 | 15.02338 | 0.0002693 |
| 92 | 16.47573 | 0.0001470 |
| 93 | 14.00941 | 0.0004145 |
| 94 | 16.38504 | 0.0001526 |
| 95 | 12.44270 | 0.0008196 |
| 96 | 12.98249 | 0.0006466 |
| 97 | 16.45955 | 0.0001480 |
| 98 | 17.03561 | 0.0001169 |
| 99 | 15.85739 | 0.0001899 |
| 100 | 14.18589 | 0.0003843 |


Table S2. Specimen information, extraction results, library preparation details, Illumina sequencing statistics, locus assembly, and raw sequence data deposition for Columbicola wing lice from small New World ground-doves.

| Louse.taxa | Extract.Code | Host.species | Host.Collection.code | Collection.location | Collection.date | DNA.extraction.date | DNA.yield..ng.uL. | Sonication.instrument | Library.preparation.kit | Sequencing.platform | Sequencing.Kit | Sequencing.cycles | Bar.code | Sequencing.pool | Read.length | Paired.end | Number.of.raw.Reads | Predicted.sequencing.depth | Read.duplication | Number.of.reads.after.pre.processing | Sequencing.depth.after.pre.processing | Number.of.genes.assembled.with.BowTie2..1.058.Genes.Targeted. | NCBI.SRA | NCBI.BioSample | NCBI.BioProject | Chosen.for.Phylonet.analyses..1.means.yes. |
| --- | --- | --- | --- | --- | --- | --- | --- | --- | --- | --- | --- | --- | --- | --- | --- | --- | --- | --- | --- | --- | --- | --- | --- | --- | --- | --- |
| Physconelloides emersoni emersoni | Pheme.Memel.5.26.2015.8 | Metriopelia melanoptera | KGM445 | Argentina: Jujay | 11-Oct-2001 | 26-May-15 | 0.853 | Covaris M220 | Kapa Biosystems | Illumina HiSeq2500 | HiSeq SBS V. 4 | 161 | ATTCCT | 4 | 160nt | yes | 118843526 | 95.07000 | 23.70 | 96244932 | 77.00000 | 1058 | SRP076185 | SAMN04633184 | PRJNA318056 | 1 |
| Physconelloides emersoni emersoni | Pheme.Memel.6.15.2015.5 | Metriopelia melanoptera | KGM511 | Bolivia: Lago Poopo, Oruro | 2-Nov-2001 | 15-Jun-15 | 0.379 | Covaris M220 | Kapa Biosystems | Illumina HiSeq2500 | HiSeq SBS V. 4 | 161 | CAGATC | 1 | 160nt | yes | 78198304 | 62.56000 | 37.67 | 50415214 | 40.33000 | 1052 | SRP076185 | SAMN04633188 | PRJNA318056 | 0 |
| Physconelloides emersoni emersoni | Pheme.Memel.3.16.2016.2 | Metriopelia melanoptera | REW273 | Peru: Puno | 21-Oct-2002 | 16-Mar-16 | 0.417 | Covaris M220 | Kapa Biosystems | Illumina HiSeq2500 | HiSeq SBS V. 4 | 161 | CGTCTCGTGA | 2 | 160nt | yes | 63517384 | 50.81000 | 17.90 | 51240446 | 40.99000 | 1057 | SRP076185 | SAMN05099560 | PRJNA318056 | 0 |
| Physconelloides emersoni emersoni | Pheme.Memel.3.16.2016.8 | Metriopelia melanoptera | ASEC004 | Ecuador: Pichincha | 23-Jan-2016 | 16-Mar-16 | 0.333 | Covaris M220 | Kapa Biosystems | Illumina HiSeq2500 | HiSeq SBS V. 4 | 161 | GAGAGTACTA | 3 | 160nt | yes | 36399076 | 29.12000 | 12.17 | 28968590 | 23.17000 | 1058 | SRP076185 | SAMN05099566 | PRJNA318056 | 0 |
| Physconelloides eurysema eurysema 3 | Pheur.Clpre.3.14.2016.5 | Claravis pretiosa | 5172 | Brazil: Para | 2-Jun-2013 | 14-Mar-16 | 0.579 | Covaris M220 | Kapa Biosystems | Illumina HiSeq2500 | HiSeq SBS V. 4 | 161 | CCGGTGGAAT | 1 | 160nt | yes | 35524478 | 28.42000 | 26.60 | 27317070 | 21.85000 | 1050 | SRP076185 | SAMN05099551 | PRJNA318056 | 0 |
| Physconelloides eurysema eurysema 3 | Pheur.Clpre.3.14.2016.11 | Claravis pretiosa | GES470 | Mexico: Campeche | 16-Mar-1998 | 14-Mar-16 | 0.212 | Covaris M220 | Kapa Biosystems | Illumina HiSeq2500 | HiSeq SBS V. 4 | 161 | TCTCGACGTT | 2 | 160nt | yes | 42423630 | 33.94000 | 27.43 | 33041298 | 26.43000 | 1058 | SRP076185 | SAMN05099557 | PRJNA318056 | 1 |
| Physconelloides eurysema eurysema 3 | Pheur.Clpre.3.16.2016.5 | Claravis pretiosa | 6414 | Brazil: Para | 1-Aug-2014 | 16-Mar-16 | 0.090 | Covaris M220 | Kapa Biosystems | Illumina HiSeq2500 | HiSeq SBS V. 4 | 161 | AGGAGAATCA | 3 | 160nt | yes | 40120794 | 32.10000 | 14.05 | 31896742 | 25.52000 | 1058 | SRP076185 | SAMN05099563 | PRJNA318056 | 0 |
| Physconelloides eurysema eurysema 3 | Pheur.Clpre.5.26.2015.6 | Claravis pretiosa | GES382 | Mexico: Campeche | 16-Mar-1998 | 26-May-15 | 0.973 | Covaris M220 | Kapa Biosystems | Illumina HiSeq2500 | HiSeq SBS V. 4 | 161 | GAGTGG | 4 | 160nt | yes | 63821562 | 51.06000 | 20.82 | 49744786 | 39.80000 | 1058 | SRP076185 | SAMN04633182 | PRJNA318056 | 0 |
| Physconelloides eurysema eurysema 3 | Pheur.Clpre.6.15.2015.8 | Claravis pretiosa | REW176 | Peru: Cajamarca | 7-Sep-2002 | 15-Jun-15 | 0.189 | Covaris M220 | Kapa Biosystems | Illumina HiSeq2500 | HiSeq SBS V. 4 | 161 | TTAGGC | 1 | 160nt | yes | 51392388 | 41.11000 | 17.67 | 37130950 | 29.70000 | 1058 | SRP076185 | SAMN04633191 | PRJNA318056 | 0 |
| Physconelloides eurysema eurysema 3 | Pheur.Cobuc.3.16.2016.4 | Columbina buckleyi | RCF998 | Peru: Cajamarca | 15-Feb-1999 | 16-Mar-16 | 0.522 | Covaris M220 | Kapa Biosystems | Illumina HiSeq2500 | HiSeq SBS V. 4 | 161 | ACTTCCGCAC | 3 | 160nt | yes | 36856494 | 29.49000 | 10.56 | 33206986 | 26.57000 | 1058 | SRP076185 | SAMN05099562 | PRJNA318056 | 0 |
| Physconelloides eurysema eurysema 3 | Pheur.Cobuc.6.15.2015.1 | Columbina buckleyi | REW170 | Peru: Cajamarca | 5-Sep-2002 | 15-Jun-15 | 0.515 | Covaris M220 | Kapa Biosystems | Illumina HiSeq2500 | HiSeq SBS V. 4 | 161 | CGATGT | 1 | 160nt | yes | 61157596 | 48.93000 | 51.75 | 36504910 | 29.20000 | 1058 | SRP076185 | SAMN04633185 | PRJNA318056 | 0 |
| Physconelloides eurysema eurysema 3 | Pheur.Cocru.3.16.2016.9 | Columbina cruziana | ASEC008 | Ecuador: Manabi | 29-Jan-2016 | 16-Mar-16 | 0.481 | Covaris M220 | Kapa Biosystems | Illumina HiSeq2500 | HiSeq SBS V. 4 | 161 | GCTCCTCTTG | 3 | 160nt | yes | 44058524 | 35.25000 | 7.95 | 35529848 | 28.42000 | 1057 | SRP076185 | SAMN05099567 | PRJNA318056 | 0 |
| Physconelloides eurysema eurysema 3 | Pheur.Cocru.6.15.2015.2 | Columbina cruziana | REW85 | Peru: Lima | 8-Aug-2002 | 15-Jun-15 | 0.887 | Covaris M220 | Kapa Biosystems | Illumina HiSeq2500 | HiSeq SBS V. 4 | 161 | TGACCA | 1 | 160nt | yes | 61326000 | 49.06000 | 71.40 | 26000756 | 20.80000 | 1046 | SRP076185 | SAMN04633186 | PRJNA318056 | 0 |
| Physconelloides eurysema eurysema 3 | Pheur.Comin.3.14.2016.9 | Columbina minuta | JK06-367 | Panama: Charco Azul | 2-Mar-2006 | 14-Mar-16 | 0.499 | Covaris M220 | Kapa Biosystems | Illumina HiSeq2500 | HiSeq SBS V. 4 | 161 | TGTTACCTAC | 2 | 160nt | yes | 32144260 | 25.72000 | 25.74 | 24668212 | 19.73000 | 1058 | SRP076185 | SAMN05099555 | PRJNA318056 | 0 |
| Physconelloides eurysema eurysema 3 | Pheur.Copas.3.14.2016.4 | Columbina passerina | 5048 | Brazil: Amazonas | 11-Jul-2013 | 14-Mar-16 | 0.520 | Covaris M220 | Kapa Biosystems | Illumina HiSeq2500 | HiSeq SBS V. 4 | 161 | ACCGCCTATT | 1 | 160nt | yes | 41362350 | 33.09000 | 13.07 | 33206986 | 26.57000 | 1056 | SRP076185 | SAMN05099550 | PRJNA318056 | 0 |
| Physconelloides eurysema eurysema 3 | Pheur.Copas.3.14.2016.7 | Columbina passerina | 5050 | Brazil: Amazonas | 11-Jul-2013 | 14-Mar-16 | 0.294 | Covaris M220 | Kapa Biosystems | Illumina HiSeq2500 | HiSeq SBS V. 4 | 161 | GCAGGCGTAT | 1 | 160nt | yes | 48354288 | 38.68000 | 17.09 | 38250780 | 30.60000 | 1050 | SRP076185 | SAMN05099553 | PRJNA318056 | 0 |
| Physconelloides eurysema eurysema 3 | Pheur.Copas.3.14.2016.10 | Columbina passerina | GES233 | Mexico: Campeche | 2-Mar-1998 | 14-Mar-16 | 0.323 | Covaris M220 | Kapa Biosystems | Illumina HiSeq2500 | HiSeq SBS V. 4 | 161 | TAGCGGCACT | 2 | 160nt | yes | 46626818 | 37.30000 | 30.13 | 35850772 | 28.68000 | 1058 | SRP076185 | SAMN05099556 | PRJNA318056 | 0 |
| Physconelloides eurysema eurysema 3 | Pheur.Copas.3.16.2016.1 | Columbina passerina | 6469 | Brazil: Rondonia | 7-Oct-2014 | 16-Mar-16 | 0.610 | Covaris M220 | Kapa Biosystems | Illumina HiSeq2500 | HiSeq SBS V. 4 | 161 | TAAGAGCGCC | 2 | 160nt | yes | 49146294 | 39.32000 | 10.47 | 39663616 | 31.73000 | 1058 | SRP076185 | SAMN05099559 | PRJNA318056 | 0 |
| Physconelloides eurysema eurysema 3 | Pheur.Copas.3.16.2016.7 | Columbina passerina | ASEC001 | Ecuador: Pichincha, Calacali | 20-Jan-2016 | 16-Mar-16 | 0.387 | Covaris M220 | Kapa Biosystems | Illumina HiSeq2500 | HiSeq SBS V. 4 | 161 | CTACAACGGC | 3 | 160nt | yes | 35742462 | 28.59000 | 10.13 | 28679204 | 22.94000 | 1058 | SRP076185 | SAMN05099565 | PRJNA318056 | 0 |
| Physconelloides eurysema eurysema 3 | Pheur.Cotal.3.14.2016.2 | Columbina talpacoti | 4741 | Brazil: Santa Catarina | 31-Oct-2013 | 14-Mar-16 | 0.197 | Covaris M220 | Kapa Biosystems | Illumina HiSeq2500 | HiSeq SBS V. 4 | 161 | TTGTTGGCGG | 1 | 160nt | yes | 24481602 | 19.59000 | 38.76 | 17876904 | 14.30000 | 1048 | SRP076185 | SAMN05099548 | PRJNA318056 | 0 |
| Physconelloides eurysema eurysema 3 | Pheur.Cotal.3.14.2016.6 | Columbina talpacoti | GMS1950 | Panama: Torti | 18-Feb-2006 | 14-Mar-16 | 0.731 | Covaris M220 | Kapa Biosystems | Illumina HiSeq2500 | HiSeq SBS V. 4 | 161 | CTAACGATCG | 1 | 160nt | yes | 49810804 | 39.85000 | 12.18 | 40316768 | 32.25000 | 1058 | SRP076185 | SAMN05099552 | PRJNA318056 | 0 |
| Physconelloides eurysema eurysema 3 | Pheur.Cotal.3.16.2016.3 | Columbina talpacoti | 6370 | Brazil: Sao Paulo | 1-Jun-2014 | 16-Mar-16 | 0.231 | Covaris M220 | Kapa Biosystems | Illumina HiSeq2500 | HiSeq SBS V. 4 | 161 | TTCTCCTTCA | 2 | 160nt | yes | 51734340 | 41.39000 | 14.29 | 41941080 | 33.55000 | 1050 | SRP076185 | SAMN05099561 | PRJNA318056 | 0 |
| Physconelloides eurysema eurysema 3 | Pheur.Cotal.3.16.2016.12 | Columbina talpacoti | SMD739 | Panama: Felipillo | 19-Feb-2006 | 16-Mar-16 | 0.207 | Covaris M220 | Kapa Biosystems | Illumina HiSeq2500 | HiSeq SBS V. 4 | 161 | ACATGGTCCA | 3 | 160nt | yes | 34980054 | 27.98000 | 11.28 | 27853084 | 22.28000 | 1057 | SRP076185 | SAMN05099570 | PRJNA318056 | 0 |
| Physconelloides eurysema eurysema 4 | Pheur.Coinc.6.15.2015.7 | Columbina inca | MCZ 01-149 | Costa Rica: Guanacaste | 10-Dec-2001 | 15-Jun-15 | 0.887 | Covaris M220 | Kapa Biosystems | Illumina HiSeq2500 | HiSeq SBS V. 4 | 161 | ATCACG | 1 | 160nt | yes | 53047444 | 42.44000 | 22.31 | 34924128 | 27.94000 | 1052 | SRP076185 | SAMN04633190 | PRJNA318056 | 1 |
| Physconelloides eurysema eurysema 1 | Pheur.Comin.6.15.2015.10 | Columbina minuta | MCZ 01-335 | Costa Rica: Guanacaste | 10-Dec-2001 | 15-Jun-15 | 0.363 | Covaris M220 | Kapa Biosystems | Illumina HiSeq2500 | HiSeq SBS V. 4 | 161 | GATCAG | 1 | 160nt | yes | 46622380 | 37.30000 | 12.96 | 31194444 | 24.96000 | 1052 | SRP076185 | SAMN04633192 | PRJNA318056 | 0 |
| Physconelloides eurysema eurysema 1 | Pheur.Copas.3.14.2016.8 | Columbina passerina | 299 | USA: Texas (captive) | 27-Apr-2000 | 14-Mar-16 | 0.967 | Covaris M220 | Kapa Biosystems | Illumina HiSeq2500 | HiSeq SBS V. 4 | 161 | GCCTTCAGGA | 2 | 160nt | yes | 44144898 | 35.32000 | 8.11 | 35767700 | 28.61000 | 1058 | SRP076185 | SAMN05099554 | PRJNA318056 | 1 |
| Physconelloides eurysema eurysema 1 | Pheur.Copas.3.16.2016.6 | Columbina passerina |  | USA: Illinois, McClean, Bloomington | 19-Oct-2015 | 16-Mar-16 | 0.105 | Covaris M220 | Kapa Biosystems | Illumina HiSeq2500 | HiSeq SBS V. 4 | 161 | AGCTCAACGA | 3 | 160nt | yes | 40646734 | 32.52000 | 19.80 | 31553360 | 25.24000 | 1050 | SRP076185 | SAMN05099564 | PRJNA318056 | 0 |
| Physconelloides eurysema eurysema 2 | Pheur.Cosqu.3.14.2016.1 | Columbina squammata | 4702 | Brazil: Para | 1-Sep-2010 | 14-Mar-16 | 0.464 | Covaris M220 | Kapa Biosystems | Illumina HiSeq2500 | HiSeq SBS V. 4 | 161 | AGTATAGCGC | 1 | 160nt | yes | 33666308 | 26.93000 | 23.60 | 25793048 | 20.63000 | 1058 | SRP076185 | SAMN05099547 | PRJNA318056 | 1 |
| Physconelloides eurysema eurysema 2 | Pheur.Cosqu.3.14.2016.12 | Columbina squammata | 4780 | Brazil: Para | 15-Aug-2013 | 14-Mar-16 | 0.162 | Covaris M220 | Kapa Biosystems | Illumina HiSeq2500 | HiSeq SBS V. 4 | 161 | GGCGAAGGTT | 2 | 160nt | yes | 49091816 | 39.27000 | 41.21 | 35989584 | 28.79000 | 1054 | SRP076185 | SAMN05099558 | PRJNA318056 | 0 |
| Physconelloides eurysema eurysema 5 | Pheur.Urcam.3.14.2016.3 | Uropelia campestris | 4887 | Brazil: Para | 20-Aug-2013 | 14-Mar-16 | 2.500 | Covaris M220 | Kapa Biosystems | Illumina HiSeq2500 | HiSeq SBS V. 4 | 161 | AACCGAGTTA | 1 | 160nt | yes | 43813494 | 35.05000 | 16.04 | 34834472 | 27.87000 | 1058 | SRP076185 | SAMN05099549 | PRJNA318056 | 1 |
| Physconelloides robbinsi robbinsi | Phrob.Mecec.5.26.2015.7 | Metriopelia ceciliae | REW153 | Peru: Caras | 28-Aug-2002 | 26-May-15 | 1.280 | Covaris M220 | Kapa Biosystems | Illumina HiSeq2500 | HiSeq SBS V. 4 | 161 | ACTGAT | 4 | 160nt | yes | 80808714 | 64.65000 | 17.87 | 63931800 | 51.15000 | 1058 | SRP076185 | SAMN04633183 | PRJNA318056 | 1 |
| Physconelloides robbinsi robbinsi | Phrob.Mecec.6.15.2015.6 | Metriopelia ceciliae | REW151 | Peru: Caras | 28-Aug-2002 | 15-Jun-15 | 0.940 | Covaris M220 | Kapa Biosystems | Illumina HiSeq2500 | HiSeq SBS V. 4 | 161 | CTTGTA | 1 | 160nt | yes | 13580582 | 10.86000 | 39.22 | 7260590 | 5.81000 | 1032 | SRP076185 | SAMN04633189 | PRJNA318056 | 0 |
| Physconelloides robbinsi robbinsi | Phrob.Mecec.3.16.2016.10 | Metriopelia ceciliae | REW150 | Peru: Caras | 28-Aug-2002 | 16-Mar-16 | 0.167 | Covaris M220 | Kapa Biosystems | Illumina HiSeq2500 | HiSeq SBS V. 4 | 161 | TAGGCATCGG | 3 | 160nt | yes | 34801206 | 27.84000 | 19.67 | 27053634 | 21.64000 | 1058 | SRP076185 | SAMN05099568 | PRJNA318056 | 0 |
| Physconelloides robbinsi robbinsi | Phrob.Mecec.3.16.2016.11 | Metriopelia ceciliae | REW294 | Peru: Tacna | 25-Oct-2002 | 16-Mar-16 | 0.257 | Covaris M220 | Kapa Biosystems | Illumina HiSeq2500 | HiSeq SBS V. 4 | 161 | CAACCTAACA | 3 | 160nt | yes | 33983950 | 27.19000 | 14.41 | 26808390 | 21.45000 | 1058 | SRP076185 | SAMN05099569 | PRJNA318056 | 0 |
| Columbicola passerinae passerinae 1 | Cosp.Coinc.11.2.2016.9 | Columbina inca | MCZ 01-149 | Costa Rica | 10-Dec-2001 | 2-Nov-2016 | 0.324 | Covaris M220 | Hyper Library (Kapa Biosystems) | Illumina HiSeq4000 | HiSeq4000 v.1 | 151 | TCCTTAACCCGG | 2 | 150 bp | yes | 45454206 | 34.09065 | 15.47 | 34876801 | 26.15760 | 1031 | SRR5998819 | SAMN07573751 | PRJNA400795 | 0 |
| Columbicola passerinae passerinae 1 | Cosp.Comin.11.2.2016.16 | Columbina minuta | JMD733 | Panama: Felipillo | 19-Feb-2006 | 2-Nov-2016 | 0.501 | Covaris M220 | Hyper Library (Kapa Biosystems) | Illumina HiSeq4000 | HiSeq4000 v.1 | 151 | GAAAGAGAGTCC | 1 | 150 bp | yes | 47902092 | 35.92657 | 31.25 | 34645037 | 25.98378 | 1037 | SRR5998828 | SAMN07573758 | PRJNA400795 | 0 |
| Columbicola passerinae passerinae 1 | Cosp.Comin.11.2.2016.4 | Columbina minuta | JLK258 | Peru: Loreto | 15-Jun-2001 | 2-Nov-2016 | 0.231 | Covaris M220 | Hyper Library (Kapa Biosystems) | Illumina HiSeq4000 | HiSeq4000 v.1 | 151 | TGATGCAACCGG | 1 | 150 bp | yes | 31447878 | 23.58591 | 15.03 | 23729175 | 17.79688 | 1037 | SRR5998812 | SAMN07573746 | PRJNA400795 | 0 |
| Columbicola passerinae passerinae 1 | Cosp.Copas.11.7.2016.12 | Columbina passerina | GES233 | Mexico: Yucatan | 2-Mar-1998 | 7-Nov-2016 | 0.268 | Covaris M220 | Hyper Library (Kapa Biosystems) | Illumina HiSeq4000 | HiSeq4000 v.1 | 151 | GGGCCAACCCTT | 1 | 150 bp | yes | 37641540 | 28.23116 | 34.47 | 26218767 | 19.66408 | 1037 | SRR5998808 | SAMN07573769 | PRJNA400795 | 0 |
| Columbicola passerinae passerinae 1 | Cosp.Copas.11.7.2016.14 | Columbina passerina | ASEC001 | Ecuador: Pichincha: Calicali | 20-Jan-2016 | 7-Nov-2016 | 0.343 | Covaris M220 | Hyper Library (Kapa Biosystems) | Illumina HiSeq4000 | HiSeq4000 v.1 | 151 | CCCGGAACCCTT | 1 | 150 bp | yes | 45388822 | 34.04162 | 28.00 | 32058302 | 24.04373 | 1036 | SRR5998802 | SAMN07573771 | PRJNA400795 | 0 |
| Columbicola passerinae passerinae 1 | Cosp.Copas.11.7.2016.15 | Columbina passerina | n/a | USA: IL: McClean: Bloomington | 19-Oct-2015 | 7-Nov-2016 | 0.402 | Covaris M220 | Hyper Library (Kapa Biosystems) | Illumina HiSeq4000 | HiSeq4000 v.1 | 151 | CTTCCAACCCGG | 2 | 150 bp | yes | 42180932 | 31.63570 | 17.18 | 32311708 | 24.23378 | 1037 | SRR5998801 | SAMN07573772 | PRJNA400795 | 0 |
| Columbicola passerinae passerinae 1 | Cosp.Copas.11.7.2016.8 | Columbina passerina | 5049 | Brazil: Amazonas | 11-Jul-2013 | 7-Nov-2016 | 0.475 | Covaris M220 | Hyper Library (Kapa Biosystems) | Illumina HiSeq4000 | HiSeq4000 v.1 | 151 | GTACTGATCCTT | 1 | 150 bp | yes | 39632690 | 29.72452 | 16.79 | 29508201 | 22.13115 | 1037 | SRR5998804 | SAMN07573765 | PRJNA400795 | 0 |
| Columbicola passerinae passerinae 1 | Cosp.Copic.11.2.2016.1 | Columbina picui | CCW880 | Bolivia: Santa Cruz | n/a | 2-Nov-2016 | 0.219 | Covaris M220 | Hyper Library (Kapa Biosystems) | Illumina HiSeq4000 | HiSeq4000 v.1 | 151 | AACGTTGTGTCC | 1 | 150 bp | yes | 32596536 | 24.44740 | 17.31 | 24402171 | 18.30163 | 1031 | SRR5998813 | SAMN07573743 | PRJNA400795 | 0 |
| Columbicola passerinae passerinae 1 | Cosp.Copic.11.2.2016.3 | Columbina picui | KGM292 | Argentina: Cordoba | 26-Apr-2001 | 2-Nov-2016 | 0.260 | Covaris M220 | Hyper Library (Kapa Biosystems) | Illumina HiSeq4000 | HiSeq4000 v.1 | 151 | GCGGAGATCCGG | 2 | 150 bp | yes | 47314896 | 35.48617 | 12.25 | 35586948 | 26.69021 | 1036 | SRR5998811 | SAMN07573745 | PRJNA400795 | 0 |
| Columbicola passerinae passerinae 1 | Cosp.Cosqu.11.7.2016.1 | Columbina squammata | 4703 | Brazil: Para | 1-Sep-2010 | 7-Nov-2016 | 0.521 | Covaris M220 | Hyper Library (Kapa Biosystems) | Illumina HiSeq4000 | HiSeq4000 v.1 | 151 | ATTCTGAAGTCC | 2 | 150 bp | yes | 48517638 | 36.38823 | 15.39 | 36766169 | 27.57463 | 1037 | SRR5998825 | SAMN07573759 | PRJNA400795 | 0 |
| Columbicola passerinae passerinae 1 | Cosp.Cosqu.11.7.2016.5 | Columbina squammata | 4781 | Brazil: Para | 15-Aug-2013 | 7-Nov-2016 | 0.156 | Covaris M220 | Hyper Library (Kapa Biosystems) | Illumina HiSeq4000 | HiSeq4000 v.1 | 151 | AACATTAGCCGG | 1 | 150 bp | yes | 72675376 | 54.50653 | 14.09 | 52631931 | 39.47395 | 1033 | SRR5998806 | SAMN07573763 | PRJNA400795 | 0 |
| Columbicola passerinae passerinae 1 | Cosp.Cotal.11.2.2016.15 | Columbina talpacoti | GMS1870 | Panama: Torti | 13-Feb-2006 | 2-Nov-2016 | 0.502 | Covaris M220 | Hyper Library (Kapa Biosystems) | Illumina HiSeq4000 | HiSeq4000 v.1 | 151 | GGCGGTTGGGCC | 2 | 150 bp | yes | 46266960 | 34.70022 | 14.42 | 35674625 | 26.75597 | 1037 | SRR5998827 | SAMN07573757 | PRJNA400795 | 0 |
| Columbicola passerinae passerinae 1 | Cosp.Cotal.11.7.2016.2 | Columbina talpacoti | JMD732 | Panama: Felipillo | 19-Feb-2006 | 7-Nov-2016 | 0.601 | Covaris M220 | Hyper Library (Kapa Biosystems) | Illumina HiSeq4000 | HiSeq4000 v.1 | 151 | GTCGTCAACCGG | 1 | 150 bp | yes | 35243764 | 26.43282 | 15.97 | 26727002 | 20.04525 | 1037 | SRR5998826 | SAMN07573760 | PRJNA400795 | 0 |
| Columbicola passerinae passerinae 1 | Cosp.Urcam.11.7.2016.4 | Uropelia campestris | 4789 | Brazil: Para | 15-Aug-2013 | 7-Nov-2016 | 0.104 | Covaris M220 | Hyper Library (Kapa Biosystems) | Illumina HiSeq4000 | HiSeq4000 v.1 | 151 | GACCGTAGCCTT | 1 | 150 bp | yes | 74749204 | 56.06190 | 17.66 | 52886962 | 39.66522 | 1036 | SRR5998830 | SAMN07573762 | PRJNA400795 | 0 |
| Columbicola passerinae passerinae 1 | Cosp.Urcam.11.7.2016.6 | Uropelia campestris | 4888 | Brazil: Para | 20-Aug-2013 | 7-Nov-2016 | 0.335 | Covaris M220 | Hyper Library (Kapa Biosystems) | Illumina HiSeq4000 | HiSeq4000 v.1 | 151 | CGTCAGTTGTCC | 2 | 150 bp | yes | 49864818 | 37.39861 | 17.87 | 39286712 | 29.46503 | 1037 | SRR5998805 | SAMN07573764 | PRJNA400795 | 0 |
| Columbicola passerinae passerinae 1 | Copas1 | Columbina picui |  | Argentina: Salta |  |  | NA |  |  |  |  | NA |  | NA |  |  | NA | NA | NA | NA | NA | 1037 | SRR3161931 | SAMN04103775 | PRJNA296666 | 1 |
| Columbicola passerinae passerinae 2 | Cosp.Clpre.11.2.2016.14 | Claravis pretiosa | REW178 | Peru: Cajamarca | 7-Sep-2002 | 2-Nov-2016 | 0.304 | Covaris M220 | Hyper Library (Kapa Biosystems) | Illumina HiSeq4000 | HiSeq4000 v.1 | 151 | CATGGCCGGTCC | 2 | 150 bp | yes | 41944386 | 31.45829 | 16.97 | 31984684 | 23.98851 | 1036 | SRR5998822 | SAMN07573756 | PRJNA400795 | 0 |
| Columbicola passerinae passerinae 2 | Cosp.Clpre.11.7.2016.11 | Claravis pretiosa | CO-14 | Mexico: Campeche | 19-Mar-1998 | 7-Nov-2016 | 0.514 | Covaris M220 | Hyper Library (Kapa Biosystems) | Illumina HiSeq4000 | HiSeq4000 v.1 | 151 | CTGGCTAGCCTT | 1 | 150 bp | yes | 45739438 | 34.30458 | 21.00 | 33977515 | 25.48314 | 1037 | SRR5998809 | SAMN07573768 | PRJNA400795 | 0 |
| Columbicola passerinae passerinae 2 | Cosp.Clpre.11.7.2016.3 | Claravis pretiosa | REW166 | Peru: Cajamarca: Jaen | 5-Sep-2002 | 7-Nov-2016 | 0.443 | Covaris M220 | Hyper Library (Kapa Biosystems) | Illumina HiSeq4000 | HiSeq4000 v.1 | 151 | TTGCATGTGGCC | 1 | 150 bp | yes | 40306124 | 30.22959 | 20.84 | 30250586 | 22.68794 | 1037 | SRR5998829 | SAMN07573761 | PRJNA400795 | 0 |
| Columbicola passerinae passerinae 2 | Cosp.Clpre.11.7.2016.9 | Claravis pretiosa | 5178 | Brazil: Para | 7-Jun-2013 | 7-Nov-2016 | 0.334 | Covaris M220 | Hyper Library (Kapa Biosystems) | Illumina HiSeq4000 | HiSeq4000 v.1 | 151 | GTCACGTACCTT | 1 | 150 bp | yes | 42960738 | 32.22055 | 44.09 | 27071367 | 20.30353 | 1037 | SRR5998803 | SAMN07573766 | PRJNA400795 | 0 |
| Columbicola passerinae passerinae 2 | Cosp.Cobuc.11.2.2016.13 | Columbina buckleyi | REW187 | Peru: Cajamarca | 7-Sep-2002 | 2-Nov-2016 | 0.638 | Covaris M220 | Hyper Library (Kapa Biosystems) | Illumina HiSeq4000 | HiSeq4000 v.1 | 151 | CCTGTTGACCGG | 2 | 150 bp | yes | 36749634 | 27.56223 | 17.85 | 28306675 | 21.23001 | 1034 | SRR5998821 | SAMN07573755 | PRJNA400795 | 0 |
| Columbicola passerinae passerinae 2 | Cosp.Cobuc.11.2.2016.2 | Columbina buckleyi | RCF998 | Peru | n/a | 2-Nov-2016 | 0.403 | Covaris M220 | Hyper Library (Kapa Biosystems) | Illumina HiSeq4000 | HiSeq4000 v.1 | 151 | GCAGTGTTGGCC | 1 | 150 bp | yes | 32833854 | 24.62539 | 25.53 | 24018454 | 18.01384 | 1037 | SRR5998814 | SAMN07573744 | PRJNA400795 | 0 |
| Columbicola passerinae passerinae 2 | Cosp.Cocru.11.2.2016.11 | Columbina cruziana | REW84 | Peru: Lima: Hatillo | 8-Aug-2002 | 2-Nov-2016 | 0.386 | Covaris M220 | Hyper Library (Kapa Biosystems) | Illumina HiSeq4000 | HiSeq4000 v.1 | 151 | CAGTACGCGGCC | 1 | 150 bp | yes | 42659324 | 31.99449 | 41.41 | 28547754 | 21.41082 | 1037 | SRR5998823 | SAMN07573753 | PRJNA400795 | 0 |
| Columbicola passerinae passerinae 2 | Cosp.Cocru.11.7.2016.16 | Columbina cruziana | ASEC006 | Ecuador: Manabi: Cerro Seco | 29-Jan-2016 | 7-Nov-2016 | 0.353 | Covaris M220 | Hyper Library (Kapa Biosystems) | Illumina HiSeq4000 | HiSeq4000 v.1 | 151 | GAAGGAACCCGG | 2 | 150 bp | yes | 49782620 | 37.33696 | 15.47 | 38438694 | 28.82902 | 1036 | SRR5998831 | SAMN07573773 | PRJNA400795 | 0 |
| Columbicola passerinae passerinae 2 | Cosp.Cotal.11.2.2016.8 | Columbina talpacoti | n/a | Bolivia | 5-May-2002 | 2-Nov-2016 | 0.482 | Covaris M220 | Hyper Library (Kapa Biosystems) | Illumina HiSeq4000 | HiSeq4000 v.1 | 151 | AAATTAACCCTT | 2 | 150 bp | yes | 48092106 | 36.06908 | 15.46 | 36474253 | 27.35569 | 1037 | SRR5998816 | SAMN07573750 | PRJNA400795 | 0 |
| Columbicola passerinae passerinae 2 | Cosp.Cotal.11.7.2016.10 | Columbina talpacoti | 6471 | Brazil: Rondonia | 7-Oct-2014 | 7-Nov-2016 | 0.304 | Covaris M220 | Hyper Library (Kapa Biosystems) | Illumina HiSeq4000 | HiSeq4000 v.1 | 151 | ACTACCAACCGG | 2 | 150 bp | yes | 45408470 | 34.05635 | 13.86 | 34842969 | 26.13223 | 1037 | SRR5998810 | SAMN07573767 | PRJNA400795 | 0 |
| Columbicola passerinae passerinae 2 | Copas2 | Columbina cruziana |  | Peru |  |  | NA |  |  |  |  | NA |  | NA |  |  | NA | NA | NA | NA | NA | 1037 | SRR3161930 | SAMN04103774 | PRJNA296666 | 1 |
| Columbicola drowni drowni | Cosp.Memel.11.2.2016.6 | Metriopelia melanoptera | KGM 443 | Argentina: Jujuy | 10-Oct-2001 | 2-Nov-2016 | 0.501 | Covaris M220 | Hyper Library (Kapa Biosystems) | Illumina HiSeq4000 | HiSeq4000 v.1 | 151 | TACCGGCGCCGG | 2 | 150 bp | yes | 46552564 | 34.91442 | 12.95 | 35613935 | 26.71045 | 1039 | SRR5998818 | SAMN07573748 | PRJNA400795 | 0 |
| Columbicola drowni drowni | Cosp.Memel.11.7.2016.13 | Metriopelia melanoptera | ASEC003 | Ecuador: Pichincha: Secas | 23-Jan-2016 | 7-Nov-2016 | 0.570 | Covaris M220 | Hyper Library (Kapa Biosystems) | Illumina HiSeq4000 | HiSeq4000 v.1 | 151 | TACTACGGCCTT | 2 | 150 bp | yes | 54926416 | 41.19481 | 19.90 | 41232463 | 30.92435 | 1039 | SRR5998807 | SAMN07573770 | PRJNA400795 | 0 |
| Columbicola drowni drowni | Codro | Metriopelia melanoptera |  | Argentina: Jujuy |  |  | NA |  |  |  |  | NA |  | NA |  |  | NA | NA | NA | NA | NA | 1039 | SRR3161922 | SAMN04103766 | PRJNA296666 | 1 |
| Columbicola gymnopeliae gymnopeliae | Cosp.Mecec.11.2.2016.10 | Metriopelia ceciliae | REW192 | Peru: Cajamarca | 13-Sep-2002 | 2-Nov-2016 | 0.494 | Covaris M220 | Hyper Library (Kapa Biosystems) | Illumina HiSeq4000 | HiSeq4000 v.1 | 151 | GCTCGGCCGGCC | 1 | 150 bp | yes | 44362564 | 33.27192 | 16.95 | 33757698 | 25.31827 | 1038 | SRR5998820 | SAMN07573752 | PRJNA400795 | 1 |
| Columbicola gymnopeliae gymnopeliae | Cosp.Mecec.11.2.2016.12 | Metriopelia ceciliae | REW150 | Peru: Caras | 28-Aug-2002 | 2-Nov-2016 | 0.561 | Covaris M220 | Hyper Library (Kapa Biosystems) | Illumina HiSeq4000 | HiSeq4000 v.1 | 151 | CATAGCTTCCGG | 2 | 150 bp | yes | 39654522 | 29.74089 | 12.77 | 30602281 | 22.95171 | 1037 | SRR5998824 | SAMN07573754 | PRJNA400795 | 0 |
| Columbicola gymnopeliae gymnopeliae | Cogym | Metriopelia ceciliae |  | Peru: Cajamarca |  |  | NA |  |  |  |  | NA |  | NA |  |  | NA | NA | NA | NA | NA | 1037 | SRR3161923 | SAMN04103767 | PRJNA296666 | 0 |
| Columbicola altamimiae altamimiae | Cosp.Meaym.11.2.2016.5 | Metriopelia aymara | KGM 431 | Argentina: Salta | 6-Oct-2001 | 2-Nov-2016 | 0.606 | Covaris M220 | Hyper Library (Kapa Biosystems) | Illumina HiSeq4000 | HiSeq4000 v.1 | 151 | TTACTTCTCCGG | 1 | 150 bp | yes | 35598232 | 26.69867 | 13.33 | 26911929 | 20.18395 | 1031 | SRR5998817 | SAMN07573747 | PRJNA400795 | 1 |
| Columbicola altamimiae altamimiae | Cosp.Meaym.11.2.2016.7 | Metriopelia aymara | KGM 510 | Bolivia: Oruro: Lago Poopo | 2-Nov-2001 | 2-Nov-2016 | 0.168 | Covaris M220 | Hyper Library (Kapa Biosystems) | Illumina HiSeq4000 | HiSeq4000 v.1 | 151 | ATGATCGGCCTT | 2 | 150 bp | yes | 46193236 | 34.64493 | 15.82 | 34544913 | 25.90868 | 1031 | SRR5998815 | SAMN07573749 | PRJNA400795 | 0 |
| Columbicola altamimiae altamimiae | Coalt | Metriopelia aymara |  | Argentina: Salta |  |  | NA |  |  |  |  | NA |  | NA |  |  | NA | NA | NA | NA | NA | 1031 | SRR3161921 | SAMN04103765 | PRJNA296666 | 0 |

### Figs. S1-S12

Click the links below (in blue) to visualize each plot. Individuals used for the phylonet analyses are asterisked.

#### Columbicola

##### Fig. S1. sppIDer results of Columbicola altamimiae samples

\*coalt (Species coverage)

```
<embed src="OUT_speciesDepth_coalt.pdf">
</embed>
```

coalt (Coverage across the genomes)

```
<embed src="OUT_sppIDerDepthPlot-covBins_coalt.pdf">
</embed>
```

coalt (Mapping quality)

```
<embed src="OUT_plotMQ_coalt.pdf">
</embed>
```

Cosp\_Meaym\_11\_2\_2016\_5 (Species coverage)

```
<embed src="OUT_speciesDepth_Cosp_Meaym_11_2_2016_5.pdf">
</embed>
```

Cosp\_Meaym\_11\_2\_2016\_5 (Coverage across the genomes)

```
<embed src="OUT_sppIDerDepthPlot-covBins_Cosp_Meaym_11_2_2016_5.pdf">
</embed>
```

Cosp\_Meaym\_11\_2\_2016\_5 (Mapping quality)

```
<embed src="OUT_plotMQ_Cosp_Meaym_11_2_2016_5.pdf">
</embed>
```

Cosp\_Meaym\_11\_2\_2016\_7 (Species coverage)

```
<embed src="OUT_speciesDepth_Cosp_Meaym_11_2_2016_7.pdf">
</embed>
```

Cosp\_Meaym\_11\_2\_2016\_7 (Coverage across the genomes)

```
<embed src="OUT_sppIDerDepthPlot-covBins_Cosp_Meaym_11_2_2016_7.pdf">
</embed>
```

Cosp\_Meaym\_11\_2\_2016\_7 (Mapping quality)

```
<embed src="OUT_plotMQ_Cosp_Meaym_11_2_2016_7.pdf">
</embed>
```

##### Fig. S2. sppIDer results of Columbicola drowni\_drowni samples

\*Codro (Species coverage)

```
<embed src="OUT_speciesDepth_Codro.pdf">
</embed>
```

Codro (Coverage across the genomes)

```
<embed src="OUT_sppIDerDepthPlot-covBins_Codro.pdf">
</embed>
```

Codro (Mapping quality)

```
<embed src="OUT_plotMQ_Codro.pdf">
</embed>
```

Cosp\_Memel\_11\_7\_2016\_13 (Species coverage)

```
<embed src="OUT_speciesDepth_Cosp_Memel_11_7_2016_13.pdf">
</embed>
```

Cosp\_Memel\_11\_7\_2016\_13 (Coverage across the genomes)

```
<embed src="OUT_sppIDerDepthPlot-covBins_Cosp_Memel_11_7_2016_13">
</embed>
```

Cosp\_Memel\_11\_7\_2016\_13 (Mapping quality)

```
<embed src="OUT_plotMQ_Cosp_Memel_11_7_2016_13.pdf">
</embed>
```

Cosp\_Memel\_11\_2\_2016\_6 (Species coverage)

```
<embed src="OUT_speciesDepth_Cosp_Memel_11_2_2016_6.pdf">
</embed>
```

Cosp\_Memel\_11\_2\_2016\_6 (Coverage across the genomes)

```
<embed src="OUT_sppIDerDepthPlot-covBins_Cosp_Memel_11_2_2016_6.pdf">
</embed>
```

Cosp\_Memel\_11\_2\_2016\_6 (Mapping quality)

```
<embed src="OUT_plotMQ_Cosp_Memel_11_2_2016_6.pdf">
</embed>
```

##### Fig. S3. sppIDer results of Columbicola gymnopeliae samples

cogym (Species coverage)

```
<embed src="OUT_speciesDepth_cogym.pdf">
</embed>
```

cogym (Coverage across the genomes)

```
<embed src="OUT_sppIDerDepthPlot-covBins_cogym.pdf">
</embed>
```

cogym (Mapping quality)

```
<embed src="OUT_plotMQ_cogym.pdf">
</embed>
```

\*Cosp\_Mecec\_11\_2\_2016\_10 (Species coverage)

```
<embed src="OUT_speciesDepth_Cosp_Mecec_11_2_2016_10.pdf">
</embed>
```

Cosp\_Mecec\_11\_2\_2016\_10 (Coverage across the genomes)

```
<embed src="OUT_sppIDerDepthPlot-covBins_Cosp_Mecec_11_2_2016_10.pdf">
</embed>
```

Cosp\_Mecec\_11\_2\_2016\_10 (Mapping quality)

```
<embed src="OUT_plotMQ_Cosp_Mecec_11_2_2016_10.pdf">
</embed>
```

Cosp\_Mecec\_11\_2\_2016\_12 (Species coverage)

```
<embed src="OUT_speciesDepth_Cosp_Mecec_11_2_2016_12.pdf">
</embed>
```

Cosp\_Mecec\_11\_2\_2016\_12 (Coverage across the genomes)

```
<embed src="OUT_sppIDerDepthPlot-covBins_Cosp_Mecec_11_2_2016_12_6">
</embed>
```

Cosp\_Mecec\_11\_2\_2016\_12 (Mapping quality)

```
<embed src="OUT_plotMQ_Cosp_Mecec_11_2_2016_12.pdf">
</embed>
```

##### Fig. S4. sppIDer results of Columbicola passerinae 1 samples

\*copas1 (Species coverage)

```
<embed src="OUT_speciesDepth_copas1.pdf">
</embed>
```

copas1 (Coverage across the genomes)

```
<embed src="OUT_sppIDerDepthPlot-covBins_copas1.pdf">
</embed>
```

copas1 (Mapping quality)

```
<embed src="OUT_plotMQ_copas1.pdf">
</embed>
```

Cosp\_Coinc\_11\_2\_2016\_9 (Species coverage)

```
<embed src="OUT_speciesDepth_Cosp_Coinc_11_2_2016_9.pdf">
</embed>
```

Cosp\_Coinc\_11\_2\_2016\_9 (Coverage across the genomes)

```
<embed src="OUT_sppIDerDepthPlot-covBins_Cosp_Coinc_11_2_2016_9.pdf">
</embed>
```

Cosp\_Coinc\_11\_2\_2016\_9 (Mapping quality)

```
<embed src="OUT_plotMQ_Cosp_Coinc_11_2_2016_9.pdf">
</embed>
```

Cosp\_Comin\_11\_2\_2016\_16 (Species coverage)

```
<embed src="OUT_speciesDepth_Cosp_Comin_11_2_2016_16.pdf">
</embed>
```

Cosp\_Comin\_11\_2\_2016\_16 (Coverage across the genomes)

```
<embed src="OUT_sppIDerDepthPlot-covBins_Cosp_Comin_11_2_2016_16.pdf">
</embed>
```

Cosp\_Comin\_11\_2\_2016\_16 (Mapping quality)

```
<embed src="OUT_plotMQ_Cosp_Comin_11_2_2016_16.pdf">
</embed>
```

Cosp\_Comin\_11\_2\_2016\_4 (Species coverage)

```
<embed src="OUT_speciesDepth_Cosp_Comin_11_2_2016_4.pdf">
</embed>
```

Cosp\_Comin\_11\_2\_2016\_4 (Coverage across the genomes)

```
<embed src="OUT_sppIDerDepthPlot-covBins_Cosp_Comin_11_2_2016_4.pdf">
</embed>
```

Cosp\_Comin\_11\_2\_2016\_4 (Mapping quality)

```
<embed src="OUT_plotMQ_Cosp_Comin_11_2_2016_4.pdf">
</embed>
```

Cosp\_Copas\_11\_7\_2016\_8 (Species coverage)

```
<embed src="OUT_speciesDepth_Cosp_Copas_11_7_2016_8.pdf">
</embed>
```

Cosp\_Copas\_11\_7\_2016\_8 (Coverage across the genomes)

```
<embed src="OUT_sppIDerDepthPlot-covBins_Cosp_Copas_11_7_2016_8.pdf">
</embed>
```

Cosp\_Copas\_11\_7\_2016\_8 (Mapping quality)

```
<embed src="OUT_plotMQ_Cosp_Copas_11_7_2016_8.pdf">
</embed>
```

Cosp\_Copas\_11\_7\_2016\_12 (Species coverage)

```
<embed src="OUT_speciesDepth_Cosp_Copas_11_7_2016_12.pdf">
</embed>
```

Cosp\_Copas\_11\_7\_2016\_12 (Coverage across the genomes)

```
<embed src="OUT_sppIDerDepthPlot-covBins_Cosp_Copas_11_7_2016_12.pdf">
</embed>
```

Cosp\_Copas\_11\_7\_2016\_12 (Mapping quality)

```
<embed src="OUT_plotMQ_Cosp_Copas_11_7_2016_12.pdf">
</embed>
```

Cosp\_Copas\_11\_7\_2016\_14 (Species coverage)

```
<embed src="OUT_speciesDepth_Cosp_Copas_11_7_2016_14.pdf">
</embed>
```

Cosp\_Copas\_11\_7\_2016\_14 (Coverage across the genomes)

```
<embed src="OUT_sppIDerDepthPlot-covBins_Cosp_Copas_11_7_2016_14.pdf">
</embed>
```

Cosp\_Copas\_11\_7\_2016\_14 (Mapping quality)

```
<embed src="OUT_plotMQ_Cosp_Copas_11_7_2016_14.pdf">
</embed>
```

Cosp\_Copas\_11\_7\_2016\_15 (Species coverage)

```
<embed src="OUT_speciesDepth_Cosp_Copas_11_7_2016_15.pdf">
</embed>
```

Cosp\_Copas\_11\_7\_2016\_15 (Coverage across the genomes)

```
<embed src="OUT_sppIDerDepthPlot-covBins_Cosp_Copas_11_7_2016_15.pdf">
</embed>
```

Cosp\_Copas\_11\_7\_2016\_15 (Mapping quality)

```
<embed src="OUT_plotMQ_Cosp_Copas_11_7_2016_15.pdf">
</embed>
```

Cosp\_Copic\_11\_2\_2016\_1 (Species coverage)

```
<embed src="OUT_speciesDepth_Cosp_Copic_11_2_2016_1.pdf">
</embed>
```

Cosp\_Copic\_11\_2\_2016\_1 (Coverage across the genomes)

```
<embed src="OUT_sppIDerDepthPlot-covBins_Cosp_Copic_11_2_2016_1.pdf">
</embed>
```

Cosp\_Copic\_11\_2\_2016\_1 (Mapping quality)

```
<embed src="OUT_plotMQ_Cosp_Copic_11_2_2016_1.pdf">
</embed>
```

Cosp\_Copic\_11\_2\_2016\_3 (Species coverage)

```
<embed src="OUT_speciesDepth_Cosp_Copic_11_2_2016_3.pdf">
</embed>
```

Cosp\_Copic\_11\_2\_2016\_3 (Coverage across the genomes)

```
<embed src="OUT_sppIDerDepthPlot-covBins_Cosp_Copic_11_2_2016_3.pdf">
</embed>
```

Cosp\_Copic\_11\_2\_2016\_3 (Mapping quality)

```
<embed src="OUT_plotMQ_Cosp_Copic_11_2_2016_3.pdf">
</embed>
```

Cosp\_Cosqu\_11\_7\_2016\_1 (Species coverage)

```
<embed src="OUT_speciesDepth_Cosp_Cosqu_11_7_2016_1.pdf">
</embed>
```

Cosp\_Cosqu\_11\_7\_2016\_1 (Coverage across the genomes)

```
<embed src="OUT_sppIDerDepthPlot-covBins_Cosp_Cosqu_11_7_2016_1.pdf">
</embed>
```

Cosp\_Cosqu\_11\_7\_2016\_1 (Mapping quality)

```
<embed src="OUT_plotMQ_Cosp_Cosqu_11_7_2016_1.pdf">
</embed>
```

Cosp\_Cosqu\_11\_7\_2016\_5 (Species coverage)

```
<embed src="OUT_speciesDepth_Cosp_Cosqu_11_7_2016_5.pdf">
</embed>
```

Cosp\_Cosqu\_11\_7\_2016\_5 (Coverage across the genomes)

```
<embed src="OUT_sppIDerDepthPlot-covBins_Cosp_Cosqu_11_7_2016_5.pdf">
</embed>
```

Cosp\_Cosqu\_11\_7\_2016\_5 (Mapping quality)

```
<embed src="OUT_plotMQ_Cosp_Cosqu_11_7_2016_5.pdf">
</embed>
```

Cosp\_Cotal\_11\_2\_2016\_15 (Species coverage)

```
<embed src="OUT_speciesDepth_Cosp_Cotal_11_2_2016_15.pdf">
</embed>
```

Cosp\_Cotal\_11\_2\_2016\_15 (Coverage across the genomes)

```
<embed src="OUT_sppIDerDepthPlot-covBins_Cosp_Cotal_11_2_2016_15.pdf">
</embed>
```

Cosp\_Cotal\_11\_2\_2016\_15 (Mapping quality)

```
<embed src="OUT_plotMQ_Cosp_Cotal_11_2_2016_15.pdf">
</embed>
```

Cosp\_Cotal\_11\_7\_2016\_2 (Species coverage)

```
<embed src="OUT_speciesDepth_Cosp_Cotal_11_7_2016_2.pdf">
</embed>
```

Cosp\_Cotal\_11\_7\_2016\_2 (Coverage across the genomes)

```
<embed src="OUT_sppIDerDepthPlot-covBins_Cosp_Cotal_11_7_2016_2.pdf">
</embed>
```

Cosp\_Cotal\_11\_7\_2016\_2 (Mapping quality)

```
<embed src="OUT_plotMQ_Cosp_Cotal_11_7_2016_2.pdf">
</embed>
```

Cosp\_Urcam\_11\_7\_2016\_6 (Species coverage)

```
<embed src="OUT_speciesDepth_Cosp_Urcam_11_7_2016_6.pdf">
</embed>
```

Cosp\_Urcam\_11\_7\_2016\_6 (Coverage across the genomes)

```
<embed src="OUT_sppIDerDepthPlot-covBins_Cosp_Urcam_11_7_2016_6.pdf">
</embed>
```

Cosp\_Urcam\_11\_7\_2016\_6 (Mapping quality)

```
<embed src="OUT_plotMQ_Cosp_Urcam_11_7_2016_6.pdf">
</embed>
```

Cosp\_Urcam\_11\_7\_2016\_4 (Species coverage)

```
<embed src="OUT_speciesDepth_Cosp_Urcam_11_7_2016_4.pdf">
</embed>
```

Cosp\_Urcam\_11\_7\_2016\_4 (Coverage across the genomes)

```
<embed src="OUT_sppIDerDepthPlot-covBins_Cosp_Urcam_11_7_2016_4.pdf">
</embed>
```

Cosp\_Urcam\_11\_7\_2016\_4 (Mapping quality)

```
<embed src="OUT_plotMQ_Cosp_Urcam_11_7_2016_4.pdf">
</embed>
```

##### Fig. S5. sppIDer results of Columbicola passerinae 2 samples

\*Copas2 (Species coverage)

```
<embed src="OUT_speciesDepth_Copas2.pdf">
</embed>
```

Copas2 (Coverage across the genomes)

```
<embed src="OUT_sppIDerDepthPlot-covBins_Copas2.pdf">
</embed>
```

Copas2 (Mapping quality)

```
<embed src="OUT_plotMQ_Copas2.pdf">
</embed>
```

Cosp\_Clpre\_11\_2\_2016\_14 (Species coverage)

```
<embed src="OUT_speciesDepth_Cosp_Clpre_11_2_2016_14.pdf">
</embed>
```

Cosp\_Clpre\_11\_2\_2016\_14 (Coverage across the genomes)

```
<embed src="OUT_sppIDerDepthPlot-covBins_Cosp_Clpre_11_2_2016_14.pdf">
</embed>
```

Cosp\_Clpre\_11\_2\_2016\_14 (Mapping quality)

```
<embed src="OUT_plotMQ_Cosp_Clpre_11_2_2016_14.pdf">
</embed>
```

Cosp\_Clpre\_11\_7\_2016\_3 (Species coverage)

```
<embed src="OUT_speciesDepth_Cosp_Clpre_11_7_2016_3.pdf">
</embed>
```

Cosp\_Clpre\_11\_7\_2016\_3 (Coverage across the genomes)

```
<embed src="OUT_sppIDerDepthPlot-covBins_Cosp_Clpre_11_7_2016_3.pdf">
</embed>
```

Cosp\_Clpre\_11\_7\_2016\_3 (Mapping quality)

```
<embed src="OUT_plotMQ_Cosp_Clpre_11_7_2016_3.pdf">
</embed>
```

Cosp\_Clpre\_11\_7\_2016\_9 (Species coverage)

```
<embed src="OUT_speciesDepth_Cosp_Clpre_11_7_2016_9.pdf">
</embed>
```

Cosp\_Clpre\_11\_7\_2016\_9 (Coverage across the genomes)

```
<embed src="OUT_sppIDerDepthPlot-covBins_Cosp_Clpre_11_7_2016_9.pdf">
</embed>
```

Cosp\_Clpre\_11\_7\_2016\_9 (Mapping quality)

```
<embed src="OUT_plotMQ_Cosp_Clpre_11_7_2016_9.pdf">
</embed>
```

Cosp\_Clpre\_11\_7\_2016\_11 (Species coverage)

```
<embed src="OUT_speciesDepth_Cosp_Clpre_11_7_2016_11.pdf">
</embed>
```

Cosp\_Clpre\_11\_7\_2016\_11 (Coverage across the genomes)

```
<embed src="OUT_sppIDerDepthPlot-covBins_Cosp_Clpre_11_7_2016_11.pdf">
</embed>
```

Cosp\_Clpre\_11\_7\_2016\_11 (Mapping quality)

```
<embed src="OUT_plotMQ_Cosp_Clpre_11_7_2016_11.pdf">
</embed>
```

Cosp\_Cobuc\_11\_2\_2016\_13 (Species coverage)

```
<embed src="OUT_speciesDepth_Cosp_Cobuc_11_2_2016_13.pdf">
</embed>
```

Cosp\_Cobuc\_11\_2\_2016\_13 (Coverage across the genomes)

```
<embed src="OUT_sppIDerDepthPlot-covBins_Cosp_Cobuc_11_2_2016_13.pdf">
</embed>
```

Cosp\_Cobuc\_11\_2\_2016\_13 (Mapping quality)

```
<embed src="OUT_plotMQ_Cosp_Cobuc_11_2_2016_13.pdf">
</embed>
```

Cosp\_Cobuc\_11\_2\_2016\_2 (Species coverage)

```
<embed src="OUT_speciesDepth_Cosp_Cobuc_11_2_2016_2.pdf">
</embed>
```

Cosp\_Cobuc\_11\_2\_2016\_2 (Coverage across the genomes)

```
<embed src="OUT_sppIDerDepthPlot-covBins_Cosp_Cobuc_11_2_2016_2.pdf">
</embed>
```

Cosp\_Cobuc\_11\_2\_2016\_2 (Mapping quality)

```
<embed src="OUT_plotMQ_Cosp_Cobuc_11_2_2016_2.pdf">
</embed>
```

Cosp\_Cocru\_11\_2\_2016\_11 (Species coverage)

```
<embed src="OUT_speciesDepth_Cosp_Cocru_11_2_2016_11.pdf">
</embed>
```

Cosp\_Cocru\_11\_2\_2016\_11 (Coverage across the genomes)

```
<embed src="OUT_sppIDerDepthPlot-covBins_Cosp_Cocru_11_2_2016_11.pdf">
</embed>
```

Cosp\_Cocru\_11\_2\_2016\_11 (Mapping quality)

```
<embed src="OUT_plotMQ_Cosp_Cocru_11_2_2016_11.pdf">
</embed>
```

Cosp\_Cocru\_11\_7\_2016\_16 (Species coverage)

```
<embed src="OUT_speciesDepth_Cosp_Cocru_11_7_2016_16.pdf">
</embed>
```

Cosp\_Cocru\_11\_7\_2016\_16 (Coverage across the genomes)

```
<embed src="OUT_sppIDerDepthPlot-covBins_Cosp_Cocru_11_7_2016_16.pdf">
</embed>
```

Cosp\_Cocru\_11\_7\_2016\_16 (Mapping quality)

```
<embed src="OUT_plotMQ_Cosp_Cocru_11_7_2016_16.pdf">
</embed>
```

Cosp\_Cotal\_11\_7\_2016\_10 (Species coverage)

```
<embed src="OUT_speciesDepth_Cosp_Cotal_11_7_2016_10.pdf">
</embed>
```

Cosp\_Cotal\_11\_7\_2016\_10 (Coverage across the genomes)

```
<embed src="OUT_sppIDerDepthPlot-covBins_Cosp_Cotal_11_7_2016_10.pdf">
</embed>
```

Cosp\_Cotal\_11\_7\_2016\_10 (Mapping quality)

```
<embed src="OUT_plotMQ_Cosp_Cotal_11_7_2016_10.pdf">
</embed>
```

Cosp\_Cotal\_11\_2\_2016\_8 (Species coverage)

```
<embed src="OUT_speciesDepth_Cosp_Cotal_11_2_2016_8.pdf">
</embed>
```

Cosp\_Cotal\_11\_2\_2016\_8 (Coverage across the genomes)

```
<embed src="OUT_sppIDerDepthPlot-covBins_Cosp_Cotal_11_2_2016_8.pdf">
</embed>
```

Cosp\_Cotal\_11\_2\_2016\_8 (Mapping quality)

```
<embed src="OUT_plotMQ_Cosp_Cotal_11_2_2016_8.pdf">
</embed>
```

#### Physconelloides

##### Fig. S6. sppIDer results of Physconelloides eurysema 1 samples

\*Pheur\_Copas\_3\_14\_2016\_8 (Species coverage)

```
<embed src="OUT_speciesDepth_Pheur_Copas_3_14_2016_8.pdf">
</embed>
```

Pheur\_Copas\_3\_14\_2016\_8 (Coverage across the genomes)

```
<embed src="OUT_sppIDerDepthPlot-covBins_Pheur_Copas_3_14_2016_8.pdf">
</embed>
```

Pheur\_Copas\_3\_14\_2016\_8 (Mapping quality)

```
<embed src="OUT_plotMQ_Pheur_Copas_3_14_2016_8.pdf">
</embed>
```

Pheur\_Copas\_3\_16\_2016\_6 (Species coverage)

```
<embed src="OUT_speciesDepth_Pheur_Copas_3_16_2016_6.pdf">
</embed>
```

Pheur\_Copas\_3\_16\_2016\_6 (Coverage across the genomes)

```
<embed src="OUT_sppIDerDepthPlot-covBins_Pheur_Copas_3_16_2016_6.pdf">
</embed>
```

Pheur\_Copas\_3\_16\_2016\_6 (Mapping quality)

```
<embed src="OUT_plotMQ_Pheur_Copas_3_16_2016_6.pdf">
</embed>
```

Pheur\_Comin\_6\_15\_2015\_10 (Species coverage)

```
<embed src="OUT_speciesDepth_Pheur.Comin.6.15.2015.10.pdf">
</embed>
```

Pheur\_Comin\_6\_15\_2015\_10 (Coverage across the genomes)

```
<embed src="OUT_sppIDerDepthPlot-covBins_Pheur.Comin.6.15.2015.10.pdf">
</embed>
```

Pheur\_Comin\_6\_15\_2015\_10 (Mapping quality)

```
<embed src="OUT_plotMQ_Pheur.Comin.6.15.2015.10.pdf">
</embed>
```

##### Fig. S7. sppIDer results of Physconelloides eurysema 2 samples

\*Pheur\_Cosqu\_3\_14\_2016\_1 (Species coverage)

```
<embed src="OUT_speciesDepth_Pheur_Cosqu_3_14_2016_1.pdf">
</embed>
```

Pheur\_Cosqu\_3\_14\_2016\_1 (Coverage across the genomes)

```
<embed src="OUT_sppIDerDepthPlot-covBins_Pheur_Cosqu_3_14_2016_1.pdf">
</embed>
```

Pheur\_Cosqu\_3\_14\_2016\_1 (Mapping quality)

```
<embed src="OUT_plotMQ_Pheur_Cosqu_3_14_2016_1.10.pdf">
</embed>
```

Pheur\_Cosqu\_3\_14\_2016\_12 (Species coverage)

```
<embed src="OUT_speciesDepth_Pheur_Cosqu_3_14_2016_12.pdf">
</embed>
```

Pheur\_Cosqu\_3\_14\_2016\_12 (Coverage across the genomes)

```
<embed src="OUT_sppIDerDepthPlot-covBins_Pheur_Cosqu_3_14_2016_12.pdf">
</embed>
```

Pheur\_Cosqu\_3\_14\_2016\_12 (Mapping quality)

```
<embed src="OUT_plotMQ_Pheur_Cosqu_3_14_2016_12.pdf">
</embed>
```

##### Fig. S8. sppIDer results of Physconelloides eurysema 3 samples

Pheur2\_Clpre\_5\_26\_2015\_6 (Species coverage)

```
<embed src="OUT_speciesDepth_Pheur2_Clpre_5_26_2015_6.pdf">
</embed>
```

Pheur2\_Clpre\_5\_26\_2015\_6 (Coverage across the genomes)

```
<embed src="OUT_sppIDerDepthPlot-covBins_Pheur2_Clpre_5_26_2015_6.pdf">
</embed>
```

Pheur2\_Clpre\_5\_26\_2015\_6 (Mapping quality)

```
<embed src="OUT_plotMQ_Pheur2_Clpre_5_26_2015_6.pdf">
</embed>
```

Pheur\_Clpre\_6\_15\_2015\_8 (Species coverage)

```
<embed src="OUT_speciesDepth_Pheur_Clpre_6_15_2015_8.pdf">
</embed>
```

Pheur\_Clpre\_6\_15\_2015\_8 (Coverage across the genomes)

```
<embed src="OUT_sppIDerDepthPlot-covBins_Pheur_Clpre_6_15_2015_8.pdf">
</embed>
```

Pheur\_Clpre\_6\_15\_2015\_8 (Mapping quality)

```
<embed src="OUT_plotMQ_Pheur_Clpre_6_15_2015_8.pdf">
</embed>
```

Pheur\_Cotal\_3\_14\_2016\_2 (Species coverage)

```
<embed src="OUT_speciesDepth_Pheur_Cotal_3_14_2016_2.pdf">
</embed>
```

Pheur\_Cotal\_3\_14\_2016\_2 (Coverage across the genomes)

```
<embed src="OUT_sppIDerDepthPlot-covBins_Pheur_Cotal_3_14_2016_2.pdf">
</embed>
```

Pheur\_Cotal\_3\_14\_2016\_2 (Mapping quality)

```
<embed src="OUT_plotMQ_Pheur_Cotal_3_14_2016_2.pdf">
</embed>
```

Pheur\_Copas\_3\_14\_2016\_4 (Species coverage)

```
<embed src="OUT_speciesDepth_Pheur_Copas_3_14_2016_4.pdf">
</embed>
```

Pheur\_Copas\_3\_14\_2016\_4 (Coverage across the genomes)

```
<embed src="OUT_sppIDerDepthPlot-covBins_Pheur_Copas_3_14_2016_4.pdf">
</embed>
```

Pheur\_Copas\_3\_14\_2016\_4 (Mapping quality)

```
<embed src="OUT_plotMQ_Pheur_Copas_3_14_2016_4.pdf">
</embed>
```

Pheur2\_Clpre\_3\_14\_2016\_5 (Species coverage)

```
<embed src="OUT_speciesDepth_Pheur2_Clpre_3_14_2016_5.pdf">
</embed>
```

Pheur2\_Clpre\_3\_14\_2016\_5 (Coverage across the genomes)

```
<embed src="OUT_sppIDerDepthPlot-covBins_Pheur2_Clpre_3_14_2016_5.pdf">
</embed>
```

Pheur2\_Clpre\_3\_14\_2016\_5 (Mapping quality)

```
<embed src="OUT_plotMQ_Pheur2_Clpre_3_14_2016_5.pdf">
</embed>
```

Pheur\_Cotal\_3\_14\_2016\_6 (Species coverage)

```
<embed src="OUT_speciesDepth_Pheur_Cotal_3_14_2016_6.pdf">
</embed>
```

Pheur\_Cotal\_3\_14\_2016\_6 (Coverage across the genomes)

```
<embed src="OUT_sppIDerDepthPlot-covBins_Pheur_Cotal_3_14_2016_6.pdf">
</embed>
```

Pheur\_Cotal\_3\_14\_2016\_6 (Mapping quality)

```
<embed src="OUT_plotMQ_Pheur_Cotal_3_14_2016_6.pdf">
</embed>
```

Pheur\_Copas\_3\_14\_2016\_7 (Species coverage)

```
<embed src="OUT_speciesDepth_Pheur_Copas_3_14_2016_7.pdf">
</embed>
```

Pheur\_Copas\_3\_14\_2016\_7 (Coverage across the genomes)

```
<embed src="OUT_sppIDerDepthPlot-covBins_Pheur_Copas_3_14_2016_7.pdf">
</embed>
```

Pheur\_Copas\_3\_14\_2016\_7 (Mapping quality)

```
<embed src="OUT_plotMQ_Pheur_Copas_3_14_2016_7.pdf">
</embed>
```

Pheur1\_Comin\_3\_14\_2016\_9 (Species coverage)

```
<embed src="OUT_speciesDepth_Pheur1_Comin_3_14_2016_9.pdf">
</embed>
```

Pheur1\_Comin\_3\_14\_2016\_9 (Coverage across the genomes)

```
<embed src="OUT_sppIDerDepthPlot-covBins_Pheur1_Comin_3_14_2016_9.pdf">
</embed>
```

Pheur1\_Comin\_3\_14\_2016\_9 (Mapping quality)

```
<embed src="OUT_plotMQ_Pheur1_Comin_3_14_2016_9.pdf">
</embed>
```

Pheur\_Copas\_3\_14\_2016\_10 (Species coverage)

```
<embed src="OUT_speciesDepth_Pheur_Copas_3_14_2016_10.pdf">
</embed>
```

Pheur\_Copas\_3\_14\_2016\_10 (Coverage across the genomes)

```
<embed src="OUT_sppIDerDepthPlot-covBins_Pheur_Copas_3_14_2016_10.pdf">
</embed>
```

Pheur\_Copas\_3\_14\_2016\_10 (Mapping quality)

```
<embed src="OUT_plotMQ_Pheur_Copas_3_14_2016_10.pdf">
</embed>
```

\*Pheur2\_Clpre\_3\_14\_2016\_11 (Species coverage)

```
<embed src="OUT_speciesDepth_Pheur2_Clpre_3_14_2016_11.pdf">
</embed>
```

Pheur2\_Clpre\_3\_14\_2016\_11 (Coverage across the genomes)

```
<embed src="OUT_sppIDerDepthPlot-covBins_Pheur2_Clpre_3_14_2016_11.pdf">
</embed>
```

Pheur2\_Clpre\_3\_14\_2016\_11 (Mapping quality)

```
<embed src="OUT_plotMQ_Pheur2_Clpre_3_14_2016_11.pdf">
</embed>
```

Pheur\_Copas\_3\_16\_2016\_1 (Species coverage)

```
<embed src="OUT_speciesDepth_Pheur_Copas_3_16_2016_1.pdf">
</embed>
```

Pheur\_Copas\_3\_16\_2016\_1 (Coverage across the genomes)

```
<embed src="OUT_sppIDerDepthPlot-covBins_Pheur_Copas_3_16_2016_1.pdf">
</embed>
```

Pheur\_Copas\_3\_16\_2016\_1 (Mapping quality)

```
<embed src="OUT_plotMQ_Pheur_Copas_3_16_2016_1.pdf">
</embed>
```

Pheur\_Cotal\_3\_16\_2016\_3 (Species coverage)

```
<embed src="OUT_speciesDepth_Pheur_Cotal_3_16_2016_3.pdf">
</embed>
```

Pheur\_Cotal\_3\_16\_2016\_3 (Coverage across the genomes)

```
<embed src="OUT_sppIDerDepthPlot-covBins_Pheur_Cotal_3_16_2016_3.pdf">
</embed>
```

Pheur\_Cotal\_3\_16\_2016\_3 (Mapping quality)

```
<embed src="OUT_plotMQ_Pheur_Cotal_3_16_2016_3.pdf">
</embed>
```

Pheur2\_Cobuc\_3\_16\_2016\_4 (Species coverage)

```
<embed src="OUT_speciesDepth_Pheur2_Cobuc_3_16_2016_4.pdf">
</embed>
```

Pheur2\_Cobuc\_3\_16\_2016\_4 (Coverage across the genomes)

```
<embed src="OUT_sppIDerDepthPlot-covBins_Pheur2_Cobuc_3_16_2016_4.pdf">
</embed>
```

Pheur2\_Cobuc\_3\_16\_2016\_4 (Mapping quality)

```
<embed src="OUT_plotMQ_Pheur2_Cobuc_3_16_2016_4.pdf">
</embed>
```

Pheur2\_Clpre\_3\_16\_2016\_5 (Species coverage)

```
<embed src="OUT_speciesDepth_Pheur2_Clpre_3_16_2016_5.pdf">
</embed>
```

Pheur2\_Clpre\_3\_16\_2016\_5 (Coverage across the genomes)

```
<embed src="OUT_sppIDerDepthPlot-covBins_Pheur2_Clpre_3_16_2016_5.pdf">
</embed>
```

Pheur2\_Clpre\_3\_16\_2016\_5 (Mapping quality)

```
<embed src="OUT_plotMQ_Pheur2_Clpre_3_16_2016_5.pdf">
</embed>
```

Pheur\_Copas\_3\_16\_2016\_7 (Species coverage)

```
<embed src="OUT_speciesDepth_Pheur_Copas_3_16_2016_7.pdf">
</embed>
```

Pheur\_Copas\_3\_16\_2016\_7 (Coverage across the genomes)

```
<embed src="OUT_sppIDerDepthPlot-covBins_Pheur_Copas_3_16_2016_7.pdf">
</embed>
```

Pheur\_Copas\_3\_16\_2016\_7 (Mapping quality)

```
<embed src="OUT_plotMQ_Pheur_Copas_3_16_2016_7.pdf">
</embed>
```

Pheur2\_Cocru\_3\_16\_2016\_9 (Species coverage)

```
<embed src="OUT_speciesDepth_Pheur2_Cocru_3_16_2016_9.pdf">
</embed>
```

Pheur2\_Cocru\_3\_16\_2016\_9 (Coverage across the genomes)

```
<embed src="OUT_sppIDerDepthPlot-covBins_Pheur2_Cocru_3_16_2016_9.pdf">
</embed>
```

Pheur2\_Cocru\_3\_16\_2016\_9 (Mapping quality)

```
<embed src="OUT_plotMQ_Pheur2_Cocru_3_16_2016_9.pdf">
</embed>
```

Pheur\_Cotal\_3\_16\_2016\_12 (Species coverage)

```
<embed src="OUT_speciesDepth_Pheur_Cotal_3_16_2016_12.pdf">
</embed>
```

Pheur\_Cotal\_3\_16\_2016\_12 (Coverage across the genomes)

```
<embed src="OUT_sppIDerDepthPlot-covBins_Pheur_Cotal_3_16_2016_12.pdf">
</embed>
```

Pheur\_Cotal\_3\_16\_2016\_12 (Mapping quality)

```
<embed src="OUT_plotMQ_Pheur_Cotal_3_16_2016_12.pdf">
</embed>
```

Pheur\_Cobuc\_6\_15\_2015\_1 (Species coverage)

```
<embed src="OUT_speciesDepth_Pheur_Cobuc_6_15_2015_1.pdf">
</embed>
```

Pheur\_Cobuc\_6\_15\_2015\_1 (Coverage across the genomes)

```
<embed src="OUT_sppIDerDepthPlot-covBins_Pheur_Cobuc_6_15_2015_1.pdf">
</embed>
```

Pheur\_Cobuc\_6\_15\_2015\_1 (Mapping quality)

```
<embed src="OUT_plotMQ_Pheur_Cobuc_6_15_2015_1.pdf">
</embed>
```

Pheur\_Cocru\_6\_15\_2015\_2 (Species coverage)

```
<embed src="OUT_speciesDepth_Pheur_Cocru_6_15_2015_2.pdf">
</embed>
```

Pheur\_Cocru\_6\_15\_2015\_2 (Coverage across the genomes)

```
<embed src="OUT_sppIDerDepthPlot-covBins_Pheur_Cocru_6_15_2015_2.pdf">
</embed>
```

Pheur\_Cocru\_6\_15\_2015\_2 (Mapping quality)

```
<embed src="OUT_plotMQ_Pheur_Cocru_6_15_2015_2.pdf">
</embed>
```

Pheur\_Cotal\_6\_15\_2015\_4 (Species coverage)

```
<embed src="OUT_speciesDepth_Pheur_Cotal_6_15_2015_4.pdf">
</embed>
```

Pheur\_Cotal\_6\_15\_2015\_4 (Coverage across the genomes)

```
<embed src="OUT_sppIDerDepthPlot-covBins_Pheur_Cotal_6_15_2015_4.pdf">
</embed>
```

Pheur\_Cotal\_6\_15\_2015\_4 (Mapping quality)

```
<embed src="OUT_plotMQ_Pheur_Cotal_6_15_2015_4.pdf">
</embed>
```

##### Fig. S9. sppIDer results of Physconelloides eurysema 4 samples

\*Pheur\_Coinc\_6\_15\_2015\_7 (Species coverage)

```
<embed src="OUT_speciesDepth_Pheur_Coinc_6_15_2015_7.pdf">
</embed>
```

Pheur\_Coinc\_6\_15\_2015\_7 (Coverage across the genomes)

```
<embed src="OUT_sppIDerDepthPlot-covBins_Pheur_Coinc_6_15_2015_7.pdf">
</embed>
```

Pheur\_Coinc\_6\_15\_2015\_7 (Mapping quality)

```
<embed src="OUT_plotMQ_Pheur_Coinc_6_15_2015_7.pdf">
</embed>
```

##### Fig. S10. sppIDer results of Physconelloides eurysema 5 samples

\*Pheur\_Urcam\_3\_14\_2016\_3 (Species coverage)

```
<embed src="OUT_speciesDepth_Pheur_Urcam_3_14_2016_3.pdf">
</embed>
```

Pheur\_Urcam\_3\_14\_2016\_3 (Coverage across the genomes)

```
<embed src="OUT_sppIDerDepthPlot-covBins_Pheur_Urcam_3_14_2016_3.pdf">
</embed>
```

Pheur\_Urcam\_3\_14\_2016\_3 (Mapping quality)

```
<embed src="OUT_plotMQ_Pheur_Urcam_3_14_2016_3.pdf">
</embed>
```

##### Fig. S11. sppIDer results of Physconelloides robbinsi samples

\*Pheme\_Memel\_5\_26\_2015\_8 (Species coverage)

```
<embed src="OUT_speciesDepth_Pheme_Memel_5_26_2015_8.pdf">
</embed>
```

Pheme\_Memel\_5\_26\_2015\_8 (Coverage across the genomes)

```
<embed src="OUT_sppIDerDepthPlot-covBins_Pheme_Memel_5_26_2015_8.pdf">
</embed>
```

Pheme\_Memel\_5\_26\_2015\_8 (Mapping quality)

```
<embed src="OUT_plotMQ_Pheme_Memel_5_26_2015_8.pdf">
</embed>
```

Phrob\_Mecec\_3\_16\_2016\_10 (Species coverage)

```
<embed src="OUT_speciesDepth_Phrob_Mecec_3_16_2016_10.pdf">
</embed>
```

Phrob\_Mecec\_3\_16\_2016\_10 (Coverage across the genomes)

```
<embed src="OUT_sppIDerDepthPlot-covBins_Phrob_Mecec_3_16_2016_10.pdf">
</embed>
```

Phrob\_Mecec\_3\_16\_2016\_10 (Mapping quality)

```
<embed src="OUT_plotMQ_Phrob_Mecec_3_16_2016_10.pdf">
</embed>
```

Phrob\_Mecec\_3\_16\_2016\_11 (Species coverage)

```
<embed src="OUT_speciesDepth_Phrob_Mecec_3_16_2016_11.pdf">
</embed>
```

Phrob\_Mecec\_3\_16\_2016\_11 (Coverage across the genomes)

```
<embed src="OUT_sppIDerDepthPlot-covBins_Phrob_Mecec_3_16_2016_11.pdf">
</embed>
```

Phrob\_Mecec\_3\_16\_2016\_11 (Mapping quality)

```
<embed src="OUT_plotMQ_Phrob_Mecec_3_16_2016_11.pdf">
</embed>
```

Phrob\_Mecec\_6\_15\_2015\_6 (Species coverage)

```
<embed src="OUT_speciesDepth_Phrob_Mecec_6_15_2015_6.pdf">
</embed>
```

Phrob\_Mecec\_6\_15\_2015\_6 (Coverage across the genomes)

```
<embed src="OUT_sppIDerDepthPlot-covBins_Phrob_Mecec_6_15_2015_6.pdf">
</embed>
```

Phrob\_Mecec\_6\_15\_2015\_6 (Mapping quality)

```
<embed src="OUT_plotMQ_Phrob_Mecec_6_15_2015_6.pdf">
</embed>
```

##### Fig. S12. sppIDer results of Physconelloides emersoni samples

\*Phrob\_Mecec\_5\_26\_2015\_7 (Species coverage)

```
<embed src="OUT_speciesDepth_Phrob_Mecec_5_26_2015_7.pdf">
</embed>
```

Phrob\_Mecec\_5\_26\_2015\_7 (Coverage across the genomes)

```
<embed src="OUT_sppIDerDepthPlot-covBins_Phrob_Mecec_5_26_2015_7.pdf">
</embed>
```

Phrob\_Mecec\_5\_26\_2015\_7 (Mapping quality)

```
<embed src="OUT_plotMQ_Phrob_Mecec_5_26_2015_7.pdf">
</embed>
```

Pheme\_Memel\_3\_16\_2016\_2 (Species coverage)

```
<embed src="OUT_speciesDepth_Pheme_Memel_3_16_2016_2.pdf">
</embed>
```

Pheme\_Memel\_3\_16\_2016\_2 (Coverage across the genomes)

```
<embed src="OUT_sppIDerDepthPlot-covBins_Pheme_Memel_3_16_2016_2.pdf">
</embed>
```

Pheme\_Memel\_3\_16\_2016\_2 (Mapping quality)

```
<embed src="OUT_plotMQ_Pheme_Memel_3_16_2016_2.pdf">
</embed>
```

Pheme\_Memel\_3\_16\_2016\_8 (Species coverage)

```
<embed src="OUT_speciesDepth_Pheme_Memel_3_16_2016_8.pdf">
</embed>
```

Pheme\_Memel\_3\_16\_2016\_8 (Coverage across the genomes)

```
<embed src="OUT_sppIDerDepthPlot-covBins_Pheme_Memel_3_16_2016_8.pdf">
</embed>
```

Pheme\_Memel\_3\_16\_2016\_8 (Mapping quality)

```
<embed src="OUT_plotMQ_Pheme_Memel_3_16_2016_8.pdf">
</embed>
```

Pheme\_Memel\_6\_15\_2015\_5 (Species coverage)

```
<embed src="OUT_speciesDepth_Pheme_Memel_6_15_2015_5.pdf">
</embed>
```

Pheme\_Memel\_6\_15\_2015\_5 (Coverage across the genomes)

```
<embed src="OUT_sppIDerDepthPlot-covBins_Pheme_Memel_6_15_2015_5.pdf">
</embed>
```

Pheme\_Memel\_6\_15\_2015\_5 (Mapping quality)

```
<embed src="OUT_plotMQ_Pheme_Memel_6_15_2015_5.pdf">
</embed>
```

### Fig. S13. Raw phylogenetic network of Columbicola

Columbicola\_network

```
<embed src="network_columbicola.pdf">
</embed>
```

### Fig. S14. Raw phylogenetic network of Physconelloides

Physconelloides\_network

```
<embed src="network_physconelloides.pdf">
</embed>
```
