## Supplementary figures and images for "Comparing rates of introgression in parasitic feather lice with differing dispersal capabilities" (from another version of this preprint: version 7, 2020-03-31)

### OUT_speciesDepth_Cosp_Cotal_11_7_2016_10.pdf

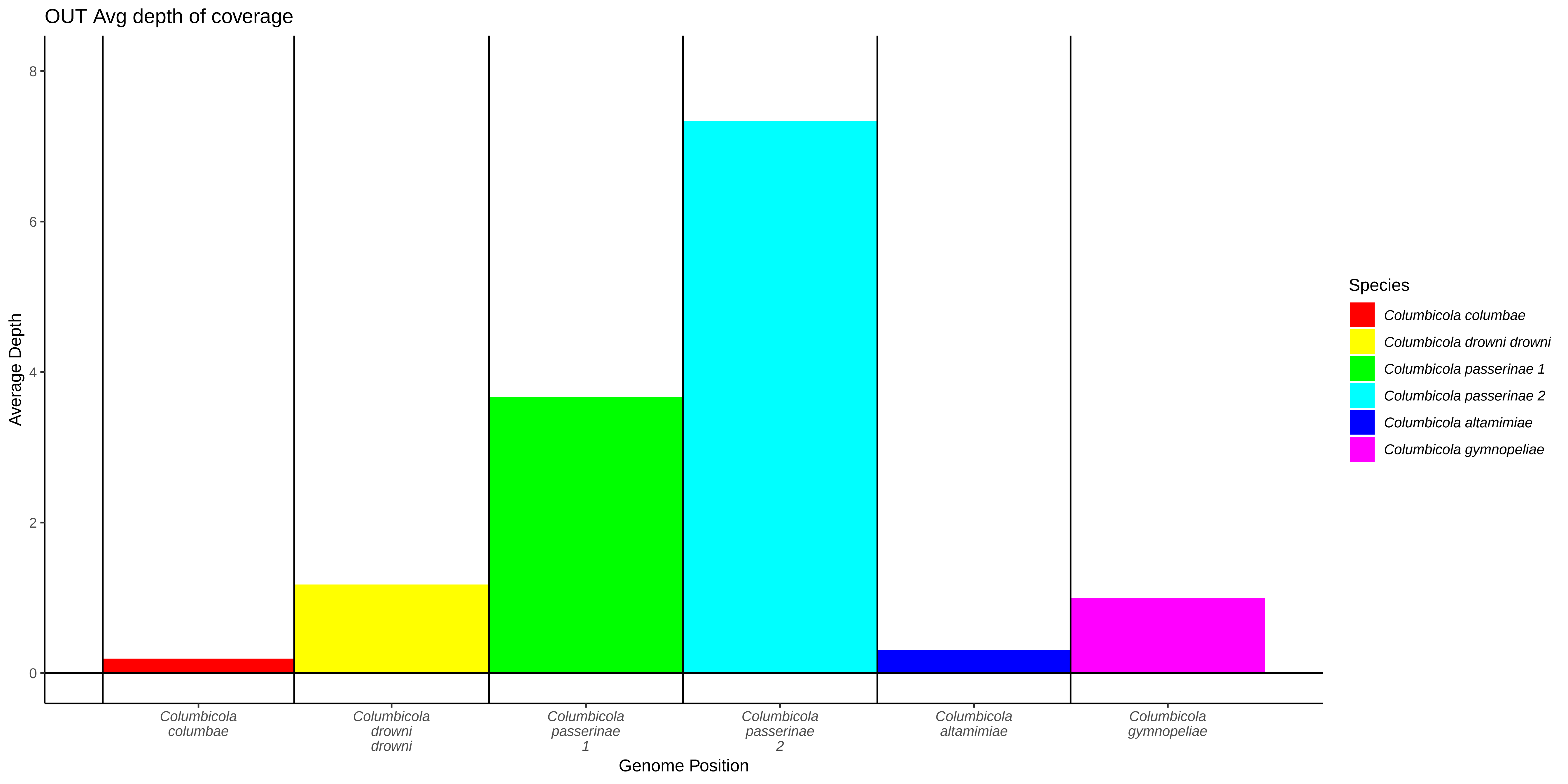

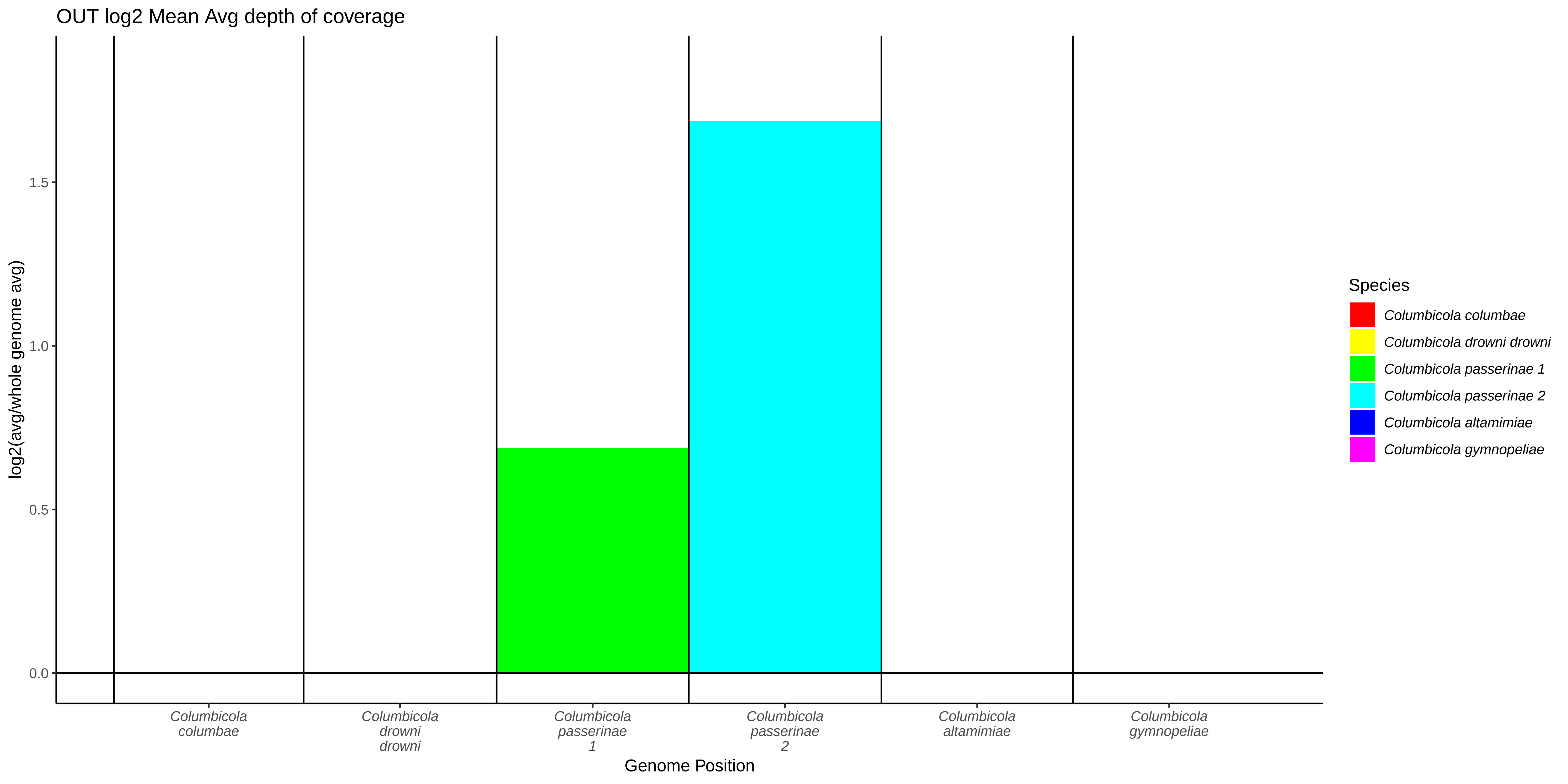

### OUT_speciesDepth_Cosp_Meaym_11_2_2016_5.pdf

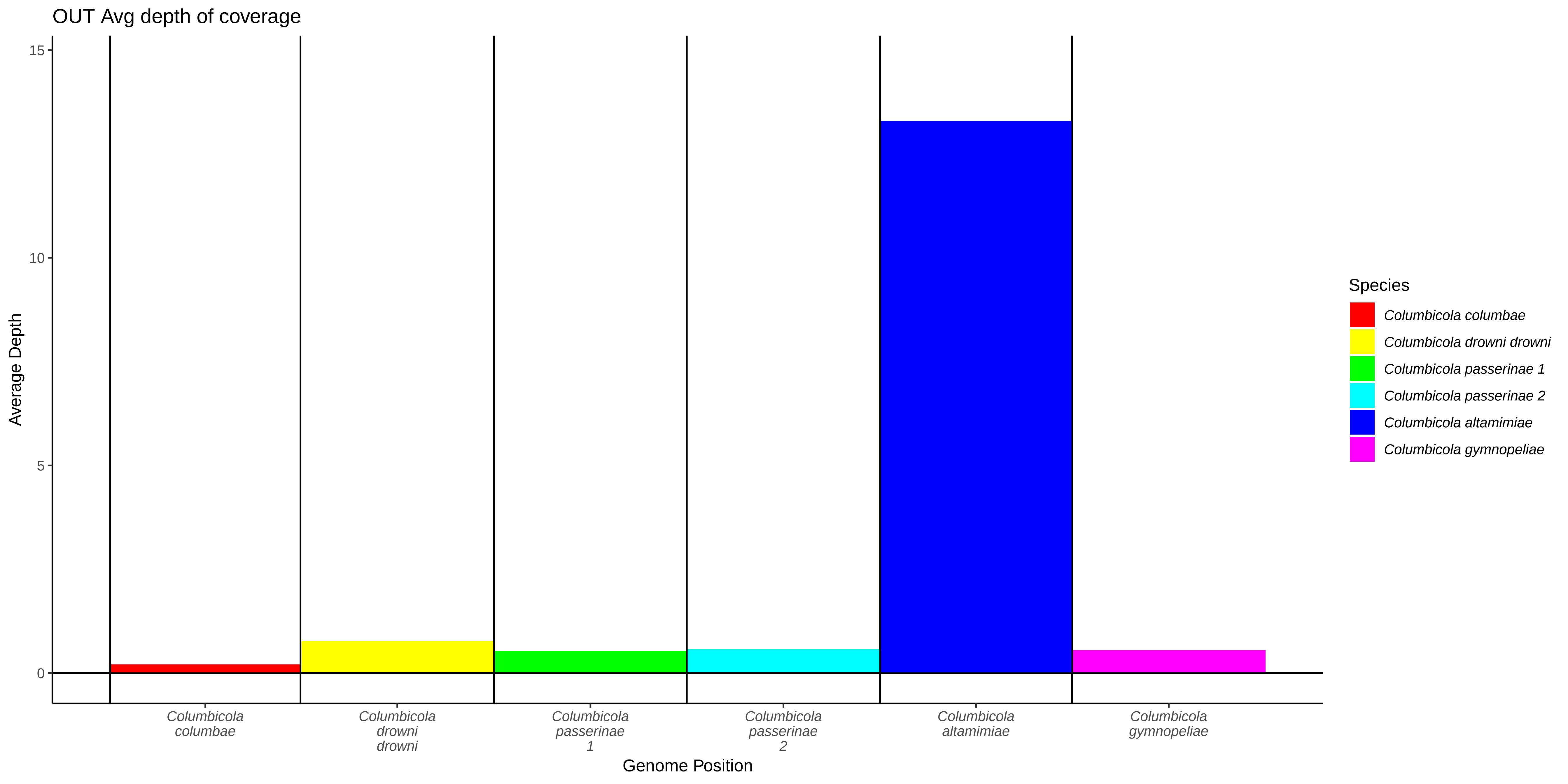

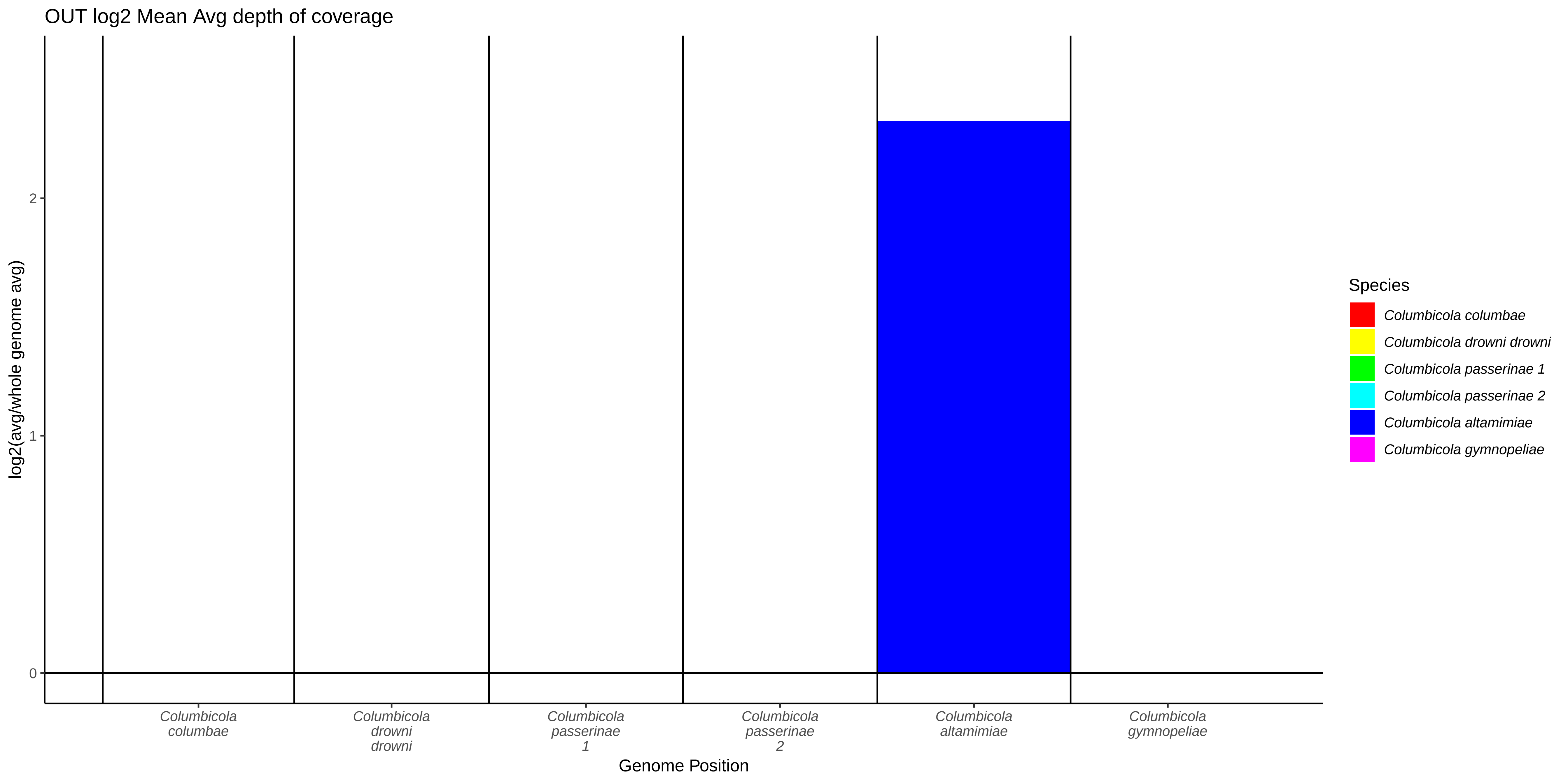

### OUT_speciesDepth_Cosp_Meaym_11_2_2016_7.pdf

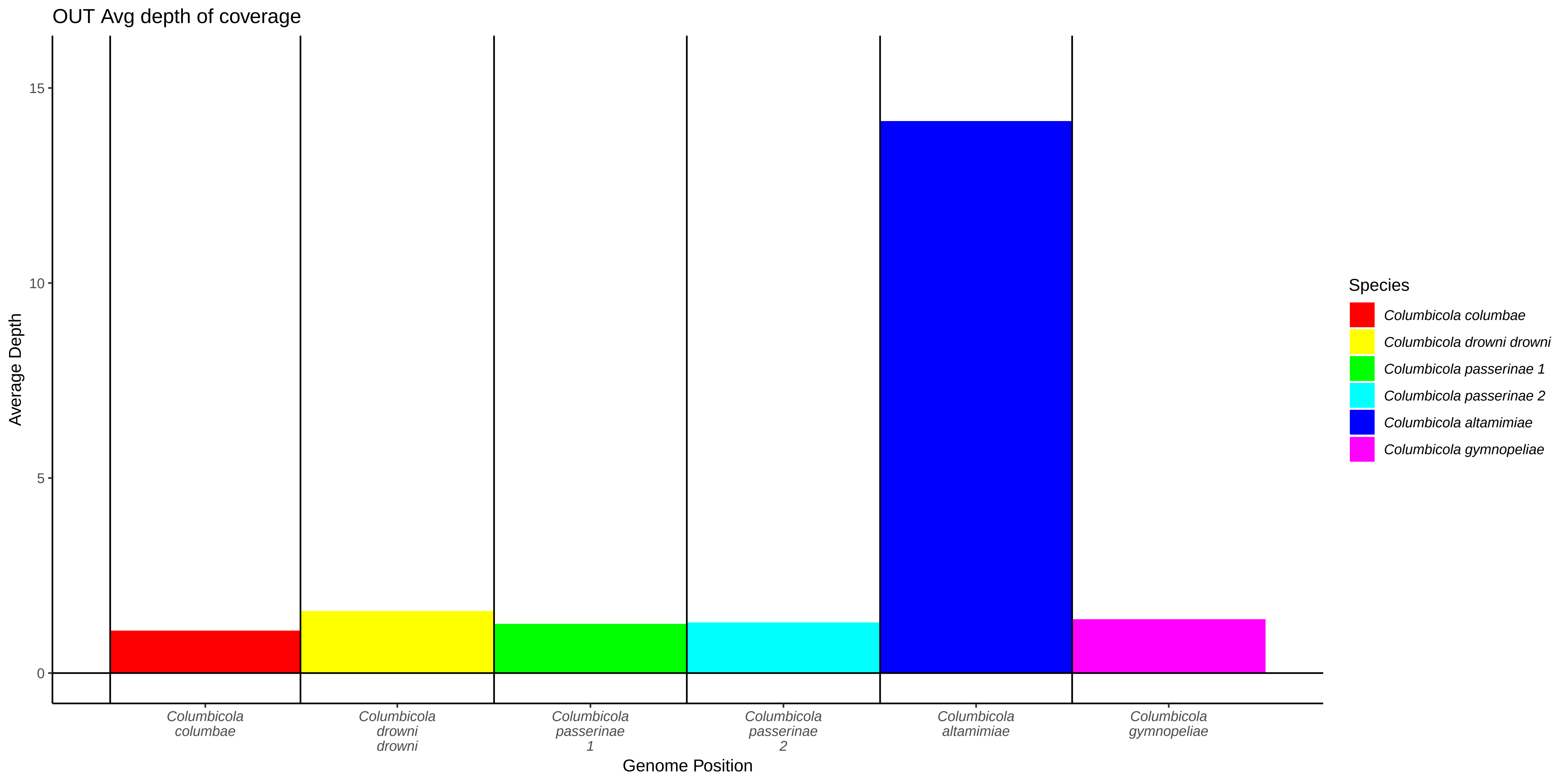

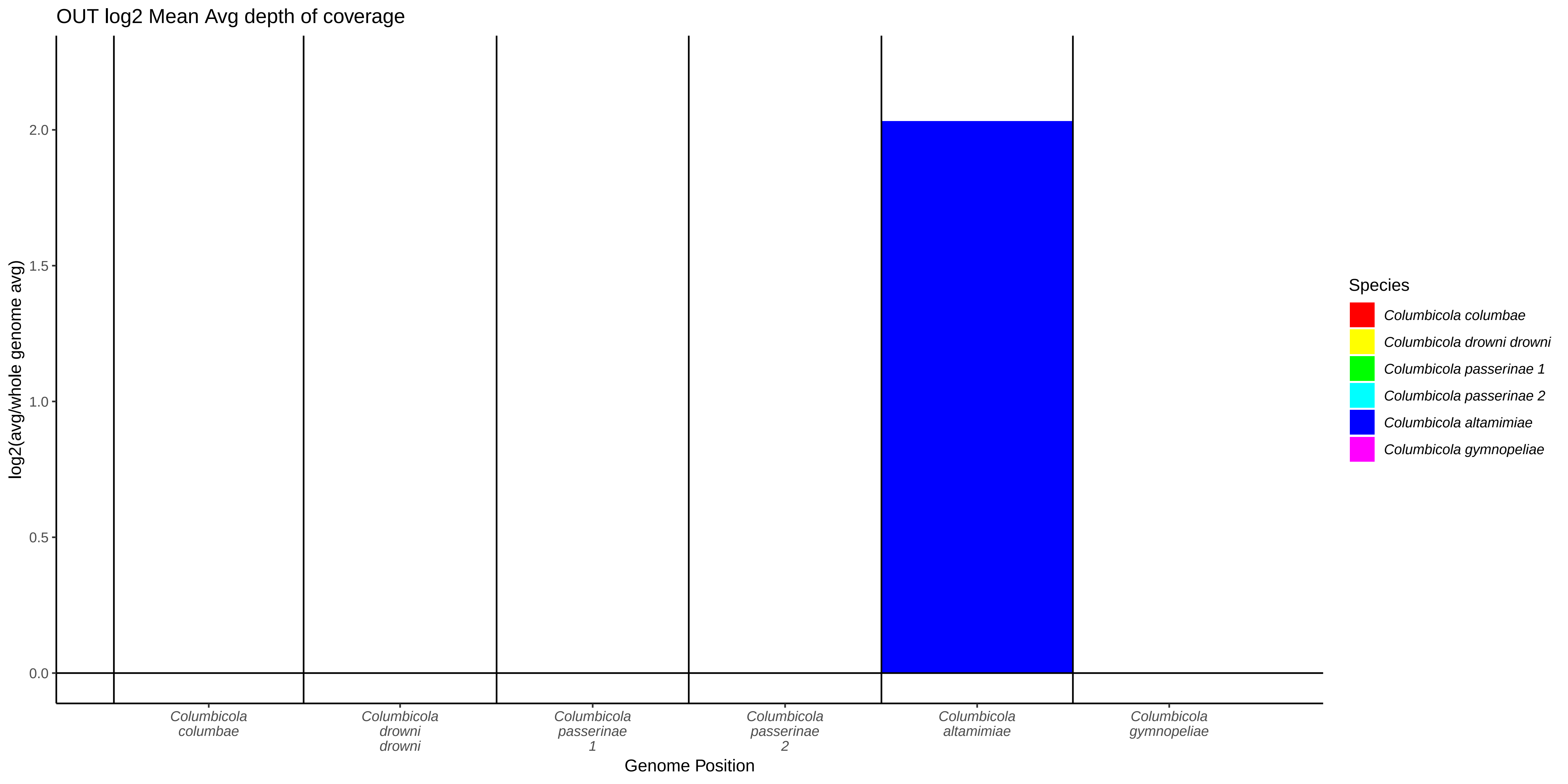

### OUT_speciesDepth_Cosp_Mecec_11_2_2016_10.pdf

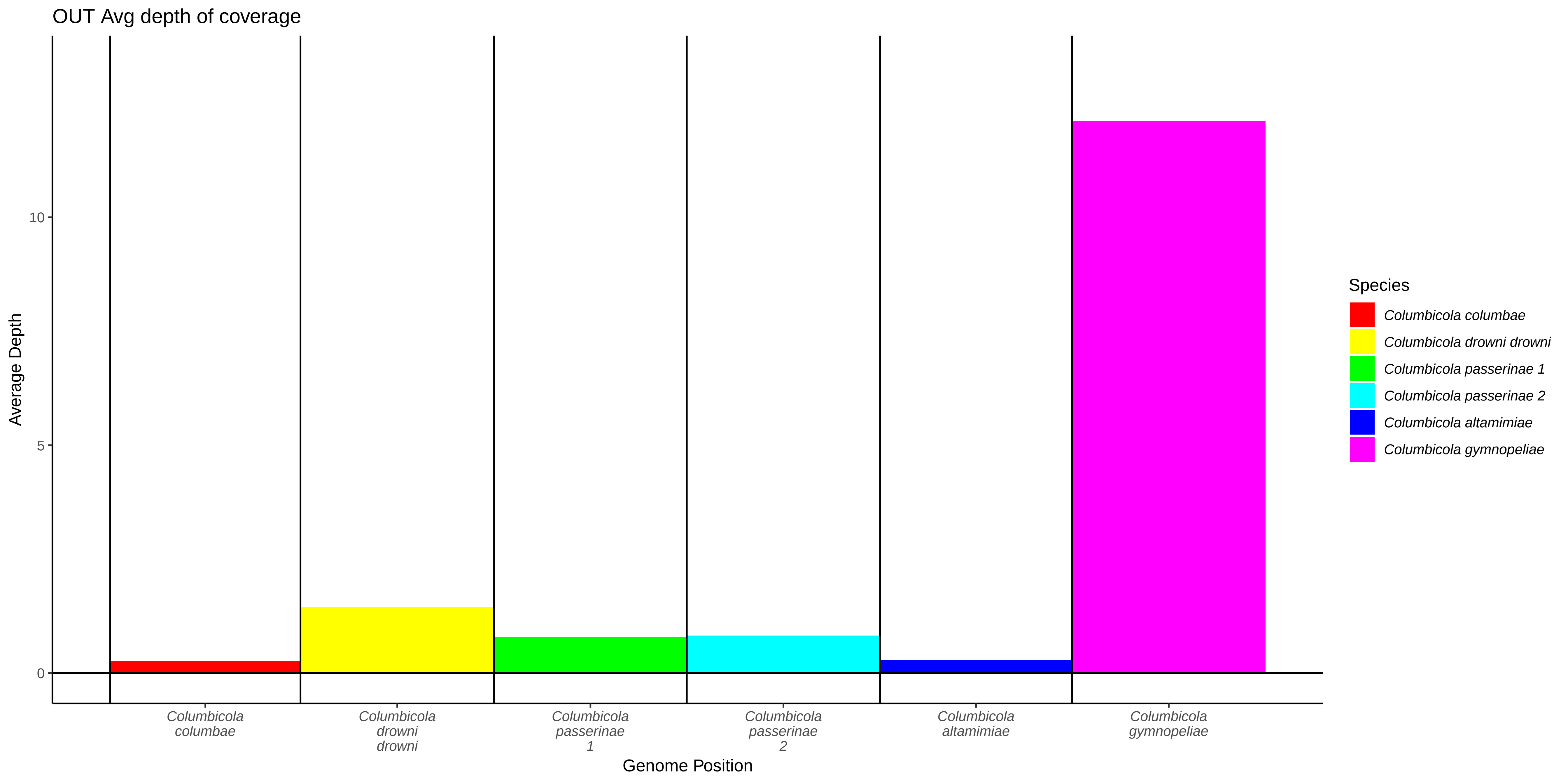

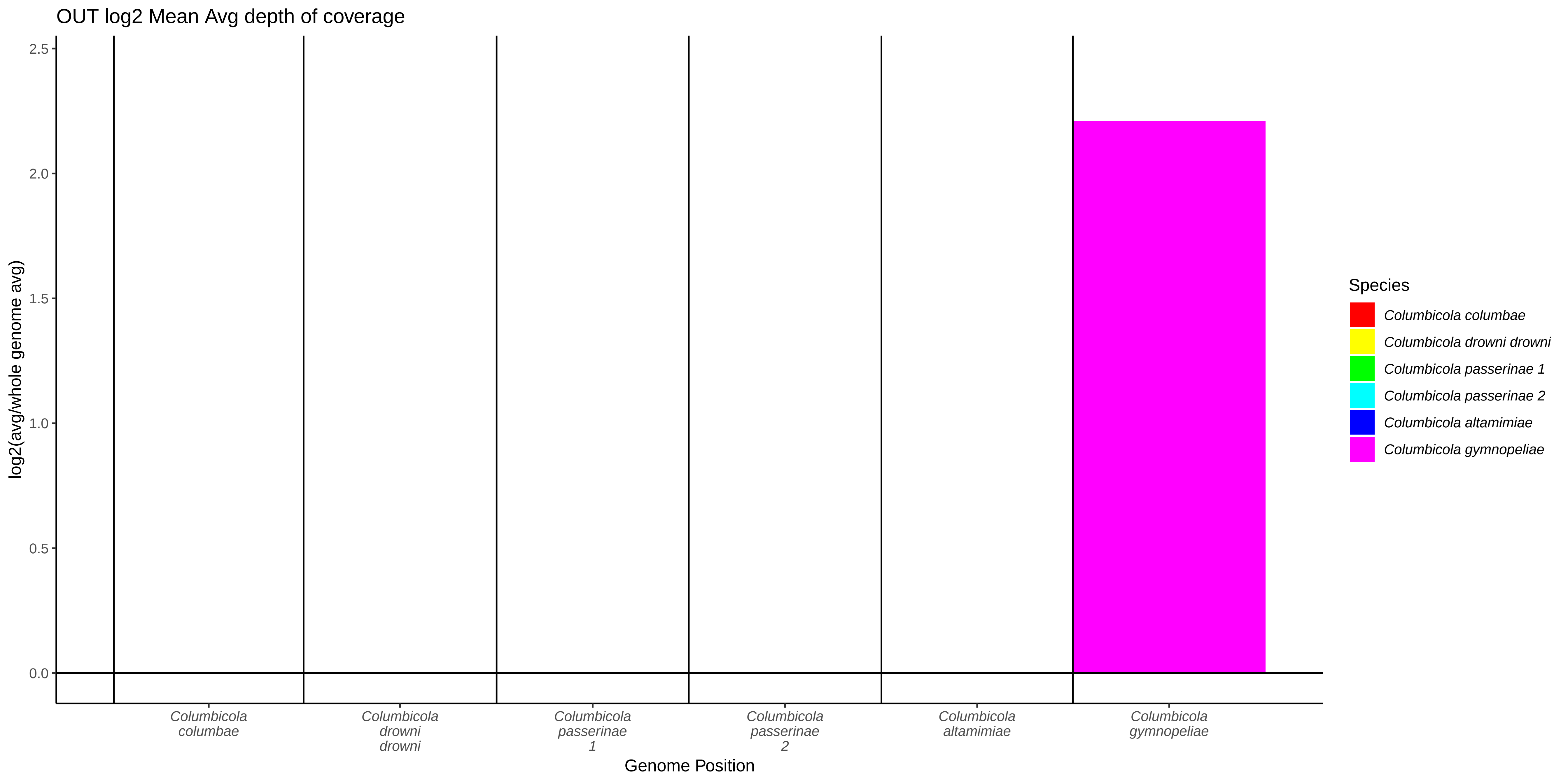

### OUT_speciesDepth_Cosp_Mecec_11_2_2016_12.pdf

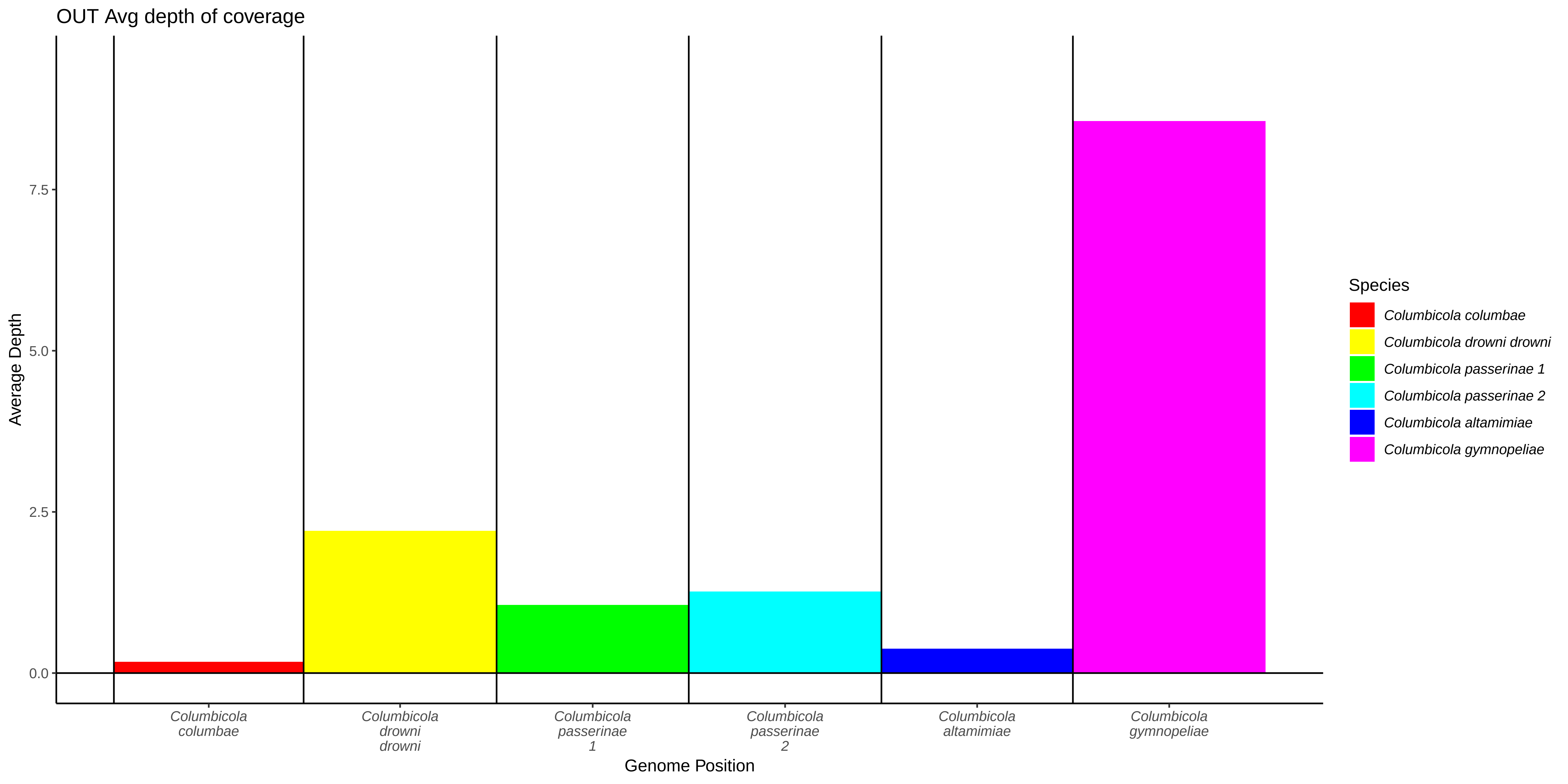

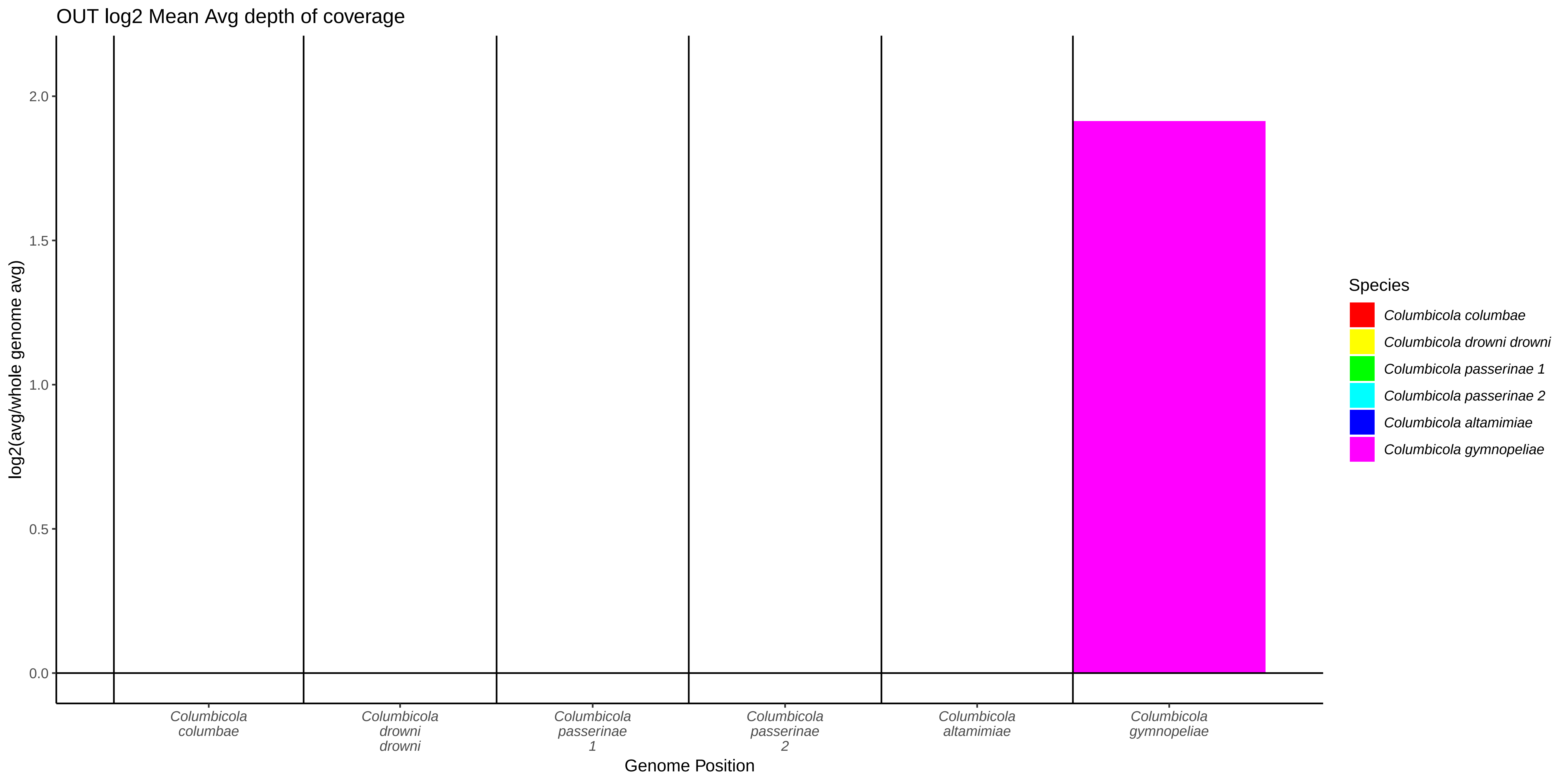

### OUT_speciesDepth_Cosp_Memel_11_2_2016_6.pdf

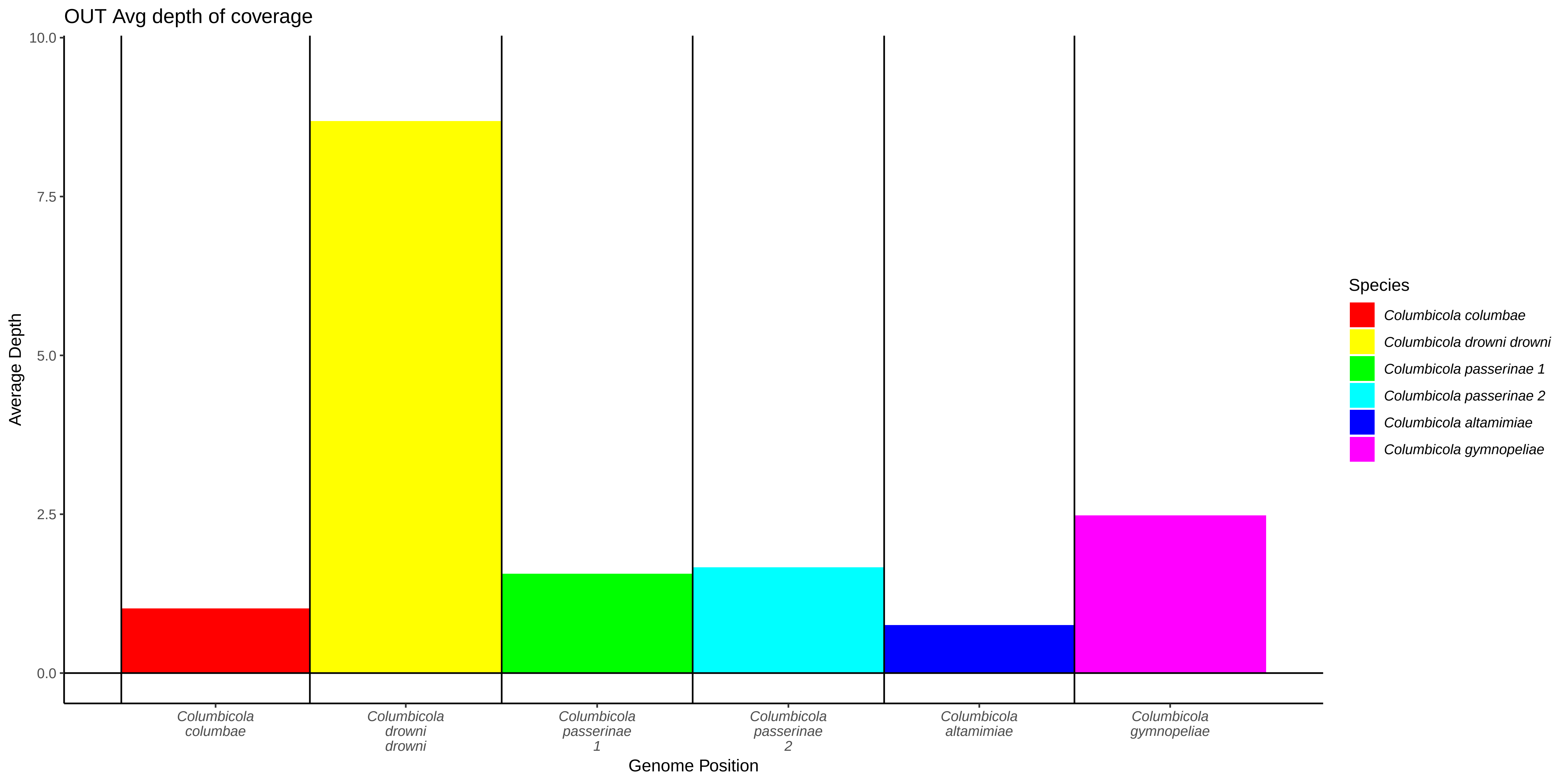

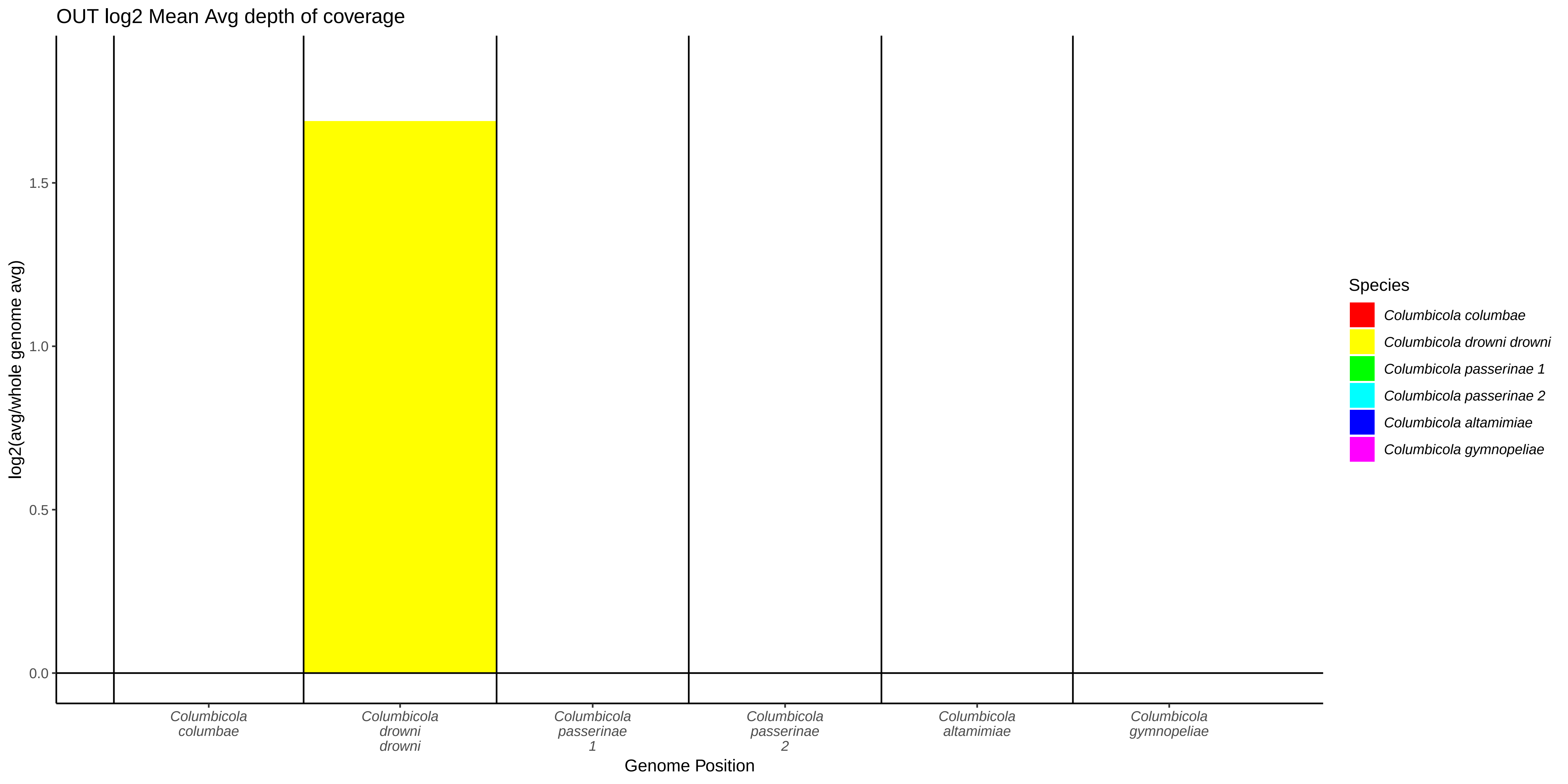

### OUT_speciesDepth_Cosp_Memel_11_7_2016_13.pdf

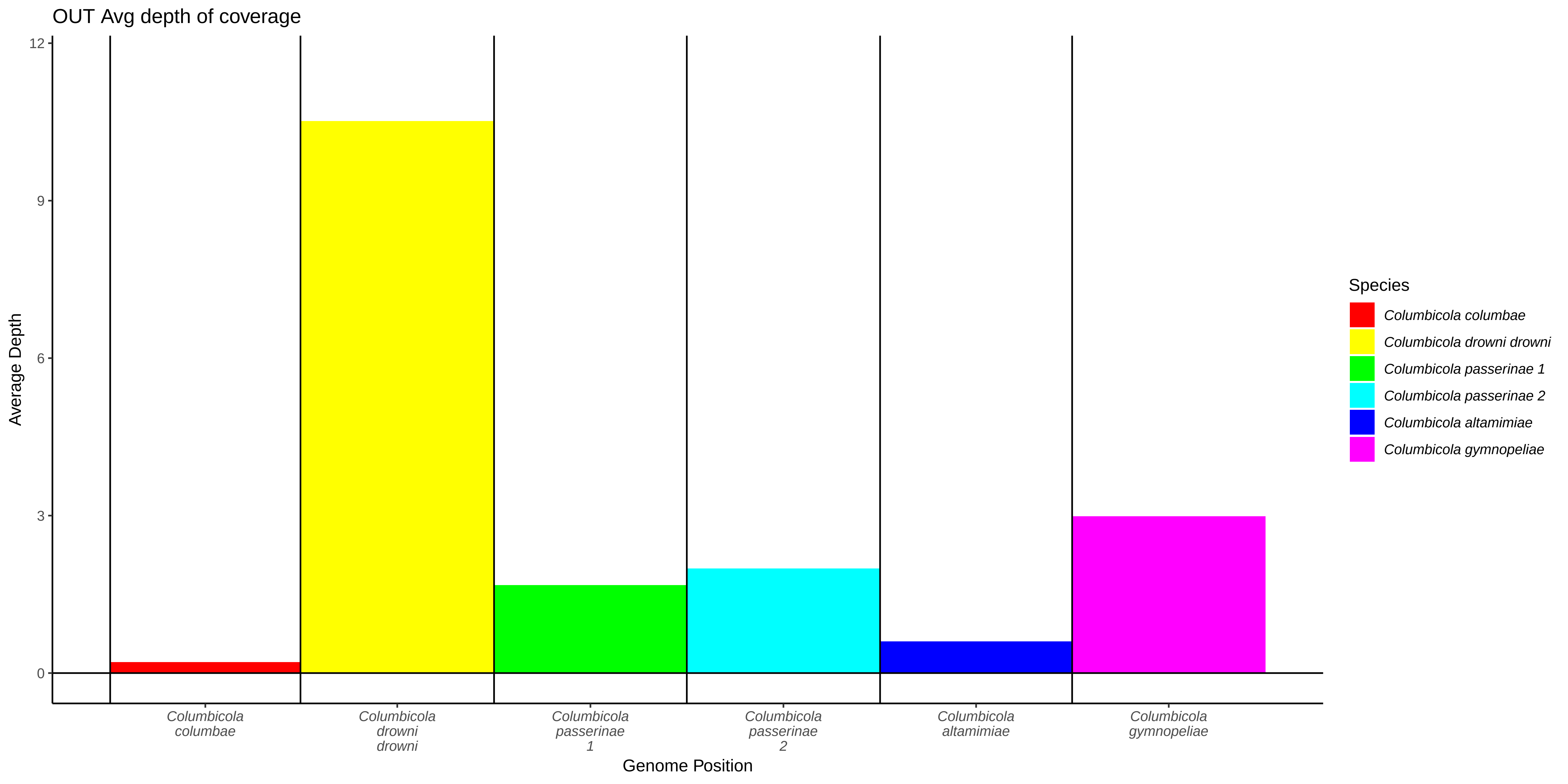

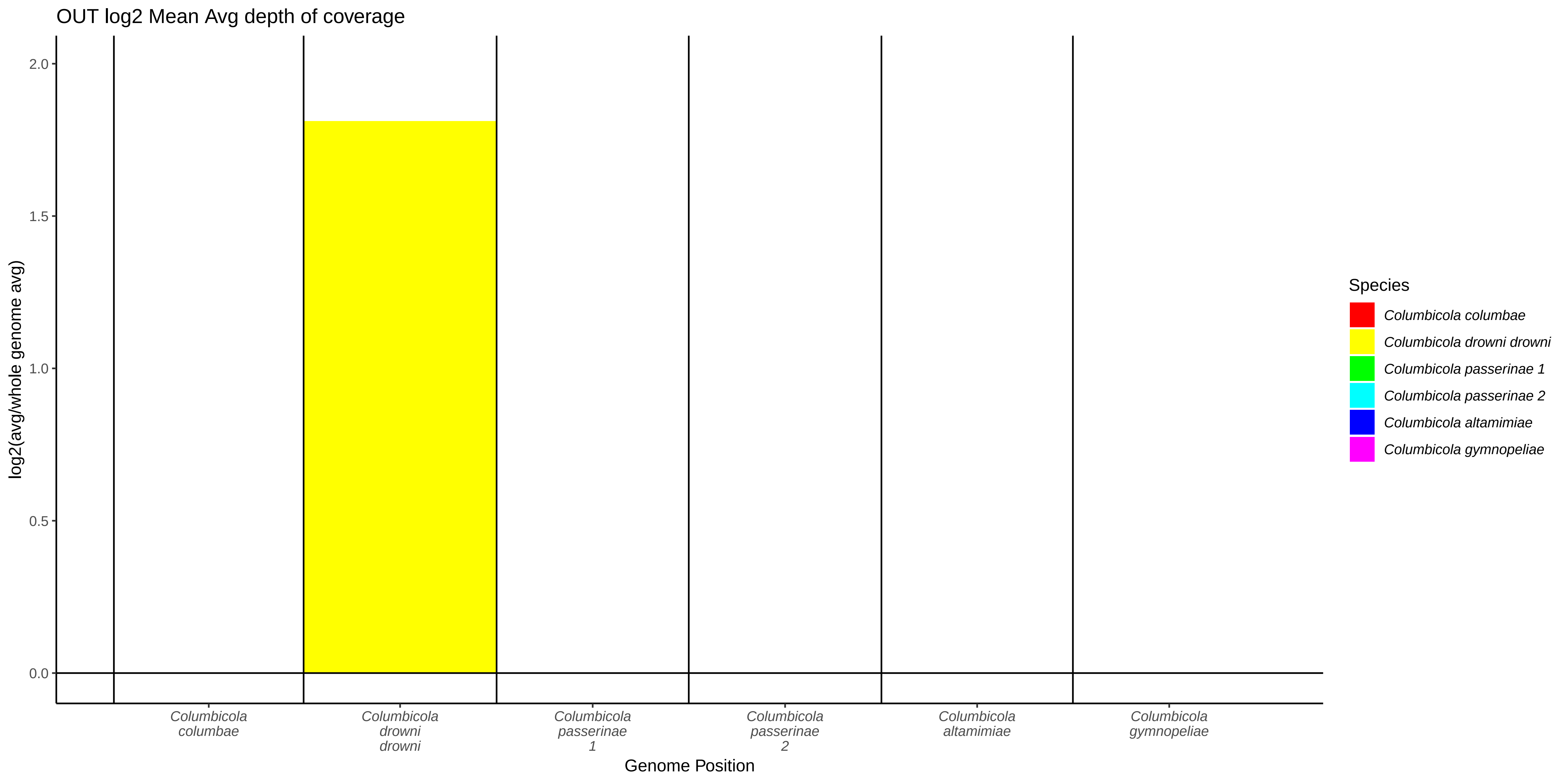

### OUT_speciesDepth_Cosp_Urcam_11_7_2016_4.pdf

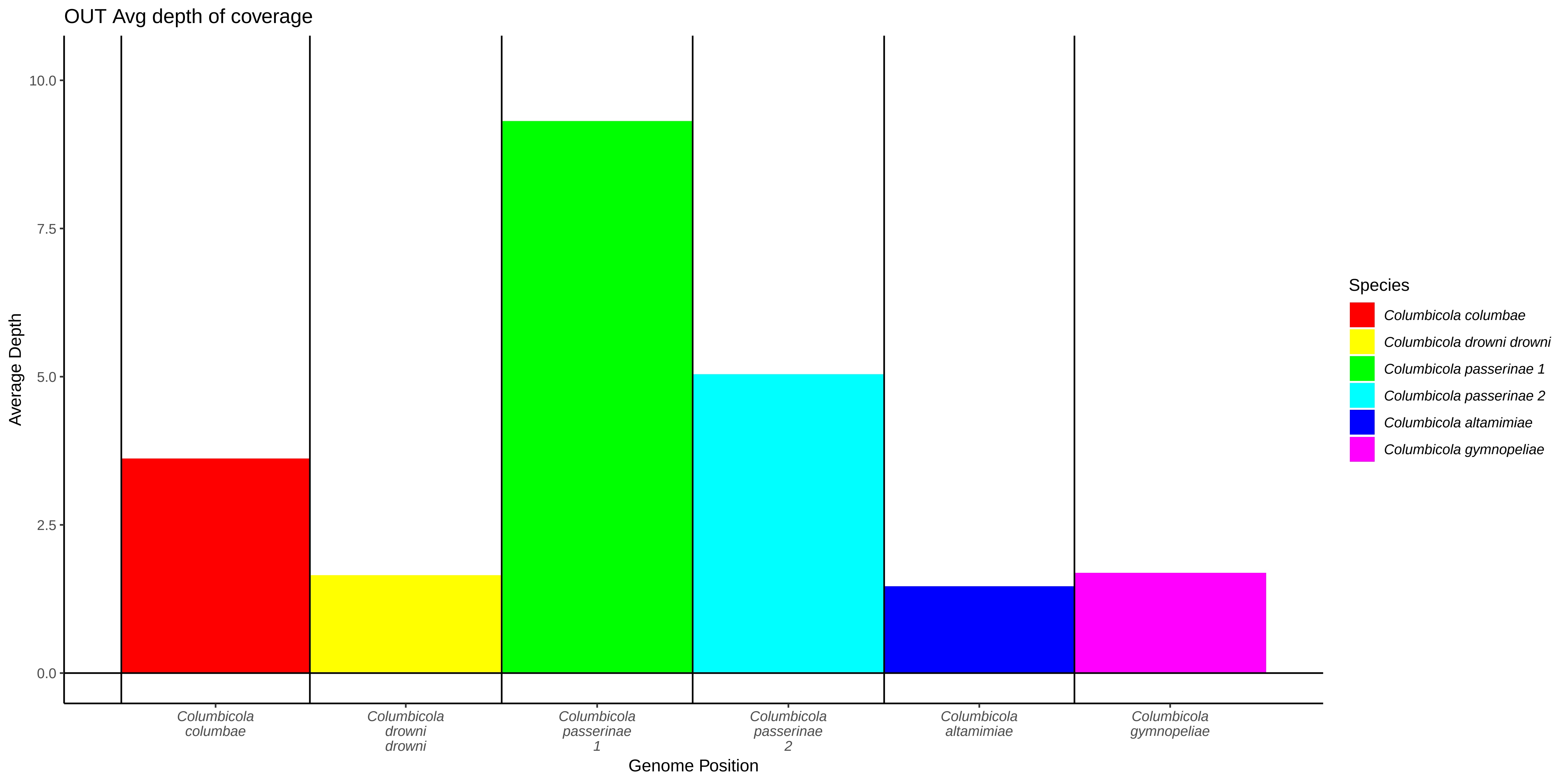

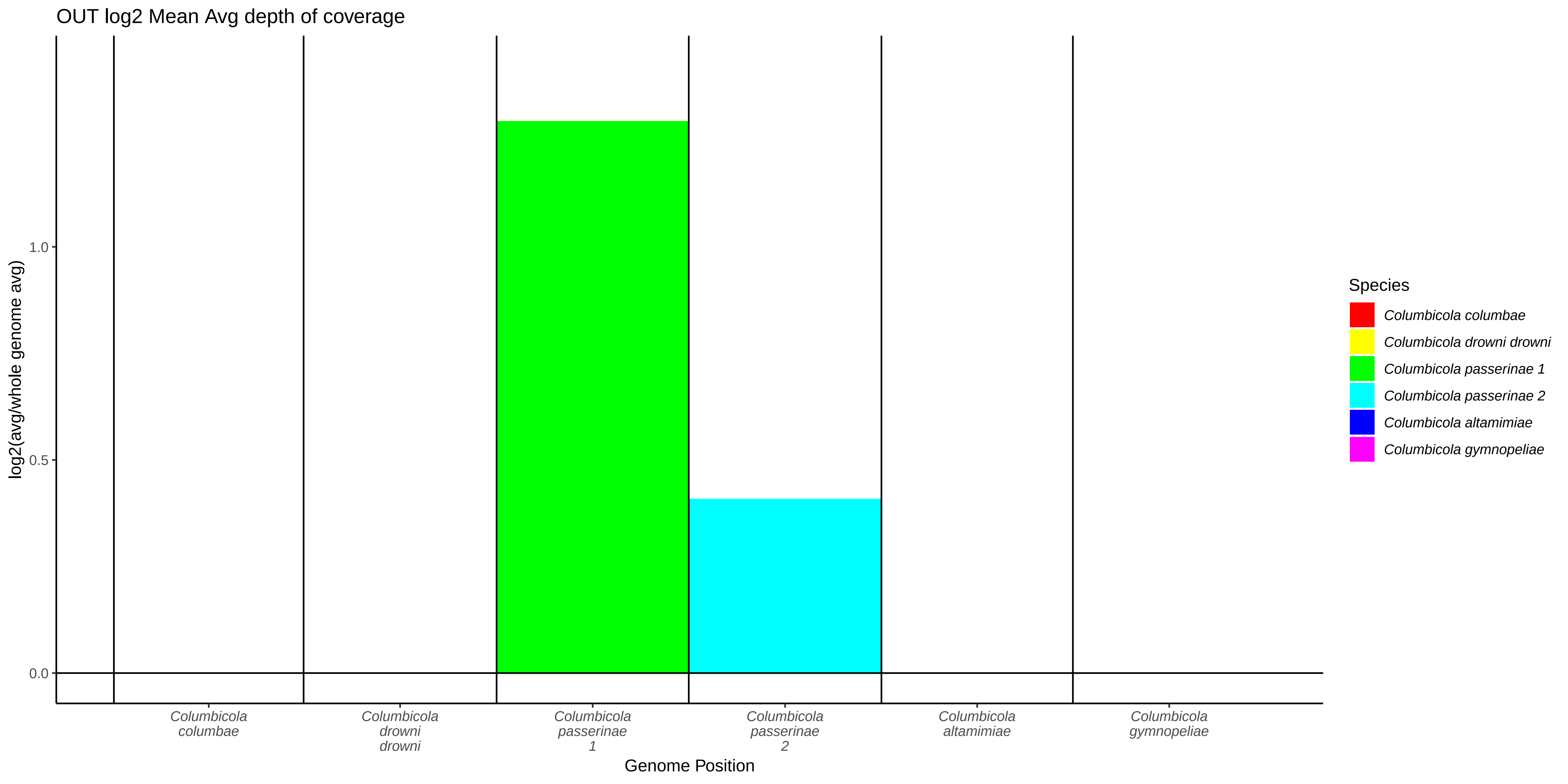

### OUT_speciesDepth_Cosp_Urcam_11_7_2016_6.pdf

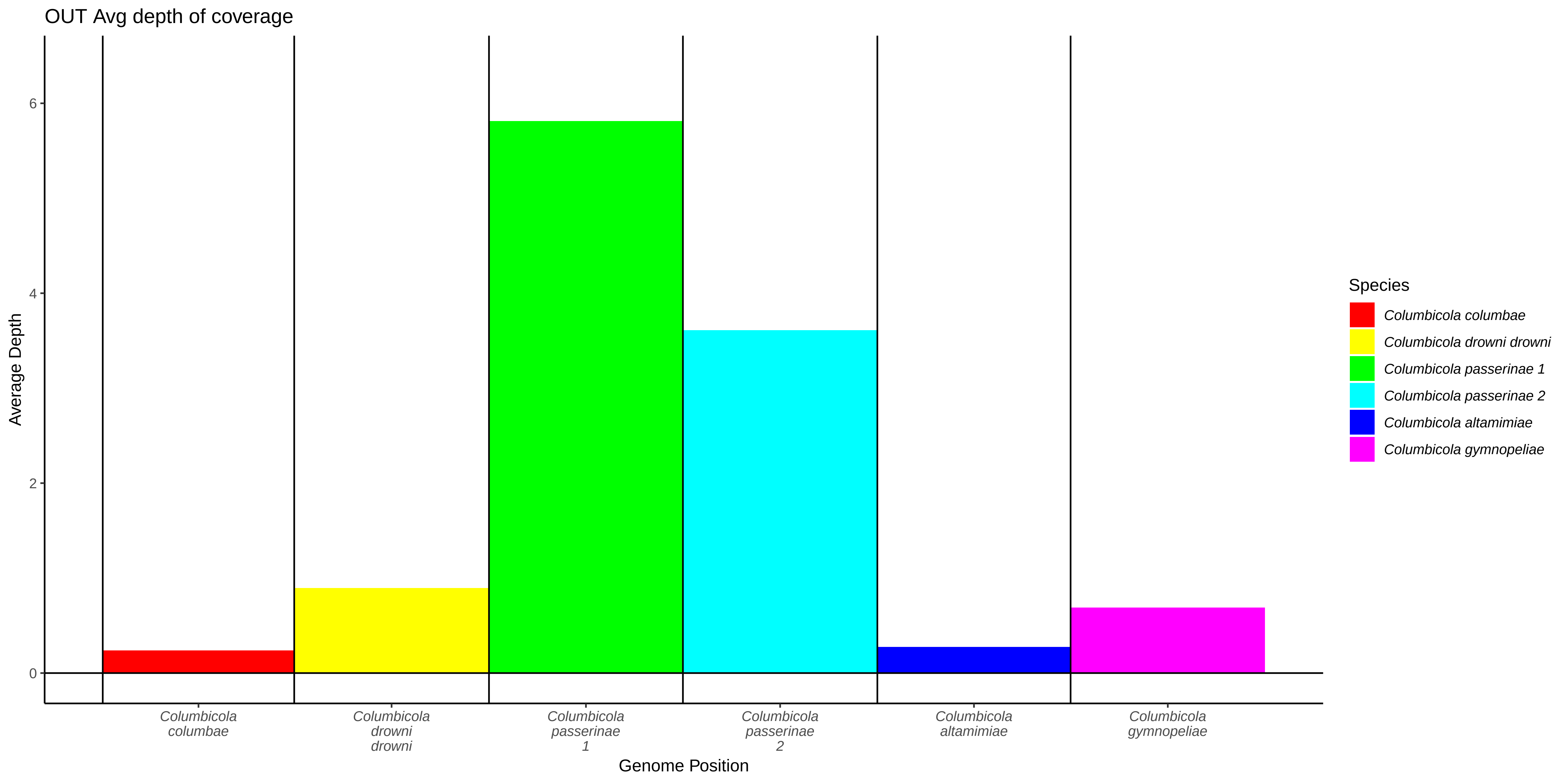

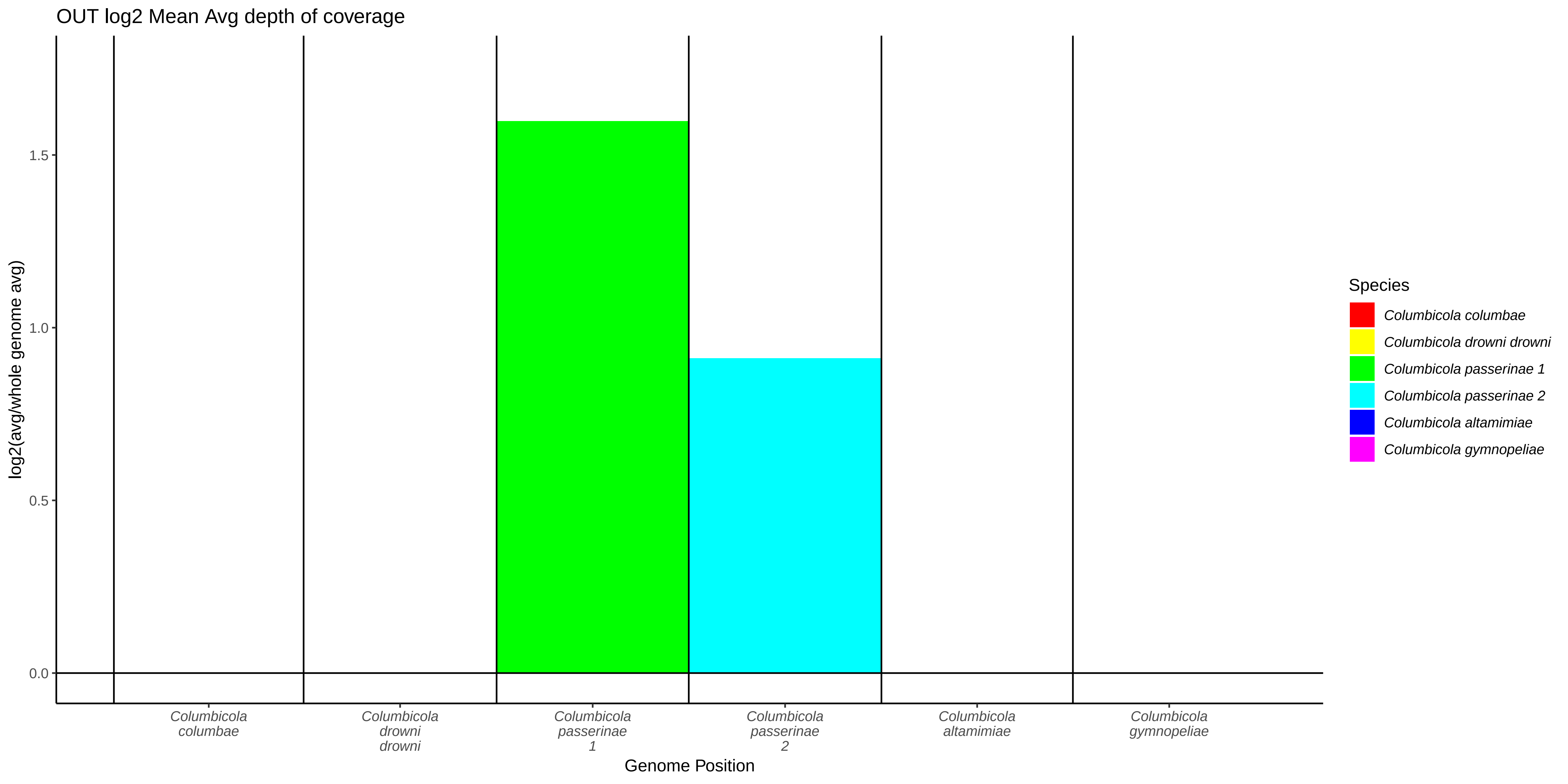

### OUT_speciesDepth_Pheme_Memel_3_16_2016_2.pdf

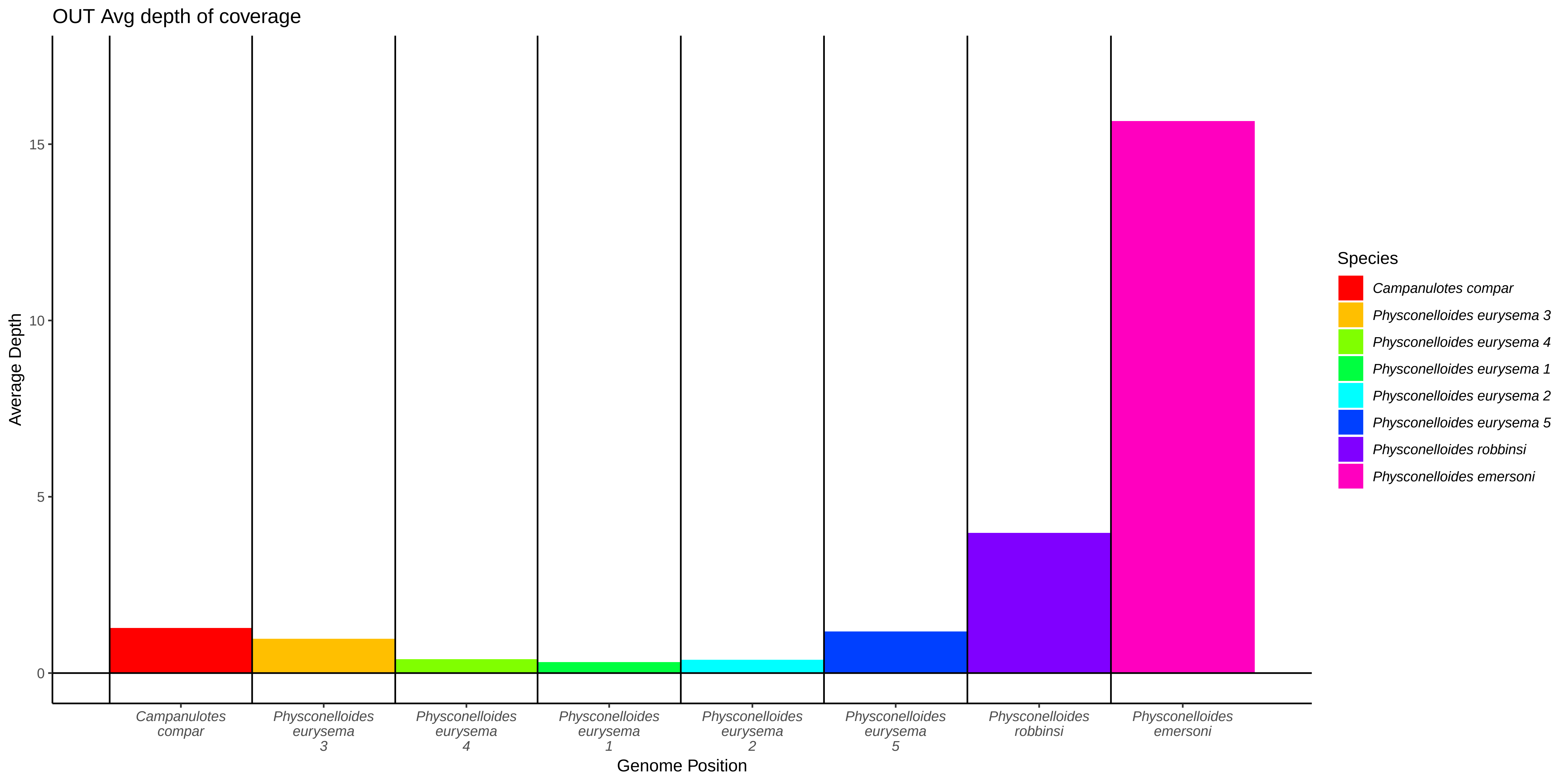

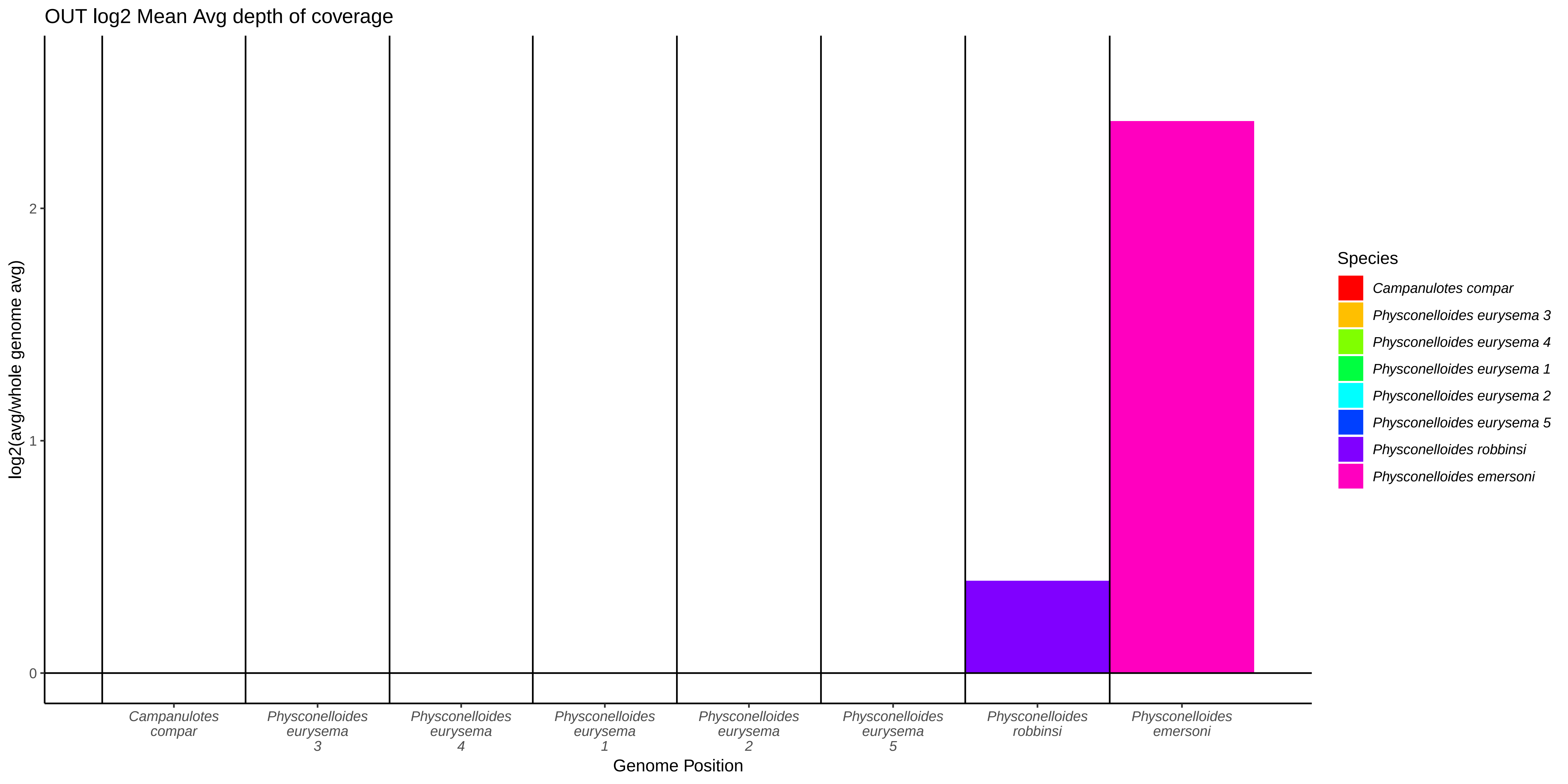

### OUT_speciesDepth_Pheme_Memel_3_16_2016_8.pdf

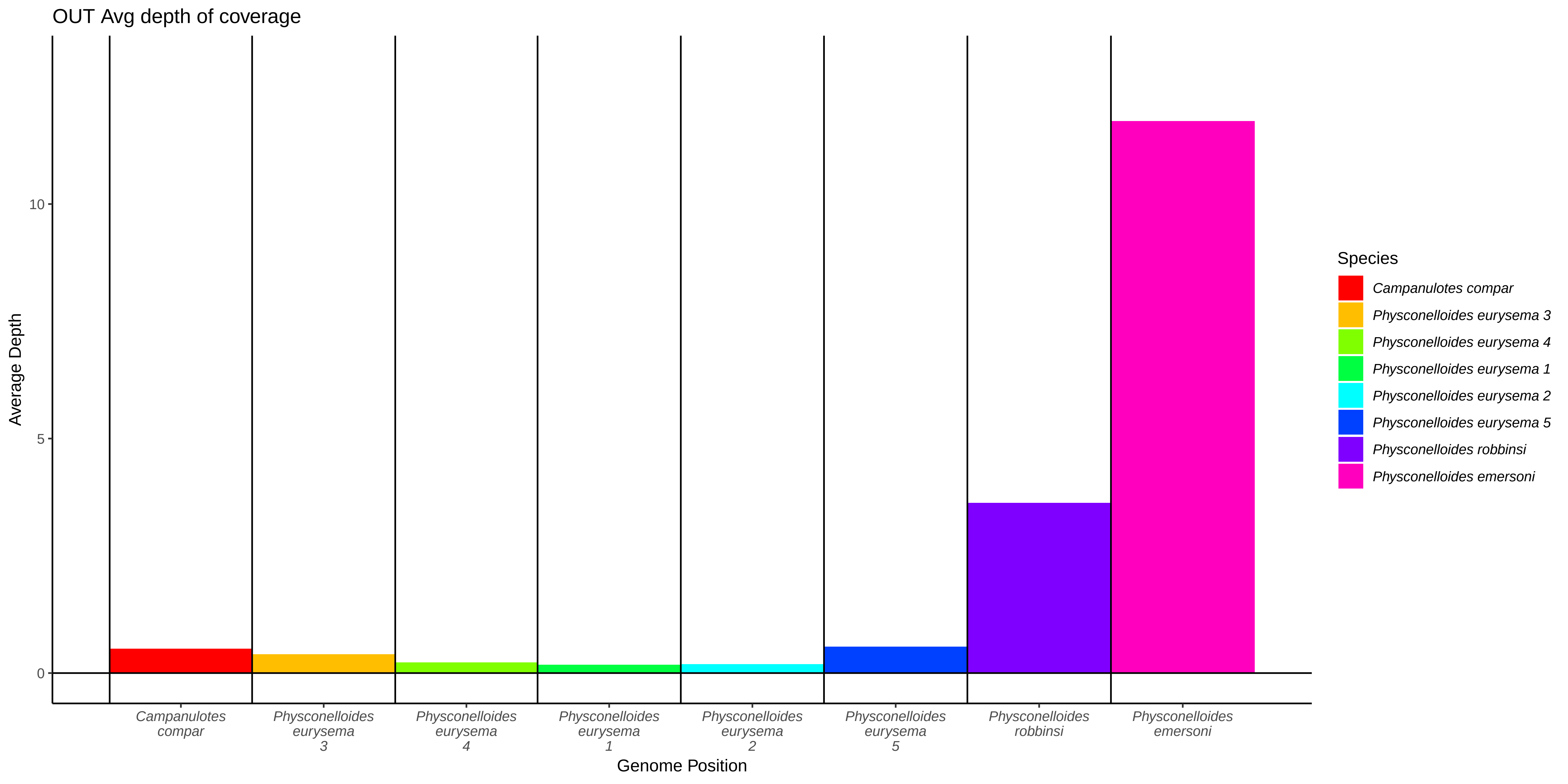

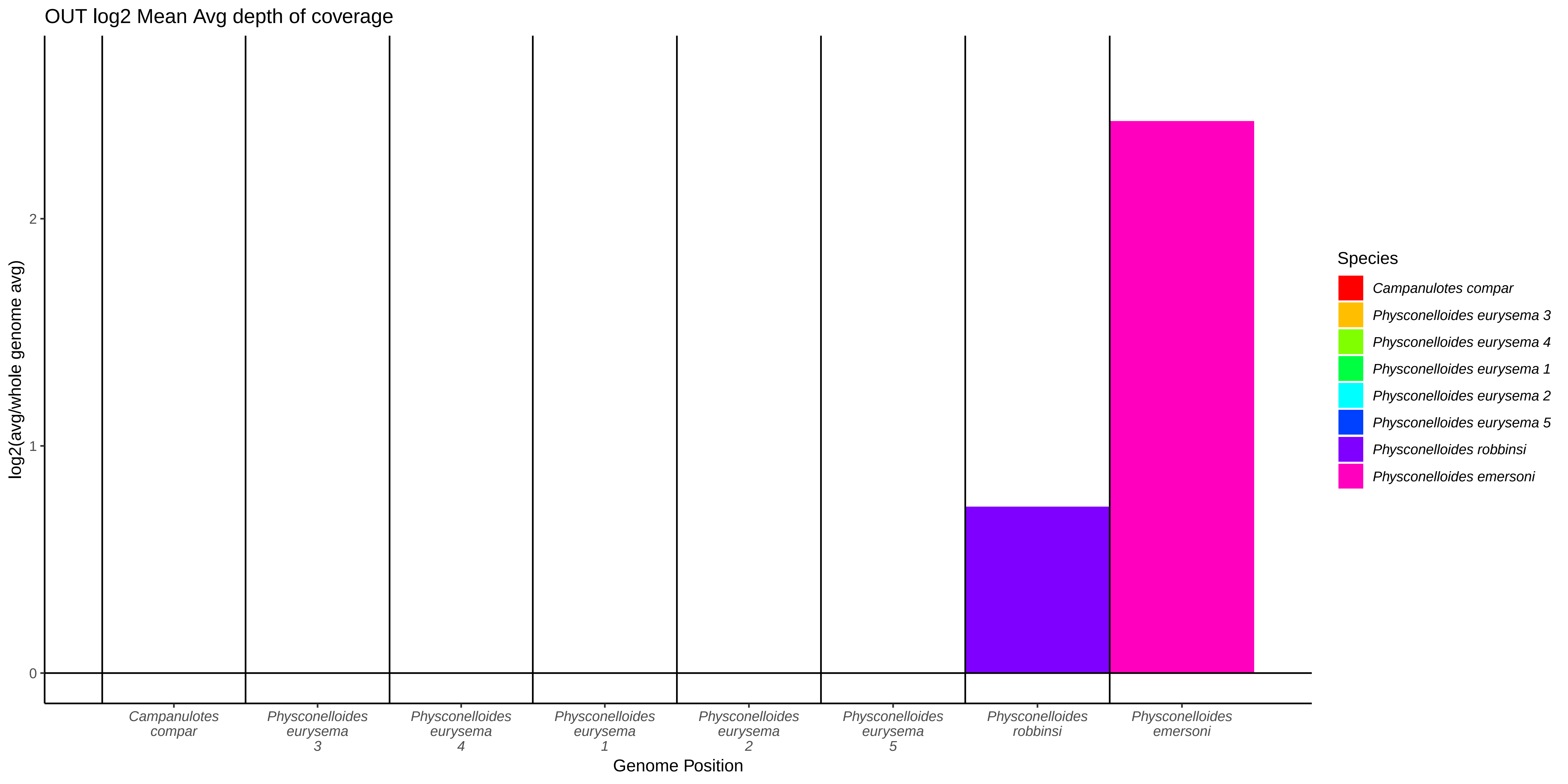

### OUT_speciesDepth_Pheme_Memel_5_26_2015_8.pdf

OUT Avg depth of coverage

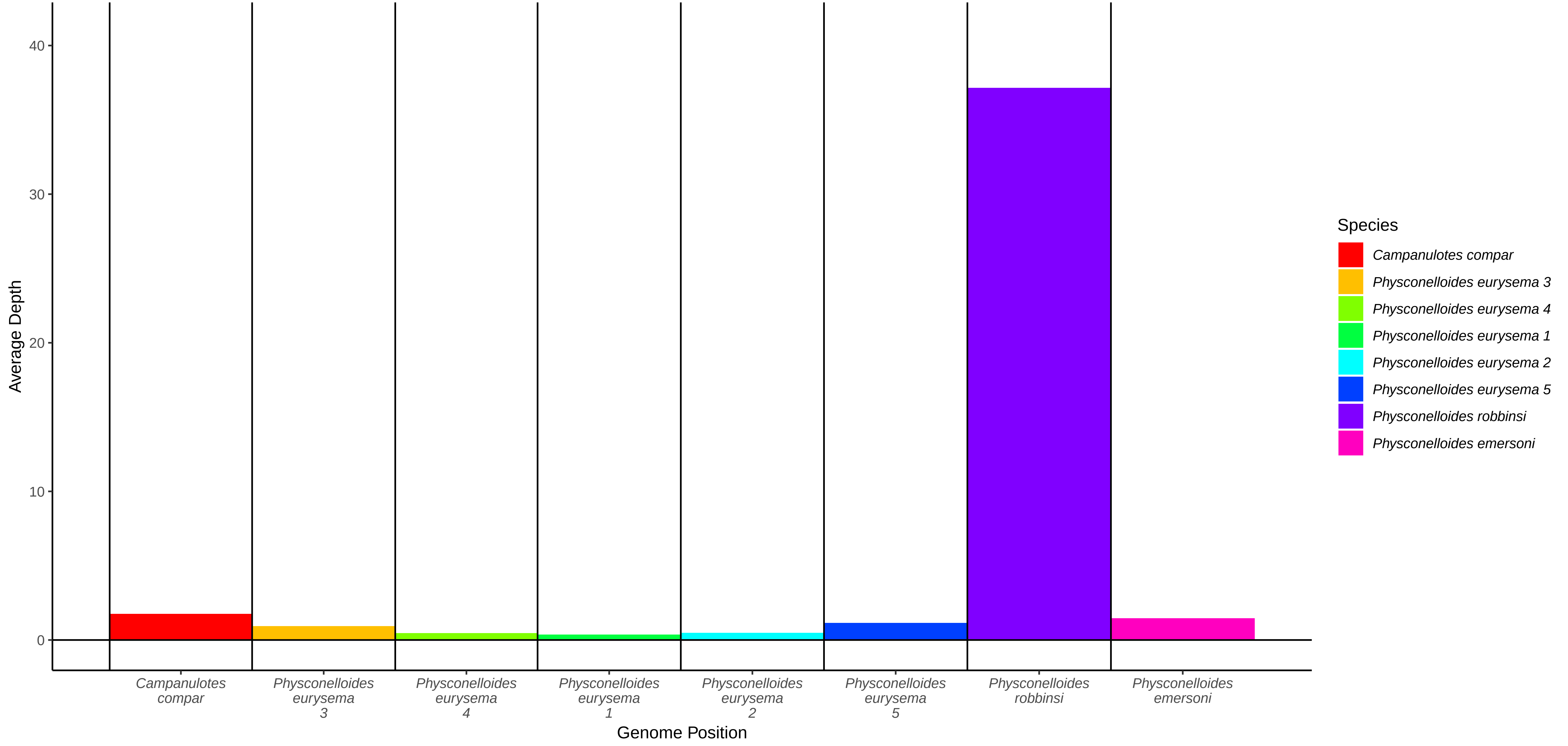

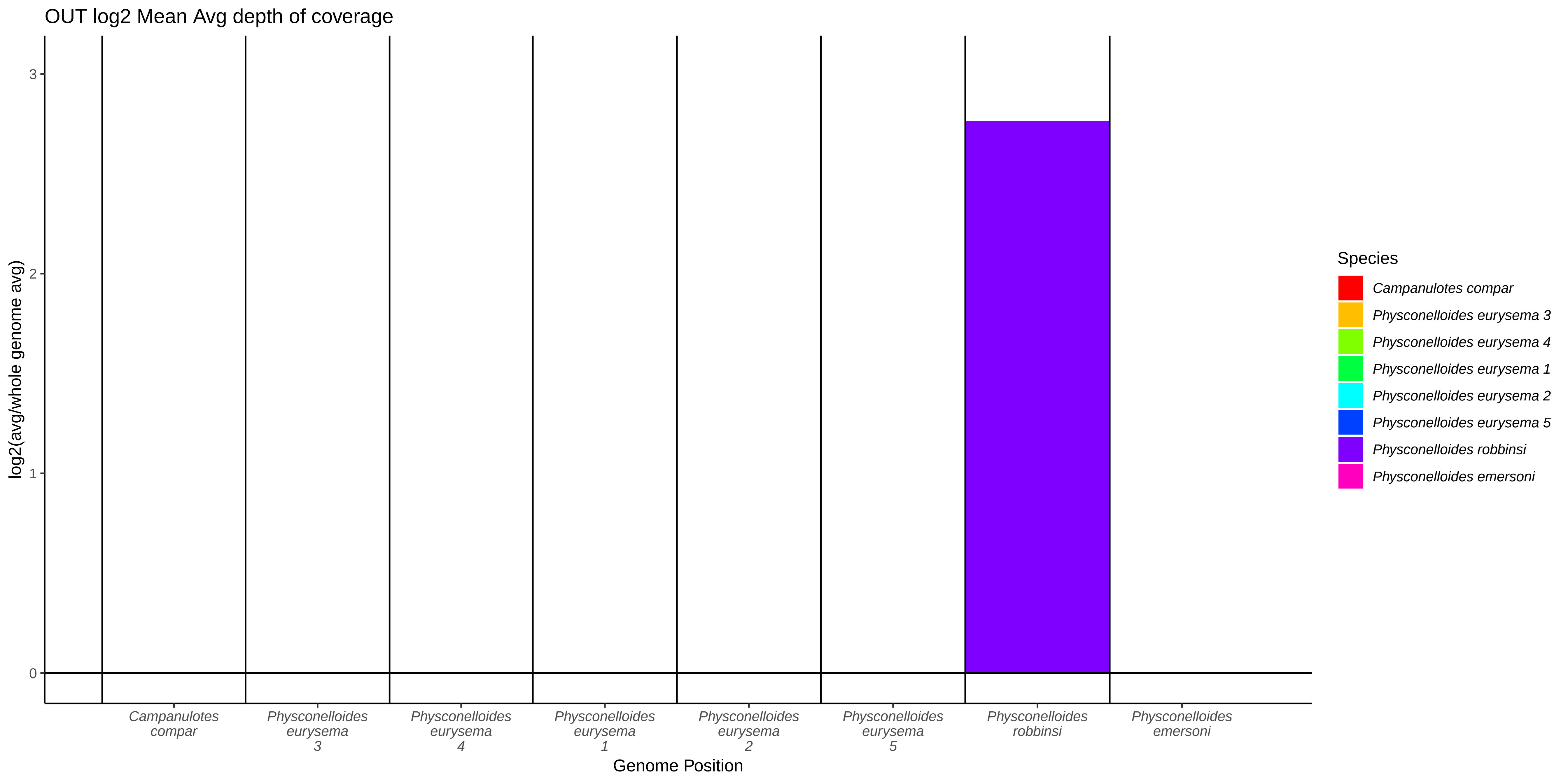

### OUT_speciesDepth_Pheme_Memel_6_15_2015_5.pdf

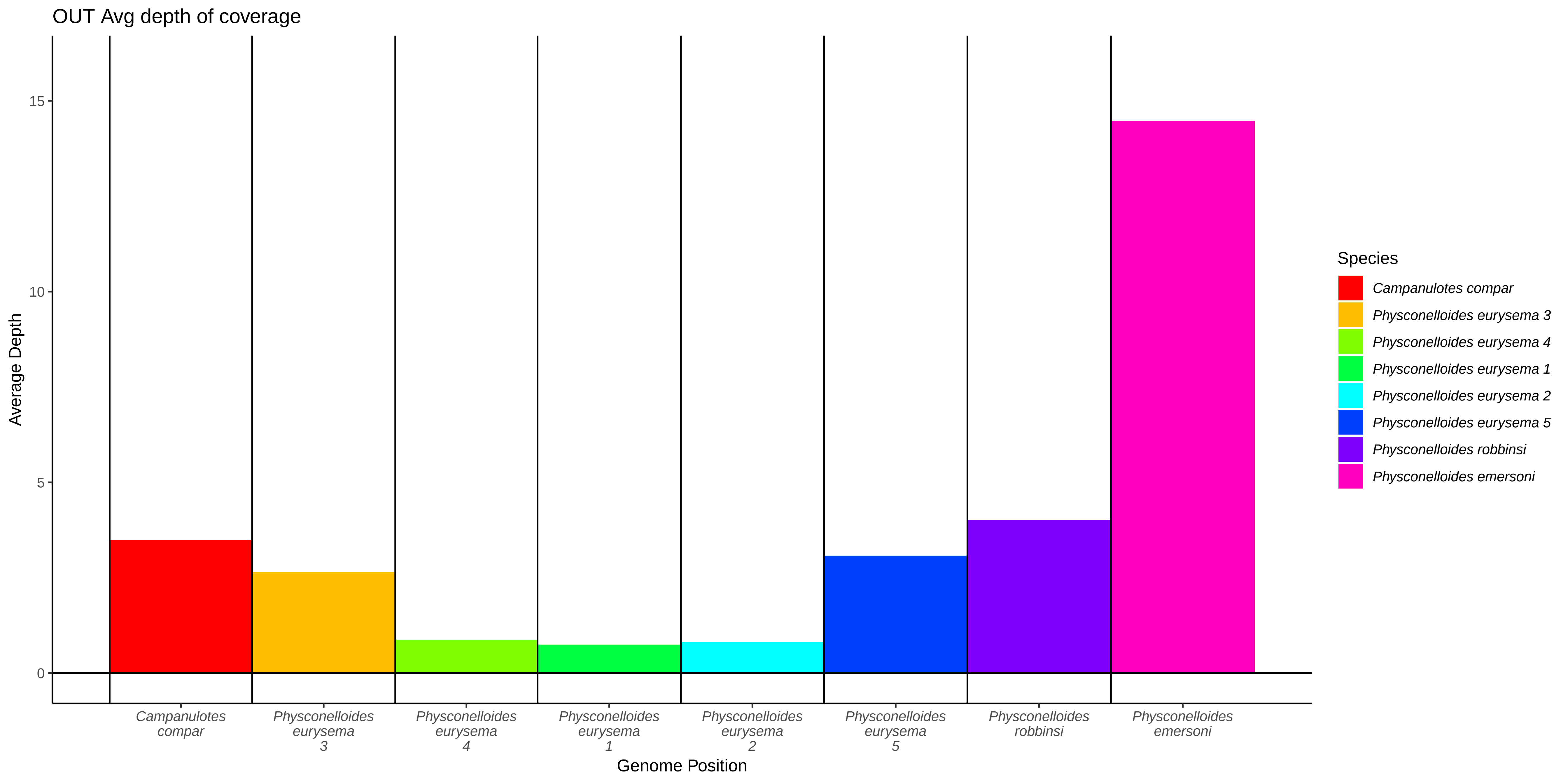

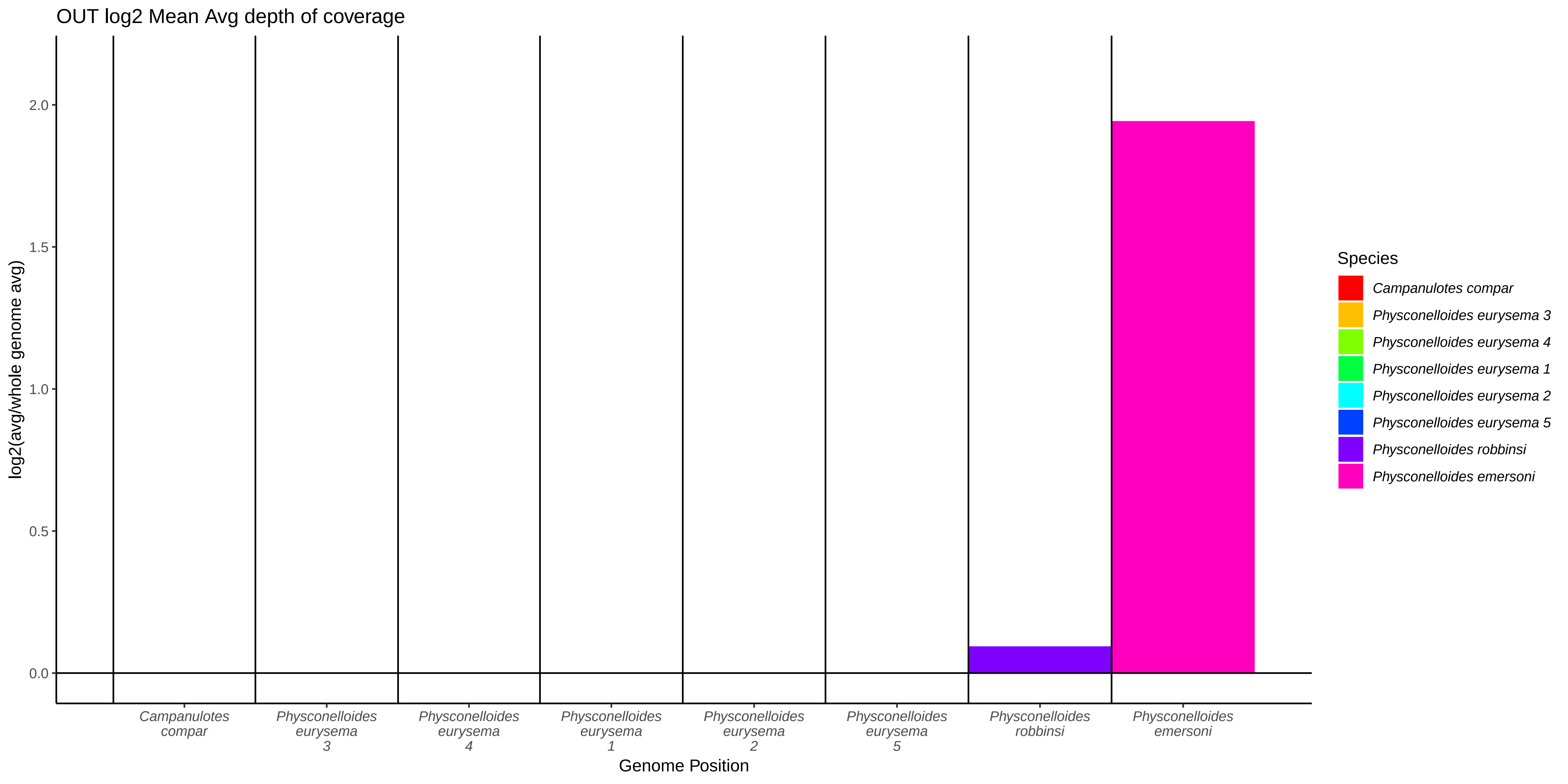

### OUT_speciesDepth_Pheur1_Comin_3_14_2016_9.pdf

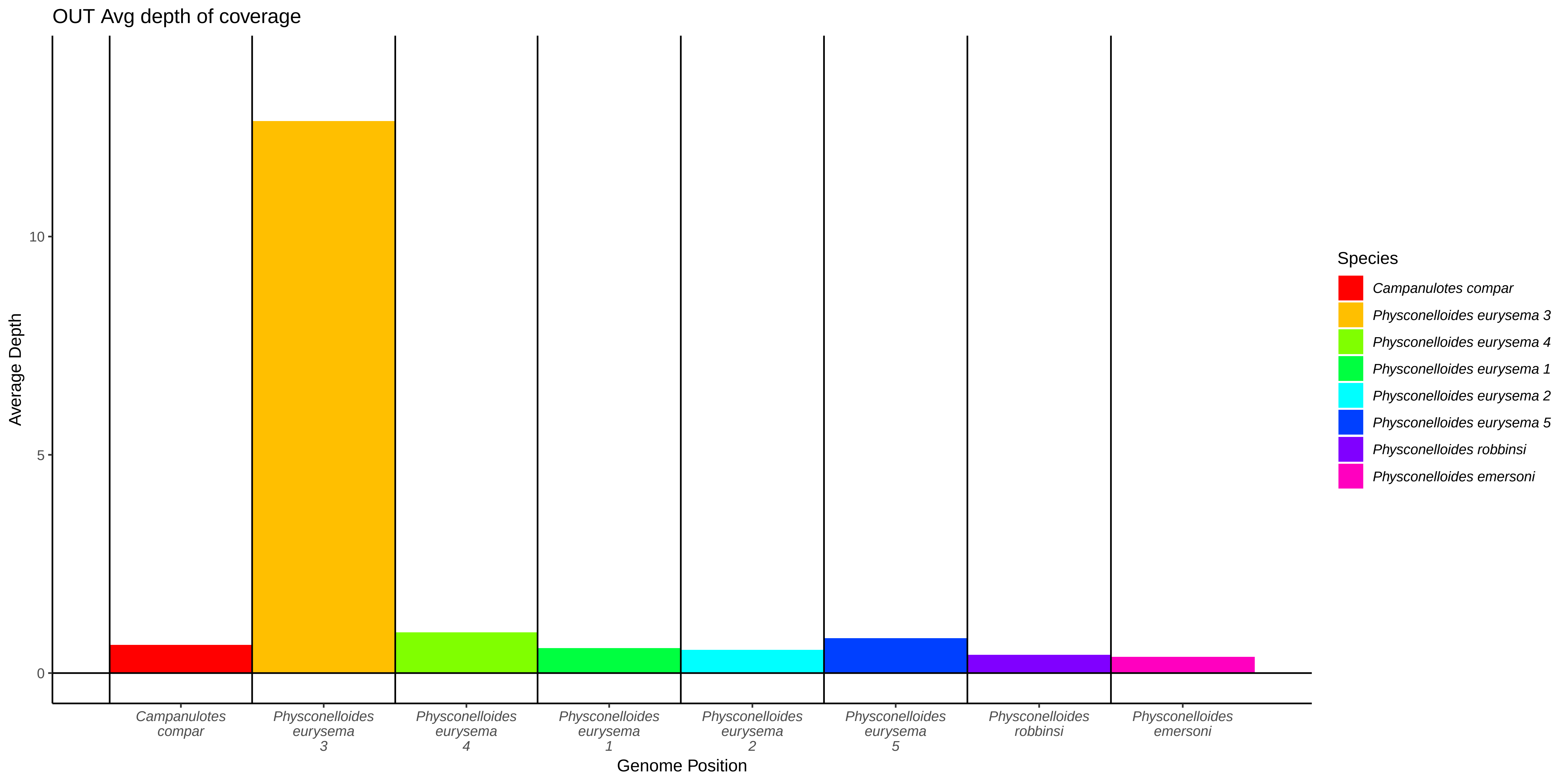

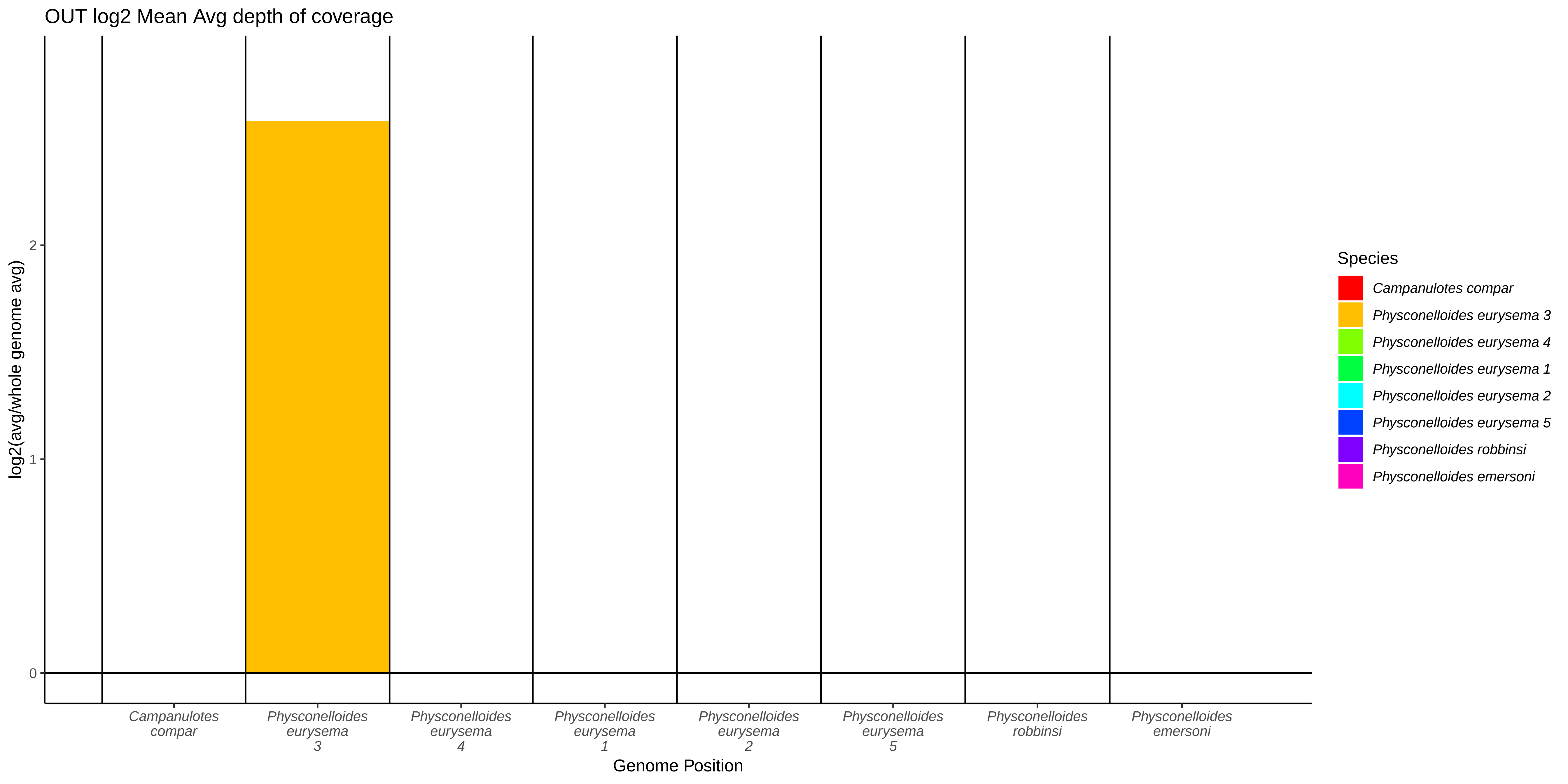

### OUT_speciesDepth_Pheur2_Clpre_3_14_2016_5.pdf

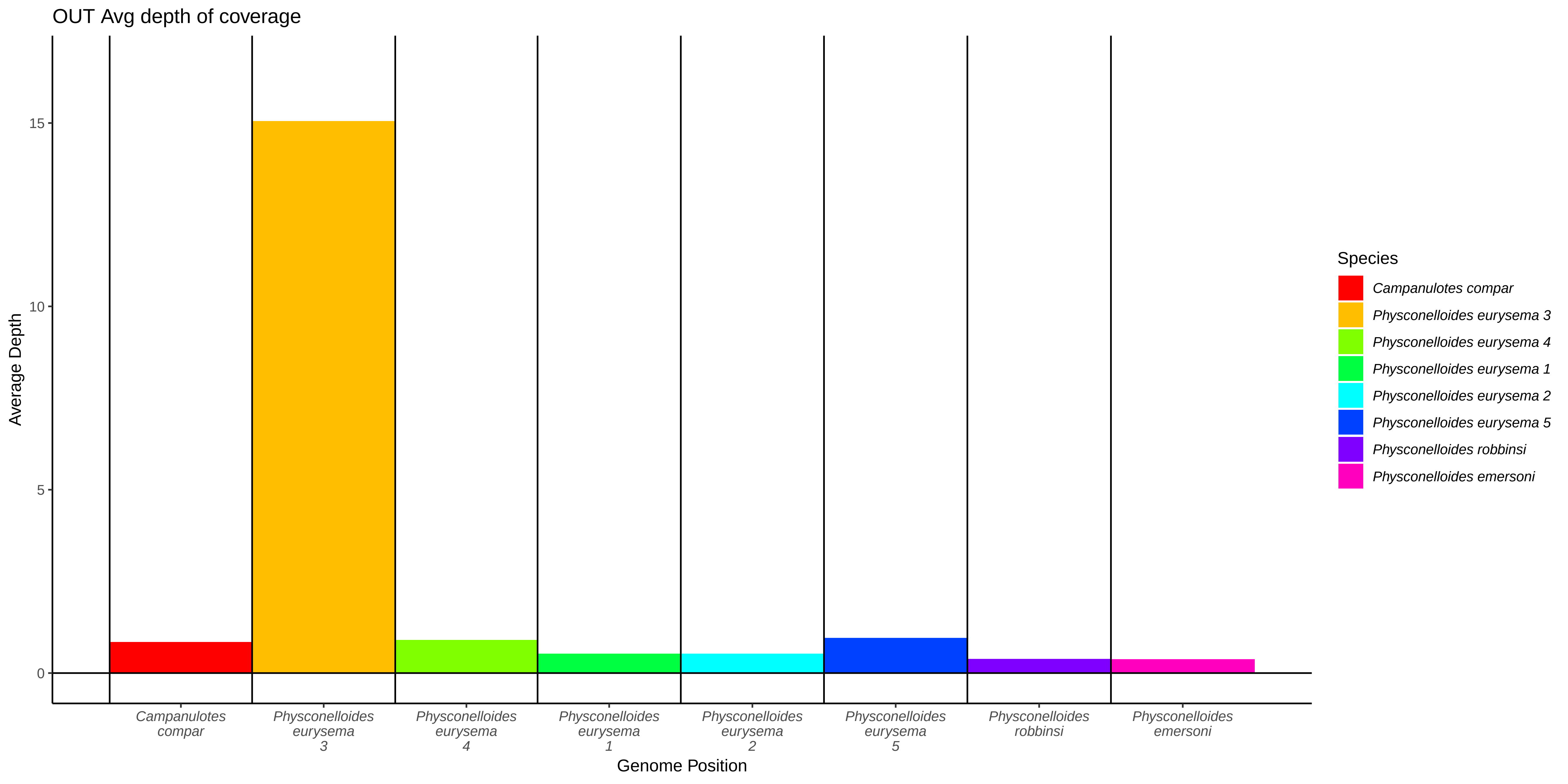

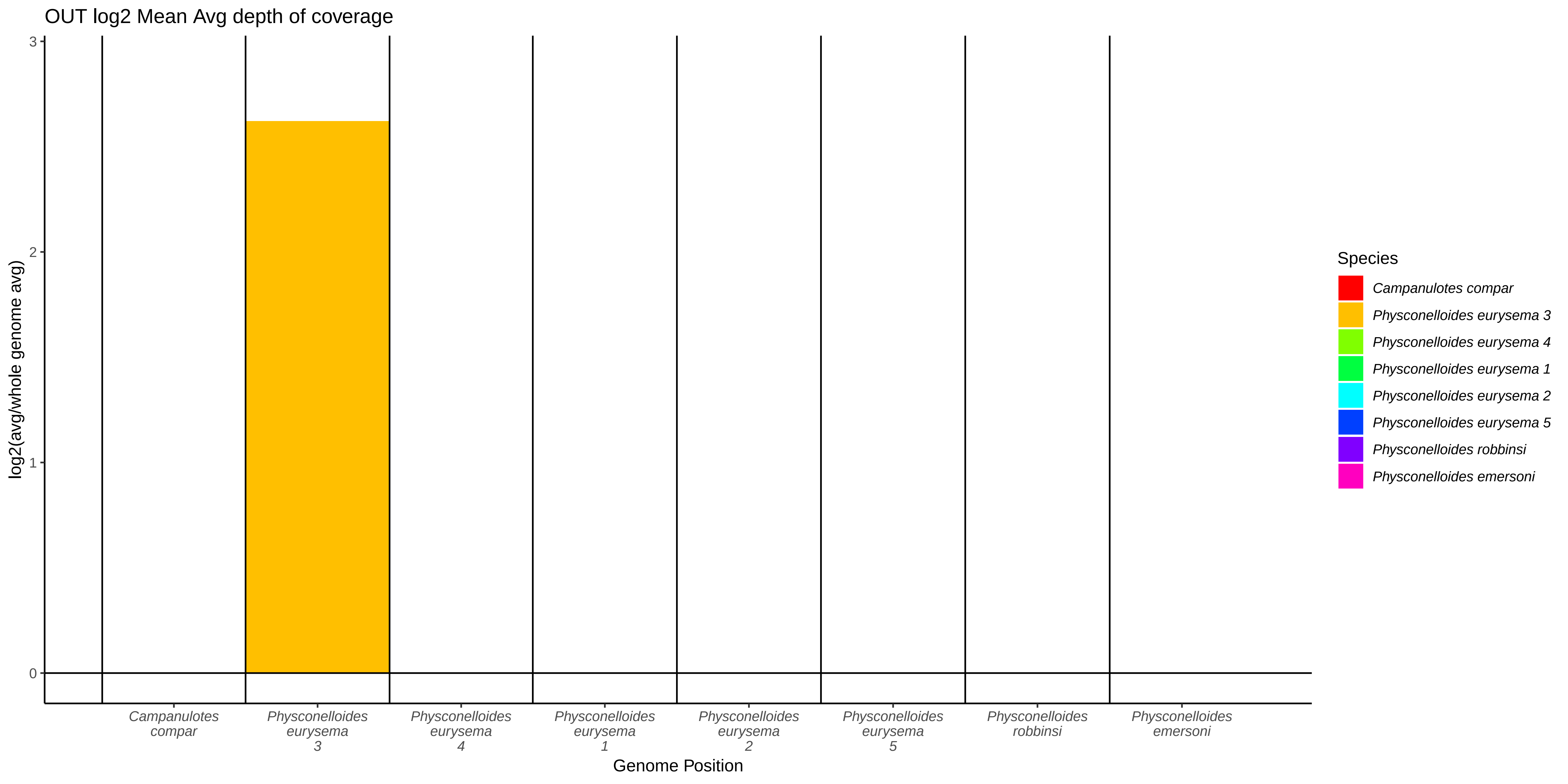

### OUT_speciesDepth_Pheur2_Clpre_3_14_2016_11.pdf

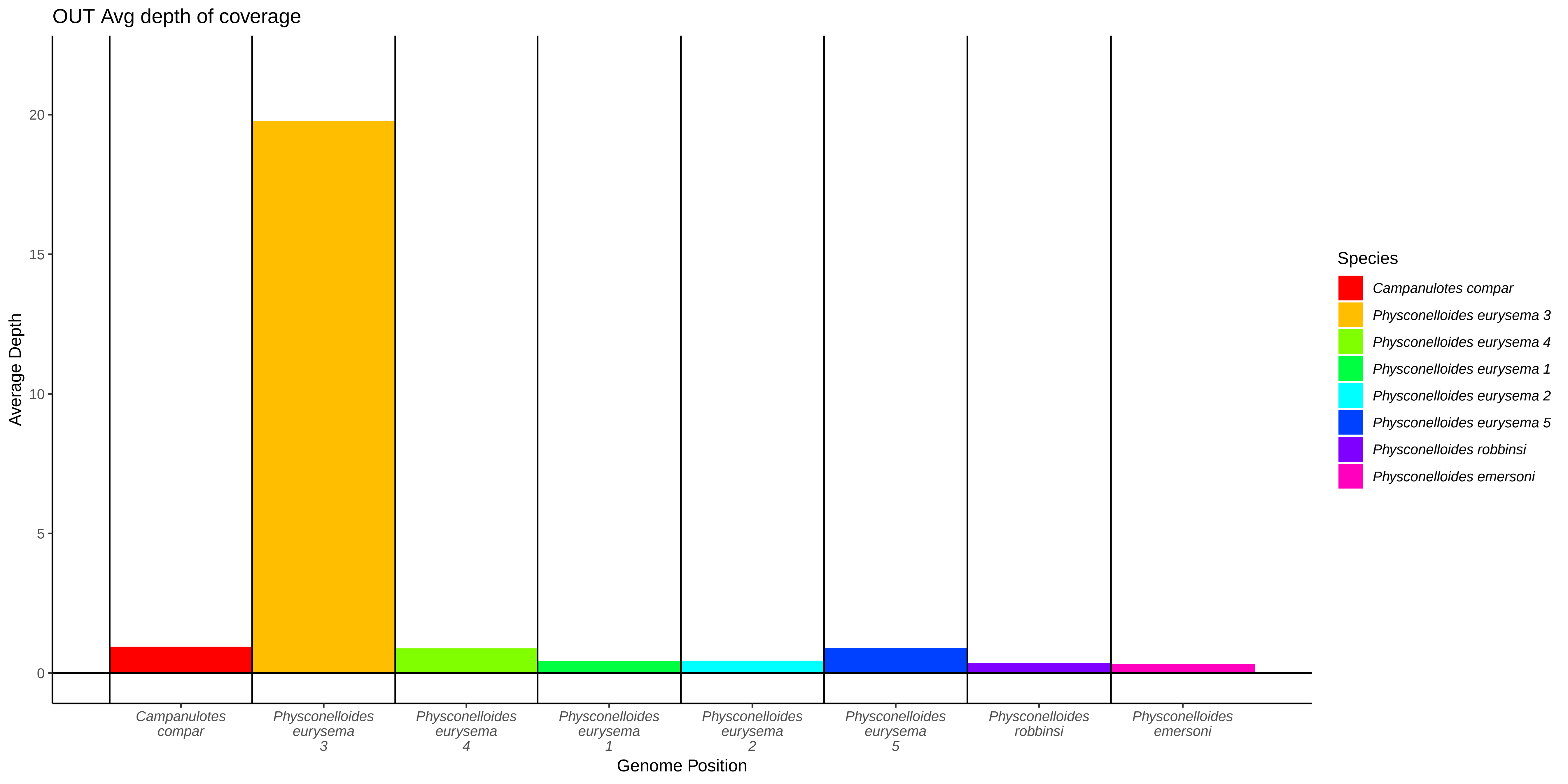

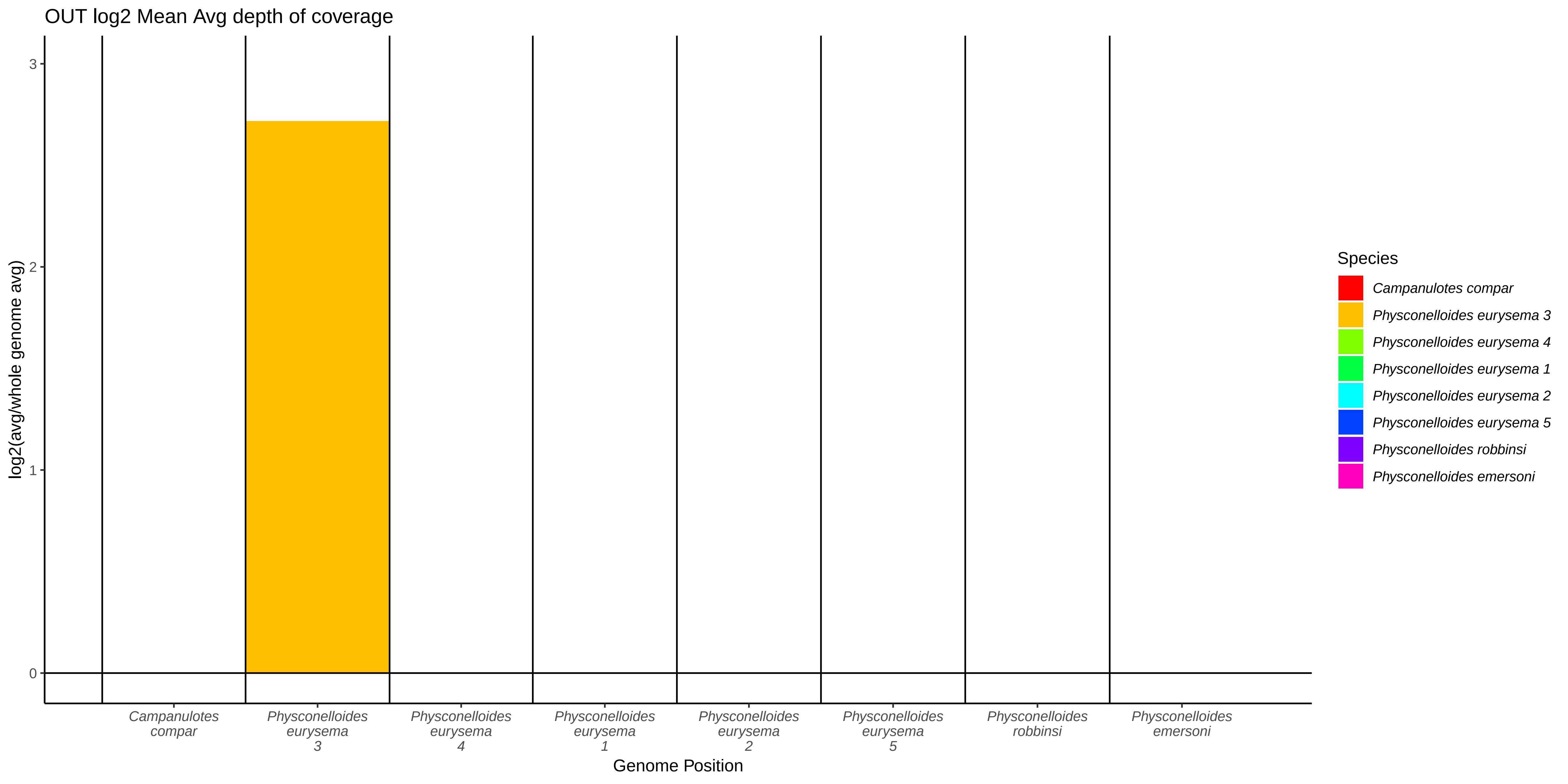

### OUT_speciesDepth_Pheur2_Clpre_3_16_2016_5.pdf

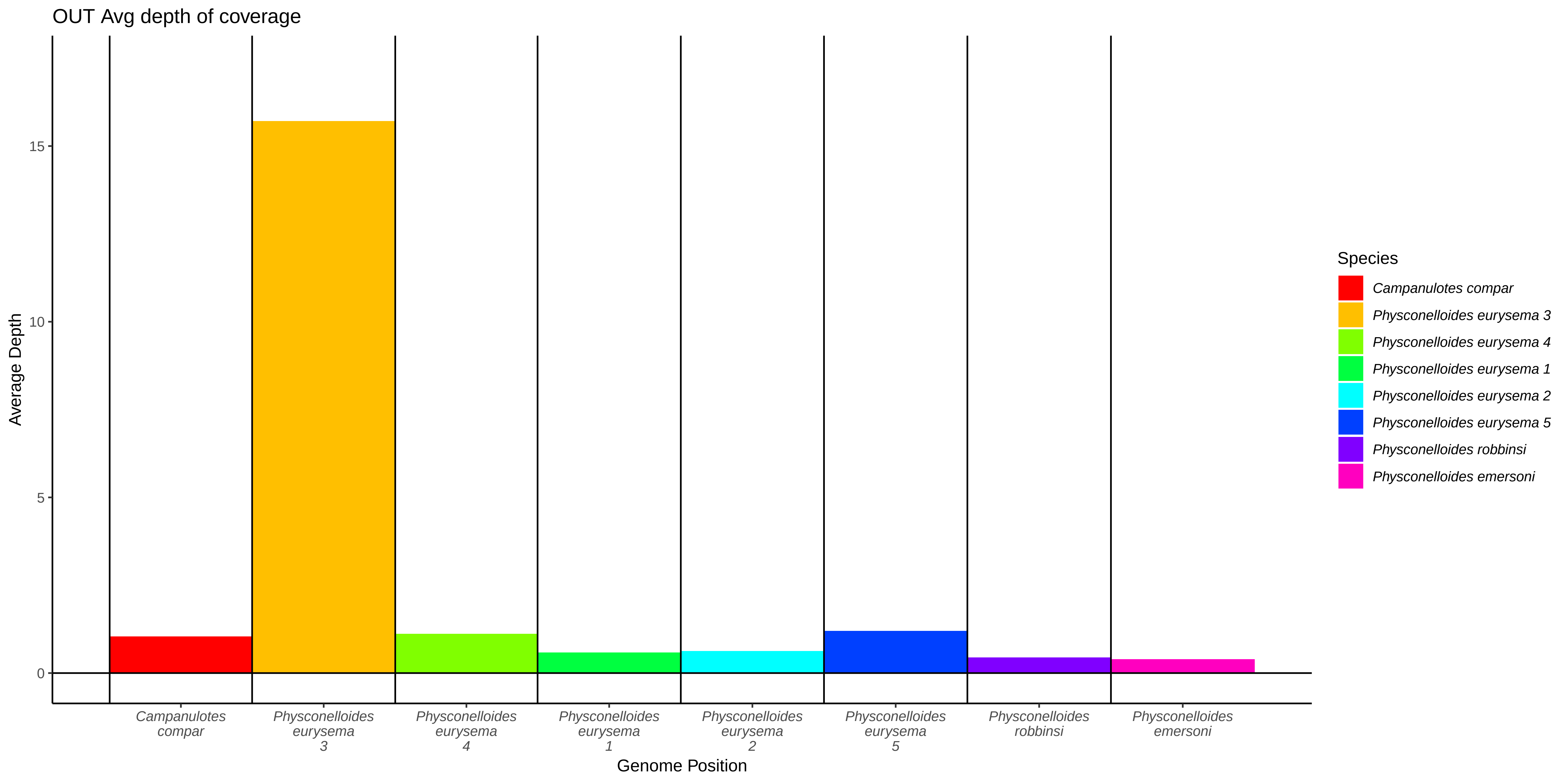

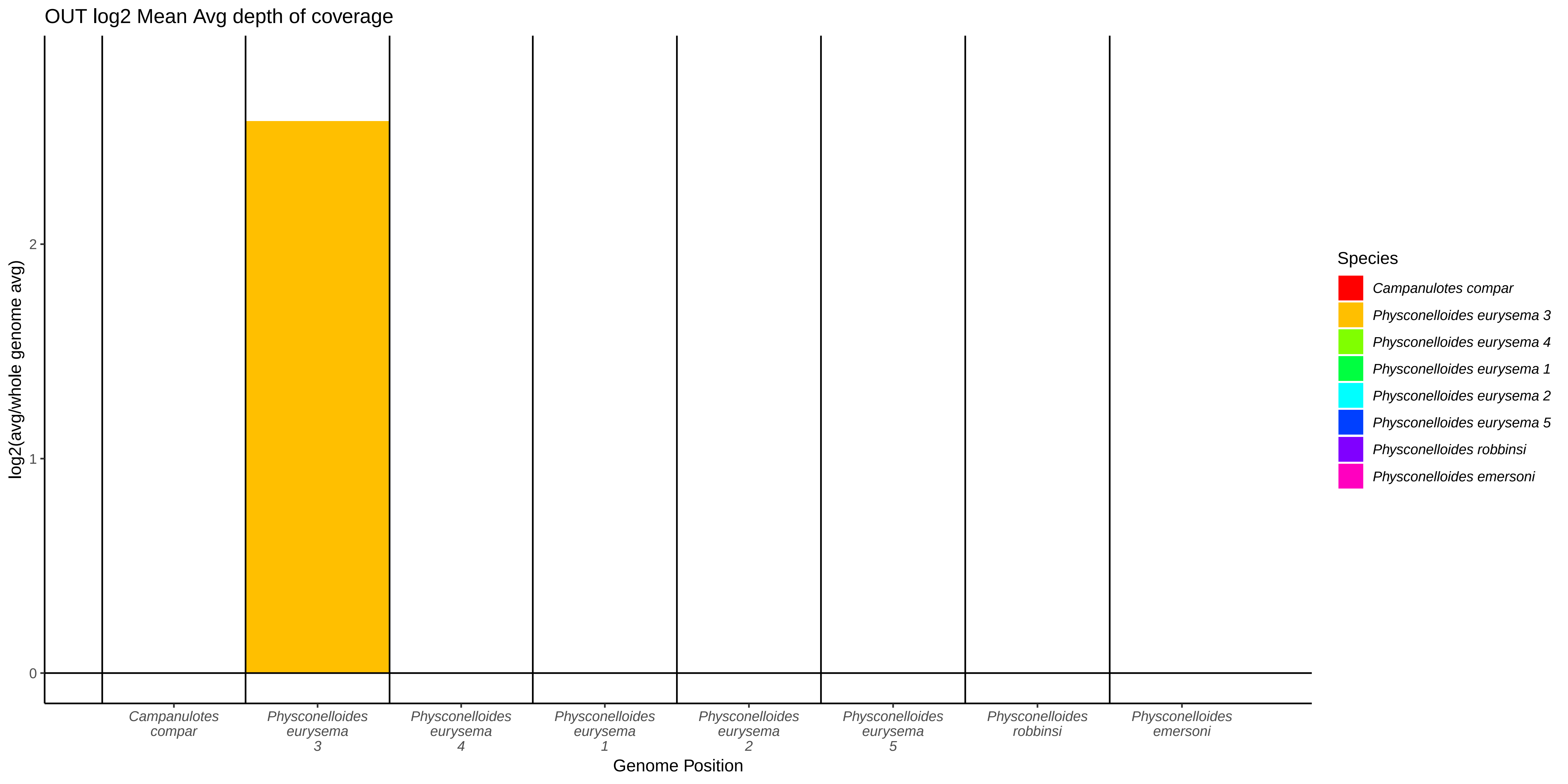

### OUT_speciesDepth_Pheur2_Clpre_5_26_2015_6.pdf

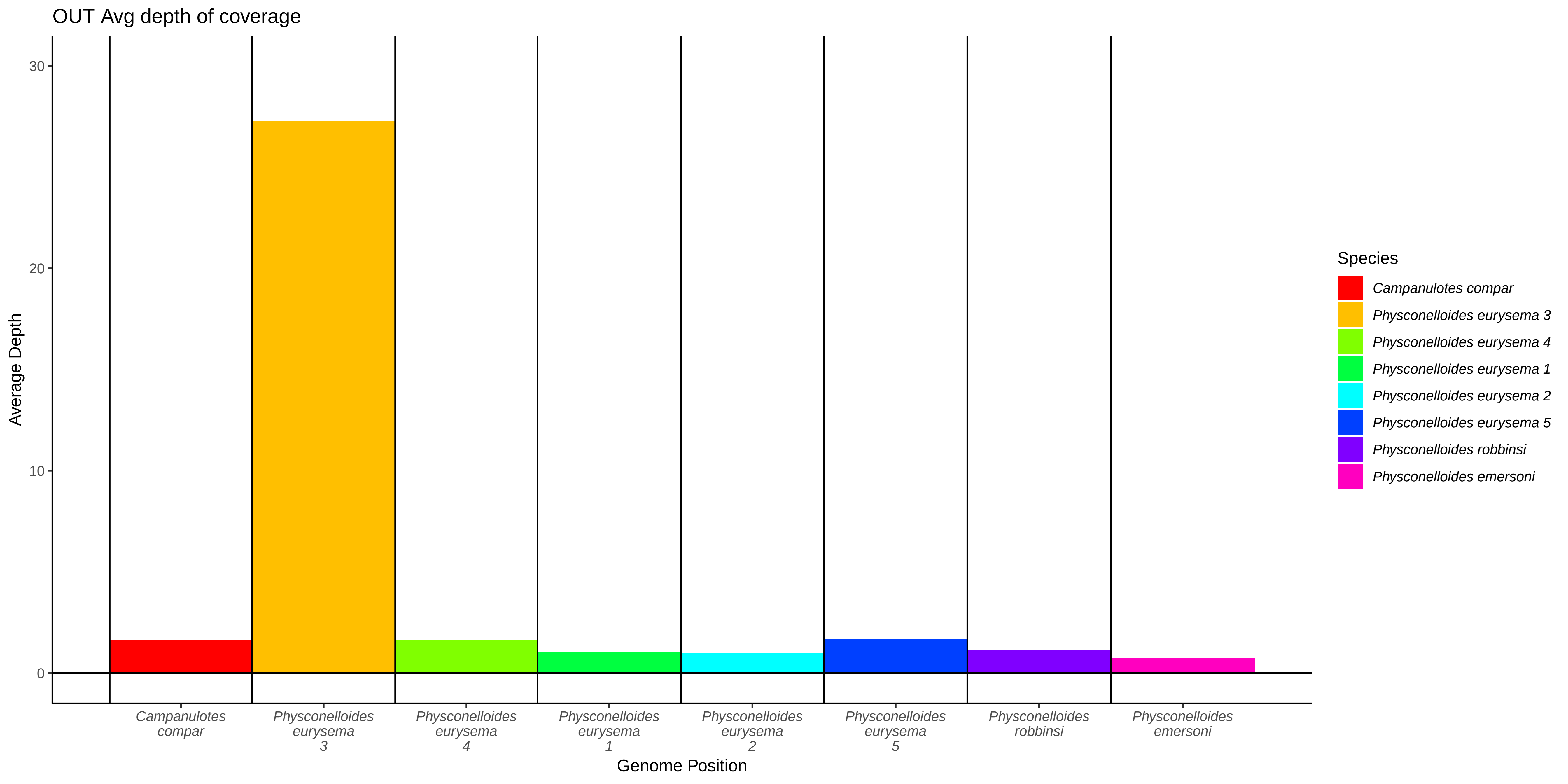

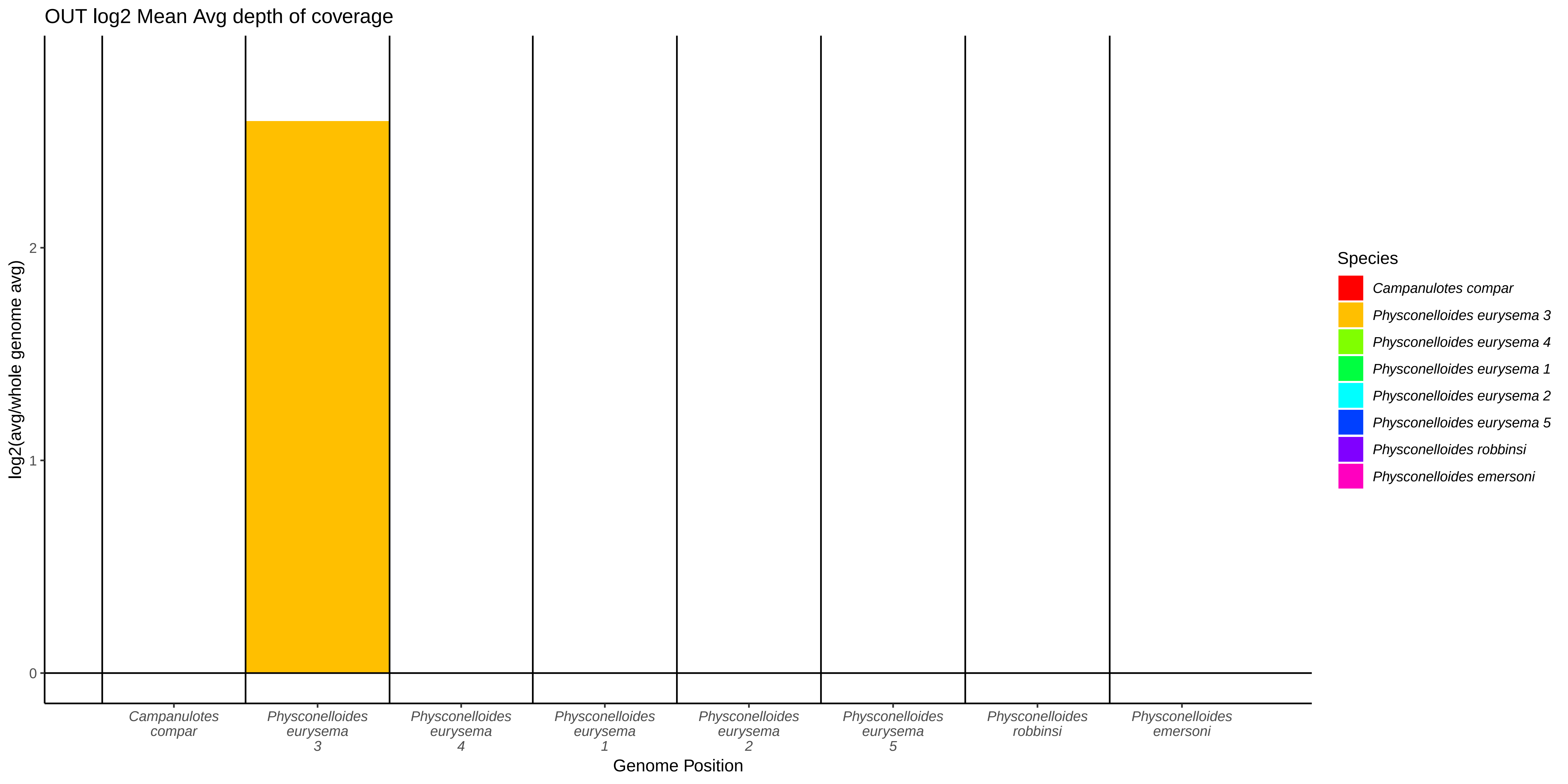

### OUT_speciesDepth_Pheur2_Cobuc_3_16_2016_4.pdf

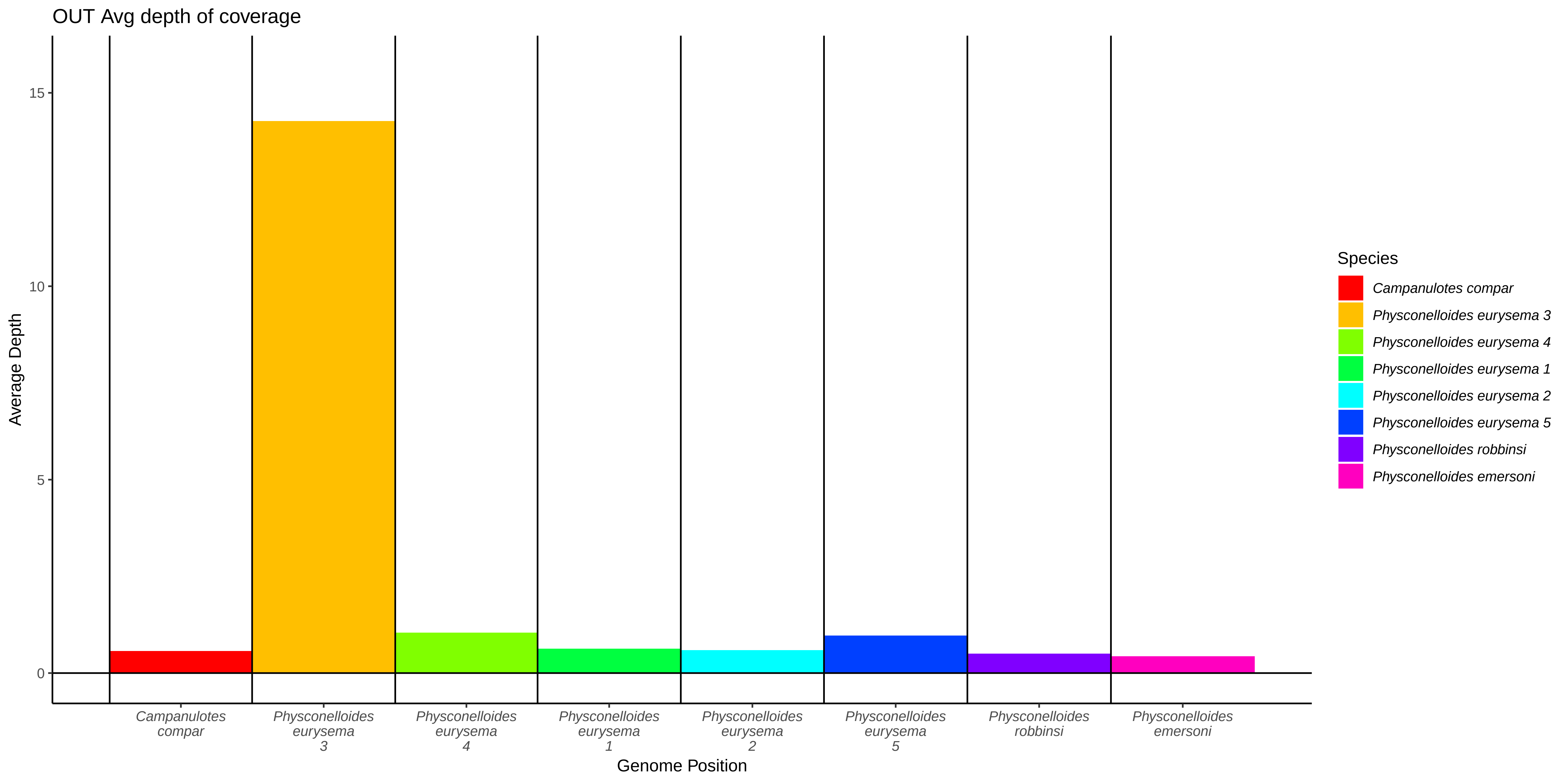

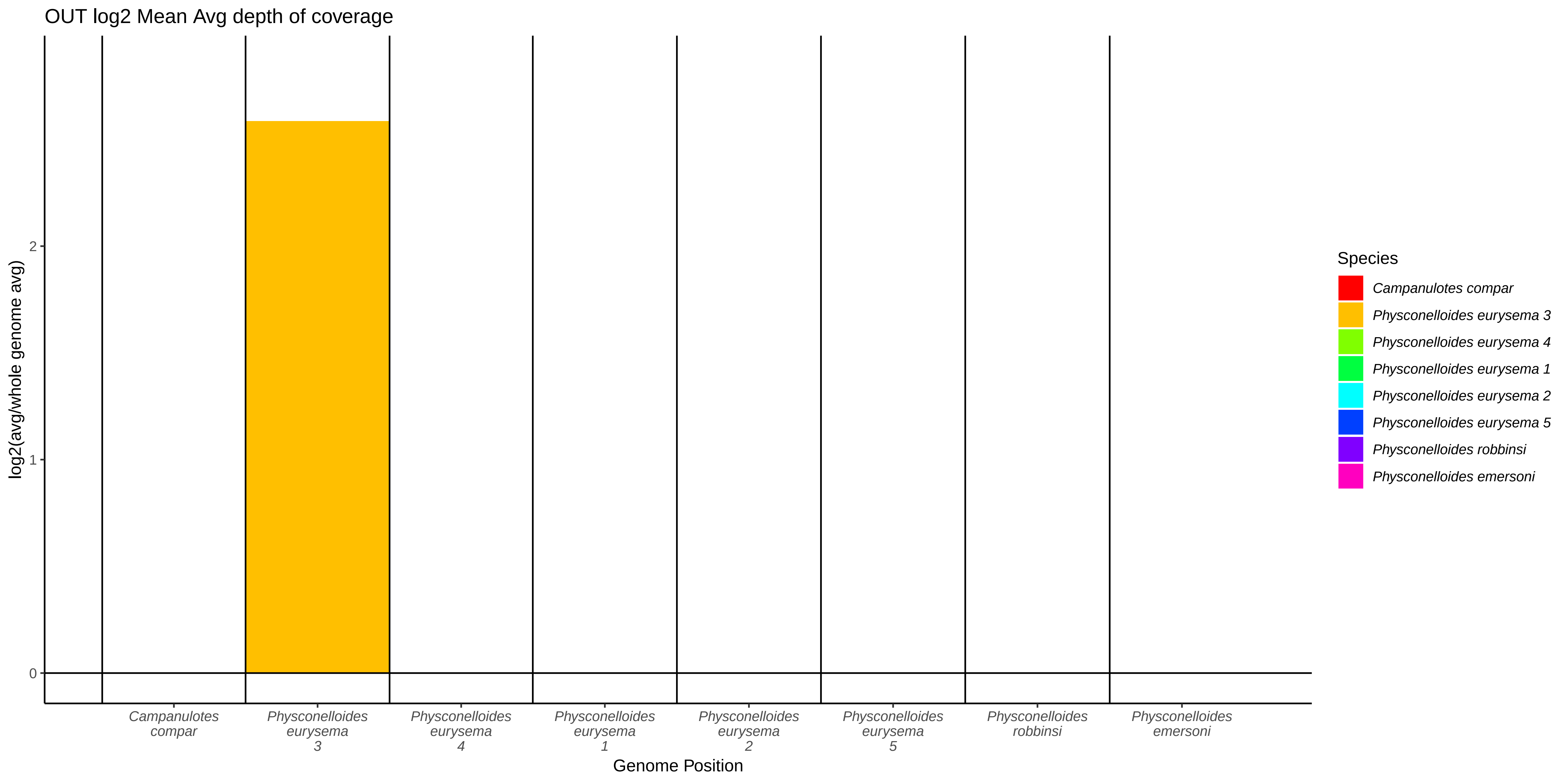

### OUT_speciesDepth_Pheur2_Cocru_3_16_2016_9.pdf

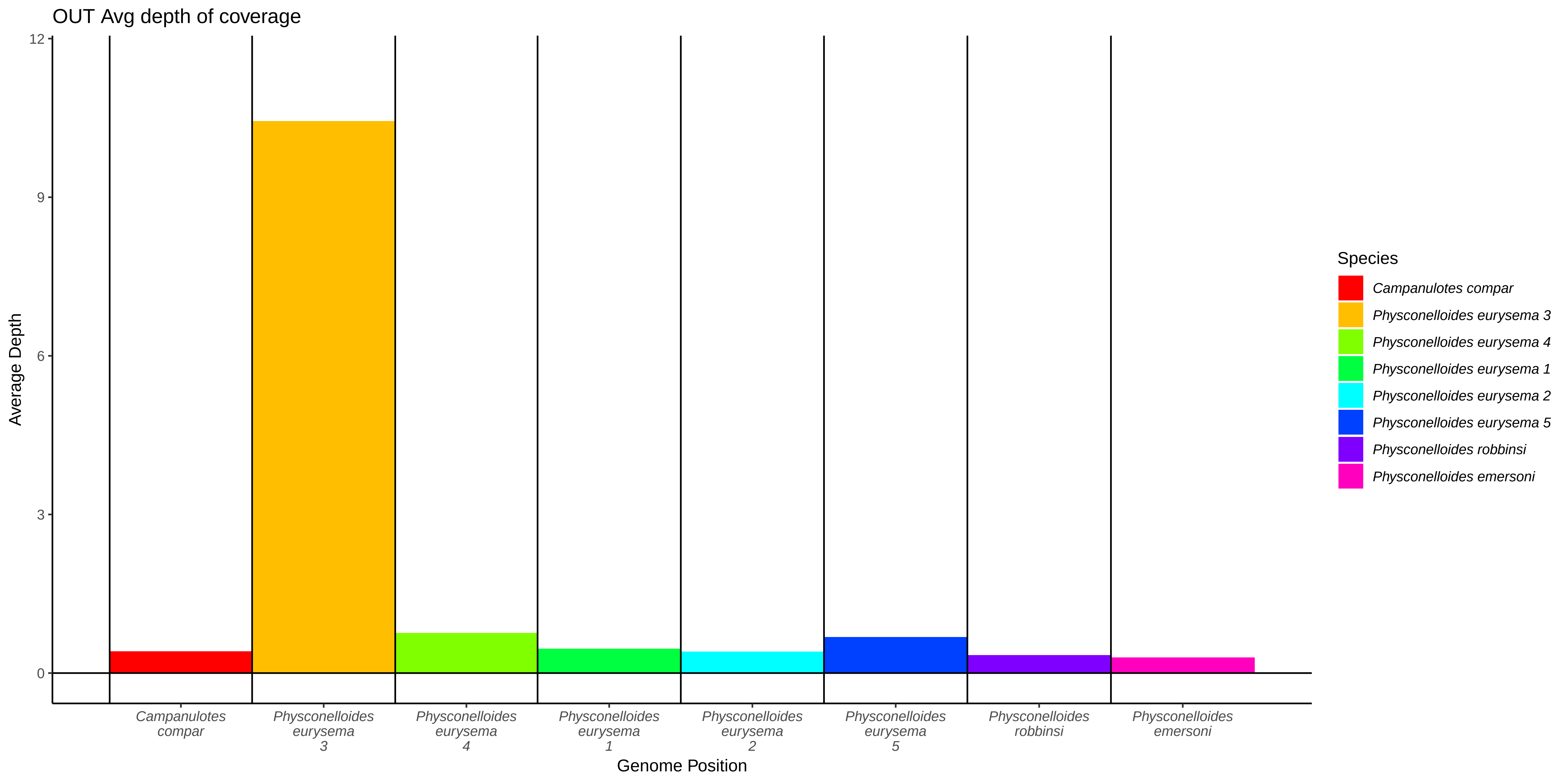

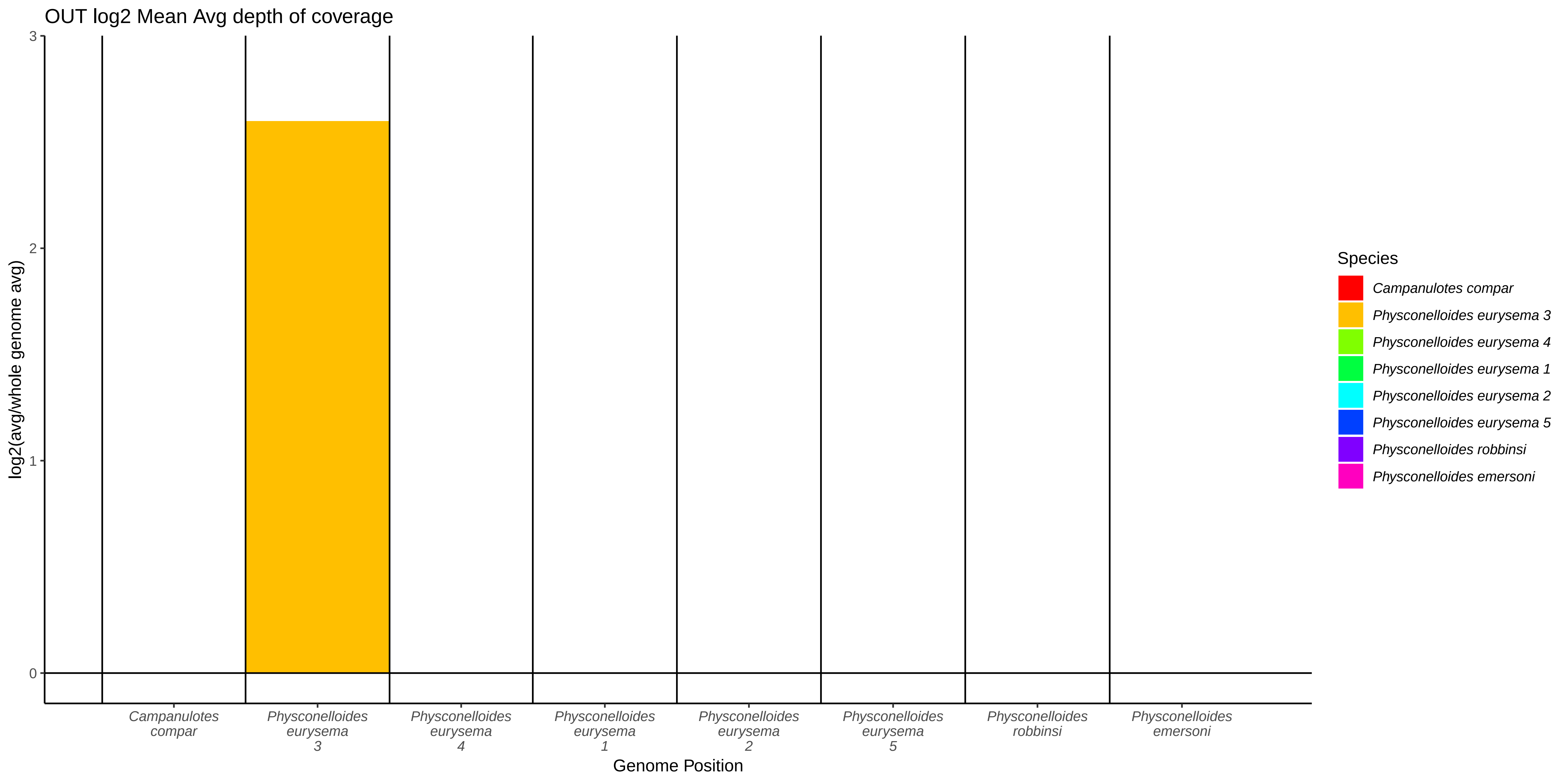

### OUT_speciesDepth_Pheur.Comin.6.15.2015.10.pdf

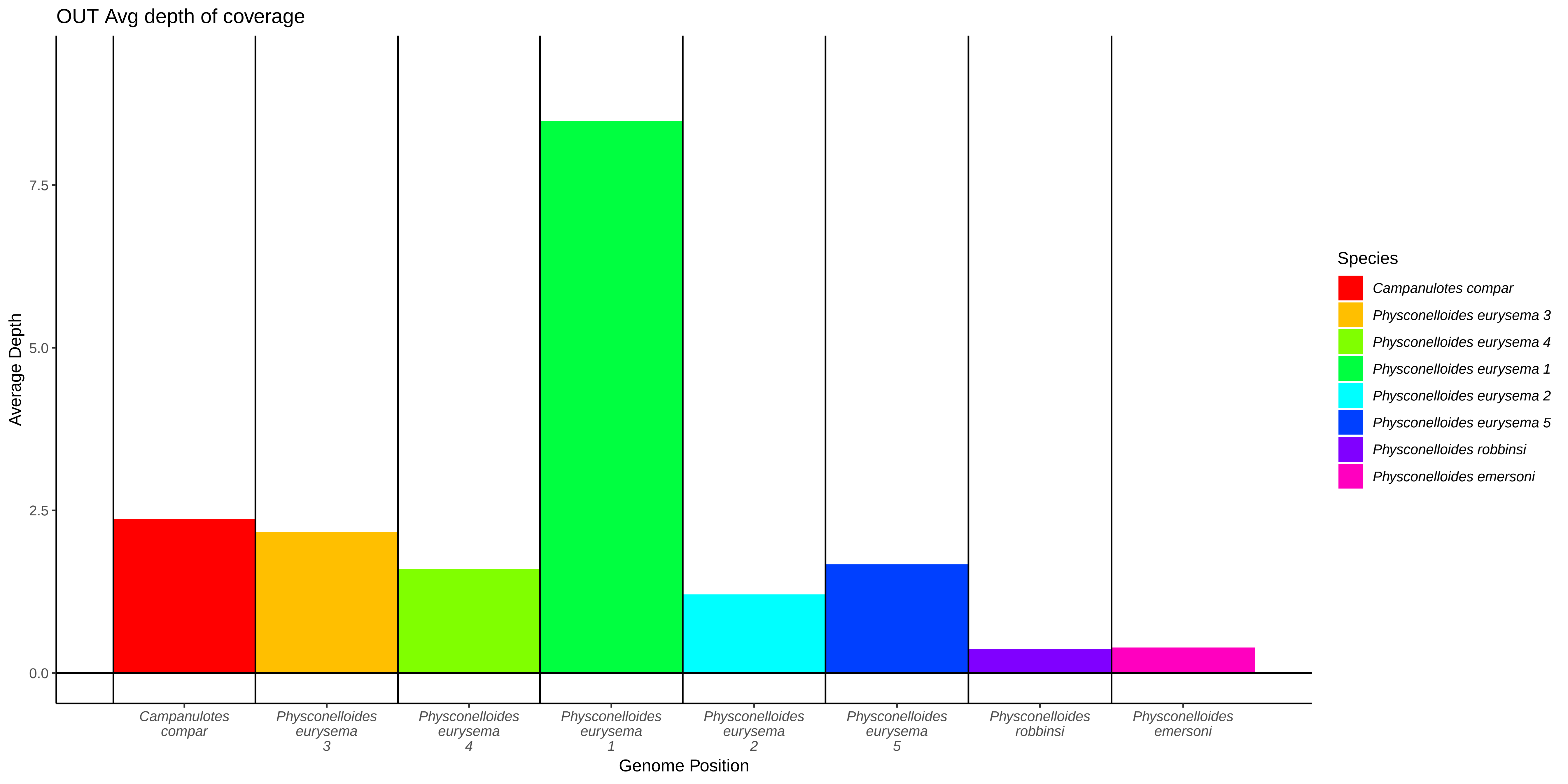

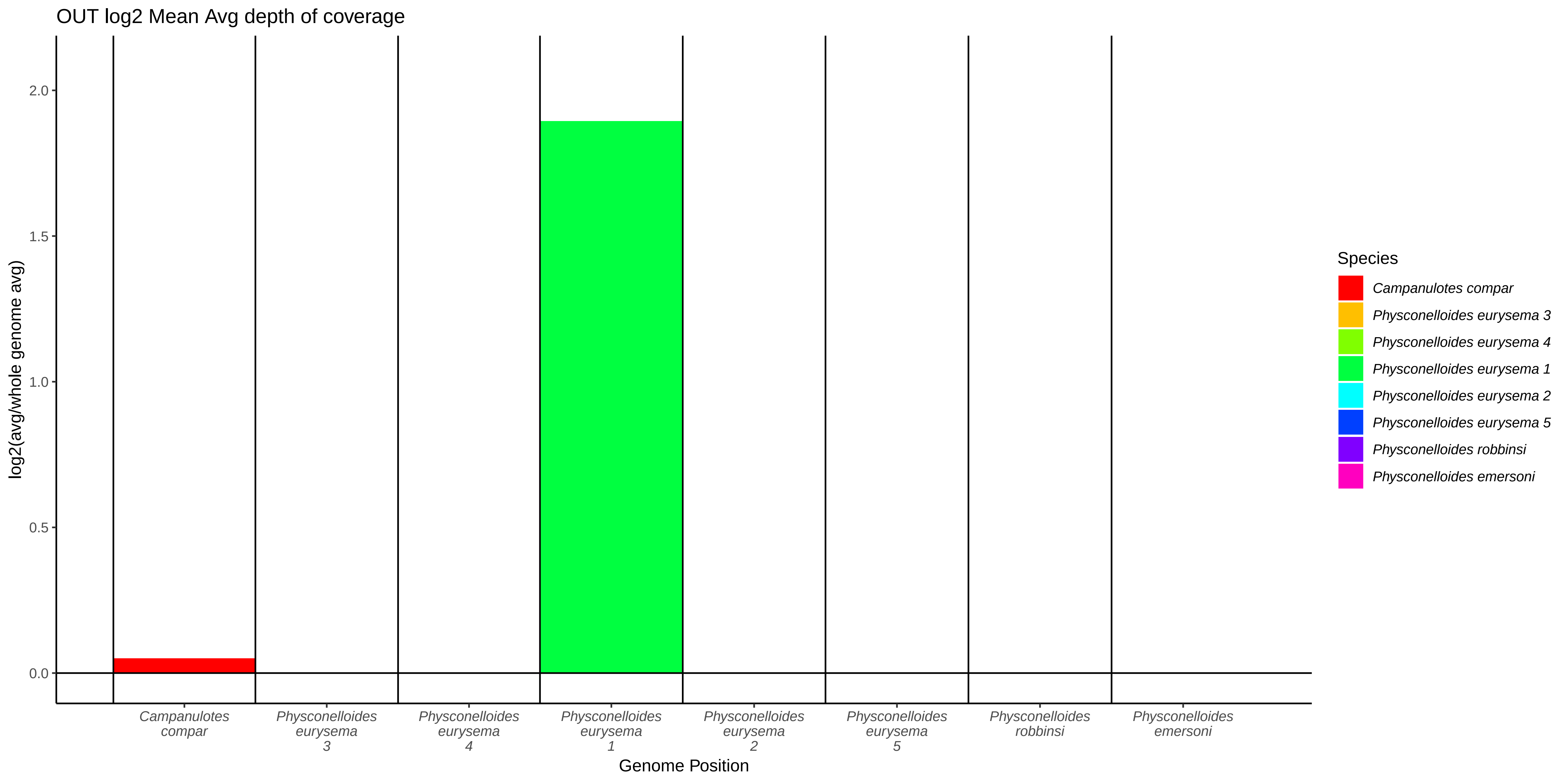

### OUT_speciesDepth_Pheur_Clpre_6_15_2015_8.pdf

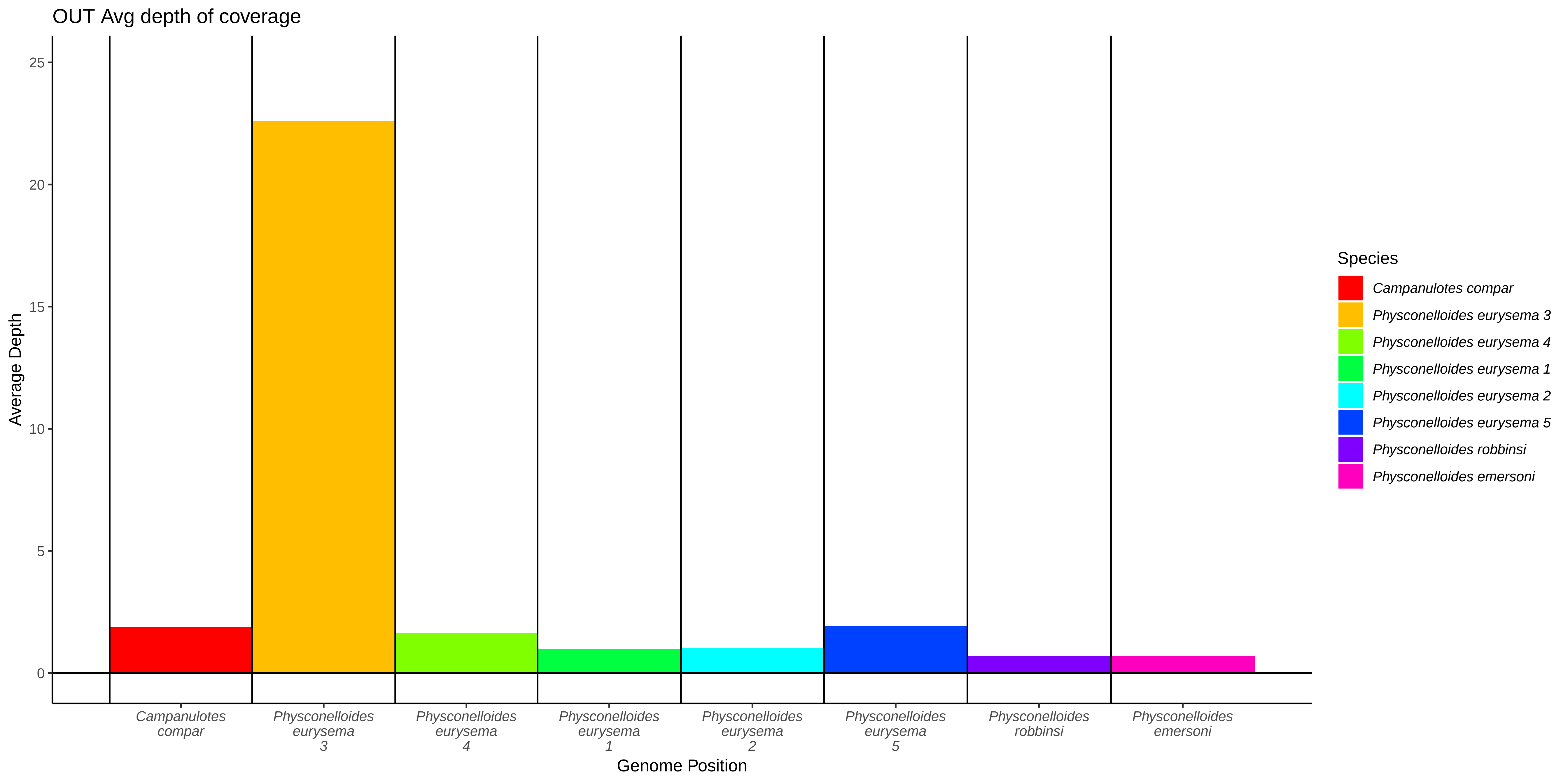

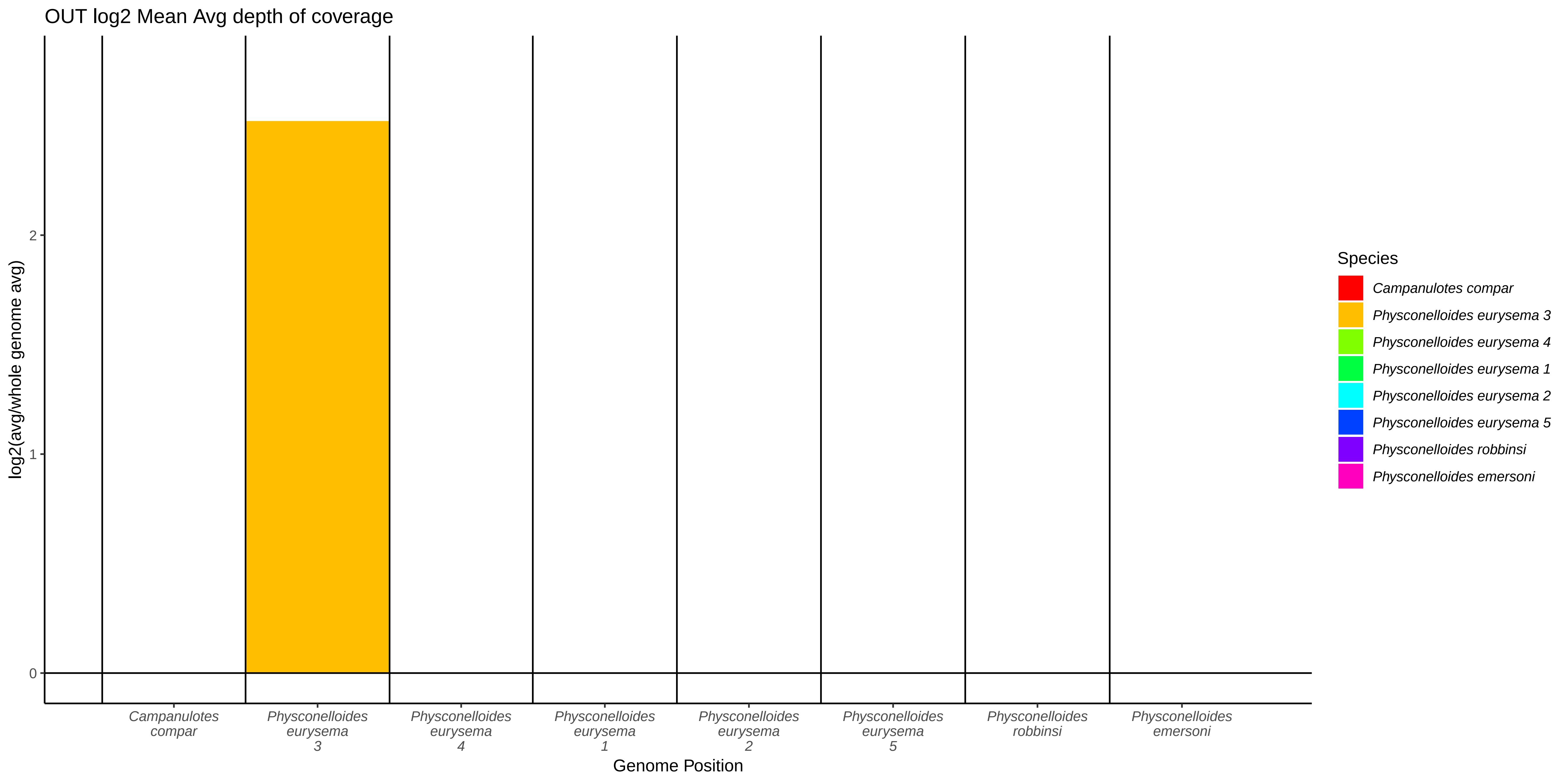

### OUT_speciesDepth_Pheur_Cobuc_6_15_2015_1.pdf

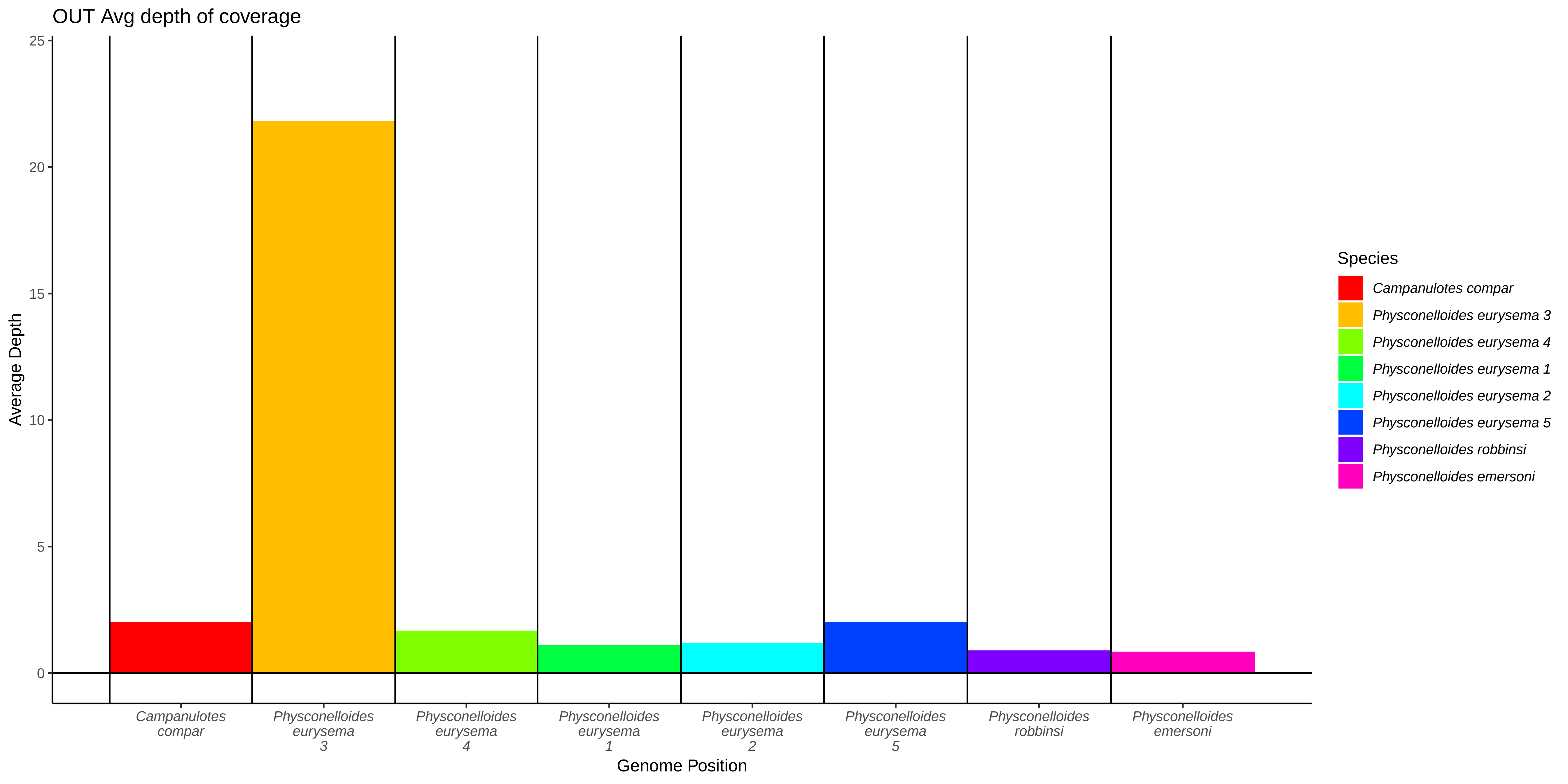

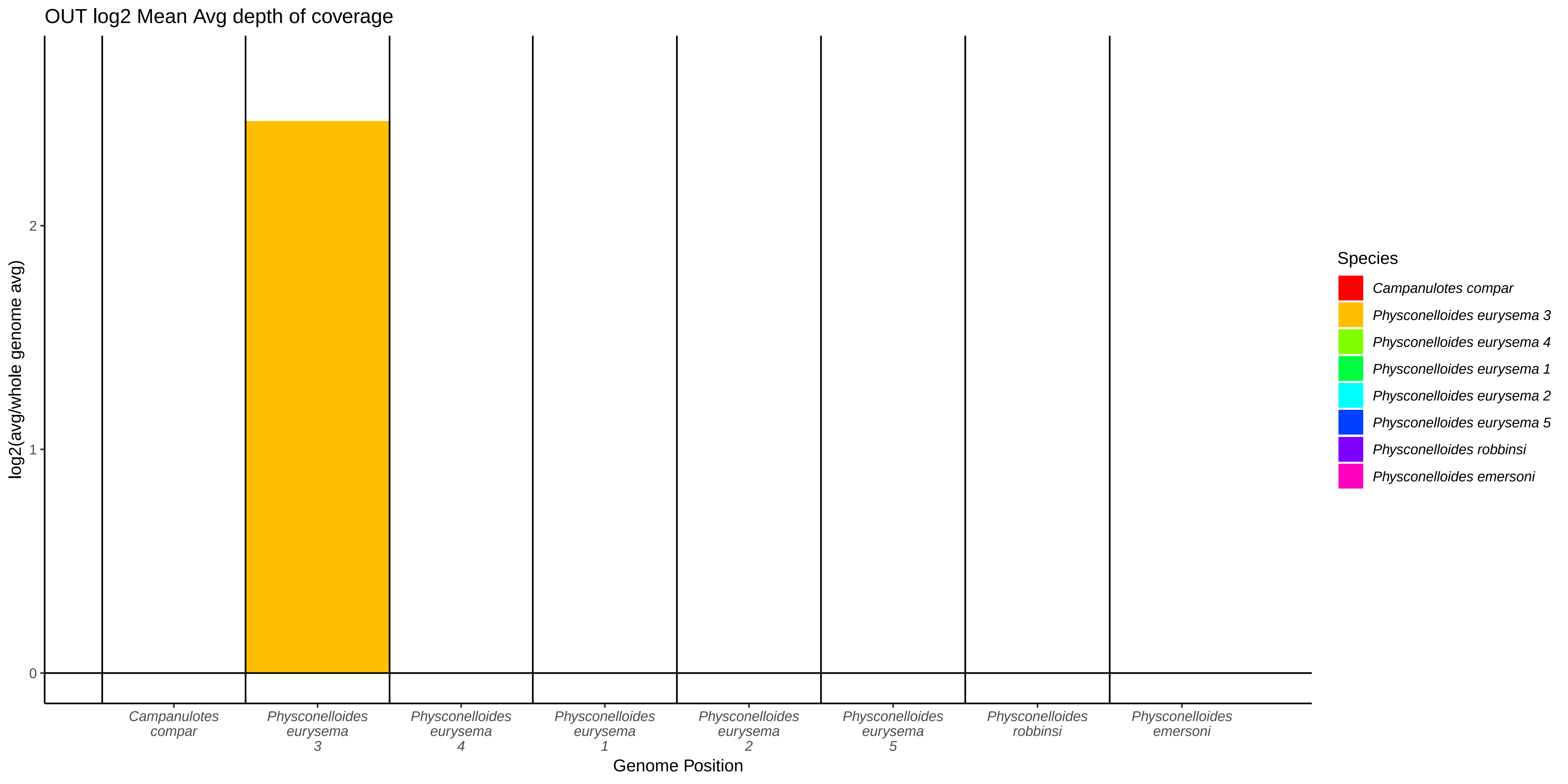

### OUT_speciesDepth_Pheur_Cocru_6_15_2015_2.pdf

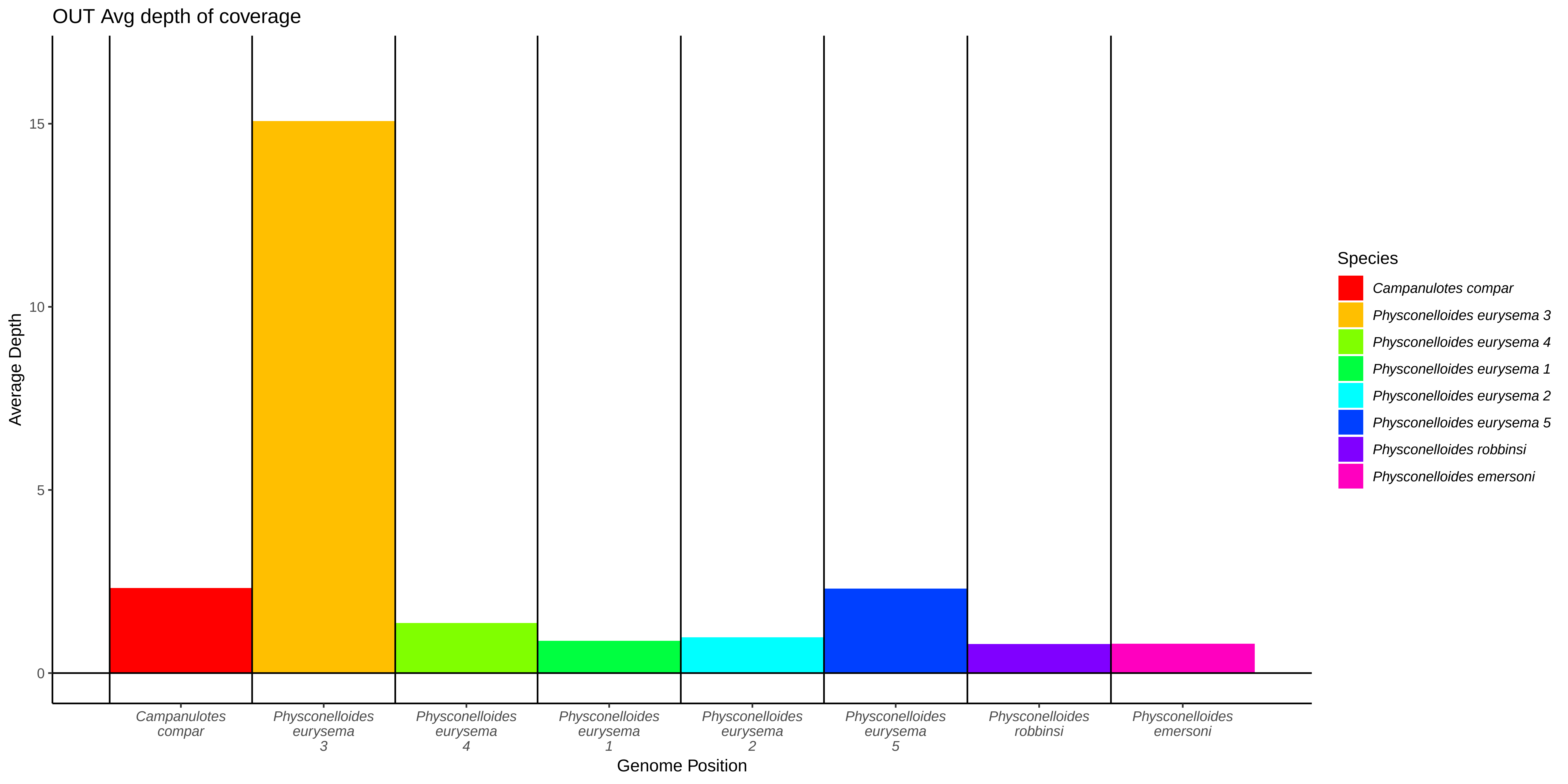

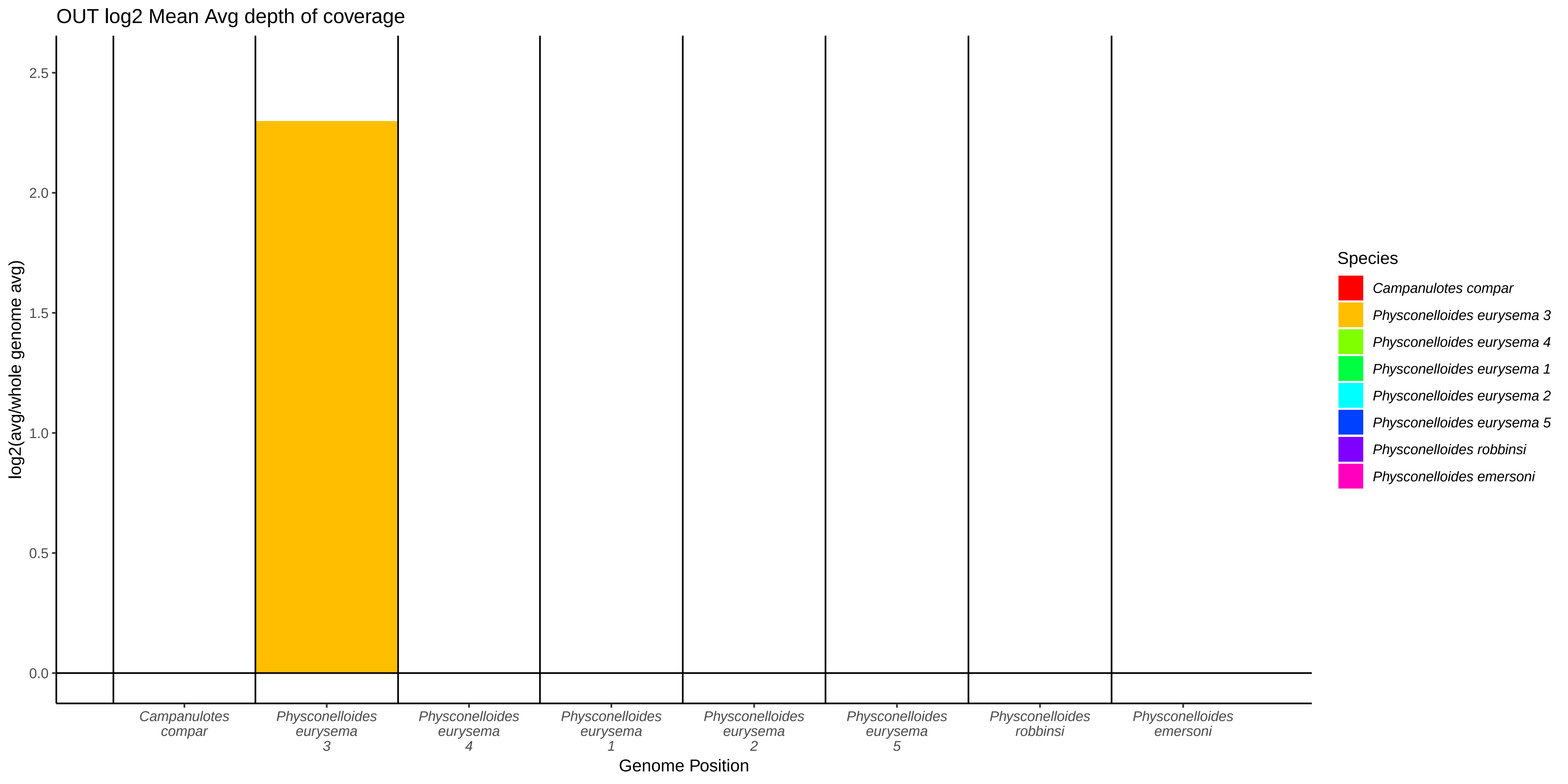

### OUT_speciesDepth_Pheur_Coinc_6_15_2015_7.pdf

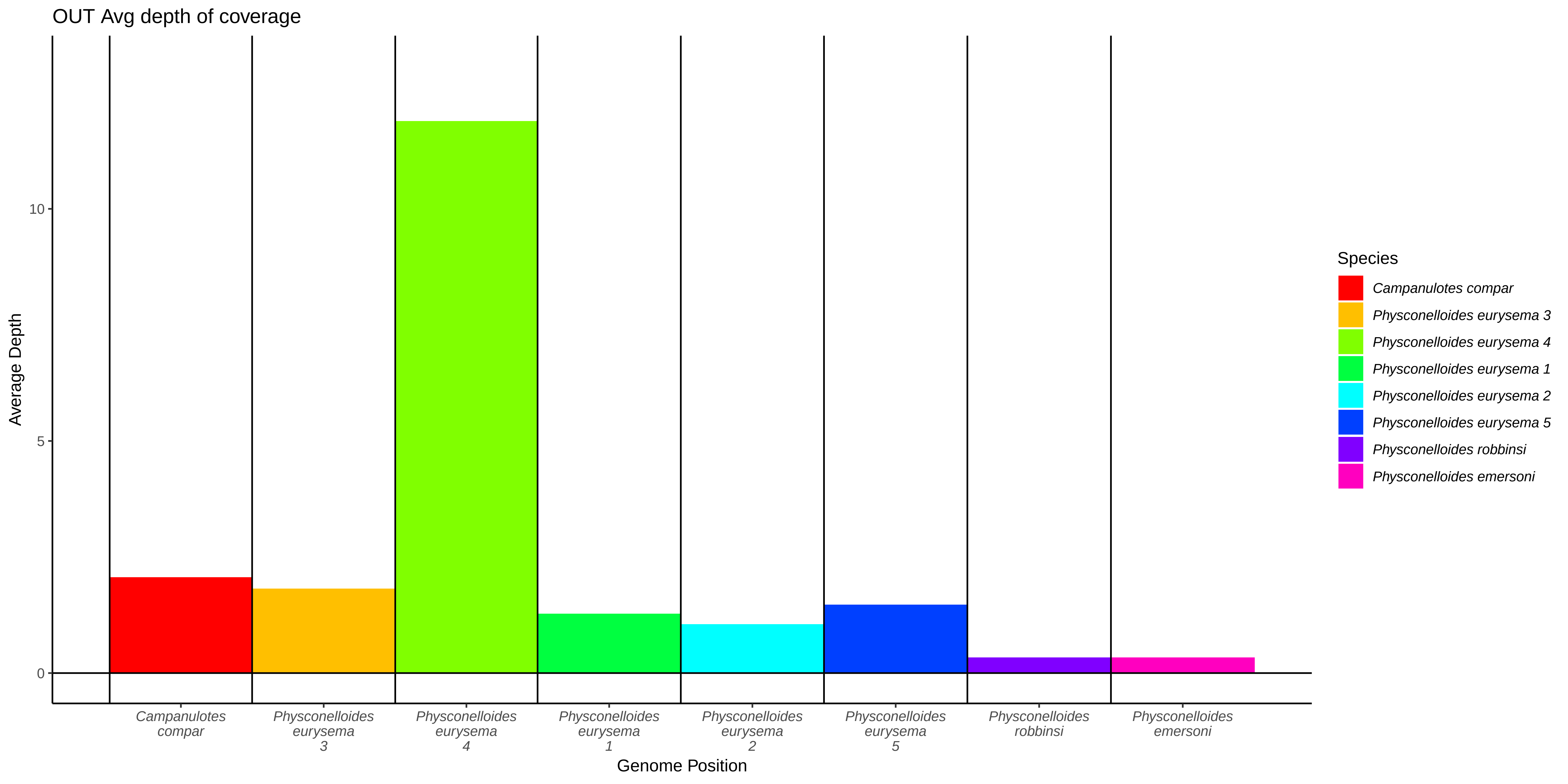

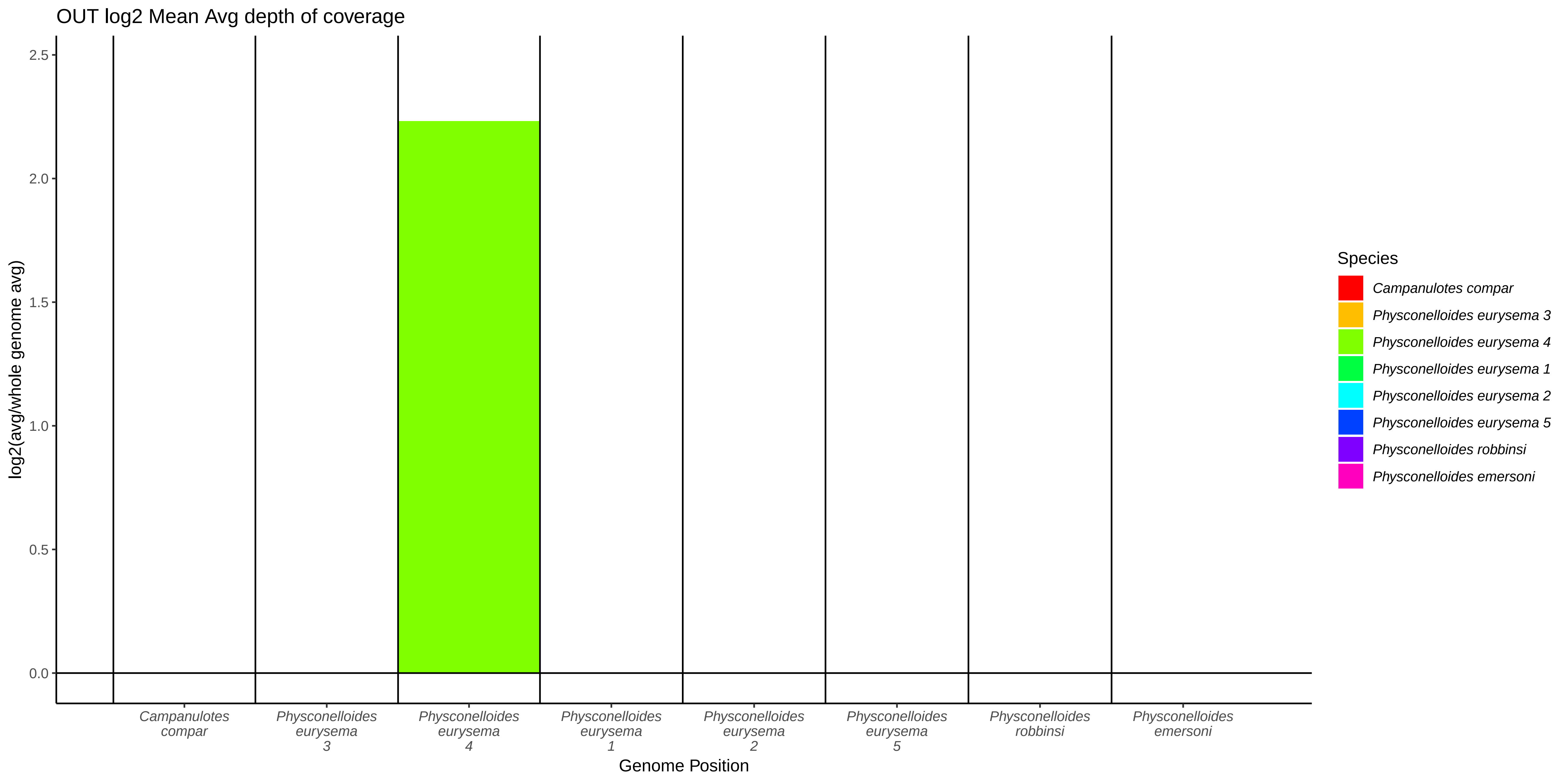

### OUT_speciesDepth_Pheur_Copas_3_14_2016_4.pdf

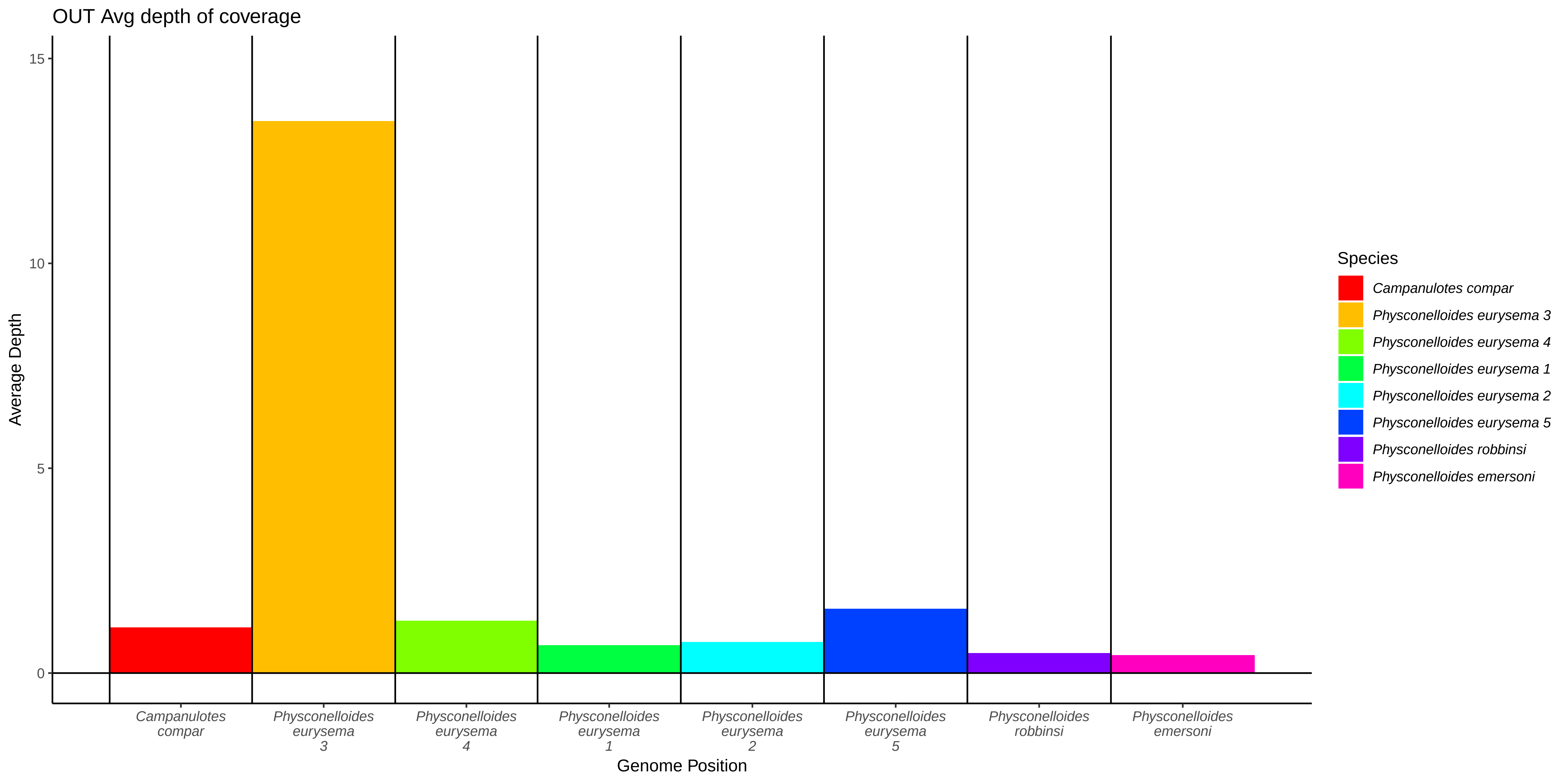

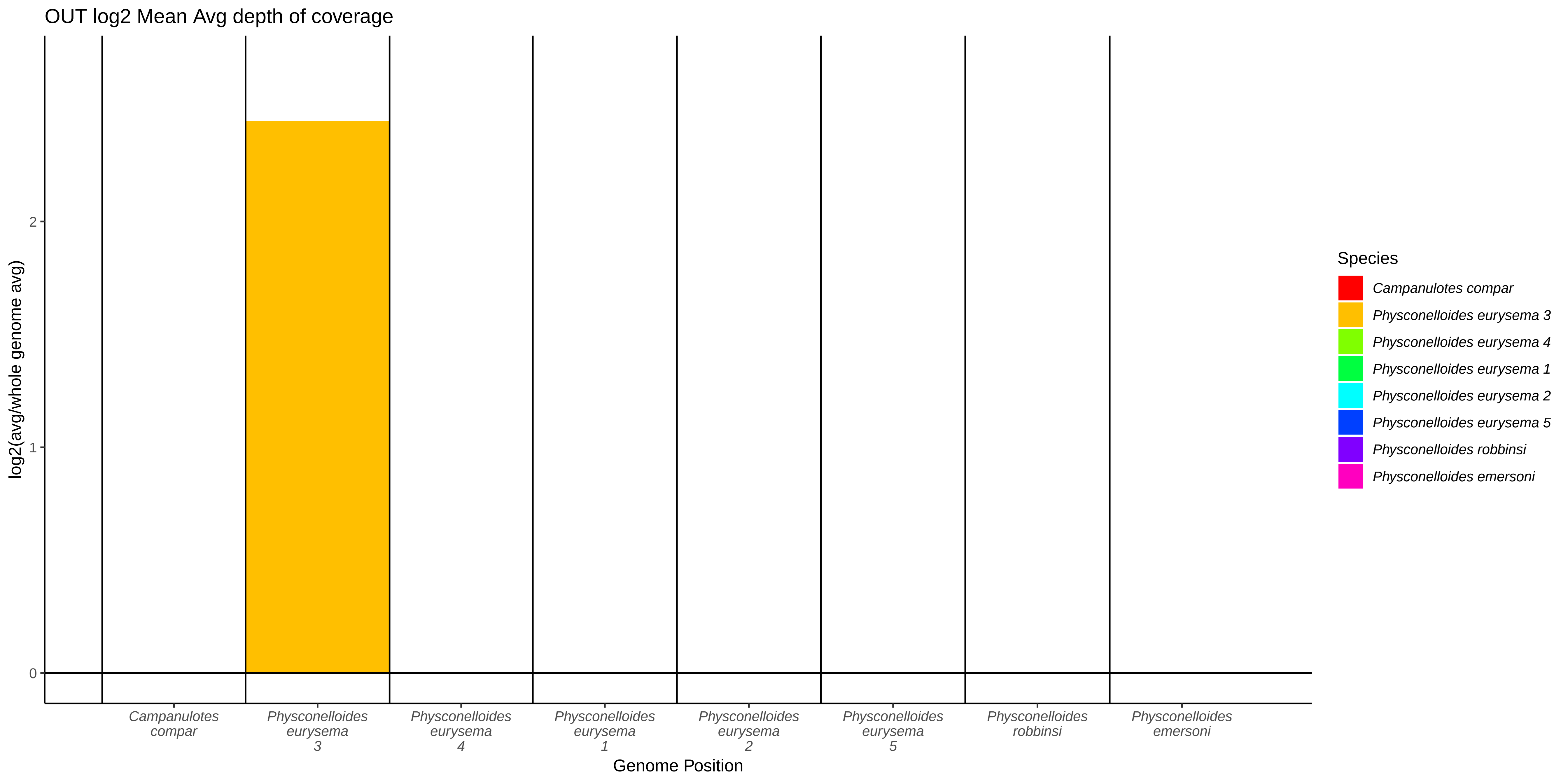

### OUT_speciesDepth_Pheur_Copas_3_14_2016_7.pdf

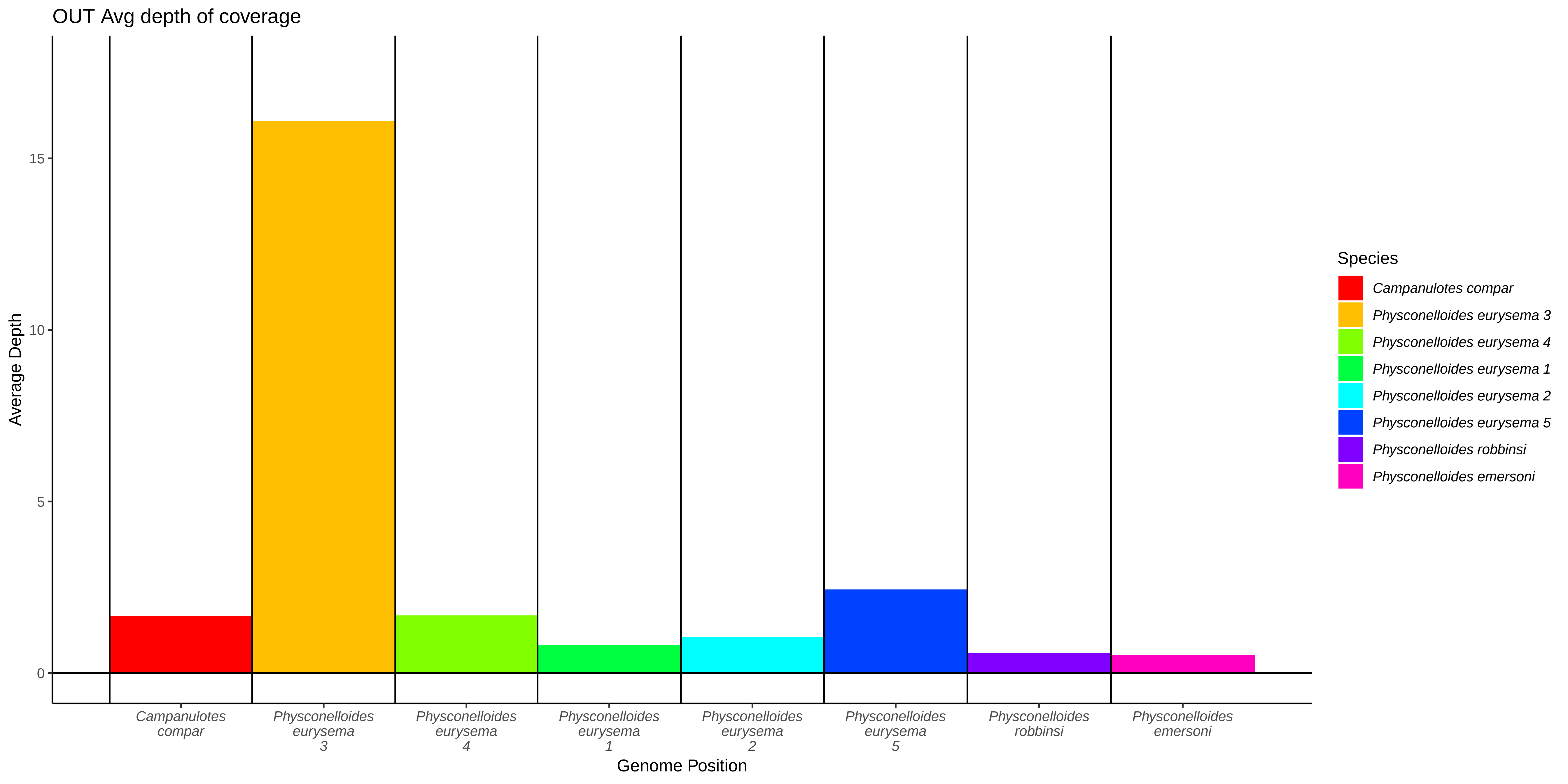

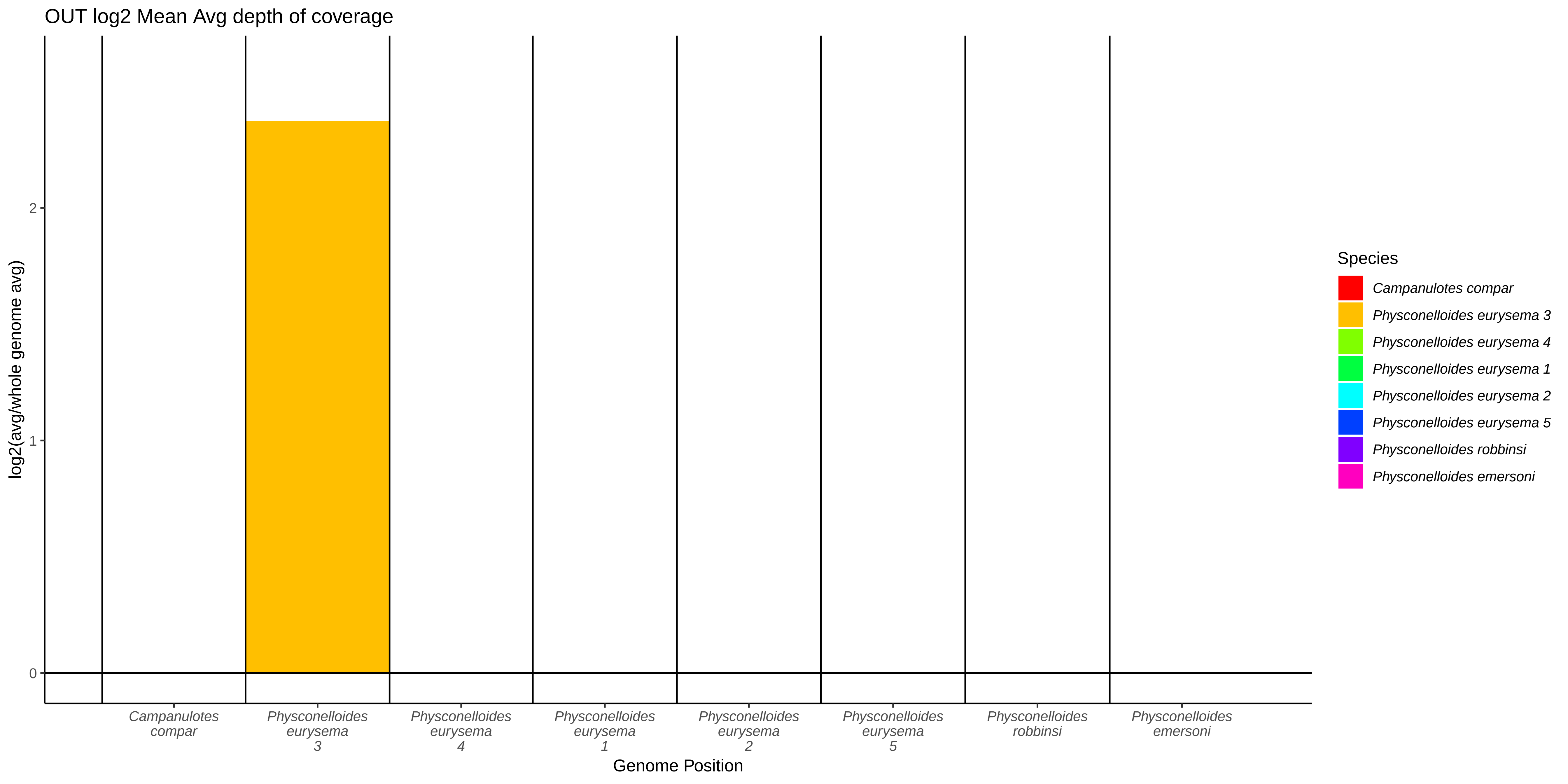

### OUT_speciesDepth_Pheur_Copas_3_14_2016_8.pdf

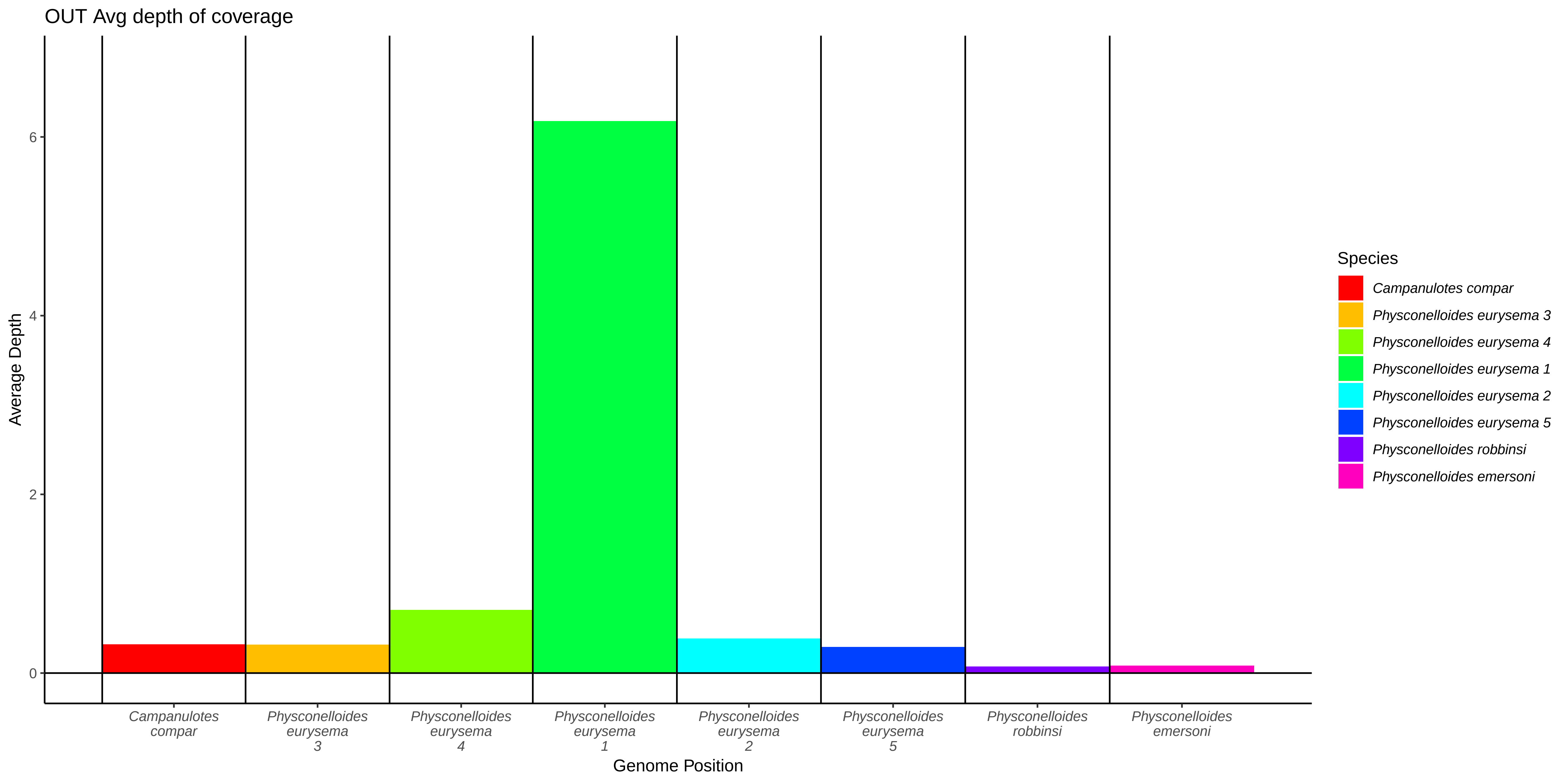

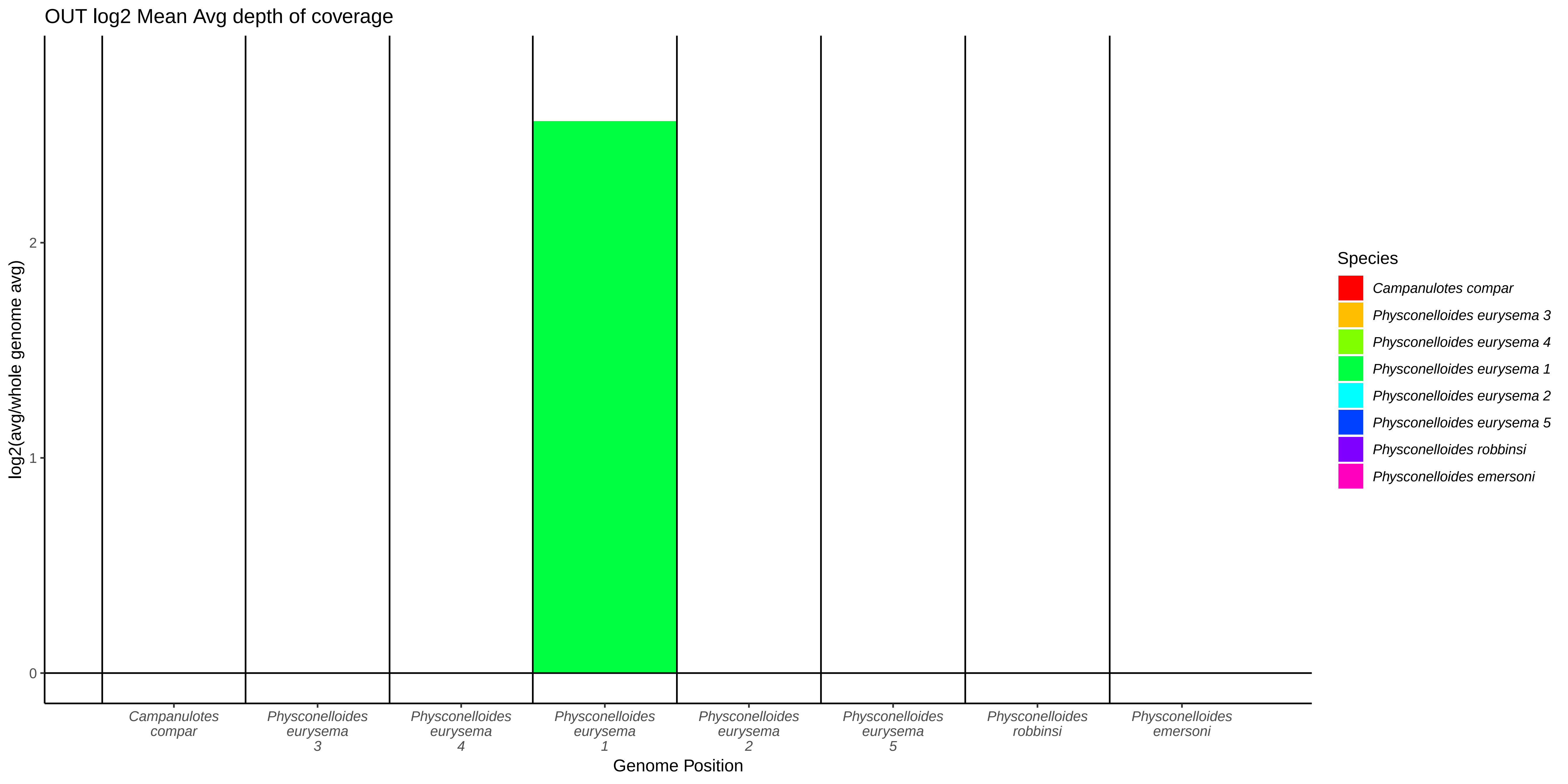

### OUT_speciesDepth_Pheur_Copas_3_14_2016_10.pdf

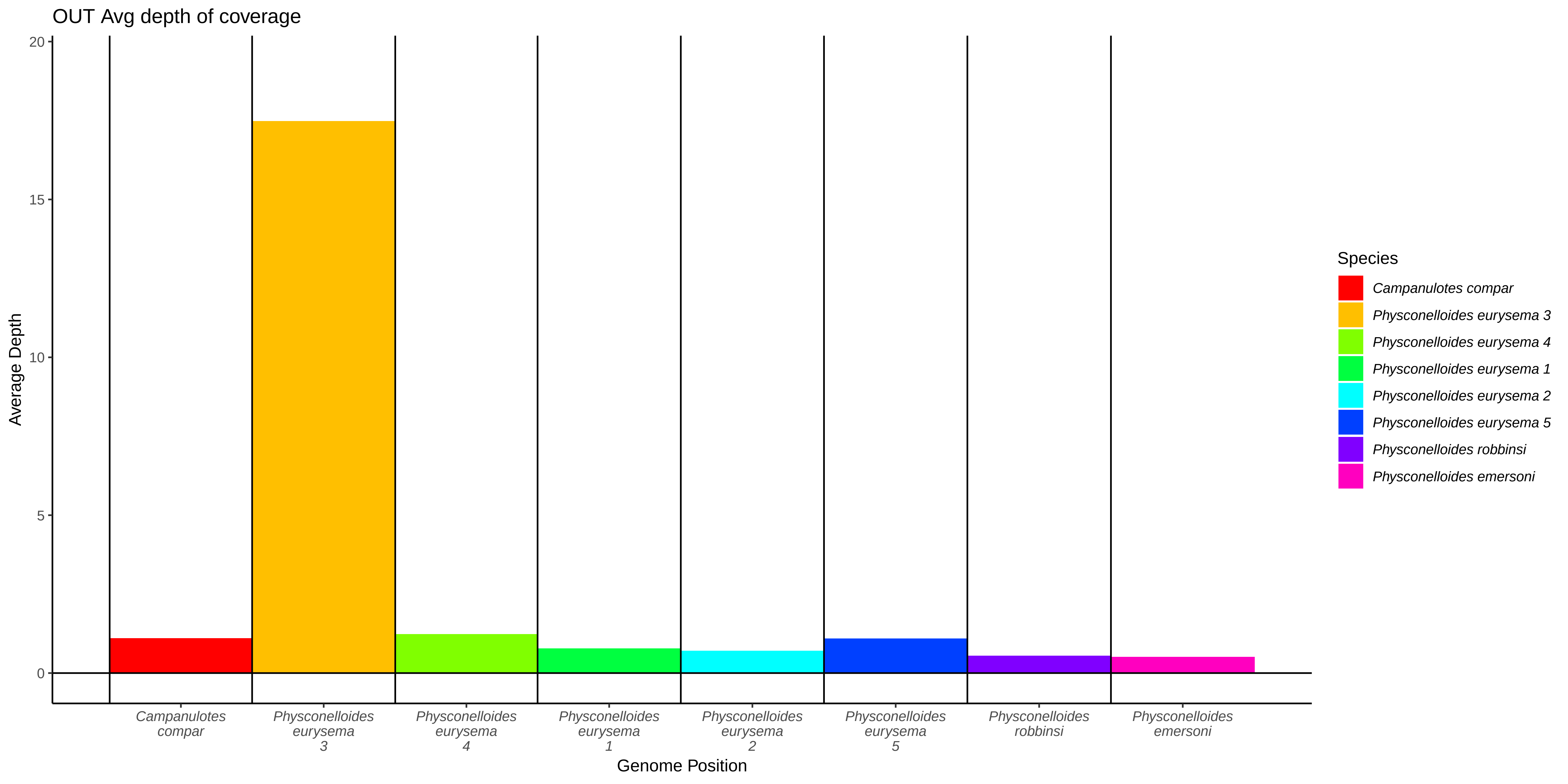

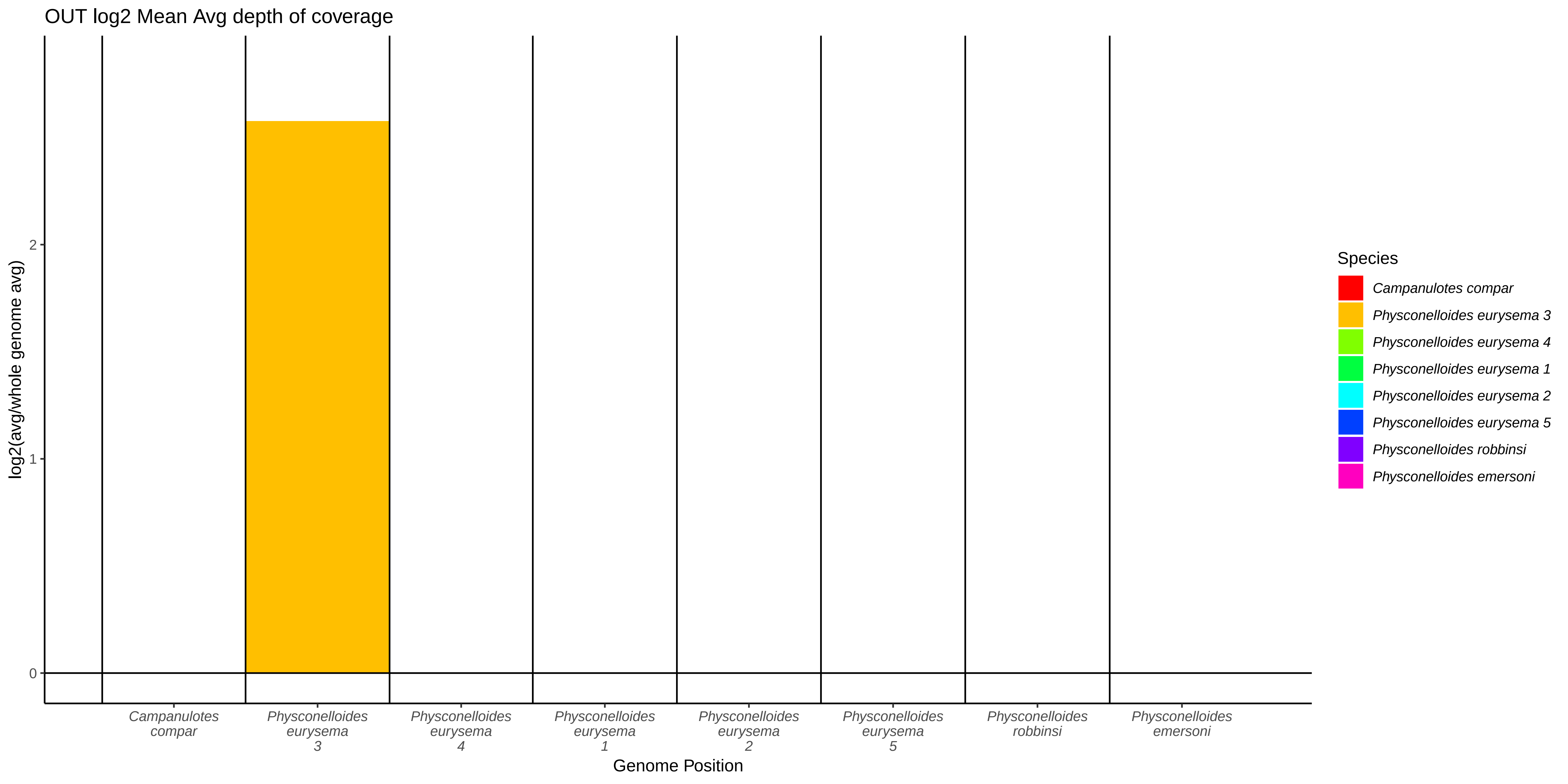

### OUT_speciesDepth_Pheur_Copas_3_16_2016_1.pdf

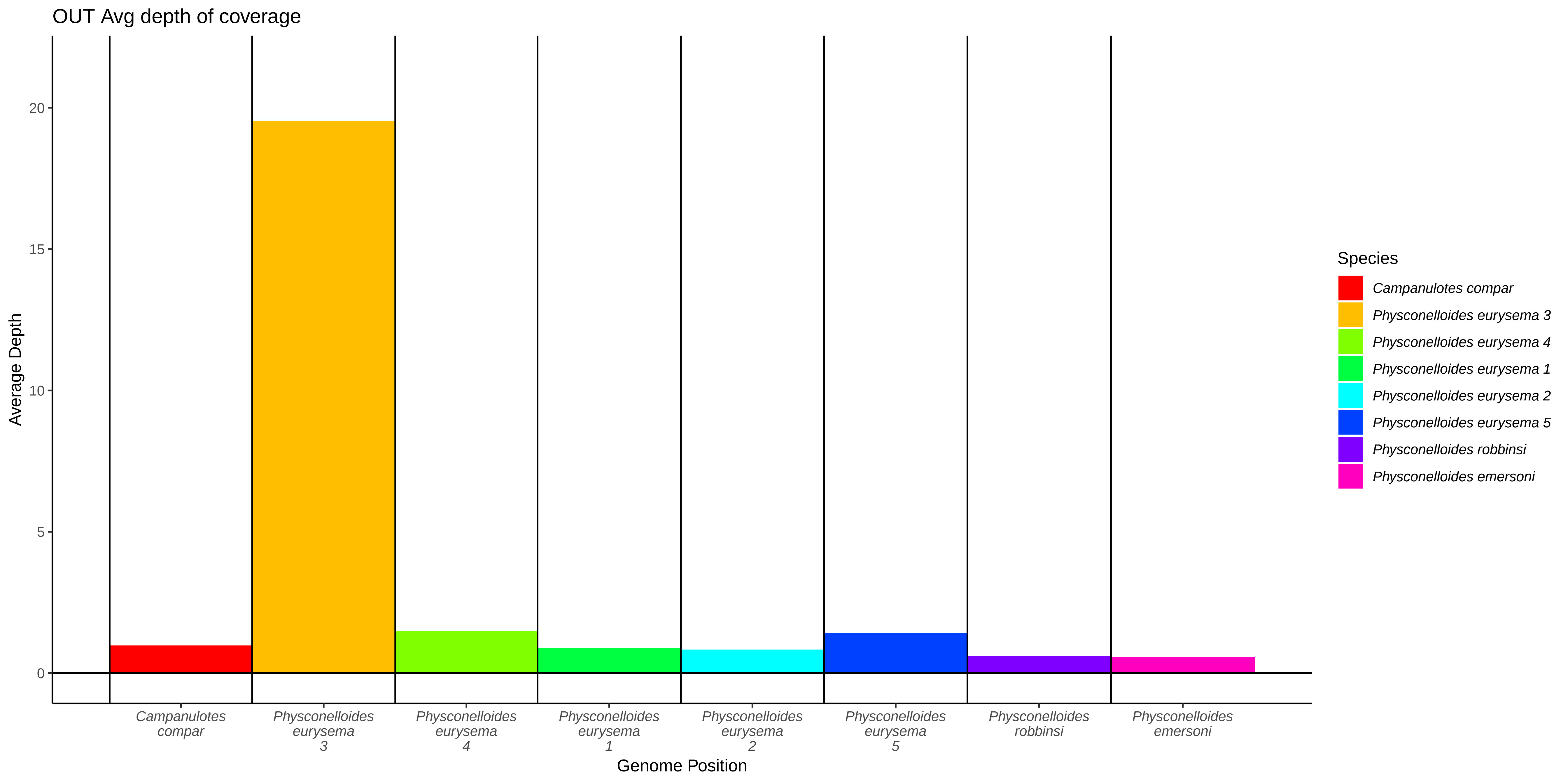

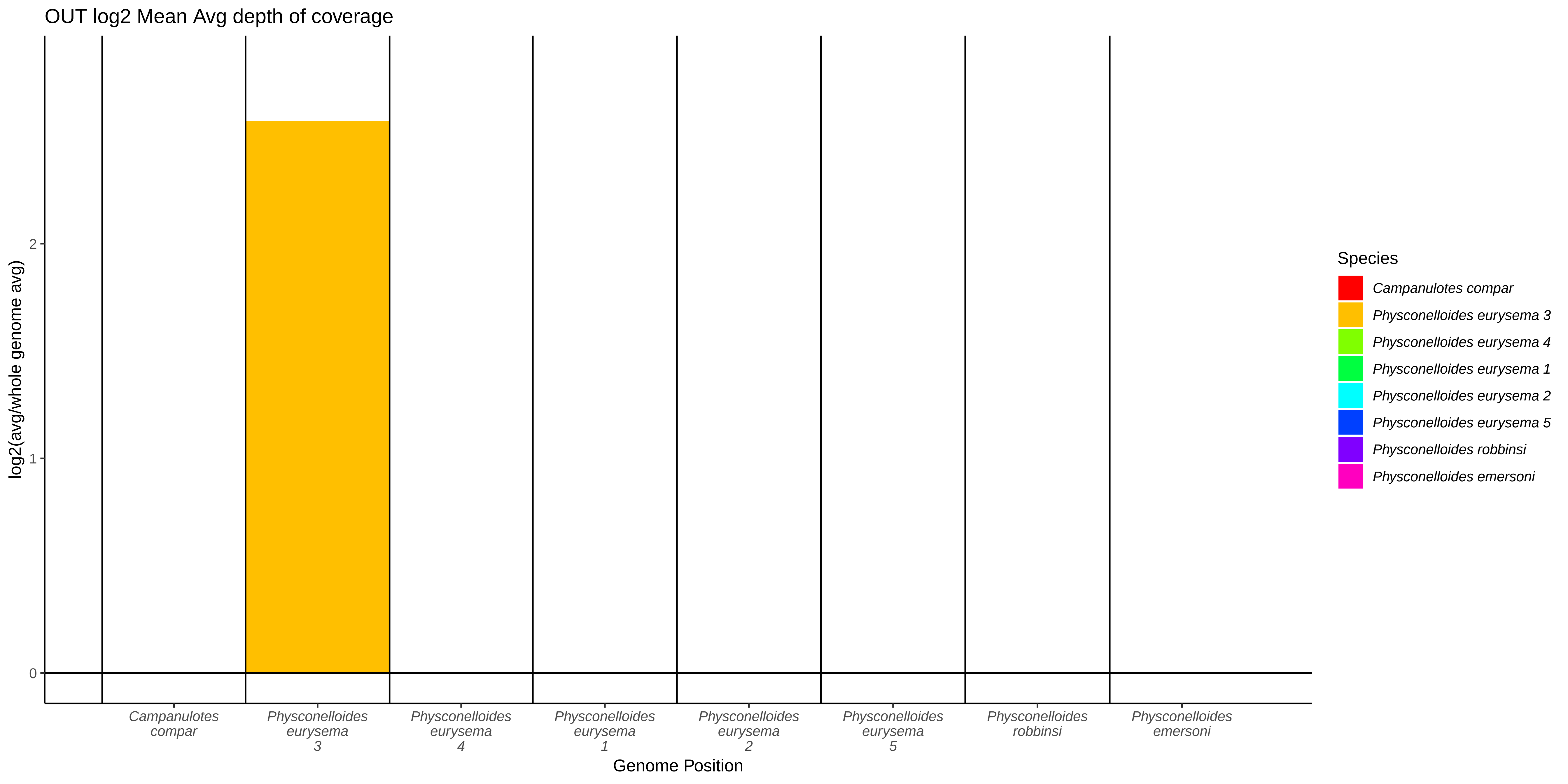

### OUT_speciesDepth_Pheur_Copas_3_16_2016_6.pdf

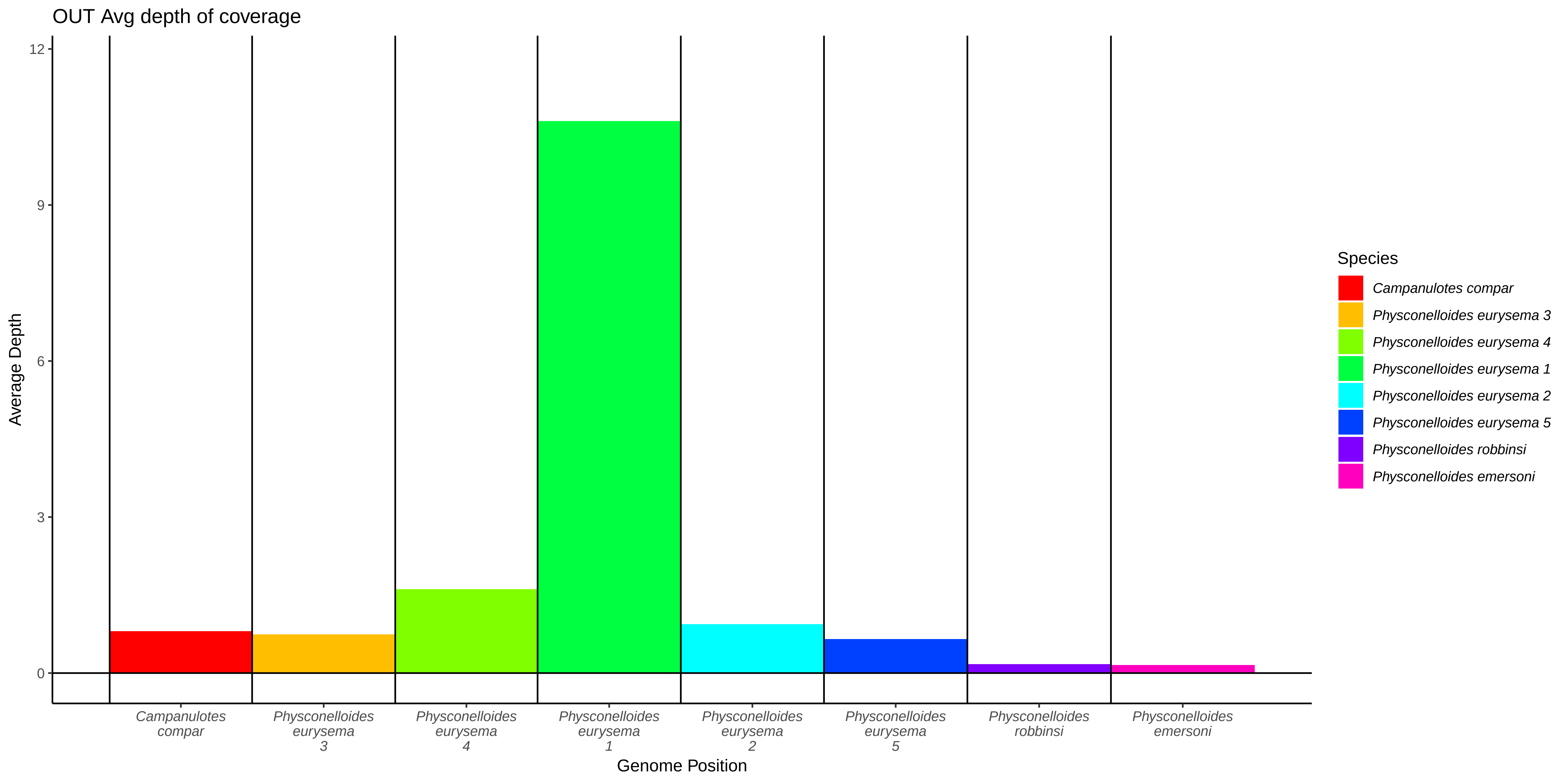

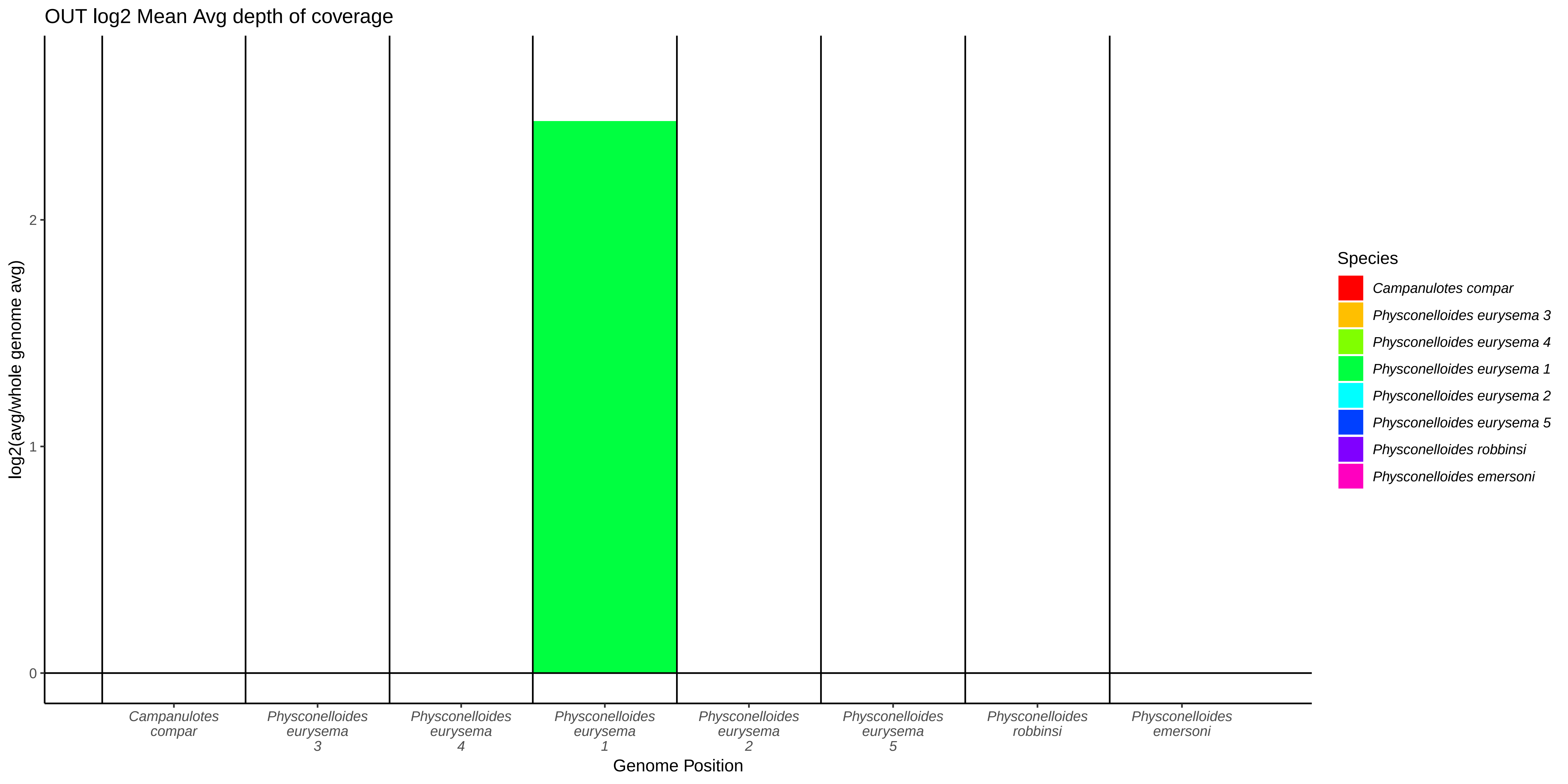

### OUT_speciesDepth_Pheur_Copas_3_16_2016_7.pdf

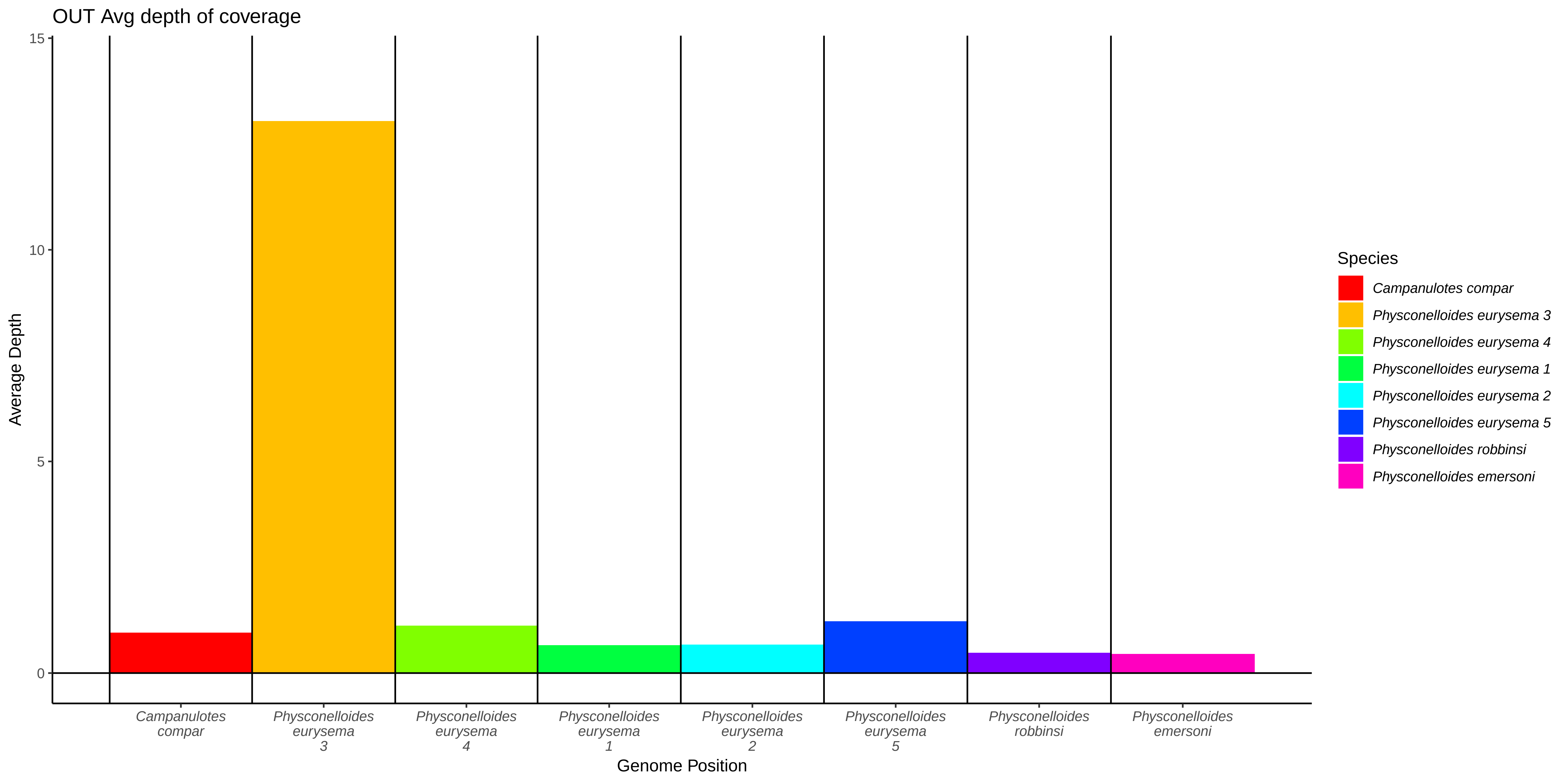

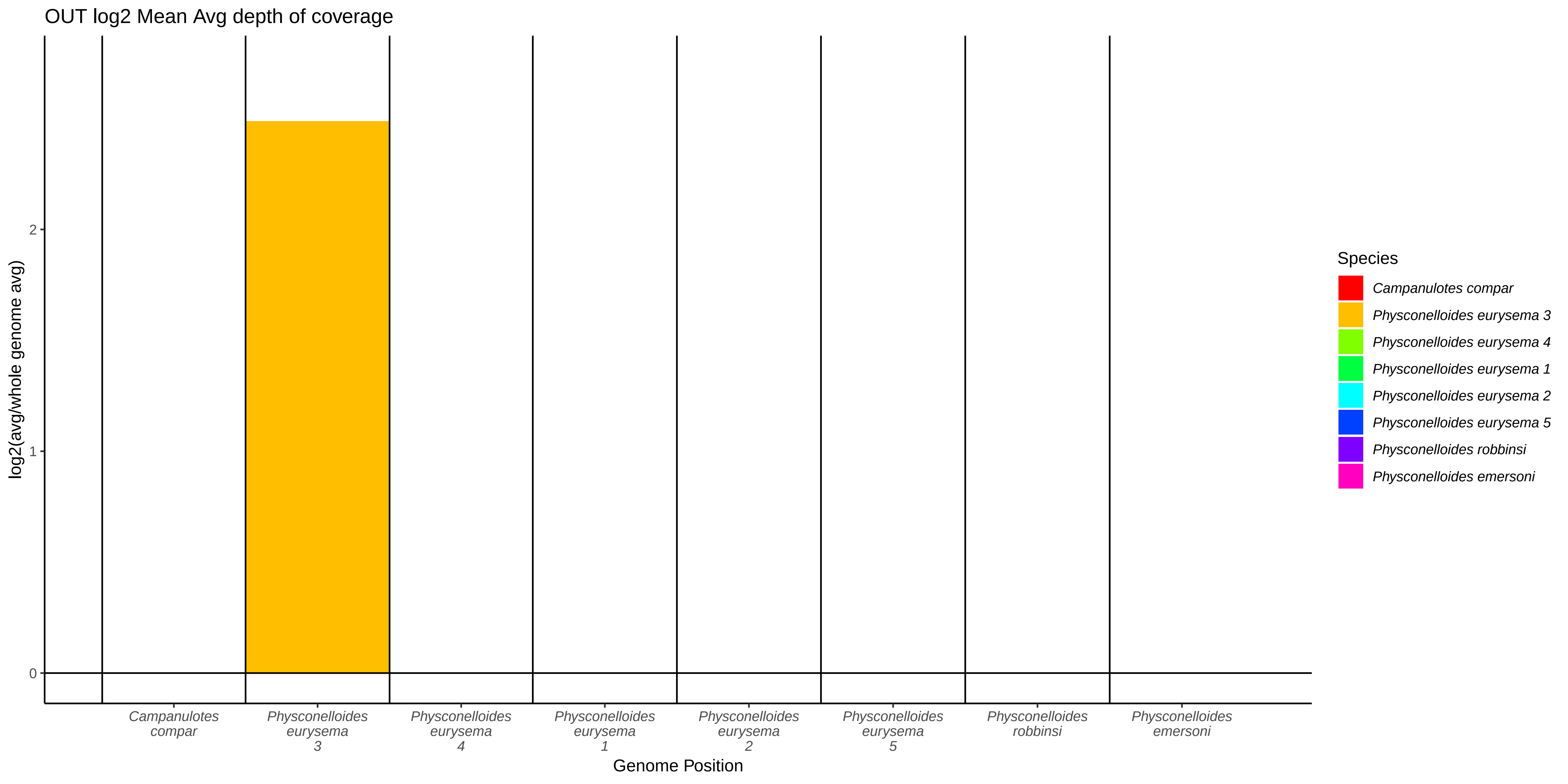

### OUT_speciesDepth_Pheur_Cosqu_3_14_2016_1.pdf

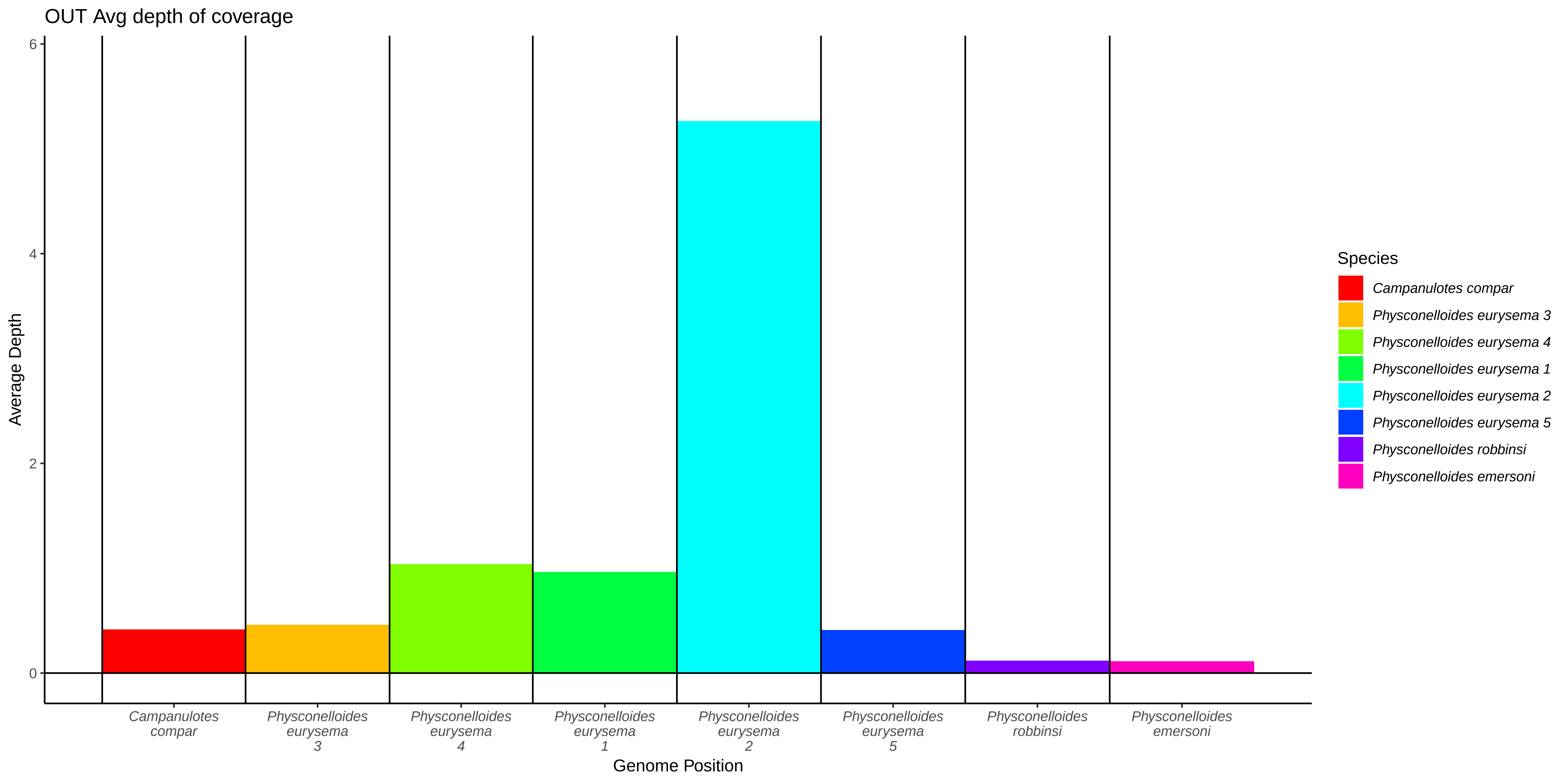

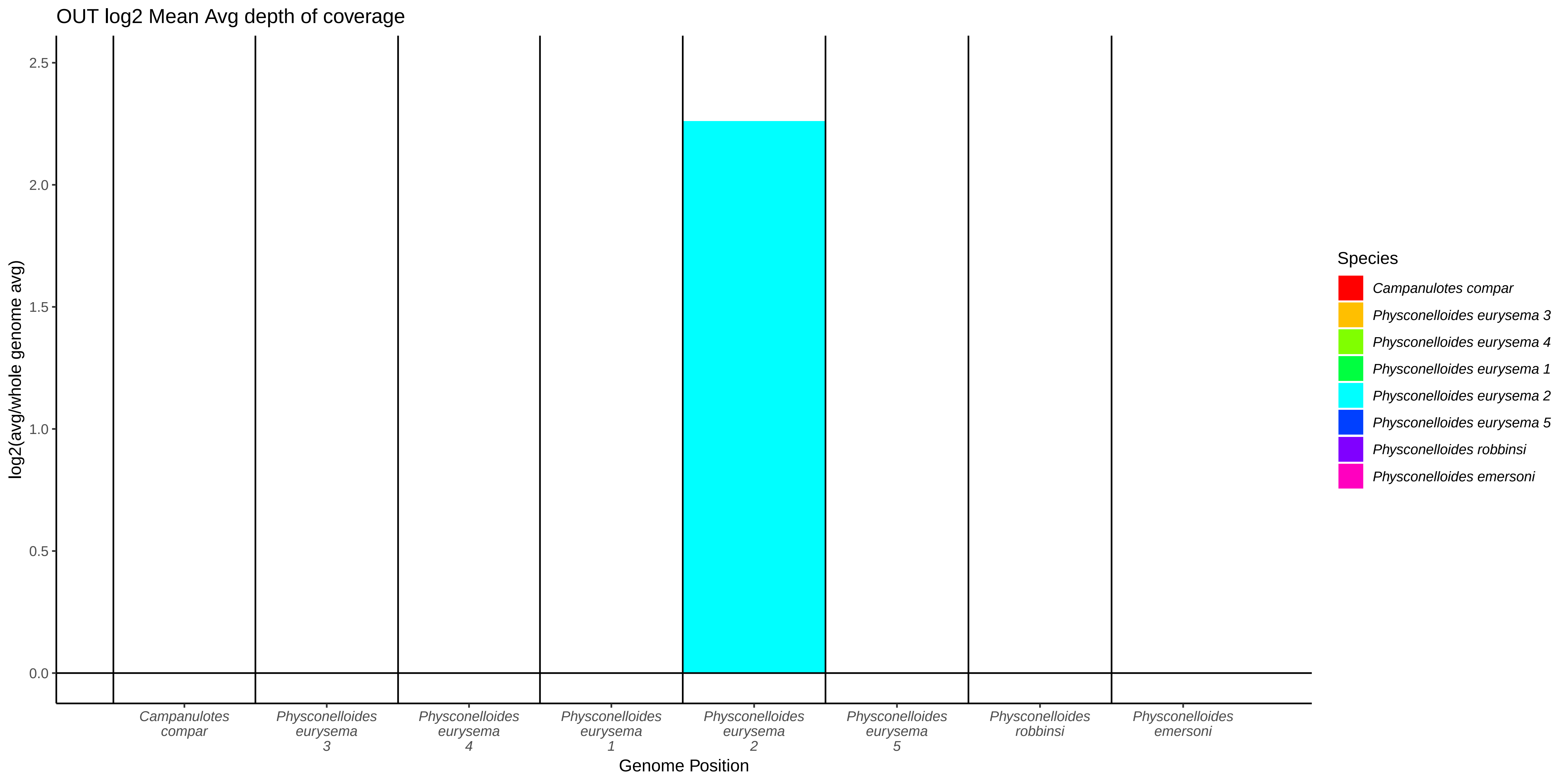

### OUT_speciesDepth_Pheur_Cosqu_3_14_2016_12.pdf

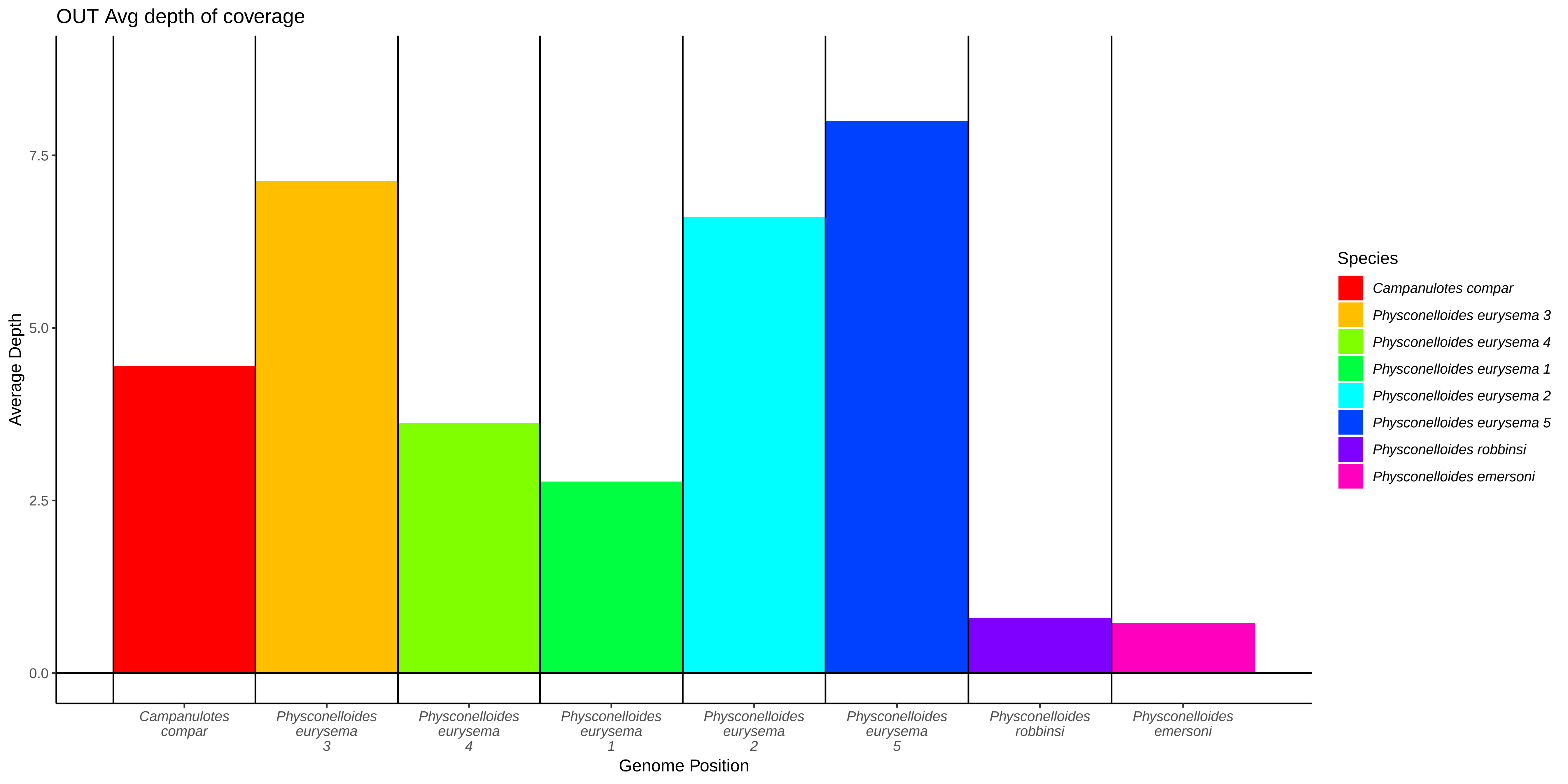

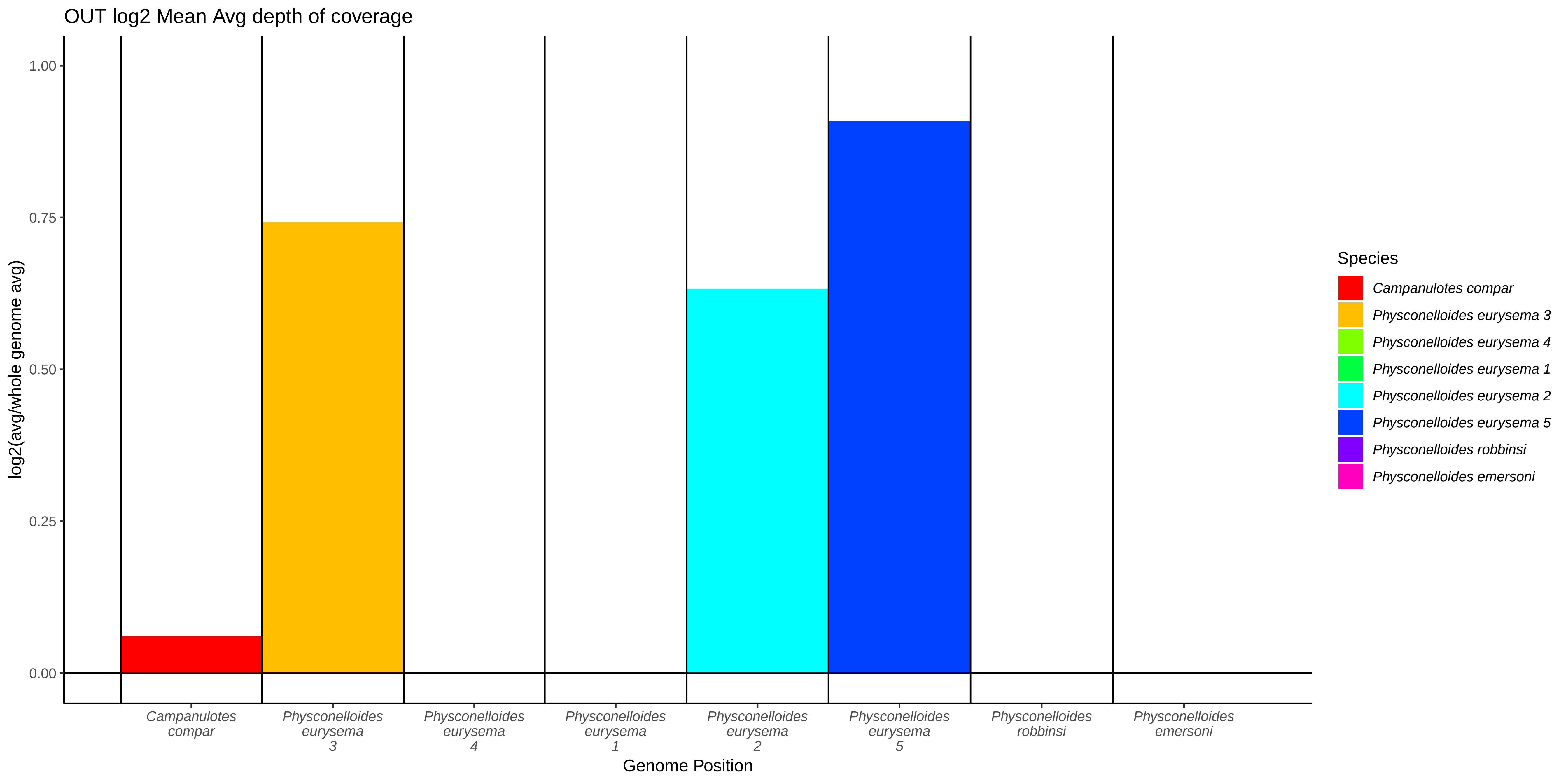

### OUT_speciesDepth_Pheur_Cotal_3_14_2016_2.pdf

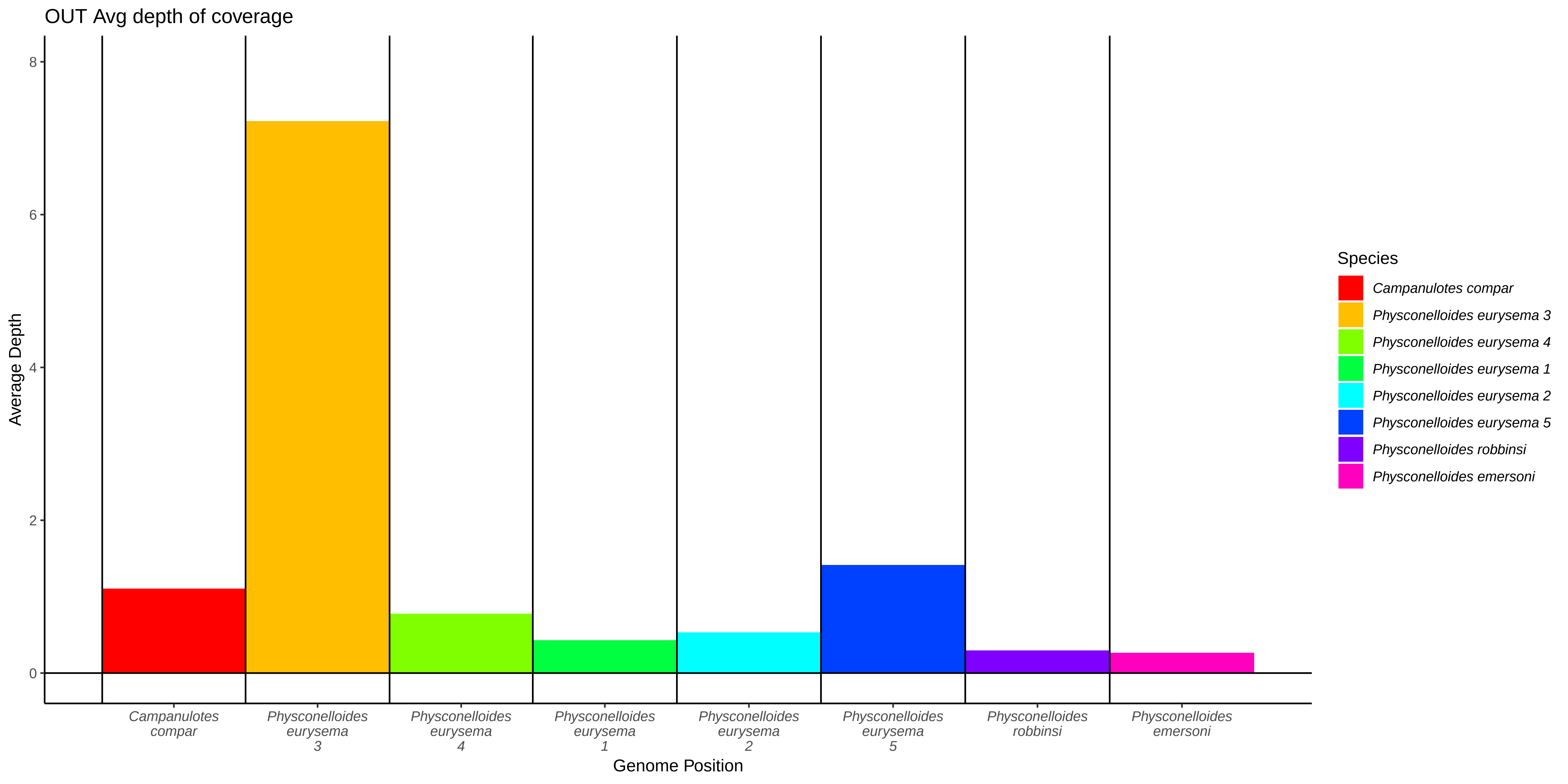

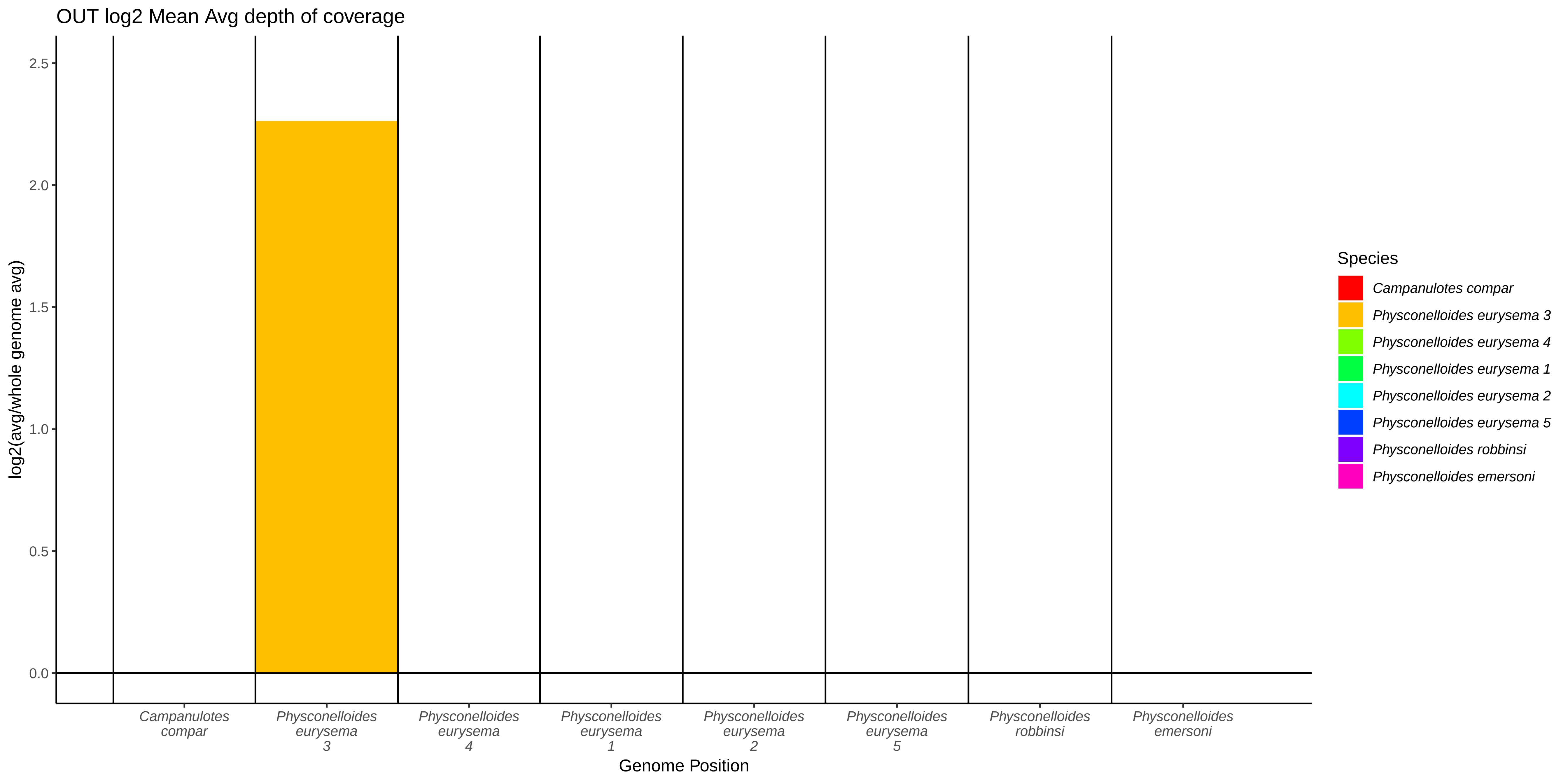

### OUT_speciesDepth_Pheur_Cotal_3_14_2016_6.pdf

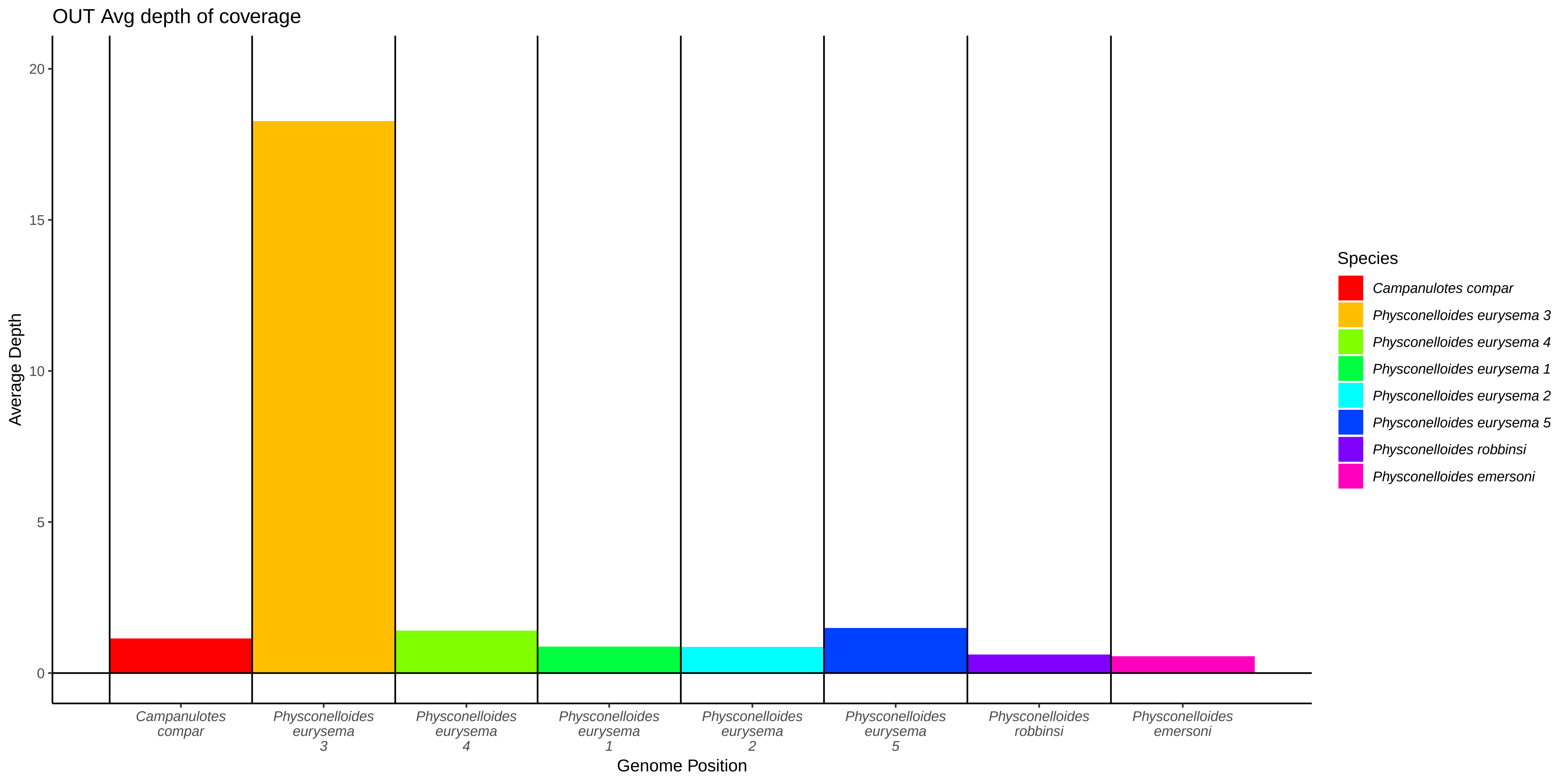

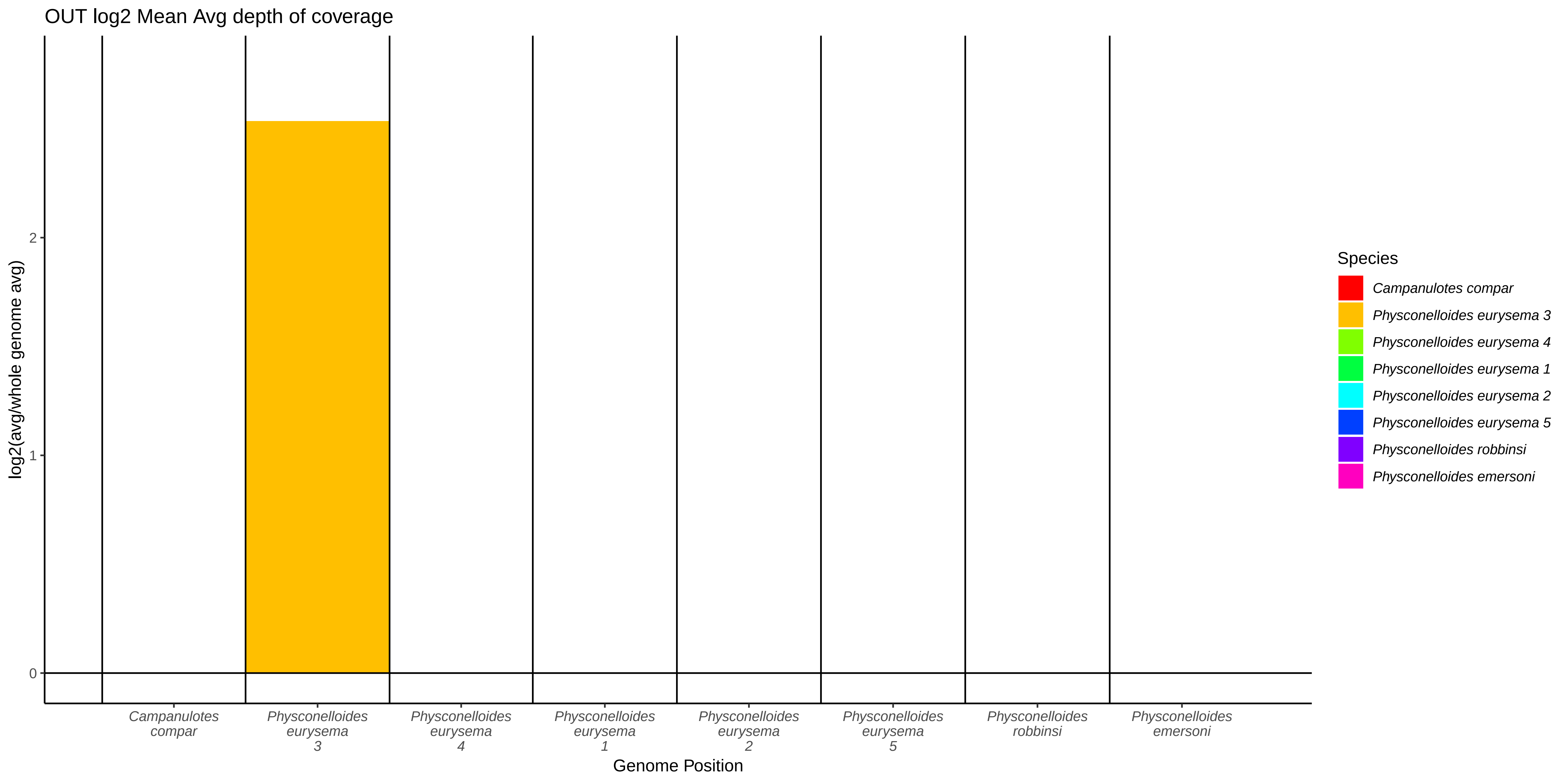

### OUT_speciesDepth_Pheur_Cotal_3_16_2016_3.pdf

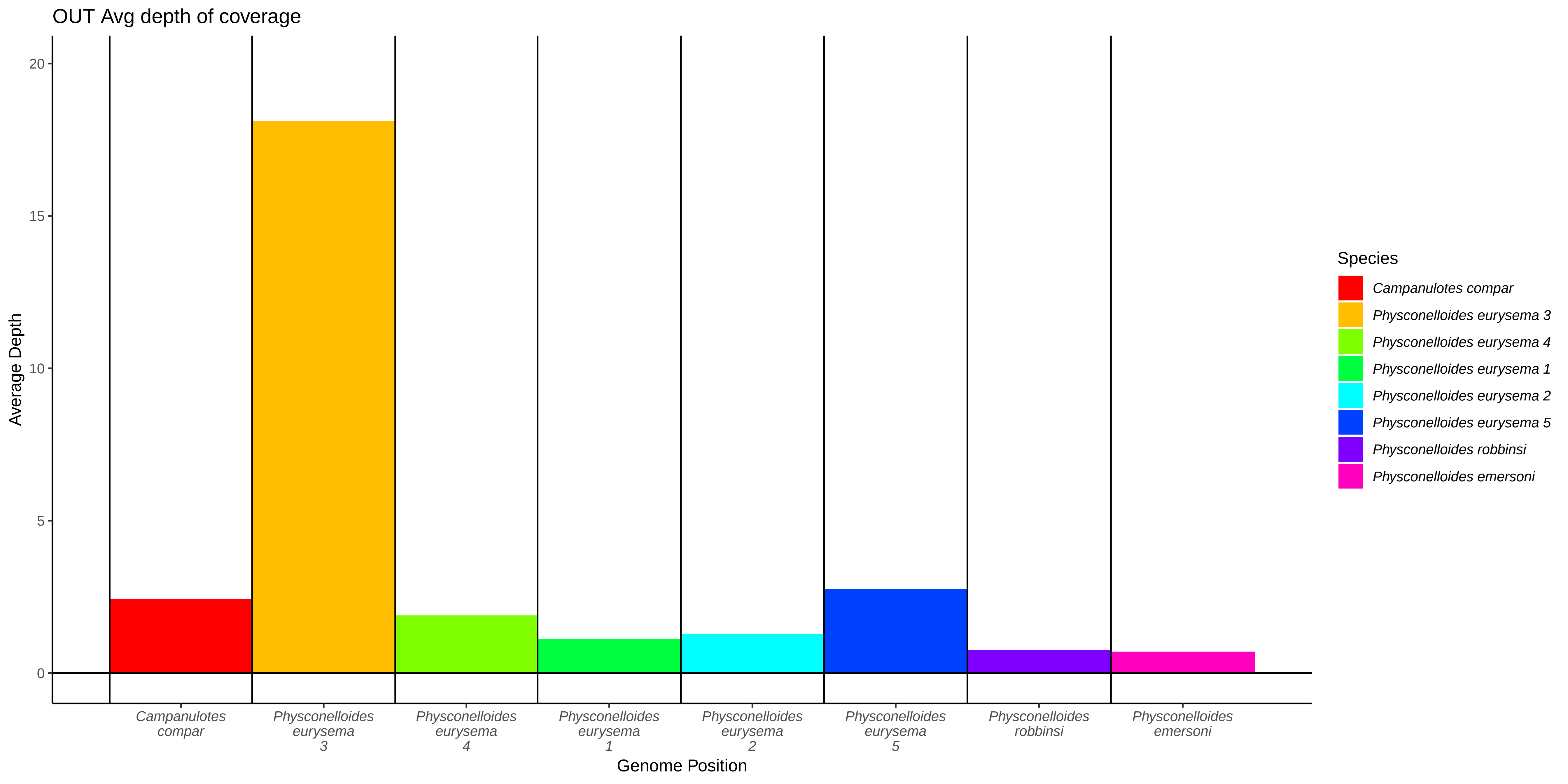

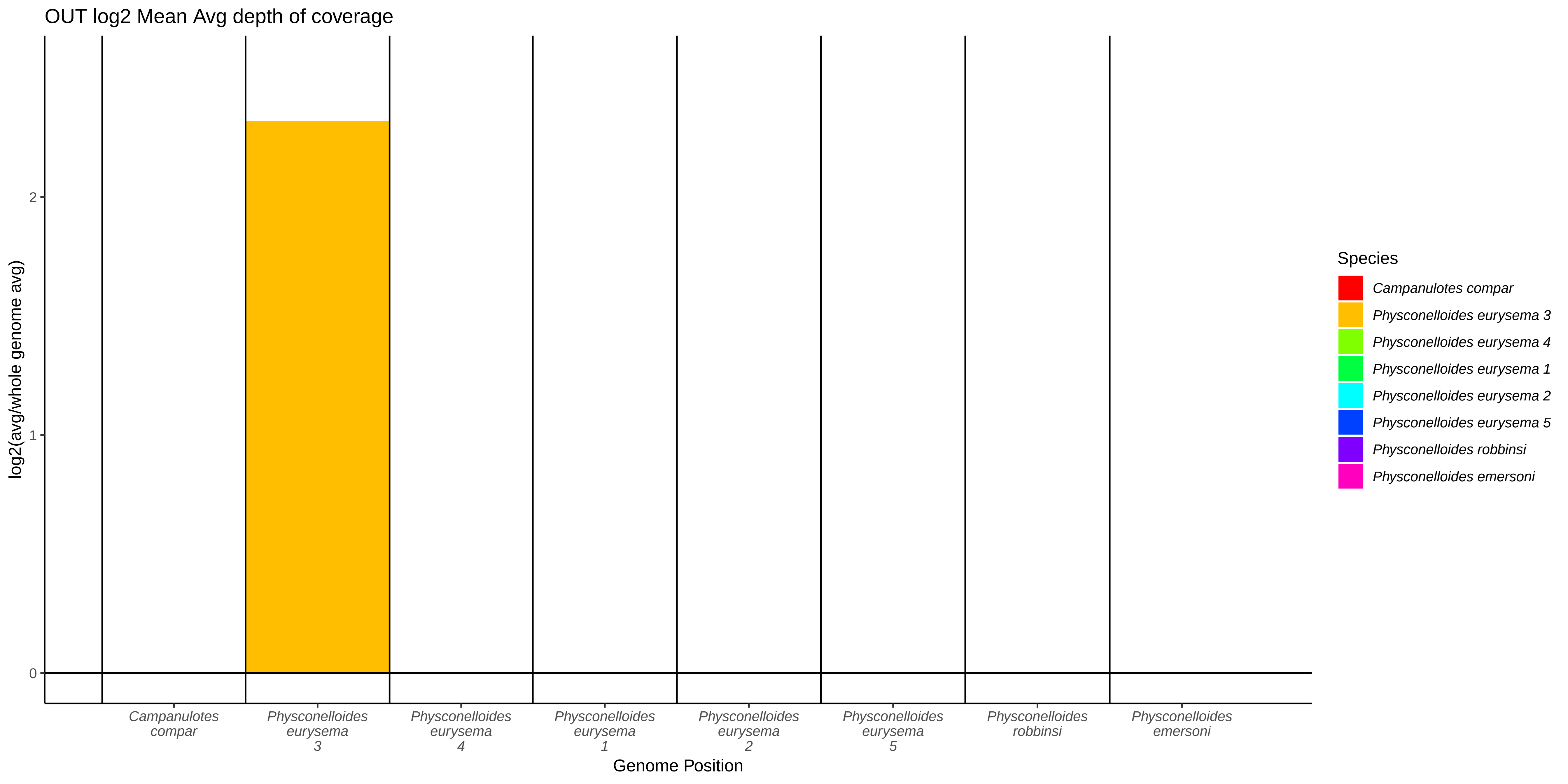

### OUT_speciesDepth_Pheur_Cotal_3_16_2016_12.pdf

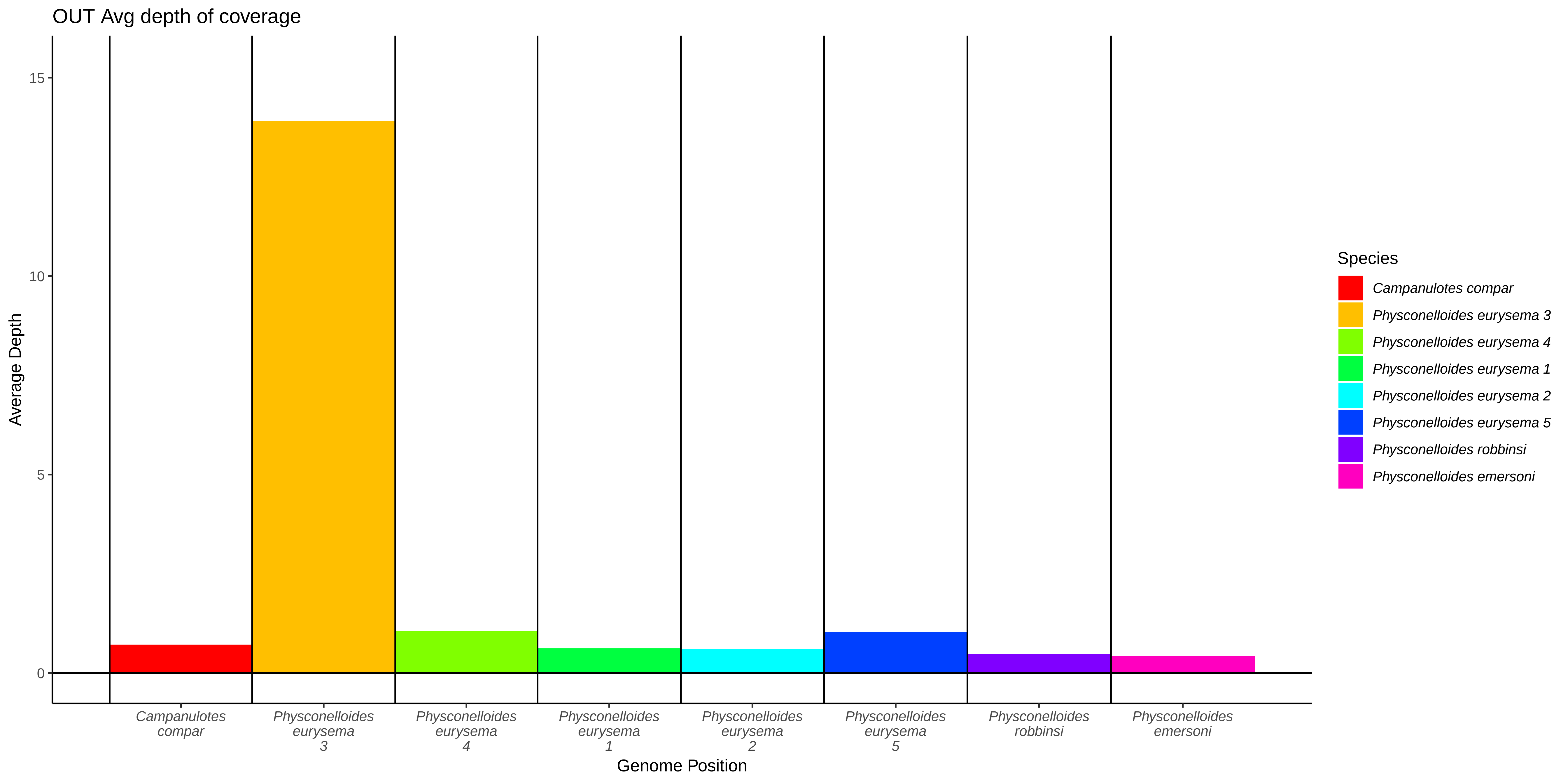

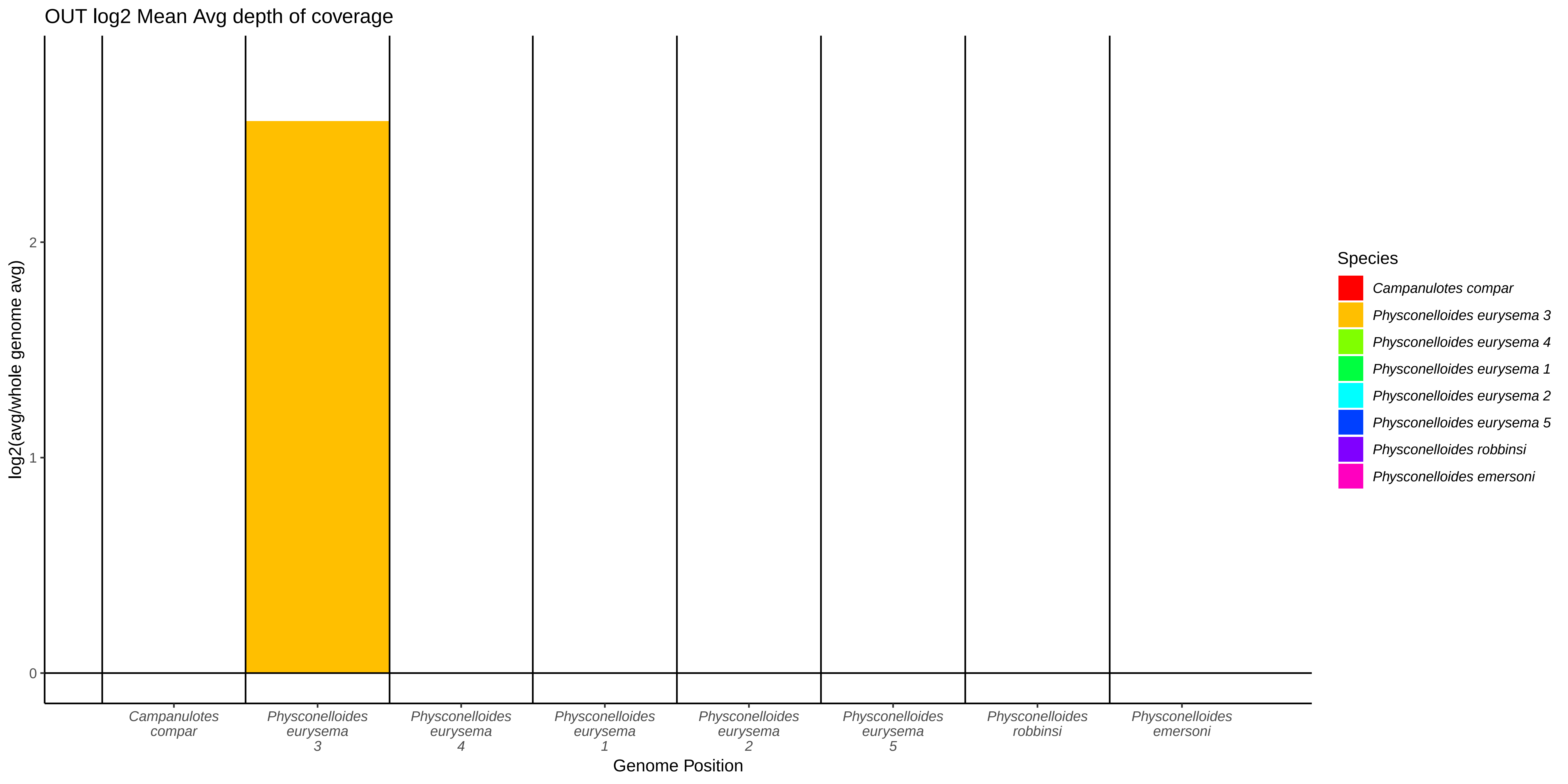

### OUT_speciesDepth_Pheur_Cotal_6_15_2015_4.pdf

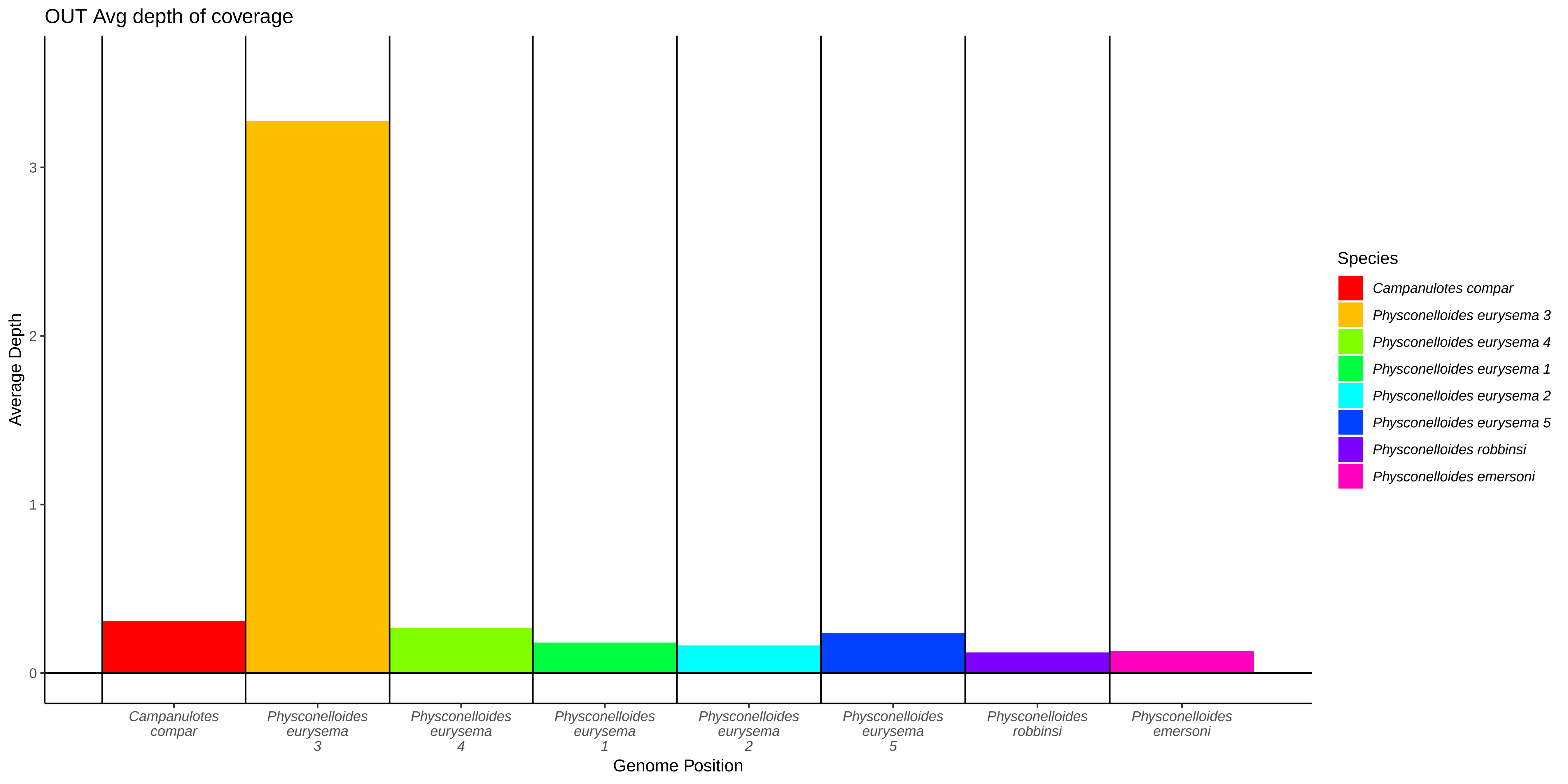

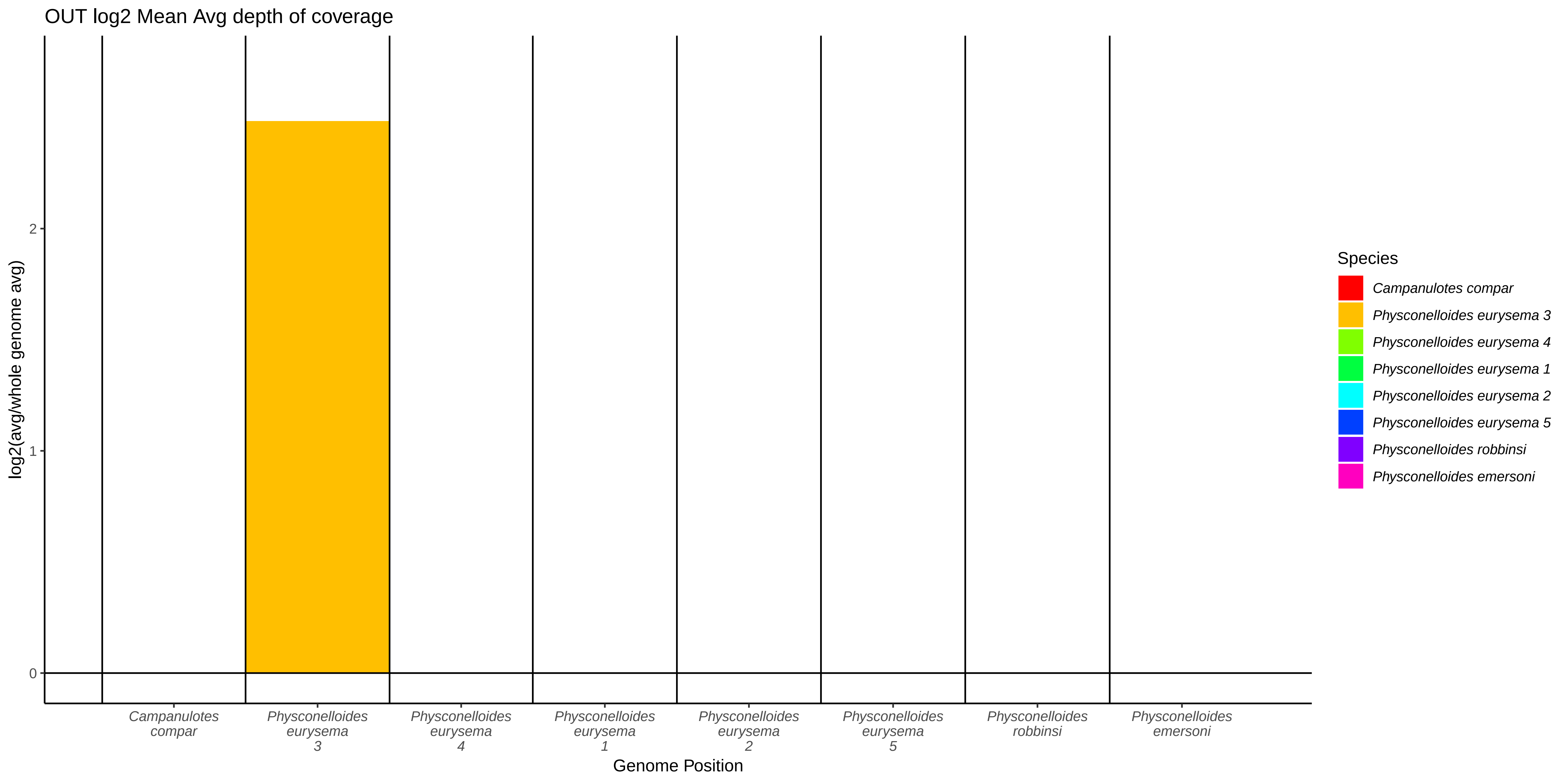

### OUT_speciesDepth_Pheur_Urcam_3_14_2016_3.pdf

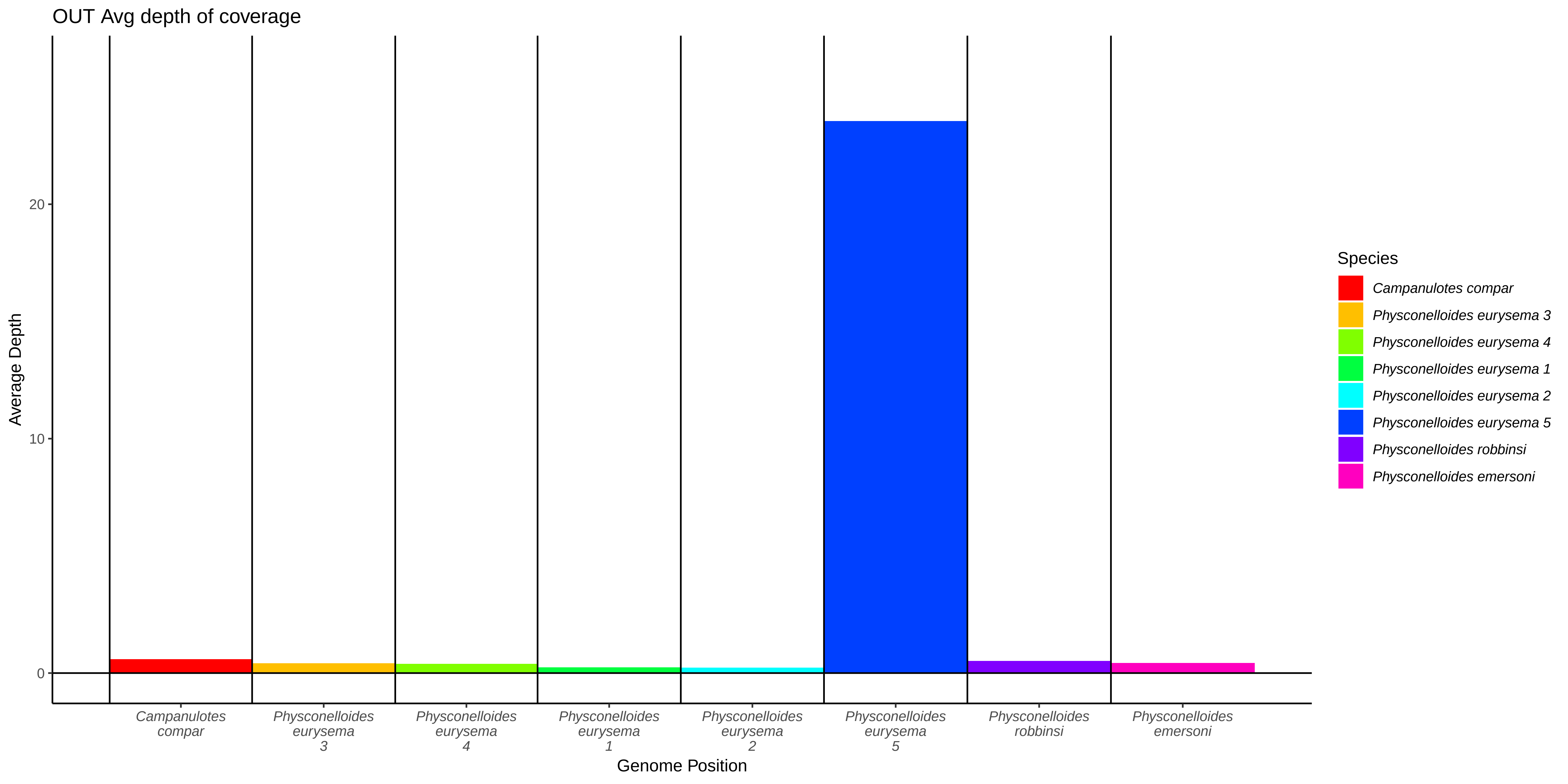

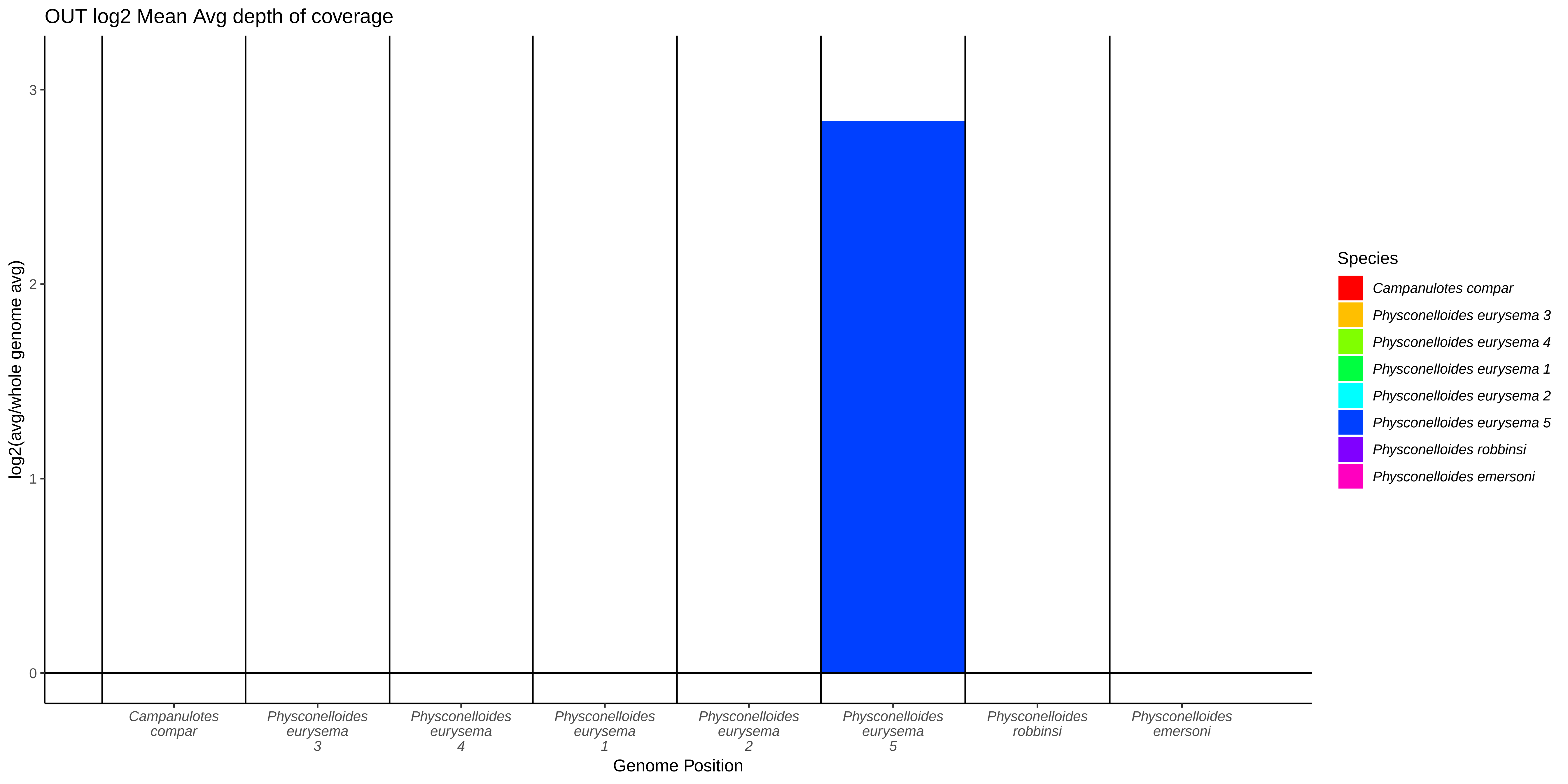

### OUT_speciesDepth_Phrob_Mecec_3_16_2016_10.pdf

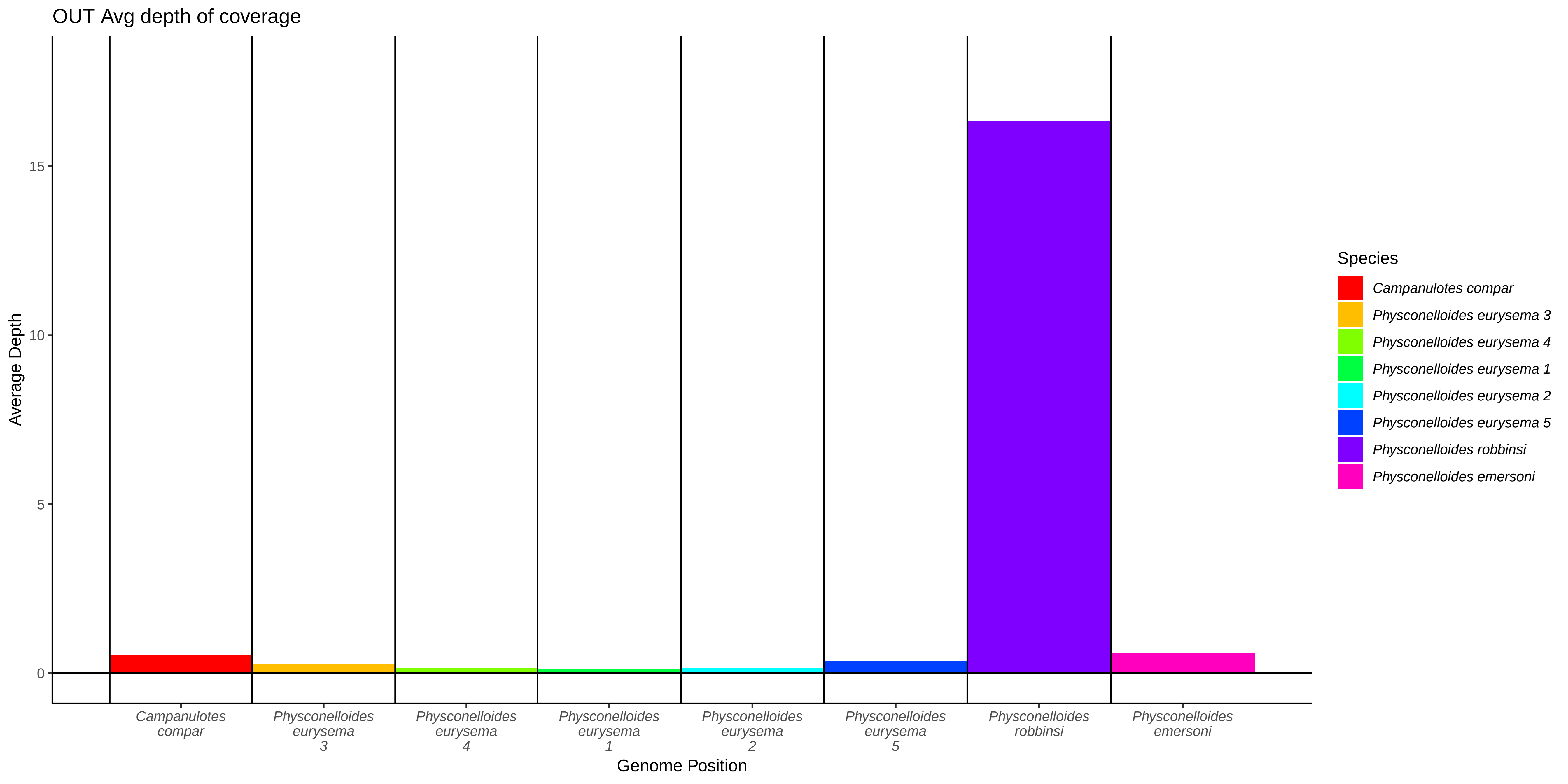

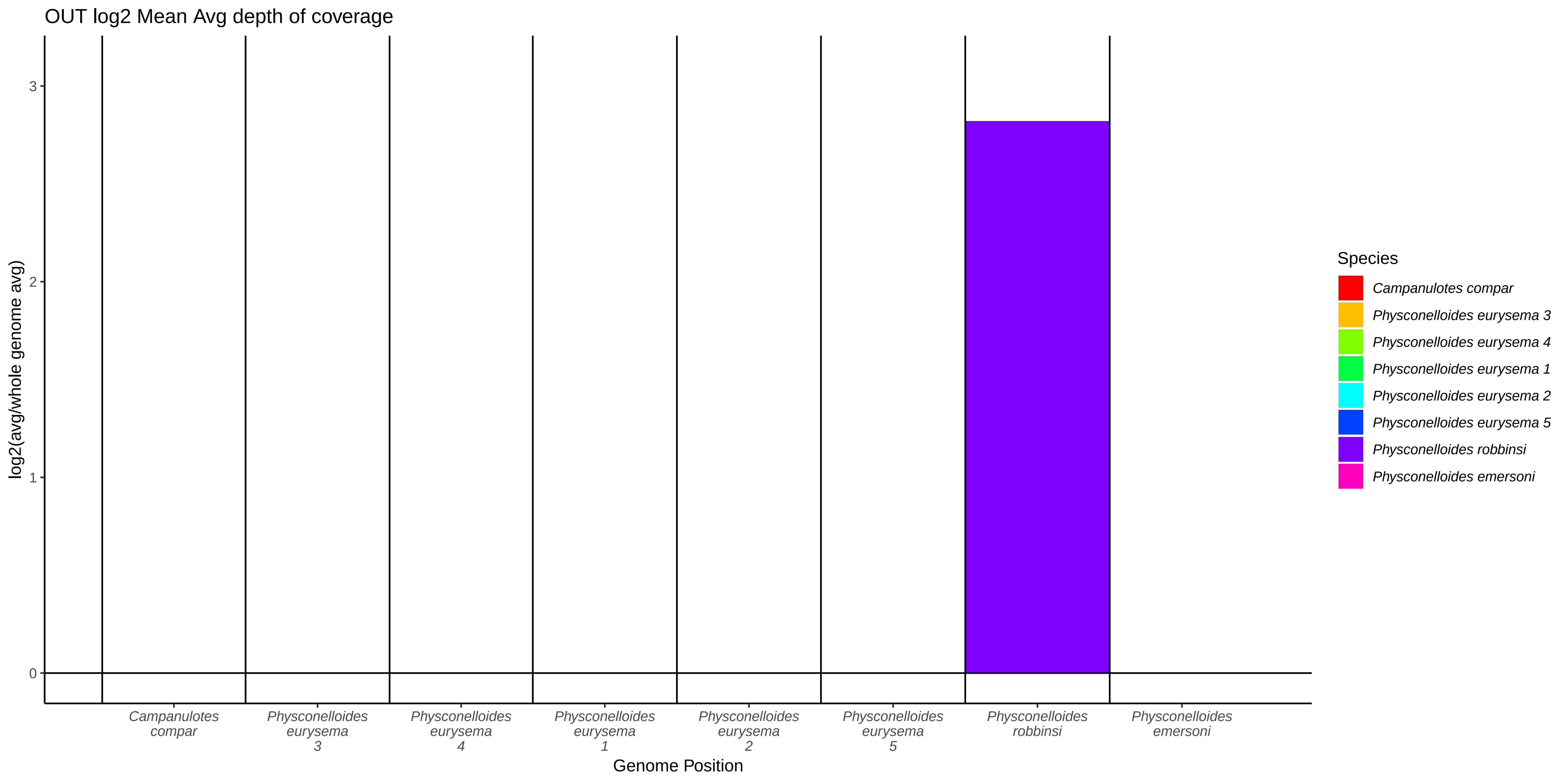

### OUT_speciesDepth_Phrob_Mecec_3_16_2016_11.pdf

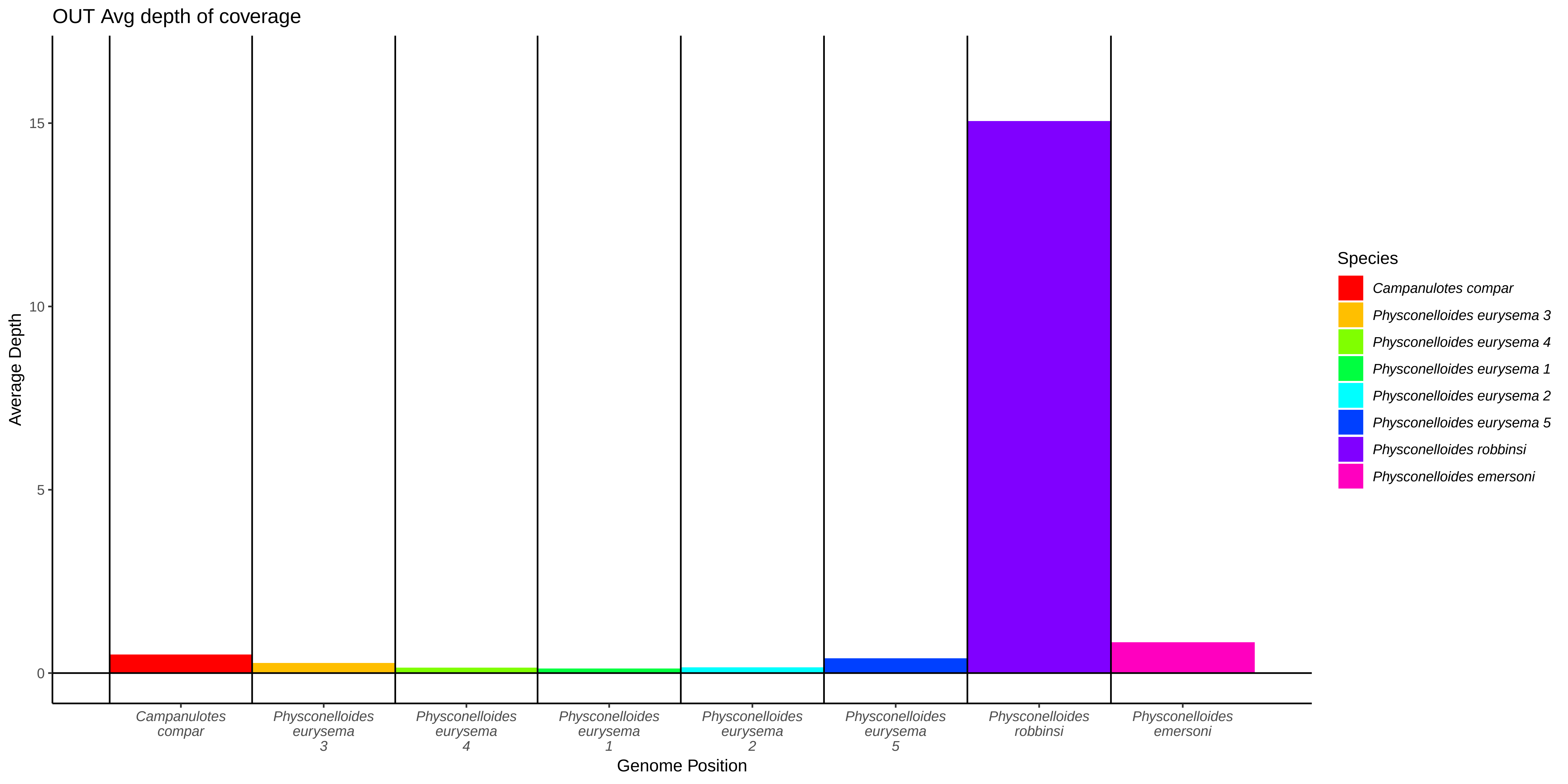

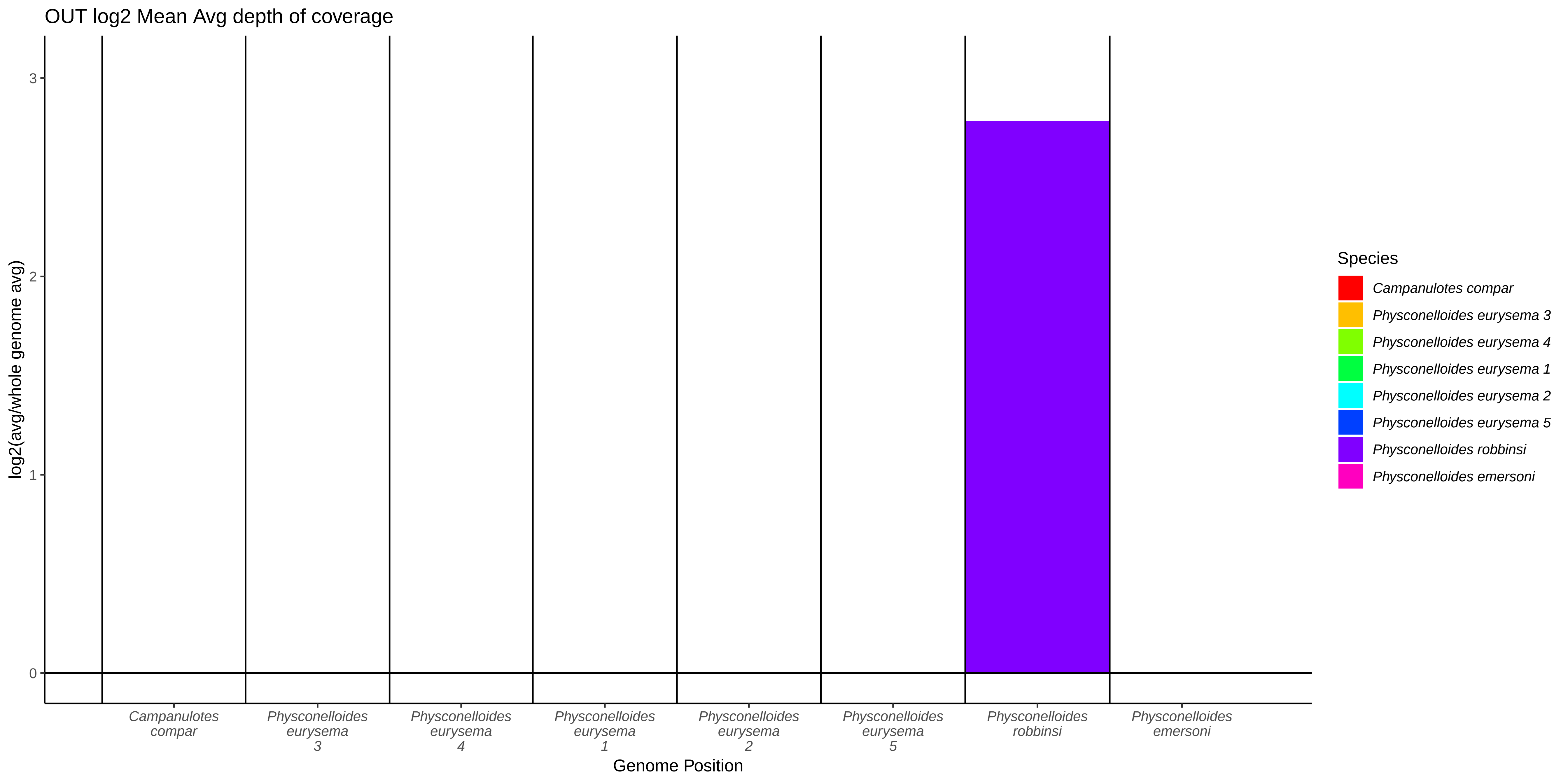

### OUT_speciesDepth_Phrob_Mecec_5_26_2015_7.pdf

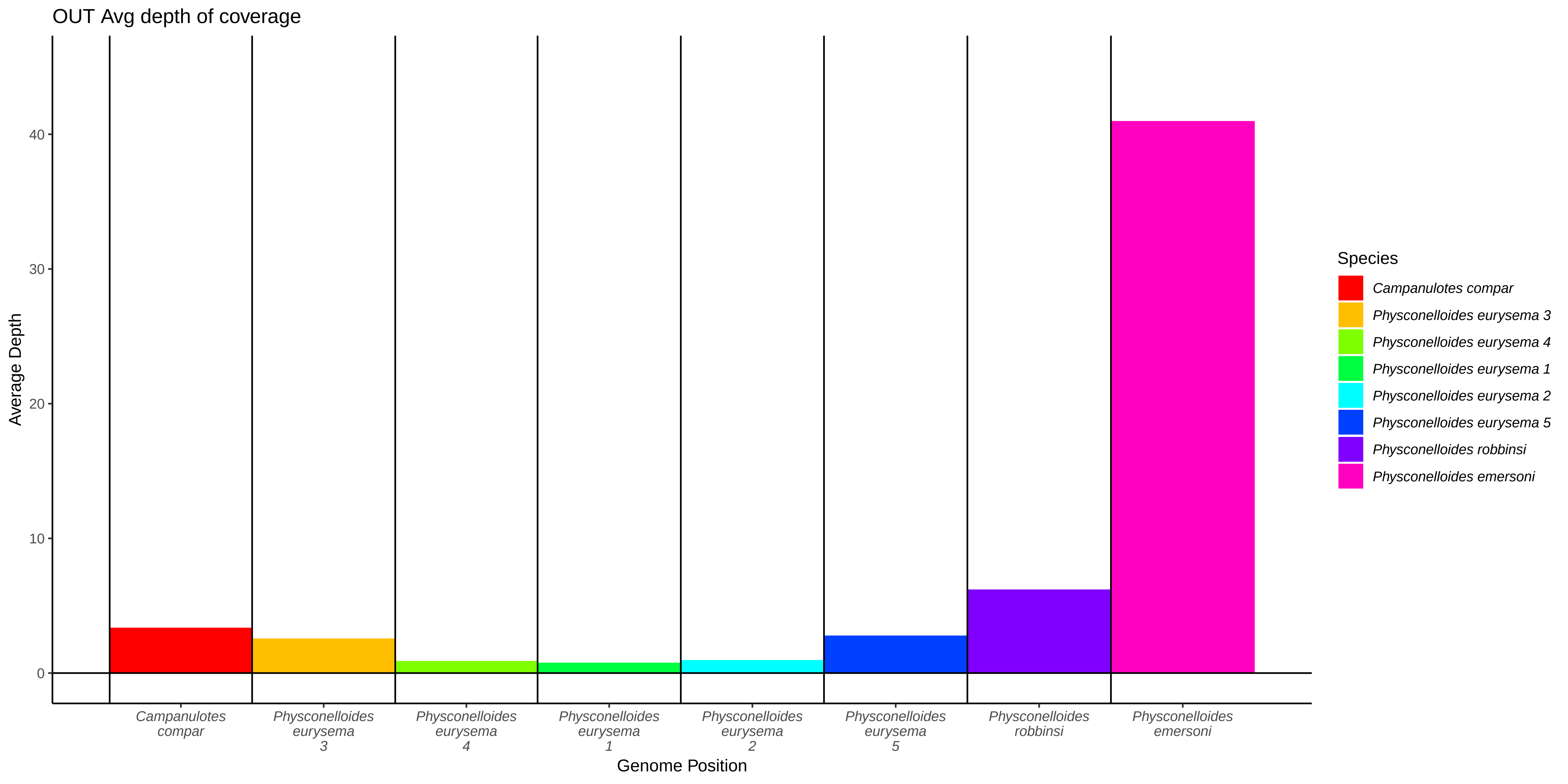

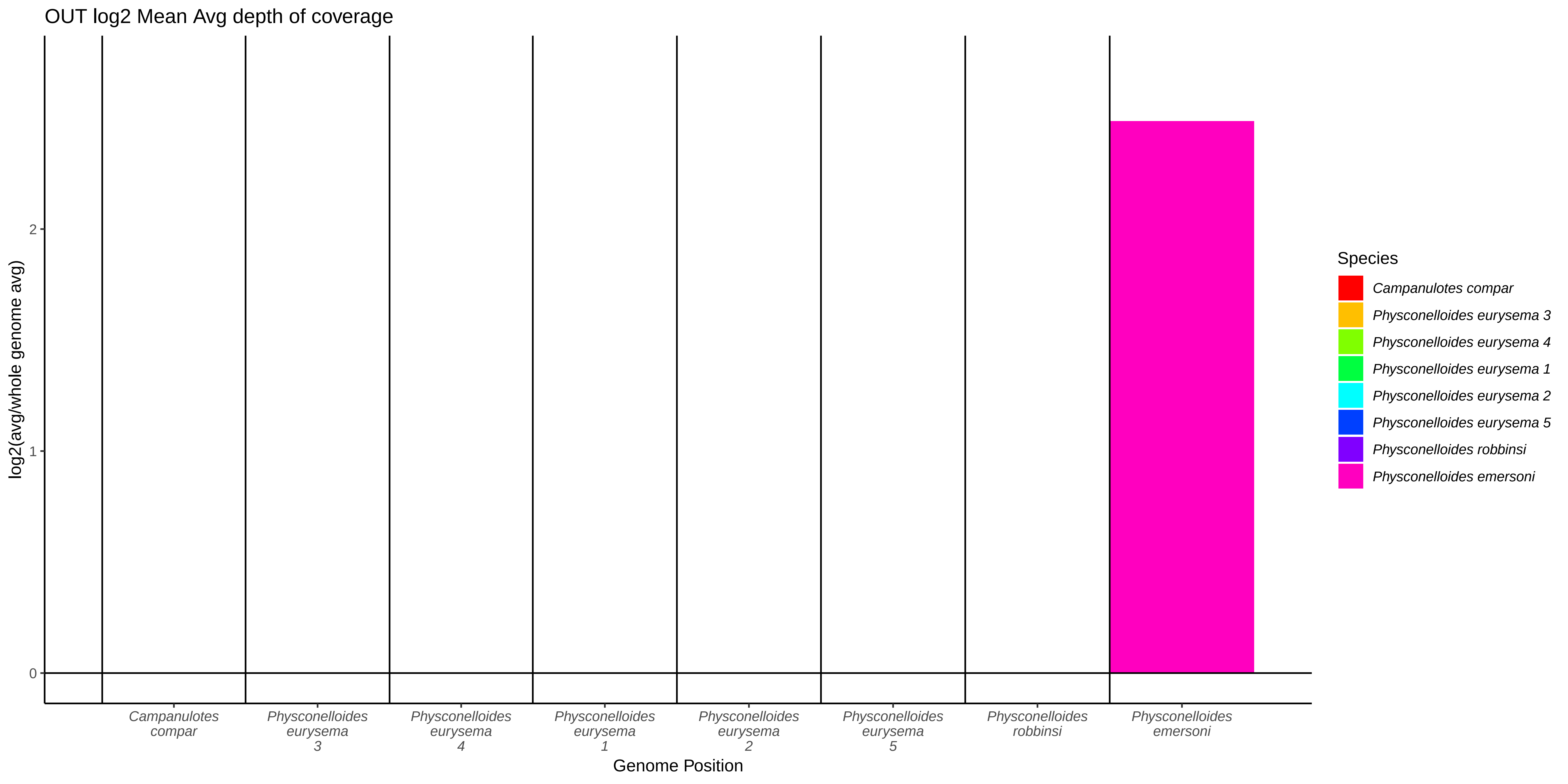

### OUT_speciesDepth_Phrob_Mecec_6_15_2015_6.pdf

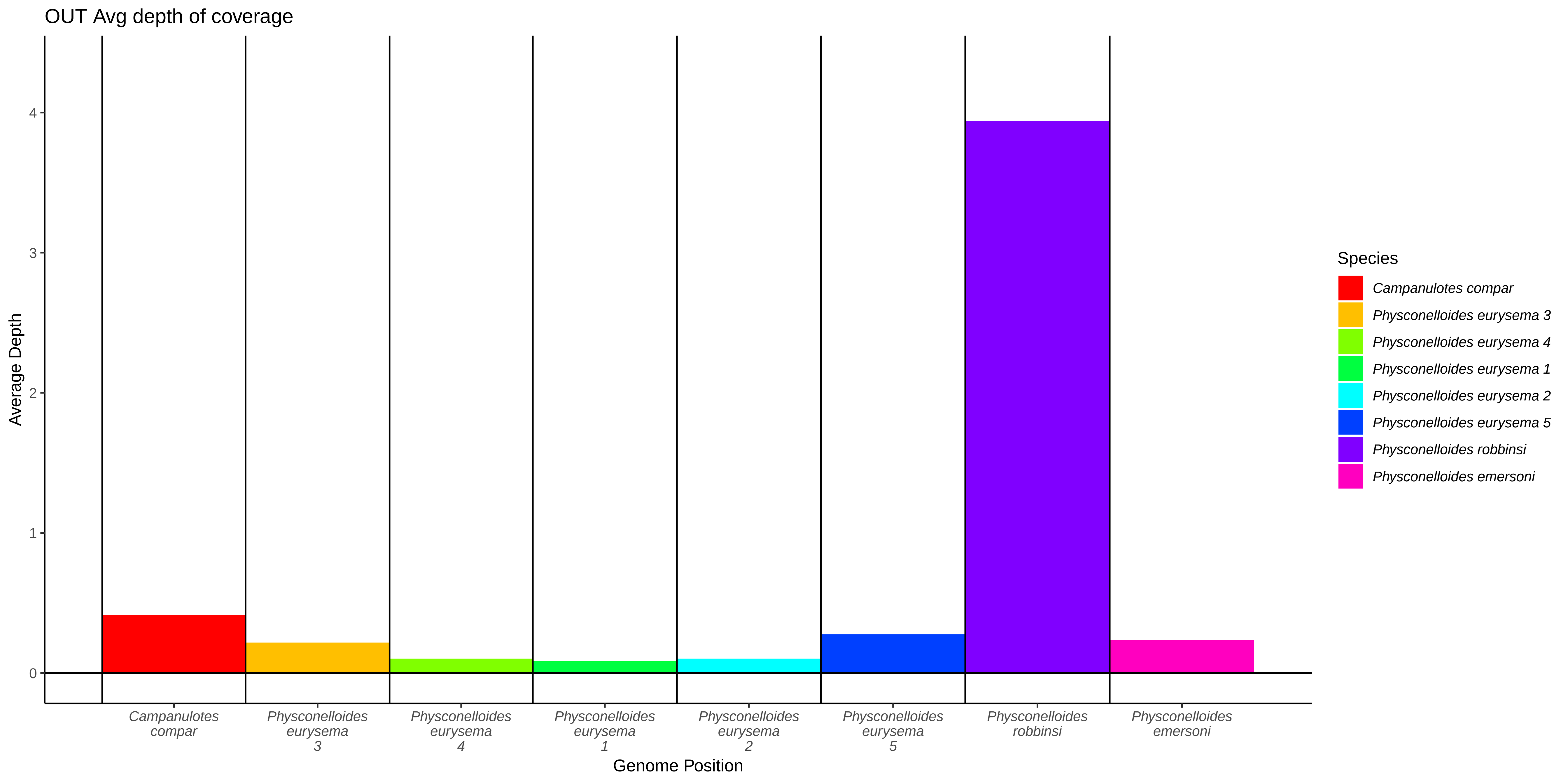

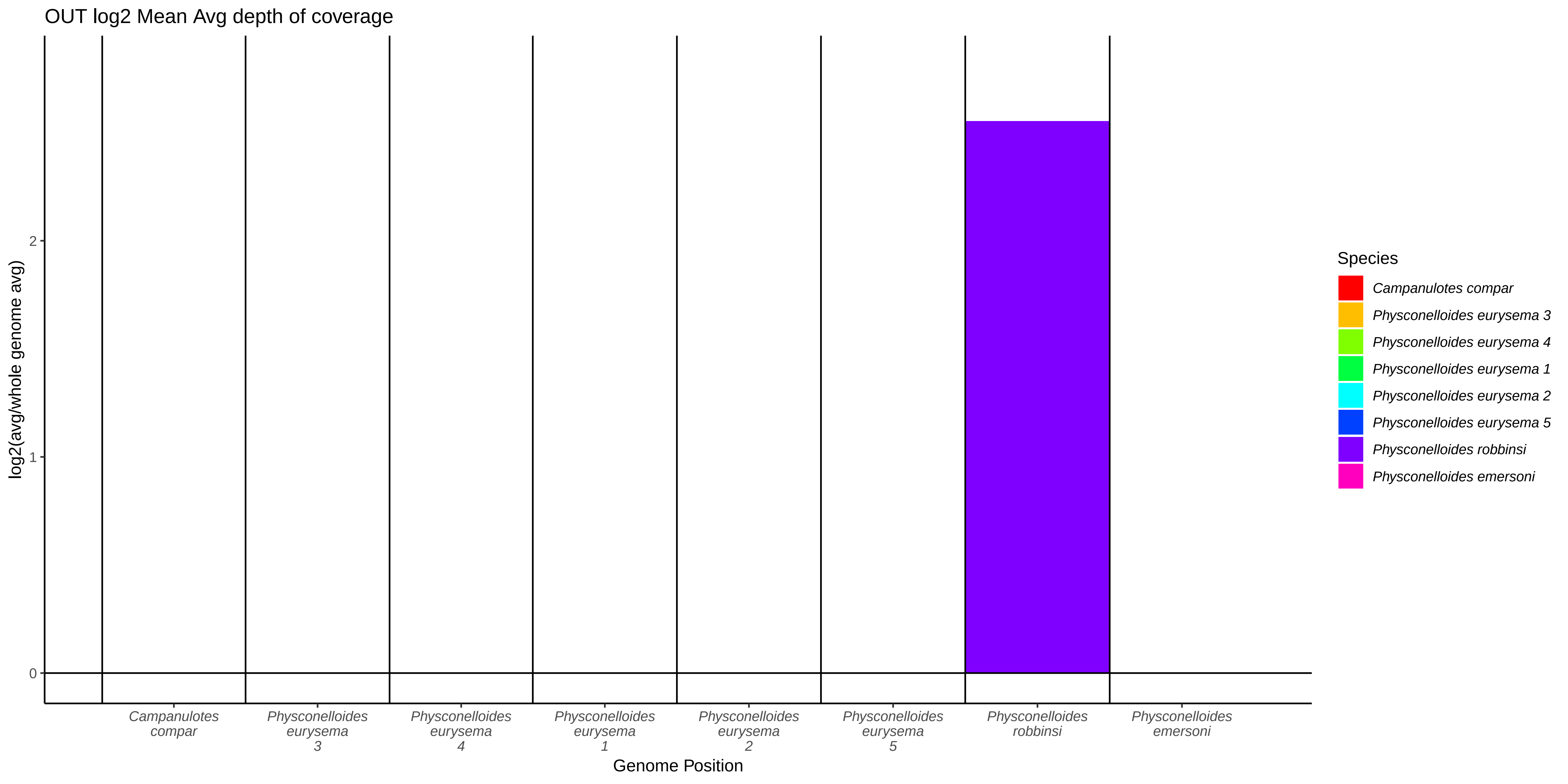
