## Supplementary figures and images for "Comparing rates of introgression in parasitic feather lice with differing dispersal capabilities" (from another version of this preprint: version 7, 2020-03-31)

### network_columbicola.pdf

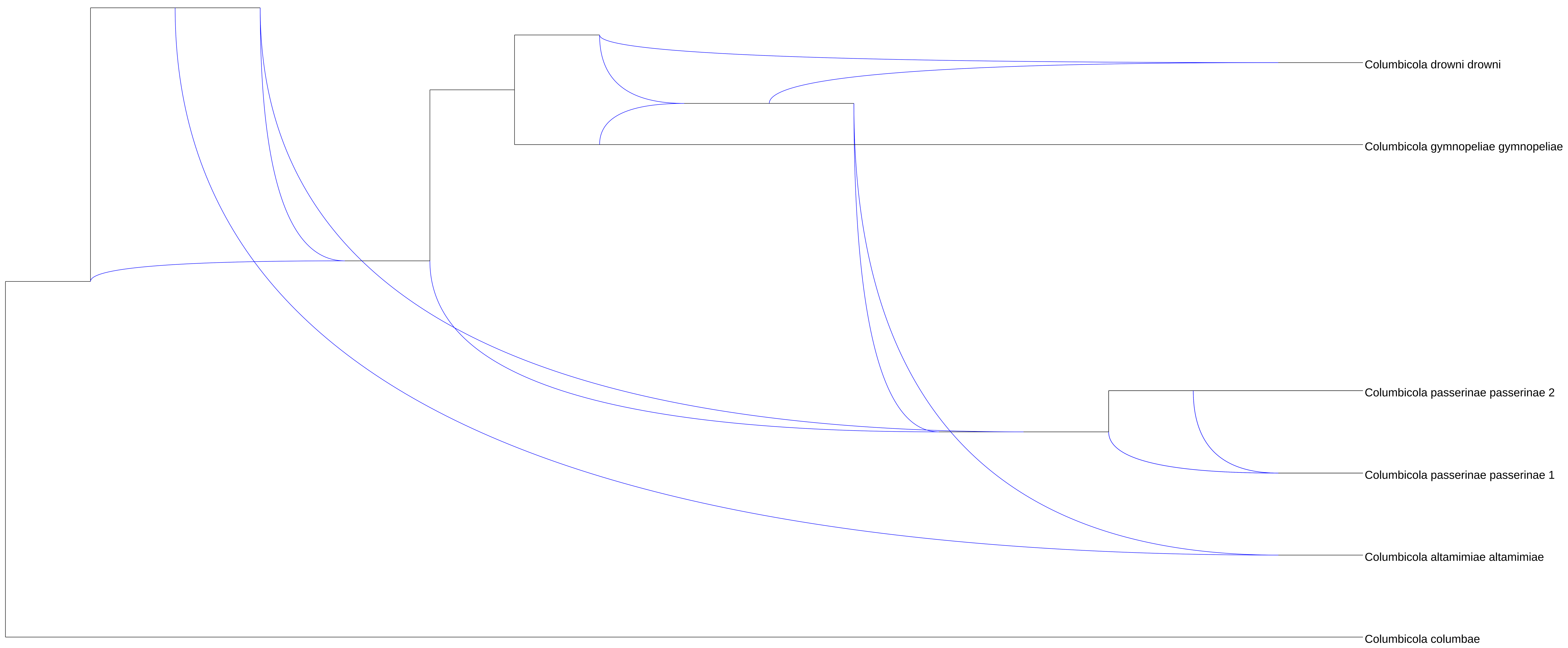

### network_physconelloides.pdf

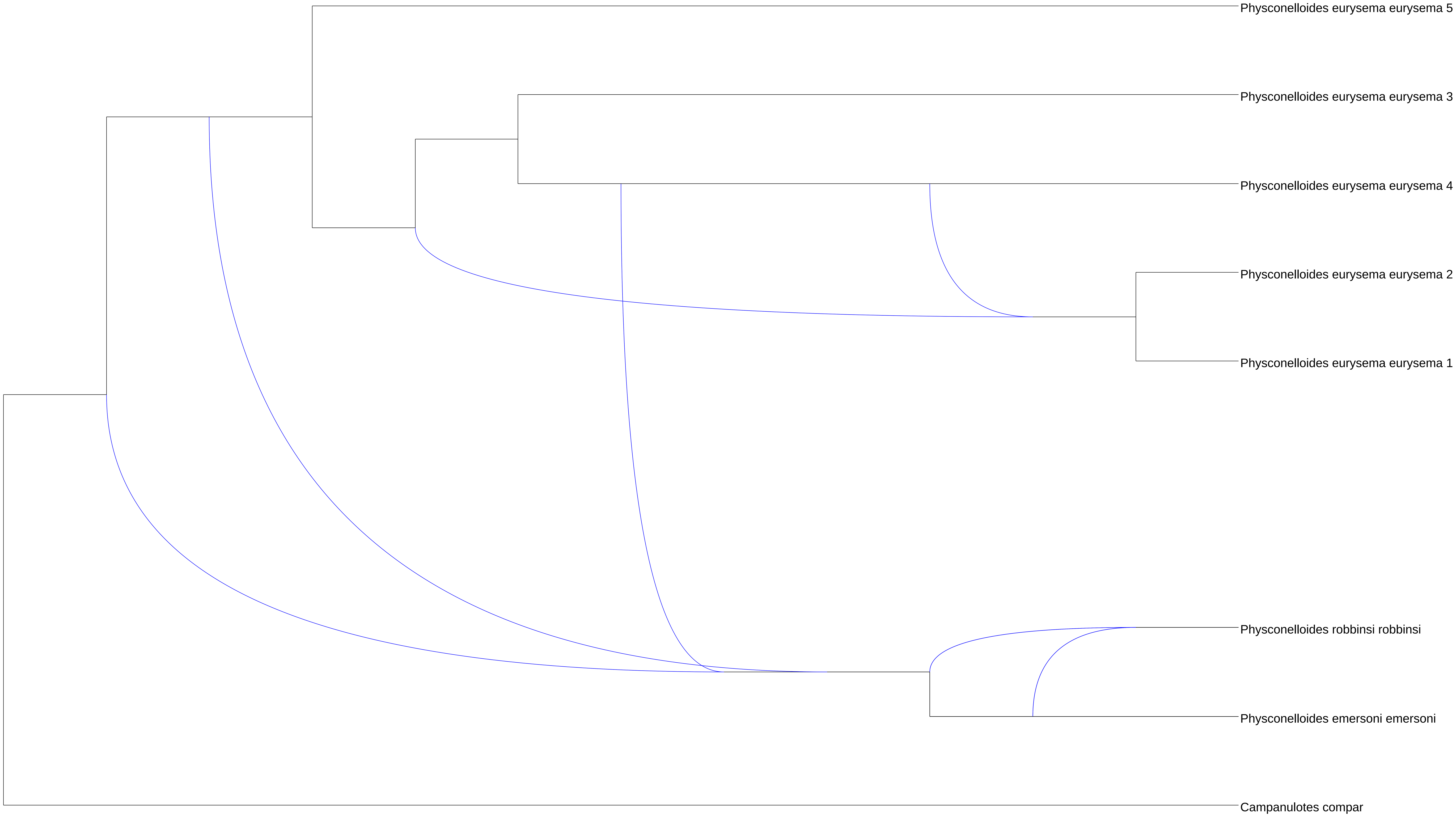

### OUT_plotMQ_coalt.pdf

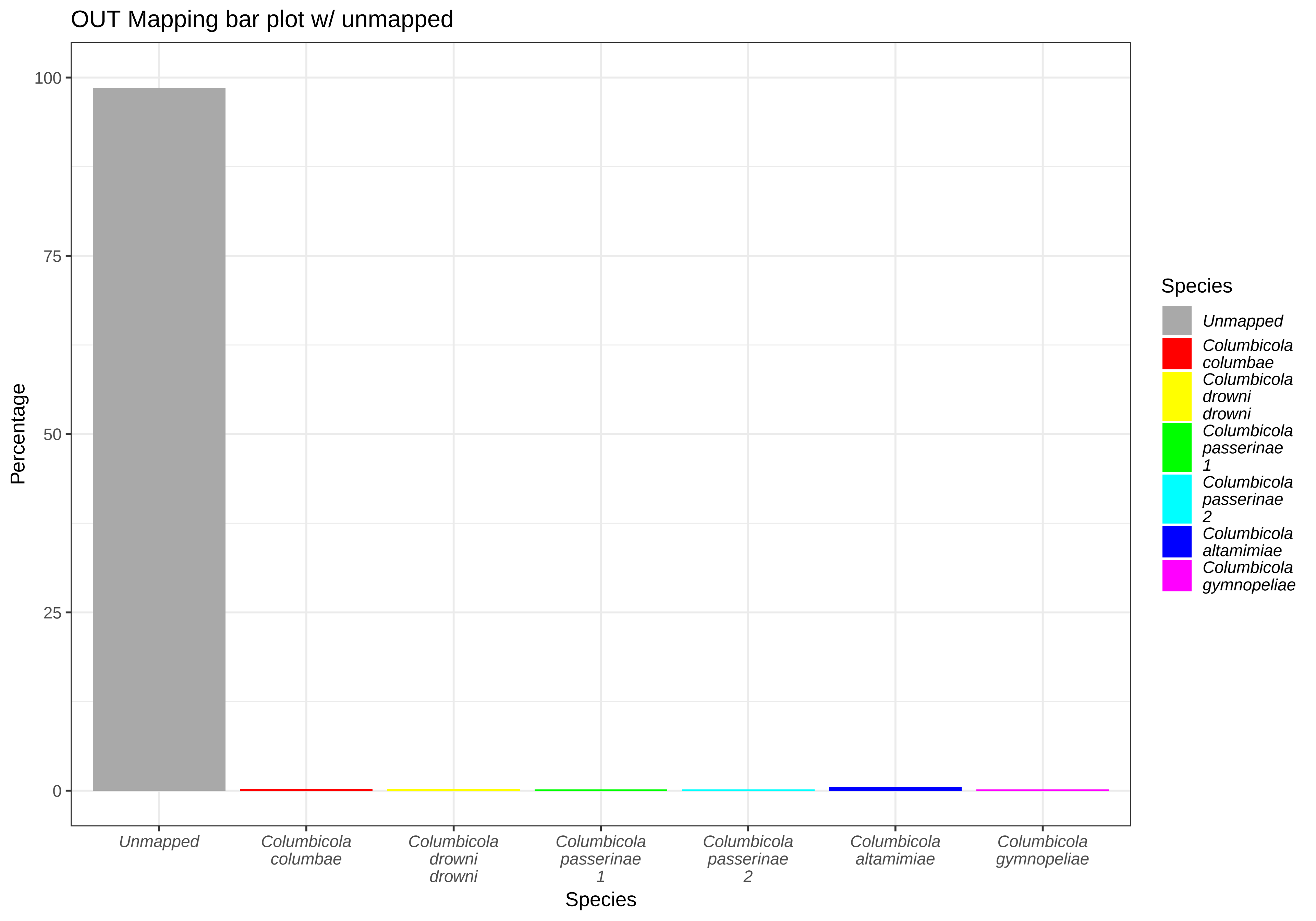

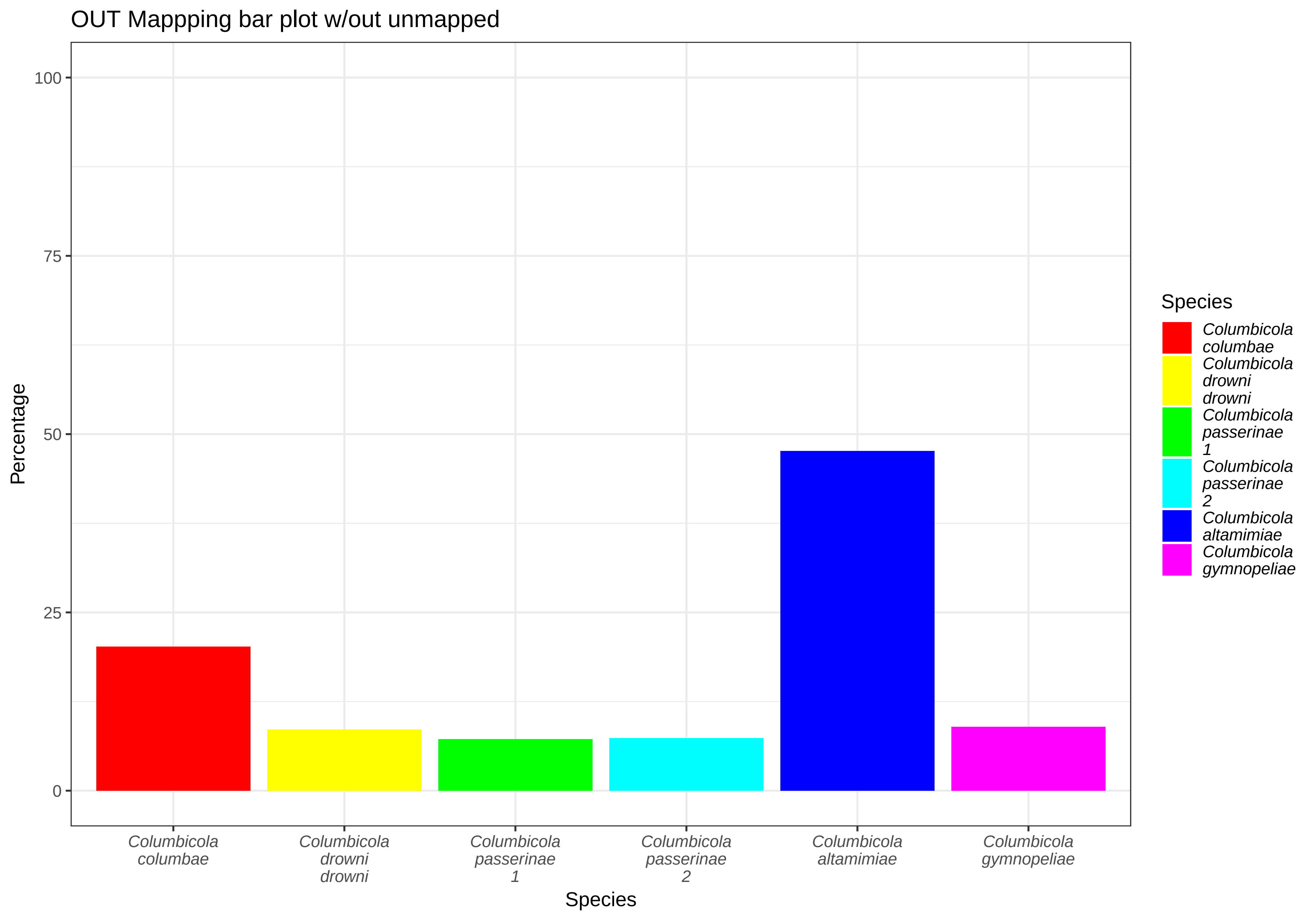

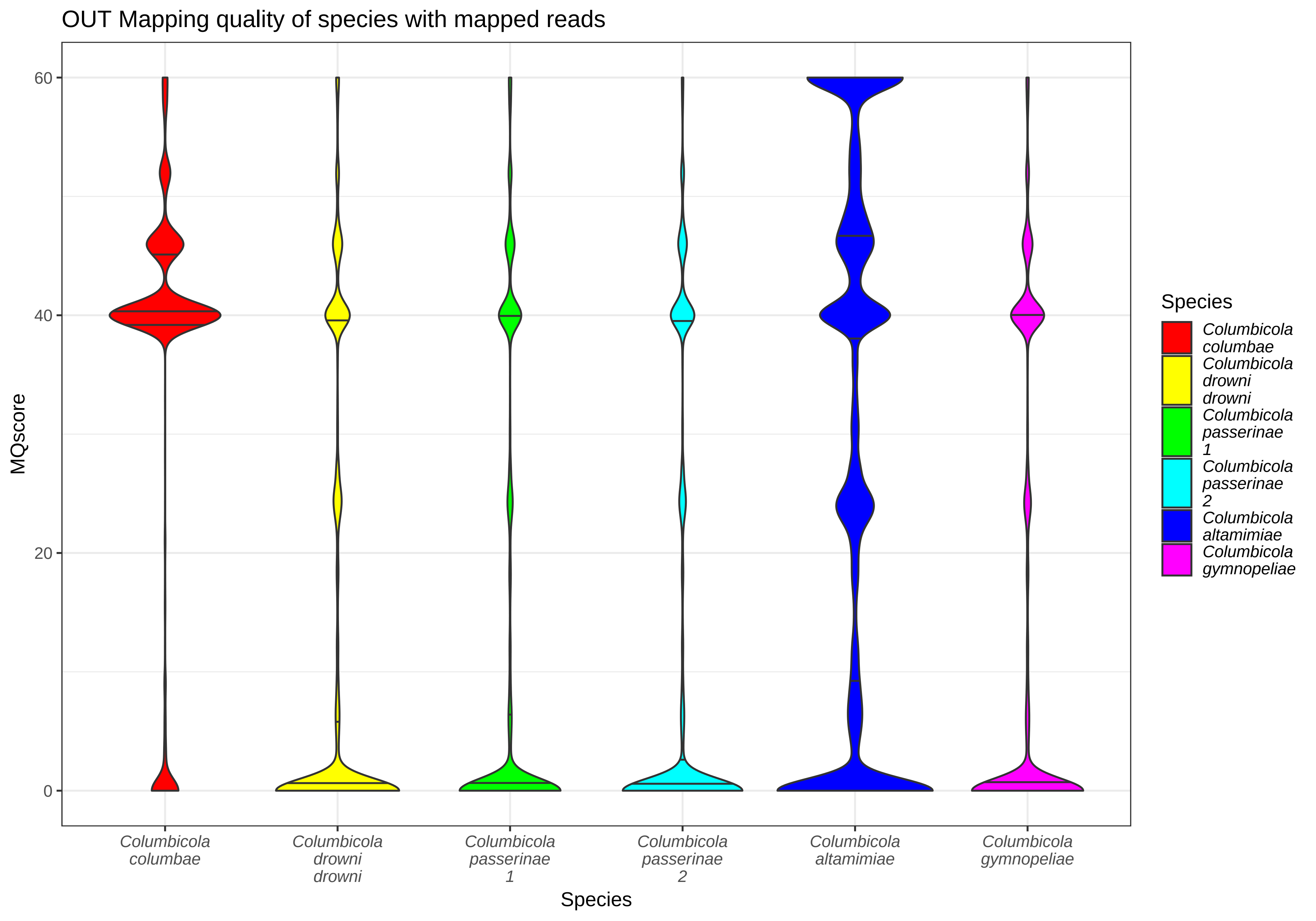

### OUT_plotMQ_Codro.pdf

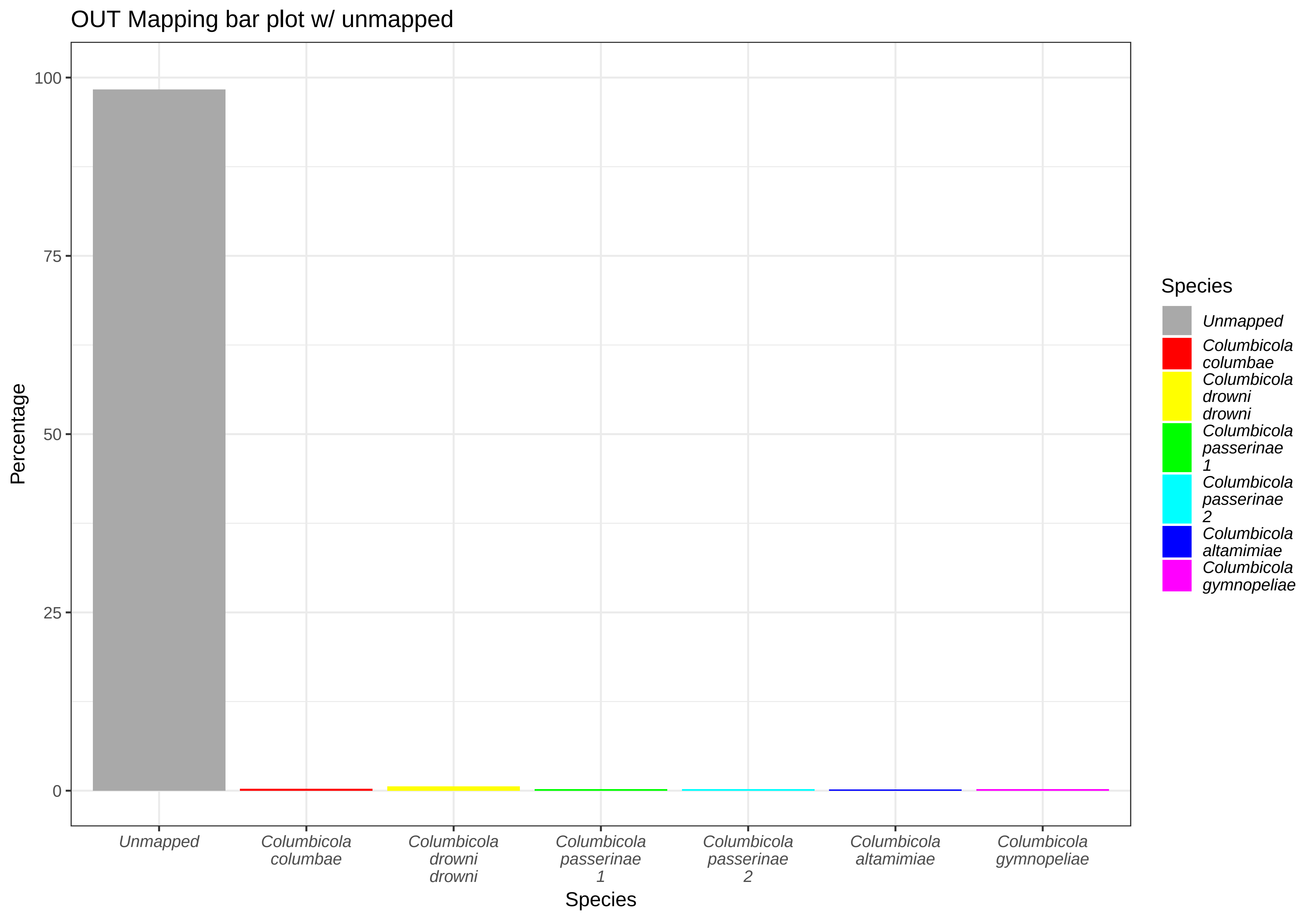

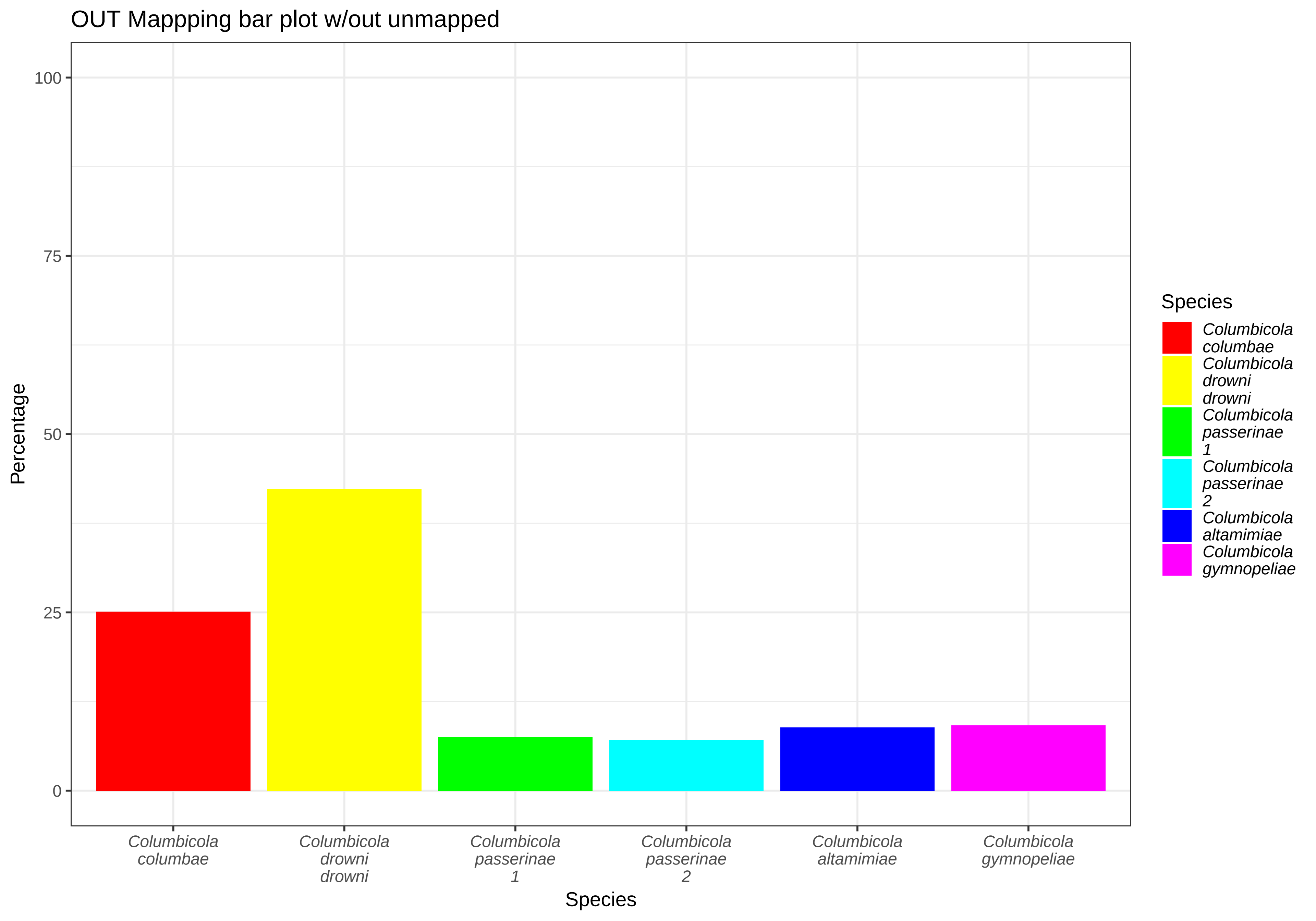

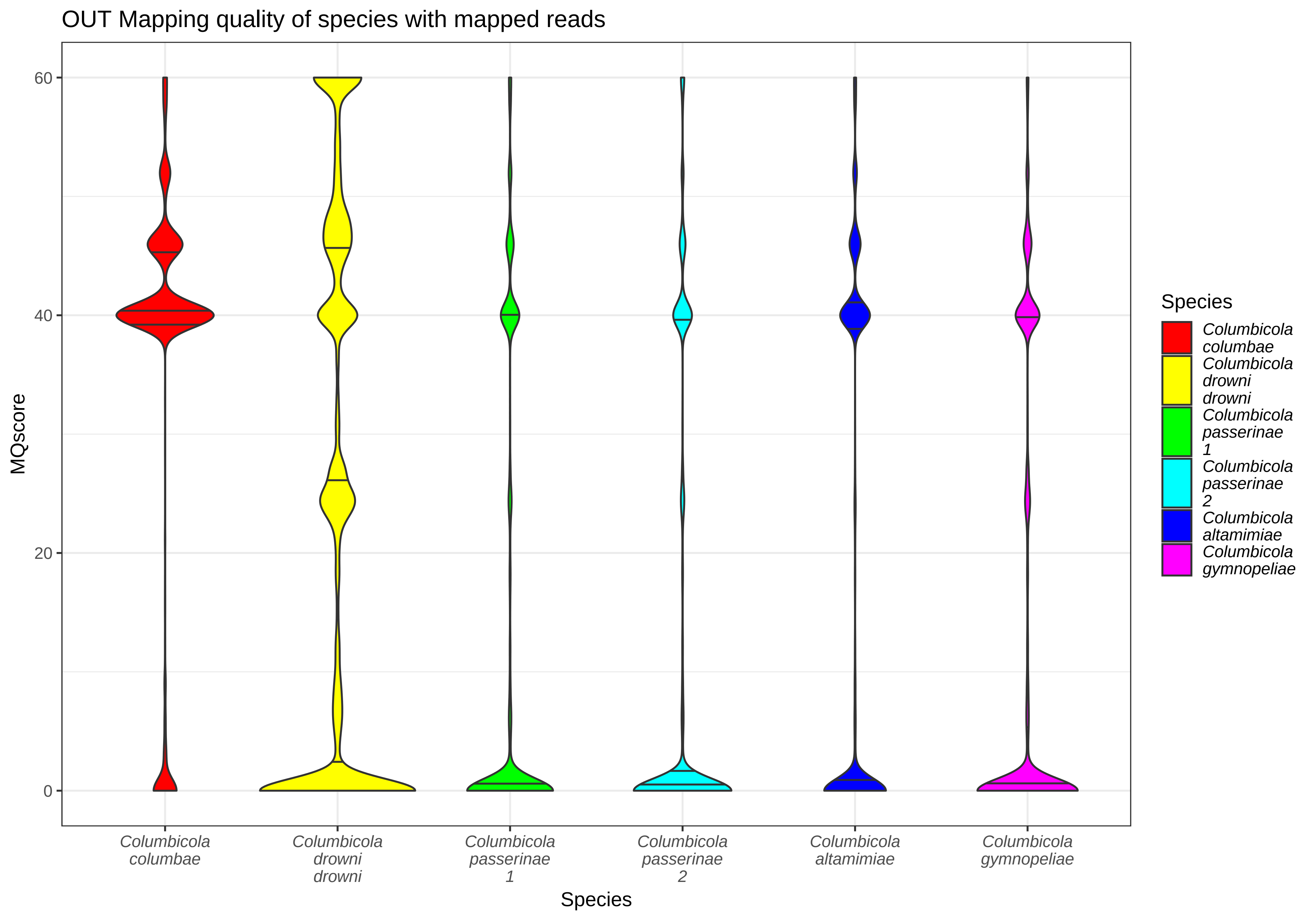

### OUT_plotMQ_cogym.pdf

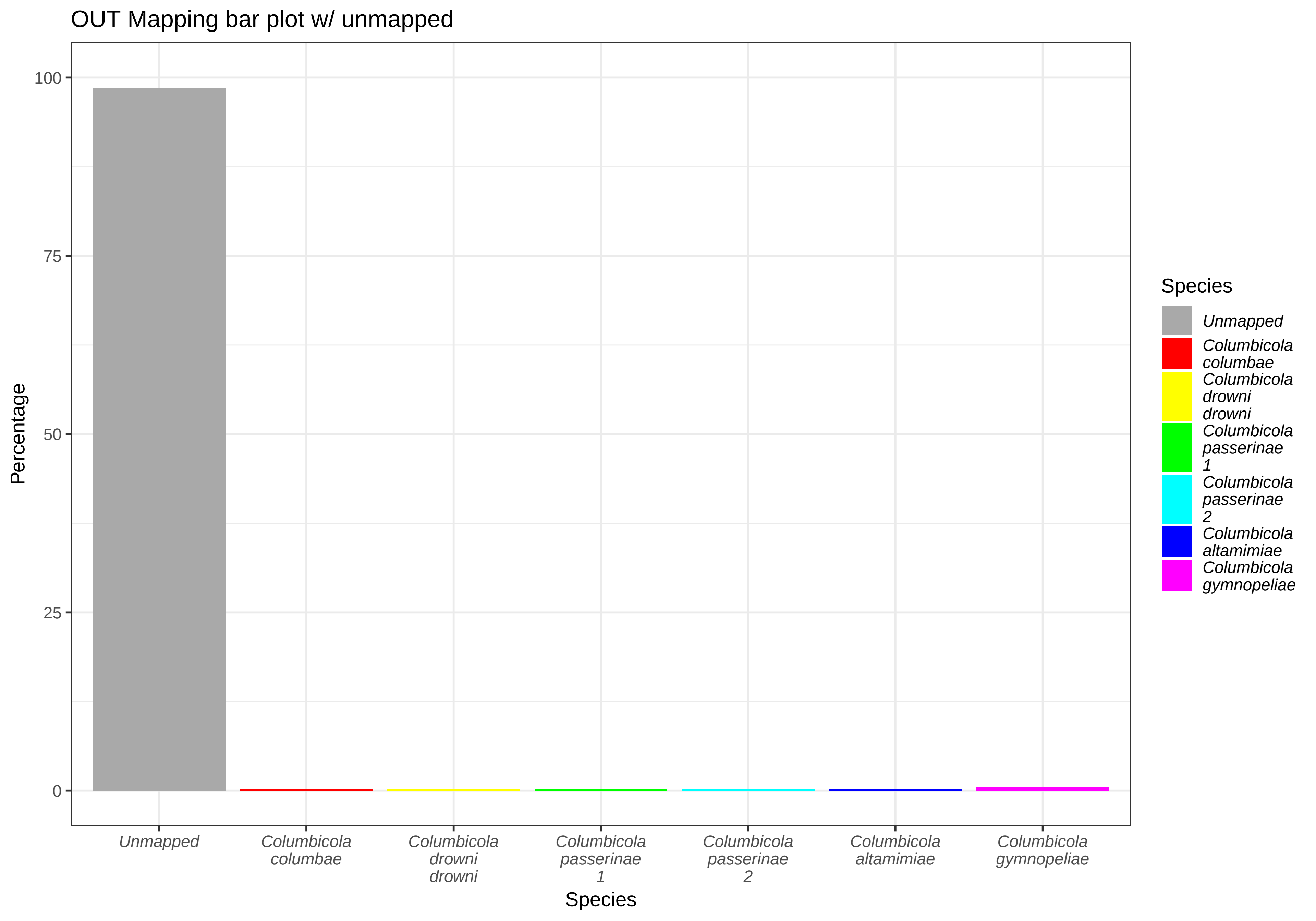

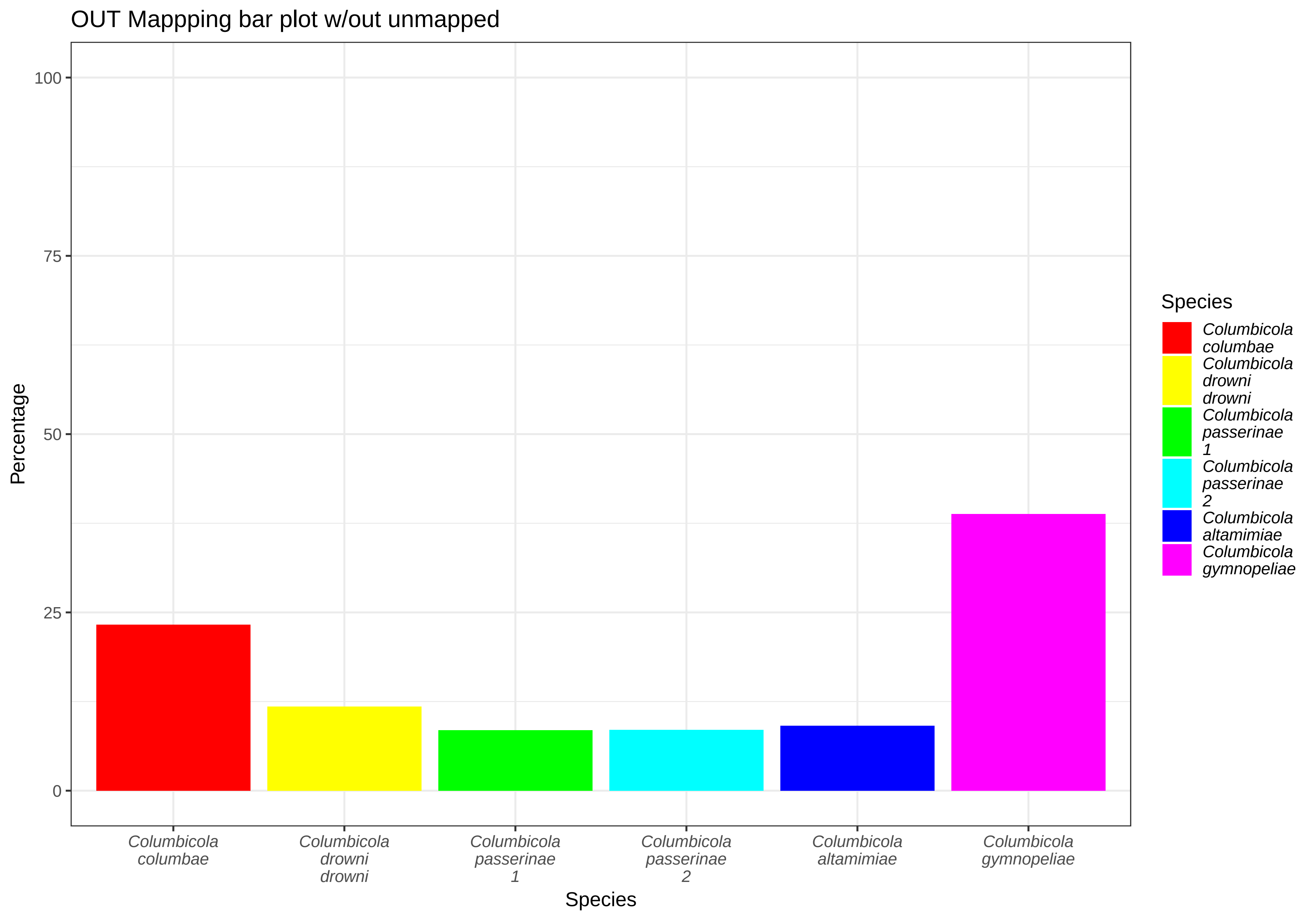

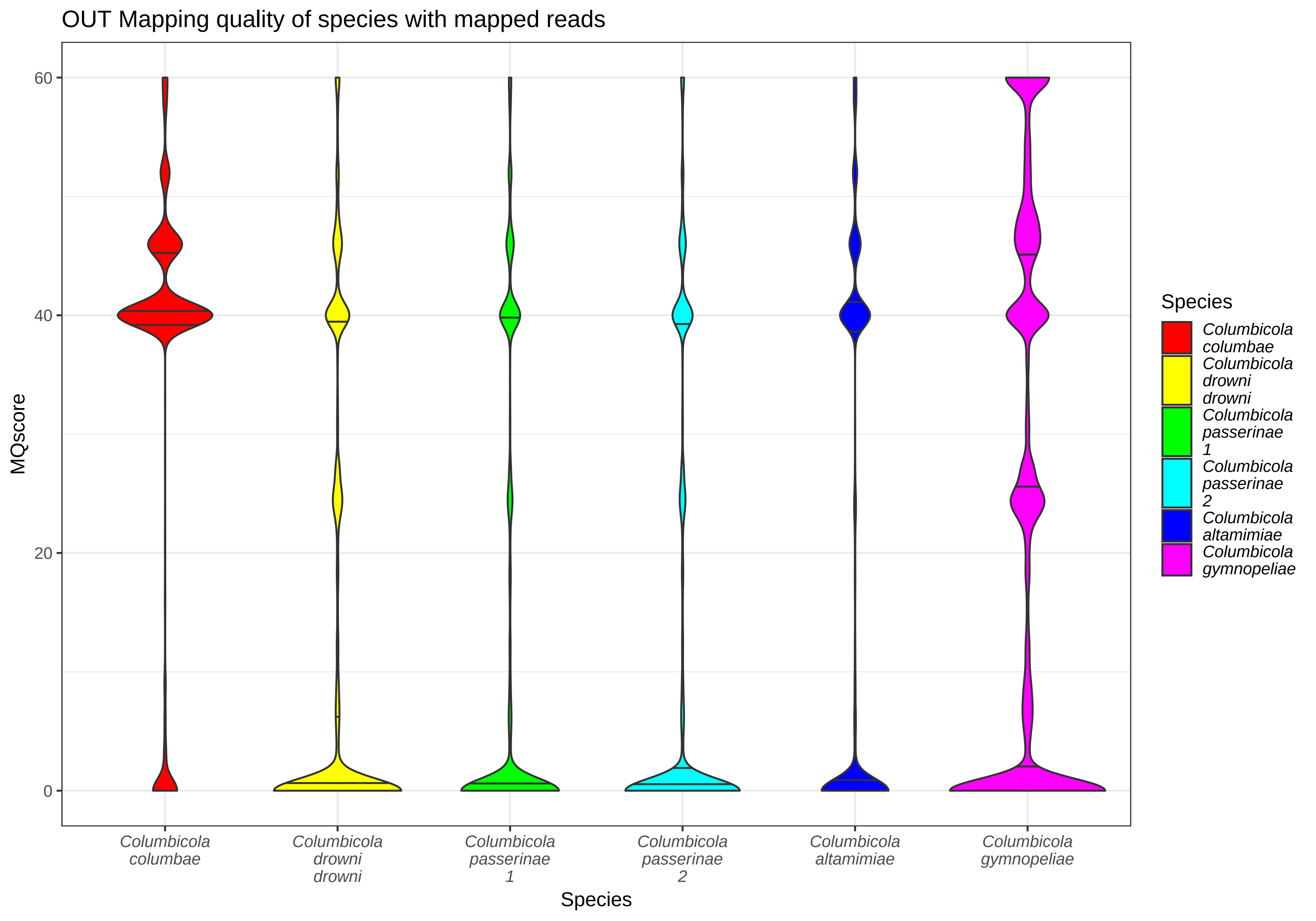

### OUT_plotMQ_copas1.pdf

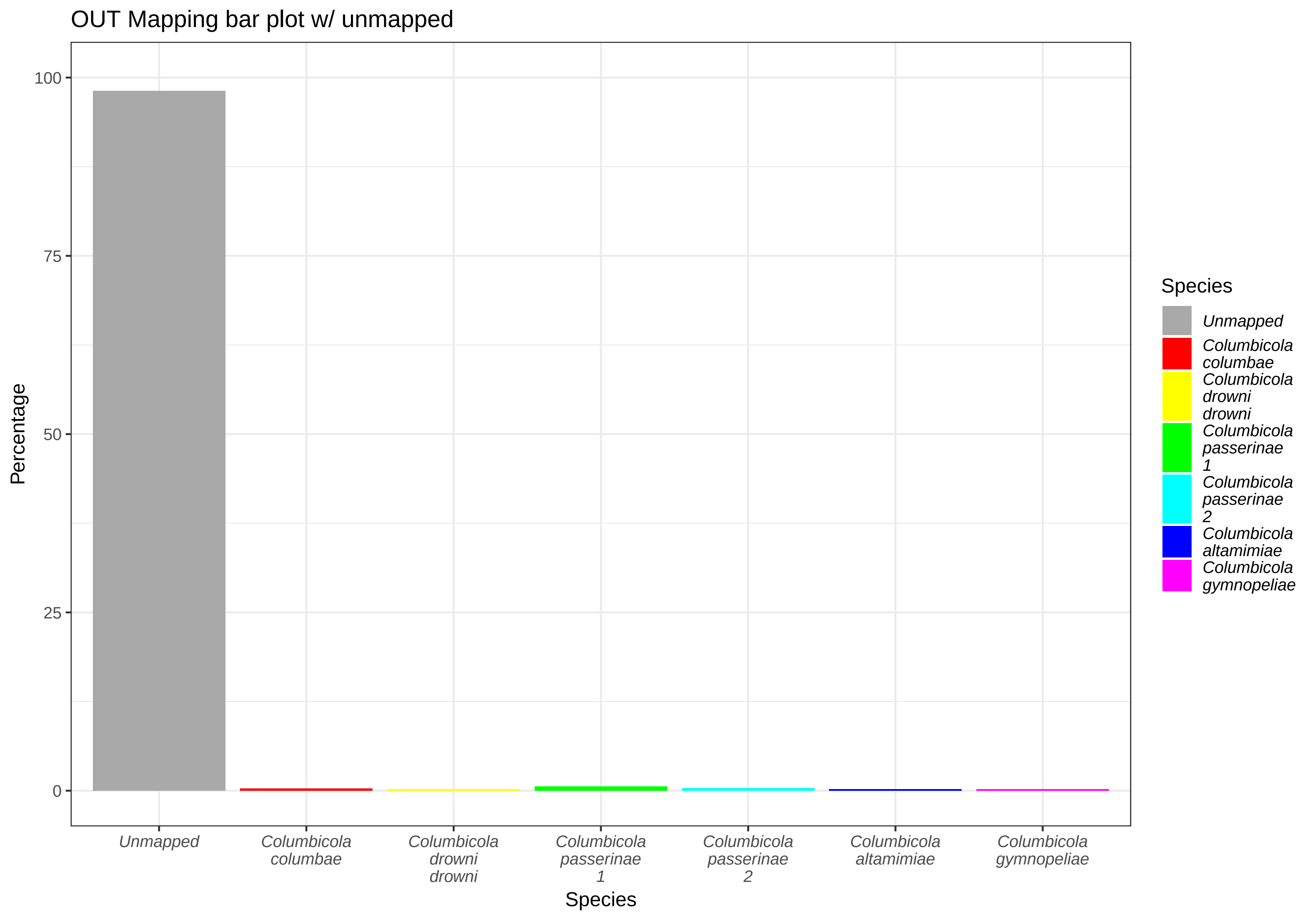

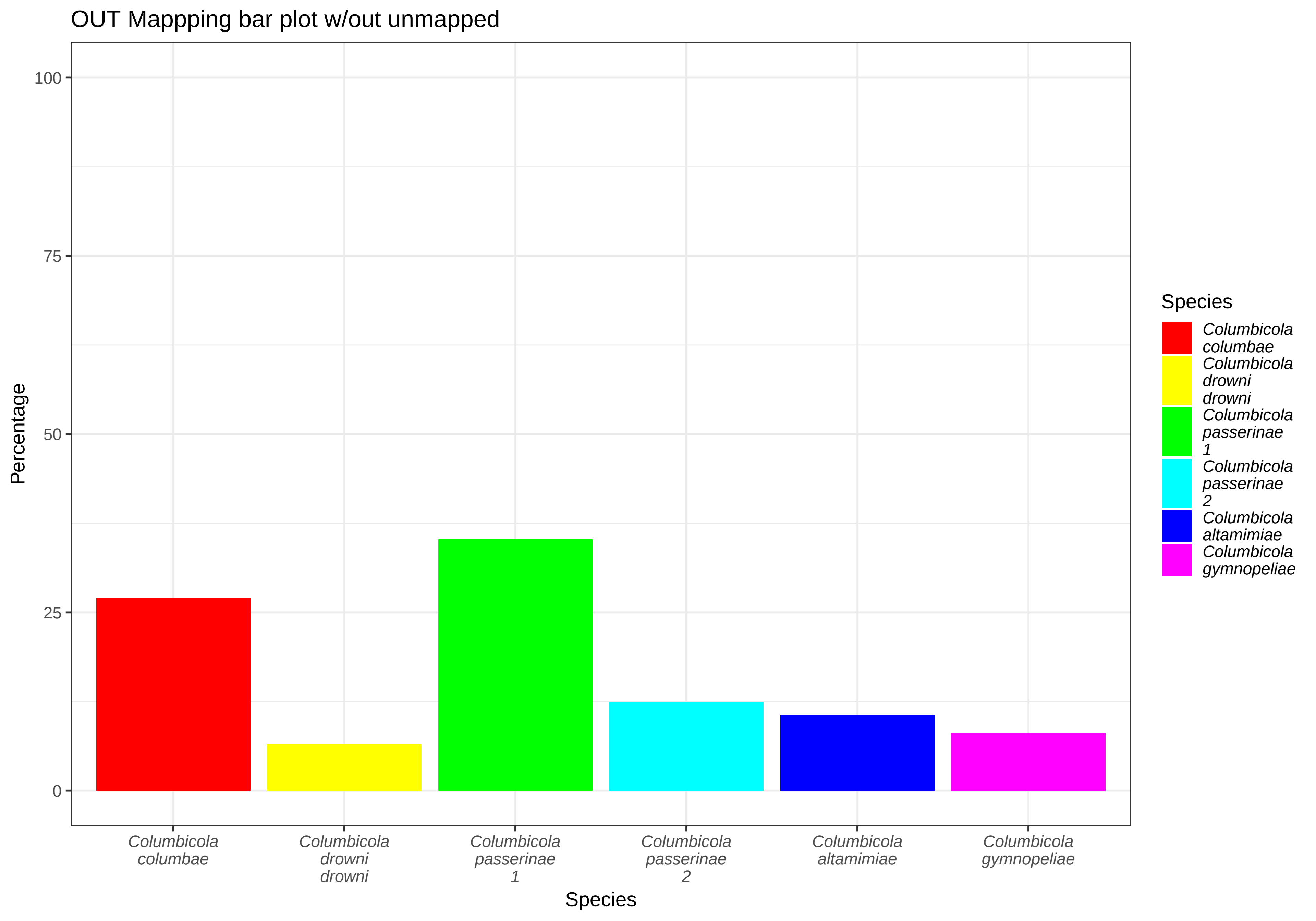

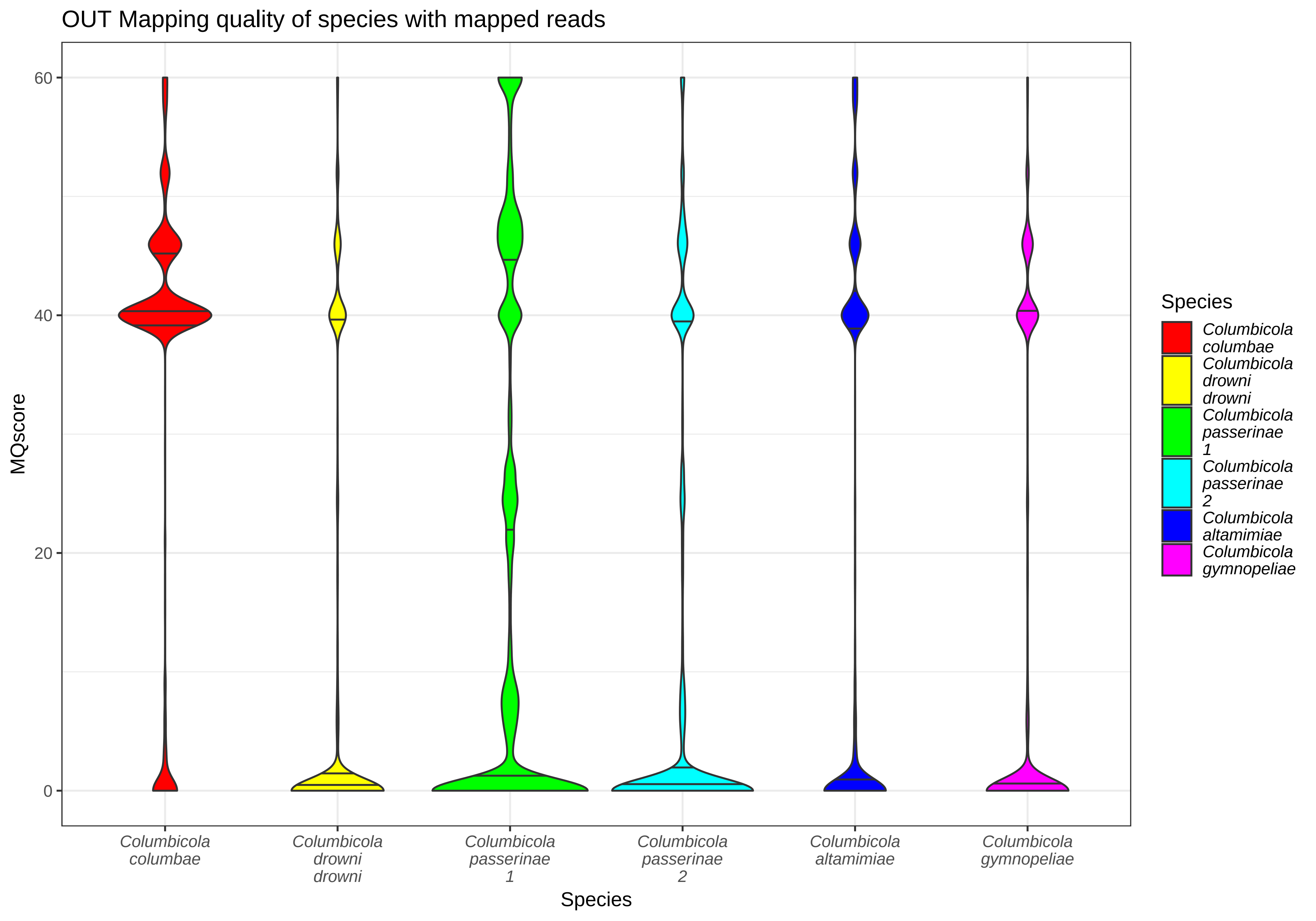

### OUT_plotMQ_Copas2.pdf

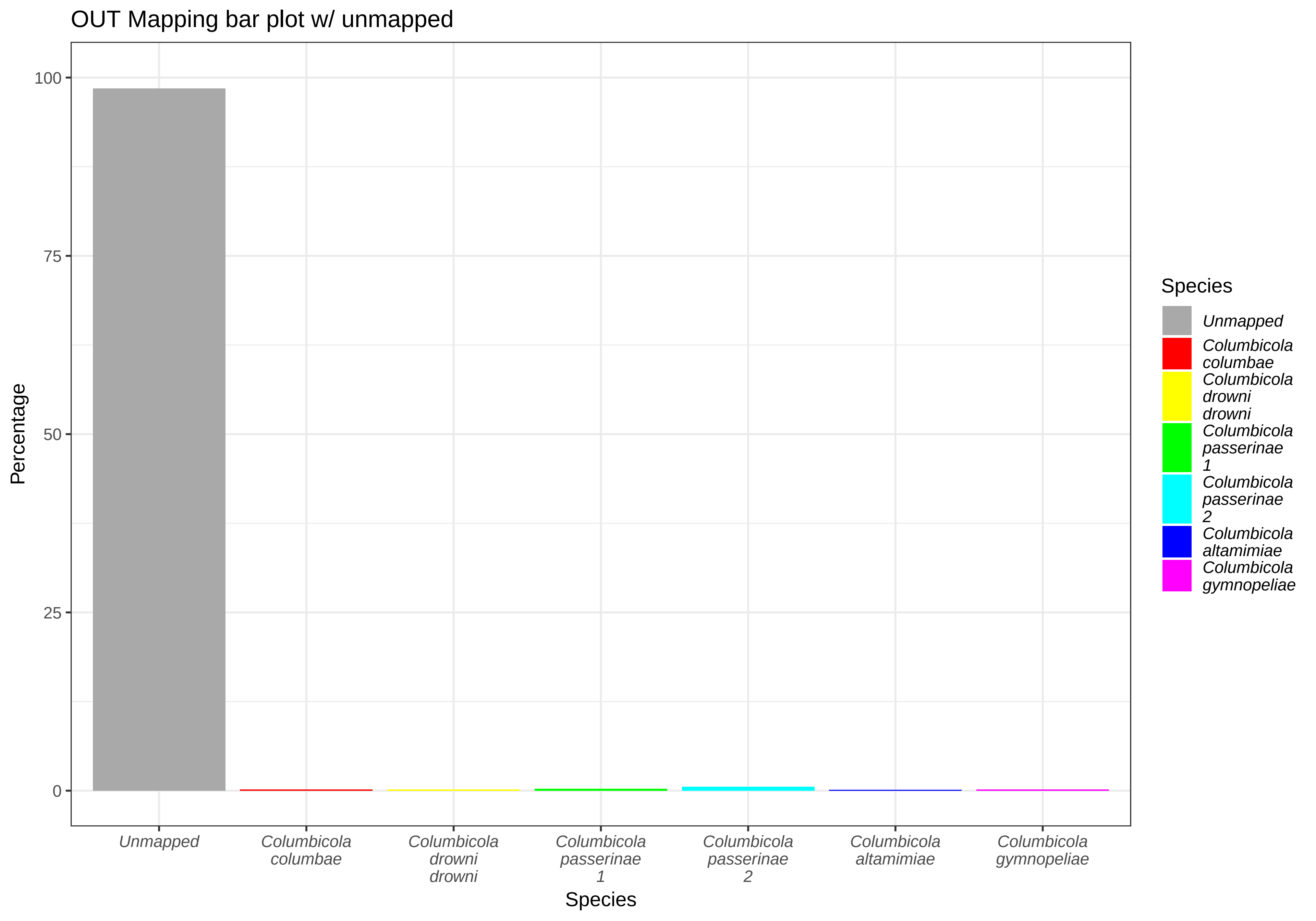

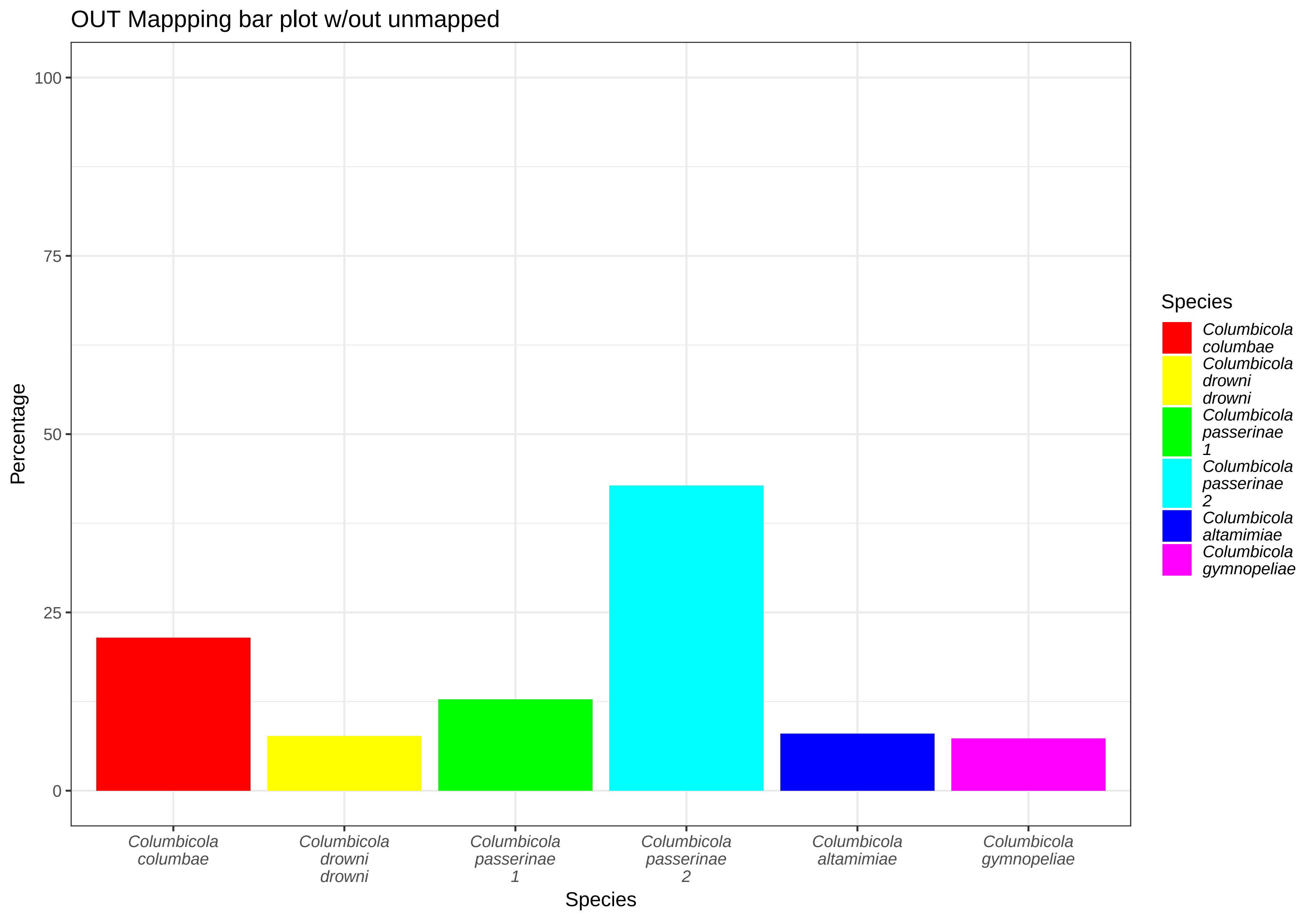

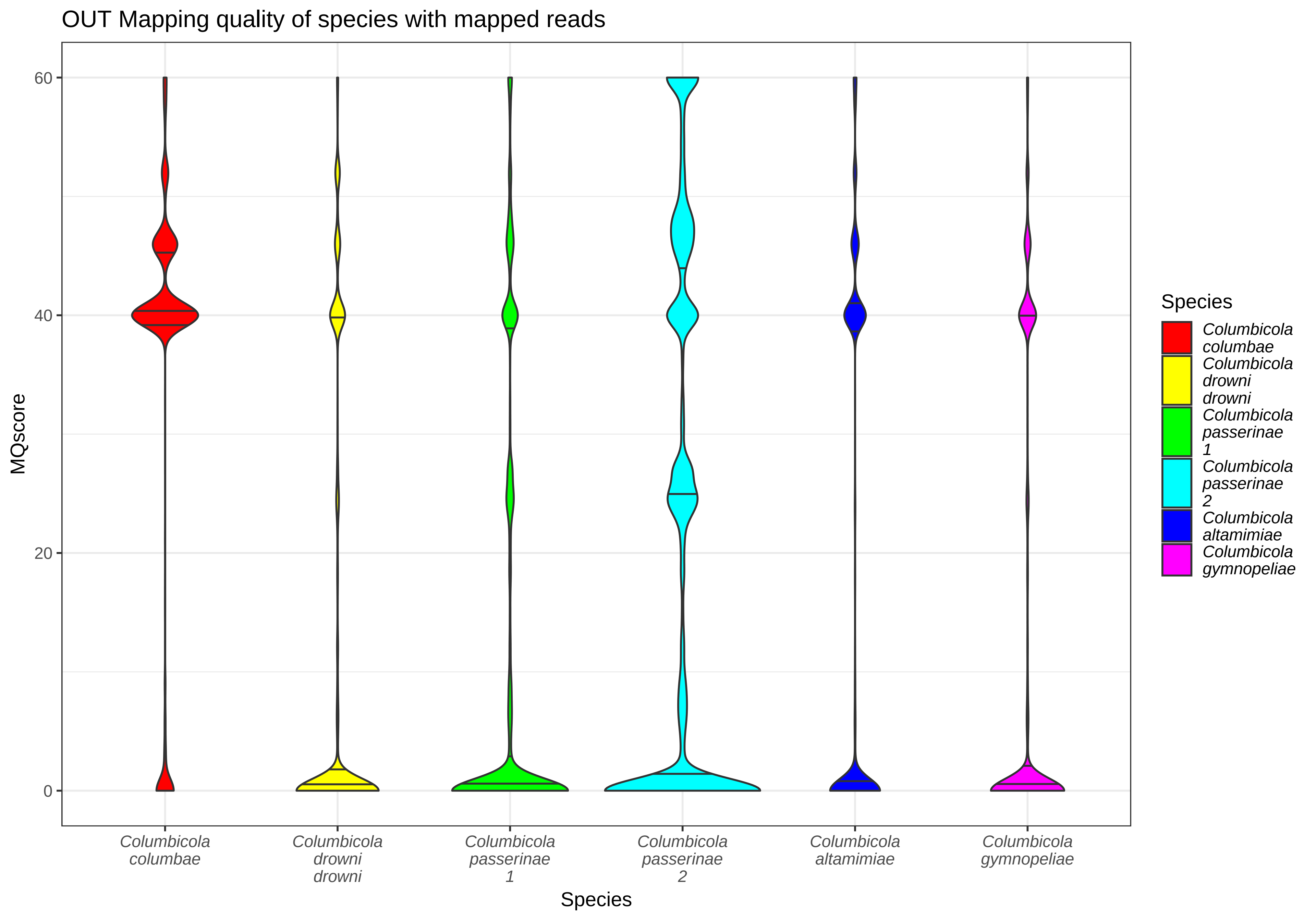

### OUT_plotMQ_Cosp_Clpre_11_2_2016_14.pdf

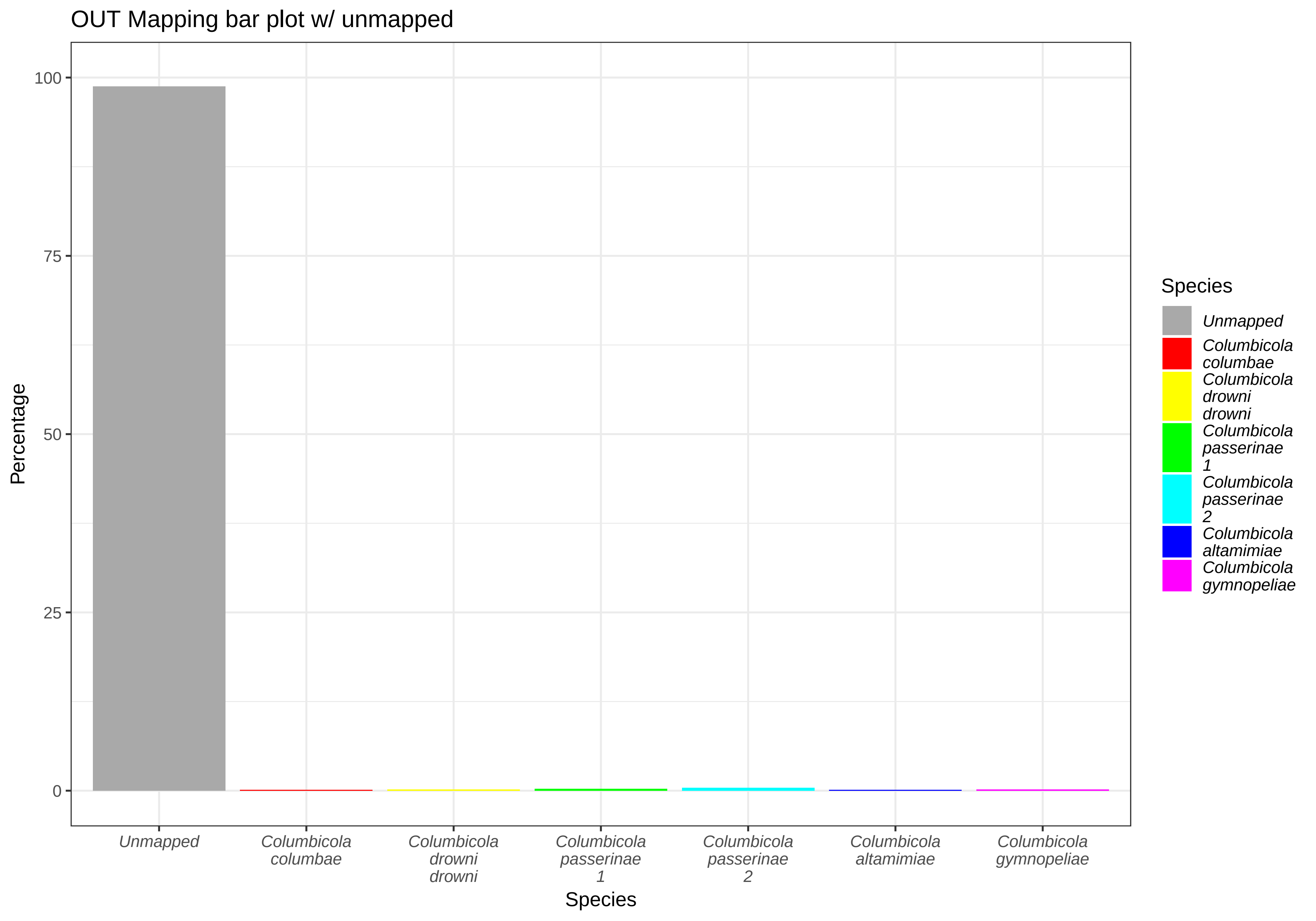

OUT Mappping bar plot w/out unmapped

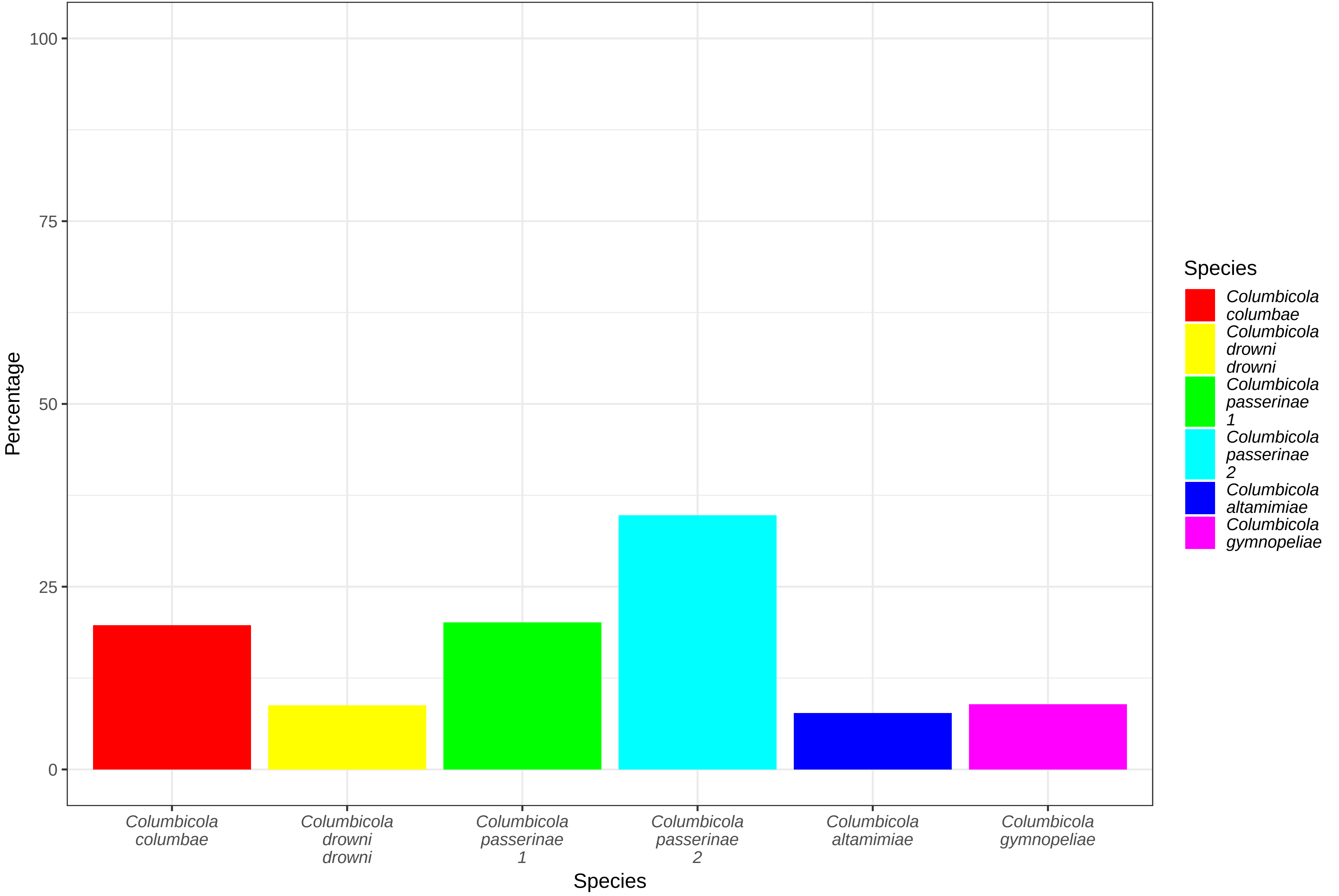

OUT Mapping quality of species with mapped reads

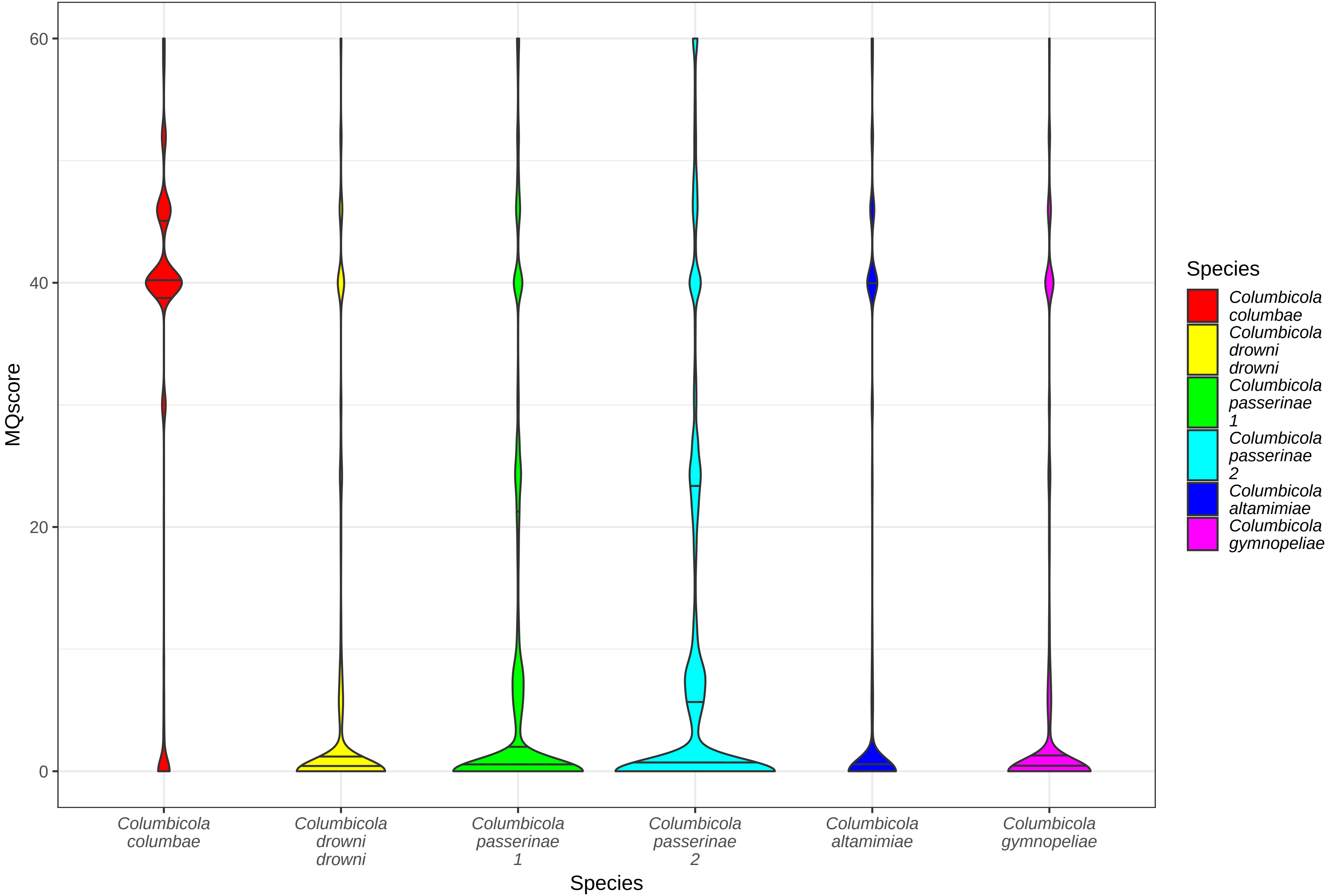

### OUT_plotMQ_Cosp_Clpre_11_7_2016_3.pdf

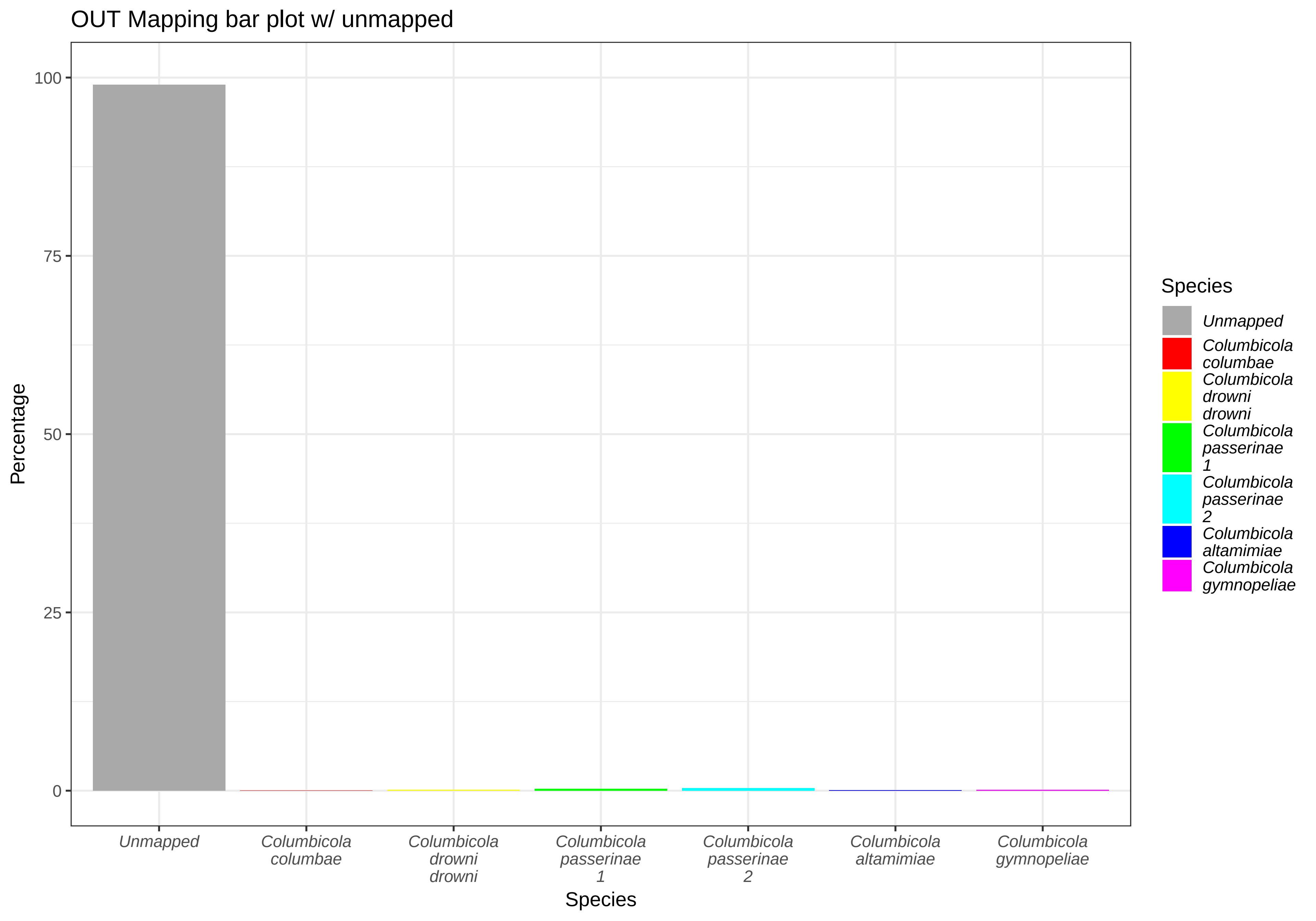

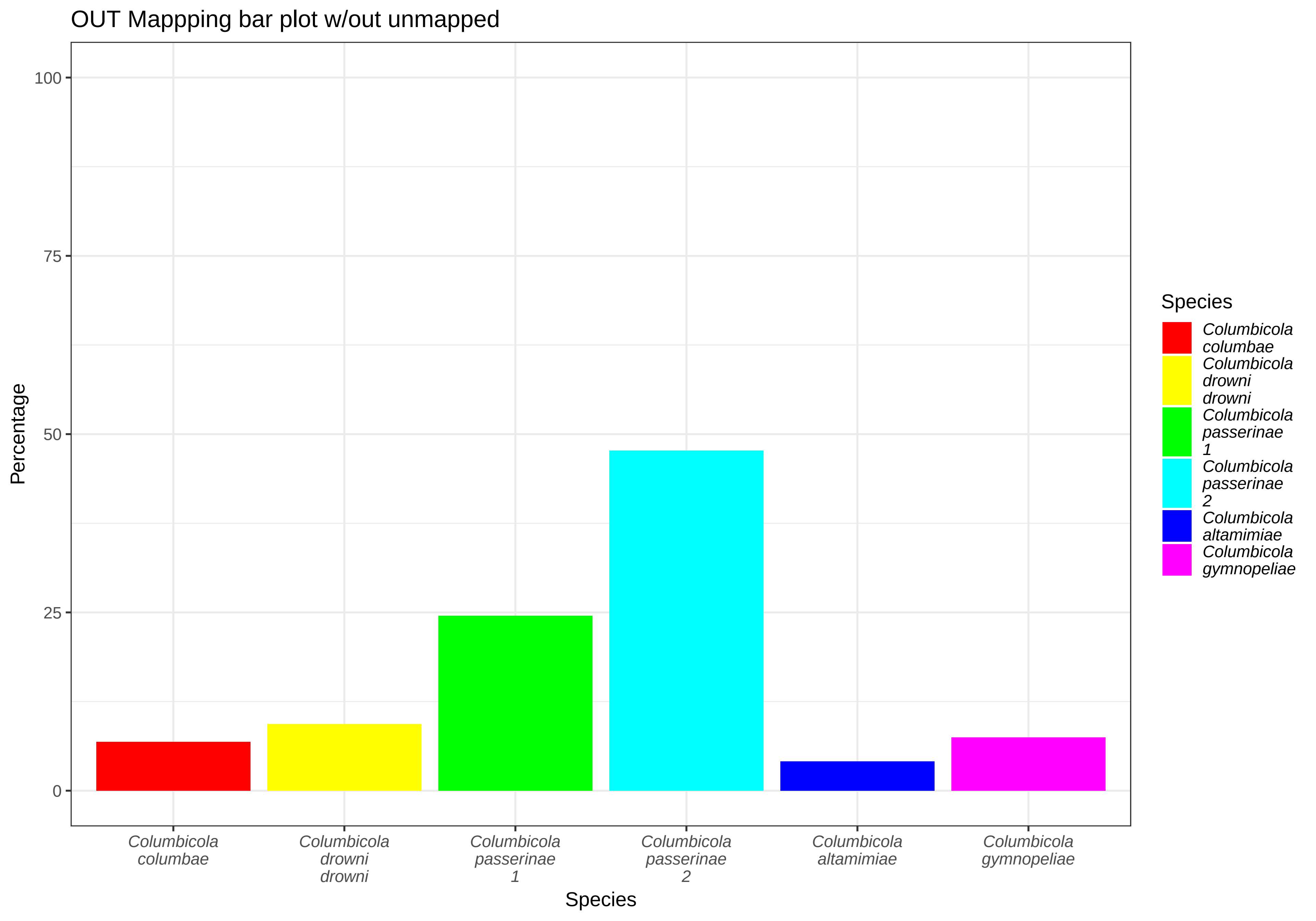

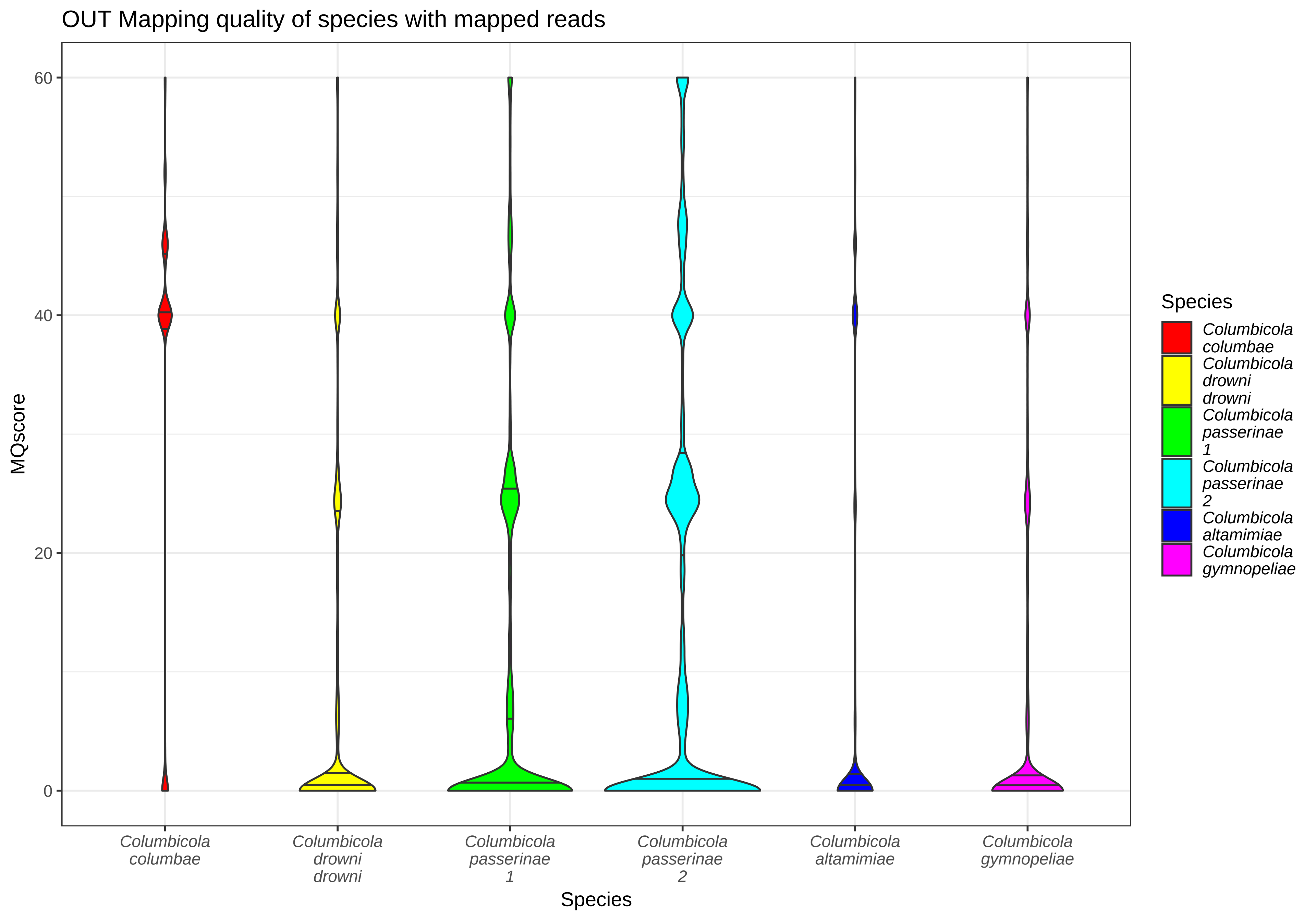

### OUT_plotMQ_Cosp_Clpre_11_7_2016_9.pdf

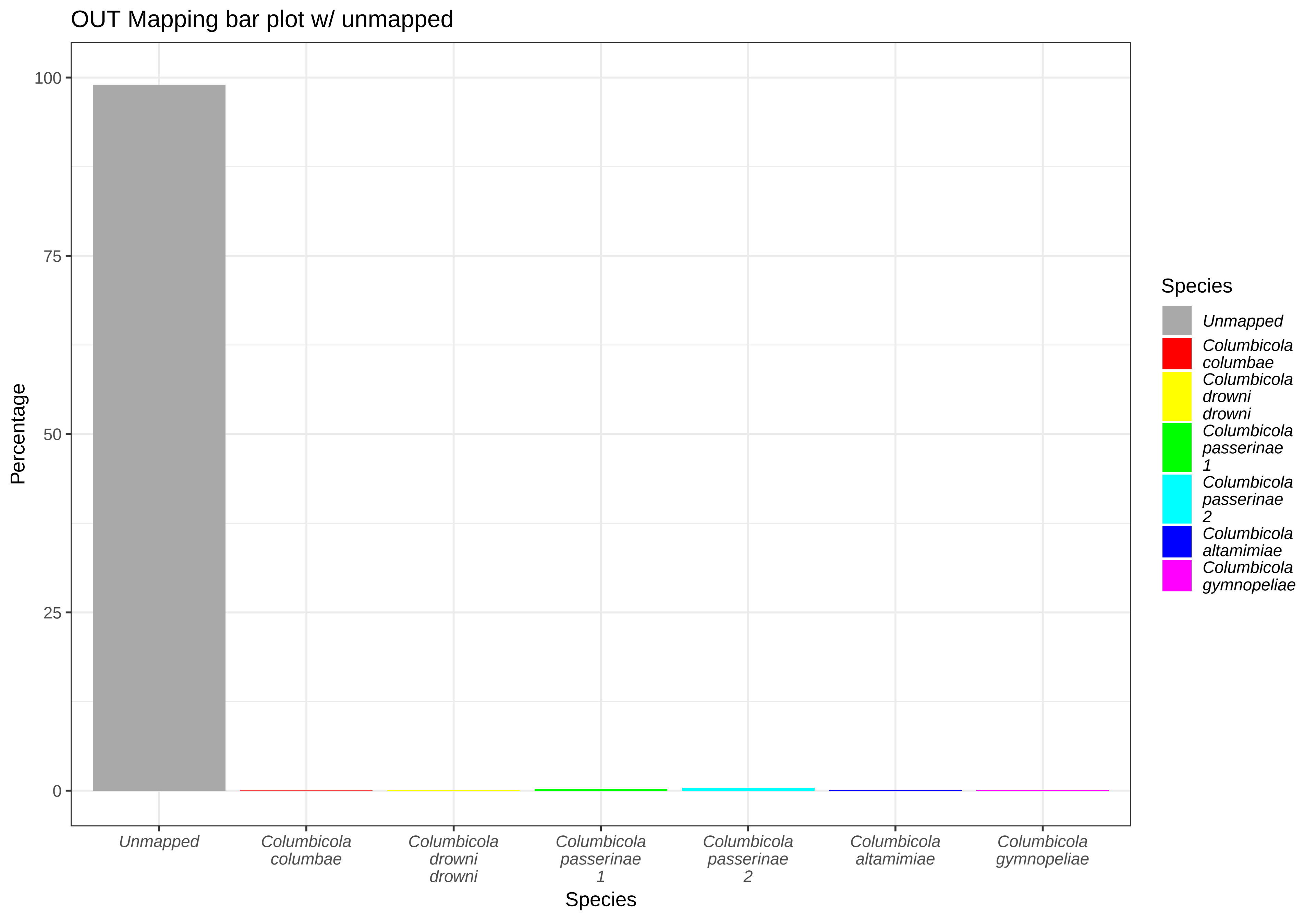

OUT Mappping bar plot w/out unmapped

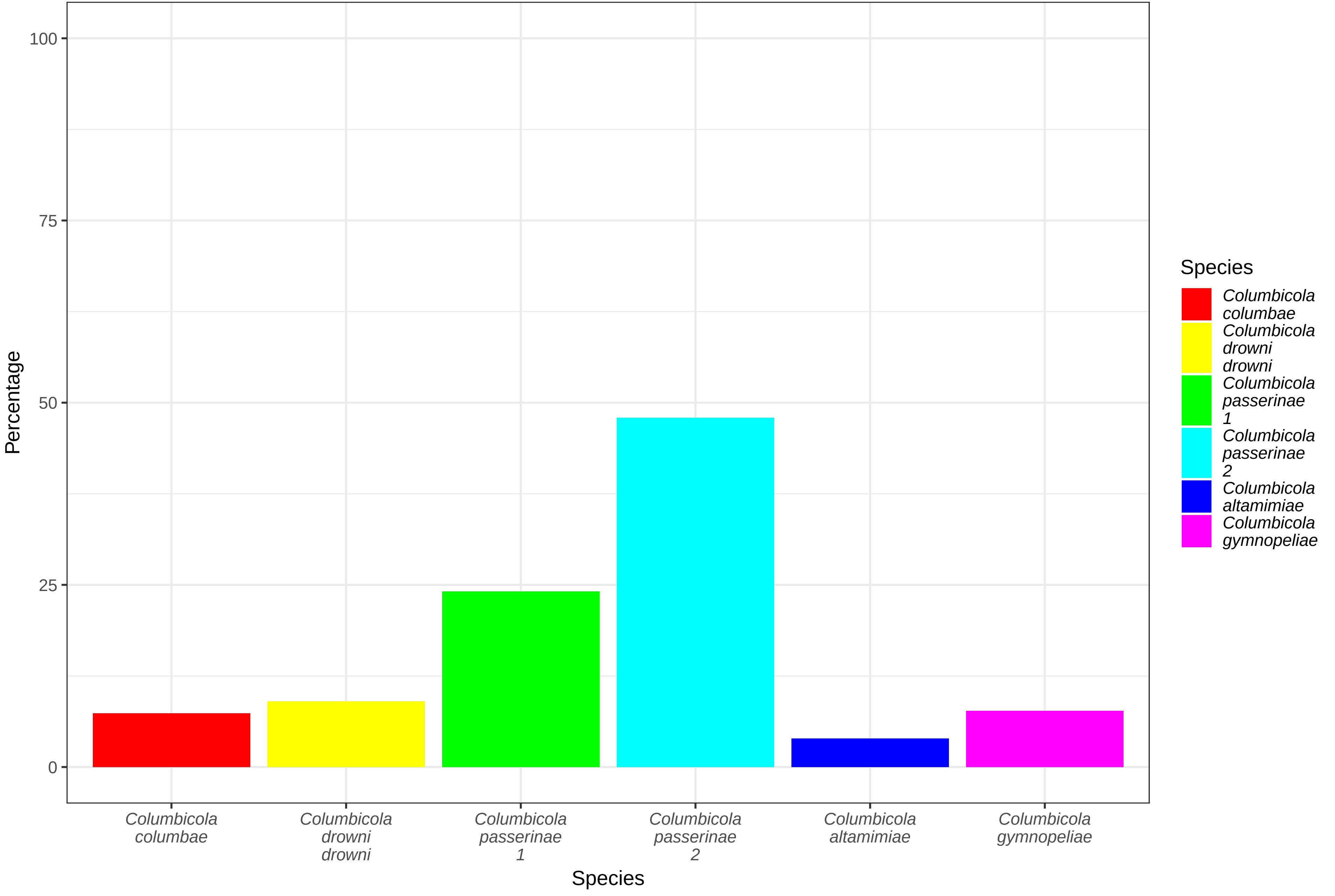

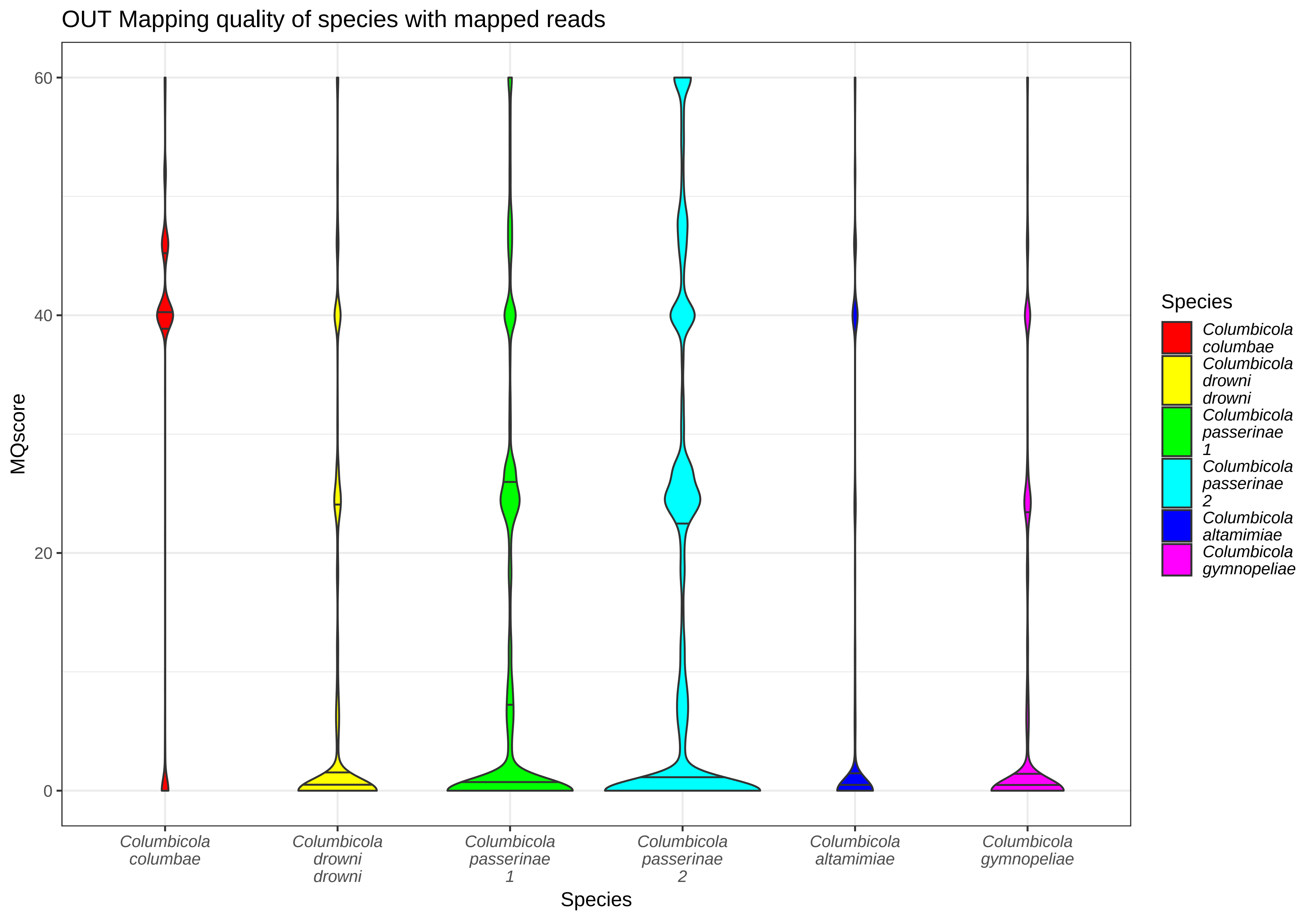

### OUT_plotMQ_Cosp_Clpre_11_7_2016_11.pdf

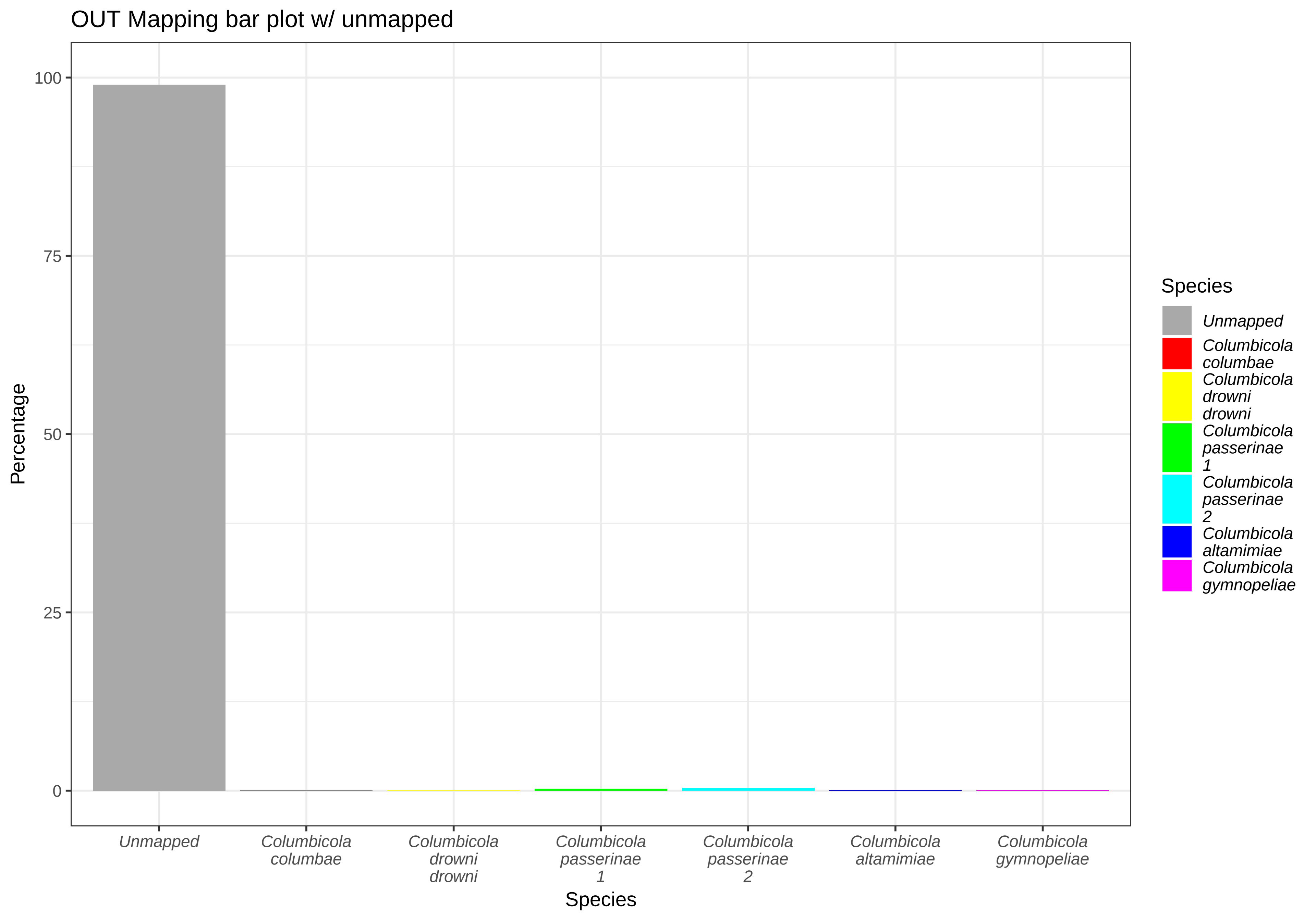

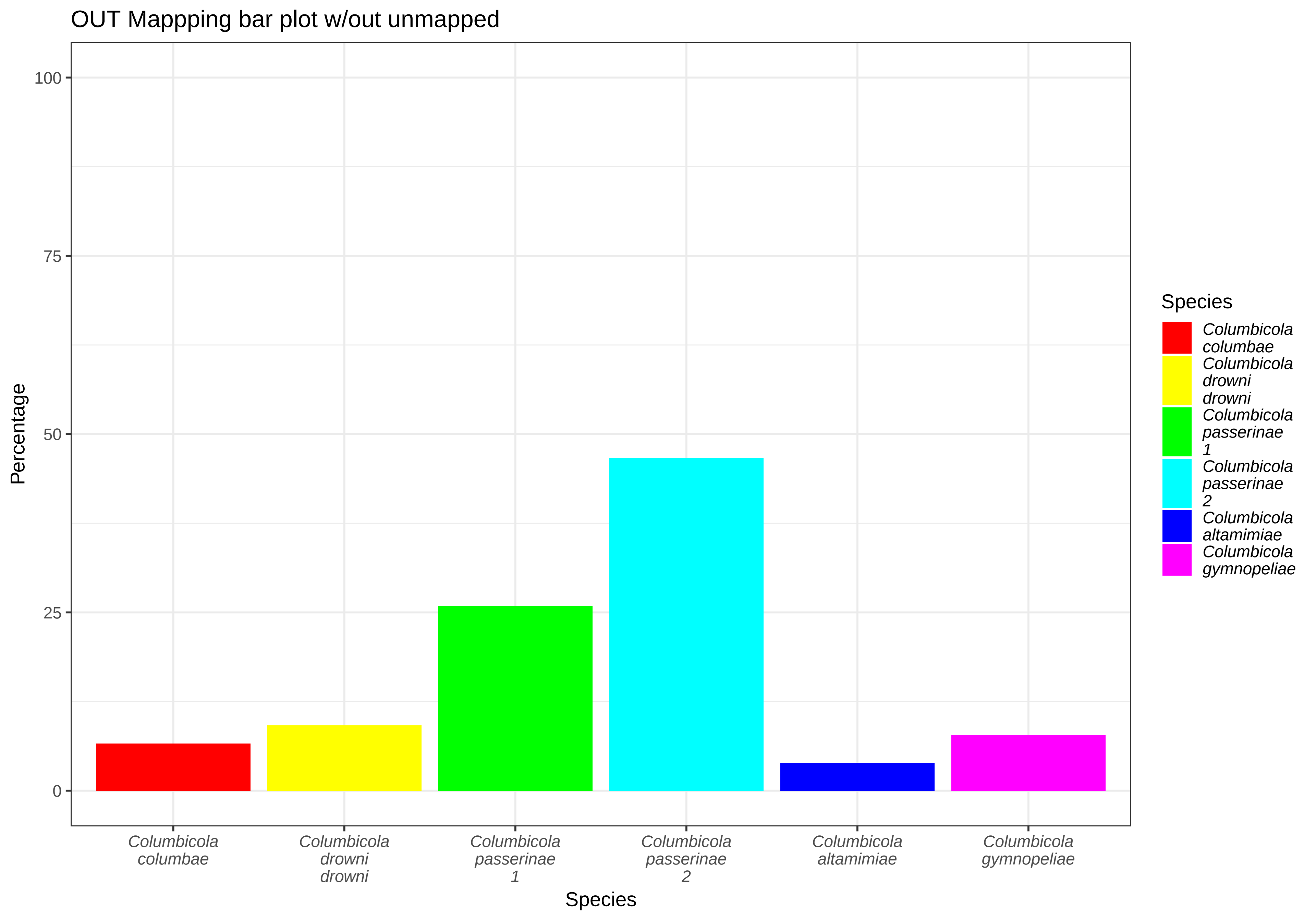

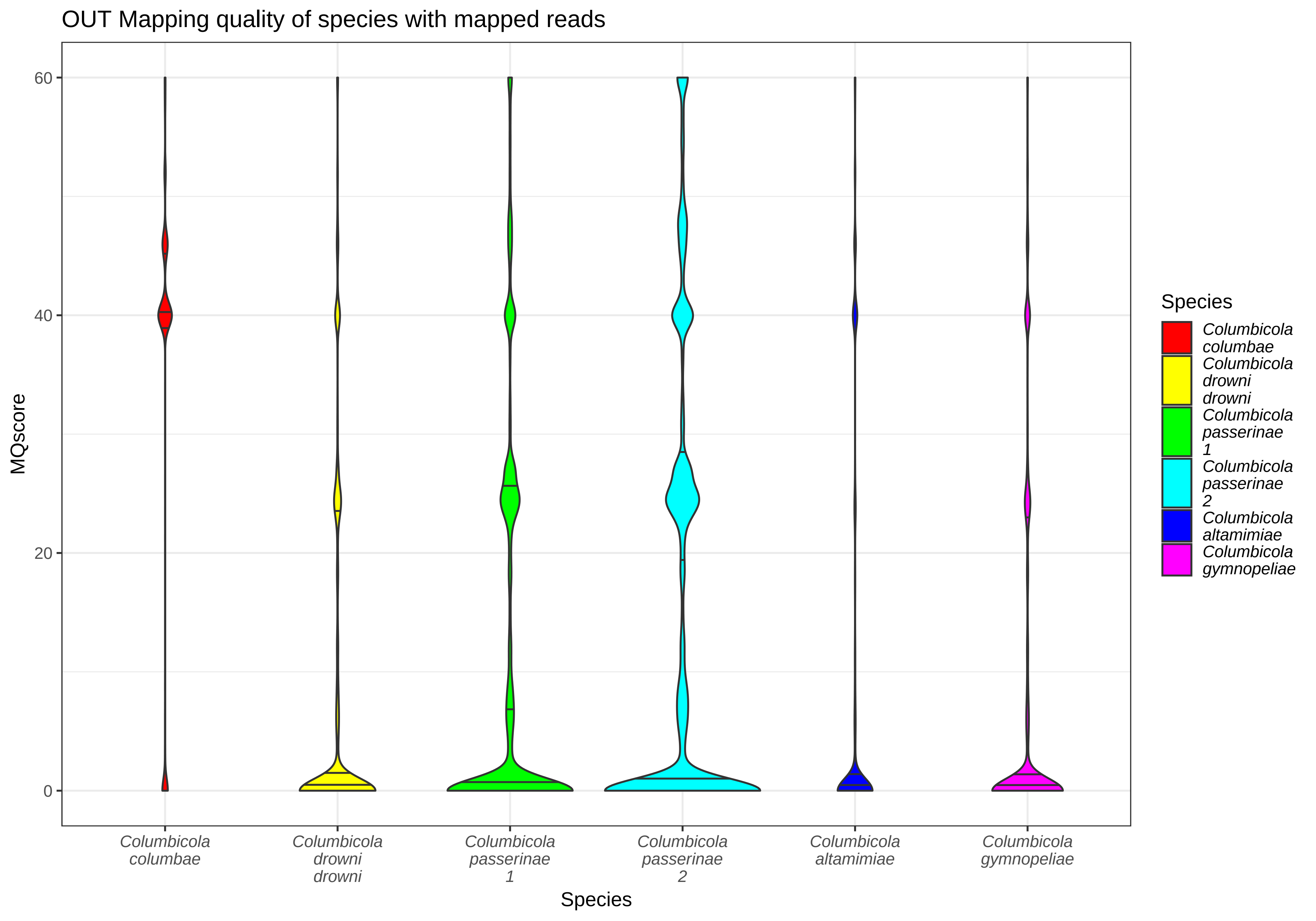

### OUT_plotMQ_Cosp_Cobuc_11_2_2016_2.pdf

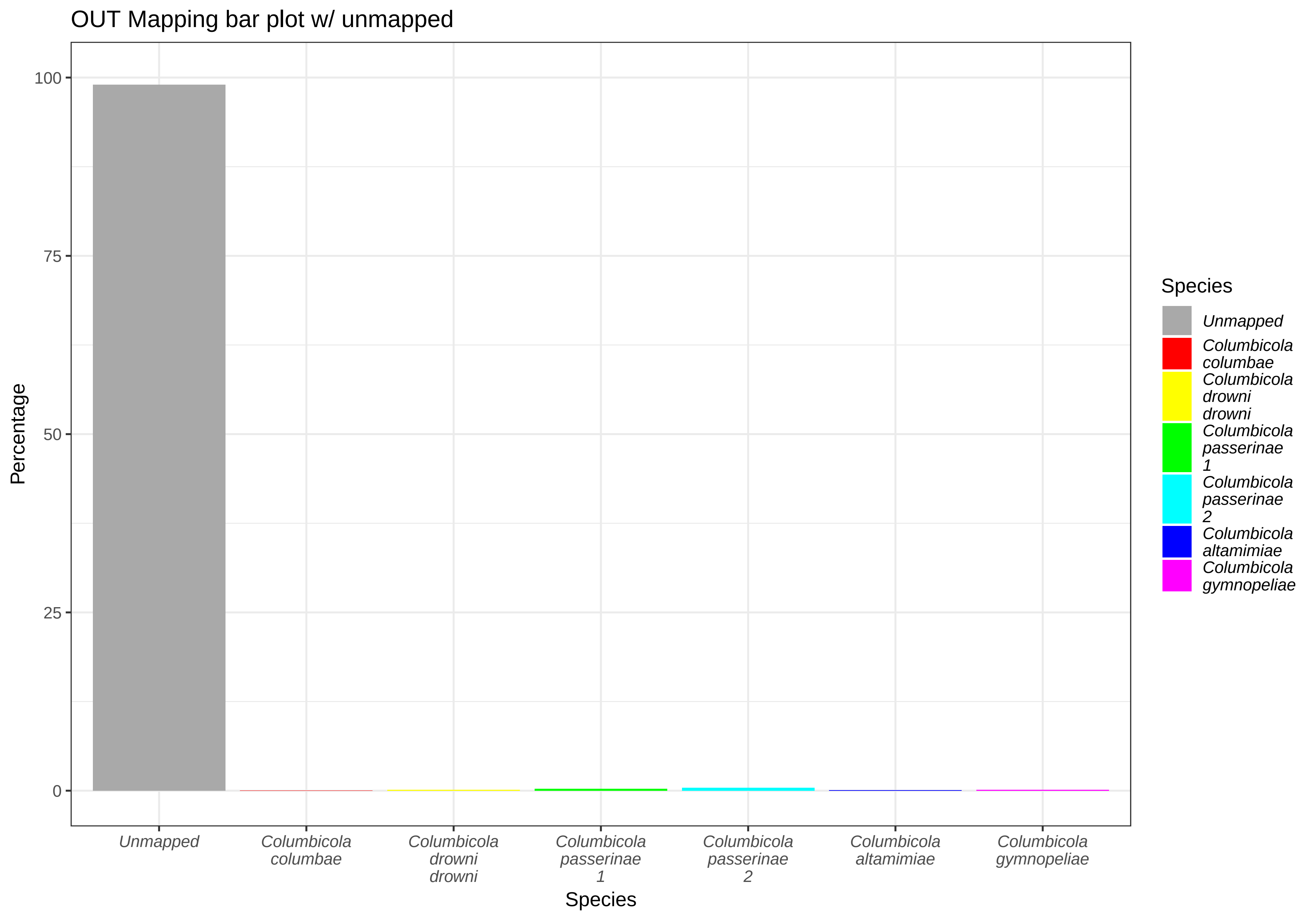

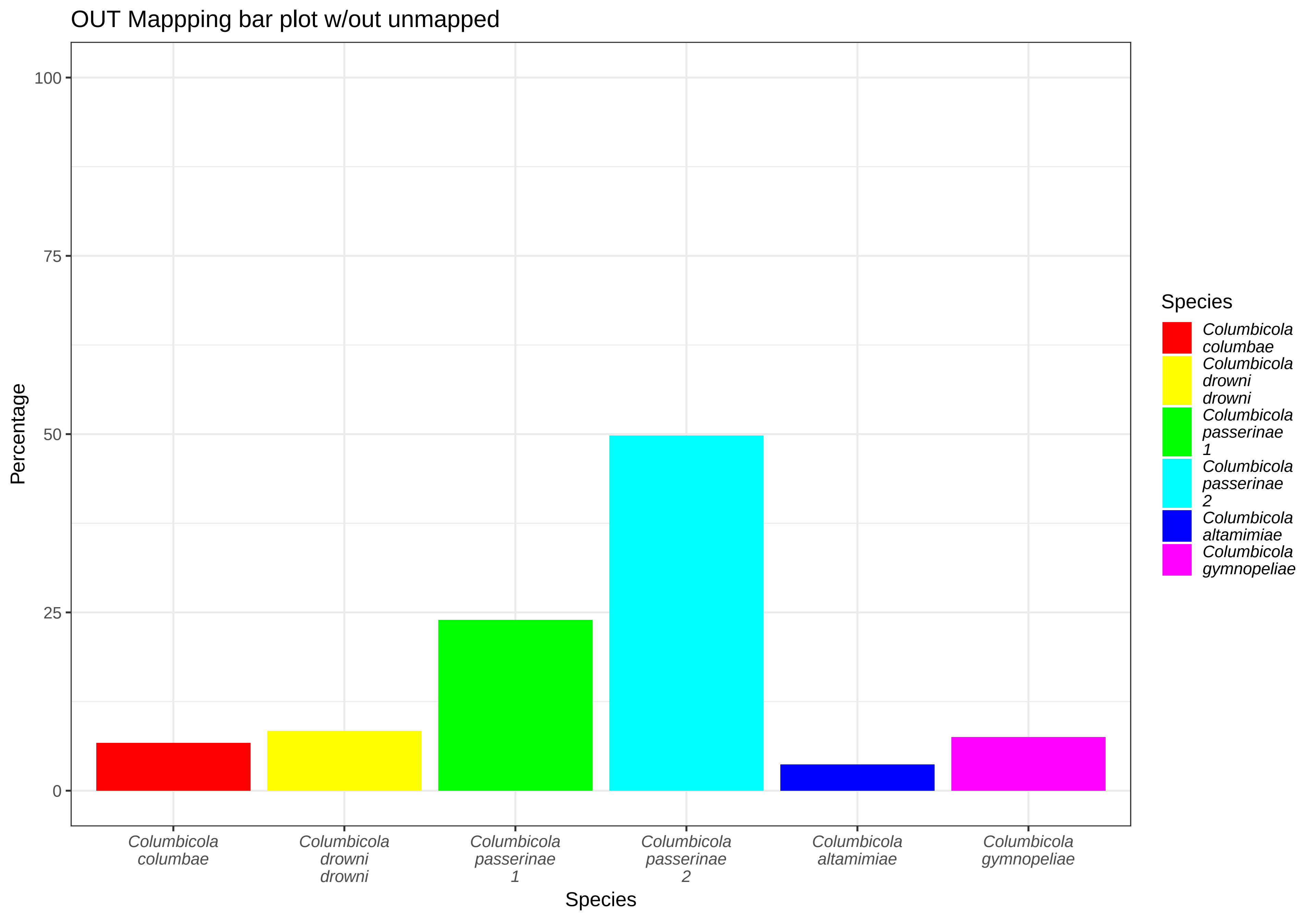

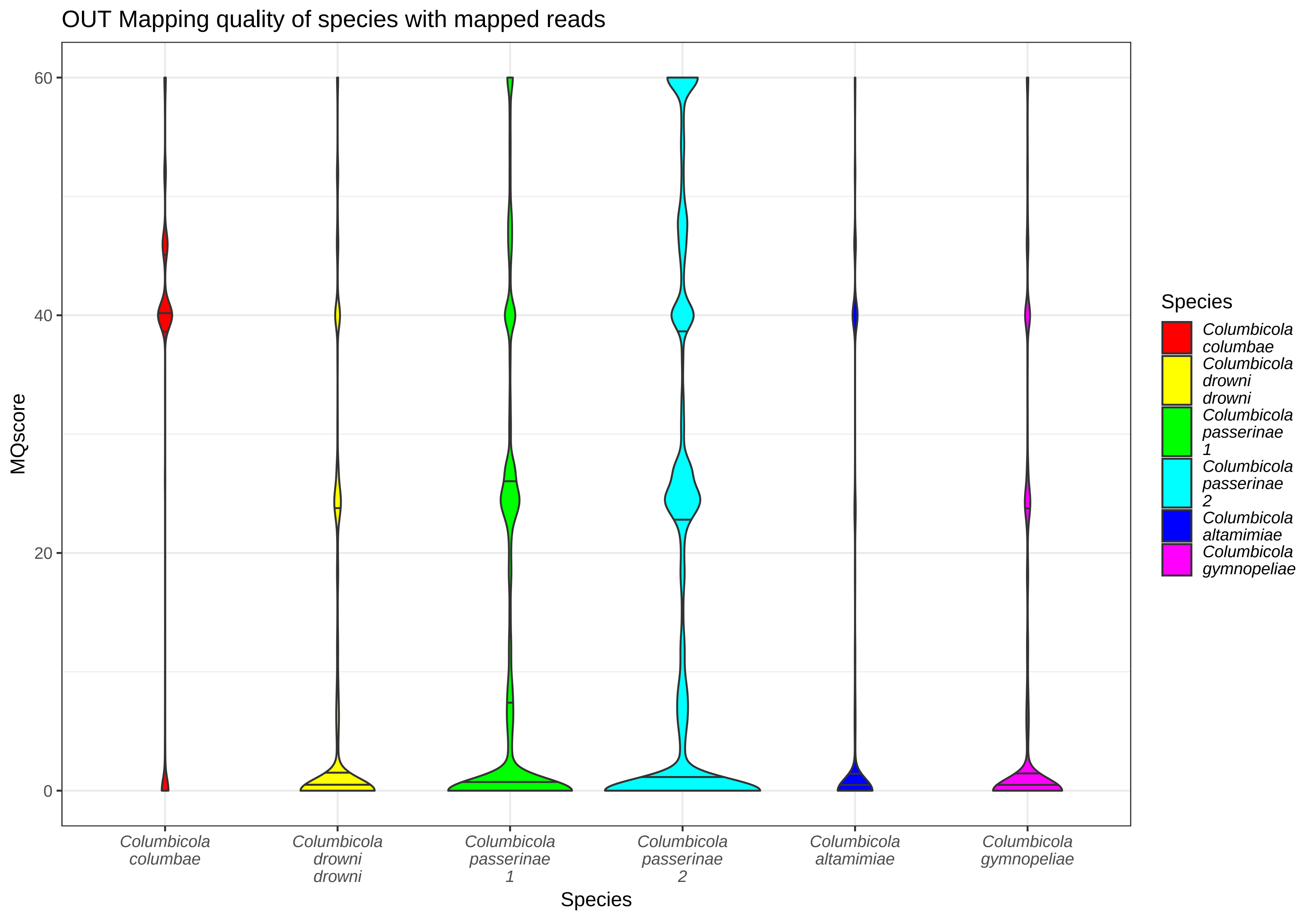

### OUT_plotMQ_Cosp_Cobuc_11_2_2016_13.pdf

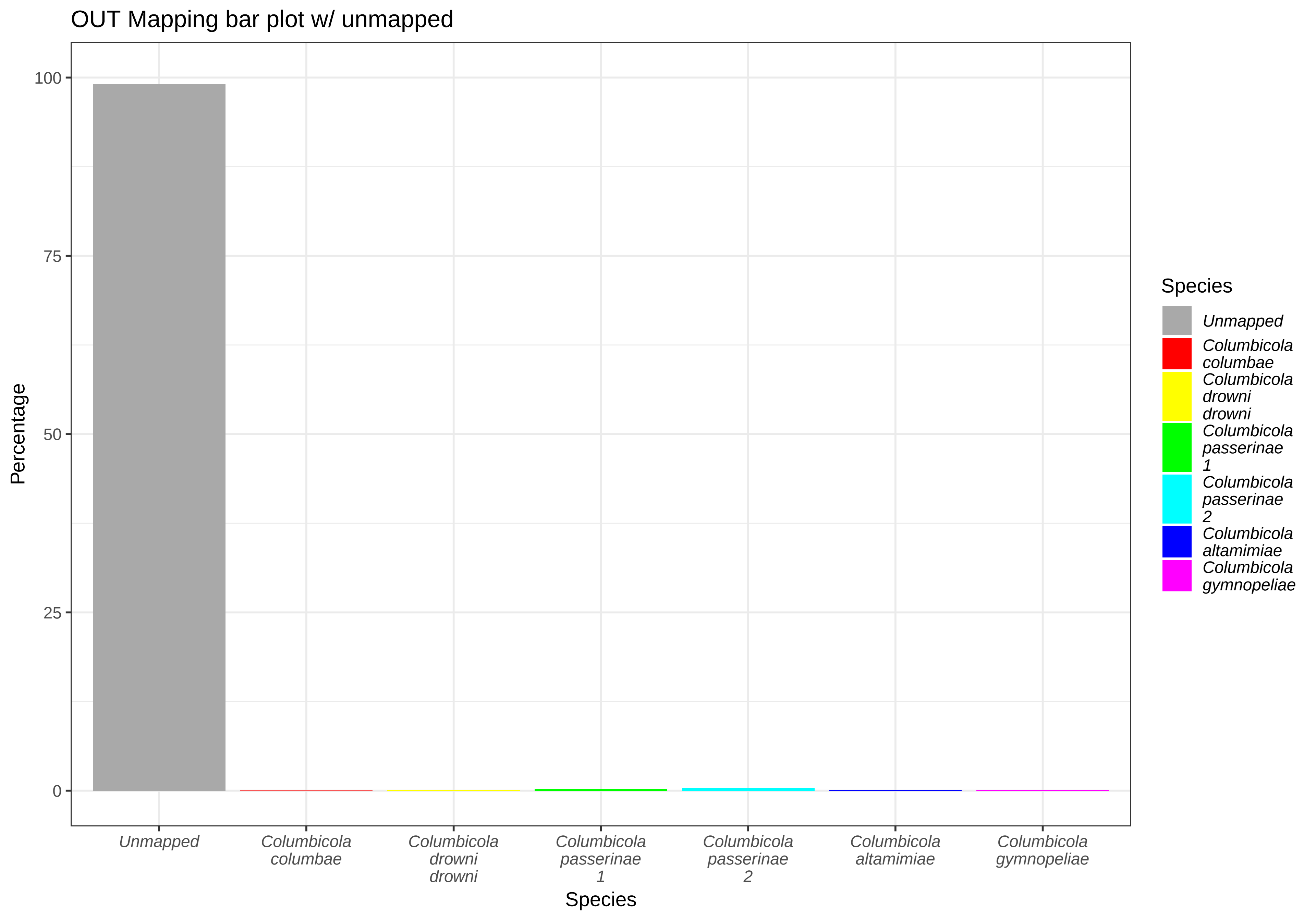

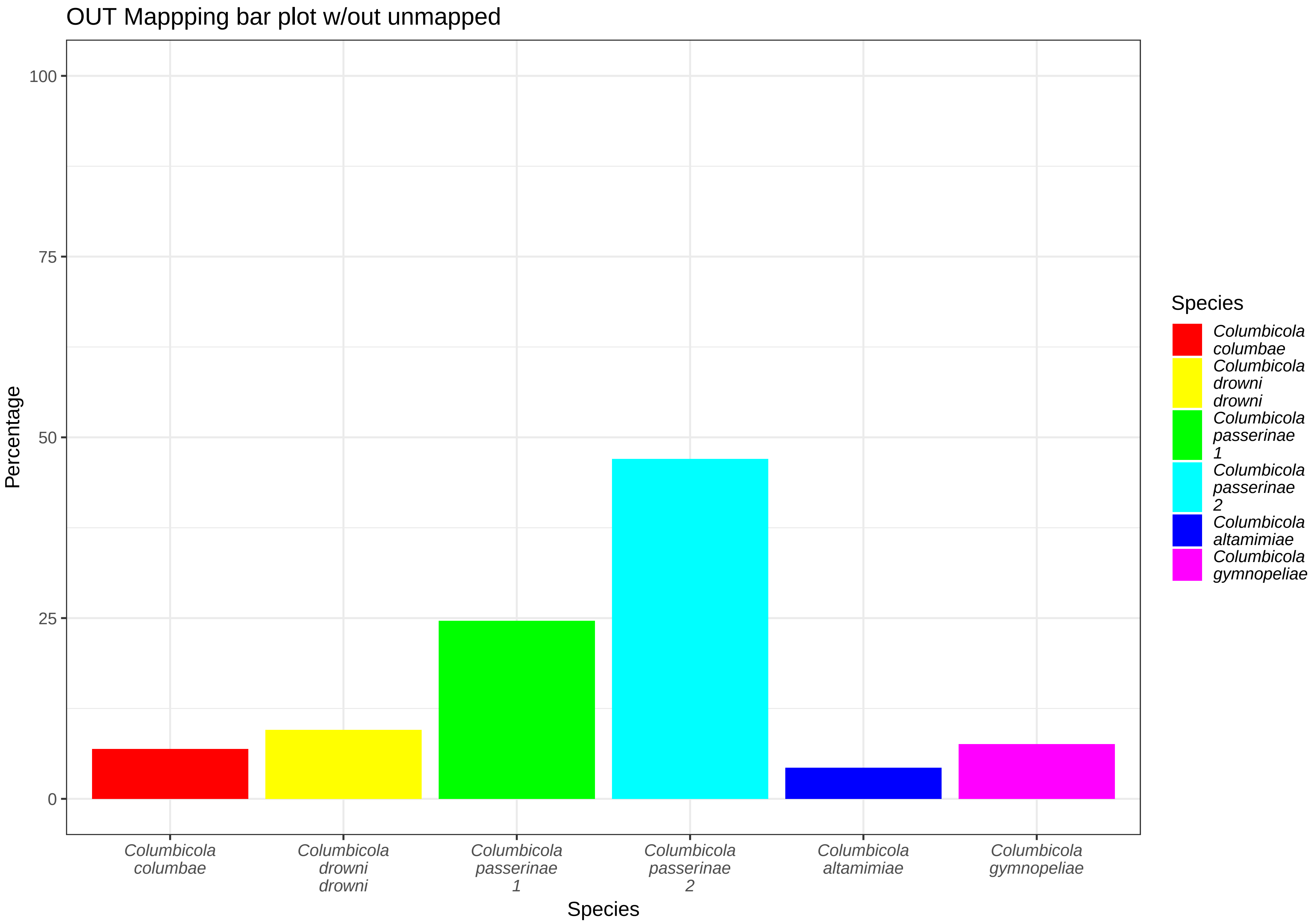

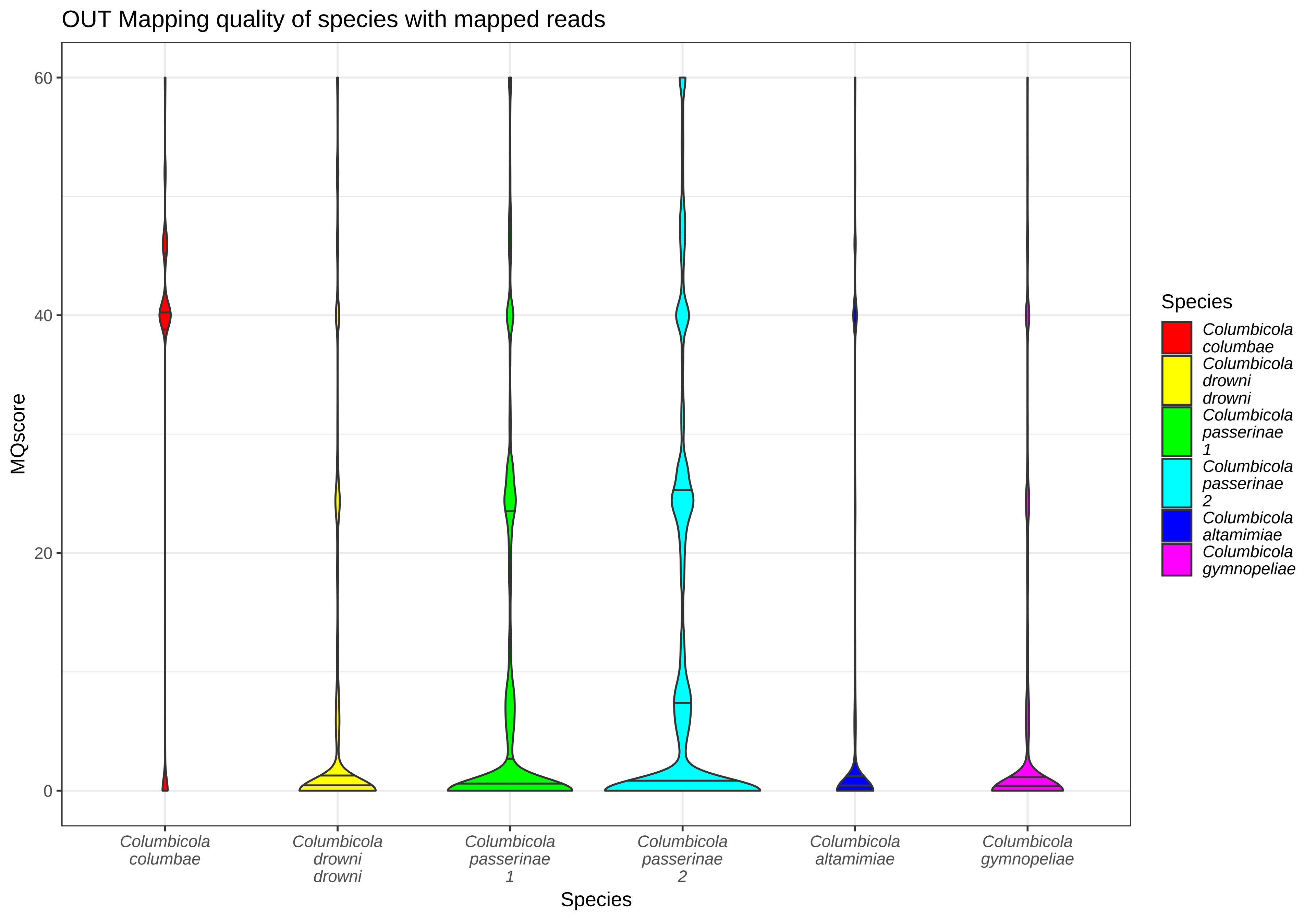

### OUT_plotMQ_Cosp_Cocru_11_2_2016_11.pdf

OUT Mapping bar plot w/ unmapped

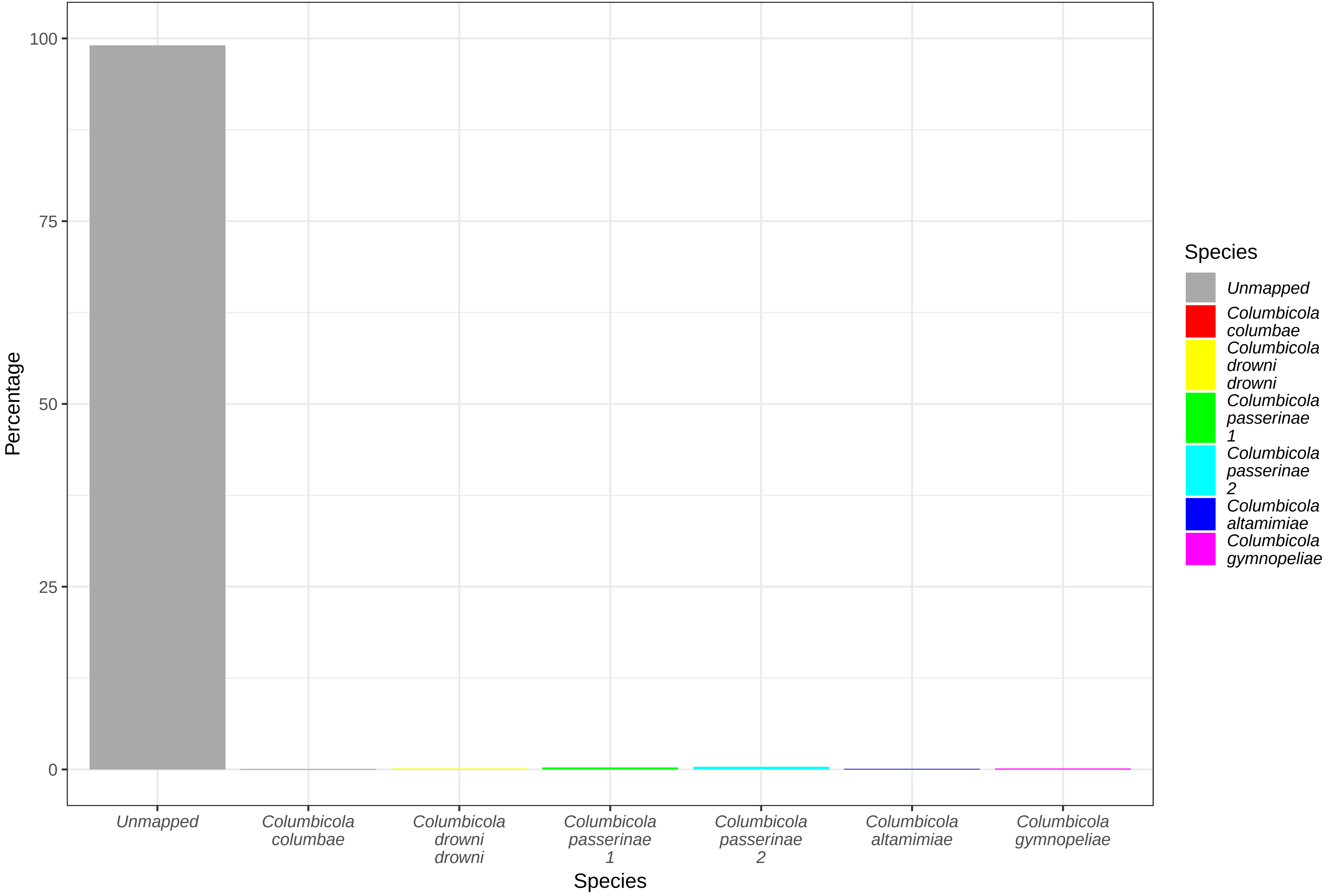

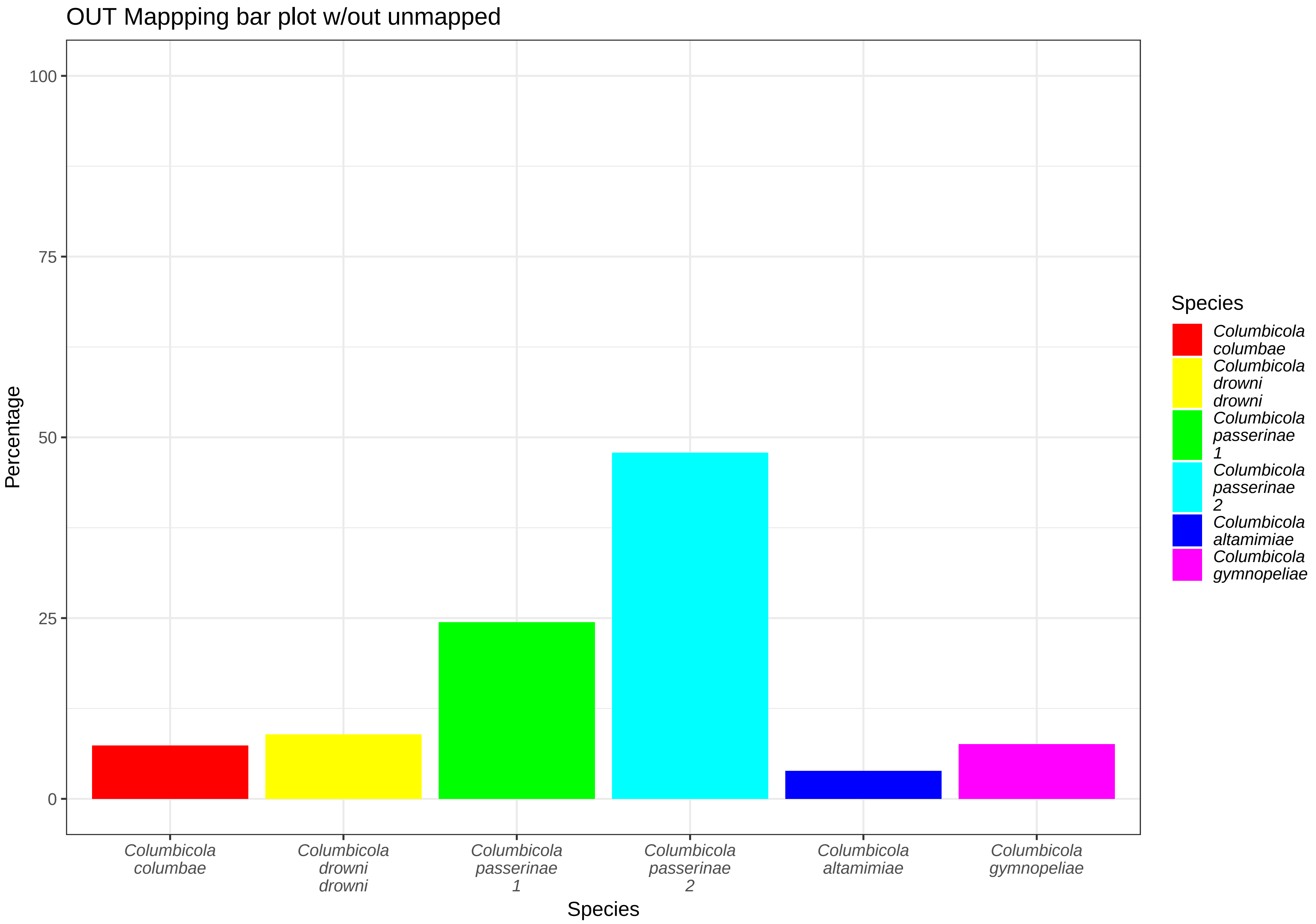

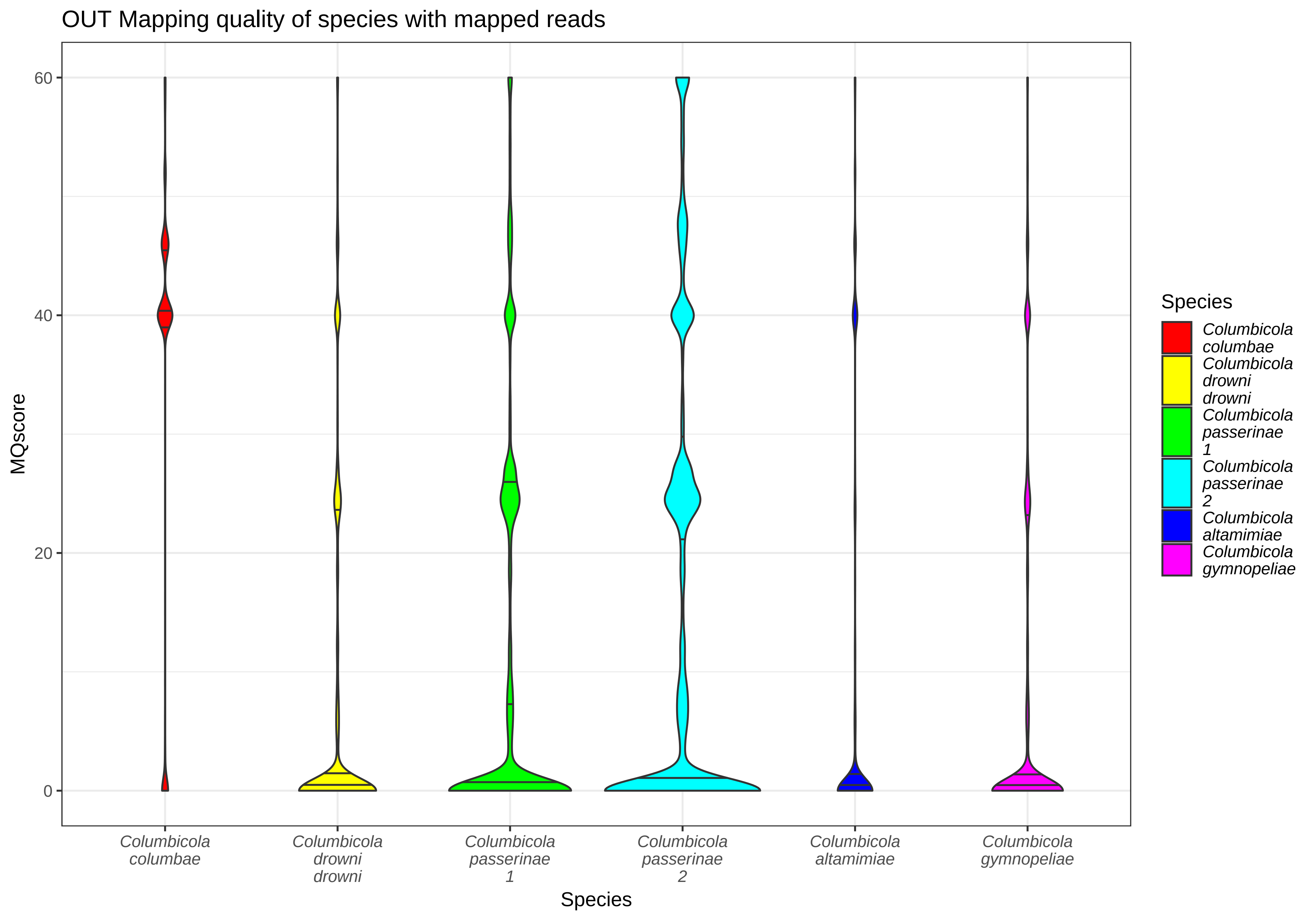

### OUT_plotMQ_Cosp_Cocru_11_7_2016_16.pdf

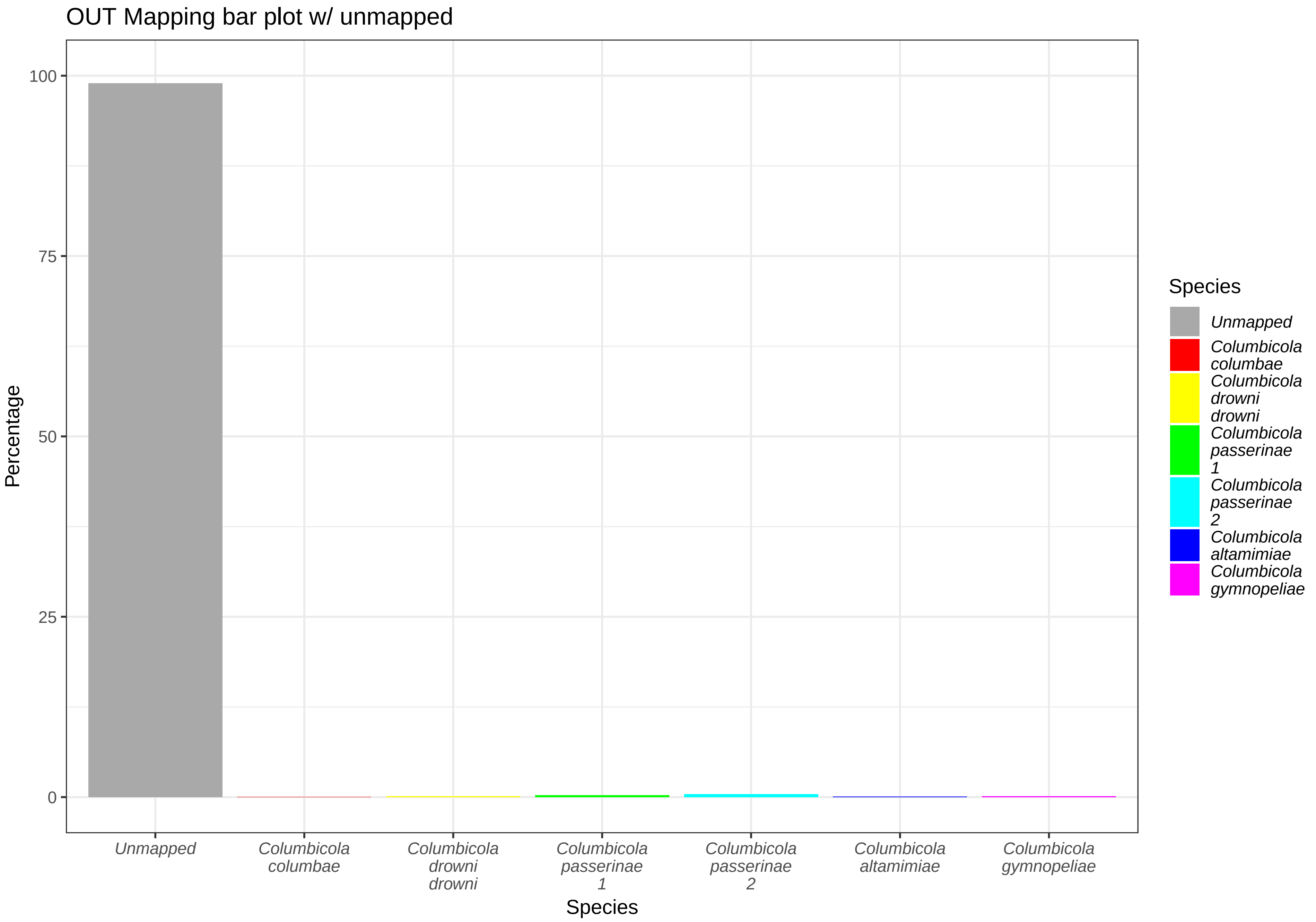

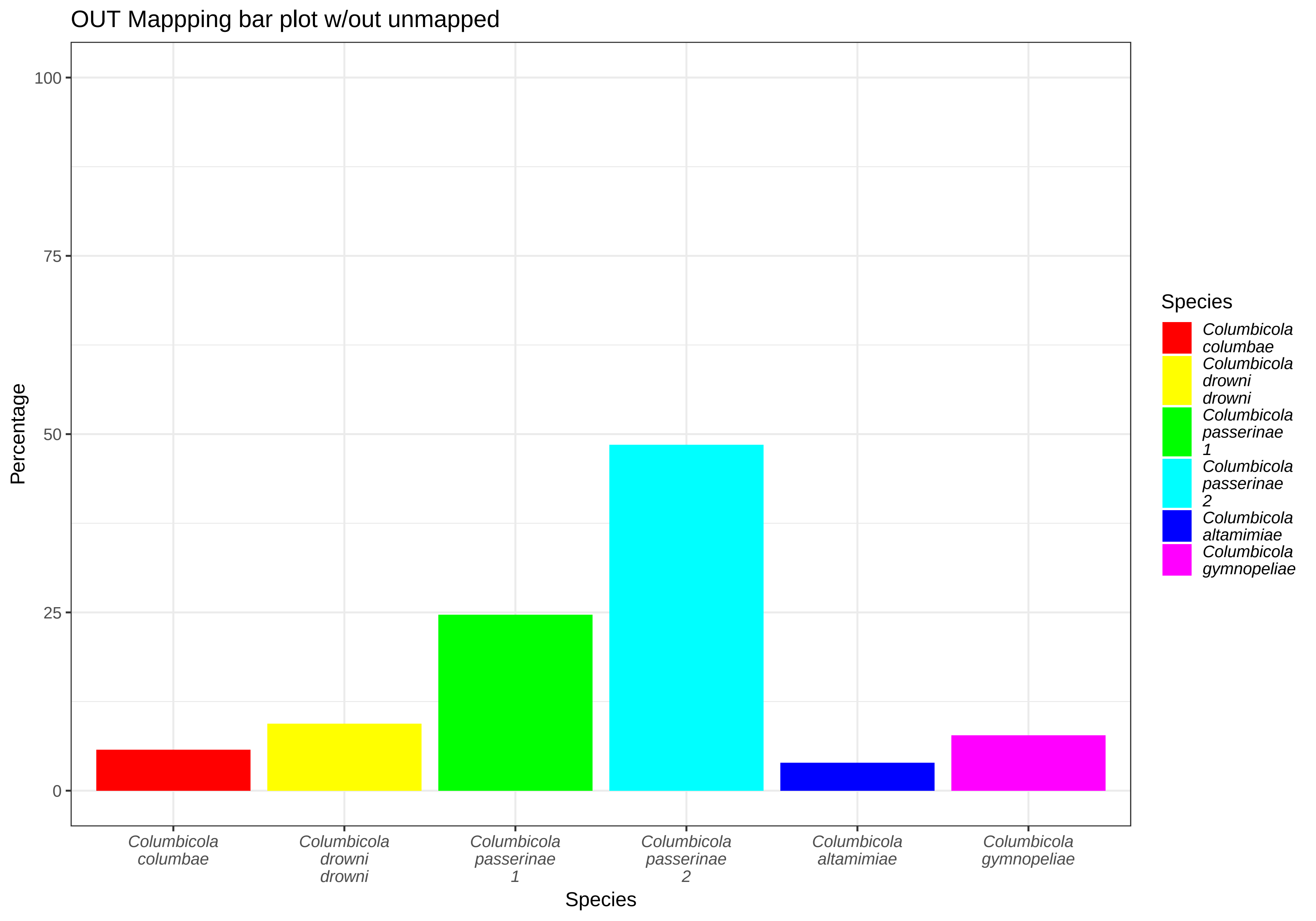

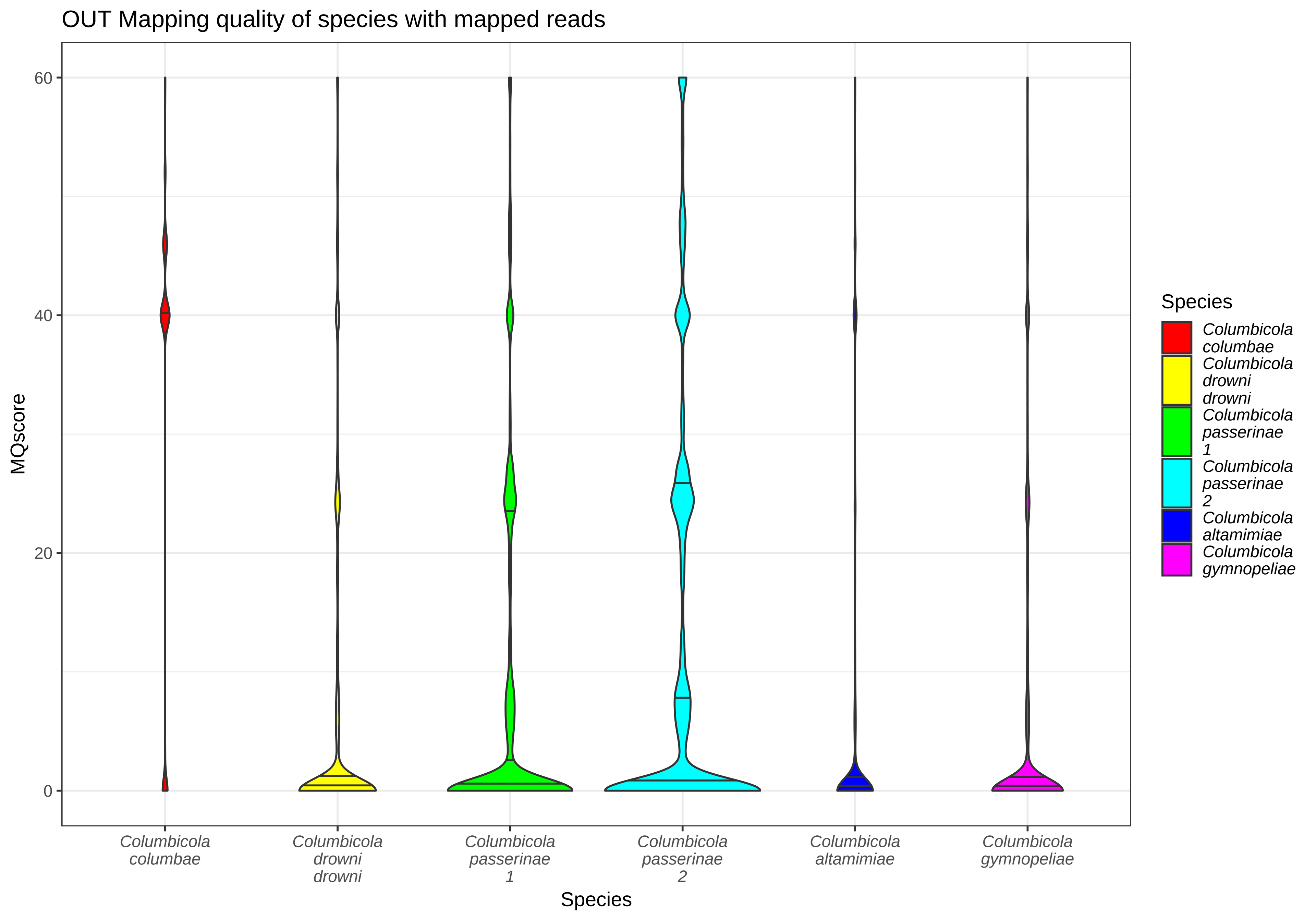

### OUT_plotMQ_Cosp_Coinc_11_2_2016_9.pdf

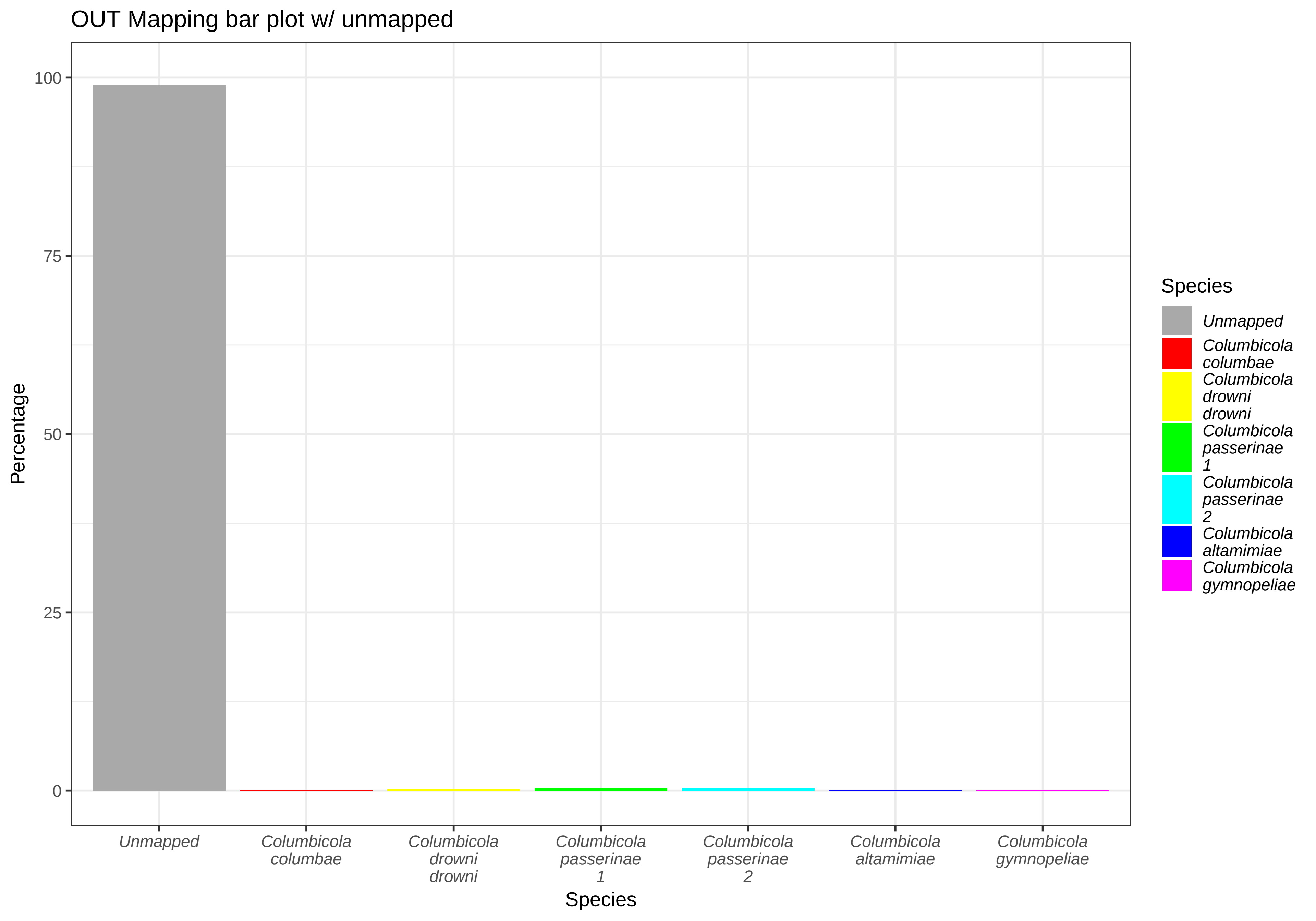

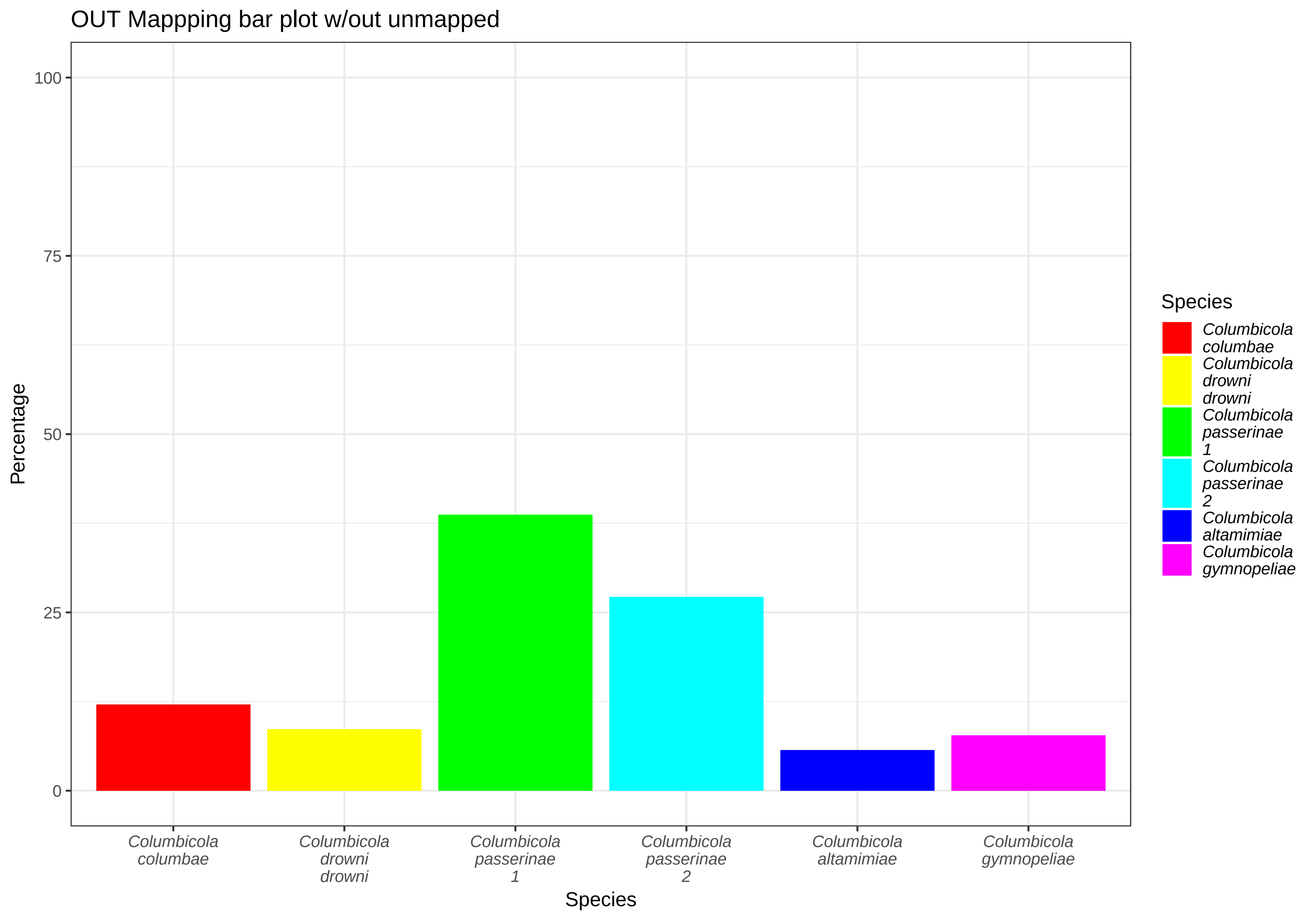

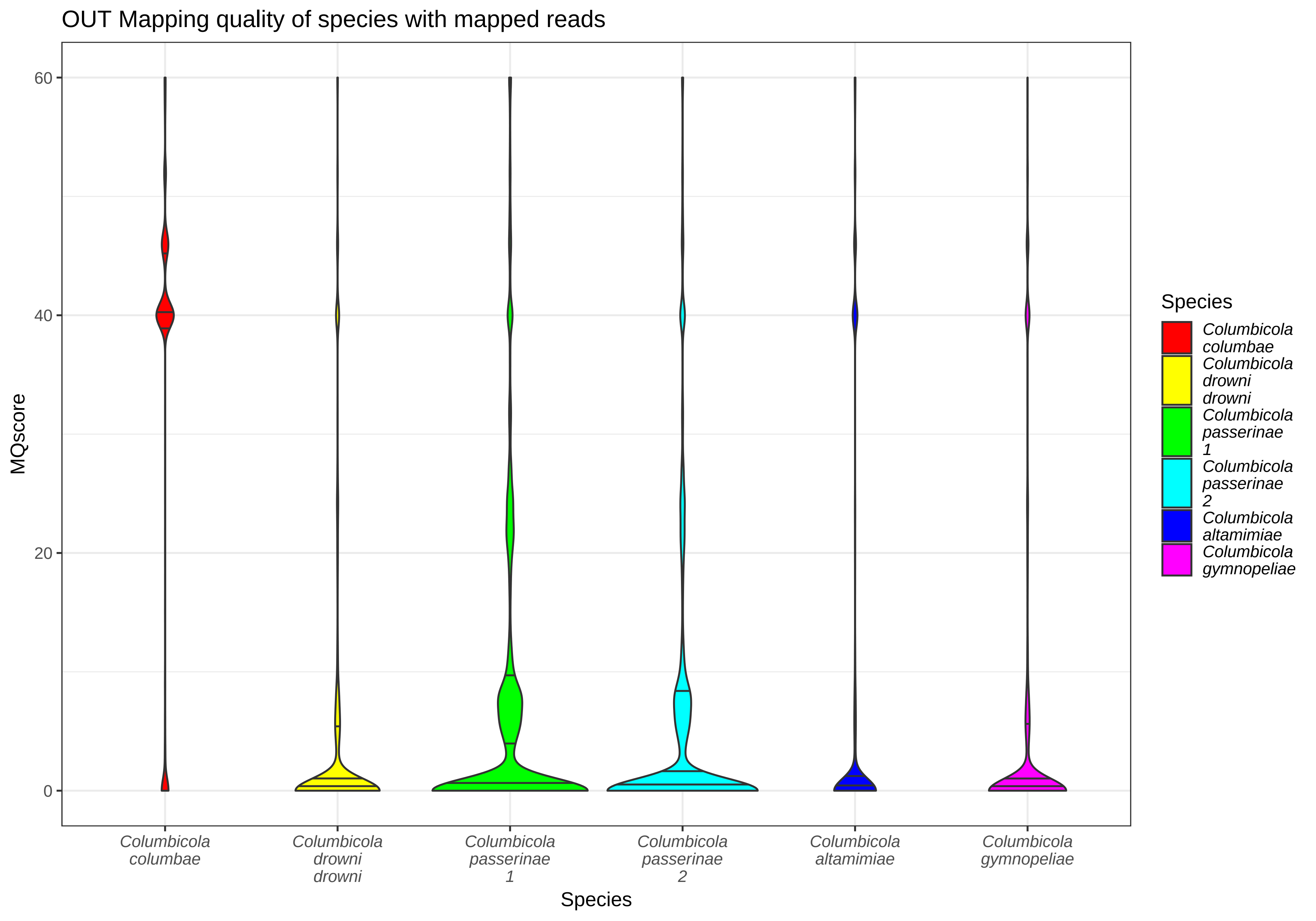

### OUT_plotMQ_Cosp_Comin_11_2_2016_4.pdf

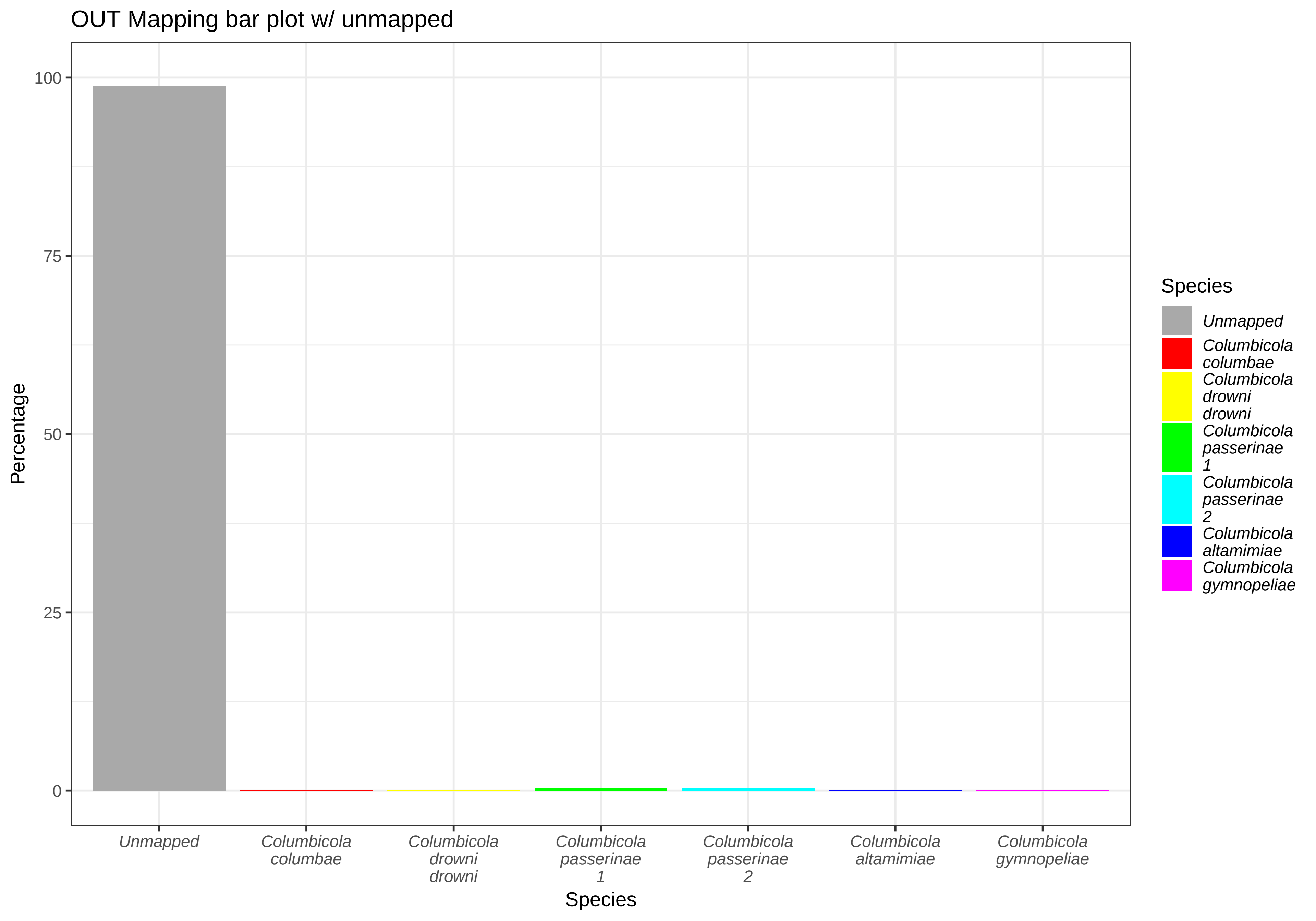

OUT Mappping bar plot w/out unmapped

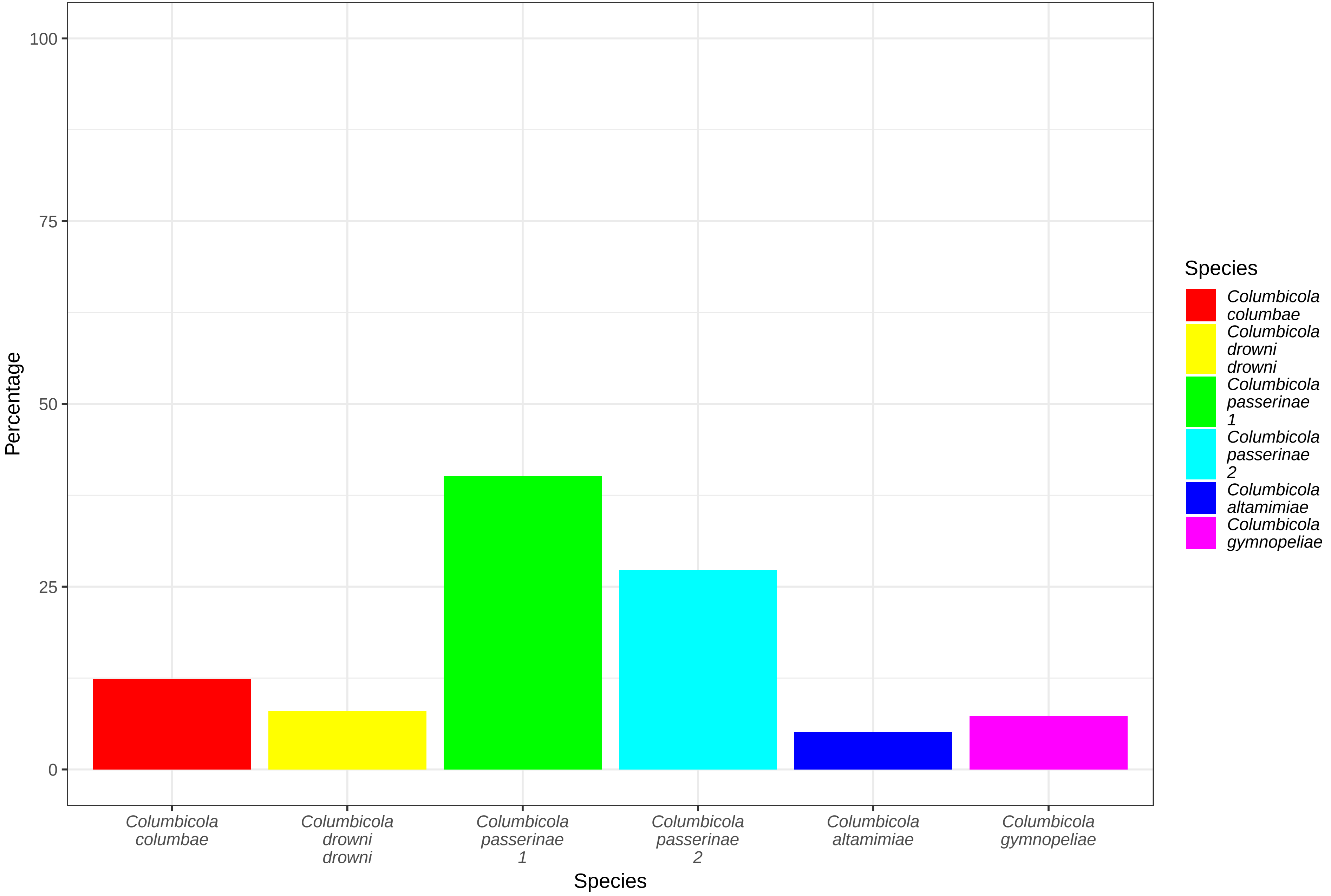

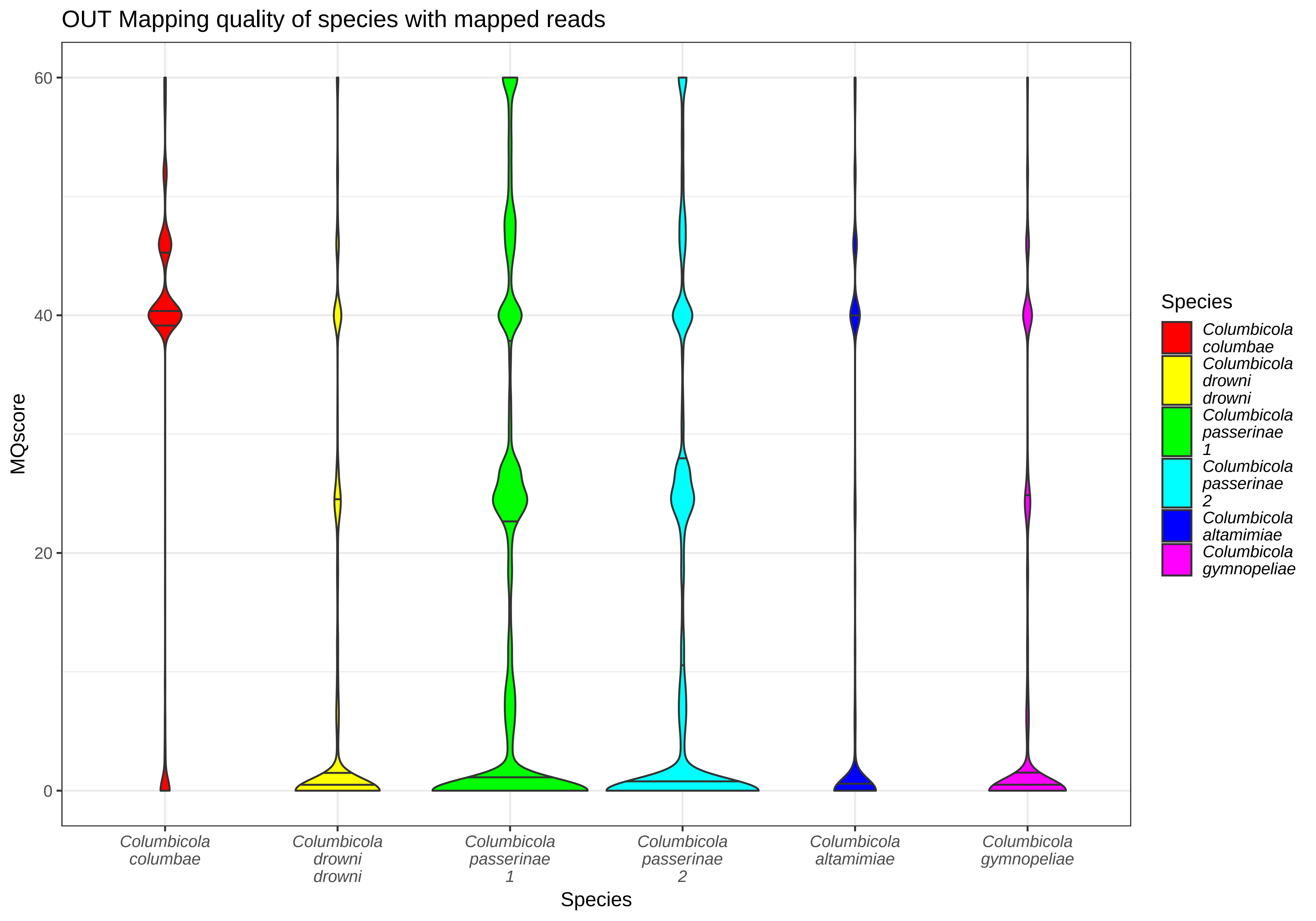

### OUT_plotMQ_Cosp_Comin_11_2_2016_16.pdf

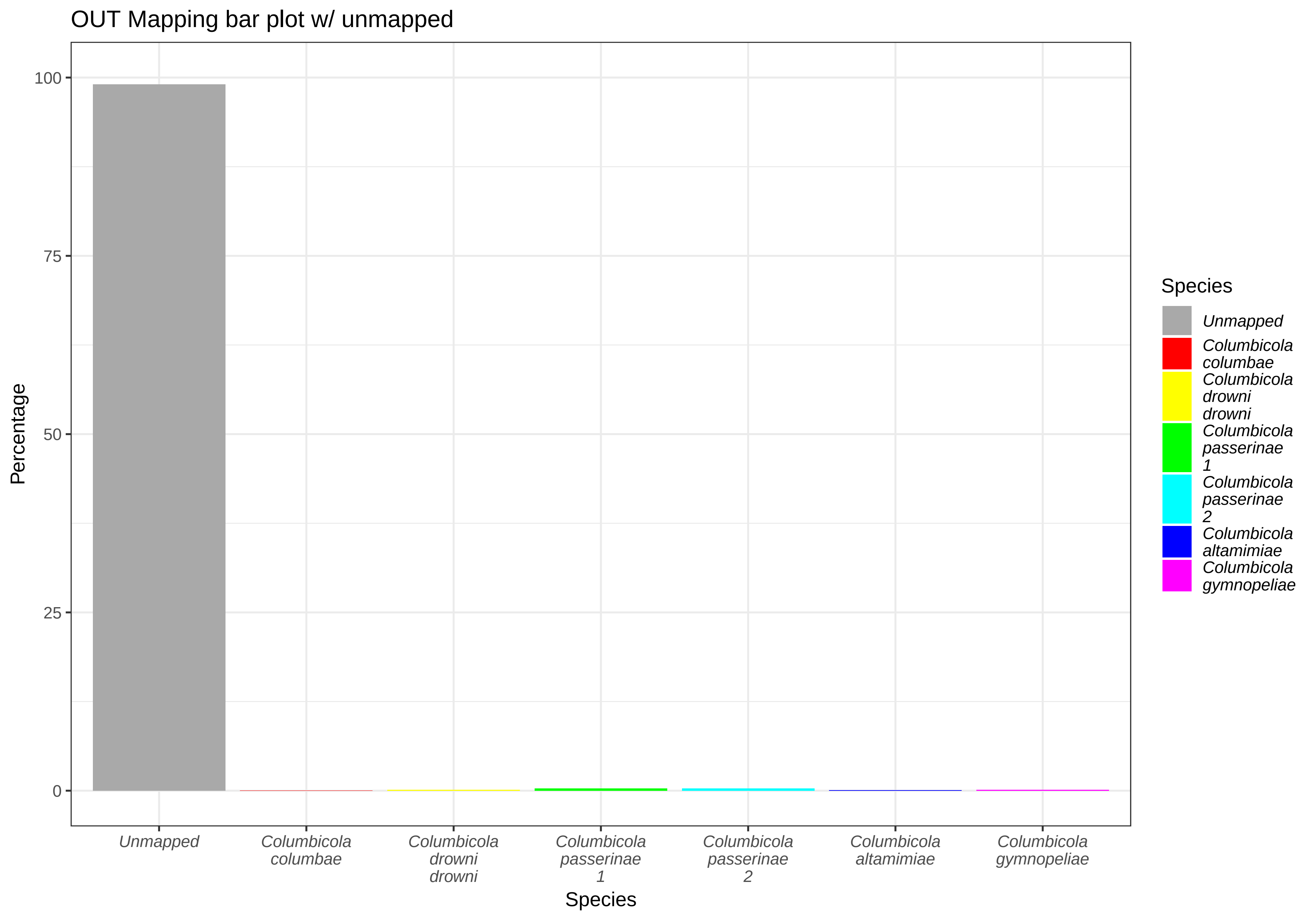

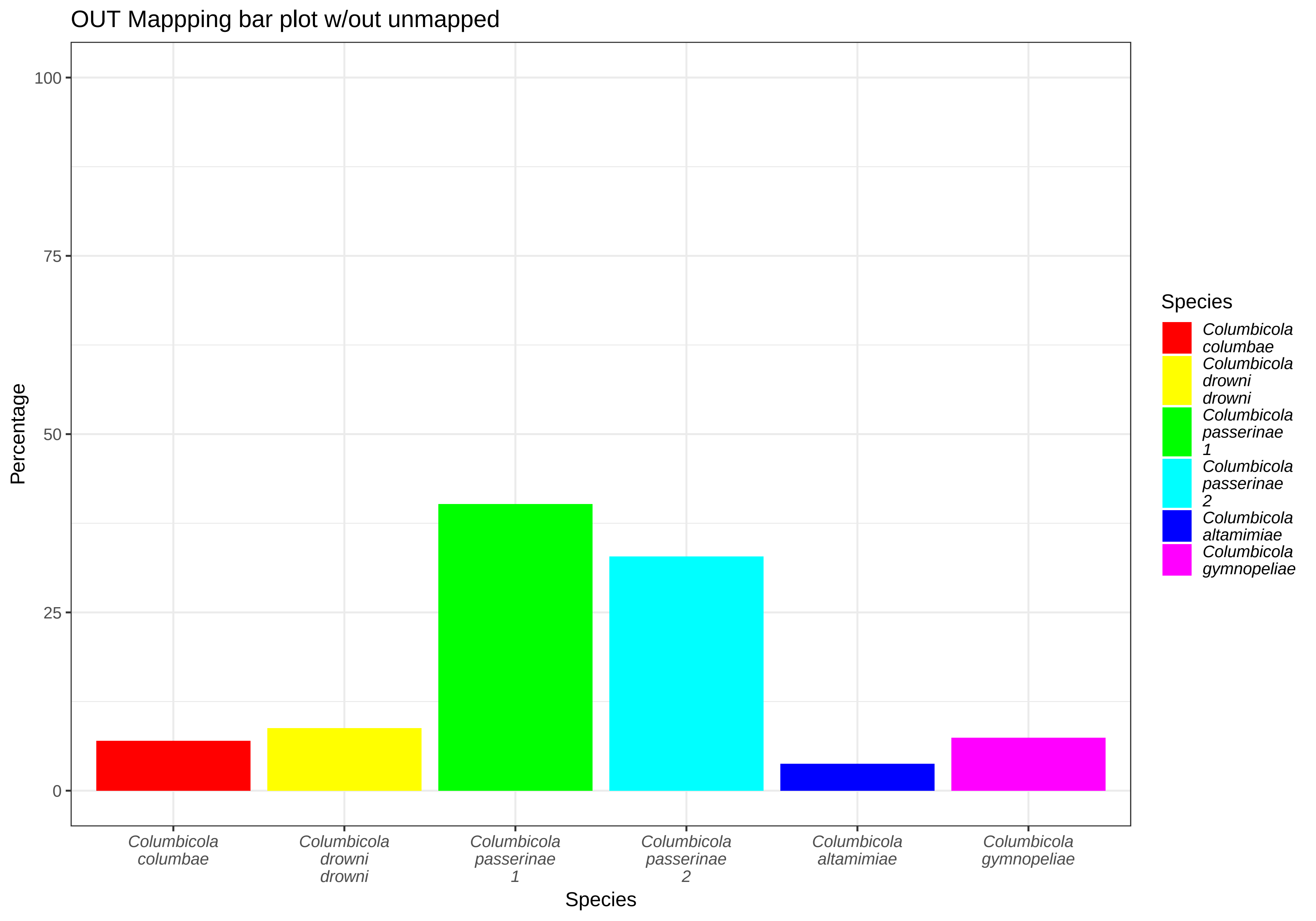

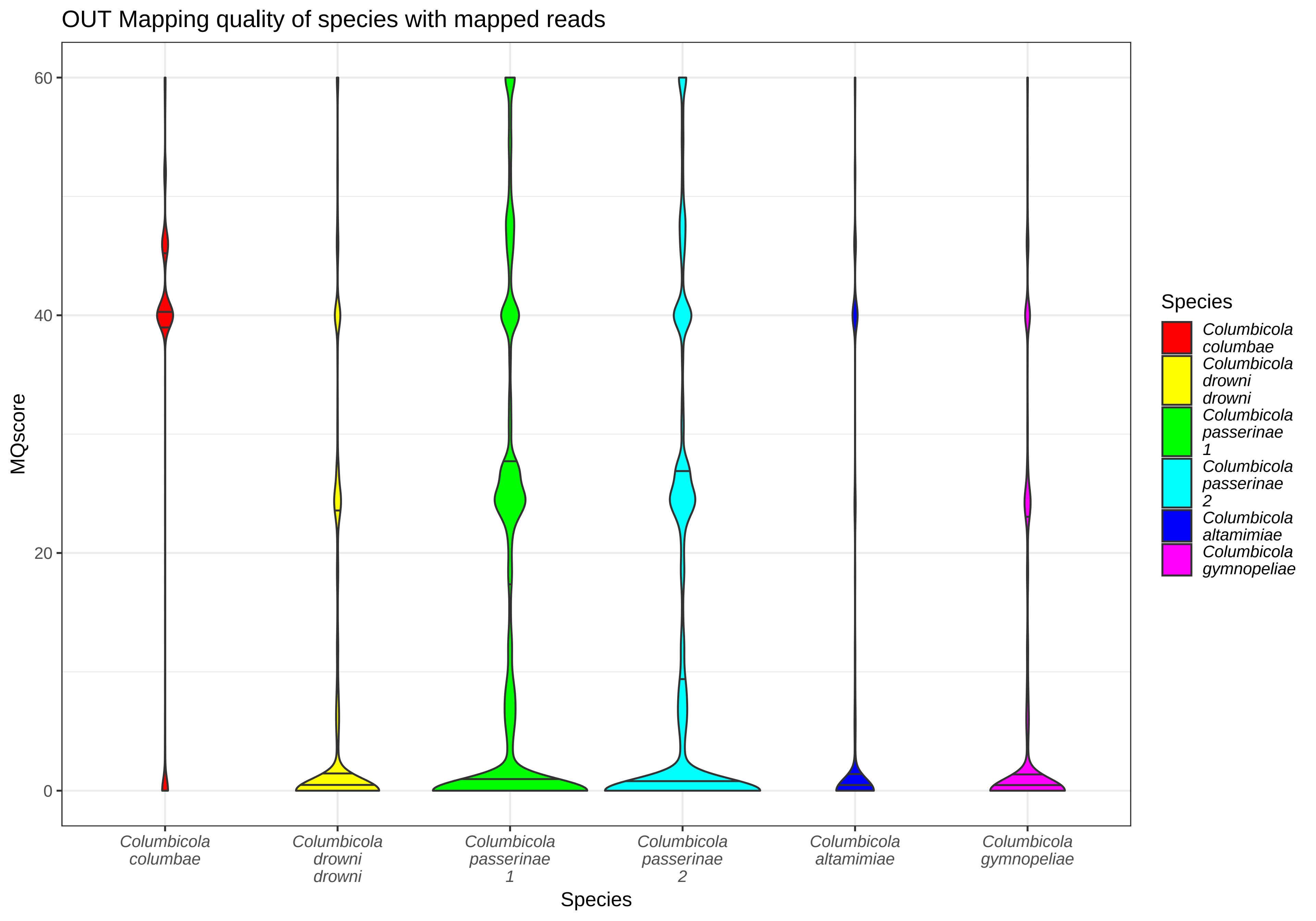

### OUT_plotMQ_Cosp_Copas_11_7_2016_8.pdf

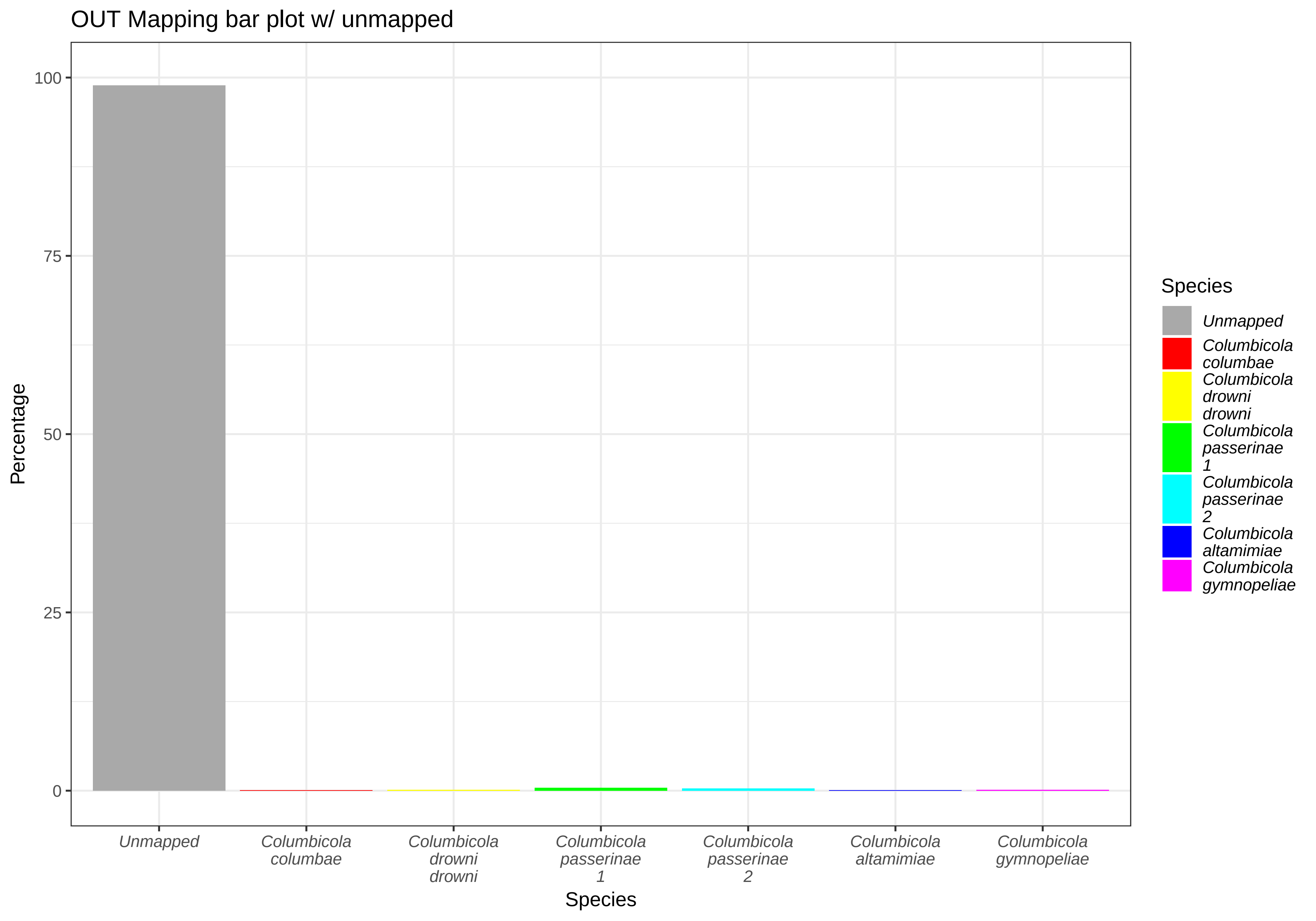

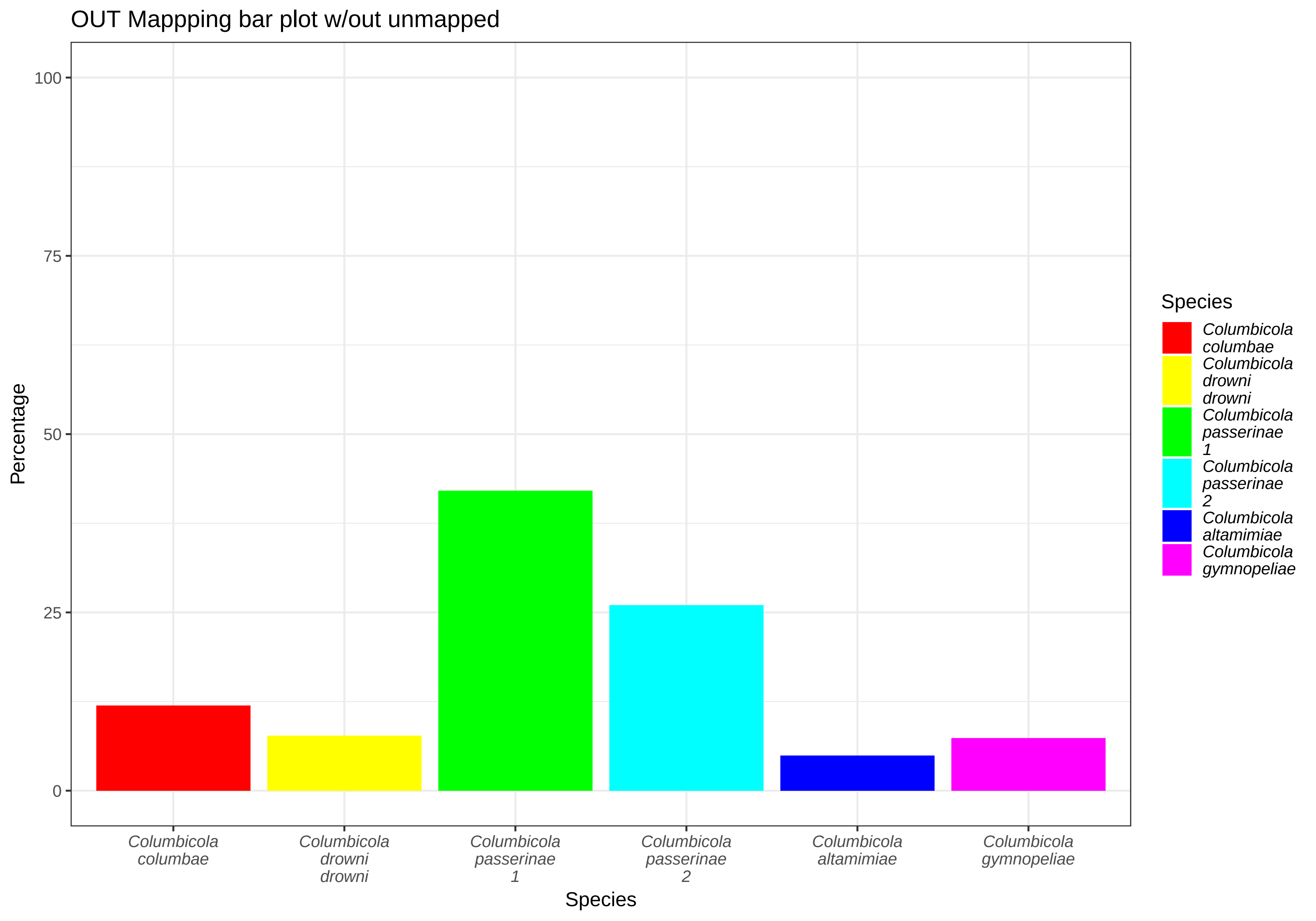

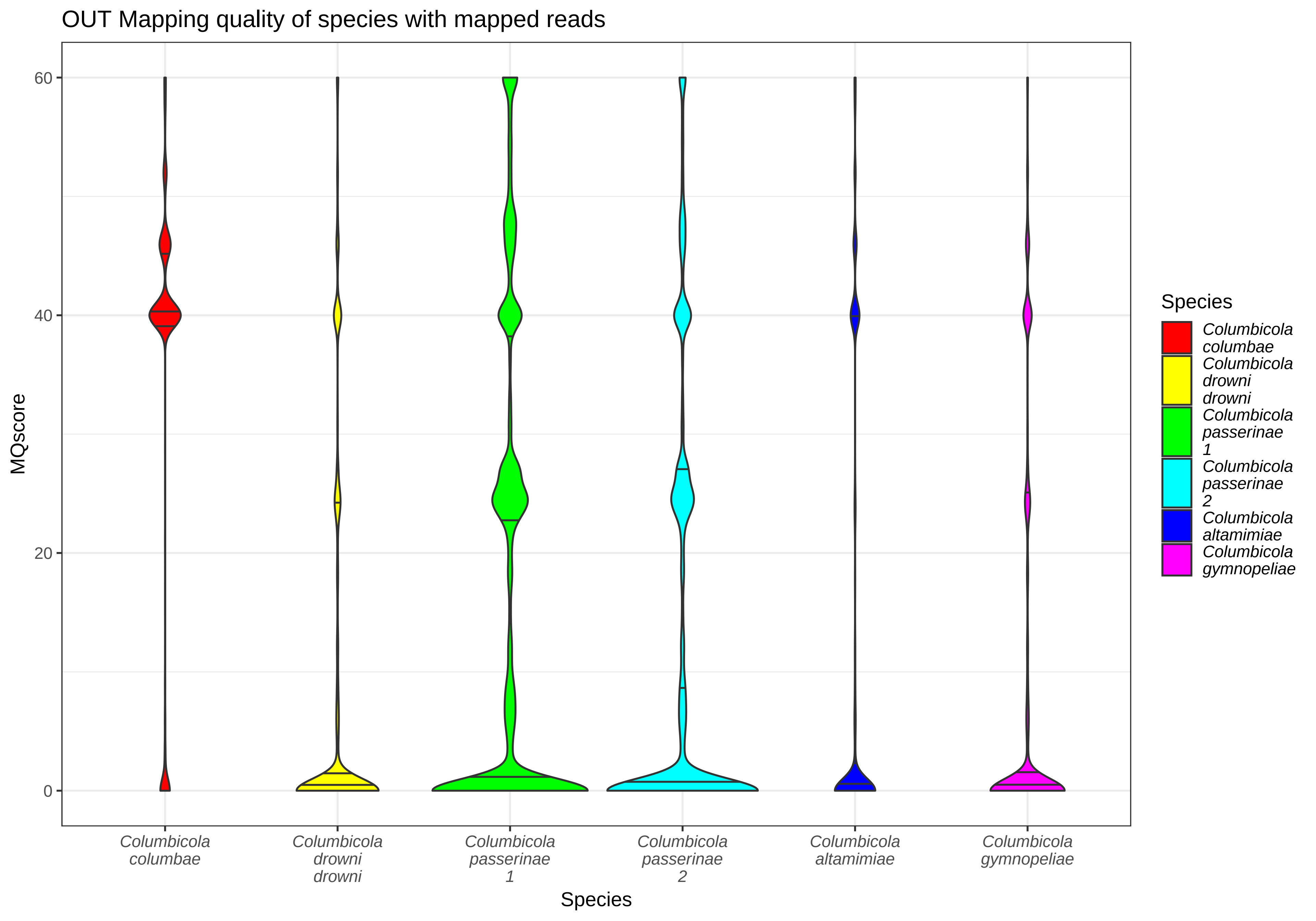

### OUT_plotMQ_Cosp_Copas_11_7_2016_12.pdf

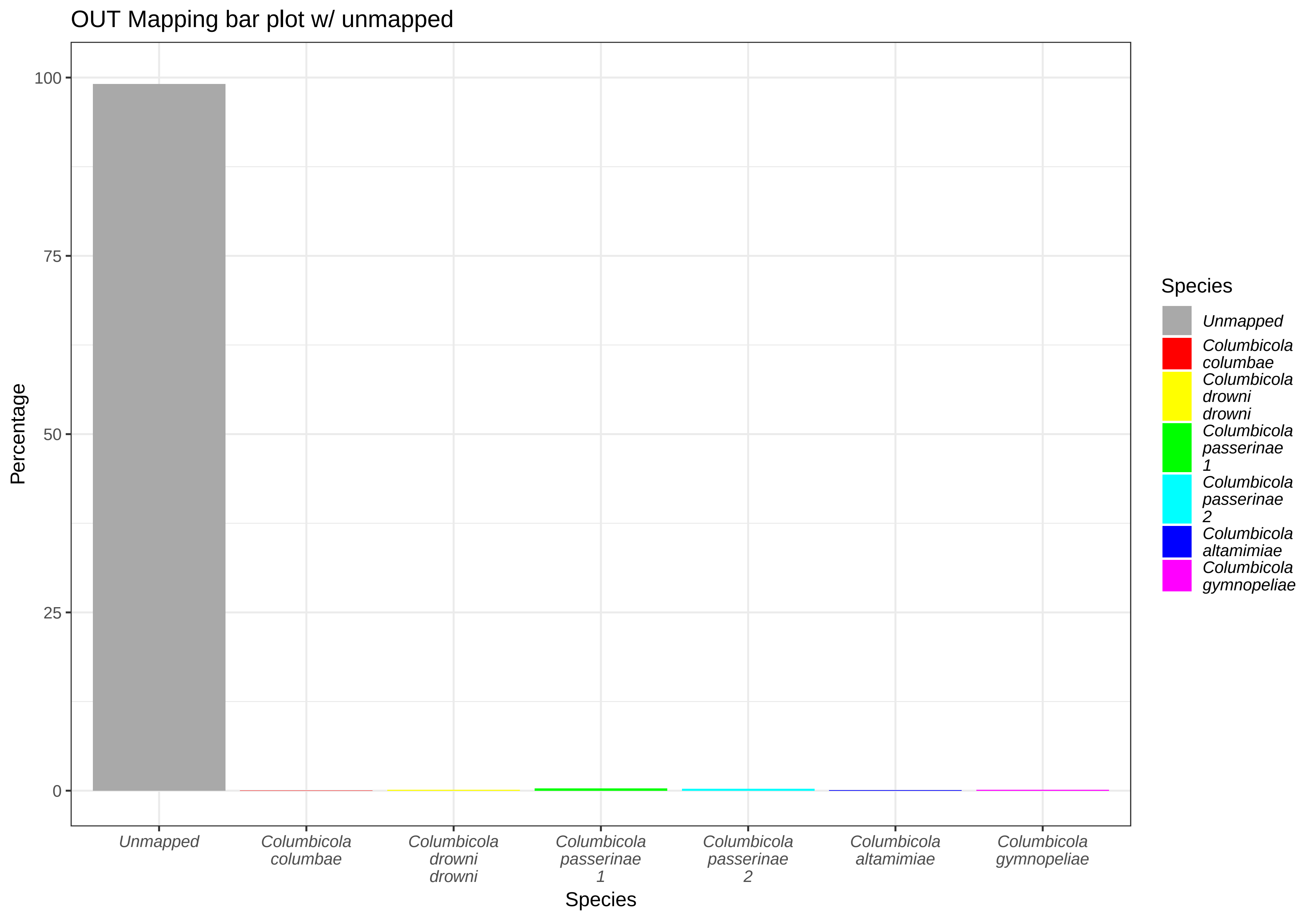

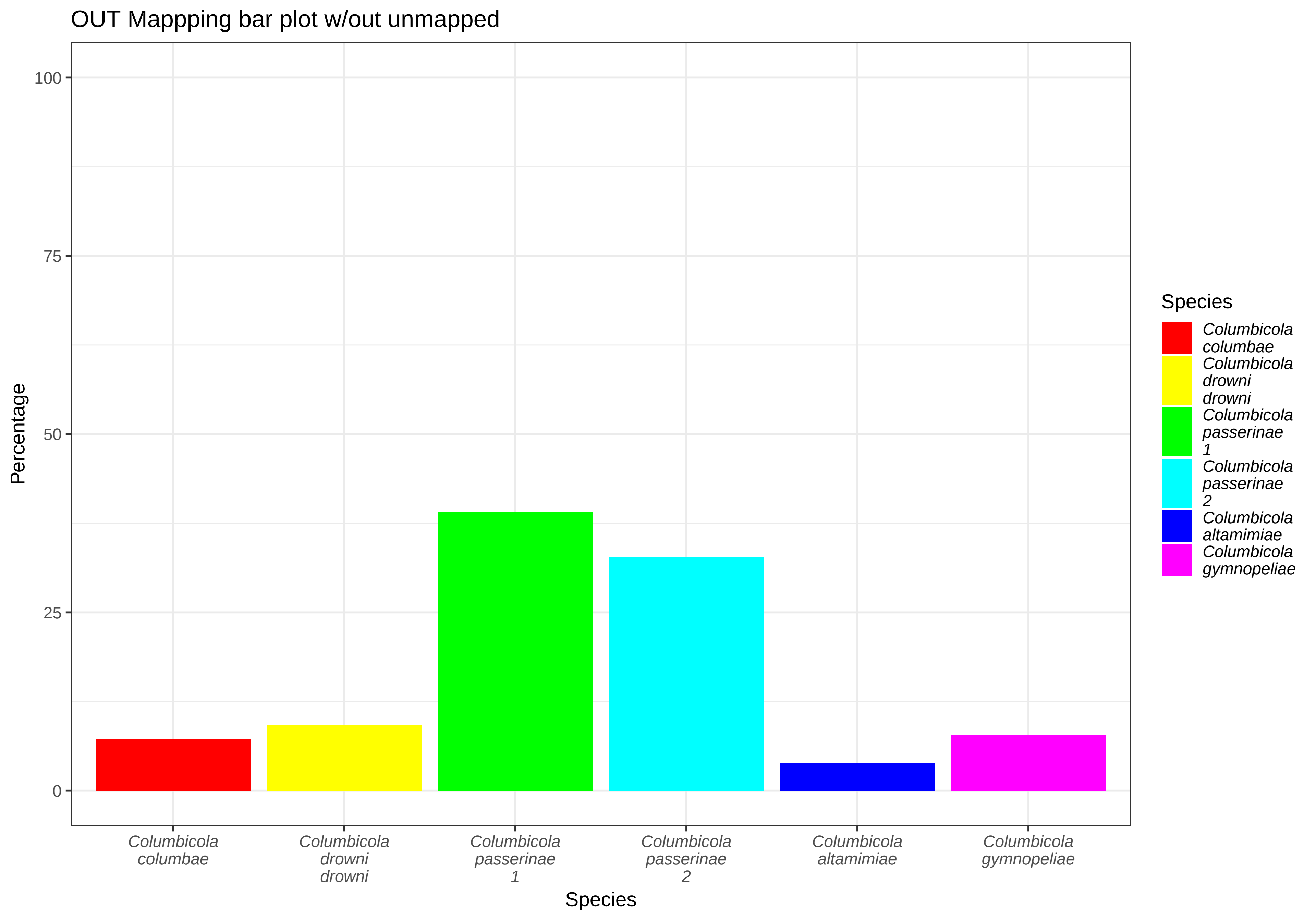

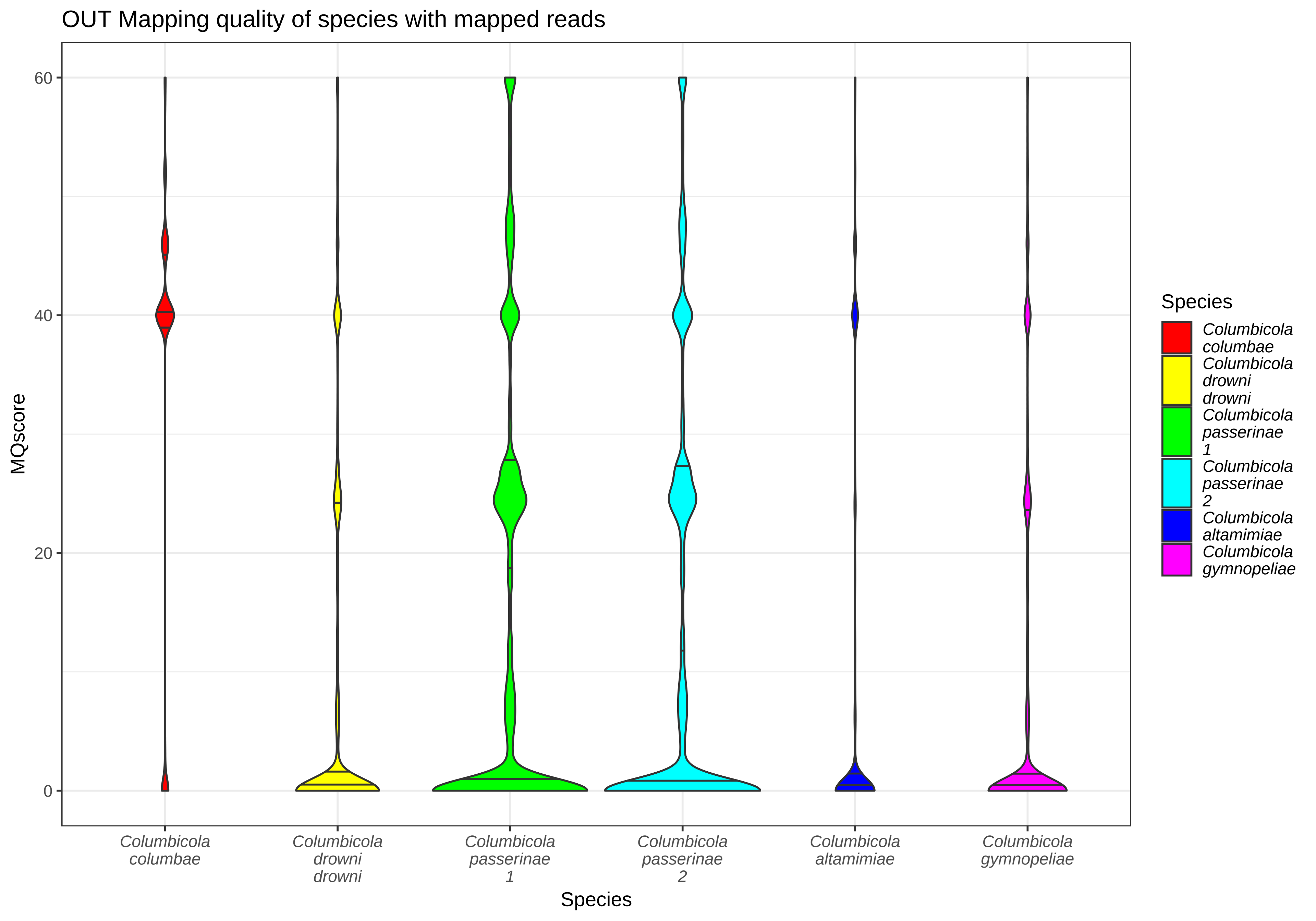

### OUT_plotMQ_Cosp_Copas_11_7_2016_14.pdf

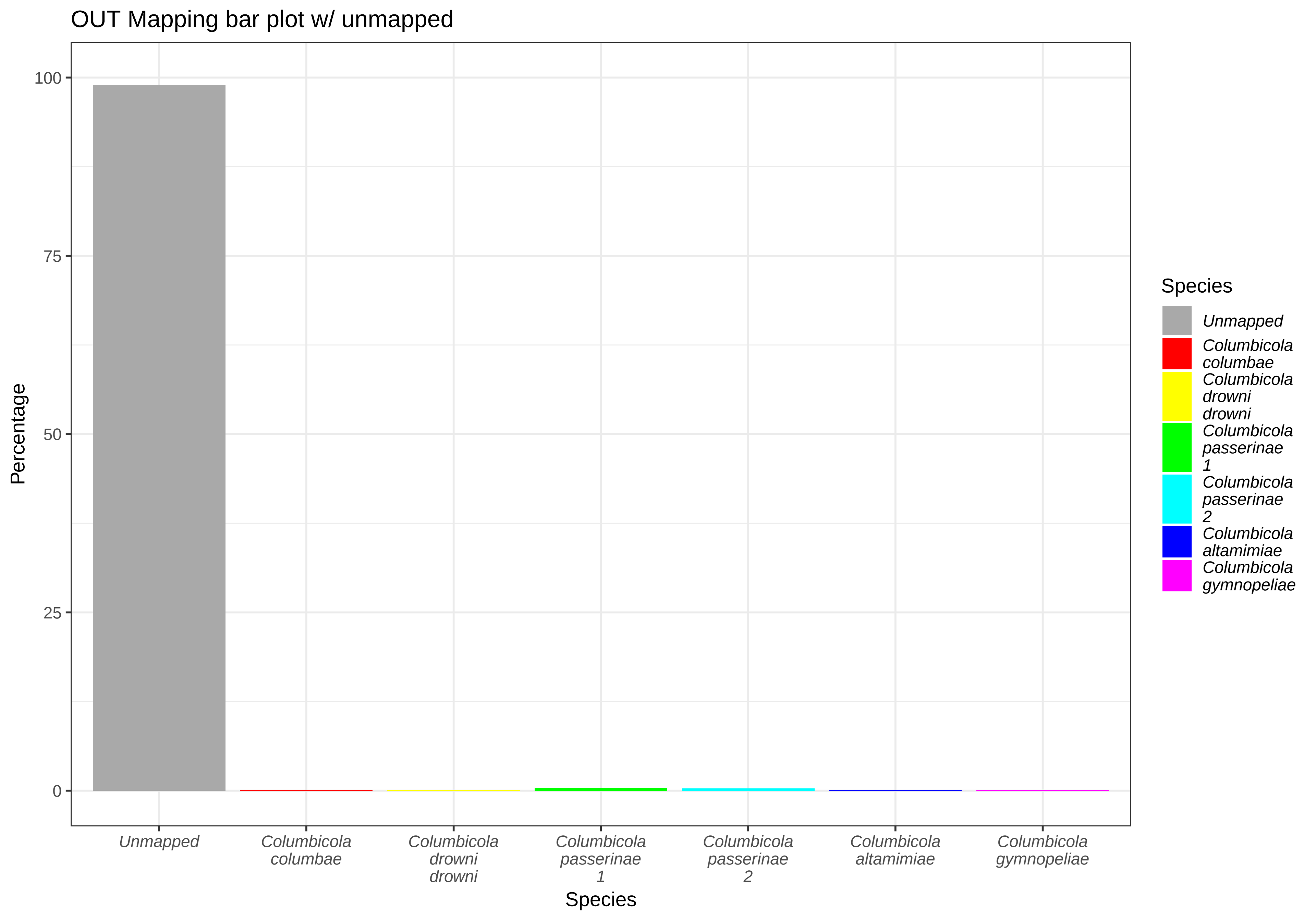

OUT Mappping bar plot w/out unmapped

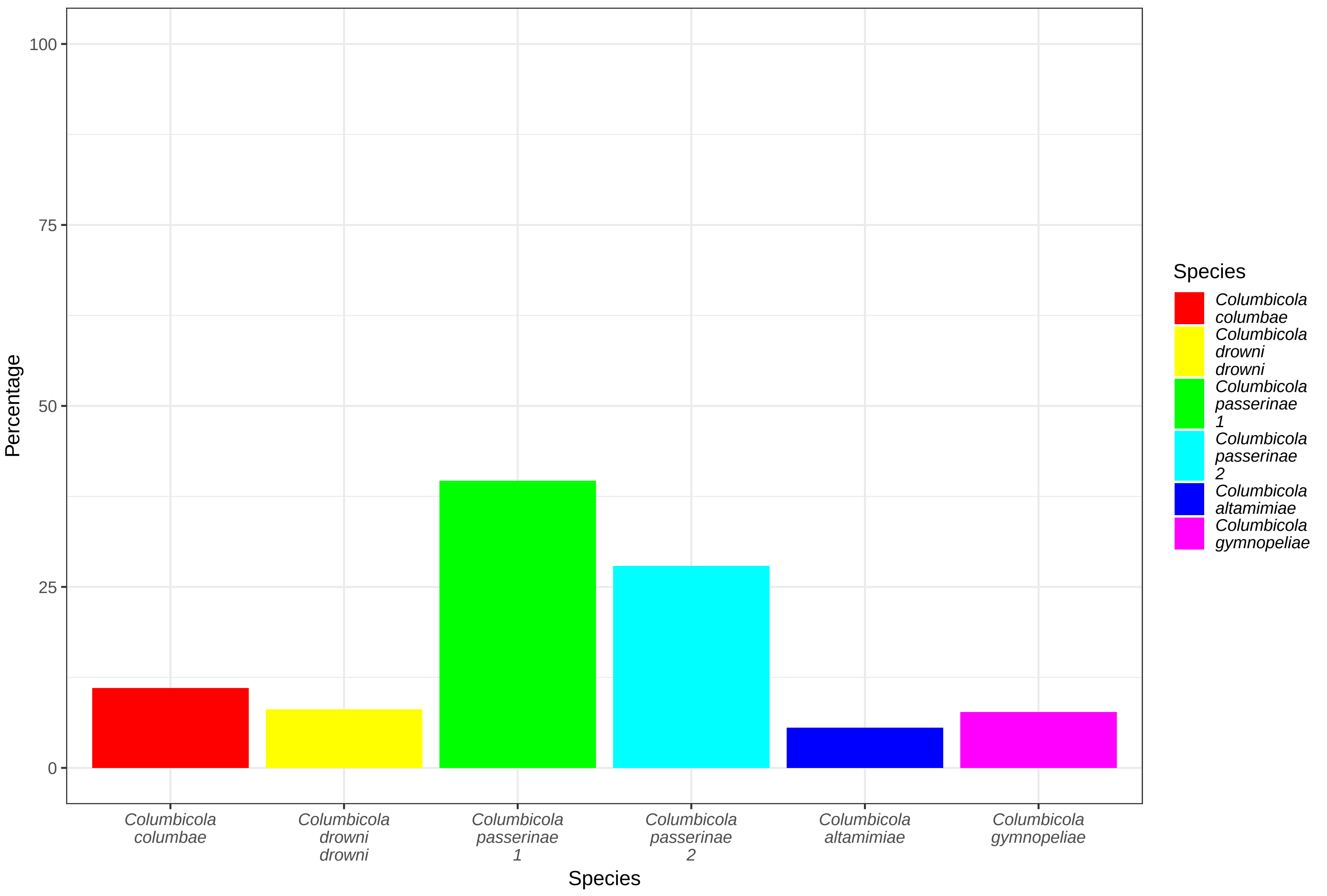

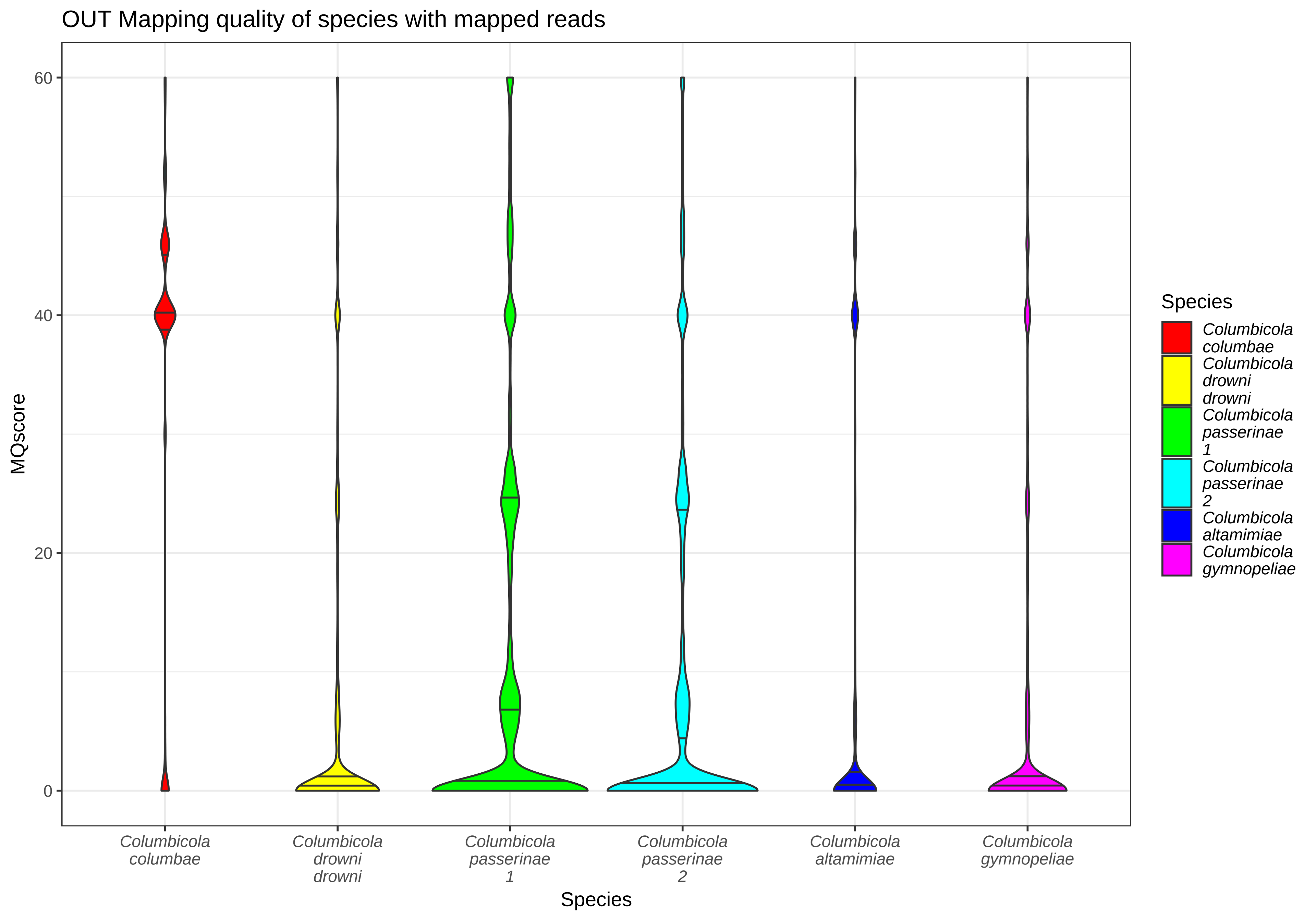

### OUT_plotMQ_Cosp_Copas_11_7_2016_15.pdf

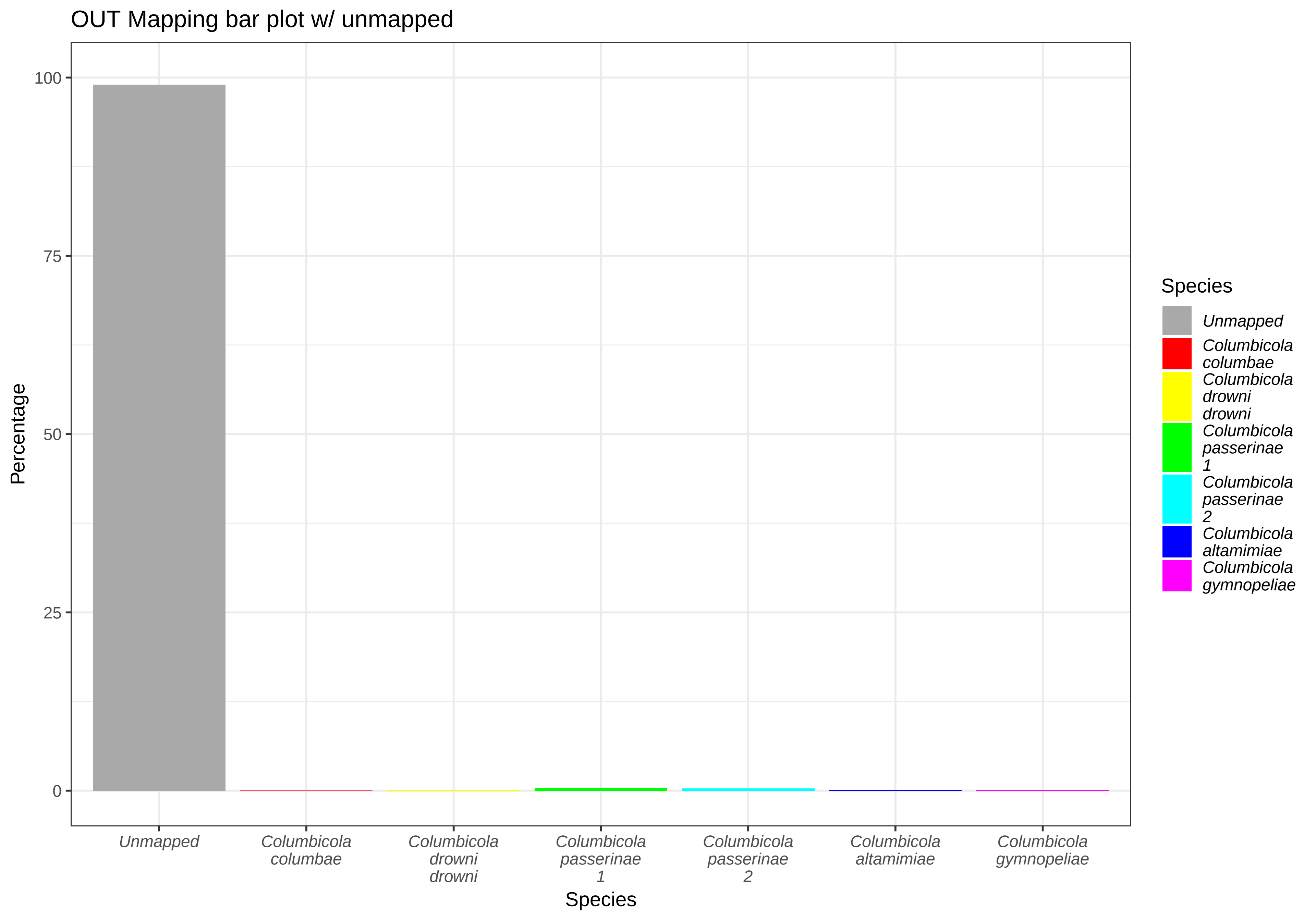

OUT Mappping bar plot w/out unmapped

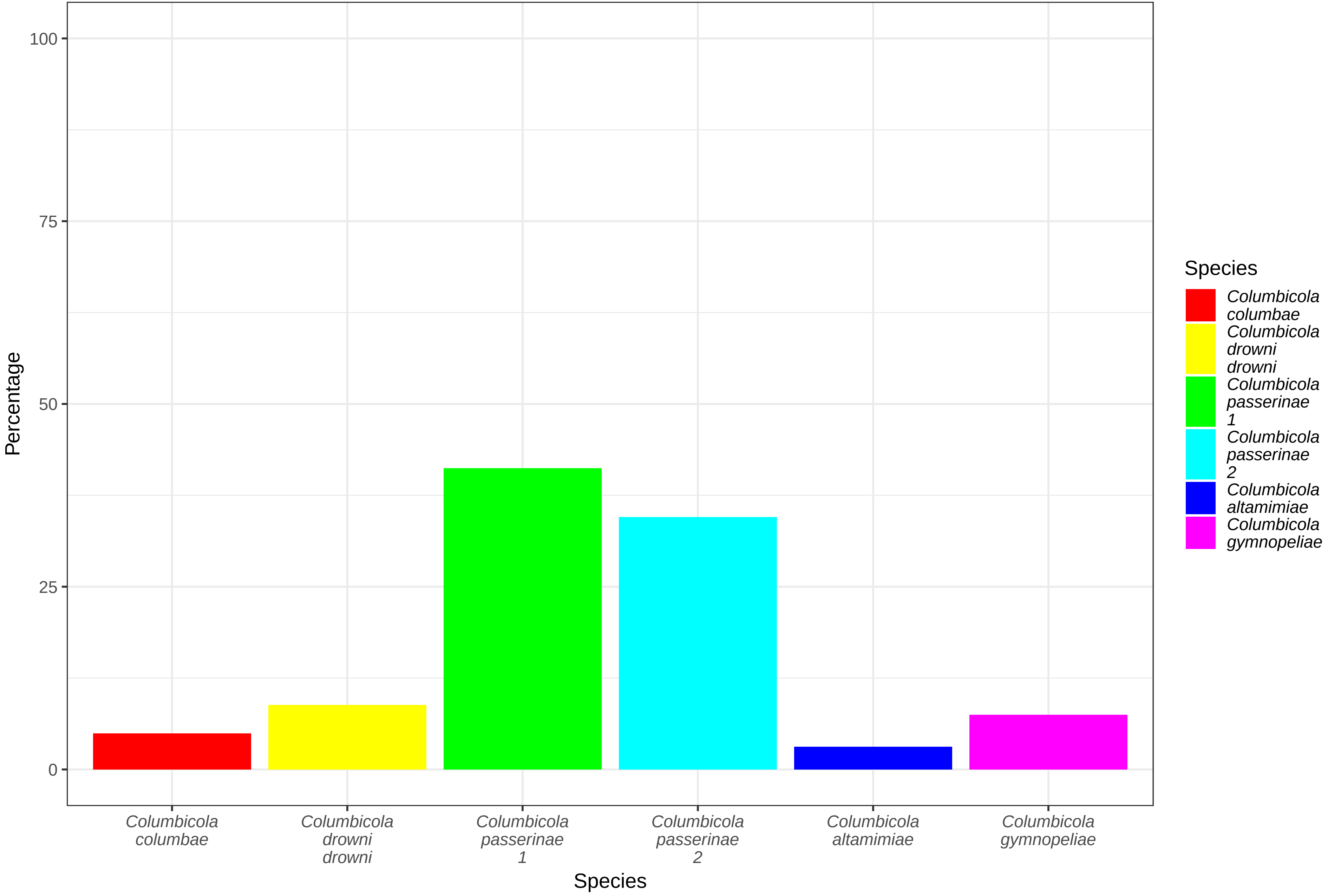

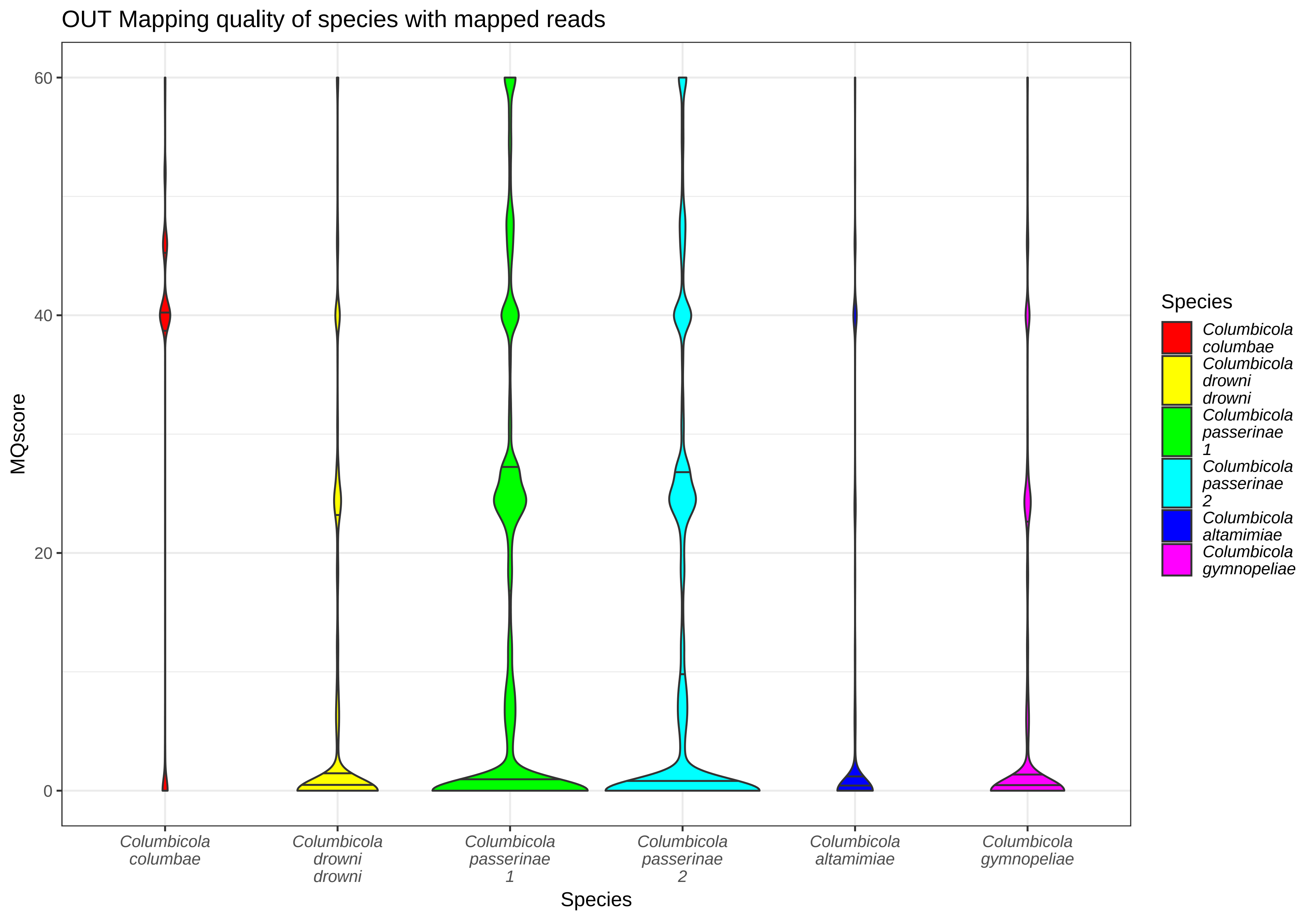

### OUT_plotMQ_Cosp_Copic_11_2_2016_1.pdf

OUT Mapping bar plot w/ unmapped

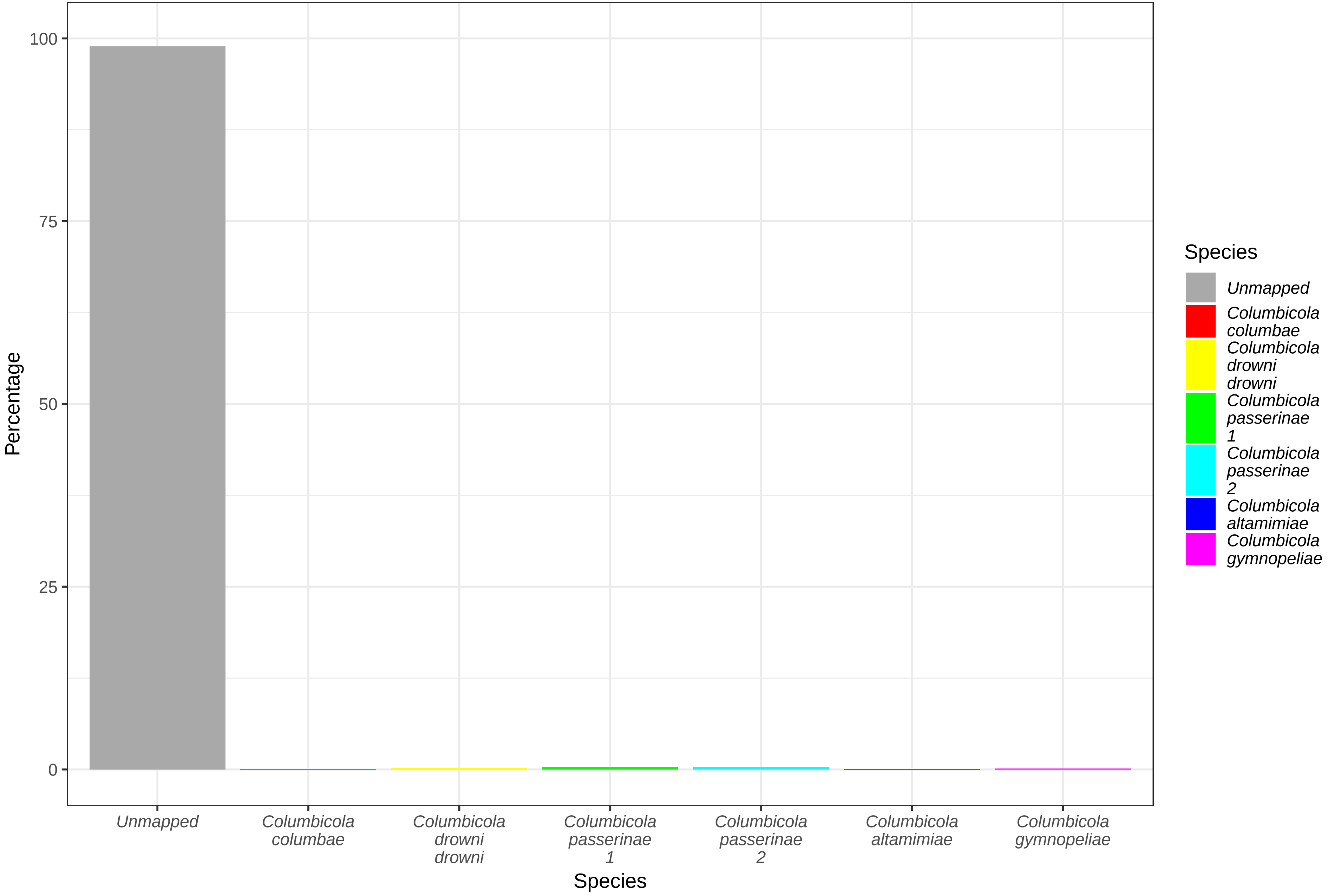

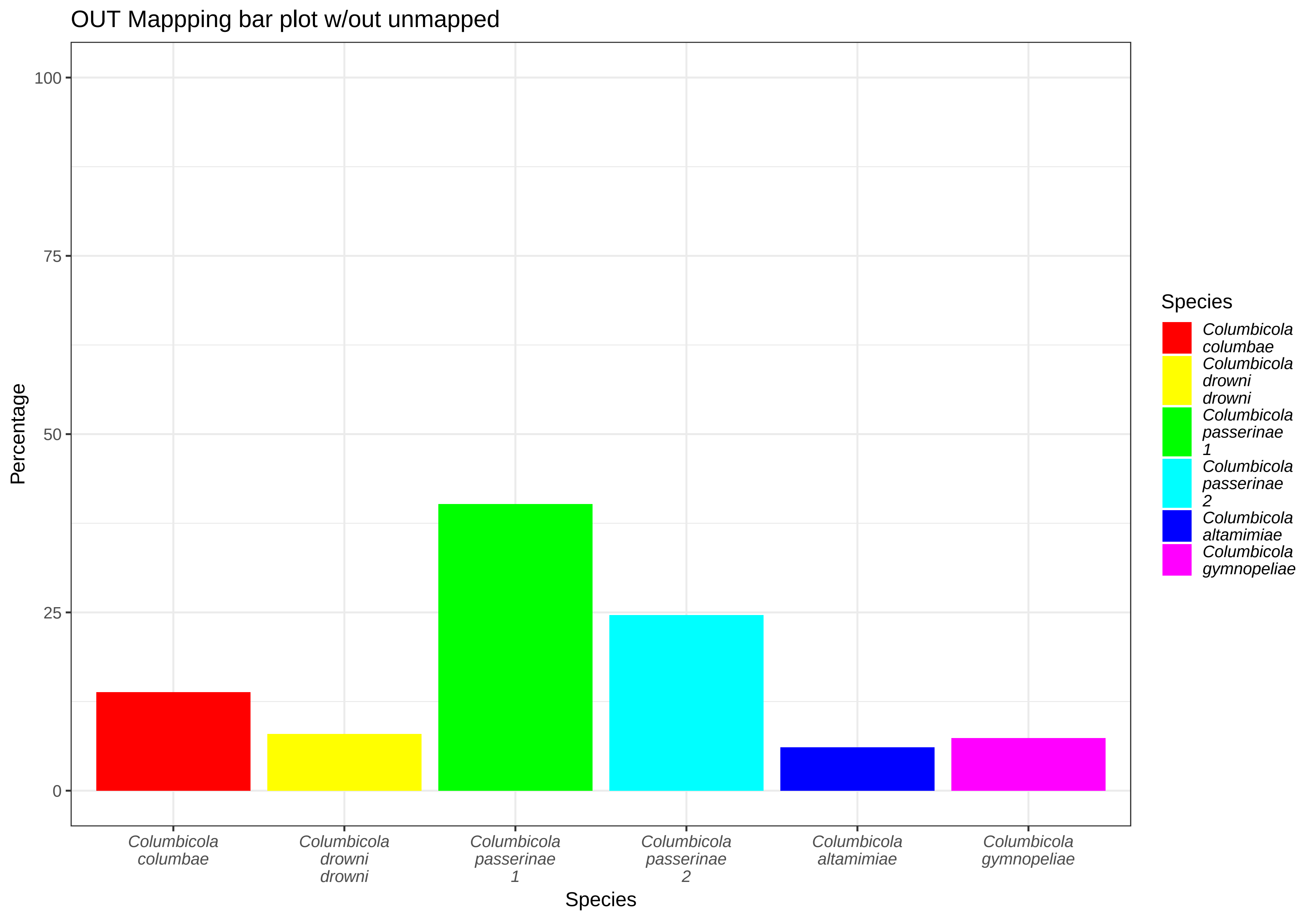

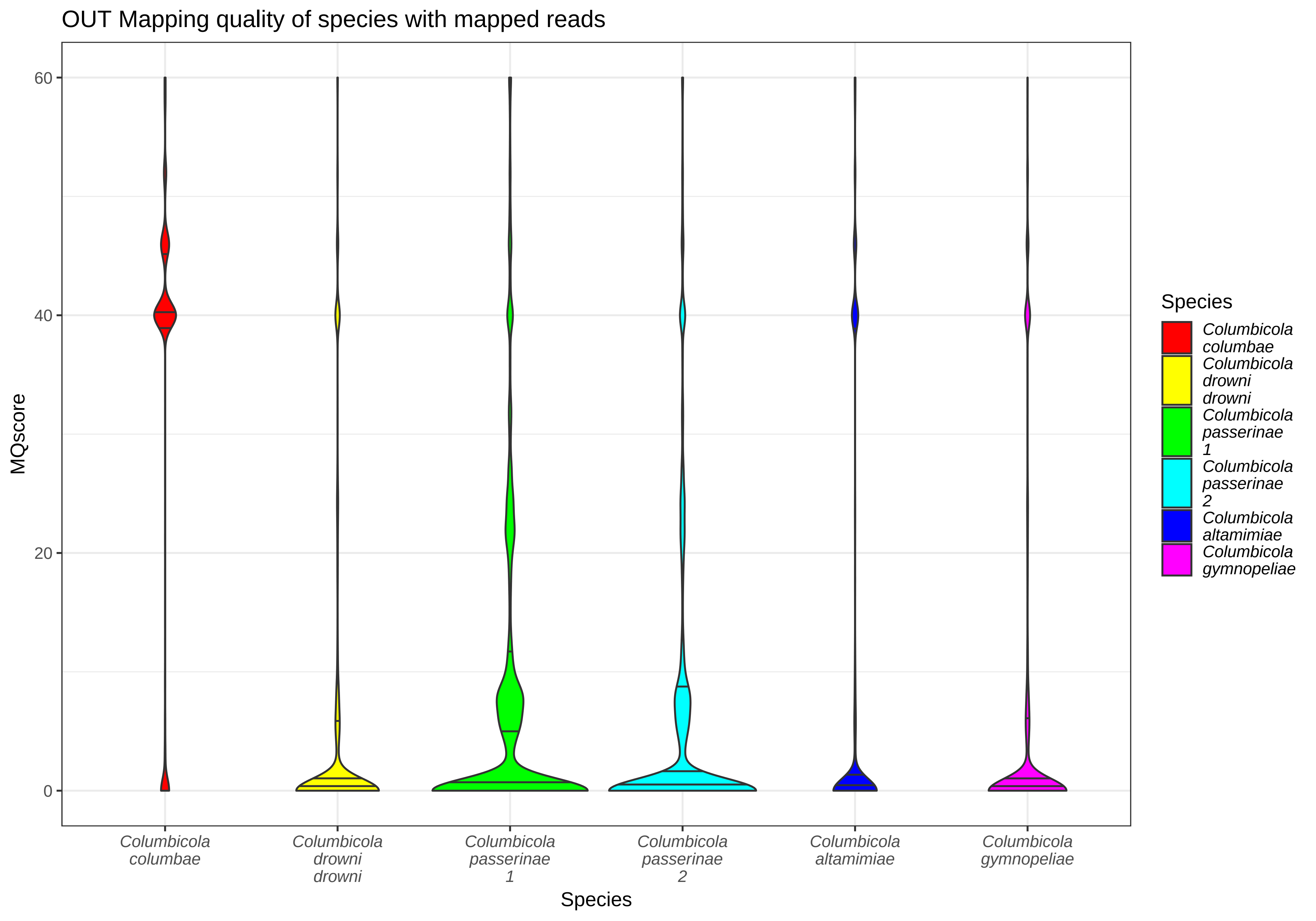

### OUT_plotMQ_Cosp_Copic_11_2_2016_3.pdf

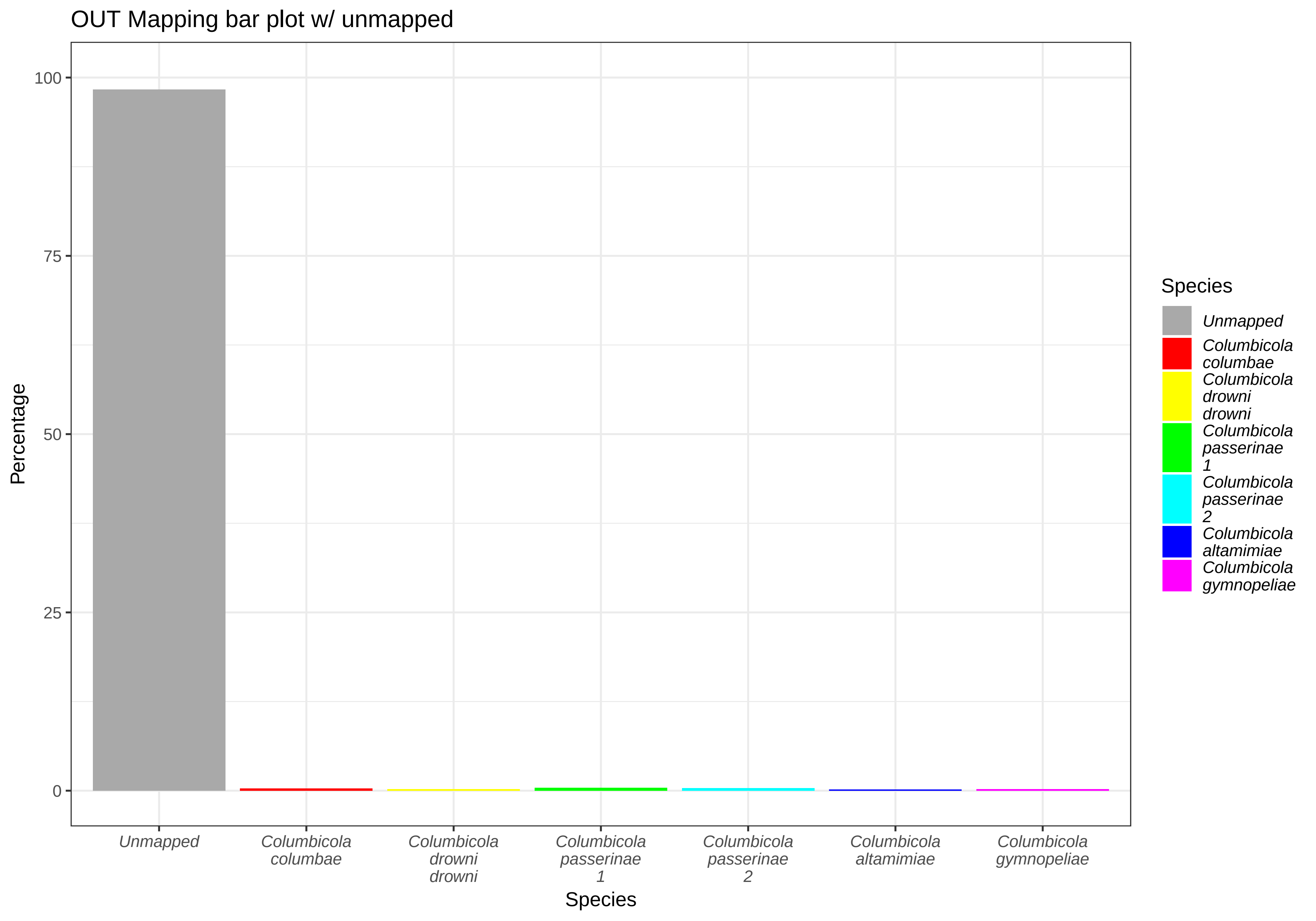

OUT Mappping bar plot w/out unmapped

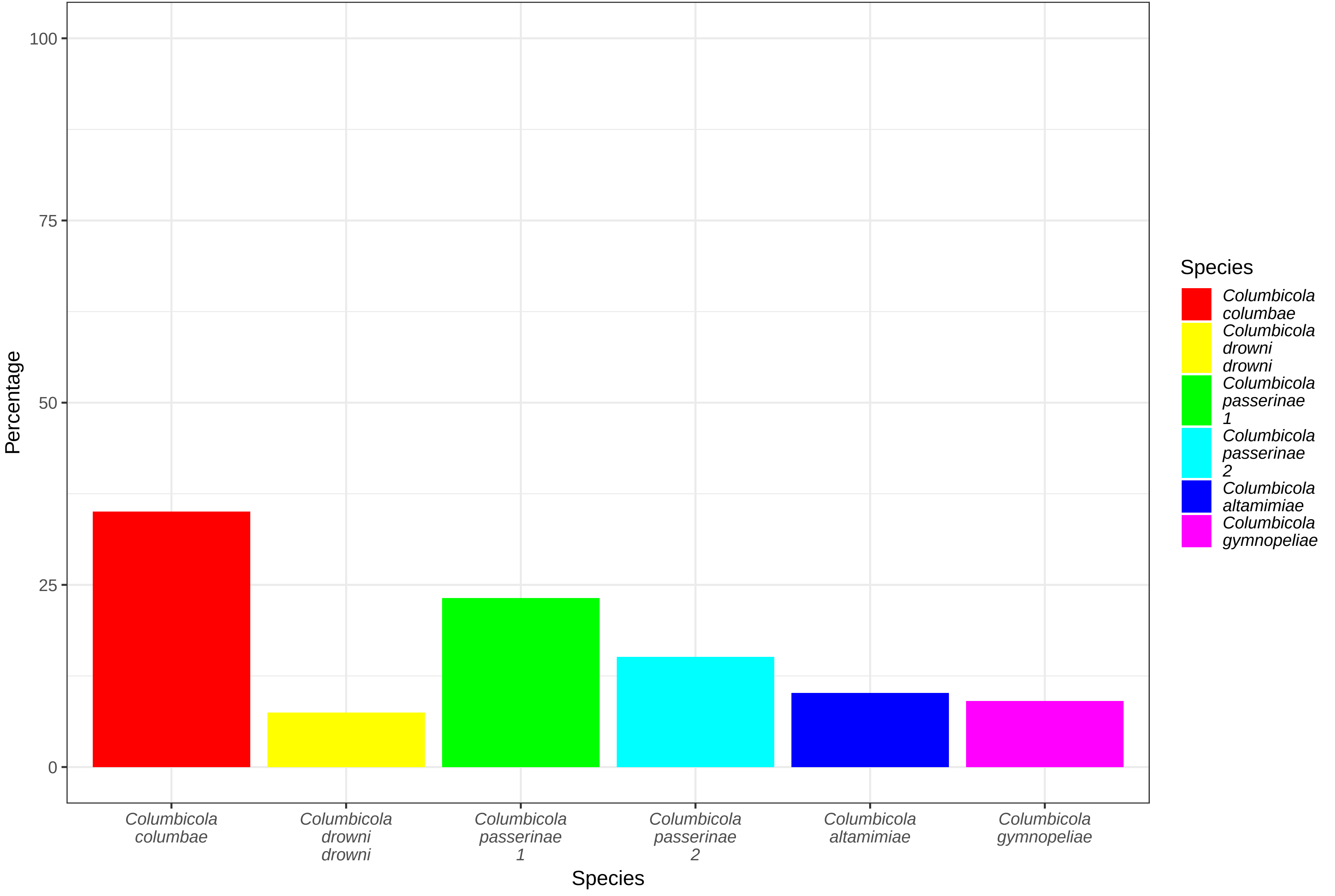

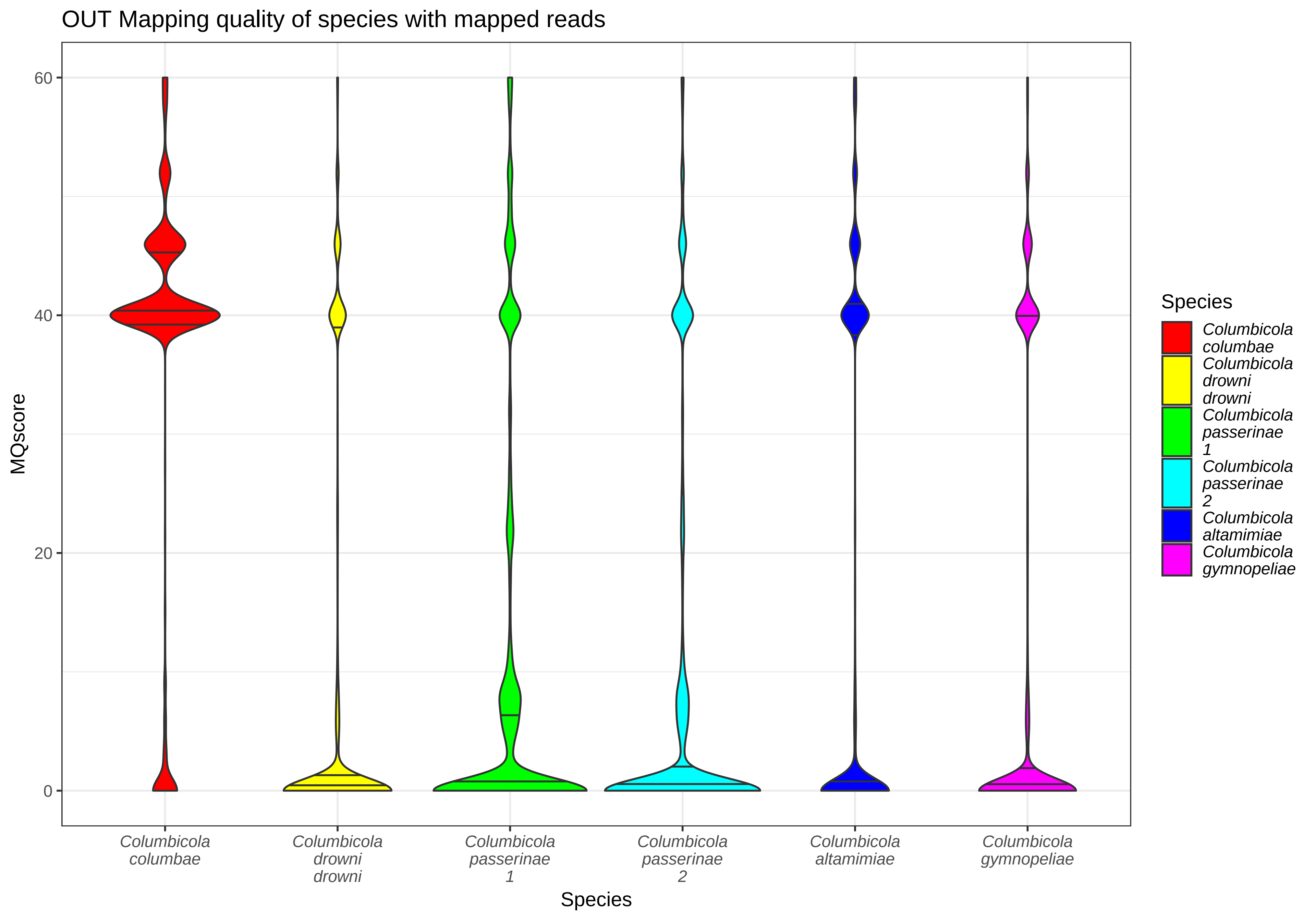

### OUT_plotMQ_Cosp_Cosqu_11_7_2016_1.pdf

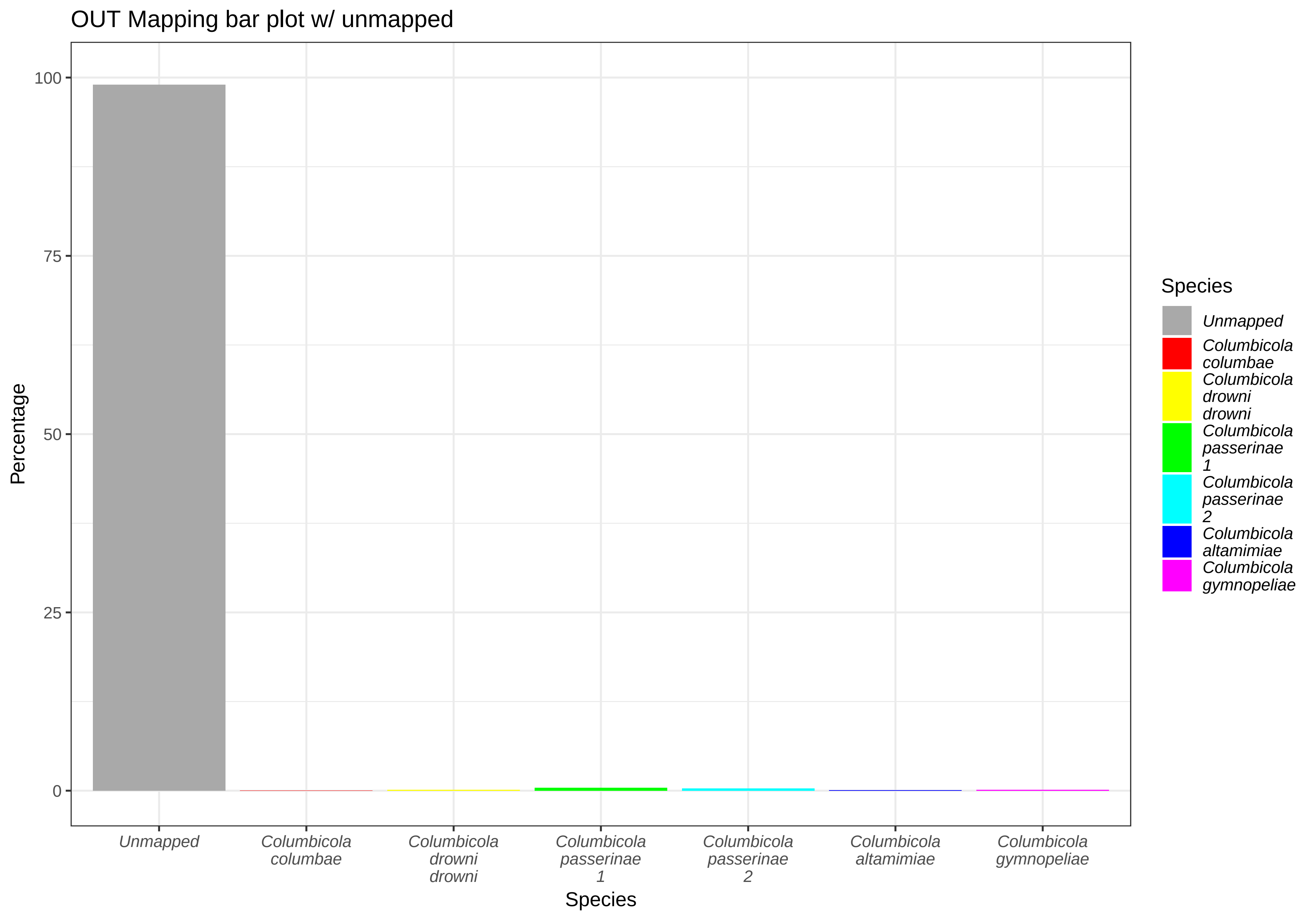

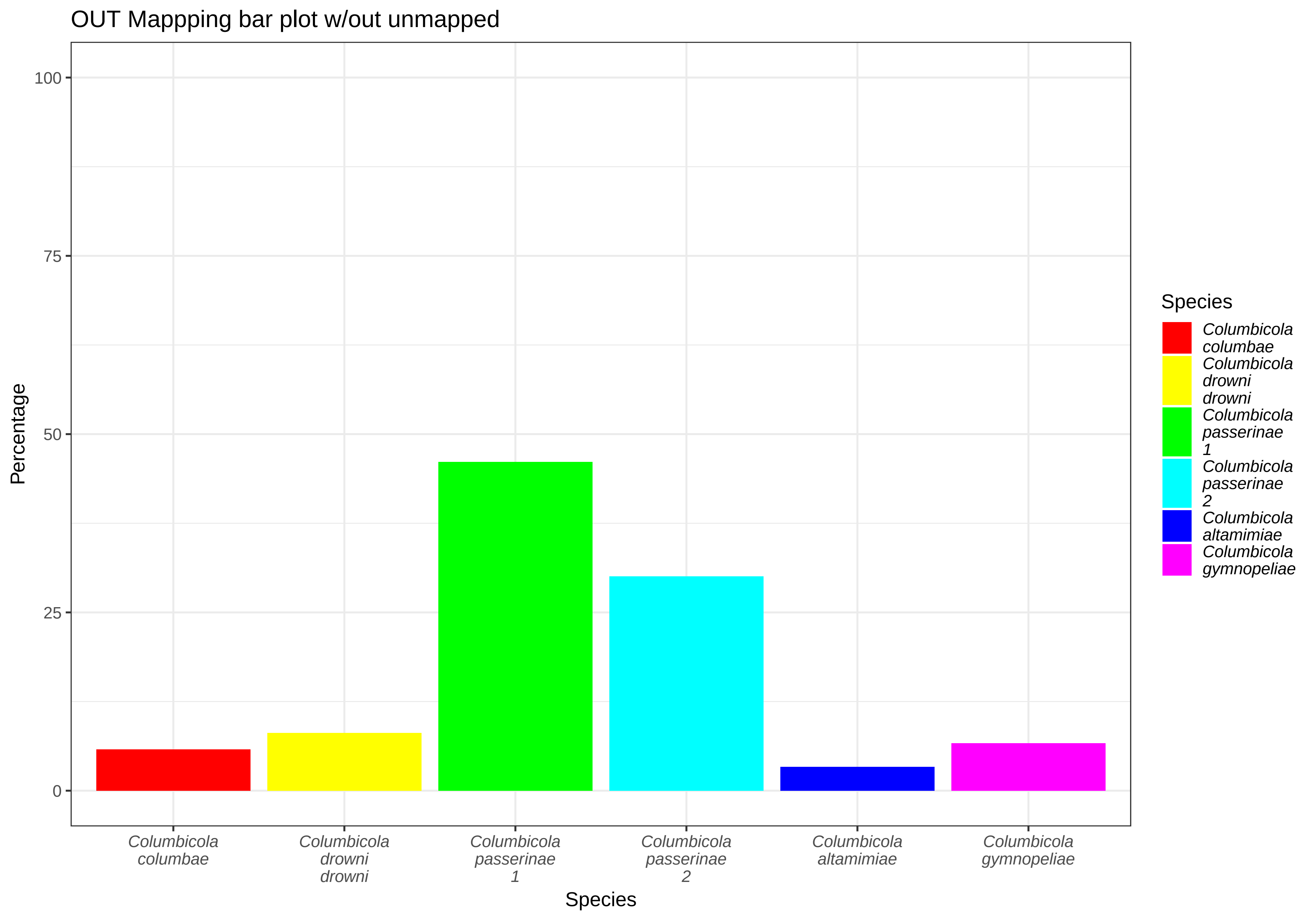

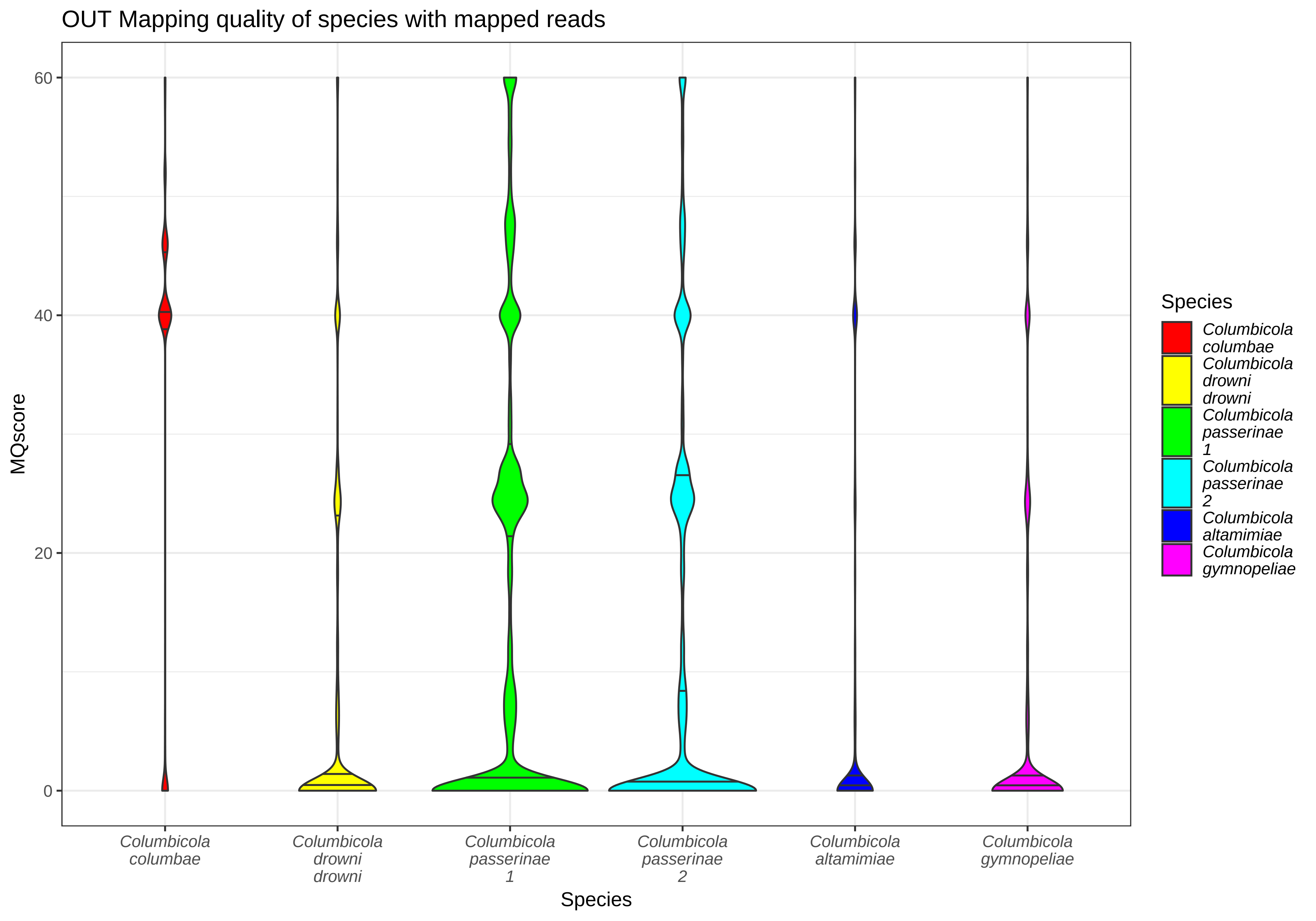

### OUT_plotMQ_Cosp_Cosqu_11_7_2016_5.pdf

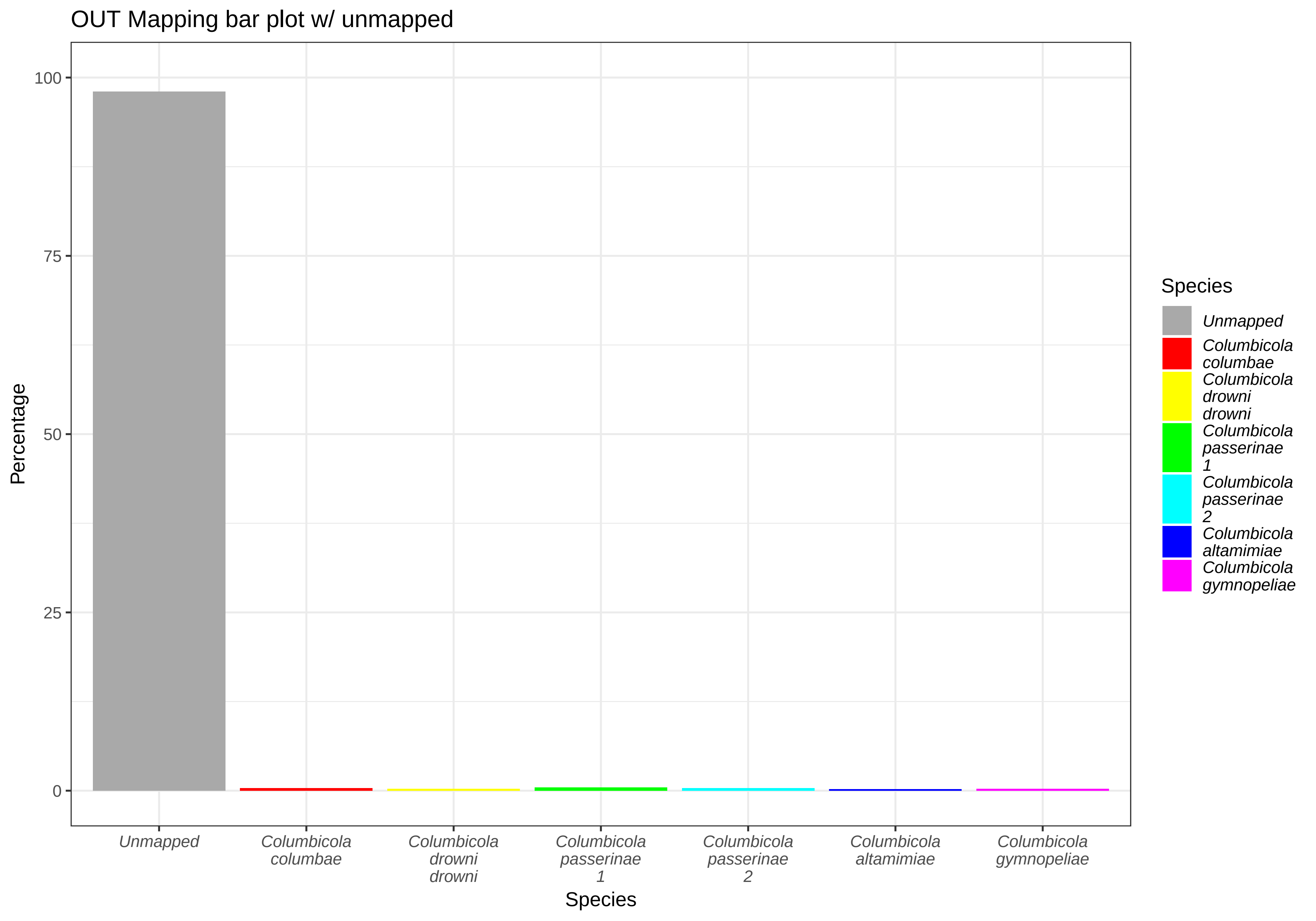

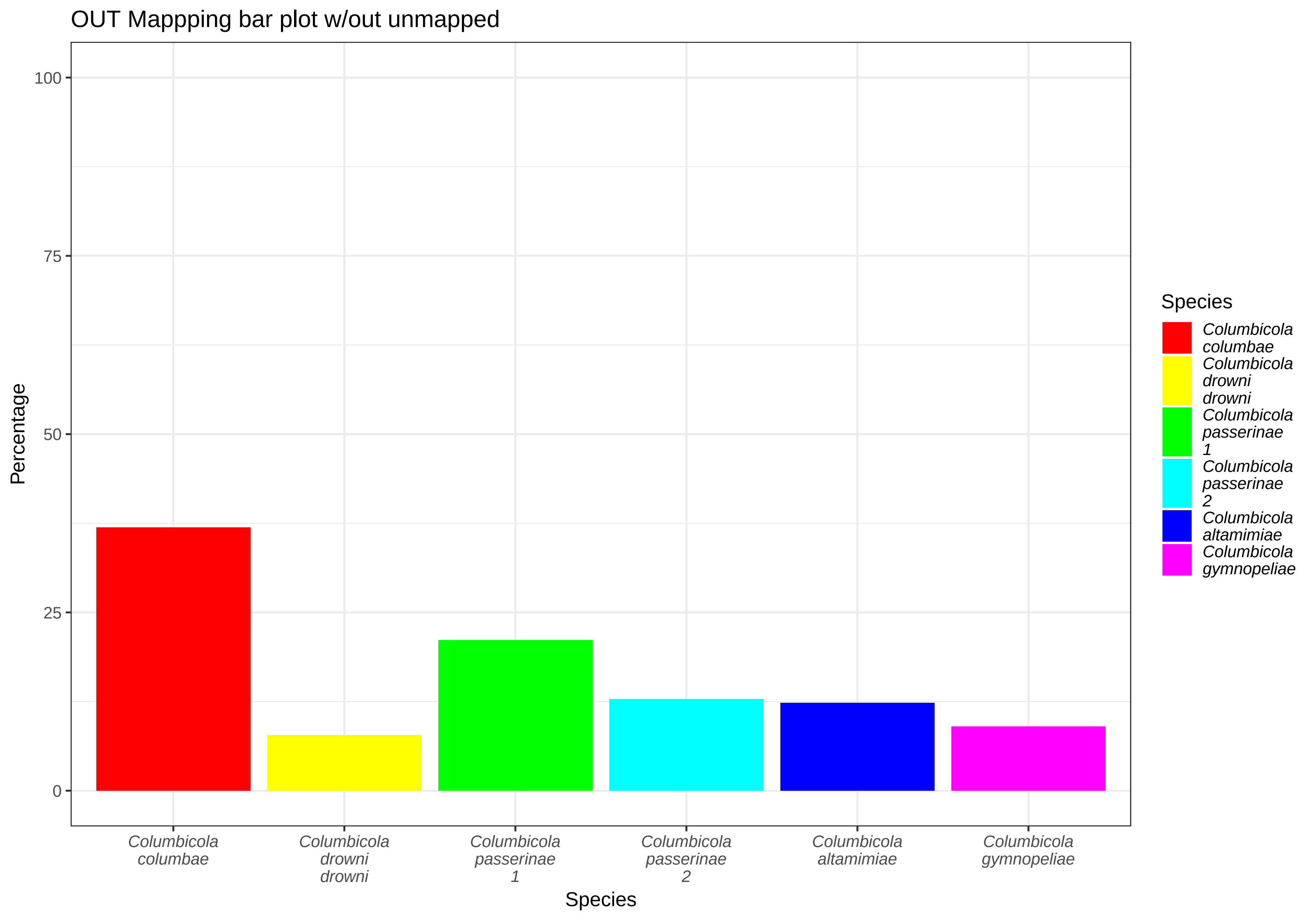

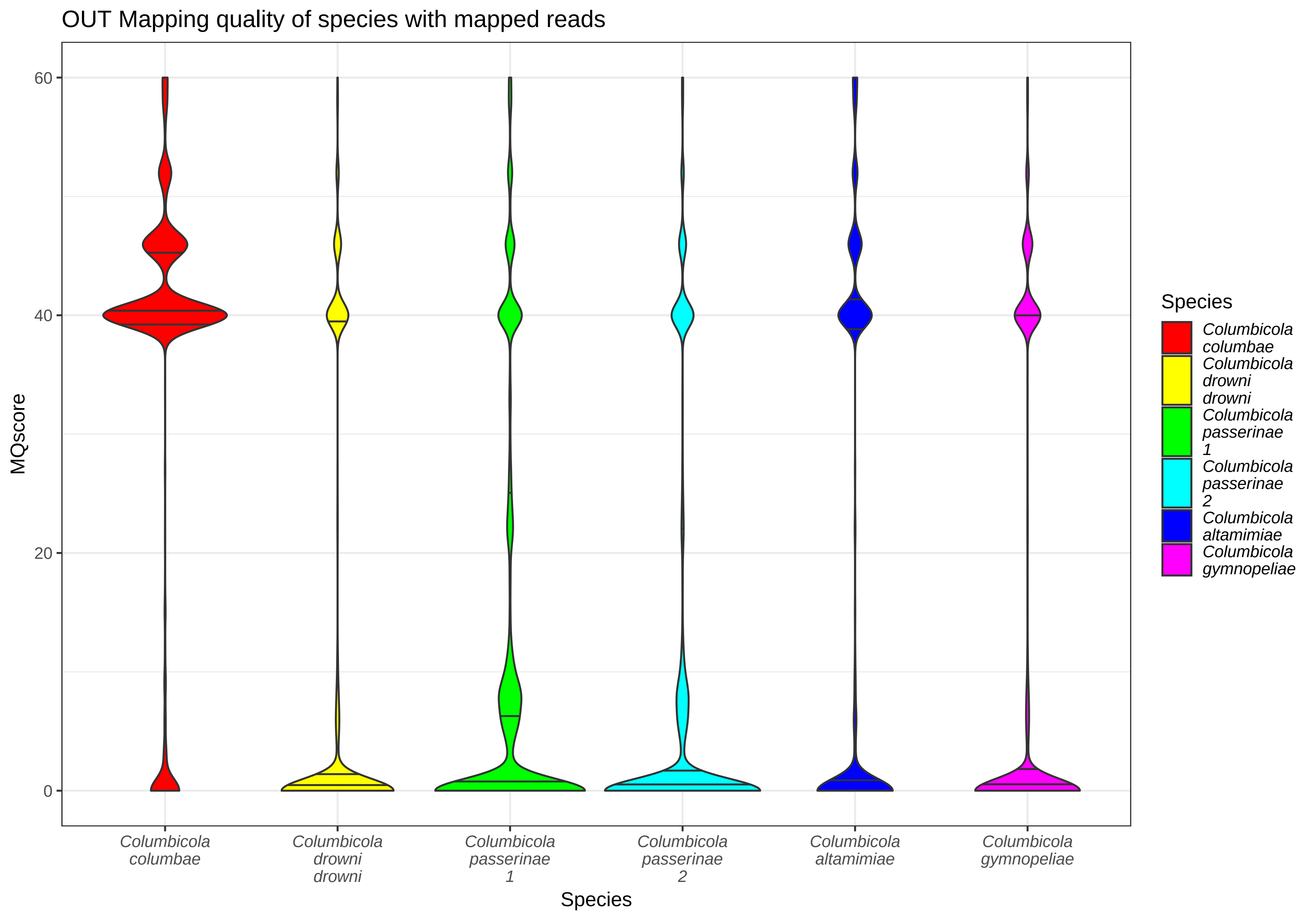

### OUT_plotMQ_Cosp_Cotal_11_2_2016_8.pdf

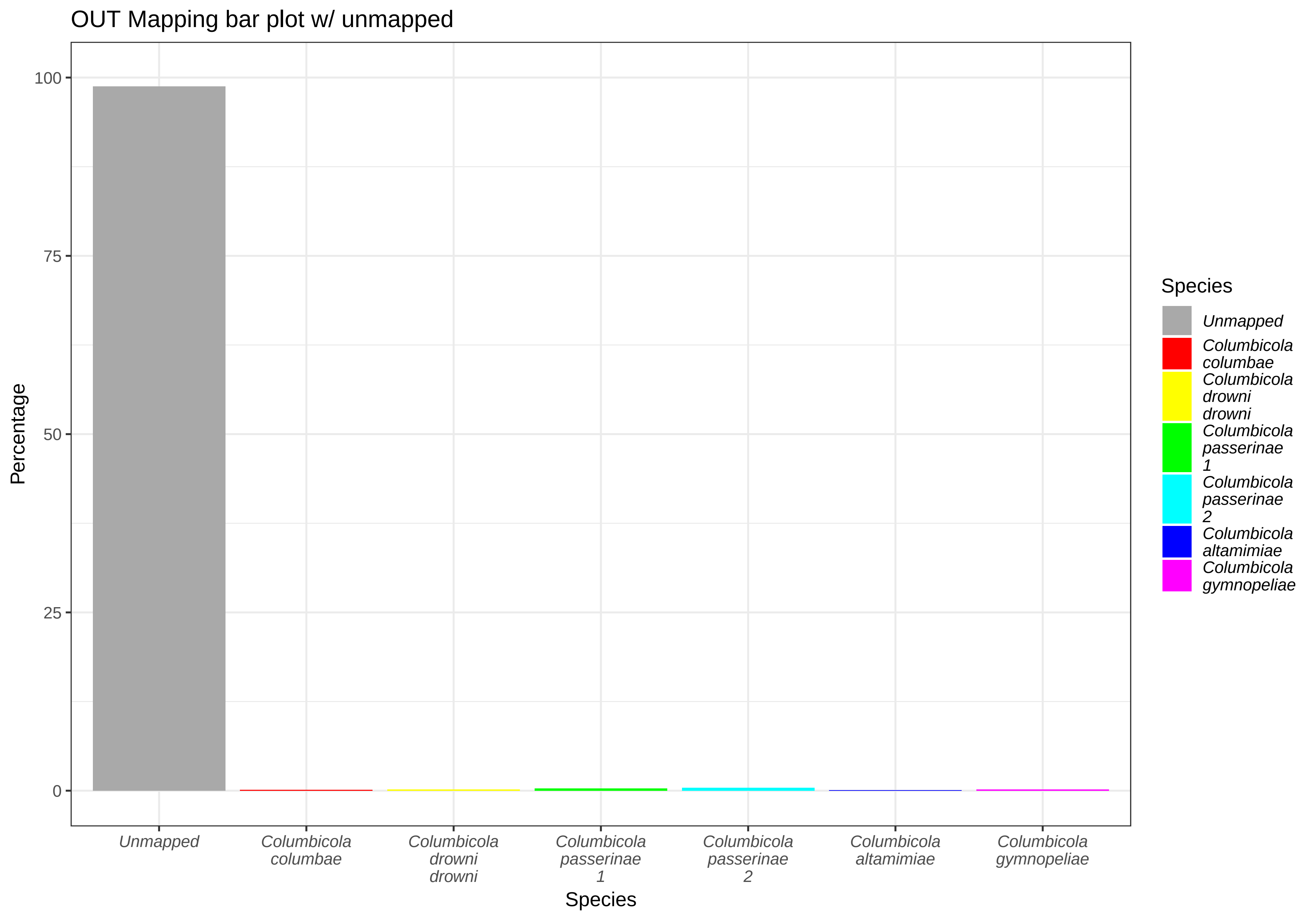

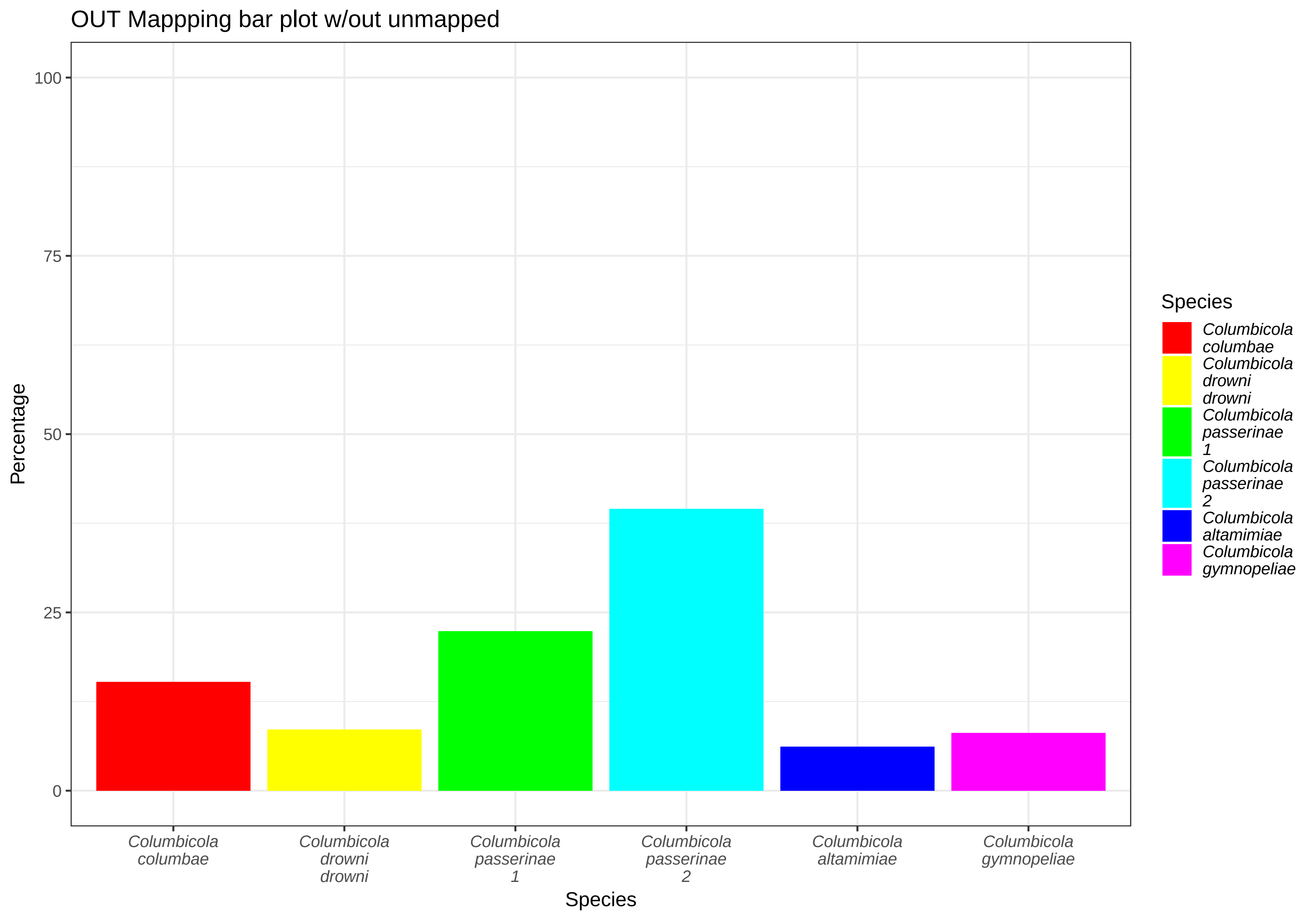

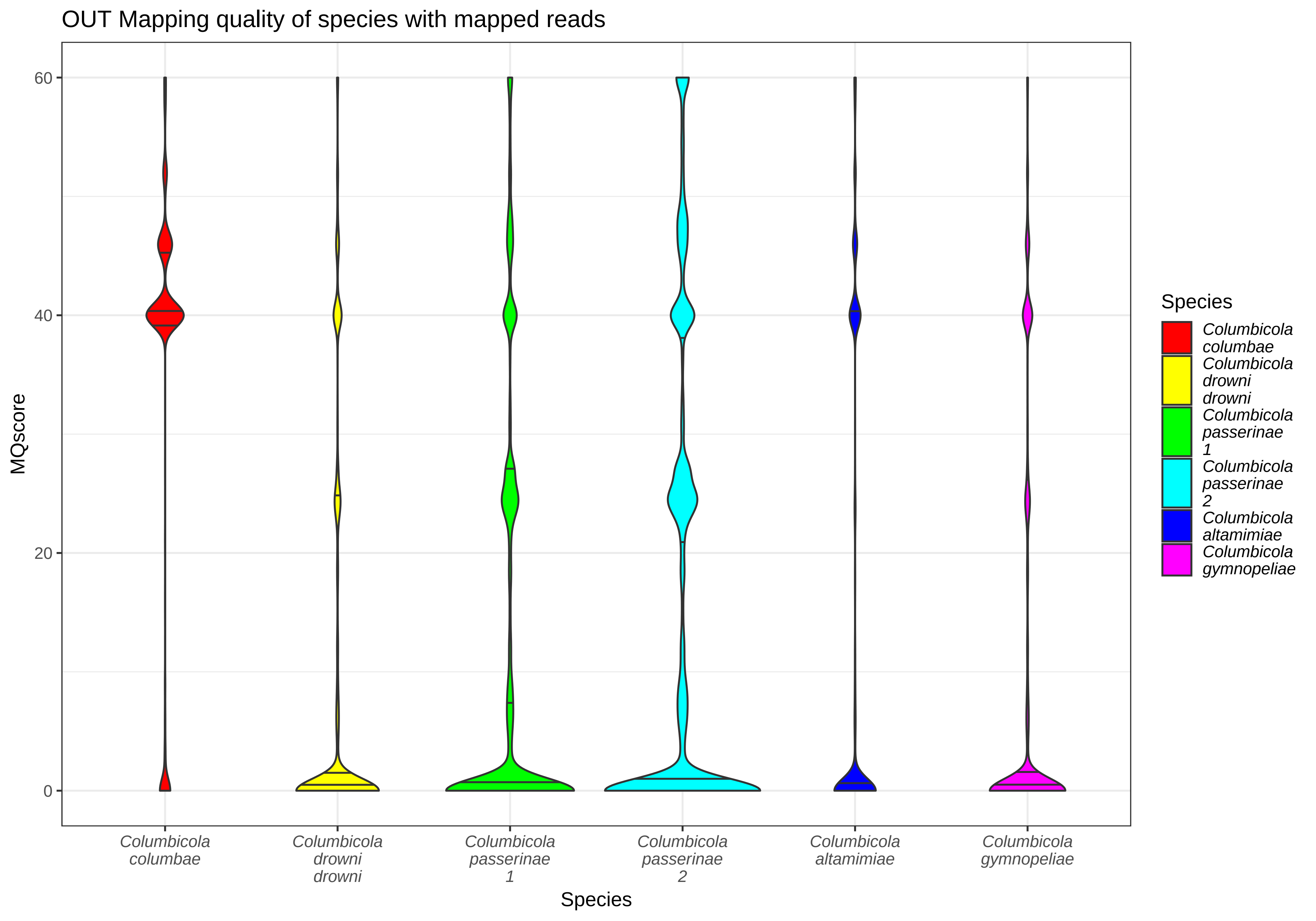

### OUT_plotMQ_Cosp_Cotal_11_2_2016_15.pdf

OUT Mapping bar plot w/ unmapped

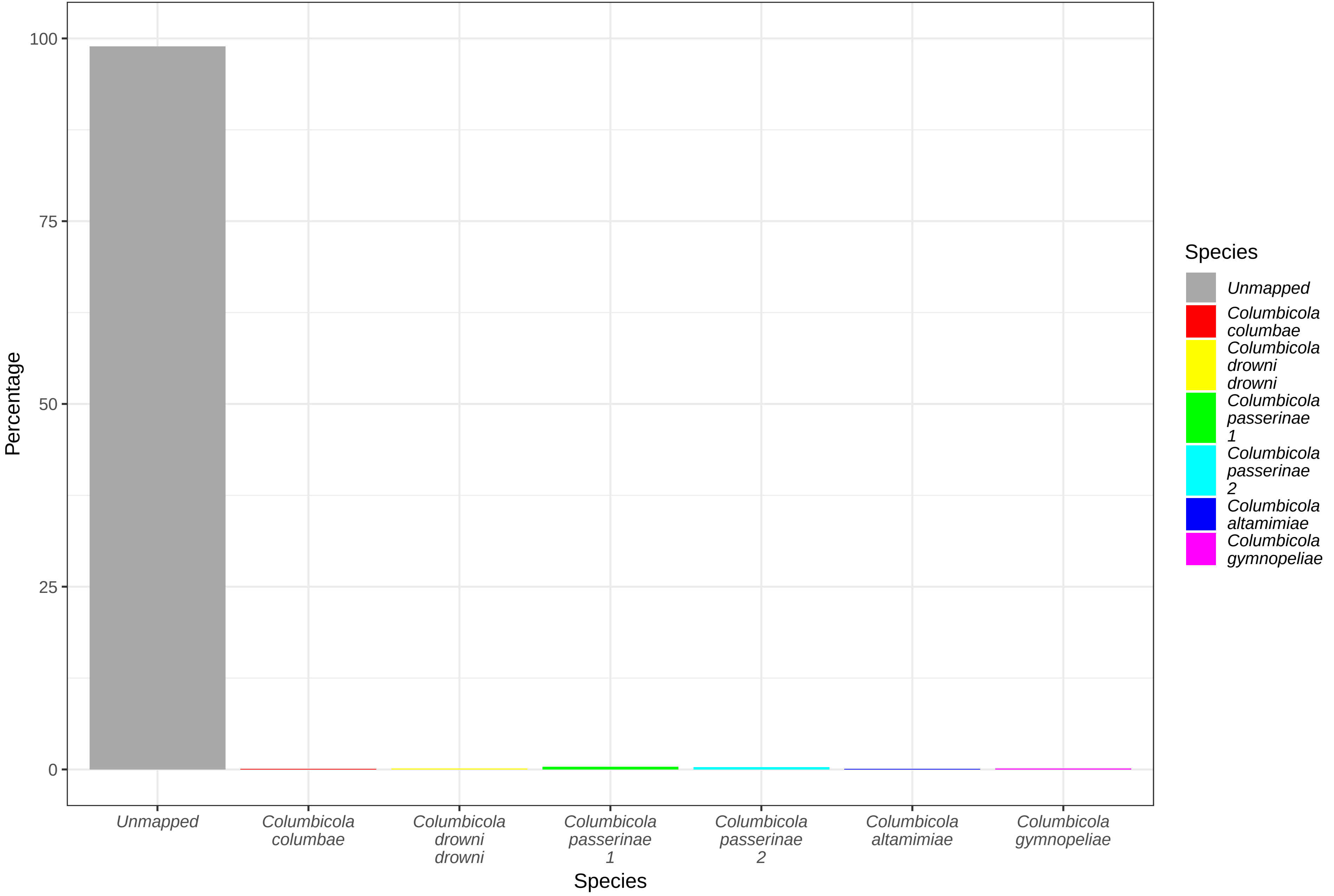

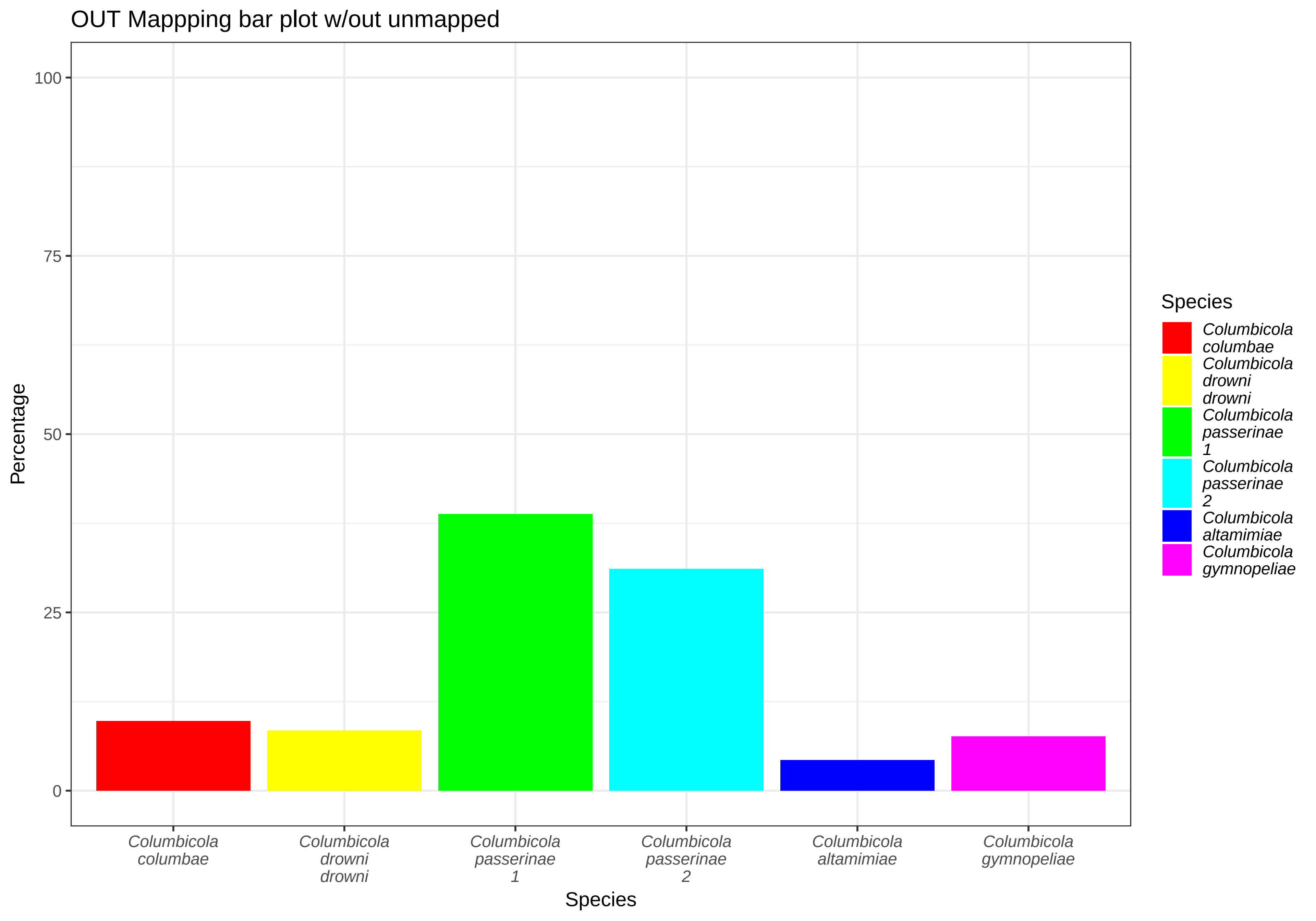

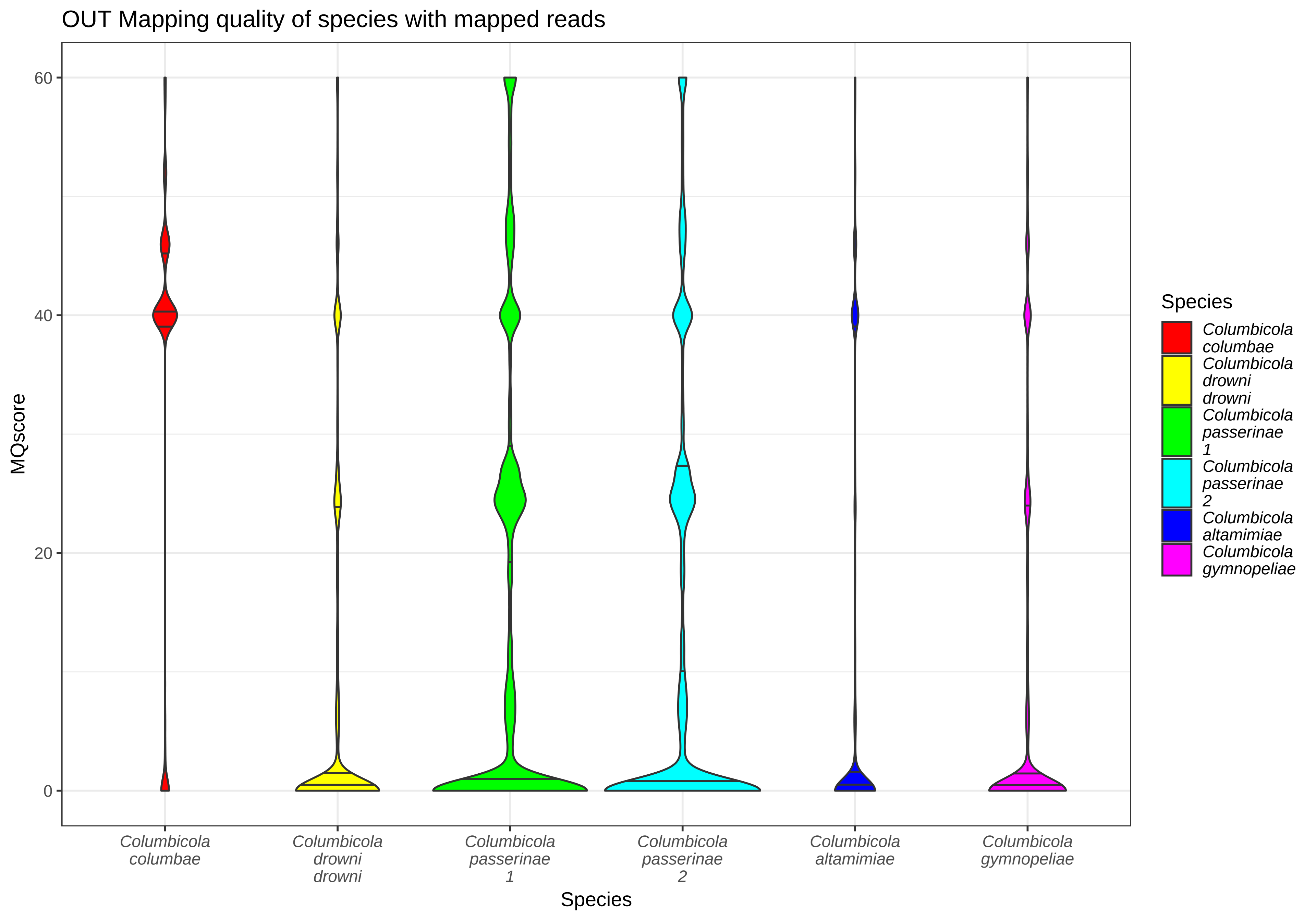

### OUT_plotMQ_Cosp_Cotal_11_7_2016_2.pdf

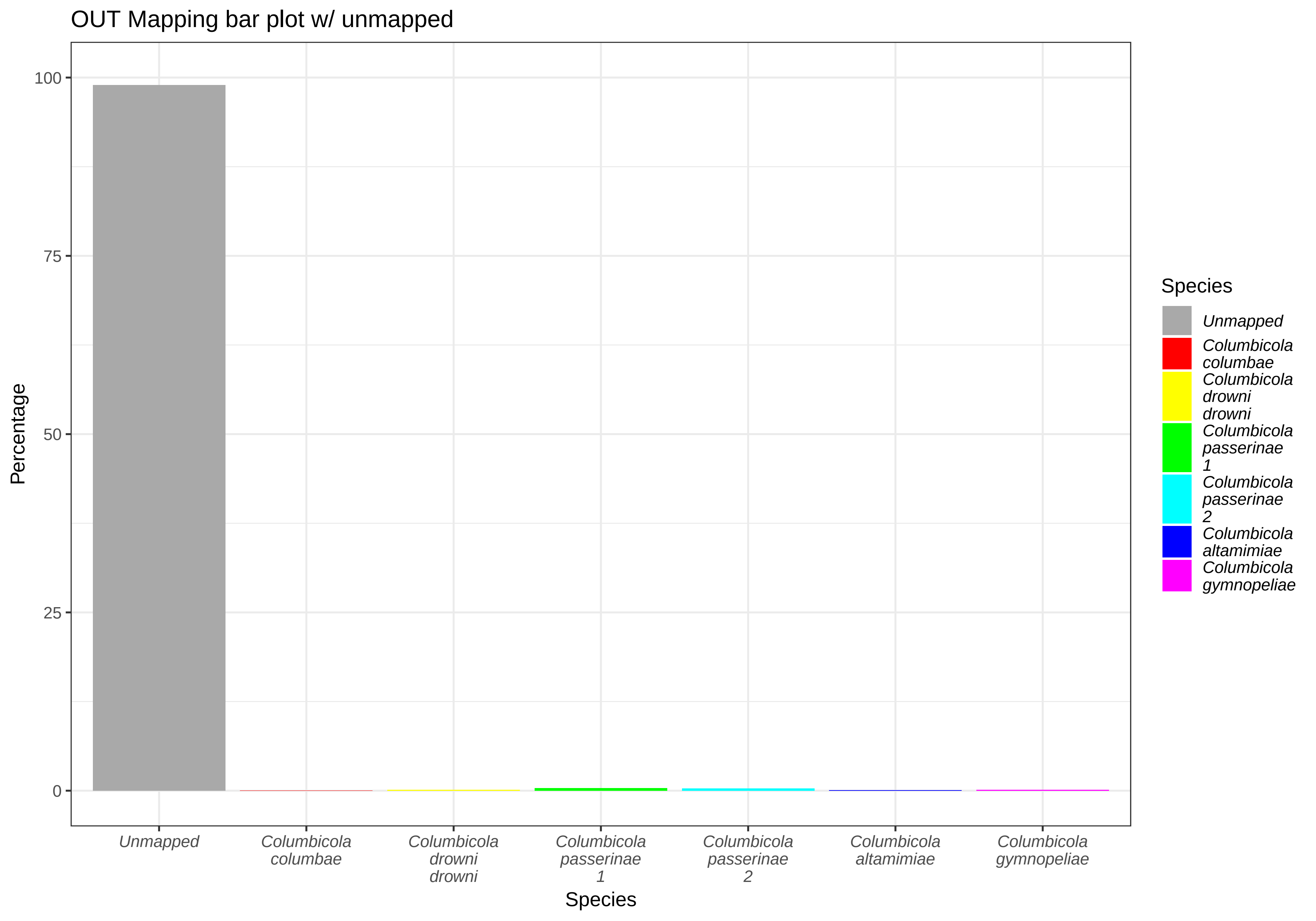

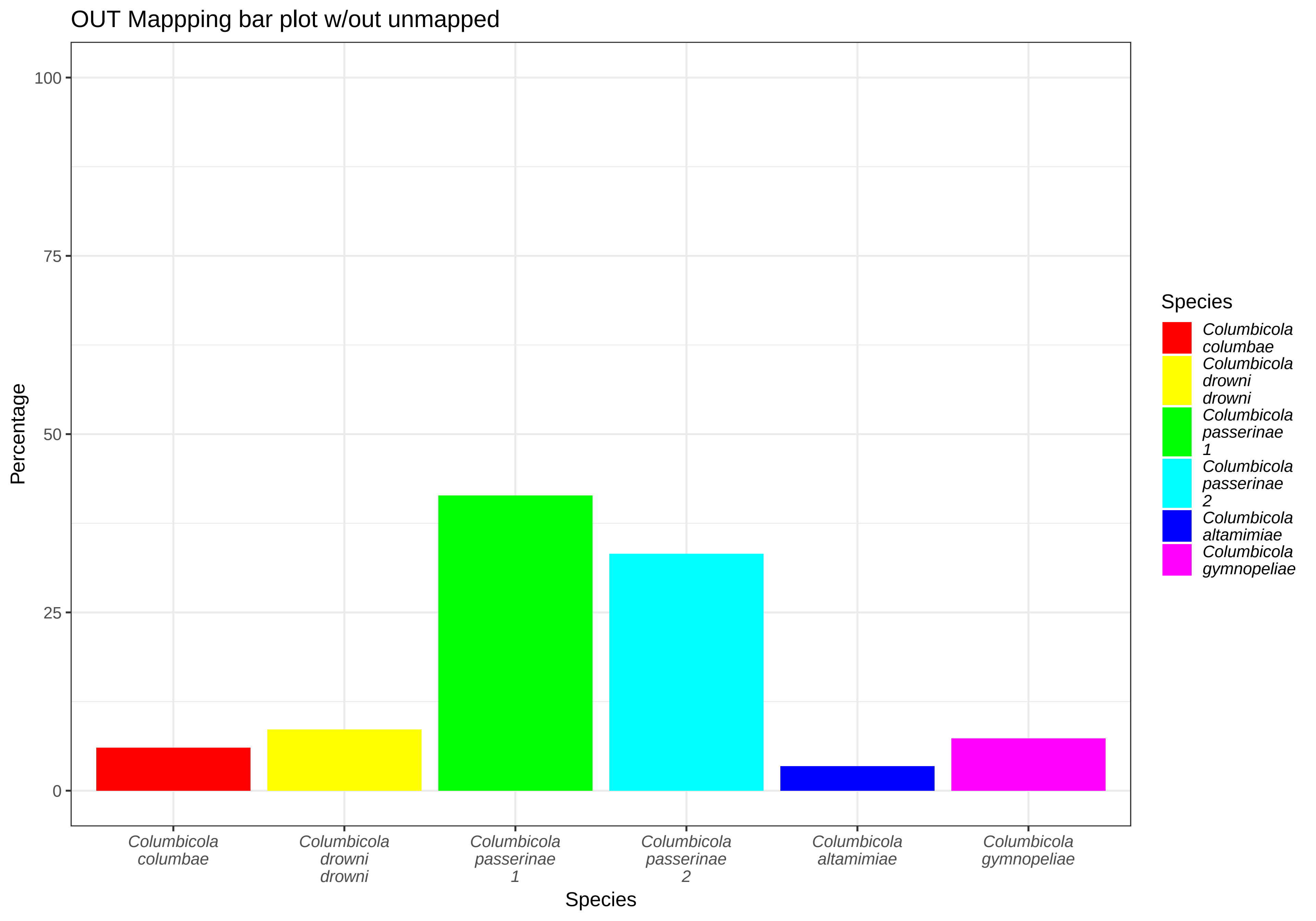

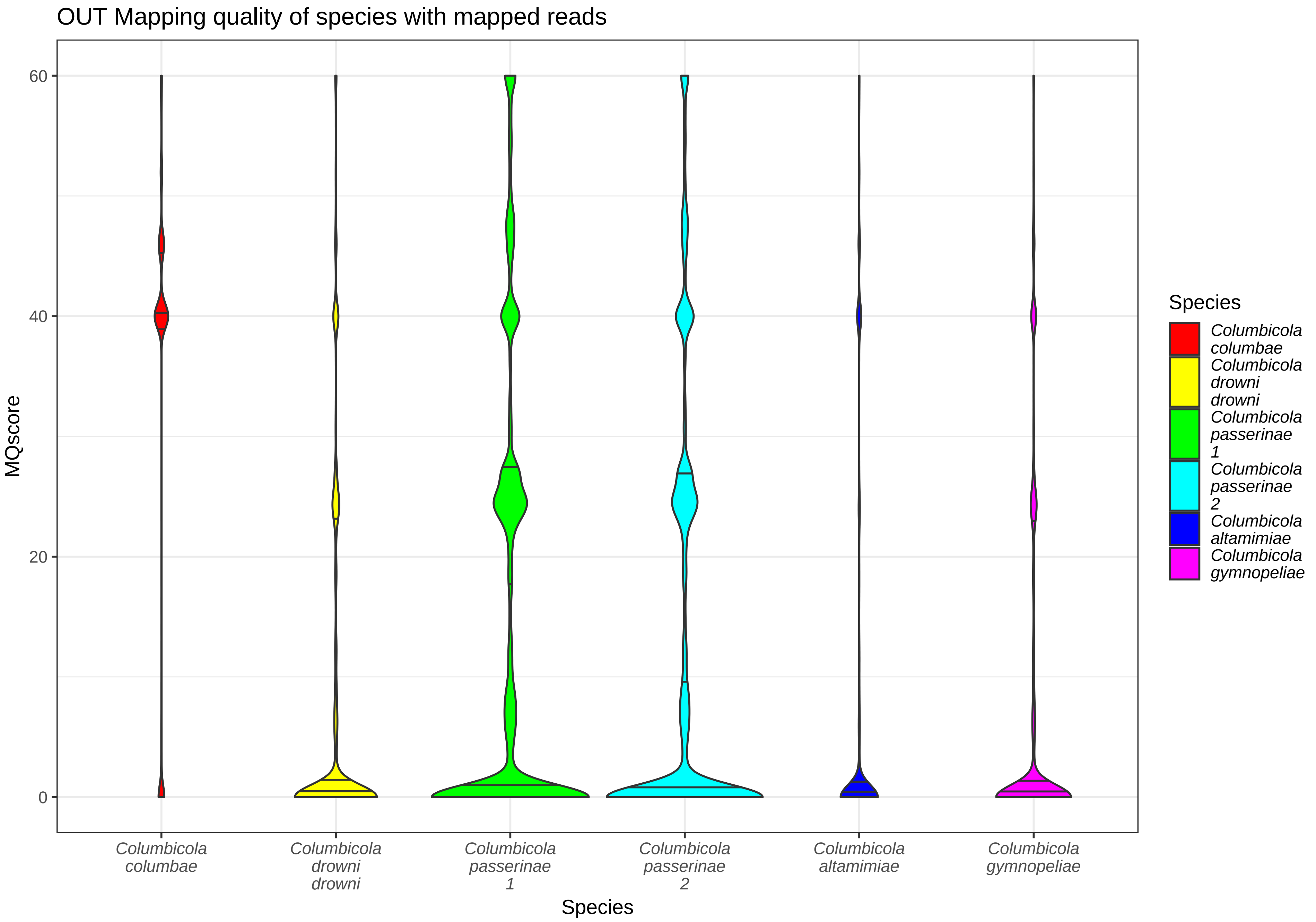

### OUT_plotMQ_Cosp_Cotal_11_7_2016_10.pdf

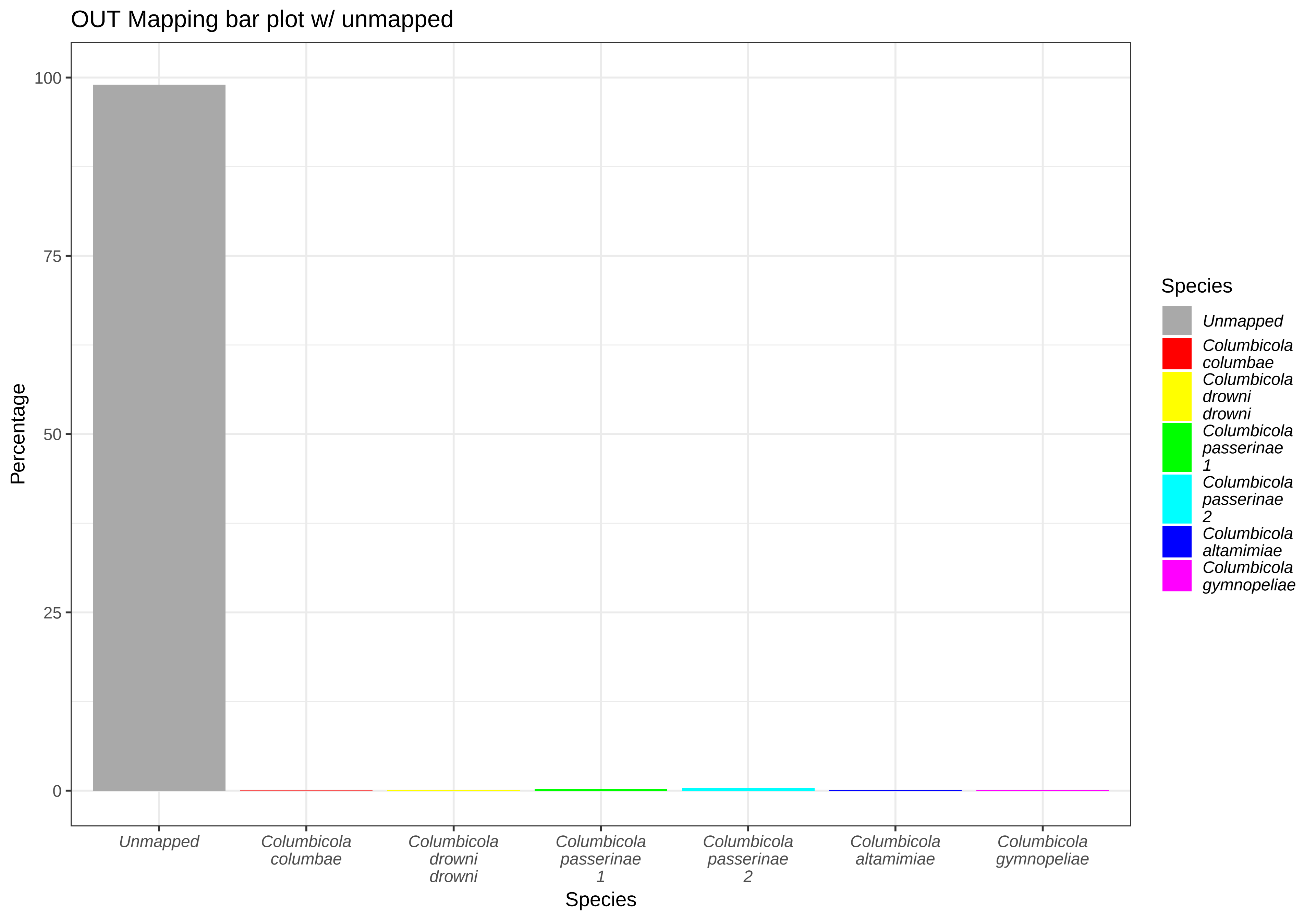

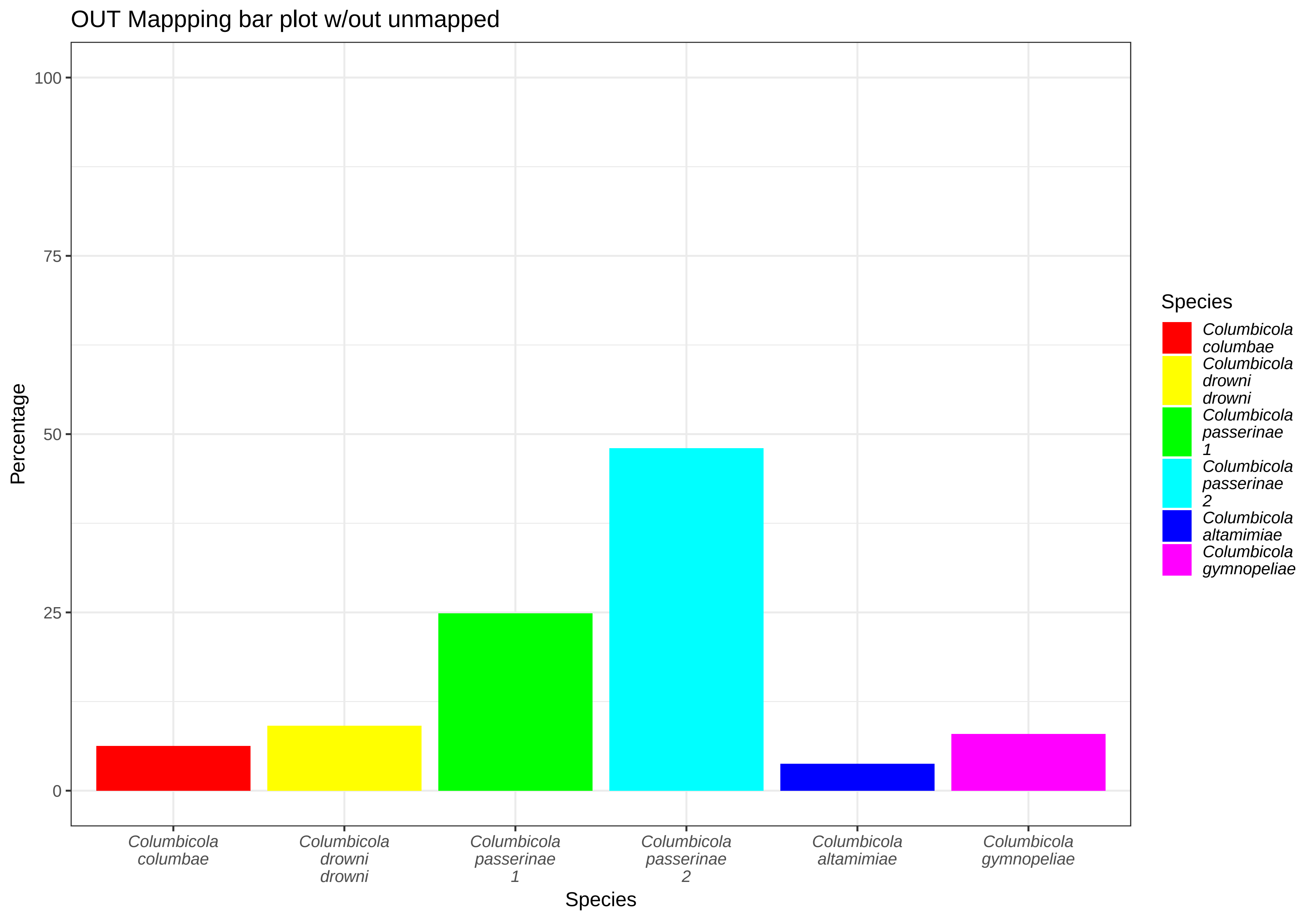

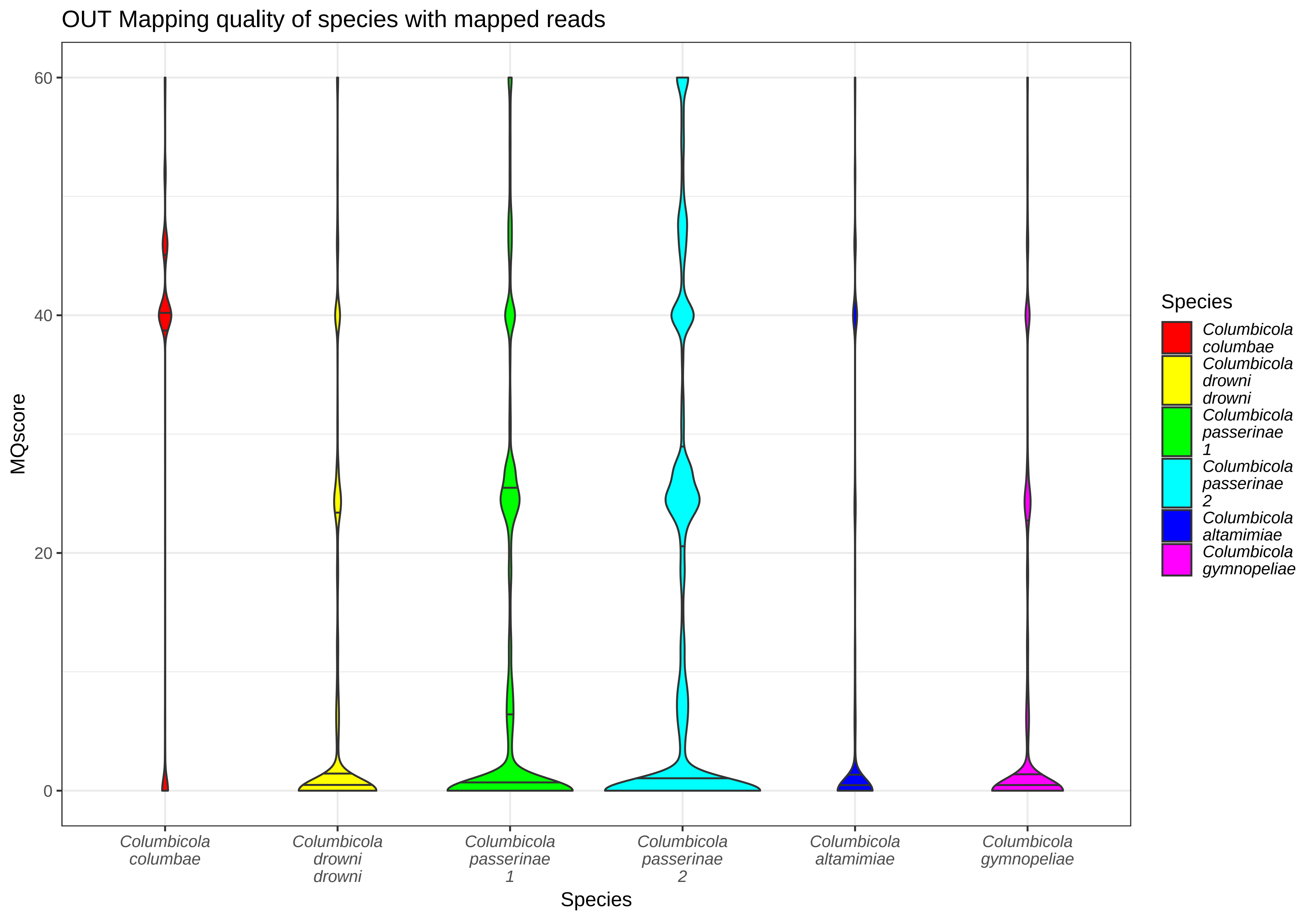

### OUT_plotMQ_Cosp_Meaym_11_2_2016_5.pdf

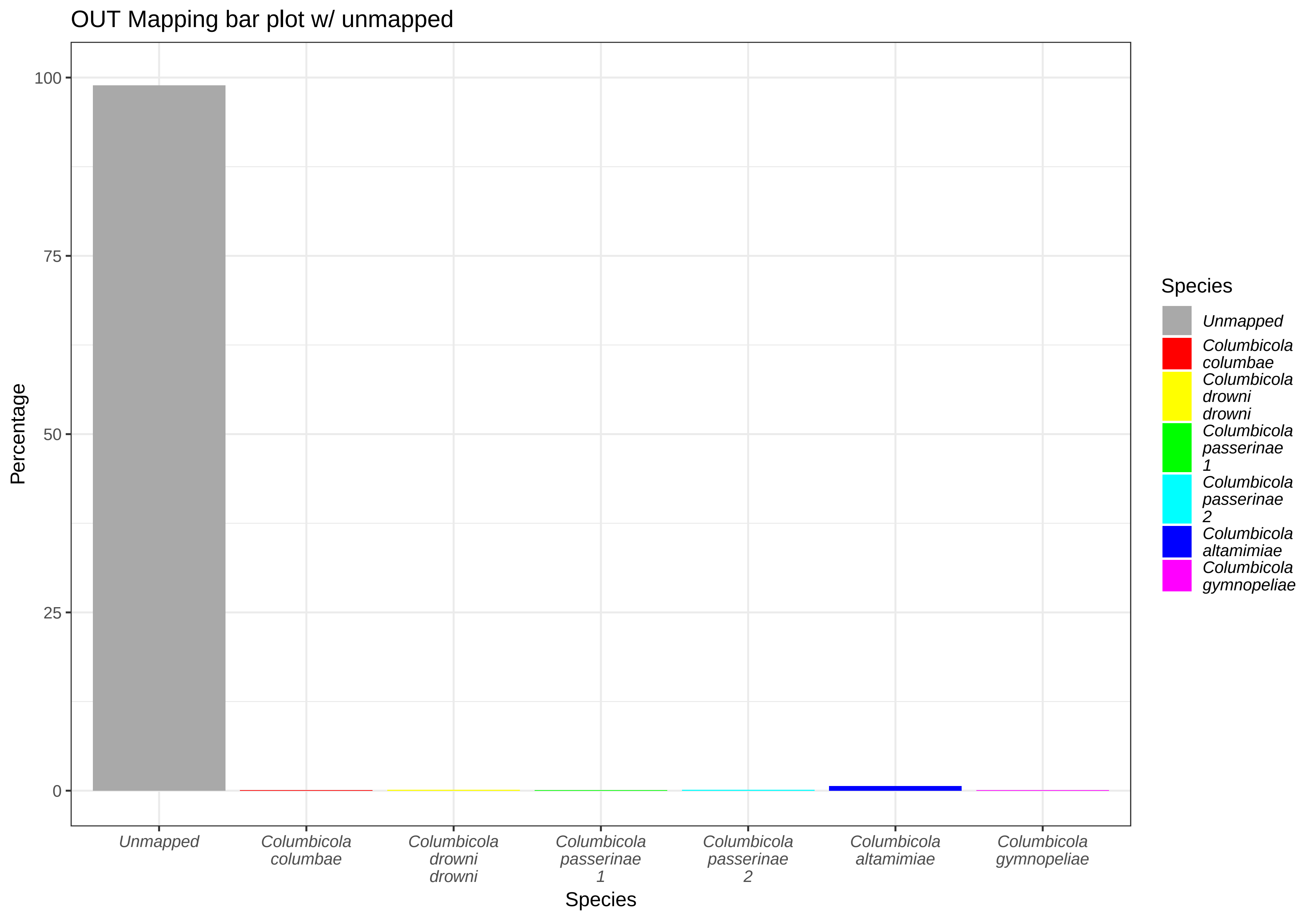

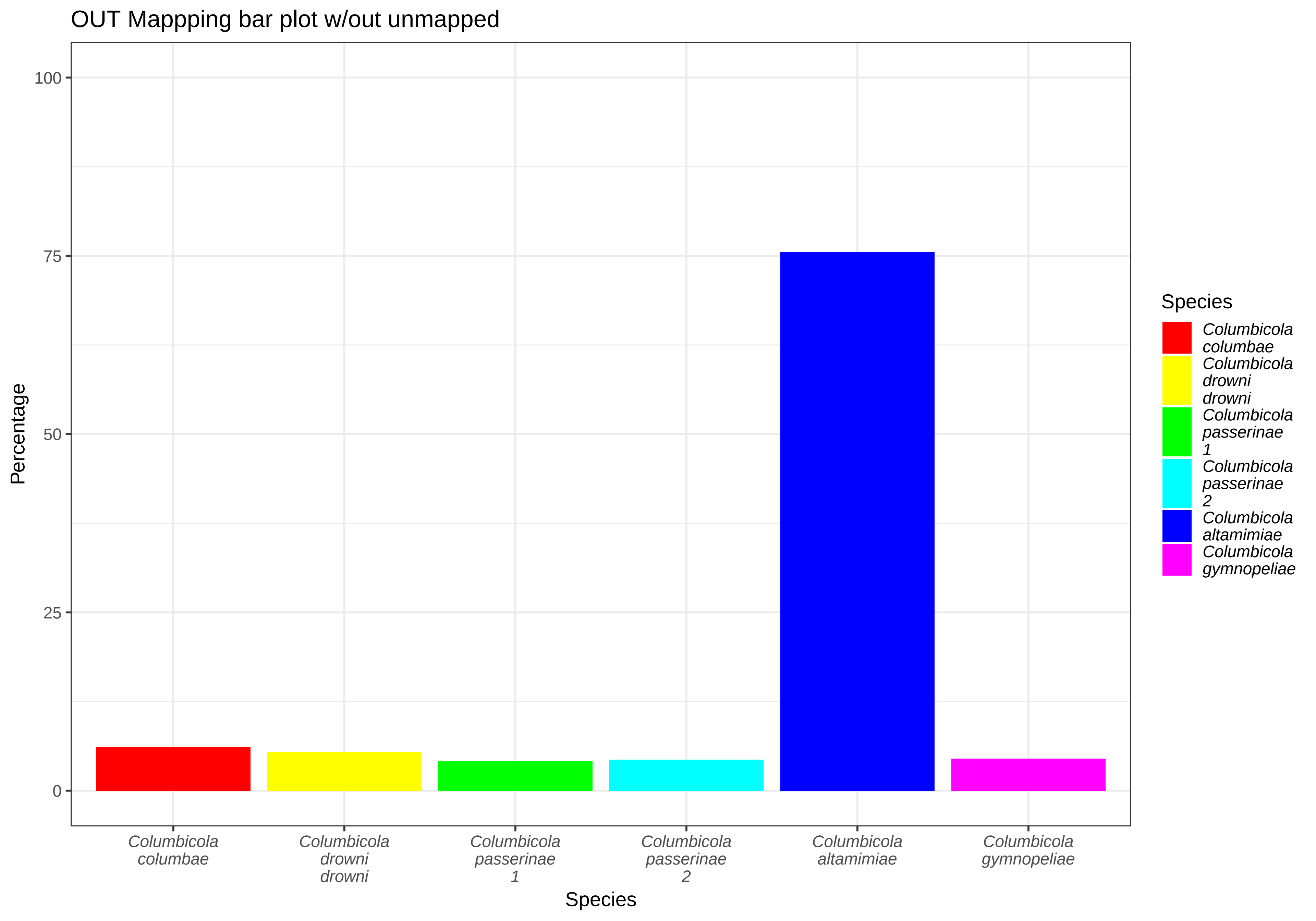

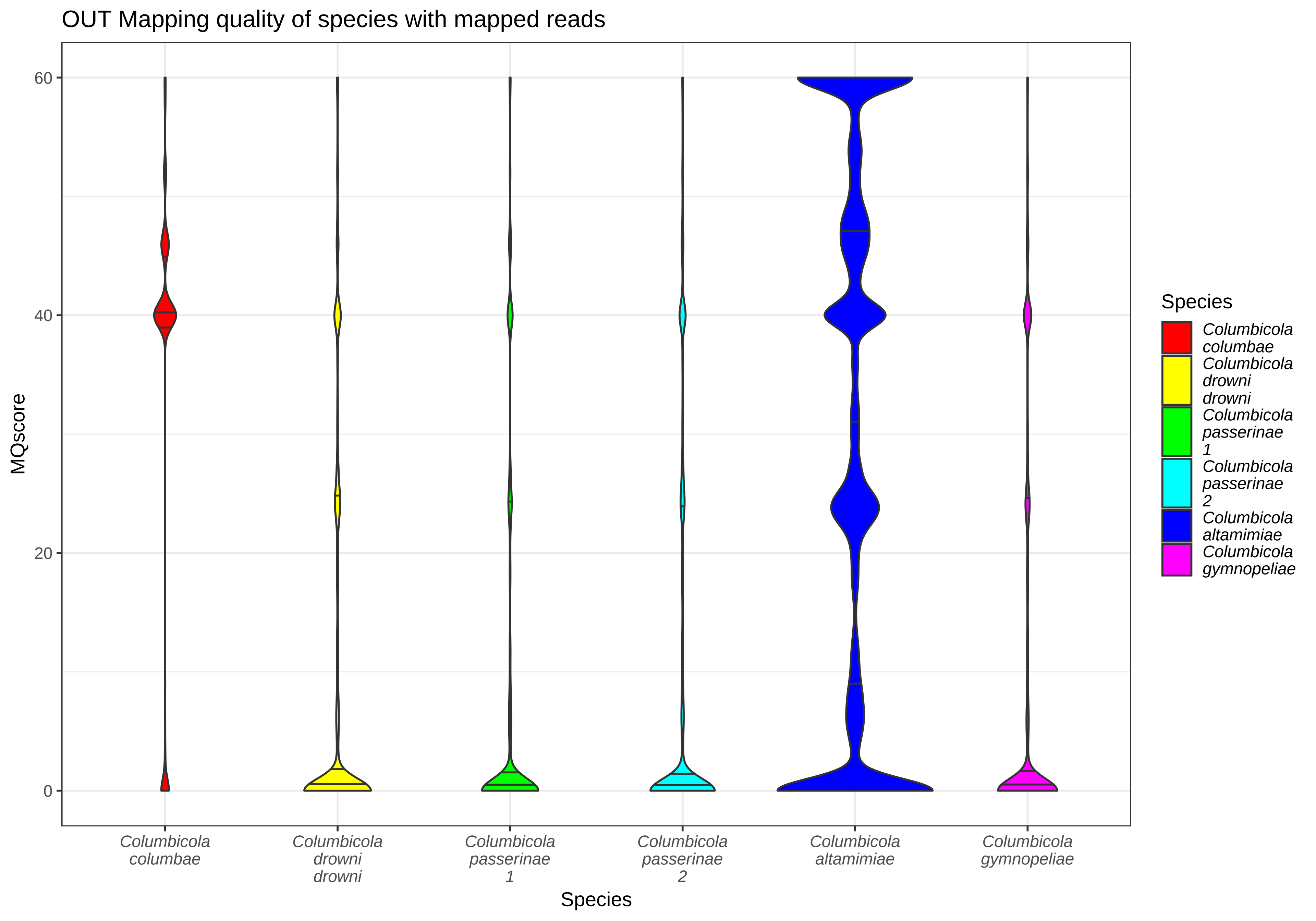

### OUT_plotMQ_Cosp_Meaym_11_2_2016_7.pdf

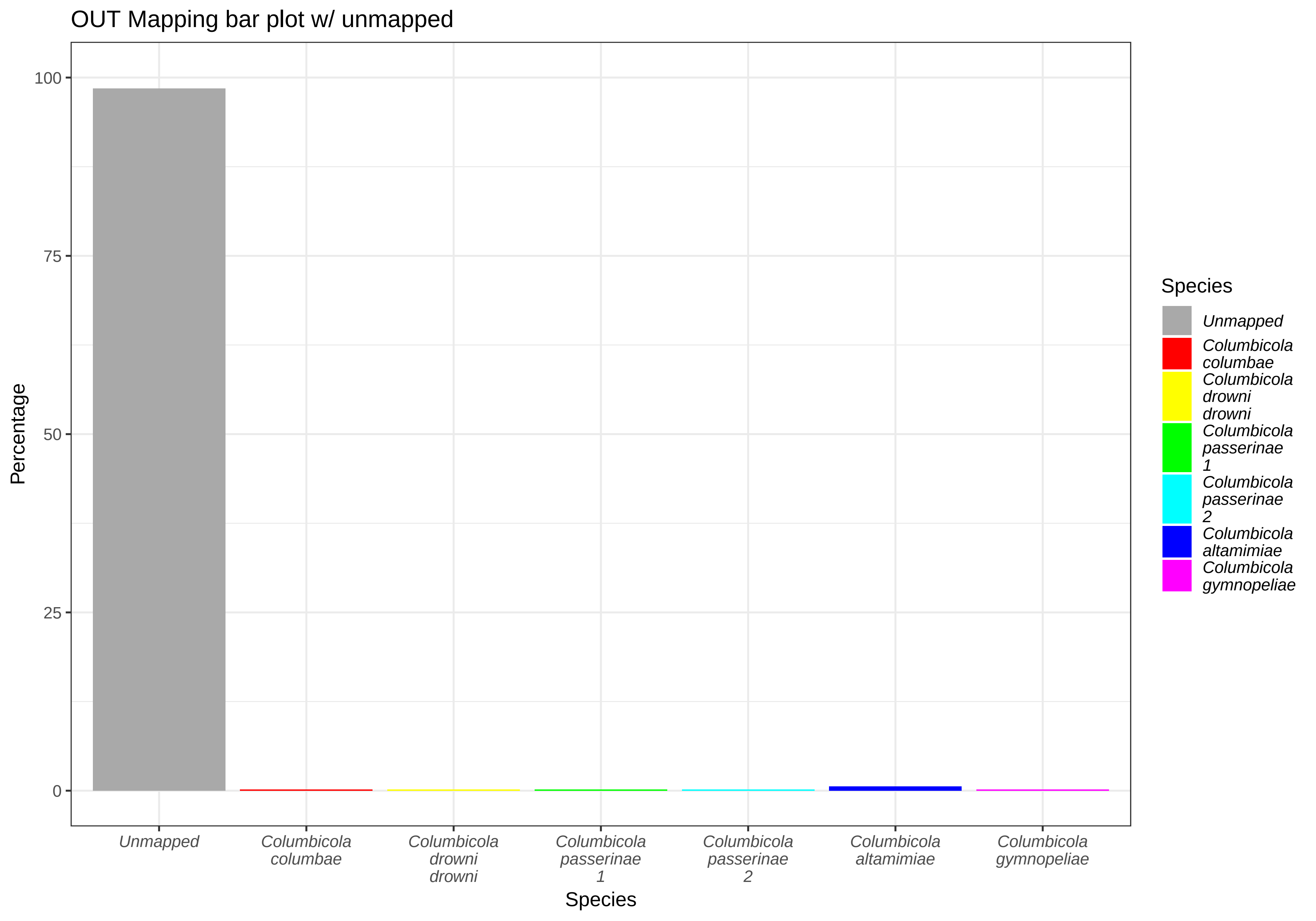

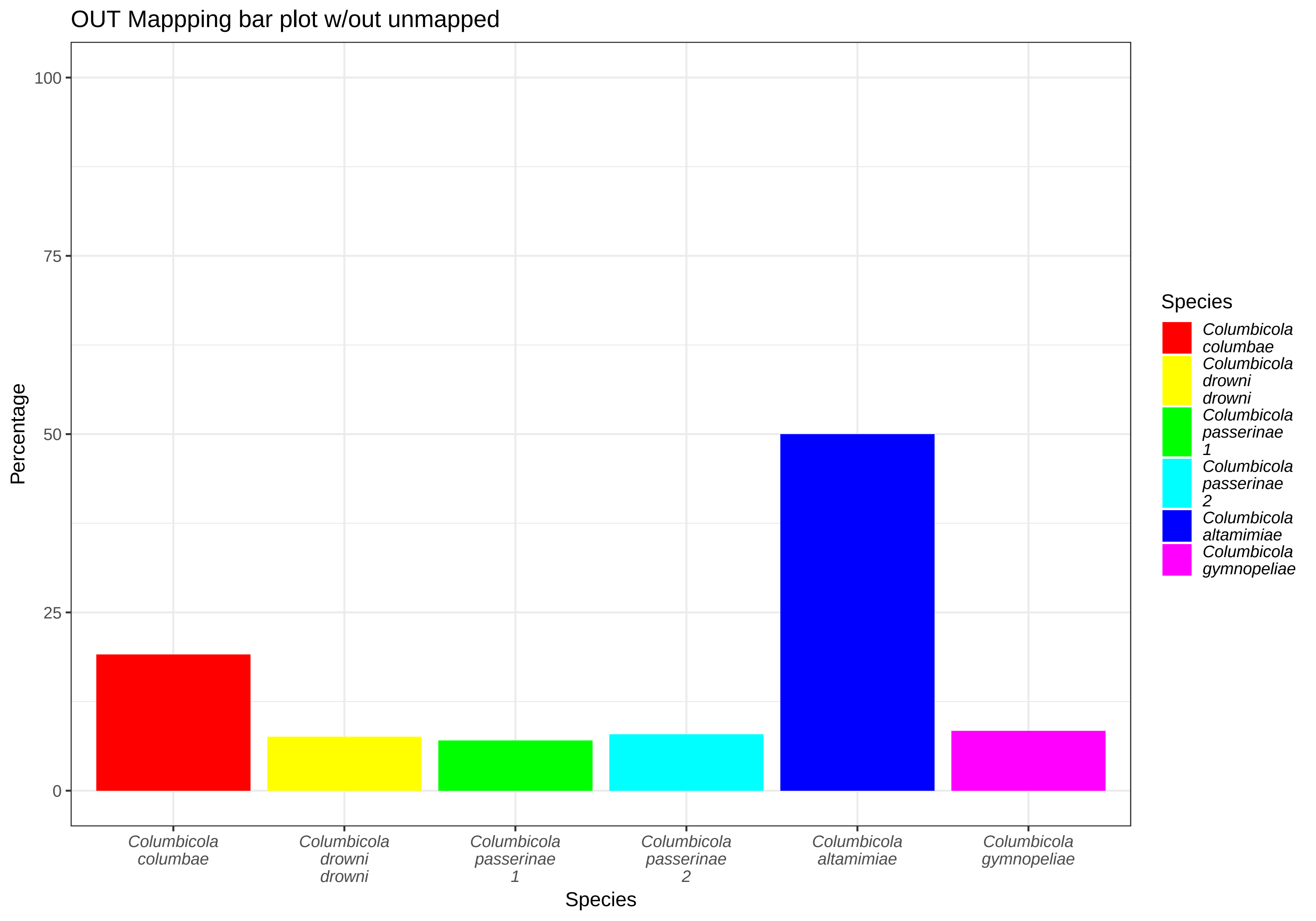

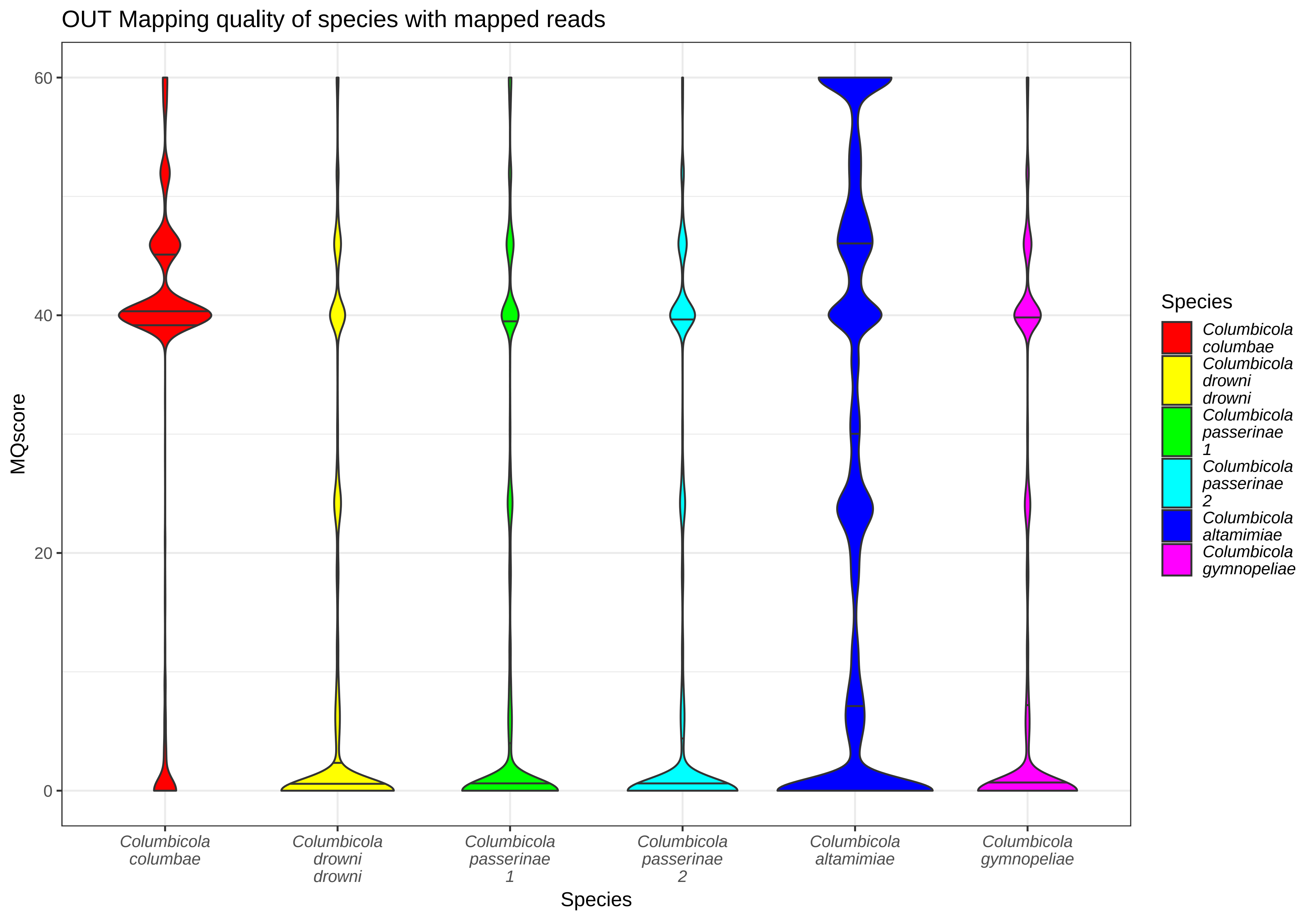

### OUT_plotMQ_Cosp_Mecec_11_2_2016_10.pdf

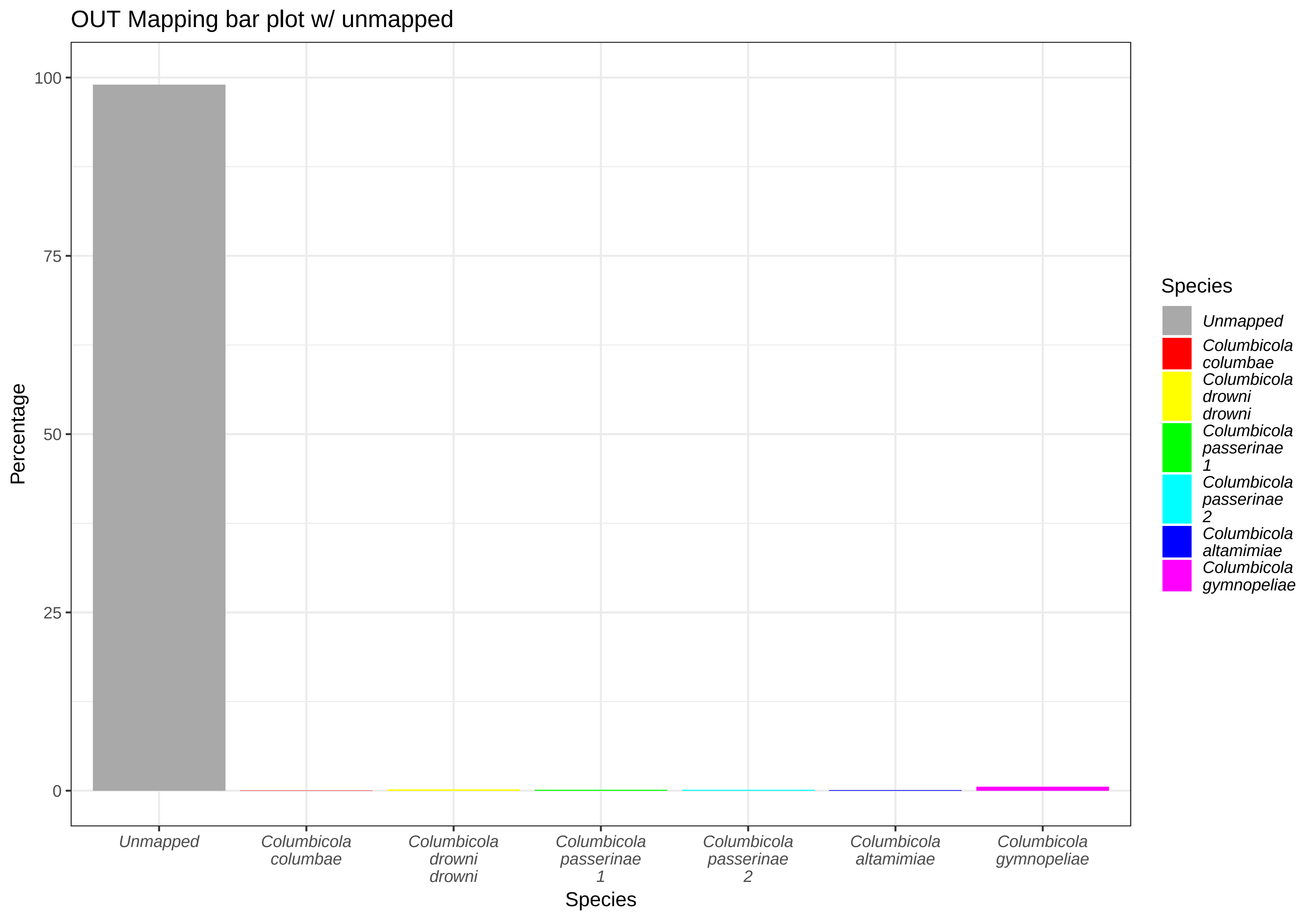

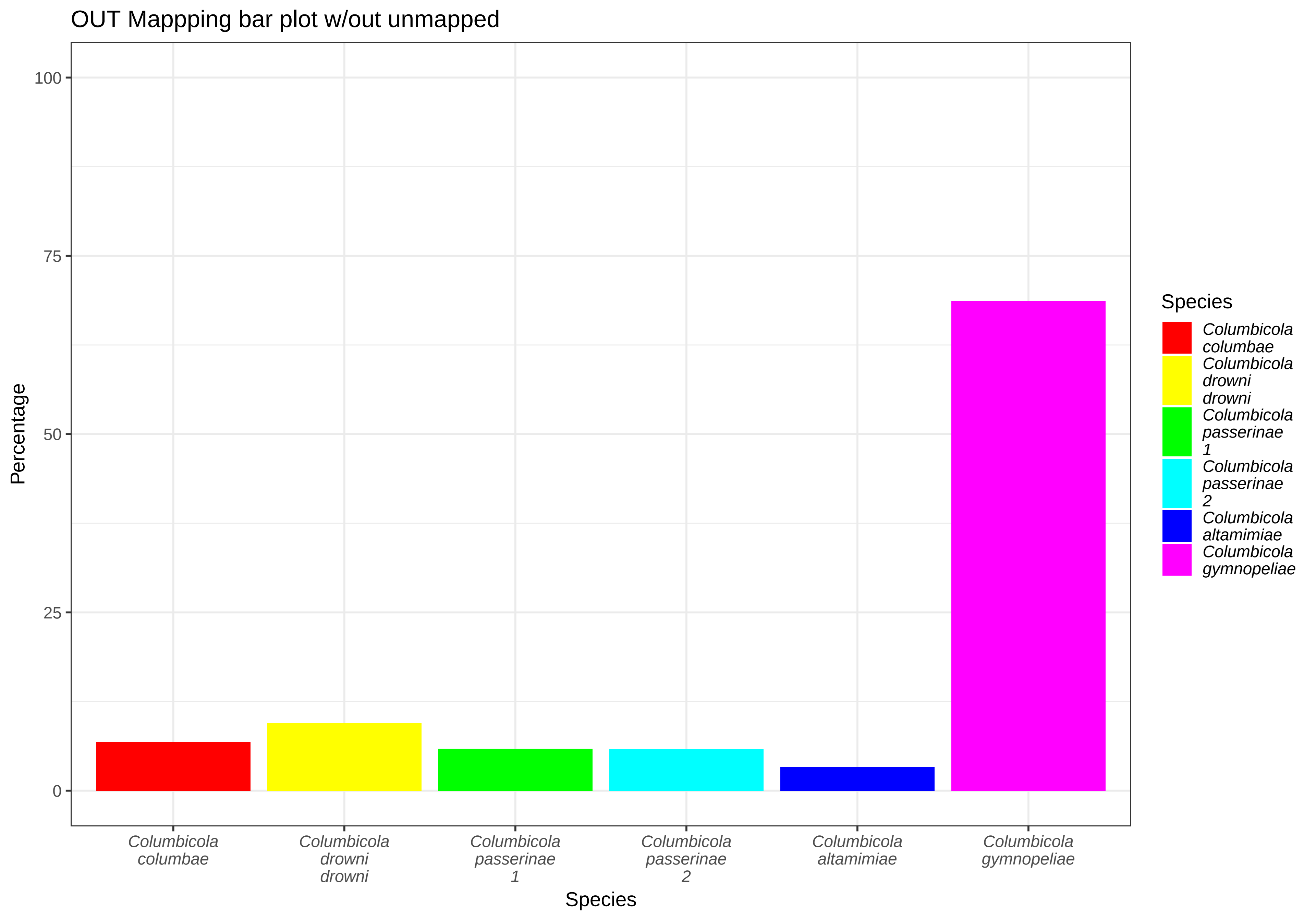

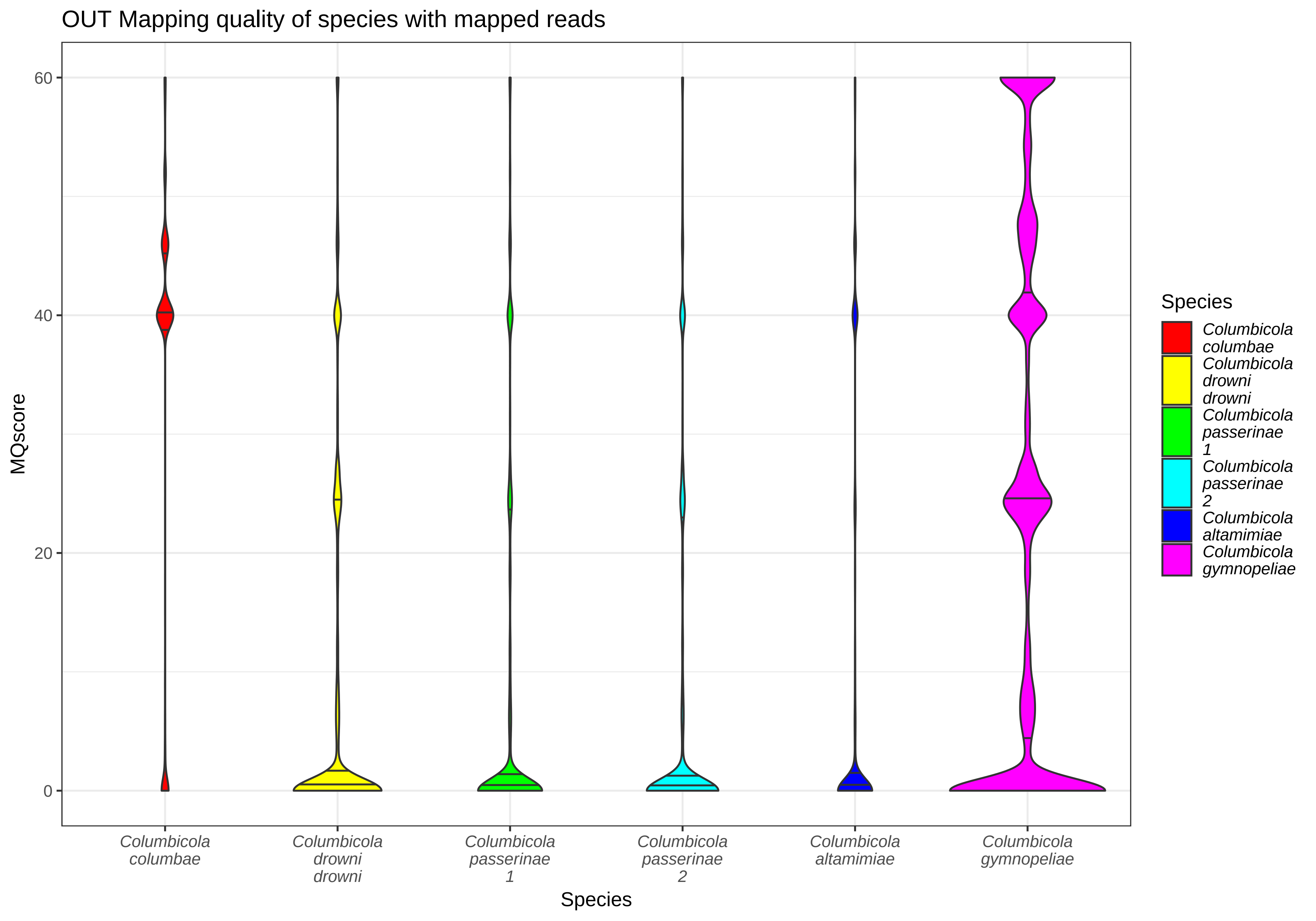

### OUT_plotMQ_Cosp_Mecec_11_2_2016_12.pdf

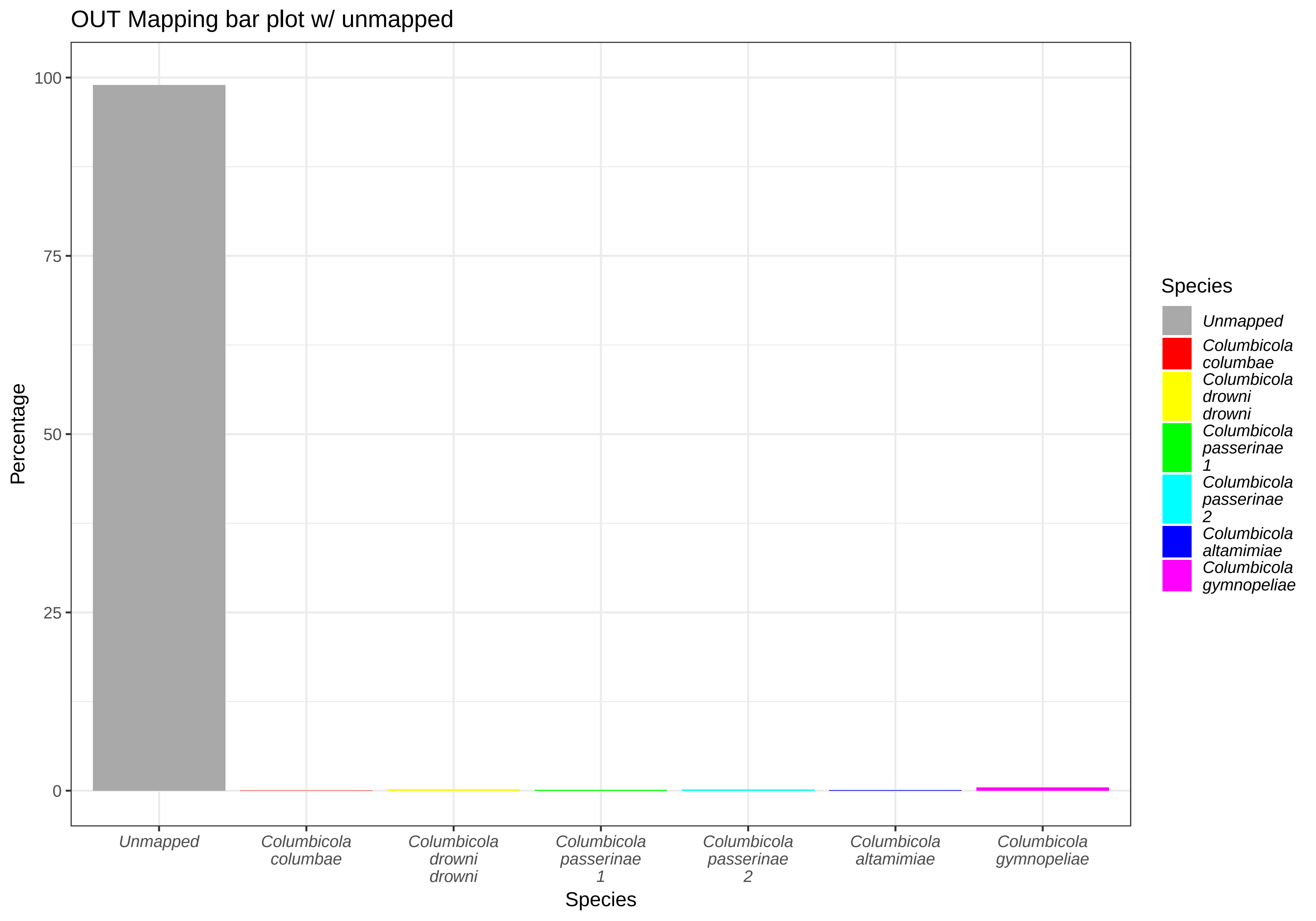

OUT Mappping bar plot w/out unmapped

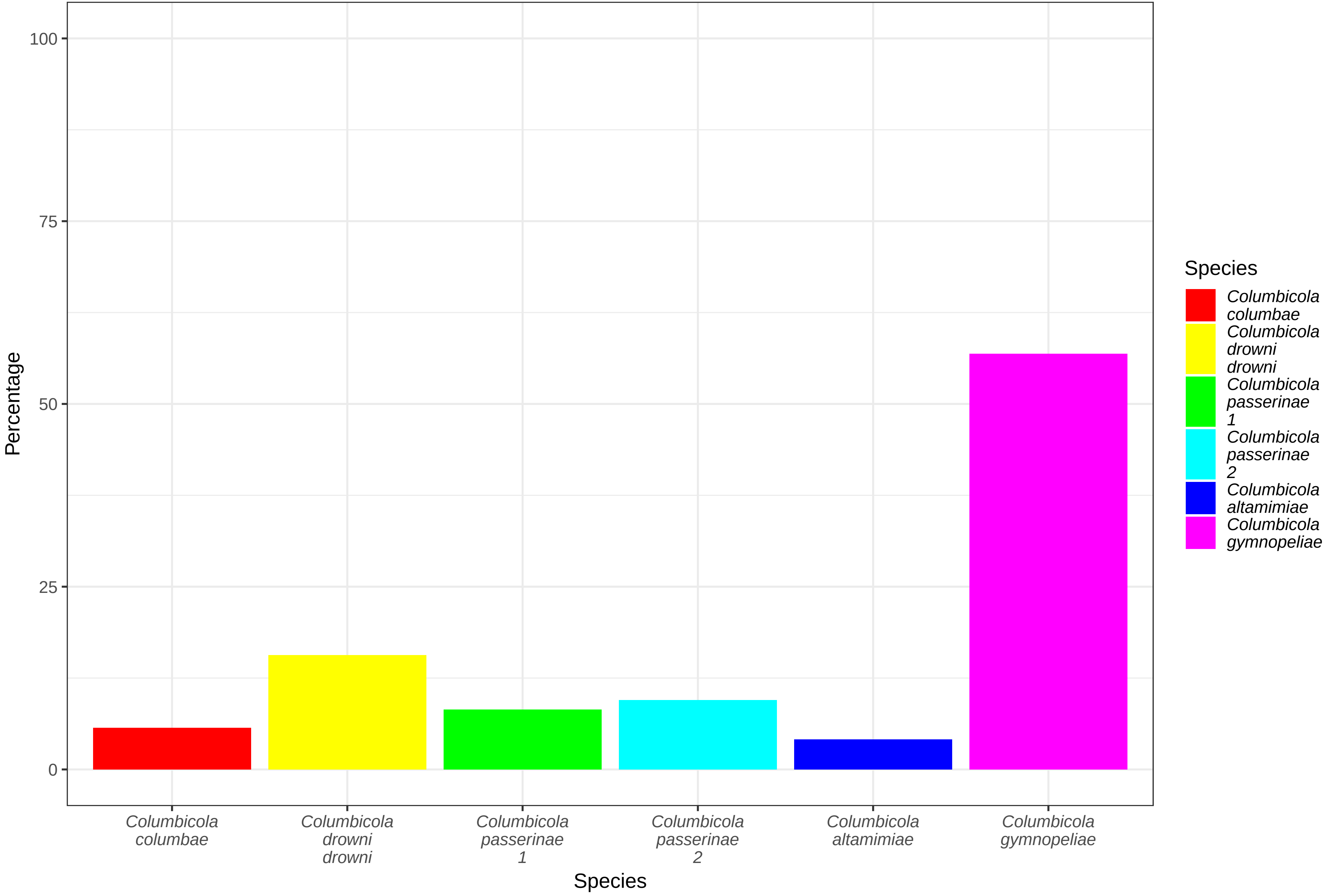

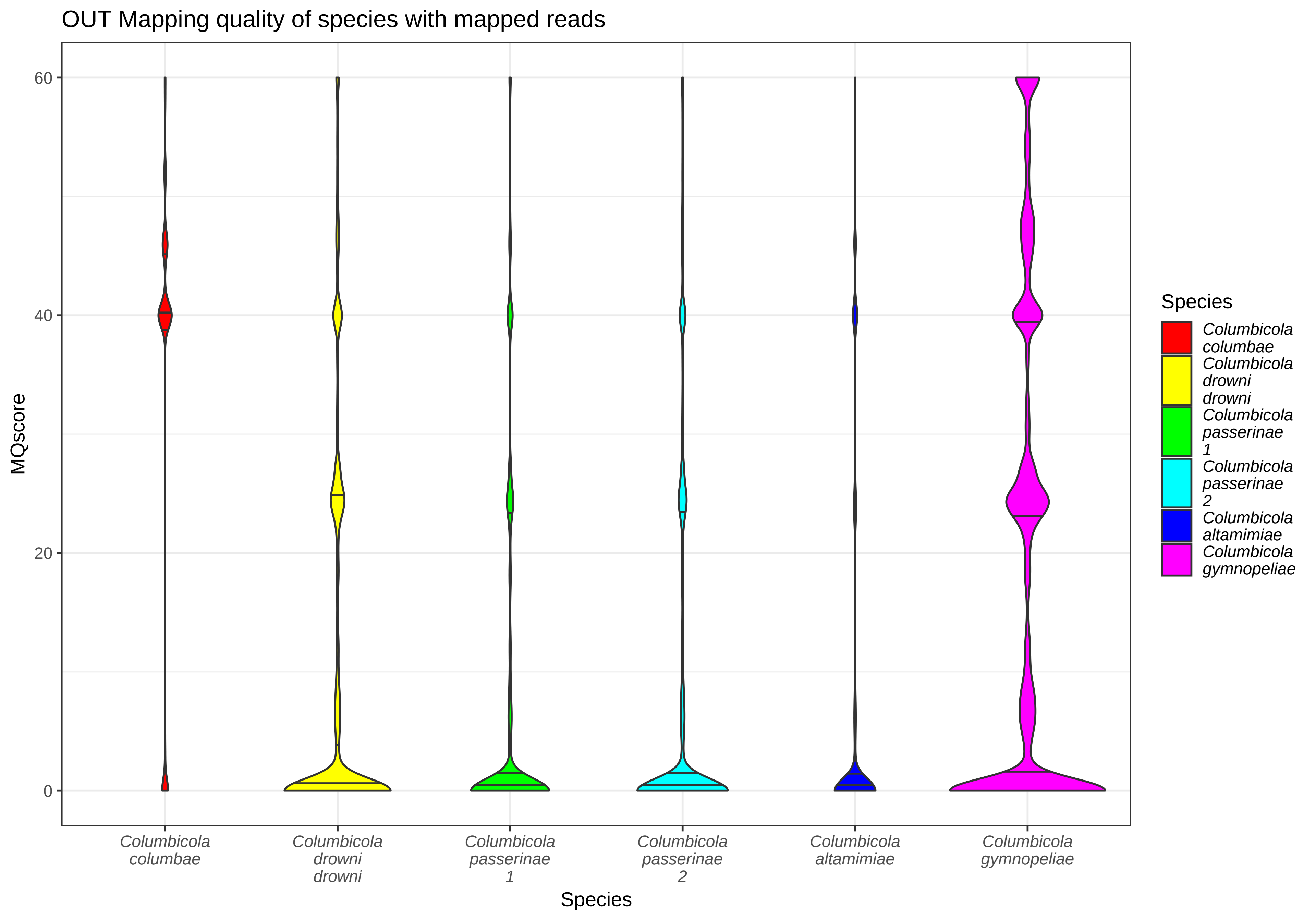

### OUT_plotMQ_Cosp_Memel_11_2_2016_6.pdf

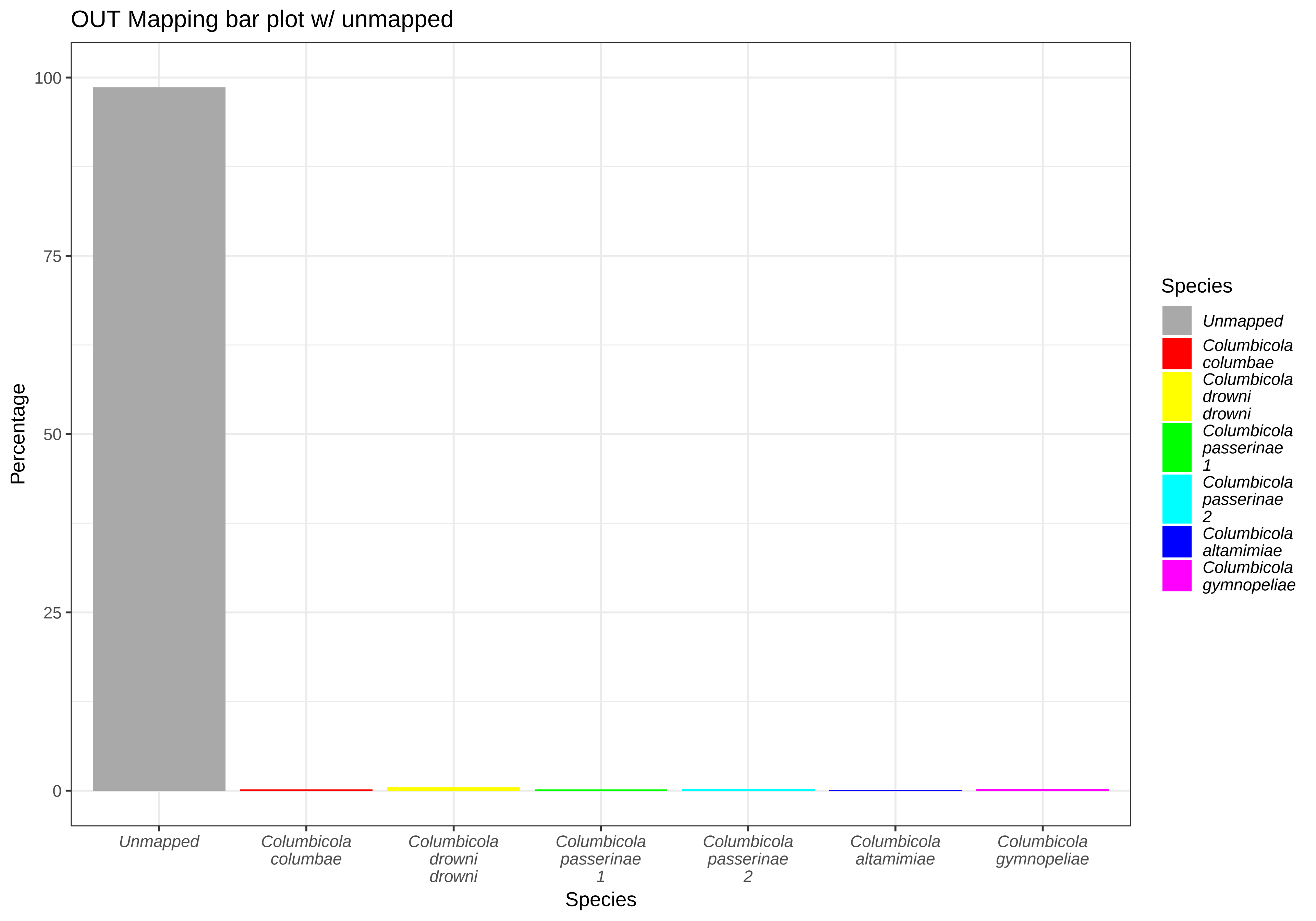

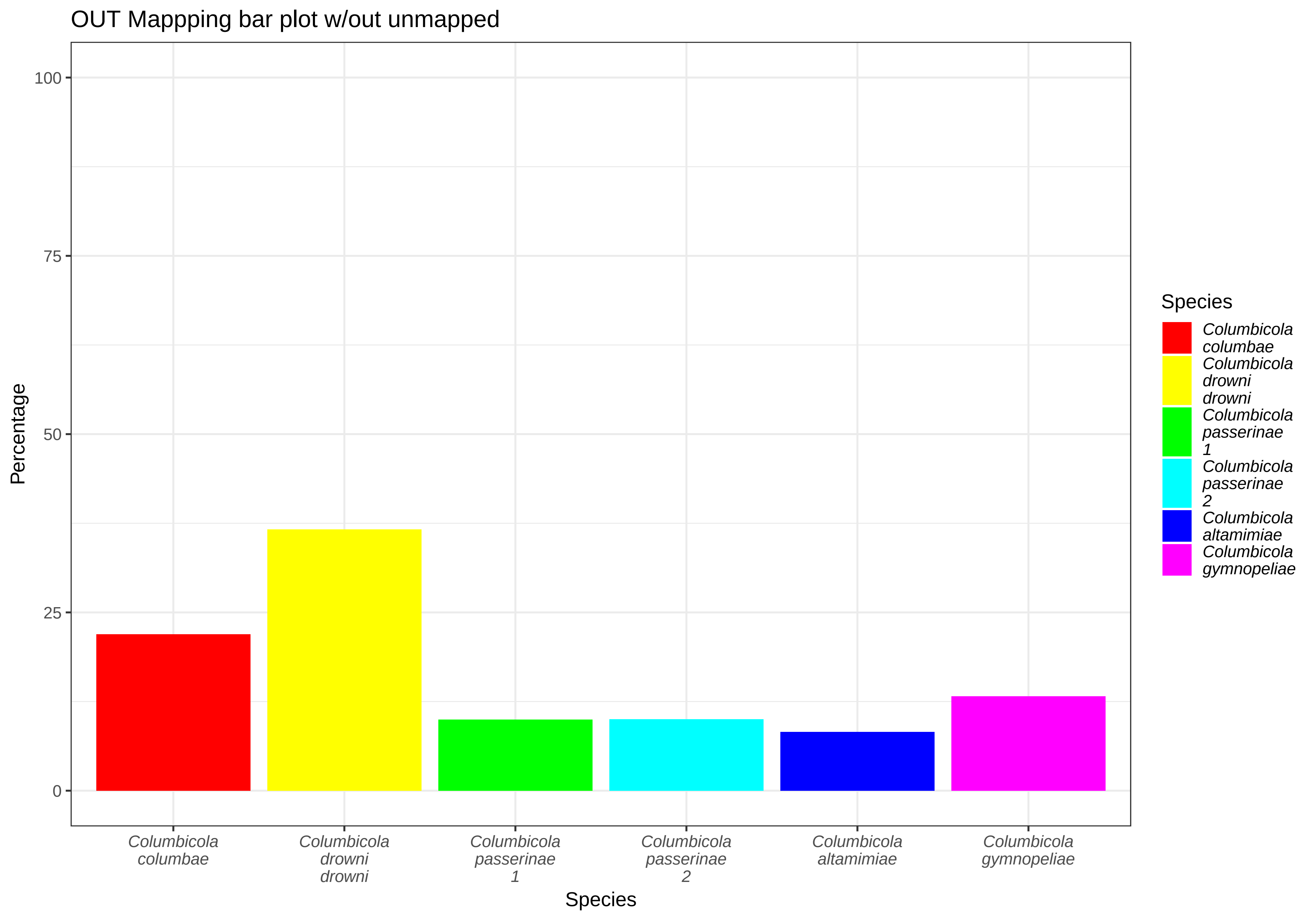

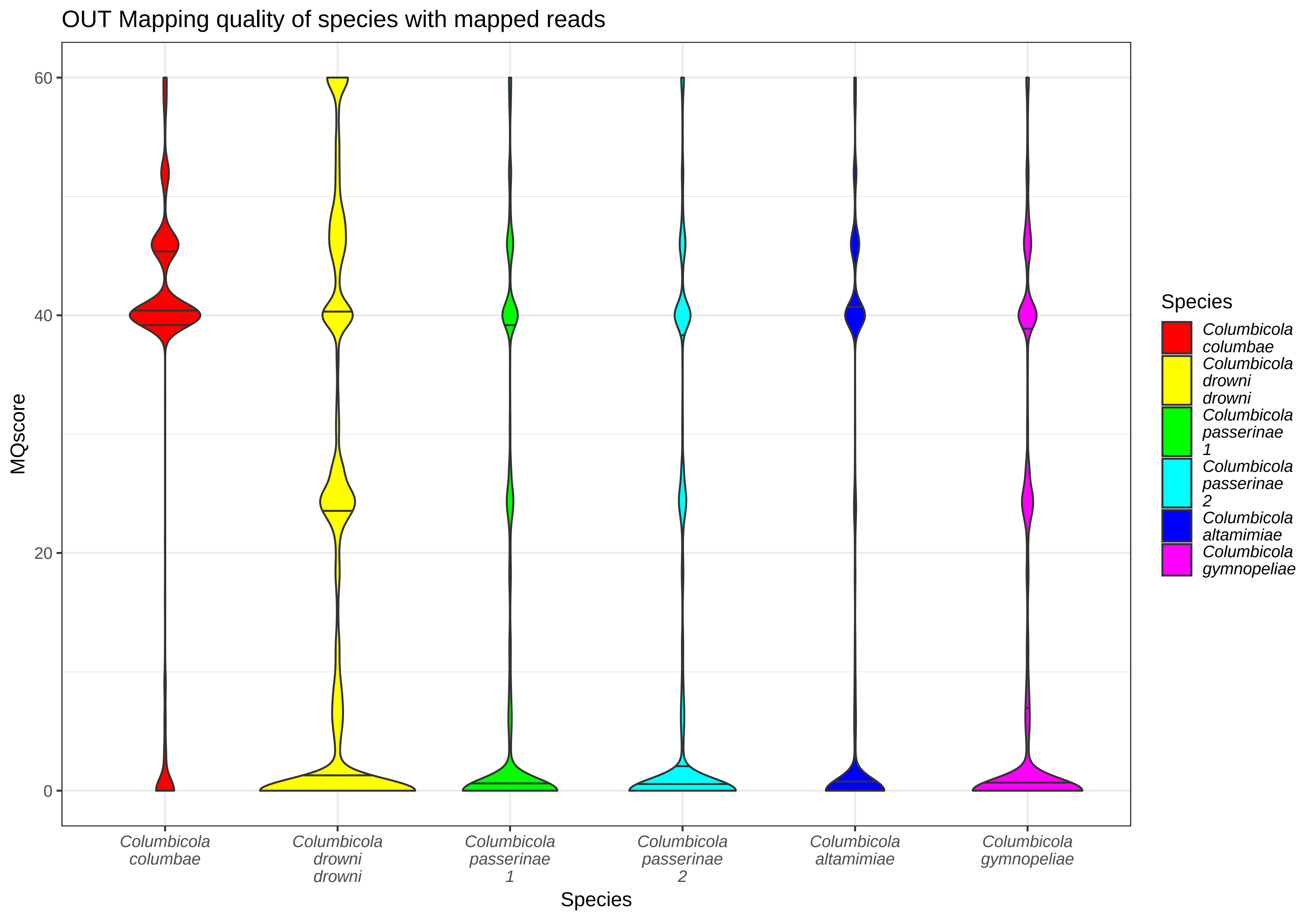

### OUT_plotMQ_Cosp_Memel_11_7_2016_13.pdf

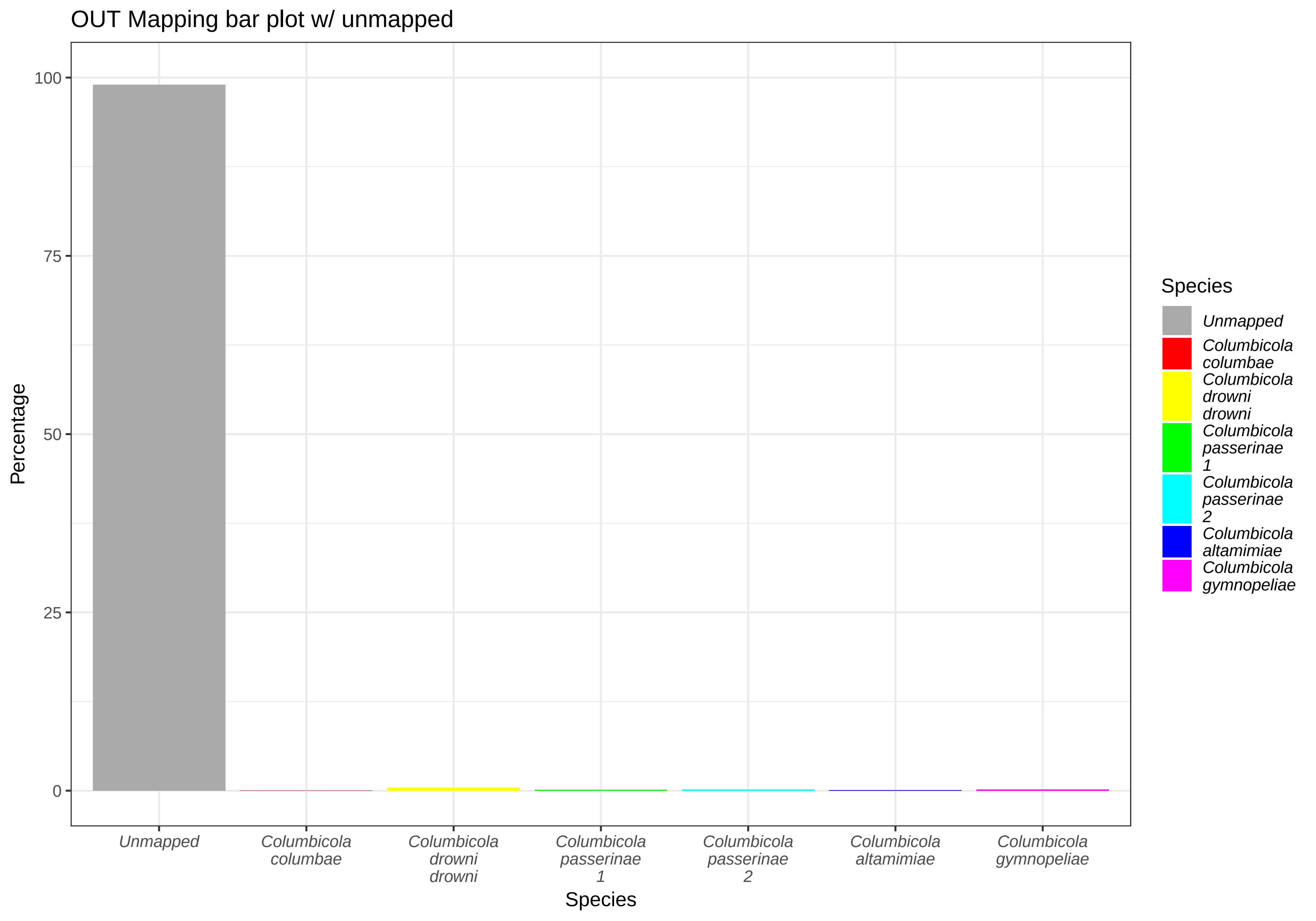

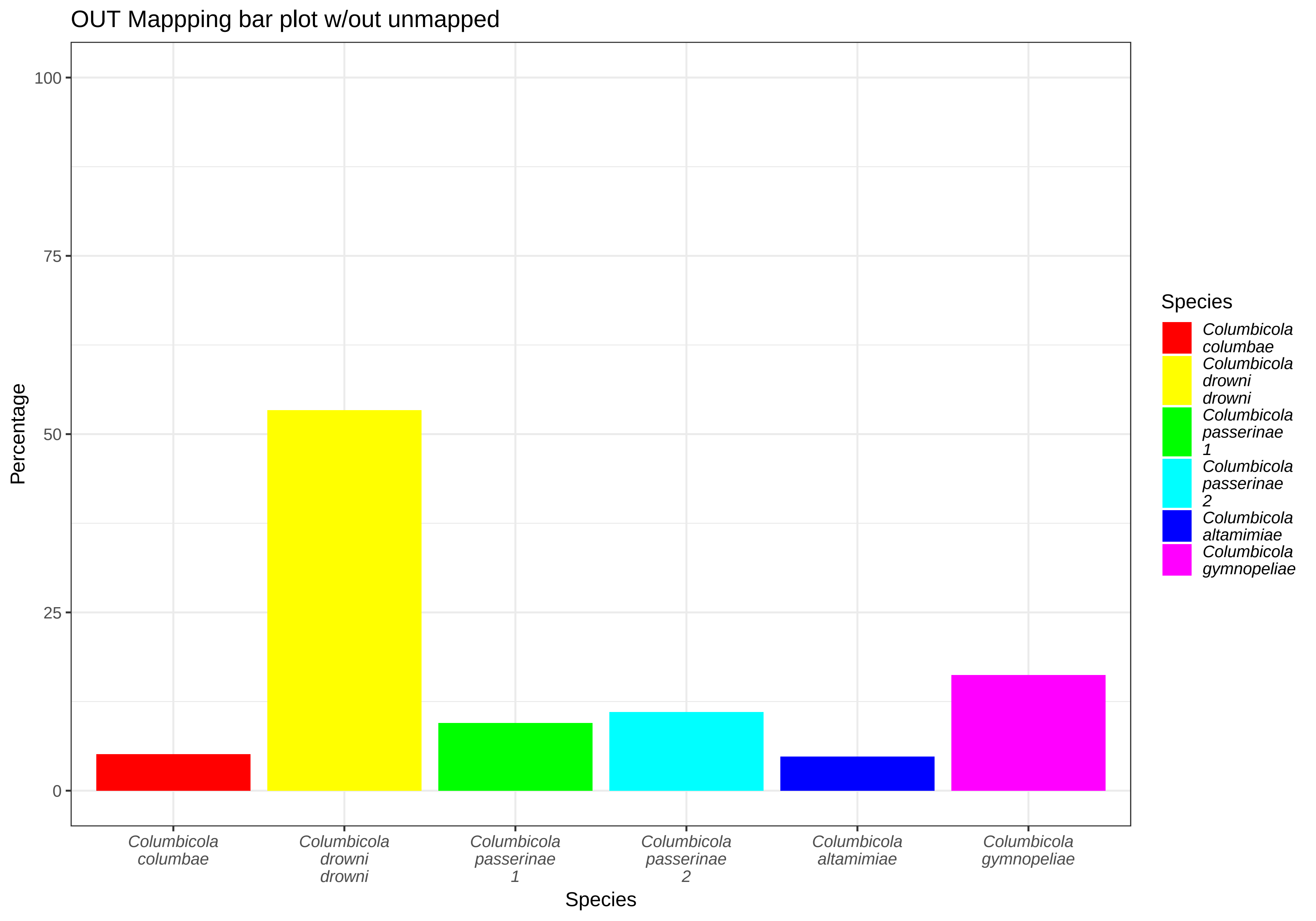

#### OUT Mapping quality of species with mapped reads

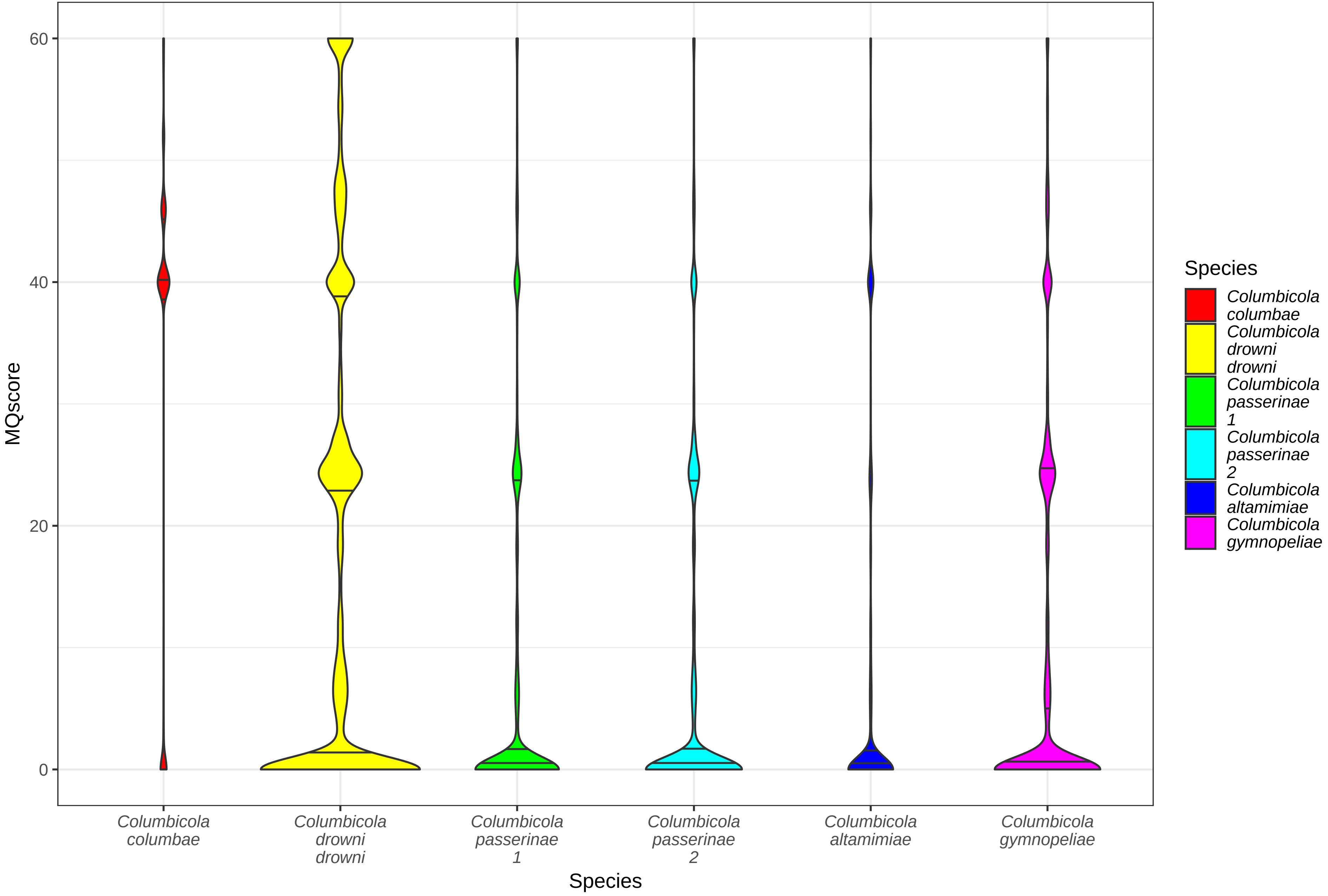

### OUT_plotMQ_Cosp_Urcam_11_7_2016_4.pdf

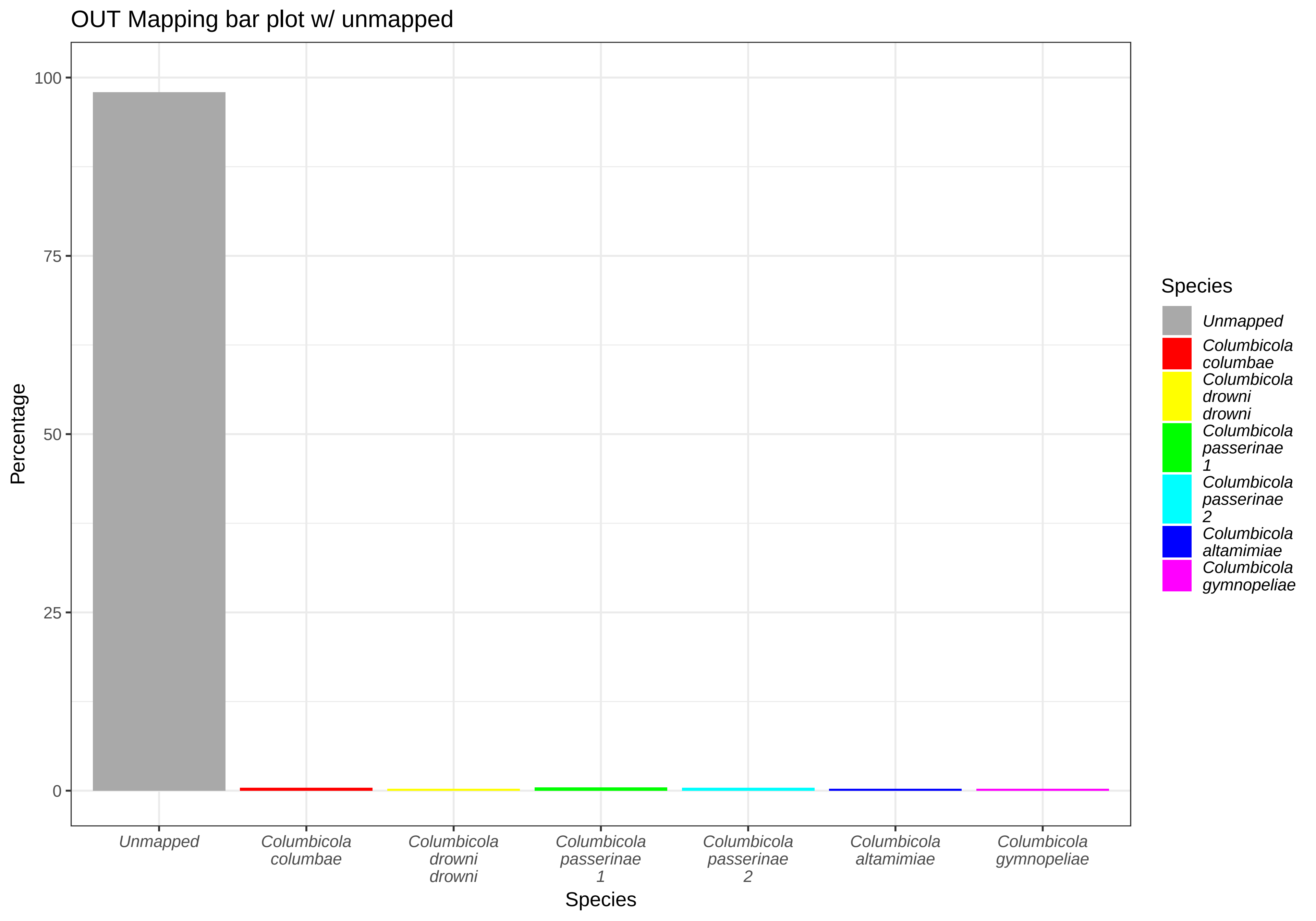

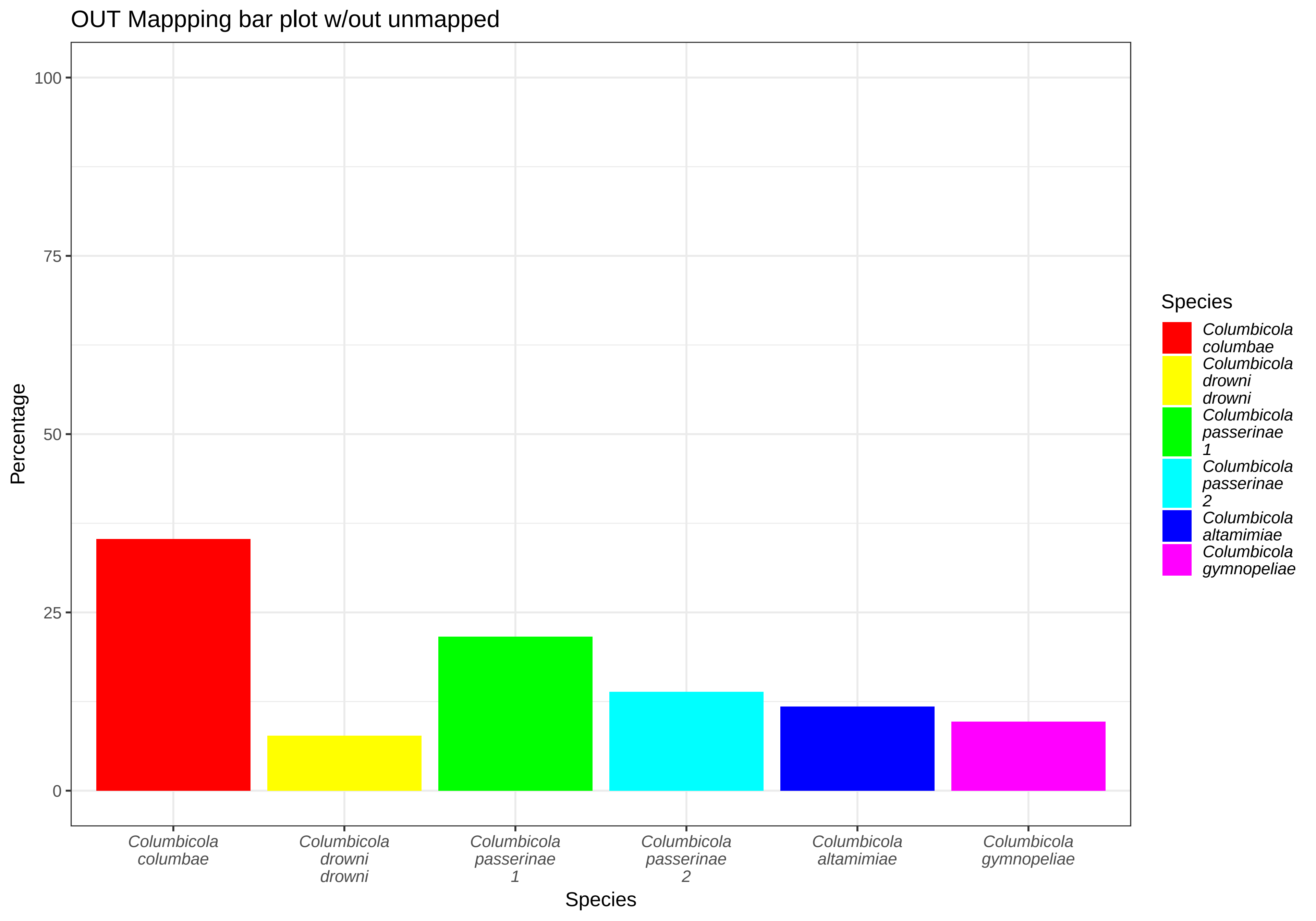

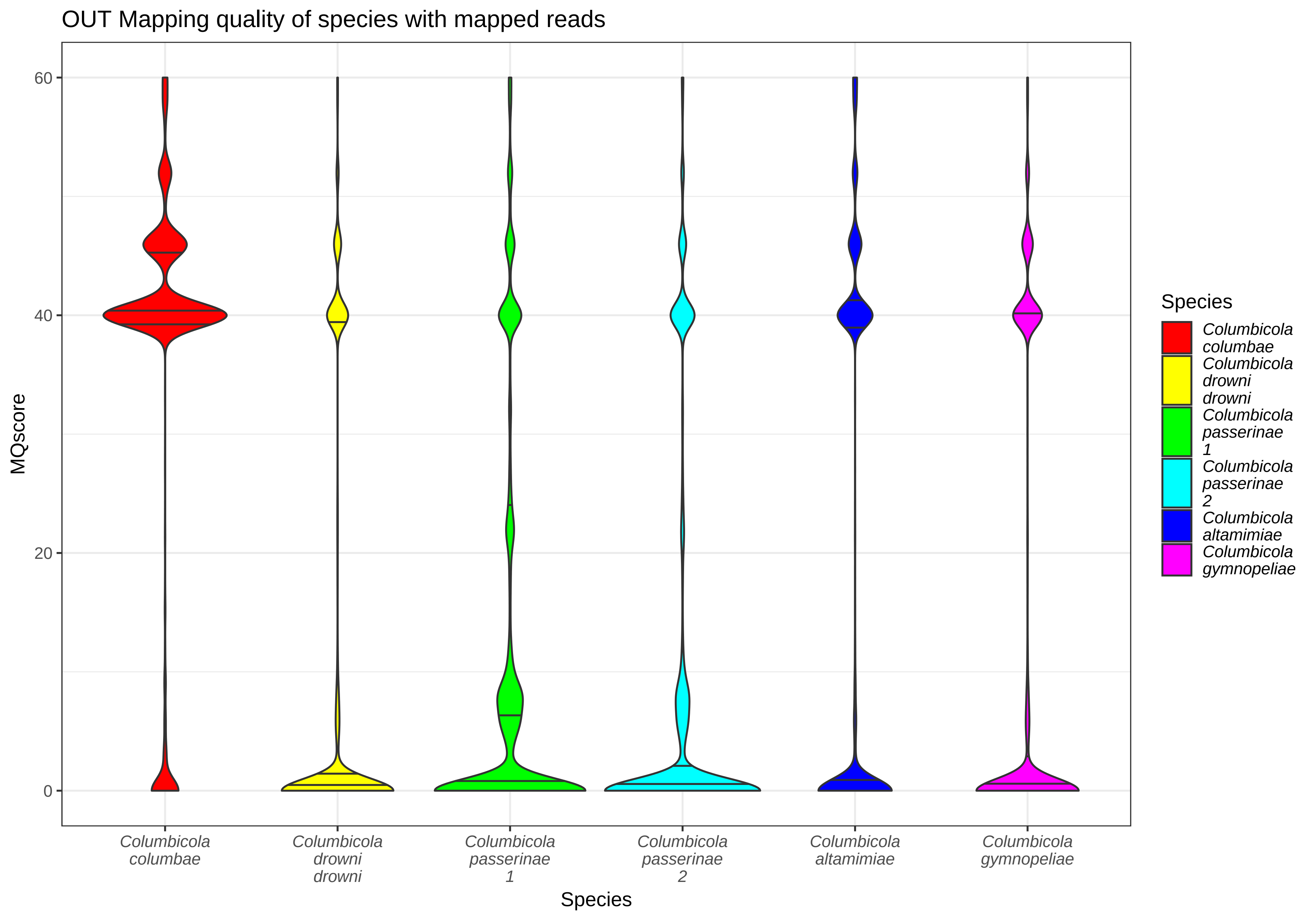

### OUT_plotMQ_Cosp_Urcam_11_7_2016_6.pdf

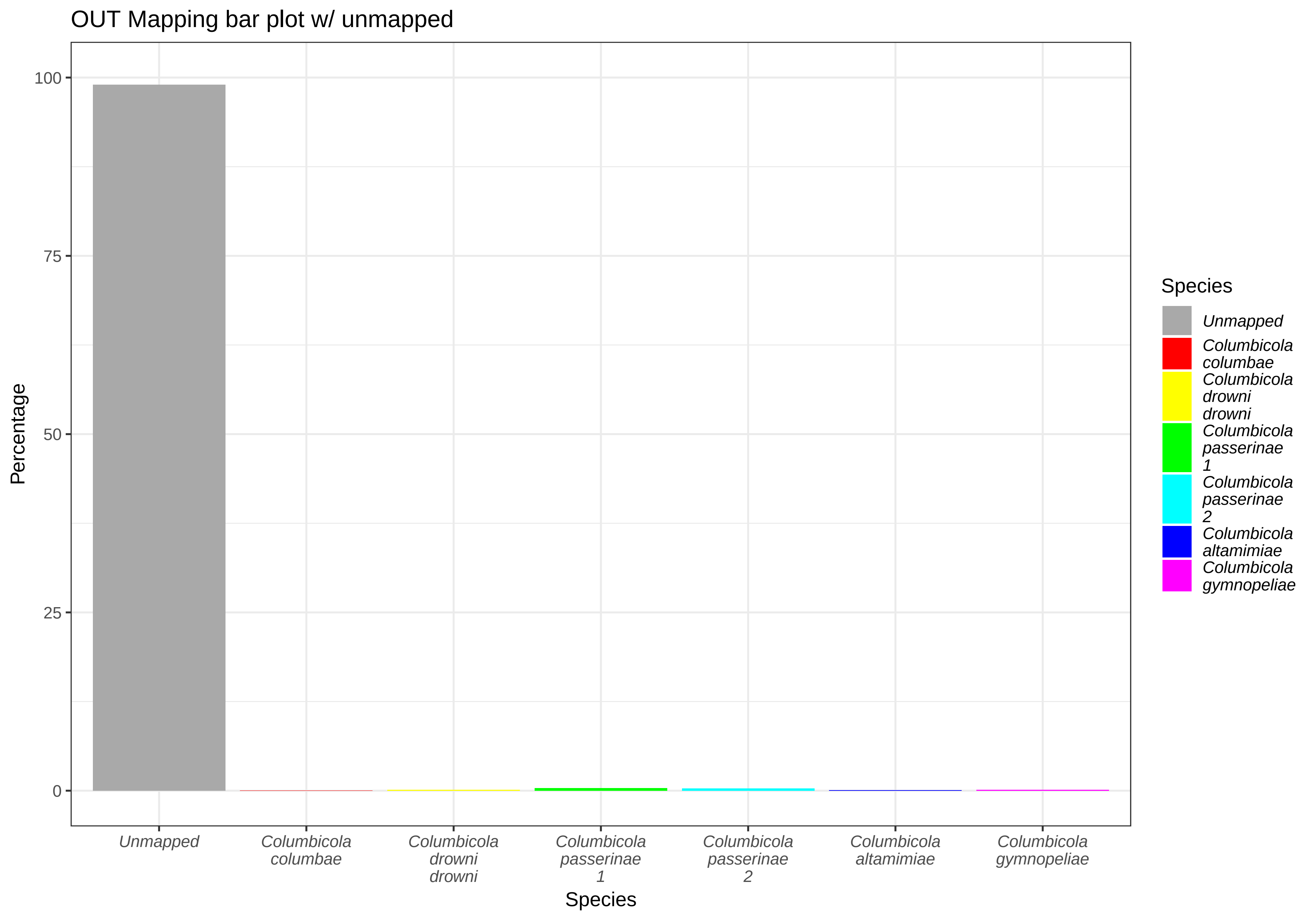

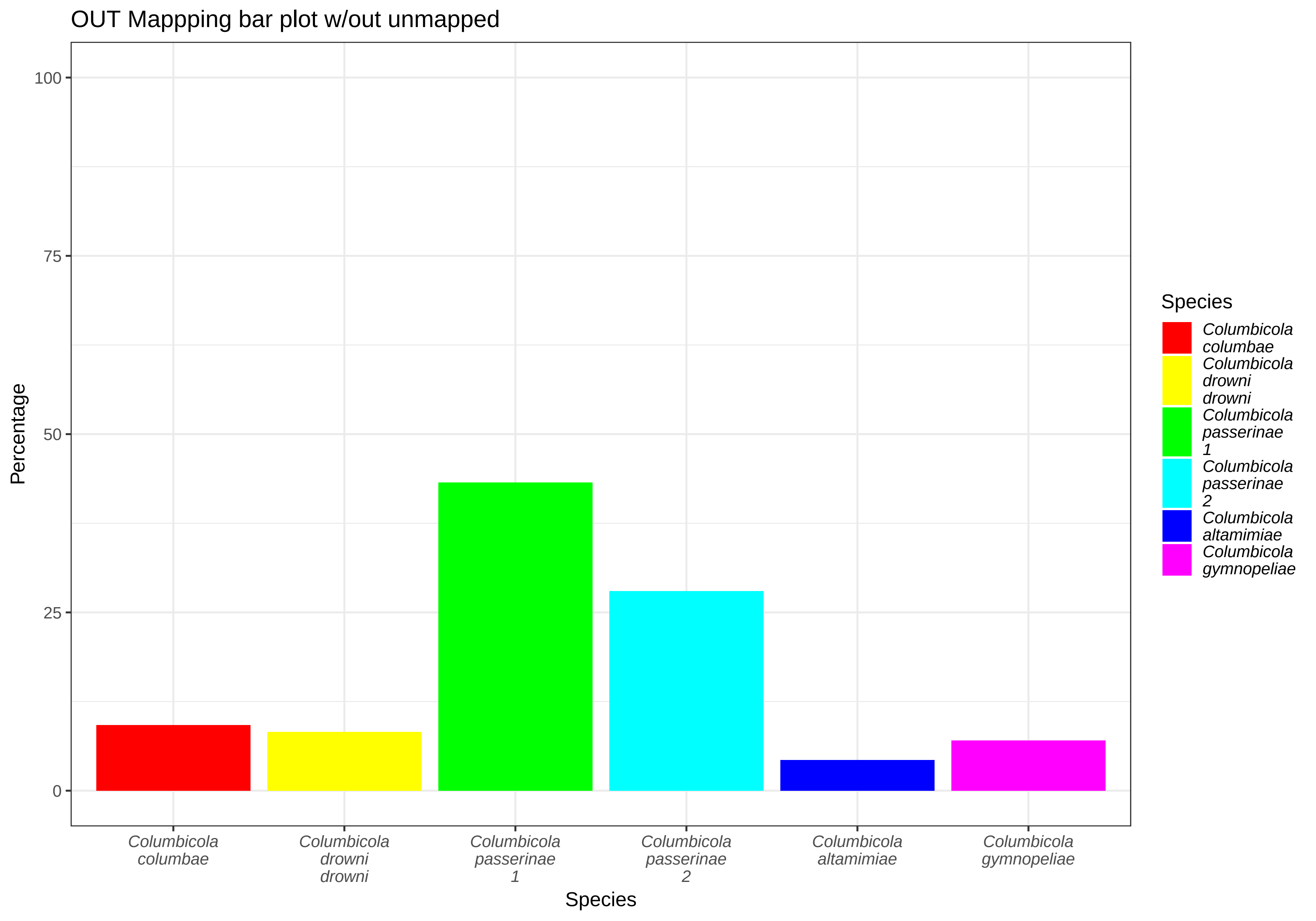

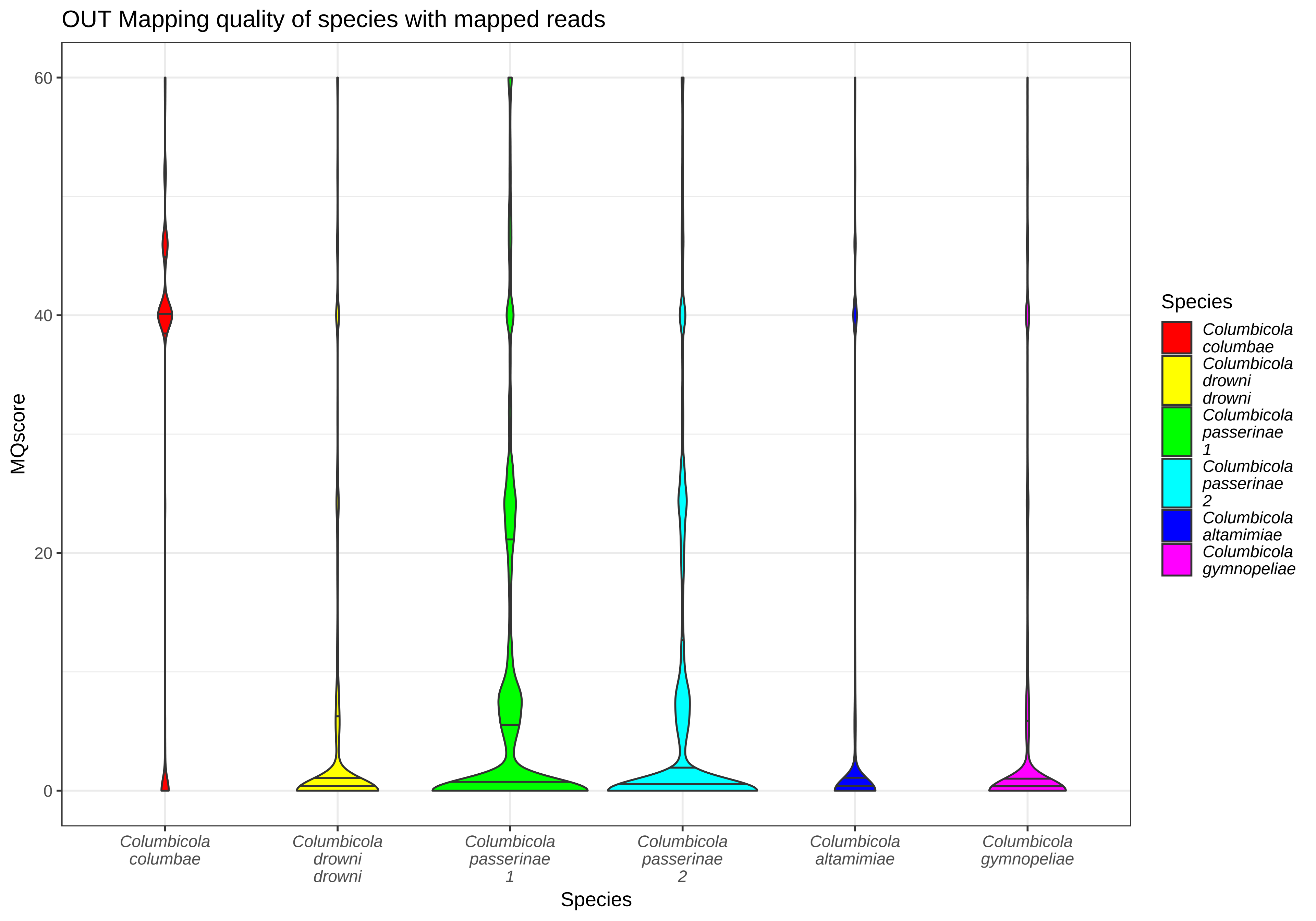

### OUT_plotMQ_Pheme_Memel_3_16_2016_2.pdf

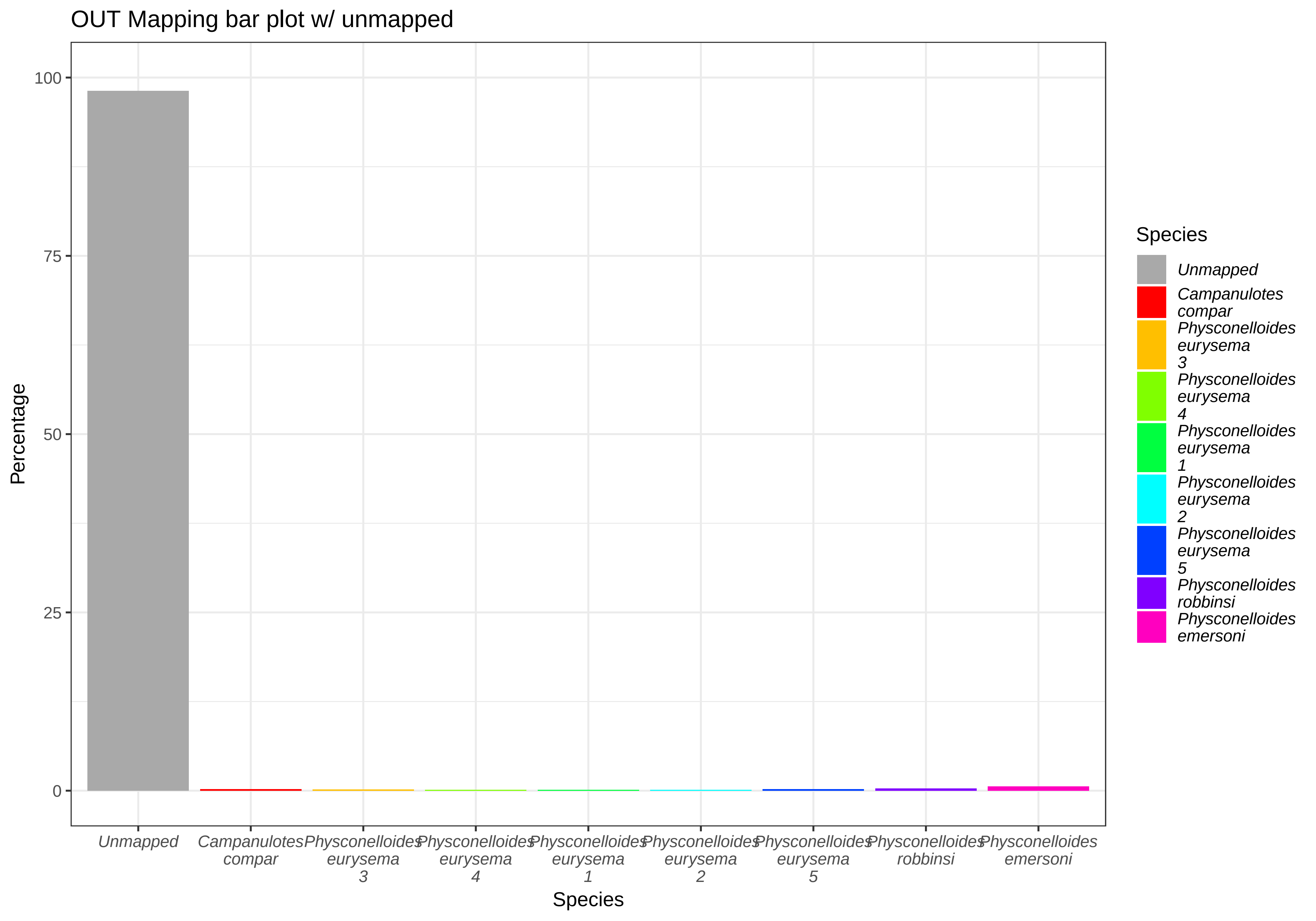

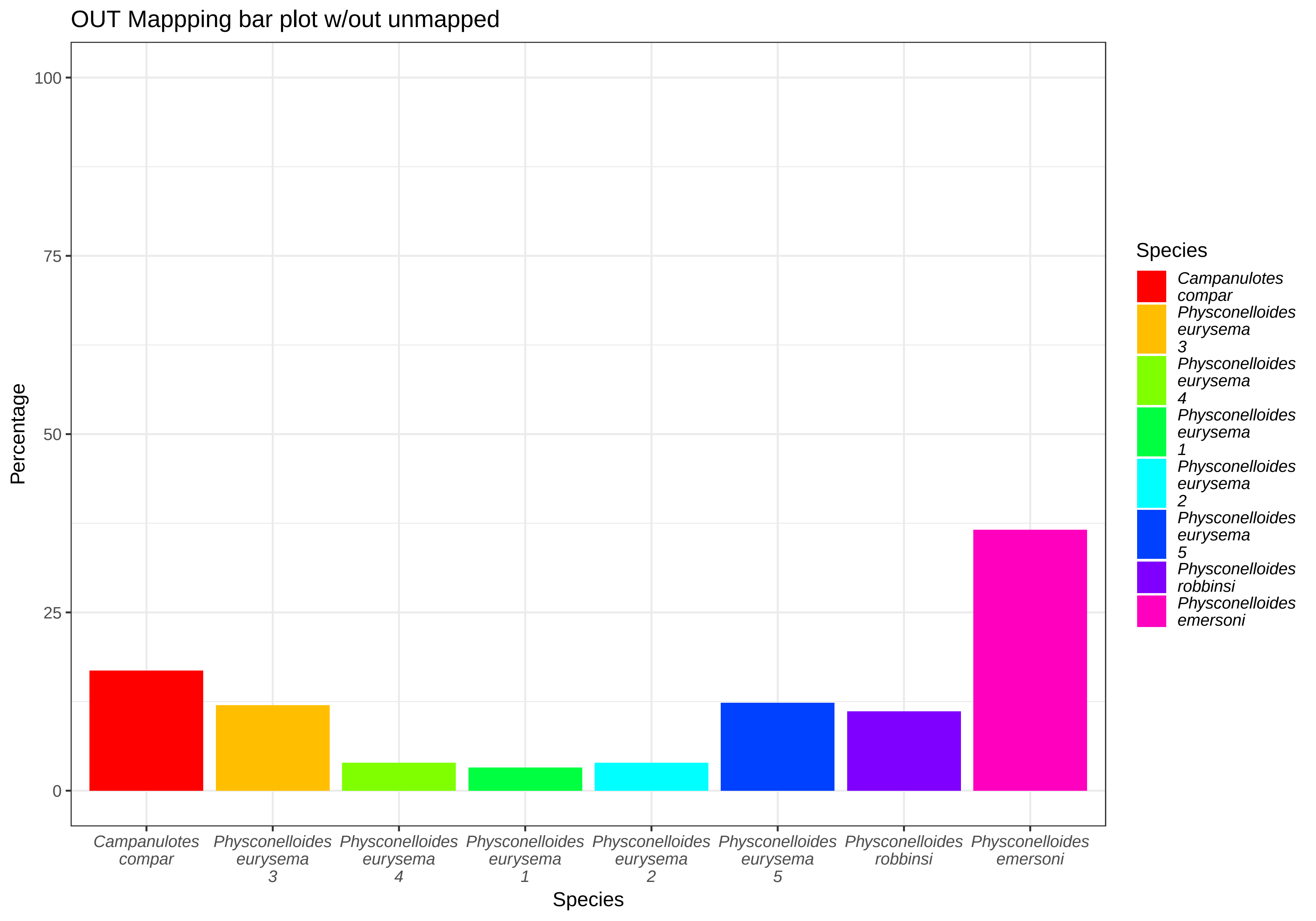

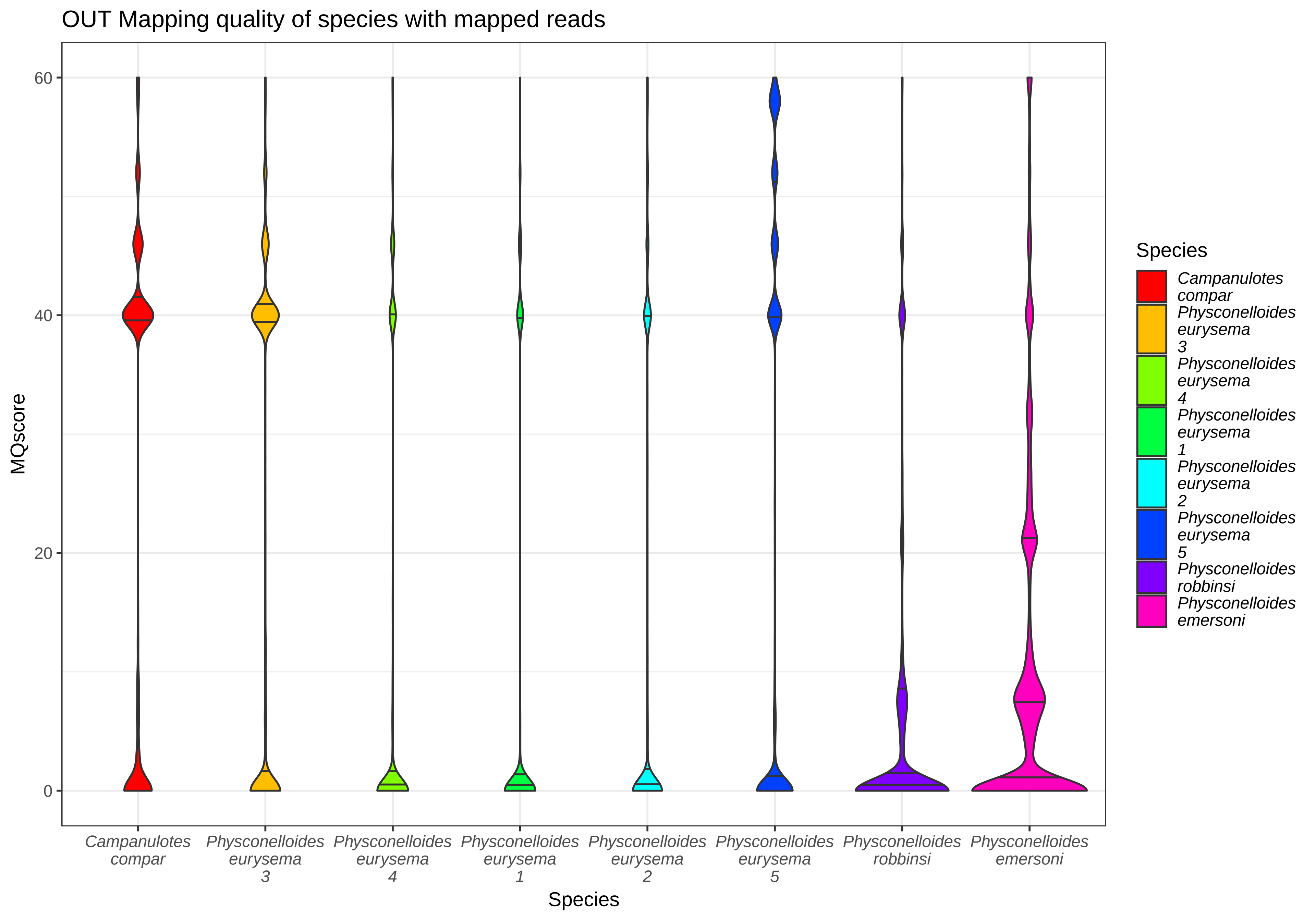

### OUT_plotMQ_Pheme_Memel_3_16_2016_8.pdf

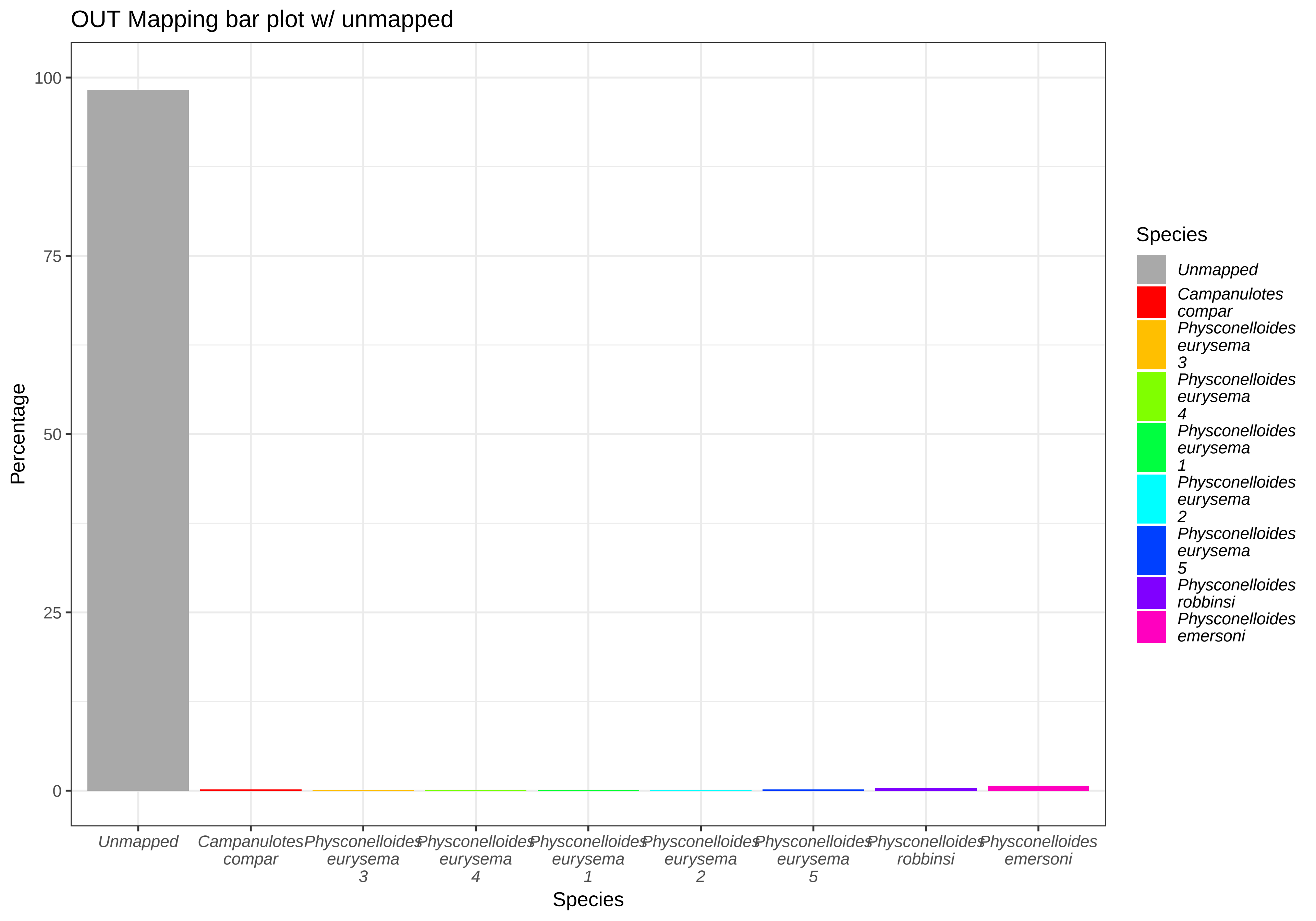

OUT Mappping bar plot w/out unmapped

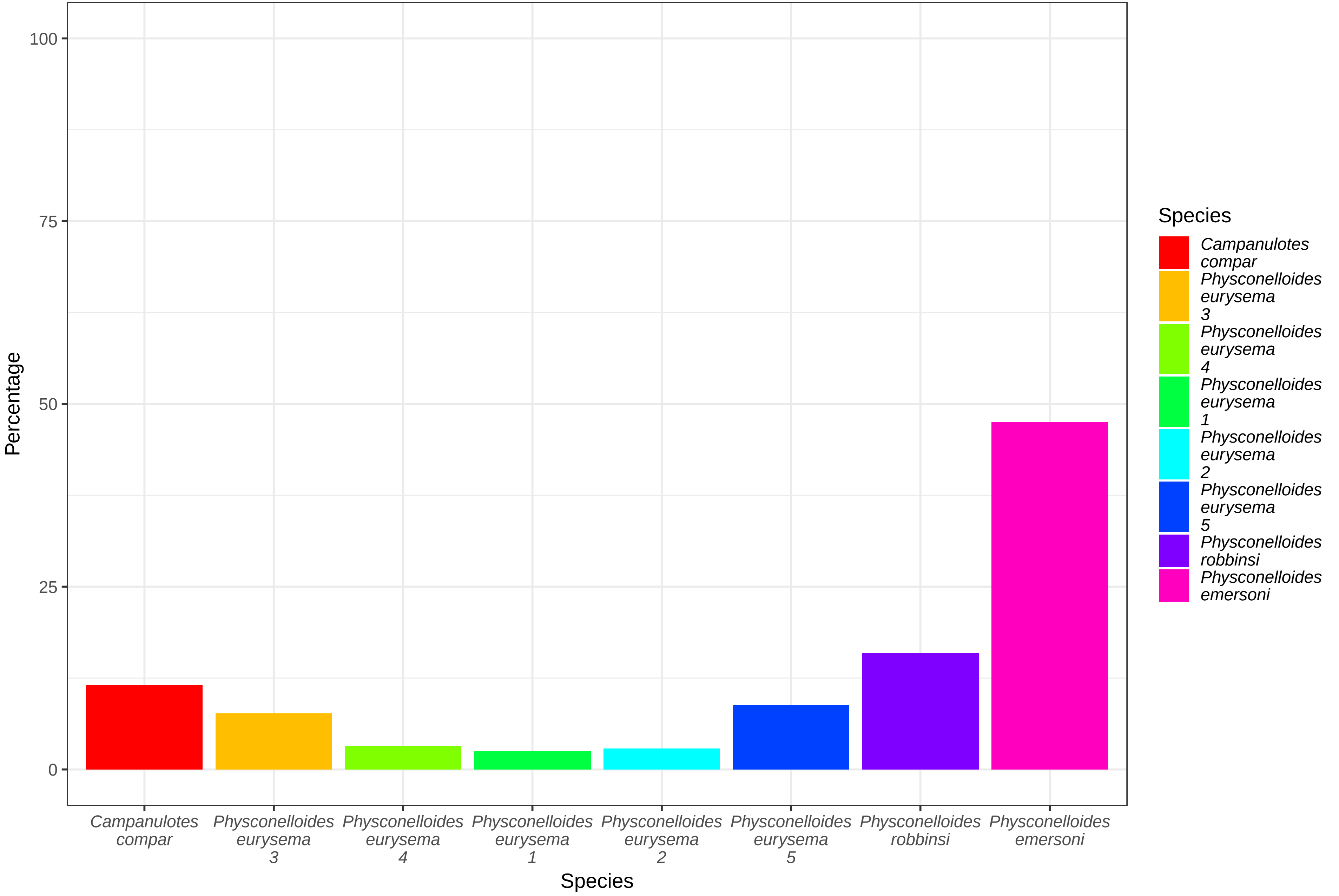

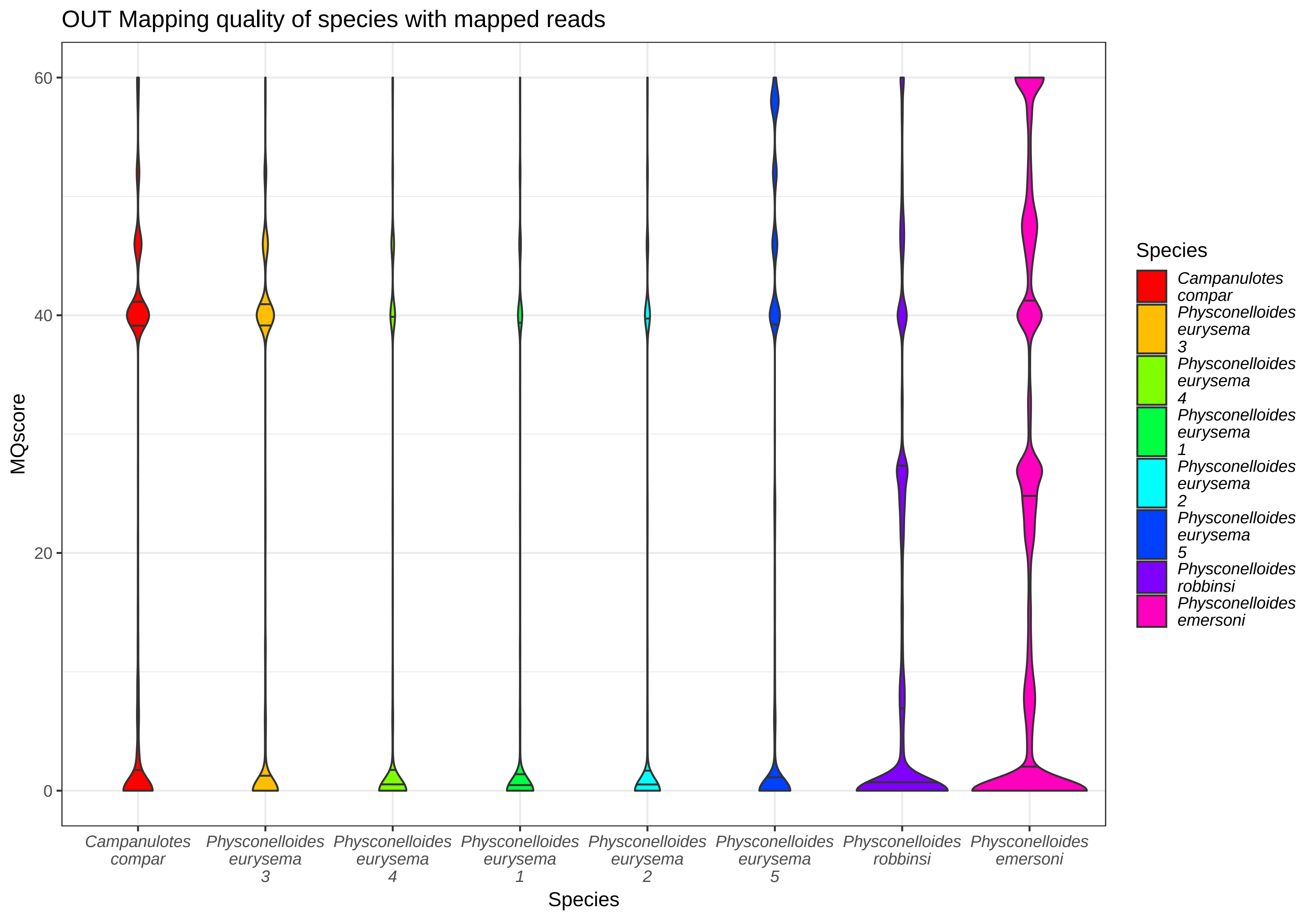

### OUT_plotMQ_Pheme_Memel_5_26_2015_8.pdf

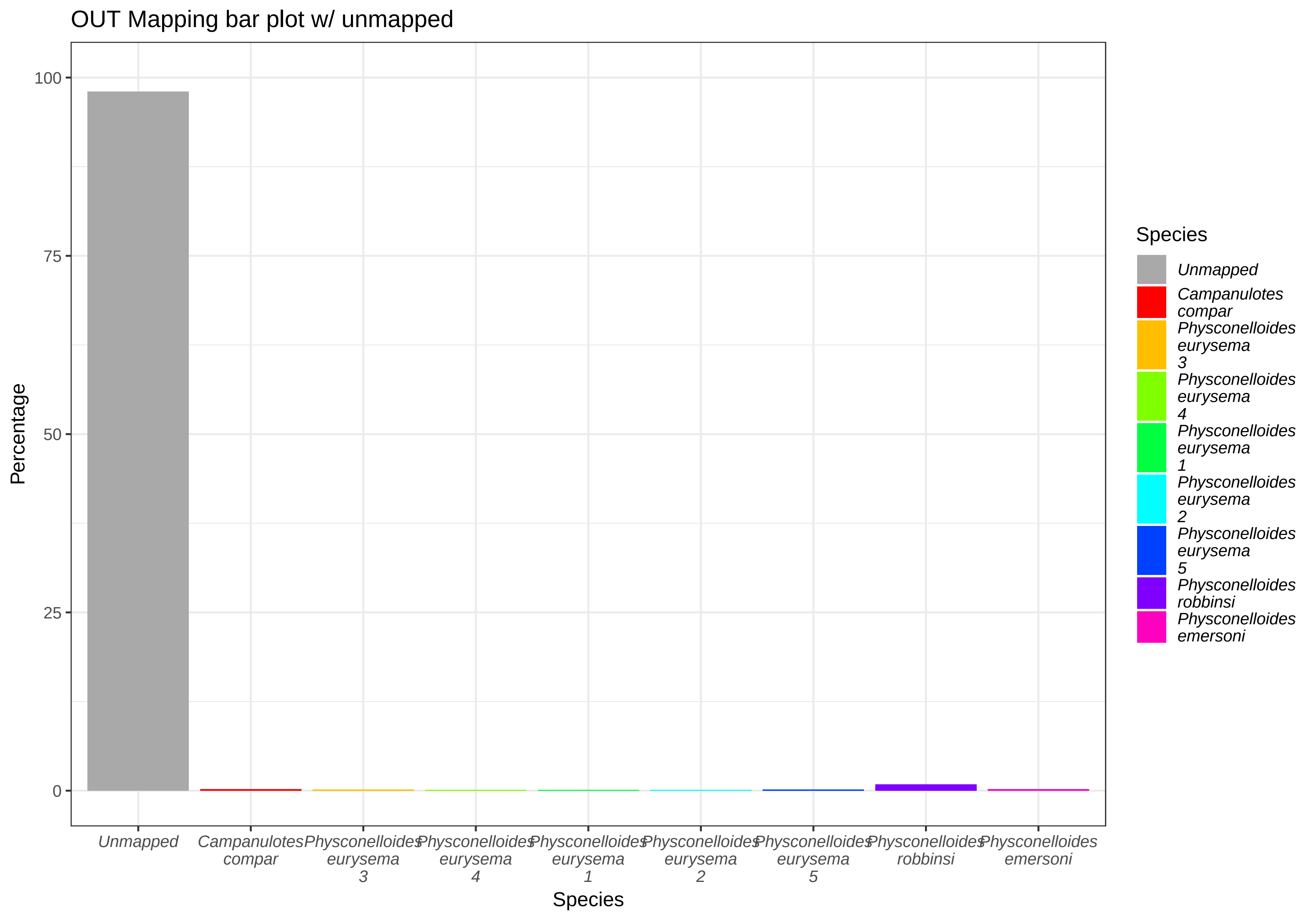

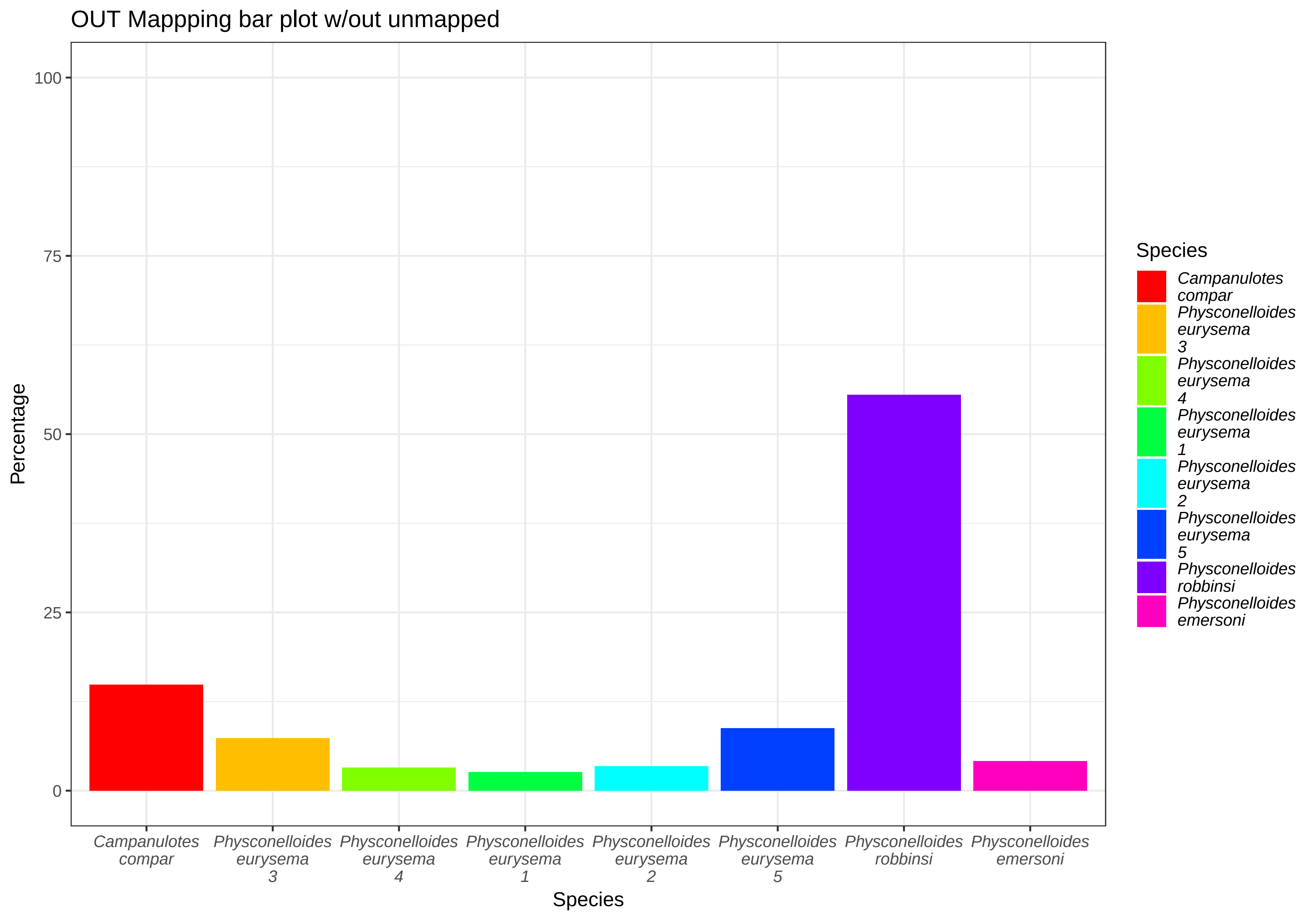

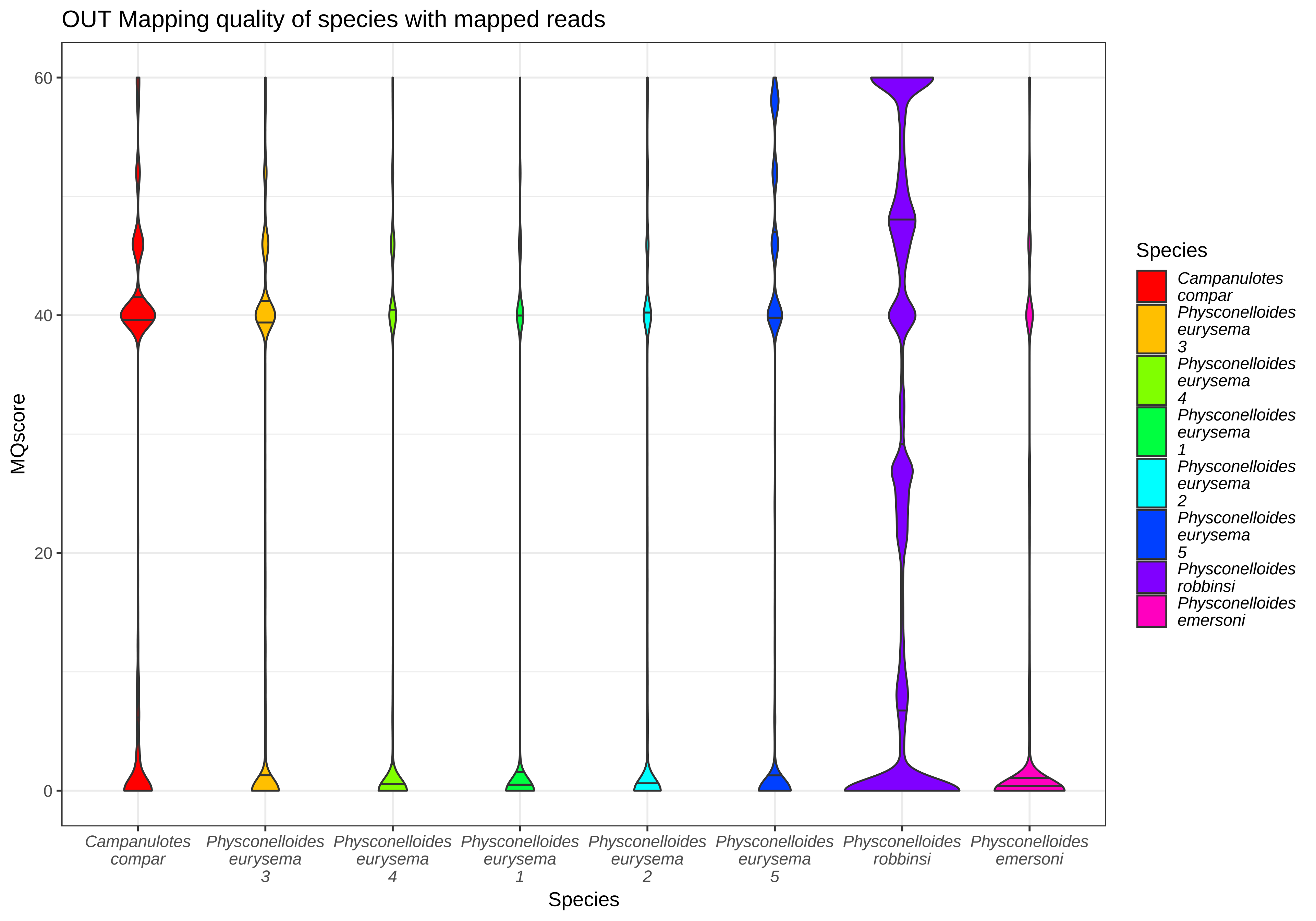

### OUT_plotMQ_Pheme_Memel_6_15_2015_5.pdf

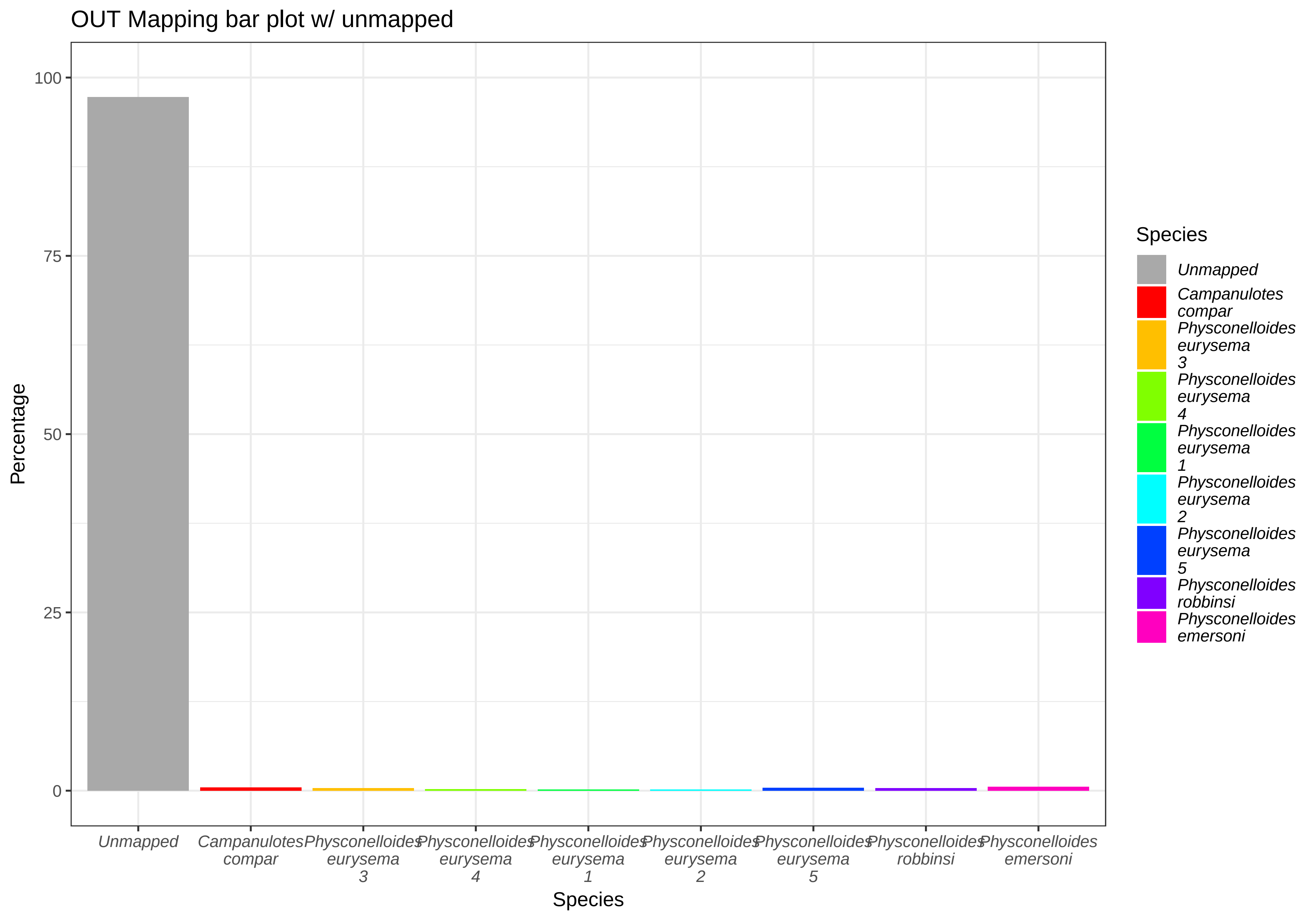

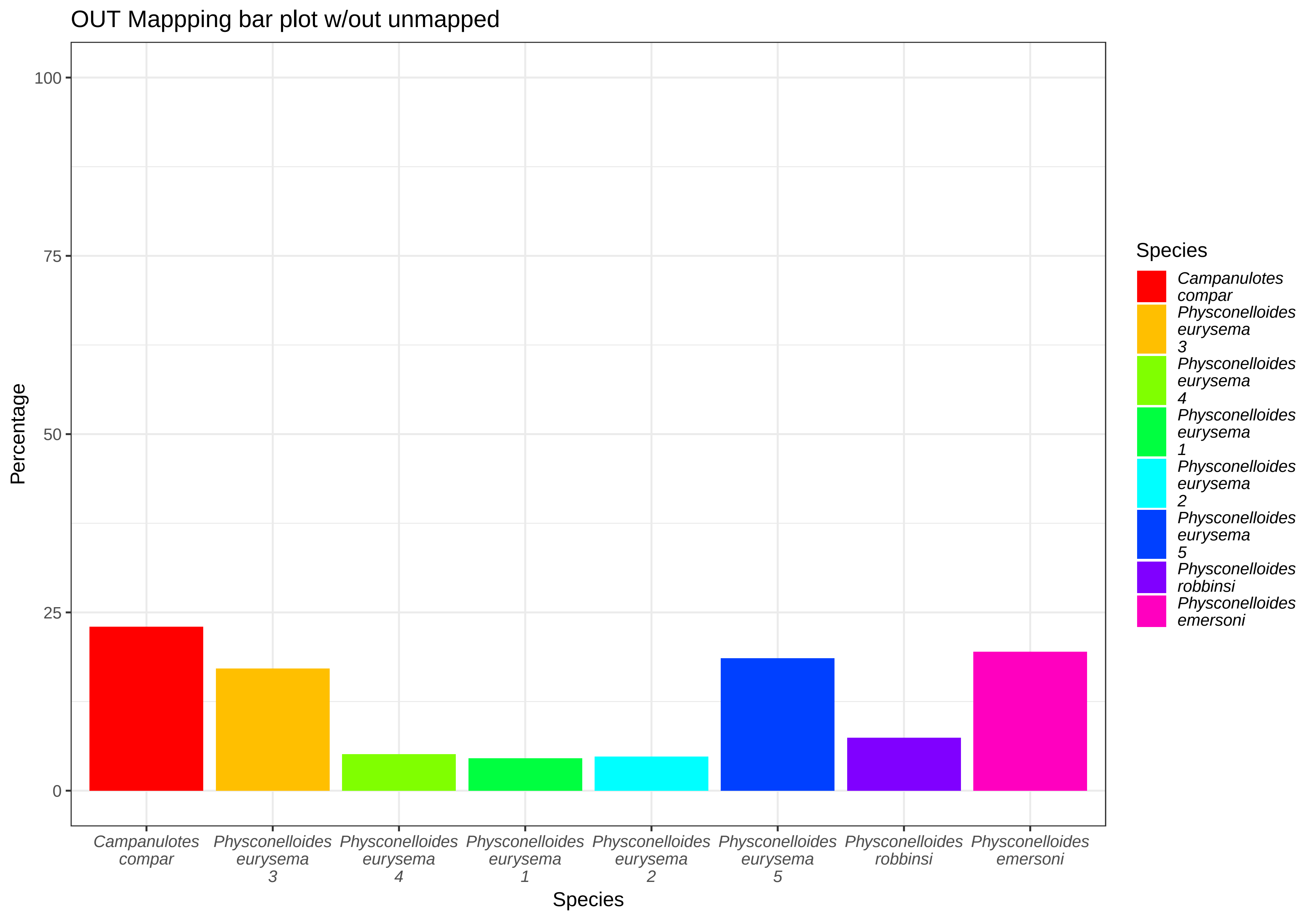

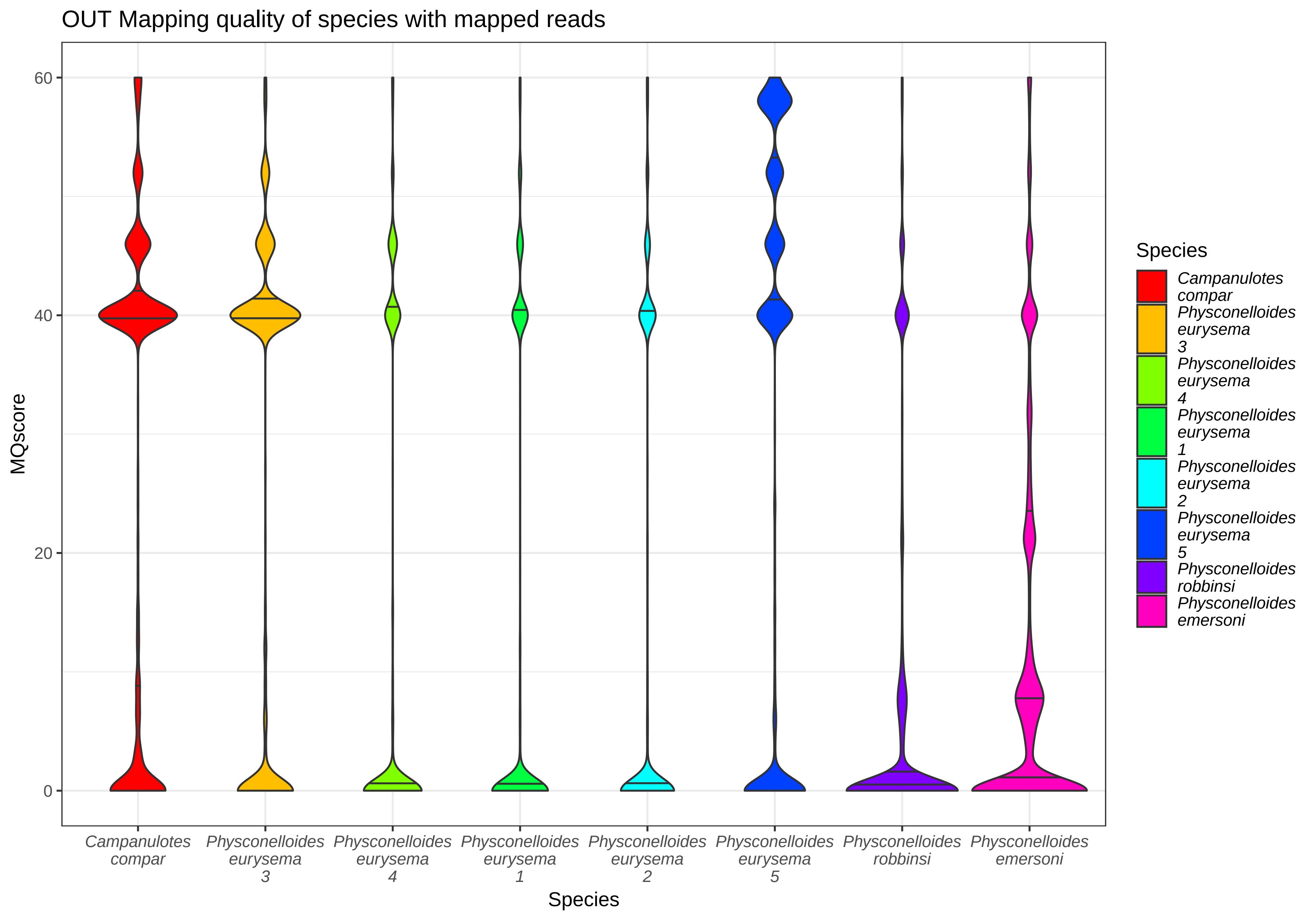

### OUT_plotMQ_Pheur1_Comin_3_14_2016_9.pdf

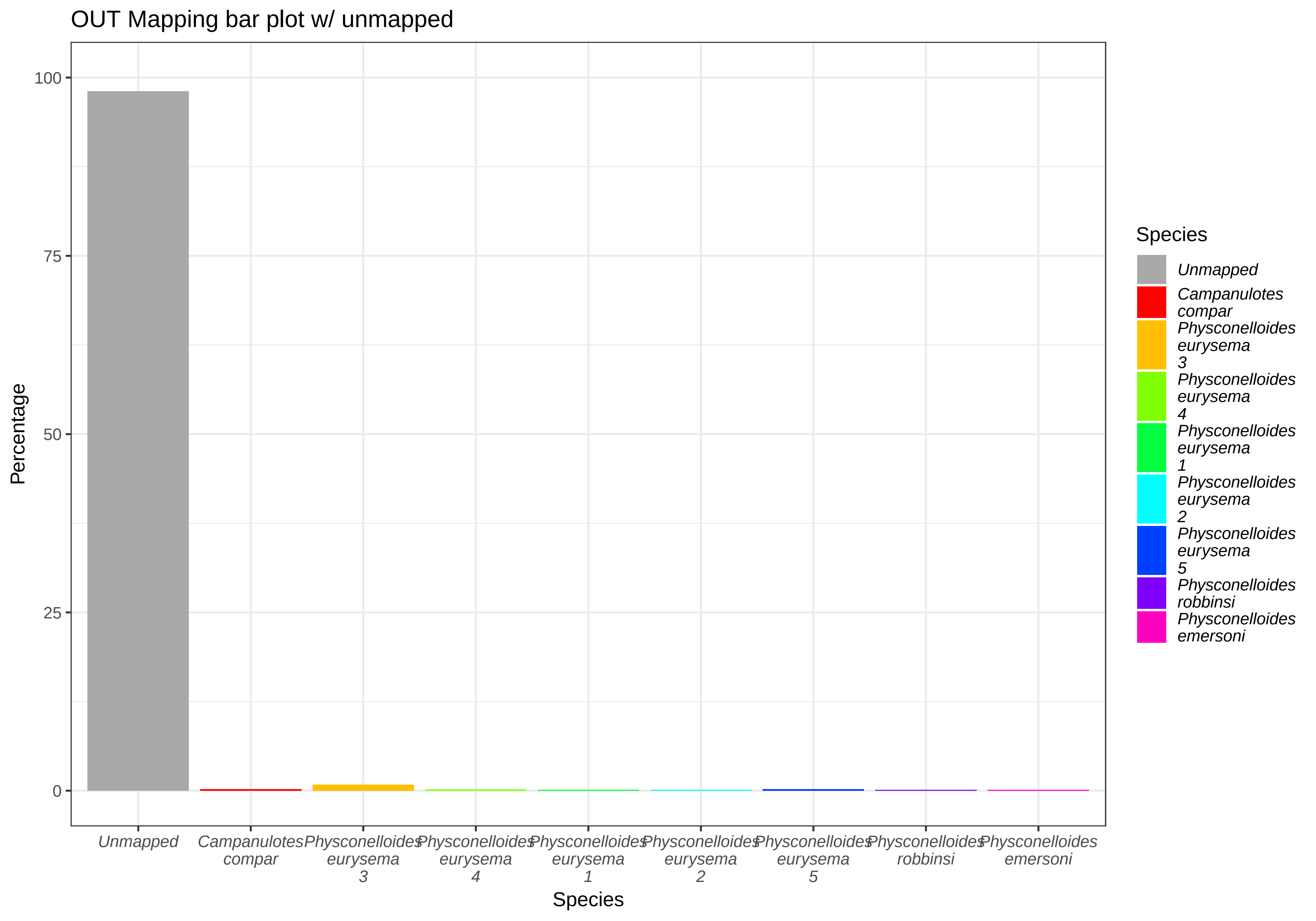

#### OUT Mappping bar plot w/out unmapped

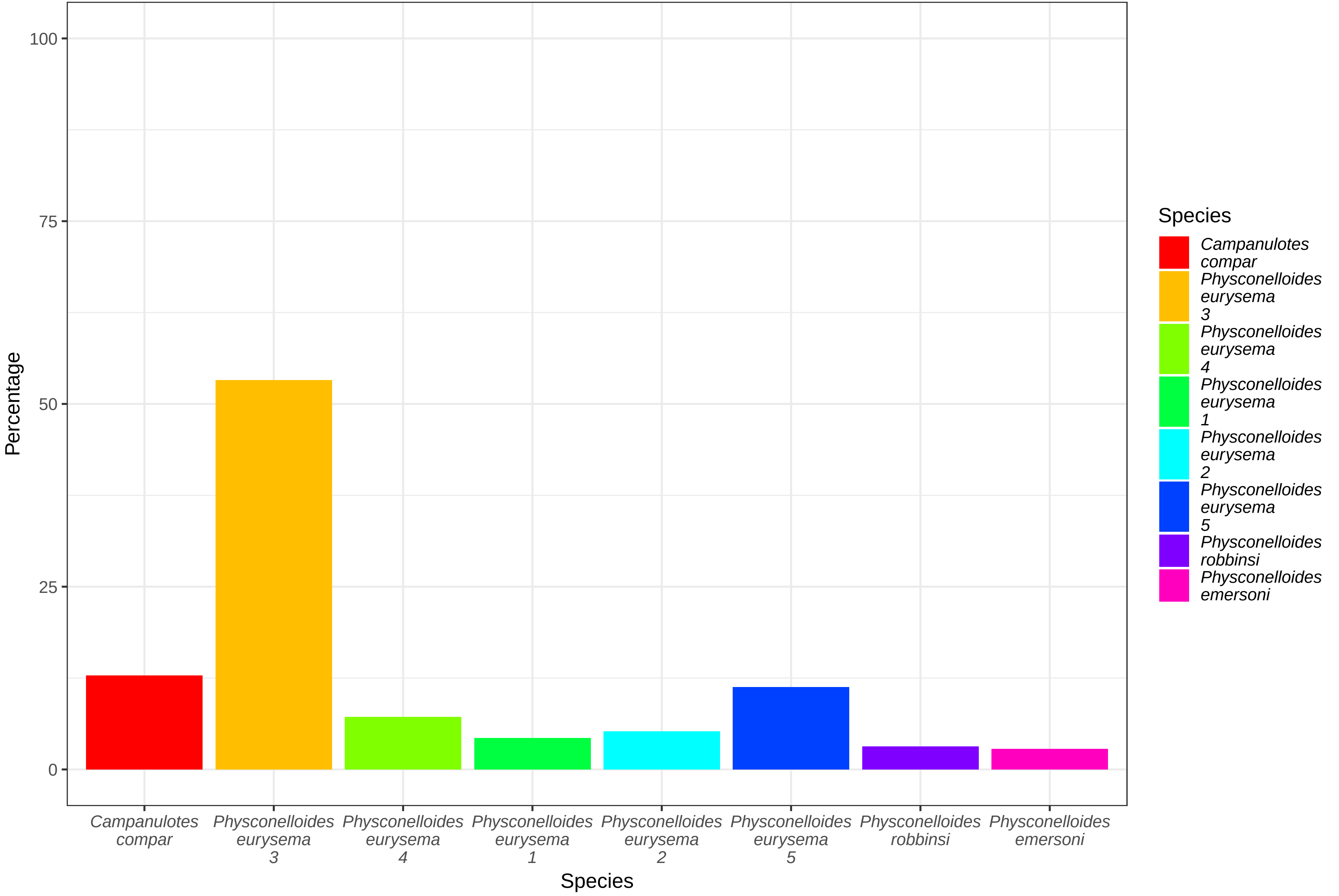

OUT Mapping quality of species with mapped reads

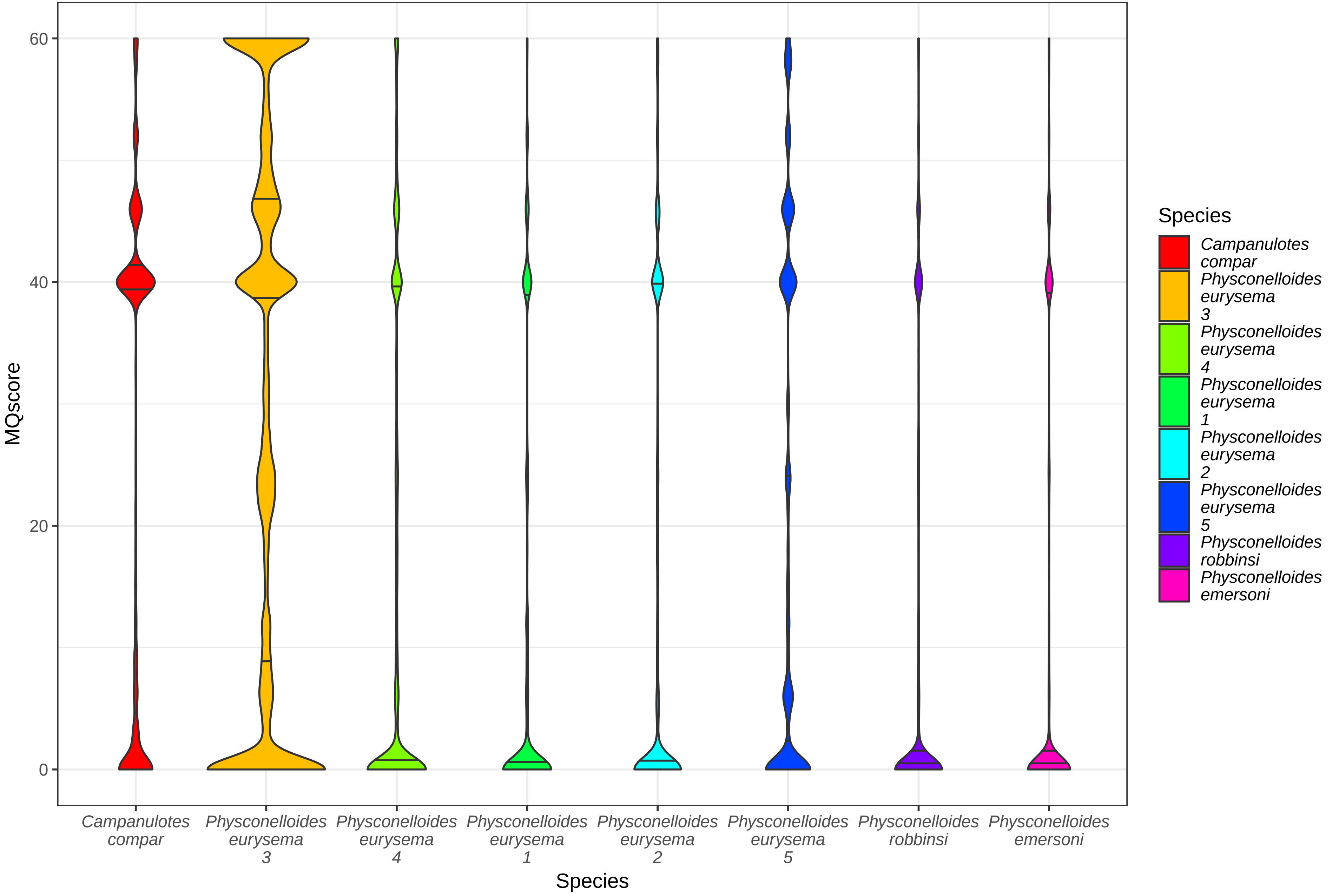

### OUT_plotMQ_Pheur2_Clpre_3_14_2016_5.pdf

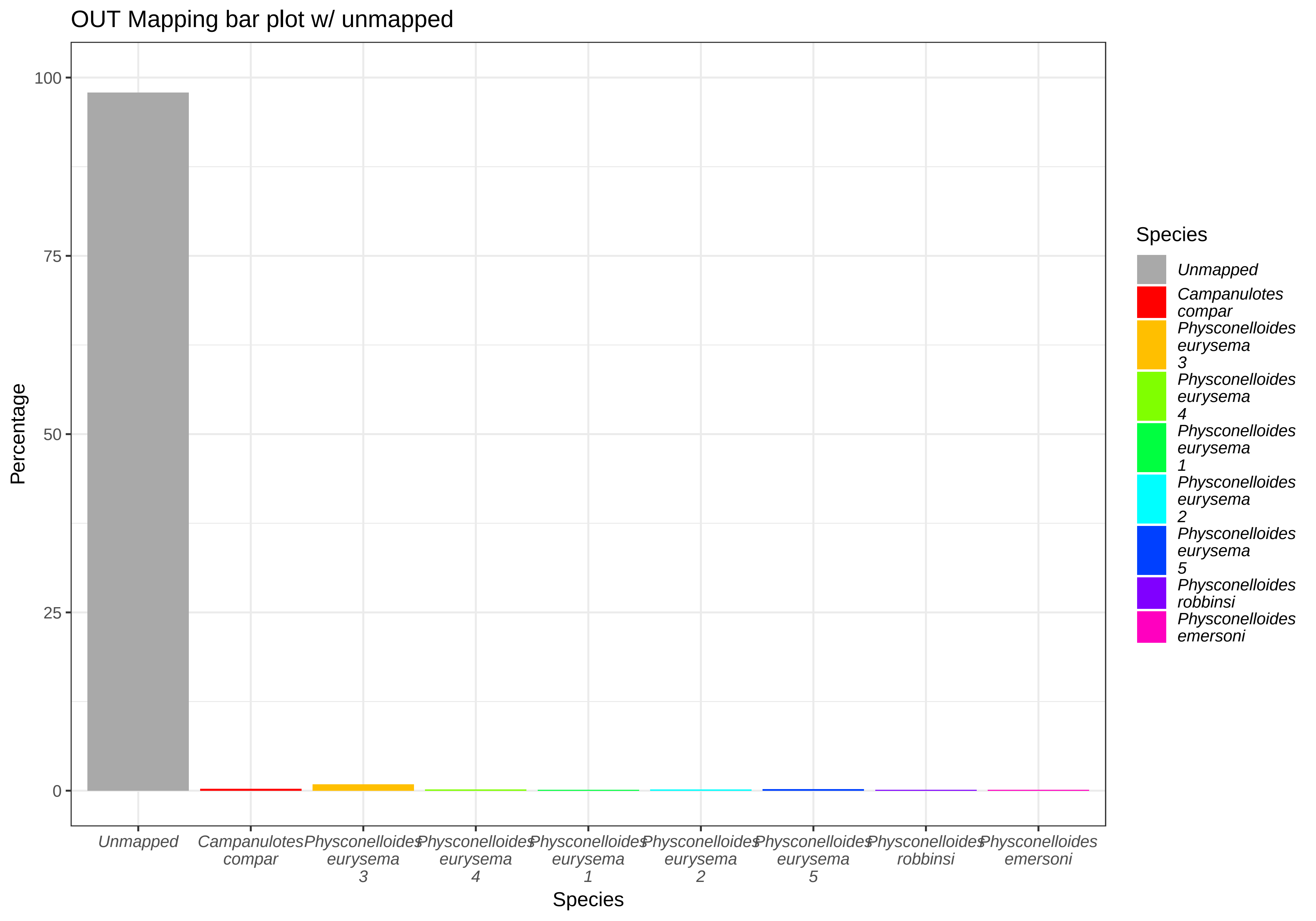

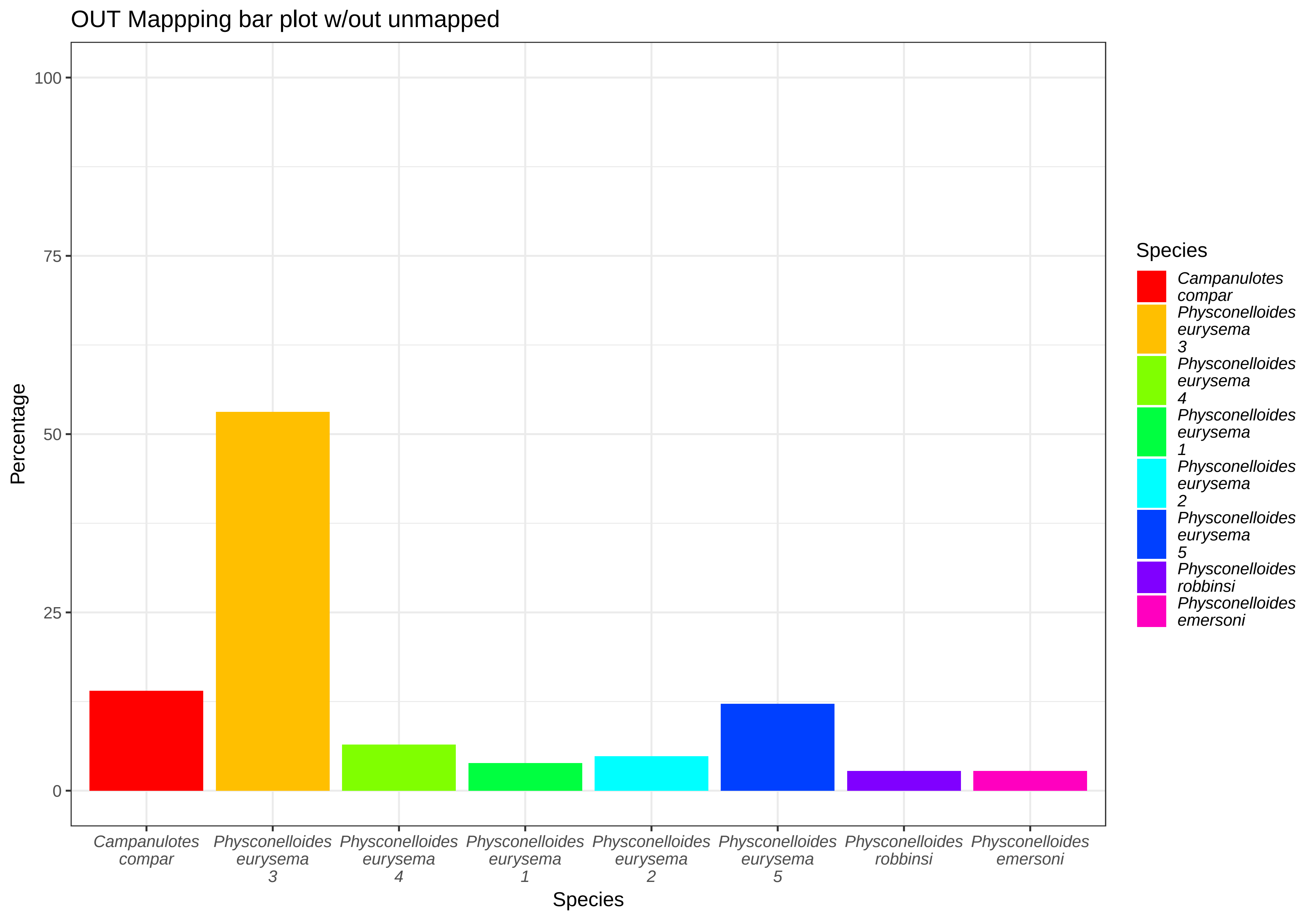

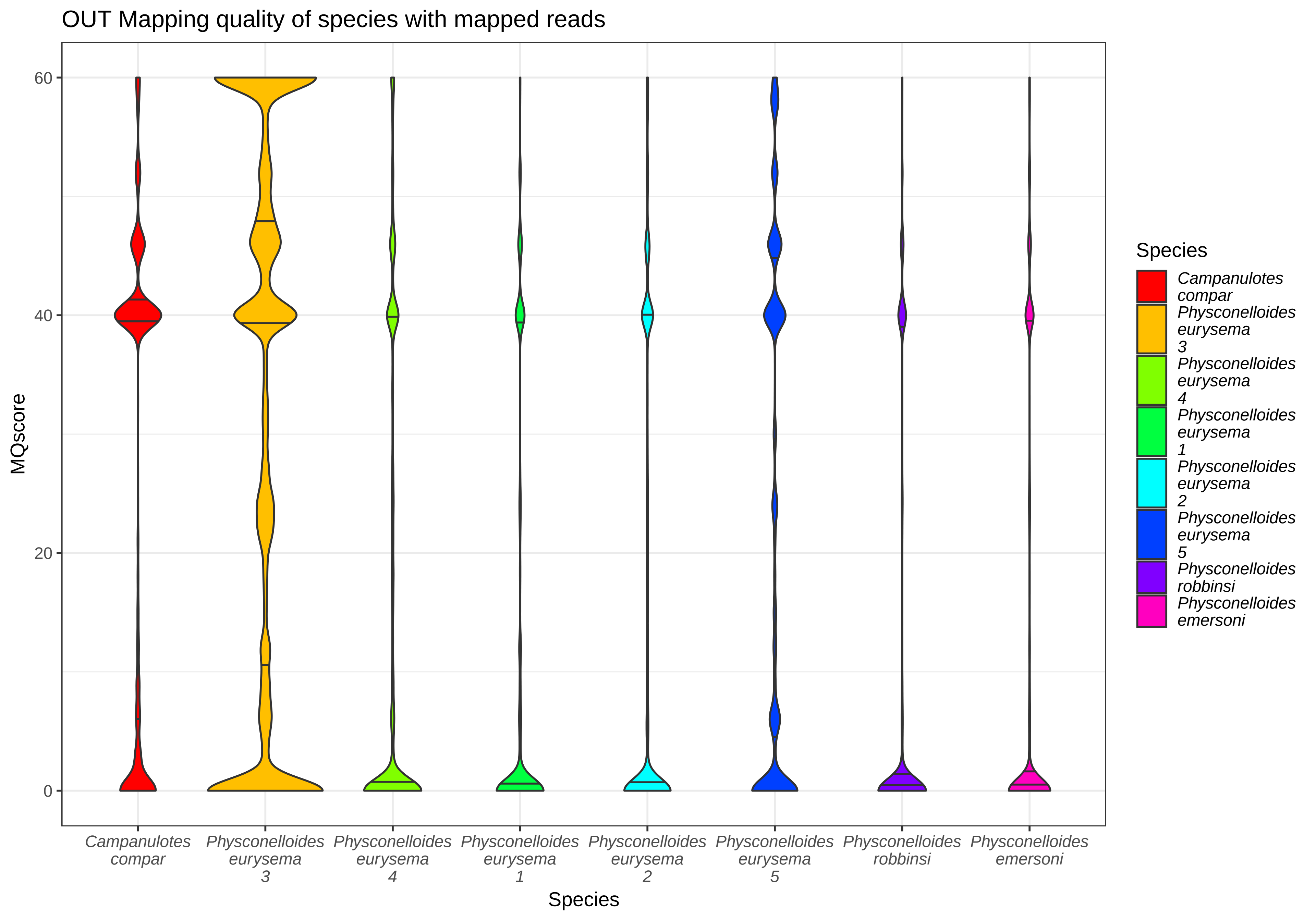

### OUT_plotMQ_Pheur2_Clpre_3_14_2016_11.pdf

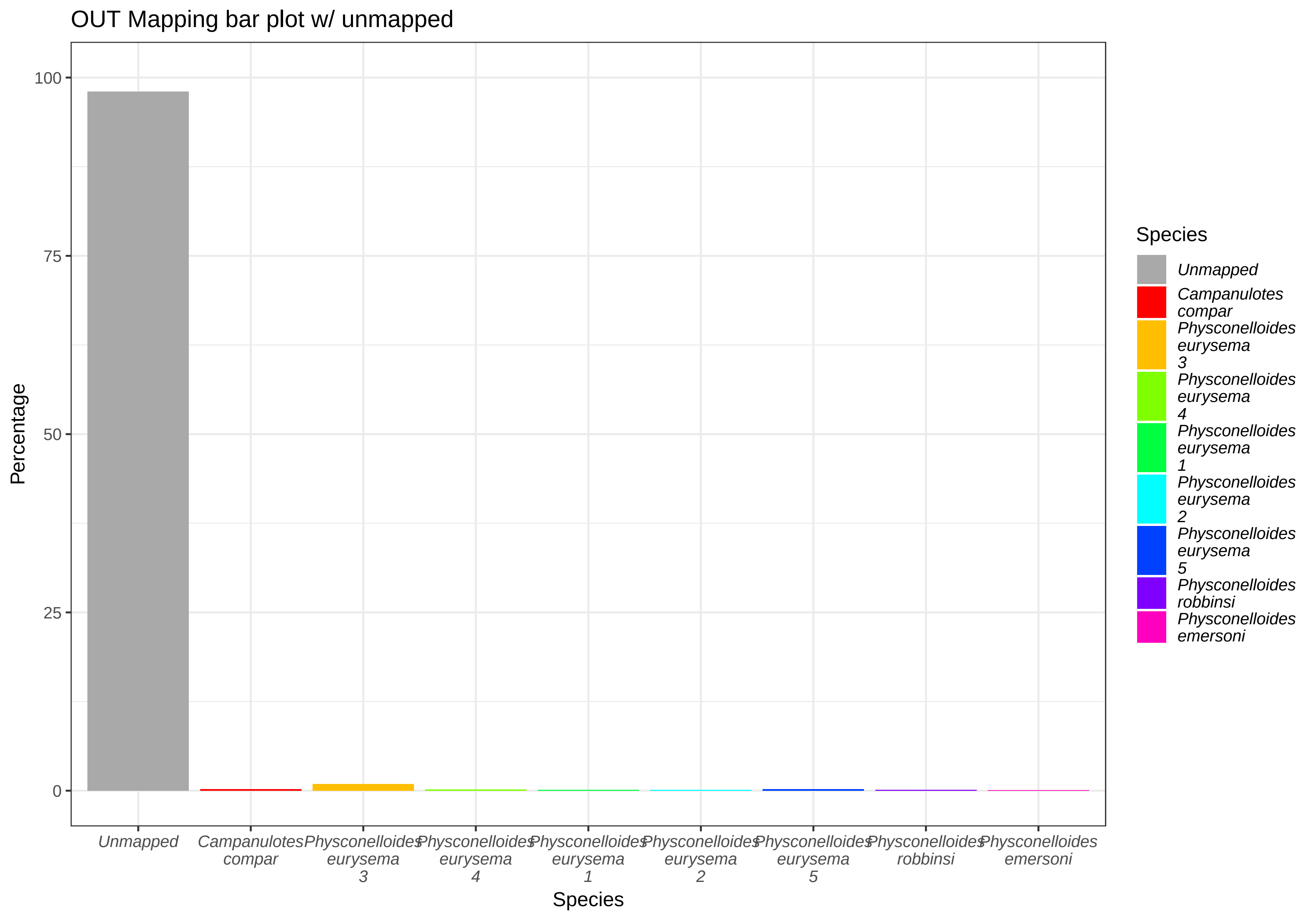

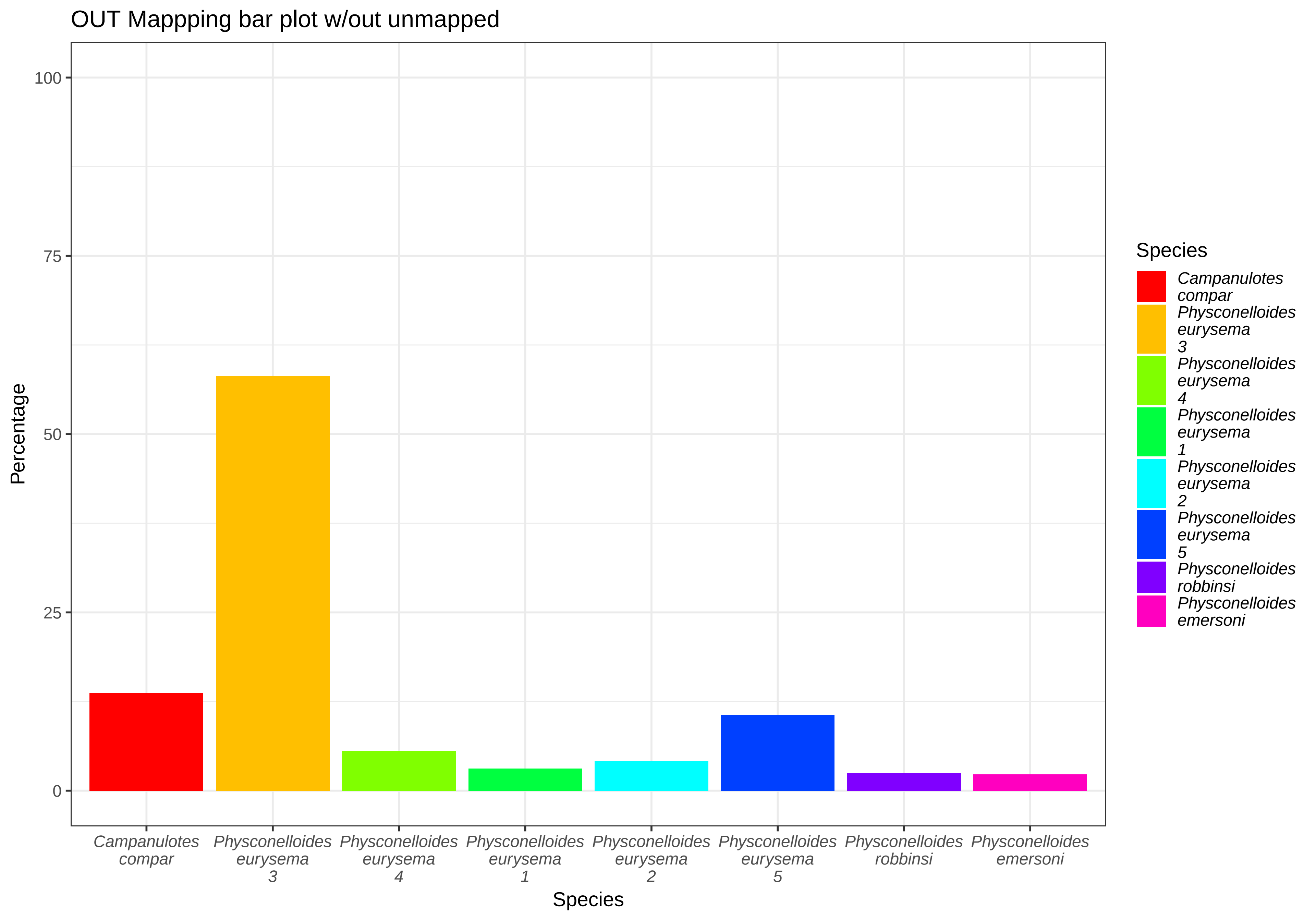

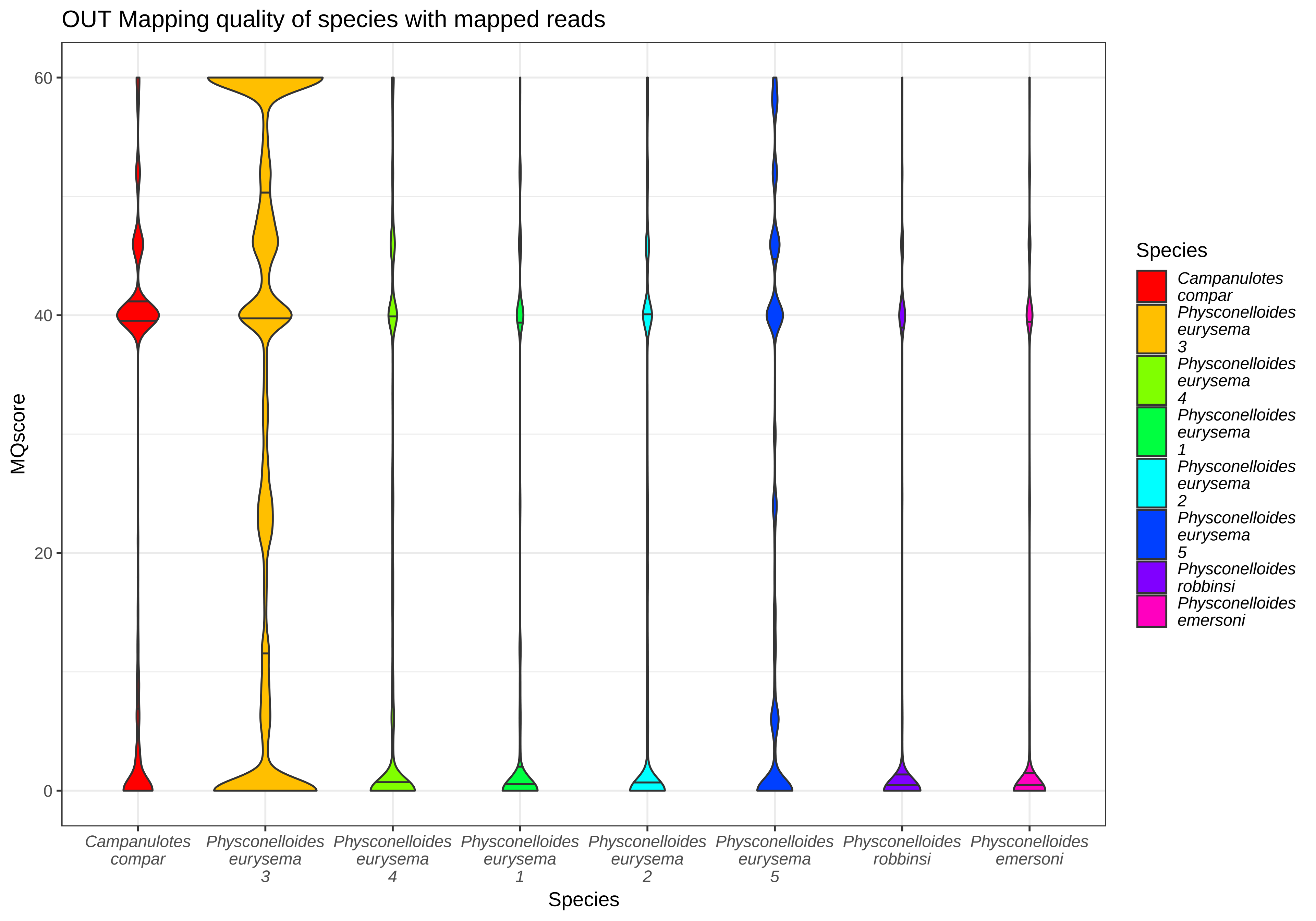

### OUT_plotMQ_Pheur2_Clpre_3_16_2016_5.pdf

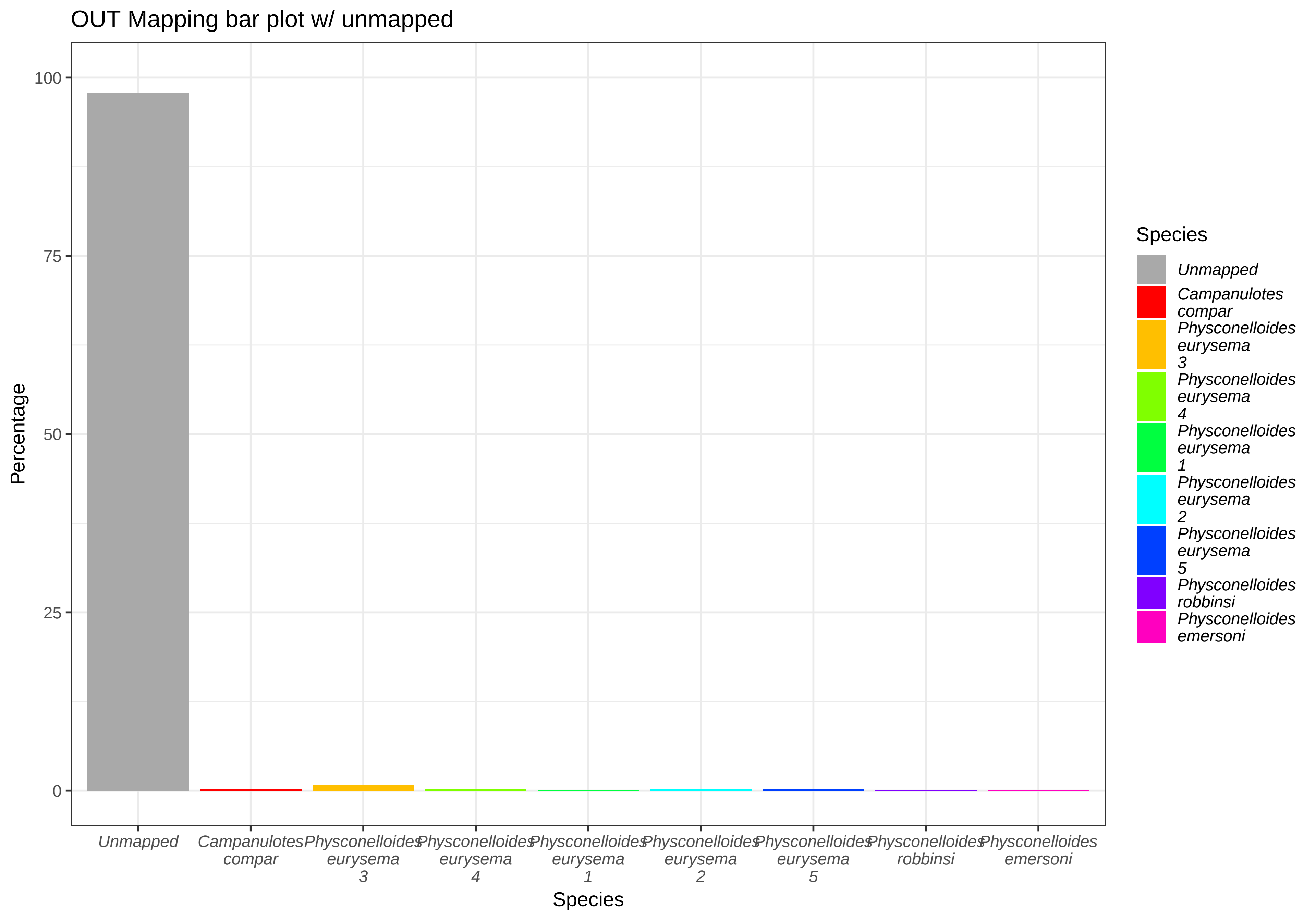

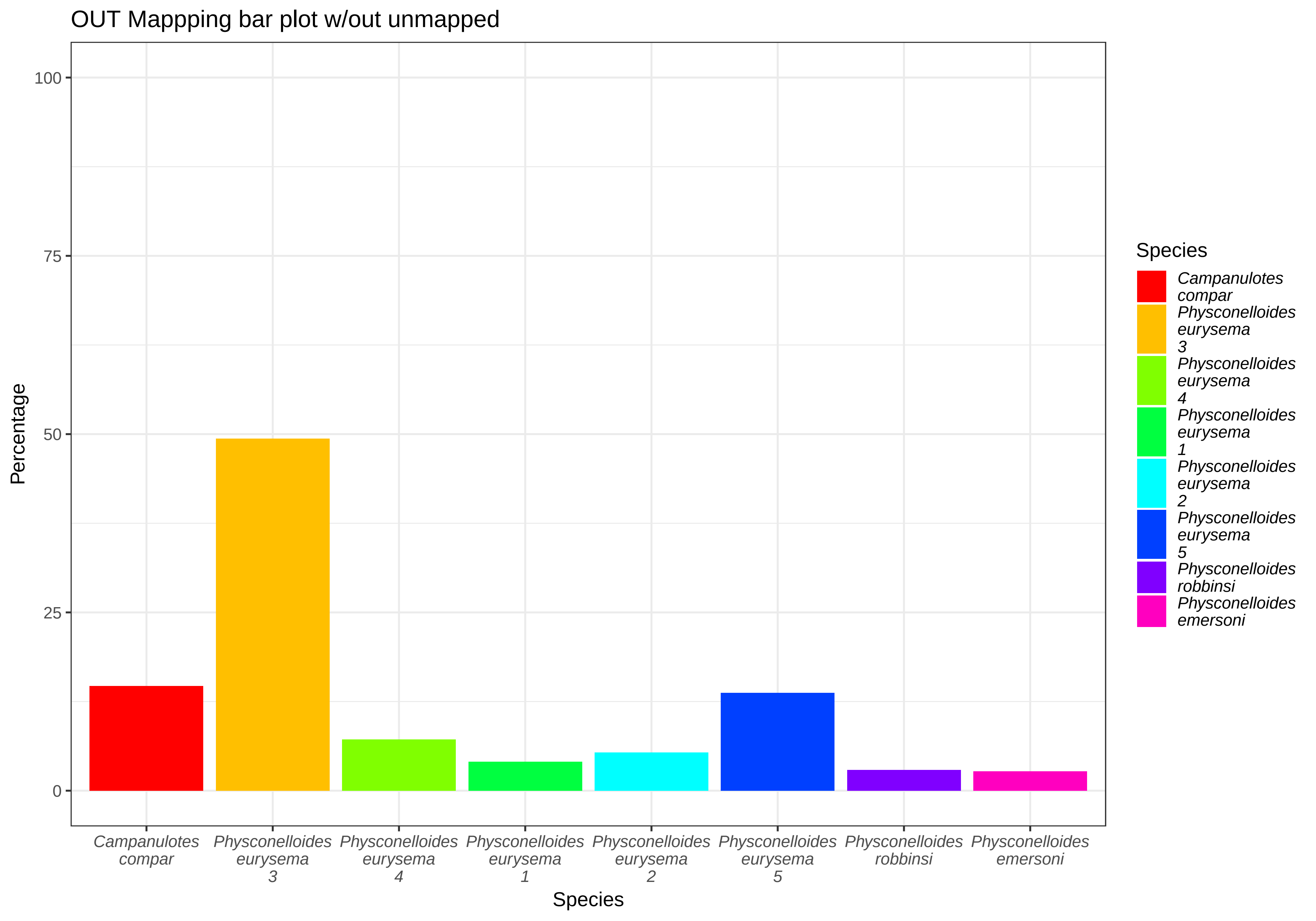

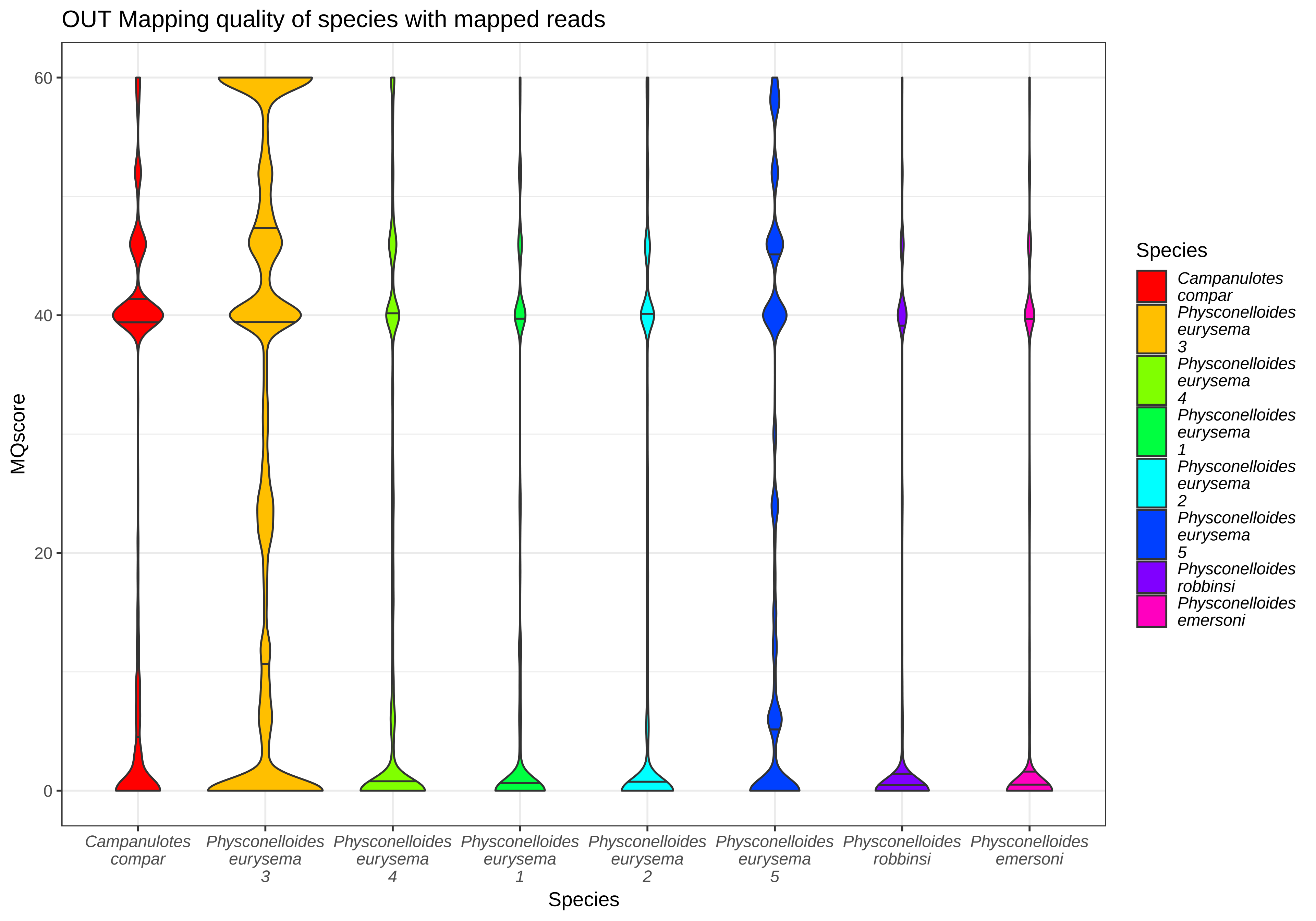

### OUT_plotMQ_Pheur2_Clpre_5_26_2015_6.pdf

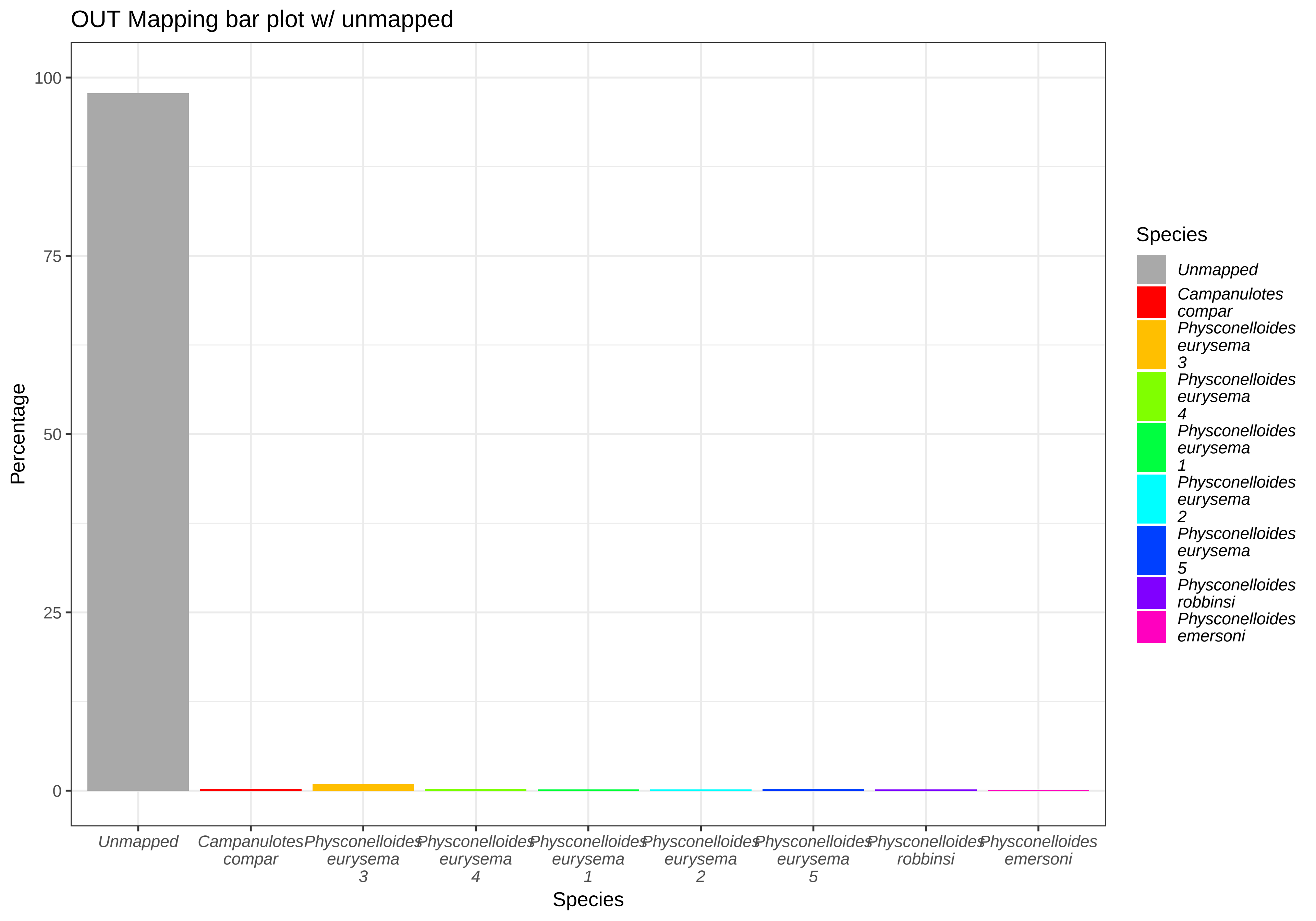

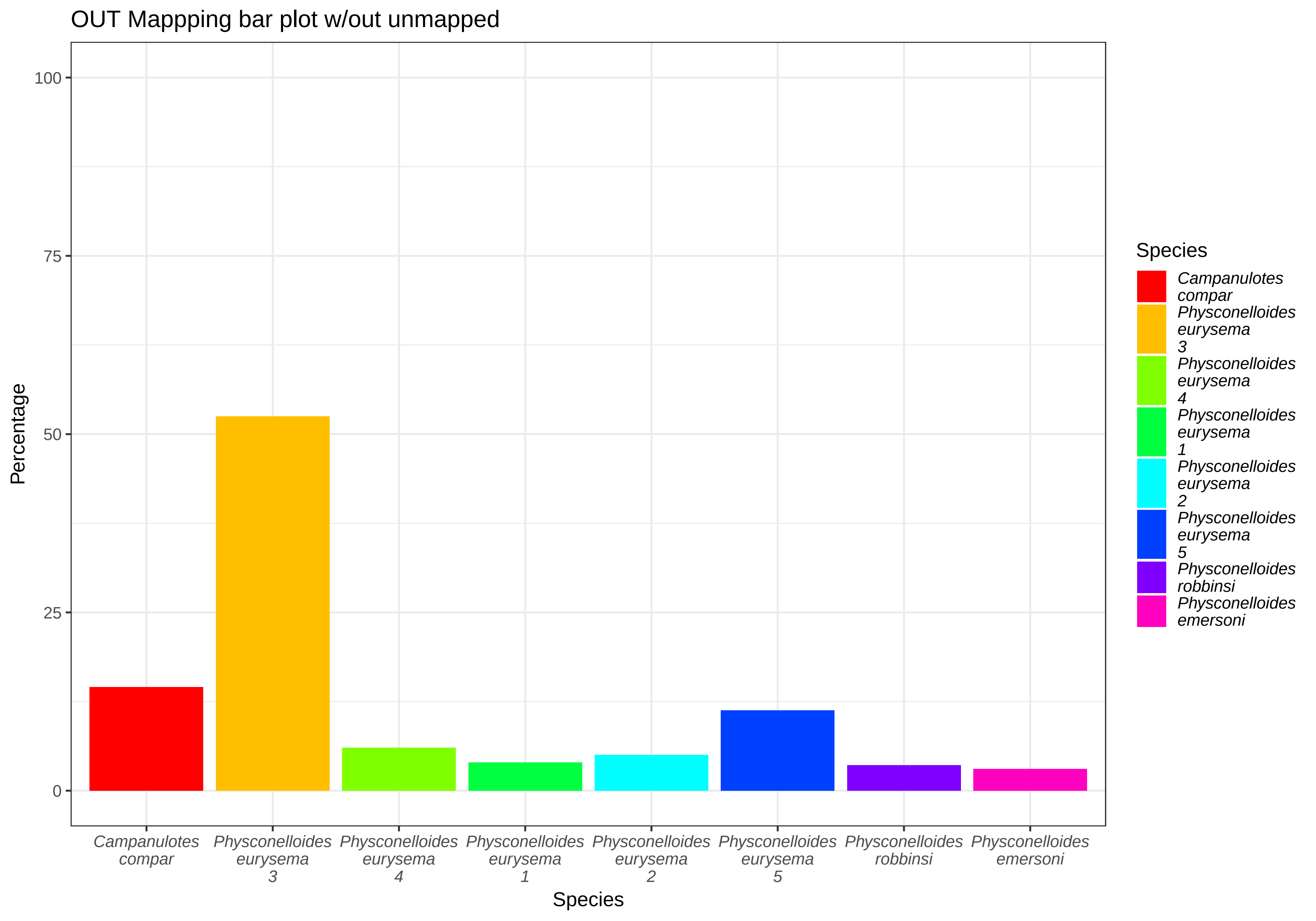

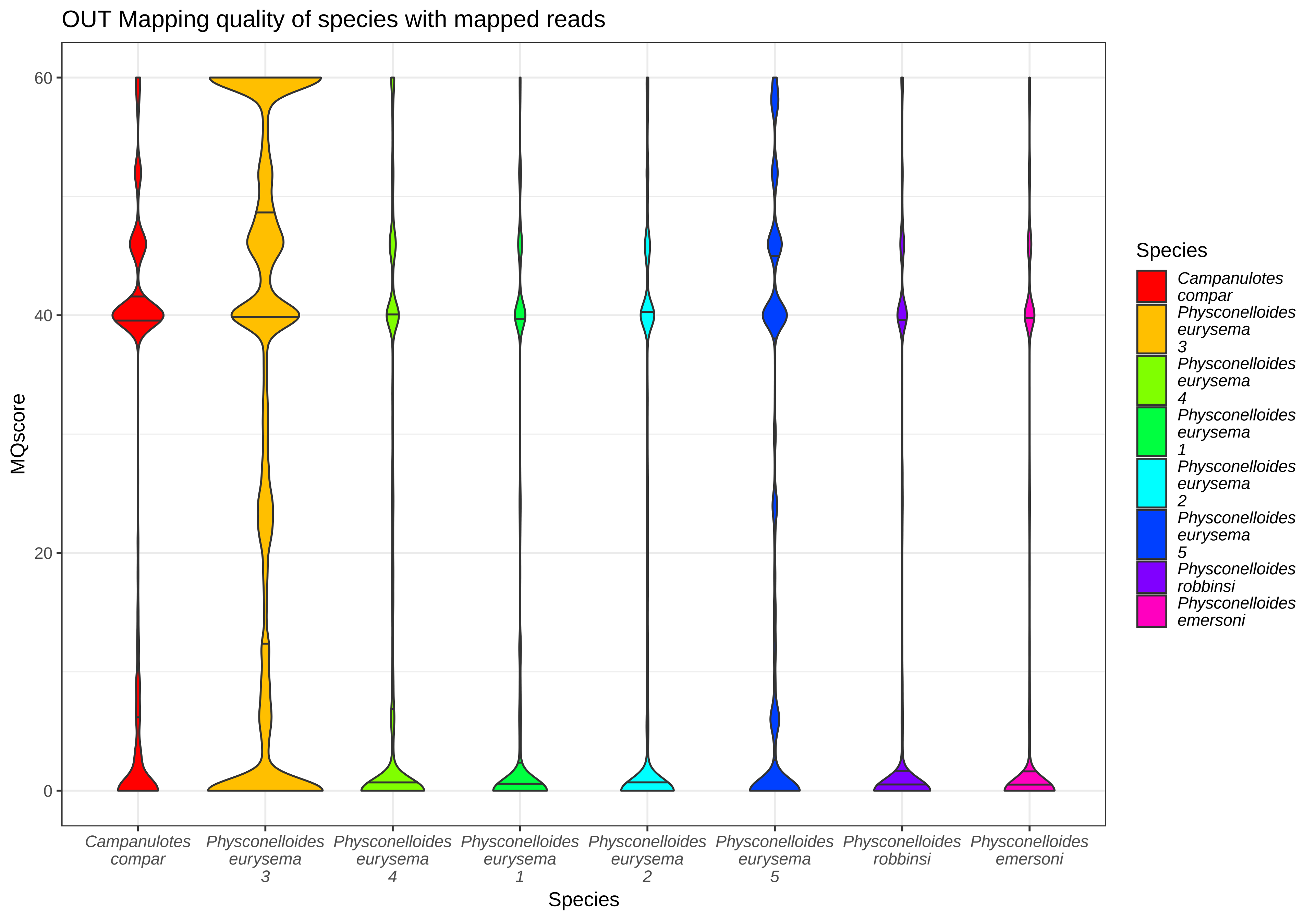

### OUT_plotMQ_Pheur2_Cobuc_3_16_2016_4.pdf

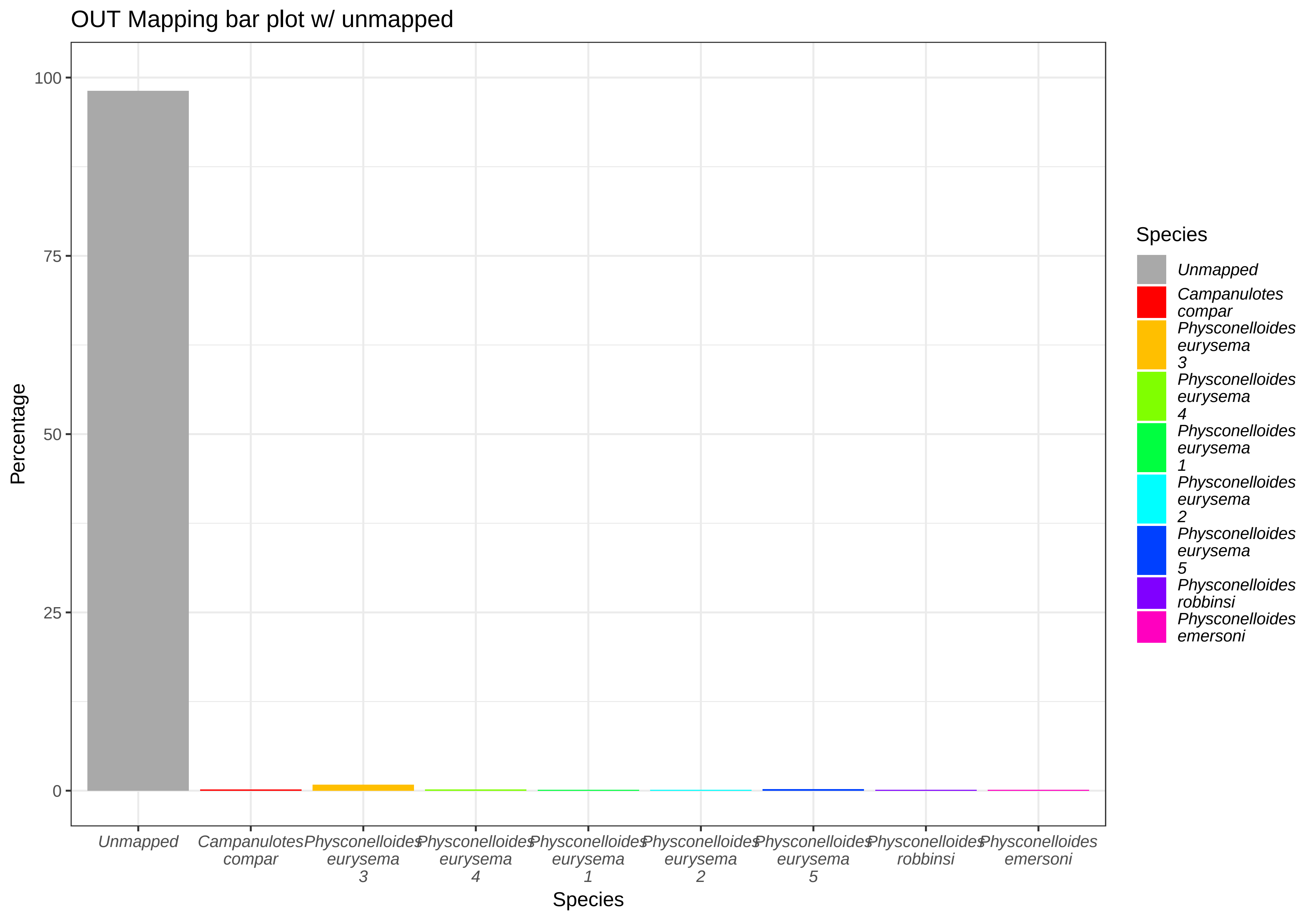

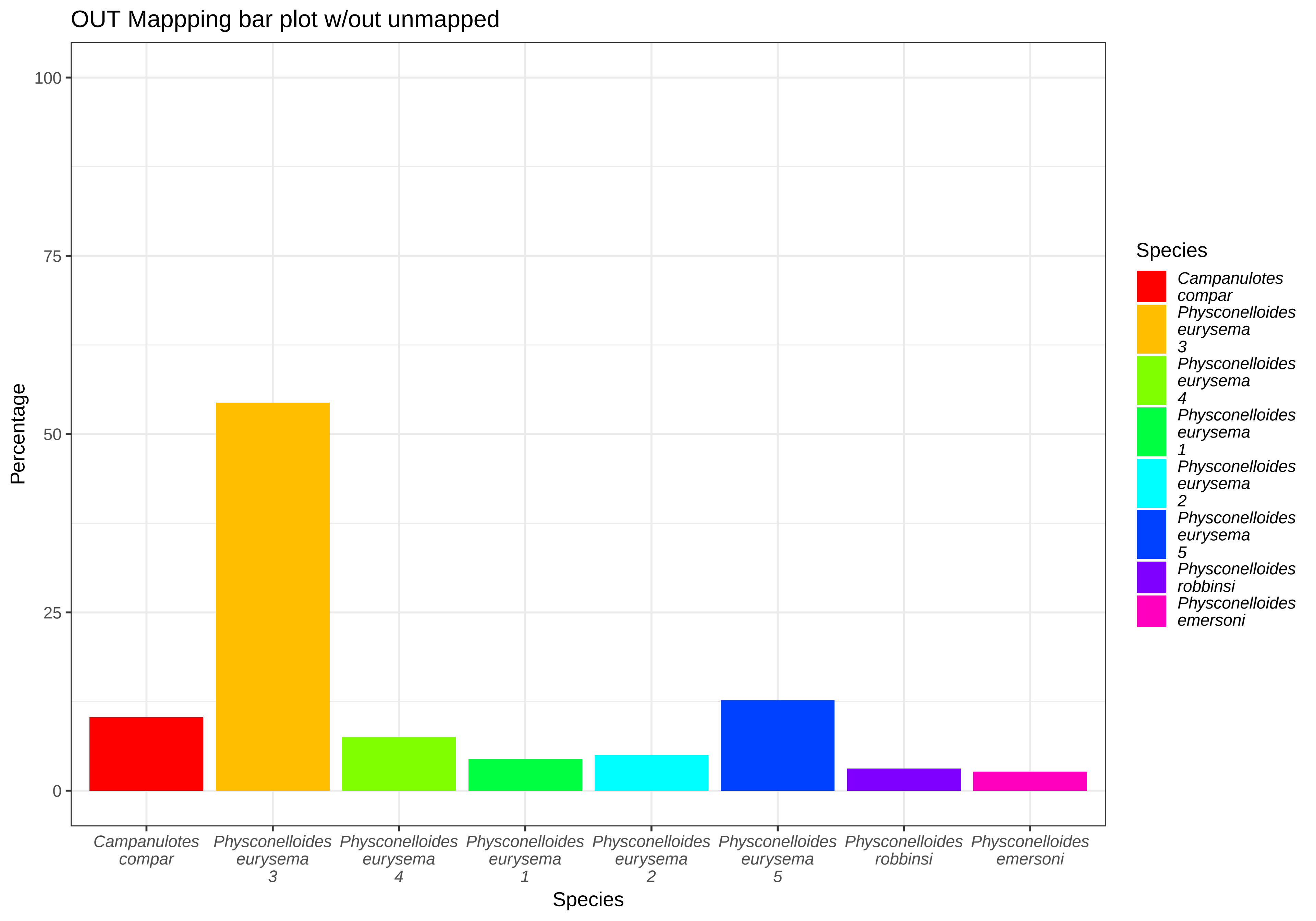

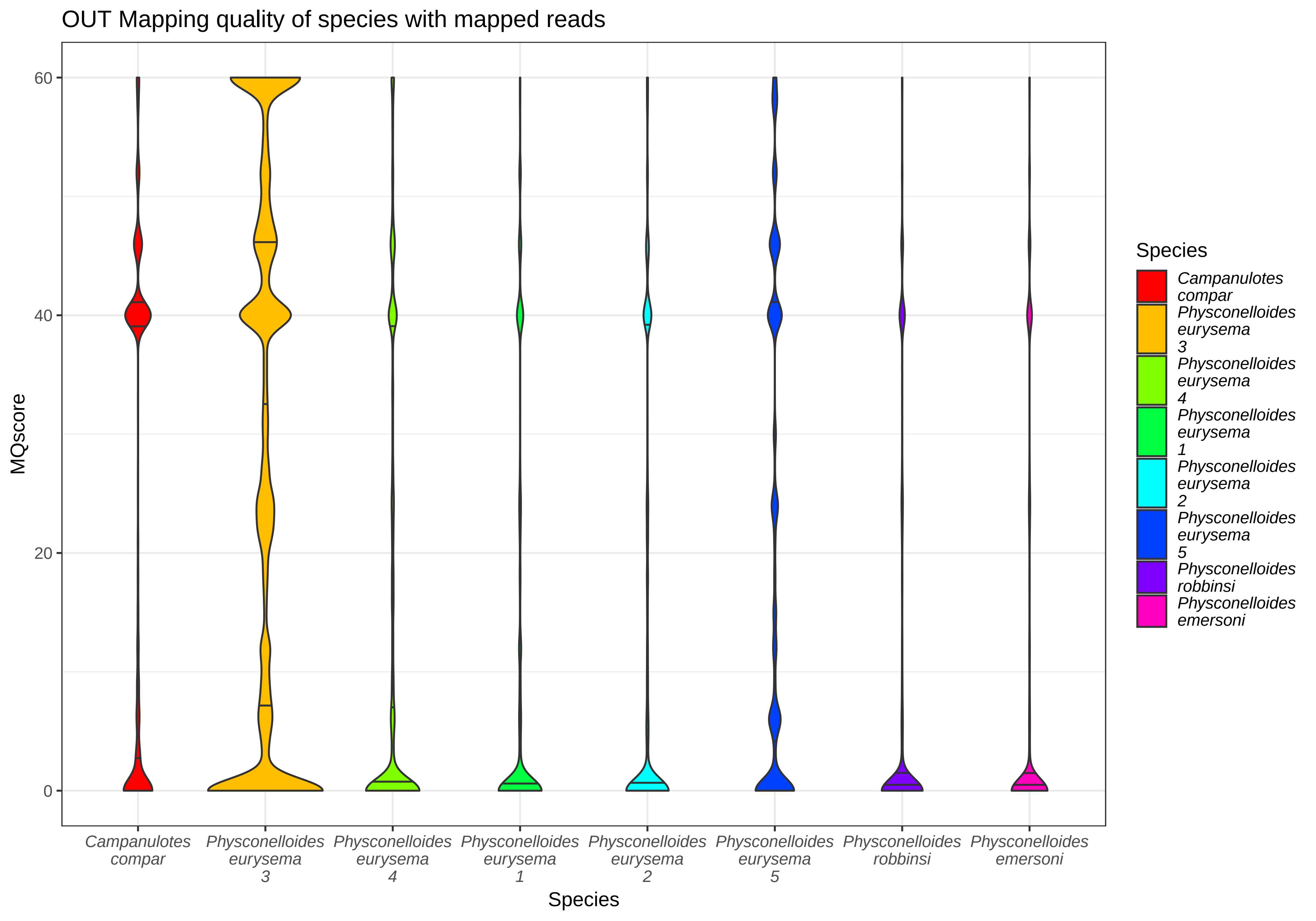

### OUT_plotMQ_Pheur2_Cocru_3_16_2016_9.pdf

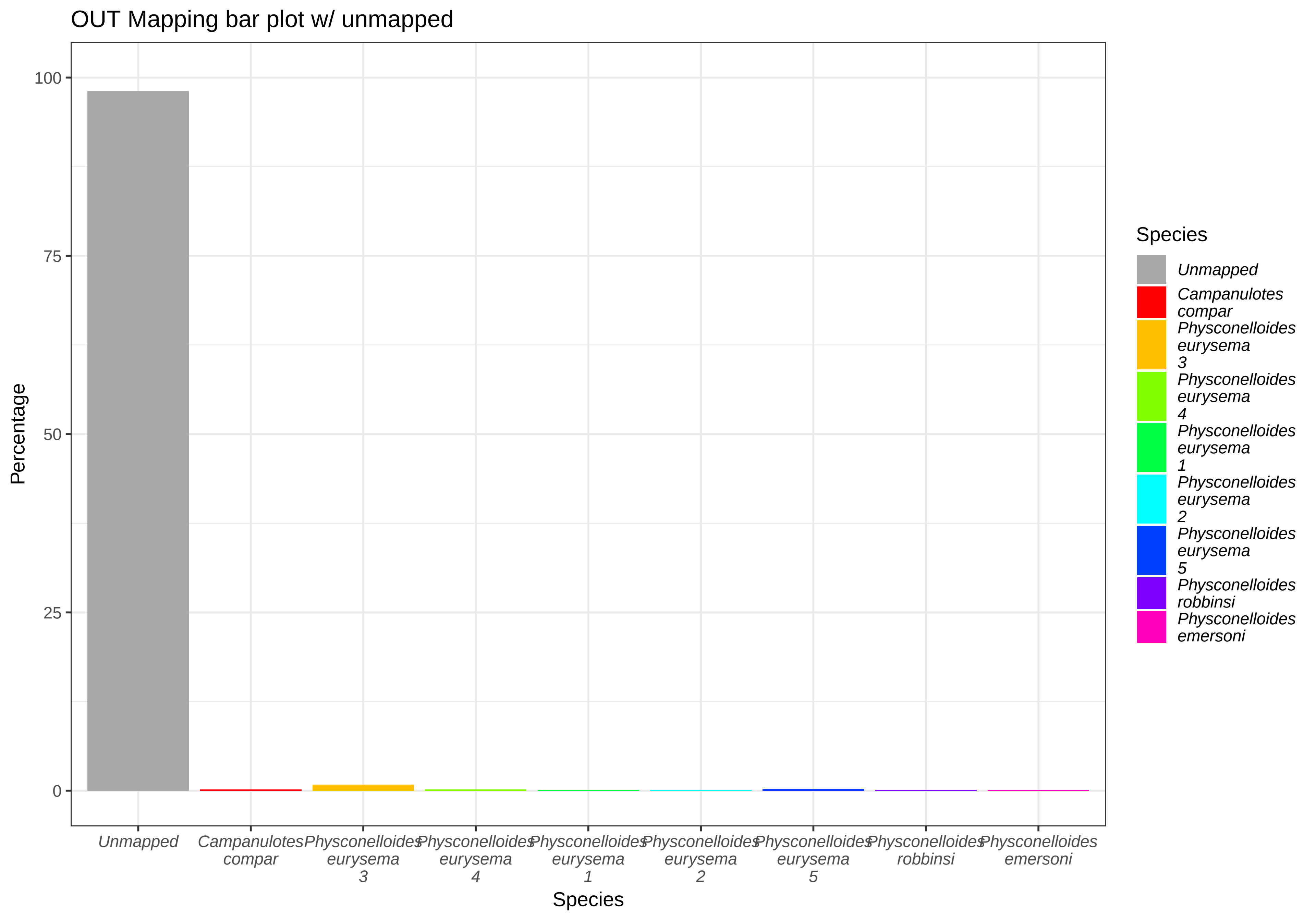

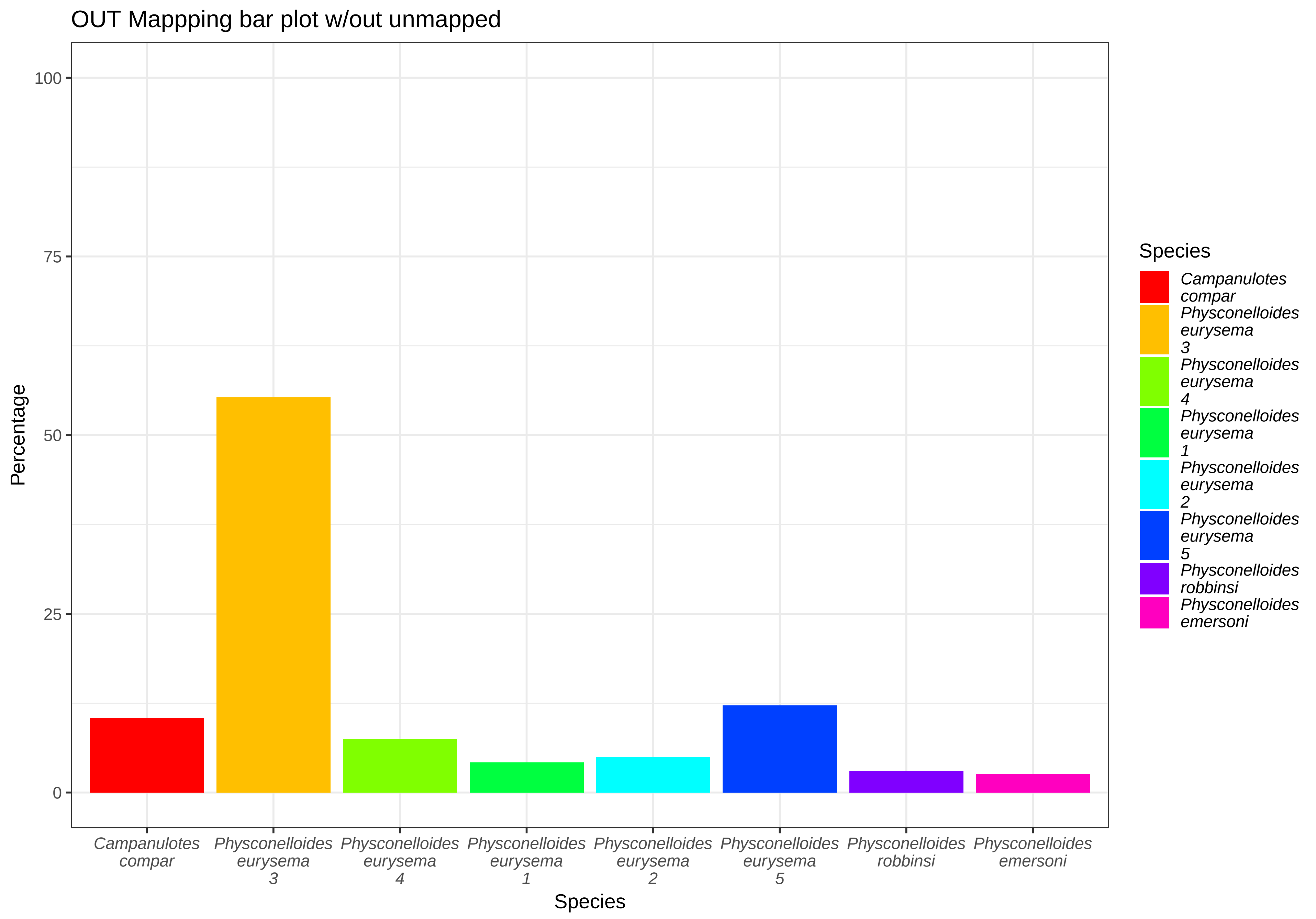

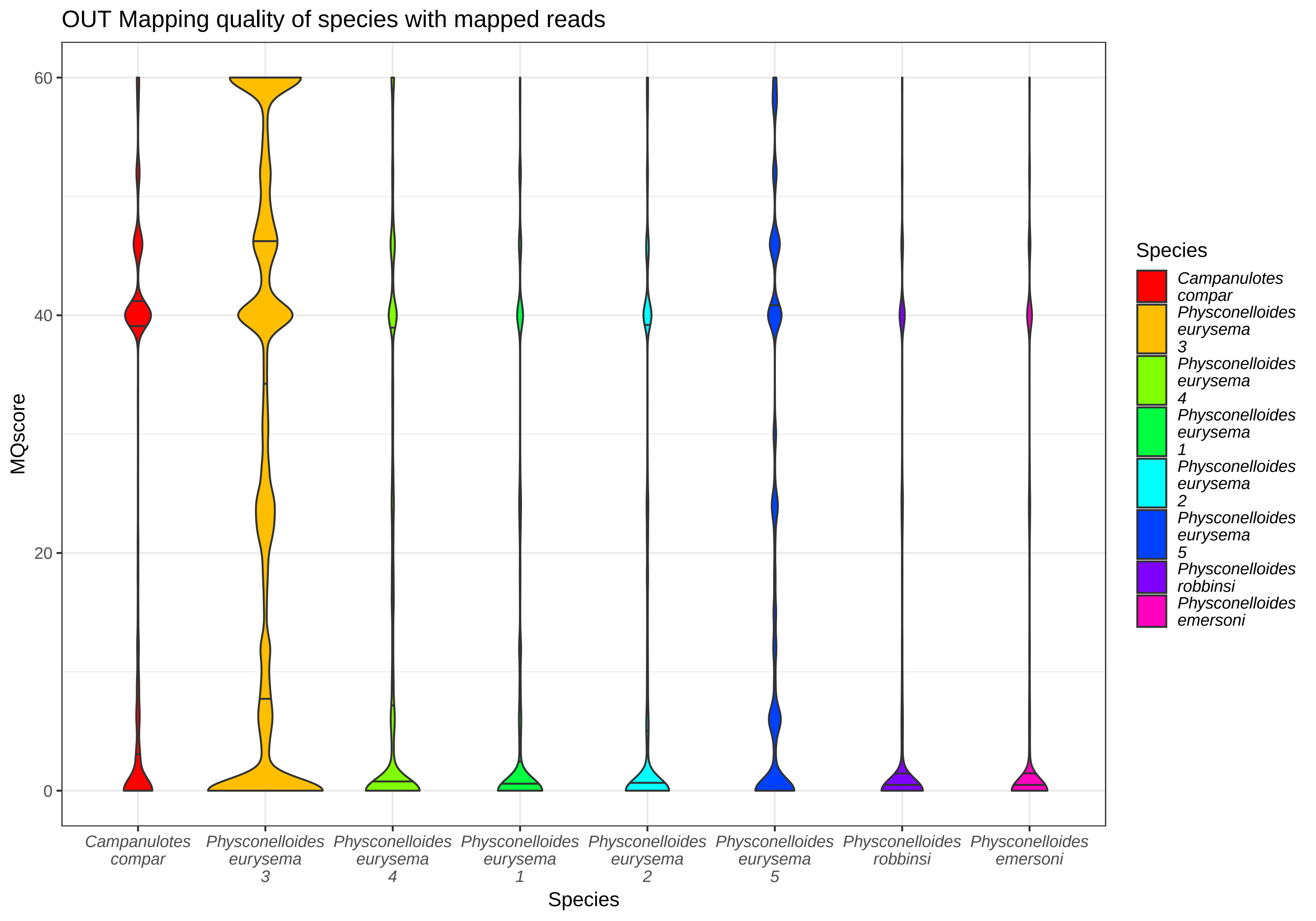

### OUT_plotMQ_Pheur.Comin.6.15.2015.10.pdf

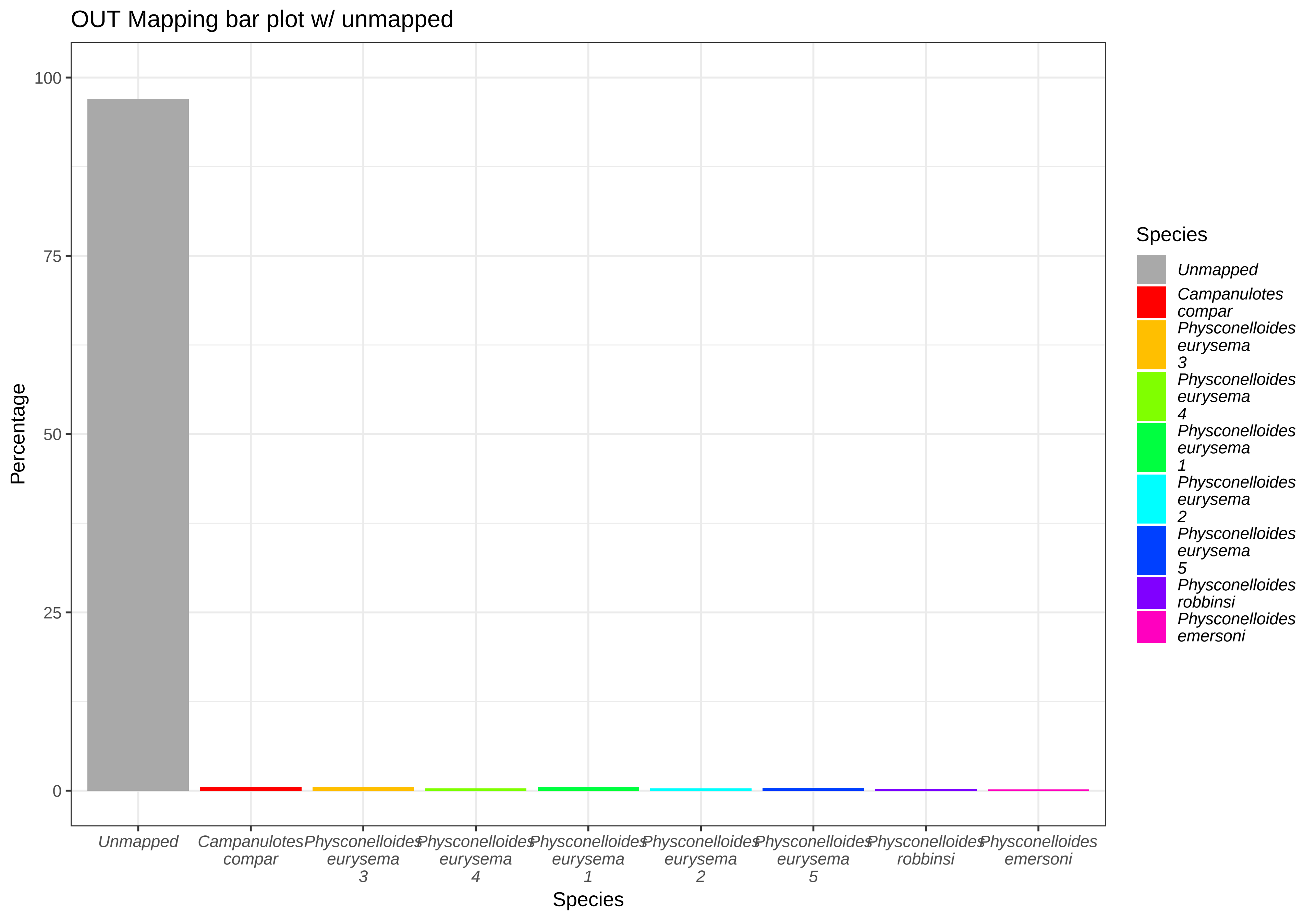

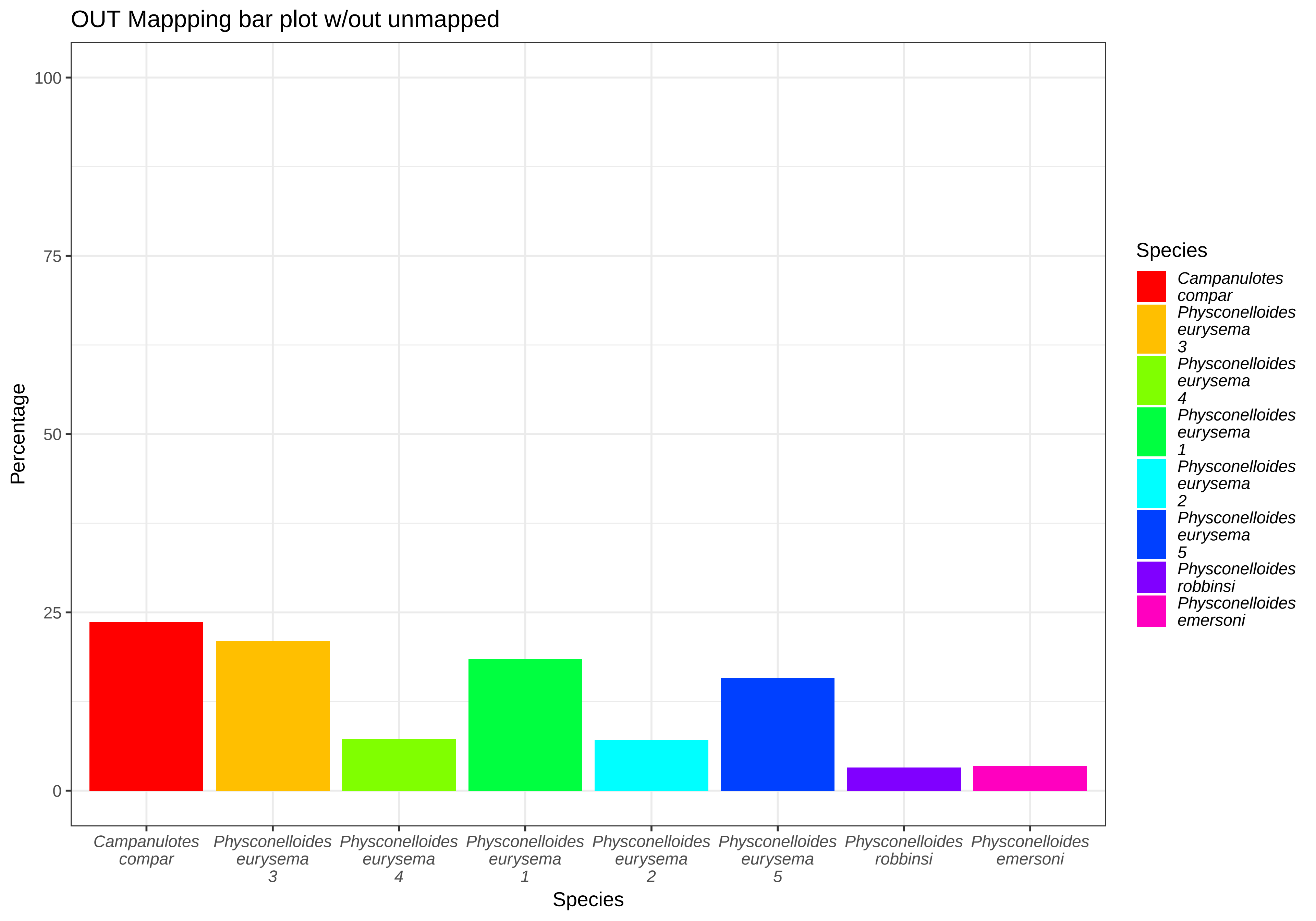

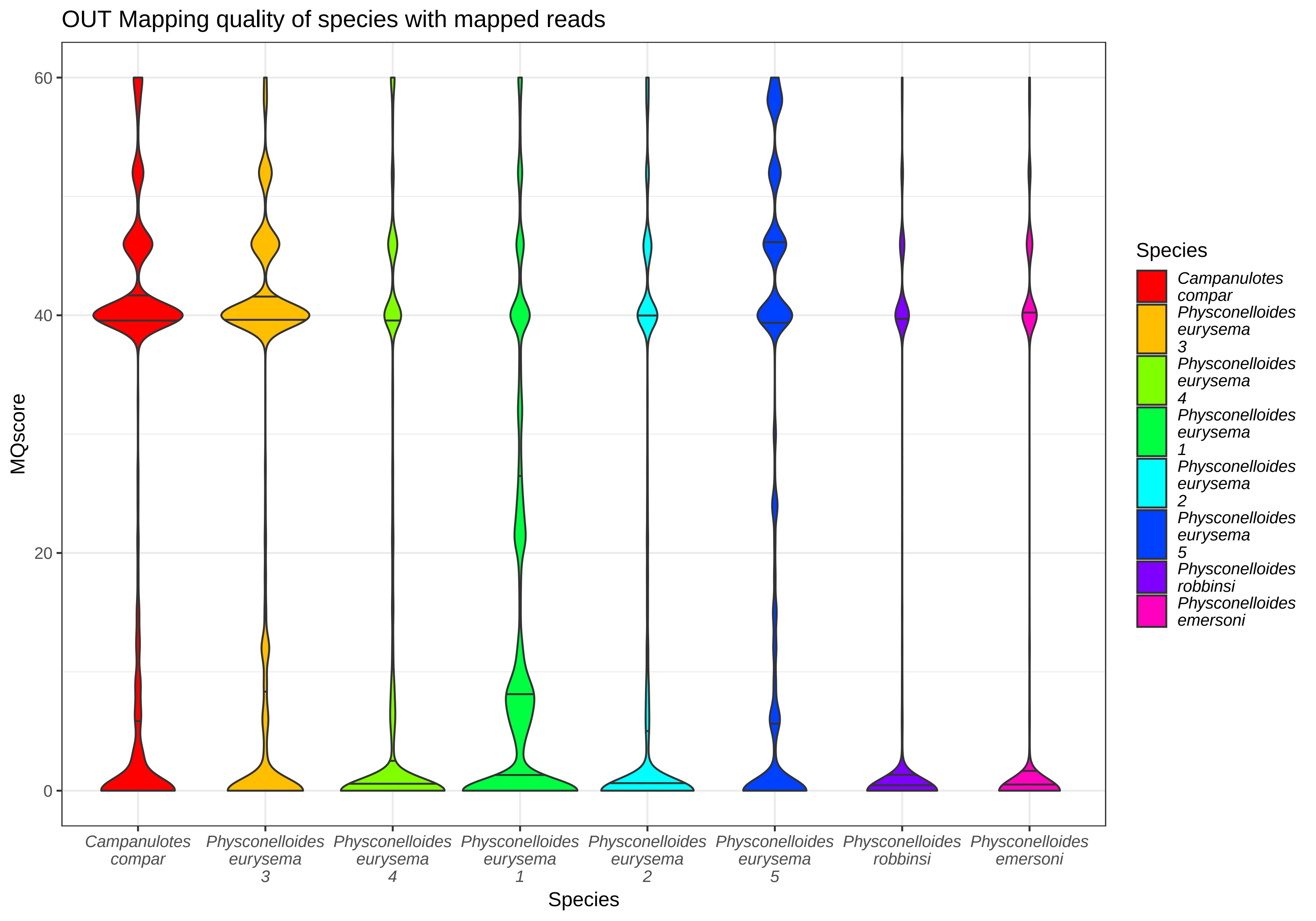

### OUT_plotMQ_Pheur_Clpre_6_15_2015_8.pdf

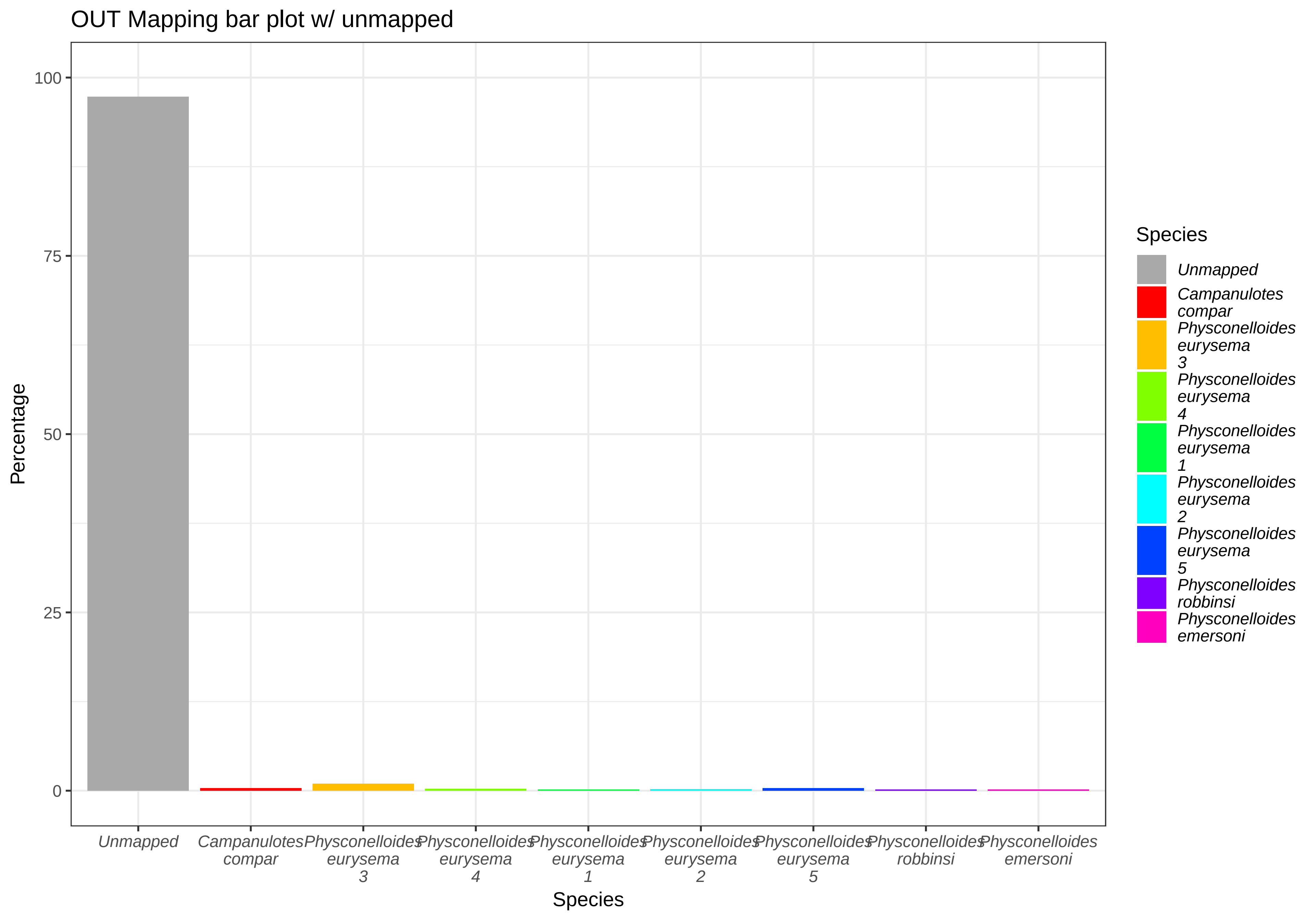

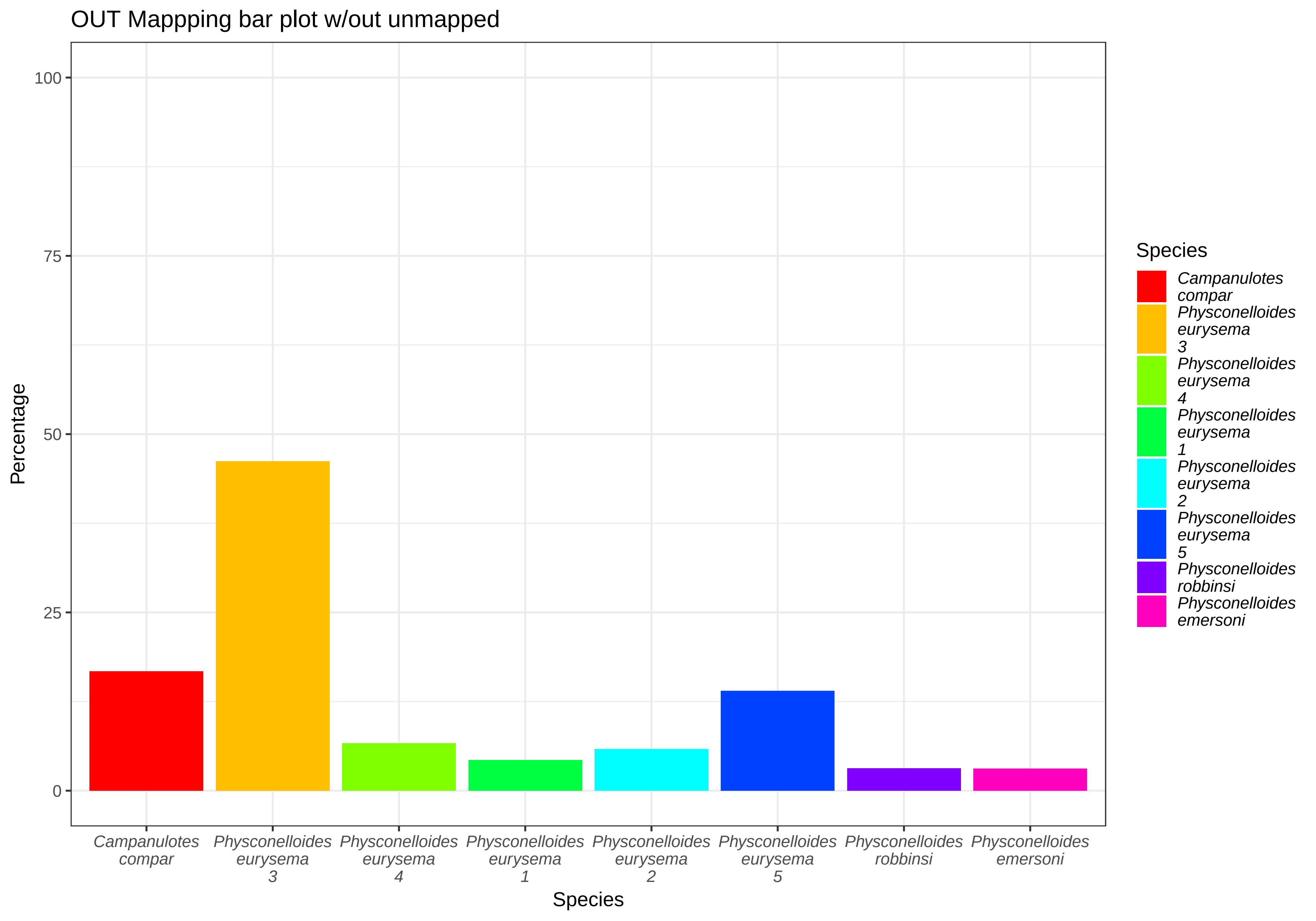

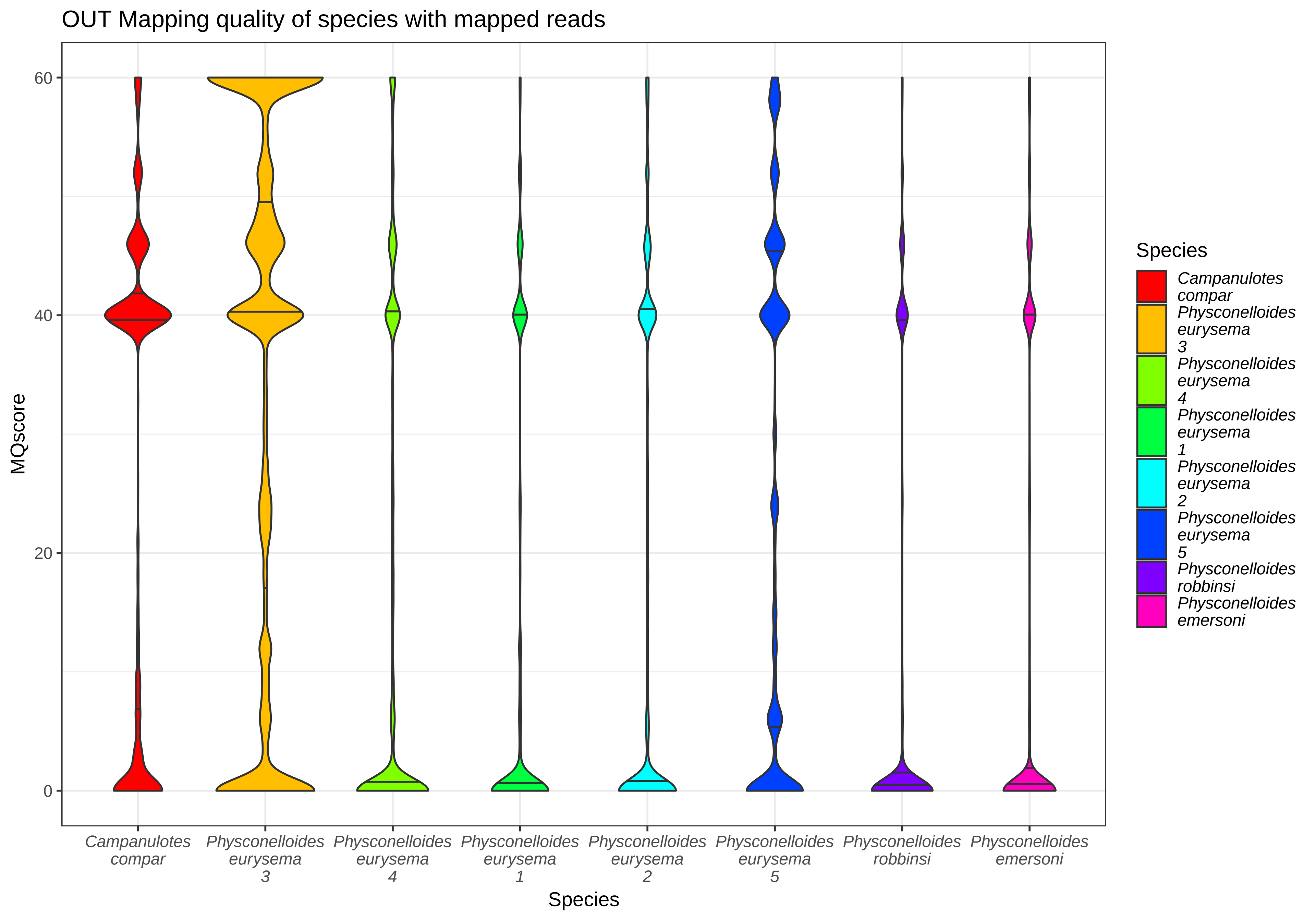

### OUT_plotMQ_Pheur_Cobuc_6_15_2015_1.pdf

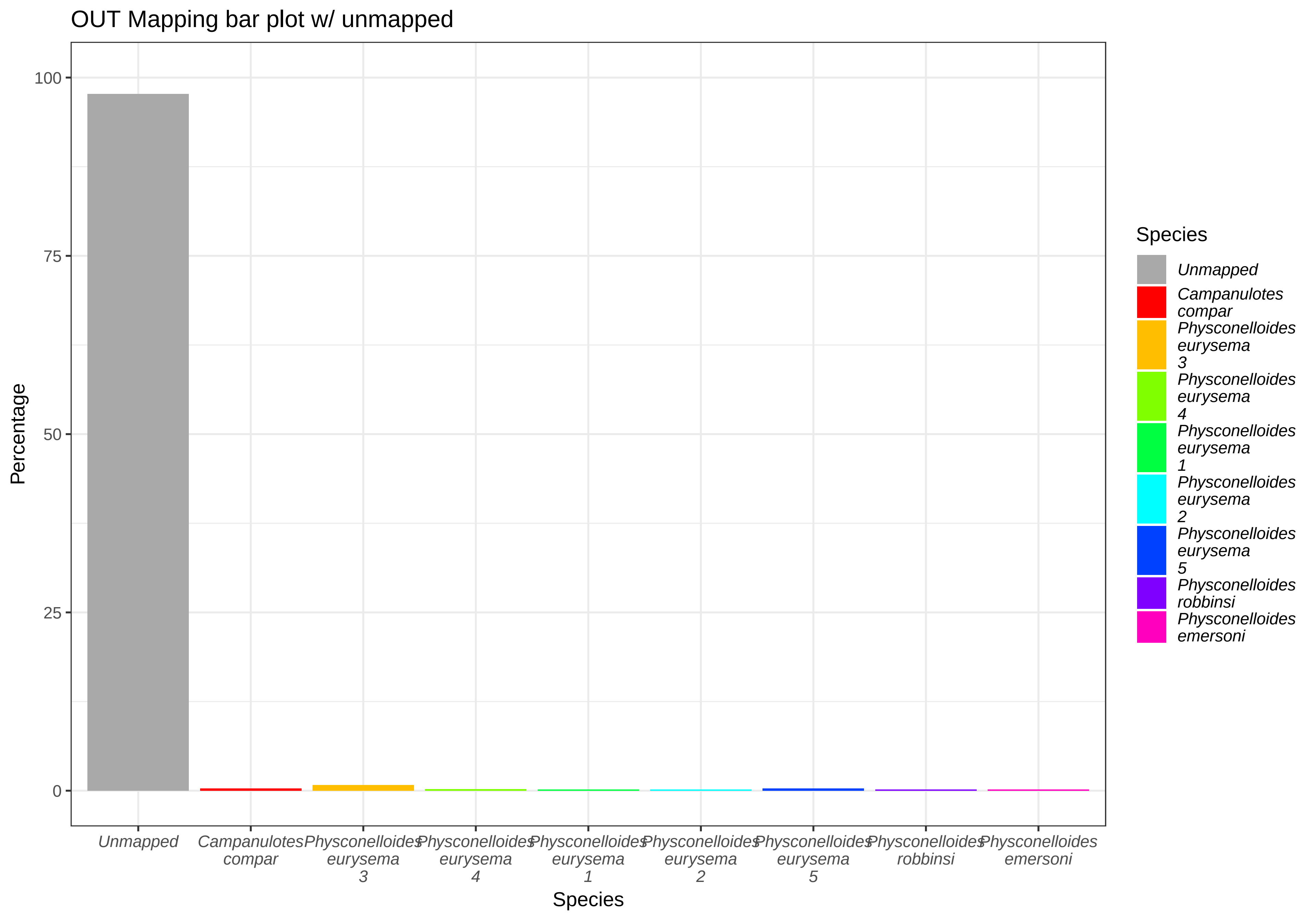

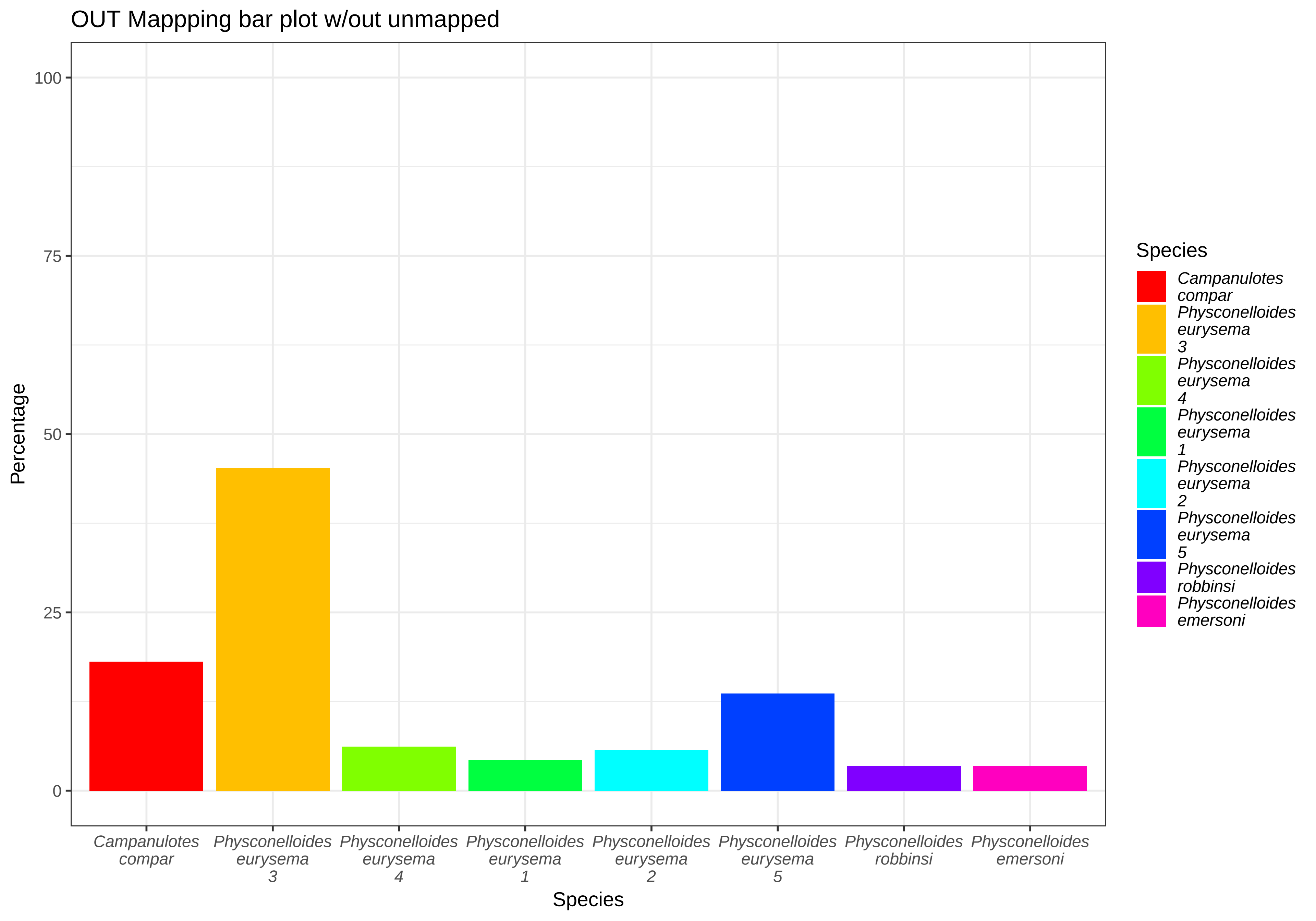

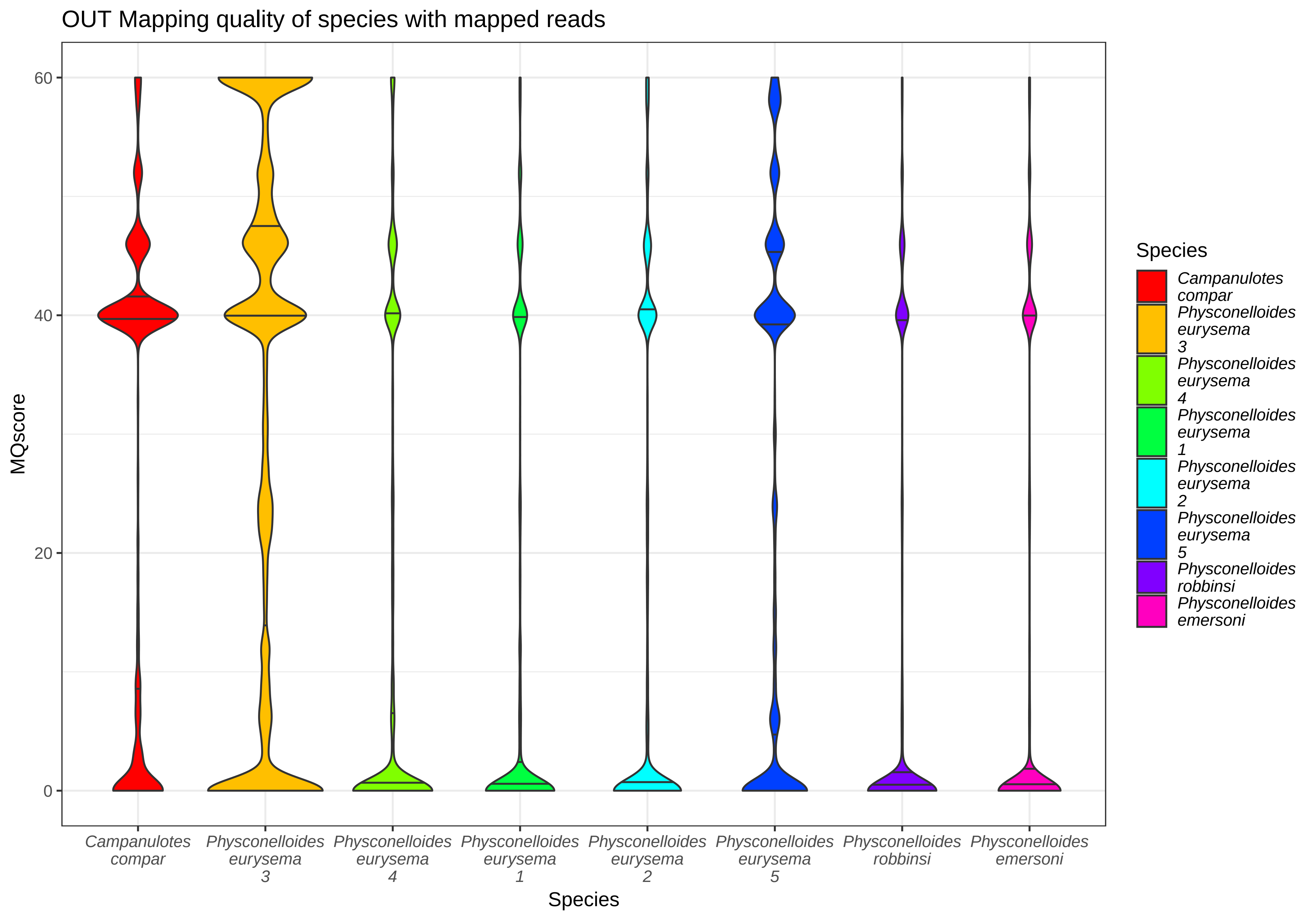

### OUT_plotMQ_Pheur_Cocru_6_15_2015_2.pdf

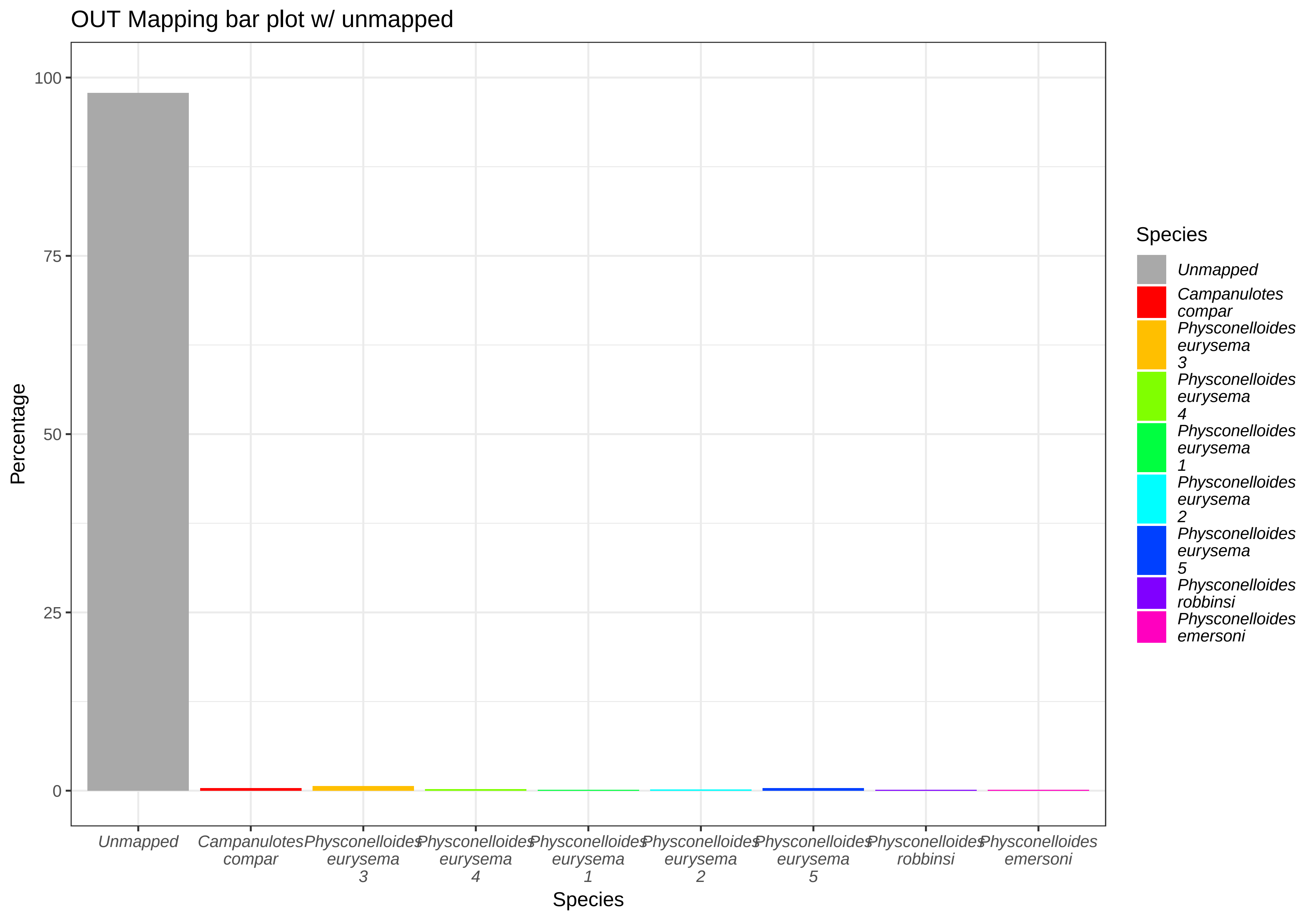

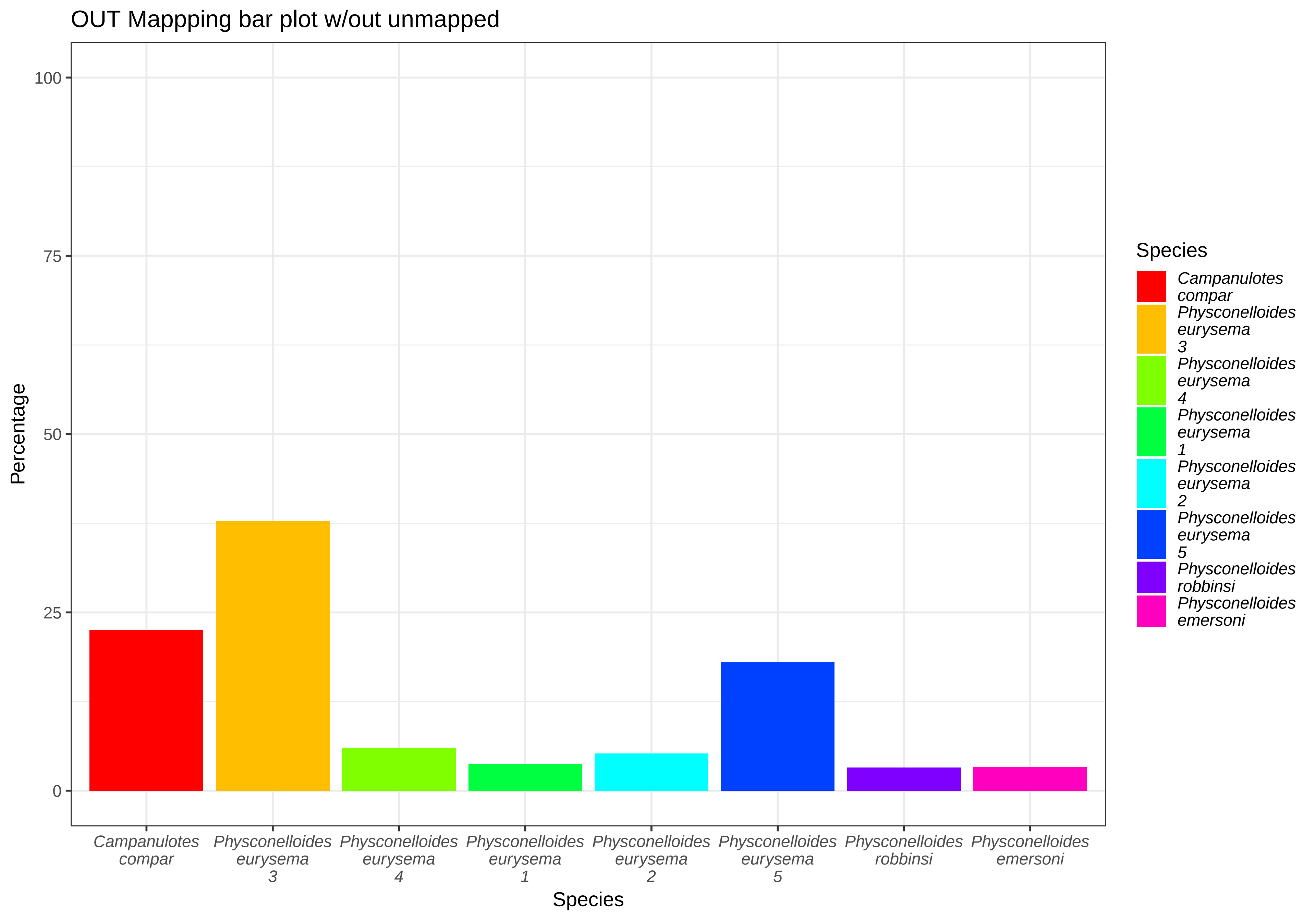

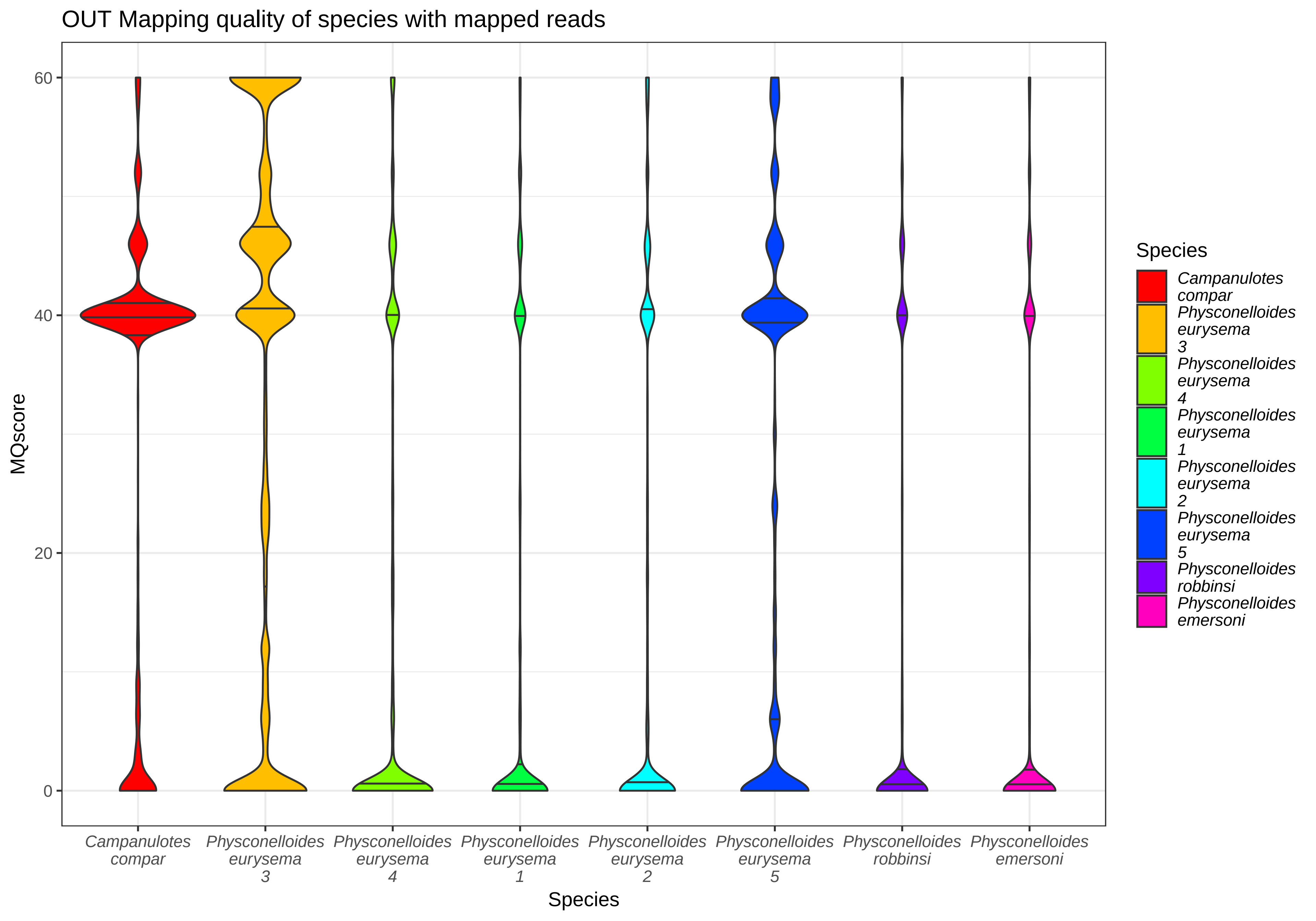

### OUT_plotMQ_Pheur_Coinc_6_15_2015_7.pdf

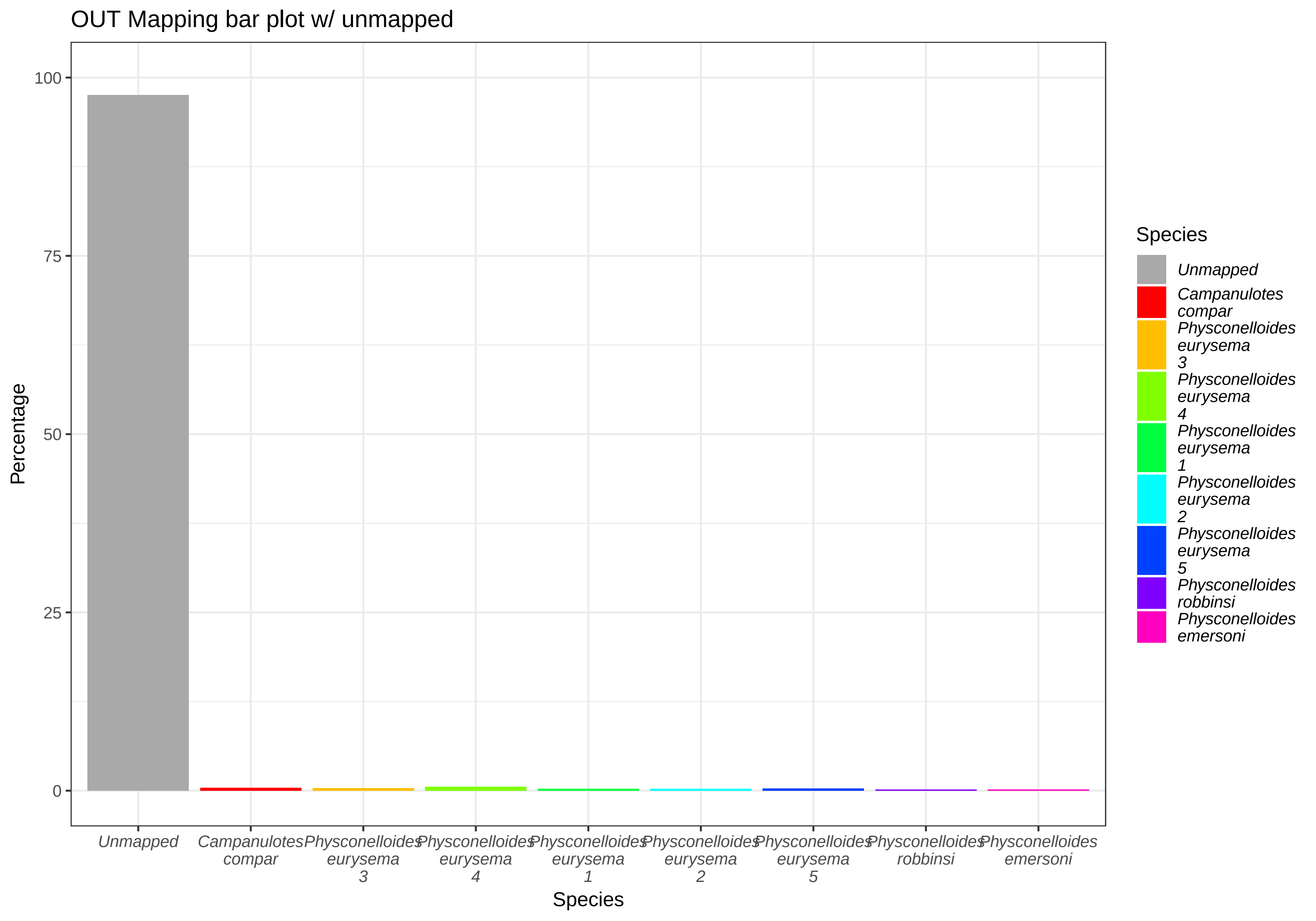

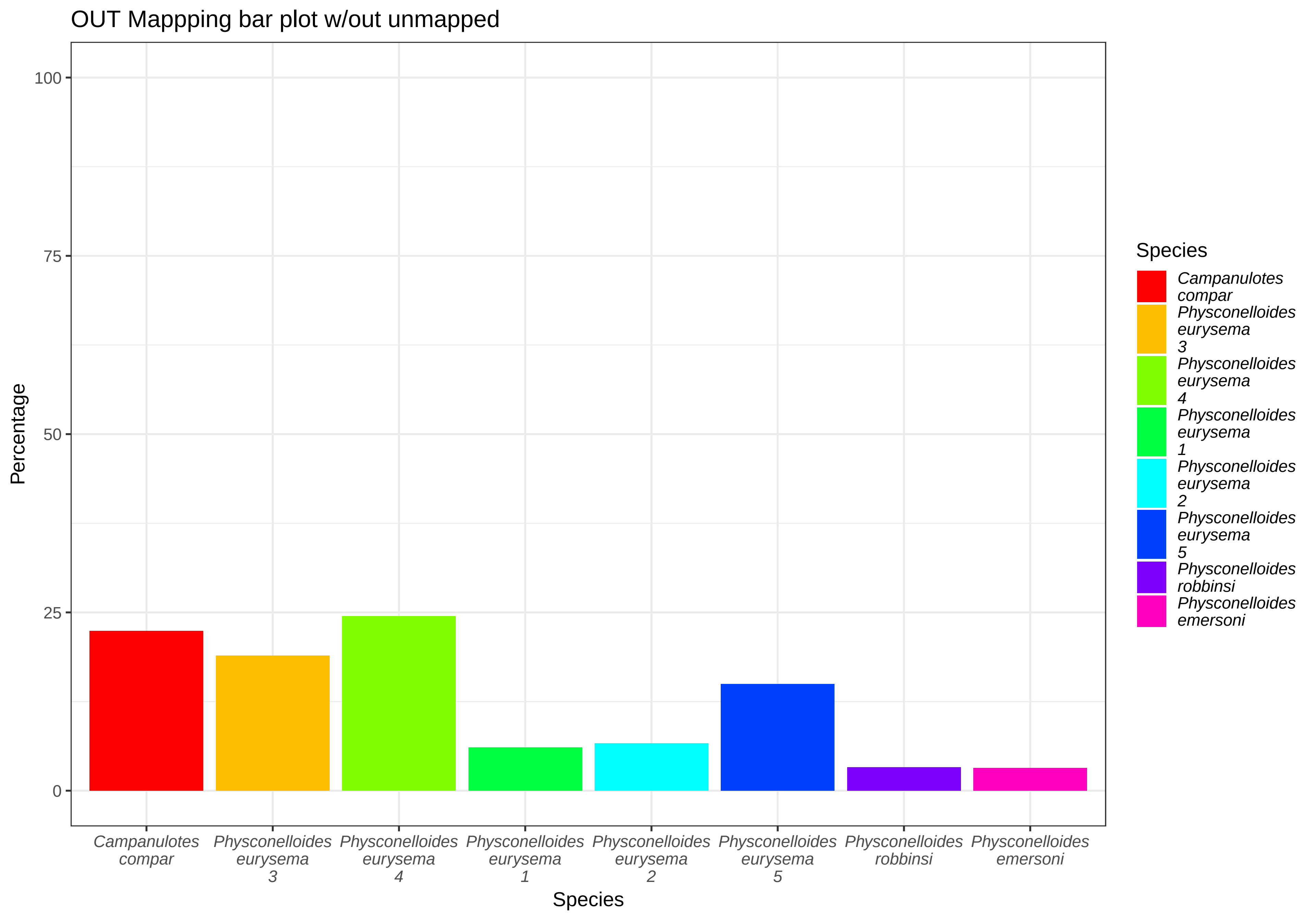

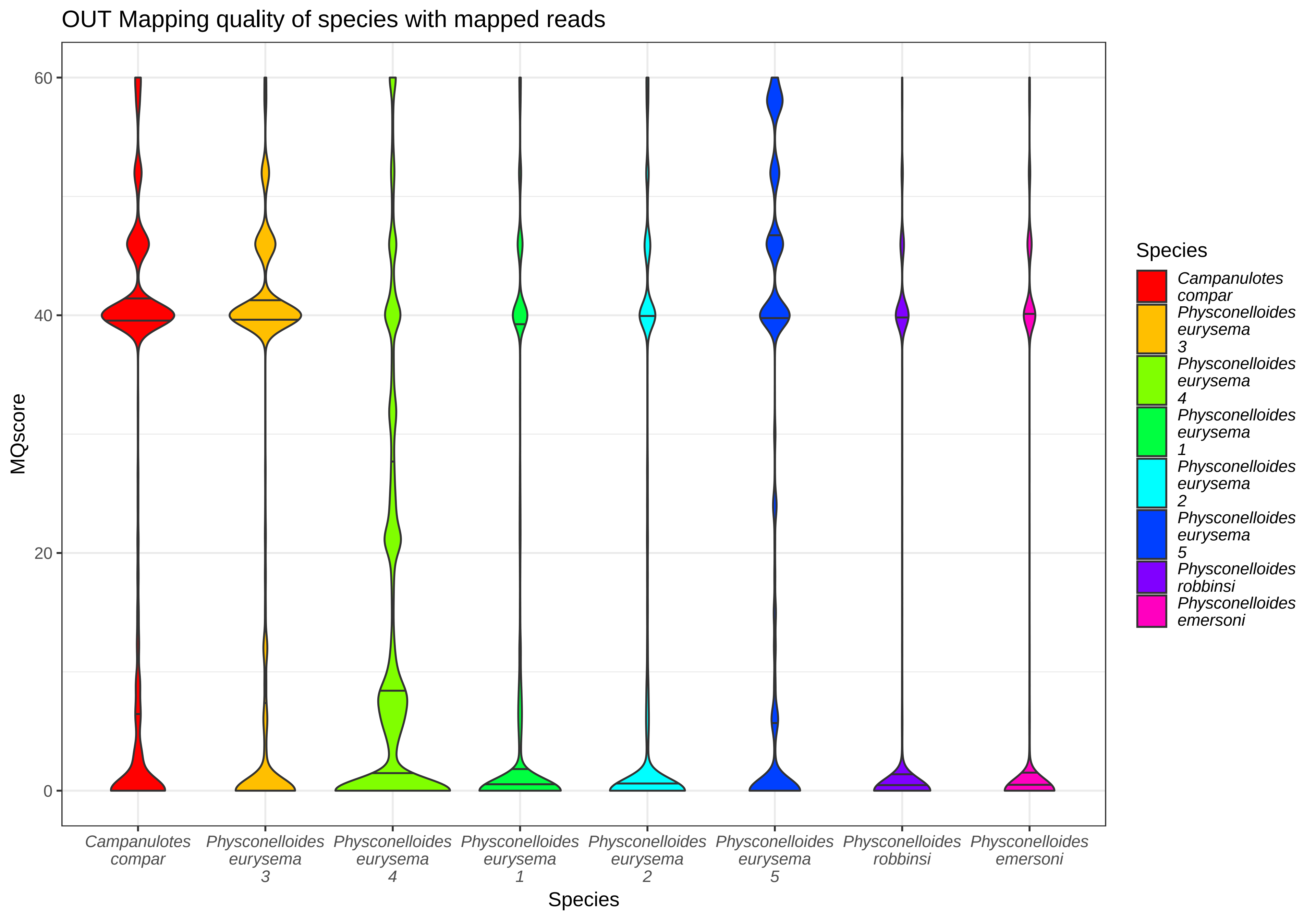

### OUT_plotMQ_Pheur_Copas_3_14_2016_4.pdf

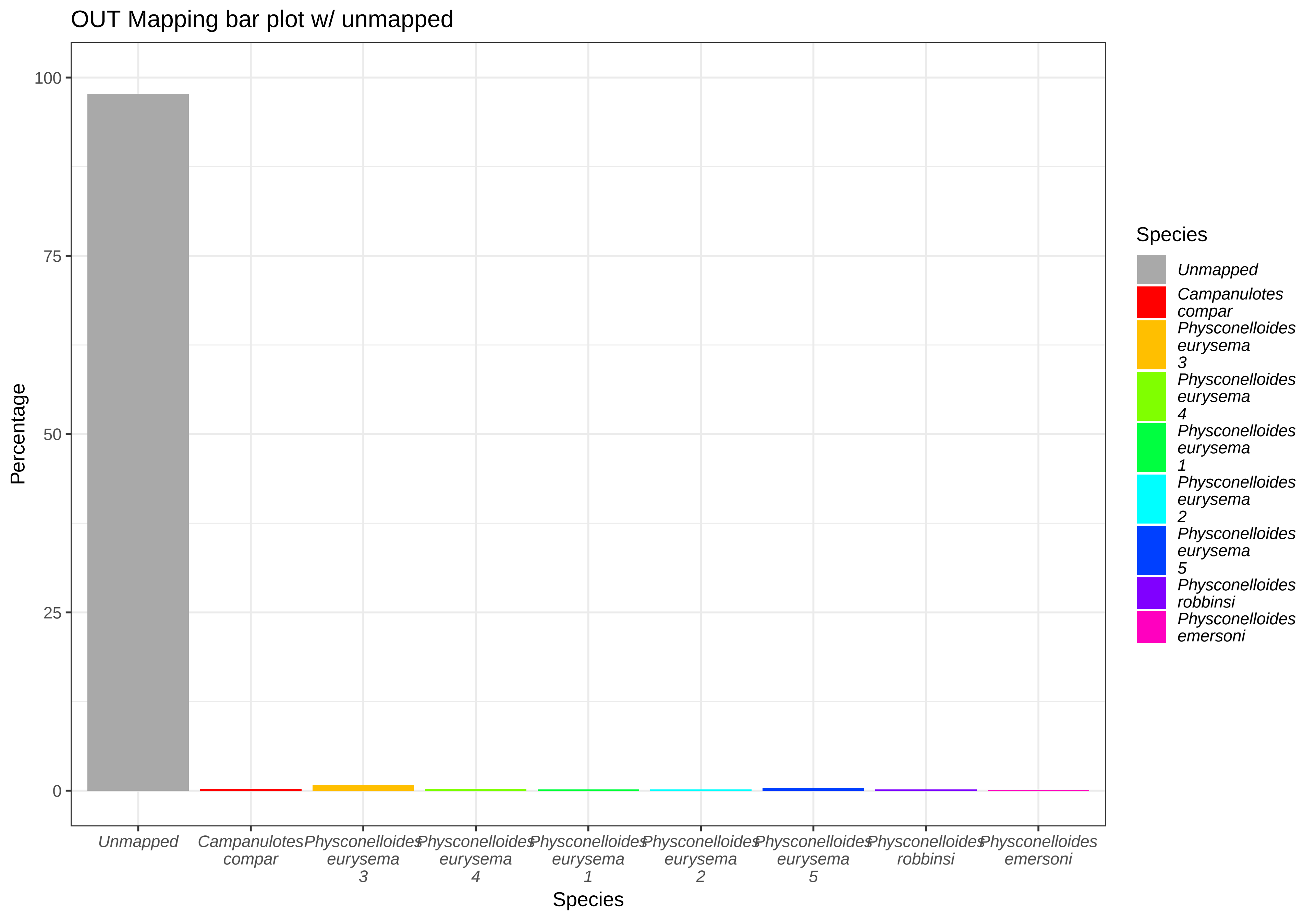

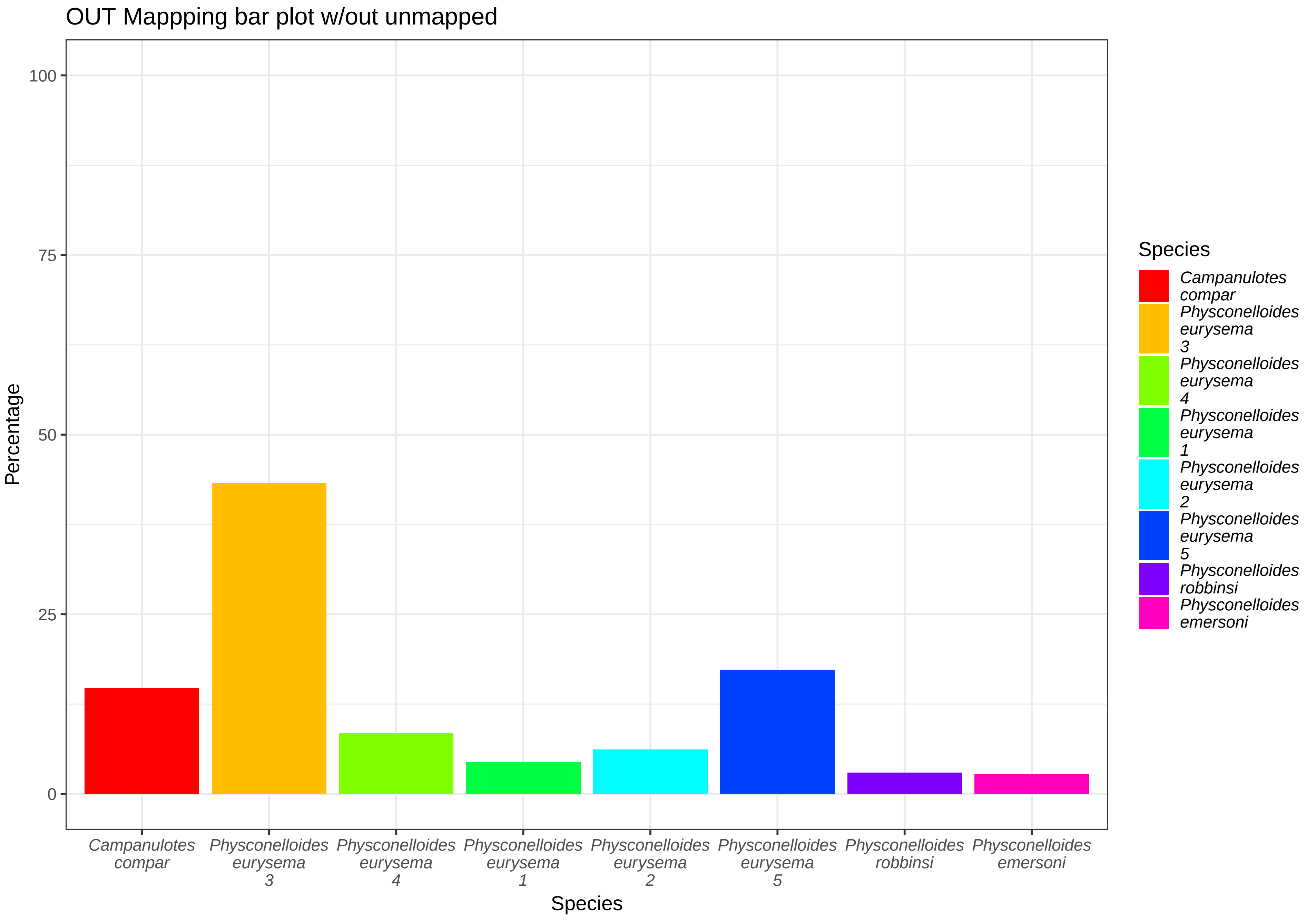

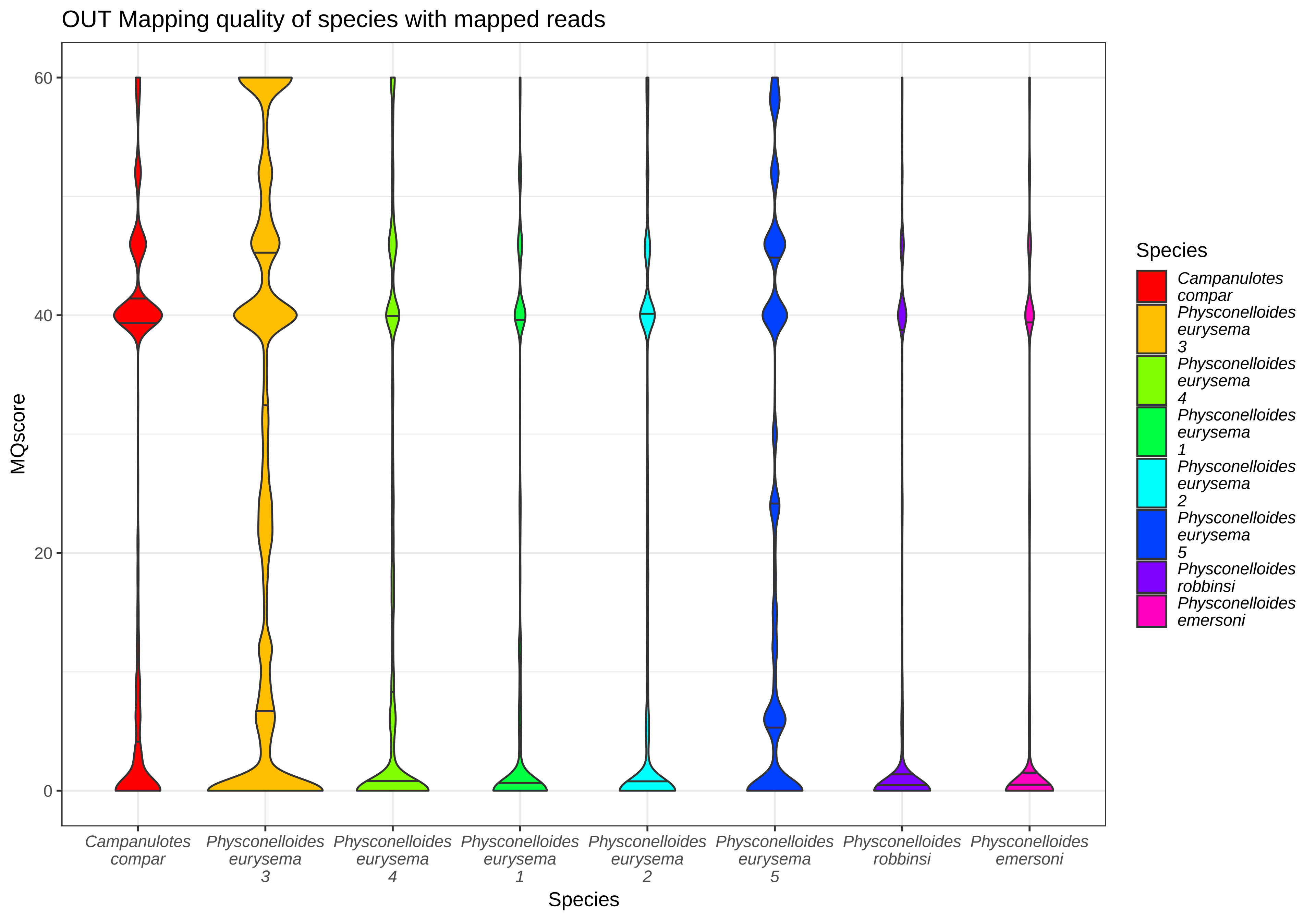

### OUT_plotMQ_Pheur_Copas_3_14_2016_7.pdf

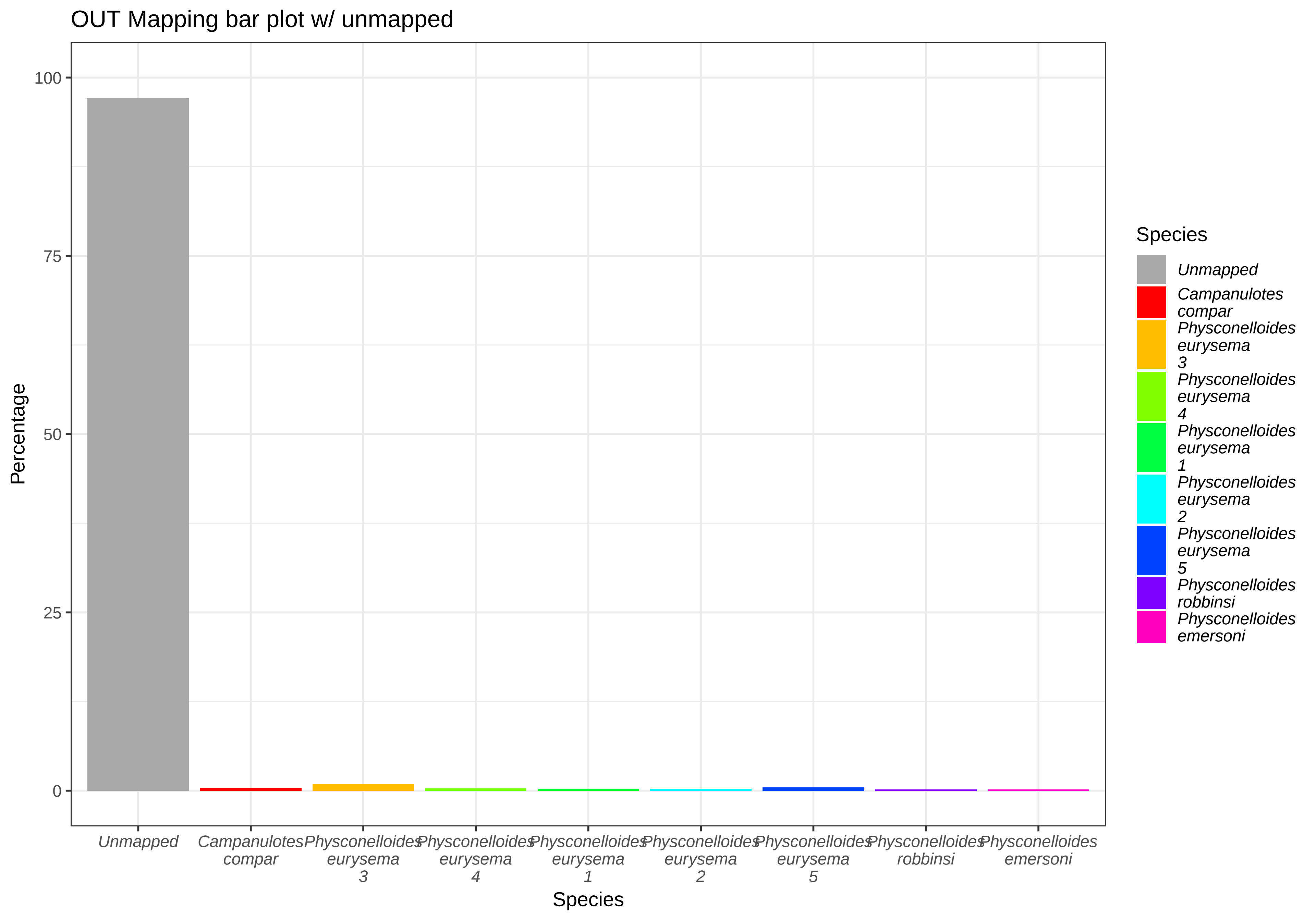

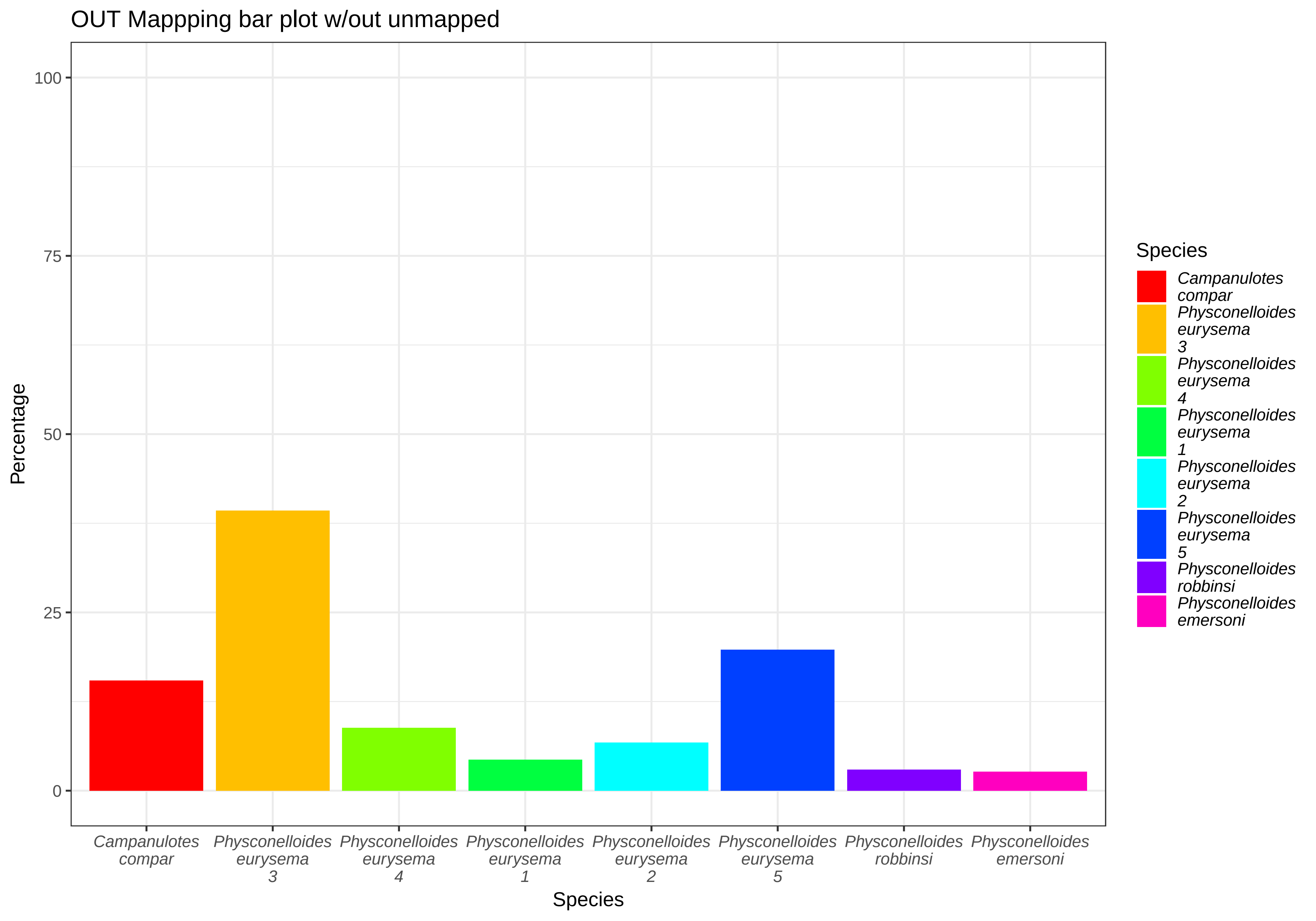

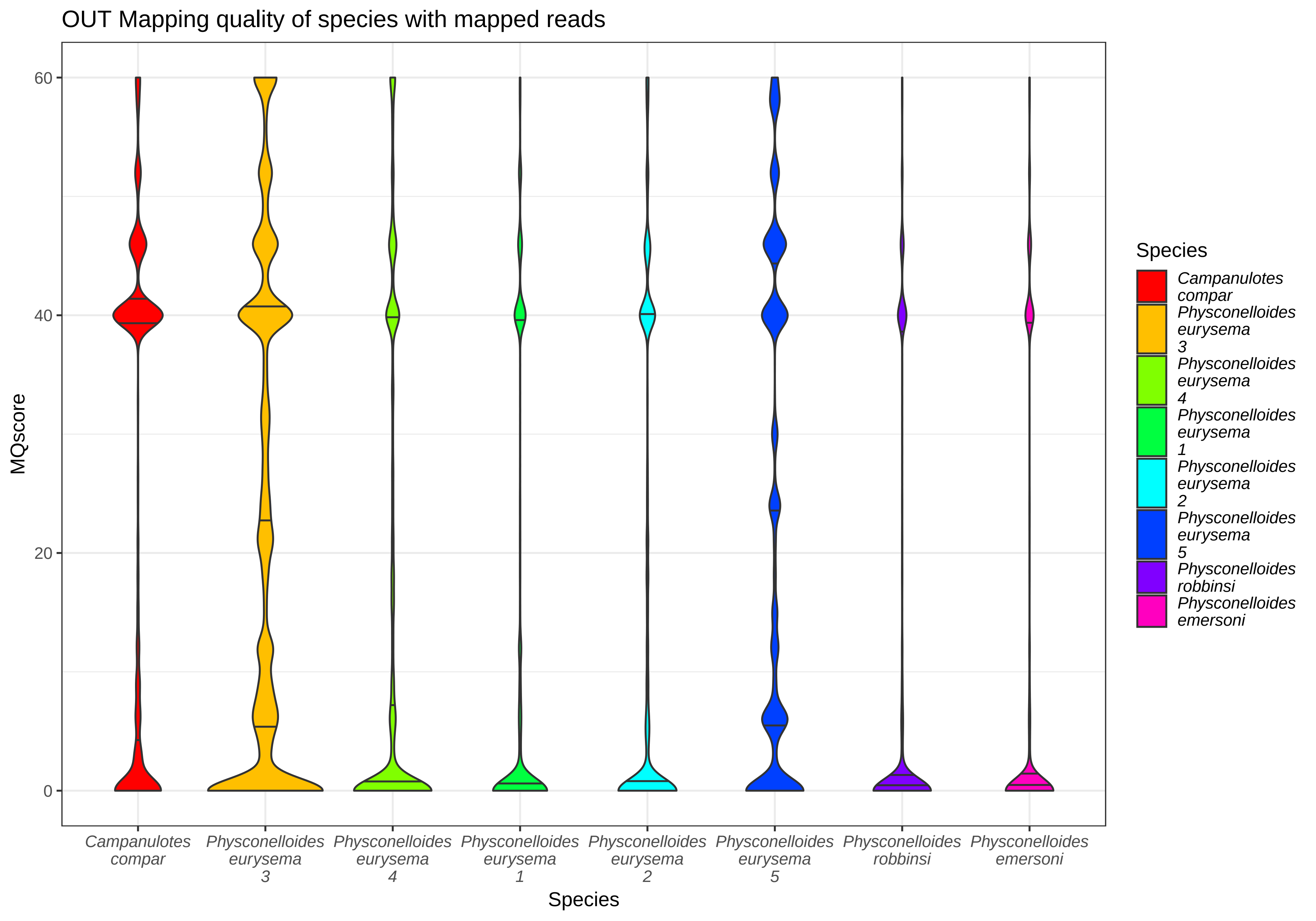

### OUT_plotMQ_Pheur_Copas_3_14_2016_8.pdf

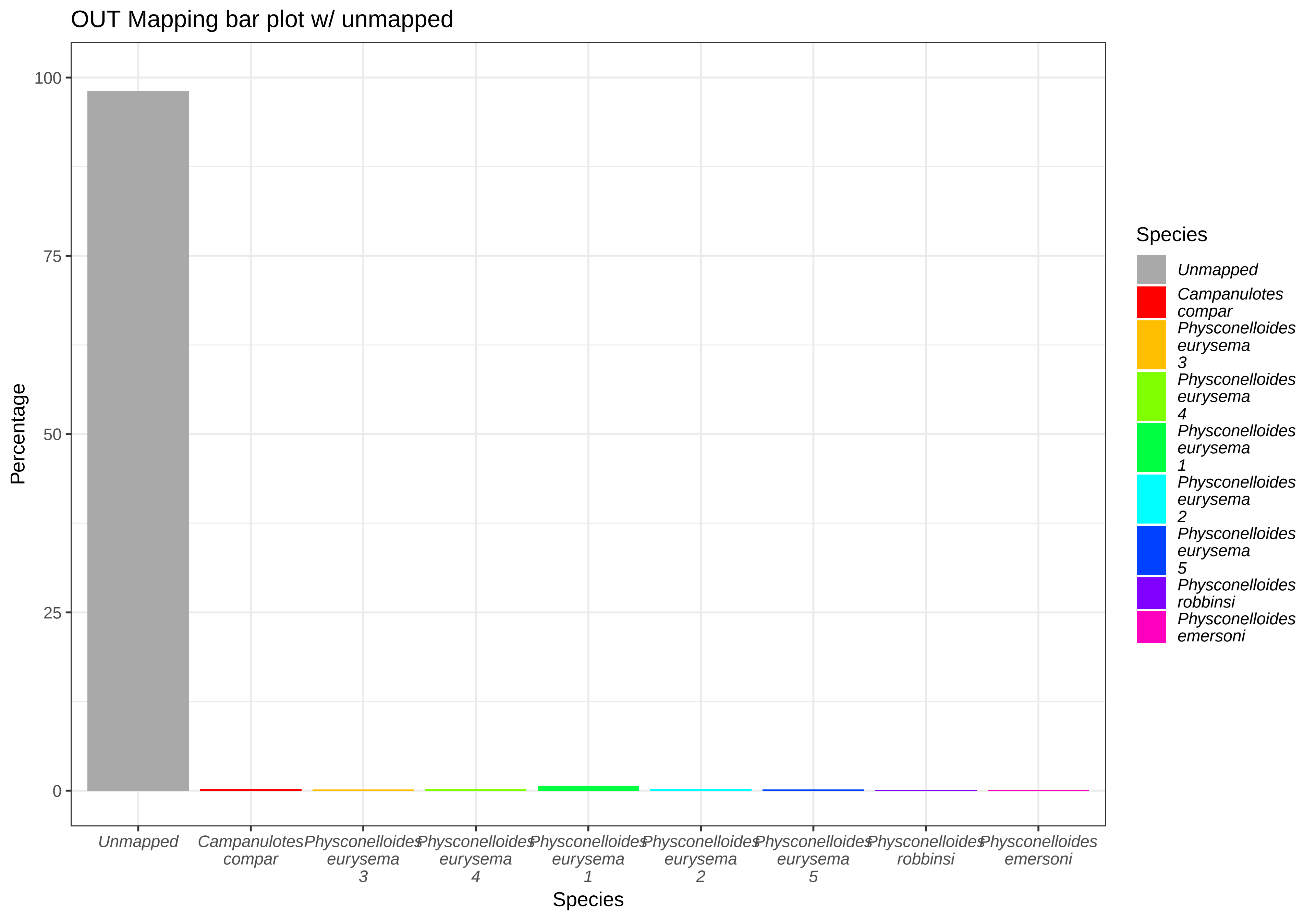

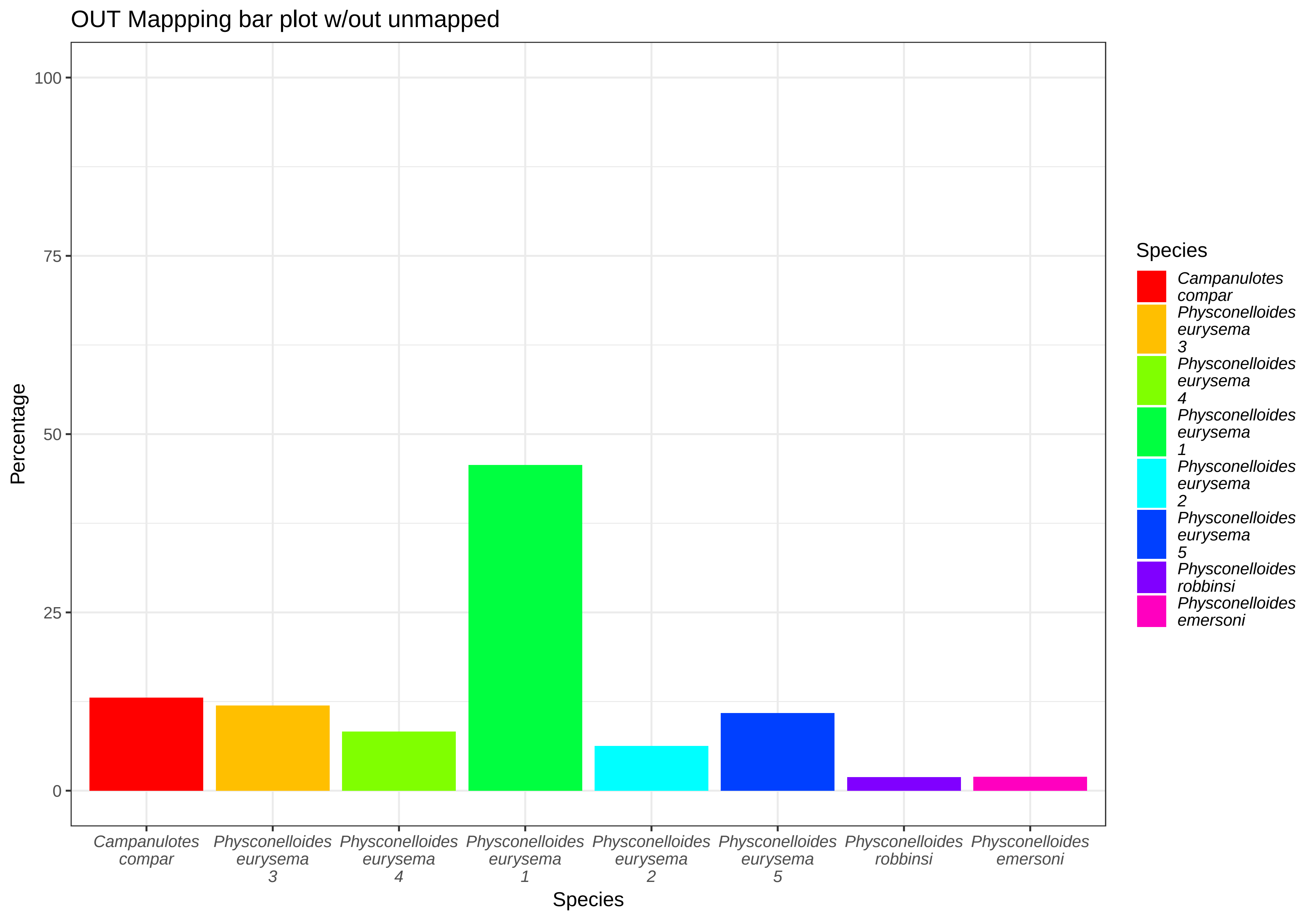

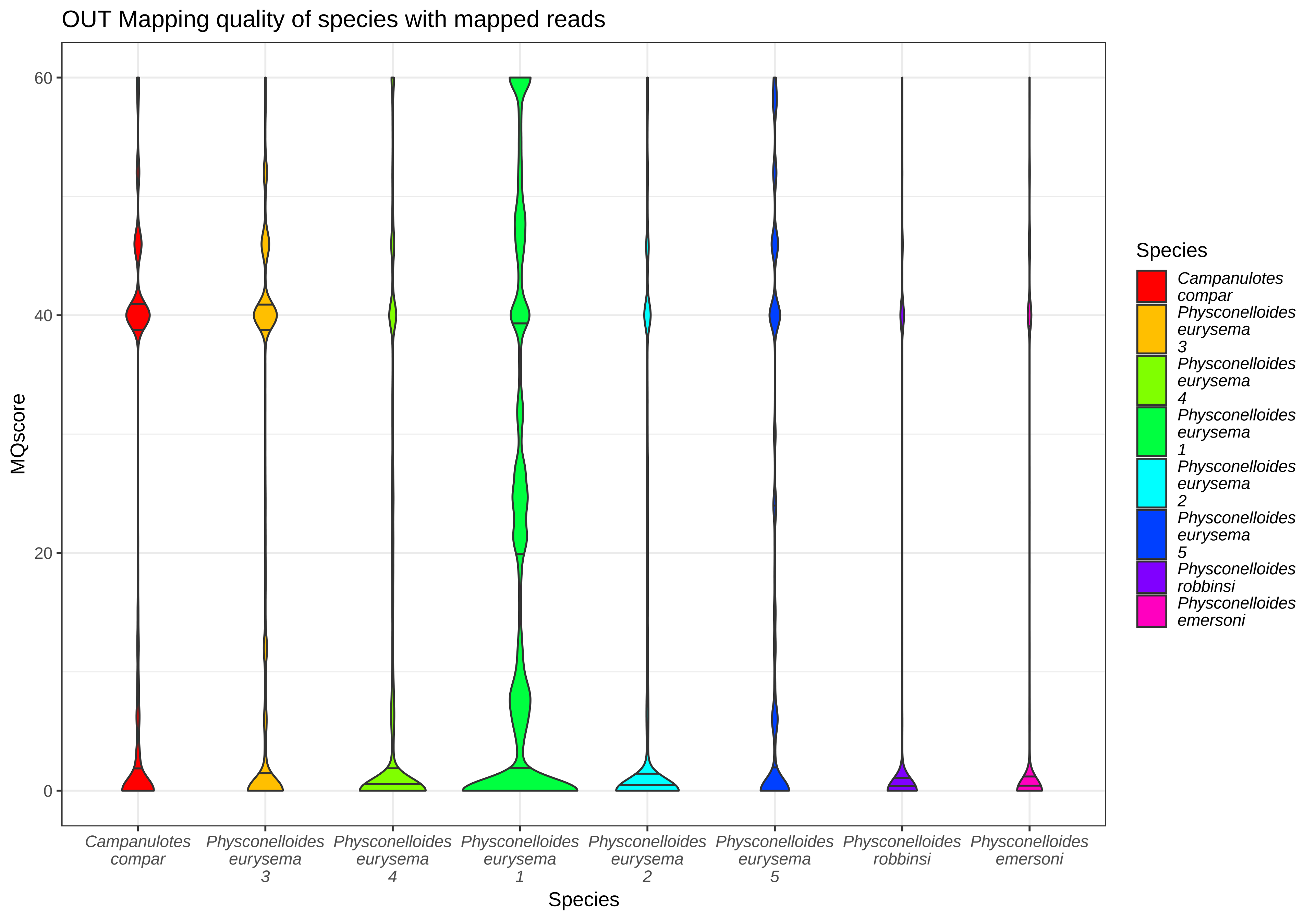

### OUT_plotMQ_Pheur_Copas_3_14_2016_10.pdf

##### OUT Mapping bar plot w/ unmapped

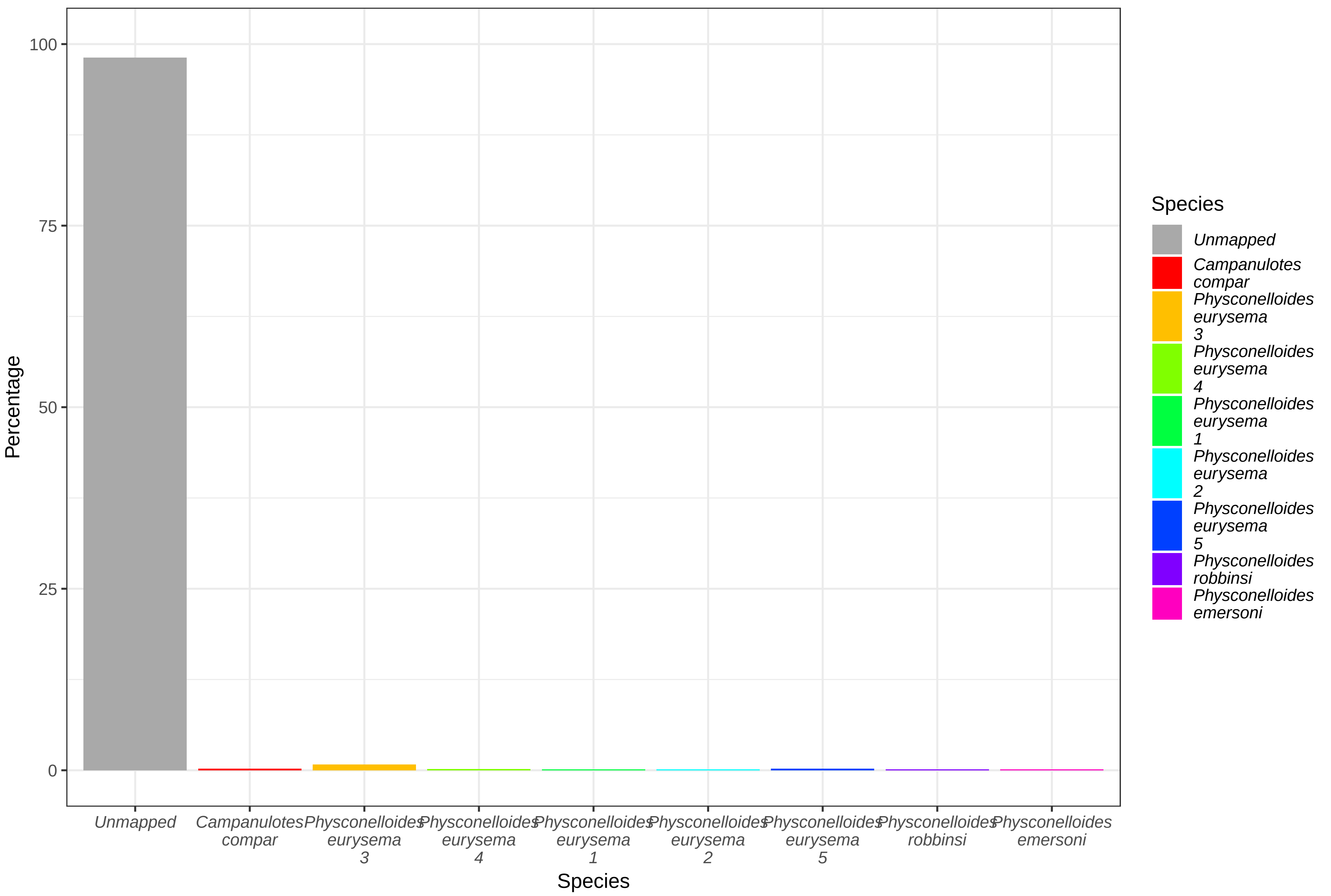

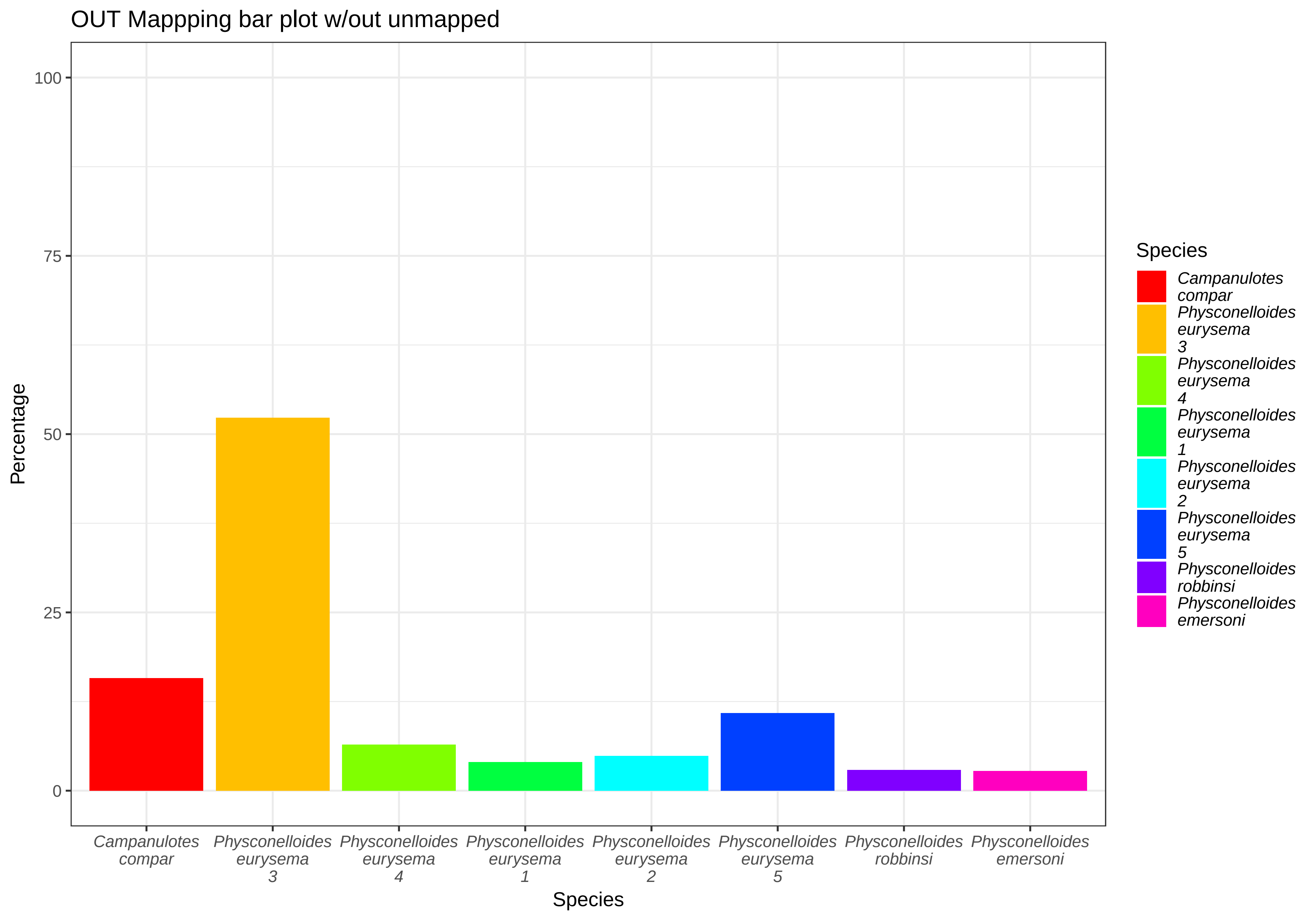

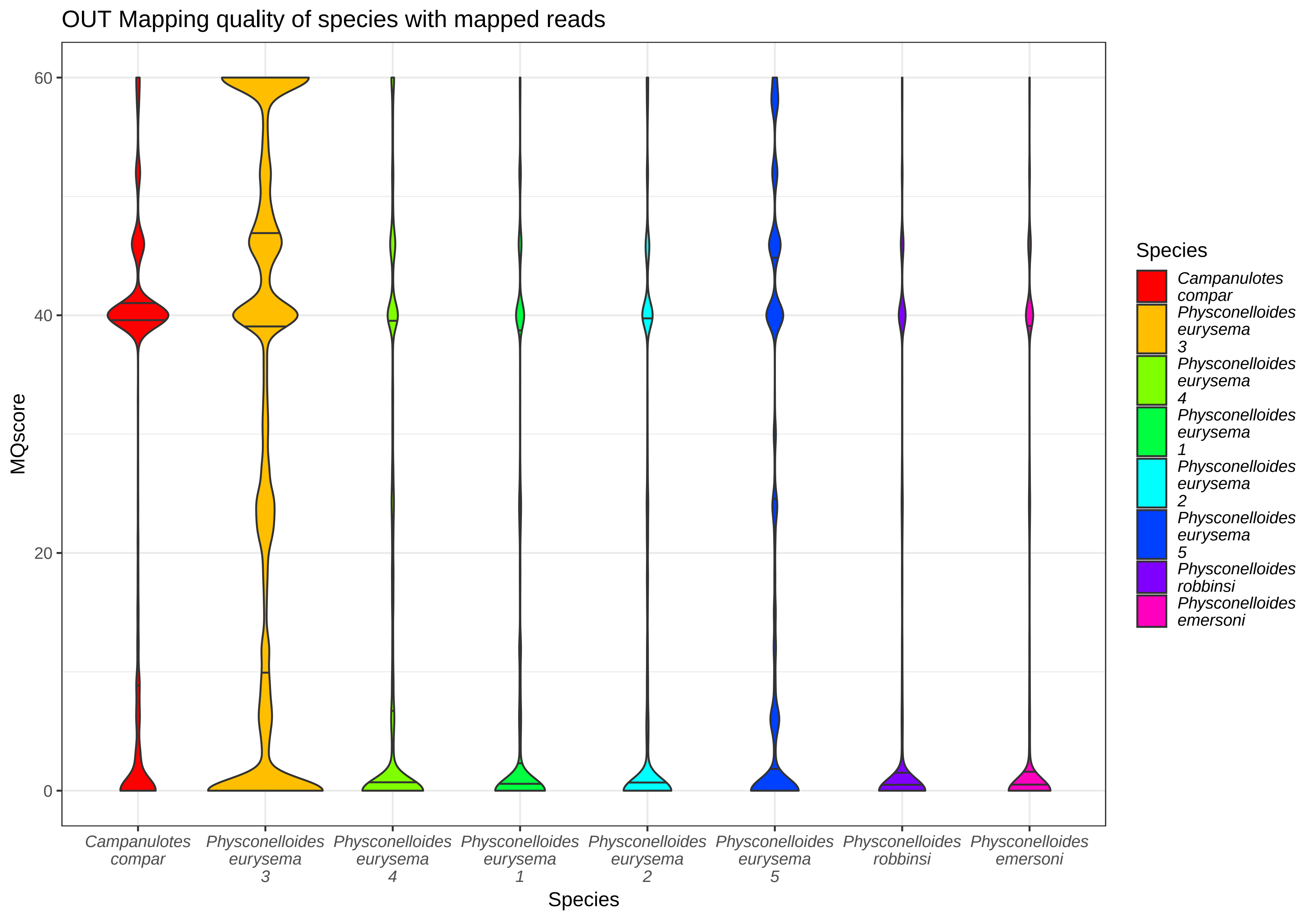

### OUT_plotMQ_Pheur_Copas_3_16_2016_1.pdf

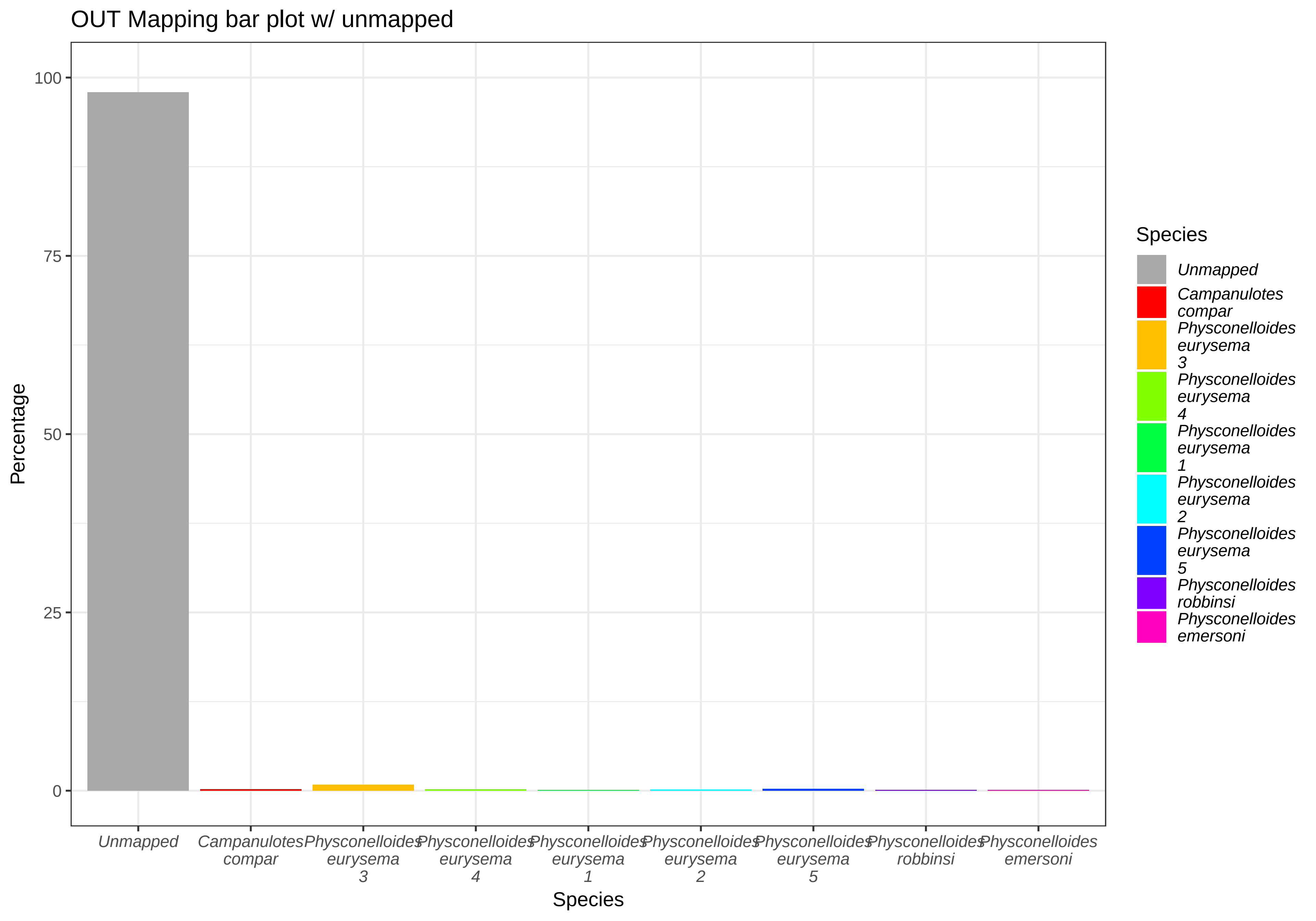

OUT Mappping bar plot w/out unmapped

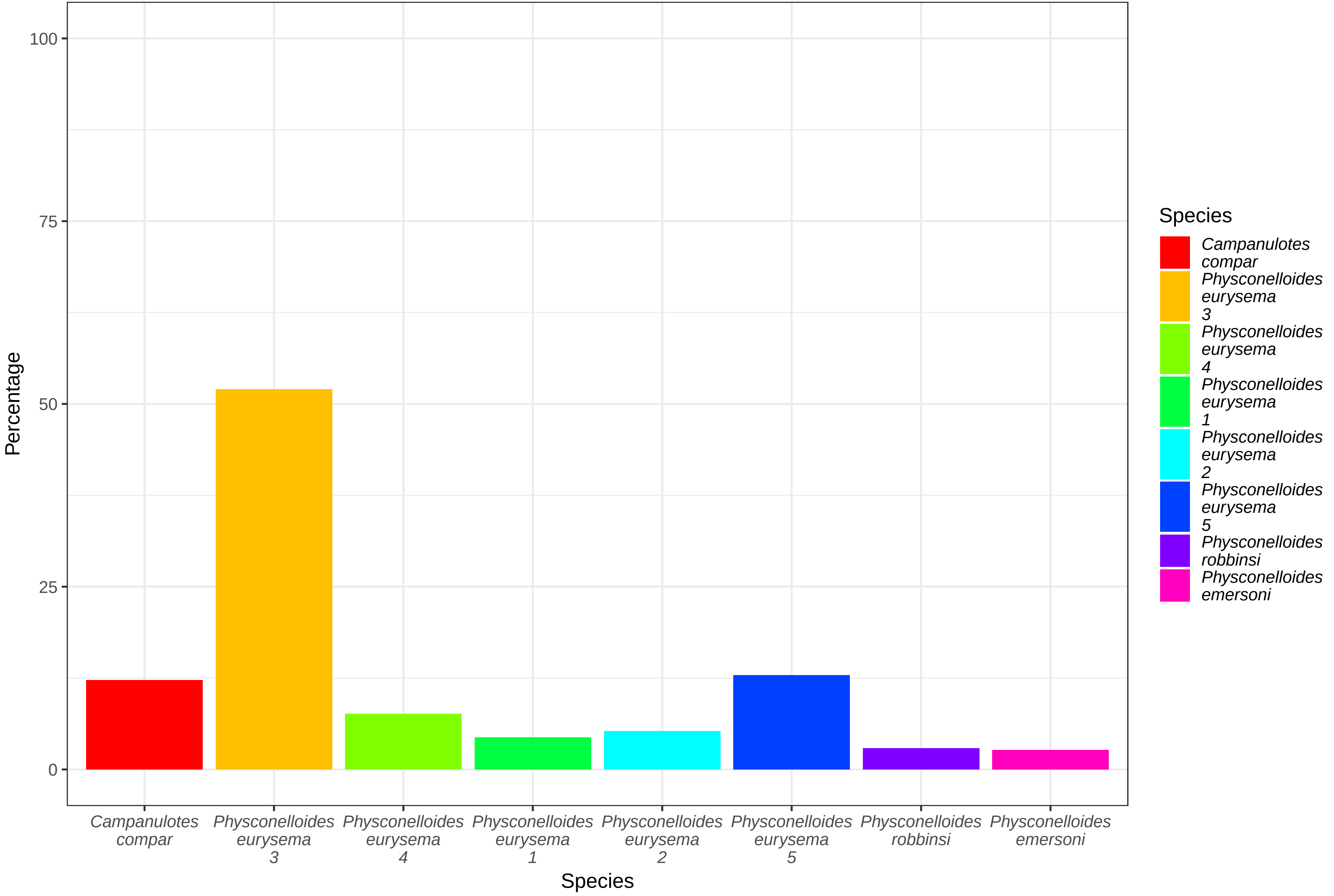

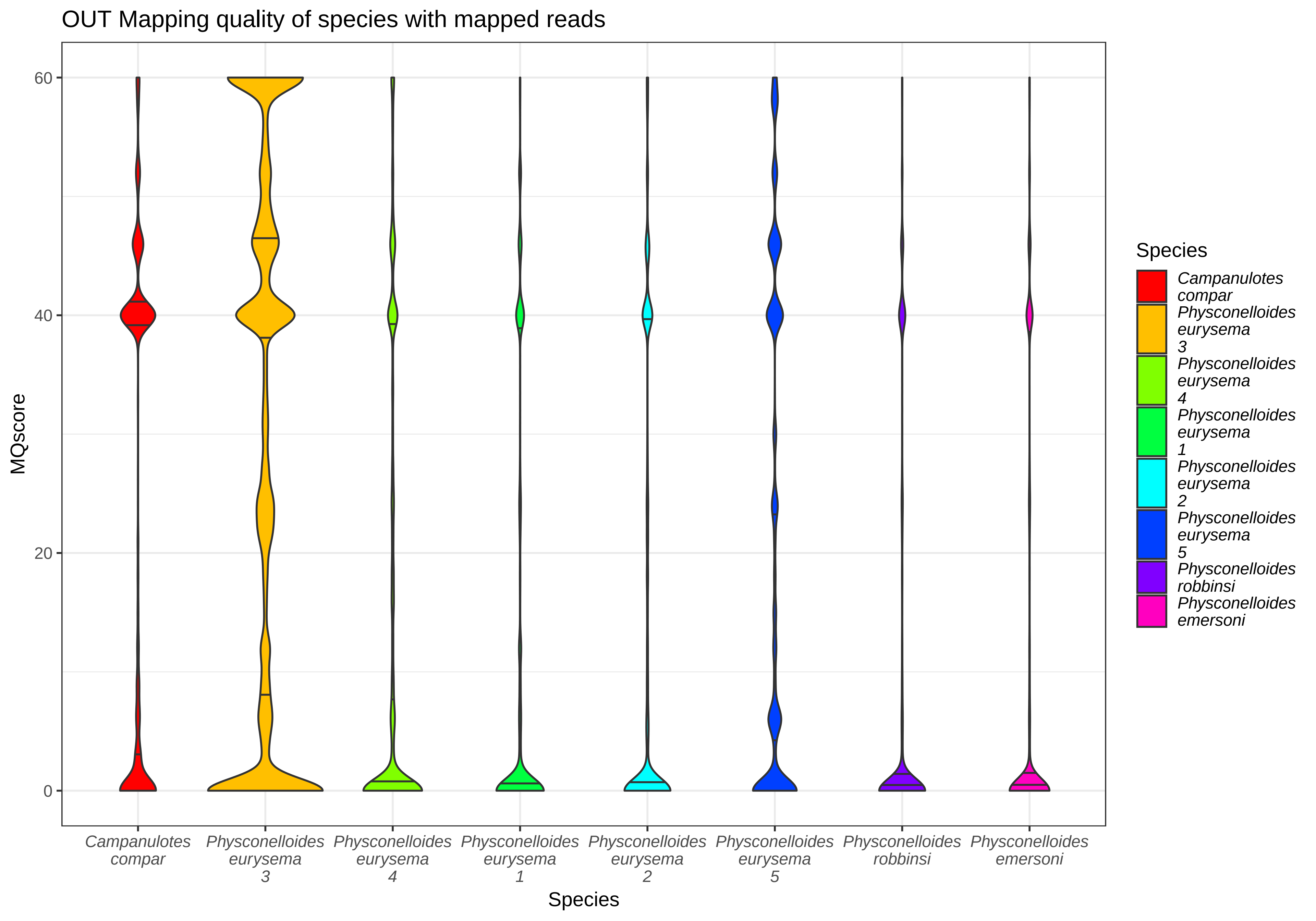

### OUT_plotMQ_Pheur_Copas_3_16_2016_6.pdf

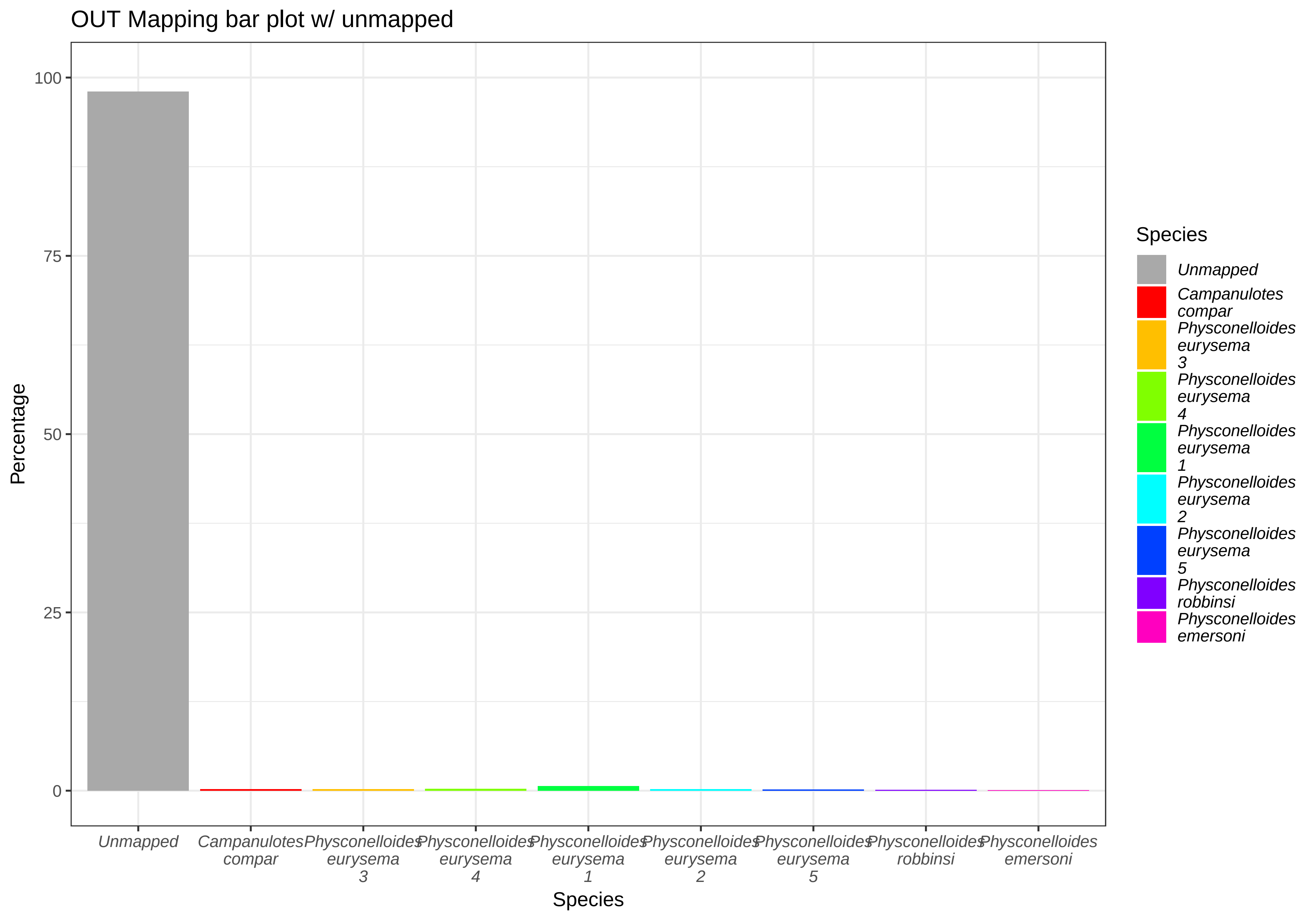

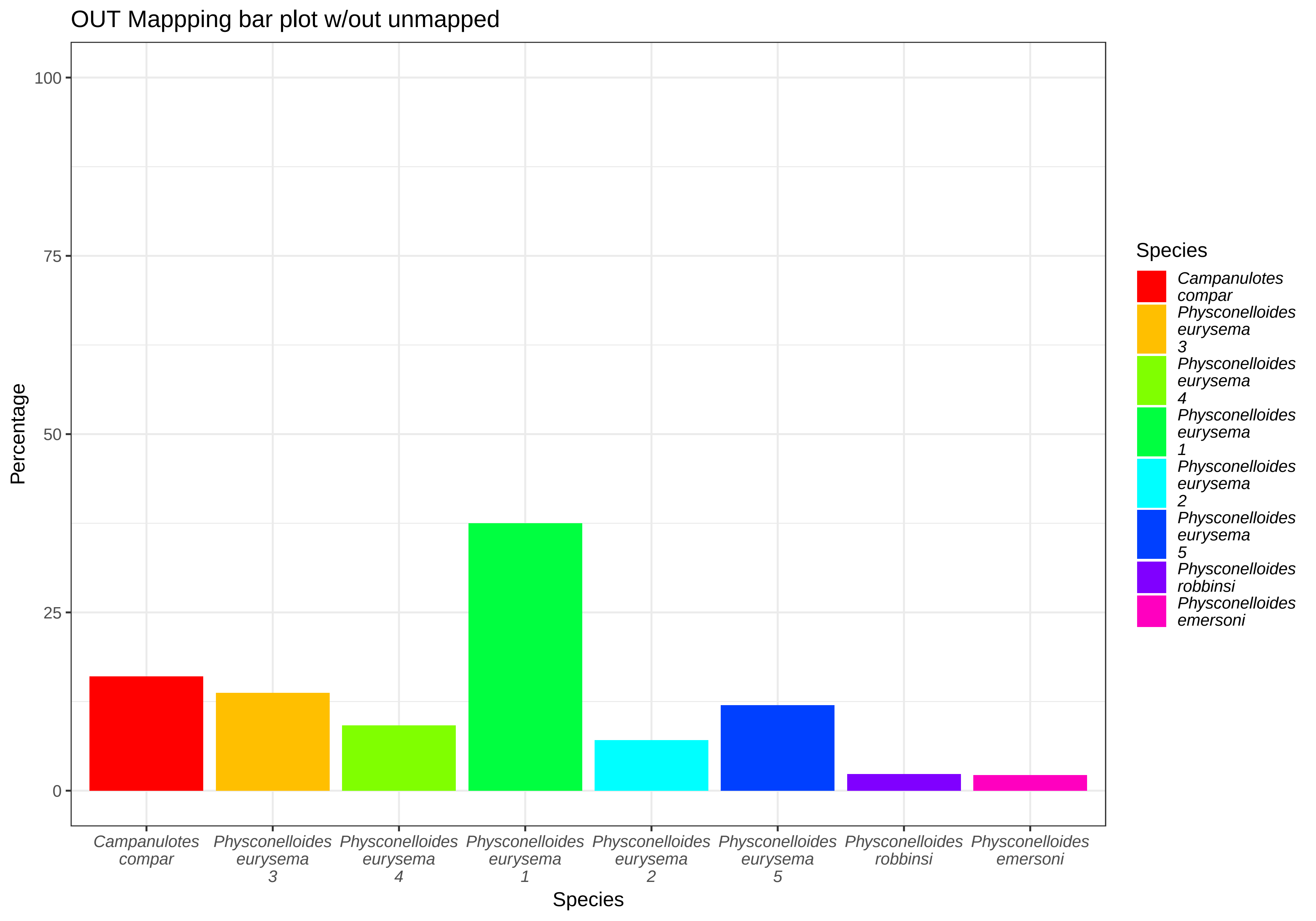

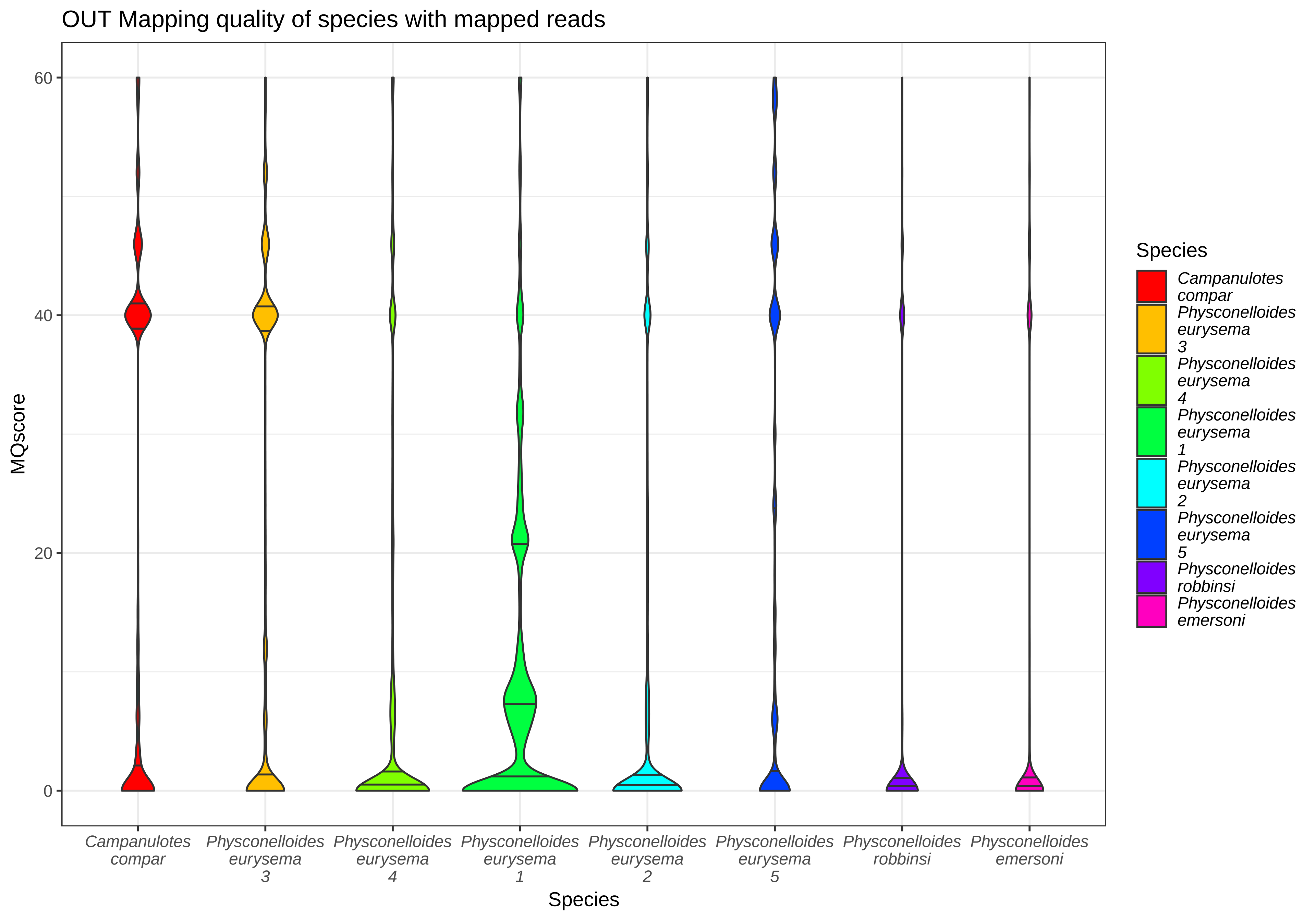

### OUT_plotMQ_Pheur_Copas_3_16_2016_7.pdf

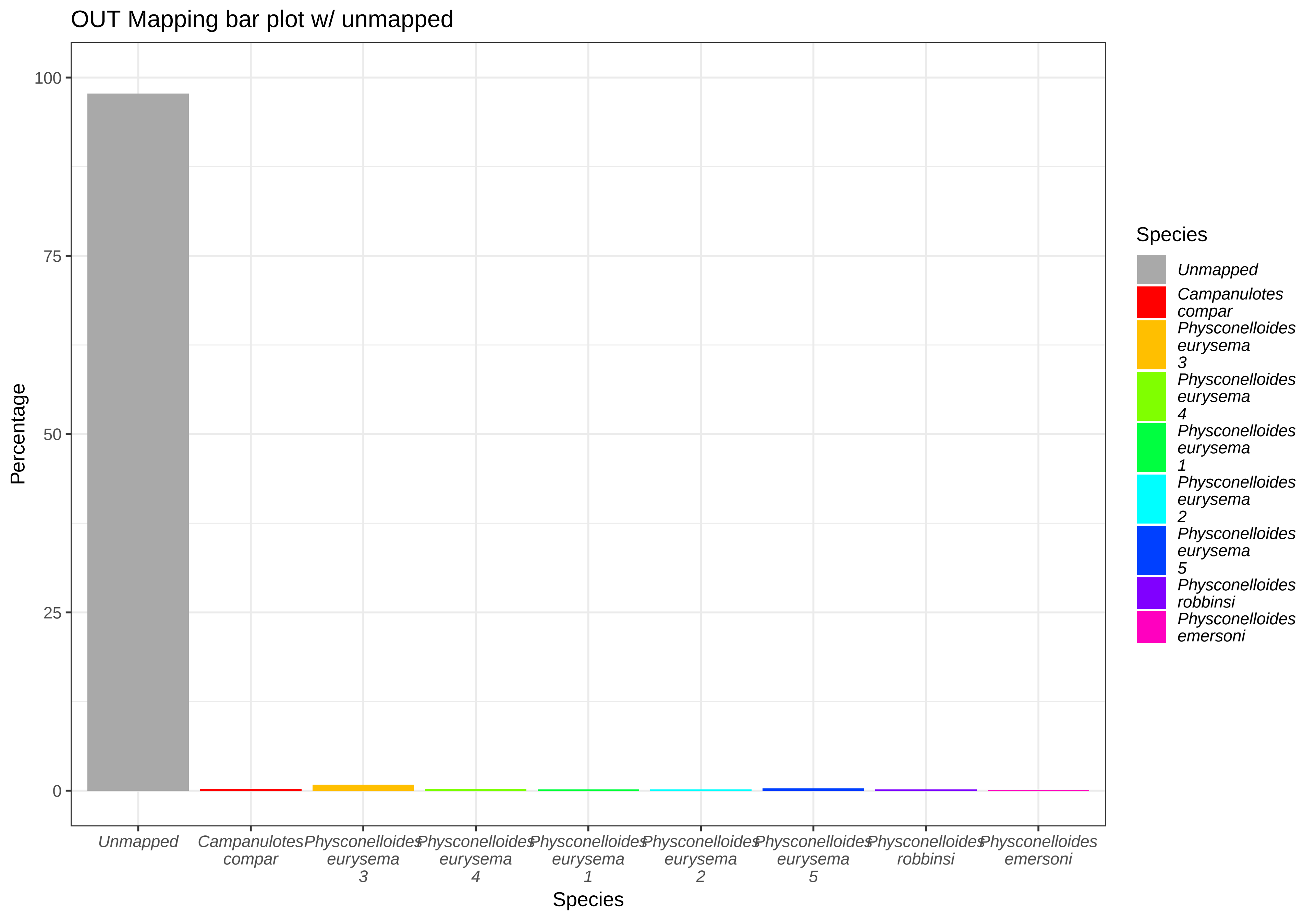

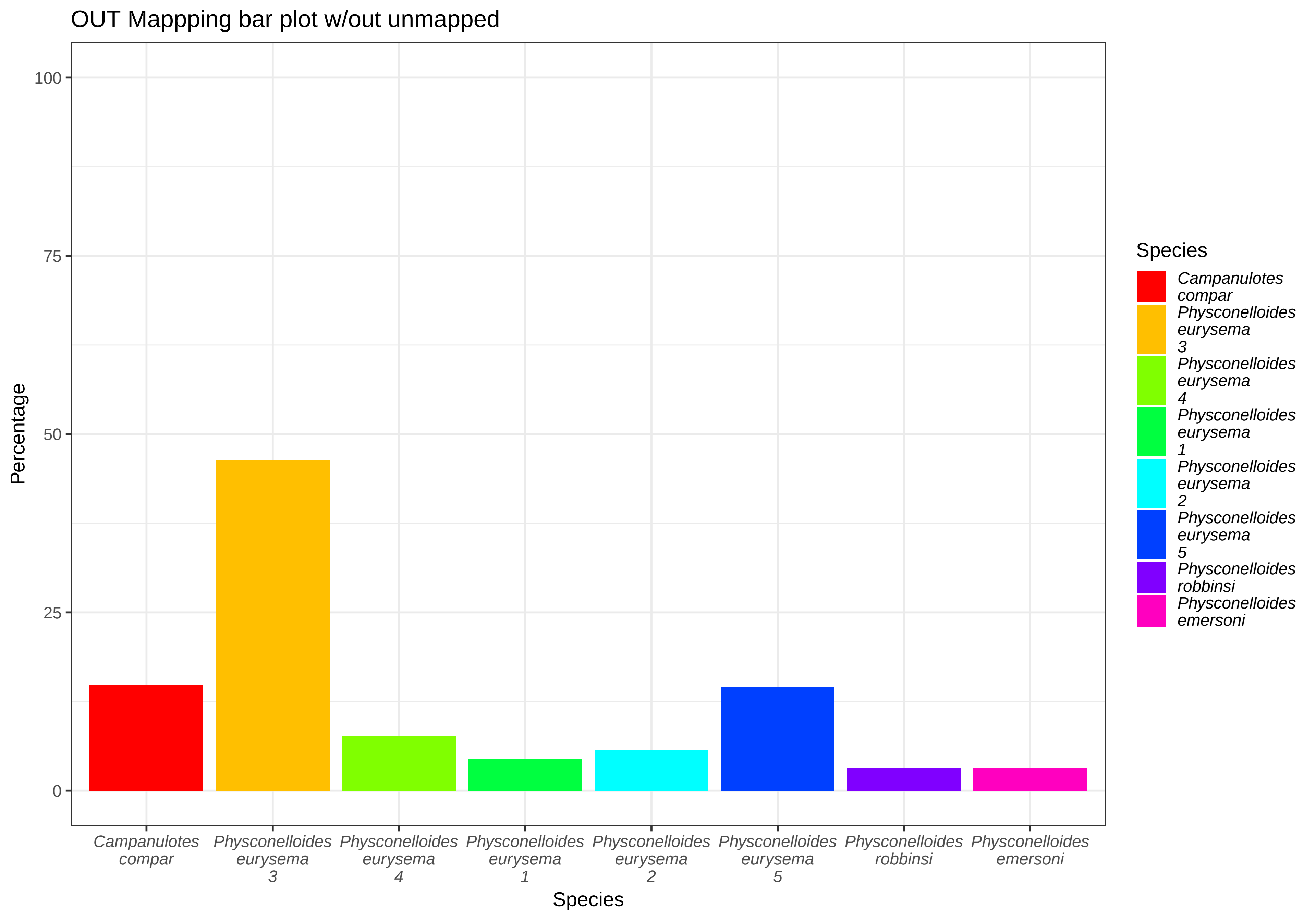

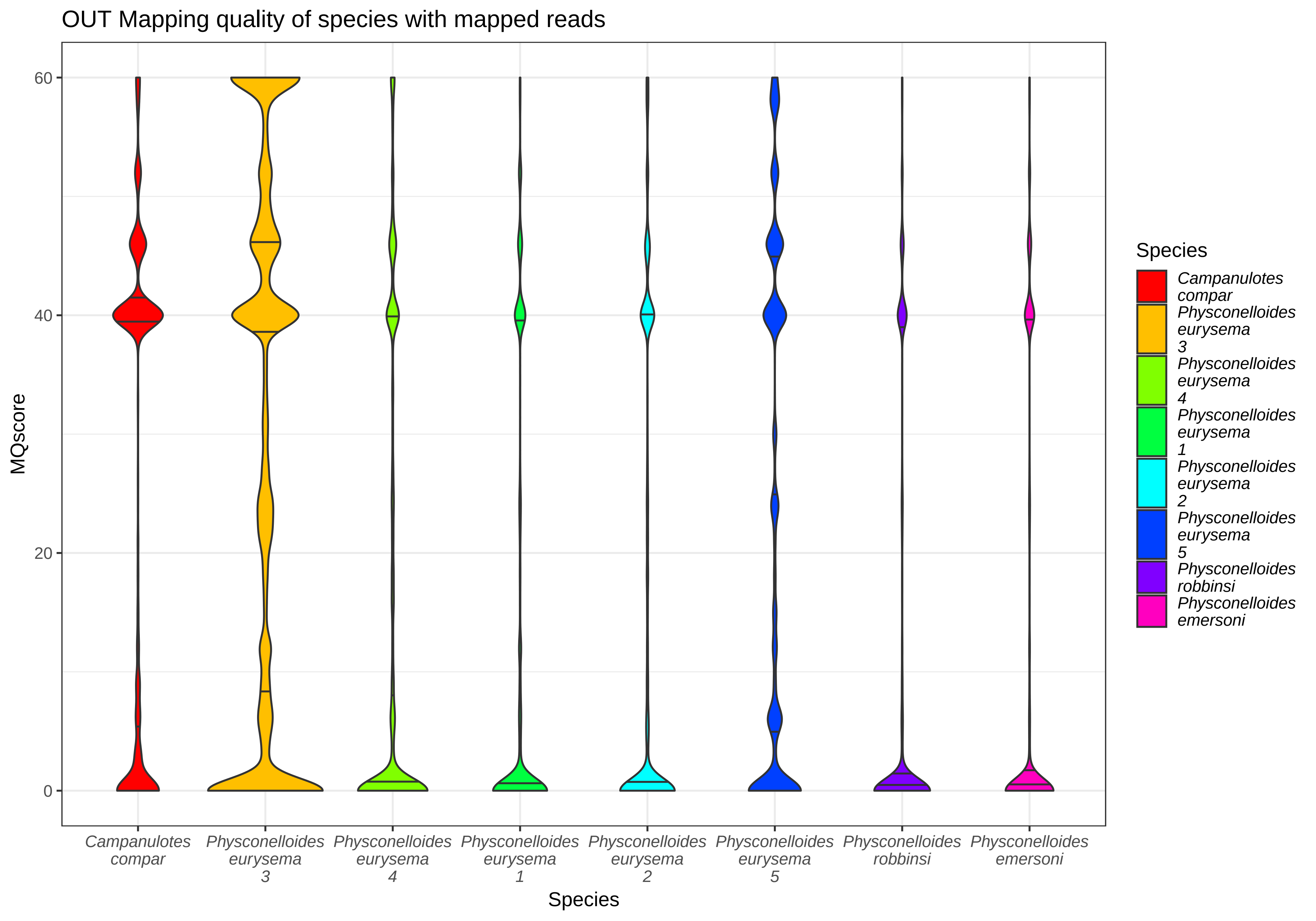

### OUT_plotMQ_Pheur_Cosqu_3_14_2016_1.pdf

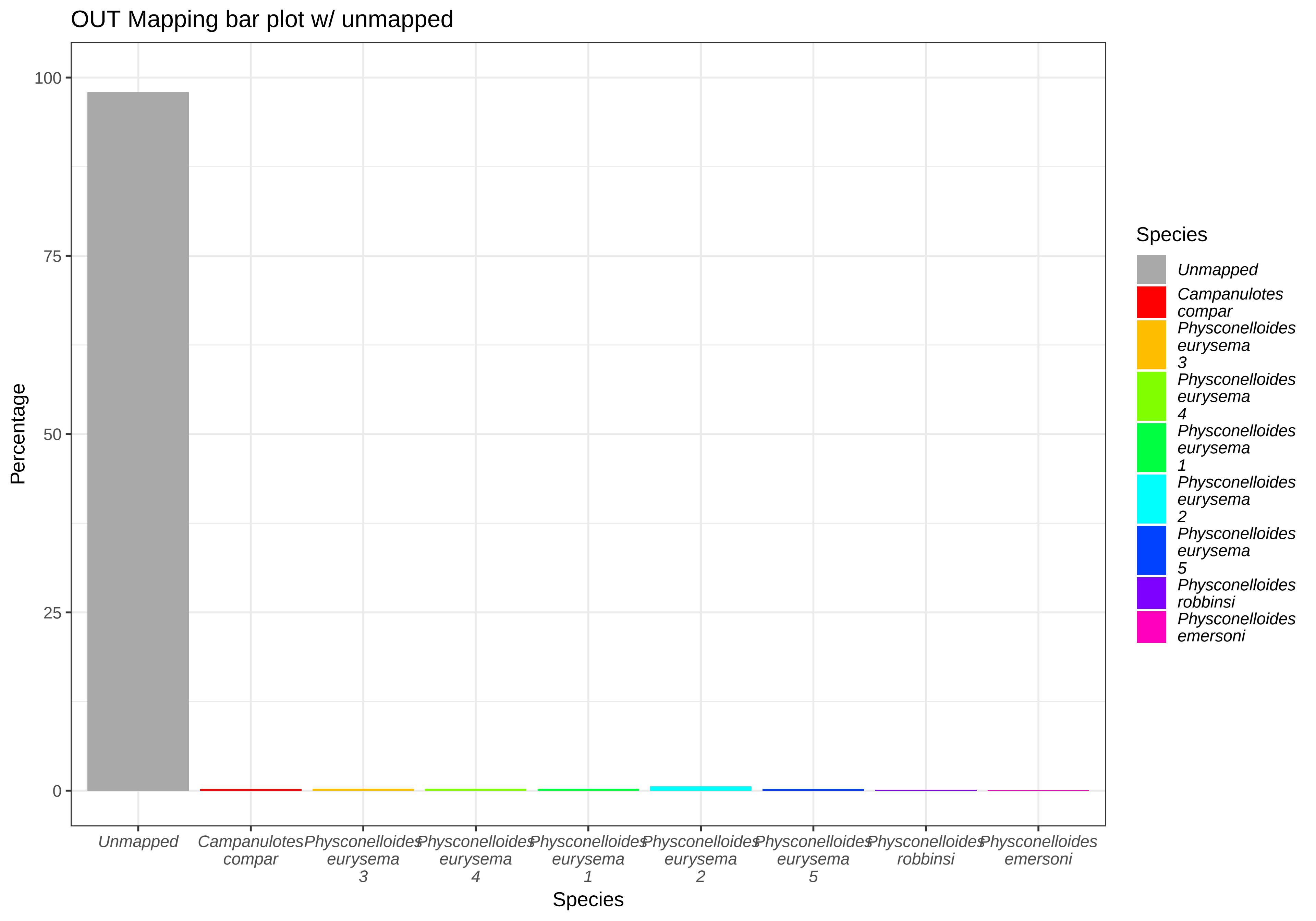

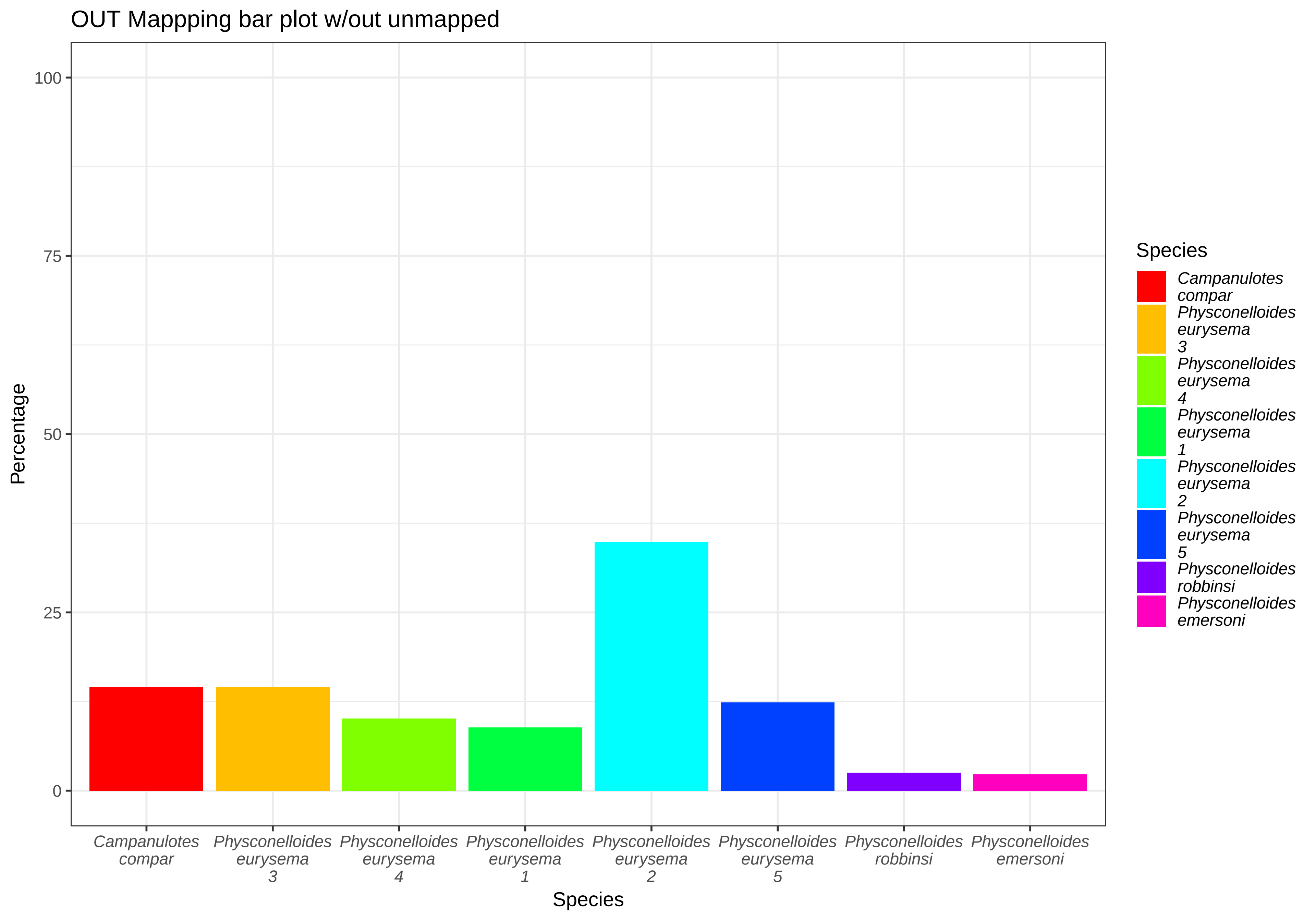

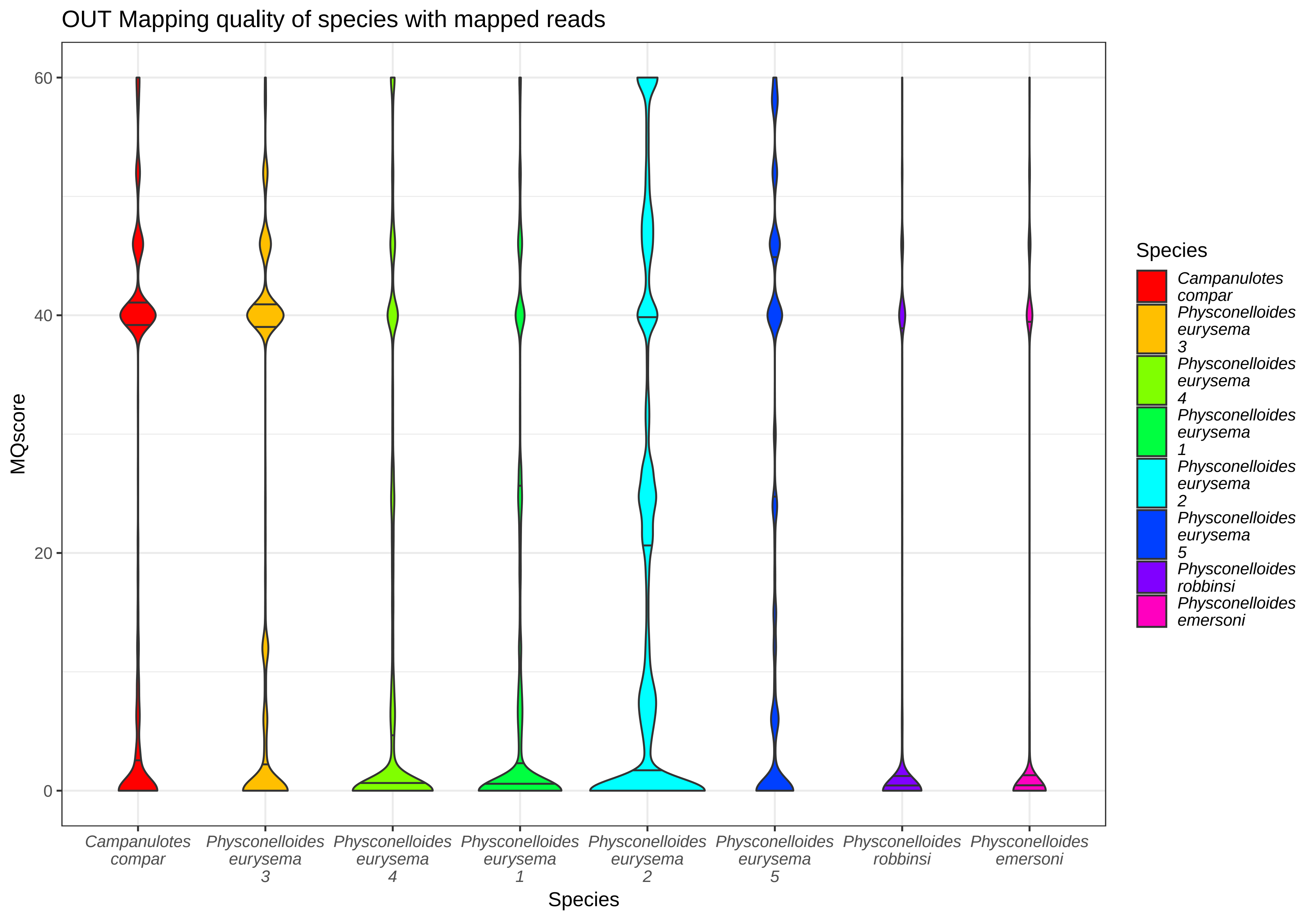

### OUT_plotMQ_Pheur_Cosqu_3_14_2016_12.pdf

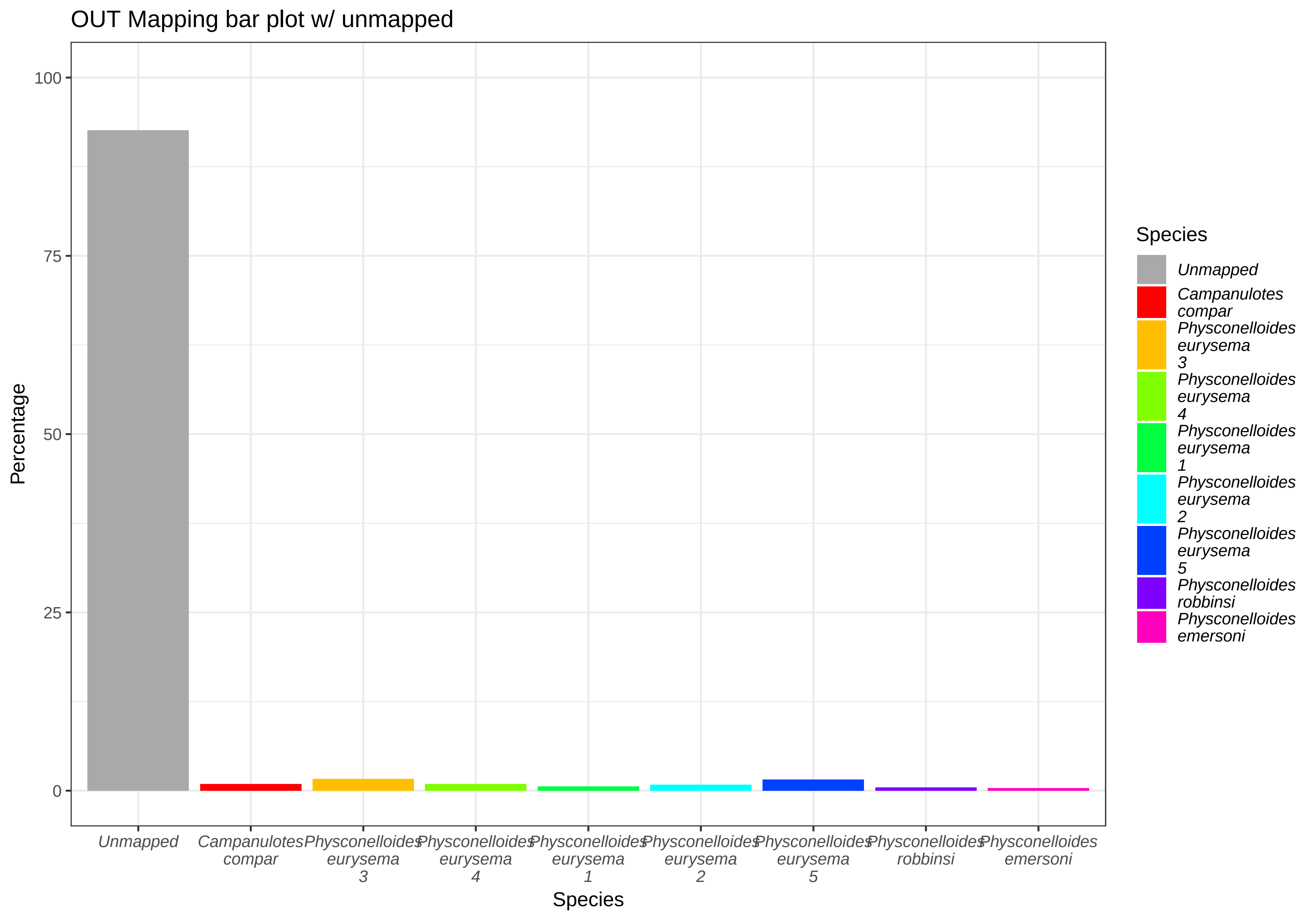

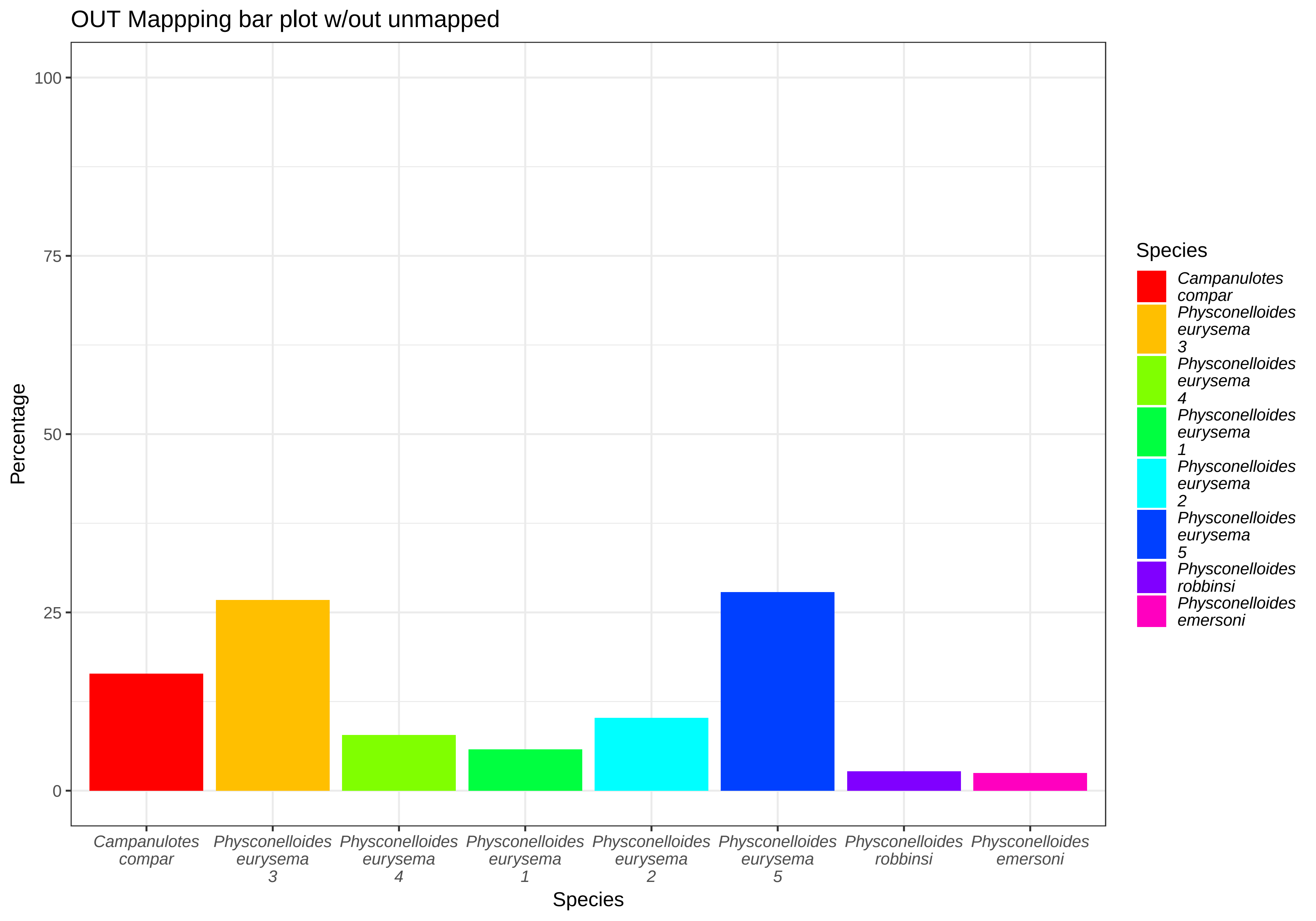

OUT Mapping quality of species with mapped reads

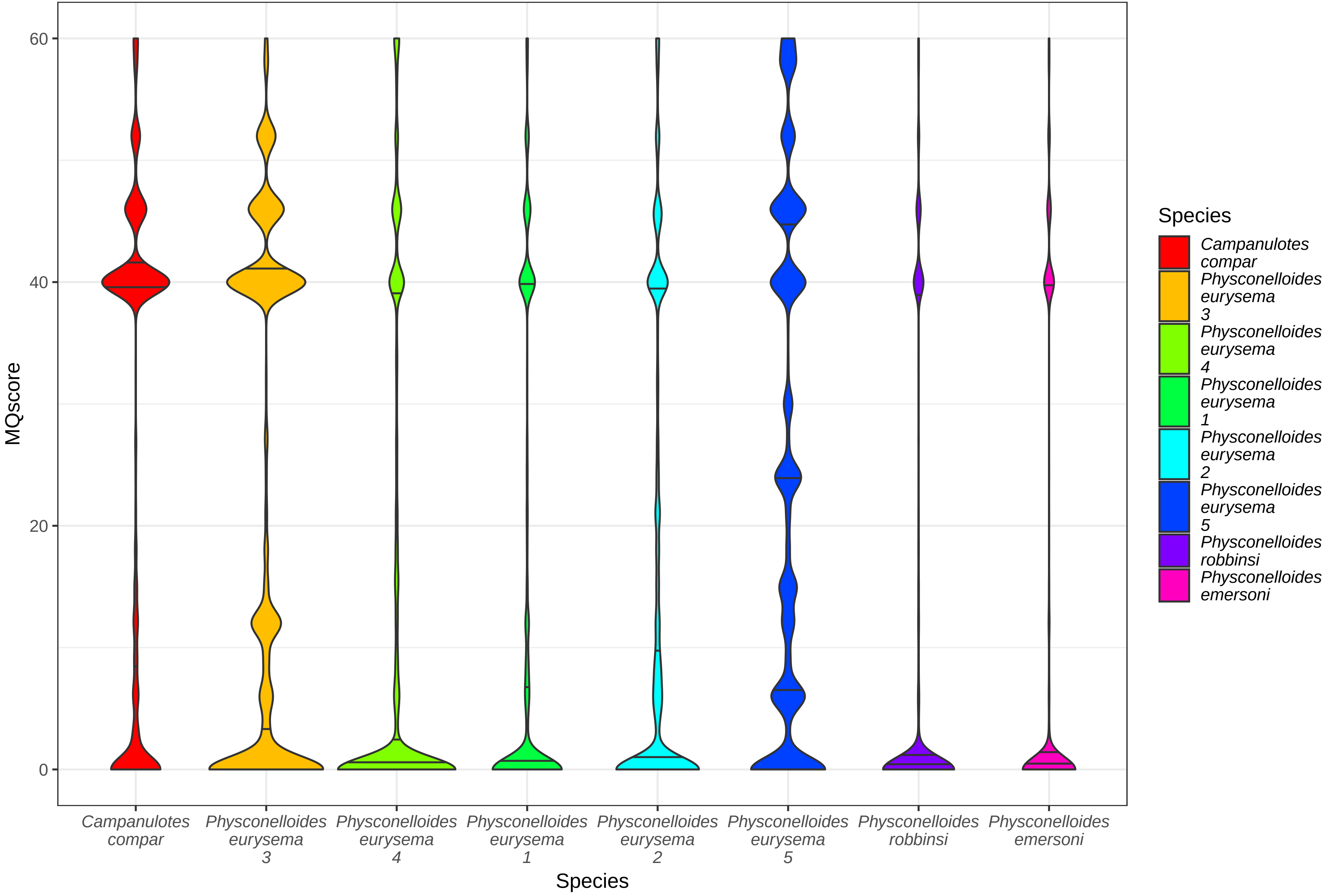

### OUT_plotMQ_Pheur_Cotal_3_14_2016_2.pdf

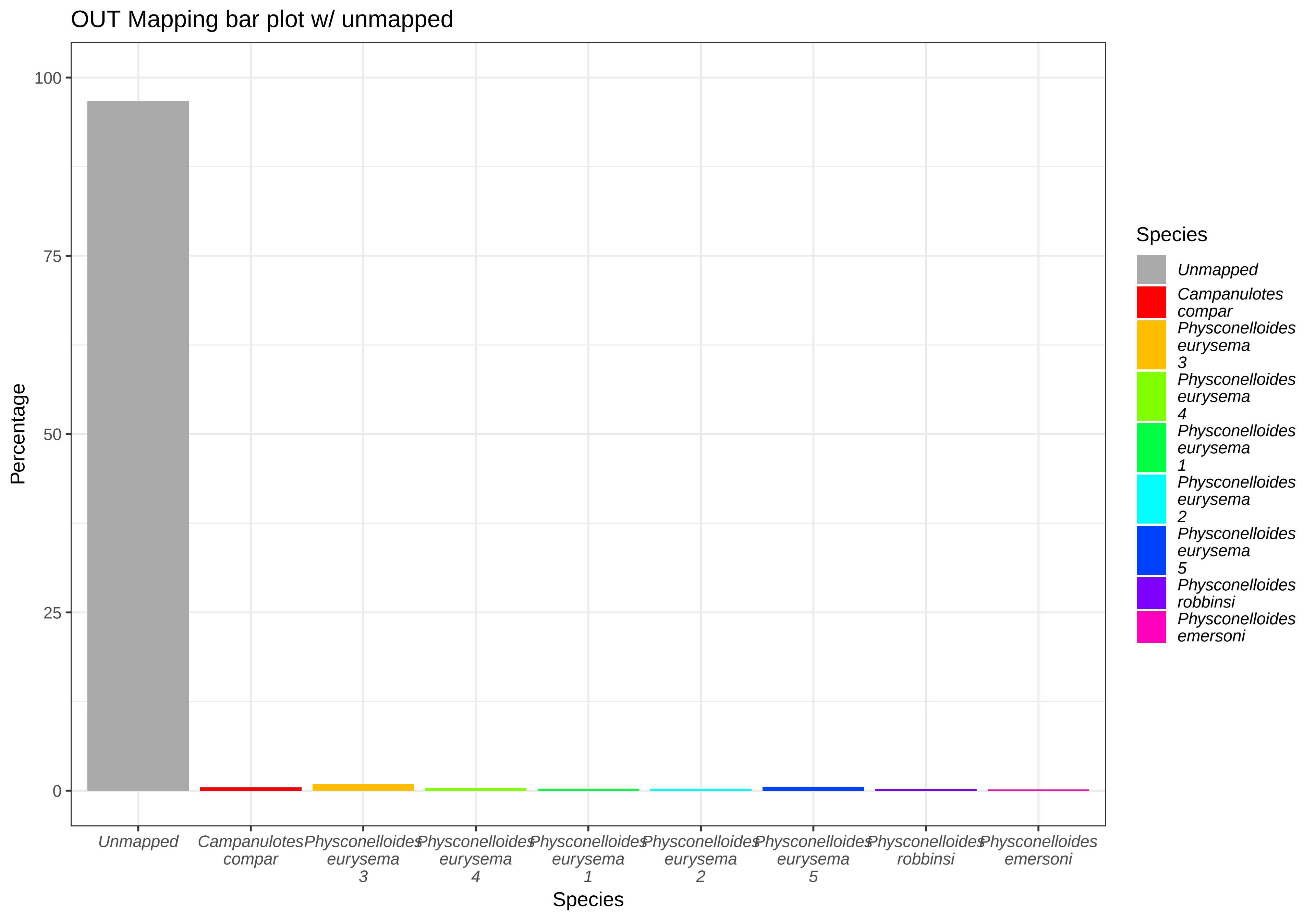

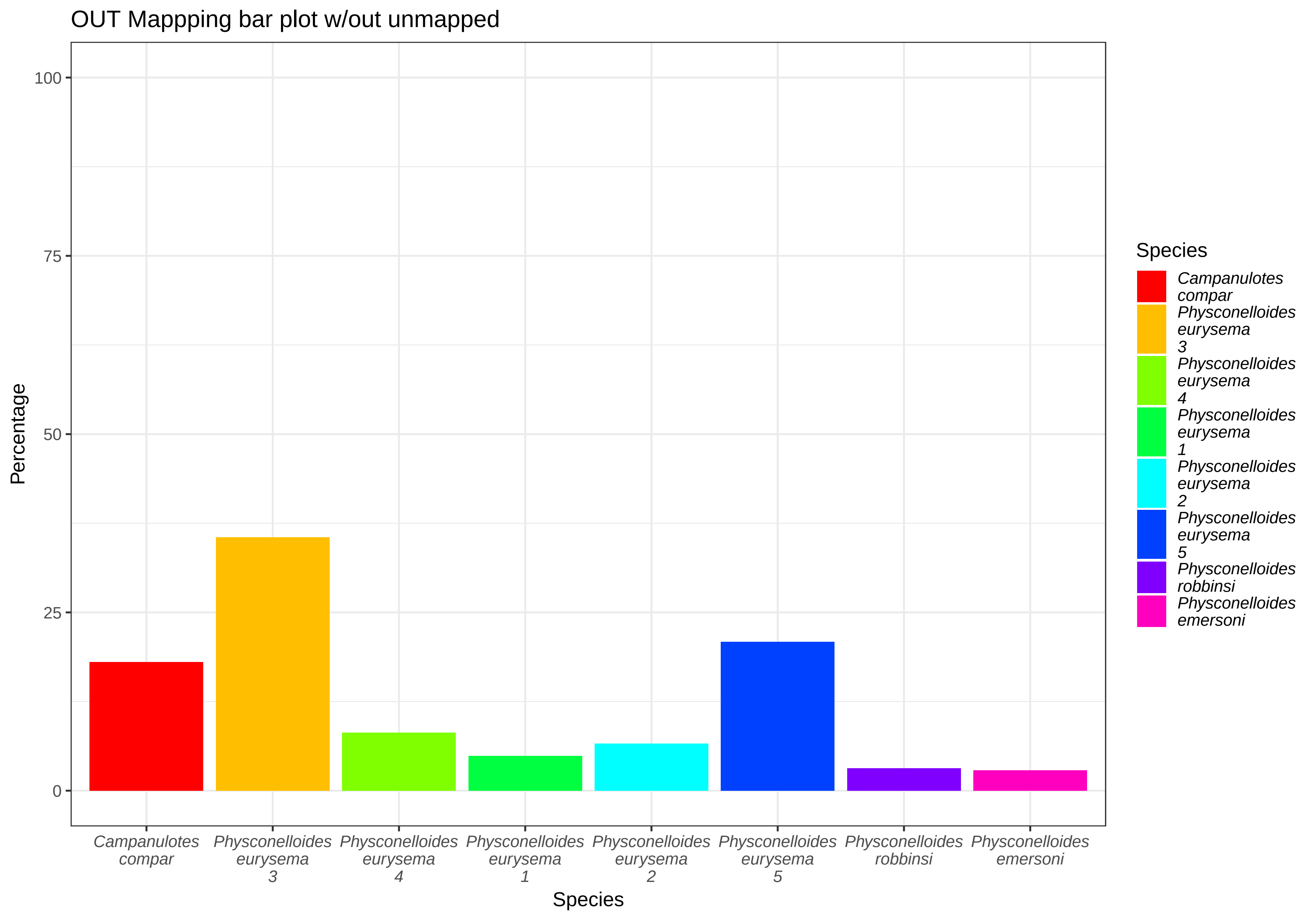

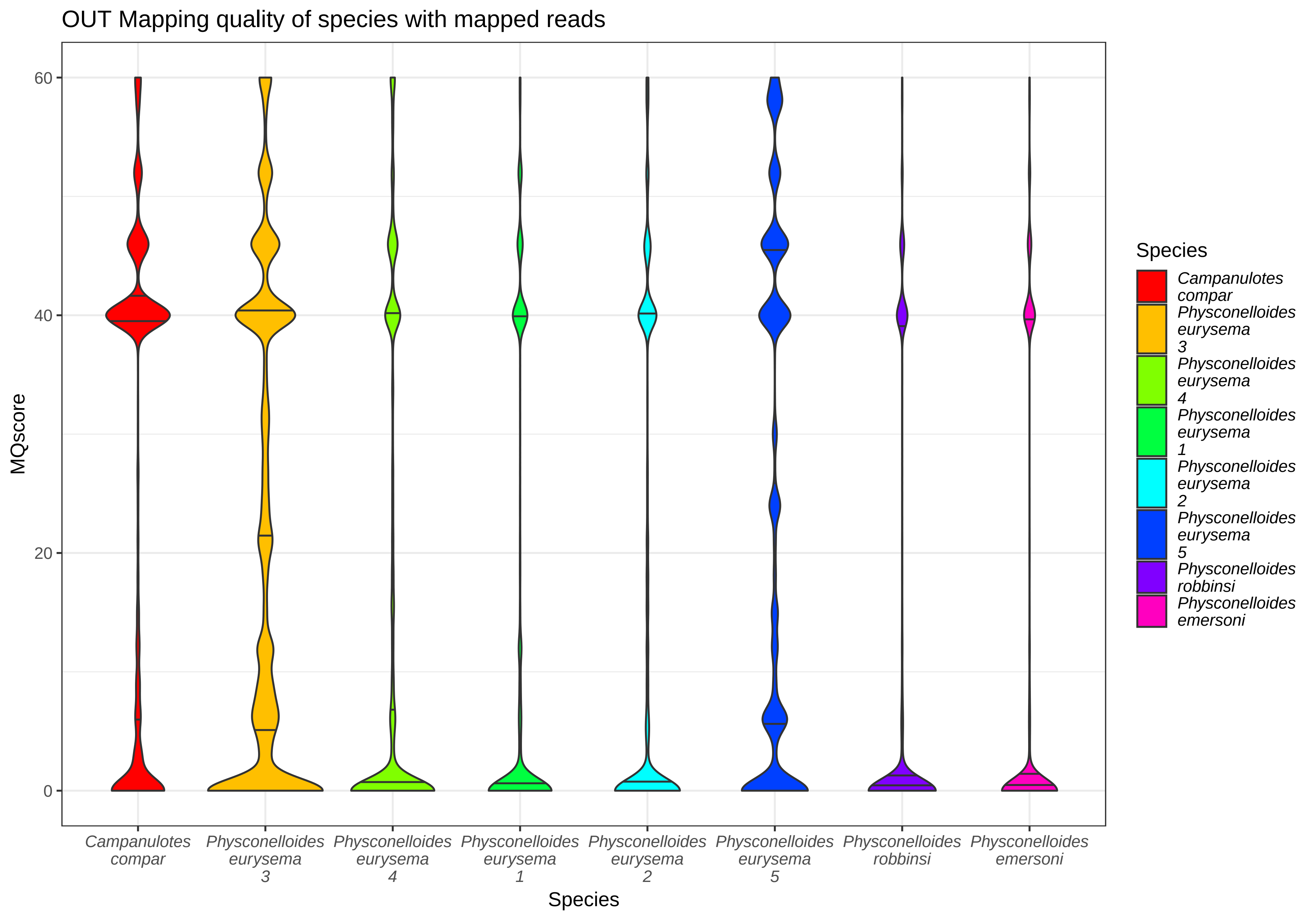

### OUT_plotMQ_Pheur_Cotal_3_14_2016_6.pdf

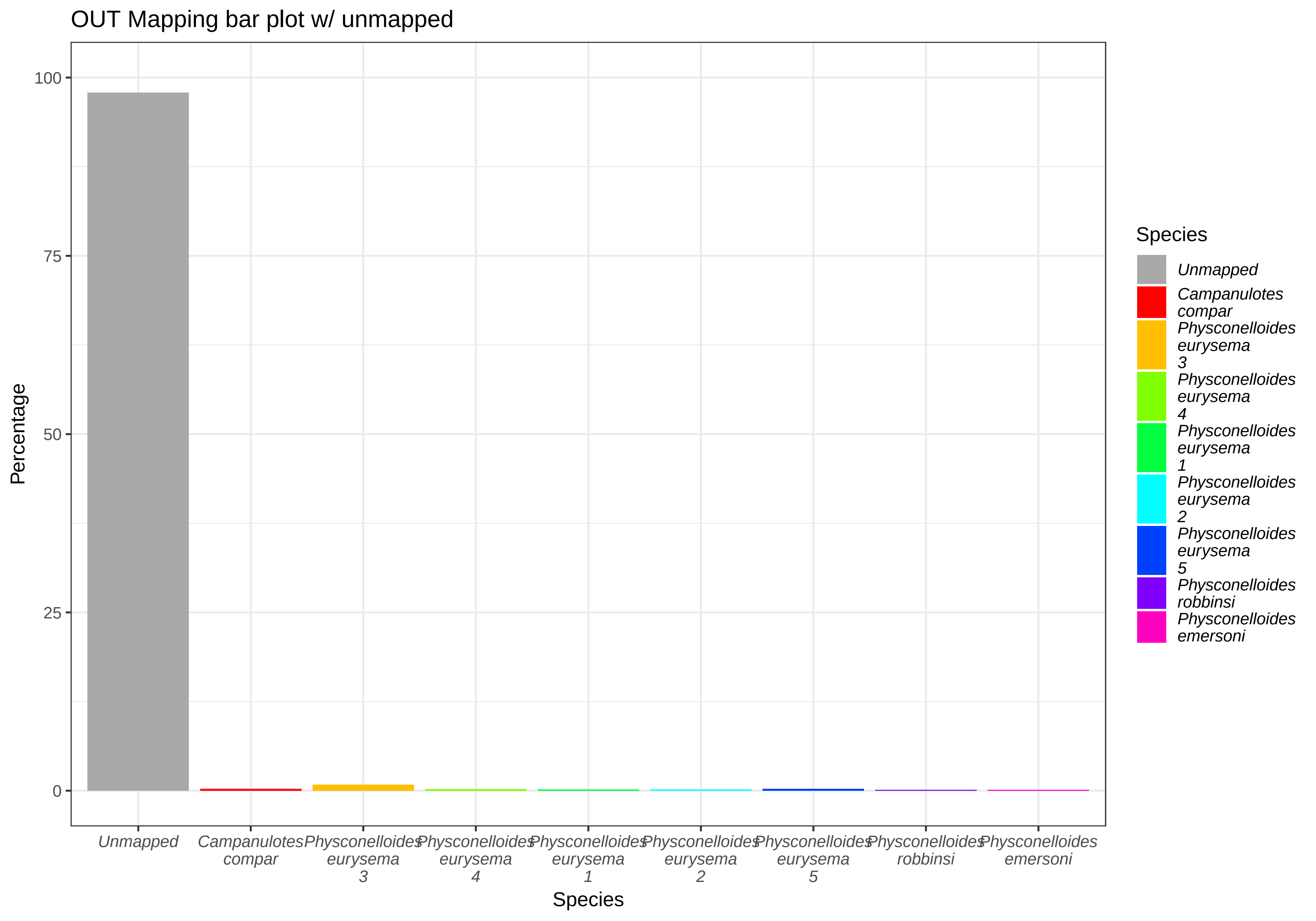

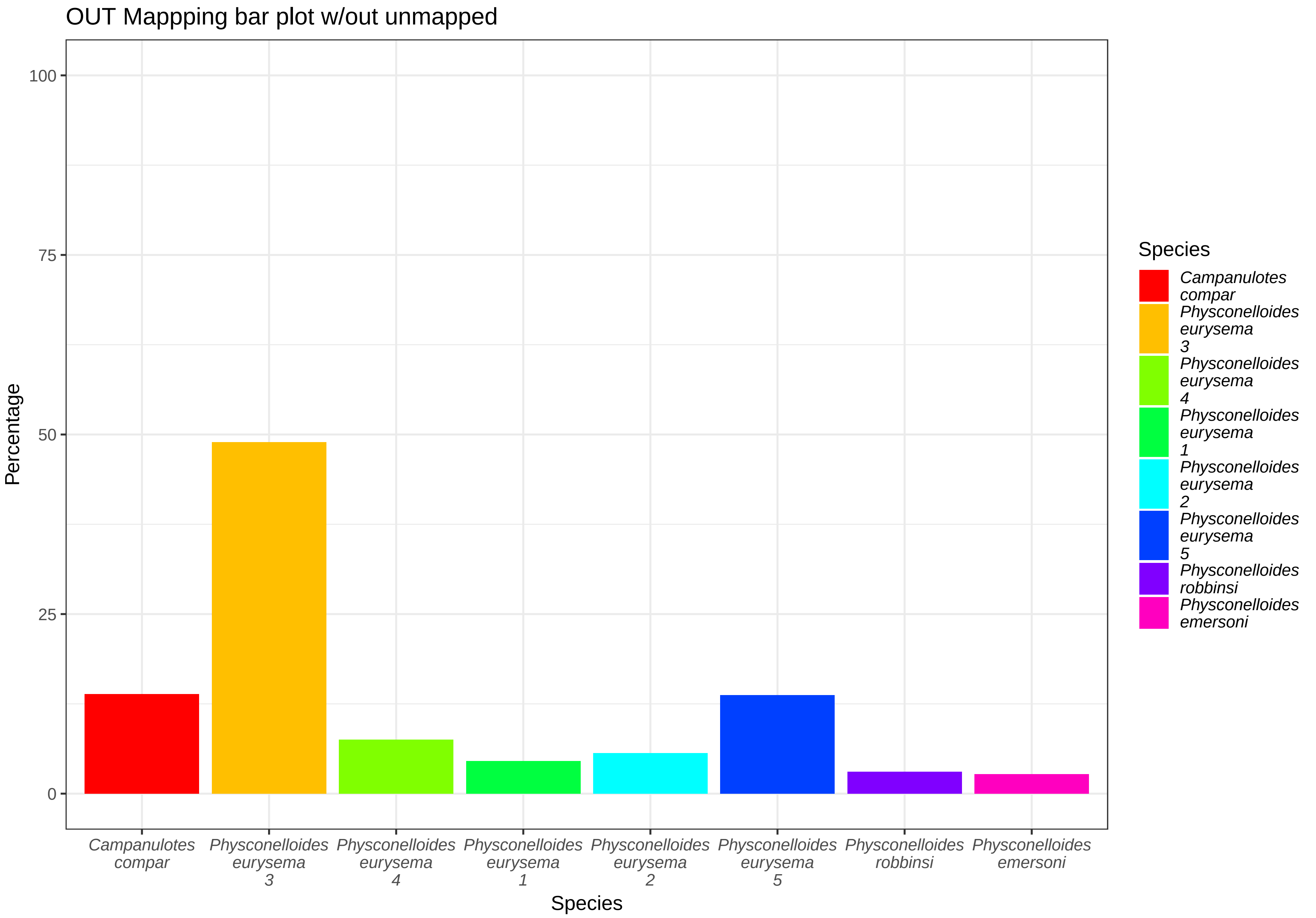

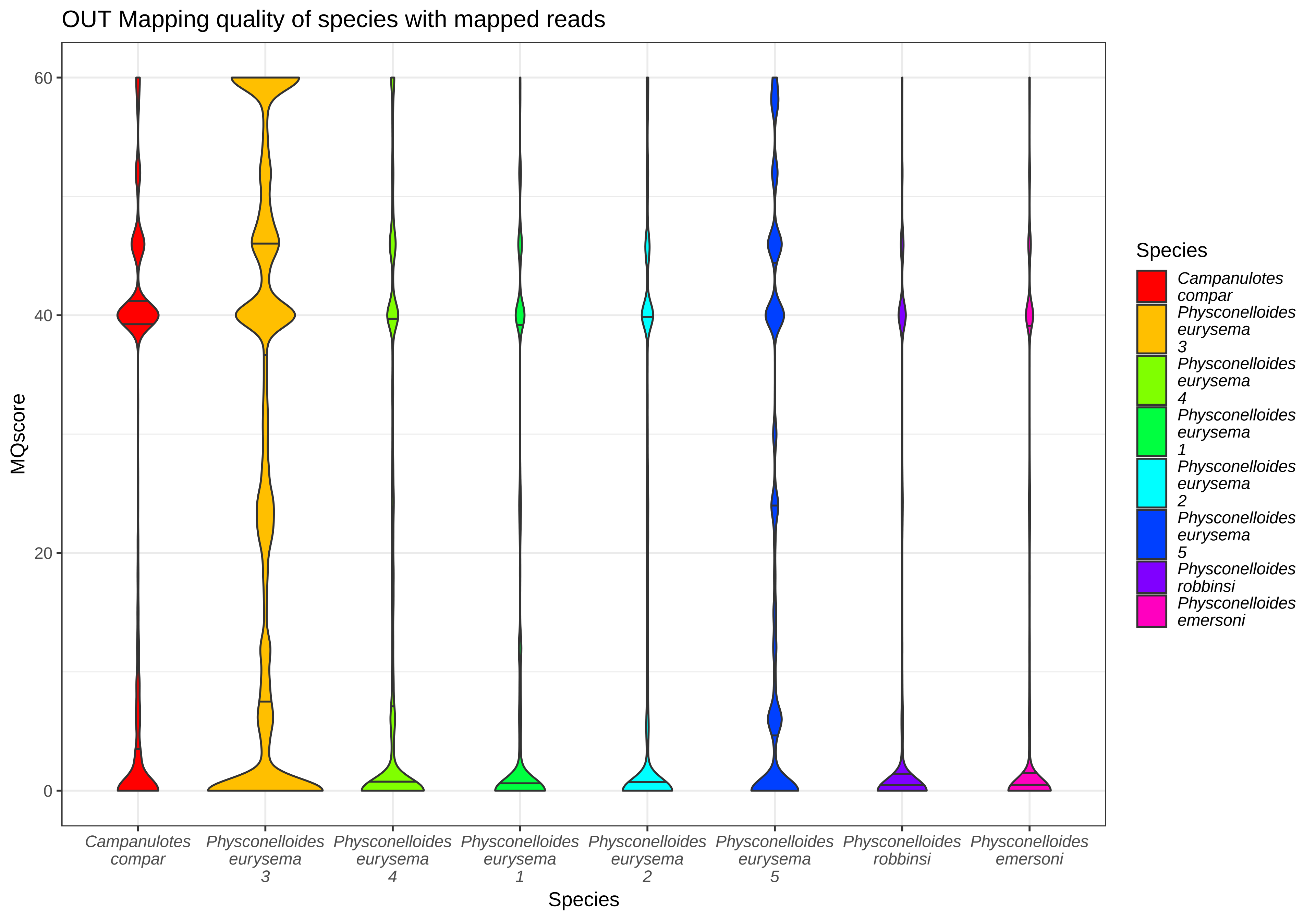

### OUT_plotMQ_Pheur_Cotal_3_16_2016_3.pdf

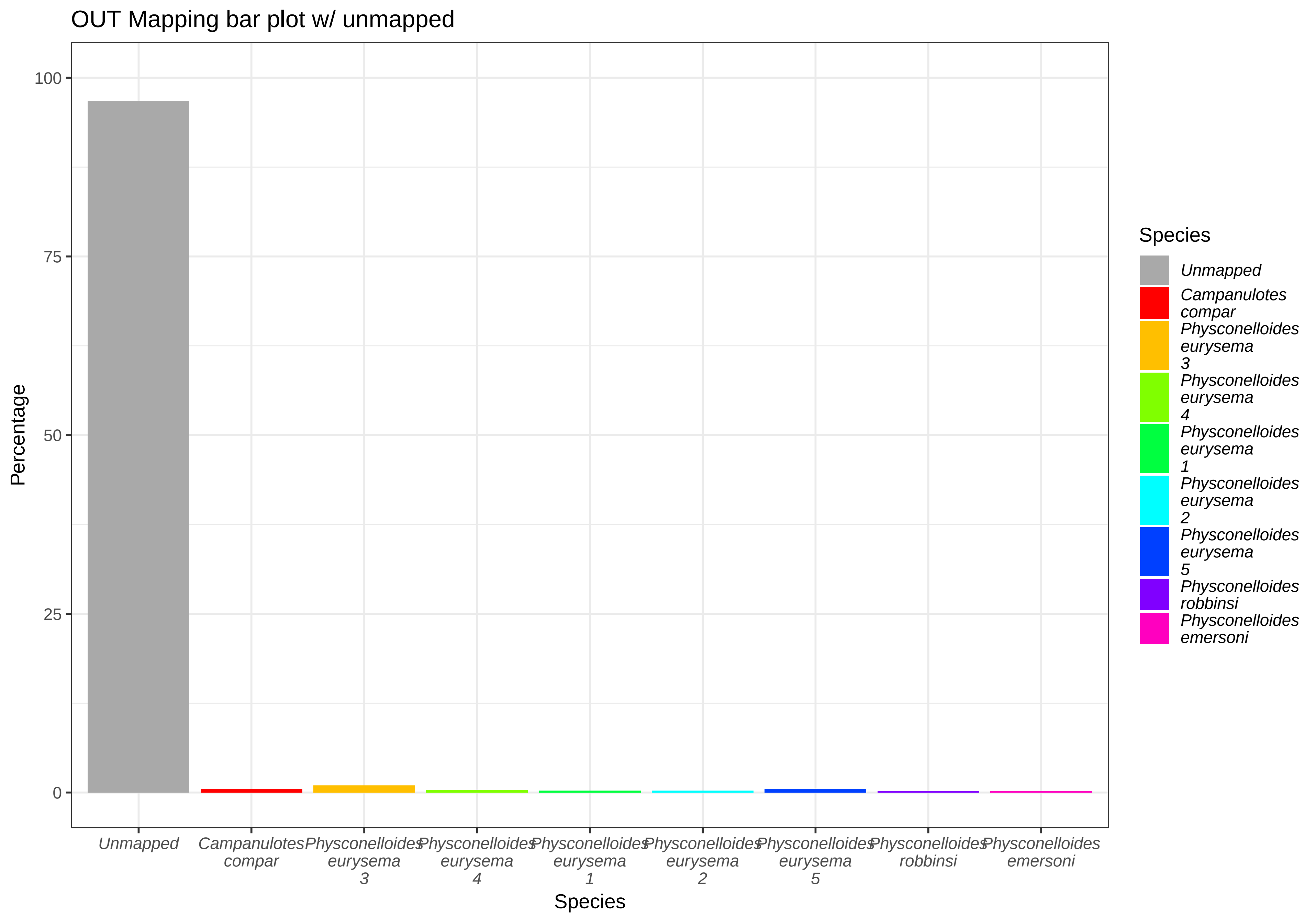

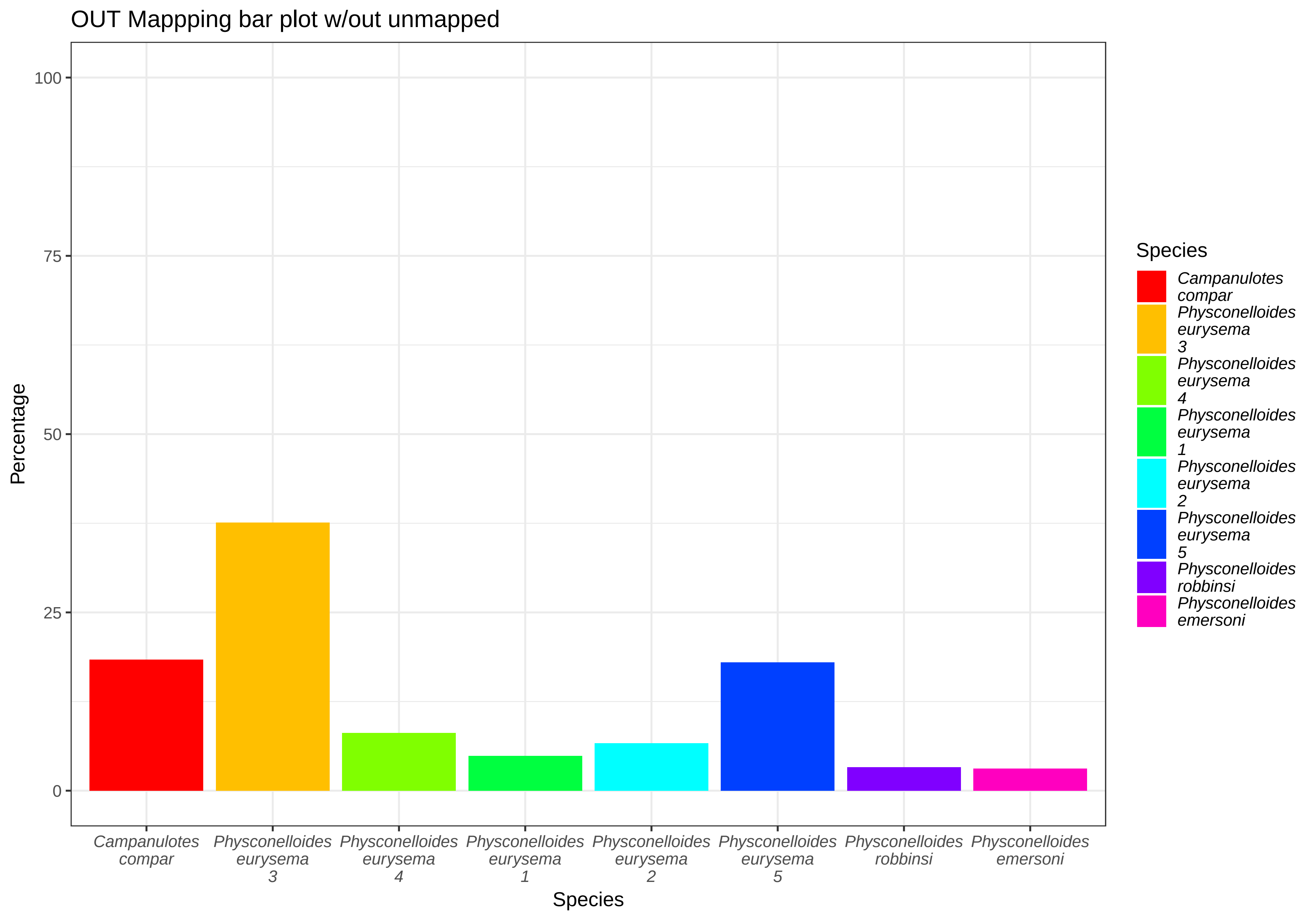

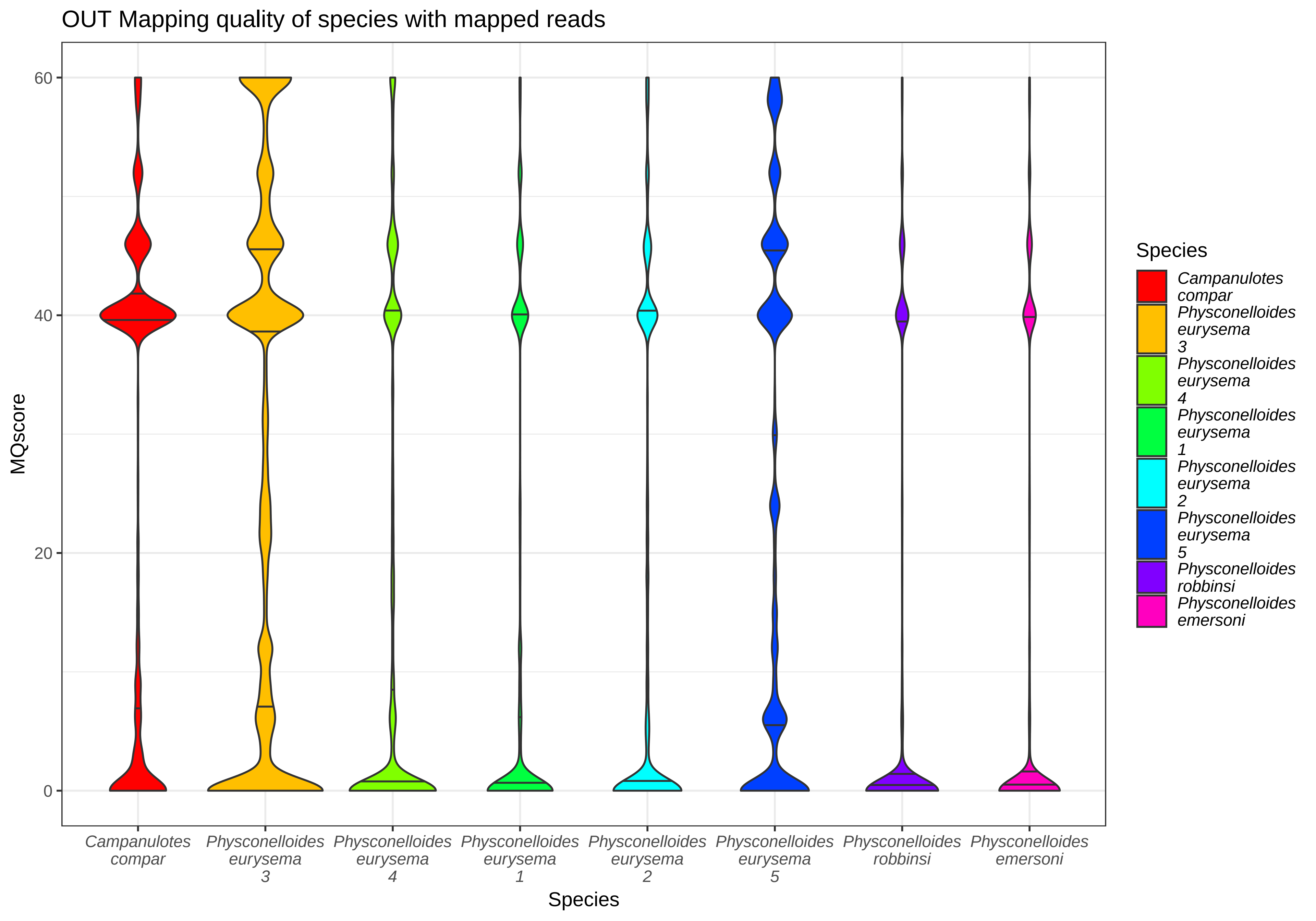

### OUT_plotMQ_Pheur_Cotal_3_16_2016_12.pdf

OUT Mapping bar plot w/ unmapped

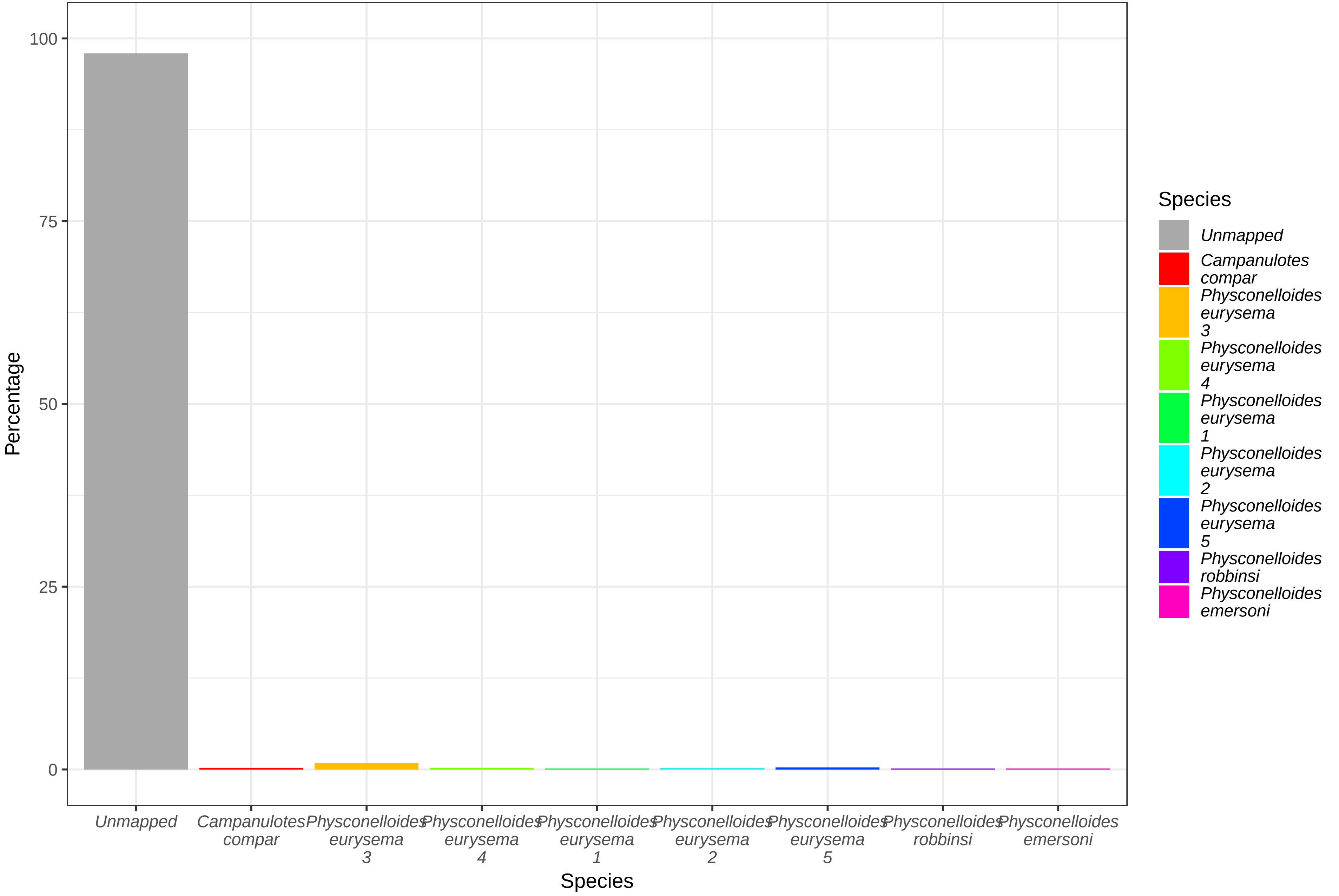

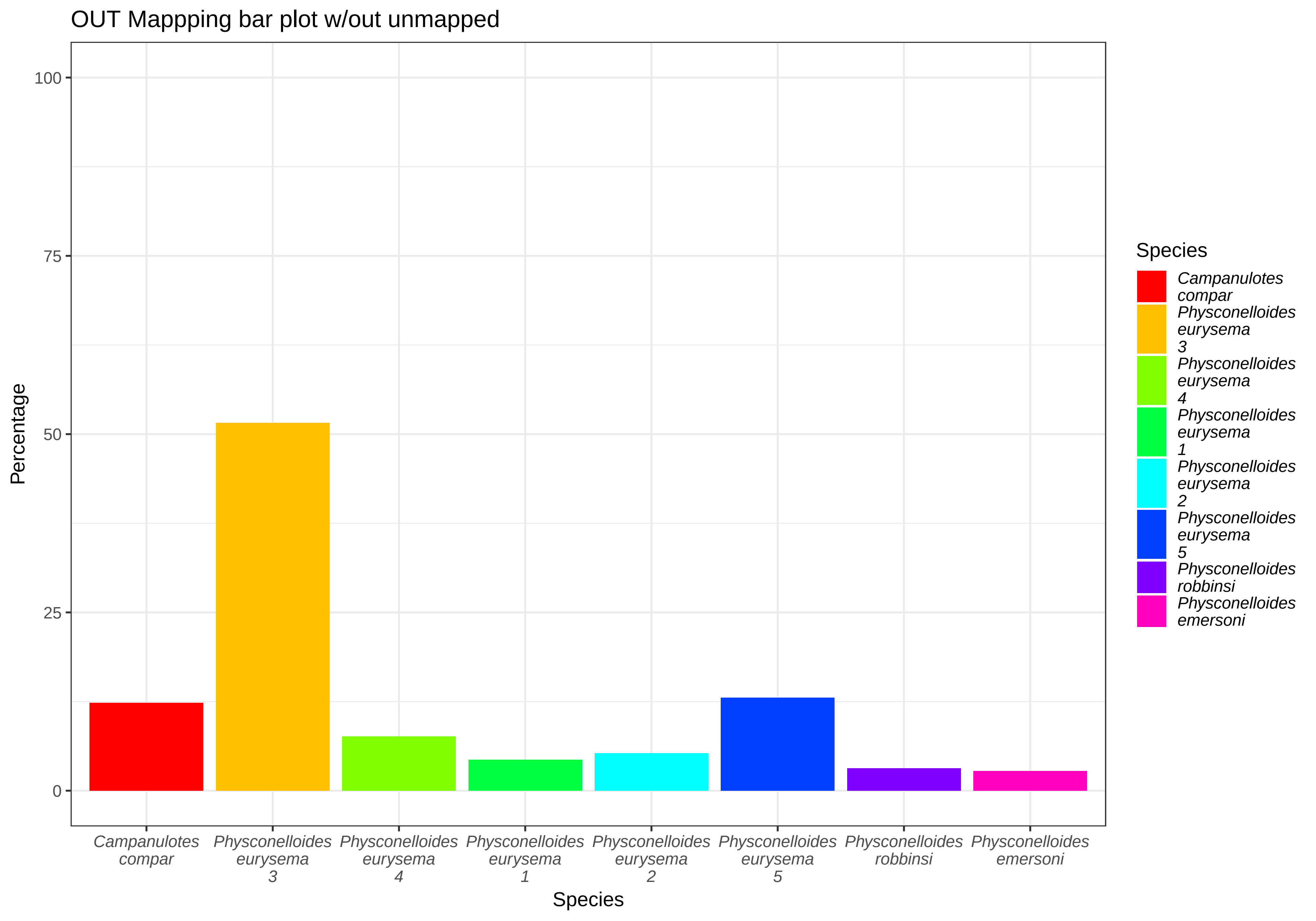

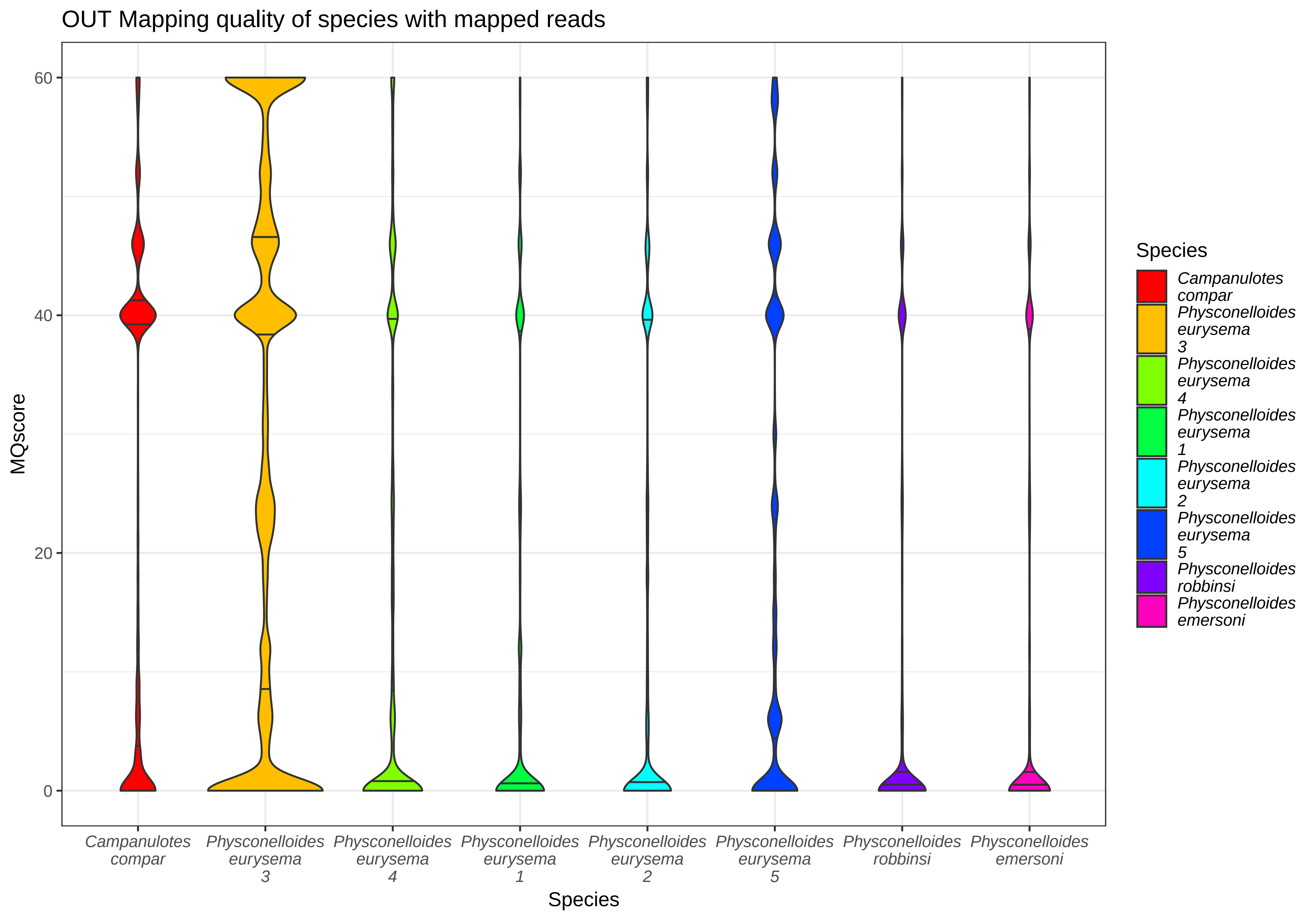

### OUT_plotMQ_Pheur_Cotal_6_15_2015_4.pdf

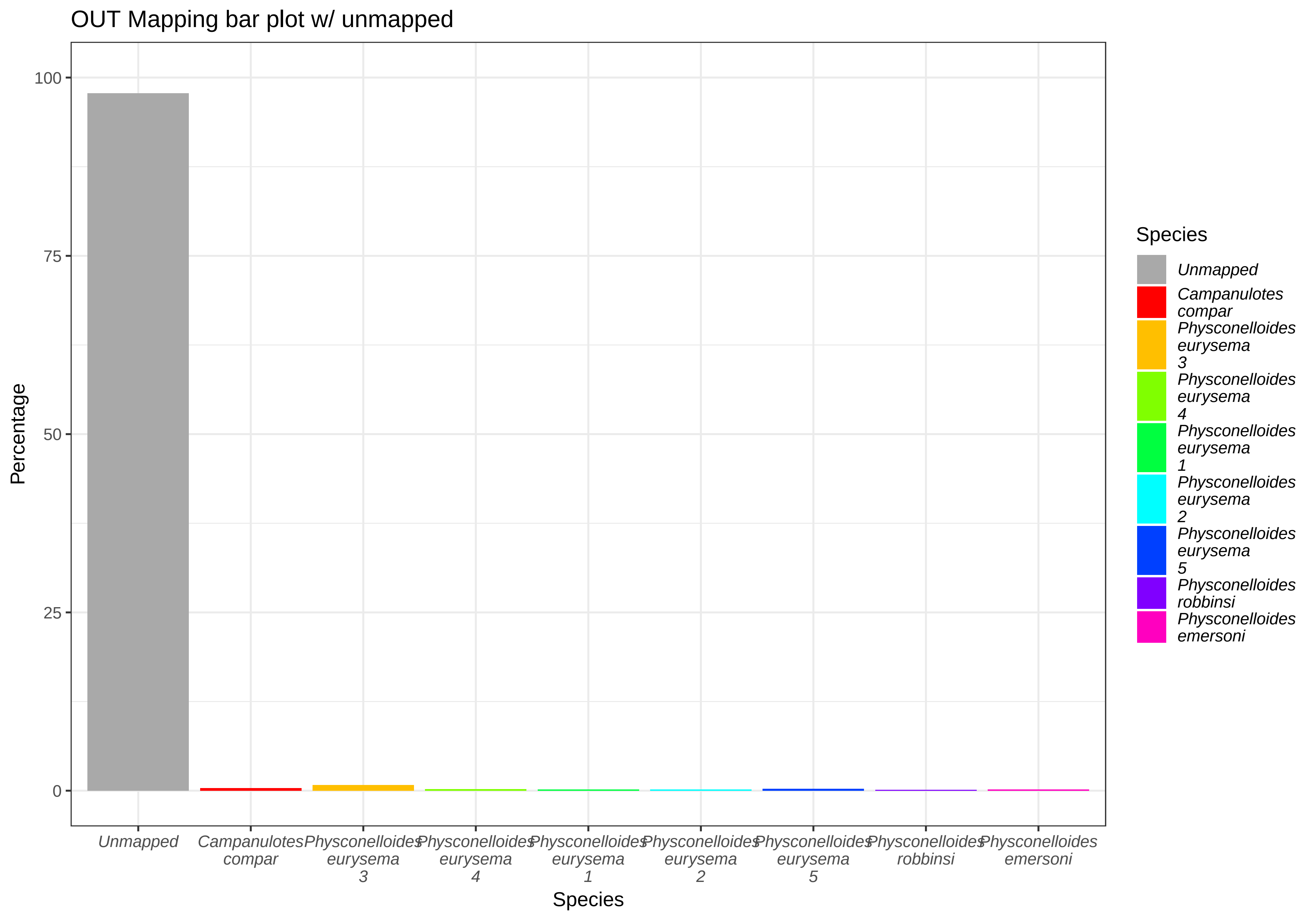

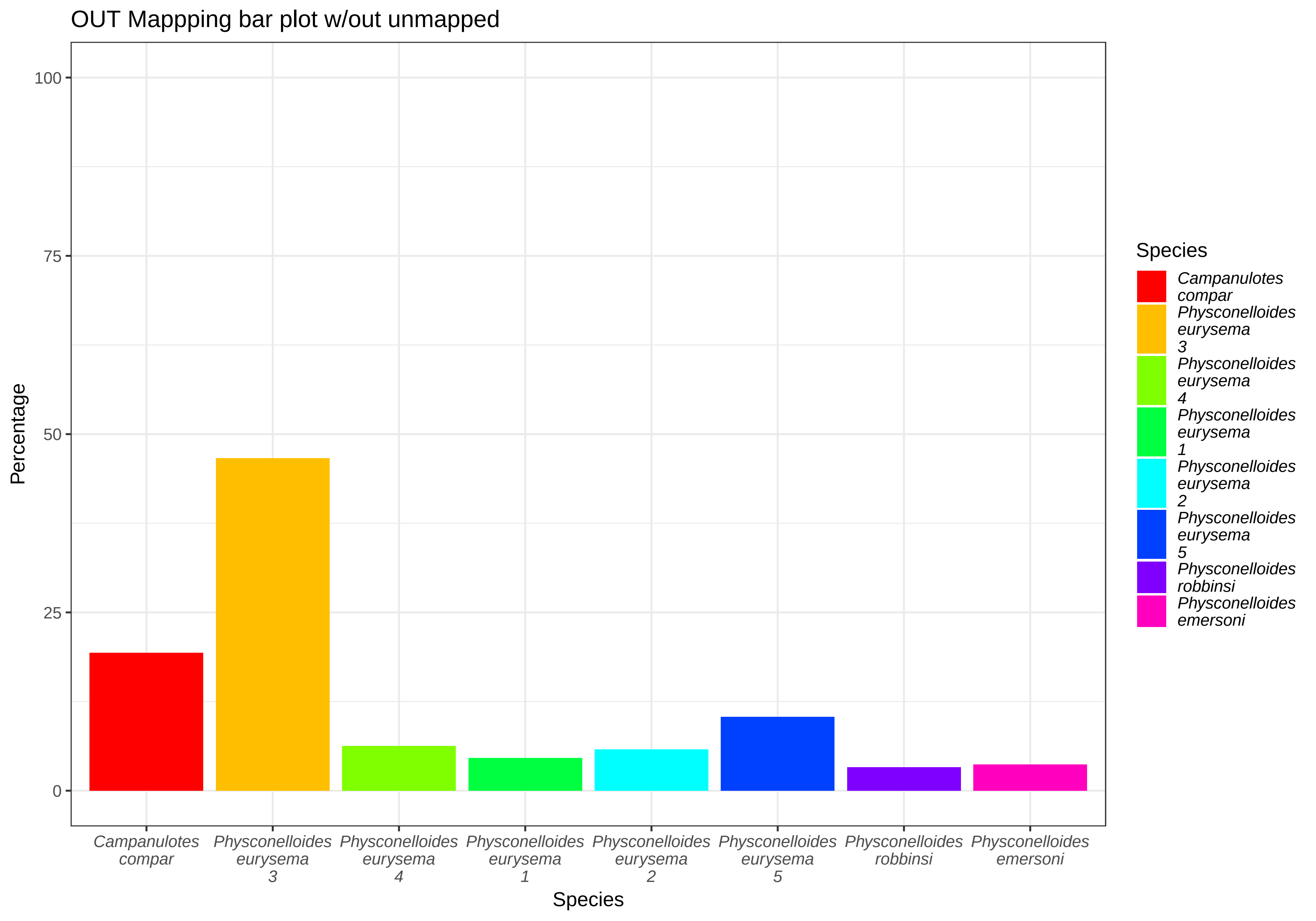

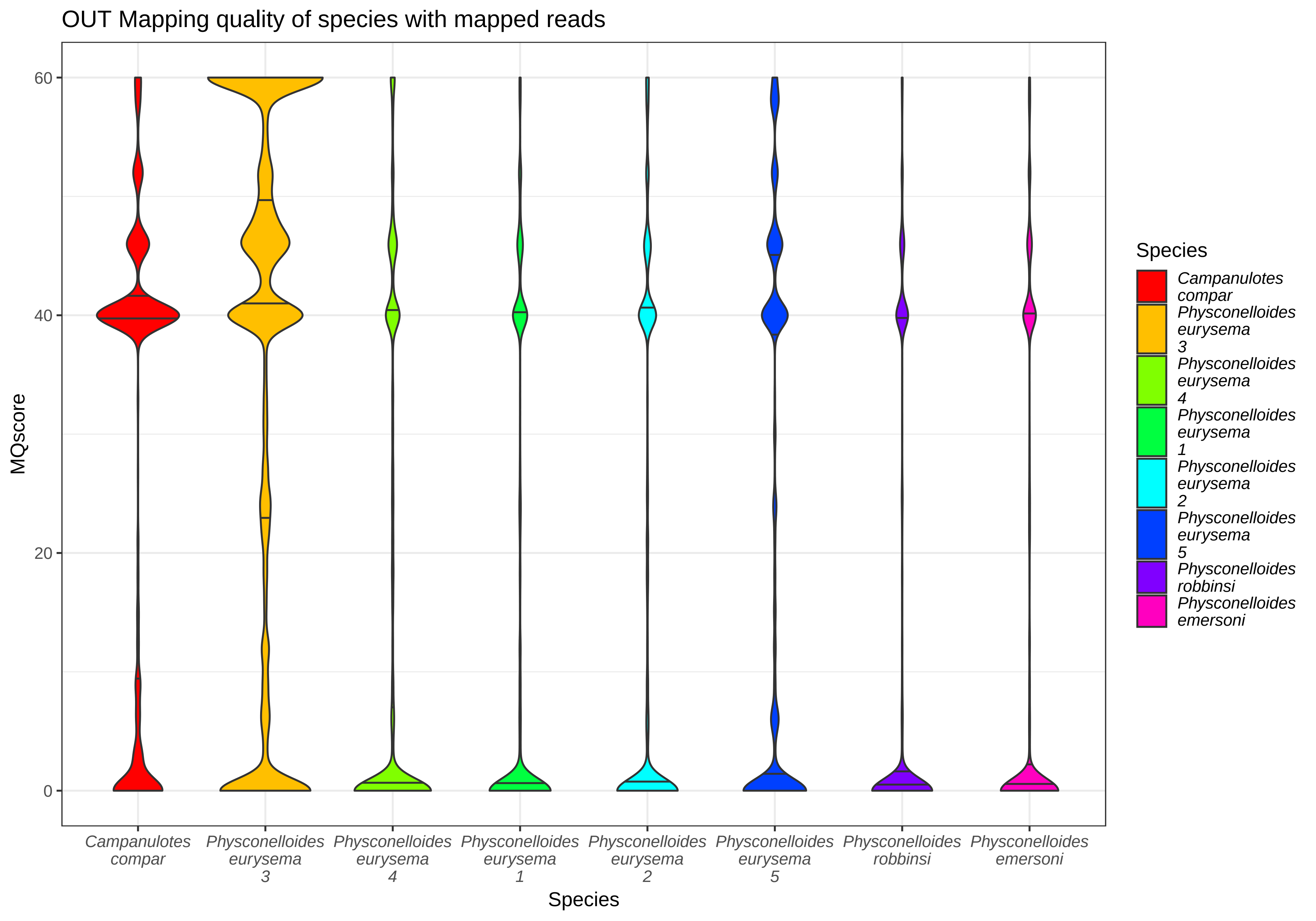

### OUT_plotMQ_Pheur_Urcam_3_14_2016_3.pdf

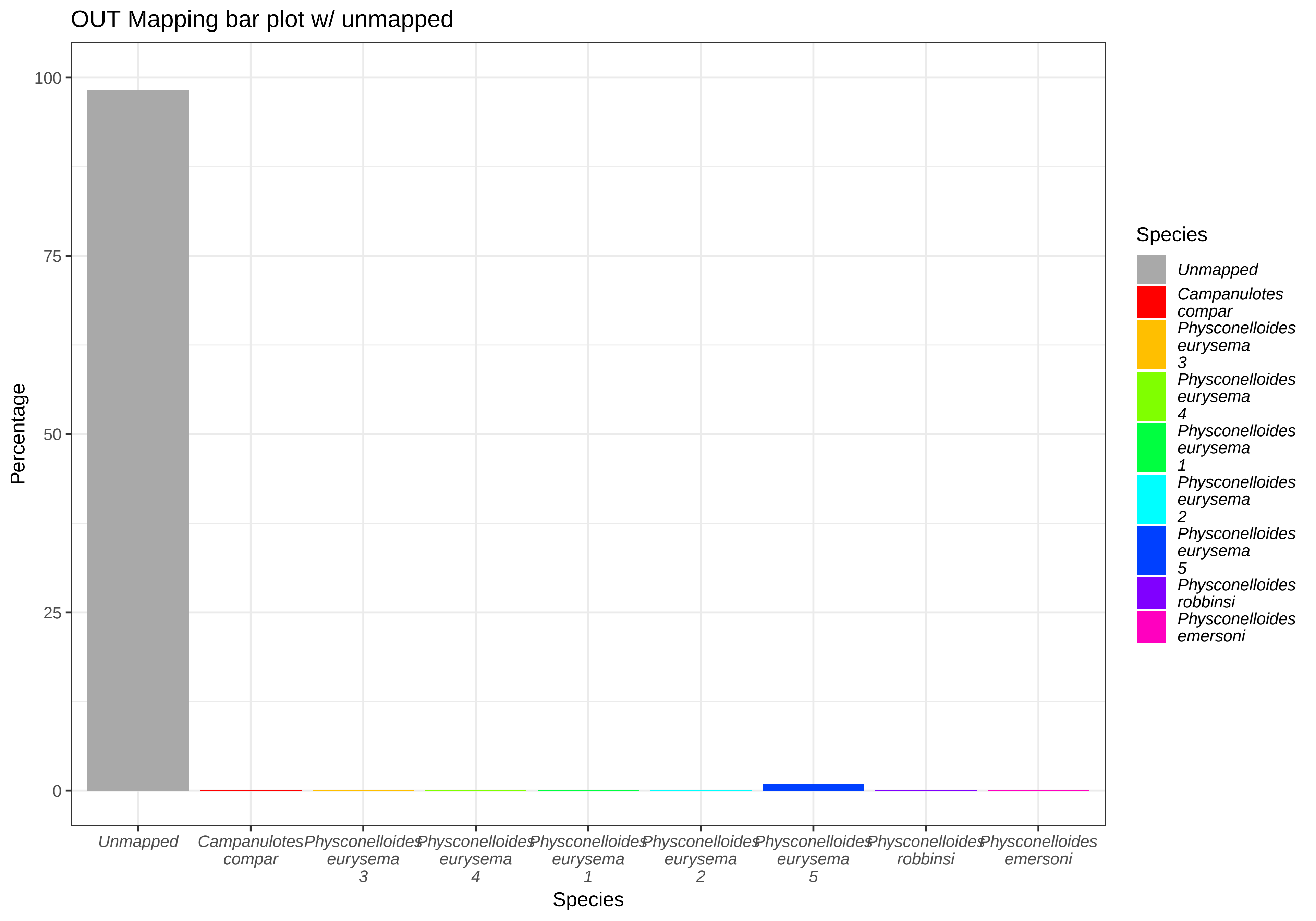

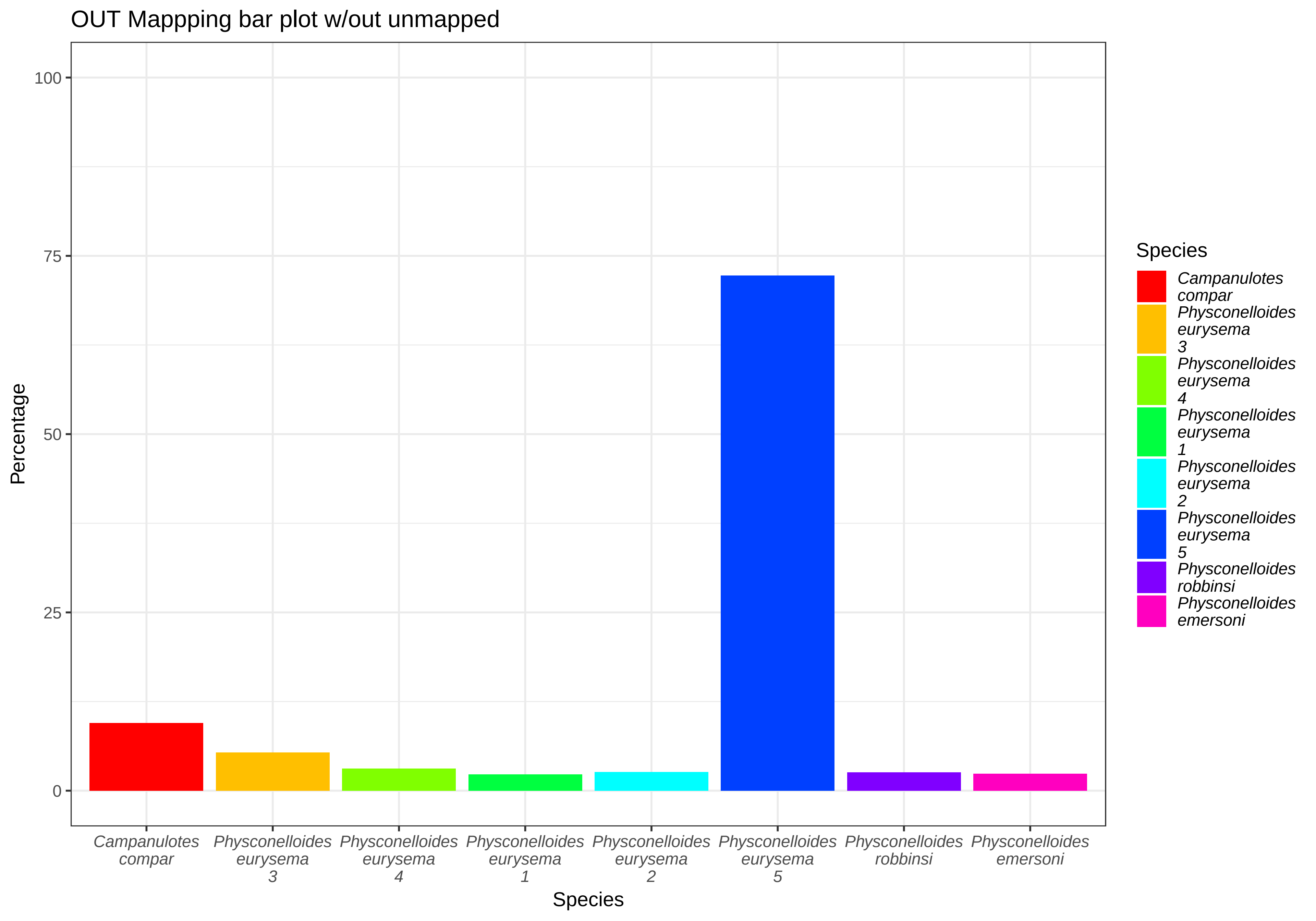

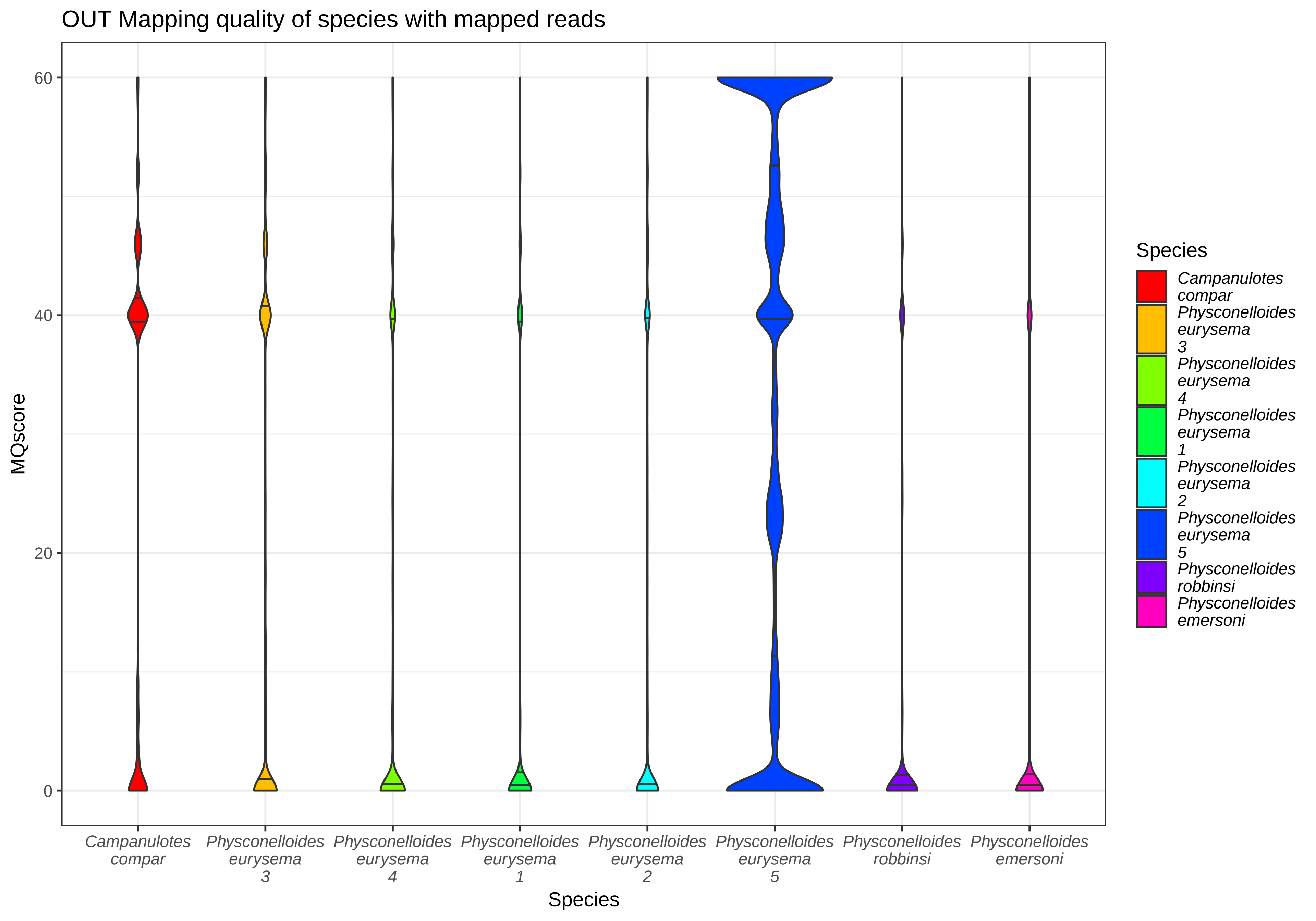

### OUT_plotMQ_Phrob_Mecec_3_16_2016_10.pdf

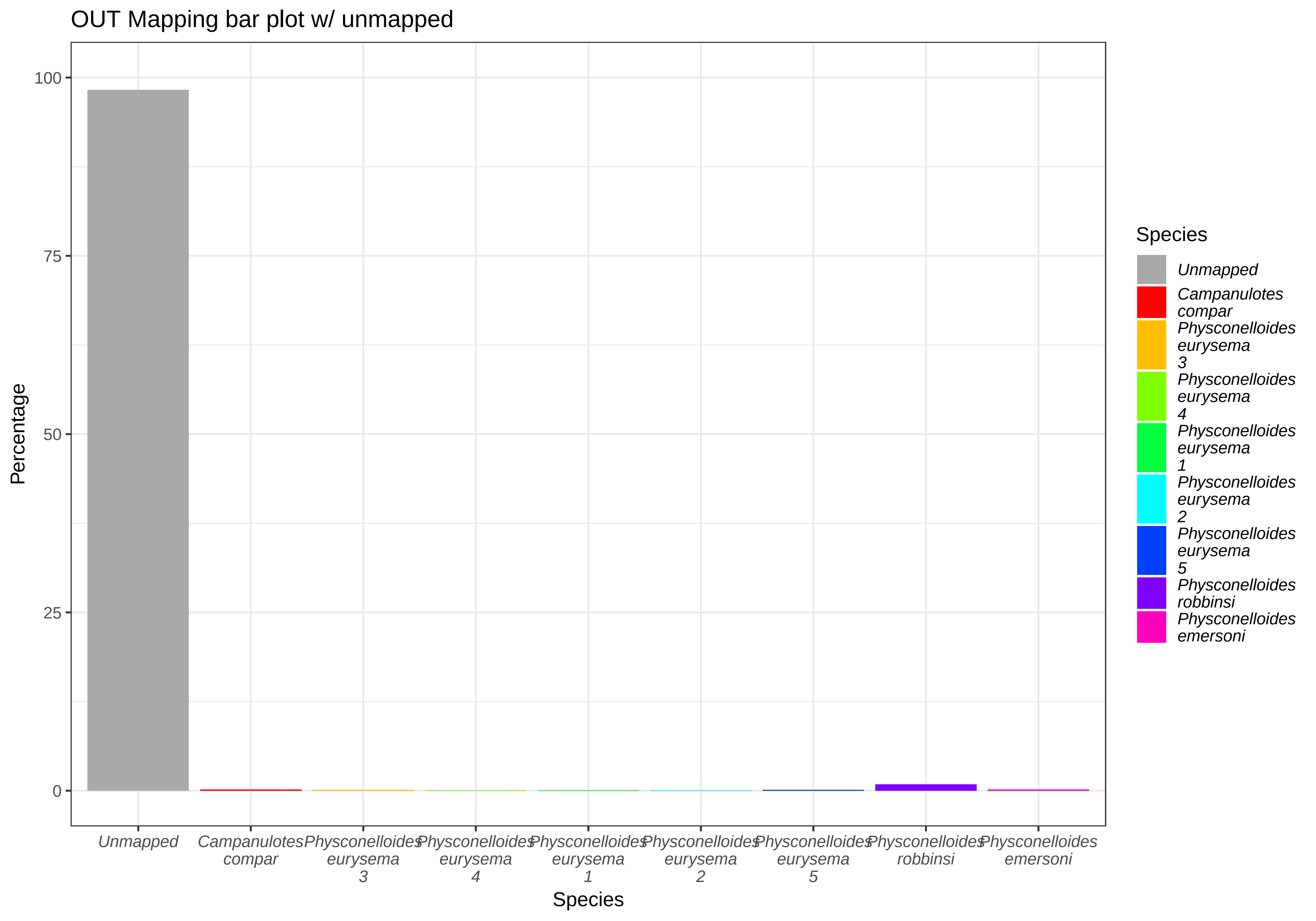

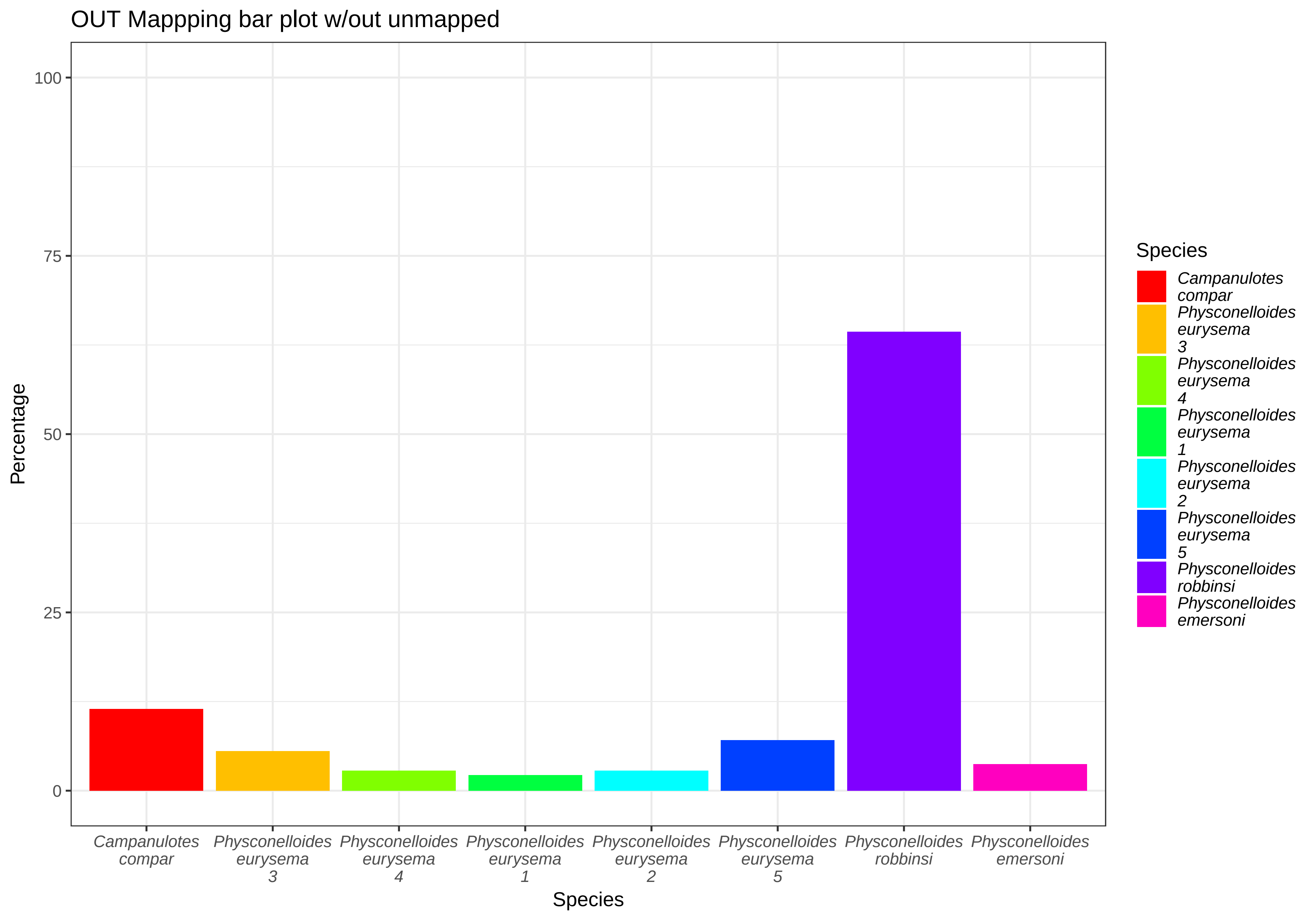

OUT Mapping quality of species with mapped reads

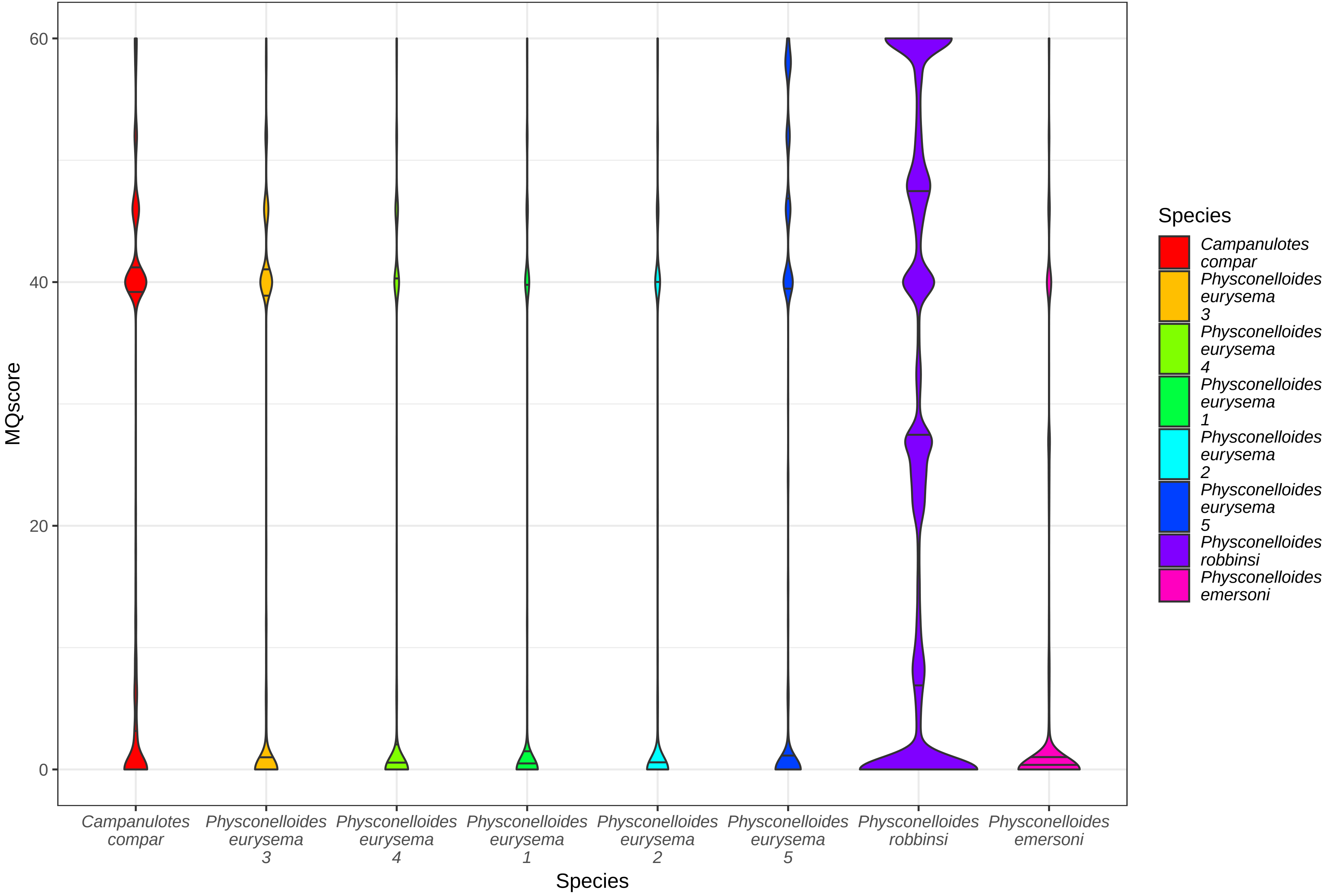

### OUT_plotMQ_Phrob_Mecec_3_16_2016_11.pdf

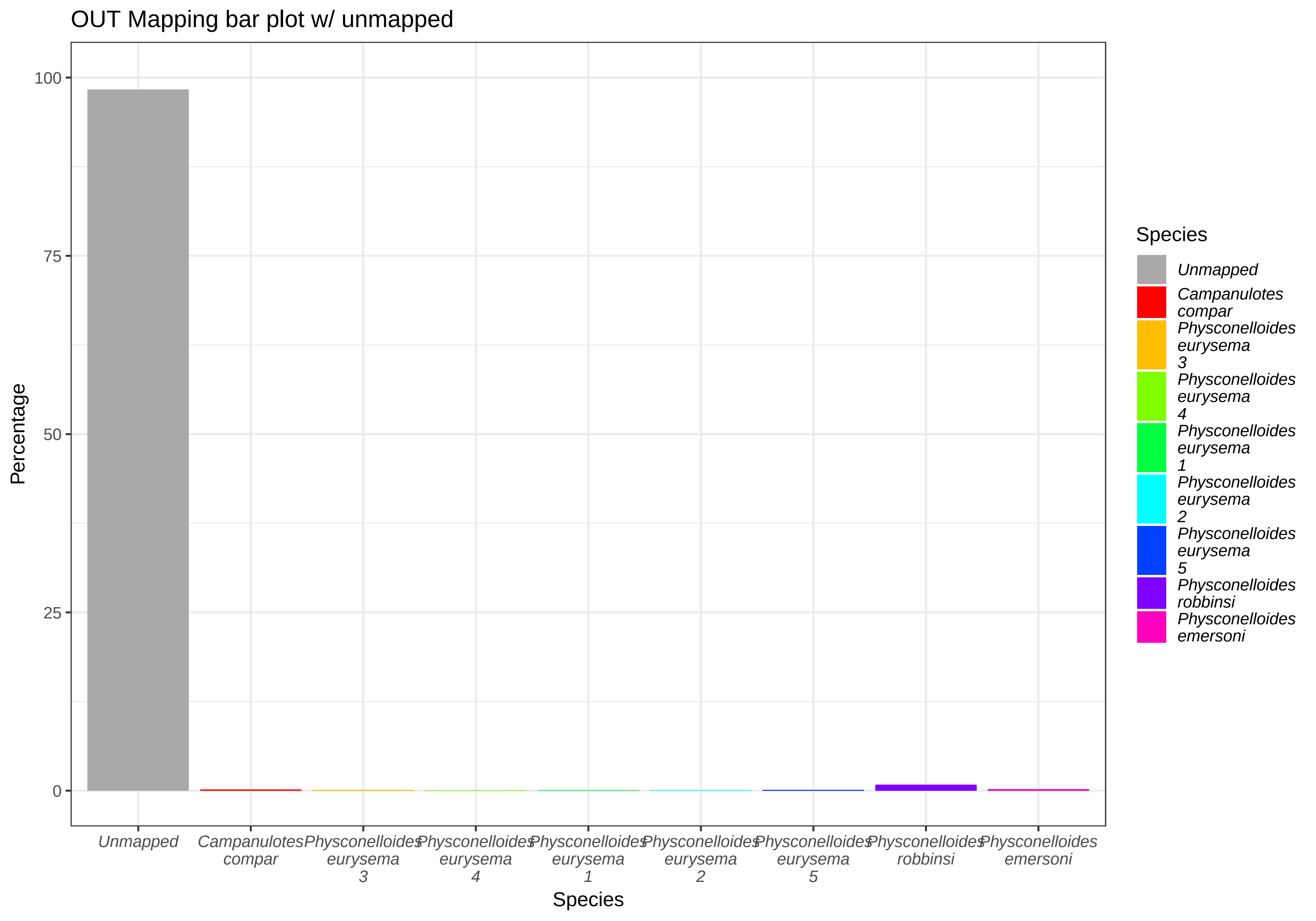

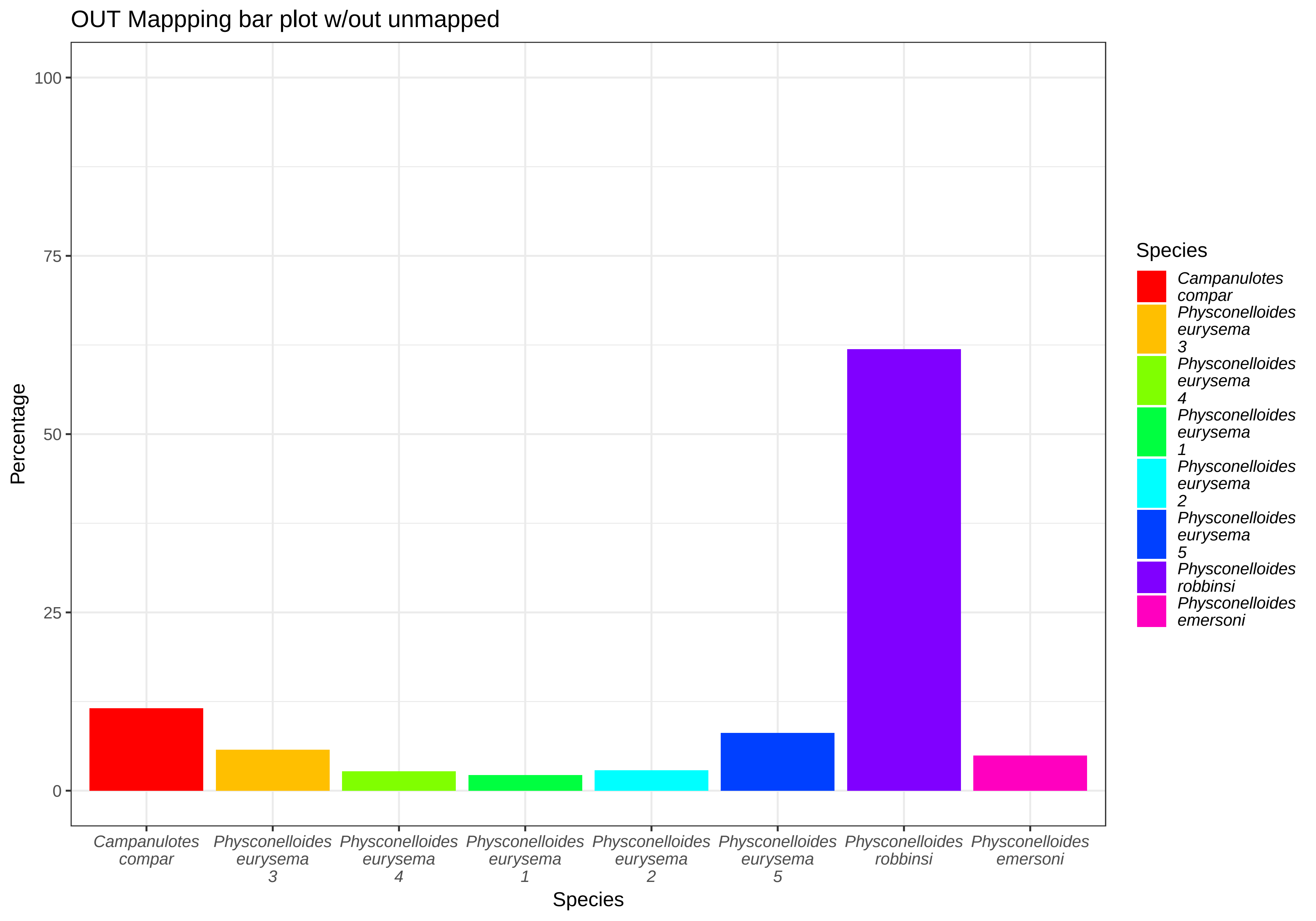

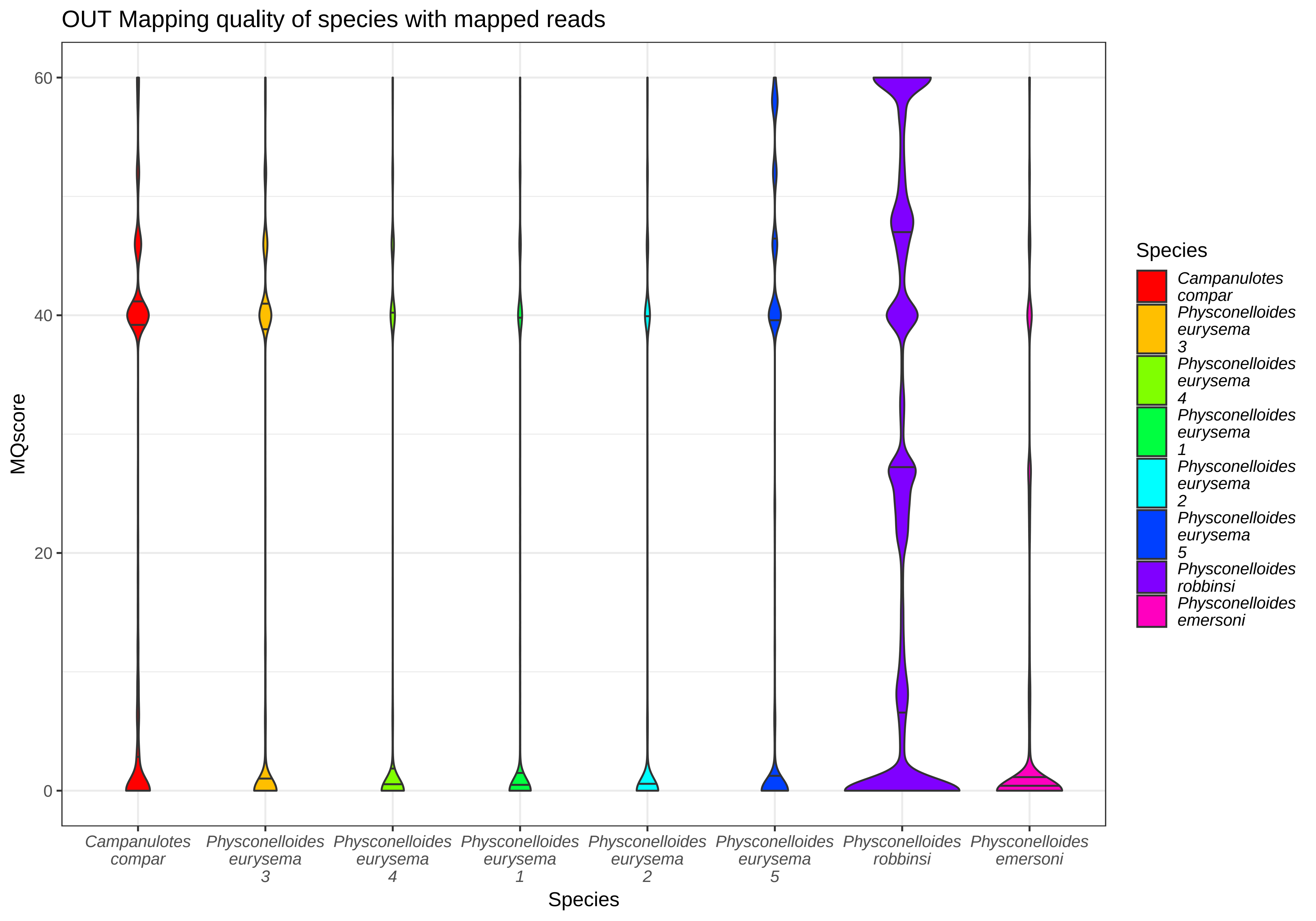

### OUT_plotMQ_Phrob_Mecec_5_26_2015_7.pdf

OUT Mapping bar plot w/ unmapped

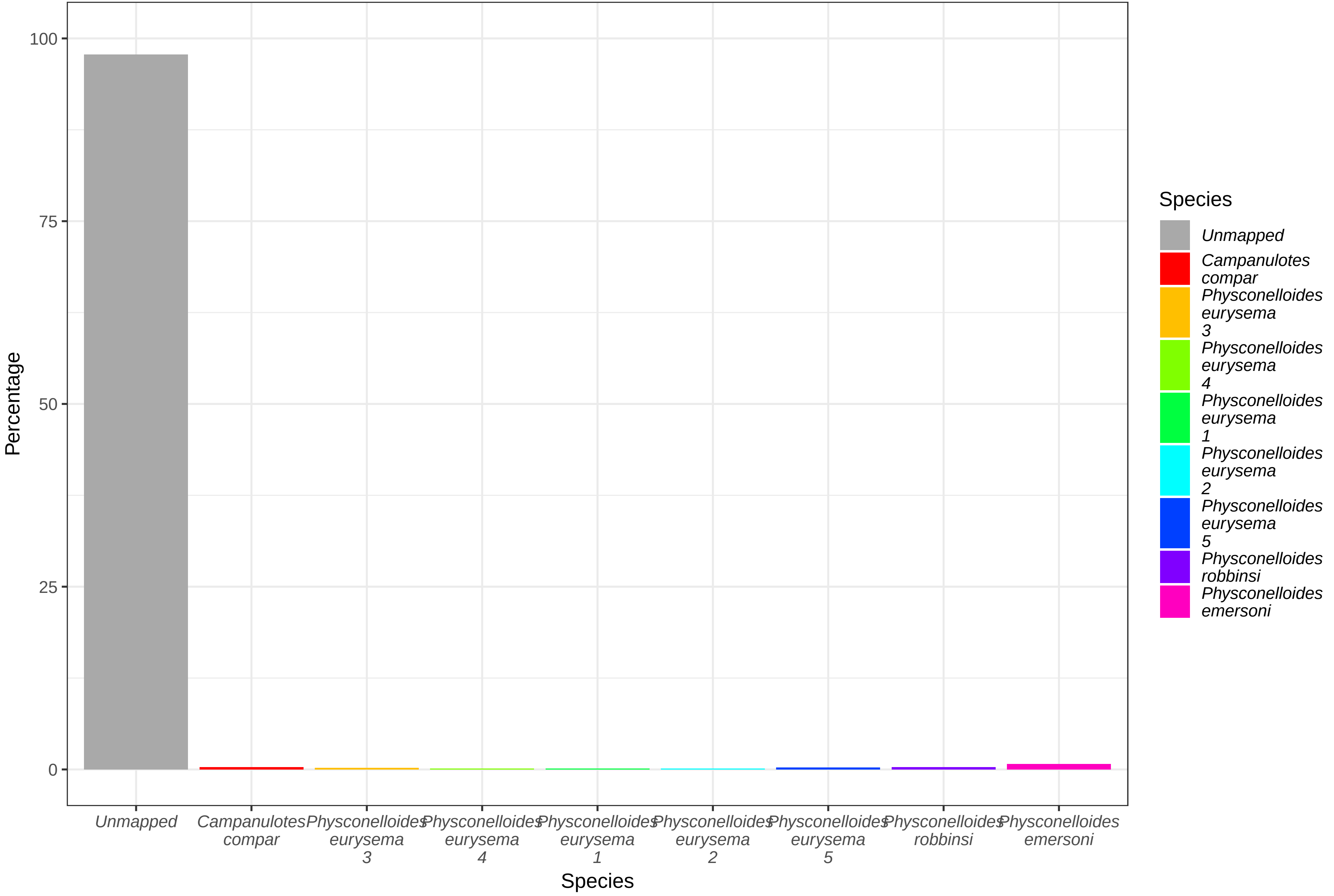

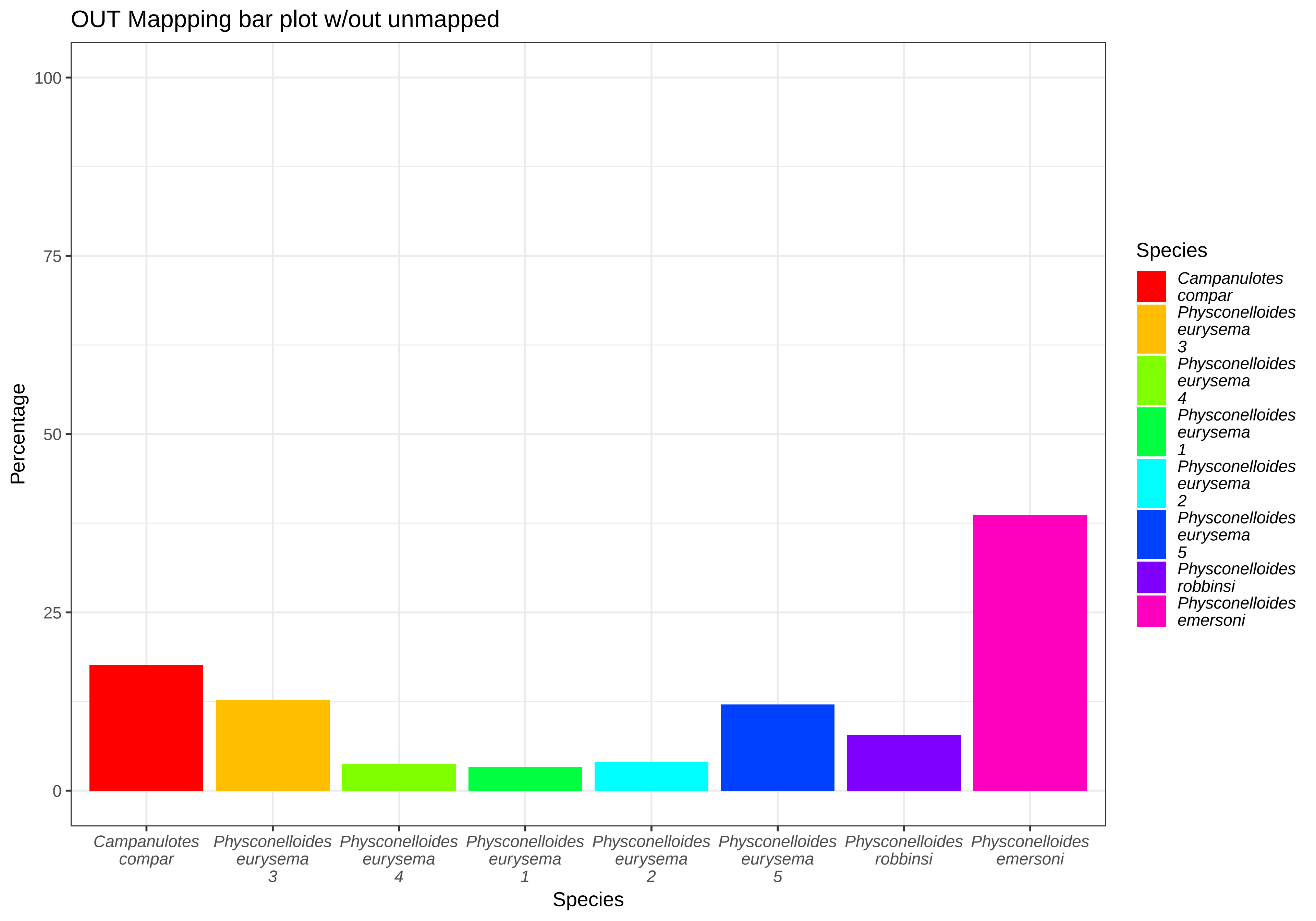

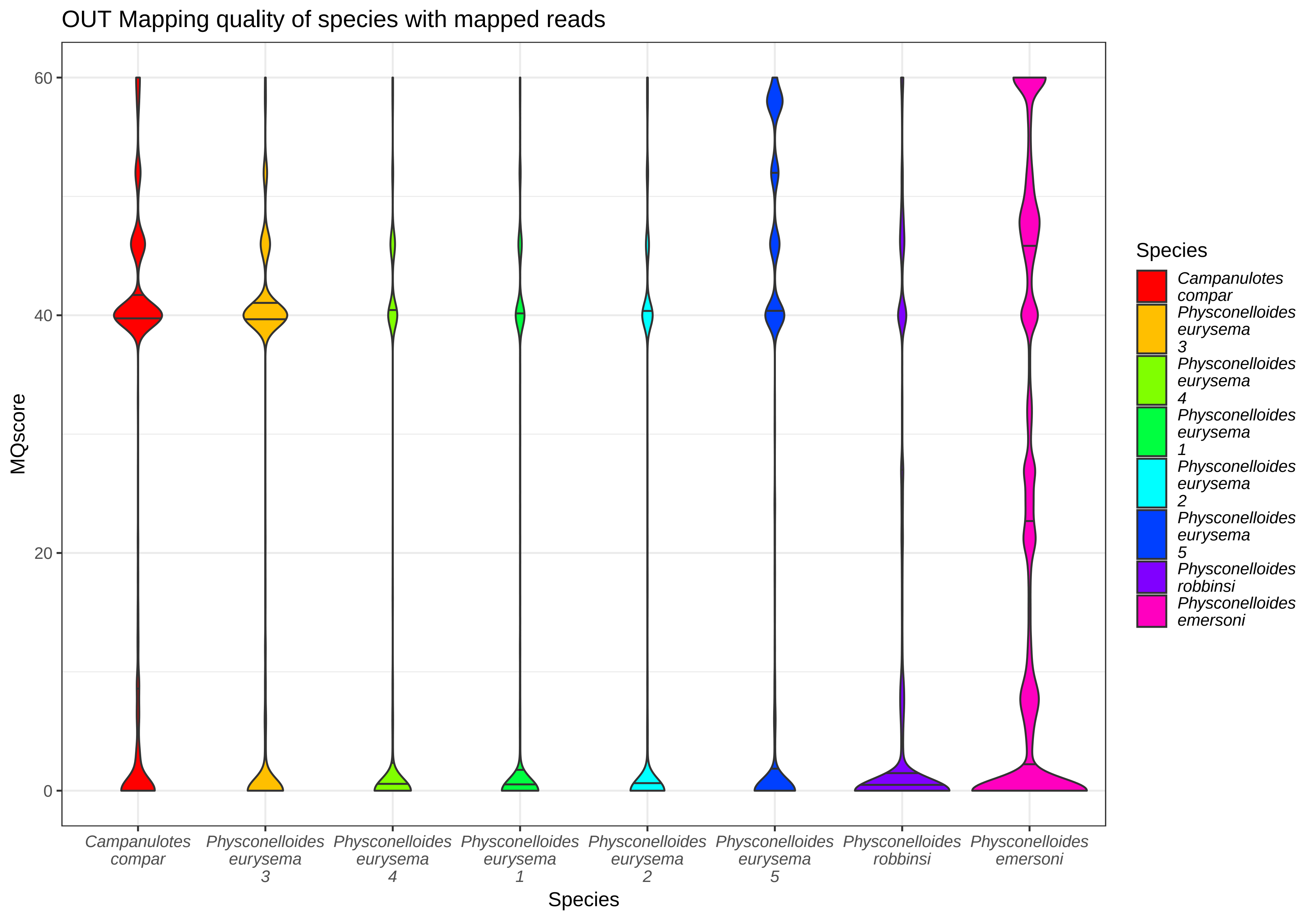

### OUT_plotMQ_Phrob_Mecec_6_15_2015_6.pdf

OUT Mapping bar plot w/ unmapped

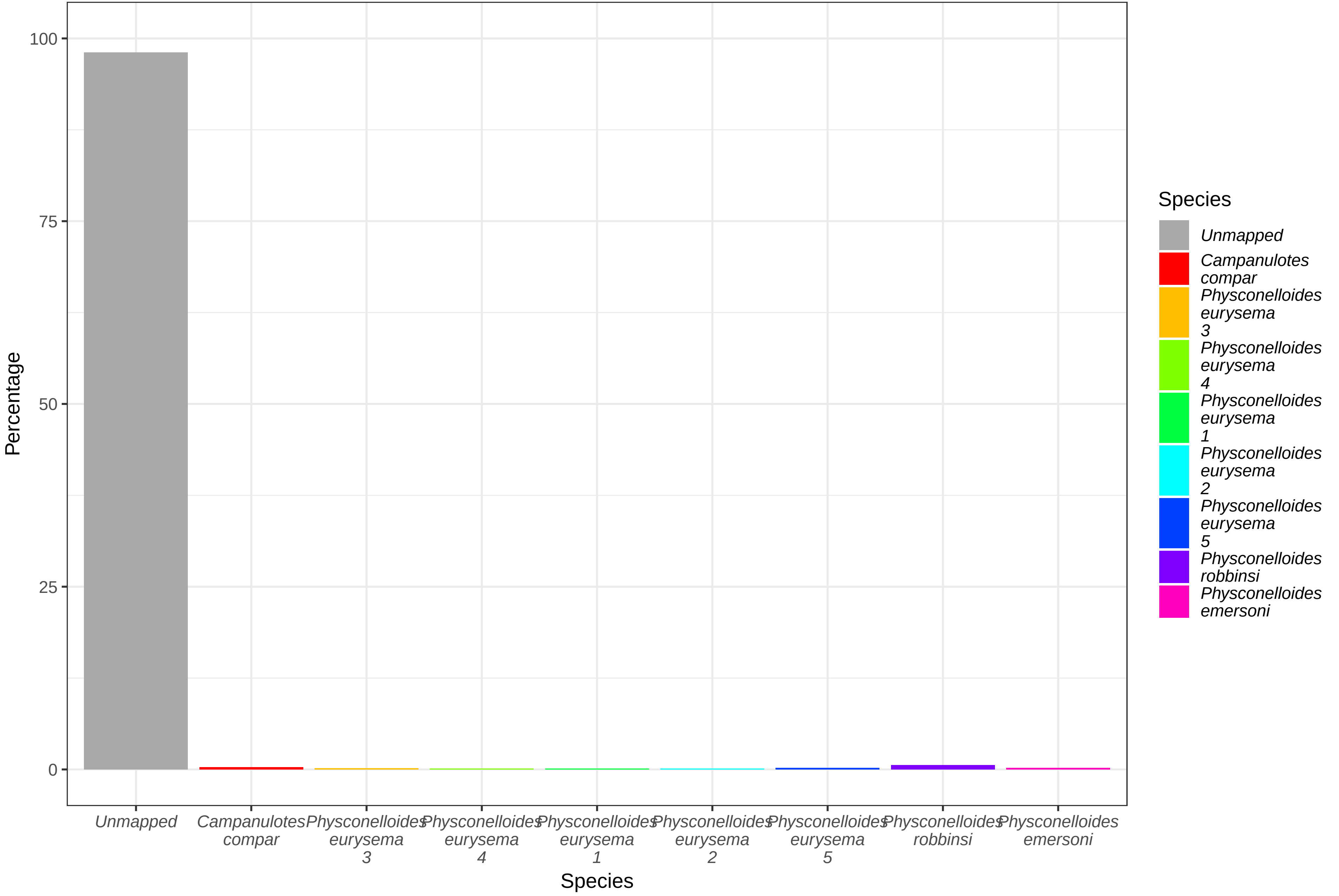

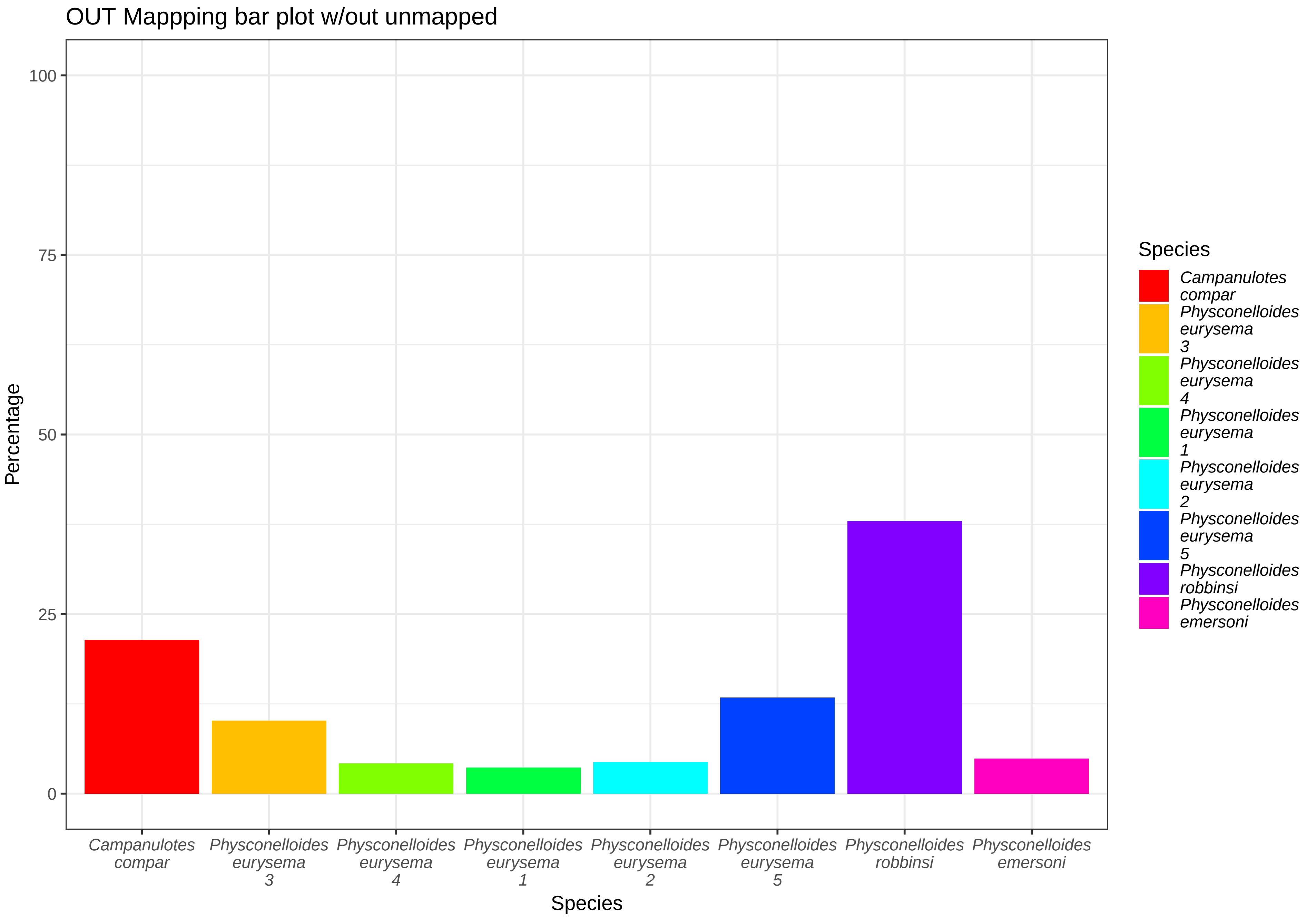

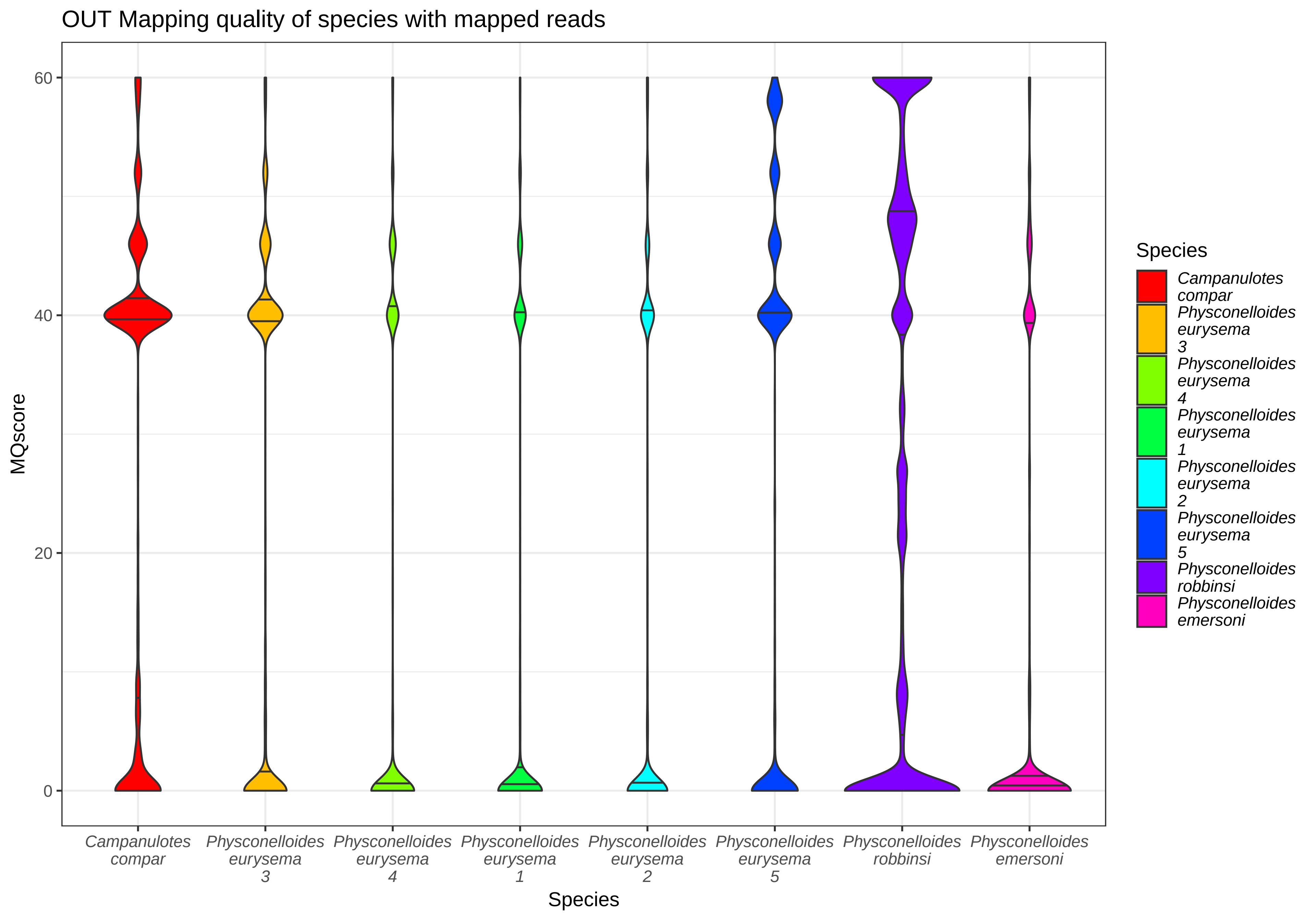

### OUT_speciesDepth_coalt.pdf

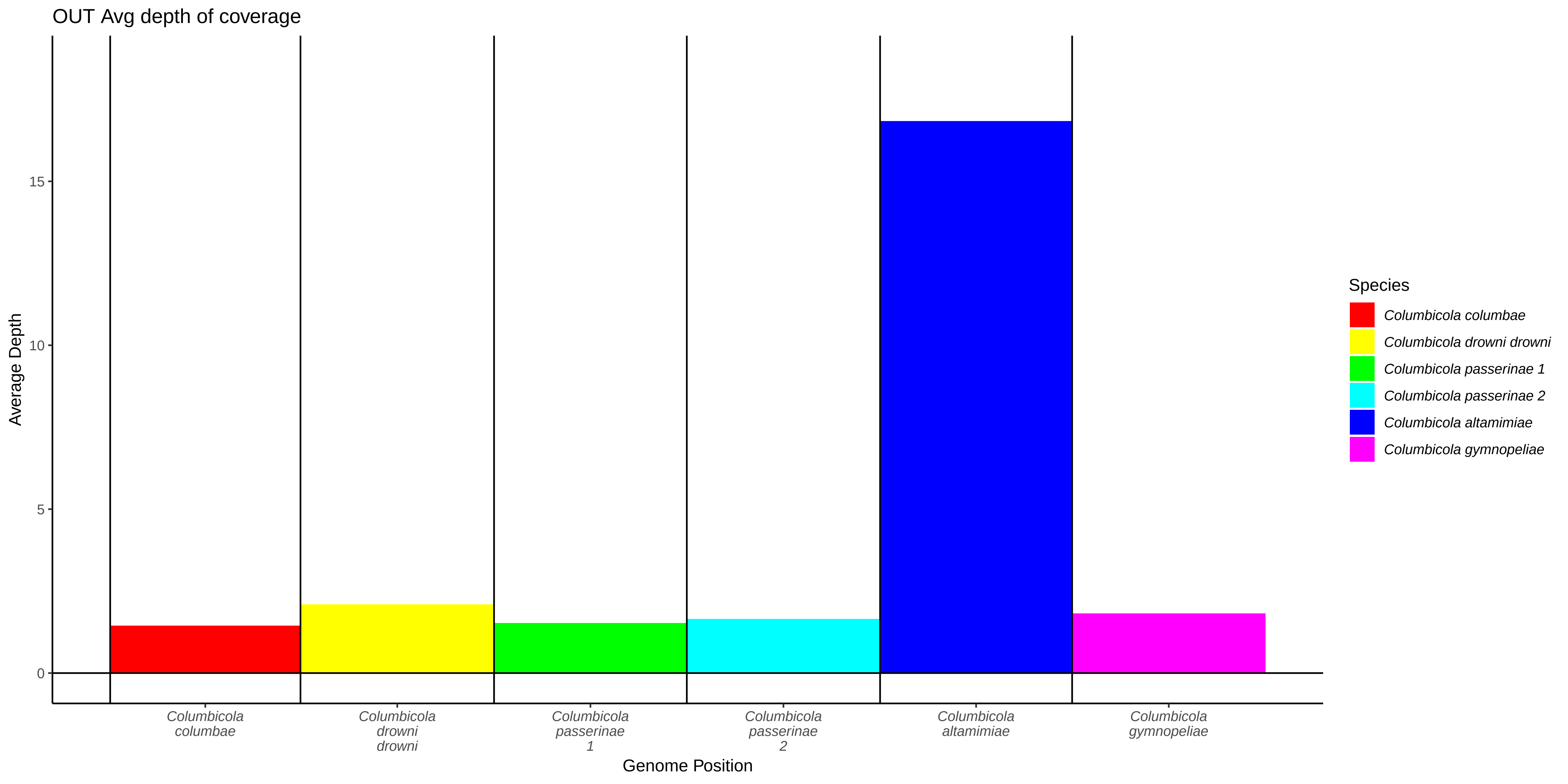

OUT log2 Mean Avg depth of coverage

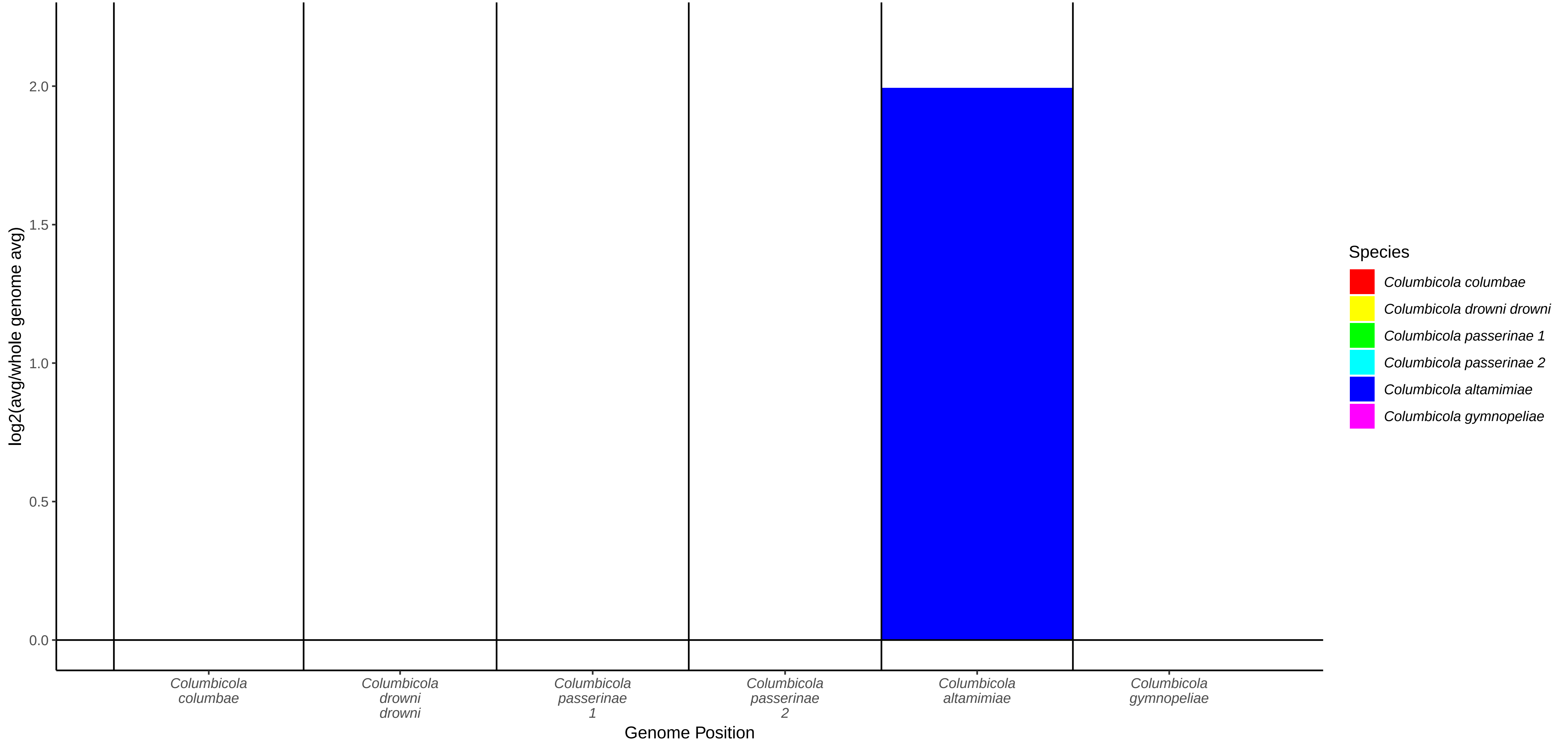

### OUT_speciesDepth_Codro.pdf

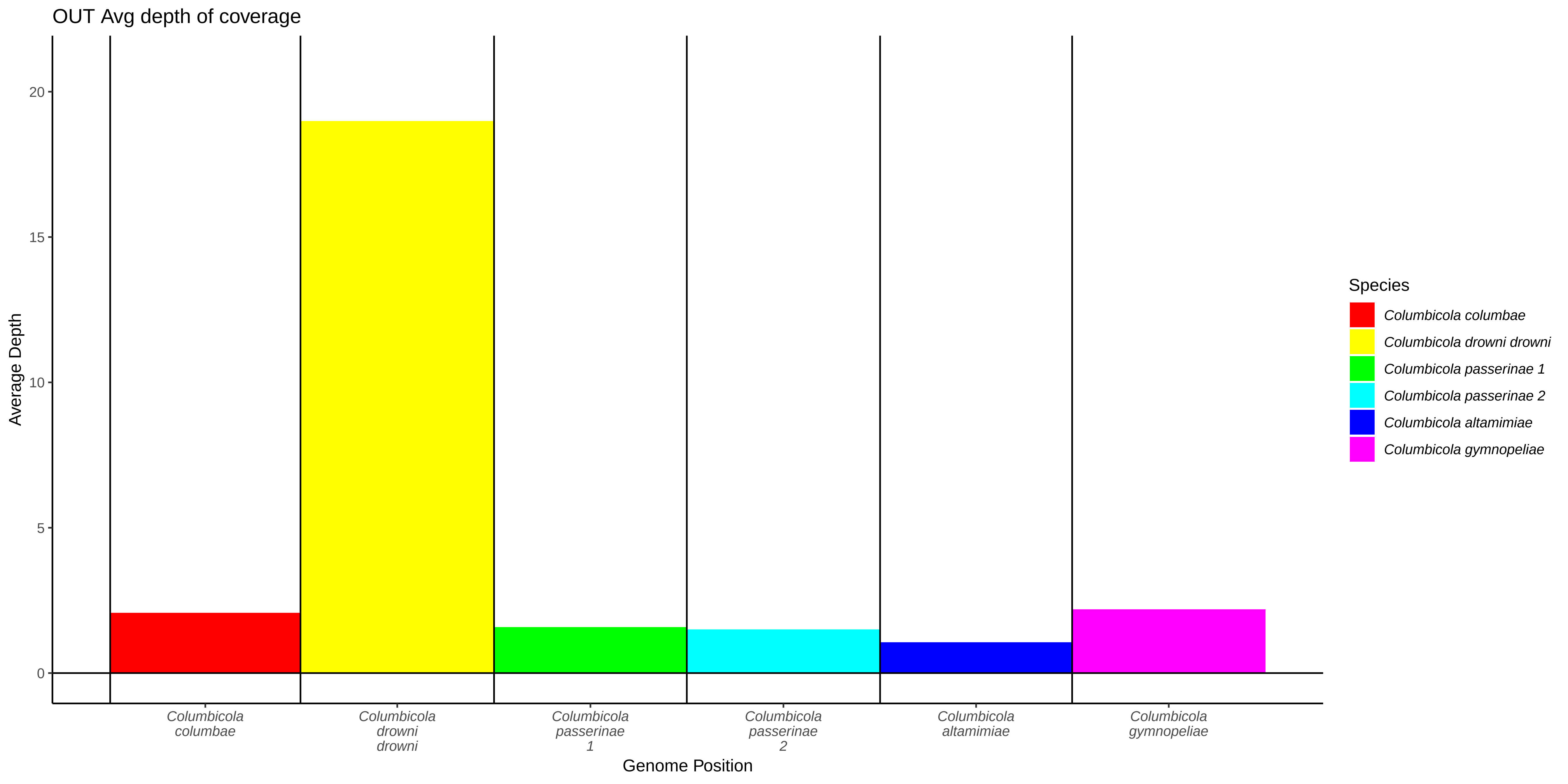

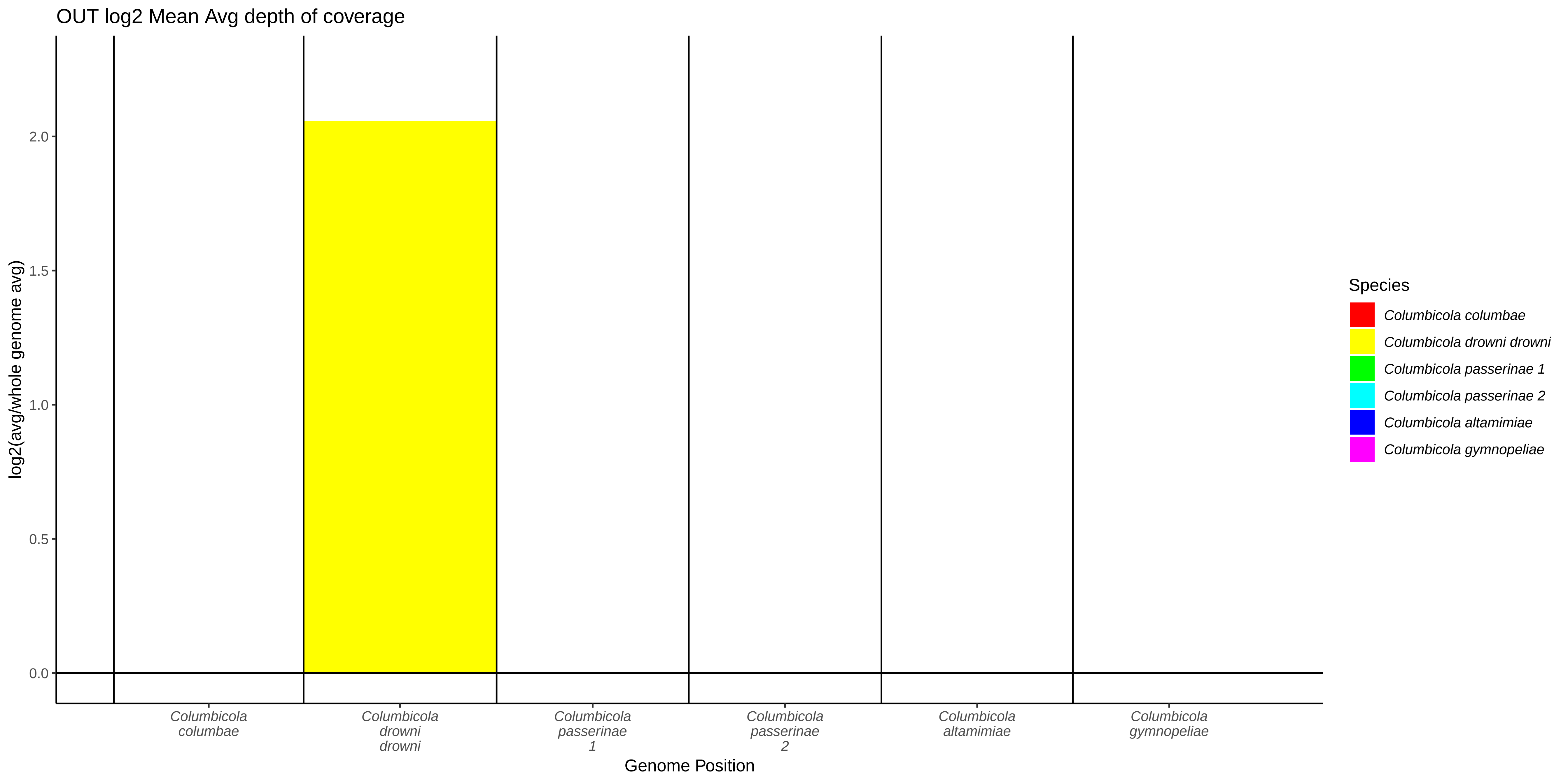

### OUT_speciesDepth_cogym.pdf

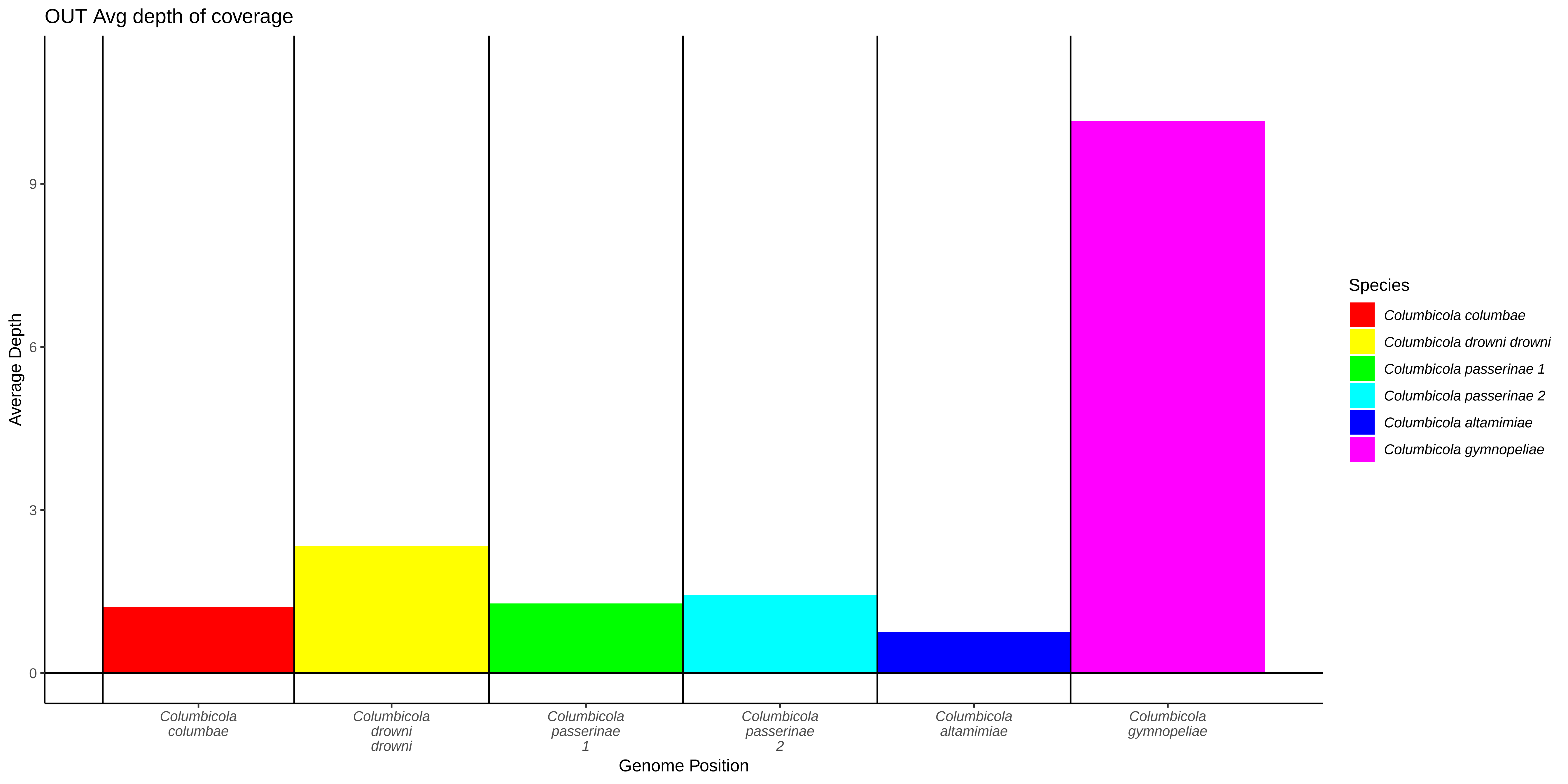

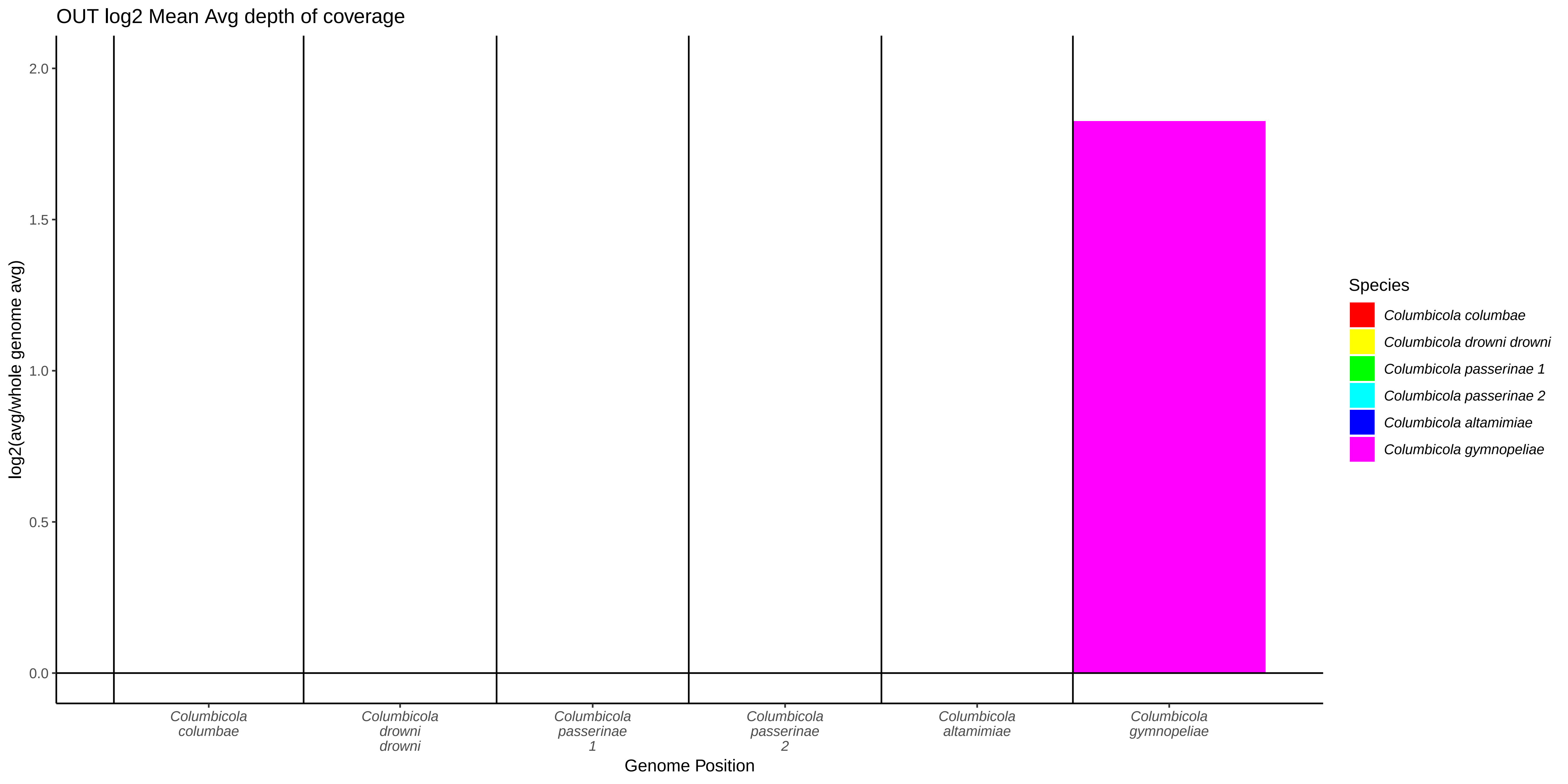

### OUT_speciesDepth_copas1.pdf

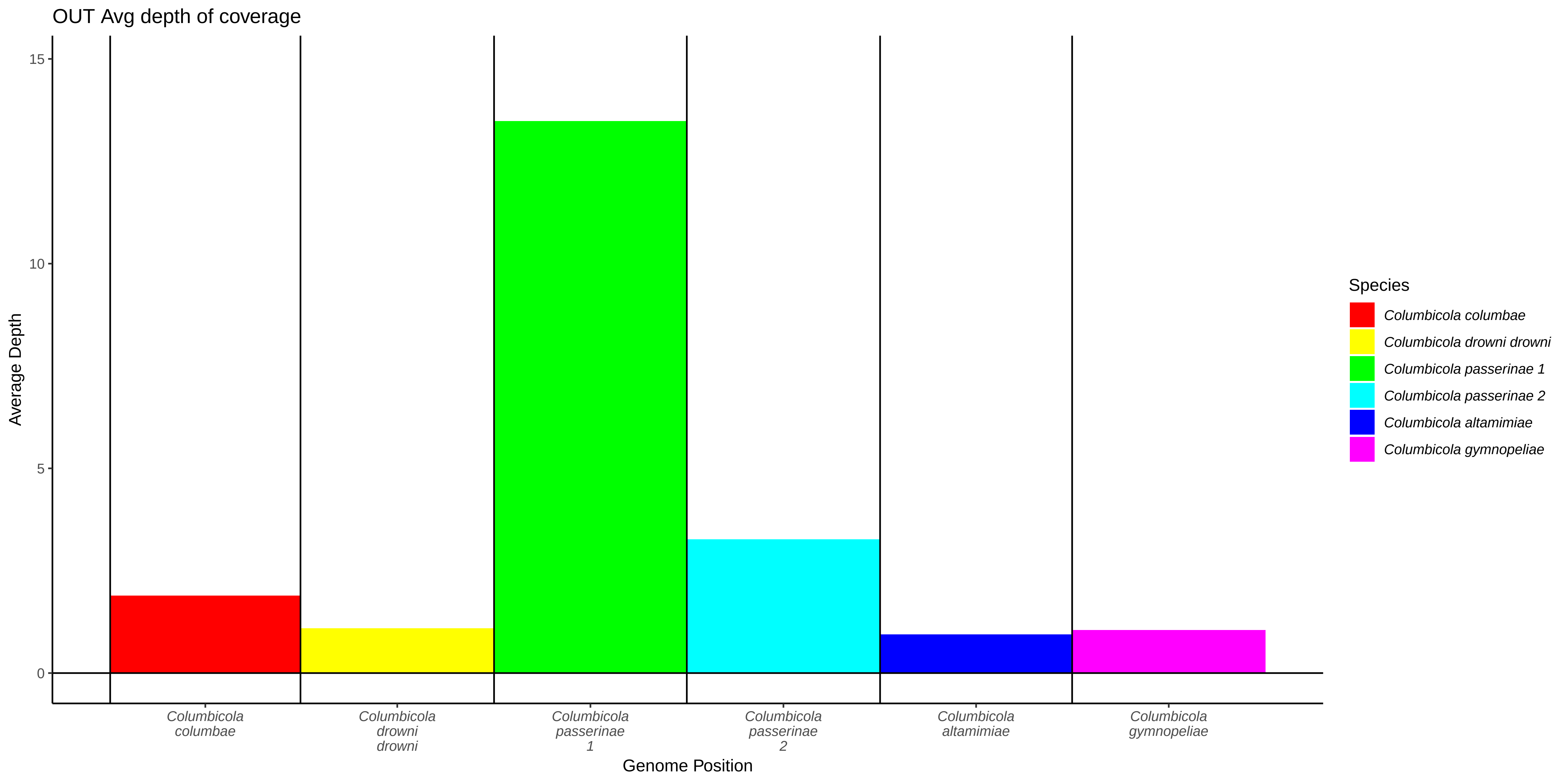

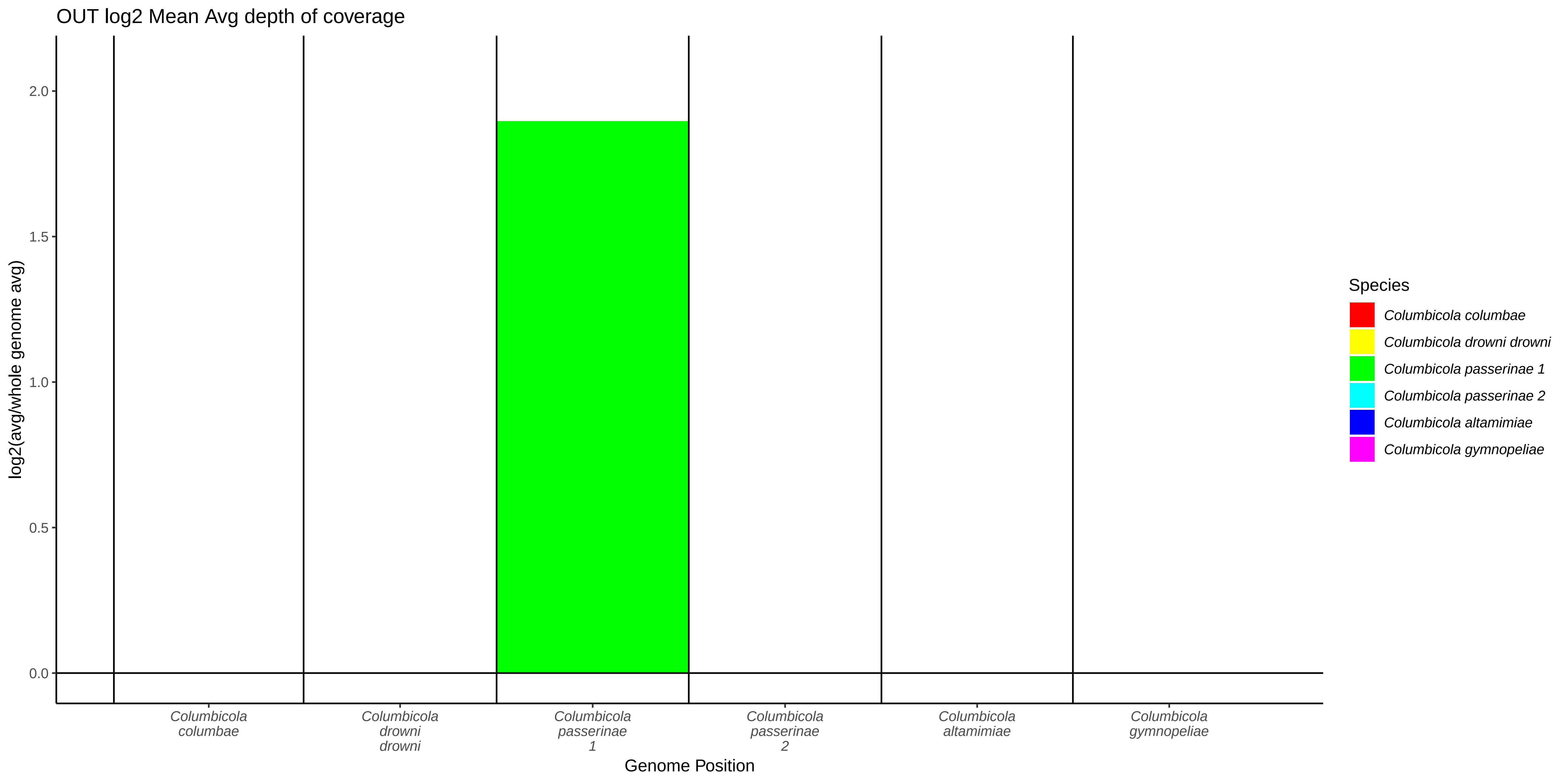

### OUT_speciesDepth_Copas2.pdf

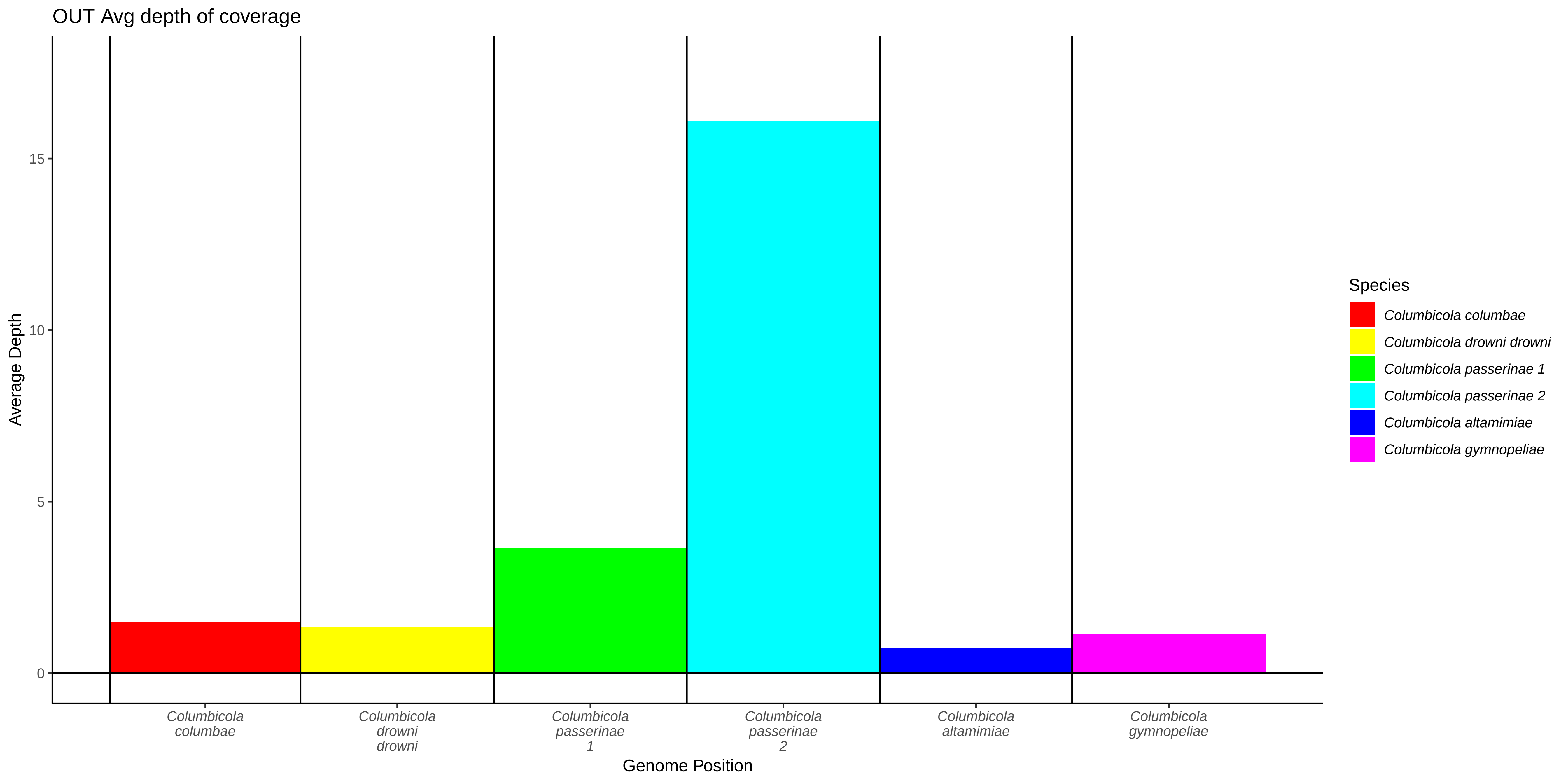

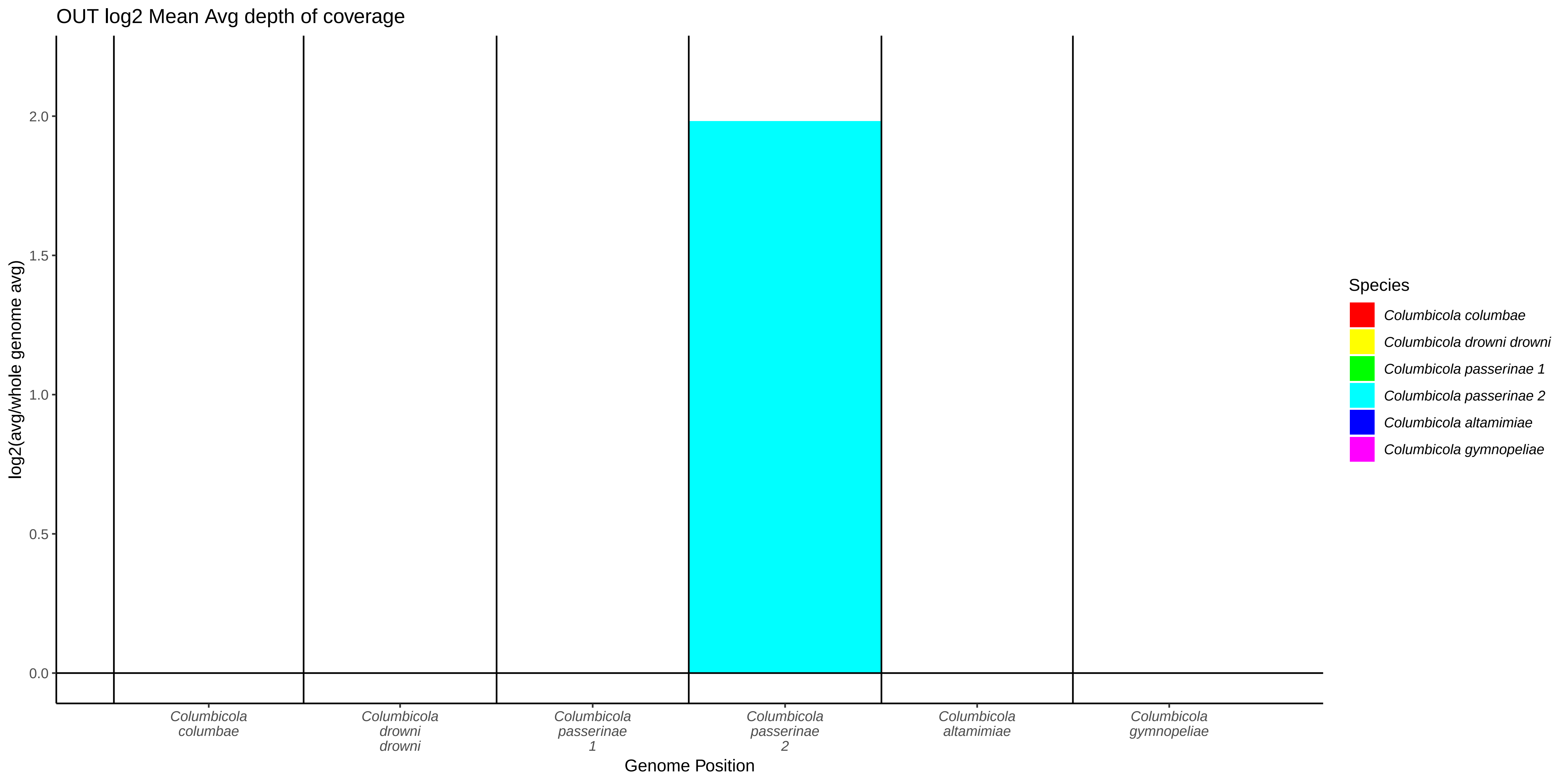

### OUT_speciesDepth_Cosp_Clpre_11_2_2016_14.pdf

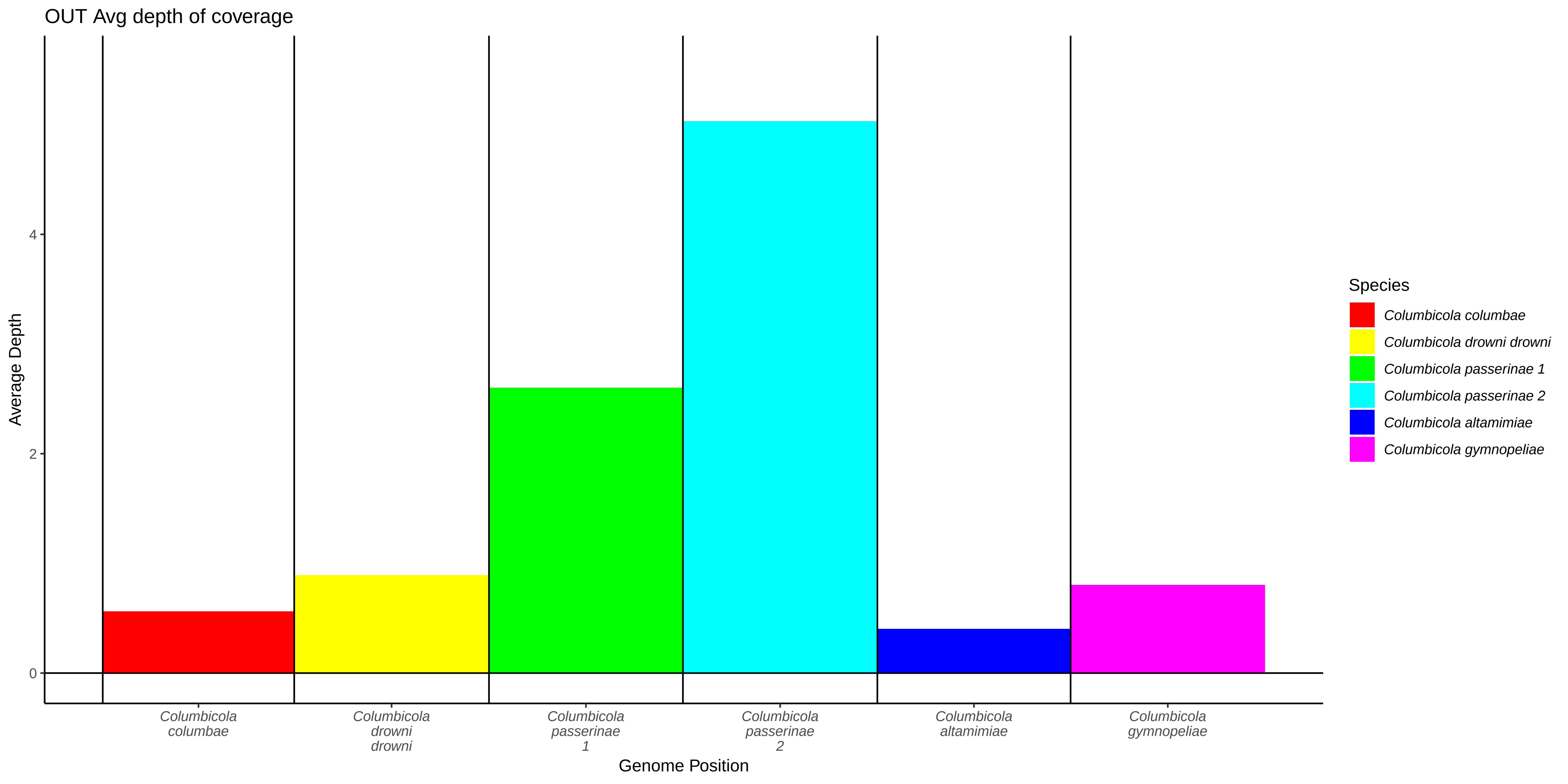

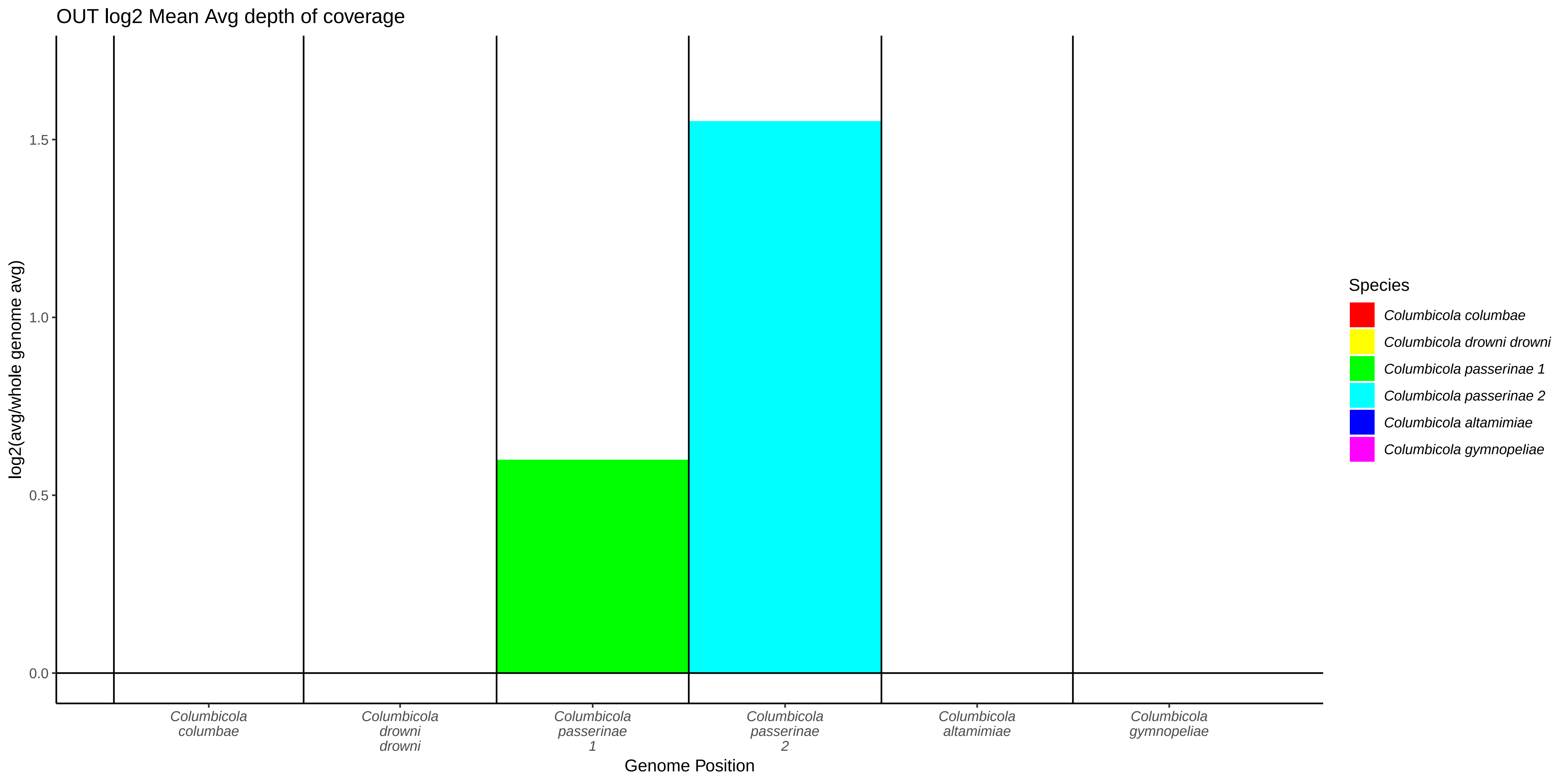

### OUT_speciesDepth_Cosp_Clpre_11_7_2016_3.pdf

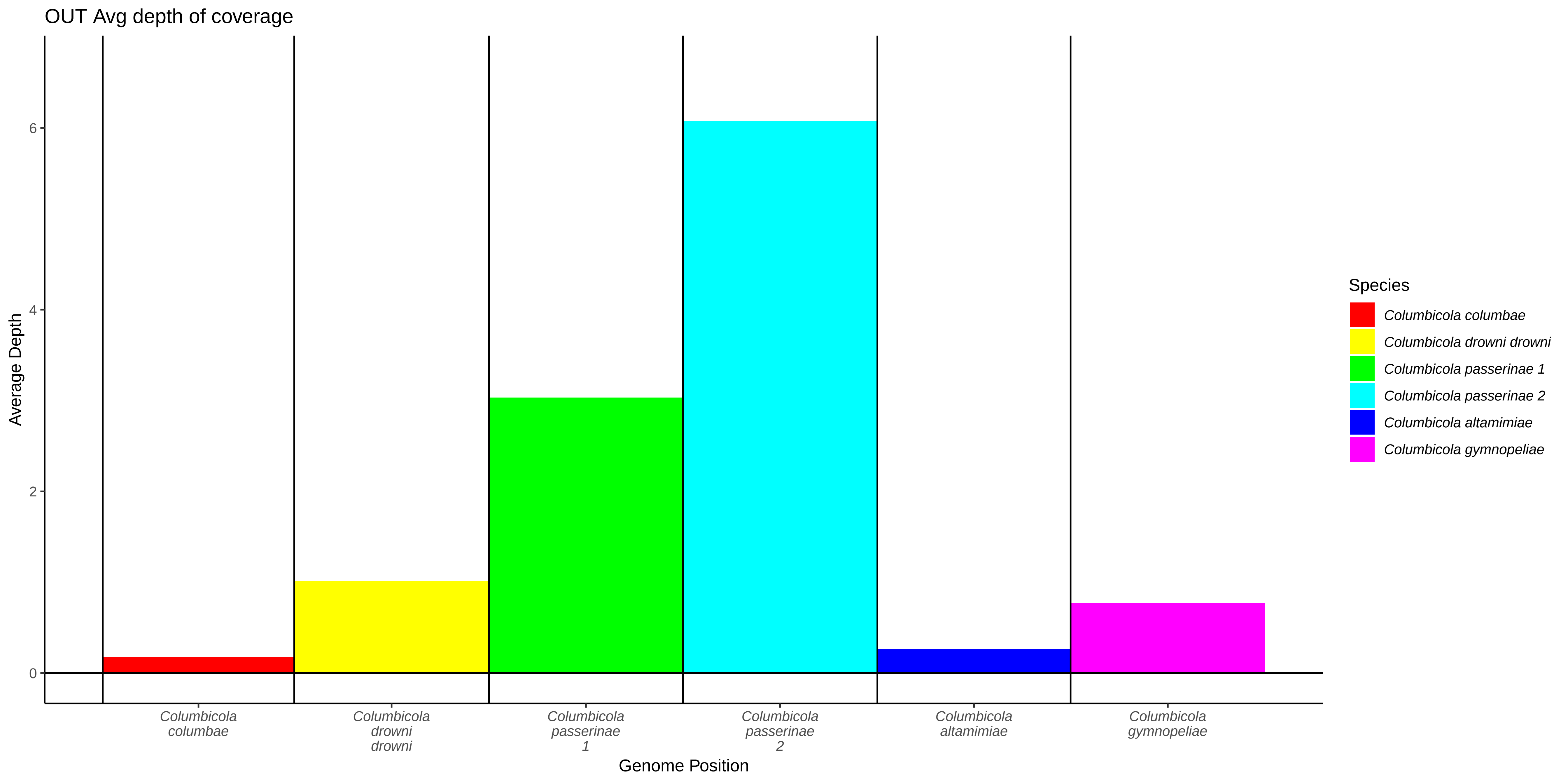

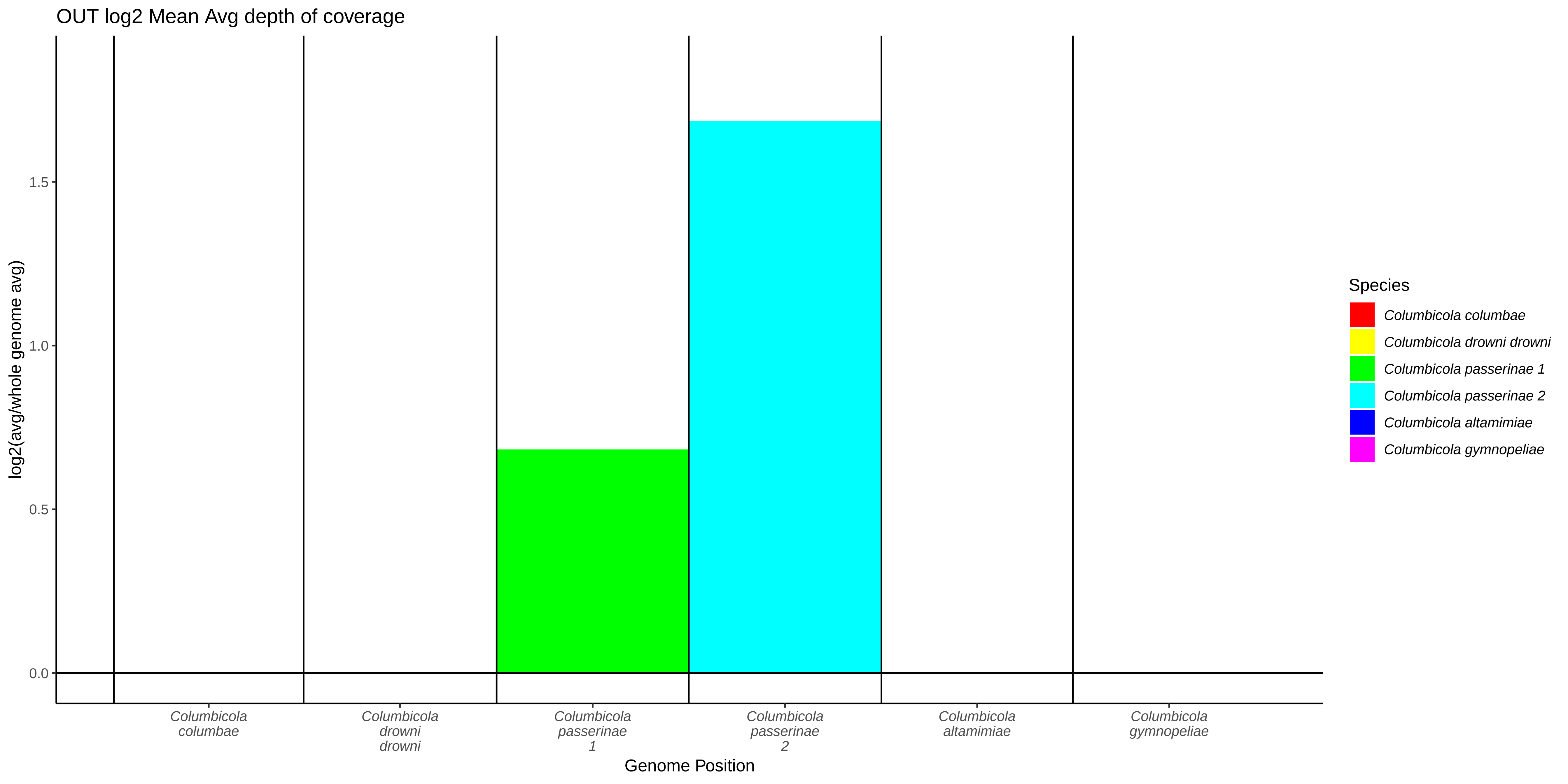

### OUT_speciesDepth_Cosp_Clpre_11_7_2016_9.pdf

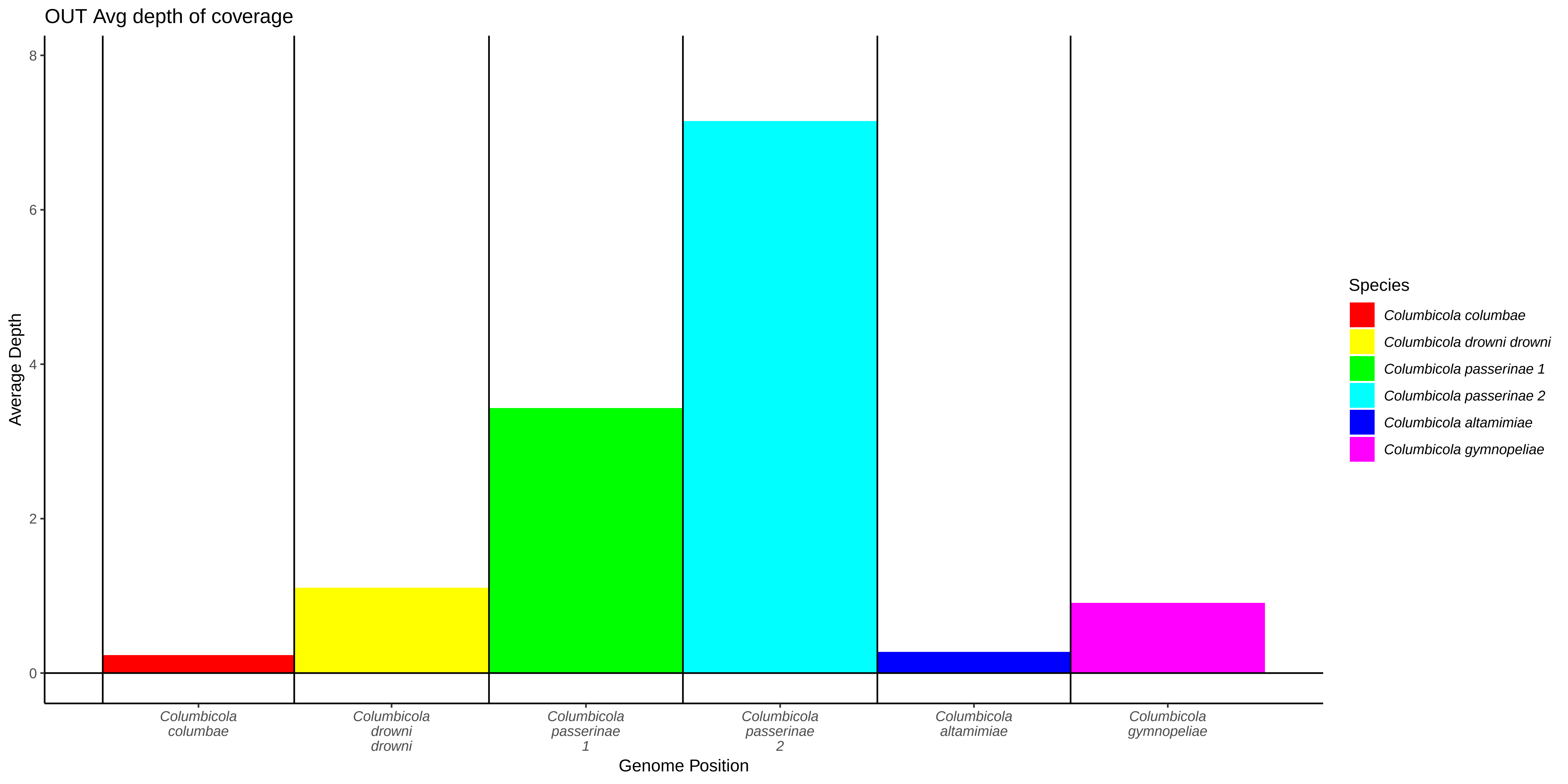

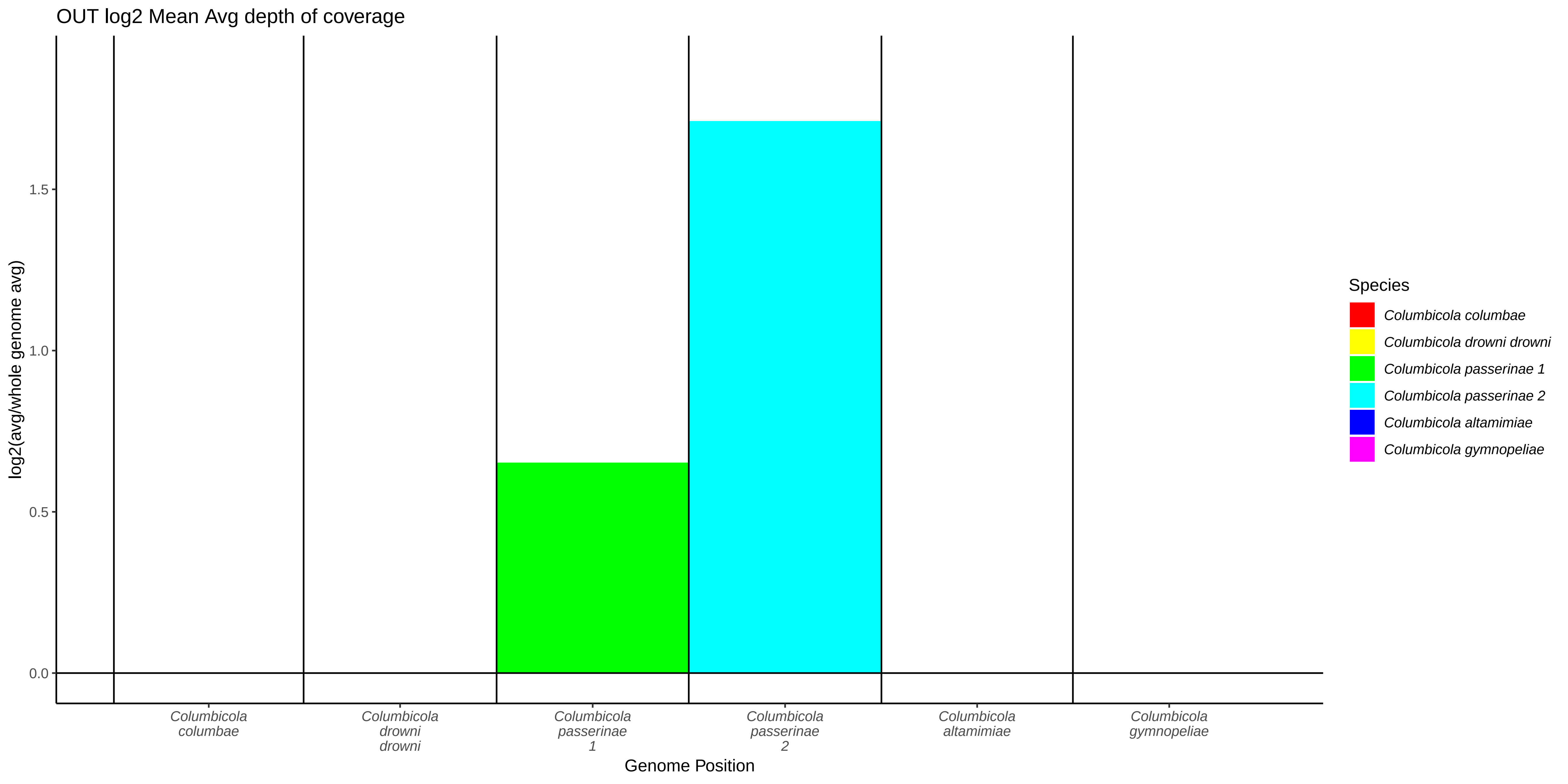

### OUT_speciesDepth_Cosp_Clpre_11_7_2016_11.pdf

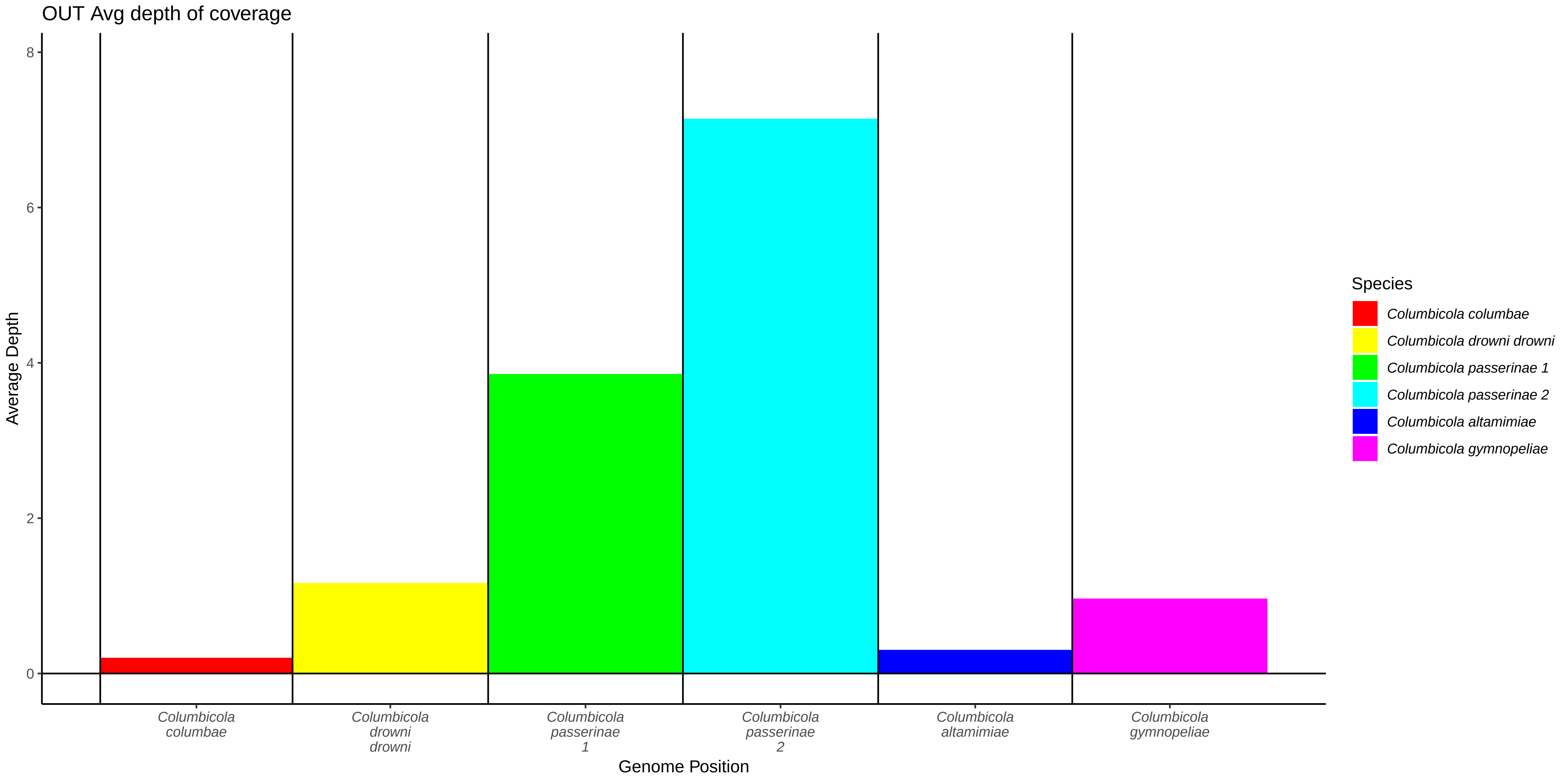

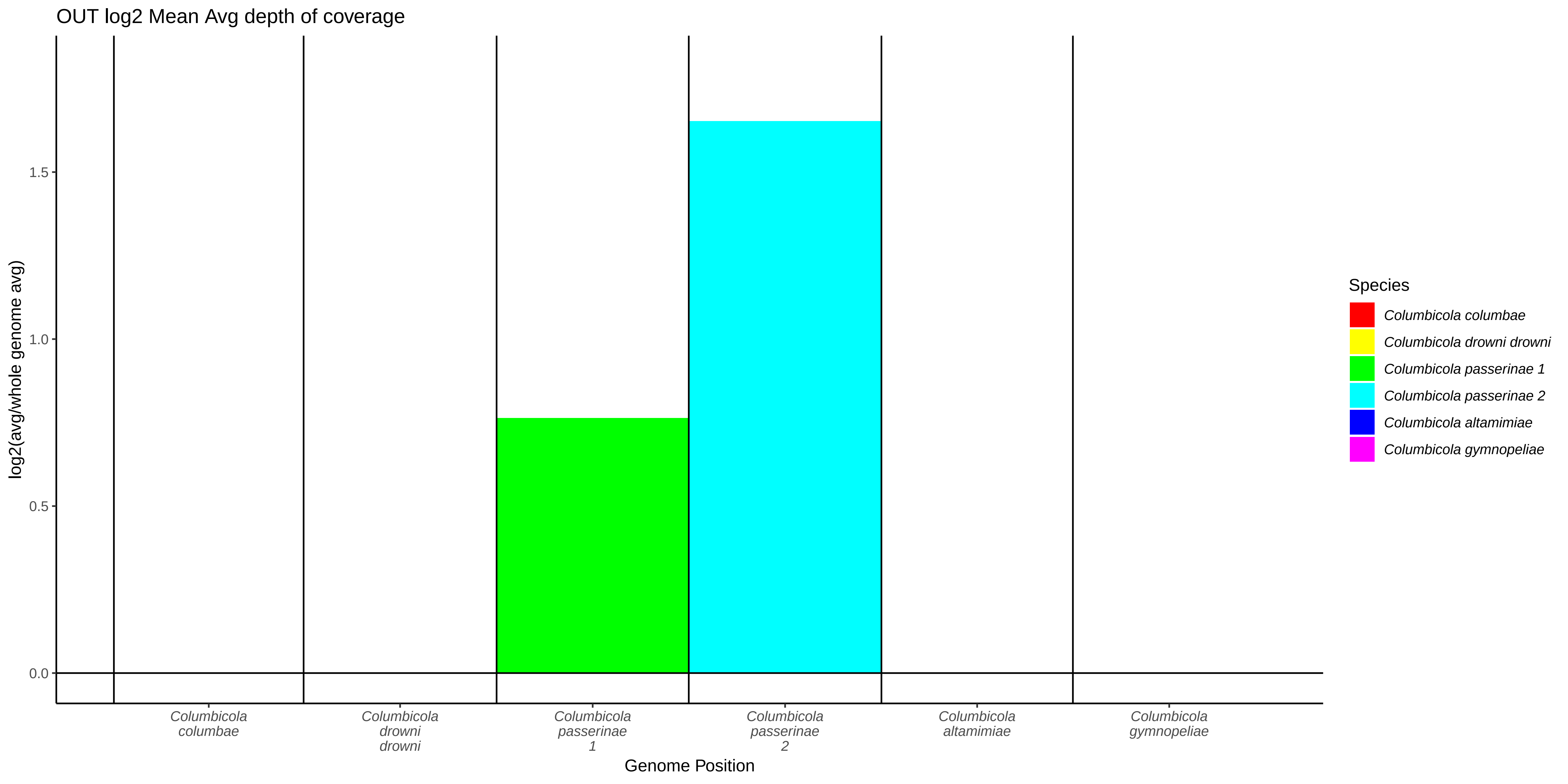

### OUT_speciesDepth_Cosp_Cobuc_11_2_2016_2.pdf

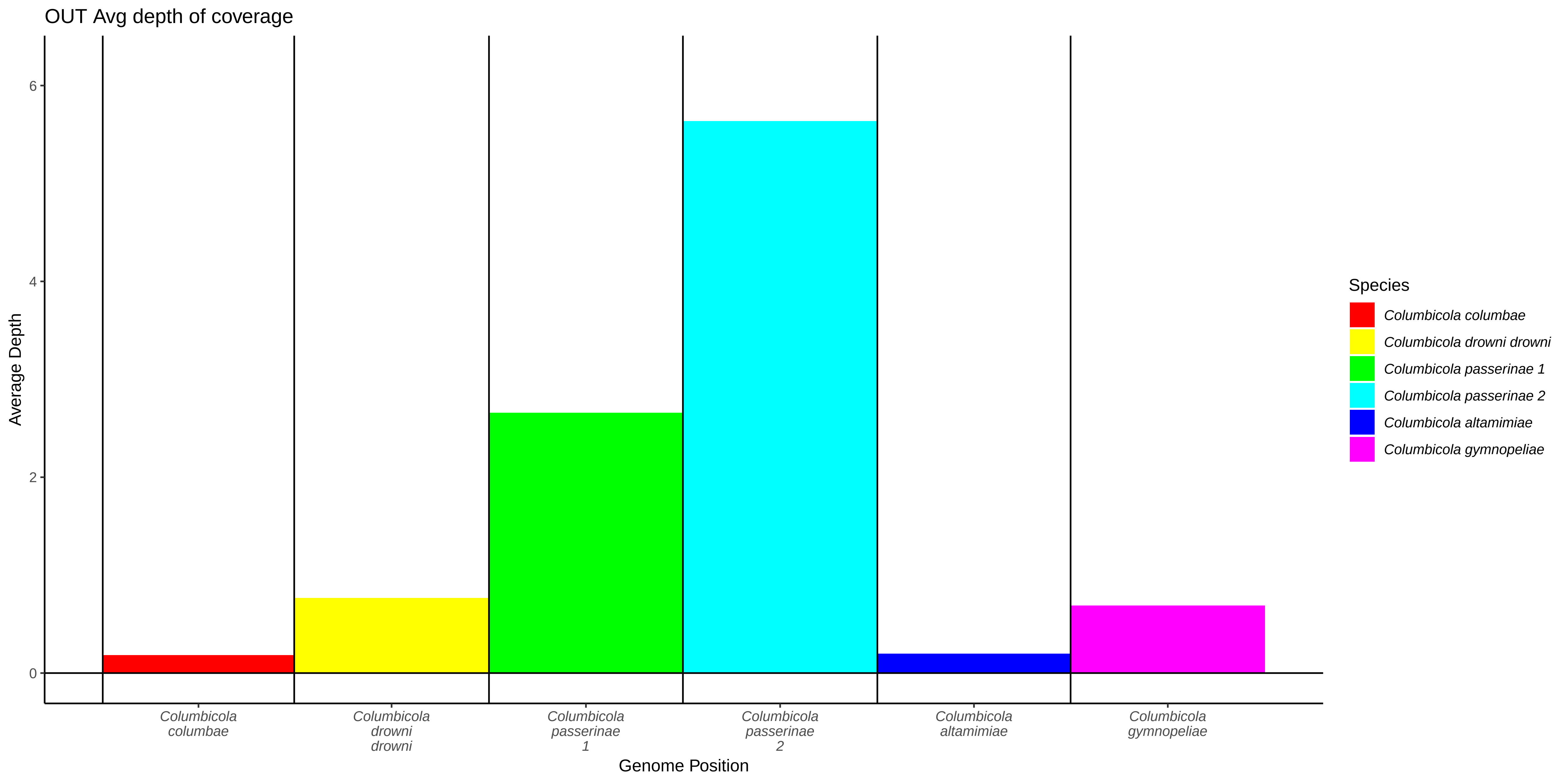

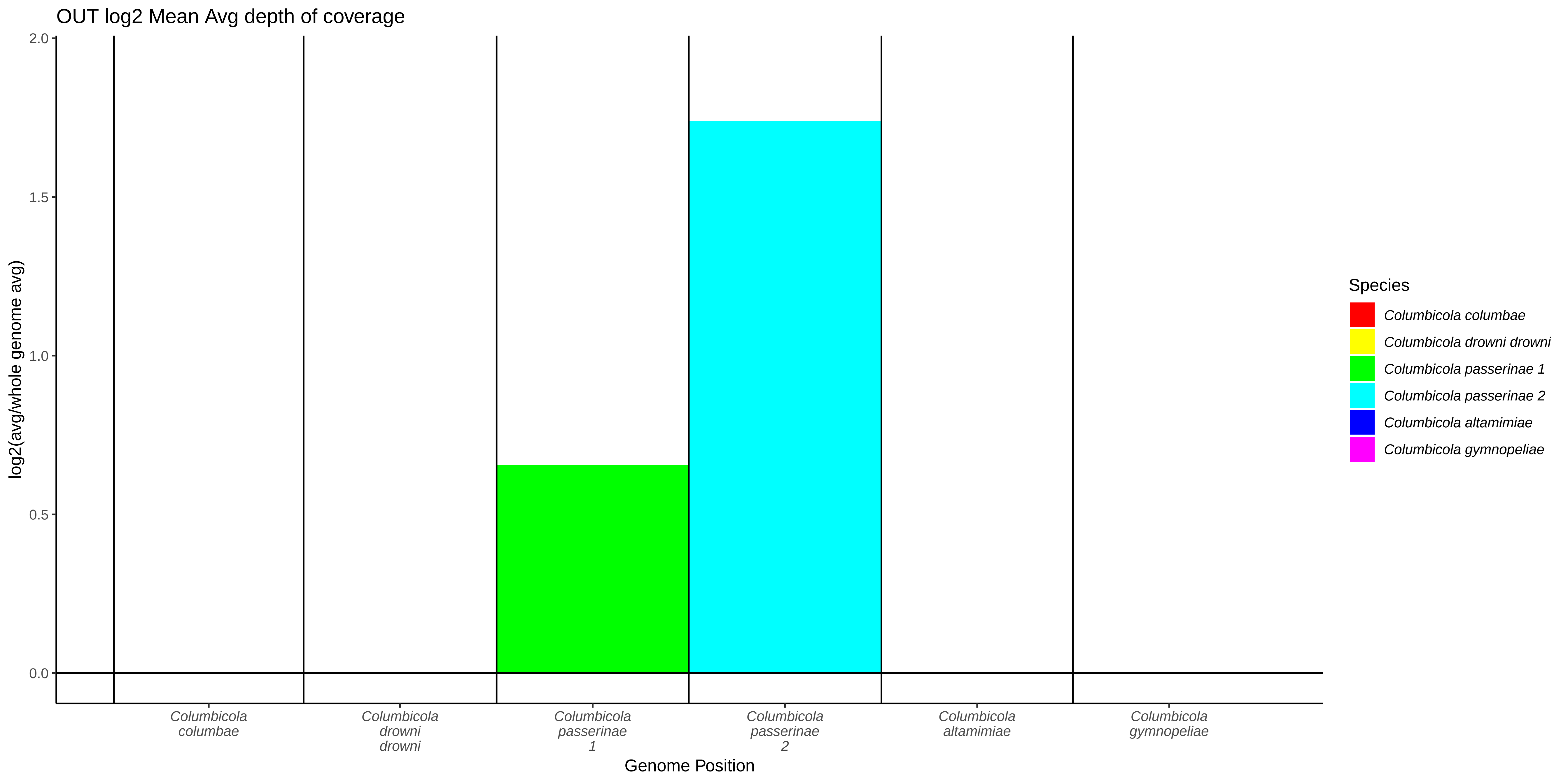

### OUT_speciesDepth_Cosp_Cobuc_11_2_2016_13.pdf

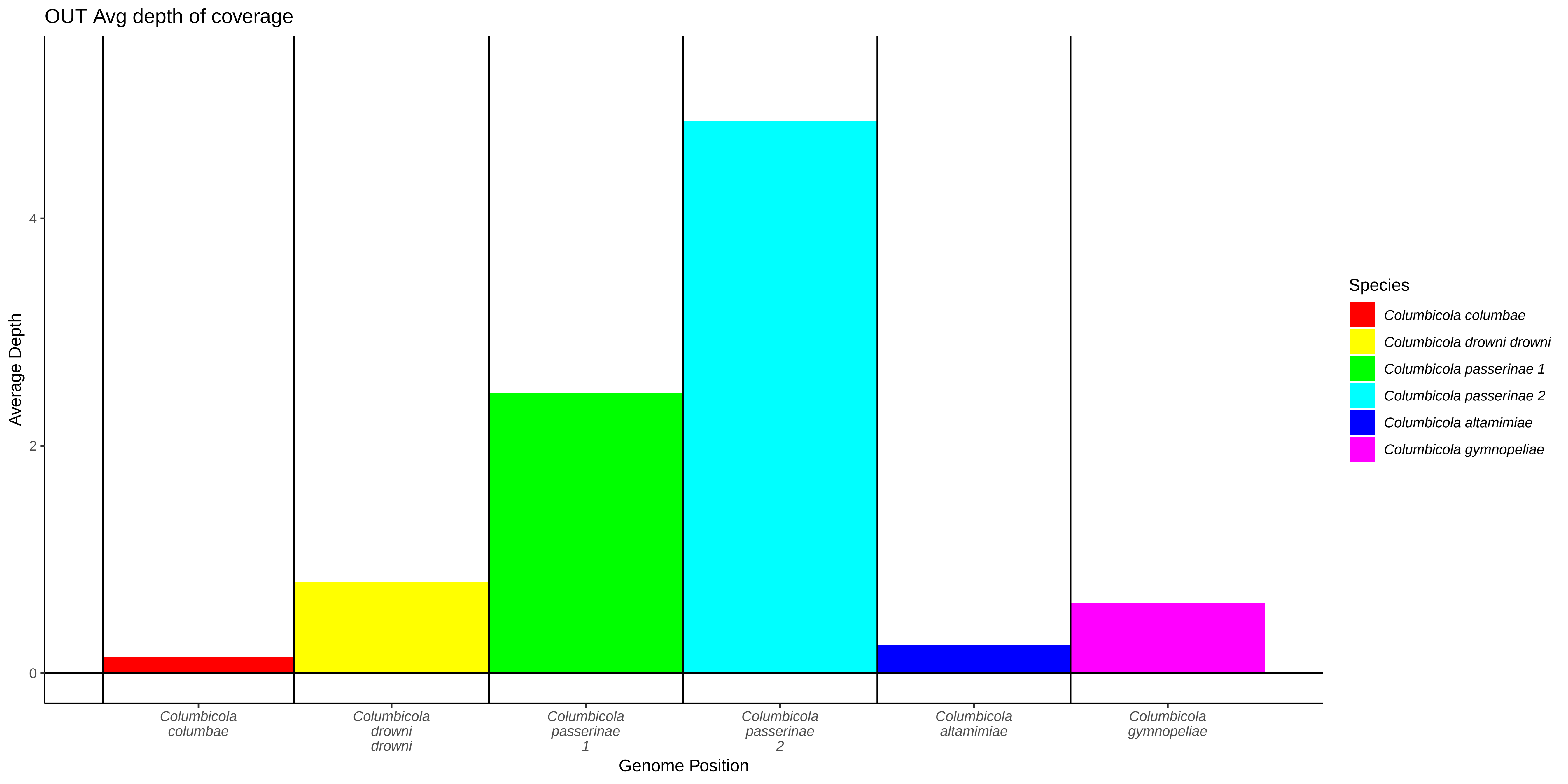

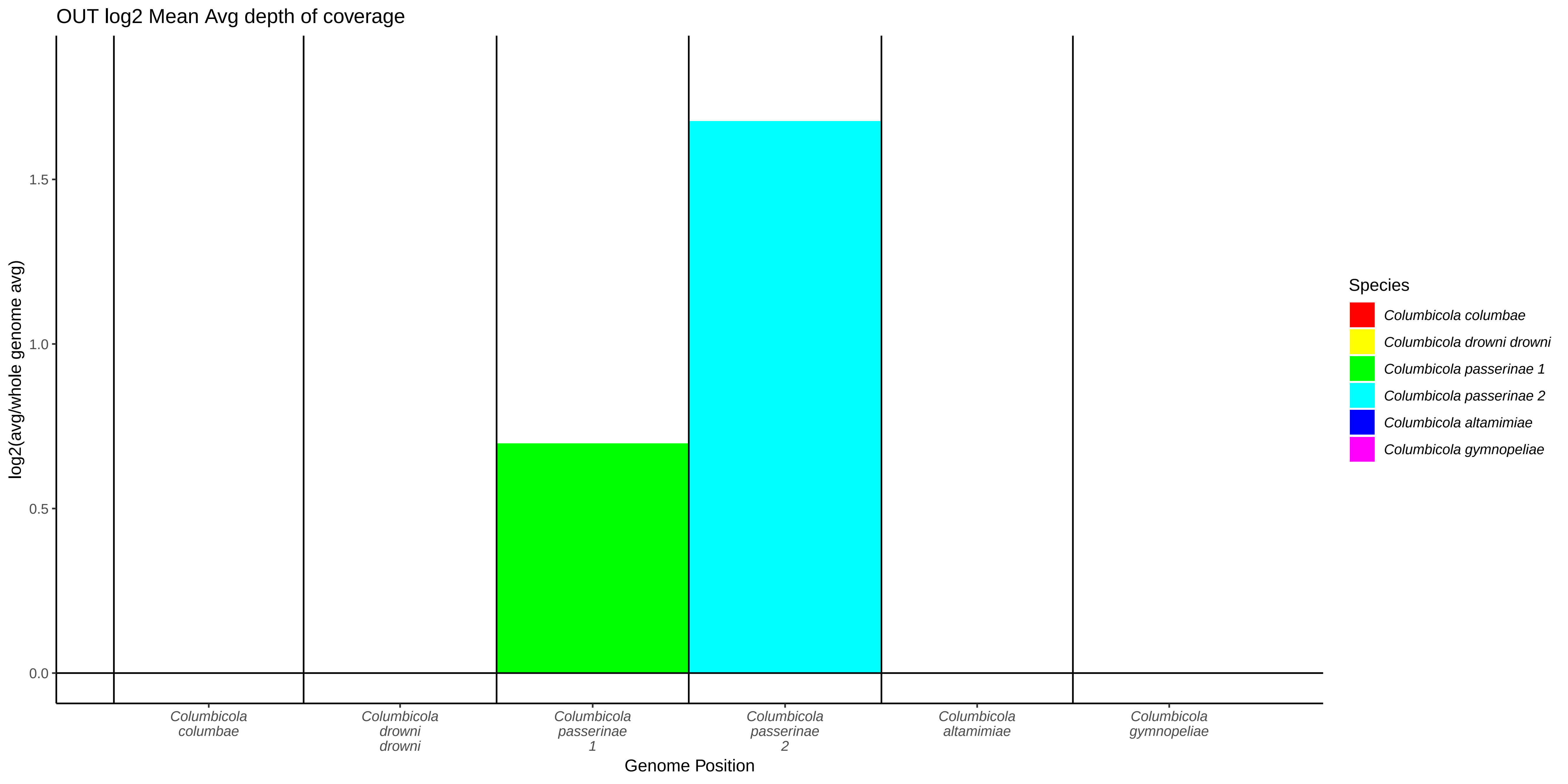

### OUT_speciesDepth_Cosp_Cocru_11_2_2016_11.pdf

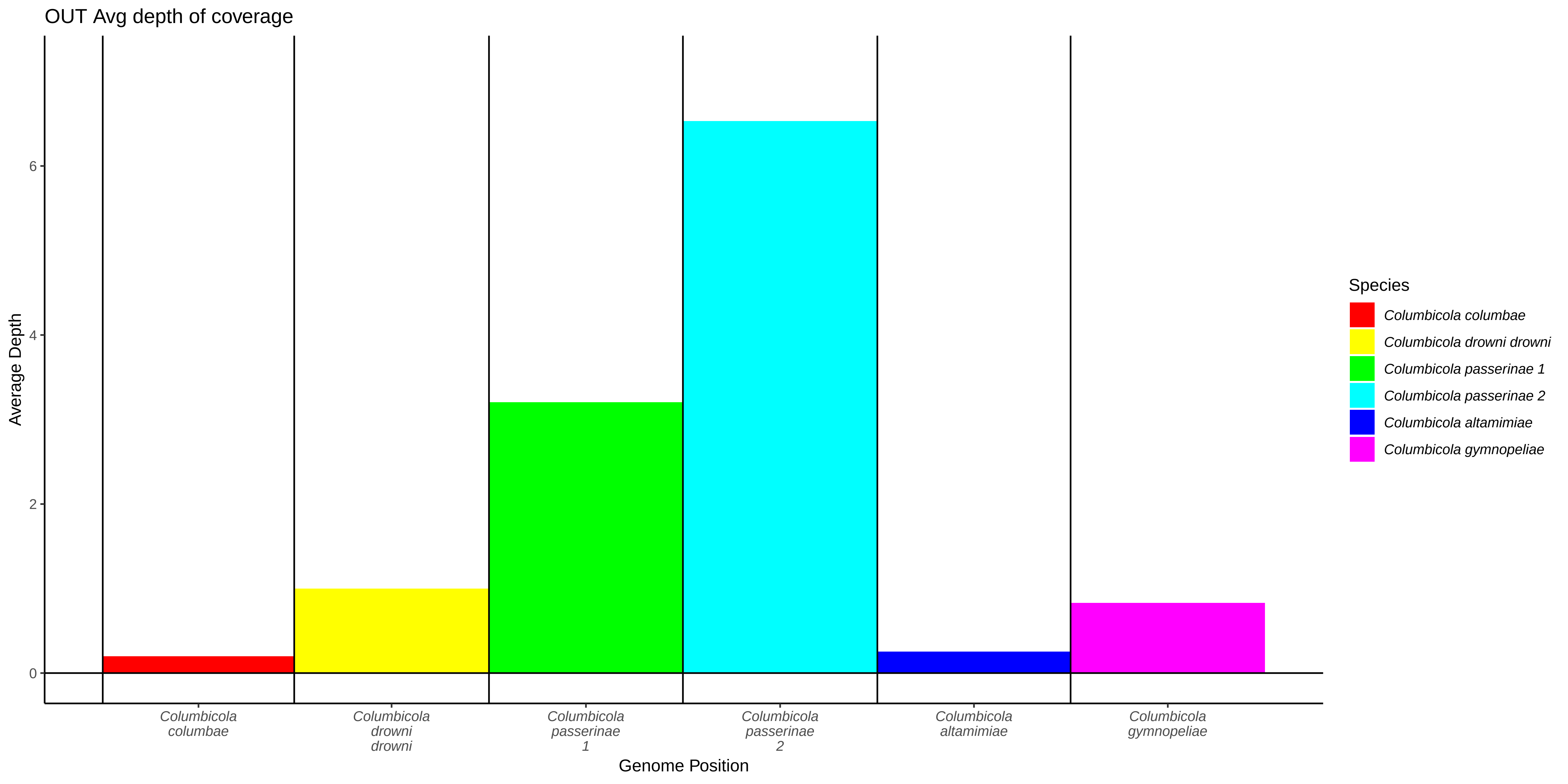

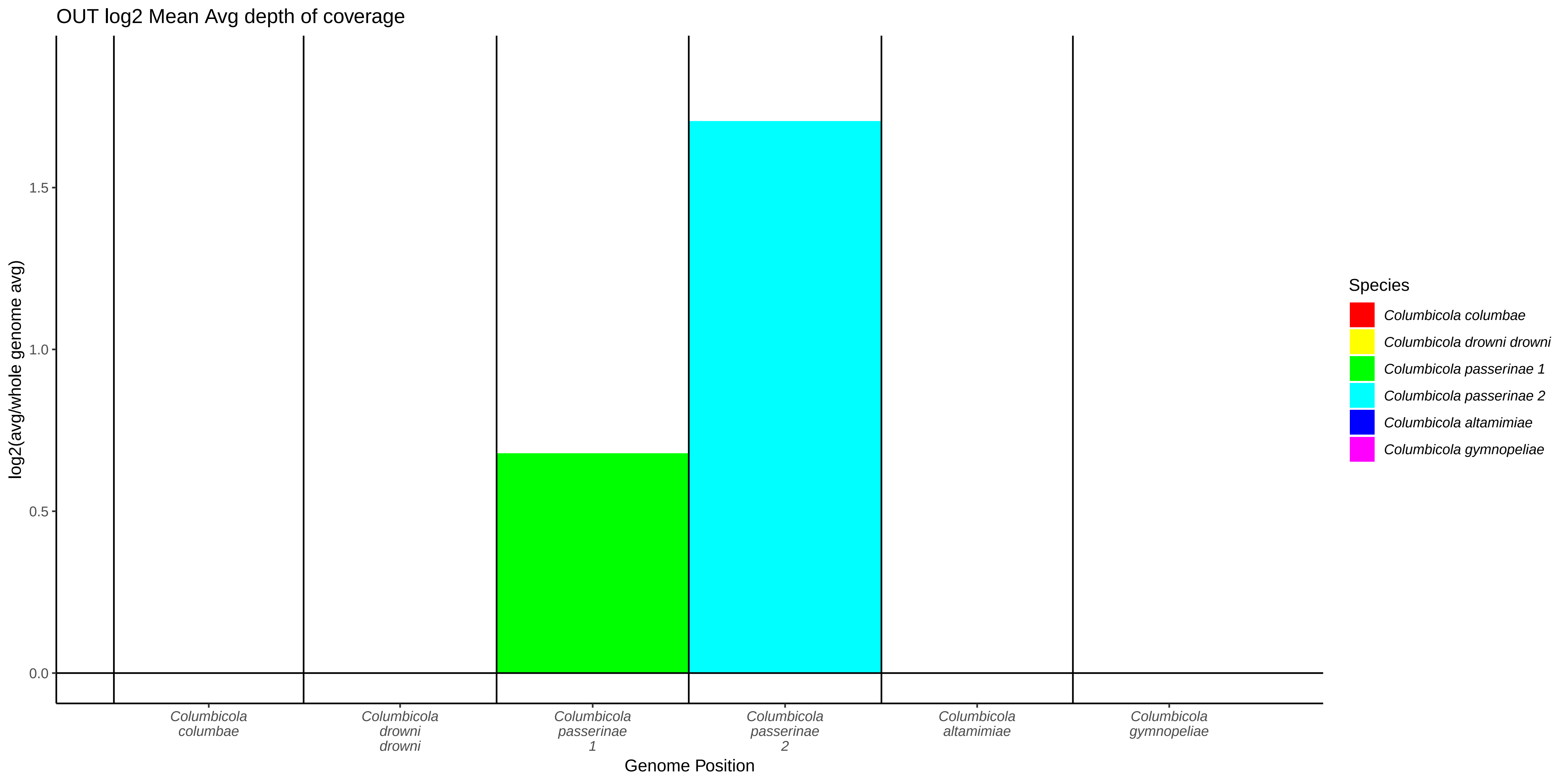

### OUT_speciesDepth_Cosp_Cocru_11_7_2016_16.pdf

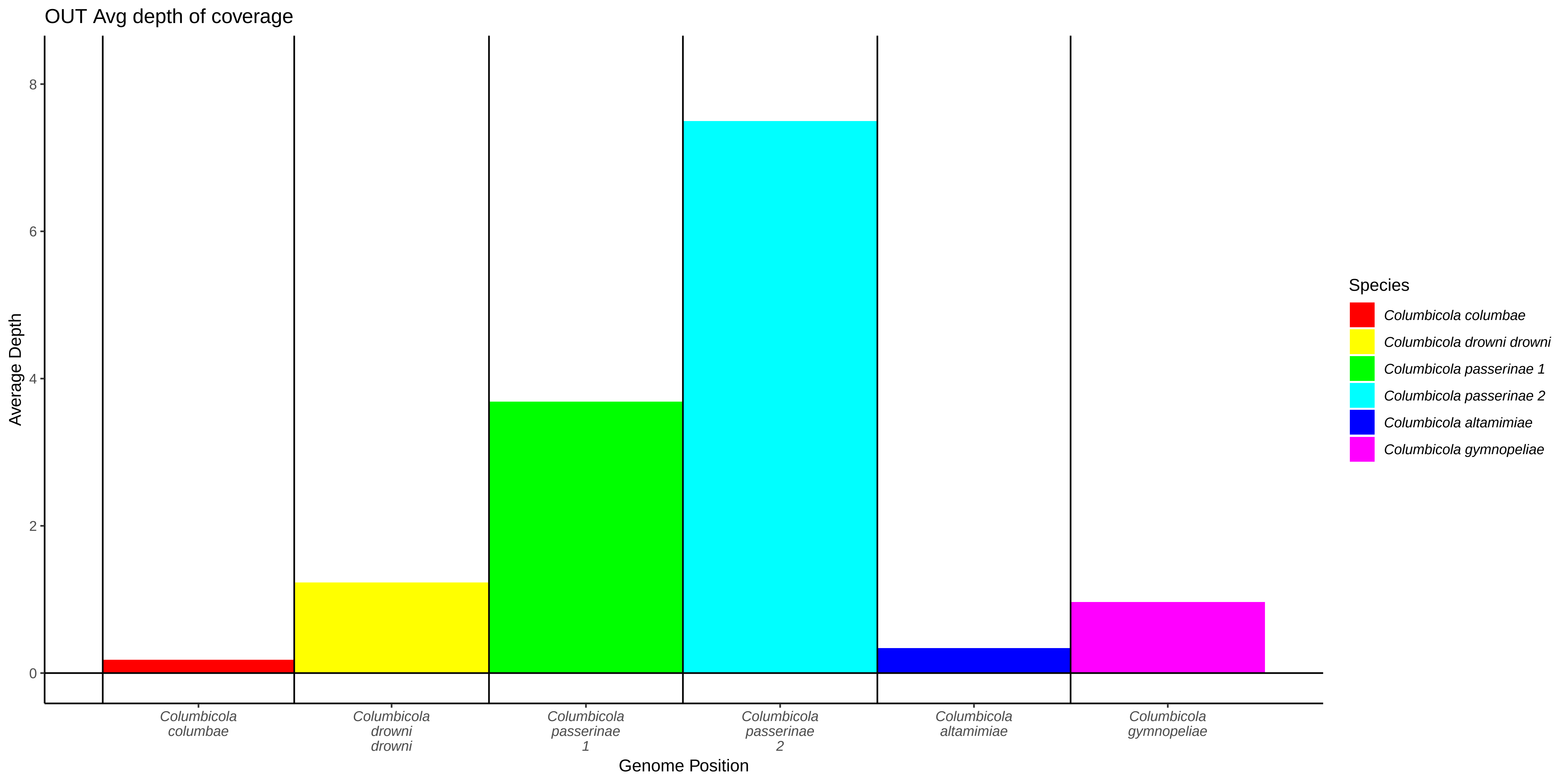

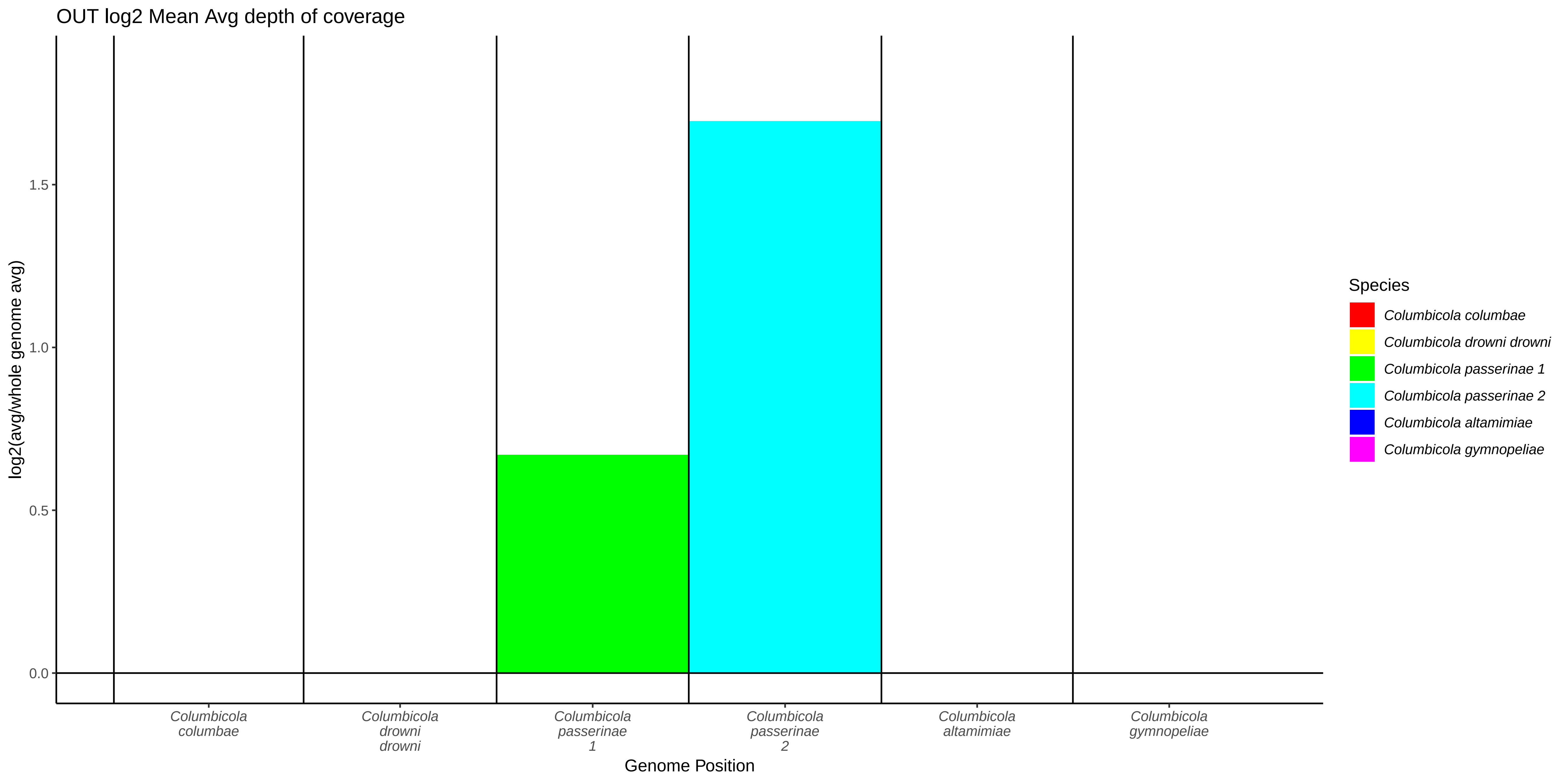

### OUT_speciesDepth_Cosp_Coinc_11_2_2016_9.pdf

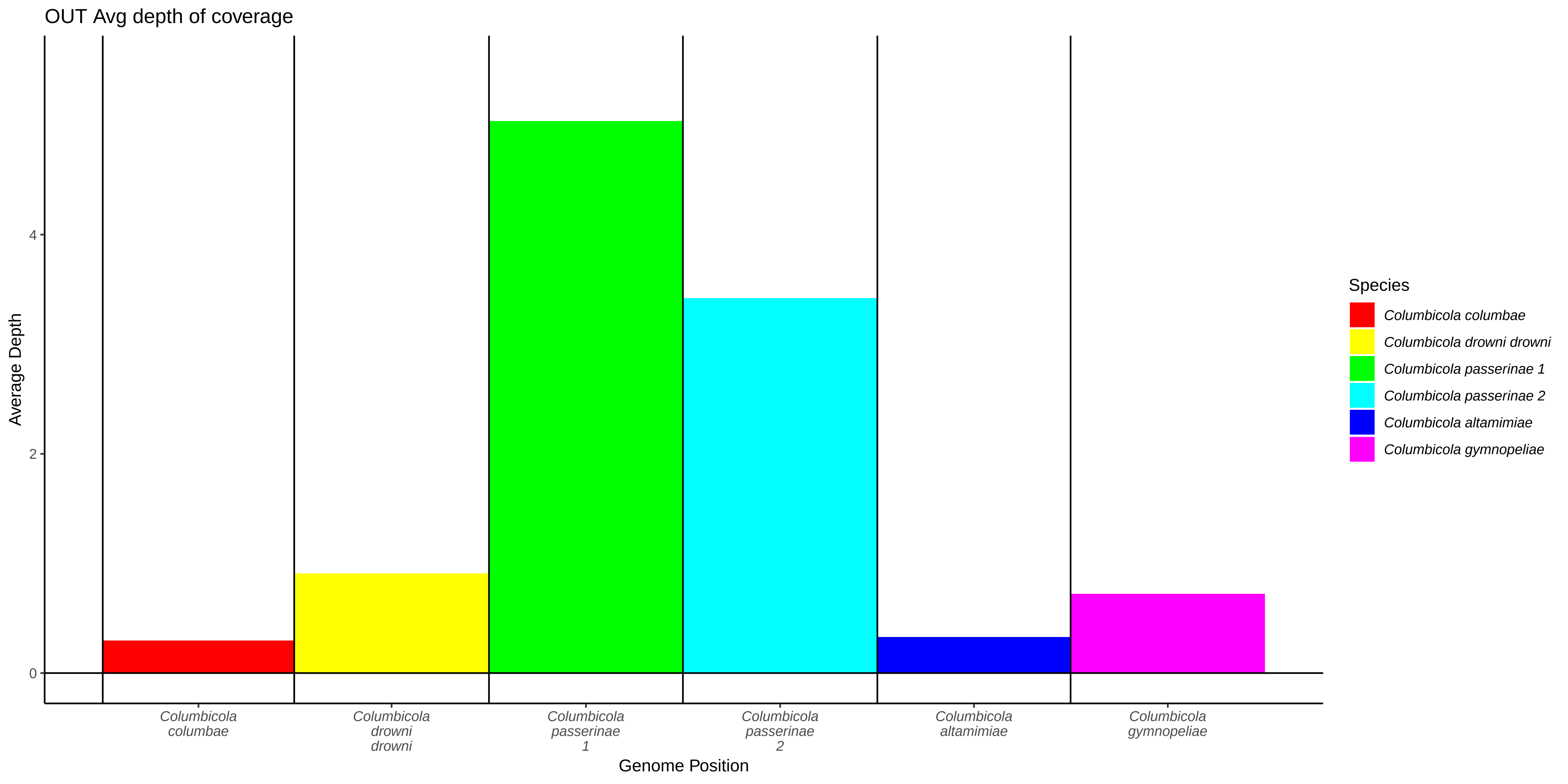

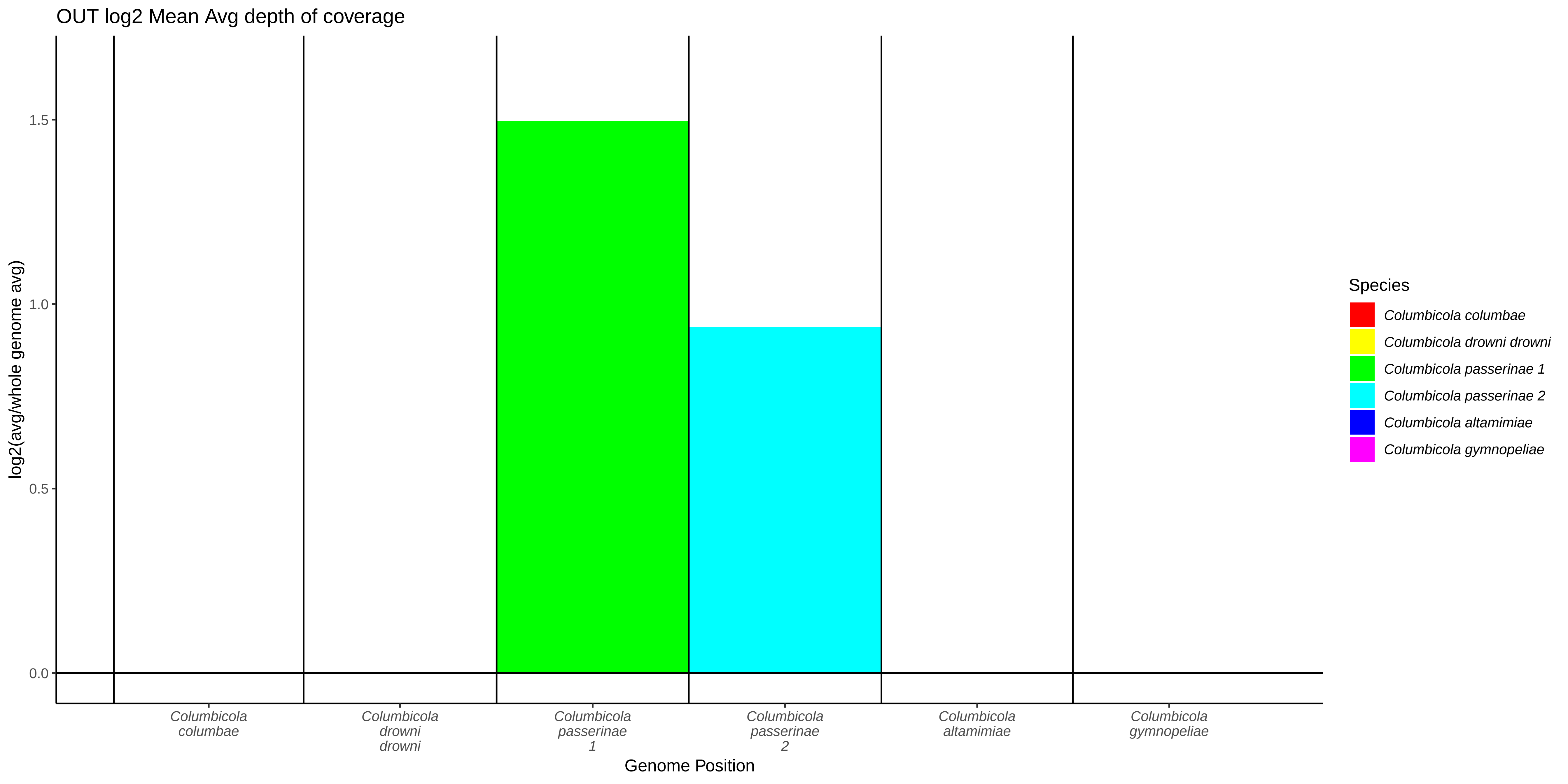

### OUT_speciesDepth_Cosp_Comin_11_2_2016_4.pdf

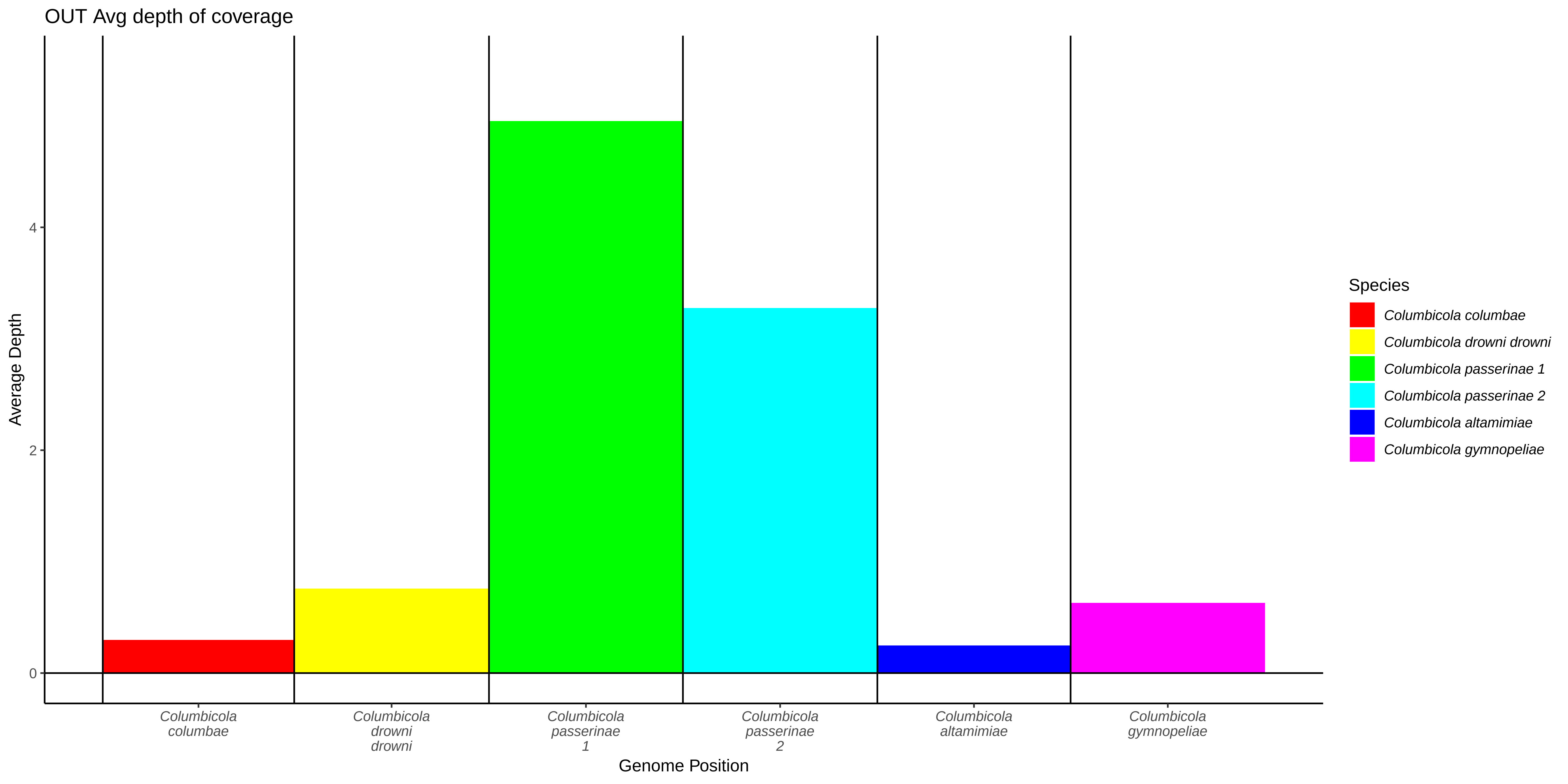

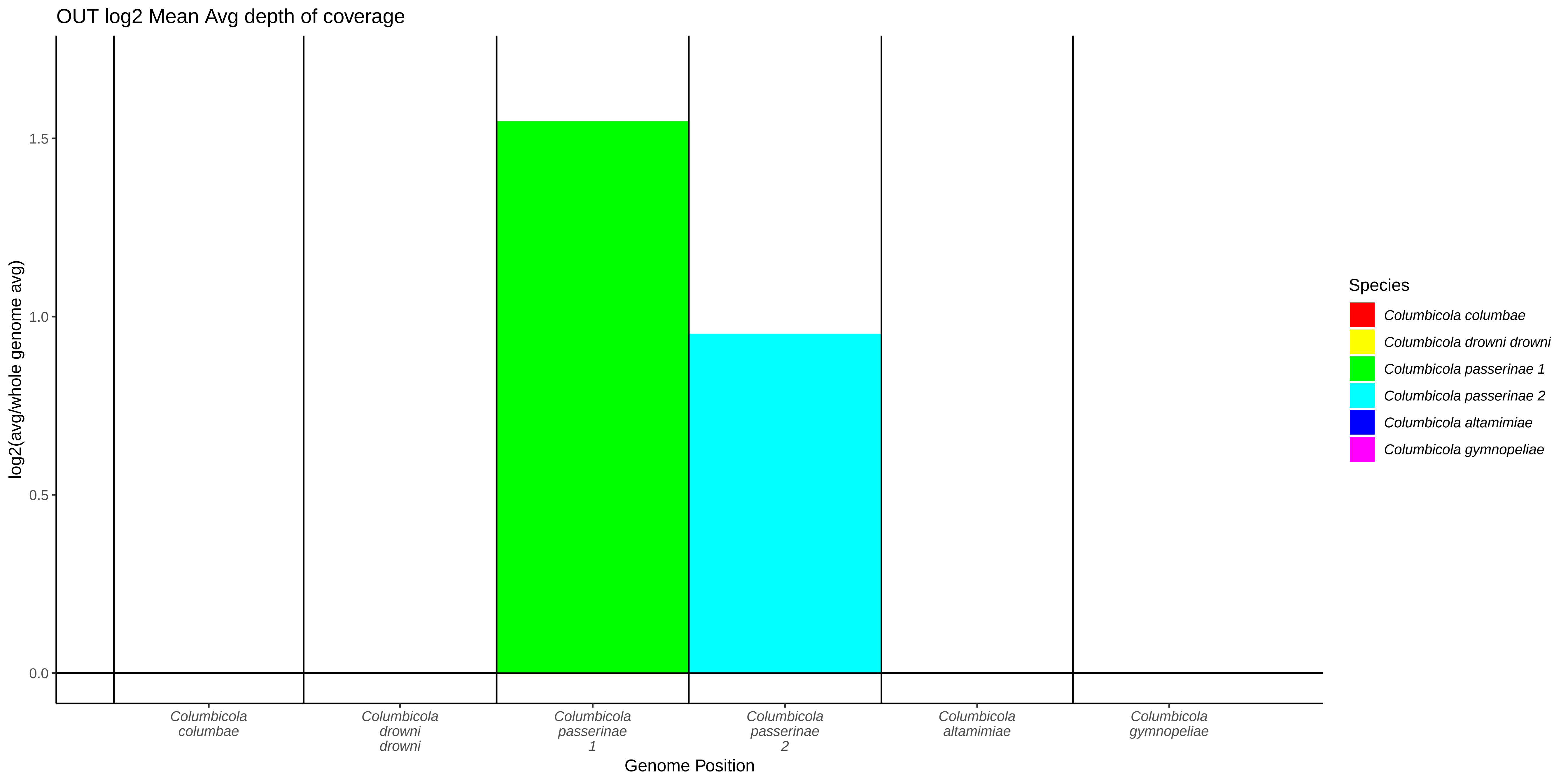

### OUT_speciesDepth_Cosp_Comin_11_2_2016_16.pdf

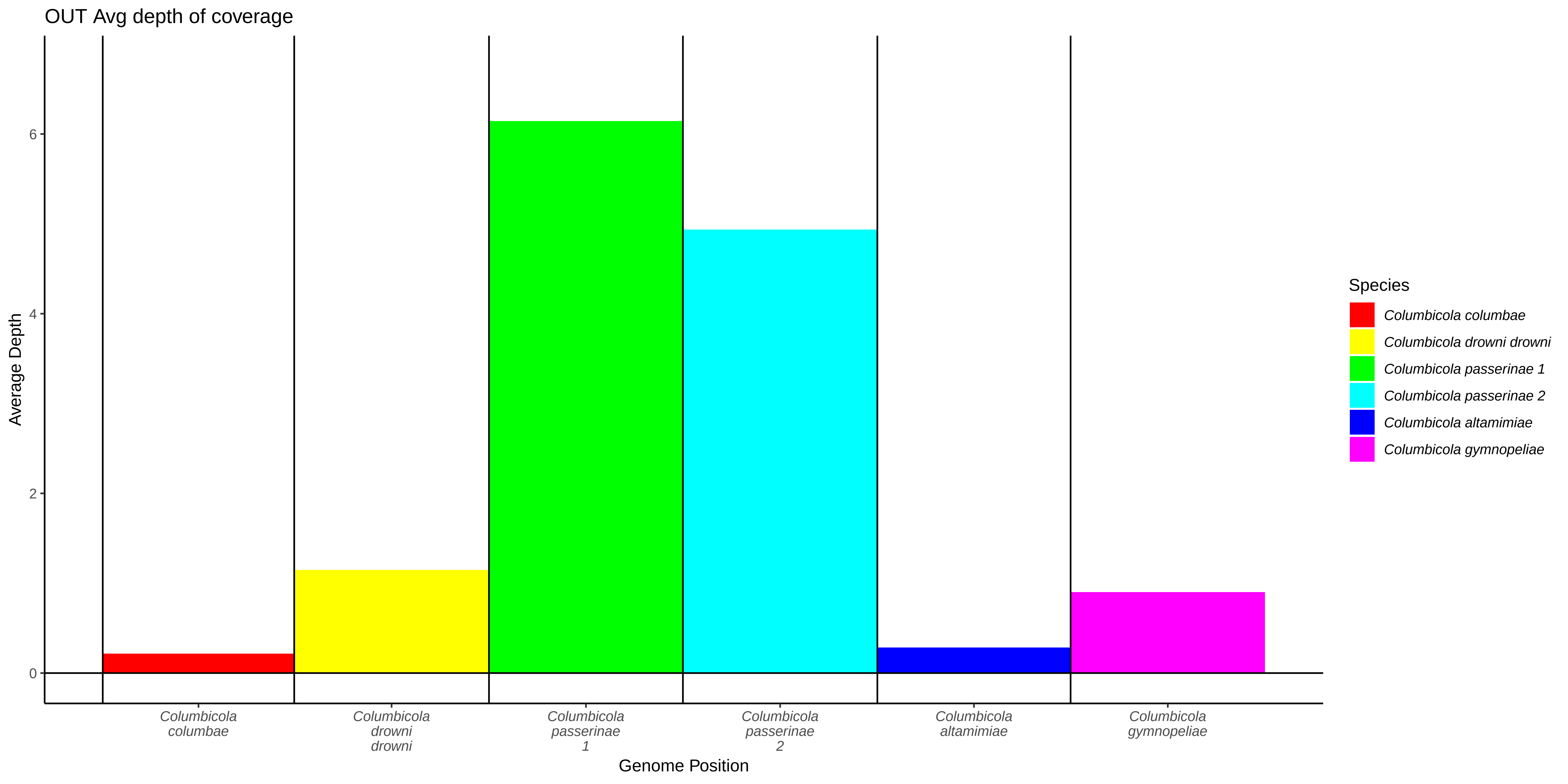

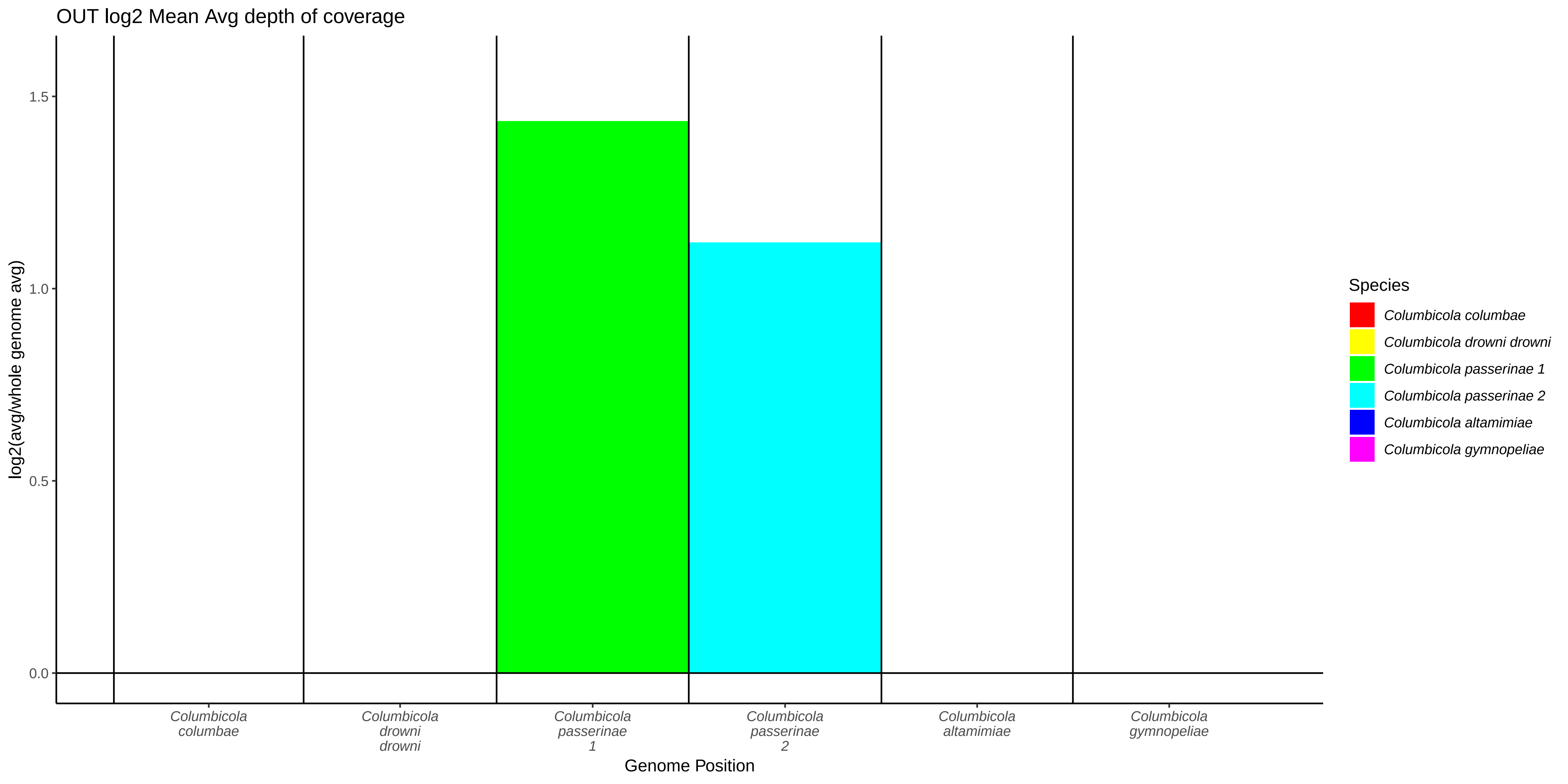

### OUT_speciesDepth_Cosp_Copas_11_7_2016_8.pdf

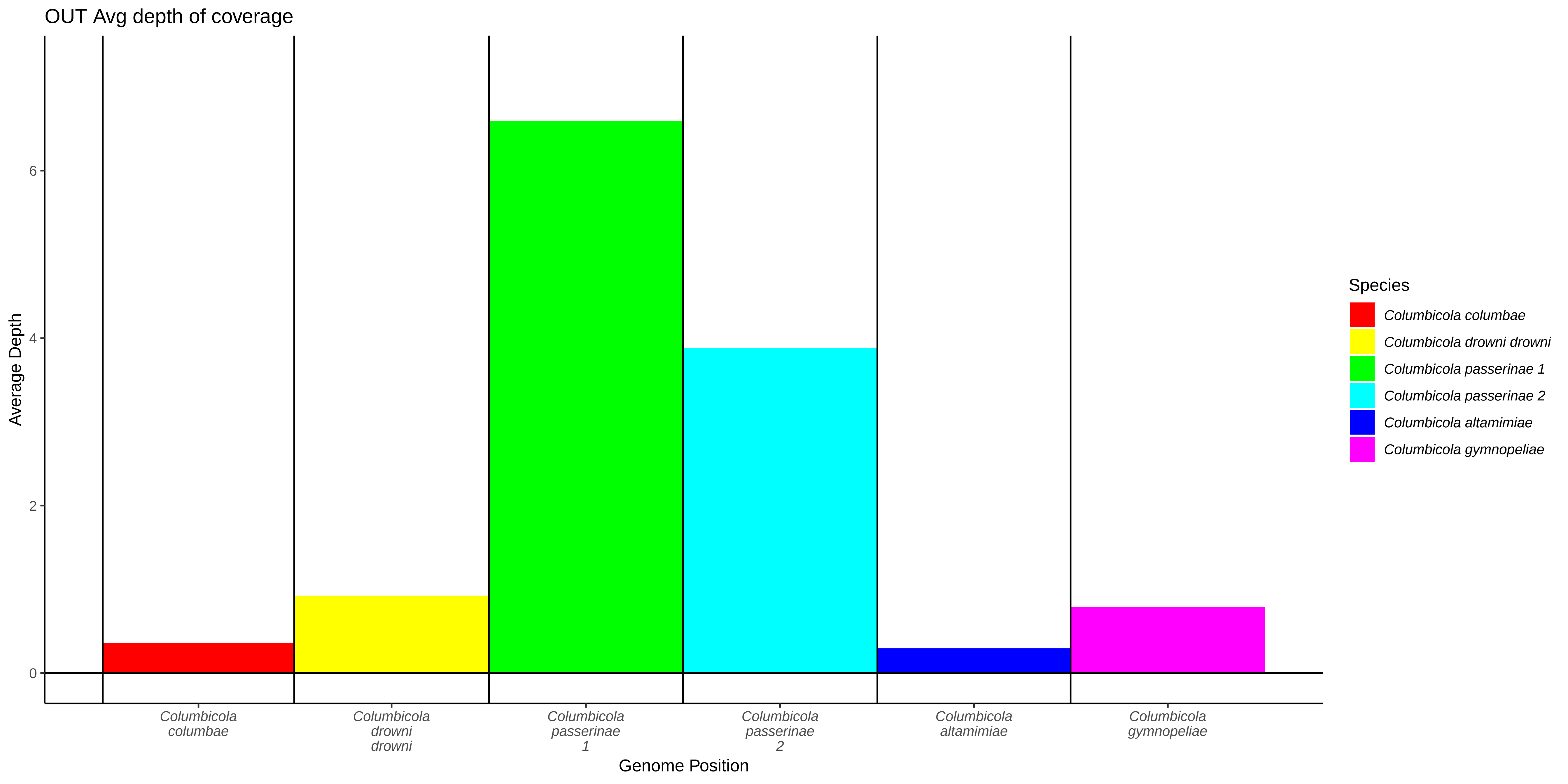

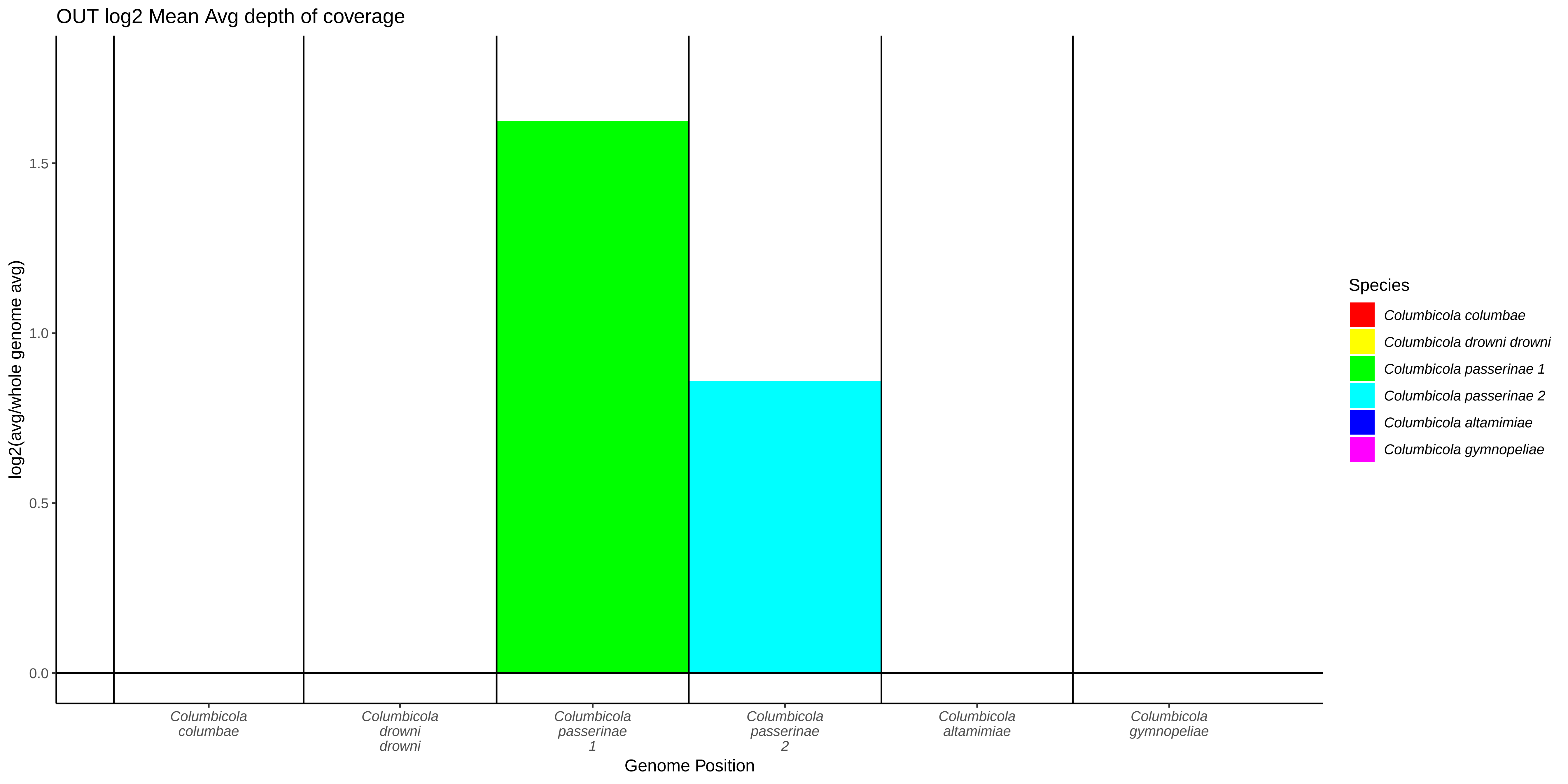

### OUT_speciesDepth_Cosp_Copas_11_7_2016_12.pdf

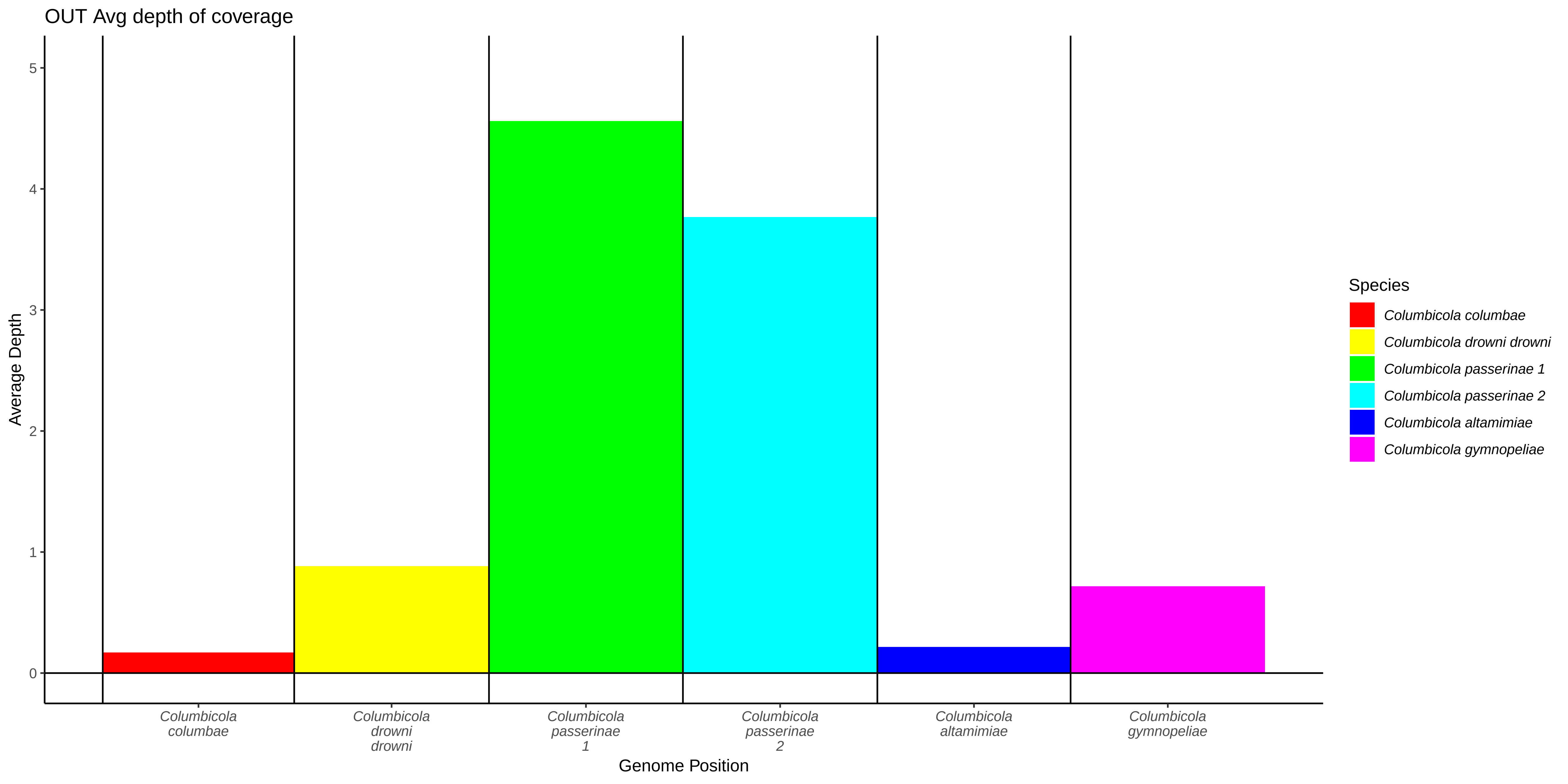

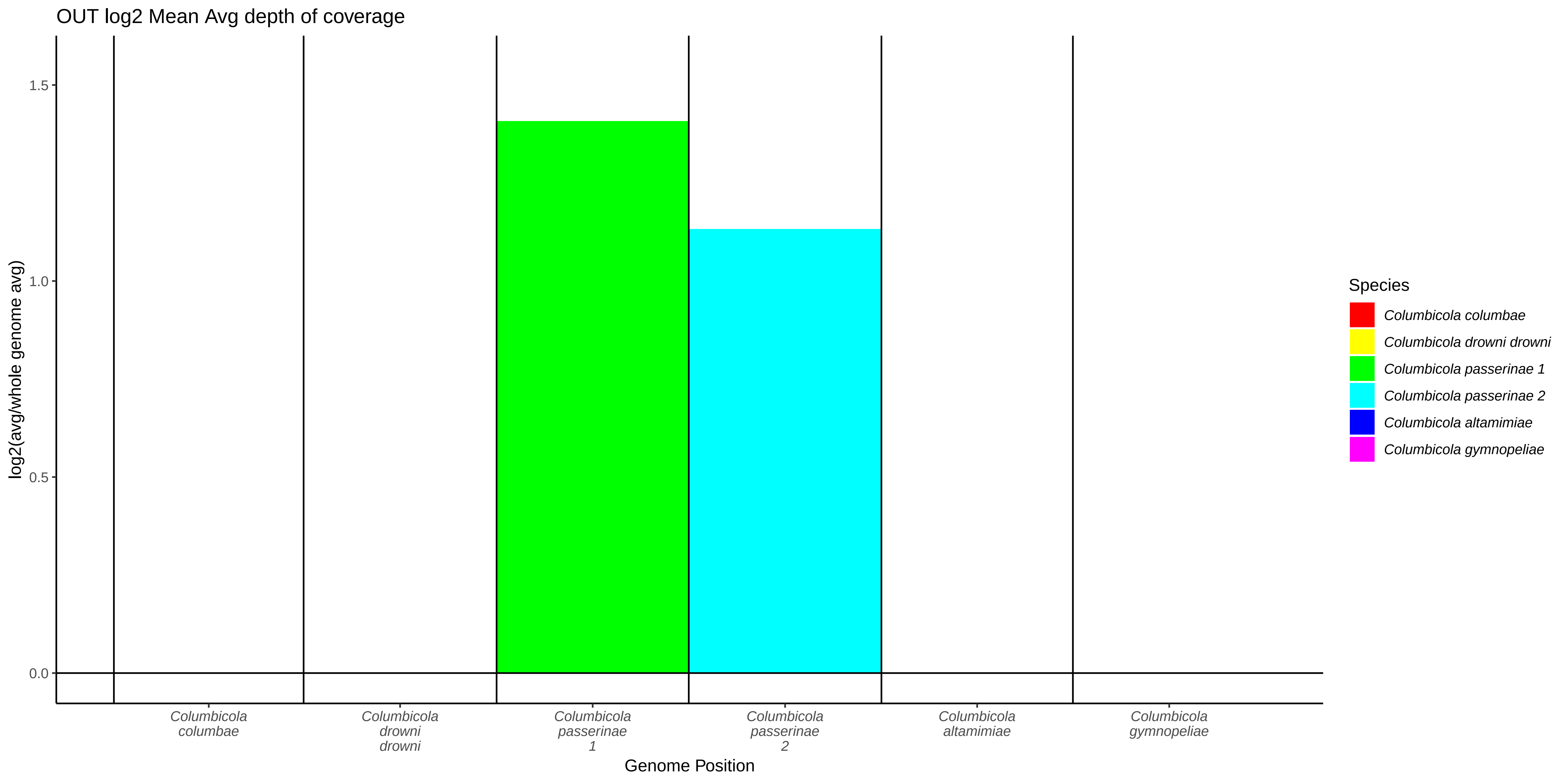

### OUT_speciesDepth_Cosp_Copas_11_7_2016_14.pdf

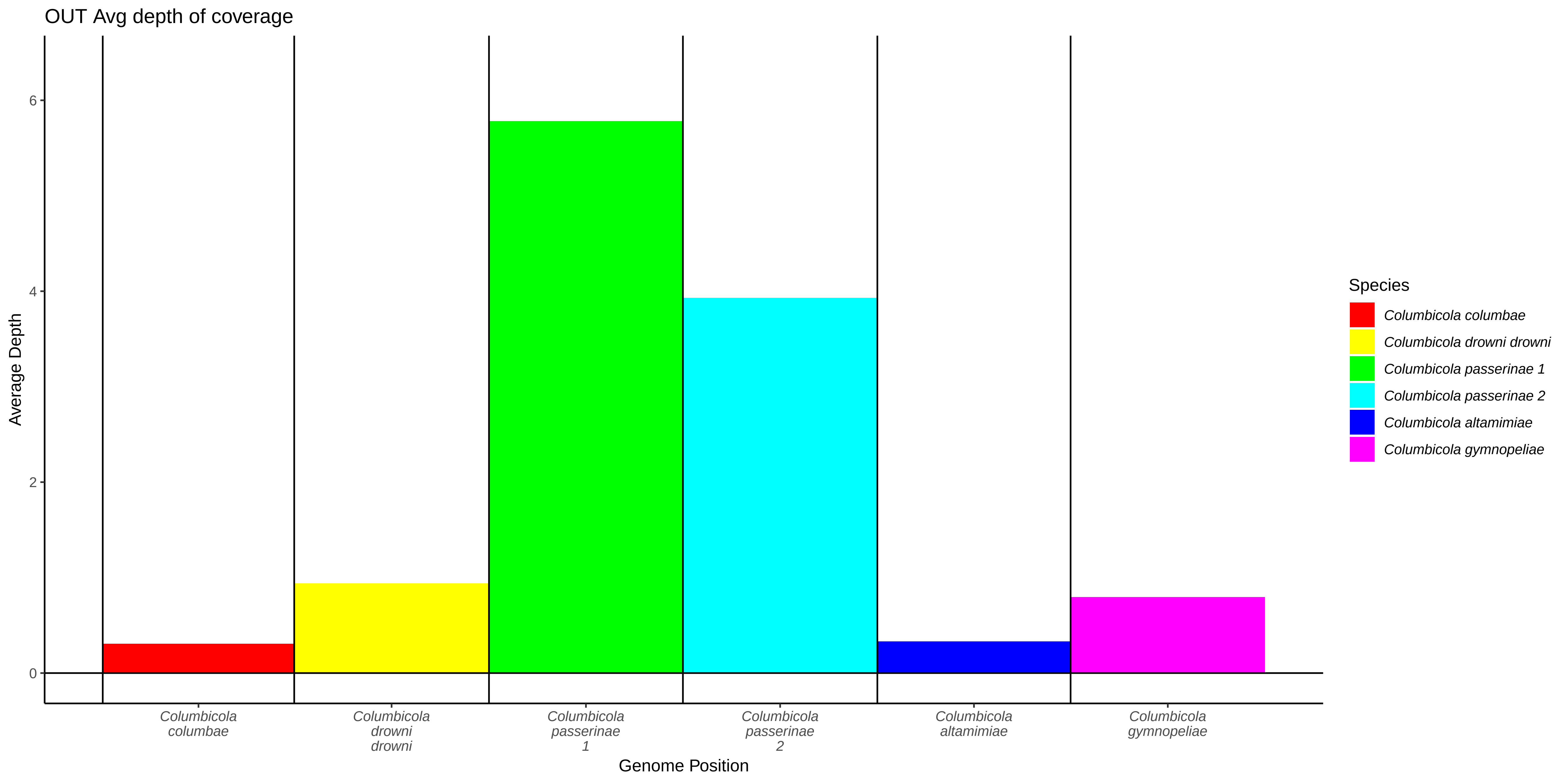

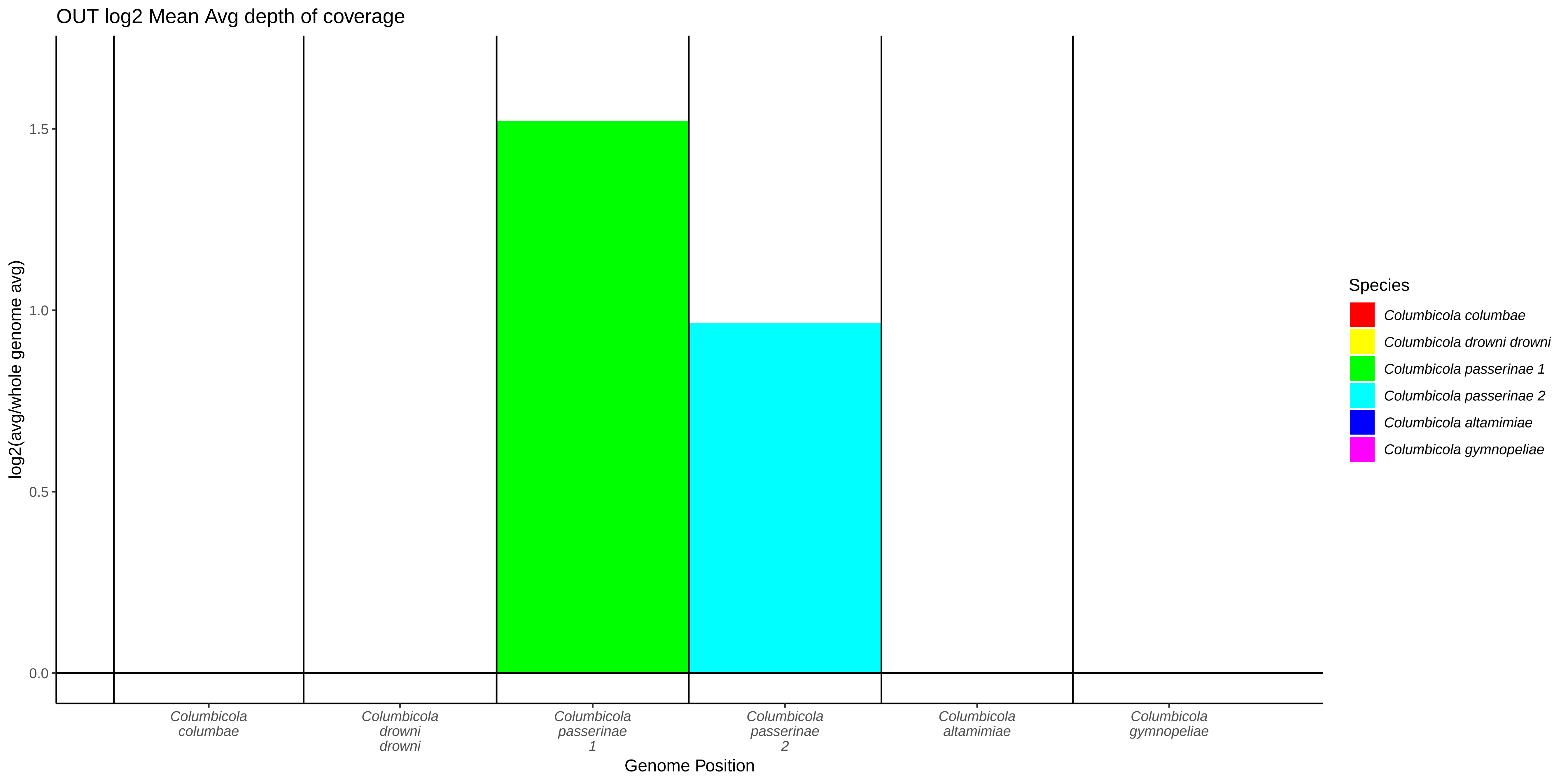

### OUT_speciesDepth_Cosp_Copas_11_7_2016_15.pdf

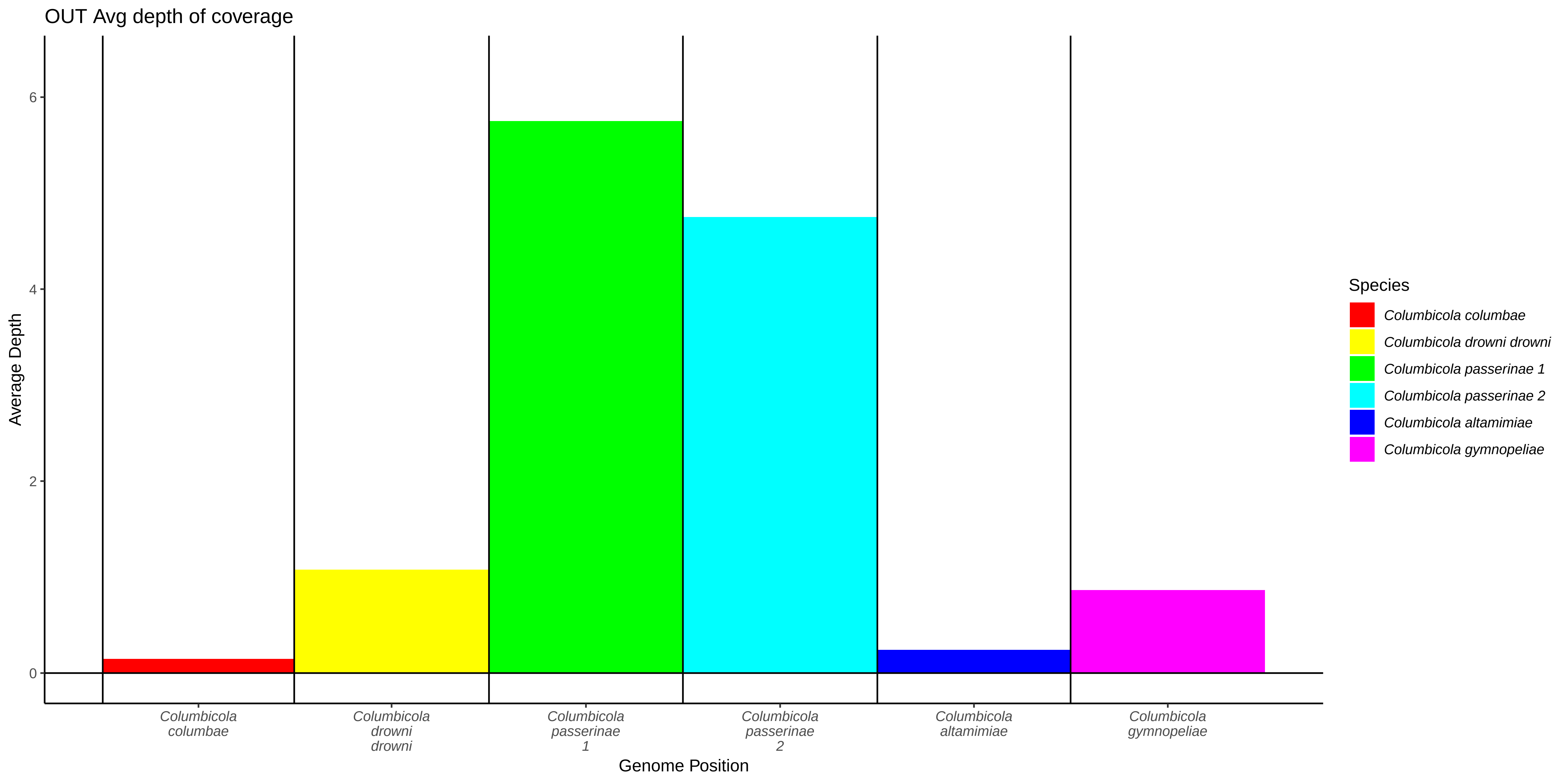

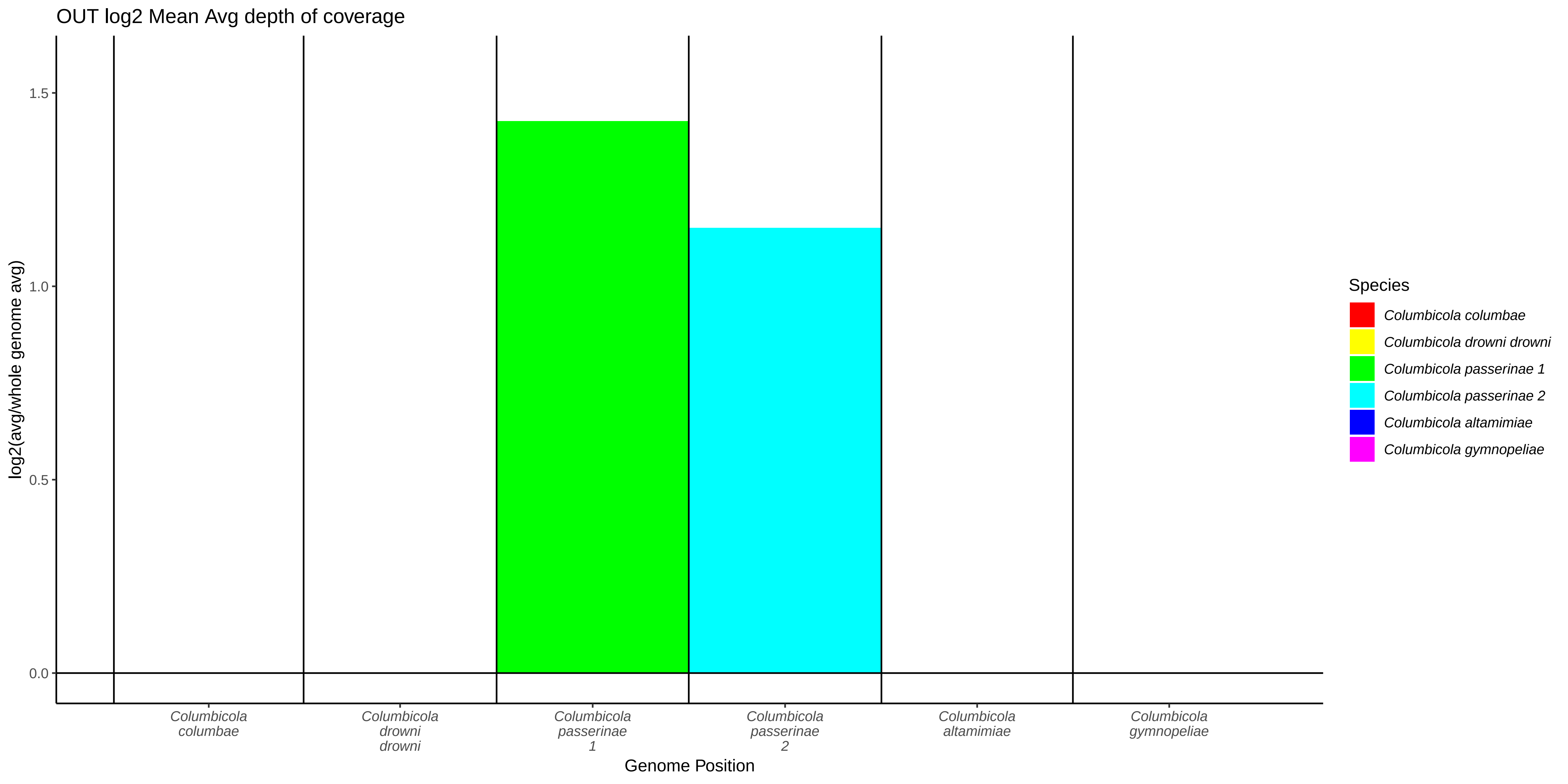

### OUT_speciesDepth_Cosp_Copic_11_2_2016_1.pdf

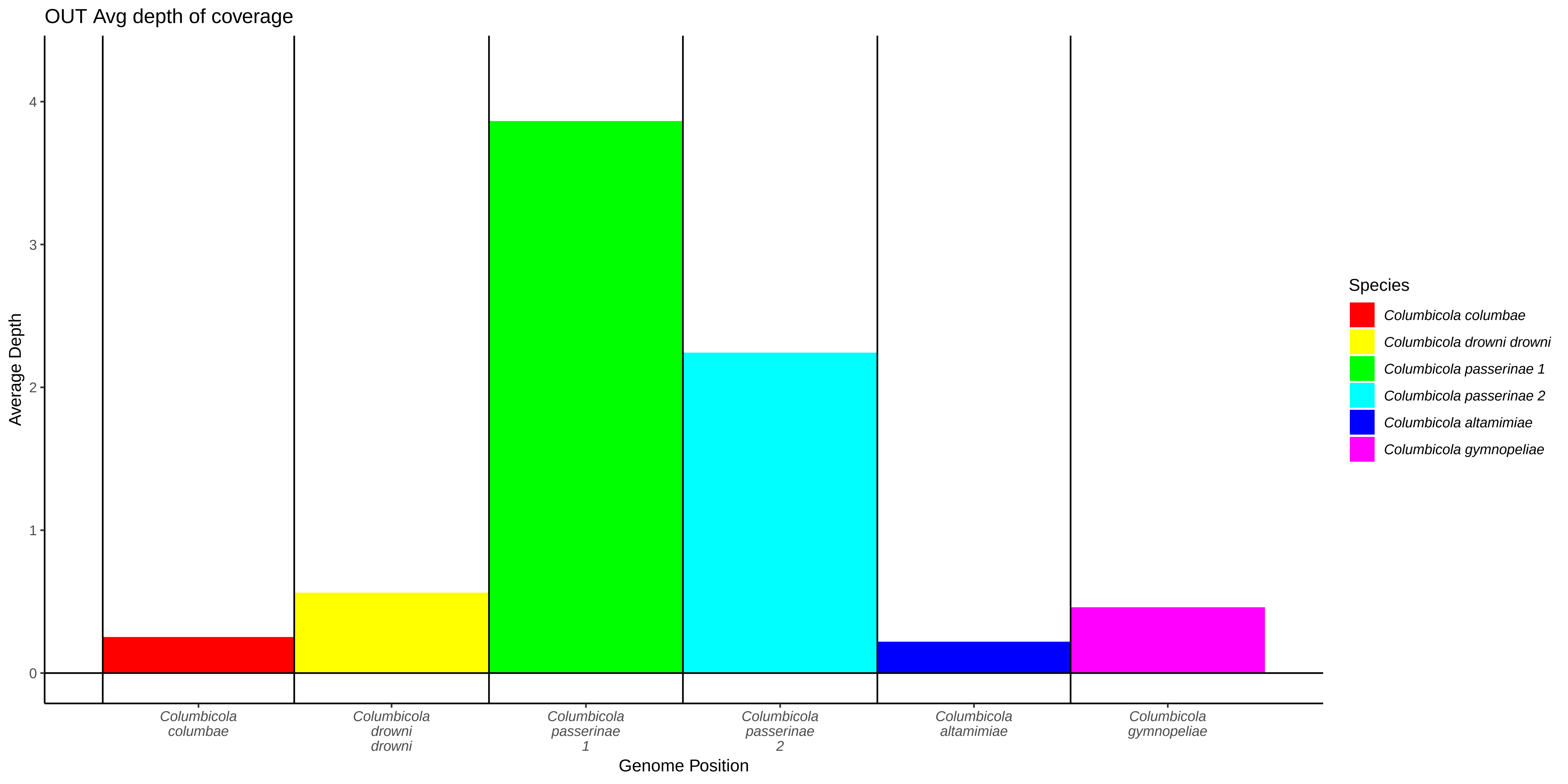

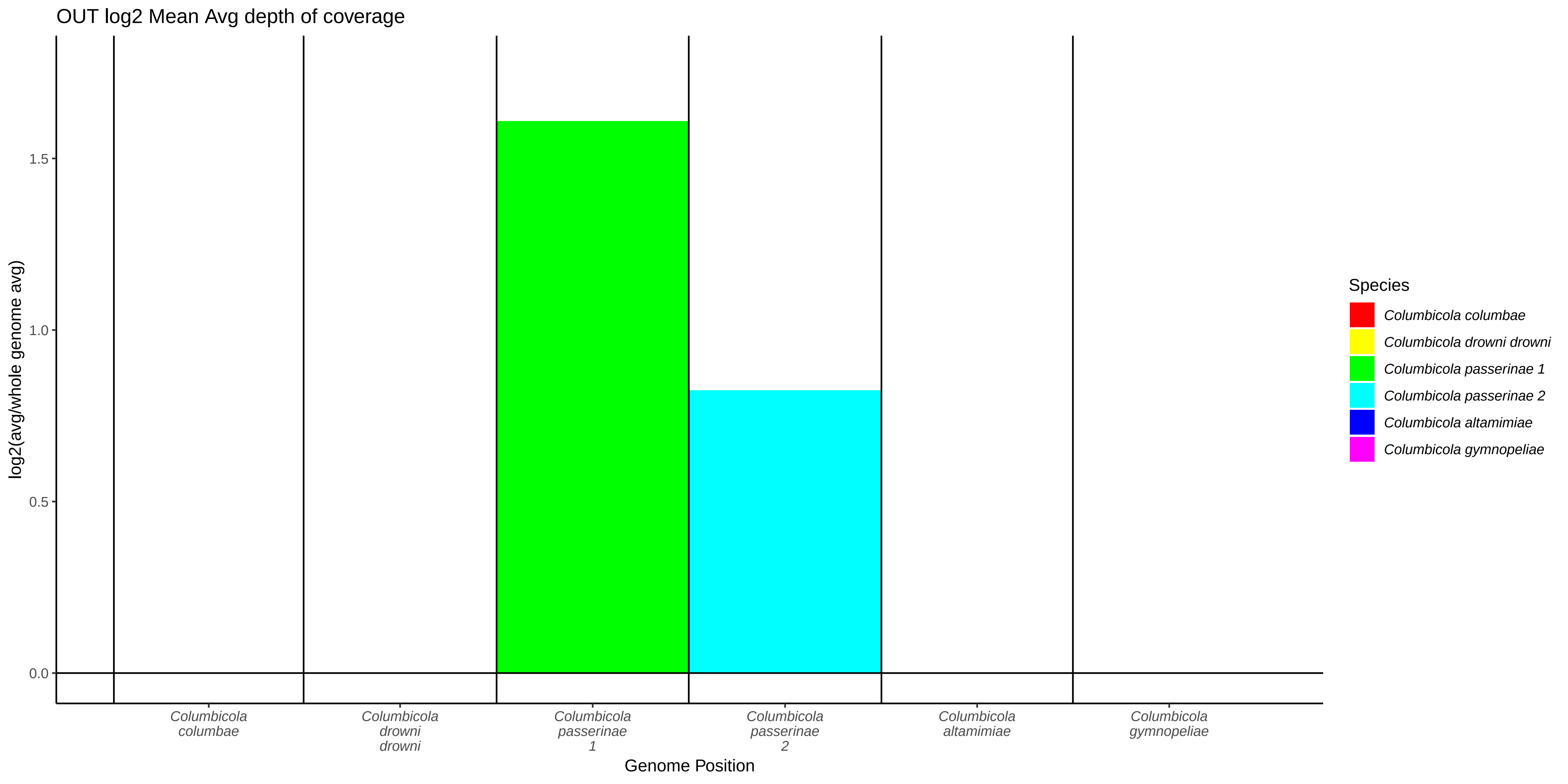

### OUT_speciesDepth_Cosp_Copic_11_2_2016_3.pdf

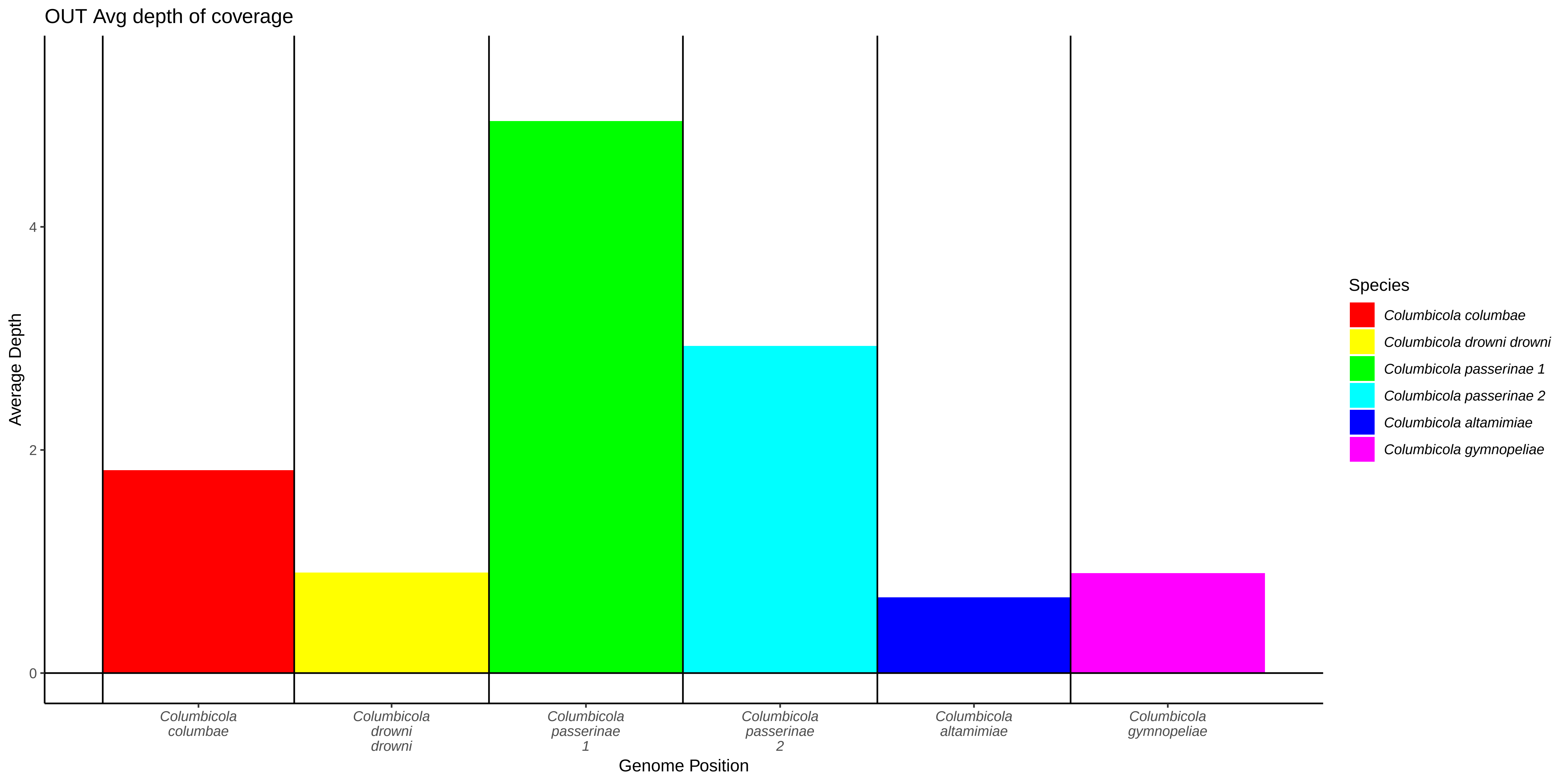

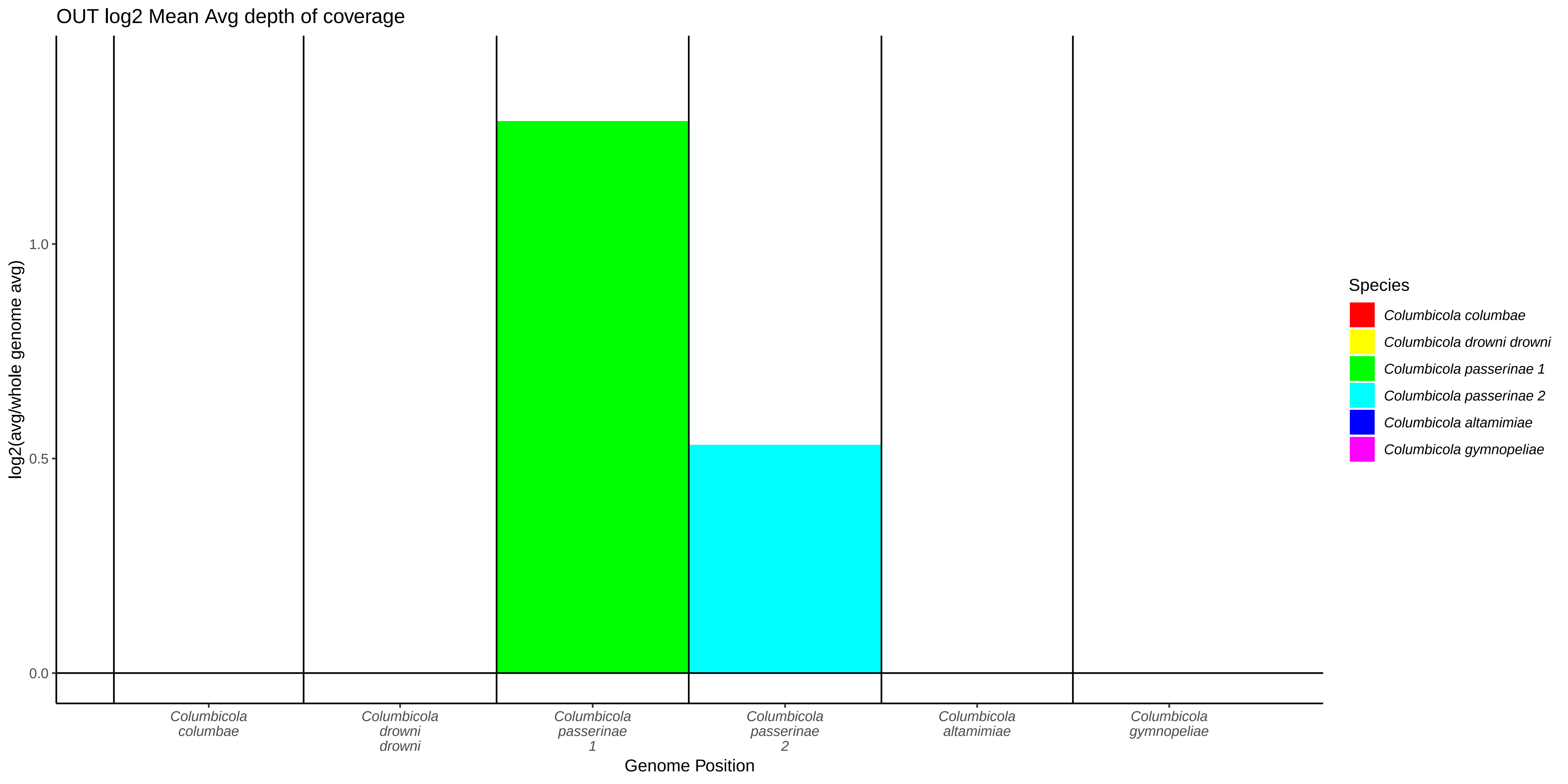

### OUT_speciesDepth_Cosp_Cosqu_11_7_2016_1.pdf

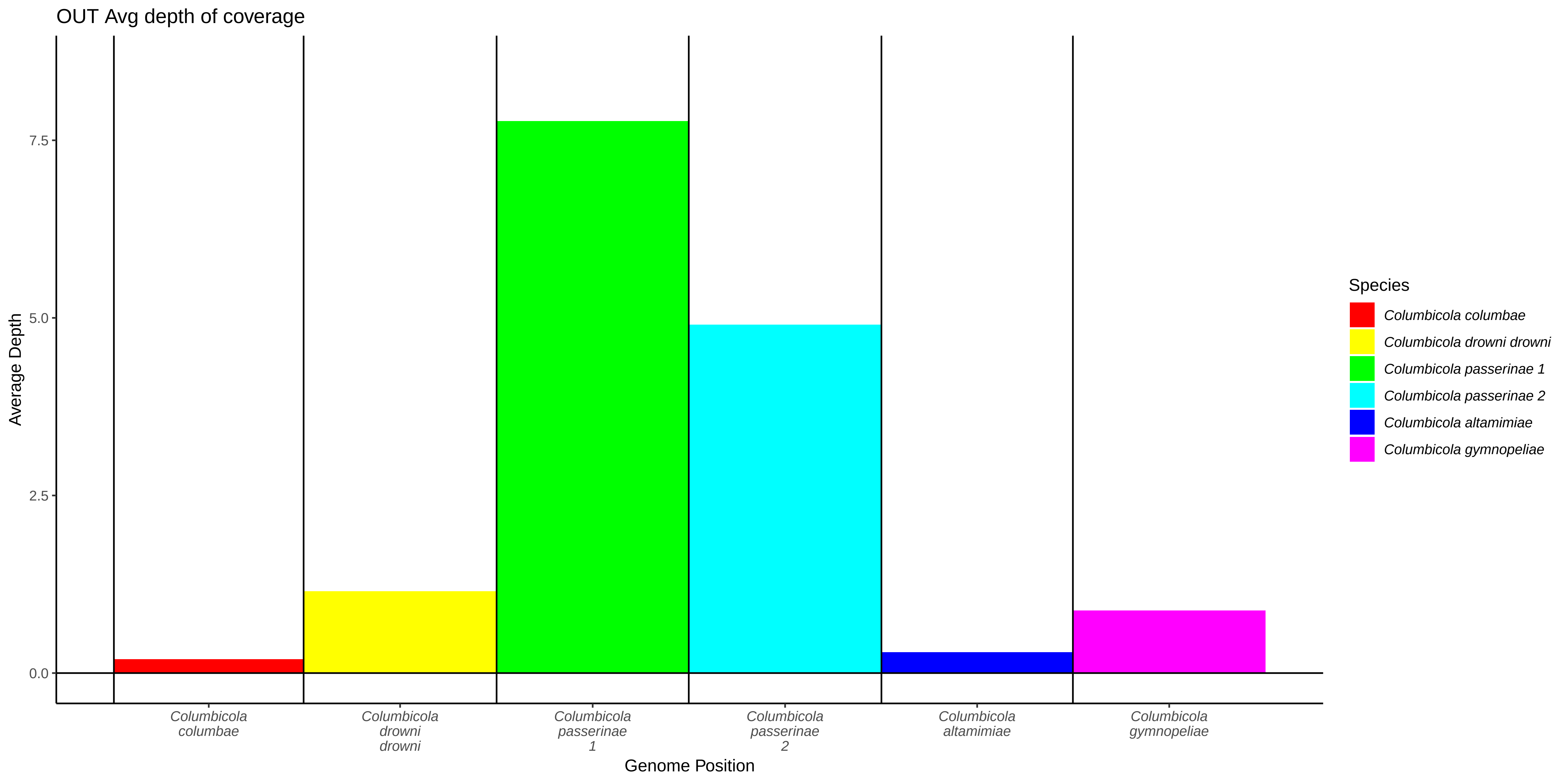

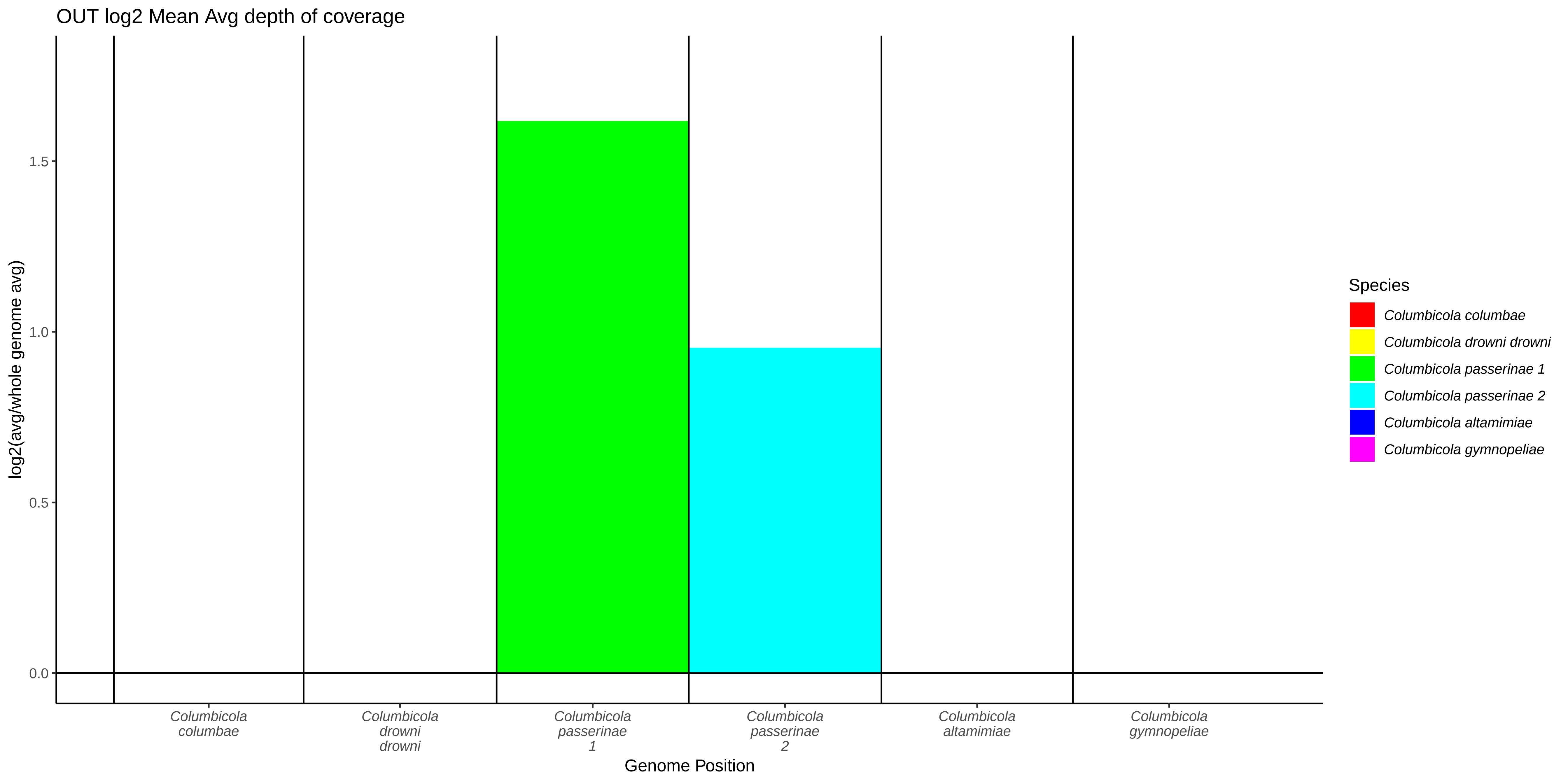

### OUT_speciesDepth_Cosp_Cosqu_11_7_2016_5.pdf

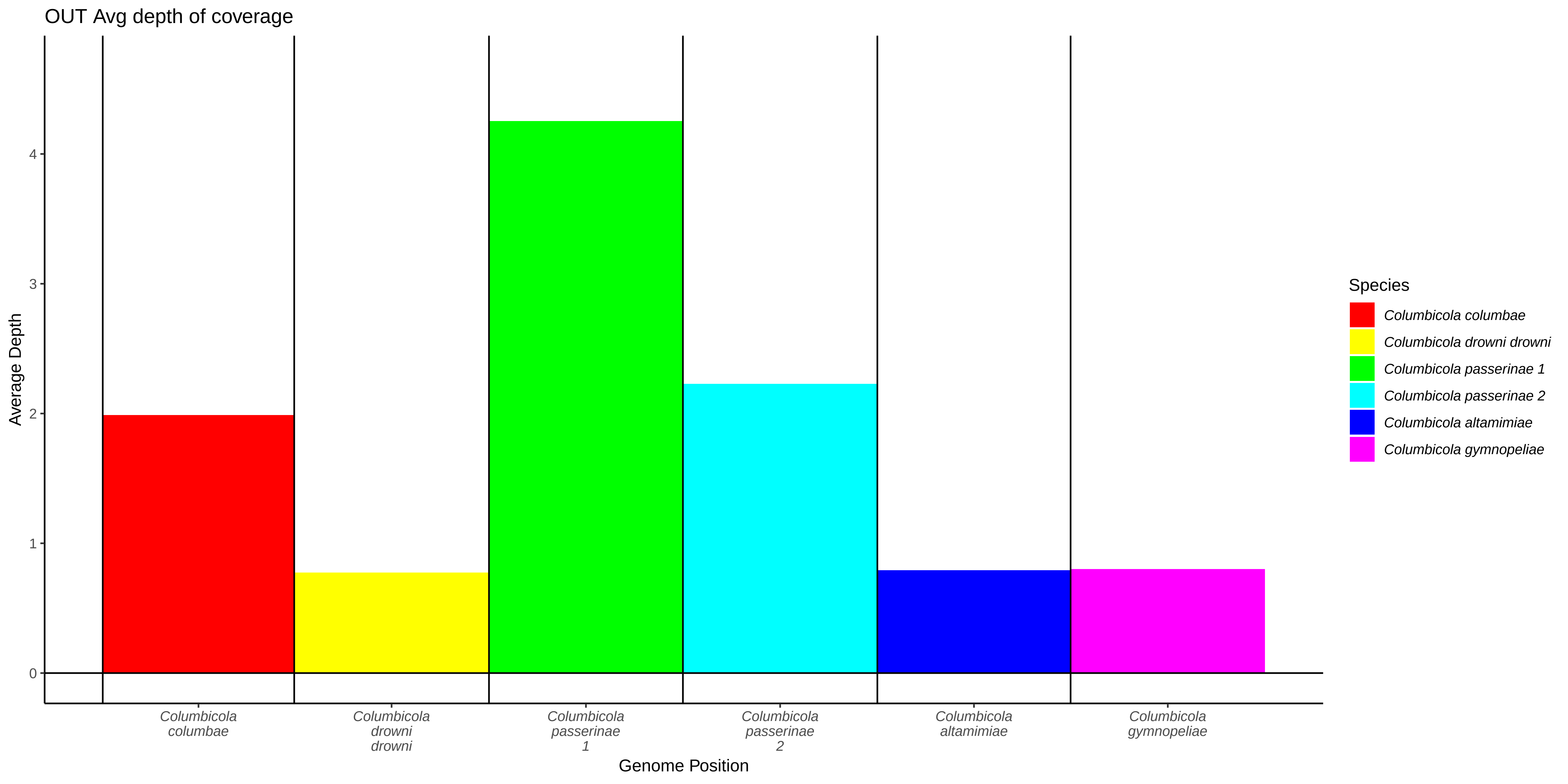

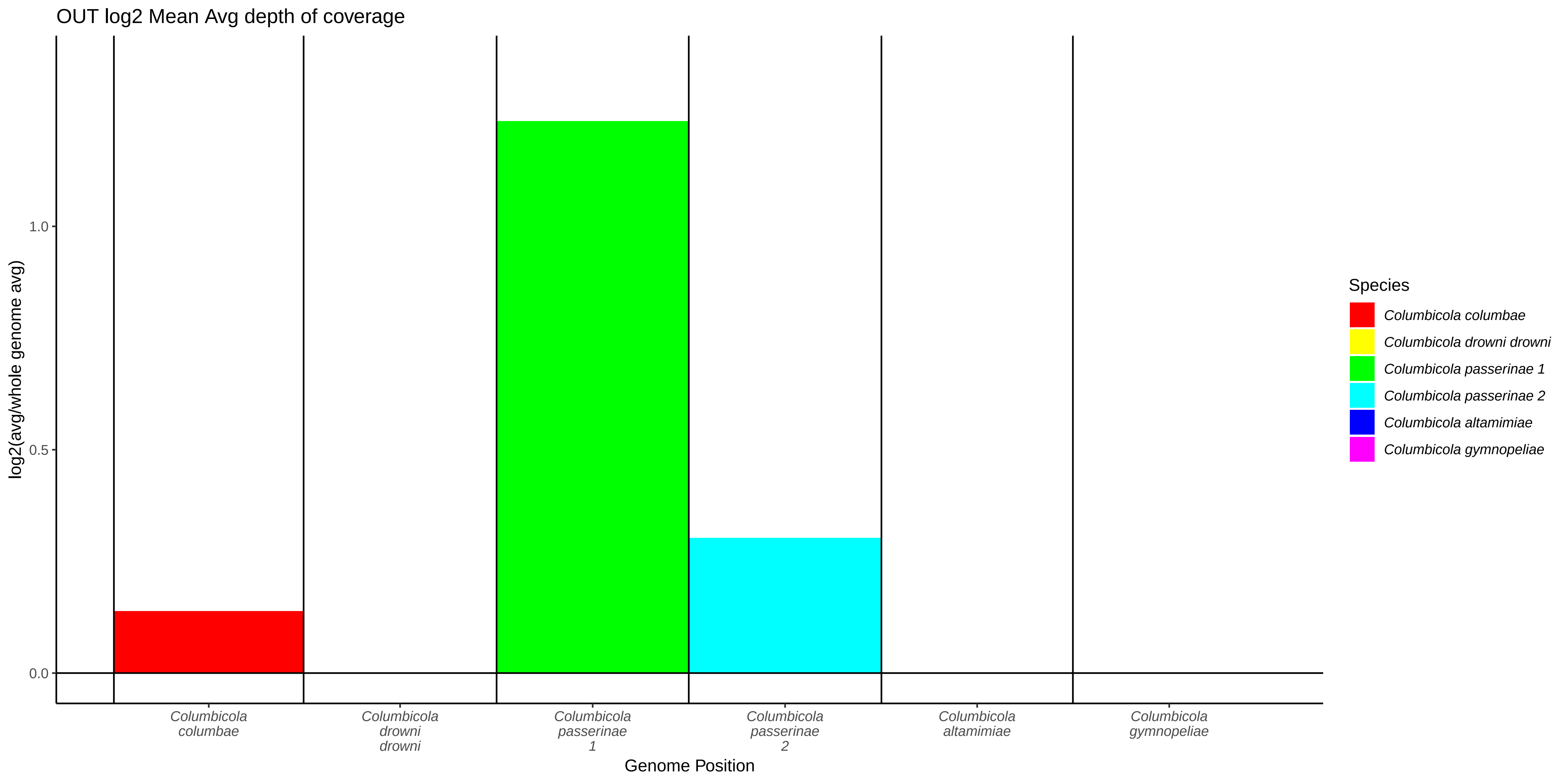

### OUT_speciesDepth_Cosp_Cotal_11_2_2016_8.pdf

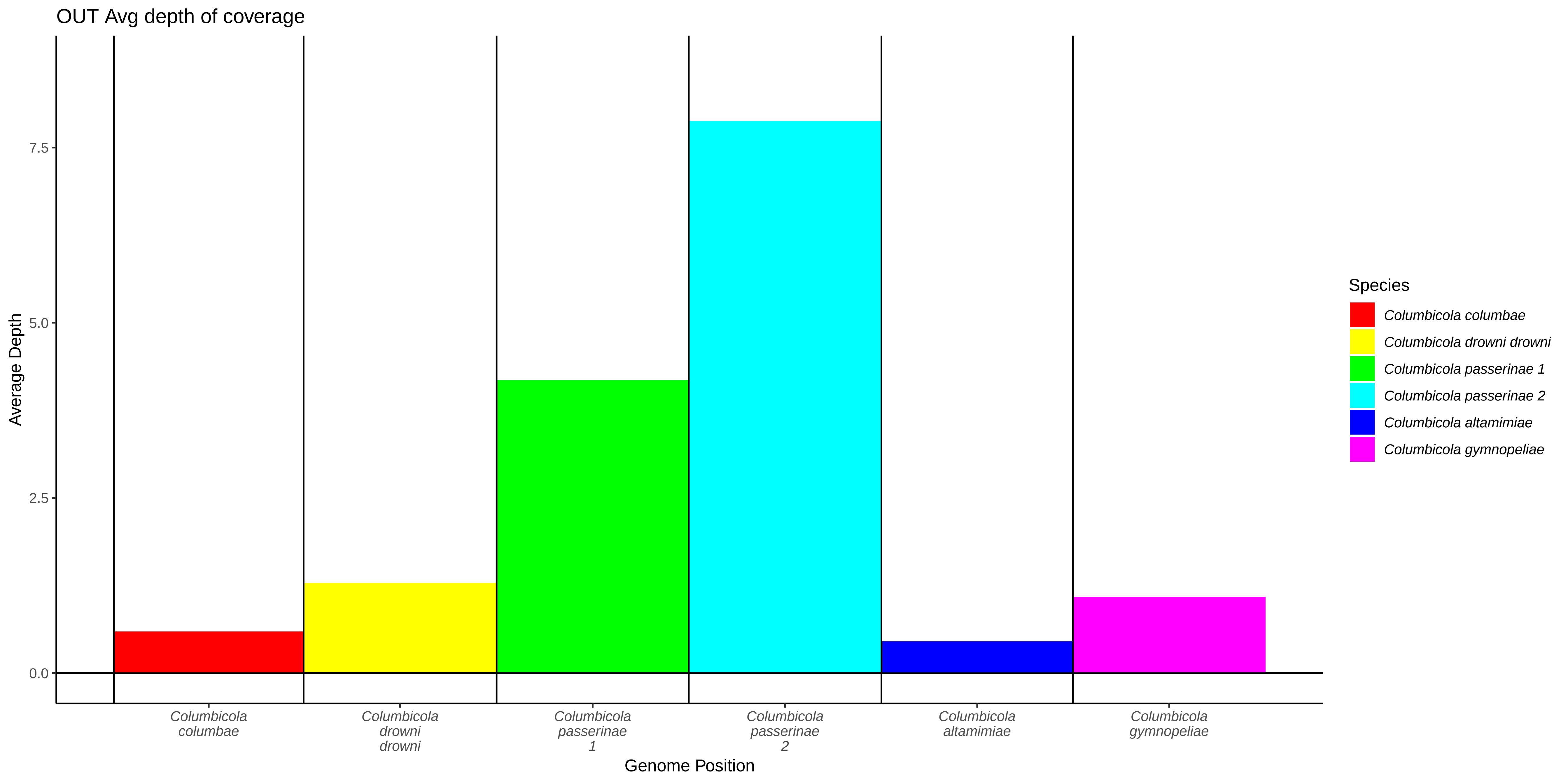

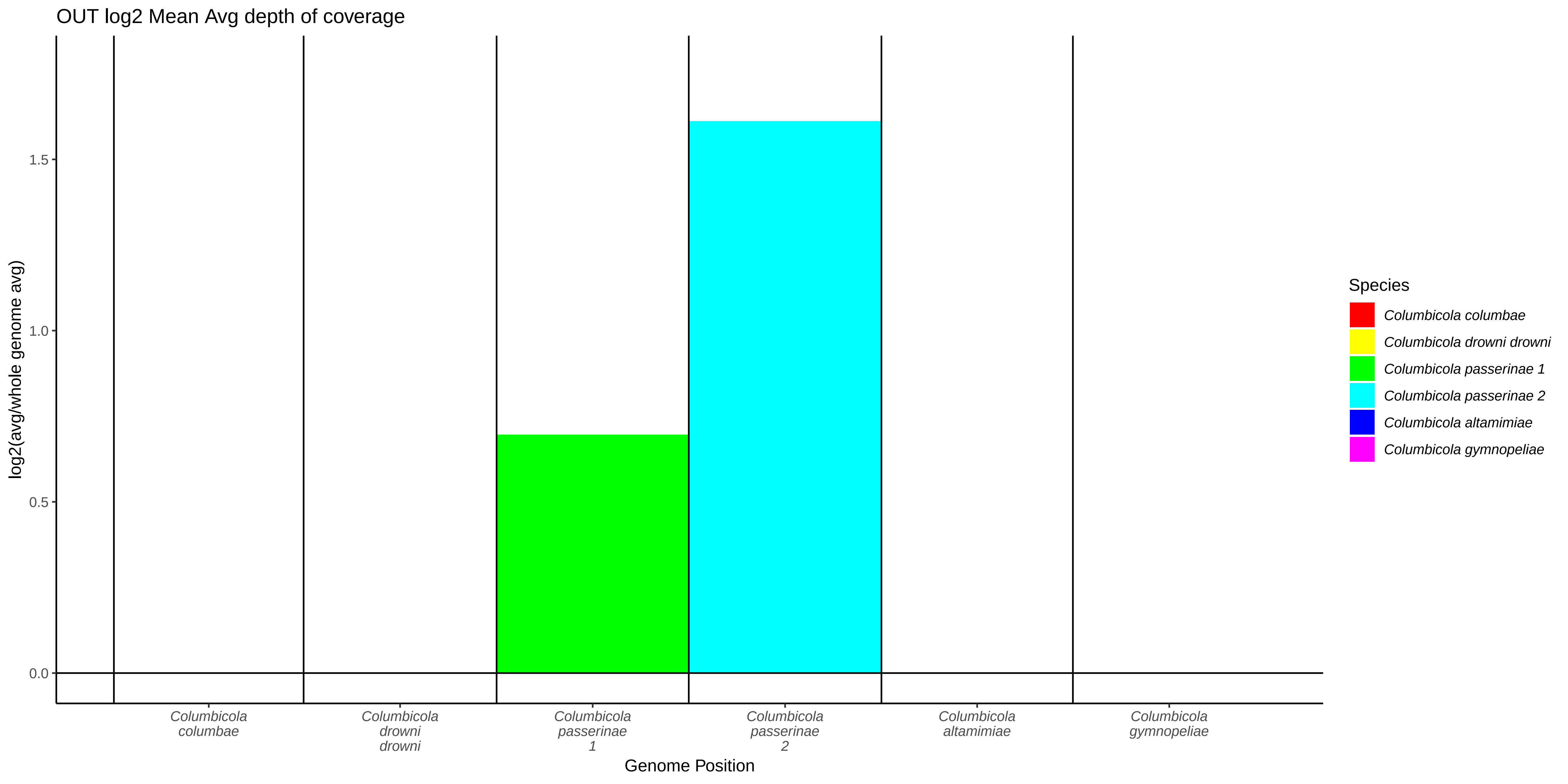

### OUT_speciesDepth_Cosp_Cotal_11_2_2016_15.pdf

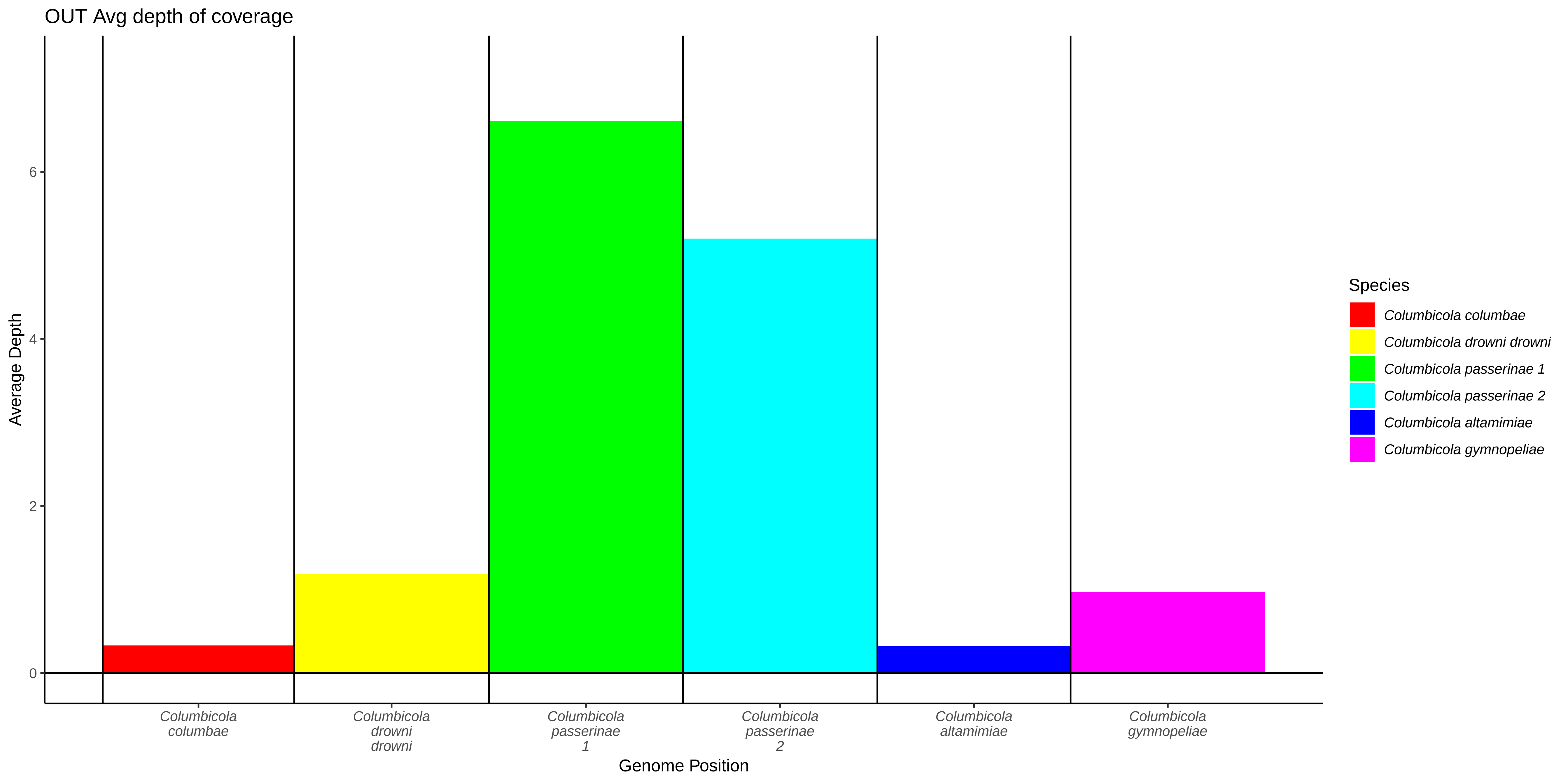

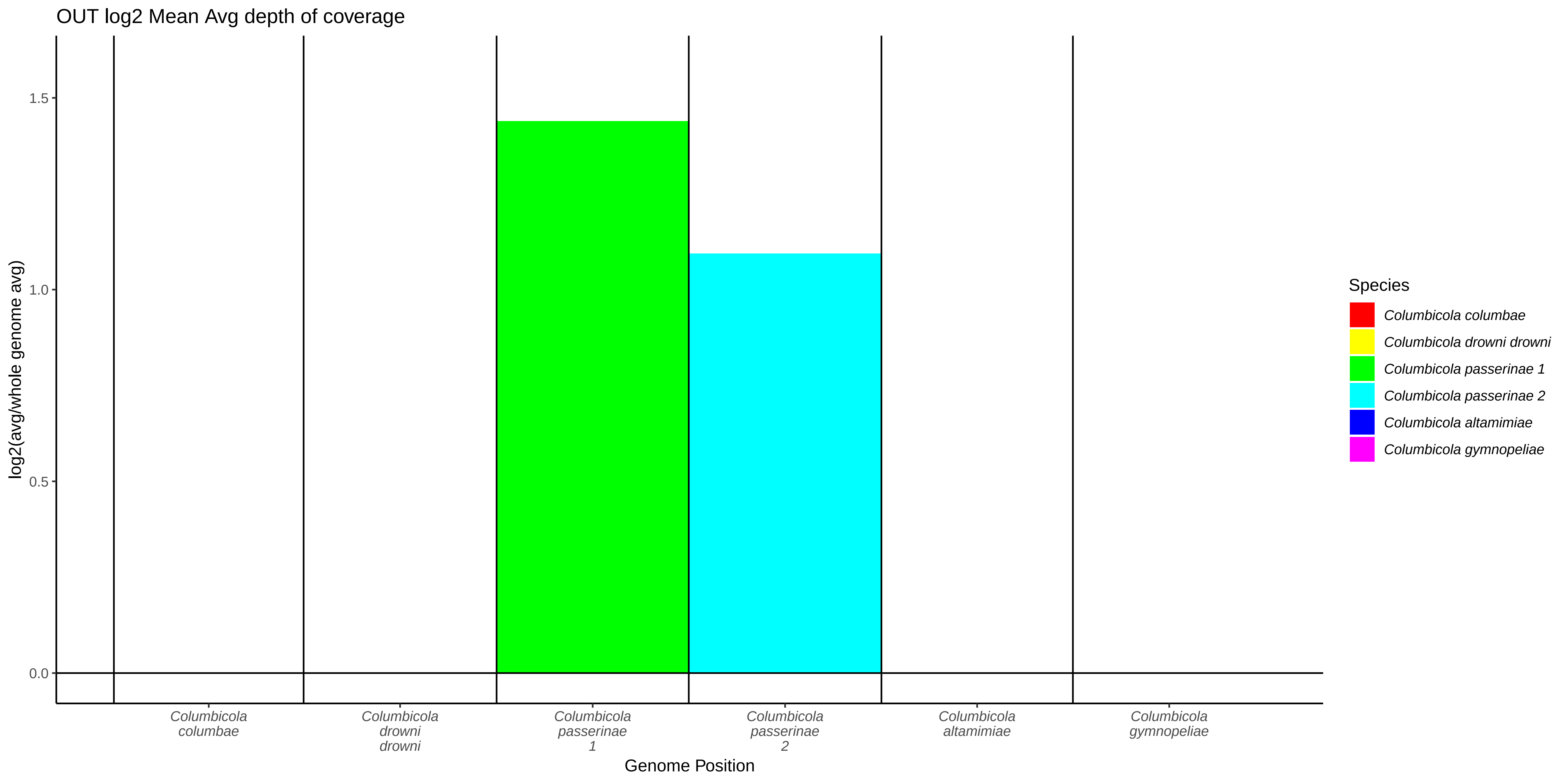

### OUT_speciesDepth_Cosp_Cotal_11_7_2016_2.pdf

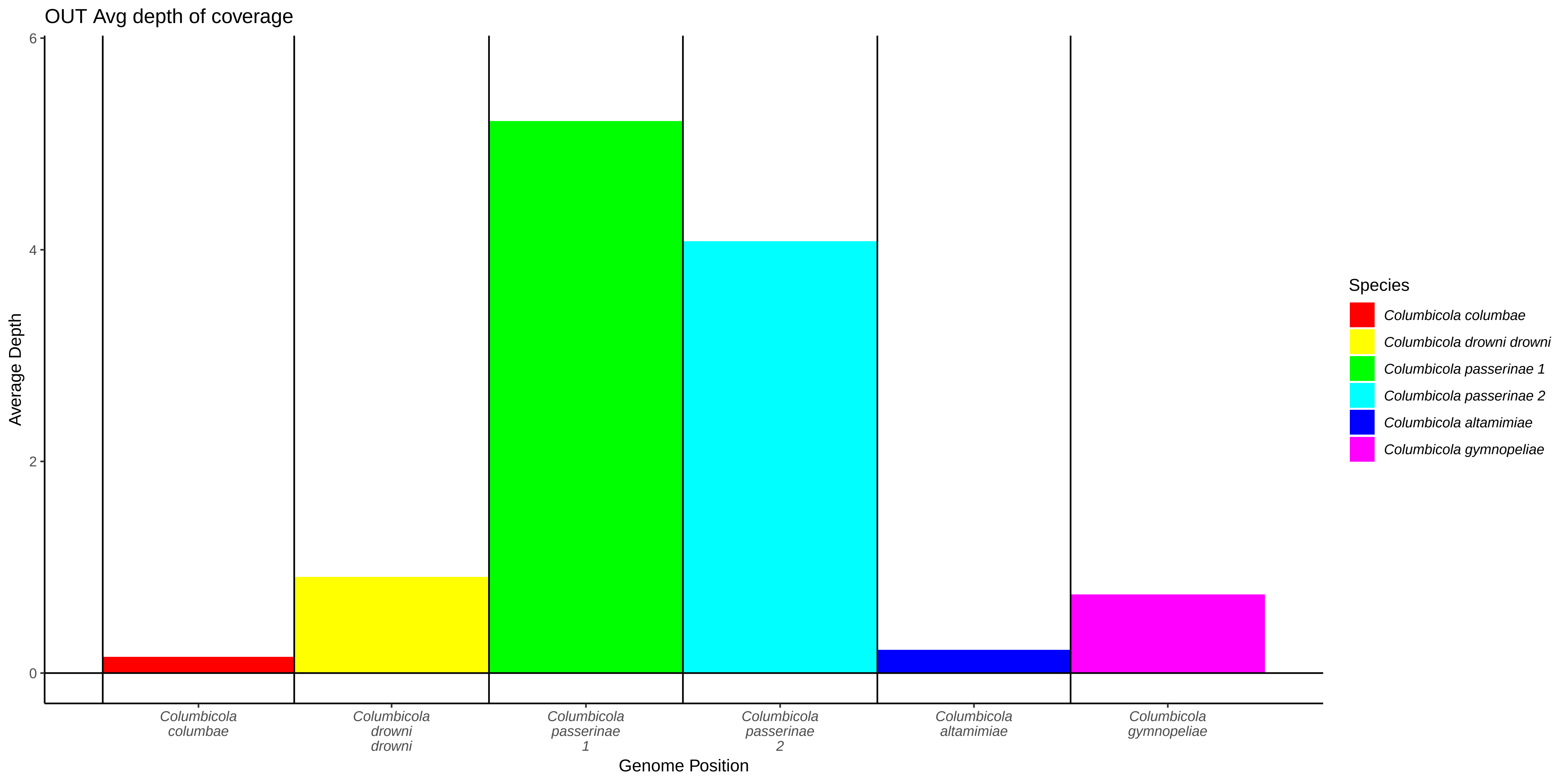

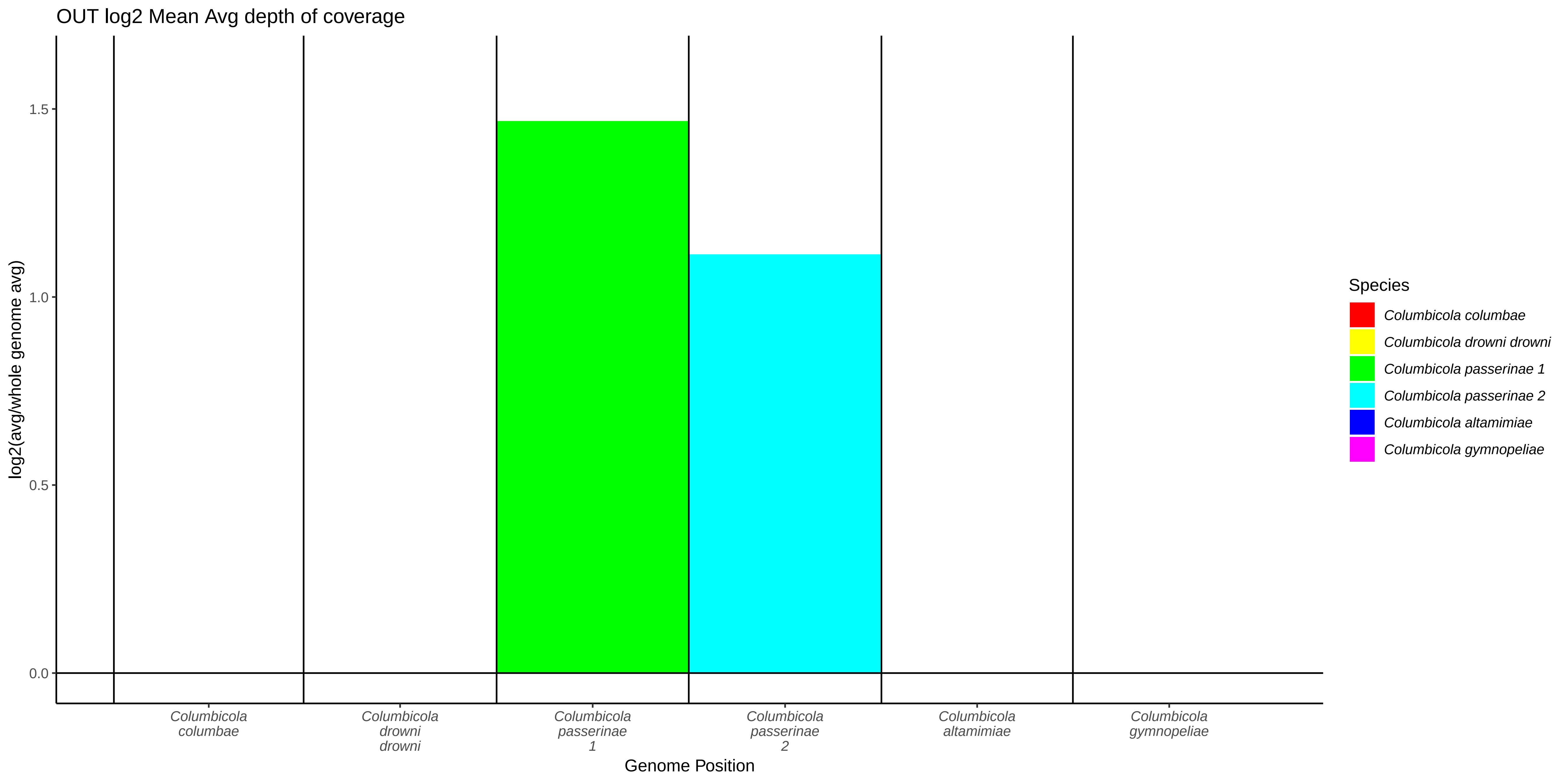
